# Uncharacterized yeast gene *YBR238C,* an effector of TORC1 signaling in a mitochondrial feedback loop, accelerates cellular aging via *HAP4*- and *RMD9*-dependent mechanisms

**DOI:** 10.1101/2023.07.04.547743

**Authors:** Mohammad Alfatah, Jolyn Jia Jia Lim, Yizhong Zhang, Arshia Naaz, Cheng Yi Ning Trishia, Sonia Yogasundaram, Nashrul Afiq Faidzinn, Jing Lin Jovian, Birgit Eisenhaber, Frank Eisenhaber

## Abstract

Uncovering the regulators of cellular aging will unravel the complexity of aging biology and identify potential therapeutic interventions to delay the onset and progress of chronic, aging-related diseases. In this work, we systematically compared gene sets involved in regulating the lifespan of *Saccharomyces cerevisiae* (a powerful model organism to study the cellular aging of humans) and those with expression changes under rapamycin treatment. Among the functionally uncharacterized genes in the overlap set, *YBR238C* stood out as the only one downregulated by rapamycin and with an increased chronological and replicative lifespan upon deletion. We show that *YBR238C* and its paralogue *RMD9* oppositely affect mitochondria and aging. *YBR238C* deletion increases the cellular lifespan by enhancing mitochondrial function. Its overexpression accelerates cellular aging via mitochondrial dysfunction. We find that the phenotypic effect of *YBR238C* is largely explained by *HAP4*- and *RMD9*-dependent mechanisms. Further, we find that genetic or chemical-based induction of mitochondrial dysfunction increases TORC1 (Target of Rapamycin Complex 1) activity that, subsequently, accelerates cellular aging. Notably, TORC1 inhibition by rapamycin (or deletion of *YBR238C*) improves the shortened lifespan under these mitochondrial dysfunction conditions in yeast and human cells. The growth of mutant cells (a proxy of TORC1 activity) with enhanced mitochondrial function is sensitive to rapamycin whereas the growth of defective mitochondrial mutants is largely resistant to rapamycin compared to wild type. Our findings demonstrate a feedback loop between TORC1 and mitochondria (the <u>TO</u>RC1-<u>MI</u>tochondria-<u>TO</u>RC1 (TOMITO) signaling process) that regulates cellular aging processes. Hereby, *YBR238C* is an effector of TORC1 modulating mitochondrial function.

## INTRODUCTION

Healthy aging is crucially determined by cellular functions, and their defective status is associated with premature dysfunction and/or depletion of critical cell populations and various aging-associated pathologies such as neurodegenerative diseases, cancer, cardiovascular disorders, diabetes, sarcopenia and maculopathy ^1–5^. Advancements in biomedical research including genome-wide screening in different aging model organisms, identified several biological pathways that support the unprecedented progress in understanding overlapping profiles between aged cells and different chronic aging-associated diseases ^6–9^. In principle, uncovering the molecular mechanisms that drive cellular aging identifies potential drug targets and can fuel the development of therapeutics to delay aging and increase healthspan ^10–13^.

Given the evolutionary conservation of many aging-related pathways, yeast is one of the aging model organisms that have been extensively used to study the biology of human cellular aging by analyzing chronological lifespan (CLS) and replicative lifespan (RLS) under various conditions ^14–17^. The CLS is the duration of time that a non-dividing cell is viable. This is a cellular aging model for post-mitotic human cells such as neurons and muscle cells. The RLS defines as the number of times a mother cell divides to form daughter cells. Such experiments provide a replicative human aging model for mitotic cells such as stem cells. Genome-wide or individual gene deletion strains’ screening identified thousands of genes affecting cellular lifespan. These lists are a rich resource to identify unique genetic regulators, functional networks, and interactions of aging hallmarks relevant to cellular lifespan ^1,4,6,10–13^. At the same, these lists are rich in genes of unknown function, a class of genes that, unfortunately, got increasingly ignored by the attention of research teams ^18–20^.

We systematically examined the available genetic data on aging and lifespan for budding yeast, *Saccharomyces cerevisiae*, from various sources. Our bioinformatics analyses revealed consensus lists of genes regulating yeast CLS and RLS. These gene sets were compared with lists of genes the expression of which changed under TORC1 inhibition with rapamycin. When we turned our attention to the functionally uncharacterized genes involved, we found *YBR238C* the only one among the latter that increases both CLS and RLS upon deletion and that is downregulated by rapamycin. Transcriptomics and biochemical experiments revealed that *YBR238C* negatively regulates mitochondrial function, largely via *HAP4*- and *RMD9*-dependent mechanisms, and thereby affects cellular lifespan. Surprisingly, *YBR238C* and its paralogue *RMD9* oppositely influence mitochondrial function and cellular aging.

Our chemical genetics and metabolic analyses unravel a feedback loop of the interaction of TORC1 with mitochondria that affect cellular aging. *YBR238C* is an effector of TORC1 that modulates mitochondrial function. We also show that mitochondrial dysfunction induces TORC1 activity enhancing cellular aging. In turn, TORC1 inhibition in yeast and human cells with mitochondrial dysfunction enhances their cellular survival.

## RESULTS

### Genome-wide survey of genes affecting cellular lifespan in yeast in accordance with literature and public databases

Lists of scientific literature instances mentioning a yeast gene as affecting lifespan were downloaded from the databases SGD (*Saccharomyces cerevisiae* genome database) ^21^, and GenAge ^22^. In most cases, the experiments referred to are gene deletion phenotype studies. After processing the files for the mentioning of ‘increase/decrease’ of ‘chronological lifespan’ (CLS) and/or ‘replicative lifespan’ (RLS) as well as the suppression of gene duplicates, we found 2399 entries with distinct yeast genes in 15 categories reported to increase/decrease CLS and/or RLS under various conditions (Table 1). We collectively call them Aging Associated Genes (AAGs). Notably, about one-third of the total yeast genome belongs to that category. Downloaded files (as of 8^th^ November 2022), description of the processing details, and all the resulting gene lists are available in ‘Additional File 1’.

**Table 1.** Numbers of known aging associated genes (AAGs) reported in the scientific literature to affect (increase/decrease CLS/RLS) under a variety of conditions. The gene lists were extracted, after processing, from downloaded files (as of 8^th^ November 2022) using the databases SGD ^21^ and GenAge ^22^. The actual gene lists are available in Additional File 1’. The signs ‘+’ and ‘-‘ indicate that the respective annotation property is present or missing in that group of genes. We present the total number of genes in each category as the number of, as a trend, severely understudied/uncharacterized genes among those. The columns ‘RUG’ and ‘RDG’ show the numbers of rapamycin-up- and -downregulated genes in each of the 15 categories.

| category | CLS<br>increase | CLS<br>decrease | RLS<br>increase | RLS<br>decrease | Number<br>of genes | Understudied<br>genes | RUG | RDG |
| --- | --- | --- | --- | --- | --- | --- | --- | --- |
| 1 | + | - | - | - | 328 | 20 | 68 | 62 |
| 2 | - | + | - | - | 615 | 22 | 108 | 73 |
| 3 | - | - | + | - | 349 | 19 | 73 | 64 |
| 4 | - | - | - | + | 318 | 15 | 43 | 56 |
| 5 | + | + | - | - | 108 | 0 | 17 | 18 |
| 6 | + | - | + | - | 72 | 2 | 12 | 26 |
| 7 | + | - | - | + | 72 | 3 | 9 | 18 |
| 8 | - | + | + | - | 120 | 3 | 14 | 28 |
| 9 | - | + | - | + | 180 | 0 | 24 | 34 |
| 10 | - | - | + | + | 59 | 0 | 8 | 14 |
| 11 | + | + | + | - | 29 | 1 | 7 | 7 |
| 12 | + | + | - | + | 45 | 1 | 7 | 9 |
| 13 | + | - | + | + | 30 | 1 | 2 | 11 |
| 14 | - | + | + | + | 55 | 1 | 5 | 11 |
| 15 | + | + | + | + | 19 | 0 | 0 | 2 |

Whereas most of the genes (1610, 67%) have been mentioned for just one of the four scenarios (“CLS increase”, “CLS decrease”, “RLS increase”, and “RLS decrease”), the remaining genes have been described for several alternative and/or even opposite/conflicting outcomes. 19 genes are brought up in context with all four scenarios. As the experimental conditions varied among the reports and the gene networks are complex, this is not necessarily a contradiction. Yet (see below) some of the cases are actually annotation errors at the database level.

We explored the potential enrichment of gene ontology (GO) terms among the genes involved in the 15 categories with the help of DAVID WWW server ^23^. The most significant result is the enrichment of ontology terms related to mitochondrial function among the genes annotated with the single qualifier “CLS decrease”. The top mitochondrial term cluster has an enrichment score 18.2 (gene-wise P-values and Benjamini-Hochberg values all below 6.e-13); a second one related to mitochondrial translation has the score 4.7. Enrichment of mitochondrial ontology terms has also been observed for “CLS decrease” in combination with “CLS increase” or “RLS increase/decrease”.

Every other signal from the ontology is much weaker. Term enrichment (with scores near 2 or better) related to autophagy and vesicular transport pop up for CLS-annotated genes whereas terms connected with translation, DNA repairs, telomeres, protein degradation and signaling are observed with RLS-tagged gene lists.

We also explored the presence of uncharacterized or severely under-characterized genes in the 15 categories of AAGs (Table 1). In total, 944 genes are annotated in SGD as coding for a protein of unknown function. Additionally, we considered genes without dedicated gene name (only with six-letter locus tag) as dramatically under-characterized. Comparison of this combined list with those in the 15 categories reveals 88 severely understudied AAGs that are candidates for enhanced attention from the scientific community. Thus, functionally insufficiently characterized genes have a great role in cellular aging-related processes.

### Rapamycin response genes overlap with the AAGs

Nutrient sensing dysregulation is one of the aging hallmarks ^10,11^. The conserved protein complex Target of Rapamycin Complex 1 (TORC1) senses nutrients such as amino acids and glucose and links metabolism with cellular growth and proliferation ^24–27^. TORC1 positively regulates aging, and its inhibition increases lifespan in various eukaryotic organisms including yeast and mammals ^12,25,26,28,29^. The drug rapamycin, initially discovered as an antifungal natural product produced by *Streptomyces hygroscopicus*, inhibits TORC1 and increases lifespan (Figures S1A and S1B) ^30,31^.

To explore the connection between nutrient signaling and cellular aging we mapped the TORC1 regulated genes with AAGs. We first identified rapamycin response genes (RRGs) by transcriptomics analysis (RNA-Seq for yeast *S. cerevisiae* BY4743 cells treated with rapamycin and DMSO control; Figure S1C). Relevant measurement results, lists of 2365 RRGs and supplementary methodical comments are available in ‘Additional File 2’.

As overlap of two RNA-Seq data analysis methods (see Methods), we identified 1155 rapamycin upregulate genes (RUG) and 1210 rapamycin downregulated genes (RDG) (Figures S1D-S1F; Additional File 2). RNA-Seq results were confirmed for some genes by qRT-PCR (Figure S2A). We also checked the differential gene expressions in prototrophic yeast *S. cerevisiae* strain CEN.PK and found similar results as with the BY4743 strain (Figure S2B). To note, our transcriptomics analyses are, as a trend, consistent with previous RNA-Seq studies carried out under partially different experimental conditions ^32,33^.

We mapped the RRGs (RUG and RDG) with AAGs. Among the 2399 AAGs names, 397 and 433 (in total 830) re-occur in the lists of RUG and RDG, respectively. Thus, the overlap with AAGs is nearly 35%. I.e., TORC1 controls about one third of the AAGs. Table 1 shows how many AAGs are up- and down-regulated by rapamycin treatment for each of the 15 categories with regard to CLS/RLS increase/decrease. Notably, the order of magnitude for the respective numbers of RUG and RDG is the same for all 15 studied subgroups.

Since rapamycin increases the lifespan by an inhibitory effect on TORC1 activity, it appears most interesting to focus on AAGs with a deletion phenotype of increased CLS/RLS and being downregulated by rapamycin application. A manual analysis of these gene lists reveals that, among the uncharacterized or severely understudied AAGs, there is a single one known to increase both CLS and RLS upon deletion and being downregulated by rapamycin treatment. This gene is *YBR238C*.

### Uncharacterized genes mapping unravels the role of *YBR238C* in cellular aging

Not much is known about *YBR238C* besides its effect on lifespan (to increase CLS and RLS), its mitochondrial localization ^34^, its transcriptional up-regulation by TORC1 and the existence of the paralogue *RMD9*. The encoded protein has 731 amino acid residues. Sequence architecture analysis with the ANNOTATOR ^35^ reveals an intrinsically unstructured region over the first ca. 130 residues (first, a long polar but uncharged run followed by a histidine/asparagine-rich region beginning with position 83) and a pentatricopetide repeat region (residues 130-675, for example, due to a HHpred ^36^ hit to structure 7A9X chain A with E-value 2.e-56). Given the sequence homology, we hypothesize that the protein encoded by *YBR238C* is involved in RNA binding as its paralogue *RMD9* with a similar globular segment^37^.

To note, *YBR238C* carries the conflicting annotations in SGD for increased and decreased RLS upon deletion. Unfortunately, there is not a single direct report about *YBR238C* listed in the scientific literature at the time of writing. There are a few genome-wide deletion strain studies that identified *YBR238C* as one of the gene that increases CLS ^38^ and RLS ^7,39–41^. However, the SGD database wrongly documented one of the latter studies as evidence for a decreased RLS phenotype of *YBR238C* ^41^. Thus, this examination allows us to claim *YBR238C* as the only uncharacterized rapamycin downregulated gene causing CLS and RLS increase upon deletion.

First, we confirmed that *YBR238C* is indeed a rapamycin response gene by qRT-PCR expression analysis in both yeast backgrounds BY4743 and CEN.PK (Figures 1A and 1B). Given that only a single genome-wide deletion strain study identified *YBR238C* as a gene that enhances CLS ^38^, we further tested the role of *YBR238C* in CLS of yeast. CLS was analysed in BY4743 and CEN.PK strains using three different outgrowth survival methods (see detail in methods section). Cell survival of wild type and *ybr238c*Δ strains was analysed at various age time points. We found higher CLS of *ybr238c*Δ cells compared to wild type cells (Figures 1C-1F). Together, these results confirmed that rapamycin inhibits the expression of *YBR238C,* and deletion of this gene indeed increases the cellular lifespan.

**Figure 1.**
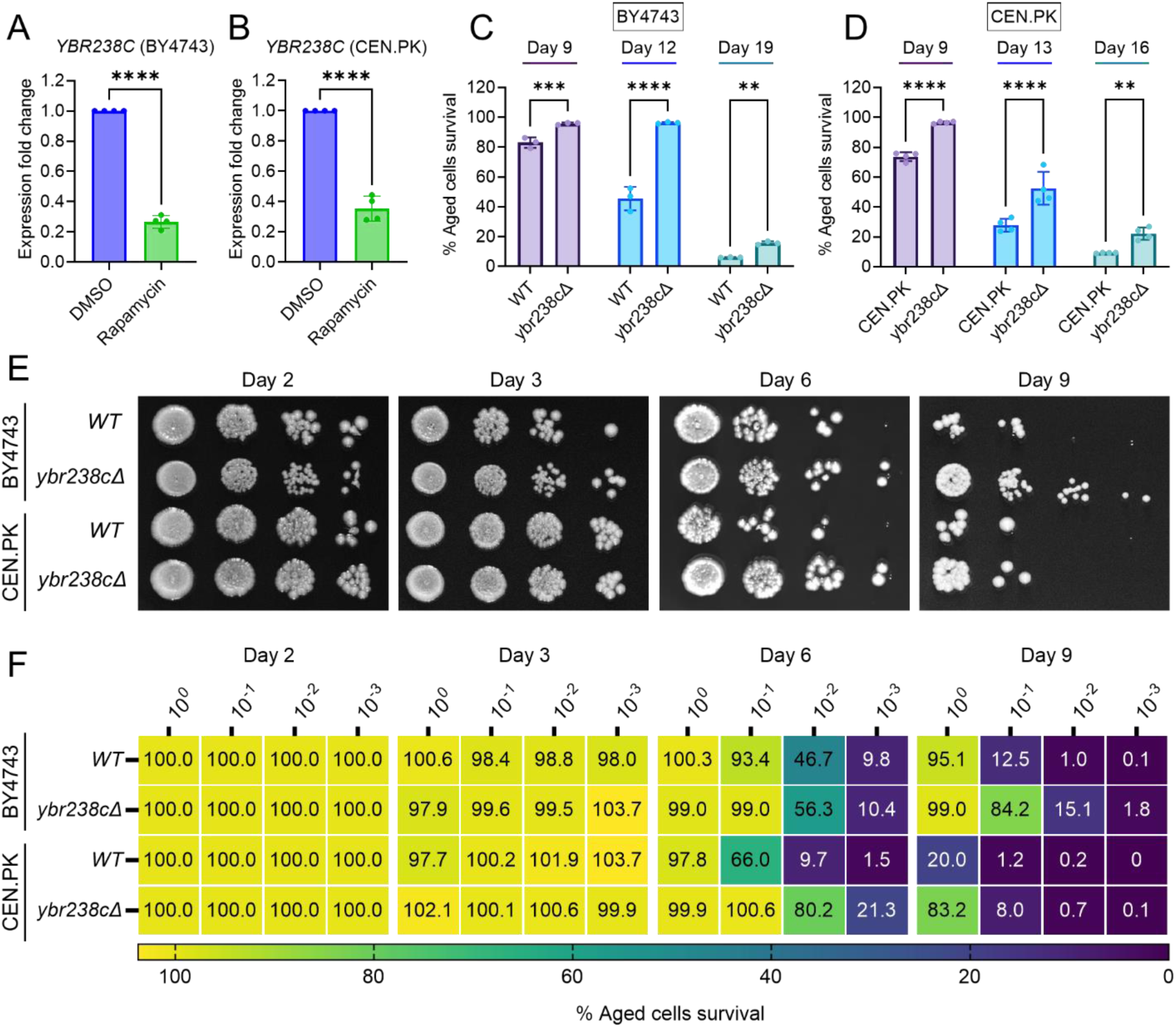
*YBR238C* deletion increases the cellular lifespan. (A and B) Expression analysis of *YBR238C* gene by qRT-PCR in yeast *Saccharomyces cerevisiae* genetic backgrounds BY4743 (A) and CEN.PK113-7D (B). qRT-PCR was performed using RNA extracted from logarithmic-phase cultures grown in synthetic defined medium supplemented with auxotrophic amino acids for BY4743 (A) and only the synthetic defined medium for CEN.PK113-7D (B). Data are represented as means ± SD (n=4). ****P < 0.0001 based on two-sided Student’s t-test. (C and D) Chronological lifespan (CLS) of the wild type and *ybr238c*Δ mutant was assessed in synthetic defined medium supplemented with auxotrophic amino acids for BY4743 (C) and only the synthetic defined medium for CEN.PK113-7D (D) strains in 96-well plate. Aged cells survival was measured relative to the outgrowth of day 2. Data are represented as means ± SD (n=3) (C) and (n=4) (D). **P < 0.01, ***P < 0.001, and ****P < 0.0001 based on two-way ANOVA followed by Šídák’s multiple comparisons test. (E and F) CLS of the wild type and *ybr238c*Δ mutant was performed in synthetic defined medium supplemented with auxotrophic amino acids for BY4743 and only the synthetic defined medium for CEN.PK113-7D strains using flasks. Outgrowth was performed for ten-fold serial diluted aged cells onto the YPD agar plate (E) and YPD medium in the 96-well plate (F). The serial outgrowth of aged cells on agar medium was imaged (E) and quantified the survival relative to outgrowth of day 2 (F). A representative of two experiments for (E) and (F) is shown.

### Transcriptomics analysis reveals the longevity gene expression signatures of *ybr238c*Δ mutants

The transcriptome of the long-lived *ybr238c*Δ mutant was compared with that of the wild type (Figure S3A). Applying standard significance criteria, we found 326 genes up- and 61 genes down-regulated in the *ybr238c*Δ mutant compared to wild type (Figure S3B; Additional File 3). Thus, the transcriptome of the *ybr238c*Δ mutant is very distinct from that of the wild type.

Notably, we see several major metabolic changes. For the *ybr238c*Δ mutant, genes in mitochondrial metabolic processes such as oxidative phosphorylation and aerobic respiration show predominant enrichment (Figures 2A and 2B; Additional File 3). Since *YBR238C* expression is regulated by TORC1 (Figures 1A and 1B), it is not surprising that we observe similarities in the profiles of upregulated DEGs for the *ybr238c*Δ mutant and for the case of treating the wild type with rapamycin (Figures S3C and S3D).

**Figure 2.**
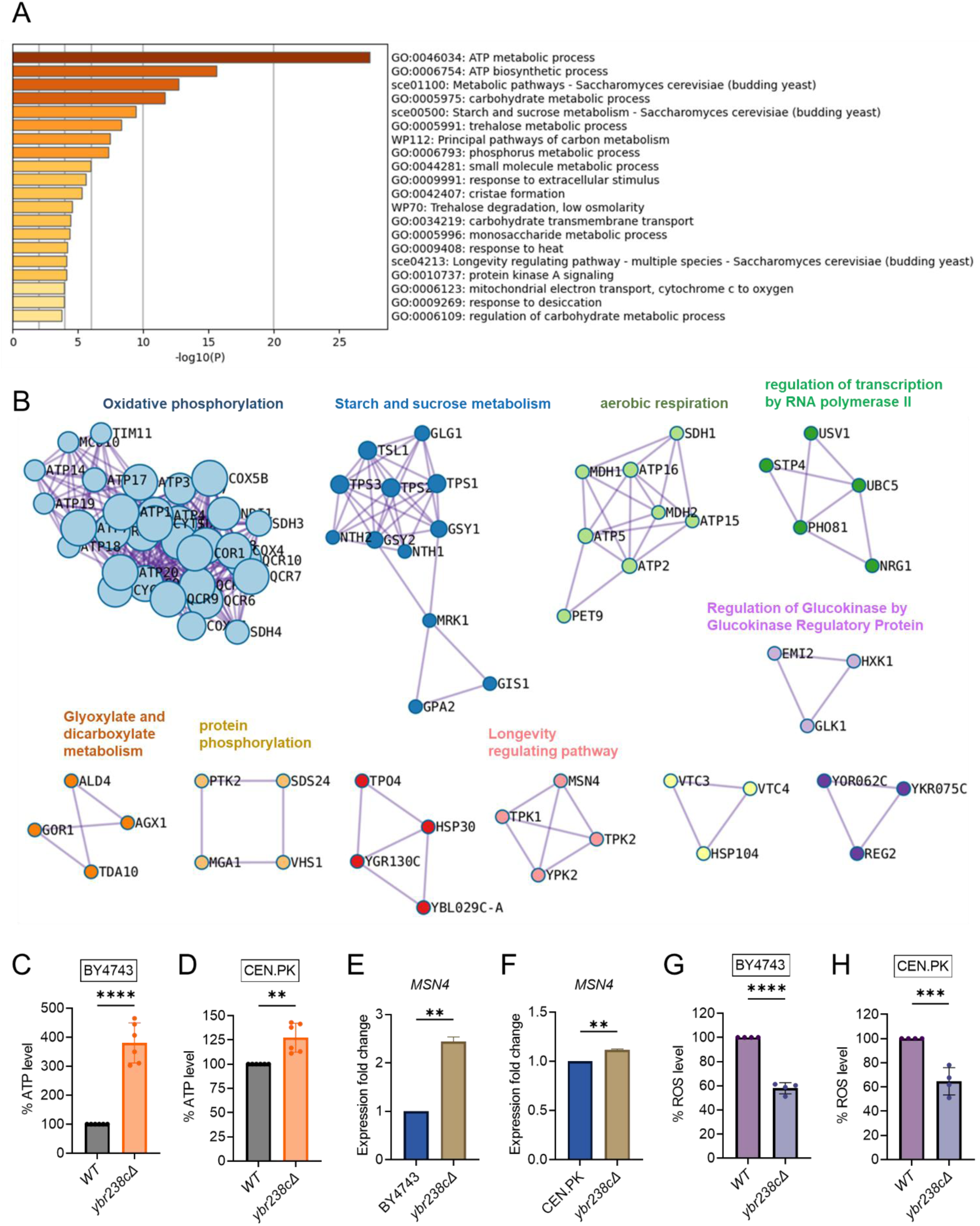
Longevity signatures of *ybr238c*Δ mutant. (A and B) Functional enrichment analysis of upregulated genes in yeast *Saccharomyces cerevisiae ybr238c*Δ mutant compared to wild type (BY4743). Bar plots showing the enriched biological process (A) and identified MCODE complexes based on the top-three ontology enriched terms (B). Full set of genes for functional enrichment analysis (A) and the MCODE complexes (B) are available in Additional File 3. (C and D) ATP analysis of wild type and *ybr238c*Δ mutant for BY4743 (C) and CEN.PK113-7D (D). Quantification was performed using ATP extracted from 72-hour stationary cultures grown in synthetic defined medium supplemented with auxotrophic amino acids for BY4743 and only the synthetic defined medium for CEN.PK113-7D strains. Data are represented as means ± SD (n=6). **P < 0.01, and ****P < 0.0001 based on two-sided Student’s t-test. (E and F) Expression analysis of *MSN4* gene by qRT-PCR in yeast *Saccharomyces cerevisiae* genetic backgrounds BY4743 (E) and CEN.PK113-7D (F). qRT-PCR was performed using RNA extracted from logarithmic-phase cultures grown in synthetic defined medium supplemented with auxotrophic amino acids for BY4743 and only the synthetic defined medium for CEN.PK113-7D strains. Data are represented as means ± SD (n=2). **P < 0.01 based on two-sided Student’s t-test. (G and H) ROS analysis of wild type and *ybr238c*Δ mutant for BY4743 (G) and CEN.PK113-7D (H). Quantification was performed for logarithmic-phase cultures grown in synthetic defined medium supplemented with auxotrophic amino acids for BY4743 and only the synthetic defined medium for CEN.PK113-7D strains. Data are represented as means ± SD (n=4). ***P < 0.001, and ****P < 0.0001 based on two-sided Student’s t-test.

Next, we experimentally tested whether the transcriptome longevity signatures is associated with enhanced mitochondrial metabolism, whether the cellular energy level has gone up and cellular stress responses are induced with a switch to oxidative metabolism ^42,43^. Indeed, our metabolic analysis revealed an increased ATP level in *ybr238c*Δ mutants compared to wild type cells (Figures 2C and 2D; S3E). ATP generation through the oxidative phosphorylation (OXPHOS) system is regulated by mitochondrial DNA (mtDNA) ^44^. We observed a higher mtDNA copy number in the *ybr238c*Δ mutant compared to wild type cells (Figure S3F). Hence, the *ybr238c*Δ mutation rewires the cellular metabolism to promote resource-saving ways of energy production as the up-regulated expression of OXPHOS machinery subunits truly boosts ATP synthesis in *ybr238c*Δ mutant cells.

The enhanced mitochondrial function in *ybr238c*Δ mutants does also improve the protection against reactive oxygen species (ROS). Among the 150 TFs that control the upregulated DEG of the *ybr238c*Δ mutant (Additional File 3), 13 TFs are significantly overrepresented within the upregulated DEGs (Additional File 3). Importantly, we found that the stress response controlling transcription factor *MSN4* is upregulated in *ybr238c*Δ mutants (Figures 2E and 2F; S4A-S4B) ^15,45^. Concomitantly, we found less ROS level in *ybr238c*Δ mutant compared to wild type cells (Figures 2G and 2H; S3G and S3H) and is resistant to H_2_O_2_ induced oxidative stress toxicity (Figure S3I).

Notably, the transcription factor regulated activation of stress response pathways (thioredoxins, molecular chaperones, etc.) ^46,47^ and the switch from fermentation to respiration are associated with delayed cellular aging ^42,43,48,49^. Our results show that the *ybr238c*Δ mutation triggers oxidative metabolism and stress protective machineries activation and, as a result increases CLS. Collectively, our findings reveal that *YBR238C* is a TORC1 regulated gene involved in mitochondrial function coupled with cellular aging. Therefore, we refer to *YBR238C* as AAG1: <u>A</u>ging <u>A</u>ssociated <u>G</u>ene 1.

### *YBR238C* negatively regulates mitochondrial function via *HAP4*-dependent and -independent mechanisms

*HAP4* is a transcription factor that controls the expression of mitochondrial electron transport chain components including OXPHOS genes ^45^. *HAP4* activity has been shown to increase lifespan by enhancing the mitochondrial respiration in cells ^45^. Intriguingly, *HAP4* is among the 13 TFs overrepresented among the upregulated DEGs in *ybr238c*Δ mutant and control mitochondrial genes (Figure 3A; Additional File 3). We validated the upregulation of *HAP4* gene expression in the *ybr238c*Δ mutant under both yeast backgrounds BY4743 and CEN.PK (Figures 3B and 3C). Consistent with these findings, transcription profile of mitochondrial genes in the *ybr238c*Δ mutant is opposite to that of the *hap4*Δ mutant (Figure 3D). For example, ETC complexes I – V genes’ expression is increased in *ybr238c*Δ, however, it is decreased by *HAP4* deletion (Figure 3D).

**Figure 3.**
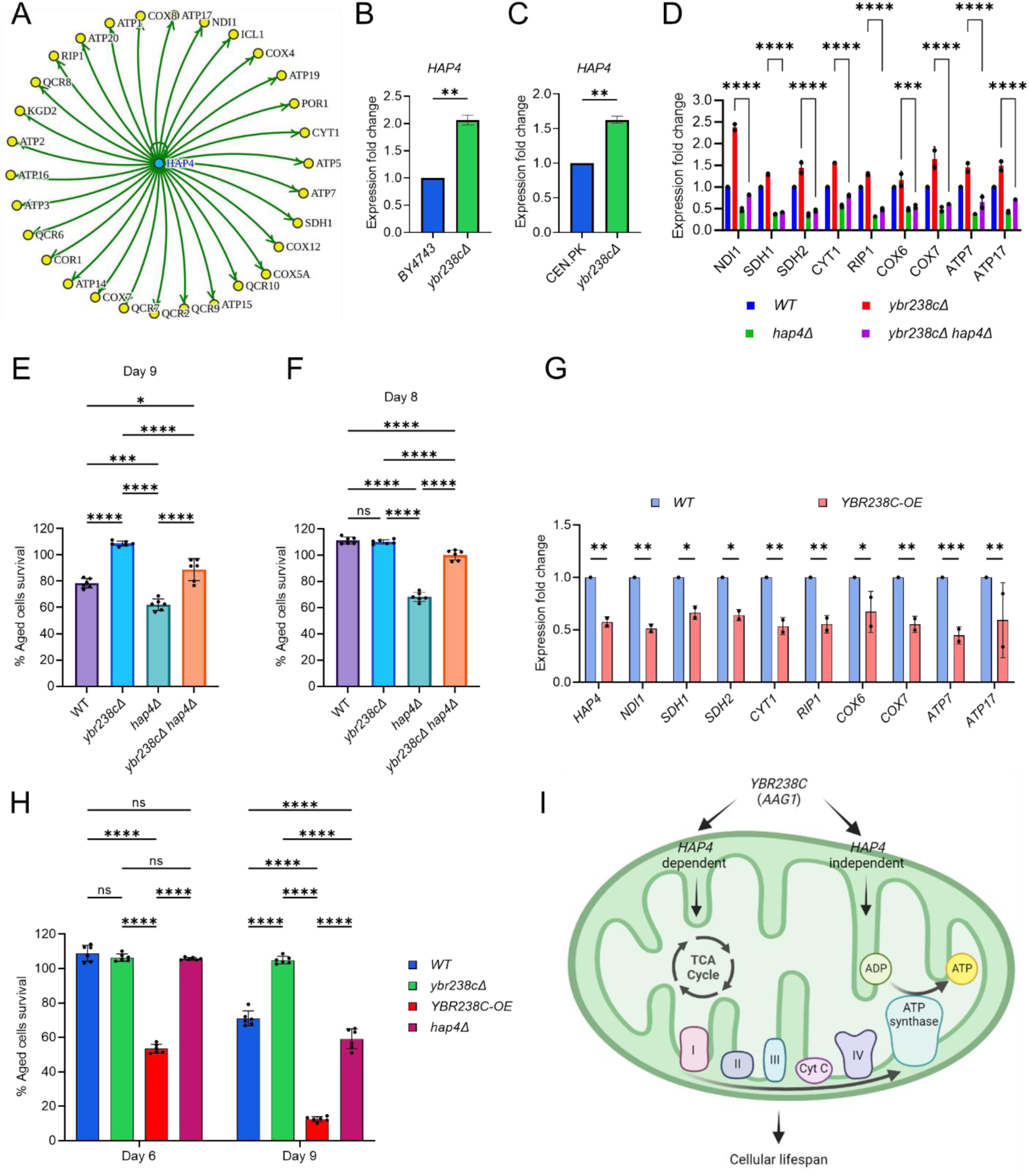
*YBR238C* affects cellular aging via *HAP4*-dependent and independent mechanisms. (A) The *HAP4* regulon within the upregulated DEGs of *ybr238c*Δ mutant. See also Additional File 3. (B and C) Expression analysis of *HAP4* gene by qRT-PCR in yeast *Saccharomyces cerevisiae* genetic backgrounds BY4743 (B) and CEN.PK113-7D (C). qRT-PCR was performed using RNA extracted from logarithmic-phase cultures grown in synthetic defined medium supplemented with auxotrophic amino acids for BY4743 and only the synthetic defined medium for CEN.PK113-7D strains. Data are represented as means ± SD (n=2). **P < 0.01 based on two-sided Student’s t-test. (D) Expression analysis of mitochondrial ETC genes by qRT-PCR in wild type, *ybr238c*Δ, *hap4*Δ, and *ybr238cΔ hap4*Δ, strains of yeast *Saccharomyces cerevisiae* genetic background CEN.PK113-7D. qRT-PCR was performed using RNA extracted from logarithmic-phase cultures grown in synthetic defined medium. Data are represented as means ± SD (n=2). Comparing data between *ybr238c*Δ and *ybr238cΔ hap4*Δ. ***P < 0.001, and ****P < 0.0001 based on two-way ANOVA followed by Šídák’s multiple comparisons test. (E, F and H) Chronological lifespan (CLS) of yeast *Saccharomyces cerevisiae* genetic background CEN.PK113-7D strains was performed in synthetic defined medium using 96-well plate. Aged cells survival was measured relative to the outgrowth of day 2. Data are represented as means ± SD (n=6). *P < 0.05, ***P < 0.001, ****P < 0.0001 and ns: non-significant. Ordinary one-way ANOVA followed by Tukey’s multiple comparisons test (E and F). Two-way ANOVA followed by Šídák’s multiple comparisons test (H). (G) Expression analysis of *HAP4* and ETC genes by qRT-PCR in wild type and *YBR238C* overexpression (*YBR238C-OE*) strains of yeast *Saccharomyces cerevisiae* genetic background CEN.PK113-7D (C). qRT-PCR was performed using RNA extracted from logarithmic-phase cultures grown in synthetic defined medium. Data are represented as means ± SD (n=2). *P < 0.05, **P < 0.01, and ***P < 0.001 based on Two-way ANOVA followed by Šídák’s multiple comparisons test. (I) Model representing regulation of cellular aging by *YBR238C* via *HAP4*-dependent and independent mechanisms.

We found that deletion of *HAP4* moderately decreases CLS at the background of the *ybr238c*Δ mutant (Figure 3E). To confirm that the decrease in lifespan is through the *HAP4* pathway, we examined the expression of mitochondrial respiratory genes. We found that *HAP4* deletion significantly decrease the ETC complex I – V genes’ expression in *ybr238c*Δ mutant (Figure 3D). To note, *HAP4* deletion at the wild-type background decreases the cellular lifespan even more (Figure 3E), which is consistent with more dramatic reduction of ETC complexes I – V genes’ expression (Figure 3D). Taken together these data suggests that *YBR238C* negatively regulates the *HAP4* activity. *HAP4*-upregulated, increased mitochondrial function contributes to the prolonged lifespan of *ybr238c*Δ cells.

*YBR238C* gene deletion rescues some loss of lifespan of *hap4*Δ cells (Figure 3F). The effect is especially pronounced until day 8 when wild-type cells are essentially 100% surviving. Yet, complete epistasis of phenotypes is not achieved. This observation parallels the above findings that *HAP4* deletion in *ybr238c*Δ mutant does not fully recover the mitochondrial ETC complexes I – V genes’ expression and CLS at day 9 compared to the wild type background (Figures 3D and 3E). Together, these results indicate that *YBR238C* affects cellular lifespan via *HAP4*-dependent and -independent mechanisms.

To confirm the existence of *HAP4*-independent mechanisms, we examined the lifespan and transcriptome under conditions of *YBR238C* overexpression (*YBR238C-OE*). As expected, *YBR238C-OE* decreases the expression of *HAP4*, of mitochondrial ETC complexes I – V genes (Figure 3G) as well as the CLS (Figure 3H). Strikingly, the lifespan of *YBR238C-OE* cells was shorter than for *hap4*Δ cells (Figure 3H). Thus, a *HAP4*-independent mechanism does exist through which *YBR238C* also affects cellular aging (Figure 3I).

### The *YBR238C* paralogue *RMD9* deletion decreases the lifespan of cells

In yeast *S. cerevisiae*, *YBR238C* has a paralogue *RMD9* that shares ∼45% amino acid identity ^50^. Given that *YBR238C* activity is tightly coupled with the lifespan, we investigated the role of its paralogue *RMD9* in cellular aging. Surprisingly, we found that *RMD9* deletion has an effect opposite to *YBR238C* deletion and it shortens CLS (Figures 4A-4C; S5A-S5C).

**Figure 4.**
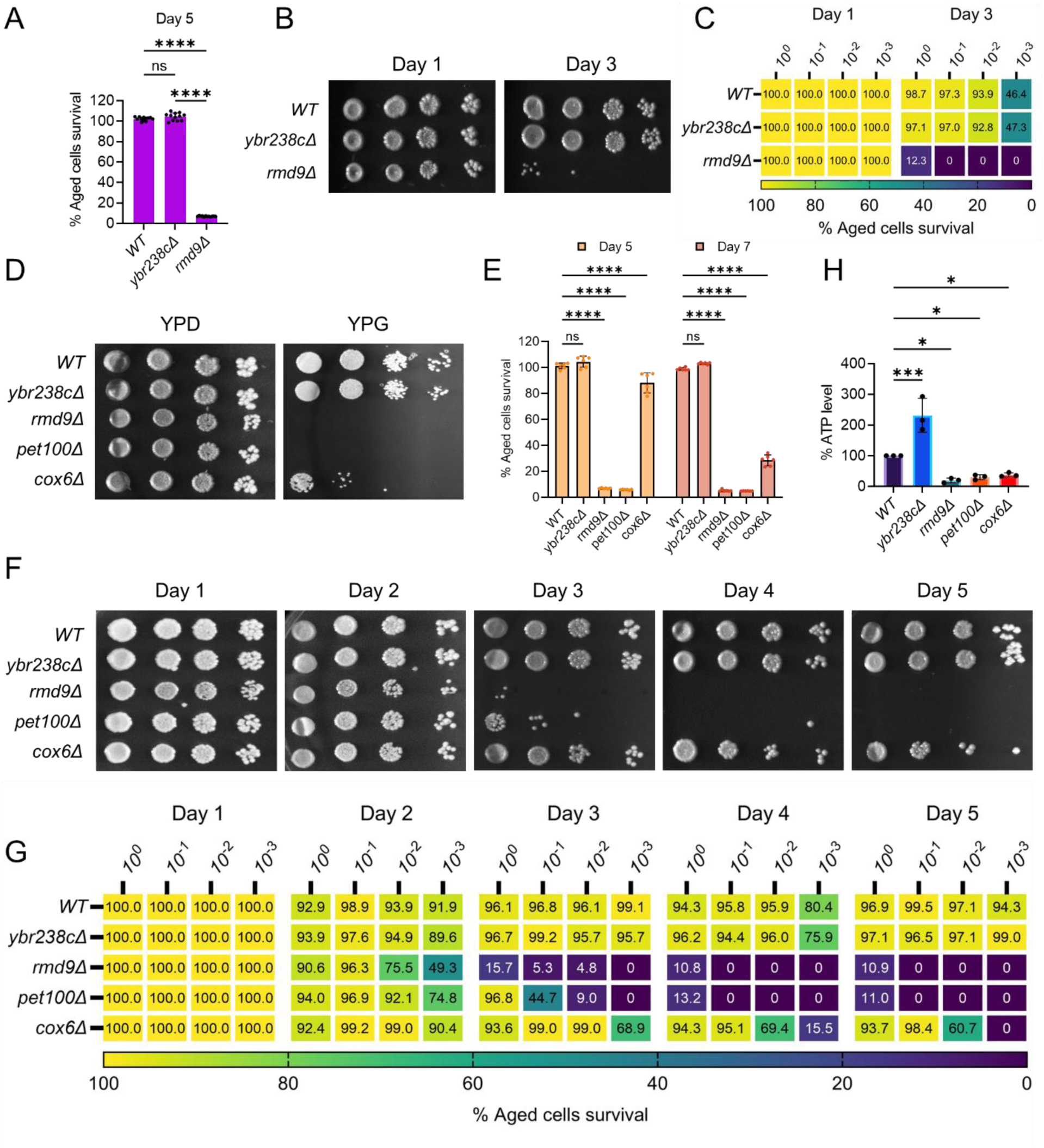
*RMD9* deletion decreases the cellular lifespan. (A and E) Chronological lifespan (CLS) of yeast *Saccharomyces cerevisiae* genetic background CEN.PK113-7D strains was performed in synthetic defined medium using 96-well plate. Aged cells survival was measured relative to the outgrowth of day 2. Data are represented as means ± SD (n=12) (A) and (n=6) (B). ****P < 0.0001 and ns: non-significant. Ordinary one-way ANOVA followed by Tukey’s multiple comparisons test (A). Two-way ANOVA followed by Dunnett’s multiple comparisons test (E). (B, C, F and G) CLS of yeast strains was performed in synthetic defined medium using flask. Outgrowth was performed for ten-fold serial diluted aged cells onto the YPD agar plate (B and F) and YPD medium in the 96-well plate (C and G). The serial outgrowth of aged cells on agar medium was imaged (B and F) and quantified the survival relative to outgrowth of day 1 (C and G). A representative of two experiments for B, C, F and G is shown. (D) Serial dilution growth assays of the yeast strains on fermentative glucose medium (YPD) and respiratory glycerol medium (YPG). (H) ATP level in yeast strains. Quantification was performed using ATP extracted from 72-hour stationary cultures grown in synthetic defined medium. Data are represented as means ± SD (n=3). *P < 0.05, and ***P < 0.001 based on ordinary one-way ANOVA followed by Dunnett’s multiple comparisons test. See also Figure S5.

*RMD9* was initially isolated during a search for genes required for meiotic nuclear division ^51^. Subsequently, its role was uncovered in controlling the expression of mitochondrial metabolism genes by stabilizing and processing mitochondrial mRNAs ^37,50,52^. We asked whether *RMD9* deletion induces mitochondrial dysfunction that, therefore, causes accelerated cellular aging. We first tested the mitochondrial activity by allowing mutant cells to grow under respiratory conditions ^53–55^. We found that *rmd9*Δ mutants could perform on glucose medium, but they failed to grow on glycerol as a carbon source (known to require functional mitochondria; Figures 4D; S5D). Our results are in line with previously observed dysfunctional mitochondria in *rmd9*Δ mutants ^37,50^.

In control experiments, we found a similar growth defect phenotype on glycerol medium for cells deficient in *PET100* and *COX6* mitochondrial genes (Figures 4D; S5D) ^55^. CLS of *pet100*Δ and *cox6*Δ mutants were reduced compared to wild type (Figures 4E-4G; S5E-S5G). We also found lowered ATP- and higher ROS-levels in *rmd9Δ, pet100Δ, cox6*Δ mutants compared to wild type cells (Figures 4H and S5H).

In contrast, long lived *ybr238c*Δ mutants efficiently grow on respiratory medium with high ATP- and low ROS-levels (Figures 4D and 4H; S5D and S5H). So far, the results indicate that deletion of *YBR238C* potentiates the mitochondrial function that, in turn, leads to CLS increase. Thus, *YBR238C* and its paralogue *RMD9* antagonistically affect mitochondrial function, CLS and cellular aging.

### *YBR238C* affects the cellular lifespan via an *RMD9*-dependent mechanism

Previously, *YBR238C* deletion was shown to increase the CLS via *HAP4*-dependent and -independent mechanisms (Figure 3) and to rescue some loss of CLS of the *hap4*Δ mutant by a *HAP4*-independent mechanism (Figures 3E and 3F). Here, we ask whether *YBR238C* deletion suppresses the shortened lifespan of *rmd9*Δ mutants. We examined the lifespan of double deletion *rmd9Δ ybr238c*Δ mutant and compared it with *rmd9*Δ. We found that the deletion of *YBR238C* largely failed to recover the lifespan of *rmd9*Δ mutant to wild type cells; however, it partially prevented their early cell death (Figures 5A-5C). These results show that *YBR238C* deletion increases the cellular lifespan via *RMD9*-dependent mechanisms. Also, we found that the *YBR238C* deletion results in increased ATP- and reduced ROS-levels in the *rmd9*Δ mutant (Figures 5D and 5E).

**Figure 5.**
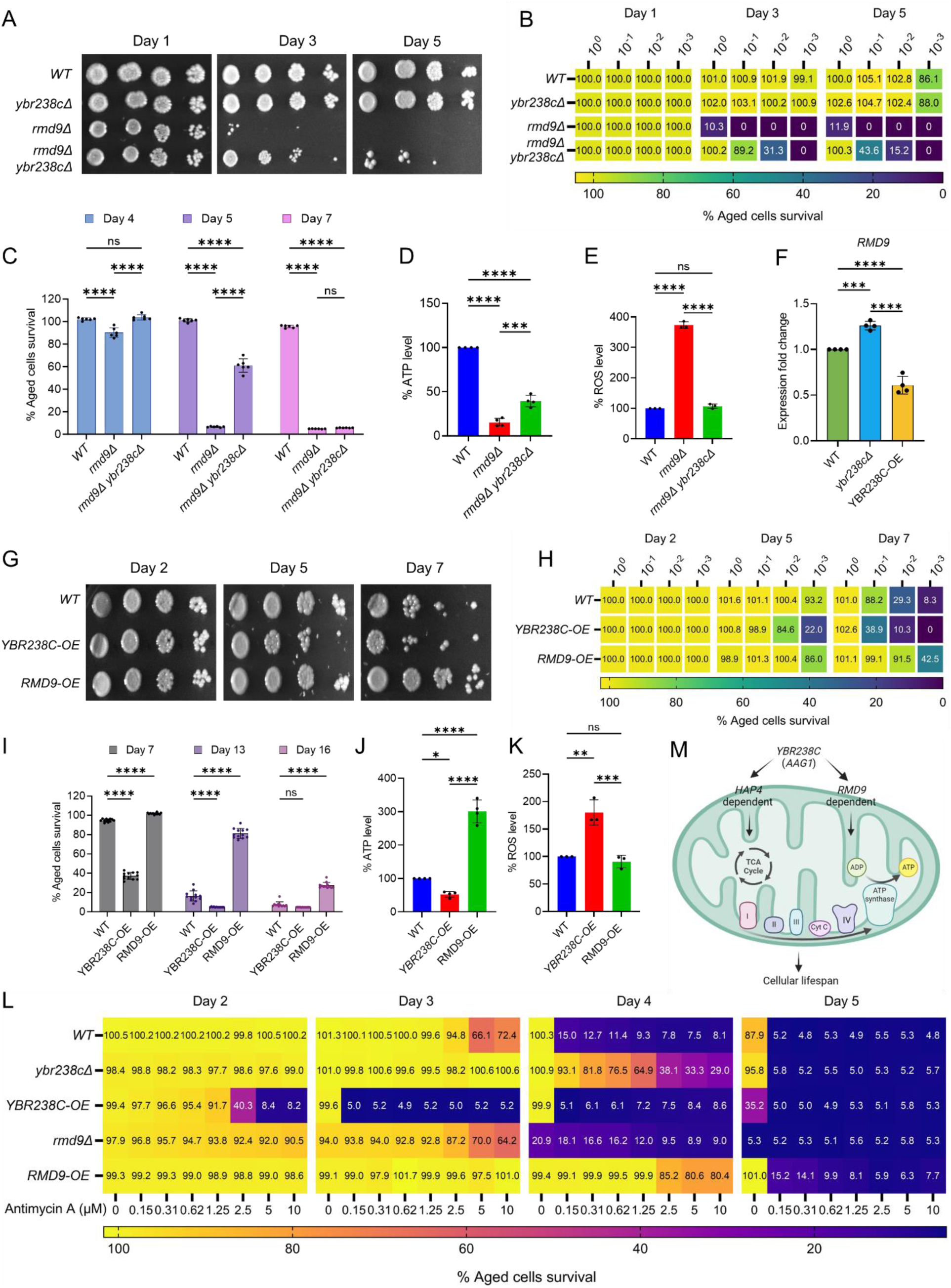
*YBR238C* affects cellular aging via *RMD9*-dependent mechanism. (A, B, G and H) Chronological lifespan (CLS) of yeast *Saccharomyces cerevisiae* genetic background CEN.PK113-7D strains was performed in synthetic defined medium using flask. Outgrowth was performed for ten-fold serial diluted aged cells onto the YPD agar plate (A and G) and YPD medium in the 96-well plate (B and H). The serial outgrowth of aged cells on agar medium was imaged (A and G) and quantified the survival relative to outgrowth of day 2 (B and H). A representative of two experiments for A, B, G and H is shown. (C and I) CLS of yeast strains was performed in synthetic defined medium using 96-well plate. Aged cells survival was measured relative to the outgrowth of day 2. Data are represented as means ± SD (n=6) (C) and (n=12) (I). ****P < 0.0001 based on two-way ANOVA followed by Tukey’s multiple comparisons test (C) and Dunnett’s multiple comparisons test (I). ns: non-significant. (D and J) ATP level in yeast strains. Quantification was performed using ATP extracted from 72-hour stationary cultures grown in synthetic defined medium. Data are represented as means ± SD (n=4). *P < 0.05, ***P < 0.001, and ****P < 0.0001 based on ordinary one-way ANOVA followed by Tukey’s multiple comparisons test. (E and K) ROS level in yeast strains. Quantification was performed for logarithmic-phase cultures grown in synthetic defined medium. Data are represented as means ± SD (n=3). **P < 0.01, ***P < 0.001, and ****P < 0.0001 based on ordinary one-way ANOVA followed by Tukey’s multiple comparisons test. ns: non-significant. (F) Expression analysis of *RMD9* gene by qRT-PCR in yeast strains. qRT-PCR was performed using RNA extracted from logarithmic-phase cultures grown in synthetic defined medium. Data are represented as means ± SD (n=4). ***P < 0.001, and ****P < 0.0001 based on ordinary one-way ANOVA followed by Tukey’s multiple comparisons test. (L) Effect of antimycin A (AMA) treatment on CLS of yeast strain was assessed in synthetic defined medium using 96-well plate. Aged cells survival was measured relative to the outgrowth of day 1. A representative heatmap data of three experiments is shown. (M) Model representing regulation of cellular aging by *YBR238C* via *HAP4* and *RMD9* dependent mechanisms.

Intriguingly, we find *RMD9* expression upregulated in *ybr238c*Δ and downregulated in *YBR238C-OE* cells, respectively (Figure 5F). So, we asked whether *RMD9* expression change is transcriptionally coupled in cellular lifespan phenotypes. Remarkably, *RMD9* overexpression increases the lifespan of cells (Figures 5G-5I) as we would have predicted from the observed changes with the *YBR238C* deletion phenotype.

To know whether this CLS increase is contributed by enhanced mitochondrial function, we quantified the ATP level in wild type, *YBR238C-OE,* and *RMD9-OE* cells. ATP level is higher in *RMD9-OE* cells than wild type cells, a result in line with *RMD9* positively regulating mitochondrial activity (Figure 5J). Consistent with the above findings, *YBR238C* overexpression decreases the ATP level (Figure 5J). Next, we assessed the oxidative stress and found that the ROS level in *RMD9-OE* cells was comparable to wild type. Yet, *YBR238C* overexpression increases the ROS level (Figure 5K).

It can be shown that the effects of *YBR238C* and *RMD9* are at least partially realized via the mitochondrial ETC pathway. Antimycin A (AMA) is an inhibitor of the ETC complex III, decreasing ATP synthesis (Figure S6A) ^56^. Also, AMA treatment reduces the lifespan of cells (Figure S6B), confirming that mitochondrial energy supply is critical to delay cellular aging. Since *YBR238C* and *RMD9* affect the ATP level, we tested the AMA effect on their deletion and overexpression strains. We found that *ybr238c*Δ and *RMD9-OE* cells with their enhanced mitochondrial function phenotype were largely resistant to AMA treatment (Figure 5L). AMA aggravates the cellular aging of mitochondrial defective *YBR238C-OE* cells (Figure 5L). Notably, mitochondrial defective *rmd9*Δ cells were not further affected by AMA treatment (Figure 5L).

Altogether, our findings reveal that *YBR238C* affects CLS and cellular aging via modulating mitochondrial function by mechanisms including *HAP4*- and *RMD9*-dependent pathways (Figure 5M).

### *YBR238C* connects TORC1 signaling with modulating mitochondrial function and cellular aging

So far, we learned that *YBR238C* is regulated by TORC1 (Figures 1A and 1B) and it affects cellular lifespan by modulating mitochondrial function. Here, we wish establish the direct connection between the TORC1 signaling and mitochondrial activity. We treated cells with the TORC1 inhibitor rapamycin and found that, subsequently, the ATP level increases in the cells (Figures 6A; S6C). To test whether TORC1 affects mitochondrial function via *YBR238C*, we analyse the expression profile of mitochondrial ETC genes. As expected, rapamycin supplementation decreased the expression of *YBR238C* (Figures S6D and S6E) but induced the expression of ETC genes (Figure 6B). Remarkably, rapamycin-induced changes in the expression of ETC genes were largely unaffected in *ybr238c*Δ cells and reduced in *YBR238C-OE* cells (Figure 6B). Our results are consistent with the hypothesis that TORC1 regulates the mitochondrial ETC genes via *YBR238C*.

**Figure 6.**
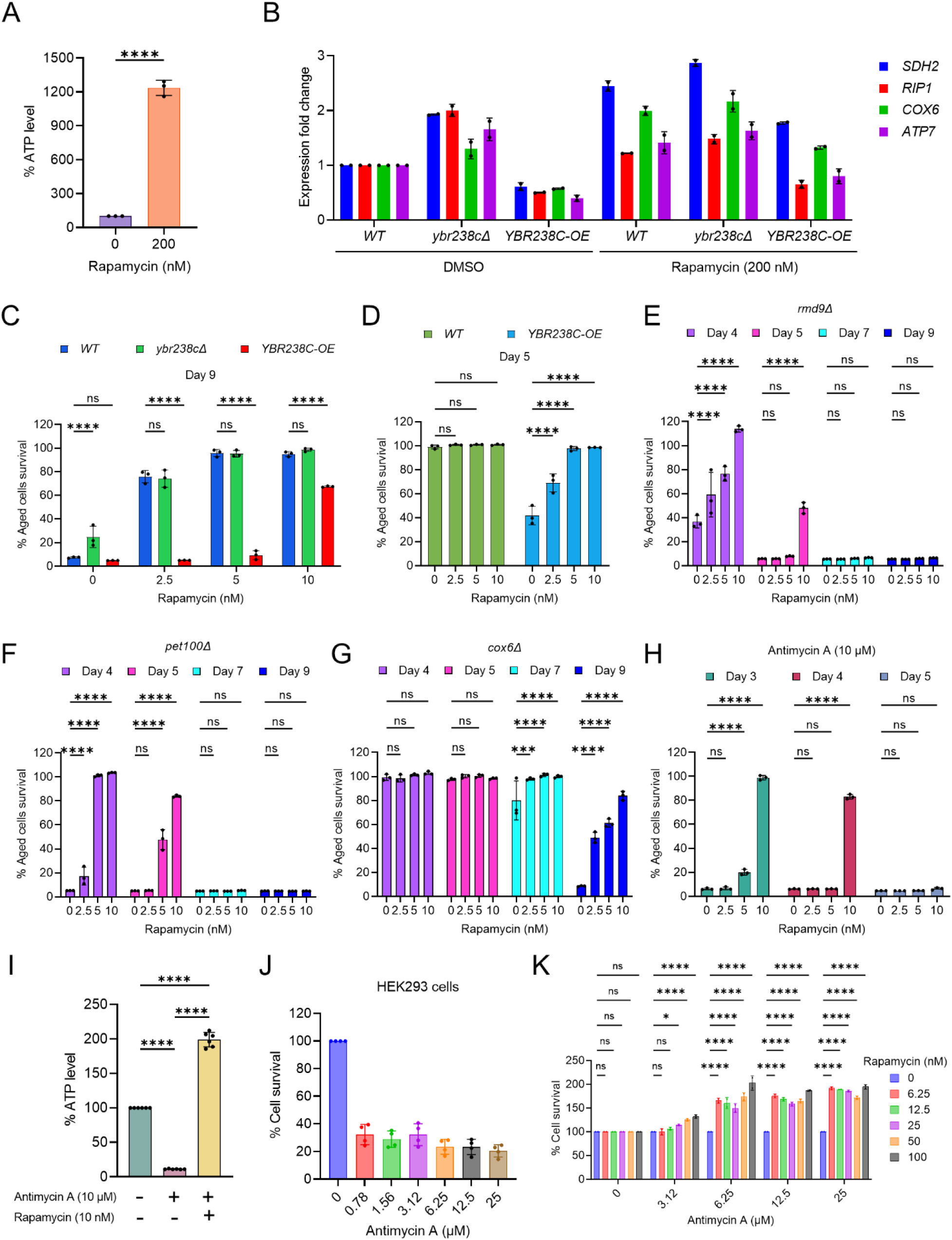
<u>TO</u>RC1-<u>MI</u>tochondria-<u>TO</u>RC1 (TOMITO) signaling axis regulate cellular aging. (A) ATP analysis of logarithmic-phase wild type CEN.PK113-7D yeast cells treated with rapamycin for 1 hour in synthetic defined medium. Data are represented as means ± SD (n=3). ****P < 0.0001 based on ordinary one-way ANOVA followed by Dunnett’s multiple comparisons test. See also Figure S6C. (B) Expression analysis of mitochondrial ETC genes by qRT-PCR. Analysis was performed using RNA extracted from logarithmic-phase wild type CEN.PK113-7D yeast cell treated with rapamycin for 1 hour in synthetic defined medium. Data are represented as means ± SD (n=2). (C-G) Chronological lifespan (CLS) of yeast strains with indicated concentrations of rapamycin was performed in synthetic defined medium using 96-well plate. Aged cells survival was measured relative to the outgrowth of day 2. Data are represented as means ± SD (n=3). ***P < 0.001, and ****P < 0.0001 based on two-way ANOVA followed by Dunnett’s multiple comparisons test. ns: non-significant. (H) CLS of wild type yeast strain with indicated concentrations of rapamycin and antimycin A (10 µM) was performed in synthetic defined medium using 96-well plate. Aged cells survival was measured relative to the outgrowth of day 1. Data are represented as means ± SD (n=3). ****P < 0.0001 based on two-way ANOVA followed by Dunnett’s multiple comparisons test. ns: non-significant. (I) ATP analysis of stationary-phase wild type CEN.PK113-7D yeast cells were incubated with rapamycin (10 nM) and antimycin A (10 µM) in synthetic defined medium. Data are represented as means ± SD (n=6). ****P < 0.0001 based on ordinary one-way ANOVA followed by Tukey’s multiple comparisons test. See also Figure S6H. (J) Determination of human HEK293 cells survival incubated with indicated concentrations of antimycin A for 6 days. Untreated control is considered 100% cell survival. Cells survival was measured relative to the control. Data are represented as means ± SD (n=6). (K) Cell survival analysis of human HEK293 cells incubated with indicated concentrations of antimycin A and rapamycin for 6 days. Antimycin A and rapamycin individual treated concentrations considered 100% cell survival control. Cells survival of combined treated cells was measured relative to the control. Data are represented as means ± SD (n=2). *P < 0.05, ****P < 0.0001, and ns: non-significant based on two-way ANOVA followed by Dunnett’s multiple comparisons test.

The strains *ybr238c*Δ and *YBR238C-OE* are associated with increased and decreased CLS, respectively. So, we ask whether TORC1 influences cellular aging via *YBR238C* by modulating mitochondrial function. We examined the effect of rapamycin supplementation on the lifespan of *ybr238c*Δ and *YBR238C-OE* cells. Strikingly, addition of rapamycin does not further increase CLS of *ybr238c*Δ cells (Figure 6C). Additionally, anti-aging effect of rapamycin is significantly reduced in *YBR238C-OE* cells (Figure 6C). These findings are consistent with the rapamycin effect on transcriptomics profiles of ETC genes in *ybr238c*Δ and *YBR238C-OE* cells (Figure 6B). Taken together, these results reveal that *YBR238C* is a downstream effector of TORC1 signaling, connecting mitochondrial function for the regulation of cellular aging.

### Mitochondrial dysfunction induces the TORC1 activity that causes accelerated cellular aging

We showed that *YBR238C-OE* cells are associated with mitochondrial dysfunction and shortened lifespan (Figure 5). Apparently, the accelerated cellular aging phenotype is primarily due to compromised cellular energy level that affects the lifespan. Genetic (*ybr238c*Δ) and chemical (rapamycin) mediated enhancement of ATP level increases the lifespan of wild type cells validate this conclusion. Notably, the rapamycin supplementation significantly rescued the shortened lifespan of mitochondrial defective *YBR238C-OE* cells (Figures 6D; and S6F). This observation suggests that, as a reaction on sensing mitochondrial dysfunction, TORC1 is activated which, in turn, leads to accelerated cellular aging. Indeed, we found that cellular growth (a proxy for TORC1 activity) of mitochondria dysfunctional *YBR238C-OE* cells was resistant to rapamycin inhibition at concentrations that were effective for wild type cells (Figure S6G).

Further, we asked what the effect of TORC1 activity under enhanced mitochondrial function conditions would be. To test this, we assessed the growth of *ybr238c*Δ cells with rapamycin. We found that the cellular growth of *ybr238c*Δ cells was reduced by rapamycin compared to wild type cells (Figure S6G). Apparently, TORC1 activity is not signalled downstream under these conditions and rapamycin further aggravates this by TORC1 inhibition. This result suggests that *YBR238C* affects TORC1 activity via modulating mitochondrial function.

We found a similar pattern of rapamycin effect on growth of mutant cells with enhanced or damaged mitochondrial function. Growth of *RMD9-OE* cells having enhanced mitochondrial function was sensitive to rapamycin (Figure S6G). Likewise, growth of defective mitochondrial *rmd9*Δ, *pet100*Δ and *cox6*Δ mutants is resistant to rapamycin compared to wild type (Figure S6G). The *YBR238C* deletion reduced the rapamycin growth resistance phenotype of mitochondrial dysfunctional *rmd9*Δ (Figure S6G) and *hap4*Δ cells (Figure S6H), apparently by enabling enhanced mitochondrial function.

Taken together, we see that *YBR238C* plays a junction role in integrating mitochondrial function and TORC1 signaling.

### TORC1 inhibition prevents accelerated cellular aging caused by mitochondrial dysfunction in yeast and human cells

Mitochondrial dysfunction increases the TORC1 activity and, subsequently, causes accelerating cellular aging. We asked whether inhibition of TORC1 activity could prevent accelerating cellular aging even under mitochondrial dysfunction conditions. Previously, we have already shown that rapamycin-mediated TORC1 inhibition partly rescued the shortened lifespan of mitochondrial defective *YBR238C-OE* cells (Figures 6D; and S6F). We examined other mitochondrial dysfunctional conditions to confirm that the suppressive effect of rapamycin is not only specific to *YBR238C-OE*. We tested defective mitochondrial mutants *rmd9Δ, pet100*Δ and *cox6*Δ. In all cases, rapamycin supplementation prevents the accelerated cellular aging (Figures 6E-6G).

Rapamycin supplementation also rescued the AMA-mediated accelerated cellular aging (Figure 6H). We also quantified the ATP level to confirm that the suppressive effect of rapamycin is specific to mitochondrial function. Rapamycin supplementation increases the ATP level of AMA-treated cells (Figures 6I; and S6I), demonstrating that TORC1 inhibition improves mitochondrial function and thus suppresses accelerated cellular aging.

We further verified the connection of TORC1 and mitochondrial function in human cells. First, we examined the effect of AMA on the survival of HEK293 cells. AMA treatment decreases the viability of HEK293 cells (Figure 6J). Cell survival reduction is due to the mitochondrial dysfunction as we see lower ATP levels in AMA treated cells compared to untreated control (Figure S6J). Subsequently, we examined the survival of AMA-treated HEK293 cells with or without rapamycin supplementation. Rapamycin supplementation suppresses AMA-mediated cellular death (Figure 6K). Further rapamycin supplementation increases the ATP level treated cells (Figure S6J).

Our findings convincingly support that TORC1 inhibition suppresses cellular aging associated with mitochondrial dysfunction across species.

## DISCUSSION

Identifying lifespan regulators, genetic networks, and connecting genetic nodes relevant to cellular aging provides functional insight into the complexity of aging biology ^1,6,10^ for the subsequent design of anti-aging interventions. Understanding the mechanism of aging will also require understanding the role of many genes of yet unknown function as *YBR238C* at the beginning of this work. Unfortunately, the group of uncharacterized genes coding proteins of unknown function has received dramatically decreasing attention during the past two decades ^18–20^.

In this study, we first summarized literature and genome database reports about genes affecting aging in the yeast model (mostly observed in gene deletion screens). We found that AAGs (Aging Associated Genes) make up about one-third of the total yeast genome (many of them are functionally uncharacterized) and can be classified into 15 categories for increase/decrease CLS and/or RLS under various conditions.

We compared the set of AAGs with the group of RRGs (Rapamycin Regulated Genes) and found that about one third of the AAGs is TORC1 activity regulated. Among the functionally uncharacterized genes in this overlap set, the gene *YBR238C* stands out as the only one that (1) is downregulated by rapamycin and (2) its deletion increases both CLS ^38^ and RLS ^7,39–41^. Thus, *YBR238C* is important among identified rapamycin-downregulated uncharacterized genes in AAG categories as it seems to rationally connect the rapamycin-induced TORC1 inhibition with the increase of both CLS and RLS.

We observed that the CLS of *ybr238c*Δ cells is higher than for wild type, regardless of various aging experimental conditions tested in this study. Transcriptional analysis of *ybr238c*Δ cells identified a longevity signature including enhanced gene expression related to mitochondrial energy metabolism and stress response genes ^8,45,57–60^. In contrast, *YBR238C-OE* cells display mitochondrial dysfunction leading to decreased lifespan. Together, these results reveal that TORC1 regulates the *YBR238C* expression which is linked to mitochondrial function and cellular lifespan.

The *YBR238C* paralogue *RMD9* has been previously shown to affect mitochondrial function ^37,50^. Surprisingly, we observed an antagonism regarding the CLS phenotype of deletion/overexpression for *YBR238C* and *RMD9*. *YBR238C* overexpression and *RMD9* deletion confer defective mitochondrial function associated with accelerated cellular aging and decrease lifespan of cells. In contrast, *YBR238C* deletion and *RMD9* overexpression enhance the mitochondrial function that increase the cells’ longevity. Consistent with the identified negative regulatory role of *YBR238C* on mitochondrial function, deletion of this gene partially suppressed the accelerated aging of *rmd9*Δ cells and AMA-treated cells. We identified that *YBR238C* influences mitochondrial function and the CLS phenotype with contributions from *HAP4*- and *RMD9*-dependent mechanisms. As (i) one of *RMD9*’s molecular functions has been shown to stabilize certain (mitochondrial) mRNAs and (ii) *YBR238C* has a has a homologous pentatricopeptide repeat region, the two paralogues might differ in the sets of protected mRNAs with opposite outcome on cellular longevity.

While Kaeberlein et al. identified *YBR238C* as a top candidate for increasing replicative lifespan (RLS) in yeast through deletion ^40^, our study builds upon their work by investigating the mechanisms and the connection with TORC1 in chronological lifespan (CLS). Results of genetic interventions (*YBR238C* deletion and overexpression) and rapamycin treatment show that *YBR238C* is a downstream effector by which TORC1 modulates mitochondrial activity. Most importantly, we found that TORC1 inhibition increases the mitochondrial function via *YBR238C* and, thereby, extends cellular lifespan. TORC1 is involved in both CLS and RLS ^15^. Whether the TORC1 - *YBR238C* axis has similar or distinct mechanisms for CLS and RLS will be interesting to identify in the future.

TORC1 is the major controller of cellular metabolism that links environmental cues and signals for cellular growth and homeostasis. TORC1 upregulates anabolic processes such as *de novo* synthesis of proteins, nucleotides, lipids, and downregulates catabolic processes such as inhibition of autophagy ^25,27^. We find that dysfunctional mitochondria lead to TORC1 activation both in yeast and in human cells with accompanying onset of accelerated cellular aging. Our results are in line with a recent report about anabolic pathways enhancement and suppression of catabolic processes in cells with defective mitochondria ^61^. On the other hand, TORC1 activity decreases under enhanced mitochondrial function environment. The cause of the mitochondrial dysfunction (whether due to deletion of a critical gene or pharmacological intervention) is irrelevant in this context. Nevertheless, despite the mitochondrial insufficiency background, inhibition of TORC1 (e.g., with rapamycin) can partially rescue the shortened cellular lifespan (Figures 6D-6H; and S6F).

Our work sheds light onto the questions (i) how mitochondrial dysfunction is linked to accelerated cellular aging and (ii) how TORC1 inhibition suppresses the mitochondrial dysfunction and prevents shortening lifespan. Apparently, mitochondrial dysfunction aberrantly signals to increase the TORC1 activity that leads to accelerated aging in cells. Remarkably, TORC1 inhibition can often suppress the accelerated cellular aging associated with impaired mitochondrial function. Yet, we found that growth of mutant strains with dysfunctional mitochondria can be resistant despite TORC1 inhibition by rapamycin with concentrations effective in wild type cells possibly due to the dramatic increase of TORC1 activity as in *YBR238C-OE*, *rmd9*Δ, *pet100*Δ, *cox6*Δ and *hap4*Δ cells (Figures S6G and S6H).

Our interpretation of the experimental results is supported by two recently published studies: (1) TORC1 activation was observed after mitochondrial ETC dysfunction in an induced pluripotent cell model ^62^. (2) The inhibition of TORC1 delays the progression of brain pathology of mice with ETC complex I NDUFS4-subunit knockout ^63^.

We think that it will be insightful to explore TORC1 activity under various conditions with compromised mitochondrial function including mitochondrial fission and fusion dynamics which is reported to affect in cell viability ^64,65^. Altogether, our findings uncover the central role of the feedback loop between mitochondrial function and TORC1 signaling (<u>TO</u>RC1-<u>MI</u>tochondria-<u>TO</u>RC1 (TOMITO) signaling process). Whereas the effector from TORC1 to mitochondria involves *YBR238C*, the other direction appears executed via metabolite sensing (e.g., α-keto-glutarate, glutamine, etc.) ^66^.

## METHODS

### Data acquisition

The gene lists that modulate cellular lifespan in aging model organism yeast *Saccharomyces cerevisiae* were extracted from database SGD ^21^ and GenAge ^22^ (as of 8^th^ November 2022). The actual gene lists are available in ‘Additional File 1’.

### Yeast strains, growth media and cell culture

The *S. cerevisiae* auxotrophic BY4743 (Euroscarf) and prototrophic CEN.PK113-7D ^67^ strains were used in this study. Deletion strains for BY4743 obtained from yeast homozygous diploid collection. Deletion strains in CEN.PK background was generated using standard PCR-based method ^68^. Yeast strains were revived from frozen glycerol stock on YPD agar (1% Bacto yeast extract, 2% Bacto peptone, 2% glucose and 2.5% Bacto agar) medium for 2-3 days at 30°C. For all experiments, CEN.PK113-7D strains were grown in synthetic defined (SD) medium contain 6.7 g/L yeast nitrogen base with ammonium sulfate without amino acids and 2% glucose. SD medium supplemented with histidine (40 mg/L), leucine (160 mg/L), and uracil (40 mg/L) for auxotrophic BY4743 strains.

### Human cell lines, growth media and cell culture

Human embryonic kidney cell line (HEK293, ATCC) was cultured in high-glucose DMEM supplemented with 10% FBS and 1% Penicillin Streptomycin Solution. All cells cultured in a humidified incubator with 5% CO_2_ at 37°C.

### Chemical treatment to cell culture

Stock solution of rapamycin and antimycin A was prepared in dimethyl sulfoxide (DMSO). The final concentration of DMSO did not exceed 1% in yeast and 0.01% in human cell lines experiments.

### Yeast aging assay

For the chronological lifespan (CLS) experiments, prototrophic CEN.PK113-7D strains were grown in synthetic defined (SD) medium contain 6.7 g/L yeast nitrogen base with ammonium sulfate without amino acids and 2% glucose. SD medium supplemented with histidine (40 mg/L), leucine (160 mg/L), and uracil (40 mg/L) for auxotrophic BY4743 strains. Chronological aging was assessed by determining the lifespan of yeast as described previously with slight modifications ^15,69^. Yeast culture grown in overnight at 30°C with 220 rpm shaking in glass flask was diluted to starting optical density at 600 nm (OD_600_) ∼ 0.2 in fresh medium to initiate the CLS experiment. CLS was performed by outgrowth method utilizing three different approaches: (i) Cells grown and aged in 96-well plates with a total 200 µL culture in SD medium at 30°C. At various age time points yeast stationary culture (2μL) were transferred to a second 96-well plate containing 200 μL YPD medium and incubated for 24 hours at 30°C without shaking. Outgrowth OD_600_ for each age point was measured by the microplate reader; (ii) Cells grown and aged in flask with a total culture volume more than 5 ml SD medium and incubated for 24 hours at 30°C with 220 rpm shaking. At different age time points yeast stationary culture washed and normalized to OD_600_ of 1.0 with YPD medium. Further, normalized yeast cells were serial 10-fold diluted with YPD medium in 96-well plates. 3 μL of diluted culture were spotted onto the YPD agar plate and incubated for 48 hours at 30°C. The outgrowth of aged cells on the YPD agar plate was photographed using the GelDoc imaging system; (iii) The above discussed serial 10-fold diluted yeast stationary culture with YPD medium in 96-well plates incubated for 24 hours at 30°C without shaking. Outgrowth OD_600_ for serial diluted aged cells was measured by the microplate reader.

### RNA extraction

Yeast cells were first mechanically lysed using the manufacturer’s disruption protocol. Total RNA from yeast cells was extracted using Qiagen RNeasy mini kit. ND-1000 UV-visible light spectrophotometer (Nanodrop Technologies) and Bioanalyzer 2100 with the RNA 6000 Nano Lab Chip kit (Agilent) was used to assess the concentration and integrity of RNA.

### RNA sequencing and bioinformatics analysis

RNA sequencing (RNA-Seq) was conducted using NovaSeq PE150. Raw Fastq files were then passed into Fastp v0.23.2 for adapter trimming and low-quality reads removal. Both single end and paired end reads were processed with default parameters with --detect adapter for pe. Raw reads that passed the quality check were then aligned using HiSat 2 v2.2.1 with the index built from *Saccharomyces cerevisiae* R-64-1-1 top-level DNA fasta file obtained from Ensembl for sequencing data with S288C strain background. Library information for all experiments were first checked with RSeQC infer_experiment.py v4.0.0 before the addition of the respective parameters for alignment and counts. The resulting SAM files were then converted and sorted to BAM files using Samtools v1.13. The BAM files were then used to generate feature counts using HTSeq v1.99.2. HTSeq counts from each experiment were then used for downstream Differentially Expressed Genes (DEG) analysis. Counts generated from HTSeq were then used for differential gene expression analysis using EdgeR v3.34.1 quasi likelihood F-test and DESeq2 v1.36.0. Principal Component Analysis (PCA) was conducted via the use of Transcript Per Million normalized counts. Samples that are separated by batches in the PCA were corrected using ComBat-Seq v3.44.0 before PCA was conducted after normalization of the corrected counts. Functional enrichment analysis was performed by metascape tool ^28^.

### qRT-PCR analysis

qRT-PCR (Quantitative Reverse Transcription-Polymerase Chain Reaction) experiments were performed as described previously using QuantiTect Reverse Transcription Kit (Qiagen) and SYBR Fast Universal qPCR Kit (Kapa Biosystems) ^33^. The abundance of each gene was determined relative to the house-keeping transcript *ACT1*.

### ATP analysis

ATP analysis was performed as described previously ^70^. Yeast cells were mixed with the final concentration of 5% trichloroacetic acid (TCA) and then kept on ice for at least 5 min. Cells were washed and resuspended in 10% TCA and lysis was performed with glass beads in a bead beater to extract the ATP. ATP extraction from human HEK293 cells was performed using Triton X-100 lysis buffer. The ATP level was quantified by PhosphoWorks™ Luminometric ATP Assay Kit (AAT Bioquest) and normalized by protein content measured Bio-Rad protein assay kit.

### Mitochondrial DNA copy number analysis

Determination of mitochondrial DNA (mtDNA) copy number was analysed by real time qPCR (Quantitative-Polymerase Chain Reaction). DNA was extrcated using Quick-DNA Midiprep Plus Kit (Zymo Research). qPCR was performed in a final volume of 20 μl containing 20 ng of total DNA using SYBR Fast Universal qPCR Kit (Kapa Biosystems) and analysed using the Quant Studio 6 Flex system (Applied Biosystems). The real-time qPCR conditions were one hold at (95 °C, 180 s), followed by 40 cycles of (95 °C, 1 s) and (60 °C, 20 s) steps. A melting-curve analysis was included in the cycle after amplification to verify PCR specificity and the absence of primer dimers. Relative mtDNA content was determined by qPCR of mitochondrial genes (*ATP6* and *COX3*) and normalized with nuclear specific gene *ACT1*.

### ROS measurement

Cells were washed and resuspended in 1× phosphate buffer saline (PBS, pH 7.4). After that cells were incubated with 40 μM H2DCFDA (Molecular probe) for 30 min at 30°C ^71^. Cells were then washed with PBS and ROS level was measured by fluorescence reading (excitation at 485 nm, emission at 524 nm) by the microplate reader. The fluorescence intensity was normalized with OD_600_.

### Transcription factors analysis

Transcription factors (TFs) enrichment analysis was performed using YEASTRACT ^72,73^. The significant p-value <0.05 considered for regulatory network analysis based on DNA-binding plus expression evidence. The TFs regulatory networks were visualized with a force-directed layout.

### Fermentative and respiratory growth assay

Yeast cells grown in YPD medium were washed and normalized to OD_600_ of 1.0 with water. Further, normalized yeast cells were serial 10-fold diluted with water. 3 μL of diluted culture were spotted onto the agar medium YPD (2% glucose) and YPG (3% glycerol) and incubated for 48 hours at 30°C. The cell growth on the agar plate was photographed using the GelDoc imaging system.

### Growth sensitivity assay

Effect of chemical compounds on cell growth was carried out in 96-well plates. At an OD_600_ of ∼ 0.2 in SD medium 200 µL yeast cells was transferred into the 96-well plate containing serially double-diluted concentrations of compounds. Cells were incubated at 30°C and the growth was measured at OD_600_ by the microplate reader.

### Quantification and statistical analysis

Data analysis of all the experimental results such as mean value, standard deviations, significance, and graphing were performed using GraphPad Prism v.9.3.1 software. The comparison of obtained results were statistically performed using the Student’s t-tests, Ordinary One-way ANOVA and Two-way ANOVA followed by multiples comparison tests. In all the graph plots, P values are shown as *P < 0.05, **P < 0.01, ***P < 0.001, and ****P < 0.0001 were considered significant. ns: non-significant.

## Data availability

Additional File 1, Additional File 2, and Additional File 3 data are deposited.

## Lead contact and materials availability

Further information and requests for resources and reagents should be directed to and will be fulfilled by the Lead Contact, Dr. Mohammad Alfatah.

## SUPPLEMENTARY FIGURE LEGENDS

**Figure S1.**
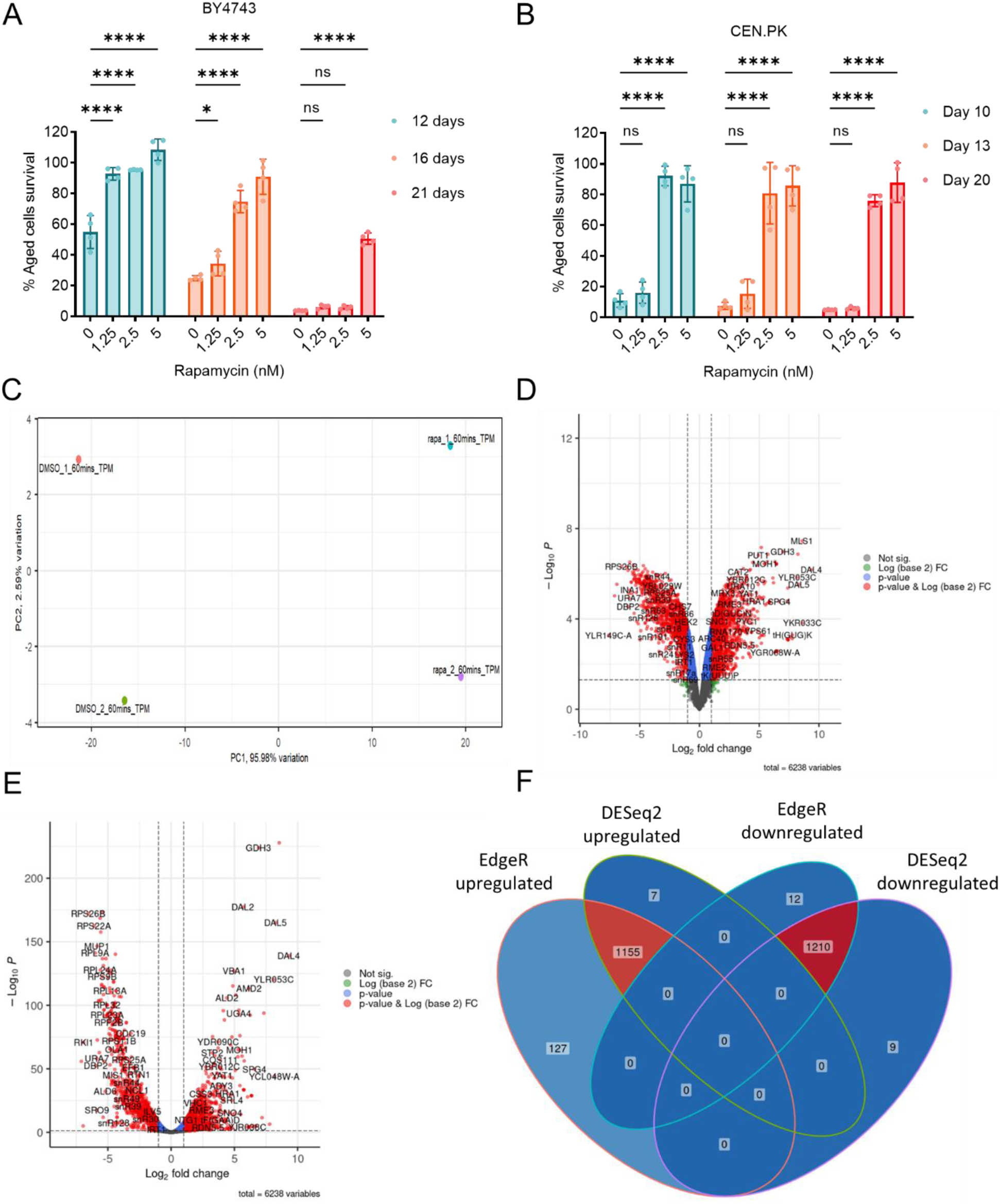
Transcriptomics analysis of rapamycin to identify the TORC1 regulated genes. (A and B) Chronological lifespan (CLS) of the wild type BY4743 (A) and CEN.PK113-7D (B) yeast strains was assessed in the synthetic defined medium supplemented with auxotrophic amino acids for BY4743 (A) and only the synthetic defined medium for CEN.PK113-7D (B) in 96-well plate. Aged cells survival was measured relative to the outgrowth of day 2. Data are represented as means ± SD (n=4). *P < 0.05, and ****P < 0.0001 based on two-way ANOVA followed by Dunnett’s multiple comparisons test. ns: non-significant. (C) Principal component analysis (PCA) of the RNA-seq data from replicates of DMSO and rapamycin (50 nM) treated logarithmic-phase yeast cells (BY4743) for 1 hour in synthetic defined medium supplemented with auxotrophic amino acids. PC1 represents 95.98% of the total variance in the experimental data, and PC2 represents 2.59% of the total variance. (D and E) Volcano plots produced by EdgeR (D) and DESeq2 (E) demonstrate the gene expression pattern and the differentially expressed genes (DEGs) with p-value <0.05, with a fold change of log2(fold change) >1 for upregulated genes and log2(fold change) <−1 for downregulated genes. (F) Overlap of identified DEGs using EdgeR and DESeq2 with the criteria for all two tools that p-value <0.05, with a fold change of log2(fold change) >1 for upregulated genes and log2(fold change) <−1 for downregulated genes. Upregulated and downregulated DEGs overlaps for both EdgeR and DESeq2 were further considered for functional GO enrichment analysis. See also Additional File 2.

**Figure S2.**
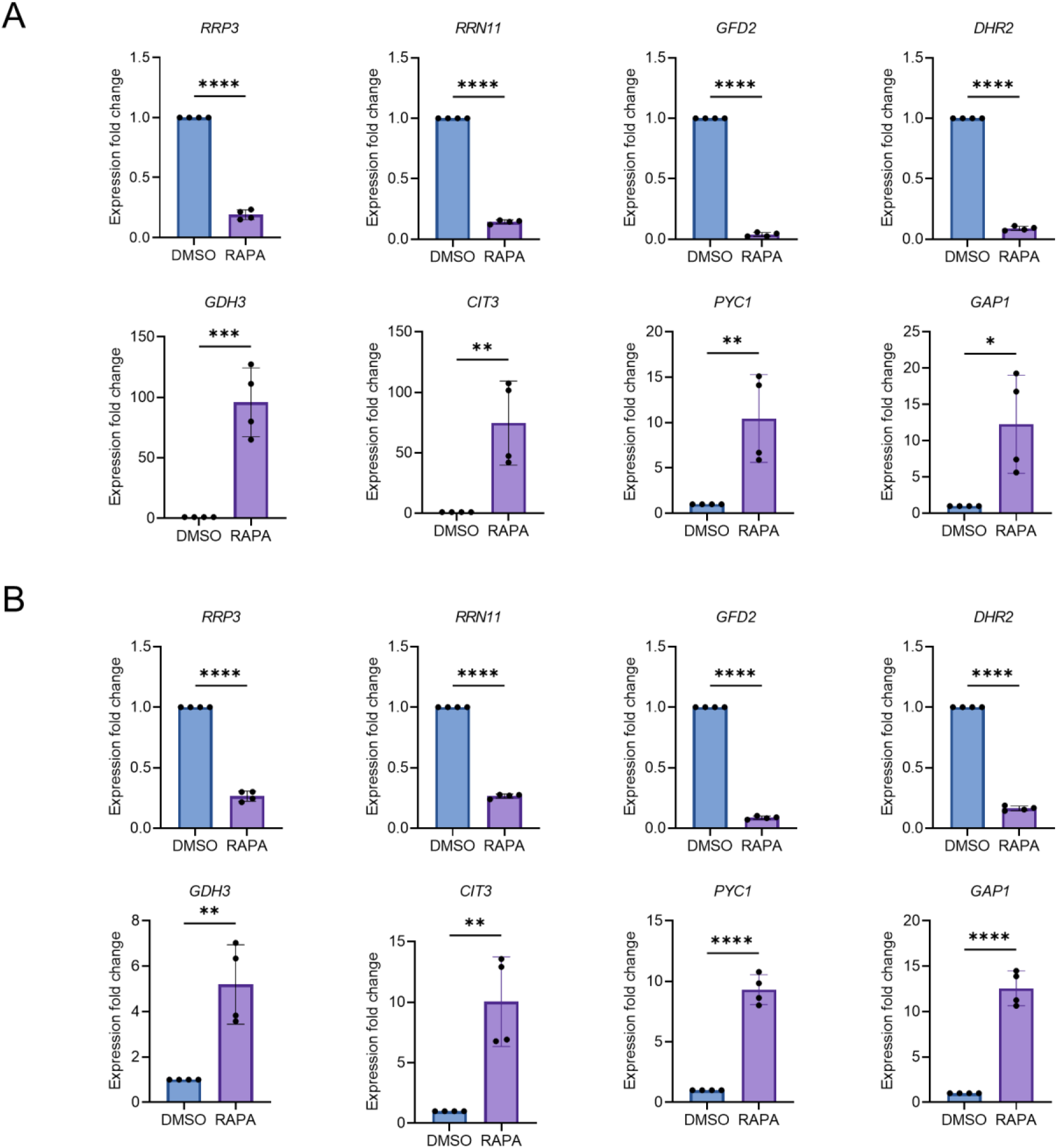
RNA sequencing data validation by qRT-PCR. (A and B) Expression analysis of selected genes of RNA sequencing (RNA-seq) data by qRT-PCR in *Saccharomyces cerevisiae* genetic backgrounds BY4743 (A) and CEN.PK113-7D (B). Logarithmic-phase wild type cells treated with DMSO and rapamycin (50 nM) in synthetic defined medium supplemented with auxotrophic amino acids for BY4743 (A) and only the synthetic defined medium for CEN.PK113-7D (B) strains. Gene expression of rapamycin treated samples were compared with DMSO control. Data are represented as means ± SD (n=4). *P < 0.05, **P < 0.01, ***P < 0.001, and ****P < 0.0001 based on two-sided Student’s t-tests.

**Figure S3.**
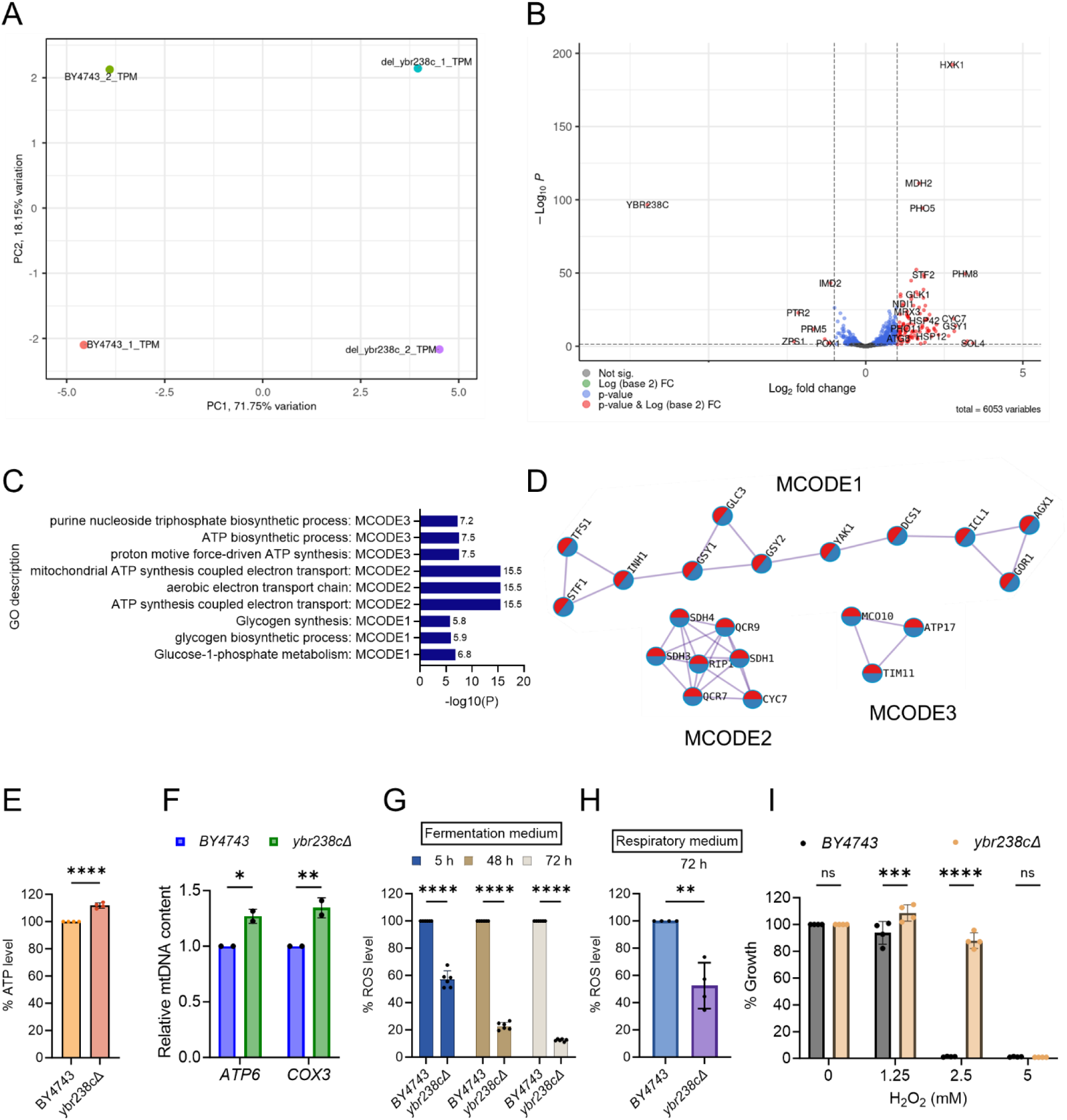
Identification of the role of uncharacterized *YBR238C* gene in cellular aging, Related to Figure 2. (A) Principal component analysis (PCA) of the RNA-seq data from replicates of logarithmic-phase wild type BY4743 and *ybr238c*Δ yeast strains in the synthetic defined medium supplemented with auxotrophic amino acids. PC1 represents 71.75% of the total variance in the experimental data, and PC2 represents 18.15% of the total variance. (B) Volcano plots produced by DESeq2 demonstrate the gene expression pattern and the differentially expressed genes (DEGs) with p-value <0.05, with a fold change of log_2_(fold change) >0.5 for upregulated genes and log_2_(fold change) <−0.5 for downregulated genes. GO enrichment analysis was performed for upregulated DEGs. See also Additional File 3. (C and D) Comparison of functional enrichment analysis for upregulated DEGs of *ybr238c*Δ mutant and rapamycin (50 nM) treated wild type BY4743 cells. Bar plots showing the common enriched biological process (C) and MCODE complexes based on the top-three ontology enriched terms (D). (E) ATP analysis of wild type BY4743 and *ybr238c*Δ mutant strains. Quantification was performed using ATP extracted from logarithmic-phase cultures grown in synthetic defined medium supplemented with auxotrophic amino acids. Data are represented as means ± SD (n=4). ****P < 0.0001 based on two-sided Student’s t-test. (F) Relative mitochondrial DNA (mtDNA) in logarithmic-phase wild type BY4743 and *ybr238c*Δ mutant in synthetic defined medium supplemented with auxotrophic amino acids. Relative mtDNA content was determined by qPCR of mitochondrial genes (*ATP6* and *COX3*) and normalized with nuclear specific gene *ACT1*. Data are represented as means ± SD (n=2). *P < 0.05, and **P < 0.01 based on two-way ANOVA followed by Šídák’s multiple comparisons test. (G and H) ROS analysis of wild type BY4743 and *ybr238c*Δ mutant in synthetic defined (G) fermentative glucose medium, and (H) respiratory glycerol medium, supplemented with auxotrophic amino acids, at indicated time points. (G) Data are represented as means ± SD (n=6). ****P < 0.0001 based on two-way ANOVA followed by Šídák’s multiple comparisons test. (H) Data are represented as means ± SD (n=4). **P < 0.01 based on based on two-sided Student’s t-tests. (I) Growth assay with H_2_O_2_ of wild type BY4743 and *ybr238c*Δ strains in synthetic defined medium supplemented with auxotrophic amino acids. Data are represented as means ± SD (n=4). ***P < 0.001, ****P < 0.0001 and ns: non-significant based on two-way ANOVA followed by Šídák’s multiple comparisons test.

**Figure S4.**
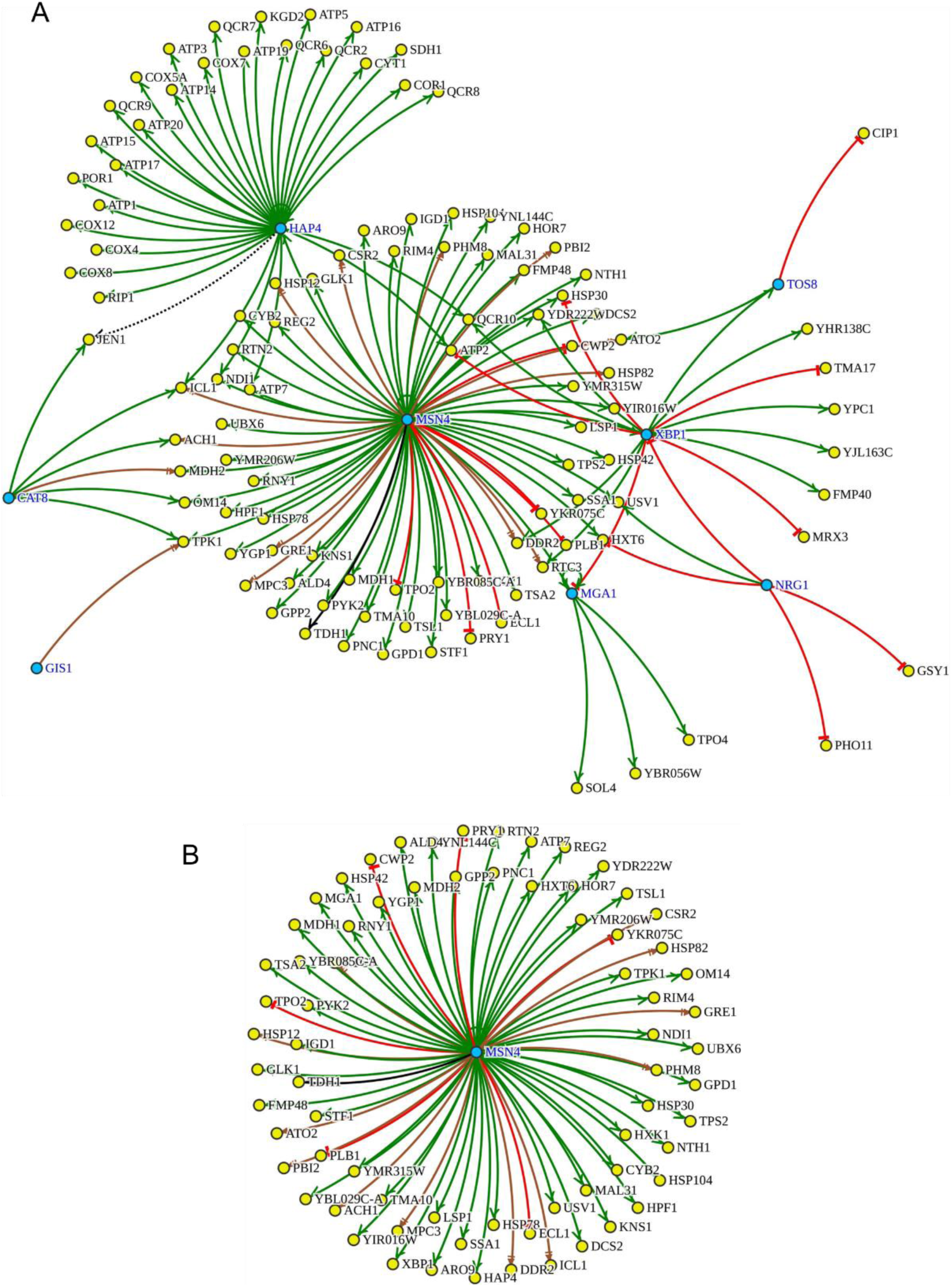
Transcription factors analysis for upregulated DEGs of *ybr238c*Δ mutant, Related to Figure 2. (A) The whole regulatory network of overrepresented top eight enriched transcription factors (blue) within the upregulated DEGs of *ybr238c*Δ mutant. Msn4 and Hap4 TFs are appearing as responsible for the highest number of regulated genes in *ybr238c*Δ mutant. (B) The Msn4 regulon within the upregulated DEGs of *ybr238c*Δ mutant. See also Additional File 3.

**Figure S5.**
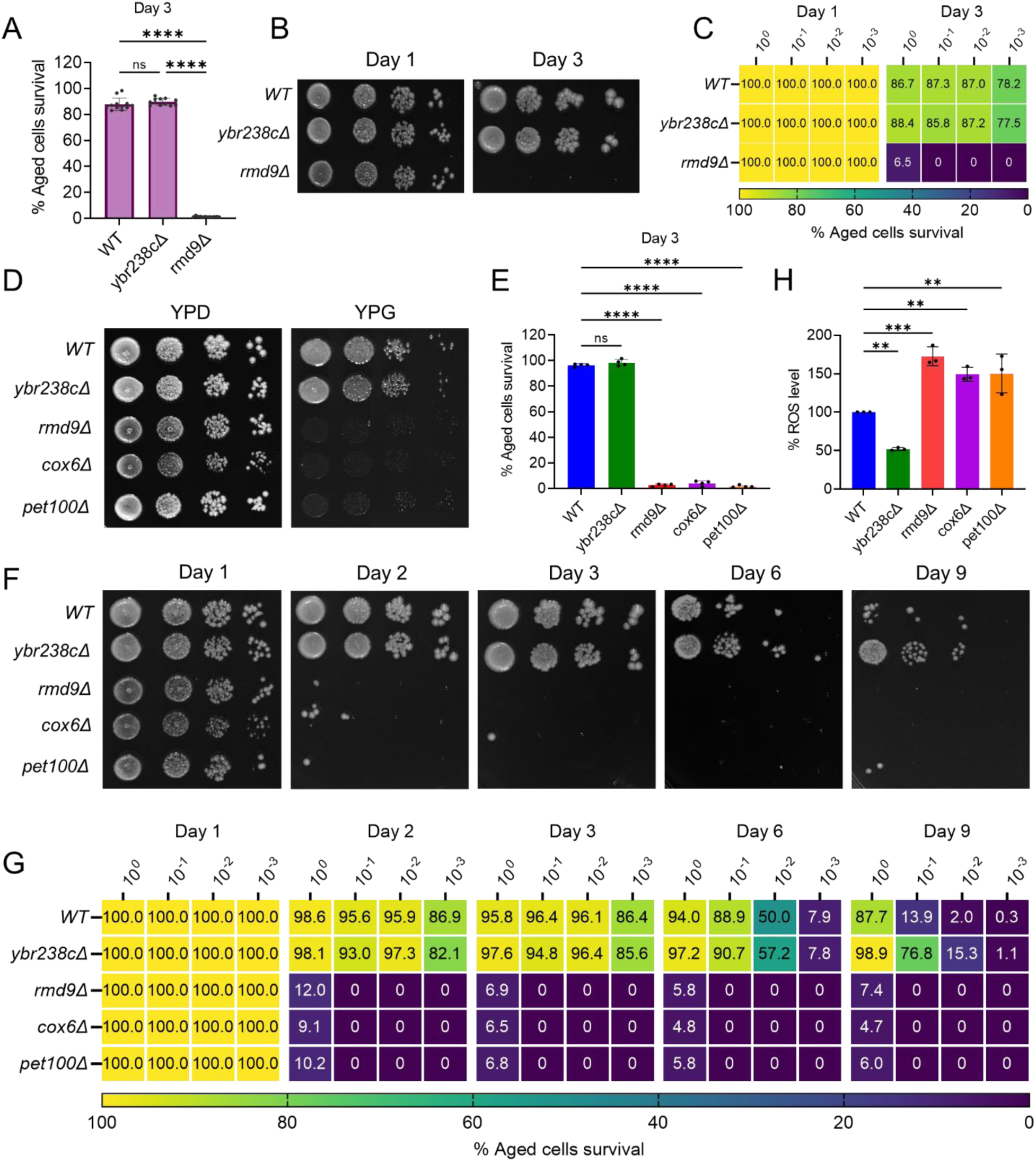
*YBR238C* homolog *RMD9* deletion leads to mitochondrial dysfunction associated with accelerated cellular aging, Related to Figure 4. (A and E) Chronological lifespan (CLS) of yeast wild type BY4743 and deletion strains was performed in synthetic defined medium supplemented with auxotrophic amino acids using 96-well plate. Aged cells survival was measured relative to the outgrowth of day 1. (A) Data are represented as means ± SD (n=12). ****P < 0.0001 based on two-way ANOVA followed by Tukey’s multiple comparisons test. (E) Data are represented as means ± SD (n=4). ****P < 0.0001 based on ordinary one-way ANOVA followed by Dunnett’s multiple comparisons test. ns: non-significant. (B, C, F and G) CLS of yeast wild type BY4743 and deletion strains was performed in synthetic defined medium supplemented with auxotrophic amino acids using the flask. Outgrowth was performed for ten-fold serial diluted aged cells onto the YPD agar plate (B and F) and YPD medium in the 96-well plate (C and G). The serial outgrowth of aged cells on agar medium was imaged (B and F) and quantified the survival relative to outgrowth of day 1 (C and G). A representative of two experiments for B, C, F and G is shown. (D) Serial dilution growth assays of the yeast strains on fermentative glucose medium (YPD) and respiratory glycerol medium (YPG). (H) ROS level of logarithmic-phase yeast wild type BY4743 and deletion strains in synthetic defined medium supplemented with auxotrophic amino acids. Data are represented as means ± SD (n=3). **P < 0.01, and ****P < 0.0001 based on ordinary one-way ANOVA followed by Dunnett’s multiple comparisons test.

**Figure S6.**
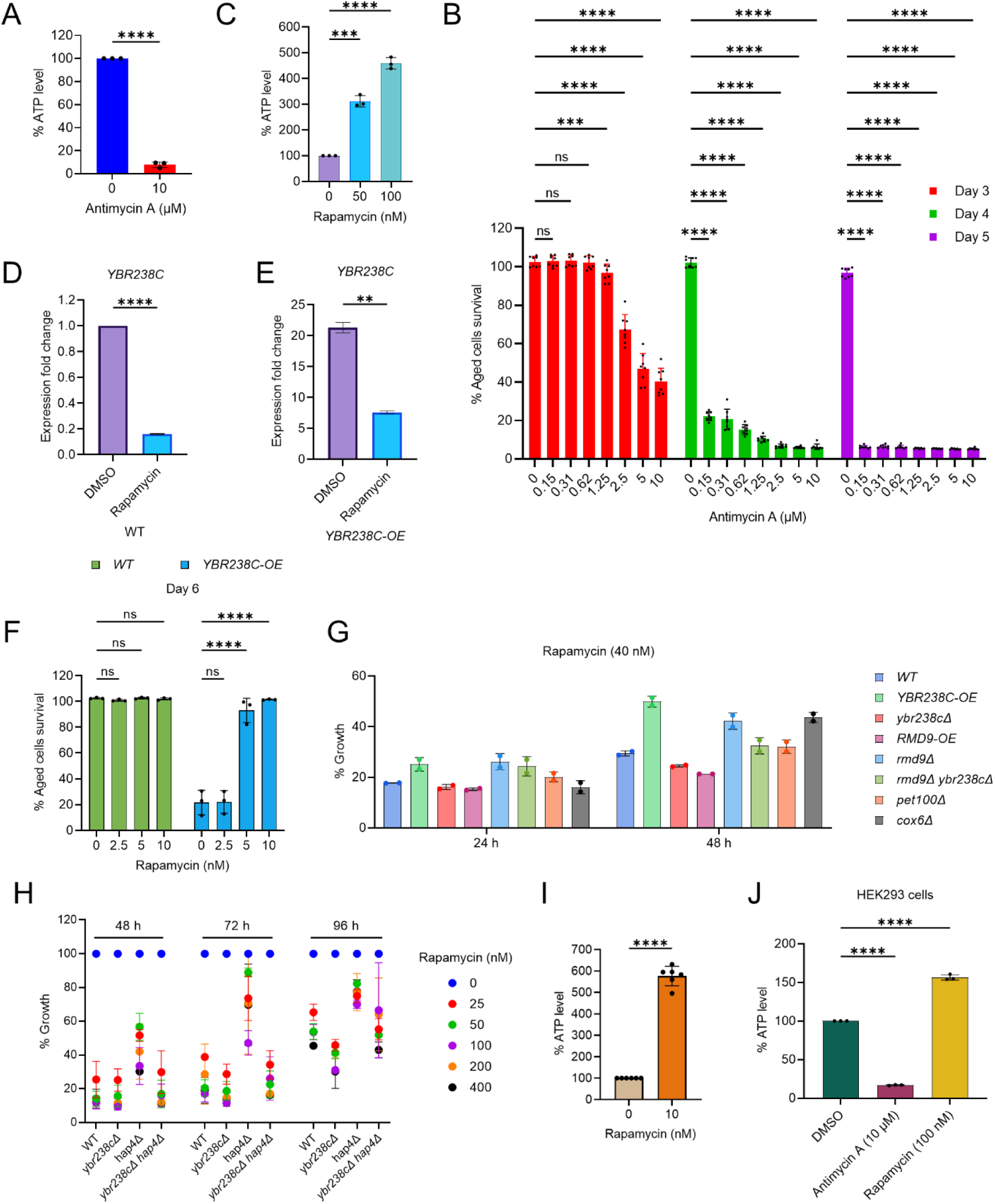
TORC1-Mitochondrial function signaling pathways control cellular aging. (A and C) ATP analysis of wild type CEN.PK113-7D yeast cells treated with (A) antimycin A (72-hour stationary-phase culture) and (C) rapamycin (logarithmic-phase culture) in synthetic defined medium. Data are represented as means ± SD (n=3). ***P < 0.001, and ****P < 0.0001 based on ordinary one-way ANOVA followed by Dunnett’s multiple comparisons test. (C) See also Figure 6A. (B) Chronological lifespan (CLS) of wild type CEN.PK113-7D yeast strain with indicated concentrations of antimycin A was performed in the synthetic defined medium in 96-well plate. Aged cells survival was measured relative to the outgrowth of day 1. Data are represented as means ± SD (n=8). ***P < 0.001, and ****P < 0.0001 based on two-way ANOVA followed by Dunnett’s multiple comparisons test. ns: non-significant. ns: non-significant. (D and E) Expression analysis by qRT-PCR for *YBR238C* of logarithmic-phase yeast CEN.PK113-7D strains treated with rapamycin (200 nM) for 1 hour in synthetic defined medium. Data are represented as means ± SD (n=2). **P < 0.01, and ****P < 0.0001 based on two-sided Student’s t-test. (F) CLS of yeast CEN.PK113-7D strains with indicated concentrations of rapamycin was performed in synthetic defined medium using 96-well plate. Aged cells survival was measured relative to the outgrowth of day 2. Data are represented as means ± SD (n=3). ****P < 0.0001 based on two-way ANOVA followed by Dunnett’s multiple comparisons test. ns: non-significant. See also Figure 6D. (G) Growth assay of yeast CEN.PK113-7D strains with rapamycin (40 nM) for 24 and 48h in synthetic defined medium. Growth was normalized with untreated control. (H) Growth assay of yeast CEN.PK113-7D strains with indicated concentrations of rapamycin for 48, 72 and 96 h in synthetic defined medium. Growth was normalized with untreated control. (I) ATP analysis of stationary-phase wild type CEN.PK113-7D yeast cells in synthetic defined medium incubated with 10 nM rapamycin. Data are represented as means ± SD (n=6). ****P < 0.0001 based on ordinary one-way ANOVA followed by Tukey’s multiple comparisons test. (J) ATP analysis of human HEK293 cells treated with antimycin A (10 µM) and rapamycin (100 nM) for 48 hours. Data are represented as means ± SD (n=3). ****P < 0.0001 based on ordinary one-way ANOVA followed by Tukey’s multiple comparisons test.

## AUTHOR CONTRIBUTIONS STATEMENT

Mohammad Alfatah: Conceiving of the project, Writing, Editing and Funding Acquisition Jolyn Jia Jia Lim: Investigation and Formal analysis

Yizhong Zhnag: Investigation and Formal analysis Arshia Naaz: Investigation and Formal analysis

Trishia Cheng Yi Ning: Investigation and Formal analysis Sonia Yogasundaram: Investigation and Formal analysis Nashrul Afiq Faidzinn: Investigation and Formal analysis Jovian Lin Jing: Investigation and Formal analysis

Birgit Eisenhaber: Investigation and Formal analysis

Frank Eisenhaber: Investigation, Formal analysis, Reviewing and Editing

## ACKNOWLEDGMENTS

We thank Sebastian Maurer-Stroh (Executive Director, BII, A*STAR), Lee Hwee Kuan (Training and Talent Deputy Director, BII, A*STAR), Chandra S Verma (Research Deputy Director, BII, A*STAR), Su Xinyi (Acting Executive Director, IMCB, A*STAR), Farid John Ghadessy (Senior Principal Scientist) and Cheok Chit Fang (Principal Investigator, IMCB, A*STAR) for backing this research.

