## Additional File 1 for "Uncharacterized yeast gene *YBR238C,* an effector of TORC1 signaling in a mitochondrial feedback loop, accelerates cellular aging via *HAP4*- and *RMD9*-dependent mechanisms": replicative_lifespan_decreased_annotations.txt

!
!Date: 11/08/2022
!From: Saccharomyces Genome Database (SGD) 
!URL: http://www.yeastgenome.org/ 
!Contact 
!Funding: NHGRI at US NIH, grant number 5-U41-HG001315 
!

Gene	Gene Systematic Name	Phenotype	Experiment Type	Experiment Type Category	Mutant Information	Strain Background	Chemical	Details	Reference
AAT1	YKL106W	replicative lifespan: decreased 	classical genetics	classical genetics	null Allele: aat1-Δ	S288C		Media: glucose carbon source, 0.5% Details: calorie restriction-mediated lifespan extension is smaller than in wild type 	Easlon E, et al. (2008) PMID:18381895
AAT2	YLR027C	replicative lifespan: decreased 	systematic mutation set 	large-scale survey	null Allele: aat2-Δ	S288C		Details: short-lived mutants, one standard deviation below the mean, were identified using an HTP sequencing-based yeast replicative lifespan screen (SEBYL), with pooled cells subject to four rounds of biotin-based sorting to enrich old vs young cells, followed by sequencing to ID mutants and abundance 	Yu R, et al. (2021) PMID:33790287
ACE2	YLR131C	replicative lifespan: decreased 	systematic mutation set 	large-scale survey	null Allele: ace2-Δ	S288C		Details: short-lived mutants, one standard deviation below the mean, were identified using an HTP sequencing-based yeast replicative lifespan screen (SEBYL), with pooled cells subject to four rounds of biotin-based sorting to enrich old vs young cells, followed by sequencing to ID mutants and abundance 	Yu R, et al. (2021) PMID:33790287
ACS2	YLR153C	replicative lifespan: decreased 	classical genetics	classical genetics	null Allele: acs2-Δ	Other		Media: rich (YPE) medium containing ethanol as carbon source Details: 17 generations relative to 55 generations for wild type cells 	Falcón AA, et al. (2010) PMID:19618123
AFG3	YER017C	replicative lifespan: decreased 	classical genetics	classical genetics	null Allele: afg3-Δ	S288C		Treatment: caloric restriction by glucose reduction, 0.05% Details: dietary restriction-induced lifespan extension is decreased relative to wt 	Schleit J, et al. (2013) PMID:23837470
AFT1	YGL071W	replicative lifespan: decreased 	classical genetics 	classical genetics	activation Allele: AFT1-1UP	S288C			Patnaik PK, et al. (2022) PMID:35858543
AGA1	YNR044W	replicative lifespan: decreased 	systematic mutation set 	large-scale survey	null Allele: aga1-Δ	S288C		Details: short-lived mutants, one standard deviation below the mean, were identified using an HTP sequencing-based yeast replicative lifespan screen (SEBYL), with pooled cells subject to four rounds of biotin-based sorting to enrich old vs young cells, followed by sequencing to ID mutants and abundance 	Yu R, et al. (2021) PMID:33790287
AIM44	YPL158C	replicative lifespan: decreased 	systematic mutation set 	large-scale survey	null Allele: aim44-Δ	S288C		Details: short-lived mutants, one standard deviation below the mean, were identified using an HTP sequencing-based yeast replicative lifespan screen (SEBYL), with pooled cells subject to four rounds of biotin-based sorting to enrich old vs young cells, followed by sequencing to ID mutants and abundance 	Yu R, et al. (2021) PMID:33790287
ALG12	YNR030W	replicative lifespan: decreased 	classical genetics	classical genetics	null Allele: alg12-Δ	S288C		Treatment: caloric restriction by glucose reduction, 0.05% Details: dietary restriction-induced lifespan extension is decreased relative to wt 	Schleit J, et al. (2013) PMID:23837470
AMD1	YML035C	replicative lifespan: decreased 	systematic mutation set 	large-scale survey	null Allele: amd1-Δ	S288C		Details: short-lived mutants, one standard deviation below the mean, were identified using an HTP sequencing-based yeast replicative lifespan screen (SEBYL), with pooled cells subject to four rounds of biotin-based sorting to enrich old vs young cells, followed by sequencing to ID mutants and abundance 	Yu R, et al. (2021) PMID:33790287
APC5	YOR249C	replicative lifespan: decreased 	classical genetics	classical genetics	reduction of function Allele: apc5-CA two bp deleted in codon encoding Phe12, generating 18 additional nonspecific aa followed by a stop	S288C			Harkness TA, et al. (2004) PMID:15514051
APC9	YLR102C	replicative lifespan: decreased 	classical genetics	classical genetics	null Allele: apc9-Δ	S288C			Harkness TA, et al. (2004) PMID:15514051
ARC18	YLR370C	replicative lifespan: decreased 	systematic mutation set 	large-scale survey	null Allele: arc18-Δ	S288C		Details: short-lived mutants, one standard deviation below the mean, were identified using an HTP sequencing-based yeast replicative lifespan screen (SEBYL), with pooled cells subject to four rounds of biotin-based sorting to enrich old vs young cells, followed by sequencing to ID mutants and abundance 	Yu R, et al. (2021) PMID:33790287
ARF1	YDL192W	replicative lifespan: decreased 	classical genetics	classical genetics	null Allele: arf1-Δ	S288C		Details: reduced lifespan; cells do not display characteristics of accelerated aging 	Ashrafi K, et al. (2000) PMID:10921902
ARO1	YDR127W	replicative lifespan: decreased 	homozygous diploid 	classical genetics	null Allele: aro1-Δ	Other		Details: dramatic reduction in median replicative lifespan determined using the Mother Enrichment Program 	Beas AO, et al. (2020) PMID:32493904
ARP1	YHR129C	replicative lifespan: decreased 	systematic mutation set 	large-scale survey	null Allele: arp1-Δ	S288C		Details: short-lived mutants, one standard deviation below the mean, were identified using an HTP sequencing-based yeast replicative lifespan screen (SEBYL), with pooled cells subject to four rounds of biotin-based sorting to enrich old vs young cells, followed by sequencing to ID mutants and abundance 	Yu R, et al. (2021) PMID:33790287
ARP4	YJL081C	replicative lifespan: decreased 	classical genetics 	classical genetics	conditional Allele: arp4-26 G187R	Other		Temperature: permissive temperature, 28 °C Details: significant reduction in the budding lifespan and the post-reproductive lifespan 	Mołoń M, et al. (2022) PMID:36078161
ASF1	YJL115W	replicative lifespan: decreased 	classical genetics	classical genetics	null Allele: asf1-Δ	S288C			Feser J, et al. (2010) PMID:20832724
ASF1	YJL115W	replicative lifespan: decreased 	classical genetics	classical genetics	null Allele: asf1-Δ	W303			Feser J, et al. (2010) PMID:20832724
ASH1	YKL185W	replicative lifespan: decreased 	systematic mutation set 	large-scale survey	null Allele: ash1-Δ	S288C		Details: short-lived mutants, one standard deviation below the mean, were identified using an HTP sequencing-based yeast replicative lifespan screen (SEBYL), with pooled cells subject to four rounds of biotin-based sorting to enrich old vs young cells, followed by sequencing to ID mutants and abundance 	Yu R, et al. (2021) PMID:33790287
ATG1	YGL180W	replicative lifespan: decreased 	classical genetics	classical genetics	null Allele: atg1-Δ	S288C		Treatment: caloric restriction by glucose reduction, 0.05% Details: dietary restriction-induced lifespan extension is decreased relative to wt 	Schleit J, et al. (2013) PMID:23837470
ATG15	YCR068W	replicative lifespan: decreased 	classical genetics	classical genetics	null Allele: atg15-Δ	S288C		Treatment: caloric restriction Details: required for caloric restriction-triggered life-span extension 	Tang F, et al. (2008) PMID:18690010
ATG17	YLR423C	replicative lifespan: decreased 	classical genetics	classical genetics	null Allele: atg17-Δ	S288C		Media: rich medium Details: mean lifespan decreased relative to wildtype 	Tang F, et al. (2008) PMID:18690010
ATG17	YLR423C	replicative lifespan: decreased 	classical genetics	classical genetics	null Allele: atg17-Δ	S288C		Treatment: caloric restriction Details: required for caloric restriction-triggered life-span extension 	Tang F, et al. (2008) PMID:18690010
ATG8	YBL078C	replicative lifespan: decreased 	classical genetics	classical genetics	null Allele: atg8-Δ	S288C		Treatment: caloric restriction by glucose reduction, 0.05% Details: dietary restriction-induced lifespan extension is decreased relative to wt 	Schleit J, et al. (2013) PMID:23837470
ATP2	YJR121W	replicative lifespan: decreased 	classical genetics	classical genetics	conditional Allele: atp2-G325A V109I	Other		Temperature: elevated temperature, 37 °C Details: young cells respire like wild type but older cells begin to lose mitochondrial membrane potential and fail to segregate mitochondria to daughters 	Lai CY, et al. (2002) PMID:12242224
ATP20	YPR020W	replicative lifespan: decreased 	systematic mutation set 	large-scale survey	null Allele: atp20-Δ	S288C		Details: short-lived mutants, one standard deviation below the mean, were identified using an HTP sequencing-based yeast replicative lifespan screen (SEBYL), with pooled cells subject to four rounds of biotin-based sorting to enrich old vs young cells, followed by sequencing to ID mutants and abundance 	Yu R, et al. (2021) PMID:33790287
ATP23	YNR020C	replicative lifespan: decreased 	classical genetics	classical genetics	null Allele: atp23-Δ	S288C			Hacioglu E, et al. (2010) PMID:20934449
AVT1	YJR001W	replicative lifespan: decreased 	homozygous diploid	classical genetics	null Allele: avt1-Δ	S288C		Details: decrease in both median and maximal replicative lifespan 	Hughes AL and Gottschling DE (2012) PMID:23172144
BCY1	YIL033C	replicative lifespan: decreased 	classical genetics	classical genetics	null Allele: bcy1-Δ	Other		Details: null mutants have reduced viability 	Sun J, et al. (1994) PMID:8034612
BEM4	YPL161C	replicative lifespan: decreased 	classical genetics	classical genetics	null Allele: bem4-Δ	S288C		Treatment: caloric restriction by glucose reduction, 0.05% Details: dietary restriction-induced lifespan extension is decreased relative to wt 	Schleit J, et al. (2013) PMID:23837470
BIL1	YOR304C-A	replicative lifespan: decreased 	systematic mutation set 	large-scale survey	null Allele: bil1-Δ	S288C		Details: short-lived mutants, one standard deviation below the mean, were identified using an HTP sequencing-based yeast replicative lifespan screen (SEBYL), with pooled cells subject to four rounds of biotin-based sorting to enrich old vs young cells, followed by sequencing to ID mutants and abundance 	Yu R, et al. (2021) PMID:33790287
BLM10	YFL007W	replicative lifespan: decreased 	classical genetics 	classical genetics	null Allele: blm10-Δ	S288C		Details: decrease in rate and number of bud scars 	Jiang TX, et al. (2021) PMID:33391545
BMH1	YER177W	replicative lifespan: decreased 	systematic mutation set 	large-scale survey	null Allele: bmh1-Δ	S288C		Details: short-lived mutants, one standard deviation below the mean, were identified using an HTP sequencing-based yeast replicative lifespan screen (SEBYL), with pooled cells subject to four rounds of biotin-based sorting to enrich old vs young cells, followed by sequencing to ID mutants and abundance 	Yu R, et al. (2021) PMID:33790287
BNA1	YJR025C	replicative lifespan: decreased 	homozygous diploid 	classical genetics	null Allele: bna1-Δ	Other		Details: significant decrease in replicative lifespan determined using the Mother Enrichment Program 	Beas AO, et al. (2020) PMID:32493904
BNA2	YJR078W	replicative lifespan: decreased 	homozygous diploid 	classical genetics	null Allele: bna2-Δ	Other		Details: significant decrease in replicative lifespan determined using the Mother Enrichment Program 	Beas AO, et al. (2020) PMID:32493904
BNA5	YLR231C	replicative lifespan: decreased 	homozygous diploid 	classical genetics	null Allele: bna5-Δ	Other		Details: significant decrease in replicative lifespan determined using the Mother Enrichment Program 	Beas AO, et al. (2020) PMID:32493904
BNA6	YFR047C	replicative lifespan: decreased 	homozygous diploid 	classical genetics	null Allele: bna6-Δ	Other		Details: significant decrease in replicative lifespan determined using the Mother Enrichment Program 	Beas AO, et al. (2020) PMID:32493904
BNA7	YDR428C	replicative lifespan: decreased 	homozygous diploid 	classical genetics	null Allele: bna7-Δ	Other		Details: significant decrease in replicative lifespan determined using the Mother Enrichment Program 	Beas AO, et al. (2020) PMID:32493904
BNI1	YNL271C	replicative lifespan: decreased 	systematic mutation set 	large-scale survey	null Allele: bni1-Δ	S288C		Details: short-lived mutants, one standard deviation below the mean, were identified using an HTP sequencing-based yeast replicative lifespan screen (SEBYL), with pooled cells subject to four rounds of biotin-based sorting to enrich old vs young cells, followed by sequencing to ID mutants and abundance 	Yu R, et al. (2021) PMID:33790287
BNI4	YNL233W	replicative lifespan: decreased 	systematic mutation set 	large-scale survey	null Allele: bni4-Δ	S288C		Details: short-lived mutants, one standard deviation below the mean, were identified using an HTP sequencing-based yeast replicative lifespan screen (SEBYL), with pooled cells subject to four rounds of biotin-based sorting to enrich old vs young cells, followed by sequencing to ID mutants and abundance 	Yu R, et al. (2021) PMID:33790287
BRE1	YDL074C	replicative lifespan: decreased 	systematic mutation set 	large-scale survey	null Allele: bre1-Δ	S288C		Details: short-lived mutants, one standard deviation below the mean, were identified using an HTP sequencing-based yeast replicative lifespan screen (SEBYL), with pooled cells subject to four rounds of biotin-based sorting to enrich old vs young cells, followed by sequencing to ID mutants and abundance 	Yu R, et al. (2021) PMID:33790287
BRE2	YLR015W	replicative lifespan: decreased 	systematic mutation set 	large-scale survey	null Allele: bre2-Δ	S288C		Details: short-lived mutants, one standard deviation below the mean, were identified using an HTP sequencing-based yeast replicative lifespan screen (SEBYL), with pooled cells subject to four rounds of biotin-based sorting to enrich old vs young cells, followed by sequencing to ID mutants and abundance 	Yu R, et al. (2021) PMID:33790287
BRE5	YNR051C	replicative lifespan: decreased 	systematic mutation set 	large-scale survey	null Allele: bre5-Δ	S288C		Details: short-lived mutants, one standard deviation below the mean, were identified using an HTP sequencing-based yeast replicative lifespan screen (SEBYL), with pooled cells subject to four rounds of biotin-based sorting to enrich old vs young cells, followed by sequencing to ID mutants and abundance 	Yu R, et al. (2021) PMID:33790287
BRP1	YGL007W	replicative lifespan: decreased 	systematic mutation set 	large-scale survey	null Allele: brp1-Δ	S288C		Details: short-lived mutants, one standard deviation below the mean, were identified using an HTP sequencing-based yeast replicative lifespan screen (SEBYL), with pooled cells subject to four rounds of biotin-based sorting to enrich old vs young cells, followed by sequencing to ID mutants and abundance 	Yu R, et al. (2021) PMID:33790287
BTS1	YPL069C	replicative lifespan: decreased 	systematic mutation set 	large-scale survey	null Allele: bts1-Δ	S288C		Details: short-lived mutants, one standard deviation below the mean, were identified using an HTP sequencing-based yeast replicative lifespan screen (SEBYL), with pooled cells subject to four rounds of biotin-based sorting to enrich old vs young cells, followed by sequencing to ID mutants and abundance 	Yu R, et al. (2021) PMID:33790287
CAC2	YML102W	replicative lifespan: decreased 	systematic mutation set 	large-scale survey	null Allele: cac2-Δ	S288C		Details: short-lived mutants, one standard deviation below the mean, were identified using an HTP sequencing-based yeast replicative lifespan screen (SEBYL), with pooled cells subject to four rounds of biotin-based sorting to enrich old vs young cells, followed by sequencing to ID mutants and abundance 	Yu R, et al. (2021) PMID:33790287
CAJ1	YER048C	replicative lifespan: decreased 	systematic mutation set 	large-scale survey	null Allele: caj1-Δ	S288C		Details: short-lived mutants, one standard deviation below the mean, were identified using an HTP sequencing-based yeast replicative lifespan screen (SEBYL), with pooled cells subject to four rounds of biotin-based sorting to enrich old vs young cells, followed by sequencing to ID mutants and abundance 	Yu R, et al. (2021) PMID:33790287
CAR2	YLR438W	replicative lifespan: decreased 	classical genetics 	classical genetics	null Allele: car2-Δ	S288C		Details: markedly shortened lifespan relative to wt 	Mukai Y, et al. (2019) PMID:31646149
CCR4	YAL021C	replicative lifespan: decreased 	systematic mutation set 	large-scale survey	null Allele: ccr4-Δ	S288C		Details: short-lived mutants, one standard deviation below the mean, were identified using an HTP sequencing-based yeast replicative lifespan screen (SEBYL), with pooled cells subject to four rounds of biotin-based sorting to enrich old vs young cells, followed by sequencing to ID mutants and abundance 	Yu R, et al. (2021) PMID:33790287
CDC26	YFR036W	replicative lifespan: decreased 	classical genetics	classical genetics	null Allele: cdc26-Δ	S288C			Harkness TA, et al. (2004) PMID:15514051
CHL1	YPL008W	replicative lifespan: decreased 	classical genetics	classical genetics	null Allele: chl1-Δ	W303		Details: ~ 1/3 reduction compared to wild type 	Das SP and Sinha P (2005) PMID:16182251
CHO2	YGR157W	replicative lifespan: decreased 	systematic mutation set 	large-scale survey	null Allele: cho2-Δ	S288C		Details: short-lived mutants, one standard deviation below the mean, were identified using an HTP sequencing-based yeast replicative lifespan screen (SEBYL), with pooled cells subject to four rounds of biotin-based sorting to enrich old vs young cells, followed by sequencing to ID mutants and abundance 	Yu R, et al. (2021) PMID:33790287
CHS3	YBR023C	replicative lifespan: decreased 	classical genetics 	classical genetics	null Allele: chs3-Δ	S288C			Molon M, et al. (2018) PMID:29189912
CHS6	YJL099W	replicative lifespan: decreased 	systematic mutation set 	large-scale survey	null Allele: chs6-Δ	S288C		Details: short-lived mutants, one standard deviation below the mean, were identified using an HTP sequencing-based yeast replicative lifespan screen (SEBYL), with pooled cells subject to four rounds of biotin-based sorting to enrich old vs young cells, followed by sequencing to ID mutants and abundance 	Yu R, et al. (2021) PMID:33790287
CKB2	YOR039W	replicative lifespan: decreased 	systematic mutation set 	large-scale survey	null Allele: ckb2-Δ	S288C		Details: short-lived mutants, one standard deviation below the mean, were identified using an HTP sequencing-based yeast replicative lifespan screen (SEBYL), with pooled cells subject to four rounds of biotin-based sorting to enrich old vs young cells, followed by sequencing to ID mutants and abundance 	Yu R, et al. (2021) PMID:33790287
CLA4	YNL298W	replicative lifespan: decreased 	systematic mutation set 	large-scale survey	null Allele: cla4-Δ	S288C		Details: short-lived mutants, one standard deviation below the mean, were identified using an HTP sequencing-based yeast replicative lifespan screen (SEBYL), with pooled cells subject to four rounds of biotin-based sorting to enrich old vs young cells, followed by sequencing to ID mutants and abundance 	Yu R, et al. (2021) PMID:33790287
CLB5	YPR120C	replicative lifespan: decreased 	systematic mutation set 	large-scale survey	null Allele: clb5-Δ	S288C		Details: short-lived mutants, one standard deviation below the mean, were identified using an HTP sequencing-based yeast replicative lifespan screen (SEBYL), with pooled cells subject to four rounds of biotin-based sorting to enrich old vs young cells, followed by sequencing to ID mutants and abundance 	Yu R, et al. (2021) PMID:33790287
CLG1	YGL215W	replicative lifespan: decreased 	classical genetics 	classical genetics	null Allele: clg1-Δ	S288C		Media: high phosphate, 11 mM Details: moderately short lived under high Pi conditions; significantly shorter in pho80 double null 	Nakajima T, et al. (2020) PMID:31797348
CMP2	YML057W	replicative lifespan: decreased 	classical genetics	classical genetics	overexpression Allele: CMP2-ΔC constitutively activated C-terminal auto-inhibitory domain-truncated catalytic subunit	Other			Tsubakiyama R, et al. (2011) PMID:21712379
CNB1	YKL190W	replicative lifespan: decreased 	classical genetics	classical genetics	null Allele: cnb1-Δ	Other			Tsubakiyama R, et al. (2011) PMID:21712379
CNL1	YDR357C	replicative lifespan: decreased 	systematic mutation set 	large-scale survey	null Allele: cnl1-Δ	S288C		Details: short-lived mutants, one standard deviation below the mean, were identified using an HTP sequencing-based yeast replicative lifespan screen (SEBYL), with pooled cells subject to four rounds of biotin-based sorting to enrich old vs young cells, followed by sequencing to ID mutants and abundance 	Yu R, et al. (2021) PMID:33790287
CSF1	YLR087C	replicative lifespan: decreased 	systematic mutation set 	large-scale survey	null Allele: csf1-Δ	S288C		Details: short-lived mutants, one standard deviation below the mean, were identified using an HTP sequencing-based yeast replicative lifespan screen (SEBYL), with pooled cells subject to four rounds of biotin-based sorting to enrich old vs young cells, followed by sequencing to ID mutants and abundance 	Yu R, et al. (2021) PMID:33790287
CSM3	YMR048W	replicative lifespan: decreased 	systematic mutation set 	large-scale survey	null Allele: csm3-Δ	S288C		Details: short-lived mutants, one standard deviation below the mean, were identified using an HTP sequencing-based yeast replicative lifespan screen (SEBYL), with pooled cells subject to four rounds of biotin-based sorting to enrich old vs young cells, followed by sequencing to ID mutants and abundance 	Yu R, et al. (2021) PMID:33790287
CST26	YBR042C	replicative lifespan: decreased 	classical genetics	classical genetics	null Allele: cst26-Δ	S288C			Hacioglu E, et al. (2010) PMID:20934449
CTF4	YPR135W	replicative lifespan: decreased 	classical genetics	classical genetics	null Allele: ctf4-Δ	Other			Hoopes LL, et al. (2002) PMID:12024027
CTF8	YHR191C	replicative lifespan: decreased 	systematic mutation set 	large-scale survey	null Allele: ctf8-Δ	S288C		Details: short-lived mutants, one standard deviation below the mean, were identified using an HTP sequencing-based yeast replicative lifespan screen (SEBYL), with pooled cells subject to four rounds of biotin-based sorting to enrich old vs young cells, followed by sequencing to ID mutants and abundance 	Yu R, et al. (2021) PMID:33790287
CTS1	YLR286C	replicative lifespan: decreased 	classical genetics 	classical genetics	null Allele: cts1-Δ	S288C			Molon M, et al. (2018) PMID:29189912
CUE1	YMR264W	replicative lifespan: decreased 	systematic mutation set 	large-scale survey	null Allele: cue1-Δ	S288C		Details: short-lived mutants, one standard deviation below the mean, were identified using an HTP sequencing-based yeast replicative lifespan screen (SEBYL), with pooled cells subject to four rounds of biotin-based sorting to enrich old vs young cells, followed by sequencing to ID mutants and abundance 	Yu R, et al. (2021) PMID:33790287
CYK3	YDL117W	replicative lifespan: decreased 	systematic mutation set 	large-scale survey	null Allele: cyk3-Δ	S288C		Details: short-lived mutants, one standard deviation below the mean, were identified using an HTP sequencing-based yeast replicative lifespan screen (SEBYL), with pooled cells subject to four rounds of biotin-based sorting to enrich old vs young cells, followed by sequencing to ID mutants and abundance 	Yu R, et al. (2021) PMID:33790287
CYR1	YJL005W	replicative lifespan: decreased 	classical genetics	classical genetics	overexpression	Other		Details: large percentage of cells unable to divide, are presumed dead and did not contribute to the survival curve 	Sun J, et al. (1994) PMID:8034612
CYS4	YGR155W	replicative lifespan: decreased 	systematic mutation set 	large-scale survey	null Allele: cys4-Δ	S288C		Details: short-lived mutants, one standard deviation below the mean, were identified using an HTP sequencing-based yeast replicative lifespan screen (SEBYL), with pooled cells subject to four rounds of biotin-based sorting to enrich old vs young cells, followed by sequencing to ID mutants and abundance 	Yu R, et al. (2021) PMID:33790287
DAT1	YML113W	replicative lifespan: decreased 	systematic mutation set 	large-scale survey	null Allele: dat1-Δ	S288C		Details: short-lived mutants, one standard deviation below the mean, were identified using an HTP sequencing-based yeast replicative lifespan screen (SEBYL), with pooled cells subject to four rounds of biotin-based sorting to enrich old vs young cells, followed by sequencing to ID mutants and abundance 	Yu R, et al. (2021) PMID:33790287
DBP3	YGL078C	replicative lifespan: decreased 	classical genetics	classical genetics	null Allele: dbp3-Δ	S288C		Treatment: caloric restriction by glucose reduction, 0.05% Details: dietary restriction-induced lifespan extension is decreased relative to wt 	Schleit J, et al. (2013) PMID:23837470
DEG1	YFL001W	replicative lifespan: decreased 	classical genetics	classical genetics	unspecified Allele: deg1-(hrm3)	Other			Defossez PA, et al. (1999) PMID:10230397
DFG10	YIL049W	replicative lifespan: decreased 	systematic mutation set 	large-scale survey	null Allele: dfg10-Δ	S288C		Details: short-lived mutants, one standard deviation below the mean, were identified using an HTP sequencing-based yeast replicative lifespan screen (SEBYL), with pooled cells subject to four rounds of biotin-based sorting to enrich old vs young cells, followed by sequencing to ID mutants and abundance 	Yu R, et al. (2021) PMID:33790287
DFG16	YOR030W	replicative lifespan: decreased 	systematic mutation set 	large-scale survey	null Allele: dfg16-Δ	S288C		Details: short-lived mutants, one standard deviation below the mean, were identified using an HTP sequencing-based yeast replicative lifespan screen (SEBYL), with pooled cells subject to four rounds of biotin-based sorting to enrich old vs young cells, followed by sequencing to ID mutants and abundance 	Yu R, et al. (2021) PMID:33790287
DHH1	YDL160C	replicative lifespan: decreased 	classical genetics	classical genetics	null Allele: dhh1-Δ	S288C		Details: shortened by 20- 30% compared with wild-type 	Duan R, et al. (2013) PMID:24358479
DHH1	YDL160C	replicative lifespan: decreased 	systematic mutation set 	large-scale survey	null Allele: dhh1-Δ	S288C		Details: short-lived mutants, one standard deviation below the mean, were identified using an HTP sequencing-based yeast replicative lifespan screen (SEBYL), with pooled cells subject to four rounds of biotin-based sorting to enrich old vs young cells, followed by sequencing to ID mutants and abundance 	Yu R, et al. (2021) PMID:33790287
DJP1	YIR004W	replicative lifespan: decreased 	systematic mutation set 	large-scale survey	null Allele: djp1-Δ	S288C		Details: short-lived mutants, one standard deviation below the mean, were identified using an HTP sequencing-based yeast replicative lifespan screen (SEBYL), with pooled cells subject to four rounds of biotin-based sorting to enrich old vs young cells, followed by sequencing to ID mutants and abundance 	Yu R, et al. (2021) PMID:33790287
DLT1	YMR126C	replicative lifespan: decreased 	classical genetics	classical genetics	null Allele: dlt1-Δ	S288C		Treatment: caloric restriction by glucose reduction, 0.05% Details: dietary restriction-induced lifespan extension is decreased relative to wt 	Schleit J, et al. (2013) PMID:23837470
DNA2	YHR164C	replicative lifespan: decreased 	classical genetics	classical genetics	conditional Allele: dna2-1 P504S; nuclease domain mutated	Other		Temperature: permissive temperature, 23 °C Details: age-related increase in sterility 	Hoopes LL, et al. (2002) PMID:12024027
DNA2	YHR164C	replicative lifespan: decreased 	classical genetics	classical genetics	unspecified Allele: dna2-2 R1253Q	Other		Details: age-dependent increase in mother cell size, nucleolar enlargement and fragmentation; similar results with dna2-20 and dna2-21 	Hoopes LL, et al. (2002) PMID:12024027
DNA2	YHR164C	replicative lifespan: decreased 	classical genetics	classical genetics	unspecified Allele: dna2-2 R1253Q	W303		Details: age-dependent hyperrecombination at the rDNA locus 	Hoopes LL, et al. (2002) PMID:12024027
DOA4	YDR069C	replicative lifespan: decreased 	systematic mutation set 	large-scale survey	null Allele: doa4-Δ	S288C		Details: short-lived mutants, one standard deviation below the mean, were identified using an HTP sequencing-based yeast replicative lifespan screen (SEBYL), with pooled cells subject to four rounds of biotin-based sorting to enrich old vs young cells, followed by sequencing to ID mutants and abundance 	Yu R, et al. (2021) PMID:33790287
DOC1	YGL240W	replicative lifespan: decreased 	classical genetics	classical genetics	null Allele: doc1-Δ	S288C			Harkness TA, et al. (2004) PMID:15514051
DPB4	YDR121W	replicative lifespan: decreased 	systematic mutation set 	large-scale survey	null Allele: dpb4-Δ	S288C		Details: short-lived mutants, one standard deviation below the mean, were identified using an HTP sequencing-based yeast replicative lifespan screen (SEBYL), with pooled cells subject to four rounds of biotin-based sorting to enrich old vs young cells, followed by sequencing to ID mutants and abundance 	Yu R, et al. (2021) PMID:33790287
DPH1	YIL103W	replicative lifespan: decreased 	systematic mutation set 	large-scale survey	null Allele: dph1-Δ	S288C		Details: short-lived mutants, one standard deviation below the mean, were identified using an HTP sequencing-based yeast replicative lifespan screen (SEBYL), with pooled cells subject to four rounds of biotin-based sorting to enrich old vs young cells, followed by sequencing to ID mutants and abundance 	Yu R, et al. (2021) PMID:33790287
DPH2	YKL191W	replicative lifespan: decreased 	systematic mutation set 	large-scale survey	null Allele: dph2-Δ	S288C		Details: short-lived mutants, one standard deviation below the mean, were identified using an HTP sequencing-based yeast replicative lifespan screen (SEBYL), with pooled cells subject to four rounds of biotin-based sorting to enrich old vs young cells, followed by sequencing to ID mutants and abundance 	Yu R, et al. (2021) PMID:33790287
DPH5	YLR172C	replicative lifespan: decreased 	systematic mutation set 	large-scale survey	null Allele: dph5-Δ	S288C		Details: short-lived mutants, one standard deviation below the mean, were identified using an HTP sequencing-based yeast replicative lifespan screen (SEBYL), with pooled cells subject to four rounds of biotin-based sorting to enrich old vs young cells, followed by sequencing to ID mutants and abundance 	Yu R, et al. (2021) PMID:33790287
DRS2	YAL026C	replicative lifespan: decreased 	systematic mutation set 	large-scale survey	null Allele: drs2-Δ	S288C		Details: short-lived mutants, one standard deviation below the mean, were identified using an HTP sequencing-based yeast replicative lifespan screen (SEBYL), with pooled cells subject to four rounds of biotin-based sorting to enrich old vs young cells, followed by sequencing to ID mutants and abundance 	Yu R, et al. (2021) PMID:33790287
DUN1	YDL101C	replicative lifespan: decreased 	systematic mutation set 	large-scale survey	null Allele: dun1-Δ	S288C		Details: short-lived mutants, one standard deviation below the mean, were identified using an HTP sequencing-based yeast replicative lifespan screen (SEBYL), with pooled cells subject to four rounds of biotin-based sorting to enrich old vs young cells, followed by sequencing to ID mutants and abundance 	Yu R, et al. (2021) PMID:33790287
EAF1	YDR359C	replicative lifespan: decreased 	systematic mutation set 	large-scale survey	null Allele: eaf1-Δ	S288C		Details: short-lived mutants, one standard deviation below the mean, were identified using an HTP sequencing-based yeast replicative lifespan screen (SEBYL), with pooled cells subject to four rounds of biotin-based sorting to enrich old vs young cells, followed by sequencing to ID mutants and abundance 	Yu R, et al. (2021) PMID:33790287
EAF7	YNL136W	replicative lifespan: decreased 	systematic mutation set 	large-scale survey	null Allele: eaf7-Δ	S288C		Details: short-lived mutants, one standard deviation below the mean, were identified using an HTP sequencing-based yeast replicative lifespan screen (SEBYL), with pooled cells subject to four rounds of biotin-based sorting to enrich old vs young cells, followed by sequencing to ID mutants and abundance 	Yu R, et al. (2021) PMID:33790287
ECM29	YHL030W	replicative lifespan: decreased 	systematic mutation set 	large-scale survey	null Allele: ecm29-Δ	S288C		Details: short-lived mutants, one standard deviation below the mean, were identified using an HTP sequencing-based yeast replicative lifespan screen (SEBYL), with pooled cells subject to four rounds of biotin-based sorting to enrich old vs young cells, followed by sequencing to ID mutants and abundance 	Yu R, et al. (2021) PMID:33790287
EFM4	YIL064W	replicative lifespan: decreased 	systematic mutation set 	large-scale survey	null Allele: efm4-Δ	S288C		Details: short-lived mutants, one standard deviation below the mean, were identified using an HTP sequencing-based yeast replicative lifespan screen (SEBYL), with pooled cells subject to four rounds of biotin-based sorting to enrich old vs young cells, followed by sequencing to ID mutants and abundance 	Yu R, et al. (2021) PMID:33790287
ELO3	YLR372W	replicative lifespan: decreased 	classical genetics	classical genetics	null Allele: elo3-Δ	S288C		Treatment: caloric restriction Details: required for caloric restriction-triggered life-span extension 	Tang F, et al. (2008) PMID:18690010
ELP3	YPL086C	replicative lifespan: decreased 	systematic mutation set 	large-scale survey	null Allele: elp3-Δ	S288C		Details: short-lived mutants, one standard deviation below the mean, were identified using an HTP sequencing-based yeast replicative lifespan screen (SEBYL), with pooled cells subject to four rounds of biotin-based sorting to enrich old vs young cells, followed by sequencing to ID mutants and abundance 	Yu R, et al. (2021) PMID:33790287
ELP4	YPL101W	replicative lifespan: decreased 	systematic mutation set 	large-scale survey	null Allele: elp4-Δ	S288C		Details: short-lived mutants, one standard deviation below the mean, were identified using an HTP sequencing-based yeast replicative lifespan screen (SEBYL), with pooled cells subject to four rounds of biotin-based sorting to enrich old vs young cells, followed by sequencing to ID mutants and abundance 	Yu R, et al. (2021) PMID:33790287
ELP6	YMR312W	replicative lifespan: decreased 	systematic mutation set 	large-scale survey	null Allele: elp6-Δ	S288C		Details: short-lived mutants, one standard deviation below the mean, were identified using an HTP sequencing-based yeast replicative lifespan screen (SEBYL), with pooled cells subject to four rounds of biotin-based sorting to enrich old vs young cells, followed by sequencing to ID mutants and abundance 	Yu R, et al. (2021) PMID:33790287
ENT4	YLL038C	replicative lifespan: decreased 	systematic mutation set 	large-scale survey	null Allele: ent4-Δ	S288C		Details: short-lived mutants, one standard deviation below the mean, were identified using an HTP sequencing-based yeast replicative lifespan screen (SEBYL), with pooled cells subject to four rounds of biotin-based sorting to enrich old vs young cells, followed by sequencing to ID mutants and abundance 	Yu R, et al. (2021) PMID:33790287
ERG2	YMR202W	replicative lifespan: decreased 	classical genetics	classical genetics	null Allele: erg2-Δ	S288C		Media: rich medium Details: mean lifespan of cells from petite colonies decreased relative to wildtype 	Tang F, et al. (2008) PMID:18690010
ERG2	YMR202W	replicative lifespan: decreased 	classical genetics	classical genetics	null Allele: erg2-Δ	S288C		Treatment: caloric restriction Details: cells from petite colonies do not display a life-span extension due to caloric restriction, as observed in wildtype cells 	Tang F, et al. (2008) PMID:18690010
ERG5	YMR015C	replicative lifespan: decreased 	classical genetics	classical genetics	null Allele: erg5-Δ	S288C		Treatment: caloric restriction Details: required for caloric restriction-triggered life-span extension 	Tang F, et al. (2008) PMID:18690010
ERG6	YML008C	replicative lifespan: decreased 	classical genetics	classical genetics	null Allele: erg6-Δ	S288C		Treatment: caloric restriction by glucose reduction, 0.05% Details: dietary restriction-induced lifespan extension is decreased relative to wt 	Schleit J, et al. (2013) PMID:23837470
ERG6	YML008C	replicative lifespan: decreased 	classical genetics	classical genetics	null Allele: erg6-Δ	S288C		Treatment: caloric restriction Details: required for caloric restriction-triggered life-span extension 	Tang F, et al. (2008) PMID:18690010
ESC2	YDR363W	replicative lifespan: decreased 	classical genetics	classical genetics	null Allele: esc2-Δ	W303		Details: mean replicative lifespan reduced from 19.0 to 9.8 generations in the mutant 	Ohya T, et al. (2008) PMID:18757937
EST2	YLR318W	replicative lifespan: decreased 	systematic mutation set 	large-scale survey	null Allele: est2-Δ	S288C		Details: short-lived mutants, one standard deviation below the mean, were identified using an HTP sequencing-based yeast replicative lifespan screen (SEBYL), with pooled cells subject to four rounds of biotin-based sorting to enrich old vs young cells, followed by sequencing to ID mutants and abundance 	Yu R, et al. (2021) PMID:33790287
EST3	YIL009C-A	replicative lifespan: decreased 	systematic mutation set 	large-scale survey	null Allele: est3-Δ	S288C		Details: short-lived mutants, one standard deviation below the mean, were identified using an HTP sequencing-based yeast replicative lifespan screen (SEBYL), with pooled cells subject to four rounds of biotin-based sorting to enrich old vs young cells, followed by sequencing to ID mutants and abundance 	Yu R, et al. (2021) PMID:33790287
EUG1	YDR518W	replicative lifespan: decreased 	classical genetics	classical genetics	null Allele: eug1-Δ	S288C			Hacioglu E, et al. (2010) PMID:20934449
FAT1	YBR041W	replicative lifespan: decreased 	systematic mutation set 	large-scale survey	null Allele: fat1-Δ	S288C		Details: short-lived mutants, one standard deviation below the mean, were identified using an HTP sequencing-based yeast replicative lifespan screen (SEBYL), with pooled cells subject to four rounds of biotin-based sorting to enrich old vs young cells, followed by sequencing to ID mutants and abundance 	Yu R, et al. (2021) PMID:33790287
FHL1	YPR104C	replicative lifespan: decreased 	heterozygous diploid	classical genetics	null Allele: fhl1-Δ	S288C		Details: lifespan considerably short relative to wt 	Kamei Y, et al. (2015) PMID:26022127
FIG4	YNL325C	replicative lifespan: decreased 	classical genetics	classical genetics	null Allele: fig4-Δ	S288C			Hacioglu E, et al. (2010) PMID:20934449
FKS1	YLR342W	replicative lifespan: decreased 	systematic mutation set 	large-scale survey	null Allele: fks1-Δ	S288C		Details: short-lived mutants, one standard deviation below the mean, were identified using an HTP sequencing-based yeast replicative lifespan screen (SEBYL), with pooled cells subject to four rounds of biotin-based sorting to enrich old vs young cells, followed by sequencing to ID mutants and abundance 	Yu R, et al. (2021) PMID:33790287
FLO8	YER109C	replicative lifespan: decreased 	systematic mutation set 	large-scale survey	null Allele: flo8-Δ	S288C		Details: short-lived mutants, one standard deviation below the mean, were identified using an HTP sequencing-based yeast replicative lifespan screen (SEBYL), with pooled cells subject to four rounds of biotin-based sorting to enrich old vs young cells, followed by sequencing to ID mutants and abundance 	Yu R, et al. (2021) PMID:33790287
FPS1	YLL043W	replicative lifespan: decreased 	systematic mutation set 	large-scale survey	null Allele: fps1-Δ	S288C		Details: short-lived mutants, one standard deviation below the mean, were identified using an HTP sequencing-based yeast replicative lifespan screen (SEBYL), with pooled cells subject to four rounds of biotin-based sorting to enrich old vs young cells, followed by sequencing to ID mutants and abundance 	Yu R, et al. (2021) PMID:33790287
FPY1	YMR178W	replicative lifespan: decreased 	systematic mutation set 	large-scale survey	null Allele: fpy1-Δ	S288C		Details: short-lived mutants, one standard deviation below the mean, were identified using an HTP sequencing-based yeast replicative lifespan screen (SEBYL), with pooled cells subject to four rounds of biotin-based sorting to enrich old vs young cells, followed by sequencing to ID mutants and abundance 	Yu R, et al. (2021) PMID:33790287
FTR1	YER145C	replicative lifespan: decreased 	systematic mutation set 	large-scale survey	null Allele: ftr1-Δ	S288C		Details: short-lived mutants, one standard deviation below the mean, were identified using an HTP sequencing-based yeast replicative lifespan screen (SEBYL), with pooled cells subject to four rounds of biotin-based sorting to enrich old vs young cells, followed by sequencing to ID mutants and abundance 	Yu R, et al. (2021) PMID:33790287
FUN26	YAL022C	replicative lifespan: decreased 	systematic mutation set 	large-scale survey	null Allele: fun26-Δ	S288C		Details: short-lived mutants, one standard deviation below the mean, were identified using an HTP sequencing-based yeast replicative lifespan screen (SEBYL), with pooled cells subject to four rounds of biotin-based sorting to enrich old vs young cells, followed by sequencing to ID mutants and abundance 	Yu R, et al. (2021) PMID:33790287
FUS3	YBL016W	replicative lifespan: decreased 	systematic mutation set 	large-scale survey	null Allele: fus3-Δ	S288C		Details: short-lived mutants, one standard deviation below the mean, were identified using an HTP sequencing-based yeast replicative lifespan screen (SEBYL), with pooled cells subject to four rounds of biotin-based sorting to enrich old vs young cells, followed by sequencing to ID mutants and abundance 	Yu R, et al. (2021) PMID:33790287
FYV10	YIL097W	replicative lifespan: decreased 	systematic mutation set 	large-scale survey	null Allele: fyv10-Δ	S288C		Details: short-lived mutants, one standard deviation below the mean, were identified using an HTP sequencing-based yeast replicative lifespan screen (SEBYL), with pooled cells subject to four rounds of biotin-based sorting to enrich old vs young cells, followed by sequencing to ID mutants and abundance 	Yu R, et al. (2021) PMID:33790287
FYV5	YCL058C	replicative lifespan: decreased 	systematic mutation set 	large-scale survey	null Allele: fyv5-Δ	S288C		Details: short-lived mutants, one standard deviation below the mean, were identified using an HTP sequencing-based yeast replicative lifespan screen (SEBYL), with pooled cells subject to four rounds of biotin-based sorting to enrich old vs young cells, followed by sequencing to ID mutants and abundance 	Yu R, et al. (2021) PMID:33790287
GAD1	YMR250W	replicative lifespan: decreased 	classical genetics	classical genetics	null Allele: gad1-Δ	S288C		Treatment: caloric restriction by glucose reduction, 0.05% Details: dietary restriction-induced lifespan extension is decreased relative to wt 	Schleit J, et al. (2013) PMID:23837470
GCN5	YGR252W	replicative lifespan: decreased 	classical genetics 	classical genetics	null Allele: gcn5-Δ	S288C	10 μg/ml (-)-epigallocatechin 3-gallate	Details: effects of epigallocatechin gallate (EGCG), a catechin and histone acetyltransferase inhibitor, on lifespan extension are abolished  	Huang B, et al. (2020) PMID:32157780
GCN5	YGR252W	replicative lifespan: decreased 	classical genetics 	classical genetics	null Allele: gcn5-Δ	S288C		Media: glucose-containing media, 0.5 % Details: caloric restriction-mediated lifespan extension present in wt cells is absent in the null mutant  	Huang B, et al. (2020) PMID:32157780
GCR2	YNL199C	replicative lifespan: decreased 	systematic mutation set 	large-scale survey	null Allele: gcr2-Δ	S288C		Details: short-lived mutants, one standard deviation below the mean, were identified using an HTP sequencing-based yeast replicative lifespan screen (SEBYL), with pooled cells subject to four rounds of biotin-based sorting to enrich old vs young cells, followed by sequencing to ID mutants and abundance 	Yu R, et al. (2021) PMID:33790287
GEP4	YHR100C	replicative lifespan: decreased 	systematic mutation set 	large-scale survey	null Allele: gep4-Δ	S288C		Details: short-lived mutants, one standard deviation below the mean, were identified using an HTP sequencing-based yeast replicative lifespan screen (SEBYL), with pooled cells subject to four rounds of biotin-based sorting to enrich old vs young cells, followed by sequencing to ID mutants and abundance 	Yu R, et al. (2021) PMID:33790287
GGC1	YDL198C	replicative lifespan: decreased 	systematic mutation set 	large-scale survey	null Allele: ggc1-Δ	S288C		Details: short-lived mutants, one standard deviation below the mean, were identified using an HTP sequencing-based yeast replicative lifespan screen (SEBYL), with pooled cells subject to four rounds of biotin-based sorting to enrich old vs young cells, followed by sequencing to ID mutants and abundance 	Yu R, et al. (2021) PMID:33790287
GIM4	YEL003W	replicative lifespan: decreased 	systematic mutation set 	large-scale survey	null Allele: gim4-Δ	S288C		Details: short-lived mutants, one standard deviation below the mean, were identified using an HTP sequencing-based yeast replicative lifespan screen (SEBYL), with pooled cells subject to four rounds of biotin-based sorting to enrich old vs young cells, followed by sequencing to ID mutants and abundance 	Yu R, et al. (2021) PMID:33790287
GIM5	YML094W	replicative lifespan: decreased 	systematic mutation set 	large-scale survey	null Allele: gim5-Δ	S288C		Details: short-lived mutants, one standard deviation below the mean, were identified using an HTP sequencing-based yeast replicative lifespan screen (SEBYL), with pooled cells subject to four rounds of biotin-based sorting to enrich old vs young cells, followed by sequencing to ID mutants and abundance 	Yu R, et al. (2021) PMID:33790287
GLO3	YER122C	replicative lifespan: decreased 	systematic mutation set 	large-scale survey	null Allele: glo3-Δ	S288C		Details: short-lived mutants, one standard deviation below the mean, were identified using an HTP sequencing-based yeast replicative lifespan screen (SEBYL), with pooled cells subject to four rounds of biotin-based sorting to enrich old vs young cells, followed by sequencing to ID mutants and abundance 	Yu R, et al. (2021) PMID:33790287
GLR1	YPL091W	replicative lifespan: decreased 	systematic mutation set 	large-scale survey	null Allele: glr1-Δ	S288C		Details: short-lived mutants, one standard deviation below the mean, were identified using an HTP sequencing-based yeast replicative lifespan screen (SEBYL), with pooled cells subject to four rounds of biotin-based sorting to enrich old vs young cells, followed by sequencing to ID mutants and abundance 	Yu R, et al. (2021) PMID:33790287
GPA2	YER020W	replicative lifespan: decreased 	systematic mutation set 	large-scale survey	null Allele: gpa2-Δ	S288C		Details: short-lived mutants, one standard deviation below the mean, were identified using an HTP sequencing-based yeast replicative lifespan screen (SEBYL), with pooled cells subject to four rounds of biotin-based sorting to enrich old vs young cells, followed by sequencing to ID mutants and abundance 	Yu R, et al. (2021) PMID:33790287
GRX5	YPL059W	replicative lifespan: decreased 	classical genetics	classical genetics	null Allele: grx5-Δ	W303			Zadrag R, et al. (2008) PMID:20021387
GTR1	YML121W	replicative lifespan: decreased 	classical genetics 	classical genetics	overexpression	S288C		Details: reduction in RLS relative to wt; counteracts the lifespan extending ability of corylin, a flavinoid from Psoralea corylifolia 	Wang TH, et al. (2022) PMID:35264584
GUT2	YIL155C	replicative lifespan: decreased 	classical genetics	classical genetics	null Allele: gut2-Δ	S288C		Media: glucose carbon source, 0.5% Details: calorie restriction-mediated lifespan extension is smaller than in wild type 	Easlon E, et al. (2008) PMID:18381895
HAC1	YFL031W	replicative lifespan: decreased 	systematic mutation set 	large-scale survey	null Allele: hac1-Δ	S288C		Details: short-lived mutants, one standard deviation below the mean, were identified using an HTP sequencing-based yeast replicative lifespan screen (SEBYL), with pooled cells subject to four rounds of biotin-based sorting to enrich old vs young cells, followed by sequencing to ID mutants and abundance 	Yu R, et al. (2021) PMID:33790287
HAP4	YKL109W	replicative lifespan: decreased 	classical genetics	classical genetics	null Allele: hap4-Δ	Other		Media: 2% glucose without nonessential amino acids 	Wang J, et al. (2010) PMID:20178842
HAP4	YKL109W	replicative lifespan: decreased 	classical genetics	classical genetics	null Allele: hap4-Δ	Other		Media: low glucose (0.1%) 	Wang J, et al. (2010) PMID:20178842
HCM1	YCR065W	replicative lifespan: decreased 	classical genetics	classical genetics	null Allele: hcm1-Δ	S288C			Maoz N, et al. (2015) PMID:24835838
HCM1	YCR065W	replicative lifespan: decreased 	systematic mutation set 	large-scale survey	null Allele: hcm1-Δ	S288C		Details: short-lived mutants, one standard deviation below the mean, were identified using an HTP sequencing-based yeast replicative lifespan screen (SEBYL), with pooled cells subject to four rounds of biotin-based sorting to enrich old vs young cells, followed by sequencing to ID mutants and abundance 	Yu R, et al. (2021) PMID:33790287
HES1	YOR237W	replicative lifespan: decreased 	classical genetics	classical genetics	overexpression	S288C			Gebre S, et al. (2012) PMID:22622083
HIR2	YOR038C	replicative lifespan: decreased 	systematic mutation set 	large-scale survey	null Allele: hir2-Δ	S288C		Details: short-lived mutants, one standard deviation below the mean, were identified using an HTP sequencing-based yeast replicative lifespan screen (SEBYL), with pooled cells subject to four rounds of biotin-based sorting to enrich old vs young cells, followed by sequencing to ID mutants and abundance 	Yu R, et al. (2021) PMID:33790287
HMF1	YER057C	replicative lifespan: decreased 	systematic mutation set 	large-scale survey	null Allele: hmf1-Δ	S288C		Details: short-lived mutants, one standard deviation below the mean, were identified using an HTP sequencing-based yeast replicative lifespan screen (SEBYL), with pooled cells subject to four rounds of biotin-based sorting to enrich old vs young cells, followed by sequencing to ID mutants and abundance 	Yu R, et al. (2021) PMID:33790287
HOR7	YMR251W-A	replicative lifespan: decreased 	classical genetics	classical genetics	null Allele: hor7-Δ	S288C		Treatment: caloric restriction by glucose reduction, 0.05% Details: dietary restriction-induced lifespan extension is decreased relative to wt 	Schleit J, et al. (2013) PMID:23837470
HSP104	YLL026W	replicative lifespan: decreased 	classical genetics	classical genetics	null Allele: hsp104-Δ	S288C		Treatment: caloric restriction by glucose reduction, 0.05% Details: dietary restriction-induced lifespan extension is decreased relative to wt 	Schleit J, et al. (2013) PMID:23837470
HSP104	YLL026W	replicative lifespan: decreased 	classical genetics 	classical genetics	null Allele: hsp104-Δ	S288C		Details: ~33 generations for null compared to ~55 for wild-type 	Erjavec N, et al. (2007) PMID:17908928
HSP12	YFL014W	replicative lifespan: decreased 	classical genetics	classical genetics	null Allele: hsp12-Δ	S288C		Treatment: caloric restriction by glucose reduction, 0.5% Details: dietary restriction-induced lifespan extension is abolished 	Herbert AP, et al. (2012) PMID:22848679
HXK2	YGL253W	replicative lifespan: decreased 	classical genetics	classical genetics	null Allele: hxk2-Δ	S288C		Treatment: caloric restriction by glucose reduction, 0.05% Details: dietary restriction-induced lifespan extension is decreased relative to wt 	Schleit J, et al. (2013) PMID:23837470
IDH2	YOR136W	replicative lifespan: decreased 	systematic mutation set 	large-scale survey	null Allele: idh2-Δ	S288C		Details: short-lived mutants, one standard deviation below the mean, were identified using an HTP sequencing-based yeast replicative lifespan screen (SEBYL), with pooled cells subject to four rounds of biotin-based sorting to enrich old vs young cells, followed by sequencing to ID mutants and abundance 	Yu R, et al. (2021) PMID:33790287
ILM1	YJR118C	replicative lifespan: decreased 	systematic mutation set 	large-scale survey	null Allele: ilm1-Δ	S288C		Details: short-lived mutants, one standard deviation below the mean, were identified using an HTP sequencing-based yeast replicative lifespan screen (SEBYL), with pooled cells subject to four rounds of biotin-based sorting to enrich old vs young cells, followed by sequencing to ID mutants and abundance 	Yu R, et al. (2021) PMID:33790287
INP1	YMR204C	replicative lifespan: decreased 	classical genetics 	classical genetics	null Allele: inp1-Δ	S288C			Kumar S, et al. (2018) PMID:29361529
INP52	YNL106C	replicative lifespan: decreased 	classical genetics	classical genetics	null Allele: inp52-Δ	S288C			Hacioglu E, et al. (2010) PMID:20934449
IRC15	YPL017C	replicative lifespan: decreased 	systematic mutation set 	large-scale survey	null Allele: irc15-Δ	S288C		Details: short-lived mutants, one standard deviation below the mean, were identified using an HTP sequencing-based yeast replicative lifespan screen (SEBYL), with pooled cells subject to four rounds of biotin-based sorting to enrich old vs young cells, followed by sequencing to ID mutants and abundance 	Yu R, et al. (2021) PMID:33790287
IRC25	YLR021W	replicative lifespan: decreased 	systematic mutation set 	large-scale survey	null Allele: irc25-Δ	S288C		Details: short-lived mutants, one standard deviation below the mean, were identified using an HTP sequencing-based yeast replicative lifespan screen (SEBYL), with pooled cells subject to four rounds of biotin-based sorting to enrich old vs young cells, followed by sequencing to ID mutants and abundance 	Yu R, et al. (2021) PMID:33790287
IRE1	YHR079C	replicative lifespan: decreased 	classical genetics	classical genetics	null Allele: ire1-Δ	S288C		Treatment: caloric restriction by glucose reduction, 0.05% Details: dietary restriction-induced lifespan extension is decreased relative to wt 	Schleit J, et al. (2013) PMID:23837470
IRE1	YHR079C	replicative lifespan: decreased 	systematic mutation set 	large-scale survey	null Allele: ire1-Δ	S288C		Details: short-lived mutants, one standard deviation below the mean, were identified using an HTP sequencing-based yeast replicative lifespan screen (SEBYL), with pooled cells subject to four rounds of biotin-based sorting to enrich old vs young cells, followed by sequencing to ID mutants and abundance 	Yu R, et al. (2021) PMID:33790287
ISC1	YER019W	replicative lifespan: decreased 	systematic mutation set 	large-scale survey	null Allele: isc1-Δ	S288C		Details: short-lived mutants, one standard deviation below the mean, were identified using an HTP sequencing-based yeast replicative lifespan screen (SEBYL), with pooled cells subject to four rounds of biotin-based sorting to enrich old vs young cells, followed by sequencing to ID mutants and abundance 	Yu R, et al. (2021) PMID:33790287
IST3	YIR005W	replicative lifespan: decreased 	systematic mutation set 	large-scale survey	null Allele: ist3-Δ	S288C		Details: short-lived mutants, one standard deviation below the mean, were identified using an HTP sequencing-based yeast replicative lifespan screen (SEBYL), with pooled cells subject to four rounds of biotin-based sorting to enrich old vs young cells, followed by sequencing to ID mutants and abundance 	Yu R, et al. (2021) PMID:33790287
JJJ3	YJR097W	replicative lifespan: decreased 	systematic mutation set 	large-scale survey	null Allele: jjj3-Δ	S288C		Details: short-lived mutants, one standard deviation below the mean, were identified using an HTP sequencing-based yeast replicative lifespan screen (SEBYL), with pooled cells subject to four rounds of biotin-based sorting to enrich old vs young cells, followed by sequencing to ID mutants and abundance 	Yu R, et al. (2021) PMID:33790287
JNM1	YMR294W	replicative lifespan: decreased 	systematic mutation set 	large-scale survey	null Allele: jnm1-Δ	S288C		Details: short-lived mutants, one standard deviation below the mean, were identified using an HTP sequencing-based yeast replicative lifespan screen (SEBYL), with pooled cells subject to four rounds of biotin-based sorting to enrich old vs young cells, followed by sequencing to ID mutants and abundance 	Yu R, et al. (2021) PMID:33790287
KAP123	YER110C	replicative lifespan: decreased 	systematic mutation set 	large-scale survey	null Allele: kap123-Δ	S288C		Details: short-lived mutants, one standard deviation below the mean, were identified using an HTP sequencing-based yeast replicative lifespan screen (SEBYL), with pooled cells subject to four rounds of biotin-based sorting to enrich old vs young cells, followed by sequencing to ID mutants and abundance 	Yu R, et al. (2021) PMID:33790287
KIN3	YAR018C	replicative lifespan: decreased 	systematic mutation set 	large-scale survey	null Allele: kin3-Δ	S288C		Details: short-lived mutants, one standard deviation below the mean, were identified using an HTP sequencing-based yeast replicative lifespan screen (SEBYL), with pooled cells subject to four rounds of biotin-based sorting to enrich old vs young cells, followed by sequencing to ID mutants and abundance 	Yu R, et al. (2021) PMID:33790287
KSP1	YHR082C	replicative lifespan: decreased 	classical genetics	classical genetics	null Allele: ksp1-Δ	S288C		Details: mean replicative lifespan decreased 21% relative to wild-type 	Zhao W, et al. (2017) PMID:28347693
KTI12	YKL110C	replicative lifespan: decreased 	systematic mutation set 	large-scale survey	null Allele: kti12-Δ	S288C		Details: short-lived mutants, one standard deviation below the mean, were identified using an HTP sequencing-based yeast replicative lifespan screen (SEBYL), with pooled cells subject to four rounds of biotin-based sorting to enrich old vs young cells, followed by sequencing to ID mutants and abundance 	Yu R, et al. (2021) PMID:33790287
LAG2	YOL025W	replicative lifespan: decreased 	classical genetics	classical genetics	null Allele: lag2-Δ	Other		Details: 50% decrease in mean and maximum lifespan relative to wild type cells 	Childress AM, et al. (1996) PMID:8760941
LAT1	YNL071W	replicative lifespan: decreased 	classical genetics	classical genetics	null Allele: lat1-Δ	S288C		Details: lifespan is not extended in response to calorie restriction 	Easlon E, et al. (2007) PMID:17200108
LCB4	YOR171C	replicative lifespan: decreased 	classical genetics	classical genetics	null Allele: lcb4-Δ	S288C		Treatment: caloric restriction Details: required for caloric restriction-triggered life-span extension 	Tang F, et al. (2008) PMID:18690010
LCB5	YLR260W	replicative lifespan: decreased 	systematic mutation set 	large-scale survey	null Allele: lcb5-Δ	S288C		Details: short-lived mutants, one standard deviation below the mean, were identified using an HTP sequencing-based yeast replicative lifespan screen (SEBYL), with pooled cells subject to four rounds of biotin-based sorting to enrich old vs young cells, followed by sequencing to ID mutants and abundance 	Yu R, et al. (2021) PMID:33790287
LDB7	YBL006C	replicative lifespan: decreased 	systematic mutation set 	large-scale survey	null Allele: ldb7-Δ	S288C		Details: short-lived mutants, one standard deviation below the mean, were identified using an HTP sequencing-based yeast replicative lifespan screen (SEBYL), with pooled cells subject to four rounds of biotin-based sorting to enrich old vs young cells, followed by sequencing to ID mutants and abundance 	Yu R, et al. (2021) PMID:33790287
LEA1	YPL213W	replicative lifespan: decreased 	systematic mutation set 	large-scale survey	null Allele: lea1-Δ	S288C		Details: short-lived mutants, one standard deviation below the mean, were identified using an HTP sequencing-based yeast replicative lifespan screen (SEBYL), with pooled cells subject to four rounds of biotin-based sorting to enrich old vs young cells, followed by sequencing to ID mutants and abundance 	Yu R, et al. (2021) PMID:33790287
LHS1	YKL073W	replicative lifespan: decreased 	systematic mutation set 	large-scale survey	null Allele: lhs1-Δ	S288C		Details: short-lived mutants, one standard deviation below the mean, were identified using an HTP sequencing-based yeast replicative lifespan screen (SEBYL), with pooled cells subject to four rounds of biotin-based sorting to enrich old vs young cells, followed by sequencing to ID mutants and abundance 	Yu R, et al. (2021) PMID:33790287
LIA1	YJR070C	replicative lifespan: decreased 	systematic mutation set 	large-scale survey	null Allele: lia1-Δ	S288C		Details: short-lived mutants, one standard deviation below the mean, were identified using an HTP sequencing-based yeast replicative lifespan screen (SEBYL), with pooled cells subject to four rounds of biotin-based sorting to enrich old vs young cells, followed by sequencing to ID mutants and abundance 	Yu R, et al. (2021) PMID:33790287
LMO1	YLL007C	replicative lifespan: decreased 	systematic mutation set 	large-scale survey	null Allele: lmo1-Δ	S288C		Details: short-lived mutants, one standard deviation below the mean, were identified using an HTP sequencing-based yeast replicative lifespan screen (SEBYL), with pooled cells subject to four rounds of biotin-based sorting to enrich old vs young cells, followed by sequencing to ID mutants and abundance 	Yu R, et al. (2021) PMID:33790287
LOC1	YFR001W	replicative lifespan: decreased 	systematic mutation set 	large-scale survey	null Allele: loc1-Δ	S288C		Details: short-lived mutants, one standard deviation below the mean, were identified using an HTP sequencing-based yeast replicative lifespan screen (SEBYL), with pooled cells subject to four rounds of biotin-based sorting to enrich old vs young cells, followed by sequencing to ID mutants and abundance 	Yu R, et al. (2021) PMID:33790287
LRP1	YHR081W	replicative lifespan: decreased 	systematic mutation set 	large-scale survey	null Allele: lrp1-Δ	S288C		Details: short-lived mutants, one standard deviation below the mean, were identified using an HTP sequencing-based yeast replicative lifespan screen (SEBYL), with pooled cells subject to four rounds of biotin-based sorting to enrich old vs young cells, followed by sequencing to ID mutants and abundance 	Yu R, et al. (2021) PMID:33790287
LSM6	YDR378C	replicative lifespan: decreased 	systematic mutation set 	large-scale survey	null Allele: lsm6-Δ	S288C		Details: short-lived mutants, one standard deviation below the mean, were identified using an HTP sequencing-based yeast replicative lifespan screen (SEBYL), with pooled cells subject to four rounds of biotin-based sorting to enrich old vs young cells, followed by sequencing to ID mutants and abundance 	Yu R, et al. (2021) PMID:33790287
LSM7	YNL147W	replicative lifespan: decreased 	systematic mutation set 	large-scale survey	null Allele: lsm7-Δ	S288C		Details: short-lived mutants, one standard deviation below the mean, were identified using an HTP sequencing-based yeast replicative lifespan screen (SEBYL), with pooled cells subject to four rounds of biotin-based sorting to enrich old vs young cells, followed by sequencing to ID mutants and abundance 	Yu R, et al. (2021) PMID:33790287
MAK31	YCR020C-A	replicative lifespan: decreased 	systematic mutation set 	large-scale survey	null Allele: mak31-Δ	S288C		Details: short-lived mutants, one standard deviation below the mean, were identified using an HTP sequencing-based yeast replicative lifespan screen (SEBYL), with pooled cells subject to four rounds of biotin-based sorting to enrich old vs young cells, followed by sequencing to ID mutants and abundance 	Yu R, et al. (2021) PMID:33790287
MAP1	YLR244C	replicative lifespan: decreased 	classical genetics	classical genetics	null Allele: map1-Δ	S288C			Hacioglu E, et al. (2010) PMID:20934449
MCM1	YMR043W	replicative lifespan: decreased 	heterozygous diploid	classical genetics	null Allele: mcm1-Δ	S288C		Details: moderately short lifespan 	Kamei Y, et al. (2015) PMID:26022127
MDH1	YKL085W	replicative lifespan: decreased 	classical genetics	classical genetics	null Allele: mdh1-Δ	S288C		Media: glucose carbon source, 0.5% Details: calorie restriction-mediated lifespan extension is smaller than in wild type 	Easlon E, et al. (2008) PMID:18381895
MDM1	YML104C	replicative lifespan: decreased 	classical genetics	classical genetics	null Allele: mdm1-Δ	S288C		Treatment: caloric restriction by glucose reduction, 0.05% Details: dietary restriction-induced lifespan extension is decreased relative to wt 	Schleit J, et al. (2013) PMID:23837470
MDM35	YKL053C-A	replicative lifespan: decreased 	systematic mutation set 	large-scale survey	null Allele: mdm35-Δ	S288C		Details: short-lived mutants, one standard deviation below the mean, were identified using an HTP sequencing-based yeast replicative lifespan screen (SEBYL), with pooled cells subject to four rounds of biotin-based sorting to enrich old vs young cells, followed by sequencing to ID mutants and abundance 	Yu R, et al. (2021) PMID:33790287
MED2	YDL005C	replicative lifespan: decreased 	classical genetics 	classical genetics	null Allele: med2-Δ	S288C		Details: RLS is 16% shorter in the null mutant relative to wt 	Zhao W, et al. (2020) PMID:33045248
MED2	YDL005C	replicative lifespan: decreased 	systematic mutation set 	large-scale survey	null Allele: med2-Δ	S288C		Details: short-lived mutants, one standard deviation below the mean, were identified using an HTP sequencing-based yeast replicative lifespan screen (SEBYL), with pooled cells subject to four rounds of biotin-based sorting to enrich old vs young cells, followed by sequencing to ID mutants and abundance 	Yu R, et al. (2021) PMID:33790287
MET12	YPL023C	replicative lifespan: decreased 	systematic mutation set 	large-scale survey	null Allele: met12-Δ	S288C		Details: short-lived mutants, one standard deviation below the mean, were identified using an HTP sequencing-based yeast replicative lifespan screen (SEBYL), with pooled cells subject to four rounds of biotin-based sorting to enrich old vs young cells, followed by sequencing to ID mutants and abundance 	Yu R, et al. (2021) PMID:33790287
MET18	YIL128W	replicative lifespan: decreased 	heterozygous diploid 	classical genetics	null Allele: met18-Δ	S288C			Chen YQ, et al. (2017) PMID:28828388
MFB1	YDR219C	replicative lifespan: decreased 	classical genetics 	classical genetics	null Allele: mfb1-Δ	S288C		Details: reduction in RLS from 26.7 ± 7.51 generations in wt cells to 12.1 ± 5.01 generations in the null mutant 	Yang EJ, et al. (2022) PMID:35281729
MFB1	YDR219C	replicative lifespan: decreased 	classical genetics 	classical genetics	null Allele: mfb1-Δ	S288C		Details: reduction in RLS from 26.7 generations in wt cells to 12.1 generations in the null mutant 	Pernice WM, et al. (2016) PMID:26839174
MFB1	YDR219C	replicative lifespan: decreased 	systematic mutation set 	large-scale survey	null Allele: mfb1-Δ	S288C		Details: short-lived mutants, one standard deviation below the mean, were identified using an HTP sequencing-based yeast replicative lifespan screen (SEBYL), with pooled cells subject to four rounds of biotin-based sorting to enrich old vs young cells, followed by sequencing to ID mutants and abundance 	Yu R, et al. (2021) PMID:33790287
MFT1	YML062C	replicative lifespan: decreased 	systematic mutation set 	large-scale survey	null Allele: mft1-Δ	S288C		Details: short-lived mutants, one standard deviation below the mean, were identified using an HTP sequencing-based yeast replicative lifespan screen (SEBYL), with pooled cells subject to four rounds of biotin-based sorting to enrich old vs young cells, followed by sequencing to ID mutants and abundance 	Yu R, et al. (2021) PMID:33790287
MGA2	YIR033W	replicative lifespan: decreased 	systematic mutation set 	large-scale survey	null Allele: mga2-Δ	S288C		Details: short-lived mutants, one standard deviation below the mean, were identified using an HTP sequencing-based yeast replicative lifespan screen (SEBYL), with pooled cells subject to four rounds of biotin-based sorting to enrich old vs young cells, followed by sequencing to ID mutants and abundance 	Yu R, et al. (2021) PMID:33790287
MGR2	YPL098C	replicative lifespan: decreased 	systematic mutation set 	large-scale survey	null Allele: mgr2-Δ	S288C		Details: short-lived mutants, one standard deviation below the mean, were identified using an HTP sequencing-based yeast replicative lifespan screen (SEBYL), with pooled cells subject to four rounds of biotin-based sorting to enrich old vs young cells, followed by sequencing to ID mutants and abundance 	Yu R, et al. (2021) PMID:33790287
MIC60	YKR016W	replicative lifespan: decreased 	systematic mutation set 	large-scale survey	null Allele: mic60-Δ	S288C		Details: short-lived mutants, one standard deviation below the mean, were identified using an HTP sequencing-based yeast replicative lifespan screen (SEBYL), with pooled cells subject to four rounds of biotin-based sorting to enrich old vs young cells, followed by sequencing to ID mutants and abundance 	Yu R, et al. (2021) PMID:33790287
MIG1	YGL035C	replicative lifespan: decreased 	classical genetics	classical genetics	overexpression	S288C			Harkness TA, et al. (2004) PMID:15514051
MIG1	YGL035C	replicative lifespan: decreased 	classical genetics	classical genetics	null Allele: mig1-Δ	S288C			Harkness TA, et al. (2004) PMID:15514051
MIH1	YMR036C	replicative lifespan: decreased 	systematic mutation set 	large-scale survey	null Allele: mih1-Δ	S288C		Details: short-lived mutants, one standard deviation below the mean, were identified using an HTP sequencing-based yeast replicative lifespan screen (SEBYL), with pooled cells subject to four rounds of biotin-based sorting to enrich old vs young cells, followed by sequencing to ID mutants and abundance 	Yu R, et al. (2021) PMID:33790287
MMR1	YLR190W	replicative lifespan: decreased 	classical genetics 	classical genetics	null Allele: mmr1-Δ	S288C		Details: reduction in RLS from 26.7 generations in wt cells to 17.8 generations in the null mutant 	Pernice WM, et al. (2016) PMID:26839174
MMS1	YPR164W	replicative lifespan: decreased 	systematic mutation set 	large-scale survey	null Allele: mms1-Δ	S288C		Details: short-lived mutants, one standard deviation below the mean, were identified using an HTP sequencing-based yeast replicative lifespan screen (SEBYL), with pooled cells subject to four rounds of biotin-based sorting to enrich old vs young cells, followed by sequencing to ID mutants and abundance 	Yu R, et al. (2021) PMID:33790287
MMS22	YLR320W	replicative lifespan: decreased 	classical genetics	classical genetics	null Allele: mms22-Δ	S288C		Treatment: caloric restriction by glucose reduction, 0.05% Details: dietary restriction-induced lifespan extension is decreased relative to wt 	Schleit J, et al. (2013) PMID:23837470
MMS4	YBR098W	replicative lifespan: decreased 	systematic mutation set 	large-scale survey	null Allele: mms4-Δ	S288C		Details: short-lived mutants, one standard deviation below the mean, were identified using an HTP sequencing-based yeast replicative lifespan screen (SEBYL), with pooled cells subject to four rounds of biotin-based sorting to enrich old vs young cells, followed by sequencing to ID mutants and abundance 	Yu R, et al. (2021) PMID:33790287
MNN9	YPL050C	replicative lifespan: decreased 	classical genetics 	classical genetics	null Allele: mnn9-Δ	S288C		Details: almost a 4-fold reduction in replicative lifespan 	Molon M, et al. (2018) PMID:29189912
MOD5	YOR274W	replicative lifespan: decreased 	systematic mutation set 	large-scale survey	null Allele: mod5-Δ	S288C		Details: short-lived mutants, one standard deviation below the mean, were identified using an HTP sequencing-based yeast replicative lifespan screen (SEBYL), with pooled cells subject to four rounds of biotin-based sorting to enrich old vs young cells, followed by sequencing to ID mutants and abundance 	Yu R, et al. (2021) PMID:33790287
MOG1	YJR074W	replicative lifespan: decreased 	systematic mutation set 	large-scale survey	null Allele: mog1-Δ	S288C		Details: short-lived mutants, one standard deviation below the mean, were identified using an HTP sequencing-based yeast replicative lifespan screen (SEBYL), with pooled cells subject to four rounds of biotin-based sorting to enrich old vs young cells, followed by sequencing to ID mutants and abundance 	Yu R, et al. (2021) PMID:33790287
MPD2	YOL088C	replicative lifespan: decreased 	classical genetics	classical genetics	null Allele: mpd2-Δ	S288C			Hacioglu E, et al. (2010) PMID:20934449
MPT5	YGL178W	replicative lifespan: decreased 	classical genetics	classical genetics	null Allele: mpt5-Δ	Other			Kennedy BK, et al. (1997) PMID:9150138
MRC1	YCL061C	replicative lifespan: decreased 	systematic mutation set 	large-scale survey	null Allele: mrc1-Δ	S288C		Details: short-lived mutants, one standard deviation below the mean, were identified using an HTP sequencing-based yeast replicative lifespan screen (SEBYL), with pooled cells subject to four rounds of biotin-based sorting to enrich old vs young cells, followed by sequencing to ID mutants and abundance 	Yu R, et al. (2021) PMID:33790287
MRN1	YPL184C	replicative lifespan: decreased 	systematic mutation set 	large-scale survey	null Allele: mrn1-Δ	S288C		Details: short-lived mutants, one standard deviation below the mean, were identified using an HTP sequencing-based yeast replicative lifespan screen (SEBYL), with pooled cells subject to four rounds of biotin-based sorting to enrich old vs young cells, followed by sequencing to ID mutants and abundance 	Yu R, et al. (2021) PMID:33790287
MSH2	YOL090W	replicative lifespan: decreased 	systematic mutation set 	large-scale survey	null Allele: msh2-Δ	S288C		Details: short-lived mutants, one standard deviation below the mean, were identified using an HTP sequencing-based yeast replicative lifespan screen (SEBYL), with pooled cells subject to four rounds of biotin-based sorting to enrich old vs young cells, followed by sequencing to ID mutants and abundance 	Yu R, et al. (2021) PMID:33790287
MSN2	YMR037C	replicative lifespan: decreased 	classical genetics 	classical genetics	overexpression	S288C		Details: markedly shortened lifespan relative to wt, which can be almost fully restored by deletion of PUT1 	Mukai Y, et al. (2019) PMID:31646149
MSN5	YDR335W	replicative lifespan: decreased 	systematic mutation set 	large-scale survey	null Allele: msn5-Δ	S288C		Details: short-lived mutants, one standard deviation below the mean, were identified using an HTP sequencing-based yeast replicative lifespan screen (SEBYL), with pooled cells subject to four rounds of biotin-based sorting to enrich old vs young cells, followed by sequencing to ID mutants and abundance 	Yu R, et al. (2021) PMID:33790287
MSY1	YPL097W	replicative lifespan: decreased 	systematic mutation set 	large-scale survey	null Allele: msy1-Δ	S288C		Details: short-lived mutants, one standard deviation below the mean, were identified using an HTP sequencing-based yeast replicative lifespan screen (SEBYL), with pooled cells subject to four rounds of biotin-based sorting to enrich old vs young cells, followed by sequencing to ID mutants and abundance 	Yu R, et al. (2021) PMID:33790287
MUP3	YHL036W	replicative lifespan: decreased 	classical genetics 	classical genetics	null Allele: mup3-Δ	S288C			Tai A, et al. (2017) PMID:28495531
MUS81	YDR386W	replicative lifespan: decreased 	systematic mutation set 	large-scale survey	null Allele: mus81-Δ	S288C		Details: short-lived mutants, one standard deviation below the mean, were identified using an HTP sequencing-based yeast replicative lifespan screen (SEBYL), with pooled cells subject to four rounds of biotin-based sorting to enrich old vs young cells, followed by sequencing to ID mutants and abundance 	Yu R, et al. (2021) PMID:33790287
MXR1	YER042W	replicative lifespan: decreased 	classical genetics	classical genetics	null Allele: mxr1-Δ	Other		Media: glucose or lactate as carbon source 	Kaya A, et al. (2010) PMID:20799725
MXR1	YER042W	replicative lifespan: decreased 	classical genetics	classical genetics	null Allele: mxr1-Δ	S288C		Details: 26% reduction in replicative lifespan 	Koc A, et al. (2004) PMID:15141092
MXR1	YER042W	replicative lifespan: decreased 	classical genetics	classical genetics	null Allele: mxr1-Δ	S288C			Le DT, et al. (2009) PMID:19049972
NAR1	YNL240C	replicative lifespan: decreased 	heterozygous diploid	classical genetics	null Allele: nar1-Δ	S288C		Details: mean replicative lifespan is 18% shorter than in wt on rich medium 	Zhao W, et al. (2014) PMID:24486555
NBA1	YOL070C	replicative lifespan: decreased 	classical genetics	classical genetics	null Allele: nba1-Δ	Other		Details: lifespan is significantly shorter than wt cells 	Meitinger F, et al. (2014) PMID:25416945
NCR1	YPL006W	replicative lifespan: decreased 	systematic mutation set 	large-scale survey	null Allele: ncr1-Δ	S288C		Details: short-lived mutants, one standard deviation below the mean, were identified using an HTP sequencing-based yeast replicative lifespan screen (SEBYL), with pooled cells subject to four rounds of biotin-based sorting to enrich old vs young cells, followed by sequencing to ID mutants and abundance 	Yu R, et al. (2021) PMID:33790287
NCS6	YGL211W	replicative lifespan: decreased 	systematic mutation set 	large-scale survey	null Allele: ncs6-Δ	S288C		Details: short-lived mutants, one standard deviation below the mean, were identified using an HTP sequencing-based yeast replicative lifespan screen (SEBYL), with pooled cells subject to four rounds of biotin-based sorting to enrich old vs young cells, followed by sequencing to ID mutants and abundance 	Yu R, et al. (2021) PMID:33790287
NDE1	YMR145C	replicative lifespan: decreased 	classical genetics	classical genetics	null Allele: nde1-Δ	S288C		Treatment: caloric restriction by glucose reduction, 0.05% Details: dietary restriction-induced lifespan extension is decreased relative to wt 	Schleit J, et al. (2013) PMID:23837470
NDI1	YML120C	replicative lifespan: decreased 	classical genetics	classical genetics	null Allele: ndi1-Δ	S288C		Treatment: caloric restriction by glucose reduction, 0.05% Details: dietary restriction-induced lifespan extension is decreased relative to wt 	Schleit J, et al. (2013) PMID:23837470
NEM1	YHR004C	replicative lifespan: decreased 	systematic mutation set 	large-scale survey	null Allele: nem1-Δ	S288C		Details: short-lived mutants, one standard deviation below the mean, were identified using an HTP sequencing-based yeast replicative lifespan screen (SEBYL), with pooled cells subject to four rounds of biotin-based sorting to enrich old vs young cells, followed by sequencing to ID mutants and abundance 	Yu R, et al. (2021) PMID:33790287
NGG1	YDR176W	replicative lifespan: decreased 	classical genetics 	classical genetics	null Allele: ngg1-Δ	S288C	10 μg/ml (-)-epigallocatechin 3-gallate	Details: effects of epigallocatechin gallate (EGCG), a catechin and histone acetyltransferase inhibitor, on lifespan extension are abolished  	Huang B, et al. (2020) PMID:32157780
NGG1	YDR176W	replicative lifespan: decreased 	classical genetics 	classical genetics	null Allele: ngg1-Δ	S288C		Media: glucose-containing media, 0.5 % Details: caloric restriction-mediated lifespan extension present in wt cells is absent in the null mutant  	Huang B, et al. (2020) PMID:32157780
NIP100	YPL174C	replicative lifespan: decreased 	systematic mutation set 	large-scale survey	null Allele: nip100-Δ	S288C		Details: short-lived mutants, one standard deviation below the mean, were identified using an HTP sequencing-based yeast replicative lifespan screen (SEBYL), with pooled cells subject to four rounds of biotin-based sorting to enrich old vs young cells, followed by sequencing to ID mutants and abundance 	Yu R, et al. (2021) PMID:33790287
NMT1	YLR195C	replicative lifespan: decreased 	classical genetics	classical genetics	conditional Allele: nmt1-451D G451D; mutant enzyme has reduced affinity for myristoyl-coA	S288C		Temperature: permissive temperature, 24 °C Details: mean lifespan is 14.5 generations as compared to 27.5 in the wild type; cells display typical characteristics of accelerated aging late in their lifespan, such as sterility, nucleolar fragmentation, abnormal distribution of the Sir complex, and accumulation of extrachromosomal rDNA circles 	Ashrafi K, et al. (2000) PMID:10921902
NPR2	YEL062W	replicative lifespan: decreased 	systematic mutation set 	large-scale survey	null Allele: npr2-Δ	S288C		Details: short-lived mutants, one standard deviation below the mean, were identified using an HTP sequencing-based yeast replicative lifespan screen (SEBYL), with pooled cells subject to four rounds of biotin-based sorting to enrich old vs young cells, followed by sequencing to ID mutants and abundance 	Yu R, et al. (2021) PMID:33790287
NSI1	YDR026C	replicative lifespan: decreased 	classical genetics	classical genetics	null Allele: nsi1-Δ	S288C			Ha CW, et al. (2012) PMID:22362748
NST1	YNL091W	replicative lifespan: decreased 	systematic mutation set 	large-scale survey	null Allele: nst1-Δ	S288C		Details: short-lived mutants, one standard deviation below the mean, were identified using an HTP sequencing-based yeast replicative lifespan screen (SEBYL), with pooled cells subject to four rounds of biotin-based sorting to enrich old vs young cells, followed by sequencing to ID mutants and abundance 	Yu R, et al. (2021) PMID:33790287
NUP170	YBL079W	replicative lifespan: decreased 	systematic mutation set 	large-scale survey	null Allele: nup170-Δ	S288C		Details: short-lived mutants, one standard deviation below the mean, were identified using an HTP sequencing-based yeast replicative lifespan screen (SEBYL), with pooled cells subject to four rounds of biotin-based sorting to enrich old vs young cells, followed by sequencing to ID mutants and abundance 	Yu R, et al. (2021) PMID:33790287
NUP60	YAR002W	replicative lifespan: decreased 	classical genetics 	classical genetics	null Allele: nup60-Δ	S288C			Meinema AC, et al. (2022) PMID:35373738
NYV1	YLR093C	replicative lifespan: decreased 	classical genetics	classical genetics	null Allele: nyv1-Δ	S288C		Treatment: caloric restriction Details: required for caloric restriction-triggered life-span extension 	Tang F, et al. (2008) PMID:18690010
OPI7	YDR360W	replicative lifespan: decreased 	systematic mutation set 	large-scale survey	null Allele: opi7-Δ	S288C		Details: short-lived mutants, one standard deviation below the mean, were identified using an HTP sequencing-based yeast replicative lifespan screen (SEBYL), with pooled cells subject to four rounds of biotin-based sorting to enrich old vs young cells, followed by sequencing to ID mutants and abundance 	Yu R, et al. (2021) PMID:33790287
ORM2	YLR350W	replicative lifespan: decreased 	systematic mutation set 	large-scale survey	null Allele: orm2-Δ	S288C		Details: short-lived mutants, one standard deviation below the mean, were identified using an HTP sequencing-based yeast replicative lifespan screen (SEBYL), with pooled cells subject to four rounds of biotin-based sorting to enrich old vs young cells, followed by sequencing to ID mutants and abundance 	Yu R, et al. (2021) PMID:33790287
PAC10	YGR078C	replicative lifespan: decreased 	systematic mutation set 	large-scale survey	null Allele: pac10-Δ	S288C		Details: short-lived mutants, one standard deviation below the mean, were identified using an HTP sequencing-based yeast replicative lifespan screen (SEBYL), with pooled cells subject to four rounds of biotin-based sorting to enrich old vs young cells, followed by sequencing to ID mutants and abundance 	Yu R, et al. (2021) PMID:33790287
PAM17	YKR065C	replicative lifespan: decreased 	systematic mutation set 	large-scale survey	null Allele: pam17-Δ	S288C		Details: short-lived mutants, one standard deviation below the mean, were identified using an HTP sequencing-based yeast replicative lifespan screen (SEBYL), with pooled cells subject to four rounds of biotin-based sorting to enrich old vs young cells, followed by sequencing to ID mutants and abundance 	Yu R, et al. (2021) PMID:33790287
PAP2	YOL115W	replicative lifespan: decreased 	systematic mutation set 	large-scale survey	null Allele: pap2-Δ	S288C		Details: short-lived mutants, one standard deviation below the mean, were identified using an HTP sequencing-based yeast replicative lifespan screen (SEBYL), with pooled cells subject to four rounds of biotin-based sorting to enrich old vs young cells, followed by sequencing to ID mutants and abundance 	Yu R, et al. (2021) PMID:33790287
PAT1	YCR077C	replicative lifespan: decreased 	classical genetics	classical genetics	null Allele: pat1-Δ	S288C		Details: shortened by 20- 30% compared with wild-type 	Duan R, et al. (2013) PMID:24358479
PCK1	YKR097W	replicative lifespan: decreased 	classical genetics 	classical genetics	null Allele: pck1-Δ	S288C		Details: deficiency shortened the RLS of the null mutant by approximately 16%; restored by deletion of PFK1 	Yuan Y, et al. (2020) PMID:32104690
PCL5	YHR071W	replicative lifespan: decreased 	classical genetics 	classical genetics	null Allele: pcl5-Δ	S288C		Media: high phosphate, 11 mM Details: moderately short lived under high Pi conditions; significantly shorter in pho80 double null 	Nakajima T, et al. (2020) PMID:31797348
PDE2	YOR360C	replicative lifespan: decreased 	classical genetics	classical genetics	null Allele: pde2-Δ	Other			Lin SJ, et al. (2000) PMID:11000115
PER1	YCR044C	replicative lifespan: decreased 	systematic mutation set 	large-scale survey	null Allele: per1-Δ	S288C		Details: short-lived mutants, one standard deviation below the mean, were identified using an HTP sequencing-based yeast replicative lifespan screen (SEBYL), with pooled cells subject to four rounds of biotin-based sorting to enrich old vs young cells, followed by sequencing to ID mutants and abundance 	Yu R, et al. (2021) PMID:33790287
PET117	YER058W	replicative lifespan: decreased 	systematic mutation set 	large-scale survey	null Allele: pet117-Δ	S288C		Details: short-lived mutants, one standard deviation below the mean, were identified using an HTP sequencing-based yeast replicative lifespan screen (SEBYL), with pooled cells subject to four rounds of biotin-based sorting to enrich old vs young cells, followed by sequencing to ID mutants and abundance 	Yu R, et al. (2021) PMID:33790287
PEX13	YLR191W	replicative lifespan: decreased 	classical genetics	classical genetics	null Allele: pex13-Δ	S288C		Treatment: caloric restriction by glucose reduction, 0.05% Details: dietary restriction-induced lifespan extension is decreased relative to wt 	Schleit J, et al. (2013) PMID:23837470
PEX19	YDL065C	replicative lifespan: decreased 	classical genetics	classical genetics	null Allele: pex19-Δ	S288C		Treatment: caloric restriction by glucose reduction, 0.05% Details: dietary restriction-induced lifespan extension is decreased relative to wt 	Schleit J, et al. (2013) PMID:23837470
PEX28	YHR150W	replicative lifespan: decreased 	systematic mutation set 	large-scale survey	null Allele: pex28-Δ	S288C		Details: short-lived mutants, one standard deviation below the mean, were identified using an HTP sequencing-based yeast replicative lifespan screen (SEBYL), with pooled cells subject to four rounds of biotin-based sorting to enrich old vs young cells, followed by sequencing to ID mutants and abundance 	Yu R, et al. (2021) PMID:33790287
PEX3	YDR329C	replicative lifespan: decreased 	classical genetics 	classical genetics	null Allele: pex3-Δ	S288C			Kumar S, et al. (2018) PMID:29361529
PFA4	YOL003C	replicative lifespan: decreased 	systematic mutation set 	large-scale survey	null Allele: pfa4-Δ	S288C		Details: short-lived mutants, one standard deviation below the mean, were identified using an HTP sequencing-based yeast replicative lifespan screen (SEBYL), with pooled cells subject to four rounds of biotin-based sorting to enrich old vs young cells, followed by sequencing to ID mutants and abundance 	Yu R, et al. (2021) PMID:33790287
PFD1	YJL179W	replicative lifespan: decreased 	systematic mutation set 	large-scale survey	null Allele: pfd1-Δ	S288C		Details: short-lived mutants, one standard deviation below the mean, were identified using an HTP sequencing-based yeast replicative lifespan screen (SEBYL), with pooled cells subject to four rounds of biotin-based sorting to enrich old vs young cells, followed by sequencing to ID mutants and abundance 	Yu R, et al. (2021) PMID:33790287
PGD1	YGL025C	replicative lifespan: decreased 	systematic mutation set 	large-scale survey	null Allele: pgd1-Δ	S288C		Details: short-lived mutants, one standard deviation below the mean, were identified using an HTP sequencing-based yeast replicative lifespan screen (SEBYL), with pooled cells subject to four rounds of biotin-based sorting to enrich old vs young cells, followed by sequencing to ID mutants and abundance 	Yu R, et al. (2021) PMID:33790287
PHB1	YGR132C	replicative lifespan: decreased 	classical genetics	classical genetics	null Allele: phb1-Δ	Other		Details: strain lacks mitochondrial genome 	Kirchman PA, et al. (2003) PMID:14739993
PHB1	YGR132C	replicative lifespan: decreased 	systematic mutation set 	large-scale survey	null Allele: phb1-Δ	S288C		Details: short-lived mutants, one standard deviation below the mean, were identified using an HTP sequencing-based yeast replicative lifespan screen (SEBYL), with pooled cells subject to four rounds of biotin-based sorting to enrich old vs young cells, followed by sequencing to ID mutants and abundance 	Yu R, et al. (2021) PMID:33790287
PHB2	YGR231C	replicative lifespan: decreased 	classical genetics	classical genetics	null Allele: phb2-Δ	Other		Details: strain lacks the mitochondrial genome 	Kirchman PA, et al. (2003) PMID:14739993
PHB2	YGR231C	replicative lifespan: decreased 	systematic mutation set 	large-scale survey	null Allele: phb2-Δ	S288C		Details: short-lived mutants, one standard deviation below the mean, were identified using an HTP sequencing-based yeast replicative lifespan screen (SEBYL), with pooled cells subject to four rounds of biotin-based sorting to enrich old vs young cells, followed by sequencing to ID mutants and abundance 	Yu R, et al. (2021) PMID:33790287
PHD1	YKL043W	replicative lifespan: decreased 	systematic mutation set 	large-scale survey	null Allele: phd1-Δ	S288C		Details: short-lived mutants, one standard deviation below the mean, were identified using an HTP sequencing-based yeast replicative lifespan screen (SEBYL), with pooled cells subject to four rounds of biotin-based sorting to enrich old vs young cells, followed by sequencing to ID mutants and abundance 	Yu R, et al. (2021) PMID:33790287
PHO23	YNL097C	replicative lifespan: decreased 	systematic mutation set 	large-scale survey	null Allele: pho23-Δ	S288C		Details: short-lived mutants, one standard deviation below the mean, were identified using an HTP sequencing-based yeast replicative lifespan screen (SEBYL), with pooled cells subject to four rounds of biotin-based sorting to enrich old vs young cells, followed by sequencing to ID mutants and abundance 	Yu R, et al. (2021) PMID:33790287
PHO4	YFR034C	replicative lifespan: decreased 	classical genetics 	classical genetics	null Allele: pho4-Δ	S288C		Media: low phosphate, 0.22 mM Details: slight but significant decrease in lifespan under low Pi conditions 	Nakajima T, et al. (2020) PMID:31797348
PHO80	YOL001W	replicative lifespan: decreased 	classical genetics 	classical genetics	null Allele: pho80-Δ	S288C		Media: high phosphate, 11 mM Details: drastically shortened lifespan under high Pi conditions; partially suppressed (80% of wt) by PHO4 deletion 	Nakajima T, et al. (2020) PMID:31797348
PHO80	YOL001W	replicative lifespan: decreased 	classical genetics 	classical genetics	null Allele: pho80-Δ	S288C		Media: low phosphate, 0.22 mM Details: drastically shortened lifespan under low Pi conditions; partially suppressed (70% of wt) by PHO4 deletion 	Nakajima T, et al. (2020) PMID:31797348
PHO84	YML123C	replicative lifespan: decreased 	classical genetics 	classical genetics	null Allele: pho84-Δ	S288C			Jiang JC, et al. (2016) PMID:27474729
PHO85	YPL031C	replicative lifespan: decreased 	classical genetics 	classical genetics	null Allele: pho85-Δ	S288C		Media: high phosphate, 11 mM Details: drastically shortened lifespan under high Pi conditions; partially suppressed (80% of wt) by PHO4 deletion 	Nakajima T, et al. (2020) PMID:31797348
PHO85	YPL031C	replicative lifespan: decreased 	classical genetics 	classical genetics	null Allele: pho85-Δ	S288C		Media: low phosphate, 0.22 mM Details: drastically shortened lifespan under low Pi conditions; partially suppressed (70% of wt) by PHO4 deletion 	Nakajima T, et al. (2020) PMID:31797348
PIM1	YBL022C	replicative lifespan: decreased 	systematic mutation set 	large-scale survey	null Allele: pim1-Δ	S288C		Details: short-lived mutants, one standard deviation below the mean, were identified using an HTP sequencing-based yeast replicative lifespan screen (SEBYL), with pooled cells subject to four rounds of biotin-based sorting to enrich old vs young cells, followed by sequencing to ID mutants and abundance 	Yu R, et al. (2021) PMID:33790287
PKH2	YOL100W	replicative lifespan: decreased 	systematic mutation set 	large-scale survey	null Allele: pkh2-Δ	S288C		Details: short-lived mutants, one standard deviation below the mean, were identified using an HTP sequencing-based yeast replicative lifespan screen (SEBYL), with pooled cells subject to four rounds of biotin-based sorting to enrich old vs young cells, followed by sequencing to ID mutants and abundance 	Yu R, et al. (2021) PMID:33790287
PMR1	YGL167C	replicative lifespan: decreased 	classical genetics	classical genetics	null Allele: pmr1-Δ	Other		Details: severe reduction in mean replicative lifespan of 64% 	Tsubakiyama R, et al. (2011) PMID:21712379
PMR1	YGL167C	replicative lifespan: decreased 	classical genetics	classical genetics	null Allele: pmr1-Δ	S288C		Treatment: caloric restriction by glucose reduction, 0.05% Details: dietary restriction-induced lifespan extension is decreased relative to wt 	Schleit J, et al. (2013) PMID:23837470
PMS1	YNL082W	replicative lifespan: decreased 	systematic mutation set 	large-scale survey	null Allele: pms1-Δ	S288C		Details: short-lived mutants, one standard deviation below the mean, were identified using an HTP sequencing-based yeast replicative lifespan screen (SEBYL), with pooled cells subject to four rounds of biotin-based sorting to enrich old vs young cells, followed by sequencing to ID mutants and abundance 	Yu R, et al. (2021) PMID:33790287
PMT1	YDL095W	replicative lifespan: decreased 	classical genetics	classical genetics	overexpression	S288C		Details: mean replicative lifespan decreased by 17% 	Cui HJ, et al. (2015) PMID:25936926
PMT2	YAL023C	replicative lifespan: decreased 	systematic mutation set 	large-scale survey	null Allele: pmt2-Δ	S288C		Details: short-lived mutants, one standard deviation below the mean, were identified using an HTP sequencing-based yeast replicative lifespan screen (SEBYL), with pooled cells subject to four rounds of biotin-based sorting to enrich old vs young cells, followed by sequencing to ID mutants and abundance 	Yu R, et al. (2021) PMID:33790287
PNC1	YGL037C	replicative lifespan: decreased 	classical genetics 	classical genetics	null Allele: pnc1-Δ	W303	0.5%  glucose		Anderson RM, et al. (2003) PMID:12736687
PNC1	YGL037C	replicative lifespan: decreased 	classical genetics 	classical genetics	null Allele: pnc1-Δ	W303		Temperature: elevated temperature, 37 °C 	Anderson RM, et al. (2003) PMID:12736687
POC4	YPL144W	replicative lifespan: decreased 	systematic mutation set 	large-scale survey	null Allele: poc4-Δ	S288C		Details: short-lived mutants, one standard deviation below the mean, were identified using an HTP sequencing-based yeast replicative lifespan screen (SEBYL), with pooled cells subject to four rounds of biotin-based sorting to enrich old vs young cells, followed by sequencing to ID mutants and abundance 	Yu R, et al. (2021) PMID:33790287
POL1	YNL102W	replicative lifespan: decreased 	classical genetics	classical genetics	conditional Allele: pol1-14	Other		Temperature: permissive temperature, 30 °C 	Hoopes LL, et al. (2002) PMID:12024027
POL12	YBL035C	replicative lifespan: decreased 	classical genetics	classical genetics	unspecified Allele: pol12-216	Other			Franke J, et al. (2008) PMID:18840651
POL32	YJR043C	replicative lifespan: decreased 	systematic mutation set 	large-scale survey	null Allele: pol32-Δ	S288C		Details: short-lived mutants, one standard deviation below the mean, were identified using an HTP sequencing-based yeast replicative lifespan screen (SEBYL), with pooled cells subject to four rounds of biotin-based sorting to enrich old vs young cells, followed by sequencing to ID mutants and abundance 	Yu R, et al. (2021) PMID:33790287
PPZ1	YML016C	replicative lifespan: decreased 	systematic mutation set 	large-scale survey	null Allele: ppz1-Δ	S288C		Details: short-lived mutants, one standard deviation below the mean, were identified using an HTP sequencing-based yeast replicative lifespan screen (SEBYL), with pooled cells subject to four rounds of biotin-based sorting to enrich old vs young cells, followed by sequencing to ID mutants and abundance 	Yu R, et al. (2021) PMID:33790287
PRE9	YGR135W	replicative lifespan: decreased 	classical genetics	classical genetics	null Allele: pre9-Δ	S288C			Kruegel U, et al. (2011) PMID:21931558
PRE9	YGR135W	replicative lifespan: decreased 	systematic mutation set 	large-scale survey	null Allele: pre9-Δ	S288C		Details: short-lived mutants, one standard deviation below the mean, were identified using an HTP sequencing-based yeast replicative lifespan screen (SEBYL), with pooled cells subject to four rounds of biotin-based sorting to enrich old vs young cells, followed by sequencing to ID mutants and abundance 	Yu R, et al. (2021) PMID:33790287
PRO1	YDR300C	replicative lifespan: decreased 	classical genetics 	classical genetics	null Allele: pro1-Δ	S288C		Details: markedly shortened lifespan relative to wt 	Mukai Y, et al. (2019) PMID:31646149
PRO2	YOR323C	replicative lifespan: decreased 	classical genetics 	classical genetics	null Allele: pro2-Δ	S288C		Details: markedly shortened lifespan relative to wt 	Mukai Y, et al. (2019) PMID:31646149
PRO2	YOR323C	replicative lifespan: decreased 	systematic mutation set 	large-scale survey	null Allele: pro2-Δ	S288C		Details: short-lived mutants, one standard deviation below the mean, were identified using an HTP sequencing-based yeast replicative lifespan screen (SEBYL), with pooled cells subject to four rounds of biotin-based sorting to enrich old vs young cells, followed by sequencing to ID mutants and abundance 	Yu R, et al. (2021) PMID:33790287
PRX1	YBL064C	replicative lifespan: decreased 	classical genetics	classical genetics	null Allele: prx1-Δ	S288C			Hacioglu E, et al. (2010) PMID:20934449
PTC1	YDL006W	replicative lifespan: decreased 	systematic mutation set 	large-scale survey	null Allele: ptc1-Δ	S288C		Details: short-lived mutants, one standard deviation below the mean, were identified using an HTP sequencing-based yeast replicative lifespan screen (SEBYL), with pooled cells subject to four rounds of biotin-based sorting to enrich old vs young cells, followed by sequencing to ID mutants and abundance 	Yu R, et al. (2021) PMID:33790287
PTC6	YCR079W	replicative lifespan: decreased 	classical genetics	classical genetics	null Allele: ptc6-Δ	S288C		Treatment: caloric restriction by glucose reduction, 0.05% Details: dietary restriction-induced lifespan extension is decreased relative to wt 	Schleit J, et al. (2013) PMID:23837470
PTC6	YCR079W	replicative lifespan: decreased 	systematic mutation set 	large-scale survey	null Allele: ptc6-Δ	S288C		Details: short-lived mutants, one standard deviation below the mean, were identified using an HTP sequencing-based yeast replicative lifespan screen (SEBYL), with pooled cells subject to four rounds of biotin-based sorting to enrich old vs young cells, followed by sequencing to ID mutants and abundance 	Yu R, et al. (2021) PMID:33790287
PUT3	YKL015W	replicative lifespan: decreased 	classical genetics	classical genetics	null Allele: put3-Δ	S288C		Treatment: caloric restriction by glucose reduction, 0.05% Details: dietary restriction-induced lifespan extension is decreased relative to wt 	Schleit J, et al. (2013) PMID:23837470
PXA1	YPL147W	replicative lifespan: decreased 	systematic mutation set 	large-scale survey	null Allele: pxa1-Δ	S288C		Details: short-lived mutants, one standard deviation below the mean, were identified using an HTP sequencing-based yeast replicative lifespan screen (SEBYL), with pooled cells subject to four rounds of biotin-based sorting to enrich old vs young cells, followed by sequencing to ID mutants and abundance 	Yu R, et al. (2021) PMID:33790287
PYC2	YBR218C	replicative lifespan: decreased 	systematic mutation set 	large-scale survey	null Allele: pyc2-Δ	S288C		Details: short-lived mutants, one standard deviation below the mean, were identified using an HTP sequencing-based yeast replicative lifespan screen (SEBYL), with pooled cells subject to four rounds of biotin-based sorting to enrich old vs young cells, followed by sequencing to ID mutants and abundance 	Yu R, et al. (2021) PMID:33790287
RAD23	YEL037C	replicative lifespan: decreased 	classical genetics	classical genetics	null Allele: rad23-Δ	S288C		Treatment: caloric restriction by glucose reduction, 0.05% Details: dietary restriction-induced lifespan extension is decreased relative to wt 	Schleit J, et al. (2013) PMID:23837470
RAD27	YKL113C	replicative lifespan: decreased 	classical genetics	classical genetics	null Allele: rad27-Δ	S288C		Details: cells cultured in buffered SC medium with 10% glucose exhibited a shorter reproductive capacity compared with wild-type cells at day 7, and the reproductive capacity of rad27 cells was reduced even further by day 13 	Weinberger M, et al. (2013) PMID:23518504
RAD27	YKL113C	replicative lifespan: decreased 	classical genetics	classical genetics	null Allele: rad27-Δ	Other		Details: hyperrecombination at the rDNA locus 	Hoopes LL, et al. (2002) PMID:12024027
RAD27	YKL113C	replicative lifespan: decreased 	systematic mutation set 	large-scale survey	null Allele: rad27-Δ	S288C		Details: short-lived mutants, one standard deviation below the mean, were identified using an HTP sequencing-based yeast replicative lifespan screen (SEBYL), with pooled cells subject to four rounds of biotin-based sorting to enrich old vs young cells, followed by sequencing to ID mutants and abundance 	Yu R, et al. (2021) PMID:33790287
RAD5	YLR032W	replicative lifespan: decreased 	systematic mutation set 	large-scale survey	null Allele: rad5-Δ	S288C		Details: short-lived mutants, one standard deviation below the mean, were identified using an HTP sequencing-based yeast replicative lifespan screen (SEBYL), with pooled cells subject to four rounds of biotin-based sorting to enrich old vs young cells, followed by sequencing to ID mutants and abundance 	Yu R, et al. (2021) PMID:33790287
RAD50	YNL250W	replicative lifespan: decreased 	systematic mutation set	large-scale survey	null Allele: rad50-Δ	S288C			Delaney JR, et al. (2013) PMID:23336757
RAD51	YER095W	replicative lifespan: decreased 	systematic mutation set	large-scale survey	null Allele: rad51-Δ	S288C			Delaney JR, et al. (2013) PMID:23336757
RAD52	YML032C	replicative lifespan: decreased 	systematic mutation set	large-scale survey	null Allele: rad52-Δ	S288C			Delaney JR, et al. (2013) PMID:23336757
RAD52	YML032C	replicative lifespan: decreased 	classical genetics 	classical genetics	null Allele: rad52-Δ	S288C	300 μM curcumin	Details: curcumin reduced reproductive potential, as opposed to wt cells where reproductive potential was increased 	Stępień K, et al. (2020) PMID:31659616
RAD53	YPL153C	replicative lifespan: decreased 	systematic mutation set	large-scale survey	null Allele: rad53-Δ	S288C			Delaney JR, et al. (2013) PMID:23336757
RAD55	YDR076W	replicative lifespan: decreased 	systematic mutation set 	large-scale survey	null Allele: rad55-Δ	S288C		Details: short-lived mutants, one standard deviation below the mean, were identified using an HTP sequencing-based yeast replicative lifespan screen (SEBYL), with pooled cells subject to four rounds of biotin-based sorting to enrich old vs young cells, followed by sequencing to ID mutants and abundance 	Yu R, et al. (2021) PMID:33790287
RAD6	YGL058W	replicative lifespan: decreased 	classical genetics	classical genetics	null Allele: rad6-Δ	S288C		Treatment: caloric restriction by glucose reduction, 0.05% Details: dietary restriction-induced lifespan extension is decreased relative to wt 	Schleit J, et al. (2013) PMID:23837470
RAP1	YNL216W	replicative lifespan: decreased 	heterozygous diploid	classical genetics	null Allele: rap1-Δ	S288C		Details: lifespan considerably short relative to wt 	Kamei Y, et al. (2015) PMID:26022127
RAS2	YNL098C	replicative lifespan: decreased 	classical genetics	classical genetics	null Allele: ras2-Δ	Other		Details: strain lacks the mitochondrial genome 	Kirchman PA, et al. (2003) PMID:14739993
RAS2	YNL098C	replicative lifespan: decreased 	classical genetics	classical genetics	null Allele: ras2-Δ	Other			Sun J, et al. (1994) PMID:8034612
RAS2	YNL098C	replicative lifespan: decreased 	classical genetics	classical genetics	activation Allele: RAS2-val19	Other			Hlavatá L, et al. (2008) PMID:18081742
RBD2	YPL246C	replicative lifespan: decreased 	systematic mutation set 	large-scale survey	null Allele: rbd2-Δ	S288C		Details: short-lived mutants, one standard deviation below the mean, were identified using an HTP sequencing-based yeast replicative lifespan screen (SEBYL), with pooled cells subject to four rounds of biotin-based sorting to enrich old vs young cells, followed by sequencing to ID mutants and abundance 	Yu R, et al. (2021) PMID:33790287
REB1	YBR049C	replicative lifespan: decreased 	heterozygous diploid	classical genetics	null Allele: reb1-Δ	S288C		Details: lifespan considerably short relative to wt 	Kamei Y, et al. (2015) PMID:26022127
REG1	YDR028C	replicative lifespan: decreased 	systematic mutation set 	large-scale survey	null Allele: reg1-Δ	S288C		Details: short-lived mutants, one standard deviation below the mean, were identified using an HTP sequencing-based yeast replicative lifespan screen (SEBYL), with pooled cells subject to four rounds of biotin-based sorting to enrich old vs young cells, followed by sequencing to ID mutants and abundance 	Yu R, et al. (2021) PMID:33790287
RGA1	YOR127W	replicative lifespan: decreased 	classical genetics	classical genetics	null Allele: rga1-Δ	Other		Details: lifespan is significantly shorter than wt cells 	Meitinger F, et al. (2014) PMID:25416945
RGD1	YBR260C	replicative lifespan: decreased 	classical genetics	classical genetics	null Allele: rgd1-Δ	S288C		Treatment: caloric restriction by glucose reduction, 0.05% Details: dietary restriction-induced lifespan extension is decreased relative to wt 	Schleit J, et al. (2013) PMID:23837470
RHO5	YNL180C	replicative lifespan: decreased 	systematic mutation set 	large-scale survey	null Allele: rho5-Δ	S288C		Details: short-lived mutants, one standard deviation below the mean, were identified using an HTP sequencing-based yeast replicative lifespan screen (SEBYL), with pooled cells subject to four rounds of biotin-based sorting to enrich old vs young cells, followed by sequencing to ID mutants and abundance 	Yu R, et al. (2021) PMID:33790287
RIF1	YBR275C	replicative lifespan: decreased 	classical genetics	classical genetics	null Allele: rif1-Δ	W303		Details: 40% decrease relative to wild type 	Austriaco NR and Guarente LP (1997) PMID:9275199
RIM20	YOR275C	replicative lifespan: decreased 	systematic mutation set 	large-scale survey	null Allele: rim20-Δ	S288C		Details: short-lived mutants, one standard deviation below the mean, were identified using an HTP sequencing-based yeast replicative lifespan screen (SEBYL), with pooled cells subject to four rounds of biotin-based sorting to enrich old vs young cells, followed by sequencing to ID mutants and abundance 	Yu R, et al. (2021) PMID:33790287
RIM8	YGL045W	replicative lifespan: decreased 	systematic mutation set 	large-scale survey	null Allele: rim8-Δ	S288C		Details: short-lived mutants, one standard deviation below the mean, were identified using an HTP sequencing-based yeast replicative lifespan screen (SEBYL), with pooled cells subject to four rounds of biotin-based sorting to enrich old vs young cells, followed by sequencing to ID mutants and abundance 	Yu R, et al. (2021) PMID:33790287
RIM9	YMR063W	replicative lifespan: decreased 	systematic mutation set 	large-scale survey	null Allele: rim9-Δ	S288C		Details: short-lived mutants, one standard deviation below the mean, were identified using an HTP sequencing-based yeast replicative lifespan screen (SEBYL), with pooled cells subject to four rounds of biotin-based sorting to enrich old vs young cells, followed by sequencing to ID mutants and abundance 	Yu R, et al. (2021) PMID:33790287
RLF2	YPR018W	replicative lifespan: decreased 	classical genetics	classical genetics	null Allele: rlf2-Δ	W303			Feser J, et al. (2010) PMID:20832724
RNR1	YER070W	replicative lifespan: decreased 	classical genetics 	classical genetics	null Allele: rnr1-Δ	S288C			Tai A, et al. (2017) PMID:28495531
RNR1	YER070W	replicative lifespan: decreased 	classical genetics 	classical genetics	reduction of function Allele: rnr1-C428A	S288C			Tai A, et al. (2017) PMID:28495531
RNY1	YPL123C	replicative lifespan: decreased 	systematic mutation set 	large-scale survey	null Allele: rny1-Δ	S288C		Details: short-lived mutants, one standard deviation below the mean, were identified using an HTP sequencing-based yeast replicative lifespan screen (SEBYL), with pooled cells subject to four rounds of biotin-based sorting to enrich old vs young cells, followed by sequencing to ID mutants and abundance 	Yu R, et al. (2021) PMID:33790287
RPB4	YJL140W	replicative lifespan: decreased 	classical genetics	classical genetics	null Allele: rpb4-Δ	S288C		Details: shortened by ~20% compared with wild-type 	Duan R, et al. (2013) PMID:24358479
RPB9	YGL070C	replicative lifespan: decreased 	classical genetics	classical genetics	null Allele: rpb9-Δ	S288C		Details: shortened by ~60% compared with wild-type 	Duan R, et al. (2013) PMID:24358479
RPL14A	YKL006W	replicative lifespan: decreased 	systematic mutation set 	large-scale survey	null Allele: rpl14a-Δ	S288C		Details: short-lived mutants, one standard deviation below the mean, were identified using an HTP sequencing-based yeast replicative lifespan screen (SEBYL), with pooled cells subject to four rounds of biotin-based sorting to enrich old vs young cells, followed by sequencing to ID mutants and abundance 	Yu R, et al. (2021) PMID:33790287
RPL19A	YBR084C-A	replicative lifespan: decreased 	systematic mutation set 	large-scale survey	null Allele: rpl19a-Δ	S288C		Details: short-lived mutants, one standard deviation below the mean, were identified using an HTP sequencing-based yeast replicative lifespan screen (SEBYL), with pooled cells subject to four rounds of biotin-based sorting to enrich old vs young cells, followed by sequencing to ID mutants and abundance 	Yu R, et al. (2021) PMID:33790287
RPL24A	YGL031C	replicative lifespan: decreased 	systematic mutation set 	large-scale survey	null Allele: rpl24a-Δ	S288C		Details: short-lived mutants, one standard deviation below the mean, were identified using an HTP sequencing-based yeast replicative lifespan screen (SEBYL), with pooled cells subject to four rounds of biotin-based sorting to enrich old vs young cells, followed by sequencing to ID mutants and abundance 	Yu R, et al. (2021) PMID:33790287
RPL2B	YIL018W	replicative lifespan: decreased 	systematic mutation set 	large-scale survey	null Allele: rpl2b-Δ	S288C		Details: short-lived mutants, one standard deviation below the mean, were identified using an HTP sequencing-based yeast replicative lifespan screen (SEBYL), with pooled cells subject to four rounds of biotin-based sorting to enrich old vs young cells, followed by sequencing to ID mutants and abundance 	Yu R, et al. (2021) PMID:33790287
RPL6A	YML073C	replicative lifespan: decreased 	classical genetics	classical genetics	null Allele: rpl6a-Δ	S288C			Steffen KK, et al. (2008) PMID:18423200
RPL8A	YHL033C	replicative lifespan: decreased 	systematic mutation set 	large-scale survey	null Allele: rpl8a-Δ	S288C		Details: short-lived mutants, one standard deviation below the mean, were identified using an HTP sequencing-based yeast replicative lifespan screen (SEBYL), with pooled cells subject to four rounds of biotin-based sorting to enrich old vs young cells, followed by sequencing to ID mutants and abundance 	Yu R, et al. (2021) PMID:33790287
RPN10	YHR200W	replicative lifespan: decreased 	systematic mutation set 	large-scale survey	null Allele: rpn10-Δ	S288C		Details: short-lived mutants, one standard deviation below the mean, were identified using an HTP sequencing-based yeast replicative lifespan screen (SEBYL), with pooled cells subject to four rounds of biotin-based sorting to enrich old vs young cells, followed by sequencing to ID mutants and abundance 	Yu R, et al. (2021) PMID:33790287
RPN4	YDL020C	replicative lifespan: decreased 	classical genetics	classical genetics	null Allele: rpn4-Δ	S288C		Details: dietary restriction (reduced glucose concentration; 0.05%) partially rescues the lifespan reduction 	Kruegel U, et al. (2011) PMID:21931558
RPN4	YDL020C	replicative lifespan: decreased 	classical genetics 	classical genetics	null Allele: rpn4-Δ	S288C		Details: shortening to the replicative lifespan based on bud scar number 	Chen LB, et al. (2020) PMID:32861419
RPN4	YDL020C	replicative lifespan: decreased 	systematic mutation set 	large-scale survey	null Allele: rpn4-Δ	S288C		Details: short-lived mutants, one standard deviation below the mean, were identified using an HTP sequencing-based yeast replicative lifespan screen (SEBYL), with pooled cells subject to four rounds of biotin-based sorting to enrich old vs young cells, followed by sequencing to ID mutants and abundance 	Yu R, et al. (2021) PMID:33790287
RPS17A	YML024W	replicative lifespan: decreased 	systematic mutation set 	large-scale survey	null Allele: rps17a-Δ	S288C		Details: short-lived mutants, one standard deviation below the mean, were identified using an HTP sequencing-based yeast replicative lifespan screen (SEBYL), with pooled cells subject to four rounds of biotin-based sorting to enrich old vs young cells, followed by sequencing to ID mutants and abundance 	Yu R, et al. (2021) PMID:33790287
RPS21A	YKR057W	replicative lifespan: decreased 	systematic mutation set 	large-scale survey	null Allele: rps21a-Δ	S288C		Details: short-lived mutants, one standard deviation below the mean, were identified using an HTP sequencing-based yeast replicative lifespan screen (SEBYL), with pooled cells subject to four rounds of biotin-based sorting to enrich old vs young cells, followed by sequencing to ID mutants and abundance 	Yu R, et al. (2021) PMID:33790287
RPS4A	YJR145C	replicative lifespan: decreased 	systematic mutation set 	large-scale survey	null Allele: rps4a-Δ	S288C		Details: short-lived mutants, one standard deviation below the mean, were identified using an HTP sequencing-based yeast replicative lifespan screen (SEBYL), with pooled cells subject to four rounds of biotin-based sorting to enrich old vs young cells, followed by sequencing to ID mutants and abundance 	Yu R, et al. (2021) PMID:33790287
RPT1	YKL145W	replicative lifespan: decreased 	classical genetics	classical genetics	reduction of function Allele: rpt1S K256S; hypomorphic ATPase active site mutation	Other			Kruegel U, et al. (2011) PMID:21931558
RPT2	YDL007W	replicative lifespan: decreased 	classical genetics	classical genetics	reduction of function Allele: rpt2RF K229S, S241F; hypomorphic ATPase active site mutation	Other			Kruegel U, et al. (2011) PMID:21931558
RPT3	YDR394W	replicative lifespan: decreased 	classical genetics	classical genetics	reduction of function Allele: rpt3R K219R; hypomorphic ATPase active site mutation	Other			Kruegel U, et al. (2011) PMID:21931558
RPT5	YOR117W	replicative lifespan: decreased 	classical genetics	classical genetics	reduction of function Allele: rpt5R K228R; hypomorphic ATPase active site mutation	Other			Kruegel U, et al. (2011) PMID:21931558
RPT6	YGL048C	replicative lifespan: decreased 	classical genetics	classical genetics	reduction of function	Other			Kruegel U, et al. (2011) PMID:21931558
RRP6	YOR001W	replicative lifespan: decreased 	systematic mutation set 	large-scale survey	null Allele: rrp6-Δ	S288C		Details: short-lived mutants, one standard deviation below the mean, were identified using an HTP sequencing-based yeast replicative lifespan screen (SEBYL), with pooled cells subject to four rounds of biotin-based sorting to enrich old vs young cells, followed by sequencing to ID mutants and abundance 	Yu R, et al. (2021) PMID:33790287
RRT12	YCR045C	replicative lifespan: decreased 	systematic mutation set 	large-scale survey	null Allele: rrt12-Δ	S288C		Details: short-lived mutants, one standard deviation below the mean, were identified using an HTP sequencing-based yeast replicative lifespan screen (SEBYL), with pooled cells subject to four rounds of biotin-based sorting to enrich old vs young cells, followed by sequencing to ID mutants and abundance 	Yu R, et al. (2021) PMID:33790287
RRT2	YBR246W	replicative lifespan: decreased 	systematic mutation set 	large-scale survey	null Allele: rrt2-Δ	S288C		Details: short-lived mutants, one standard deviation below the mean, were identified using an HTP sequencing-based yeast replicative lifespan screen (SEBYL), with pooled cells subject to four rounds of biotin-based sorting to enrich old vs young cells, followed by sequencing to ID mutants and abundance 	Yu R, et al. (2021) PMID:33790287
RSM22	YKL155C	replicative lifespan: decreased 	systematic mutation set 	large-scale survey	null Allele: rsm22-Δ	S288C		Details: short-lived mutants, one standard deviation below the mean, were identified using an HTP sequencing-based yeast replicative lifespan screen (SEBYL), with pooled cells subject to four rounds of biotin-based sorting to enrich old vs young cells, followed by sequencing to ID mutants and abundance 	Yu R, et al. (2021) PMID:33790287
RSR1	YGR152C	replicative lifespan: decreased 	classical genetics 	classical genetics	null Allele: rsr1-Δ	S288C		Details: reduction in RLS from 26.7 ± 7.51 generations in wt cells to 17.15 ± 4.94 generations in the null mutant 	Yang EJ, et al. (2022) PMID:35281729
RTC1	YOL138C	replicative lifespan: decreased 	systematic mutation set 	large-scale survey	null Allele: rtc1-Δ	S288C		Details: short-lived mutants, one standard deviation below the mean, were identified using an HTP sequencing-based yeast replicative lifespan screen (SEBYL), with pooled cells subject to four rounds of biotin-based sorting to enrich old vs young cells, followed by sequencing to ID mutants and abundance 	Yu R, et al. (2021) PMID:33790287
RTG2	YGL252C	replicative lifespan: decreased 	classical genetics 	classical genetics	null Allele: rtg2-Δ	Other			Borghouts C, et al. (2004) PMID:15020466
RTG2	YGL252C	replicative lifespan: decreased 	classical genetics 	classical genetics	reduction of function Allele: rtg2-T138A	Other			Rios-Anjos RM, et al. (2017) PMID:28472157
RTG2	YGL252C	replicative lifespan: decreased 	classical genetics 	classical genetics	reduction of function Allele: rtg2-R109E	Other			Rios-Anjos RM, et al. (2017) PMID:28472157
RTG2	YGL252C	replicative lifespan: decreased 	classical genetics 	classical genetics	reduction of function Allele: rtg2-E106A	Other			Rios-Anjos RM, et al. (2017) PMID:28472157
RTG2	YGL252C	replicative lifespan: decreased 	classical genetics 	classical genetics	reduction of function Allele: rtg2-E137A	Other			Rios-Anjos RM, et al. (2017) PMID:28472157
RTG2	YGL252C	replicative lifespan: decreased 	classical genetics 	classical genetics	reduction of function Allele: rtg2-D158A	Other			Rios-Anjos RM, et al. (2017) PMID:28472157
RTR1	YER139C	replicative lifespan: decreased 	systematic mutation set 	large-scale survey	null Allele: rtr1-Δ	S288C		Details: short-lived mutants, one standard deviation below the mean, were identified using an HTP sequencing-based yeast replicative lifespan screen (SEBYL), with pooled cells subject to four rounds of biotin-based sorting to enrich old vs young cells, followed by sequencing to ID mutants and abundance 	Yu R, et al. (2021) PMID:33790287
RTS1	YOR014W	replicative lifespan: decreased 	systematic mutation set 	large-scale survey	null Allele: rts1-Δ	S288C		Details: short-lived mutants, one standard deviation below the mean, were identified using an HTP sequencing-based yeast replicative lifespan screen (SEBYL), with pooled cells subject to four rounds of biotin-based sorting to enrich old vs young cells, followed by sequencing to ID mutants and abundance 	Yu R, et al. (2021) PMID:33790287
RTT101	YJL047C	replicative lifespan: decreased 	systematic mutation set 	large-scale survey	null Allele: rtt101-Δ	S288C		Details: short-lived mutants, one standard deviation below the mean, were identified using an HTP sequencing-based yeast replicative lifespan screen (SEBYL), with pooled cells subject to four rounds of biotin-based sorting to enrich old vs young cells, followed by sequencing to ID mutants and abundance 	Yu R, et al. (2021) PMID:33790287
RTT103	YDR289C	replicative lifespan: decreased 	systematic mutation set 	large-scale survey	null Allele: rtt103-Δ	S288C		Details: short-lived mutants, one standard deviation below the mean, were identified using an HTP sequencing-based yeast replicative lifespan screen (SEBYL), with pooled cells subject to four rounds of biotin-based sorting to enrich old vs young cells, followed by sequencing to ID mutants and abundance 	Yu R, et al. (2021) PMID:33790287
RXT2	YBR095C	replicative lifespan: decreased 	systematic mutation set 	large-scale survey	null Allele: rxt2-Δ	S288C		Details: short-lived mutants, one standard deviation below the mean, were identified using an HTP sequencing-based yeast replicative lifespan screen (SEBYL), with pooled cells subject to four rounds of biotin-based sorting to enrich old vs young cells, followed by sequencing to ID mutants and abundance 	Yu R, et al. (2021) PMID:33790287
SCH9	YHR205W	replicative lifespan: decreased 	classical genetics	classical genetics	null Allele: sch9-Δ	S288C		Treatment: caloric restriction by glucose reduction, 0.05% Details: dietary restriction-induced lifespan extension is decreased relative to wt 	Schleit J, et al. (2013) PMID:23837470
SCH9	YHR205W	replicative lifespan: decreased 	classical genetics	classical genetics	unspecified Allele: sch9-(hrm2)	Other			Defossez PA, et al. (1999) PMID:10230397
SCP160	YJL080C	replicative lifespan: decreased 	systematic mutation set 	large-scale survey	null Allele: scp160-Δ	S288C		Details: short-lived mutants, one standard deviation below the mean, were identified using an HTP sequencing-based yeast replicative lifespan screen (SEBYL), with pooled cells subject to four rounds of biotin-based sorting to enrich old vs young cells, followed by sequencing to ID mutants and abundance 	Yu R, et al. (2021) PMID:33790287
SDC1	YDR469W	replicative lifespan: decreased 	systematic mutation set 	large-scale survey	null Allele: sdc1-Δ	S288C		Details: short-lived mutants, one standard deviation below the mean, were identified using an HTP sequencing-based yeast replicative lifespan screen (SEBYL), with pooled cells subject to four rounds of biotin-based sorting to enrich old vs young cells, followed by sequencing to ID mutants and abundance 	Yu R, et al. (2021) PMID:33790287
SEC72	YLR292C	replicative lifespan: decreased 	systematic mutation set 	large-scale survey	null Allele: sec72-Δ	S288C		Details: short-lived mutants, one standard deviation below the mean, were identified using an HTP sequencing-based yeast replicative lifespan screen (SEBYL), with pooled cells subject to four rounds of biotin-based sorting to enrich old vs young cells, followed by sequencing to ID mutants and abundance 	Yu R, et al. (2021) PMID:33790287
SEM1	YDR363W-A	replicative lifespan: decreased 	systematic mutation set 	large-scale survey	null Allele: sem1-Δ	S288C		Details: short-lived mutants, one standard deviation below the mean, were identified using an HTP sequencing-based yeast replicative lifespan screen (SEBYL), with pooled cells subject to four rounds of biotin-based sorting to enrich old vs young cells, followed by sequencing to ID mutants and abundance 	Yu R, et al. (2021) PMID:33790287
SET1	YHR119W	replicative lifespan: decreased 	classical genetics 	classical genetics	null Allele: set1-Δ	S288C		Details: ~1/3 decrease in RLS 	Lord CL, et al. (2017) PMID:27932586
SGF73	YGL066W	replicative lifespan: decreased 	systematic mutation set 	large-scale survey	null Allele: sgf73-Δ	S288C		Details: short-lived mutants, one standard deviation below the mean, were identified using an HTP sequencing-based yeast replicative lifespan screen (SEBYL), with pooled cells subject to four rounds of biotin-based sorting to enrich old vs young cells, followed by sequencing to ID mutants and abundance 	Yu R, et al. (2021) PMID:33790287
SGO1	YOR073W	replicative lifespan: decreased 	systematic mutation set 	large-scale survey	null Allele: sgo1-Δ	S288C		Details: short-lived mutants, one standard deviation below the mean, were identified using an HTP sequencing-based yeast replicative lifespan screen (SEBYL), with pooled cells subject to four rounds of biotin-based sorting to enrich old vs young cells, followed by sequencing to ID mutants and abundance 	Yu R, et al. (2021) PMID:33790287
SGS1	YMR190C	replicative lifespan: decreased 	classical genetics	classical genetics	null Allele: sgs1-Δ	W303		Details: lifespan about 40% that of wild type cells 	Sinclair DA, et al. (1997) PMID:9271578
SGS1	YMR190C	replicative lifespan: decreased 	classical genetics	classical genetics	null Allele: sgs1-Δ	S288C			Harkness TA, et al. (2004) PMID:15514051
SGS1	YMR190C	replicative lifespan: decreased 	classical genetics	classical genetics	null Allele: sgs1-Δ	W303		Details: lifespan reduction from ~32 to ~18 generations; cells that stop dividing early have a large-budded terminal phenotype, compared to unbudded and small budded in wt, arresting at G2/M with bow-tie nuclei and short spindles extending through the bud neck 	McVey M, et al. (2001) PMID:11290710
SGS1	YMR190C	replicative lifespan: decreased 	classical genetics	classical genetics	null Allele: sgs1-Δ	W303		Details: hyperrecombination at the rDNA locus and accumulation of extrachromosomal rDNA circles 	Hoopes LL, et al. (2002) PMID:12024027
SGS1	YMR190C	replicative lifespan: decreased 	classical genetics 	classical genetics	null Allele: sgs1-Δ	Other			Heo SJ, et al. (1999) PMID:10620009
SGS1	YMR190C	replicative lifespan: decreased 	systematic mutation set 	large-scale survey	null Allele: sgs1-Δ	S288C		Details: short-lived mutants, one standard deviation below the mean, were identified using an HTP sequencing-based yeast replicative lifespan screen (SEBYL), with pooled cells subject to four rounds of biotin-based sorting to enrich old vs young cells, followed by sequencing to ID mutants and abundance 	Yu R, et al. (2021) PMID:33790287
SHE3	YBR130C	replicative lifespan: decreased 	systematic mutation set 	large-scale survey	null Allele: she3-Δ	S288C		Details: short-lived mutants, one standard deviation below the mean, were identified using an HTP sequencing-based yeast replicative lifespan screen (SEBYL), with pooled cells subject to four rounds of biotin-based sorting to enrich old vs young cells, followed by sequencing to ID mutants and abundance 	Yu R, et al. (2021) PMID:33790287
SHR5	YOL110W	replicative lifespan: decreased 	systematic mutation set 	large-scale survey	null Allele: shr5-Δ	S288C		Details: short-lived mutants, one standard deviation below the mean, were identified using an HTP sequencing-based yeast replicative lifespan screen (SEBYL), with pooled cells subject to four rounds of biotin-based sorting to enrich old vs young cells, followed by sequencing to ID mutants and abundance 	Yu R, et al. (2021) PMID:33790287
SIC1	YLR079W	replicative lifespan: decreased 	systematic mutation set 	large-scale survey	null Allele: sic1-Δ	S288C		Details: short-lived mutants, one standard deviation below the mean, were identified using an HTP sequencing-based yeast replicative lifespan screen (SEBYL), with pooled cells subject to four rounds of biotin-based sorting to enrich old vs young cells, followed by sequencing to ID mutants and abundance 	Yu R, et al. (2021) PMID:33790287
SIP1	YDR422C	replicative lifespan: decreased 	classical genetics	classical genetics	null Allele: sip1-Δ	S288C		Details: mean lifespan was reduced to 82% that of the wild type 	Ashrafi K, et al. (2000) PMID:10921902
SIP2	YGL208W	replicative lifespan: decreased 	classical genetics	classical genetics	null Allele: sip2-Δ	S288C		Details: mean lifespan is 60% that of wild type cells; cells display characteristics of accelerated aging, such as sterility, nucleolar fragmentation, abnormal distribution of the Sir complex, accumulation of extrachromosomal rDNA circles, and changes indicative of altered glucose metabolism 	Ashrafi K, et al. (2000) PMID:10921902
SIP2	YGL208W	replicative lifespan: decreased 	classical genetics	classical genetics	null Allele: sip2-Δ	W303		Details: mean lifespan is 71% that of wild type cells 	Ashrafi K, et al. (2000) PMID:10921902
SIR2	YDL042C	replicative lifespan: decreased 	homozygous diploid	classical genetics	null Allele: sir2-Δ	SK1		Details: replicative lifespan shortened from 19.55 generations to 8.87 generations 	Boselli M, et al. (2009) PMID:19531355
SIR2	YDL042C	replicative lifespan: decreased 	classical genetics	classical genetics	null Allele: sir2-Δ	W303		Details: 50% reduction in mean replicative lifespan relative to wildtype; associated with increased ERC (extrachromosomal rDNA circle) formation 	Kaeberlein M, et al. (1999) PMID:10521401
SIR2	YDL042C	replicative lifespan: decreased 	homozygous diploid	classical genetics	null Allele: sir2-Δ	W303			Kaeberlein M, et al. (1999) PMID:10521401
SIR2	YDL042C	replicative lifespan: decreased 	heterozygous diploid	classical genetics	null Allele: sir2-Δ	W303		Details: mean replicative lifespan intermediate between the wildtype diploid and the homozygous null mutant 	Kaeberlein M, et al. (1999) PMID:10521401
SIR2	YDL042C	replicative lifespan: decreased 	classical genetics	classical genetics	null Allele: sir2-Δ	S288C			Feser J, et al. (2010) PMID:20832724
SIR2	YDL042C	replicative lifespan: decreased 	classical genetics	classical genetics	null Allele: sir2-Δ	S288C		Treatment: caloric restriction by glucose reduction, 0.05% Details: dietary restriction-induced lifespan extension is decreased relative to wt 	Schleit J, et al. (2013) PMID:23837470
SIR2	YDL042C	replicative lifespan: decreased 	systematic mutation set	large-scale survey	null Allele: sir2-Δ	S288C			Delaney JR, et al. (2013) PMID:23336757
SIR2	YDL042C	replicative lifespan: decreased 	classical genetics 	classical genetics	null Allele: sir2-Δ	S288C			Kaeberlein M, et al. (2004) PMID:15328540
SIR2	YDL042C	replicative lifespan: decreased 	classical genetics	classical genetics	unspecified Allele: sir2-S473E phosphomimetic mutant; site regulated by cAMP-PKA and CK2 signaling	S288C		Details: RLS decreased by 10% 	Kang WK, et al. (2015) PMID:26329457
SIR2	YDL042C	replicative lifespan: decreased 	systematic mutation set 	large-scale survey	null Allele: sir2-Δ	S288C		Details: short-lived mutants, one standard deviation below the mean, were identified using an HTP sequencing-based yeast replicative lifespan screen (SEBYL), with pooled cells subject to four rounds of biotin-based sorting to enrich old vs young cells, followed by sequencing to ID mutants and abundance 	Yu R, et al. (2021) PMID:33790287
SIR2	YDL042C	replicative lifespan: decreased 	classical genetics 	classical genetics	null Allele: sir2-Δ	S288C		Treatment: simulated microgravity, 10 rpm Details: further reduces the RLS that is lowered as the result of simulated microgravity in a uniaxial clinostat, and assayed using the Mother Enrichment Program 	Fukuda APM, et al. (2021) PMID:33612178
SIR3	YLR442C	replicative lifespan: decreased 	classical genetics	classical genetics	null Allele: sir3-Δ	W303		Details: 40% decrease relative to wild type 	Austriaco NR and Guarente LP (1997) PMID:9275199
SIR3	YLR442C	replicative lifespan: decreased 	classical genetics	classical genetics	null Allele: sir3-Δ	W303		Details: 20% reduction in mean replicative lifespan relative to wildtype; associated with increased ERC (extrachromosomal rDNA circle) formation 	Kaeberlein M, et al. (1999) PMID:10521401
SIR4	YDR227W	replicative lifespan: decreased 	classical genetics	classical genetics	null Allele: sir4-Δ	W303		Details: 20% reduction in mean replicative lifespan relative to wildtype; associated with increased ERC (extrachromosomal rDNA circle) formation 	Kaeberlein M, et al. (1999) PMID:10521401
SIR4	YDR227W	replicative lifespan: decreased 	systematic mutation set 	large-scale survey	null Allele: sir4-Δ	S288C		Details: short-lived mutants, one standard deviation below the mean, were identified using an HTP sequencing-based yeast replicative lifespan screen (SEBYL), with pooled cells subject to four rounds of biotin-based sorting to enrich old vs young cells, followed by sequencing to ID mutants and abundance 	Yu R, et al. (2021) PMID:33790287
SIT4	YDL047W	replicative lifespan: decreased 	classical genetics 	classical genetics	null Allele: sit4-Δ	W303			Hayashi N, et al. (2005) PMID:15843932
SKI3	YPR189W	replicative lifespan: decreased 	systematic mutation set 	large-scale survey	null Allele: ski3-Δ	S288C		Details: short-lived mutants, one standard deviation below the mean, were identified using an HTP sequencing-based yeast replicative lifespan screen (SEBYL), with pooled cells subject to four rounds of biotin-based sorting to enrich old vs young cells, followed by sequencing to ID mutants and abundance 	Yu R, et al. (2021) PMID:33790287
SKN7	YHR206W	replicative lifespan: decreased 	classical genetics	classical genetics	null Allele: skn7-Δ	W303	3 μM nolinospiroside F	Details: mutant abrogates the increase in replicative lifespan observed in wt cells treated with steroidal saponin nolinospiroside F 	Sun K, et al. (2013) PMID:23439553
SKN7	YHR206W	replicative lifespan: decreased 	systematic mutation set 	large-scale survey	null Allele: skn7-Δ	S288C		Details: short-lived mutants, one standard deviation below the mean, were identified using an HTP sequencing-based yeast replicative lifespan screen (SEBYL), with pooled cells subject to four rounds of biotin-based sorting to enrich old vs young cells, followed by sequencing to ID mutants and abundance 	Yu R, et al. (2021) PMID:33790287
SKO1	YNL167C	replicative lifespan: decreased 	systematic mutation set 	large-scale survey	null Allele: sko1-Δ	S288C		Details: short-lived mutants, one standard deviation below the mean, were identified using an HTP sequencing-based yeast replicative lifespan screen (SEBYL), with pooled cells subject to four rounds of biotin-based sorting to enrich old vs young cells, followed by sequencing to ID mutants and abundance 	Yu R, et al. (2021) PMID:33790287
SKT5	YBL061C	replicative lifespan: decreased 	systematic mutation set 	large-scale survey	null Allele: skt5-Δ	S288C		Details: short-lived mutants, one standard deviation below the mean, were identified using an HTP sequencing-based yeast replicative lifespan screen (SEBYL), with pooled cells subject to four rounds of biotin-based sorting to enrich old vs young cells, followed by sequencing to ID mutants and abundance 	Yu R, et al. (2021) PMID:33790287
SLA1	YBL007C	replicative lifespan: decreased 	systematic mutation set 	large-scale survey	null Allele: sla1-Δ	S288C		Details: short-lived mutants, one standard deviation below the mean, were identified using an HTP sequencing-based yeast replicative lifespan screen (SEBYL), with pooled cells subject to four rounds of biotin-based sorting to enrich old vs young cells, followed by sequencing to ID mutants and abundance 	Yu R, et al. (2021) PMID:33790287
SLX9	YGR081C	replicative lifespan: decreased 	systematic mutation set 	large-scale survey	null Allele: slx9-Δ	S288C		Details: short-lived mutants, one standard deviation below the mean, were identified using an HTP sequencing-based yeast replicative lifespan screen (SEBYL), with pooled cells subject to four rounds of biotin-based sorting to enrich old vs young cells, followed by sequencing to ID mutants and abundance 	Yu R, et al. (2021) PMID:33790287
SNF1	YDR477W	replicative lifespan: decreased 	classical genetics	classical genetics	overexpression	S288C		Details: lifespan is 74% that of wild type; cells display typical characteristics of accelerated aging late in their lifespan, such as sterility, nucleolar fragmentation, abnormal distribution of the Sir complex, and accumulation of extrachromosomal rDNA circles 	Ashrafi K, et al. (2000) PMID:10921902
SNF1	YDR477W	replicative lifespan: decreased 	classical genetics	classical genetics	null Allele: snf1-Δ	S288C		Details: lifespan is 79% that of wild type; cells do not display all characteristics of accelerated aging; lifespan decrease is apparently due to a reduction in fitness 	Ashrafi K, et al. (2000) PMID:10921902
SNF7	YLR025W	replicative lifespan: decreased 	systematic mutation set 	large-scale survey	null Allele: snf7-Δ	S288C		Details: short-lived mutants, one standard deviation below the mean, were identified using an HTP sequencing-based yeast replicative lifespan screen (SEBYL), with pooled cells subject to four rounds of biotin-based sorting to enrich old vs young cells, followed by sequencing to ID mutants and abundance 	Yu R, et al. (2021) PMID:33790287
SNF8	YPL002C	replicative lifespan: decreased 	systematic mutation set 	large-scale survey	null Allele: snf8-Δ	S288C		Details: short-lived mutants, one standard deviation below the mean, were identified using an HTP sequencing-based yeast replicative lifespan screen (SEBYL), with pooled cells subject to four rounds of biotin-based sorting to enrich old vs young cells, followed by sequencing to ID mutants and abundance 	Yu R, et al. (2021) PMID:33790287
SOD1	YJR104C	replicative lifespan: decreased 	classical genetics	classical genetics	null Allele: sod1-Δ	Other			Zadrag R, et al. (2008) PMID:20021387
SOD1	YJR104C	replicative lifespan: decreased 	classical genetics	classical genetics	null Allele: sod1-Δ	S288C		Details: lifespan reduction on 2% glucose suppressed by antioxidant ascorbic acid 	Schleit J, et al. (2013) PMID:23837470
SOD1	YJR104C	replicative lifespan: decreased 	classical genetics	classical genetics	null Allele: sod1-Δ	S288C	3% glycerol	Media: nonfermentable carbon source Details: reduction in lifespan during respiratory relative to fermentative growth 	Schleit J, et al. (2013) PMID:23837470
SOD1	YJR104C	replicative lifespan: decreased 	classical genetics	classical genetics	null Allele: sod1-Δ	S288C		Treatment: caloric restriction by glucose reduction, 0.05% Details: dietary restriction-induced lifespan extension is decreased relative to wt; suppressed by antioxidant ascorbic acid 	Schleit J, et al. (2013) PMID:23837470
SOD1	YJR104C	replicative lifespan: decreased 	classical genetics	classical genetics	null Allele: sod1-Δ	W303	3 μM nolinospiroside F	Details: mutant abrogates the increase in replicative lifespan observed in wt cells treated with steroidal saponin nolinospiroside F 	Sun K, et al. (2013) PMID:23439553
SOD1	YJR104C	replicative lifespan: decreased 	classical genetics 	classical genetics	null Allele: sod1-Δ	S288C	200 μM curcumin	Details: curcumin reduced reproductive potential, as opposed to wt cells where reproductive potential was increased 	Stępień K, et al. (2020) PMID:31659616
SOD2	YHR008C	replicative lifespan: decreased 	classical genetics	classical genetics	null Allele: sod2-Δ	S288C		Details: lifespan reduction on 2% glucose suppressed by antioxidant ascorbic acid 	Schleit J, et al. (2013) PMID:23837470
SOD2	YHR008C	replicative lifespan: decreased 	classical genetics	classical genetics	null Allele: sod2-Δ	S288C	3% glycerol	Media: nonfermentable carbon source Details: reduction in lifespan during respiratory relative to fermentative growth 	Schleit J, et al. (2013) PMID:23837470
SOD2	YHR008C	replicative lifespan: decreased 	classical genetics	classical genetics	null Allele: sod2-Δ	S288C		Treatment: caloric restriction by glucose reduction, 0.05% Details: dietary restriction-induced lifespan extension is decreased relative to wt; suppressed by antioxidant ascorbic acid 	Schleit J, et al. (2013) PMID:23837470
SOD2	YHR008C	replicative lifespan: decreased 	classical genetics	classical genetics	null Allele: sod2-Δ	W303	3 μM nolinospiroside F	Details: mutant abrogates the increase in replicative lifespan observed in wt cells treated with steroidal saponin nolinospiroside F 	Sun K, et al. (2013) PMID:23439553
SPI1	YER150W	replicative lifespan: decreased 	systematic mutation set 	large-scale survey	null Allele: spi1-Δ	S288C		Details: short-lived mutants, one standard deviation below the mean, were identified using an HTP sequencing-based yeast replicative lifespan screen (SEBYL), with pooled cells subject to four rounds of biotin-based sorting to enrich old vs young cells, followed by sequencing to ID mutants and abundance 	Yu R, et al. (2021) PMID:33790287
SPO21	YOL091W	replicative lifespan: decreased 	systematic mutation set 	large-scale survey	null Allele: spo21-Δ	S288C		Details: short-lived mutants, one standard deviation below the mean, were identified using an HTP sequencing-based yeast replicative lifespan screen (SEBYL), with pooled cells subject to four rounds of biotin-based sorting to enrich old vs young cells, followed by sequencing to ID mutants and abundance 	Yu R, et al. (2021) PMID:33790287
SPO7	YAL009W	replicative lifespan: decreased 	systematic mutation set 	large-scale survey	null Allele: spo7-Δ	S288C		Details: short-lived mutants, one standard deviation below the mean, were identified using an HTP sequencing-based yeast replicative lifespan screen (SEBYL), with pooled cells subject to four rounds of biotin-based sorting to enrich old vs young cells, followed by sequencing to ID mutants and abundance 	Yu R, et al. (2021) PMID:33790287
SPT21	YMR179W	replicative lifespan: decreased 	systematic mutation set 	large-scale survey	null Allele: spt21-Δ	S288C		Details: short-lived mutants, one standard deviation below the mean, were identified using an HTP sequencing-based yeast replicative lifespan screen (SEBYL), with pooled cells subject to four rounds of biotin-based sorting to enrich old vs young cells, followed by sequencing to ID mutants and abundance 	Yu R, et al. (2021) PMID:33790287
SPT8	YLR055C	replicative lifespan: decreased 	systematic mutation set 	large-scale survey	null Allele: spt8-Δ	S288C		Details: short-lived mutants, one standard deviation below the mean, were identified using an HTP sequencing-based yeast replicative lifespan screen (SEBYL), with pooled cells subject to four rounds of biotin-based sorting to enrich old vs young cells, followed by sequencing to ID mutants and abundance 	Yu R, et al. (2021) PMID:33790287
SRB8	YCR081W	replicative lifespan: decreased 	systematic mutation set 	large-scale survey	null Allele: srb8-Δ	S288C		Details: short-lived mutants, one standard deviation below the mean, were identified using an HTP sequencing-based yeast replicative lifespan screen (SEBYL), with pooled cells subject to four rounds of biotin-based sorting to enrich old vs young cells, followed by sequencing to ID mutants and abundance 	Yu R, et al. (2021) PMID:33790287
SRO7	YPR032W	replicative lifespan: decreased 	systematic mutation set 	large-scale survey	null Allele: sro7-Δ	S288C		Details: short-lived mutants, one standard deviation below the mean, were identified using an HTP sequencing-based yeast replicative lifespan screen (SEBYL), with pooled cells subject to four rounds of biotin-based sorting to enrich old vs young cells, followed by sequencing to ID mutants and abundance 	Yu R, et al. (2021) PMID:33790287
SRS2	YJL092W	replicative lifespan: decreased 	classical genetics	classical genetics	null Allele: srs2-Δ	W303		Details: lifespan reduction from ~32 to ~18 generations; cells that stop divining early have a large-budded terminal phenotype, compared to unbudded/small budded in wt, arresting at G2/M with bow-tie nuclei and short spindles extending through the bud neck 	McVey M, et al. (2001) PMID:11290710
SRS2	YJL092W	replicative lifespan: decreased 	systematic mutation set 	large-scale survey	null Allele: srs2-Δ	S288C		Details: short-lived mutants, one standard deviation below the mean, were identified using an HTP sequencing-based yeast replicative lifespan screen (SEBYL), with pooled cells subject to four rounds of biotin-based sorting to enrich old vs young cells, followed by sequencing to ID mutants and abundance 	Yu R, et al. (2021) PMID:33790287
SSA3	YBL075C	replicative lifespan: decreased 	classical genetics 	classical genetics	null Allele: ssa3-Δ	S288C		Media: high phosphate, 11 mM Details: slight reduction in replicative lifespan, further reduced in a ssa3 ssa4 double null 	Nakajima T, et al. (2020) PMID:31797348
SSA4	YER103W	replicative lifespan: decreased 	classical genetics 	classical genetics	null Allele: ssa4-Δ	S288C		Media: high phosphate, 11 mM Details: slight reduction in replicative lifespan, further reduced in a ssa3 ssa4 double null 	Nakajima T, et al. (2020) PMID:31797348
SSB2	YNL209W	replicative lifespan: decreased 	classical genetics 	classical genetics	null Allele: ssb2-Δ	S288C			Zhao W, et al. (2019) PMID:30382435
SSD1	YDR293C	replicative lifespan: decreased 	classical genetics 	classical genetics	repressible	Other			Hu Z, et al. (2018) PMID:30117416
SSD1	YDR293C	replicative lifespan: decreased 	classical genetics	classical genetics	reduction of function Allele: ssd1-d deletion of C-terminal 554 residues including a conserved RNA-binding domain	Other			Kaeberlein M, et al. (2004) PMID:15126388
SSF1	YHR066W	replicative lifespan: decreased 	systematic mutation set 	large-scale survey	null Allele: ssf1-Δ	S288C		Details: short-lived mutants, one standard deviation below the mean, were identified using an HTP sequencing-based yeast replicative lifespan screen (SEBYL), with pooled cells subject to four rounds of biotin-based sorting to enrich old vs young cells, followed by sequencing to ID mutants and abundance 	Yu R, et al. (2021) PMID:33790287
SSM4	YIL030C	replicative lifespan: decreased 	systematic mutation set 	large-scale survey	null Allele: ssm4-Δ	S288C		Details: short-lived mutants, one standard deviation below the mean, were identified using an HTP sequencing-based yeast replicative lifespan screen (SEBYL), with pooled cells subject to four rounds of biotin-based sorting to enrich old vs young cells, followed by sequencing to ID mutants and abundance 	Yu R, et al. (2021) PMID:33790287
SST2	YLR452C	replicative lifespan: decreased 	systematic mutation set 	large-scale survey	null Allele: sst2-Δ	S288C		Details: short-lived mutants, one standard deviation below the mean, were identified using an HTP sequencing-based yeast replicative lifespan screen (SEBYL), with pooled cells subject to four rounds of biotin-based sorting to enrich old vs young cells, followed by sequencing to ID mutants and abundance 	Yu R, et al. (2021) PMID:33790287
STE20	YHL007C	replicative lifespan: decreased 	systematic mutation set 	large-scale survey	null Allele: ste20-Δ	S288C		Details: short-lived mutants, one standard deviation below the mean, were identified using an HTP sequencing-based yeast replicative lifespan screen (SEBYL), with pooled cells subject to four rounds of biotin-based sorting to enrich old vs young cells, followed by sequencing to ID mutants and abundance 	Yu R, et al. (2021) PMID:33790287
STE24	YJR117W	replicative lifespan: decreased 	systematic mutation set 	large-scale survey	null Allele: ste24-Δ	S288C		Details: short-lived mutants, one standard deviation below the mean, were identified using an HTP sequencing-based yeast replicative lifespan screen (SEBYL), with pooled cells subject to four rounds of biotin-based sorting to enrich old vs young cells, followed by sequencing to ID mutants and abundance 	Yu R, et al. (2021) PMID:33790287
STE5	YDR103W	replicative lifespan: decreased 	systematic mutation set 	large-scale survey	null Allele: ste5-Δ	S288C		Details: short-lived mutants, one standard deviation below the mean, were identified using an HTP sequencing-based yeast replicative lifespan screen (SEBYL), with pooled cells subject to four rounds of biotin-based sorting to enrich old vs young cells, followed by sequencing to ID mutants and abundance 	Yu R, et al. (2021) PMID:33790287
STE7	YDL159W	replicative lifespan: decreased 	systematic mutation set 	large-scale survey	null Allele: ste7-Δ	S288C		Details: short-lived mutants, one standard deviation below the mean, were identified using an HTP sequencing-based yeast replicative lifespan screen (SEBYL), with pooled cells subject to four rounds of biotin-based sorting to enrich old vs young cells, followed by sequencing to ID mutants and abundance 	Yu R, et al. (2021) PMID:33790287
STF2	YGR008C	replicative lifespan: decreased 	classical genetics	classical genetics	null Allele: stf2-Δ	S288C		Treatment: caloric restriction by glucose reduction, 0.05% Details: dietary restriction-induced lifespan extension is decreased relative to wt 	Schleit J, et al. (2013) PMID:23837470
SUT1	YGL162W	replicative lifespan: decreased 	systematic mutation set 	large-scale survey	null Allele: sut1-Δ	S288C		Details: short-lived mutants, one standard deviation below the mean, were identified using an HTP sequencing-based yeast replicative lifespan screen (SEBYL), with pooled cells subject to four rounds of biotin-based sorting to enrich old vs young cells, followed by sequencing to ID mutants and abundance 	Yu R, et al. (2021) PMID:33790287
SVF1	YDR346C	replicative lifespan: decreased 	systematic mutation set 	large-scale survey	null Allele: svf1-Δ	S288C		Details: short-lived mutants, one standard deviation below the mean, were identified using an HTP sequencing-based yeast replicative lifespan screen (SEBYL), with pooled cells subject to four rounds of biotin-based sorting to enrich old vs young cells, followed by sequencing to ID mutants and abundance 	Yu R, et al. (2021) PMID:33790287
SWD3	YBR175W	replicative lifespan: decreased 	systematic mutation set 	large-scale survey	null Allele: swd3-Δ	S288C		Details: short-lived mutants, one standard deviation below the mean, were identified using an HTP sequencing-based yeast replicative lifespan screen (SEBYL), with pooled cells subject to four rounds of biotin-based sorting to enrich old vs young cells, followed by sequencing to ID mutants and abundance 	Yu R, et al. (2021) PMID:33790287
SWI4	YER111C	replicative lifespan: decreased 	systematic mutation set 	large-scale survey	null Allele: swi4-Δ	S288C		Details: short-lived mutants, one standard deviation below the mean, were identified using an HTP sequencing-based yeast replicative lifespan screen (SEBYL), with pooled cells subject to four rounds of biotin-based sorting to enrich old vs young cells, followed by sequencing to ID mutants and abundance 	Yu R, et al. (2021) PMID:33790287
SWM2	YNR004W	replicative lifespan: decreased 	systematic mutation set 	large-scale survey	null Allele: swm2-Δ	S288C		Details: short-lived mutants, one standard deviation below the mean, were identified using an HTP sequencing-based yeast replicative lifespan screen (SEBYL), with pooled cells subject to four rounds of biotin-based sorting to enrich old vs young cells, followed by sequencing to ID mutants and abundance 	Yu R, et al. (2021) PMID:33790287
TAT2	YOL020W	replicative lifespan: decreased 	classical genetics	classical genetics	null Allele: tat2-5KR mutant protein more stable than wild-type	S288C			He C, et al. (2014) PMID:25521617
TED1	YIL039W	replicative lifespan: decreased 	systematic mutation set 	large-scale survey	null Allele: ted1-Δ	S288C		Details: short-lived mutants, one standard deviation below the mean, were identified using an HTP sequencing-based yeast replicative lifespan screen (SEBYL), with pooled cells subject to four rounds of biotin-based sorting to enrich old vs young cells, followed by sequencing to ID mutants and abundance 	Yu R, et al. (2021) PMID:33790287
TGS1	YPL157W	replicative lifespan: decreased 	classical genetics	classical genetics	null Allele: tgs1-Δ	Other			Franke J, et al. (2008) PMID:18840651
THI7	YLR237W	replicative lifespan: decreased 	systematic mutation set 	large-scale survey	null Allele: thi7-Δ	S288C		Details: short-lived mutants, one standard deviation below the mean, were identified using an HTP sequencing-based yeast replicative lifespan screen (SEBYL), with pooled cells subject to four rounds of biotin-based sorting to enrich old vs young cells, followed by sequencing to ID mutants and abundance 	Yu R, et al. (2021) PMID:33790287
THP2	YHR167W	replicative lifespan: decreased 	systematic mutation set 	large-scale survey	null Allele: thp2-Δ	S288C		Details: short-lived mutants, one standard deviation below the mean, were identified using an HTP sequencing-based yeast replicative lifespan screen (SEBYL), with pooled cells subject to four rounds of biotin-based sorting to enrich old vs young cells, followed by sequencing to ID mutants and abundance 	Yu R, et al. (2021) PMID:33790287
TLC1	YNCB0010W	replicative lifespan: decreased 	classical genetics	classical genetics	null Allele: tlc1-Δ	S288C			Peng J, et al. (2015) PMID:25822194
TOM5	YPR133W-A	replicative lifespan: decreased 	systematic mutation set 	large-scale survey	null Allele: tom5-Δ	S288C		Details: short-lived mutants, one standard deviation below the mean, were identified using an HTP sequencing-based yeast replicative lifespan screen (SEBYL), with pooled cells subject to four rounds of biotin-based sorting to enrich old vs young cells, followed by sequencing to ID mutants and abundance 	Yu R, et al. (2021) PMID:33790287
TOR1	YJR066W	replicative lifespan: decreased 	classical genetics	classical genetics	null Allele: tor1-Δ	S288C		Treatment: caloric restriction by glucose reduction, 0.05% Details: dietary restriction-induced lifespan extension is decreased relative to wt 	Schleit J, et al. (2013) PMID:23837470
TRM10	YOL093W	replicative lifespan: decreased 	systematic mutation set 	large-scale survey	null Allele: trm10-Δ	S288C		Details: short-lived mutants, one standard deviation below the mean, were identified using an HTP sequencing-based yeast replicative lifespan screen (SEBYL), with pooled cells subject to four rounds of biotin-based sorting to enrich old vs young cells, followed by sequencing to ID mutants and abundance 	Yu R, et al. (2021) PMID:33790287
TRM13	YOL125W	replicative lifespan: decreased 	systematic mutation set 	large-scale survey	null Allele: trm13-Δ	S288C		Details: short-lived mutants, one standard deviation below the mean, were identified using an HTP sequencing-based yeast replicative lifespan screen (SEBYL), with pooled cells subject to four rounds of biotin-based sorting to enrich old vs young cells, followed by sequencing to ID mutants and abundance 	Yu R, et al. (2021) PMID:33790287
TRM9	YML014W	replicative lifespan: decreased 	systematic mutation set 	large-scale survey	null Allele: trm9-Δ	S288C		Details: short-lived mutants, one standard deviation below the mean, were identified using an HTP sequencing-based yeast replicative lifespan screen (SEBYL), with pooled cells subject to four rounds of biotin-based sorting to enrich old vs young cells, followed by sequencing to ID mutants and abundance 	Yu R, et al. (2021) PMID:33790287
TRR1	YDR353W	replicative lifespan: decreased 	classical genetics	classical genetics	null Allele: trr1-Δ	S288C			Hacioglu E, et al. (2010) PMID:20934449
TSA1	YML028W	replicative lifespan: decreased 	classical genetics	classical genetics	null Allele: tsa1-Δ	S288C			Hacioglu E, et al. (2010) PMID:20934449
TSA1	YML028W	replicative lifespan: decreased 	classical genetics	classical genetics	gain of function Allele: TSA1-B7 P182L	W303		Details: lifespan is 13 generations compared to 17 in the parent strain 	Timmermann B, et al. (2010) PMID:20729566
TSA1	YML028W	replicative lifespan: decreased 	systematic mutation set 	large-scale survey	null Allele: tsa1-Δ	S288C		Details: short-lived mutants, one standard deviation below the mean, were identified using an HTP sequencing-based yeast replicative lifespan screen (SEBYL), with pooled cells subject to four rounds of biotin-based sorting to enrich old vs young cells, followed by sequencing to ID mutants and abundance 	Yu R, et al. (2021) PMID:33790287
TUP1	YCR084C	replicative lifespan: decreased 	classical genetics 	classical genetics	null Allele: tup1-Δ	S288C		Details: slight reduction in RLS assayed using a microfluidic chip-based lifespan measurement 	Yu R, et al. (2021) PMID:33790287
TUS1	YLR425W	replicative lifespan: decreased 	systematic mutation set 	large-scale survey	null Allele: tus1-Δ	S288C		Details: short-lived mutants, one standard deviation below the mean, were identified using an HTP sequencing-based yeast replicative lifespan screen (SEBYL), with pooled cells subject to four rounds of biotin-based sorting to enrich old vs young cells, followed by sequencing to ID mutants and abundance 	Yu R, et al. (2021) PMID:33790287
UBA4	YHR111W	replicative lifespan: decreased 	systematic mutation set 	large-scale survey	null Allele: uba4-Δ	S288C		Details: short-lived mutants, one standard deviation below the mean, were identified using an HTP sequencing-based yeast replicative lifespan screen (SEBYL), with pooled cells subject to four rounds of biotin-based sorting to enrich old vs young cells, followed by sequencing to ID mutants and abundance 	Yu R, et al. (2021) PMID:33790287
UBC7	YMR022W	replicative lifespan: decreased 	systematic mutation set 	large-scale survey	null Allele: ubc7-Δ	S288C		Details: short-lived mutants, one standard deviation below the mean, were identified using an HTP sequencing-based yeast replicative lifespan screen (SEBYL), with pooled cells subject to four rounds of biotin-based sorting to enrich old vs young cells, followed by sequencing to ID mutants and abundance 	Yu R, et al. (2021) PMID:33790287
UBI4	YLL039C	replicative lifespan: decreased 	classical genetics 	classical genetics	null Allele: ubi4-Δ	S288C			Zhao W, et al. (2018) PMID:29116578
UBP14	YBR058C	replicative lifespan: decreased 	systematic mutation set 	large-scale survey	null Allele: ubp14-Δ	S288C		Details: short-lived mutants, one standard deviation below the mean, were identified using an HTP sequencing-based yeast replicative lifespan screen (SEBYL), with pooled cells subject to four rounds of biotin-based sorting to enrich old vs young cells, followed by sequencing to ID mutants and abundance 	Yu R, et al. (2021) PMID:33790287
UBP6	YFR010W	replicative lifespan: decreased 	classical genetics	classical genetics	null Allele: ubp6-Δ	S288C			Kruegel U, et al. (2011) PMID:21931558
UMP1	YBR173C	replicative lifespan: decreased 	classical genetics	classical genetics	null Allele: ump1-Δ	S288C			Kruegel U, et al. (2011) PMID:21931558
UTH1	YKR042W	replicative lifespan: decreased 	classical genetics	classical genetics	null Allele: uth1-Δ	W303	3 μM nolinospiroside F	Details: mutant abrogates the increase in replicative lifespan observed in wt cells treated with steroidal saponin nolinospiroside F 	Sun K, et al. (2013) PMID:23439553
UTH1	YKR042W	replicative lifespan: decreased 	systematic mutation set 	large-scale survey	null Allele: uth1-Δ	S288C		Details: short-lived mutants, one standard deviation below the mean, were identified using an HTP sequencing-based yeast replicative lifespan screen (SEBYL), with pooled cells subject to four rounds of biotin-based sorting to enrich old vs young cells, followed by sequencing to ID mutants and abundance 	Yu R, et al. (2021) PMID:33790287
UTR1	YJR049C	replicative lifespan: decreased 	systematic mutation set 	large-scale survey	null Allele: utr1-Δ	S288C		Details: short-lived mutants, one standard deviation below the mean, were identified using an HTP sequencing-based yeast replicative lifespan screen (SEBYL), with pooled cells subject to four rounds of biotin-based sorting to enrich old vs young cells, followed by sequencing to ID mutants and abundance 	Yu R, et al. (2021) PMID:33790287
VAC14	YLR386W	replicative lifespan: decreased 	systematic mutation set 	large-scale survey	null Allele: vac14-Δ	S288C		Details: short-lived mutants, one standard deviation below the mean, were identified using an HTP sequencing-based yeast replicative lifespan screen (SEBYL), with pooled cells subject to four rounds of biotin-based sorting to enrich old vs young cells, followed by sequencing to ID mutants and abundance 	Yu R, et al. (2021) PMID:33790287
VAC8	YEL013W	replicative lifespan: decreased 	classical genetics	classical genetics	null Allele: vac8-Δ	S288C		Treatment: caloric restriction Details: required for caloric restriction-triggered life-span extension 	Tang F, et al. (2008) PMID:18690010
VAC8	YEL013W	replicative lifespan: decreased 	classical genetics	classical genetics	null Allele: vac8-Δ	S288C		Media: rich medium Details: mean lifespan decreased relative to wildtype 	Tang F, et al. (2008) PMID:18690010
VAC8	YEL013W	replicative lifespan: decreased 	systematic mutation set 	large-scale survey	null Allele: vac8-Δ	S288C		Details: short-lived mutants, one standard deviation below the mean, were identified using an HTP sequencing-based yeast replicative lifespan screen (SEBYL), with pooled cells subject to four rounds of biotin-based sorting to enrich old vs young cells, followed by sequencing to ID mutants and abundance 	Yu R, et al. (2021) PMID:33790287
VAM3	YOR106W	replicative lifespan: decreased 	classical genetics	classical genetics	null Allele: vam3-Δ	S288C		Treatment: caloric restriction Details: required for caloric restriction-triggered life-span extension 	Tang F, et al. (2008) PMID:18690010
VAM7	YGL212W	replicative lifespan: decreased 	classical genetics	classical genetics	null Allele: vam7-Δ	S288C		Media: rich medium Details: mean lifespan decreased relative to wildtype 	Tang F, et al. (2008) PMID:18690010
VAM7	YGL212W	replicative lifespan: decreased 	classical genetics	classical genetics	null Allele: vam7-Δ	S288C		Treatment: caloric restriction Details: required for caloric restriction-triggered life-span extension 	Tang F, et al. (2008) PMID:18690010
VCX1	YDL128W	replicative lifespan: decreased 	classical genetics	classical genetics	null Allele: vcx1-Δ	S288C		Treatment: caloric restriction by glucose reduction, 0.05% Details: dietary restriction-induced lifespan extension is decreased relative to wt 	Schleit J, et al. (2013) PMID:23837470
VHR1	YIL056W	replicative lifespan: decreased 	systematic mutation set 	large-scale survey	null Allele: vhr1-Δ	S288C		Details: short-lived mutants, one standard deviation below the mean, were identified using an HTP sequencing-based yeast replicative lifespan screen (SEBYL), with pooled cells subject to four rounds of biotin-based sorting to enrich old vs young cells, followed by sequencing to ID mutants and abundance 	Yu R, et al. (2021) PMID:33790287
VID27	YNL212W	replicative lifespan: decreased 	systematic mutation set 	large-scale survey	null Allele: vid27-Δ	S288C		Details: short-lived mutants, one standard deviation below the mean, were identified using an HTP sequencing-based yeast replicative lifespan screen (SEBYL), with pooled cells subject to four rounds of biotin-based sorting to enrich old vs young cells, followed by sequencing to ID mutants and abundance 	Yu R, et al. (2021) PMID:33790287
VIP1	YLR410W	replicative lifespan: decreased 	systematic mutation set 	large-scale survey	null Allele: vip1-Δ	S288C		Details: short-lived mutants, one standard deviation below the mean, were identified using an HTP sequencing-based yeast replicative lifespan screen (SEBYL), with pooled cells subject to four rounds of biotin-based sorting to enrich old vs young cells, followed by sequencing to ID mutants and abundance 	Yu R, et al. (2021) PMID:33790287
VMA1	YDL185W	replicative lifespan: decreased 	systematic mutation set 	large-scale survey	null Allele: vma1-Δ	S288C		Details: short-lived mutants, one standard deviation below the mean, were identified using an HTP sequencing-based yeast replicative lifespan screen (SEBYL), with pooled cells subject to four rounds of biotin-based sorting to enrich old vs young cells, followed by sequencing to ID mutants and abundance 	Yu R, et al. (2021) PMID:33790287
VMA13	YPR036W	replicative lifespan: decreased 	classical genetics	classical genetics	null Allele: vma13-Δ	S288C		Treatment: caloric restriction by glucose reduction, 0.05% Details: dietary restriction-induced lifespan extension is decreased relative to wt 	Schleit J, et al. (2013) PMID:23837470
VMA13	YPR036W	replicative lifespan: decreased 	systematic mutation set 	large-scale survey	null Allele: vma13-Δ	S288C		Details: short-lived mutants, one standard deviation below the mean, were identified using an HTP sequencing-based yeast replicative lifespan screen (SEBYL), with pooled cells subject to four rounds of biotin-based sorting to enrich old vs young cells, followed by sequencing to ID mutants and abundance 	Yu R, et al. (2021) PMID:33790287
VMA16	YHR026W	replicative lifespan: decreased 	systematic mutation set 	large-scale survey	null Allele: vma16-Δ	S288C		Details: short-lived mutants, one standard deviation below the mean, were identified using an HTP sequencing-based yeast replicative lifespan screen (SEBYL), with pooled cells subject to four rounds of biotin-based sorting to enrich old vs young cells, followed by sequencing to ID mutants and abundance 	Yu R, et al. (2021) PMID:33790287
VMA2	YBR127C	replicative lifespan: decreased 	classical genetics 	classical genetics	null Allele: vma2-Δ	S288C		Details: short median lifespan of five divisions compared with 19 divisions for wt cells 	Yoon SY, et al. (2022) PMID:35216480
VMA2	YBR127C	replicative lifespan: decreased 	homozygous diploid	classical genetics	null Allele: vma2-Δ	S288C		Treatment: caloric restriction Details: prevents lifespan extension by caloric restriction 	Hughes AL and Gottschling DE (2012) PMID:23172144
VMA2	YBR127C	replicative lifespan: decreased 	classical genetics	classical genetics	null Allele: vma2-Δ	S288C		Treatment: caloric restriction by glucose reduction, 0.05% Details: dietary restriction-induced lifespan extension is decreased relative to wt 	Schleit J, et al. (2013) PMID:23837470
VMA2	YBR127C	replicative lifespan: decreased 	classical genetics	classical genetics	null Allele: vma2-Δ	S288C			Henderson KA, et al. (2014) PMID:25190112
VMA21	YGR105W	replicative lifespan: decreased 	classical genetics	classical genetics	null Allele: vma21-Δ	S288C		Treatment: caloric restriction by glucose reduction, 0.05% Details: dietary restriction-induced lifespan extension is decreased relative to wt 	Schleit J, et al. (2013) PMID:23837470
VMA5	YKL080W	replicative lifespan: decreased 	classical genetics	classical genetics	null Allele: vma5-Δ	S288C		Treatment: caloric restriction by glucose reduction, 0.05% Details: dietary restriction-induced lifespan extension is decreased relative to wt 	Schleit J, et al. (2013) PMID:23837470
VMA6	YLR447C	replicative lifespan: decreased 	classical genetics	classical genetics	null Allele: vma6-Δ	S288C		Treatment: caloric restriction by glucose reduction, 0.05% Details: dietary restriction-induced lifespan extension is decreased relative to wt 	Schleit J, et al. (2013) PMID:23837470
VPS24	YKL041W	replicative lifespan: decreased 	systematic mutation set 	large-scale survey	null Allele: vps24-Δ	S288C		Details: short-lived mutants, one standard deviation below the mean, were identified using an HTP sequencing-based yeast replicative lifespan screen (SEBYL), with pooled cells subject to four rounds of biotin-based sorting to enrich old vs young cells, followed by sequencing to ID mutants and abundance 	Yu R, et al. (2021) PMID:33790287
VPS36	YLR417W	replicative lifespan: decreased 	systematic mutation set 	large-scale survey	null Allele: vps36-Δ	S288C		Details: short-lived mutants, one standard deviation below the mean, were identified using an HTP sequencing-based yeast replicative lifespan screen (SEBYL), with pooled cells subject to four rounds of biotin-based sorting to enrich old vs young cells, followed by sequencing to ID mutants and abundance 	Yu R, et al. (2021) PMID:33790287
VPS4	YPR173C	replicative lifespan: decreased 	classical genetics	classical genetics	null Allele: vps4-Δ	S288C		Treatment: caloric restriction by glucose reduction, 0.05% Details: dietary restriction-induced lifespan extension is decreased relative to wt 	Schleit J, et al. (2013) PMID:23837470
VPS41	YDR080W	replicative lifespan: decreased 	classical genetics 	classical genetics	null Allele: vps41-Δ	S288C		Details: relatively short median lifespan of 13 divisions compared with 19 divisions for wt cells 	Yoon SY, et al. (2022) PMID:35216480
VPS65	YLR322W	replicative lifespan: decreased 	systematic mutation set 	large-scale survey	null Allele: vps65-Δ	S288C		Details: short-lived mutants, one standard deviation below the mean, were identified using an HTP sequencing-based yeast replicative lifespan screen (SEBYL), with pooled cells subject to four rounds of biotin-based sorting to enrich old vs young cells, followed by sequencing to ID mutants and abundance 	Yu R, et al. (2021) PMID:33790287
VPS69	YPR087W	replicative lifespan: decreased 	systematic mutation set 	large-scale survey	null Allele: vps69-Δ	S288C		Details: short-lived mutants, one standard deviation below the mean, were identified using an HTP sequencing-based yeast replicative lifespan screen (SEBYL), with pooled cells subject to four rounds of biotin-based sorting to enrich old vs young cells, followed by sequencing to ID mutants and abundance 	Yu R, et al. (2021) PMID:33790287
VPS71	YML041C	replicative lifespan: decreased 	classical genetics	classical genetics	null Allele: vps71-Δ	S288C			Hacioglu E, et al. (2010) PMID:20934449
VPS9	YML097C	replicative lifespan: decreased 	systematic mutation set 	large-scale survey	null Allele: vps9-Δ	S288C		Details: short-lived mutants, one standard deviation below the mean, were identified using an HTP sequencing-based yeast replicative lifespan screen (SEBYL), with pooled cells subject to four rounds of biotin-based sorting to enrich old vs young cells, followed by sequencing to ID mutants and abundance 	Yu R, et al. (2021) PMID:33790287
VTS1	YOR359W	replicative lifespan: decreased 	systematic mutation set 	large-scale survey	null Allele: vts1-Δ	S288C		Details: short-lived mutants, one standard deviation below the mean, were identified using an HTP sequencing-based yeast replicative lifespan screen (SEBYL), with pooled cells subject to four rounds of biotin-based sorting to enrich old vs young cells, followed by sequencing to ID mutants and abundance 	Yu R, et al. (2021) PMID:33790287
WHI3	YNL197C	replicative lifespan: decreased 	systematic mutation set 	large-scale survey	null Allele: whi3-Δ	S288C		Details: short-lived mutants, one standard deviation below the mean, were identified using an HTP sequencing-based yeast replicative lifespan screen (SEBYL), with pooled cells subject to four rounds of biotin-based sorting to enrich old vs young cells, followed by sequencing to ID mutants and abundance 	Yu R, et al. (2021) PMID:33790287
WSS1	YHR134W	replicative lifespan: decreased 	systematic mutation set 	large-scale survey	null Allele: wss1-Δ	S288C		Details: short-lived mutants, one standard deviation below the mean, were identified using an HTP sequencing-based yeast replicative lifespan screen (SEBYL), with pooled cells subject to four rounds of biotin-based sorting to enrich old vs young cells, followed by sequencing to ID mutants and abundance 	Yu R, et al. (2021) PMID:33790287
YAP1	YML007W	replicative lifespan: decreased 	classical genetics	classical genetics	null Allele: yap1-Δ	S288C			Kruegel U, et al. (2011) PMID:21931558
YBL062W	YBL062W	replicative lifespan: decreased 	systematic mutation set 	large-scale survey	null Allele: ybl062w-Δ	S288C		Details: short-lived mutants, one standard deviation below the mean, were identified using an HTP sequencing-based yeast replicative lifespan screen (SEBYL), with pooled cells subject to four rounds of biotin-based sorting to enrich old vs young cells, followed by sequencing to ID mutants and abundance 	Yu R, et al. (2021) PMID:33790287
YBL071C	YBL071C	replicative lifespan: decreased 	systematic mutation set 	large-scale survey	null Allele: ybl071c-Δ	S288C		Details: short-lived mutants, one standard deviation below the mean, were identified using an HTP sequencing-based yeast replicative lifespan screen (SEBYL), with pooled cells subject to four rounds of biotin-based sorting to enrich old vs young cells, followed by sequencing to ID mutants and abundance 	Yu R, et al. (2021) PMID:33790287
YBL100C	YBL100C	replicative lifespan: decreased 	systematic mutation set 	large-scale survey	null Allele: ybl100c-Δ	S288C		Details: short-lived mutants, one standard deviation below the mean, were identified using an HTP sequencing-based yeast replicative lifespan screen (SEBYL), with pooled cells subject to four rounds of biotin-based sorting to enrich old vs young cells, followed by sequencing to ID mutants and abundance 	Yu R, et al. (2021) PMID:33790287
YBR062C	YBR062C	replicative lifespan: decreased 	systematic mutation set 	large-scale survey	null Allele: ybr062c-Δ	S288C		Details: short-lived mutants, one standard deviation below the mean, were identified using an HTP sequencing-based yeast replicative lifespan screen (SEBYL), with pooled cells subject to four rounds of biotin-based sorting to enrich old vs young cells, followed by sequencing to ID mutants and abundance 	Yu R, et al. (2021) PMID:33790287
YBR174C	YBR174C	replicative lifespan: decreased 	systematic mutation set 	large-scale survey	null Allele: ybr174c-Δ	S288C		Details: short-lived mutants, one standard deviation below the mean, were identified using an HTP sequencing-based yeast replicative lifespan screen (SEBYL), with pooled cells subject to four rounds of biotin-based sorting to enrich old vs young cells, followed by sequencing to ID mutants and abundance 	Yu R, et al. (2021) PMID:33790287
YBR238C	YBR238C	replicative lifespan: decreased 	classical genetics	classical genetics	null Allele: ybr238c-Δ	S288C		Treatment: caloric restriction by glucose reduction, 0.05% Details: dietary restriction-induced lifespan extension is decreased relative to wt 	Schleit J, et al. (2013) PMID:23837470
YCK3	YER123W	replicative lifespan: decreased 	systematic mutation set 	large-scale survey	null Allele: yck3-Δ	S288C		Details: short-lived mutants, one standard deviation below the mean, were identified using an HTP sequencing-based yeast replicative lifespan screen (SEBYL), with pooled cells subject to four rounds of biotin-based sorting to enrich old vs young cells, followed by sequencing to ID mutants and abundance 	Yu R, et al. (2021) PMID:33790287
YCL060C	YCL060C	replicative lifespan: decreased 	systematic mutation set 	large-scale survey	null Allele: ycl060c-Δ	S288C		Details: short-lived mutants, one standard deviation below the mean, were identified using an HTP sequencing-based yeast replicative lifespan screen (SEBYL), with pooled cells subject to four rounds of biotin-based sorting to enrich old vs young cells, followed by sequencing to ID mutants and abundance 	Yu R, et al. (2021) PMID:33790287
YCR051W	YCR051W	replicative lifespan: decreased 	systematic mutation set 	large-scale survey	null Allele: ycr051w-Δ	S288C		Details: short-lived mutants, one standard deviation below the mean, were identified using an HTP sequencing-based yeast replicative lifespan screen (SEBYL), with pooled cells subject to four rounds of biotin-based sorting to enrich old vs young cells, followed by sequencing to ID mutants and abundance 	Yu R, et al. (2021) PMID:33790287
YDL041W	YDL041W	replicative lifespan: decreased 	systematic mutation set 	large-scale survey	null Allele: ydl041w-Δ	S288C		Details: short-lived mutants, one standard deviation below the mean, were identified using an HTP sequencing-based yeast replicative lifespan screen (SEBYL), with pooled cells subject to four rounds of biotin-based sorting to enrich old vs young cells, followed by sequencing to ID mutants and abundance 	Yu R, et al. (2021) PMID:33790287
YDL158C	YDL158C	replicative lifespan: decreased 	systematic mutation set 	large-scale survey	null Allele: ydl158c-Δ	S288C		Details: short-lived mutants, one standard deviation below the mean, were identified using an HTP sequencing-based yeast replicative lifespan screen (SEBYL), with pooled cells subject to four rounds of biotin-based sorting to enrich old vs young cells, followed by sequencing to ID mutants and abundance 	Yu R, et al. (2021) PMID:33790287
YDR149C	YDR149C	replicative lifespan: decreased 	systematic mutation set 	large-scale survey	null Allele: ydr149c-Δ	S288C		Details: short-lived mutants, one standard deviation below the mean, were identified using an HTP sequencing-based yeast replicative lifespan screen (SEBYL), with pooled cells subject to four rounds of biotin-based sorting to enrich old vs young cells, followed by sequencing to ID mutants and abundance 	Yu R, et al. (2021) PMID:33790287
YDR193W	YDR193W	replicative lifespan: decreased 	systematic mutation set 	large-scale survey	null Allele: ydr193w-Δ	S288C		Details: short-lived mutants, one standard deviation below the mean, were identified using an HTP sequencing-based yeast replicative lifespan screen (SEBYL), with pooled cells subject to four rounds of biotin-based sorting to enrich old vs young cells, followed by sequencing to ID mutants and abundance 	Yu R, et al. (2021) PMID:33790287
YDR417C	YDR417C	replicative lifespan: decreased 	systematic mutation set 	large-scale survey	null Allele: ydr417c-Δ	S288C		Details: short-lived mutants, one standard deviation below the mean, were identified using an HTP sequencing-based yeast replicative lifespan screen (SEBYL), with pooled cells subject to four rounds of biotin-based sorting to enrich old vs young cells, followed by sequencing to ID mutants and abundance 	Yu R, et al. (2021) PMID:33790287
YET3	YDL072C	replicative lifespan: decreased 	systematic mutation set 	large-scale survey	null Allele: yet3-Δ	S288C		Details: short-lived mutants, one standard deviation below the mean, were identified using an HTP sequencing-based yeast replicative lifespan screen (SEBYL), with pooled cells subject to four rounds of biotin-based sorting to enrich old vs young cells, followed by sequencing to ID mutants and abundance 	Yu R, et al. (2021) PMID:33790287
YFH1	YDL120W	replicative lifespan: decreased 	classical genetics	classical genetics	null Allele: yfh1-Δ	W303		Details: 49% shorter than wildtype 	Desmyter L, et al. (2004) PMID:15130665
YGL046W	YGL046W	replicative lifespan: decreased 	systematic mutation set 	large-scale survey	null Allele: ygl046w-Δ	S288C		Details: short-lived mutants, one standard deviation below the mean, were identified using an HTP sequencing-based yeast replicative lifespan screen (SEBYL), with pooled cells subject to four rounds of biotin-based sorting to enrich old vs young cells, followed by sequencing to ID mutants and abundance 	Yu R, et al. (2021) PMID:33790287
YGR151C	YGR151C	replicative lifespan: decreased 	systematic mutation set 	large-scale survey	null Allele: ygr151c-Δ	S288C		Details: short-lived mutants, one standard deviation below the mean, were identified using an HTP sequencing-based yeast replicative lifespan screen (SEBYL), with pooled cells subject to four rounds of biotin-based sorting to enrich old vs young cells, followed by sequencing to ID mutants and abundance 	Yu R, et al. (2021) PMID:33790287
YHL005C	YHL005C	replicative lifespan: decreased 	systematic mutation set 	large-scale survey	null Allele: yhl005c-Δ	S288C		Details: short-lived mutants, one standard deviation below the mean, were identified using an HTP sequencing-based yeast replicative lifespan screen (SEBYL), with pooled cells subject to four rounds of biotin-based sorting to enrich old vs young cells, followed by sequencing to ID mutants and abundance 	Yu R, et al. (2021) PMID:33790287
YHR130C	YHR130C	replicative lifespan: decreased 	systematic mutation set 	large-scale survey	null Allele: yhr130c-Δ	S288C		Details: short-lived mutants, one standard deviation below the mean, were identified using an HTP sequencing-based yeast replicative lifespan screen (SEBYL), with pooled cells subject to four rounds of biotin-based sorting to enrich old vs young cells, followed by sequencing to ID mutants and abundance 	Yu R, et al. (2021) PMID:33790287
YHR140W	YHR140W	replicative lifespan: decreased 	systematic mutation set 	large-scale survey	null Allele: yhr140w-Δ	S288C		Details: short-lived mutants, one standard deviation below the mean, were identified using an HTP sequencing-based yeast replicative lifespan screen (SEBYL), with pooled cells subject to four rounds of biotin-based sorting to enrich old vs young cells, followed by sequencing to ID mutants and abundance 	Yu R, et al. (2021) PMID:33790287
YJL120W	YJL120W	replicative lifespan: decreased 	systematic mutation set 	large-scale survey	null Allele: yjl120w-Δ	S288C		Details: short-lived mutants, one standard deviation below the mean, were identified using an HTP sequencing-based yeast replicative lifespan screen (SEBYL), with pooled cells subject to four rounds of biotin-based sorting to enrich old vs young cells, followed by sequencing to ID mutants and abundance 	Yu R, et al. (2021) PMID:33790287
YJR087W	YJR087W	replicative lifespan: decreased 	systematic mutation set 	large-scale survey	null Allele: yjr087w-Δ	S288C		Details: short-lived mutants, one standard deviation below the mean, were identified using an HTP sequencing-based yeast replicative lifespan screen (SEBYL), with pooled cells subject to four rounds of biotin-based sorting to enrich old vs young cells, followed by sequencing to ID mutants and abundance 	Yu R, et al. (2021) PMID:33790287
YKL053W	YKL053W	replicative lifespan: decreased 	systematic mutation set 	large-scale survey	null Allele: ykl053w-Δ	S288C		Details: short-lived mutants, one standard deviation below the mean, were identified using an HTP sequencing-based yeast replicative lifespan screen (SEBYL), with pooled cells subject to four rounds of biotin-based sorting to enrich old vs young cells, followed by sequencing to ID mutants and abundance 	Yu R, et al. (2021) PMID:33790287
YKL069W	YKL069W	replicative lifespan: decreased 	classical genetics	classical genetics	null Allele: ykl069w-Δ	S288C			Le DT, et al. (2009) PMID:19049972
YKU70	YMR284W	replicative lifespan: decreased 	classical genetics	classical genetics	null Allele: yku70-Δ	W303			Kaeberlein M, et al. (1999) PMID:10521401
YKU80	YMR106C	replicative lifespan: decreased 	classical genetics	classical genetics	null Allele: yku80-Δ	W303			Kaeberlein M, et al. (1999) PMID:10521401
YLR374C	YLR374C	replicative lifespan: decreased 	systematic mutation set 	large-scale survey	null Allele: ylr374c-Δ	S288C		Details: short-lived mutants, one standard deviation below the mean, were identified using an HTP sequencing-based yeast replicative lifespan screen (SEBYL), with pooled cells subject to four rounds of biotin-based sorting to enrich old vs young cells, followed by sequencing to ID mutants and abundance 	Yu R, et al. (2021) PMID:33790287
YME1	YPR024W	replicative lifespan: decreased 	systematic mutation set 	large-scale survey	null Allele: yme1-Δ	S288C		Details: short-lived mutants, one standard deviation below the mean, were identified using an HTP sequencing-based yeast replicative lifespan screen (SEBYL), with pooled cells subject to four rounds of biotin-based sorting to enrich old vs young cells, followed by sequencing to ID mutants and abundance 	Yu R, et al. (2021) PMID:33790287
YML012C-A	YML012C-A	replicative lifespan: decreased 	systematic mutation set 	large-scale survey	null Allele: yml012c-a-Δ	S288C		Details: short-lived mutants, one standard deviation below the mean, were identified using an HTP sequencing-based yeast replicative lifespan screen (SEBYL), with pooled cells subject to four rounds of biotin-based sorting to enrich old vs young cells, followed by sequencing to ID mutants and abundance 	Yu R, et al. (2021) PMID:33790287
YML090W	YML090W	replicative lifespan: decreased 	systematic mutation set 	large-scale survey	null Allele: yml090w-Δ	S288C		Details: short-lived mutants, one standard deviation below the mean, were identified using an HTP sequencing-based yeast replicative lifespan screen (SEBYL), with pooled cells subject to four rounds of biotin-based sorting to enrich old vs young cells, followed by sequencing to ID mutants and abundance 	Yu R, et al. (2021) PMID:33790287
YOL107W	YOL107W	replicative lifespan: decreased 	systematic mutation set 	large-scale survey	null Allele: yol107w-Δ	S288C		Details: short-lived mutants, one standard deviation below the mean, were identified using an HTP sequencing-based yeast replicative lifespan screen (SEBYL), with pooled cells subject to four rounds of biotin-based sorting to enrich old vs young cells, followed by sequencing to ID mutants and abundance 	Yu R, et al. (2021) PMID:33790287
YPK1	YKL126W	replicative lifespan: decreased 	systematic mutation set 	large-scale survey	null Allele: ypk1-Δ	S288C		Details: short-lived mutants, one standard deviation below the mean, were identified using an HTP sequencing-based yeast replicative lifespan screen (SEBYL), with pooled cells subject to four rounds of biotin-based sorting to enrich old vs young cells, followed by sequencing to ID mutants and abundance 	Yu R, et al. (2021) PMID:33790287
YPK9	YOR291W	replicative lifespan: decreased 	classical genetics 	classical genetics	null Allele: ypk9-Δ	S288C		Details: RLS of the null mutant is 32% shorter than that of wt cells 	Zhao W, et al. (2021) PMID:34524607
YPL102C	YPL102C	replicative lifespan: decreased 	systematic mutation set 	large-scale survey	null Allele: ypl102c-Δ	S288C		Details: short-lived mutants, one standard deviation below the mean, were identified using an HTP sequencing-based yeast replicative lifespan screen (SEBYL), with pooled cells subject to four rounds of biotin-based sorting to enrich old vs young cells, followed by sequencing to ID mutants and abundance 	Yu R, et al. (2021) PMID:33790287
YPL150W	YPL150W	replicative lifespan: decreased 	systematic mutation set 	large-scale survey	null Allele: ypl150w-Δ	S288C		Details: short-lived mutants, one standard deviation below the mean, were identified using an HTP sequencing-based yeast replicative lifespan screen (SEBYL), with pooled cells subject to four rounds of biotin-based sorting to enrich old vs young cells, followed by sequencing to ID mutants and abundance 	Yu R, et al. (2021) PMID:33790287
YPL182C	YPL182C	replicative lifespan: decreased 	systematic mutation set 	large-scale survey	null Allele: ypl182c-Δ	S288C		Details: short-lived mutants, one standard deviation below the mean, were identified using an HTP sequencing-based yeast replicative lifespan screen (SEBYL), with pooled cells subject to four rounds of biotin-based sorting to enrich old vs young cells, followed by sequencing to ID mutants and abundance 	Yu R, et al. (2021) PMID:33790287
YPL205C	YPL205C	replicative lifespan: decreased 	systematic mutation set 	large-scale survey	null Allele: ypl205c-Δ	S288C		Details: short-lived mutants, one standard deviation below the mean, were identified using an HTP sequencing-based yeast replicative lifespan screen (SEBYL), with pooled cells subject to four rounds of biotin-based sorting to enrich old vs young cells, followed by sequencing to ID mutants and abundance 	Yu R, et al. (2021) PMID:33790287
YPL225W	YPL225W	replicative lifespan: decreased 	systematic mutation set 	large-scale survey	null Allele: ypl225w-Δ	S288C		Details: short-lived mutants, one standard deviation below the mean, were identified using an HTP sequencing-based yeast replicative lifespan screen (SEBYL), with pooled cells subject to four rounds of biotin-based sorting to enrich old vs young cells, followed by sequencing to ID mutants and abundance 	Yu R, et al. (2021) PMID:33790287
YPT7	YML001W	replicative lifespan: decreased 	classical genetics	classical genetics	null Allele: ypt7-Δ	S288C		Treatment: caloric restriction Details: required for caloric restriction-triggered life-span extension 	Tang F, et al. (2008) PMID:18690010
YPT7	YML001W	replicative lifespan: decreased 	systematic mutation set 	large-scale survey	null Allele: ypt7-Δ	S288C		Details: short-lived mutants, one standard deviation below the mean, were identified using an HTP sequencing-based yeast replicative lifespan screen (SEBYL), with pooled cells subject to four rounds of biotin-based sorting to enrich old vs young cells, followed by sequencing to ID mutants and abundance 	Yu R, et al. (2021) PMID:33790287
ZDS1	YMR273C	replicative lifespan: decreased 	classical genetics	classical genetics	null Allele: zds1-Δ	Other		Details: mean replicative lifespan, measured in terms of numbers of cell divisions, is 19.6 compared with 24.3 for wild-type 	Tsubakiyama R, et al. (2011) PMID:21712379
ZDS2	YML109W	replicative lifespan: decreased 	classical genetics	classical genetics	null Allele: zds2-Δ	W303		Details: 20% decrease in mean life span 	Roy N and Runge KW (2000) PMID:10662670
