## Additional File 1 for "Uncharacterized yeast gene *YBR238C,* an effector of TORC1 signaling in a mitochondrial feedback loop, accelerates cellular aging via *HAP4*- and *RMD9*-dependent mechanisms": replicative_lifespan_increased_annotations.txt

!
!Date: 11/08/2022
!From: Saccharomyces Genome Database (SGD) 
!URL: http://www.yeastgenome.org/ 
!Contact 
!Funding: NHGRI at US NIH, grant number 5-U41-HG001315 
!

Gene	Gene Systematic Name	Phenotype	Experiment Type	Experiment Type Category	Mutant Information	Strain Background	Chemical	Details	Reference
AAC3	YBR085W	replicative lifespan: increased 	classical genetics	classical genetics	null Allele: aac3-Δ	S288C			McCormick MA, et al. (2015) PMID:26456335
AAN1	YKL075C	replicative lifespan: increased 	classical genetics 	classical genetics	null Allele: aan1-Δ	S288C		Details: null mutant displays an extended lifespan relative to wt with an increase in the mean RLS from 19 to 26 generations 	Sing CN, et al. (2022) PMID:35577788
AAT1	YKL106W	replicative lifespan: increased 	classical genetics	classical genetics	overexpression	Other		Media: glucose carbon source, 2% Details: 25% increase in replicative lifespan 	Easlon E, et al. (2008) PMID:18381895
ABM1	YJR108W	replicative lifespan: increased 	systematic mutation set 	large-scale survey	null Allele: abm1-Δ	S288C		Details: long-lived mutants, one standard deviation above the mean, were identified using an HTP sequencing-based yeast replicative lifespan screen (SEBYL), with pooled cells subject to four rounds of biotin-based sorting to enrich old vs young cells, followed by sequencing to ID mutants and abundance 	Yu R, et al. (2021) PMID:33790287
ABP1	YCR088W	replicative lifespan: increased 	large-scale survey	large-scale survey	null Allele: abp1-Δ	S288C			Managbanag JR, et al. (2008) PMID:19030232
ABP1	YCR088W	replicative lifespan: increased 	systematic mutation set 	large-scale survey	null Allele: abp1-Δ	S288C		Details: long-lived mutants, one standard deviation above the mean, were identified using an HTP sequencing-based yeast replicative lifespan screen (SEBYL), with pooled cells subject to four rounds of biotin-based sorting to enrich old vs young cells, followed by sequencing to ID mutants and abundance 	Yu R, et al. (2021) PMID:33790287
ACB1	YGR037C	replicative lifespan: increased 	classical genetics	classical genetics	null Allele: acb1-Δ	S288C		Treatment: caloric restriction by glucose reduction, 0.05% Details: dietary restriction-induced lifespan extension is increased relative to wt 	Schleit J, et al. (2013) PMID:23837470
ADH1	YOL086C	replicative lifespan: increased 	large-scale survey	large-scale survey	null Allele: adh1-Δ	S288C			Smith ED, et al. (2008) PMID:18340043
ADH1	YOL086C	replicative lifespan: increased 	classical genetics 	classical genetics	overexpression	Other			Reverter-Branchat G, et al. (2007) PMID:17975074
ADH3	YMR083W	replicative lifespan: increased 	classical genetics	classical genetics	null Allele: adh3-Δ	S288C		Treatment: caloric restriction by glucose reduction, 0.05% Details: dietary restriction-induced lifespan extension is increased relative to wt 	Schleit J, et al. (2013) PMID:23837470
AEP1	YMR064W	replicative lifespan: increased 	classical genetics	classical genetics	null Allele: aep1-Δ	S288C			Caballero A, et al. (2011) PMID:21549315
AFG3	YER017C	replicative lifespan: increased 	large-scale survey	large-scale survey	null Allele: afg3-Δ	S288C			Smith ED, et al. (2008) PMID:18340043
AFG3	YER017C	replicative lifespan: increased 	classical genetics	classical genetics	null Allele: afg3-Δ	S288C			Delaney JR, et al. (2011) PMID:21902802
AFG3	YER017C	replicative lifespan: increased 	competitive growth	large-scale survey	null Allele: afg3-Δ	S288C			Delaney JR, et al. (2011) PMID:21191185
AFG3	YER017C	replicative lifespan: increased 	homozygous diploid 	classical genetics	null Allele: afg3-Δ	S288C		Details: 40% extended lifespan 	Muid KA, et al. (2019) PMID:31082499
AFG3	YER017C	replicative lifespan: increased 	classical genetics	classical genetics	null Allele: afg3-Δ	S288C			McCormick MA, et al. (2015) PMID:26456335
AFT1	YGL071W	replicative lifespan: increased 	systematic mutation set 	large-scale survey	null Allele: aft1-Δ	S288C		Details: long-lived mutants, one standard deviation above the mean, were identified using an HTP sequencing-based yeast replicative lifespan screen (SEBYL), with pooled cells subject to four rounds of biotin-based sorting to enrich old vs young cells, followed by sequencing to ID mutants and abundance 	Yu R, et al. (2021) PMID:33790287
AGA2	YGL032C	replicative lifespan: increased 	systematic mutation set 	large-scale survey	null Allele: aga2-Δ	S288C		Details: long-lived mutants, one standard deviation above the mean, were identified using an HTP sequencing-based yeast replicative lifespan screen (SEBYL), with pooled cells subject to four rounds of biotin-based sorting to enrich old vs young cells, followed by sequencing to ID mutants and abundance 	Yu R, et al. (2021) PMID:33790287
AIM11	YER093C-A	replicative lifespan: increased 	systematic mutation set 	large-scale survey	null Allele: aim11-Δ	S288C		Details: long-lived mutants, one standard deviation above the mean, were identified using an HTP sequencing-based yeast replicative lifespan screen (SEBYL), with pooled cells subject to four rounds of biotin-based sorting to enrich old vs young cells, followed by sequencing to ID mutants and abundance 	Yu R, et al. (2021) PMID:33790287
AIM29	YKR074W	replicative lifespan: increased 	classical genetics	classical genetics	null Allele: aim29-Δ	S288C		Treatment: caloric restriction by glucose reduction, 0.05% Details: dietary restriction-induced lifespan extension is increased relative to wt 	Schleit J, et al. (2013) PMID:23837470
ALG12	YNR030W	replicative lifespan: increased 	large-scale survey	large-scale survey	null Allele: alg12-Δ	S288C			Smith ED, et al. (2008) PMID:18340043
ALG12	YNR030W	replicative lifespan: increased 	classical genetics	classical genetics	null Allele: alg12-Δ	S288C			Delaney JR, et al. (2011) PMID:21902802
ALG12	YNR030W	replicative lifespan: increased 	competitive growth	large-scale survey	null Allele: alg12-Δ	S288C			Delaney JR, et al. (2011) PMID:21191185
ALG12	YNR030W	replicative lifespan: increased 	classical genetics	classical genetics	null Allele: alg12-Δ	S288C			Labunskyy VM, et al. (2014) PMID:24391512
ALG12	YNR030W	replicative lifespan: increased 	classical genetics	classical genetics	null Allele: alg12-Δ	S288C			McCormick MA, et al. (2015) PMID:26456335
ALG12	YNR030W	replicative lifespan: increased 	systematic mutation set 	large-scale survey	null Allele: alg12-Δ	S288C		Details: long-lived mutants, one standard deviation above the mean, were identified using an HTP sequencing-based yeast replicative lifespan screen (SEBYL), with pooled cells subject to four rounds of biotin-based sorting to enrich old vs young cells, followed by sequencing to ID mutants and abundance 	Yu R, et al. (2021) PMID:33790287
ALG3	YBL082C	replicative lifespan: increased 	classical genetics	classical genetics	null Allele: alg3-Δ	S288C			Labunskyy VM, et al. (2014) PMID:24391512
ALG3	YBL082C	replicative lifespan: increased 	systematic mutation set 	large-scale survey	null Allele: alg3-Δ	S288C		Details: long-lived mutants, one standard deviation above the mean, were identified using an HTP sequencing-based yeast replicative lifespan screen (SEBYL), with pooled cells subject to four rounds of biotin-based sorting to enrich old vs young cells, followed by sequencing to ID mutants and abundance 	Yu R, et al. (2021) PMID:33790287
ALG6	YOR002W	replicative lifespan: increased 	classical genetics	classical genetics	null Allele: alg6-Δ	S288C			McCormick MA, et al. (2015) PMID:26456335
ALG8	YOR067C	replicative lifespan: increased 	systematic mutation set 	large-scale survey	null Allele: alg8-Δ	S288C		Details: long-lived mutants, one standard deviation above the mean, were identified using an HTP sequencing-based yeast replicative lifespan screen (SEBYL), with pooled cells subject to four rounds of biotin-based sorting to enrich old vs young cells, followed by sequencing to ID mutants and abundance 	Yu R, et al. (2021) PMID:33790287
ALO1	YML086C	replicative lifespan: increased 	classical genetics	classical genetics	null Allele: alo1-Δ	S288C		Treatment: caloric restriction by glucose reduction, 0.05% Details: dietary restriction-induced lifespan extension is increased relative to wt 	Schleit J, et al. (2013) PMID:23837470
ALY1	YKR021W	replicative lifespan: increased 	systematic mutation set 	large-scale survey	null Allele: aly1-Δ	S288C		Details: long-lived mutants, one standard deviation above the mean, were identified using an HTP sequencing-based yeast replicative lifespan screen (SEBYL), with pooled cells subject to four rounds of biotin-based sorting to enrich old vs young cells, followed by sequencing to ID mutants and abundance 	Yu R, et al. (2021) PMID:33790287
APE4	YHR113W	replicative lifespan: increased 	systematic mutation set 	large-scale survey	null Allele: ape4-Δ	S288C		Details: long-lived mutants, one standard deviation above the mean, were identified using an HTP sequencing-based yeast replicative lifespan screen (SEBYL), with pooled cells subject to four rounds of biotin-based sorting to enrich old vs young cells, followed by sequencing to ID mutants and abundance 	Yu R, et al. (2021) PMID:33790287
APL6	YGR261C	replicative lifespan: increased 	classical genetics 	classical genetics	null Allele: apl6-Δ	S288C		Details: long-lived mutant assayed using a microfluidic chip-based lifespan measurement with automatic imaging to validate SEBYL 	Yu R, et al. (2021) PMID:33790287
APM2	YHL019C	replicative lifespan: increased 	systematic mutation set 	large-scale survey	null Allele: apm2-Δ	S288C		Details: long-lived mutants, one standard deviation above the mean, were identified using an HTP sequencing-based yeast replicative lifespan screen (SEBYL), with pooled cells subject to four rounds of biotin-based sorting to enrich old vs young cells, followed by sequencing to ID mutants and abundance 	Yu R, et al. (2021) PMID:33790287
ARN1	YHL040C	replicative lifespan: increased 	systematic mutation set 	large-scale survey	null Allele: arn1-Δ	S288C		Details: long-lived mutants, one standard deviation above the mean, were identified using an HTP sequencing-based yeast replicative lifespan screen (SEBYL), with pooled cells subject to four rounds of biotin-based sorting to enrich old vs young cells, followed by sequencing to ID mutants and abundance 	Yu R, et al. (2021) PMID:33790287
ASG1	YIL130W	replicative lifespan: increased 	systematic mutation set 	large-scale survey	null Allele: asg1-Δ	S288C		Details: long-lived mutants, one standard deviation above the mean, were identified using an HTP sequencing-based yeast replicative lifespan screen (SEBYL), with pooled cells subject to four rounds of biotin-based sorting to enrich old vs young cells, followed by sequencing to ID mutants and abundance 	Yu R, et al. (2021) PMID:33790287
ATG1	YGL180W	replicative lifespan: increased 	classical genetics	classical genetics	null Allele: atg1-Δ	S288C			McCormick MA, et al. (2015) PMID:26456335
ATG1	YGL180W	replicative lifespan: increased 	systematic mutation set 	large-scale survey	null Allele: atg1-Δ	S288C		Details: long-lived mutants, one standard deviation above the mean, were identified using an HTP sequencing-based yeast replicative lifespan screen (SEBYL), with pooled cells subject to four rounds of biotin-based sorting to enrich old vs young cells, followed by sequencing to ID mutants and abundance 	Yu R, et al. (2021) PMID:33790287
ATG32	YIL146C	replicative lifespan: increased 	classical genetics	classical genetics	null Allele: atg32-Δ	S288C		Treatment: caloric restriction by glucose reduction, 0.05% Details: dietary restriction-induced lifespan extension is increased relative to wt 	Schleit J, et al. (2013) PMID:23837470
ATG8	YBL078C	replicative lifespan: increased 	classical genetics	classical genetics	null Allele: atg8-Δ	S288C			McCormick MA, et al. (2015) PMID:26456335
ATO2	YNR002C	replicative lifespan: increased 	classical genetics	classical genetics	null Allele: ato2-Δ	S288C			McCormick MA, et al. (2015) PMID:26456335
ATP1	YBL099W	replicative lifespan: increased 	classical genetics	classical genetics	null Allele: atp1-Δ	S288C		Treatment: caloric restriction by glucose reduction, 0.05% Details: dietary restriction-induced lifespan extension is increased relative to wt 	Schleit J, et al. (2013) PMID:23837470
ATP17	YDR377W	replicative lifespan: increased 	classical genetics	classical genetics	null Allele: atp17-Δ	S288C			McCormick MA, et al. (2015) PMID:26456335
AVT1	YJR001W	replicative lifespan: increased 	homozygous diploid	classical genetics	overexpression	S288C		Details: increase in both median and maximal replicative lifespan 	Hughes AL and Gottschling DE (2012) PMID:23172144
BAS1	YKR099W	replicative lifespan: increased 	large-scale survey	large-scale survey	null Allele: bas1-Δ	S288C			Kaeberlein M, et al. (2005) PMID:16293764
BCK2	YER167W	replicative lifespan: increased 	systematic mutation set 	large-scale survey	null Allele: bck2-Δ	S288C		Details: long-lived mutants, one standard deviation above the mean, were identified using an HTP sequencing-based yeast replicative lifespan screen (SEBYL), with pooled cells subject to four rounds of biotin-based sorting to enrich old vs young cells, followed by sequencing to ID mutants and abundance 	Yu R, et al. (2021) PMID:33790287
BLM10	YFL007W	replicative lifespan: increased 	classical genetics 	classical genetics	overexpression	S288C		Details: increase in rate and number of bud scars 	Jiang TX, et al. (2021) PMID:33391545
BMH2	YDR099W	replicative lifespan: increased 	classical genetics	classical genetics	null Allele: bmh2-Δ	S288C			McCormick MA, et al. (2015) PMID:26456335
BNA2	YJR078W	replicative lifespan: increased 	homozygous diploid 	classical genetics	overexpression	Other		Details: >15% increase in replicative lifespan, determined by both microdissection and the Mother Enrichment Program 	Beas AO, et al. (2020) PMID:32493904
BOI1	YBL085W	replicative lifespan: increased 	systematic mutation set 	large-scale survey	null Allele: boi1-Δ	S288C		Details: long-lived mutants, one standard deviation above the mean, were identified using an HTP sequencing-based yeast replicative lifespan screen (SEBYL), with pooled cells subject to four rounds of biotin-based sorting to enrich old vs young cells, followed by sequencing to ID mutants and abundance 	Yu R, et al. (2021) PMID:33790287
BOI2	YER114C	replicative lifespan: increased 	large-scale survey	large-scale survey	null Allele: boi2-Δ	S288C			Managbanag JR, et al. (2008) PMID:19030232
BOI2	YER114C	replicative lifespan: increased 	classical genetics	classical genetics	null Allele: boi2-Δ	S288C			McCormick MA, et al. (2015) PMID:26456335
BOL2	YGL220W	replicative lifespan: increased 	systematic mutation set 	large-scale survey	null Allele: bol2-Δ	S288C		Details: long-lived mutants, one standard deviation above the mean, were identified using an HTP sequencing-based yeast replicative lifespan screen (SEBYL), with pooled cells subject to four rounds of biotin-based sorting to enrich old vs young cells, followed by sequencing to ID mutants and abundance 	Yu R, et al. (2021) PMID:33790287
BRE5	YNR051C	replicative lifespan: increased 	classical genetics	classical genetics	null Allele: bre5-Δ	S288C			Kaeberlein M, et al. (2005) PMID:16293764
BRE5	YNR051C	replicative lifespan: increased 	large-scale survey	large-scale survey	null Allele: bre5-Δ	S288C			Kaeberlein M, et al. (2005) PMID:16293764
BSC2	YDR275W	replicative lifespan: increased 	systematic mutation set 	large-scale survey	null Allele: bsc2-Δ	S288C		Details: long-lived mutants, one standard deviation above the mean, were identified using an HTP sequencing-based yeast replicative lifespan screen (SEBYL), with pooled cells subject to four rounds of biotin-based sorting to enrich old vs young cells, followed by sequencing to ID mutants and abundance 	Yu R, et al. (2021) PMID:33790287
BST1	YFL025C	replicative lifespan: increased 	classical genetics	classical genetics	null Allele: bst1-Δ	S288C			Labunskyy VM, et al. (2014) PMID:24391512
BST1	YFL025C	replicative lifespan: increased 	classical genetics	classical genetics	null Allele: bst1-Δ	S288C			McCormick MA, et al. (2015) PMID:26456335
BUD7	YOR299W	replicative lifespan: increased 	systematic mutation set 	large-scale survey	null Allele: bud7-Δ	S288C		Details: long-lived mutants, one standard deviation above the mean, were identified using an HTP sequencing-based yeast replicative lifespan screen (SEBYL), with pooled cells subject to four rounds of biotin-based sorting to enrich old vs young cells, followed by sequencing to ID mutants and abundance 	Yu R, et al. (2021) PMID:33790287
BUD8	YLR353W	replicative lifespan: increased 	classical genetics	classical genetics	null Allele: bud8-Δ	S288C			McCormick MA, et al. (2015) PMID:26456335
BUG1	YDL099W	replicative lifespan: increased 	systematic mutation set 	large-scale survey	null Allele: bug1-Δ	S288C		Details: long-lived mutants, one standard deviation above the mean, were identified using an HTP sequencing-based yeast replicative lifespan screen (SEBYL), with pooled cells subject to four rounds of biotin-based sorting to enrich old vs young cells, followed by sequencing to ID mutants and abundance 	Yu R, et al. (2021) PMID:33790287
BUL1	YMR275C	replicative lifespan: increased 	classical genetics	classical genetics	null Allele: bul1-Δ	S288C			McCormick MA, et al. (2015) PMID:26456335
BXI1	YNL305C	replicative lifespan: increased 	classical genetics	classical genetics	null Allele: bxi1-Δ	S288C			Büttner S, et al. (2011) PMID:21673659
CAF20	YOR276W	replicative lifespan: increased 	classical genetics	classical genetics	null Allele: caf20-Δ	S288C			McCormick MA, et al. (2015) PMID:26456335
CAF40	YNL288W	replicative lifespan: increased 	classical genetics	classical genetics	null Allele: caf40-Δ	S288C			McCormick MA, et al. (2015) PMID:26456335
CAN1	YEL063C	replicative lifespan: increased 	classical genetics	classical genetics	null Allele: can1-Δ	S288C			McCormick MA, et al. (2015) PMID:26456335
CBP6	YBR120C	replicative lifespan: increased 	classical genetics	classical genetics	null Allele: cbp6-Δ	S288C			Caballero A, et al. (2011) PMID:21549315
CBS1	YDL069C	replicative lifespan: increased 	classical genetics	classical genetics	null Allele: cbs1-Δ	S288C			Caballero A, et al. (2011) PMID:21549315
CBS1	YDL069C	replicative lifespan: increased 	classical genetics	classical genetics	null Allele: cbs1-Δ	S288C			McCormick MA, et al. (2015) PMID:26456335
CCS1	YMR038C	replicative lifespan: increased 	systematic mutation set 	large-scale survey	null Allele: ccs1-Δ	S288C		Details: long-lived mutants, one standard deviation above the mean, were identified using an HTP sequencing-based yeast replicative lifespan screen (SEBYL), with pooled cells subject to four rounds of biotin-based sorting to enrich old vs young cells, followed by sequencing to ID mutants and abundance 	Yu R, et al. (2021) PMID:33790287
CCW12	YLR110C	replicative lifespan: increased 	systematic mutation set 	large-scale survey	null Allele: ccw12-Δ	S288C		Details: long-lived mutants, one standard deviation above the mean, were identified using an HTP sequencing-based yeast replicative lifespan screen (SEBYL), with pooled cells subject to four rounds of biotin-based sorting to enrich old vs young cells, followed by sequencing to ID mutants and abundance 	Yu R, et al. (2021) PMID:33790287
CDC25	YLR310C	replicative lifespan: increased 	classical genetics	classical genetics	conditional Allele: cdc25-10 E1328V	W303		Temperature: permissive temperature, 30 °C Details: caloric restriction (0.5% glucose) fails to further increase lifespan 	Lin SJ, et al. (2000) PMID:11000115
CEM1	YER061C	replicative lifespan: increased 	systematic mutation set 	large-scale survey	null Allele: cem1-Δ	S288C		Details: long-lived mutants, one standard deviation above the mean, were identified using an HTP sequencing-based yeast replicative lifespan screen (SEBYL), with pooled cells subject to four rounds of biotin-based sorting to enrich old vs young cells, followed by sequencing to ID mutants and abundance 	Yu R, et al. (2021) PMID:33790287
CEX1	YOR112W	replicative lifespan: increased 	systematic mutation set 	large-scale survey	null Allele: cex1-Δ	S288C		Details: long-lived mutants, one standard deviation above the mean, were identified using an HTP sequencing-based yeast replicative lifespan screen (SEBYL), with pooled cells subject to four rounds of biotin-based sorting to enrich old vs young cells, followed by sequencing to ID mutants and abundance 	Yu R, et al. (2021) PMID:33790287
CGI121	YML036W	replicative lifespan: increased 	classical genetics	classical genetics	null Allele: cgi121-Δ	S288C			Peng J, et al. (2015) PMID:25822194
CHD1	YER164W	replicative lifespan: increased 	classical genetics	classical genetics	null Allele: chd1-Δ	S288C			McCormick MA, et al. (2015) PMID:26456335
CKA2	YOR061W	replicative lifespan: increased 	classical genetics	classical genetics	null Allele: cka2-Δ	S288C			McCormick MA, et al. (2015) PMID:26456335
CLB2	YPR119W	replicative lifespan: increased 	large-scale survey	large-scale survey	null Allele: clb2-Δ	S288C			Managbanag JR, et al. (2008) PMID:19030232
CLB3	YDL155W	replicative lifespan: increased 	systematic mutation set 	large-scale survey	null Allele: clb3-Δ	S288C		Details: long-lived mutants, one standard deviation above the mean, were identified using an HTP sequencing-based yeast replicative lifespan screen (SEBYL), with pooled cells subject to four rounds of biotin-based sorting to enrich old vs young cells, followed by sequencing to ID mutants and abundance 	Yu R, et al. (2021) PMID:33790287
CLB6	YGR109C	replicative lifespan: increased 	systematic mutation set 	large-scale survey	null Allele: clb6-Δ	S288C		Details: long-lived mutants, one standard deviation above the mean, were identified using an HTP sequencing-based yeast replicative lifespan screen (SEBYL), with pooled cells subject to four rounds of biotin-based sorting to enrich old vs young cells, followed by sequencing to ID mutants and abundance 	Yu R, et al. (2021) PMID:33790287
CLD1	YGR110W	replicative lifespan: increased 	systematic mutation set 	large-scale survey	null Allele: cld1-Δ	S288C		Details: long-lived mutants, one standard deviation above the mean, were identified using an HTP sequencing-based yeast replicative lifespan screen (SEBYL), with pooled cells subject to four rounds of biotin-based sorting to enrich old vs young cells, followed by sequencing to ID mutants and abundance 	Yu R, et al. (2021) PMID:33790287
CNE1	YAL058W	replicative lifespan: increased 	systematic mutation set 	large-scale survey	null Allele: cne1-Δ	S288C		Details: long-lived mutants, one standard deviation above the mean, were identified using an HTP sequencing-based yeast replicative lifespan screen (SEBYL), with pooled cells subject to four rounds of biotin-based sorting to enrich old vs young cells, followed by sequencing to ID mutants and abundance 	Yu R, et al. (2021) PMID:33790287
COQ3	YOL096C	replicative lifespan: increased 	classical genetics	classical genetics	null Allele: coq3-Δ	S288C		Treatment: caloric restriction by glucose reduction, 0.05% Details: dietary restriction-induced lifespan extension is increased relative to wt 	Schleit J, et al. (2013) PMID:23837470
COQ4	YDR204W	replicative lifespan: increased 	classical genetics	classical genetics	null Allele: coq4-Δ	S288C		Treatment: caloric restriction by glucose reduction, 0.05% Details: dietary restriction-induced lifespan extension is increased relative to wt 	Schleit J, et al. (2013) PMID:23837470
COS12	YGL263W	replicative lifespan: increased 	classical genetics	classical genetics	null Allele: cos12-Δ	S288C			McCormick MA, et al. (2015) PMID:26456335
CPR2	YHR057C	replicative lifespan: increased 	systematic mutation set 	large-scale survey	null Allele: cpr2-Δ	S288C		Details: long-lived mutants, one standard deviation above the mean, were identified using an HTP sequencing-based yeast replicative lifespan screen (SEBYL), with pooled cells subject to four rounds of biotin-based sorting to enrich old vs young cells, followed by sequencing to ID mutants and abundance 	Yu R, et al. (2021) PMID:33790287
CRC1	YOR100C	replicative lifespan: increased 	large-scale survey	large-scale survey	null Allele: crc1-Δ	S288C			Kaeberlein M, et al. (2005) PMID:16293764
CRH1	YGR189C	replicative lifespan: increased 	systematic mutation set 	large-scale survey	null Allele: crh1-Δ	S288C		Details: long-lived mutants, one standard deviation above the mean, were identified using an HTP sequencing-based yeast replicative lifespan screen (SEBYL), with pooled cells subject to four rounds of biotin-based sorting to enrich old vs young cells, followed by sequencing to ID mutants and abundance 	Yu R, et al. (2021) PMID:33790287
CRS5	YOR031W	replicative lifespan: increased 	systematic mutation set 	large-scale survey	null Allele: crs5-Δ	S288C		Details: long-lived mutants, one standard deviation above the mean, were identified using an HTP sequencing-based yeast replicative lifespan screen (SEBYL), with pooled cells subject to four rounds of biotin-based sorting to enrich old vs young cells, followed by sequencing to ID mutants and abundance 	Yu R, et al. (2021) PMID:33790287
CSI1	YMR025W	replicative lifespan: increased 	systematic mutation set 	large-scale survey	null Allele: csi1-Δ	S288C		Details: long-lived mutants, one standard deviation above the mean, were identified using an HTP sequencing-based yeast replicative lifespan screen (SEBYL), with pooled cells subject to four rounds of biotin-based sorting to enrich old vs young cells, followed by sequencing to ID mutants and abundance 	Yu R, et al. (2021) PMID:33790287
CSR2	YPR030W	replicative lifespan: increased 	classical genetics	classical genetics	null Allele: csr2-Δ	Other		Details: replicative lifespan is 34 generations relative to 28 for wild type 	Kharade SV, et al. (2005) PMID:16336970
CST26	YBR042C	replicative lifespan: increased 	classical genetics	classical genetics	null Allele: cst26-Δ	S288C			McCormick MA, et al. (2015) PMID:26456335
CTH1	YDR151C	replicative lifespan: increased 	classical genetics	classical genetics	null Allele: cth1-Δ	S288C			McCormick MA, et al. (2015) PMID:26456335
CTH1	YDR151C	replicative lifespan: increased 	systematic mutation set 	large-scale survey	null Allele: cth1-Δ	S288C		Details: long-lived mutants, one standard deviation above the mean, were identified using an HTP sequencing-based yeast replicative lifespan screen (SEBYL), with pooled cells subject to four rounds of biotin-based sorting to enrich old vs young cells, followed by sequencing to ID mutants and abundance 	Yu R, et al. (2021) PMID:33790287
CUE5	YOR042W	replicative lifespan: increased 	systematic mutation set 	large-scale survey	null Allele: cue5-Δ	S288C		Details: long-lived mutants, one standard deviation above the mean, were identified using an HTP sequencing-based yeast replicative lifespan screen (SEBYL), with pooled cells subject to four rounds of biotin-based sorting to enrich old vs young cells, followed by sequencing to ID mutants and abundance 	Yu R, et al. (2021) PMID:33790287
CUP9	YPL177C	replicative lifespan: increased 	classical genetics	classical genetics	null Allele: cup9-Δ	S288C			McCormick MA, et al. (2015) PMID:26456335
CUR1	YPR158W	replicative lifespan: increased 	classical genetics	classical genetics	null Allele: cur1-Δ	S288C			McCormick MA, et al. (2015) PMID:26456335
CUZ1	YNL155W	replicative lifespan: increased 	systematic mutation set 	large-scale survey	null Allele: cuz1-Δ	S288C		Details: long-lived mutants, one standard deviation above the mean, were identified using an HTP sequencing-based yeast replicative lifespan screen (SEBYL), with pooled cells subject to four rounds of biotin-based sorting to enrich old vs young cells, followed by sequencing to ID mutants and abundance 	Yu R, et al. (2021) PMID:33790287
CYC1	YJR048W	replicative lifespan: increased 	classical genetics	classical genetics	null Allele: cyc1-Δ	S288C			McCormick MA, et al. (2015) PMID:26456335
CYR1	YJL005W	replicative lifespan: increased 	classical genetics	classical genetics	conditional Allele: cyr1-(cdc35-1)	S288C		Temperature: permissive temperature, 25 °C 	Lin SJ, et al. (2000) PMID:11000115
CYT2	YKL087C	replicative lifespan: increased 	classical genetics	classical genetics	null Allele: cyt2-Δ	S288C		Treatment: caloric restriction by glucose reduction, 0.05% Details: dietary restriction-induced lifespan extension is increased relative to wt 	Schleit J, et al. (2013) PMID:23837470
DAK1	YML070W	replicative lifespan: increased 	systematic mutation set 	large-scale survey	null Allele: dak1-Δ	S288C		Details: long-lived mutants, one standard deviation above the mean, were identified using an HTP sequencing-based yeast replicative lifespan screen (SEBYL), with pooled cells subject to four rounds of biotin-based sorting to enrich old vs young cells, followed by sequencing to ID mutants and abundance 	Yu R, et al. (2021) PMID:33790287
DAS2	YDR020C	replicative lifespan: increased 	classical genetics	classical genetics	null Allele: das2-Δ	S288C			McCormick MA, et al. (2015) PMID:26456335
DBF20	YPR111W	replicative lifespan: increased 	classical genetics	classical genetics	null Allele: dbf20-Δ	S288C			McCormick MA, et al. (2015) PMID:26456335
DBP3	YGL078C	replicative lifespan: increased 	large-scale survey	large-scale survey	null Allele: dbp3-Δ	S288C			Smith ED, et al. (2008) PMID:18340043
DBP3	YGL078C	replicative lifespan: increased 	classical genetics	classical genetics	null Allele: dbp3-Δ	S288C			Delaney JR, et al. (2011) PMID:21902802
DBP3	YGL078C	replicative lifespan: increased 	competitive growth	large-scale survey	null Allele: dbp3-Δ	S288C			Delaney JR, et al. (2011) PMID:21191185
DBP3	YGL078C	replicative lifespan: increased 	classical genetics	classical genetics	null Allele: dbp3-Δ	S288C			McCormick MA, et al. (2015) PMID:26456335
DCK1	YLR422W	replicative lifespan: increased 	large-scale survey	large-scale survey	null Allele: dck1-Δ	S288C			Kaeberlein M, et al. (2005) PMID:16293764
DER1	YBR201W	replicative lifespan: increased 	classical genetics	classical genetics	null Allele: der1-Δ	S288C			Labunskyy VM, et al. (2014) PMID:24391512
DFG5	YMR238W	replicative lifespan: increased 	systematic mutation set 	large-scale survey	null Allele: dfg5-Δ	S288C		Details: long-lived mutants, one standard deviation above the mean, were identified using an HTP sequencing-based yeast replicative lifespan screen (SEBYL), with pooled cells subject to four rounds of biotin-based sorting to enrich old vs young cells, followed by sequencing to ID mutants and abundance 	Yu R, et al. (2021) PMID:33790287
DGA1	YOR245C	replicative lifespan: increased 	systematic mutation set 	large-scale survey	null Allele: dga1-Δ	S288C		Details: long-lived mutants, one standard deviation above the mean, were identified using an HTP sequencing-based yeast replicative lifespan screen (SEBYL), with pooled cells subject to four rounds of biotin-based sorting to enrich old vs young cells, followed by sequencing to ID mutants and abundance 	Yu R, et al. (2021) PMID:33790287
DGK1	YOR311C	replicative lifespan: increased 	classical genetics	classical genetics	null Allele: dgk1-Δ	S288C			McCormick MA, et al. (2015) PMID:26456335
DID2	YKR035W-A	replicative lifespan: increased 	classical genetics	classical genetics	null Allele: did2-Δ	S288C			McCormick MA, et al. (2015) PMID:26456335
DIT1	YDR403W	replicative lifespan: increased 	classical genetics 	classical genetics	null Allele: dit1-Δ	S288C		Details: long-lived mutant assayed using a microfluidic chip-based lifespan measurement with automatic imaging to validate SEBYL 	Yu R, et al. (2021) PMID:33790287
DIT2	YDR402C	replicative lifespan: increased 	systematic mutation set 	large-scale survey	null Allele: dit2-Δ	S288C		Details: long-lived mutants, one standard deviation above the mean, were identified using an HTP sequencing-based yeast replicative lifespan screen (SEBYL), with pooled cells subject to four rounds of biotin-based sorting to enrich old vs young cells, followed by sequencing to ID mutants and abundance 	Yu R, et al. (2021) PMID:33790287
DLT1	YMR126C	replicative lifespan: increased 	classical genetics	classical genetics	null Allele: dlt1-Δ	S288C			McCormick MA, et al. (2015) PMID:26456335
DMO2	YDL157C	replicative lifespan: increased 	systematic mutation set 	large-scale survey	null Allele: ydl157c-Δ	S288C		Details: long-lived mutants, one standard deviation above the mean, were identified using an HTP sequencing-based yeast replicative lifespan screen (SEBYL), with pooled cells subject to four rounds of biotin-based sorting to enrich old vs young cells, followed by sequencing to ID mutants and abundance 	Yu R, et al. (2021) PMID:33790287
DOM34	YNL001W	replicative lifespan: increased 	classical genetics	classical genetics	null Allele: dom34-Δ	S288C			McCormick MA, et al. (2015) PMID:26456335
DPC29	YGR021W	replicative lifespan: increased 	systematic mutation set 	large-scale survey	null Allele: dpc29-Δ	S288C		Details: long-lived mutants, one standard deviation above the mean, were identified using an HTP sequencing-based yeast replicative lifespan screen (SEBYL), with pooled cells subject to four rounds of biotin-based sorting to enrich old vs young cells, followed by sequencing to ID mutants and abundance 	Yu R, et al. (2021) PMID:33790287
DPP1	YDR284C	replicative lifespan: increased 	systematic mutation set 	large-scale survey	null Allele: dpp1-Δ	S288C		Details: long-lived mutants, one standard deviation above the mean, were identified using an HTP sequencing-based yeast replicative lifespan screen (SEBYL), with pooled cells subject to four rounds of biotin-based sorting to enrich old vs young cells, followed by sequencing to ID mutants and abundance 	Yu R, et al. (2021) PMID:33790287
DSE3	YOR264W	replicative lifespan: increased 	systematic mutation set 	large-scale survey	null Allele: dse3-Δ	S288C		Details: long-lived mutants, one standard deviation above the mean, were identified using an HTP sequencing-based yeast replicative lifespan screen (SEBYL), with pooled cells subject to four rounds of biotin-based sorting to enrich old vs young cells, followed by sequencing to ID mutants and abundance 	Yu R, et al. (2021) PMID:33790287
DSF2	YBR007C	replicative lifespan: increased 	classical genetics	classical genetics	null Allele: dsf2-Δ	S288C			McCormick MA, et al. (2015) PMID:26456335
DSK2	YMR276W	replicative lifespan: increased 	classical genetics	classical genetics	null Allele: dsk2-Δ	S288C			McCormick MA, et al. (2015) PMID:26456335
DSS1	YMR287C	replicative lifespan: increased 	classical genetics	classical genetics	null Allele: dss1-Δ	S288C			Caballero A, et al. (2011) PMID:21549315
DSS1	YMR287C	replicative lifespan: increased 	homozygous diploid 	classical genetics	null Allele: dss1-Δ	S288C		Details: 20% extended lifespan 	Muid KA, et al. (2019) PMID:31082499
DUG1	YFR044C	replicative lifespan: increased 	systematic mutation set 	large-scale survey	null Allele: dug1-Δ	S288C		Details: long-lived mutants, one standard deviation above the mean, were identified using an HTP sequencing-based yeast replicative lifespan screen (SEBYL), with pooled cells subject to four rounds of biotin-based sorting to enrich old vs young cells, followed by sequencing to ID mutants and abundance 	Yu R, et al. (2021) PMID:33790287
DUR3	YHL016C	replicative lifespan: increased 	systematic mutation set 	large-scale survey	null Allele: dur3-Δ	S288C		Details: long-lived mutants, one standard deviation above the mean, were identified using an HTP sequencing-based yeast replicative lifespan screen (SEBYL), with pooled cells subject to four rounds of biotin-based sorting to enrich old vs young cells, followed by sequencing to ID mutants and abundance 	Yu R, et al. (2021) PMID:33790287
EAF6	YJR082C	replicative lifespan: increased 	systematic mutation set 	large-scale survey	null Allele: eaf6-Δ	S288C		Details: long-lived mutants, one standard deviation above the mean, were identified using an HTP sequencing-based yeast replicative lifespan screen (SEBYL), with pooled cells subject to four rounds of biotin-based sorting to enrich old vs young cells, followed by sequencing to ID mutants and abundance 	Yu R, et al. (2021) PMID:33790287
EFT2	YDR385W	replicative lifespan: increased 	systematic mutation set 	large-scale survey	null Allele: eft2-Δ	S288C		Details: long-lived mutants, one standard deviation above the mean, were identified using an HTP sequencing-based yeast replicative lifespan screen (SEBYL), with pooled cells subject to four rounds of biotin-based sorting to enrich old vs young cells, followed by sequencing to ID mutants and abundance 	Yu R, et al. (2021) PMID:33790287
EGD2	YHR193C	replicative lifespan: increased 	heterozygous diploid	classical genetics	null Allele: egd2-Δ	Other			Perić M, et al. (2016) PMID:27346163
ELP4	YPL101W	replicative lifespan: increased 	large-scale survey	large-scale survey	null Allele: elp4-Δ	S288C			Managbanag JR, et al. (2008) PMID:19030232
ELP4	YPL101W	replicative lifespan: increased 	classical genetics	classical genetics	null Allele: elp4-Δ	S288C			Delaney JR, et al. (2011) PMID:21902802
ELP4	YPL101W	replicative lifespan: increased 	competitive growth	large-scale survey	null Allele: elp4-Δ	S288C			Delaney JR, et al. (2011) PMID:21191185
ELP4	YPL101W	replicative lifespan: increased 	classical genetics	classical genetics	null Allele: elp4-Δ	S288C			McCormick MA, et al. (2015) PMID:26456335
EMP24	YGL200C	replicative lifespan: increased 	classical genetics	classical genetics	null Allele: emp24-Δ	S288C			McCormick MA, et al. (2015) PMID:26456335
EMP70	YLR083C	replicative lifespan: increased 	systematic mutation set 	large-scale survey	null Allele: emp70-Δ	S288C		Details: long-lived mutants, one standard deviation above the mean, were identified using an HTP sequencing-based yeast replicative lifespan screen (SEBYL), with pooled cells subject to four rounds of biotin-based sorting to enrich old vs young cells, followed by sequencing to ID mutants and abundance 	Yu R, et al. (2021) PMID:33790287
ENO1	YGR254W	replicative lifespan: increased 	classical genetics	classical genetics	null Allele: eno1-Δ	S288C			McCormick MA, et al. (2015) PMID:26456335
ENT2	YLR206W	replicative lifespan: increased 	systematic mutation set 	large-scale survey	null Allele: ent2-Δ	S288C		Details: long-lived mutants, one standard deviation above the mean, were identified using an HTP sequencing-based yeast replicative lifespan screen (SEBYL), with pooled cells subject to four rounds of biotin-based sorting to enrich old vs young cells, followed by sequencing to ID mutants and abundance 	Yu R, et al. (2021) PMID:33790287
ENT3	YJR125C	replicative lifespan: increased 	systematic mutation set 	large-scale survey	null Allele: ent3-Δ	S288C		Details: long-lived mutants, one standard deviation above the mean, were identified using an HTP sequencing-based yeast replicative lifespan screen (SEBYL), with pooled cells subject to four rounds of biotin-based sorting to enrich old vs young cells, followed by sequencing to ID mutants and abundance 	Yu R, et al. (2021) PMID:33790287
ENT5	YDR153C	replicative lifespan: increased 	systematic mutation set 	large-scale survey	null Allele: ent5-Δ	S288C		Details: long-lived mutants, one standard deviation above the mean, were identified using an HTP sequencing-based yeast replicative lifespan screen (SEBYL), with pooled cells subject to four rounds of biotin-based sorting to enrich old vs young cells, followed by sequencing to ID mutants and abundance 	Yu R, et al. (2021) PMID:33790287
EOS1	YNL080C	replicative lifespan: increased 	classical genetics	classical genetics	null Allele: eos1-Δ	S288C			McCormick MA, et al. (2015) PMID:26456335
EPS1	YIL005W	replicative lifespan: increased 	systematic mutation set 	large-scale survey	null Allele: eps1-Δ	S288C		Details: long-lived mutants, one standard deviation above the mean, were identified using an HTP sequencing-based yeast replicative lifespan screen (SEBYL), with pooled cells subject to four rounds of biotin-based sorting to enrich old vs young cells, followed by sequencing to ID mutants and abundance 	Yu R, et al. (2021) PMID:33790287
ERC1	YHR032W	replicative lifespan: increased 	systematic mutation set 	large-scale survey	null Allele: erc1-Δ	S288C		Details: long-lived mutants, one standard deviation above the mean, were identified using an HTP sequencing-based yeast replicative lifespan screen (SEBYL), with pooled cells subject to four rounds of biotin-based sorting to enrich old vs young cells, followed by sequencing to ID mutants and abundance 	Yu R, et al. (2021) PMID:33790287
ERO1	YML130C	replicative lifespan: increased 	classical genetics	classical genetics	reduction of function	S288C			Labunskyy VM, et al. (2014) PMID:24391512
ERT1	YBR239C	replicative lifespan: increased 	systematic mutation set 	large-scale survey	null Allele: ert1-Δ	S288C		Details: long-lived mutants, one standard deviation above the mean, were identified using an HTP sequencing-based yeast replicative lifespan screen (SEBYL), with pooled cells subject to four rounds of biotin-based sorting to enrich old vs young cells, followed by sequencing to ID mutants and abundance 	Yu R, et al. (2021) PMID:33790287
ERV14	YGL054C	replicative lifespan: increased 	systematic mutation set 	large-scale survey	null Allele: erv14-Δ	S288C		Details: long-lived mutants, one standard deviation above the mean, were identified using an HTP sequencing-based yeast replicative lifespan screen (SEBYL), with pooled cells subject to four rounds of biotin-based sorting to enrich old vs young cells, followed by sequencing to ID mutants and abundance 	Yu R, et al. (2021) PMID:33790287
ERV25	YML012W	replicative lifespan: increased 	classical genetics	classical genetics	null Allele: erv25-Δ	S288C			McCormick MA, et al. (2015) PMID:26456335
ERV41	YML067C	replicative lifespan: increased 	classical genetics	classical genetics	null Allele: erv41-Δ	S288C			McCormick MA, et al. (2015) PMID:26456335
EUG1	YDR518W	replicative lifespan: increased 	classical genetics	classical genetics	null Allele: eug1-Δ	S288C			Labunskyy VM, et al. (2014) PMID:24391512
EUG1	YDR518W	replicative lifespan: increased 	classical genetics	classical genetics	null Allele: eug1-Δ	S288C			McCormick MA, et al. (2015) PMID:26456335
FCY2	YER056C	replicative lifespan: increased 	classical genetics	classical genetics	null Allele: fcy2-Δ	S288C			McCormick MA, et al. (2015) PMID:26456335
FET3	YMR058W	replicative lifespan: increased 	classical genetics	classical genetics	null Allele: fet3-Δ	S288C			McCormick MA, et al. (2015) PMID:26456335
FIT2	YOR382W	replicative lifespan: increased 	classical genetics	classical genetics	null Allele: fit2-Δ	S288C			McCormick MA, et al. (2015) PMID:26456335
FLC2	YAL053W	replicative lifespan: increased 	systematic mutation set 	large-scale survey	null Allele: flc2-Δ	S288C		Details: long-lived mutants, one standard deviation above the mean, were identified using an HTP sequencing-based yeast replicative lifespan screen (SEBYL), with pooled cells subject to four rounds of biotin-based sorting to enrich old vs young cells, followed by sequencing to ID mutants and abundance 	Yu R, et al. (2021) PMID:33790287
FLO1	YAR050W	replicative lifespan: increased 	systematic mutation set 	large-scale survey	null Allele: flo1-Δ	S288C		Details: long-lived mutants, one standard deviation above the mean, were identified using an HTP sequencing-based yeast replicative lifespan screen (SEBYL), with pooled cells subject to four rounds of biotin-based sorting to enrich old vs young cells, followed by sequencing to ID mutants and abundance 	Yu R, et al. (2021) PMID:33790287
FMP16	YDR070C	replicative lifespan: increased 	systematic mutation set 	large-scale survey	null Allele: fmp16-Δ	S288C		Details: long-lived mutants, one standard deviation above the mean, were identified using an HTP sequencing-based yeast replicative lifespan screen (SEBYL), with pooled cells subject to four rounds of biotin-based sorting to enrich old vs young cells, followed by sequencing to ID mutants and abundance 	Yu R, et al. (2021) PMID:33790287
FMP40	YPL222W	replicative lifespan: increased 	systematic mutation set 	large-scale survey	null Allele: fmp40-Δ	S288C		Details: long-lived mutants, one standard deviation above the mean, were identified using an HTP sequencing-based yeast replicative lifespan screen (SEBYL), with pooled cells subject to four rounds of biotin-based sorting to enrich old vs young cells, followed by sequencing to ID mutants and abundance 	Yu R, et al. (2021) PMID:33790287
FMP40	YPL222W	replicative lifespan: increased 	classical genetics 	classical genetics	null Allele: fmp40-Δ	S288C		Details: long-lived mutant assayed using a microfluidic chip-based lifespan measurement with automatic imaging to validate SEBYL 	Yu R, et al. (2021) PMID:33790287
FMP42	YMR221C	replicative lifespan: increased 	classical genetics	classical genetics	null Allele: fmp42-Δ	S288C			McCormick MA, et al. (2015) PMID:26456335
FMP52	YER004W	replicative lifespan: increased 	systematic mutation set 	large-scale survey	null Allele: fmp52-Δ	S288C		Details: long-lived mutants, one standard deviation above the mean, were identified using an HTP sequencing-based yeast replicative lifespan screen (SEBYL), with pooled cells subject to four rounds of biotin-based sorting to enrich old vs young cells, followed by sequencing to ID mutants and abundance 	Yu R, et al. (2021) PMID:33790287
FMS1	YMR020W	replicative lifespan: increased 	systematic mutation set 	large-scale survey	null Allele: fms1-Δ	S288C		Details: long-lived mutants, one standard deviation above the mean, were identified using an HTP sequencing-based yeast replicative lifespan screen (SEBYL), with pooled cells subject to four rounds of biotin-based sorting to enrich old vs young cells, followed by sequencing to ID mutants and abundance 	Yu R, et al. (2021) PMID:33790287
FOB1	YDR110W	replicative lifespan: increased 	classical genetics	classical genetics	null Allele: fob1-Δ	S288C			Kaeberlein M, et al. (2005) PMID:16293764
FOB1	YDR110W	replicative lifespan: increased 	large-scale survey	large-scale survey	null Allele: fob1-Δ	S288C			Kaeberlein M, et al. (2005) PMID:16293764
FOB1	YDR110W	replicative lifespan: increased 	homozygous diploid	classical genetics	null Allele: fob1-Δ	SK1		Details: replicative lifespan increased from 19.55 generations to 23.57 generations 	Boselli M, et al. (2009) PMID:19531355
FOB1	YDR110W	replicative lifespan: increased 	classical genetics 	classical genetics	null Allele: fob1-Δ	S288C		Treatment: simulated microgravity, 10 rpm Details: abrogation of the reduced RLS resulting from simulated microgravity in a uniaxial clinostat, and assayed using the Mother Enrichment Program 	Fukuda APM, et al. (2021) PMID:33612178
FOB1	YDR110W	replicative lifespan: increased 	classical genetics	classical genetics	null Allele: fob1-Δ	Other		Details: 70% increase in average replicative lifespan; mutant displays decreased ERC (extrachromosomal rDNA circle) formation 	Defossez PA, et al. (1999) PMID:10230397
FOB1	YDR110W	replicative lifespan: increased 	classical genetics	classical genetics	null Allele: fob1-Δ	S288C			Delaney JR, et al. (2011) PMID:21902802
FOB1	YDR110W	replicative lifespan: increased 	competitive growth	large-scale survey	null Allele: fob1-Δ	S288C			Delaney JR, et al. (2011) PMID:21191185
FOB1	YDR110W	replicative lifespan: increased 	classical genetics	classical genetics	null Allele: fob1-Δ	S288C		Treatment: caloric restriction by glucose reduction, 0.05% Details: dietary restriction-induced lifespan extension is increased relative to wt 	Schleit J, et al. (2013) PMID:23837470
FOB1	YDR110W	replicative lifespan: increased 	systematic mutation set	large-scale survey	null Allele: fob1-Δ	S288C			Delaney JR, et al. (2013) PMID:23336757
FOB1	YDR110W	replicative lifespan: increased 	classical genetics	classical genetics	null Allele: fob1-Δ	W303			Zaman S, et al. (2016) PMID:26951198
FOB1	YDR110W	replicative lifespan: increased 	classical genetics	classical genetics	null Allele: fob1-Δ	Other		Details: decreased recombination at rDNA locus 	Hoopes LL, et al. (2002) PMID:12024027
FOB1	YDR110W	replicative lifespan: increased 	classical genetics	classical genetics	null Allele: fob1-Δ	S288C			Peng J, et al. (2015) PMID:25822194
FOB1	YDR110W	replicative lifespan: increased 	classical genetics	classical genetics	null Allele: fob1-Δ	S288C			McCormick MA, et al. (2015) PMID:26456335
FOB1	YDR110W	replicative lifespan: increased 	classical genetics 	classical genetics	null Allele: fob1-Δ	S288C			Kaeberlein M, et al. (2004) PMID:15328540
FOB1	YDR110W	replicative lifespan: increased 	classical genetics	classical genetics	unspecified Allele: fob1-(hrm1-1) C246R	Other		Details: 70% increase in average replicative lifespan; mutant displays decreased ERC (extrachromosomal rDNA circle) formation 	Defossez PA, et al. (1999) PMID:10230397
FOB1	YDR110W	replicative lifespan: increased 	classical genetics	classical genetics	reduction of function Allele: fob1-AAA S467A, S468A, S519A	W303			Zaman S, et al. (2016) PMID:26951198
FRE6	YLL051C	replicative lifespan: increased 	classical genetics	classical genetics	null Allele: fre6-Δ	S288C		Details: replicative lifespan is decreased during dietary restriction (0.5% glucose) 	Wuttke D, et al. (2012) PMID:22912585
FRE8	YLR047C	replicative lifespan: increased 	systematic mutation set 	large-scale survey	null Allele: fre8-Δ	S288C		Details: long-lived mutants, one standard deviation above the mean, were identified using an HTP sequencing-based yeast replicative lifespan screen (SEBYL), with pooled cells subject to four rounds of biotin-based sorting to enrich old vs young cells, followed by sequencing to ID mutants and abundance 	Yu R, et al. (2021) PMID:33790287
FUN30	YAL019W	replicative lifespan: increased 	systematic mutation set 	large-scale survey	null Allele: fun30-Δ	S288C		Details: long-lived mutants, one standard deviation above the mean, were identified using an HTP sequencing-based yeast replicative lifespan screen (SEBYL), with pooled cells subject to four rounds of biotin-based sorting to enrich old vs young cells, followed by sequencing to ID mutants and abundance 	Yu R, et al. (2021) PMID:33790287
FZF1	YGL254W	replicative lifespan: increased 	classical genetics	classical genetics	null Allele: fzf1-Δ	S288C			Yoshida R, et al. (2010) PMID:20550517
FZO1	YBR179C	replicative lifespan: increased 	classical genetics	classical genetics	null Allele: fzo1-Δ	S288C		Treatment: caloric restriction by glucose reduction, 0.05% Details: dietary restriction-induced lifespan extension is increased relative to wt 	Schleit J, et al. (2013) PMID:23837470
FZO1	YBR179C	replicative lifespan: increased 	classical genetics	classical genetics	null Allele: fzo1-Δ	Other		Details: mutant mother cell has increased replicative lifespan compared to its 7th generation daughter and to wild type; age asymmetry between mother and 7th daughter not seen in wild type 	Jiang JC, et al. (2019) PMID:30298458
GAD1	YMR250W	replicative lifespan: increased 	classical genetics	classical genetics	null Allele: gad1-Δ	S288C			Kamei Y, et al. (2011) PMID:21371425
GAS4	YOL132W	replicative lifespan: increased 	classical genetics	classical genetics	null Allele: gas4-Δ	S288C			McCormick MA, et al. (2015) PMID:26456335
GCN2	YDR283C	replicative lifespan: increased 	classical genetics	classical genetics	null Allele: gcn2-Δ	S288C		Treatment: caloric restriction by glucose reduction, 0.05% Details: dietary restriction-induced lifespan extension is increased relative to wt 	Schleit J, et al. (2013) PMID:23837470
GCN4	YEL009C	replicative lifespan: increased 	large-scale survey	large-scale survey	null Allele: gcn4-Δ	S288C			Managbanag JR, et al. (2008) PMID:19030232
GCN4	YEL009C	replicative lifespan: increased 	classical genetics 	classical genetics	overexpression	Other			Hu Z, et al. (2018) PMID:30117416
GCN5	YGR252W	replicative lifespan: increased 	heterozygous diploid 	classical genetics	null Allele: gcn5-Δ	S288C		Details: significantly extended replicative lifespan 	Huang B, et al. (2020) PMID:32157780
GEM1	YAL048C	replicative lifespan: increased 	classical genetics	classical genetics	null Allele: gem1-Δ	S288C			McCormick MA, et al. (2015) PMID:26456335
GIR2	YDR152W	replicative lifespan: increased 	systematic mutation set 	large-scale survey	null Allele: gir2-Δ	S288C		Details: long-lived mutants, one standard deviation above the mean, were identified using an HTP sequencing-based yeast replicative lifespan screen (SEBYL), with pooled cells subject to four rounds of biotin-based sorting to enrich old vs young cells, followed by sequencing to ID mutants and abundance 	Yu R, et al. (2021) PMID:33790287
GPA2	YER020W	replicative lifespan: increased 	classical genetics	classical genetics	null Allele: gpa2-Δ	Other			Lin SJ, et al. (2000) PMID:11000115
GPA2	YER020W	replicative lifespan: increased 	classical genetics	classical genetics	null Allele: gpa2-Δ	S288C			Delaney JR, et al. (2011) PMID:21902802
GPA2	YER020W	replicative lifespan: increased 	competitive growth	large-scale survey	null Allele: gpa2-Δ	S288C			Delaney JR, et al. (2011) PMID:21191185
GPA2	YER020W	replicative lifespan: increased 	classical genetics	classical genetics	null Allele: gpa2-Δ	S288C			McCormick MA, et al. (2015) PMID:26456335
GPA2	YER020W	replicative lifespan: increased 	classical genetics 	classical genetics	null Allele: gpa2-Δ	S288C			Kaeberlein M, et al. (2004) PMID:15328540
GPM2	YDL021W	replicative lifespan: increased 	classical genetics	classical genetics	null Allele: gpm2-Δ	S288C			McCormick MA, et al. (2015) PMID:26456335
GPR1	YDL035C	replicative lifespan: increased 	classical genetics	classical genetics	null Allele: gpr1-Δ	Other			Lin SJ, et al. (2000) PMID:11000115
GPR1	YDL035C	replicative lifespan: increased 	competitive growth	large-scale survey	null Allele: gpr1-Δ	S288C			Delaney JR, et al. (2011) PMID:21191185
GPR1	YDL035C	replicative lifespan: increased 	systematic mutation set	large-scale survey	null Allele: gpr1-Δ	S288C			Delaney JR, et al. (2013) PMID:23336757
GPR1	YDL035C	replicative lifespan: increased 	classical genetics	classical genetics	null Allele: gpr1-Δ	S288C			McCormick MA, et al. (2015) PMID:26456335
GPR1	YDL035C	replicative lifespan: increased 	classical genetics 	classical genetics	null Allele: gpr1-Δ	S288C			Kaeberlein M, et al. (2004) PMID:15328540
GPR1	YDL035C	replicative lifespan: increased 	systematic mutation set 	large-scale survey	null Allele: gpr1-Δ	S288C		Details: long-lived mutants, one standard deviation above the mean, were identified using an HTP sequencing-based yeast replicative lifespan screen (SEBYL), with pooled cells subject to four rounds of biotin-based sorting to enrich old vs young cells, followed by sequencing to ID mutants and abundance 	Yu R, et al. (2021) PMID:33790287
GPX1	YKL026C	replicative lifespan: increased 	classical genetics	classical genetics	null Allele: gpx1-Δ	S288C		Treatment: caloric restriction by glucose reduction, 0.05% Details: dietary restriction-induced lifespan extension is increased relative to wt 	Schleit J, et al. (2013) PMID:23837470
GSH2	YOL049W	replicative lifespan: increased 	classical genetics 	classical genetics	null Allele: gsh2-Δ	S288C		Details: long-lived mutant assayed using a microfluidic chip-based lifespan measurement with automatic imaging to validate SEBYL 	Yu R, et al. (2021) PMID:33790287
GSH2	YOL049W	replicative lifespan: increased 	systematic mutation set 	large-scale survey	null Allele: gsh2-Δ	S288C		Details: long-lived mutants, one standard deviation above the mean, were identified using an HTP sequencing-based yeast replicative lifespan screen (SEBYL), with pooled cells subject to four rounds of biotin-based sorting to enrich old vs young cells, followed by sequencing to ID mutants and abundance 	Yu R, et al. (2021) PMID:33790287
GSY1	YFR015C	replicative lifespan: increased 	classical genetics	classical genetics	null Allele: gsy1-Δ	S288C			McCormick MA, et al. (2015) PMID:26456335
GTA1	YEL043W	replicative lifespan: increased 	systematic mutation set 	large-scale survey	null Allele: gta1-Δ	S288C		Details: long-lived mutants, one standard deviation above the mean, were identified using an HTP sequencing-based yeast replicative lifespan screen (SEBYL), with pooled cells subject to four rounds of biotin-based sorting to enrich old vs young cells, followed by sequencing to ID mutants and abundance 	Yu R, et al. (2021) PMID:33790287
GTO3	YMR251W	replicative lifespan: increased 	classical genetics	classical genetics	null Allele: gto3-Δ	S288C			McCormick MA, et al. (2015) PMID:26456335
GTR1	YML121W	replicative lifespan: increased 	classical genetics	classical genetics	null Allele: gtr1-Δ	S288C			McCormick MA, et al. (2015) PMID:26456335
GTR1	YML121W	replicative lifespan: increased 	classical genetics 	classical genetics	null Allele: gtr1-Δ	S288C		Details: increase in RLS relative to wt; corylin, a flavinoid from Psoralea corylifolia was unable to further increase RLS 	Wang TH, et al. (2022) PMID:35264584
GTR1	YML121W	replicative lifespan: increased 	classical genetics 	classical genetics	repressible	S288C		Details: increase in RLS relative to wt; corylin, a flavinoid from Psoralea corylifolia was unable to further increase RLS 	Wang TH, et al. (2022) PMID:35264584
GTT2	YLL060C	replicative lifespan: increased 	systematic mutation set 	large-scale survey	null Allele: gtt2-Δ	S288C		Details: long-lived mutants, one standard deviation above the mean, were identified using an HTP sequencing-based yeast replicative lifespan screen (SEBYL), with pooled cells subject to four rounds of biotin-based sorting to enrich old vs young cells, followed by sequencing to ID mutants and abundance 	Yu R, et al. (2021) PMID:33790287
GUK1	YDR454C	replicative lifespan: increased 	classical genetics 	classical genetics	reduction of function Allele: guk1-1 G12S (G34A)	S288C		Details: significant extension with an increase in average lifespan from 23.4 days to 28.3 days 	Liu P, et al. (2020) PMID:32232569
GUT2	YIL155C	replicative lifespan: increased 	classical genetics	classical genetics	overexpression	Other		Media: glucose carbon source, 2% Details: replicative lifespan is 25% increased 	Easlon E, et al. (2008) PMID:18381895
GYP5	YPL249C	replicative lifespan: increased 	systematic mutation set 	large-scale survey	null Allele: gyp5-Δ	S288C		Details: long-lived mutants, one standard deviation above the mean, were identified using an HTP sequencing-based yeast replicative lifespan screen (SEBYL), with pooled cells subject to four rounds of biotin-based sorting to enrich old vs young cells, followed by sequencing to ID mutants and abundance 	Yu R, et al. (2021) PMID:33790287
HAP4	YKL109W	replicative lifespan: increased 	classical genetics	classical genetics	null Allele: hap4-Δ	S288C		Treatment: caloric restriction by glucose reduction, 0.05% Details: dietary restriction-induced lifespan extension is increased relative to wt 	Schleit J, et al. (2013) PMID:23837470
HAT2	YEL056W	replicative lifespan: increased 	classical genetics	classical genetics	null Allele: hat2-Δ	S288C			Rosaleny LE, et al. (2005) PMID:16023114
HBS1	YKR084C	replicative lifespan: increased 	systematic mutation set 	large-scale survey	null Allele: hbs1-Δ	S288C		Details: long-lived mutants, one standard deviation above the mean, were identified using an HTP sequencing-based yeast replicative lifespan screen (SEBYL), with pooled cells subject to four rounds of biotin-based sorting to enrich old vs young cells, followed by sequencing to ID mutants and abundance 	Yu R, et al. (2021) PMID:33790287
HBS1	YKR084C	replicative lifespan: increased 	classical genetics	classical genetics	null Allele: hbs1-Δ	S288C			McCormick MA, et al. (2015) PMID:26456335
HCM1	YCR065W	replicative lifespan: increased 	classical genetics	classical genetics	overexpression	S288C			Maoz N, et al. (2015) PMID:24835838
HDA1	YNL021W	replicative lifespan: increased 	systematic mutation set 	large-scale survey	null Allele: hda1-Δ	S288C		Details: long-lived mutants, one standard deviation above the mean, were identified using an HTP sequencing-based yeast replicative lifespan screen (SEBYL), with pooled cells subject to four rounds of biotin-based sorting to enrich old vs young cells, followed by sequencing to ID mutants and abundance 	Yu R, et al. (2021) PMID:33790287
HDA1	YNL021W	replicative lifespan: increased 	classical genetics 	classical genetics	null Allele: hda1-Δ	S288C		Details: long-lived mutant validated using both a microfluidic chip-based lifespan measurement with automatic imaging and a microscopic dissection assay 	Yu R, et al. (2021) PMID:33790287
HDA2	YDR295C	replicative lifespan: increased 	systematic mutation set 	large-scale survey	null Allele: hda2-Δ	S288C		Details: long-lived mutants, one standard deviation above the mean, were identified using an HTP sequencing-based yeast replicative lifespan screen (SEBYL), with pooled cells subject to four rounds of biotin-based sorting to enrich old vs young cells, followed by sequencing to ID mutants and abundance 	Yu R, et al. (2021) PMID:33790287
HDA2	YDR295C	replicative lifespan: increased 	classical genetics 	classical genetics	null Allele: hda2-Δ	S288C		Details: long-lived mutant validated using both a microfluidic chip-based lifespan measurement with automatic imaging and a microscopic dissection assay 	Yu R, et al. (2021) PMID:33790287
HDA3	YPR179C	replicative lifespan: increased 	classical genetics 	classical genetics	null Allele: hda3-Δ	S288C		Details: long-lived mutant validated using both a microfluidic chip-based lifespan measurement with automatic imaging and a microscopic dissection assay 	Yu R, et al. (2021) PMID:33790287
HEL2	YDR266C	replicative lifespan: increased 	systematic mutation set 	large-scale survey	null Allele: hel2-Δ	S288C		Details: long-lived mutants, one standard deviation above the mean, were identified using an HTP sequencing-based yeast replicative lifespan screen (SEBYL), with pooled cells subject to four rounds of biotin-based sorting to enrich old vs young cells, followed by sequencing to ID mutants and abundance 	Yu R, et al. (2021) PMID:33790287
HFA1	YMR207C	replicative lifespan: increased 	systematic mutation set 	large-scale survey	null Allele: hfa1-Δ	S288C		Details: long-lived mutants, one standard deviation above the mean, were identified using an HTP sequencing-based yeast replicative lifespan screen (SEBYL), with pooled cells subject to four rounds of biotin-based sorting to enrich old vs young cells, followed by sequencing to ID mutants and abundance 	Yu R, et al. (2021) PMID:33790287
HIR1	YBL008W	replicative lifespan: increased 	classical genetics	classical genetics	null Allele: hir1-Δ	S288C		Details: increased 25-35% 	Feser J, et al. (2010) PMID:20832724
HIR1	YBL008W	replicative lifespan: increased 	systematic mutation set 	large-scale survey	null Allele: hir1-Δ	S288C		Details: long-lived mutants, one standard deviation above the mean, were identified using an HTP sequencing-based yeast replicative lifespan screen (SEBYL), with pooled cells subject to four rounds of biotin-based sorting to enrich old vs young cells, followed by sequencing to ID mutants and abundance 	Yu R, et al. (2021) PMID:33790287
HIR2	YOR038C	replicative lifespan: increased 	classical genetics	classical genetics	null Allele: hir2-Δ	S288C		Details: increased 25-35% 	Feser J, et al. (2010) PMID:20832724
HIR3	YJR140C	replicative lifespan: increased 	classical genetics	classical genetics	null Allele: hir3-Δ	S288C		Details: increased 25-35% 	Feser J, et al. (2010) PMID:20832724
HMT1	YBR034C	replicative lifespan: increased 	classical genetics	classical genetics	null Allele: hmt1-Δ	S288C			McCormick MA, et al. (2015) PMID:26456335
HMT1	YBR034C	replicative lifespan: increased 	systematic mutation set 	large-scale survey	null Allele: hmt1-Δ	S288C		Details: long-lived mutants, one standard deviation above the mean, were identified using an HTP sequencing-based yeast replicative lifespan screen (SEBYL), with pooled cells subject to four rounds of biotin-based sorting to enrich old vs young cells, followed by sequencing to ID mutants and abundance 	Yu R, et al. (2021) PMID:33790287
HO	YDL227C	replicative lifespan: increased 	systematic mutation set 	large-scale survey	null Allele: ho-Δ	S288C		Details: long-lived mutants, one standard deviation above the mean, were identified using an HTP sequencing-based yeast replicative lifespan screen (SEBYL), with pooled cells subject to four rounds of biotin-based sorting to enrich old vs young cells, followed by sequencing to ID mutants and abundance 	Yu R, et al. (2021) PMID:33790287
HOR7	YMR251W-A	replicative lifespan: increased 	classical genetics	classical genetics	null Allele: hor7-Δ	S288C			McCormick MA, et al. (2015) PMID:26456335
HOS2	YGL194C	replicative lifespan: increased 	systematic mutation set 	large-scale survey	null Allele: hos2-Δ	S288C		Details: long-lived mutants, one standard deviation above the mean, were identified using an HTP sequencing-based yeast replicative lifespan screen (SEBYL), with pooled cells subject to four rounds of biotin-based sorting to enrich old vs young cells, followed by sequencing to ID mutants and abundance 	Yu R, et al. (2021) PMID:33790287
HPC2	YBR215W	replicative lifespan: increased 	classical genetics	classical genetics	null Allele: hpc2-Δ	S288C		Details: increased 25-35% 	Feser J, et al. (2010) PMID:20832724
HRD1	YOL013C	replicative lifespan: increased 	classical genetics	classical genetics	null Allele: hrd1-Δ	S288C			Labunskyy VM, et al. (2014) PMID:24391512
HRK1	YOR267C	replicative lifespan: increased 	systematic mutation set 	large-scale survey	null Allele: hrk1-Δ	S288C		Details: long-lived mutants, one standard deviation above the mean, were identified using an HTP sequencing-based yeast replicative lifespan screen (SEBYL), with pooled cells subject to four rounds of biotin-based sorting to enrich old vs young cells, followed by sequencing to ID mutants and abundance 	Yu R, et al. (2021) PMID:33790287
HRQ1	YDR291W	replicative lifespan: increased 	systematic mutation set 	large-scale survey	null Allele: hrq1-Δ	S288C		Details: long-lived mutants, one standard deviation above the mean, were identified using an HTP sequencing-based yeast replicative lifespan screen (SEBYL), with pooled cells subject to four rounds of biotin-based sorting to enrich old vs young cells, followed by sequencing to ID mutants and abundance 	Yu R, et al. (2021) PMID:33790287
HSE1	YHL002W	replicative lifespan: increased 	large-scale survey	large-scale survey	null Allele: hse1-Δ	S288C			Smith ED, et al. (2008) PMID:18340043
HSE1	YHL002W	replicative lifespan: increased 	competitive growth	large-scale survey	null Allele: hse1-Δ	S288C			Delaney JR, et al. (2011) PMID:21191185
HSE1	YHL002W	replicative lifespan: increased 	classical genetics	classical genetics	null Allele: hse1-Δ	S288C			McCormick MA, et al. (2015) PMID:26456335
HSE1	YHL002W	replicative lifespan: increased 	systematic mutation set 	large-scale survey	null Allele: hse1-Δ	S288C		Details: long-lived mutants, one standard deviation above the mean, were identified using an HTP sequencing-based yeast replicative lifespan screen (SEBYL), with pooled cells subject to four rounds of biotin-based sorting to enrich old vs young cells, followed by sequencing to ID mutants and abundance 	Yu R, et al. (2021) PMID:33790287
HSP26	YBR072W	replicative lifespan: increased 	classical genetics 	classical genetics	overexpression	S288C		Details: significantly longer replicative lifespan 	Campion R, et al. (2021) PMID:34661239
HTB2	YBL002W	replicative lifespan: increased 	systematic mutation set 	large-scale survey	null Allele: htb2-Δ	S288C		Details: long-lived mutants, one standard deviation above the mean, were identified using an HTP sequencing-based yeast replicative lifespan screen (SEBYL), with pooled cells subject to four rounds of biotin-based sorting to enrich old vs young cells, followed by sequencing to ID mutants and abundance 	Yu R, et al. (2021) PMID:33790287
HUA1	YGR268C	replicative lifespan: increased 	classical genetics	classical genetics	null Allele: hua1-Δ	S288C			McCormick MA, et al. (2015) PMID:26456335
HXK2	YGL253W	replicative lifespan: increased 	classical genetics	classical genetics	null Allele: hxk2-Δ	Other			Lin SJ, et al. (2000) PMID:11000115
HXK2	YGL253W	replicative lifespan: increased 	classical genetics	classical genetics	null Allele: hxk2-Δ	S288C			Delaney JR, et al. (2011) PMID:21902802
HXK2	YGL253W	replicative lifespan: increased 	classical genetics	classical genetics	null Allele: hxk2-Δ	S288C			McCormick MA, et al. (2015) PMID:26456335
HXK2	YGL253W	replicative lifespan: increased 	classical genetics 	classical genetics	null Allele: hxk2-Δ	S288C			Kaeberlein M, et al. (2004) PMID:15328540
HXK2	YGL253W	replicative lifespan: increased 	classical genetics 	classical genetics	null Allele: hxk2-Δ	S288C		Treatment: calorie restriction 	Maslanka R and Zadrag-Tecza R (2020) PMID:33022992
HXK2	YGL253W	replicative lifespan: increased 	classical genetics 	classical genetics	null Allele: hxk2-Δ	S288C		Treatment: optimal conditions 	Maslanka R and Zadrag-Tecza R (2020) PMID:33022992
HXK2	YGL253W	replicative lifespan: increased 	classical genetics 	classical genetics	null Allele: hxk2-Δ	S288C		Treatment: calorie excess 	Maslanka R and Zadrag-Tecza R (2020) PMID:33022992
HXT2	YMR011W	replicative lifespan: increased 	systematic mutation set 	large-scale survey	null Allele: hxt2-Δ	S288C		Details: long-lived mutants, one standard deviation above the mean, were identified using an HTP sequencing-based yeast replicative lifespan screen (SEBYL), with pooled cells subject to four rounds of biotin-based sorting to enrich old vs young cells, followed by sequencing to ID mutants and abundance 	Yu R, et al. (2021) PMID:33790287
HXT4	YHR092C	replicative lifespan: increased 	systematic mutation set 	large-scale survey	null Allele: hxt4-Δ	S288C		Details: long-lived mutants, one standard deviation above the mean, were identified using an HTP sequencing-based yeast replicative lifespan screen (SEBYL), with pooled cells subject to four rounds of biotin-based sorting to enrich old vs young cells, followed by sequencing to ID mutants and abundance 	Yu R, et al. (2021) PMID:33790287
ICS3	YJL077C	replicative lifespan: increased 	systematic mutation set 	large-scale survey	null Allele: ics3-Δ	S288C		Details: long-lived mutants, one standard deviation above the mean, were identified using an HTP sequencing-based yeast replicative lifespan screen (SEBYL), with pooled cells subject to four rounds of biotin-based sorting to enrich old vs young cells, followed by sequencing to ID mutants and abundance 	Yu R, et al. (2021) PMID:33790287
IDH1	YNL037C	replicative lifespan: increased 	large-scale survey	large-scale survey	null Allele: idh1-Δ	S288C			Smith ED, et al. (2008) PMID:18340043
IDH1	YNL037C	replicative lifespan: increased 	classical genetics	classical genetics	null Allele: idh1-Δ	S288C			Delaney JR, et al. (2011) PMID:21902802
IDH1	YNL037C	replicative lifespan: increased 	competitive growth	large-scale survey	null Allele: idh1-Δ	S288C			Delaney JR, et al. (2011) PMID:21191185
IDH1	YNL037C	replicative lifespan: increased 	classical genetics	classical genetics	null Allele: idh1-Δ	S288C			McCormick MA, et al. (2015) PMID:26456335
IDH2	YOR136W	replicative lifespan: increased 	classical genetics	classical genetics	null Allele: idh2-Δ	S288C			Kaeberlein M, et al. (2005) PMID:16293764
IDH2	YOR136W	replicative lifespan: increased 	large-scale survey	large-scale survey	null Allele: idh2-Δ	S288C			Smith ED, et al. (2008) PMID:18340043
IDH2	YOR136W	replicative lifespan: increased 	large-scale survey	large-scale survey	null Allele: idh2-Δ	S288C			Kaeberlein M, et al. (2005) PMID:16293764
IDH2	YOR136W	replicative lifespan: increased 	classical genetics	classical genetics	null Allele: idh2-Δ	S288C			Delaney JR, et al. (2011) PMID:21902802
IDH2	YOR136W	replicative lifespan: increased 	competitive growth	large-scale survey	null Allele: idh2-Δ	S288C			Delaney JR, et al. (2011) PMID:21191185
IDH2	YOR136W	replicative lifespan: increased 	systematic mutation set	large-scale survey	null Allele: idh2-Δ	S288C			Delaney JR, et al. (2013) PMID:23336757
IDH2	YOR136W	replicative lifespan: increased 	classical genetics	classical genetics	null Allele: idh2-Δ	S288C			McCormick MA, et al. (2015) PMID:26456335
IDP1	YDL066W	replicative lifespan: increased 	classical genetics	classical genetics	null Allele: idp1-Δ	S288C			McCormick MA, et al. (2015) PMID:26456335
IDP3	YNL009W	replicative lifespan: increased 	large-scale survey	large-scale survey	null Allele: idp3-Δ	S288C			Managbanag JR, et al. (2008) PMID:19030232
IES4	YOR189W	replicative lifespan: increased 	systematic mutation set 	large-scale survey	null Allele: ies4-Δ	S288C		Details: long-lived mutants, one standard deviation above the mean, were identified using an HTP sequencing-based yeast replicative lifespan screen (SEBYL), with pooled cells subject to four rounds of biotin-based sorting to enrich old vs young cells, followed by sequencing to ID mutants and abundance 	Yu R, et al. (2021) PMID:33790287
IFH1	YLR223C	replicative lifespan: increased 	classical genetics	classical genetics	reduction of function Allele: ifh1-S969A mutation blocks Ifh1p phosphorylation	S288C			Cai L, et al. (2013) PMID:24035395
IMG1	YCR046C	replicative lifespan: increased 	classical genetics	classical genetics	null Allele: img1-Δ	S288C			McCormick MA, et al. (2015) PMID:26456335
IMG2	YCR071C	replicative lifespan: increased 	classical genetics	classical genetics	null Allele: img2-Δ	S288C			McCormick MA, et al. (2015) PMID:26456335
IMT4	YNCE0006W	replicative lifespan: increased 	classical genetics 	classical genetics	overexpression	S288C			Hu Z, et al. (2018) PMID:30117416
INM1	YHR046C	replicative lifespan: increased 	large-scale survey	large-scale survey	null Allele: inm1-Δ	S288C			Kaeberlein M, et al. (2005) PMID:16293764
INP51	YIL002C	replicative lifespan: increased 	large-scale survey	large-scale survey	null Allele: inp51-Δ	S288C			Smith ED, et al. (2008) PMID:18340043
INP51	YIL002C	replicative lifespan: increased 	large-scale survey	large-scale survey	null Allele: inp51-Δ	S288C			Managbanag JR, et al. (2008) PMID:19030232
INP51	YIL002C	replicative lifespan: increased 	classical genetics	classical genetics	null Allele: inp51-Δ	S288C			Delaney JR, et al. (2011) PMID:21902802
INP51	YIL002C	replicative lifespan: increased 	competitive growth	large-scale survey	null Allele: inp51-Δ	S288C			Delaney JR, et al. (2011) PMID:21191185
INP51	YIL002C	replicative lifespan: increased 	classical genetics	classical genetics	null Allele: inp51-Δ	S288C			McCormick MA, et al. (2015) PMID:26456335
INP51	YIL002C	replicative lifespan: increased 	systematic mutation set 	large-scale survey	null Allele: inp51-Δ	S288C		Details: long-lived mutants, one standard deviation above the mean, were identified using an HTP sequencing-based yeast replicative lifespan screen (SEBYL), with pooled cells subject to four rounds of biotin-based sorting to enrich old vs young cells, followed by sequencing to ID mutants and abundance 	Yu R, et al. (2021) PMID:33790287
INP53	YOR109W	replicative lifespan: increased 	large-scale survey	large-scale survey	null Allele: inp53-Δ	S288C			Smith ED, et al. (2008) PMID:18340043
INP53	YOR109W	replicative lifespan: increased 	large-scale survey	large-scale survey	null Allele: inp53-Δ	S288C			Kaeberlein M, et al. (2005) PMID:16293764
INP53	YOR109W	replicative lifespan: increased 	competitive growth	large-scale survey	null Allele: inp53-Δ	S288C			Delaney JR, et al. (2011) PMID:21191185
INP53	YOR109W	replicative lifespan: increased 	classical genetics	classical genetics	null Allele: inp53-Δ	S288C			McCormick MA, et al. (2015) PMID:26456335
IPK1	YDR315C	replicative lifespan: increased 	large-scale survey	large-scale survey	null Allele: ipk1-Δ	S288C			Kaeberlein M, et al. (2005) PMID:16293764
IRC10	YOL015W	replicative lifespan: increased 	systematic mutation set 	large-scale survey	null Allele: irc10-Δ	S288C		Details: long-lived mutants, one standard deviation above the mean, were identified using an HTP sequencing-based yeast replicative lifespan screen (SEBYL), with pooled cells subject to four rounds of biotin-based sorting to enrich old vs young cells, followed by sequencing to ID mutants and abundance 	Yu R, et al. (2021) PMID:33790287
IRC10	YOL015W	replicative lifespan: increased 	classical genetics 	classical genetics	null Allele: irc10-Δ	S288C		Details: long-lived mutant assayed using a microfluidic chip-based lifespan measurement with automatic imaging to validate SEBYL 	Yu R, et al. (2021) PMID:33790287
IRC14	YOR135C	replicative lifespan: increased 	classical genetics	classical genetics	null Allele: irc14-Δ	S288C			McCormick MA, et al. (2015) PMID:26456335
IRC14	YOR135C	replicative lifespan: increased 	systematic mutation set 	large-scale survey	null Allele: irc14-Δ	S288C		Details: long-lived mutants, one standard deviation above the mean, were identified using an HTP sequencing-based yeast replicative lifespan screen (SEBYL), with pooled cells subject to four rounds of biotin-based sorting to enrich old vs young cells, followed by sequencing to ID mutants and abundance 	Yu R, et al. (2021) PMID:33790287
IRC21	YMR073C	replicative lifespan: increased 	classical genetics	classical genetics	null Allele: irc21-Δ	S288C			McCormick MA, et al. (2015) PMID:26456335
IRC21	YMR073C	replicative lifespan: increased 	systematic mutation set 	large-scale survey	null Allele: irc21-Δ	S288C		Details: long-lived mutants, one standard deviation above the mean, were identified using an HTP sequencing-based yeast replicative lifespan screen (SEBYL), with pooled cells subject to four rounds of biotin-based sorting to enrich old vs young cells, followed by sequencing to ID mutants and abundance 	Yu R, et al. (2021) PMID:33790287
ISW2	YOR304W	replicative lifespan: increased 	classical genetics	classical genetics	null Allele: isw2-Δ	S288C			McCormick MA, et al. (2015) PMID:26456335
ITR1	YDR497C	replicative lifespan: increased 	classical genetics	classical genetics	null Allele: itr1-Δ	S288C			McCormick MA, et al. (2015) PMID:26456335
IZH2	YOL002C	replicative lifespan: increased 	systematic mutation set 	large-scale survey	null Allele: izh2-Δ	S288C		Details: long-lived mutants, one standard deviation above the mean, were identified using an HTP sequencing-based yeast replicative lifespan screen (SEBYL), with pooled cells subject to four rounds of biotin-based sorting to enrich old vs young cells, followed by sequencing to ID mutants and abundance 	Yu R, et al. (2021) PMID:33790287
JSN1	YJR091C	replicative lifespan: increased 	systematic mutation set 	large-scale survey	null Allele: jsn1-Δ	S288C		Details: long-lived mutants, one standard deviation above the mean, were identified using an HTP sequencing-based yeast replicative lifespan screen (SEBYL), with pooled cells subject to four rounds of biotin-based sorting to enrich old vs young cells, followed by sequencing to ID mutants and abundance 	Yu R, et al. (2021) PMID:33790287
KAP120	YPL125W	replicative lifespan: increased 	classical genetics	classical genetics	null Allele: kap120-Δ	S288C			McCormick MA, et al. (2015) PMID:26456335
KIN2	YLR096W	replicative lifespan: increased 	systematic mutation set 	large-scale survey	null Allele: kin2-Δ	S288C		Details: long-lived mutants, one standard deviation above the mean, were identified using an HTP sequencing-based yeast replicative lifespan screen (SEBYL), with pooled cells subject to four rounds of biotin-based sorting to enrich old vs young cells, followed by sequencing to ID mutants and abundance 	Yu R, et al. (2021) PMID:33790287
KIN4	YOR233W	replicative lifespan: increased 	systematic mutation set 	large-scale survey	null Allele: kin4-Δ	S288C		Details: long-lived mutants, one standard deviation above the mean, were identified using an HTP sequencing-based yeast replicative lifespan screen (SEBYL), with pooled cells subject to four rounds of biotin-based sorting to enrich old vs young cells, followed by sequencing to ID mutants and abundance 	Yu R, et al. (2021) PMID:33790287
KIP2	YPL155C	replicative lifespan: increased 	classical genetics	classical genetics	null Allele: kip2-Δ	S288C			McCormick MA, et al. (2015) PMID:26456335
KXD1	YGL079W	replicative lifespan: increased 	classical genetics	classical genetics	null Allele: kxd1-Δ	S288C			McCormick MA, et al. (2015) PMID:26456335
LAG1	YHL003C	replicative lifespan: increased 	classical genetics	classical genetics	null Allele: lag1-Δ	Other		Details: 47% increase in mean life span 	D'mello NP, et al. (1994) PMID:8195187
LAG2	YOL025W	replicative lifespan: increased 	classical genetics	classical genetics	overexpression	Other			Childress AM, et al. (1996) PMID:8760941
LAT1	YNL071W	replicative lifespan: increased 	classical genetics	classical genetics	overexpression	S288C		Details: replicative lifespan is increased by 30% 	Easlon E, et al. (2007) PMID:17200108
LHS1	YKL073W	replicative lifespan: increased 	heterozygous diploid	classical genetics	null Allele: lhs1-Δ	Other			Perić M, et al. (2016) PMID:27346163
LIN1	YHR156C	replicative lifespan: increased 	large-scale survey	large-scale survey	null Allele: lin1-Δ	S288C			Kaeberlein M, et al. (2005) PMID:16293764
LIN1	YHR156C	replicative lifespan: increased 	classical genetics	classical genetics	null Allele: lin1-Δ	S288C			McCormick MA, et al. (2015) PMID:26456335
LOC1	YFR001W	replicative lifespan: increased 	large-scale survey	large-scale survey	null Allele: loc1-Δ	S288C			Managbanag JR, et al. (2008) PMID:19030232
LOC1	YFR001W	replicative lifespan: increased 	classical genetics	classical genetics	null Allele: loc1-Δ	S288C			Steffen KK, et al. (2008) PMID:18423200
LOS1	YKL205W	replicative lifespan: increased 	classical genetics	classical genetics	null Allele: los1-Δ	S288C			McCormick MA, et al. (2015) PMID:26456335
LOS1	YKL205W	replicative lifespan: increased 	classical genetics 	classical genetics	null Allele: los1-Δ	S288C		Details: ~20% increase in RLS 	Lord CL, et al. (2017) PMID:27932586
LOS1	YKL205W	replicative lifespan: increased 	systematic mutation set 	large-scale survey	null Allele: los1-Δ	S288C		Details: long-lived mutants, one standard deviation above the mean, were identified using an HTP sequencing-based yeast replicative lifespan screen (SEBYL), with pooled cells subject to four rounds of biotin-based sorting to enrich old vs young cells, followed by sequencing to ID mutants and abundance 	Yu R, et al. (2021) PMID:33790287
LPD1	YFL018C	replicative lifespan: increased 	classical genetics	classical genetics	null Allele: lpd1-Δ	S288C			McCormick MA, et al. (2015) PMID:26456335
LRG1	YDL240W	replicative lifespan: increased 	systematic mutation set 	large-scale survey	null Allele: lrg1-Δ	S288C		Details: long-lived mutants, one standard deviation above the mean, were identified using an HTP sequencing-based yeast replicative lifespan screen (SEBYL), with pooled cells subject to four rounds of biotin-based sorting to enrich old vs young cells, followed by sequencing to ID mutants and abundance 	Yu R, et al. (2021) PMID:33790287
LSB1	YGR136W	replicative lifespan: increased 	classical genetics	classical genetics	null Allele: lsb1-Δ	S288C			McCormick MA, et al. (2015) PMID:26456335
LSC1	YOR142W	replicative lifespan: increased 	systematic mutation set 	large-scale survey	null Allele: lsc1-Δ	S288C		Details: long-lived mutants, one standard deviation above the mean, were identified using an HTP sequencing-based yeast replicative lifespan screen (SEBYL), with pooled cells subject to four rounds of biotin-based sorting to enrich old vs young cells, followed by sequencing to ID mutants and abundance 	Yu R, et al. (2021) PMID:33790287
LTE1	YAL024C	replicative lifespan: increased 	systematic mutation set 	large-scale survey	null Allele: lte1-Δ	S288C		Details: long-lived mutants, one standard deviation above the mean, were identified using an HTP sequencing-based yeast replicative lifespan screen (SEBYL), with pooled cells subject to four rounds of biotin-based sorting to enrich old vs young cells, followed by sequencing to ID mutants and abundance 	Yu R, et al. (2021) PMID:33790287
LTV1	YKL143W	replicative lifespan: increased 	classical genetics	classical genetics	null Allele: ltv1-Δ	S288C			McCormick MA, et al. (2015) PMID:26456335
LYP1	YNL268W	replicative lifespan: increased 	classical genetics	classical genetics	null Allele: lyp1-Δ	S288C			McCormick MA, et al. (2015) PMID:26456335
LYS12	YIL094C	replicative lifespan: increased 	large-scale survey	large-scale survey	null Allele: lys12-Δ	S288C			Kaeberlein M, et al. (2005) PMID:16293764
LYS9	YNR050C	replicative lifespan: increased 	large-scale survey	large-scale survey	null Allele: lys9-Δ	S288C			Kaeberlein M, et al. (2005) PMID:16293764
MAD3	YJL013C	replicative lifespan: increased 	classical genetics	classical genetics	null Allele: mad3-Δ	S288C			McCormick MA, et al. (2015) PMID:26456335
MAP2	YBL091C	replicative lifespan: increased 	classical genetics	classical genetics	null Allele: map2-Δ	S288C			McCormick MA, et al. (2015) PMID:26456335
MCA1	YOR197W	replicative lifespan: increased 	classical genetics	classical genetics	overexpression	S288C			Hill SM, et al. (2014) PMID:24855027
MDH1	YKL085W	replicative lifespan: increased 	classical genetics	classical genetics	overexpression	Other		Media: glucose carbon source, 2% Details: replicative lifespan is 25% increased 	Easlon E, et al. (2008) PMID:18381895
MED2	YDL005C	replicative lifespan: increased 	classical genetics 	classical genetics	overexpression	S288C		Details: RLS is 12.5% longer relative to wt 	Zhao W, et al. (2020) PMID:33045248
MEF1	YLR069C	replicative lifespan: increased 	classical genetics	classical genetics	null Allele: mef1-Δ	S288C		Treatment: caloric restriction by glucose reduction, 0.05% Details: dietary restriction-induced lifespan extension is increased relative to wt 	Schleit J, et al. (2013) PMID:23837470
MEF2	YJL102W	replicative lifespan: increased 	classical genetics	classical genetics	null Allele: mef2-Δ	S288C			McCormick MA, et al. (2015) PMID:26456335
MEP2	YNL142W	replicative lifespan: increased 	classical genetics	classical genetics	null Allele: mep2-Δ	S288C			McCormick MA, et al. (2015) PMID:26456335
MET14	YKL001C	replicative lifespan: increased 	systematic mutation set 	large-scale survey	null Allele: met14-Δ	S288C		Details: long-lived mutants, one standard deviation above the mean, were identified using an HTP sequencing-based yeast replicative lifespan screen (SEBYL), with pooled cells subject to four rounds of biotin-based sorting to enrich old vs young cells, followed by sequencing to ID mutants and abundance 	Yu R, et al. (2021) PMID:33790287
MET3	YJR010W	replicative lifespan: increased 	classical genetics	classical genetics	null Allele: met3-Δ	S288C			McCormick MA, et al. (2015) PMID:26456335
MET6	YER091C	replicative lifespan: increased 	systematic mutation set 	large-scale survey	null Allele: met6-Δ	S288C		Details: long-lived mutants, one standard deviation above the mean, were identified using an HTP sequencing-based yeast replicative lifespan screen (SEBYL), with pooled cells subject to four rounds of biotin-based sorting to enrich old vs young cells, followed by sequencing to ID mutants and abundance 	Yu R, et al. (2021) PMID:33790287
MFA1	YDR461W	replicative lifespan: increased 	classical genetics 	classical genetics	null Allele: mfa1-Δ	S288C		Details: long-lived mutant assayed using a microfluidic chip-based lifespan measurement with automatic imaging to validate SEBYL 	Yu R, et al. (2021) PMID:33790287
MGM1	YOR211C	replicative lifespan: increased 	classical genetics	classical genetics	null Allele: mgm1-Δ	S288C		Treatment: caloric restriction by glucose reduction, 0.05% Details: dietary restriction-induced lifespan extension is increased relative to wt 	Schleit J, et al. (2013) PMID:23837470
MHO1	YJR008W	replicative lifespan: increased 	systematic mutation set 	large-scale survey	null Allele: mho1-Δ	S288C		Details: long-lived mutants, one standard deviation above the mean, were identified using an HTP sequencing-based yeast replicative lifespan screen (SEBYL), with pooled cells subject to four rounds of biotin-based sorting to enrich old vs young cells, followed by sequencing to ID mutants and abundance 	Yu R, et al. (2021) PMID:33790287
MIG1	YGL035C	replicative lifespan: increased 	classical genetics	classical genetics	null Allele: mig1-Δ	S288C			McCormick MA, et al. (2015) PMID:26456335
MIG2	YGL209W	replicative lifespan: increased 	classical genetics	classical genetics	null Allele: mig2-Δ	S288C			Harkness TA, et al. (2004) PMID:15514051
MIR1	YJR077C	replicative lifespan: increased 	systematic mutation set 	large-scale survey	null Allele: mir1-Δ	S288C		Details: long-lived mutants, one standard deviation above the mean, were identified using an HTP sequencing-based yeast replicative lifespan screen (SEBYL), with pooled cells subject to four rounds of biotin-based sorting to enrich old vs young cells, followed by sequencing to ID mutants and abundance 	Yu R, et al. (2021) PMID:33790287
MIY2	YPL191C	replicative lifespan: increased 	systematic mutation set 	large-scale survey	null Allele: miy2-Δ	S288C		Details: long-lived mutants, one standard deviation above the mean, were identified using an HTP sequencing-based yeast replicative lifespan screen (SEBYL), with pooled cells subject to four rounds of biotin-based sorting to enrich old vs young cells, followed by sequencing to ID mutants and abundance 	Yu R, et al. (2021) PMID:33790287
MKC7	YDR144C	replicative lifespan: increased 	systematic mutation set 	large-scale survey	null Allele: mkc7-Δ	S288C		Details: long-lived mutants, one standard deviation above the mean, were identified using an HTP sequencing-based yeast replicative lifespan screen (SEBYL), with pooled cells subject to four rounds of biotin-based sorting to enrich old vs young cells, followed by sequencing to ID mutants and abundance 	Yu R, et al. (2021) PMID:33790287
MKK1	YOR231W	replicative lifespan: increased 	systematic mutation set 	large-scale survey	null Allele: mkk1-Δ	S288C		Details: long-lived mutants, one standard deviation above the mean, were identified using an HTP sequencing-based yeast replicative lifespan screen (SEBYL), with pooled cells subject to four rounds of biotin-based sorting to enrich old vs young cells, followed by sequencing to ID mutants and abundance 	Yu R, et al. (2021) PMID:33790287
MLH3	YPL164C	replicative lifespan: increased 	systematic mutation set 	large-scale survey	null Allele: mlh3-Δ	S288C		Details: long-lived mutants, one standard deviation above the mean, were identified using an HTP sequencing-based yeast replicative lifespan screen (SEBYL), with pooled cells subject to four rounds of biotin-based sorting to enrich old vs young cells, followed by sequencing to ID mutants and abundance 	Yu R, et al. (2021) PMID:33790287
MLO50	YLR049C	replicative lifespan: increased 	systematic mutation set 	large-scale survey	null Allele: mlo50-Δ	S288C		Details: long-lived mutants, one standard deviation above the mean, were identified using an HTP sequencing-based yeast replicative lifespan screen (SEBYL), with pooled cells subject to four rounds of biotin-based sorting to enrich old vs young cells, followed by sequencing to ID mutants and abundance 	Yu R, et al. (2021) PMID:33790287
MLP2	YIL149C	replicative lifespan: increased 	systematic mutation set 	large-scale survey	null Allele: mlp2-Δ	S288C		Details: long-lived mutants, one standard deviation above the mean, were identified using an HTP sequencing-based yeast replicative lifespan screen (SEBYL), with pooled cells subject to four rounds of biotin-based sorting to enrich old vs young cells, followed by sequencing to ID mutants and abundance 	Yu R, et al. (2021) PMID:33790287
MNE1	YOR350C	replicative lifespan: increased 	systematic mutation set 	large-scale survey	null Allele: mne1-Δ	S288C		Details: long-lived mutants, one standard deviation above the mean, were identified using an HTP sequencing-based yeast replicative lifespan screen (SEBYL), with pooled cells subject to four rounds of biotin-based sorting to enrich old vs young cells, followed by sequencing to ID mutants and abundance 	Yu R, et al. (2021) PMID:33790287
MNN1	YER001W	replicative lifespan: increased 	classical genetics	classical genetics	null Allele: mnn1-Δ	S288C			McCormick MA, et al. (2015) PMID:26456335
MNN2	YBR015C	replicative lifespan: increased 	classical genetics	classical genetics	null Allele: mnn2-Δ	S288C			McCormick MA, et al. (2015) PMID:26456335
MPC3	YGR243W	replicative lifespan: increased 	classical genetics	classical genetics	null Allele: mpc3-Δ	S288C			McCormick MA, et al. (2015) PMID:26456335
MPD1	YOR288C	replicative lifespan: increased 	classical genetics	classical genetics	null Allele: mpd1-Δ	S288C			Labunskyy VM, et al. (2014) PMID:24391512
MPD1	YOR288C	replicative lifespan: increased 	classical genetics	classical genetics	null Allele: mpd1-Δ	S288C			McCormick MA, et al. (2015) PMID:26456335
MPT5	YGL178W	replicative lifespan: increased 	classical genetics	classical genetics	unspecified Allele: mpt5-(uth4-326)	Other			Kennedy BK, et al. (1995) PMID:7859289
MRE11	YMR224C	replicative lifespan: increased 	classical genetics	classical genetics	null Allele: mre11-Δ	S288C		Treatment: caloric restriction by glucose reduction, 0.05% Details: dietary restriction-induced lifespan extension is increased relative to wt 	Schleit J, et al. (2013) PMID:23837470
MRPL16	YBL038W	replicative lifespan: increased 	systematic mutation set 	large-scale survey	null Allele: mrpl16-Δ	S288C		Details: long-lived mutants, one standard deviation above the mean, were identified using an HTP sequencing-based yeast replicative lifespan screen (SEBYL), with pooled cells subject to four rounds of biotin-based sorting to enrich old vs young cells, followed by sequencing to ID mutants and abundance 	Yu R, et al. (2021) PMID:33790287
MRPL22	YNL177C	replicative lifespan: increased 	systematic mutation set 	large-scale survey	null Allele: mrpl22-Δ	S288C		Details: long-lived mutants, one standard deviation above the mean, were identified using an HTP sequencing-based yeast replicative lifespan screen (SEBYL), with pooled cells subject to four rounds of biotin-based sorting to enrich old vs young cells, followed by sequencing to ID mutants and abundance 	Yu R, et al. (2021) PMID:33790287
MRPL25	YGR076C	replicative lifespan: increased 	classical genetics	classical genetics	null Allele: mrpl25-Δ	S288C		Details: 60% increase in median lifespan; 71% increase in maximum lifespan 	Heeren G, et al. (2009) PMID:20157544
MRPL33	YMR286W	replicative lifespan: increased 	classical genetics	classical genetics	null Allele: mrpl33-Δ	S288C			McCormick MA, et al. (2015) PMID:26456335
MRPL4	YLR439W	replicative lifespan: increased 	classical genetics	classical genetics	null Allele: mrpl4-Δ	S288C		Treatment: caloric restriction by glucose reduction, 0.05% Details: dietary restriction-induced lifespan extension is increased relative to wt 	Schleit J, et al. (2013) PMID:23837470
MRPL40	YPL173W	replicative lifespan: increased 	classical genetics	classical genetics	null Allele: mrpl40-Δ	S288C			McCormick MA, et al. (2015) PMID:26456335
MRPL49	YJL096W	replicative lifespan: increased 	classical genetics	classical genetics	null Allele: mrpl49-Δ	S288C			McCormick MA, et al. (2015) PMID:26456335
MRT4	YKL009W	replicative lifespan: increased 	classical genetics	classical genetics	null Allele: mrt4-Δ	S288C			McCormick MA, et al. (2015) PMID:26456335
MRX14	YDR115W	replicative lifespan: increased 	classical genetics	classical genetics	null Allele: mrx14-Δ	S288C			McCormick MA, et al. (2015) PMID:26456335
MSK1	YNL073W	replicative lifespan: increased 	classical genetics	classical genetics	null Allele: msk1-Δ	S288C			McCormick MA, et al. (2015) PMID:26456335
MSN5	YDR335W	replicative lifespan: increased 	classical genetics 	classical genetics	null Allele: msn5-Δ	S288C		Details: ~20% increase in RLS 	Lord CL, et al. (2017) PMID:27932586
MSP1	YGR028W	replicative lifespan: increased 	systematic mutation set 	large-scale survey	null Allele: msp1-Δ	S288C		Details: long-lived mutants, one standard deviation above the mean, were identified using an HTP sequencing-based yeast replicative lifespan screen (SEBYL), with pooled cells subject to four rounds of biotin-based sorting to enrich old vs young cells, followed by sequencing to ID mutants and abundance 	Yu R, et al. (2021) PMID:33790287
MSW1	YDR268W	replicative lifespan: increased 	large-scale survey	large-scale survey	null Allele: msw1-Δ	S288C			Managbanag JR, et al. (2008) PMID:19030232
MSW1	YDR268W	replicative lifespan: increased 	classical genetics	classical genetics	null Allele: msw1-Δ	S288C			Delaney JR, et al. (2011) PMID:21902802
MSW1	YDR268W	replicative lifespan: increased 	competitive growth	large-scale survey	null Allele: msw1-Δ	S288C			Delaney JR, et al. (2011) PMID:21191185
MSW1	YDR268W	replicative lifespan: increased 	classical genetics	classical genetics	null Allele: msw1-Δ	S288C			McCormick MA, et al. (2015) PMID:26456335
MTC2	YKL098W	replicative lifespan: increased 	classical genetics	classical genetics	null Allele: mtc2-Δ	S288C			McCormick MA, et al. (2015) PMID:26456335
MTC4	YBR255W	replicative lifespan: increased 	classical genetics	classical genetics	null Allele: mtc4-Δ	S288C			Kaeberlein M, et al. (2005) PMID:16293764
MTC4	YBR255W	replicative lifespan: increased 	large-scale survey	large-scale survey	null Allele: mtc4-Δ	S288C			Kaeberlein M, et al. (2005) PMID:16293764
MTC4	YBR255W	replicative lifespan: increased 	classical genetics	classical genetics	null Allele: mtc4-Δ	S288C			Delaney JR, et al. (2011) PMID:21902802
MTC4	YBR255W	replicative lifespan: increased 	competitive growth	large-scale survey	null Allele: mtc4-Δ	S288C			Delaney JR, et al. (2011) PMID:21191185
MTC4	YBR255W	replicative lifespan: increased 	classical genetics	classical genetics	null Allele: mtc4-Δ	S288C			McCormick MA, et al. (2015) PMID:26456335
MTO1	YGL236C	replicative lifespan: increased 	systematic mutation set 	large-scale survey	null Allele: mto1-Δ	S288C		Details: long-lived mutants, one standard deviation above the mean, were identified using an HTP sequencing-based yeast replicative lifespan screen (SEBYL), with pooled cells subject to four rounds of biotin-based sorting to enrich old vs young cells, followed by sequencing to ID mutants and abundance 	Yu R, et al. (2021) PMID:33790287
MTR10	YOR160W	replicative lifespan: increased 	classical genetics	classical genetics	overexpression	S288C			McCormick MA, et al. (2015) PMID:26456335
MUB1	YMR100W	replicative lifespan: increased 	classical genetics	classical genetics	null Allele: mub1-Δ	S288C			Kruegel U, et al. (2011) PMID:21931558
MUB1	YMR100W	replicative lifespan: increased 	classical genetics	classical genetics	null Allele: mub1-Δ	S288C			McCormick MA, et al. (2015) PMID:26456335
MVP1	YMR004W	replicative lifespan: increased 	systematic mutation set 	large-scale survey	null Allele: mvp1-Δ	S288C		Details: long-lived mutants, one standard deviation above the mean, were identified using an HTP sequencing-based yeast replicative lifespan screen (SEBYL), with pooled cells subject to four rounds of biotin-based sorting to enrich old vs young cells, followed by sequencing to ID mutants and abundance 	Yu R, et al. (2021) PMID:33790287
MXR1	YER042W	replicative lifespan: increased 	classical genetics	classical genetics	overexpression	S288C		Details: 25% increase in replicative lifespan 	Koc A, et al. (2004) PMID:15141092
MXR2	YCL033C	replicative lifespan: increased 	classical genetics	classical genetics	overexpression	S288C		Details: caloric restriction via growth in reduced glucose concentration; 0.5% 	Koc A, et al. (2004) PMID:15141092
MXR2	YCL033C	replicative lifespan: increased 	classical genetics	classical genetics	null Allele: mxr2-Δ	Other	  lactate	Details: lifespan extended; 25% 	Kaya A, et al. (2010) PMID:20799725
MYO5	YMR109W	replicative lifespan: increased 	systematic mutation set 	large-scale survey	null Allele: myo5-Δ	S288C		Details: long-lived mutants, one standard deviation above the mean, were identified using an HTP sequencing-based yeast replicative lifespan screen (SEBYL), with pooled cells subject to four rounds of biotin-based sorting to enrich old vs young cells, followed by sequencing to ID mutants and abundance 	Yu R, et al. (2021) PMID:33790287
NBA1	YOL070C	replicative lifespan: increased 	systematic mutation set 	large-scale survey	null Allele: nba1-Δ	S288C		Details: long-lived mutants, one standard deviation above the mean, were identified using an HTP sequencing-based yeast replicative lifespan screen (SEBYL), with pooled cells subject to four rounds of biotin-based sorting to enrich old vs young cells, followed by sequencing to ID mutants and abundance 	Yu R, et al. (2021) PMID:33790287
NCR1	YPL006W	replicative lifespan: increased 	classical genetics	classical genetics	null Allele: ncr1-Δ	S288C		Treatment: caloric restriction by glucose reduction, 0.05% Details: dietary restriction-induced lifespan extension is increased relative to wt 	Schleit J, et al. (2013) PMID:23837470
NCS2	YNL119W	replicative lifespan: increased 	classical genetics	classical genetics	null Allele: ncs2-Δ	S288C			McCormick MA, et al. (2015) PMID:26456335
NGG1	YDR176W	replicative lifespan: increased 	heterozygous diploid 	classical genetics	null Allele: ngg1-Δ	S288C		Details: significantly extended replicative lifespan 	Huang B, et al. (2020) PMID:32157780
NHX1	YDR456W	replicative lifespan: increased 	classical genetics	classical genetics	null Allele: nhx1-Δ	S288C		Treatment: caloric restriction by glucose reduction, 0.05% Details: dietary restriction-induced lifespan extension is increased relative to wt 	Schleit J, et al. (2013) PMID:23837470
NHX1	YDR456W	replicative lifespan: increased 	systematic mutation set 	large-scale survey	null Allele: nhx1-Δ	S288C		Details: long-lived mutants, one standard deviation above the mean, were identified using an HTP sequencing-based yeast replicative lifespan screen (SEBYL), with pooled cells subject to four rounds of biotin-based sorting to enrich old vs young cells, followed by sequencing to ID mutants and abundance 	Yu R, et al. (2021) PMID:33790287
NIS1	YNL078W	replicative lifespan: increased 	systematic mutation set 	large-scale survey	null Allele: nis1-Δ	S288C		Details: long-lived mutants, one standard deviation above the mean, were identified using an HTP sequencing-based yeast replicative lifespan screen (SEBYL), with pooled cells subject to four rounds of biotin-based sorting to enrich old vs young cells, followed by sequencing to ID mutants and abundance 	Yu R, et al. (2021) PMID:33790287
NMD4	YLR363C	replicative lifespan: increased 	large-scale survey	large-scale survey	null Allele: nmd4-Δ	S288C			Kaeberlein M, et al. (2005) PMID:16293764
NNT1	YLR285W	replicative lifespan: increased 	classical genetics	classical genetics	overexpression	W303		Details: replicative lifespan is increased but not further enhanced by caloric restriction 	Anderson RM, et al. (2003) PMID:12736687
NOP12	YOL041C	replicative lifespan: increased 	classical genetics	classical genetics	null Allele: nop12-Δ	S288C			Steffen KK, et al. (2008) PMID:18423200
NUP100	YKL068W	replicative lifespan: increased 	classical genetics 	classical genetics	null Allele: nup100-Δ	S288C		Details: ~20% increase in RLS 	Lord CL, et al. (2015) PMID:25778920
NUP170	YBL079W	replicative lifespan: increased 	classical genetics	classical genetics	null Allele: nup170-Δ	S288C		Treatment: caloric restriction by glucose reduction, 0.05% Details: dietary restriction-induced lifespan extension is increased relative to wt 	Schleit J, et al. (2013) PMID:23837470
NUP60	YAR002W	replicative lifespan: increased 	classical genetics 	classical genetics	reduction of function Allele: nup60-K467R non-acetylatable mutant	S288C			Meinema AC, et al. (2022) PMID:35373738
OCA1	YNL099C	replicative lifespan: increased 	systematic mutation set 	large-scale survey	null Allele: oca1-Δ	S288C		Details: long-lived mutants, one standard deviation above the mean, were identified using an HTP sequencing-based yeast replicative lifespan screen (SEBYL), with pooled cells subject to four rounds of biotin-based sorting to enrich old vs young cells, followed by sequencing to ID mutants and abundance 	Yu R, et al. (2021) PMID:33790287
OCA2	YNL056W	replicative lifespan: increased 	systematic mutation set 	large-scale survey	null Allele: oca2-Δ	S288C		Details: long-lived mutants, one standard deviation above the mean, were identified using an HTP sequencing-based yeast replicative lifespan screen (SEBYL), with pooled cells subject to four rounds of biotin-based sorting to enrich old vs young cells, followed by sequencing to ID mutants and abundance 	Yu R, et al. (2021) PMID:33790287
OCA4	YCR095C	replicative lifespan: increased 	systematic mutation set 	large-scale survey	null Allele: oca4-Δ	S288C		Details: long-lived mutants, one standard deviation above the mean, were identified using an HTP sequencing-based yeast replicative lifespan screen (SEBYL), with pooled cells subject to four rounds of biotin-based sorting to enrich old vs young cells, followed by sequencing to ID mutants and abundance 	Yu R, et al. (2021) PMID:33790287
OCA5	YHL029C	replicative lifespan: increased 	systematic mutation set 	large-scale survey	null Allele: oca5-Δ	S288C		Details: long-lived mutants, one standard deviation above the mean, were identified using an HTP sequencing-based yeast replicative lifespan screen (SEBYL), with pooled cells subject to four rounds of biotin-based sorting to enrich old vs young cells, followed by sequencing to ID mutants and abundance 	Yu R, et al. (2021) PMID:33790287
OCA6	YDR067C	replicative lifespan: increased 	systematic mutation set 	large-scale survey	null Allele: oca6-Δ	S288C		Details: long-lived mutants, one standard deviation above the mean, were identified using an HTP sequencing-based yeast replicative lifespan screen (SEBYL), with pooled cells subject to four rounds of biotin-based sorting to enrich old vs young cells, followed by sequencing to ID mutants and abundance 	Yu R, et al. (2021) PMID:33790287
OM45	YIL136W	replicative lifespan: increased 	large-scale survey	large-scale survey	null Allele: om45-Δ	S288C			Kaeberlein M, et al. (2005) PMID:16293764
OPI3	YJR073C	replicative lifespan: increased 	classical genetics	classical genetics	null Allele: opi3-Δ	S288C			McCormick MA, et al. (2015) PMID:26456335
OPT2	YPR194C	replicative lifespan: increased 	classical genetics	classical genetics	null Allele: opt2-Δ	S288C		Details: replicative lifespan is decreased during dietary restriction (0.5% glucose) 	Wuttke D, et al. (2012) PMID:22912585
OSH3	YHR073W	replicative lifespan: increased 	systematic mutation set 	large-scale survey	null Allele: osh3-Δ	S288C		Details: long-lived mutants, one standard deviation above the mean, were identified using an HTP sequencing-based yeast replicative lifespan screen (SEBYL), with pooled cells subject to four rounds of biotin-based sorting to enrich old vs young cells, followed by sequencing to ID mutants and abundance 	Yu R, et al. (2021) PMID:33790287
OSH6	YKR003W	replicative lifespan: increased 	classical genetics	classical genetics	misexpression	S288C			Gebre S, et al. (2012) PMID:22622083
OST3	YOR085W	replicative lifespan: increased 	systematic mutation set 	large-scale survey	null Allele: ost3-Δ	S288C		Details: long-lived mutants, one standard deviation above the mean, were identified using an HTP sequencing-based yeast replicative lifespan screen (SEBYL), with pooled cells subject to four rounds of biotin-based sorting to enrich old vs young cells, followed by sequencing to ID mutants and abundance 	Yu R, et al. (2021) PMID:33790287
OST3	YOR085W	replicative lifespan: increased 	classical genetics	classical genetics	null Allele: ost3-Δ	S288C			Labunskyy VM, et al. (2014) PMID:24391512
PAF1	YBR279W	replicative lifespan: increased 	classical genetics	classical genetics	null Allele: paf1-Δ	S288C			McCormick MA, et al. (2015) PMID:26456335
PAN2	YGL094C	replicative lifespan: increased 	systematic mutation set 	large-scale survey	null Allele: pan2-Δ	S288C		Details: long-lived mutants, one standard deviation above the mean, were identified using an HTP sequencing-based yeast replicative lifespan screen (SEBYL), with pooled cells subject to four rounds of biotin-based sorting to enrich old vs young cells, followed by sequencing to ID mutants and abundance 	Yu R, et al. (2021) PMID:33790287
PAU5	YFL020C	replicative lifespan: increased 	systematic mutation set 	large-scale survey	null Allele: pau5-Δ	S288C		Details: long-lived mutants, one standard deviation above the mean, were identified using an HTP sequencing-based yeast replicative lifespan screen (SEBYL), with pooled cells subject to four rounds of biotin-based sorting to enrich old vs young cells, followed by sequencing to ID mutants and abundance 	Yu R, et al. (2021) PMID:33790287
PCK1	YKR097W	replicative lifespan: increased 	classical genetics 	classical genetics	overexpression	S288C		Details: significantly extended the RLS of the null mutant by approximately 21% 	Yuan Y, et al. (2020) PMID:32104690
PCL1	YNL289W	replicative lifespan: increased 	systematic mutation set 	large-scale survey	null Allele: pcl1-Δ	S288C		Details: long-lived mutants, one standard deviation above the mean, were identified using an HTP sequencing-based yeast replicative lifespan screen (SEBYL), with pooled cells subject to four rounds of biotin-based sorting to enrich old vs young cells, followed by sequencing to ID mutants and abundance 	Yu R, et al. (2021) PMID:33790287
PCL1	YNL289W	replicative lifespan: increased 	classical genetics 	classical genetics	null Allele: pcl1-Δ	S288C		Details: long-lived mutant assayed using a microfluidic chip-based lifespan measurement with automatic imaging to validate SEBYL 	Yu R, et al. (2021) PMID:33790287
PDA1	YER178W	replicative lifespan: increased 	classical genetics	classical genetics	null Allele: pda1-Δ	S288C		Treatment: caloric restriction by glucose reduction, 0.05% Details: dietary restriction-induced lifespan extension is increased relative to wt 	Schleit J, et al. (2013) PMID:23837470
PDB1	YBR221C	replicative lifespan: increased 	classical genetics	classical genetics	null Allele: pdb1-Δ	S288C		Treatment: caloric restriction by glucose reduction, 0.05% Details: dietary restriction-induced lifespan extension is increased relative to wt 	Schleit J, et al. (2013) PMID:23837470
PDR17	YNL264C	replicative lifespan: increased 	systematic mutation set 	large-scale survey	null Allele: pdr17-Δ	S288C		Details: long-lived mutants, one standard deviation above the mean, were identified using an HTP sequencing-based yeast replicative lifespan screen (SEBYL), with pooled cells subject to four rounds of biotin-based sorting to enrich old vs young cells, followed by sequencing to ID mutants and abundance 	Yu R, et al. (2021) PMID:33790287
PER33	YLR064W	replicative lifespan: increased 	systematic mutation set 	large-scale survey	null Allele: per33-Δ	S288C		Details: long-lived mutants, one standard deviation above the mean, were identified using an HTP sequencing-based yeast replicative lifespan screen (SEBYL), with pooled cells subject to four rounds of biotin-based sorting to enrich old vs young cells, followed by sequencing to ID mutants and abundance 	Yu R, et al. (2021) PMID:33790287
PET127	YOR017W	replicative lifespan: increased 	classical genetics	classical genetics	null Allele: pet127-Δ	S288C			Caballero A, et al. (2011) PMID:21549315
PET130	YJL023C	replicative lifespan: increased 	classical genetics	classical genetics	null Allele: pet130-Δ	S288C			McCormick MA, et al. (2015) PMID:26456335
PET309	YLR067C	replicative lifespan: increased 	classical genetics	classical genetics	null Allele: pet309-Δ	S288C			Caballero A, et al. (2011) PMID:21549315
PEX19	YDL065C	replicative lifespan: increased 	classical genetics	classical genetics	null Allele: pex19-Δ	S288C			McCormick MA, et al. (2015) PMID:26456335
PEX27	YOR193W	replicative lifespan: increased 	systematic mutation set 	large-scale survey	null Allele: pex27-Δ	S288C		Details: long-lived mutants, one standard deviation above the mean, were identified using an HTP sequencing-based yeast replicative lifespan screen (SEBYL), with pooled cells subject to four rounds of biotin-based sorting to enrich old vs young cells, followed by sequencing to ID mutants and abundance 	Yu R, et al. (2021) PMID:33790287
PEX31	YGR004W	replicative lifespan: increased 	systematic mutation set 	large-scale survey	null Allele: pex31-Δ	S288C		Details: long-lived mutants, one standard deviation above the mean, were identified using an HTP sequencing-based yeast replicative lifespan screen (SEBYL), with pooled cells subject to four rounds of biotin-based sorting to enrich old vs young cells, followed by sequencing to ID mutants and abundance 	Yu R, et al. (2021) PMID:33790287
PEX7	YDR142C	replicative lifespan: increased 	systematic mutation set 	large-scale survey	null Allele: pex7-Δ	S288C		Details: long-lived mutants, one standard deviation above the mean, were identified using an HTP sequencing-based yeast replicative lifespan screen (SEBYL), with pooled cells subject to four rounds of biotin-based sorting to enrich old vs young cells, followed by sequencing to ID mutants and abundance 	Yu R, et al. (2021) PMID:33790287
PEX9	YMR018W	replicative lifespan: increased 	classical genetics	classical genetics	null Allele: pex9-Δ	S288C			McCormick MA, et al. (2015) PMID:26456335
PGA3	YML125C	replicative lifespan: increased 	classical genetics	classical genetics	overexpression	S288C			Jiménez-Hidalgo M, et al. (2009) PMID:19239415
PGC1	YPL206C	replicative lifespan: increased 	systematic mutation set 	large-scale survey	null Allele: pgc1-Δ	S288C		Details: long-lived mutants, one standard deviation above the mean, were identified using an HTP sequencing-based yeast replicative lifespan screen (SEBYL), with pooled cells subject to four rounds of biotin-based sorting to enrich old vs young cells, followed by sequencing to ID mutants and abundance 	Yu R, et al. (2021) PMID:33790287
PGM2	YMR105C	replicative lifespan: increased 	classical genetics	classical genetics	null Allele: pgm2-Δ	S288C			McCormick MA, et al. (2015) PMID:26456335
PHB1	YGR132C	replicative lifespan: increased 	classical genetics	classical genetics	null Allele: phb1-Δ	S288C		Treatment: caloric restriction by glucose reduction, 0.05% Details: dietary restriction-induced lifespan extension is increased relative to wt 	Schleit J, et al. (2013) PMID:23837470
PHB2	YGR231C	replicative lifespan: increased 	classical genetics	classical genetics	null Allele: phb2-Δ	S288C		Treatment: caloric restriction by glucose reduction, 0.05% Details: dietary restriction-induced lifespan extension is increased relative to wt 	Schleit J, et al. (2013) PMID:23837470
PHD1	YKL043W	replicative lifespan: increased 	large-scale survey	large-scale survey	null Allele: phd1-Δ	S288C			Managbanag JR, et al. (2008) PMID:19030232
PHM7	YOL084W	replicative lifespan: increased 	systematic mutation set 	large-scale survey	null Allele: phm7-Δ	S288C		Details: long-lived mutants, one standard deviation above the mean, were identified using an HTP sequencing-based yeast replicative lifespan screen (SEBYL), with pooled cells subject to four rounds of biotin-based sorting to enrich old vs young cells, followed by sequencing to ID mutants and abundance 	Yu R, et al. (2021) PMID:33790287
PHO2	YDL106C	replicative lifespan: increased 	systematic mutation set 	large-scale survey	null Allele: pho2-Δ	S288C		Details: long-lived mutants, one standard deviation above the mean, were identified using an HTP sequencing-based yeast replicative lifespan screen (SEBYL), with pooled cells subject to four rounds of biotin-based sorting to enrich old vs young cells, followed by sequencing to ID mutants and abundance 	Yu R, et al. (2021) PMID:33790287
PHO87	YCR037C	replicative lifespan: increased 	classical genetics	classical genetics	null Allele: pho87-Δ	S288C			McCormick MA, et al. (2015) PMID:26456335
PHO87	YCR037C	replicative lifespan: increased 	systematic mutation set 	large-scale survey	null Allele: pho87-Δ	S288C		Details: long-lived mutants, one standard deviation above the mean, were identified using an HTP sequencing-based yeast replicative lifespan screen (SEBYL), with pooled cells subject to four rounds of biotin-based sorting to enrich old vs young cells, followed by sequencing to ID mutants and abundance 	Yu R, et al. (2021) PMID:33790287
PIB1	YDR313C	replicative lifespan: increased 	large-scale survey	large-scale survey	null Allele: pib1-Δ	S288C			Kaeberlein M, et al. (2005) PMID:16293764
PIB1	YDR313C	replicative lifespan: increased 	classical genetics	classical genetics	null Allele: pib1-Δ	S288C			McCormick MA, et al. (2015) PMID:26456335
PIP2	YOR363C	replicative lifespan: increased 	systematic mutation set 	large-scale survey	null Allele: pip2-Δ	S288C		Details: long-lived mutants, one standard deviation above the mean, were identified using an HTP sequencing-based yeast replicative lifespan screen (SEBYL), with pooled cells subject to four rounds of biotin-based sorting to enrich old vs young cells, followed by sequencing to ID mutants and abundance 	Yu R, et al. (2021) PMID:33790287
PIR1	YKL164C	replicative lifespan: increased 	systematic mutation set 	large-scale survey	null Allele: pir1-Δ	S288C		Details: long-lived mutants, one standard deviation above the mean, were identified using an HTP sequencing-based yeast replicative lifespan screen (SEBYL), with pooled cells subject to four rounds of biotin-based sorting to enrich old vs young cells, followed by sequencing to ID mutants and abundance 	Yu R, et al. (2021) PMID:33790287
PIR5	YJL160C	replicative lifespan: increased 	systematic mutation set 	large-scale survey	null Allele: pir5-Δ	S288C		Details: long-lived mutants, one standard deviation above the mean, were identified using an HTP sequencing-based yeast replicative lifespan screen (SEBYL), with pooled cells subject to four rounds of biotin-based sorting to enrich old vs young cells, followed by sequencing to ID mutants and abundance 	Yu R, et al. (2021) PMID:33790287
PKH2	YOL100W	replicative lifespan: increased 	large-scale survey	large-scale survey	null Allele: pkh2-Δ	S288C			Smith ED, et al. (2008) PMID:18340043
PKH2	YOL100W	replicative lifespan: increased 	competitive growth	large-scale survey	null Allele: pkh2-Δ	S288C			Delaney JR, et al. (2011) PMID:21191185
PKH2	YOL100W	replicative lifespan: increased 	classical genetics	classical genetics	null Allele: pkh2-Δ	S288C			McCormick MA, et al. (2015) PMID:26456335
PLM2	YDR501W	replicative lifespan: increased 	systematic mutation set 	large-scale survey	null Allele: plm2-Δ	S288C		Details: long-lived mutants, one standard deviation above the mean, were identified using an HTP sequencing-based yeast replicative lifespan screen (SEBYL), with pooled cells subject to four rounds of biotin-based sorting to enrich old vs young cells, followed by sequencing to ID mutants and abundance 	Yu R, et al. (2021) PMID:33790287
PMA1	YGL008C	replicative lifespan: increased 	classical genetics	classical genetics	reduction of function Allele: pma1-105 S368F; catalytic domain mutated; reduced Km and Vmax; vanadate insensitive	S288C			Henderson KA, et al. (2014) PMID:25190112
PMC1	YGL006W	replicative lifespan: increased 	classical genetics	classical genetics	null Allele: pmc1-Δ	S288C			McCormick MA, et al. (2015) PMID:26456335
PMC1	YGL006W	replicative lifespan: increased 	systematic mutation set 	large-scale survey	null Allele: pmc1-Δ	S288C		Details: long-lived mutants, one standard deviation above the mean, were identified using an HTP sequencing-based yeast replicative lifespan screen (SEBYL), with pooled cells subject to four rounds of biotin-based sorting to enrich old vs young cells, followed by sequencing to ID mutants and abundance 	Yu R, et al. (2021) PMID:33790287
PML39	YML107C	replicative lifespan: increased 	classical genetics	classical genetics	null Allele: pml39-Δ	S288C			McCormick MA, et al. (2015) PMID:26456335
PMR1	YGL167C	replicative lifespan: increased 	large-scale survey	large-scale survey	null Allele: pmr1-Δ	S288C			Smith ED, et al. (2008) PMID:18340043
PMR1	YGL167C	replicative lifespan: increased 	competitive growth	large-scale survey	null Allele: pmr1-Δ	S288C			Delaney JR, et al. (2011) PMID:21191185
PMT1	YDL095W	replicative lifespan: increased 	classical genetics	classical genetics	null Allele: pmt1-Δ	S288C			McCormick MA, et al. (2015) PMID:26456335
PMT1	YDL095W	replicative lifespan: increased 	classical genetics	classical genetics	null Allele: pmt1-Δ	S288C		Details: mean replicative lifespan increased by 20% 	Cui HJ, et al. (2015) PMID:25936926
PMT3	YOR321W	replicative lifespan: increased 	classical genetics	classical genetics	null Allele: pmt3-Δ	S288C			McCormick MA, et al. (2015) PMID:26456335
PMT5	YDL093W	replicative lifespan: increased 	classical genetics	classical genetics	null Allele: pmt5-Δ	S288C			McCormick MA, et al. (2015) PMID:26456335
PMT7	YDR307W	replicative lifespan: increased 	large-scale survey	large-scale survey	null Allele: pmt7-Δ	S288C			Kaeberlein M, et al. (2005) PMID:16293764
PNC1	YGL037C	replicative lifespan: increased 	classical genetics 	classical genetics	overexpression	Other	5 mM nicotinamide	Details: PNC1 overexpression partially restores replicative lifespan that is decreased as a result of treatment with nicotinamide 	Gallo CM, et al. (2004) PMID:14729974
POC4	YPL144W	replicative lifespan: increased 	classical genetics	classical genetics	null Allele: poc4-Δ	S288C			McCormick MA, et al. (2015) PMID:26456335
POR1	YNL055C	replicative lifespan: increased 	classical genetics	classical genetics	null Allele: por1-Δ	S288C		Treatment: caloric restriction by glucose reduction, 0.05% Details: dietary restriction-induced lifespan extension is increased relative to wt 	Schleit J, et al. (2013) PMID:23837470
POS5	YPL188W	replicative lifespan: increased 	classical genetics	classical genetics	null Allele: pos5-Δ	S288C		Treatment: caloric restriction by glucose reduction, 0.05% Details: dietary restriction-induced lifespan extension is increased relative to wt 	Schleit J, et al. (2013) PMID:23837470
PPA2	YMR267W	replicative lifespan: increased 	homozygous diploid 	classical genetics	null Allele: ppa2-Δ	S288C		Details: 28% extended lifespan 	Muid KA, et al. (2019) PMID:31082499
PRO1	YDR300C	replicative lifespan: increased 	classical genetics 	classical genetics	activation Allele: PRO1-I150T I150T (T449C); variant located in the kinase domain between the allosteric and substrate binding sites; may increase substrate affinity; feedback inhibition insensitive	S288C		Details: exhibits markedly longer lifespan compared to wt 	Mukai Y, et al. (2019) PMID:31646149
PRR1	YKL116C	replicative lifespan: increased 	systematic mutation set 	large-scale survey	null Allele: prr1-Δ	S288C		Details: long-lived mutants, one standard deviation above the mean, were identified using an HTP sequencing-based yeast replicative lifespan screen (SEBYL), with pooled cells subject to four rounds of biotin-based sorting to enrich old vs young cells, followed by sequencing to ID mutants and abundance 	Yu R, et al. (2021) PMID:33790287
PRS5	YOL061W	replicative lifespan: increased 	systematic mutation set 	large-scale survey	null Allele: prs5-Δ	S288C		Details: long-lived mutants, one standard deviation above the mean, were identified using an HTP sequencing-based yeast replicative lifespan screen (SEBYL), with pooled cells subject to four rounds of biotin-based sorting to enrich old vs young cells, followed by sequencing to ID mutants and abundance 	Yu R, et al. (2021) PMID:33790287
PSP2	YML017W	replicative lifespan: increased 	classical genetics	classical genetics	null Allele: psp2-Δ	S288C			McCormick MA, et al. (2015) PMID:26456335
PTP2	YOR208W	replicative lifespan: increased 	systematic mutation set 	large-scale survey	null Allele: ptp2-Δ	S288C		Details: long-lived mutants, one standard deviation above the mean, were identified using an HTP sequencing-based yeast replicative lifespan screen (SEBYL), with pooled cells subject to four rounds of biotin-based sorting to enrich old vs young cells, followed by sequencing to ID mutants and abundance 	Yu R, et al. (2021) PMID:33790287
PUF3	YLL013C	replicative lifespan: increased 	systematic mutation set 	large-scale survey	null Allele: puf3-Δ	S288C		Details: long-lived mutants, one standard deviation above the mean, were identified using an HTP sequencing-based yeast replicative lifespan screen (SEBYL), with pooled cells subject to four rounds of biotin-based sorting to enrich old vs young cells, followed by sequencing to ID mutants and abundance 	Yu R, et al. (2021) PMID:33790287
PUS2	YGL063W	replicative lifespan: increased 	systematic mutation set 	large-scale survey	null Allele: pus2-Δ	S288C		Details: long-lived mutants, one standard deviation above the mean, were identified using an HTP sequencing-based yeast replicative lifespan screen (SEBYL), with pooled cells subject to four rounds of biotin-based sorting to enrich old vs young cells, followed by sequencing to ID mutants and abundance 	Yu R, et al. (2021) PMID:33790287
PUT1	YLR142W	replicative lifespan: increased 	classical genetics 	classical genetics	null Allele: put1-Δ	S288C		Details: exhibits markedly longer lifespan compared to wt 	Mukai Y, et al. (2019) PMID:31646149
PUT2	YHR037W	replicative lifespan: increased 	systematic mutation set 	large-scale survey	null Allele: put2-Δ	S288C		Details: long-lived mutants, one standard deviation above the mean, were identified using an HTP sequencing-based yeast replicative lifespan screen (SEBYL), with pooled cells subject to four rounds of biotin-based sorting to enrich old vs young cells, followed by sequencing to ID mutants and abundance 	Yu R, et al. (2021) PMID:33790287
PXP1	YEL020C	replicative lifespan: increased 	classical genetics	classical genetics	null Allele: pxp1-Δ	S288C			McCormick MA, et al. (2015) PMID:26456335
RAD34	YDR314C	replicative lifespan: increased 	classical genetics	classical genetics	null Allele: rad34-Δ	S288C			McCormick MA, et al. (2015) PMID:26456335
RAD51	YER095W	replicative lifespan: increased 	classical genetics	classical genetics	null Allele: rad51-Δ	S288C		Treatment: caloric restriction by glucose reduction, 0.05% Details: dietary restriction-induced lifespan extension is increased relative to wt 	Schleit J, et al. (2013) PMID:23837470
RAD55	YDR076W	replicative lifespan: increased 	classical genetics	classical genetics	null Allele: rad55-Δ	S288C		Treatment: caloric restriction by glucose reduction, 0.05% Details: dietary restriction-induced lifespan extension is increased relative to wt 	Schleit J, et al. (2013) PMID:23837470
RAI1	YGL246C	replicative lifespan: increased 	classical genetics	classical genetics	null Allele: rai1-Δ	S288C			McCormick MA, et al. (2015) PMID:26456335
RAS1	YOR101W	replicative lifespan: increased 	classical genetics	classical genetics	null Allele: ras1-Δ	Other			Sun J, et al. (1994) PMID:8034612
RAS1	YOR101W	replicative lifespan: increased 	large-scale survey	large-scale survey	null Allele: ras1-Δ	S288C			Kaeberlein M, et al. (2005) PMID:16293764
RAS2	YNL098C	replicative lifespan: increased 	classical genetics	classical genetics	overexpression	Other		Details: 20-40% increase in mean and maximum lifespan; delays the aging related increase in generation time 	Sun J, et al. (1994) PMID:8034612
RAS2	YNL098C	replicative lifespan: increased 	classical genetics	classical genetics	overexpression Allele: ras2ser-42 T42S; reduced activity, effector domain mutant	Other		Details: 30% increase in average lifespan; postpones aging related increase in generation time 	Sun J, et al. (1994) PMID:8034612
RAX2	YLR084C	replicative lifespan: increased 	systematic mutation set 	large-scale survey	null Allele: rax2-Δ	S288C		Details: long-lived mutants, one standard deviation above the mean, were identified using an HTP sequencing-based yeast replicative lifespan screen (SEBYL), with pooled cells subject to four rounds of biotin-based sorting to enrich old vs young cells, followed by sequencing to ID mutants and abundance 	Yu R, et al. (2021) PMID:33790287
RAX2	YLR084C	replicative lifespan: increased 	classical genetics 	classical genetics	null Allele: rax2-Δ	S288C		Details: long-lived mutant assayed using a microfluidic chip-based lifespan measurement with automatic imaging to validate SEBYL 	Yu R, et al. (2021) PMID:33790287
RBH2	YJR030C	replicative lifespan: increased 	systematic mutation set 	large-scale survey	null Allele: rbh2-Δ	S288C		Details: long-lived mutants, one standard deviation above the mean, were identified using an HTP sequencing-based yeast replicative lifespan screen (SEBYL), with pooled cells subject to four rounds of biotin-based sorting to enrich old vs young cells, followed by sequencing to ID mutants and abundance 	Yu R, et al. (2021) PMID:33790287
RCR2	YDR003W	replicative lifespan: increased 	classical genetics	classical genetics	null Allele: rcr2-Δ	S288C		Details: replicative lifespan is decreased during dietary restriction (0.5% glucose) 	Wuttke D, et al. (2012) PMID:22912585
RDH54	YBR073W	replicative lifespan: increased 	systematic mutation set 	large-scale survey	null Allele: rdh54-Δ	S288C		Details: long-lived mutants, one standard deviation above the mean, were identified using an HTP sequencing-based yeast replicative lifespan screen (SEBYL), with pooled cells subject to four rounds of biotin-based sorting to enrich old vs young cells, followed by sequencing to ID mutants and abundance 	Yu R, et al. (2021) PMID:33790287
RDS1	YCR106W	replicative lifespan: increased 	classical genetics	classical genetics	null Allele: rds1-Δ	S288C			McCormick MA, et al. (2015) PMID:26456335
REC107	YJR021C	replicative lifespan: increased 	systematic mutation set 	large-scale survey	null Allele: rec107-Δ	S288C		Details: long-lived mutants, one standard deviation above the mean, were identified using an HTP sequencing-based yeast replicative lifespan screen (SEBYL), with pooled cells subject to four rounds of biotin-based sorting to enrich old vs young cells, followed by sequencing to ID mutants and abundance 	Yu R, et al. (2021) PMID:33790287
REC8	YPR007C	replicative lifespan: increased 	classical genetics	classical genetics	null Allele: rec8-Δ	S288C			Ayyadevara S, et al. (2014) PMID:25136348
REC8	YPR007C	replicative lifespan: increased 	classical genetics	classical genetics	null Allele: rec8-Δ	S288C			McCormick MA, et al. (2015) PMID:26456335
REI1	YBR267W	replicative lifespan: increased 	classical genetics	classical genetics	null Allele: rei1-Δ	S288C			Kaeberlein M, et al. (2005) PMID:16293764
REI1	YBR267W	replicative lifespan: increased 	large-scale survey	large-scale survey	null Allele: rei1-Δ	S288C			Kaeberlein M, et al. (2005) PMID:16293764
REI1	YBR267W	replicative lifespan: increased 	classical genetics	classical genetics	null Allele: rei1-Δ	S288C			Delaney JR, et al. (2011) PMID:21902802
REI1	YBR267W	replicative lifespan: increased 	competitive growth	large-scale survey	null Allele: rei1-Δ	S288C			Delaney JR, et al. (2011) PMID:21191185
REI1	YBR267W	replicative lifespan: increased 	classical genetics	classical genetics	null Allele: rei1-Δ	S288C			McCormick MA, et al. (2015) PMID:26456335
RER1	YCL001W	replicative lifespan: increased 	systematic mutation set 	large-scale survey	null Allele: rer1-Δ	S288C		Details: long-lived mutants, one standard deviation above the mean, were identified using an HTP sequencing-based yeast replicative lifespan screen (SEBYL), with pooled cells subject to four rounds of biotin-based sorting to enrich old vs young cells, followed by sequencing to ID mutants and abundance 	Yu R, et al. (2021) PMID:33790287
REV1	YOR346W	replicative lifespan: increased 	systematic mutation set 	large-scale survey	null Allele: rev1-Δ	S288C		Details: long-lived mutants, one standard deviation above the mean, were identified using an HTP sequencing-based yeast replicative lifespan screen (SEBYL), with pooled cells subject to four rounds of biotin-based sorting to enrich old vs young cells, followed by sequencing to ID mutants and abundance 	Yu R, et al. (2021) PMID:33790287
REV1	YOR346W	replicative lifespan: increased 	classical genetics 	classical genetics	null Allele: rev1-Δ	S288C		Details: long-lived mutant assayed using a microfluidic chip-based lifespan measurement with automatic imaging to validate SEBYL 	Yu R, et al. (2021) PMID:33790287
RFX1	YLR176C	replicative lifespan: increased 	classical genetics	classical genetics	null Allele: rfx1-Δ	S288C			McCormick MA, et al. (2015) PMID:26456335
RGD3	YHR182W	replicative lifespan: increased 	systematic mutation set 	large-scale survey	null Allele: rgd3-Δ	S288C		Details: long-lived mutants, one standard deviation above the mean, were identified using an HTP sequencing-based yeast replicative lifespan screen (SEBYL), with pooled cells subject to four rounds of biotin-based sorting to enrich old vs young cells, followed by sequencing to ID mutants and abundance 	Yu R, et al. (2021) PMID:33790287
RGS2	YOR107W	replicative lifespan: increased 	large-scale survey	large-scale survey	null Allele: rgs2-Δ	S288C			Kaeberlein M, et al. (2005) PMID:16293764
RIM1	YCR028C-A	replicative lifespan: increased 	large-scale survey	large-scale survey	null Allele: rim1-Δ	S288C			Managbanag JR, et al. (2008) PMID:19030232
RIM1	YCR028C-A	replicative lifespan: increased 	classical genetics	classical genetics	null Allele: rim1-Δ	S288C			Delaney JR, et al. (2011) PMID:21902802
RIM1	YCR028C-A	replicative lifespan: increased 	classical genetics	classical genetics	null Allele: rim1-Δ	S288C			McCormick MA, et al. (2015) PMID:26456335
RIM1	YCR028C-A	replicative lifespan: increased 	competitive growth	large-scale survey	null Allele: rim1-Δ	S288C			Delaney JR, et al. (2011) PMID:21191185
RIP1	YEL024W	replicative lifespan: increased 	systematic mutation set 	large-scale survey	null Allele: rip1-Δ	S288C		Details: long-lived mutants, one standard deviation above the mean, were identified using an HTP sequencing-based yeast replicative lifespan screen (SEBYL), with pooled cells subject to four rounds of biotin-based sorting to enrich old vs young cells, followed by sequencing to ID mutants and abundance 	Yu R, et al. (2021) PMID:33790287
RIP1	YEL024W	replicative lifespan: increased 	classical genetics	classical genetics	null Allele: rip1-Δ	S288C		Treatment: caloric restriction by glucose reduction, 0.05% Details: dietary restriction-induced lifespan extension is increased relative to wt 	Schleit J, et al. (2013) PMID:23837470
RIT1	YMR283C	replicative lifespan: increased 	classical genetics	classical genetics	null Allele: rit1-Δ	S288C			McCormick MA, et al. (2015) PMID:26456335
RKM3	YBR030W	replicative lifespan: increased 	systematic mutation set 	large-scale survey	null Allele: rkm3-Δ	S288C		Details: long-lived mutants, one standard deviation above the mean, were identified using an HTP sequencing-based yeast replicative lifespan screen (SEBYL), with pooled cells subject to four rounds of biotin-based sorting to enrich old vs young cells, followed by sequencing to ID mutants and abundance 	Yu R, et al. (2021) PMID:33790287
RML2	YEL050C	replicative lifespan: increased 	systematic mutation set 	large-scale survey	null Allele: rml2-Δ	S288C		Details: long-lived mutants, one standard deviation above the mean, were identified using an HTP sequencing-based yeast replicative lifespan screen (SEBYL), with pooled cells subject to four rounds of biotin-based sorting to enrich old vs young cells, followed by sequencing to ID mutants and abundance 	Yu R, et al. (2021) PMID:33790287
RMT2	YDR465C	replicative lifespan: increased 	classical genetics	classical genetics	null Allele: rmt2-Δ	S288C			McCormick MA, et al. (2015) PMID:26456335
ROM2	YLR371W	replicative lifespan: increased 	classical genetics	classical genetics	null Allele: rom2-Δ	S288C			Kaeberlein M, et al. (2005) PMID:16293764
ROM2	YLR371W	replicative lifespan: increased 	large-scale survey	large-scale survey	null Allele: rom2-Δ	S288C			Kaeberlein M, et al. (2005) PMID:16293764
ROM2	YLR371W	replicative lifespan: increased 	classical genetics	classical genetics	null Allele: rom2-Δ	S288C			McCormick MA, et al. (2015) PMID:26456335
RPD3	YNL330C	replicative lifespan: increased 	classical genetics 	classical genetics	null Allele: rpd3-Δ	Other			Kim S, et al. (1999) PMID:10512855
RPH1	YER169W	replicative lifespan: increased 	classical genetics	classical genetics	null Allele: rph1-Δ	S288C			McCormick MA, et al. (2015) PMID:26456335
RPL12A	YEL054C	replicative lifespan: increased 	classical genetics	classical genetics	null Allele: rpl12a-Δ	S288C			McCormick MA, et al. (2015) PMID:26456335
RPL12B	YDR418W	replicative lifespan: increased 	classical genetics	classical genetics	null Allele: rpl12b-Δ	S288C			Steffen KK, et al. (2012) PMID:22377630
RPL12B	YDR418W	replicative lifespan: increased 	classical genetics	classical genetics	null Allele: rpl12b-Δ	S288C			McCormick MA, et al. (2015) PMID:26456335
RPL13A	YDL082W	replicative lifespan: increased 	classical genetics	classical genetics	null Allele: rpl13a-Δ	S288C			Steffen KK, et al. (2008) PMID:18423200
RPL13A	YDL082W	replicative lifespan: increased 	classical genetics	classical genetics	null Allele: rpl13a-Δ	S288C			Delaney JR, et al. (2011) PMID:21902802
RPL13A	YDL082W	replicative lifespan: increased 	competitive growth	large-scale survey	null Allele: rpl13a-Δ	S288C			Delaney JR, et al. (2011) PMID:21191185
RPL13A	YDL082W	replicative lifespan: increased 	classical genetics	classical genetics	null Allele: rpl13a-Δ	S288C			McCormick MA, et al. (2015) PMID:26456335
RPL13A	YDL082W	replicative lifespan: increased 	systematic mutation set 	large-scale survey	null Allele: rpl13a-Δ	S288C		Details: long-lived mutants, one standard deviation above the mean, were identified using an HTP sequencing-based yeast replicative lifespan screen (SEBYL), with pooled cells subject to four rounds of biotin-based sorting to enrich old vs young cells, followed by sequencing to ID mutants and abundance 	Yu R, et al. (2021) PMID:33790287
RPL13B	YMR142C	replicative lifespan: increased 	classical genetics	classical genetics	null Allele: rpl13b-Δ	S288C			Steffen KK, et al. (2012) PMID:22377630
RPL13B	YMR142C	replicative lifespan: increased 	classical genetics	classical genetics	null Allele: rpl13b-Δ	S288C			McCormick MA, et al. (2015) PMID:26456335
RPL16A	YIL133C	replicative lifespan: increased 	systematic mutation set 	large-scale survey	null Allele: rpl16a-Δ	S288C		Details: long-lived mutants, one standard deviation above the mean, were identified using an HTP sequencing-based yeast replicative lifespan screen (SEBYL), with pooled cells subject to four rounds of biotin-based sorting to enrich old vs young cells, followed by sequencing to ID mutants and abundance 	Yu R, et al. (2021) PMID:33790287
RPL16B	YNL069C	replicative lifespan: increased 	classical genetics	classical genetics	null Allele: rpl16b-Δ	S288C			Steffen KK, et al. (2012) PMID:22377630
RPL16B	YNL069C	replicative lifespan: increased 	classical genetics	classical genetics	null Allele: rpl16b-Δ	S288C			McCormick MA, et al. (2015) PMID:26456335
RPL16B	YNL069C	replicative lifespan: increased 	systematic mutation set 	large-scale survey	null Allele: rpl16b-Δ	S288C		Details: long-lived mutants, one standard deviation above the mean, were identified using an HTP sequencing-based yeast replicative lifespan screen (SEBYL), with pooled cells subject to four rounds of biotin-based sorting to enrich old vs young cells, followed by sequencing to ID mutants and abundance 	Yu R, et al. (2021) PMID:33790287
RPL19A	YBR084C-A	replicative lifespan: increased 	large-scale survey	large-scale survey	null Allele: rpl19a-Δ	S288C			Smith ED, et al. (2008) PMID:18340043
RPL19A	YBR084C-A	replicative lifespan: increased 	classical genetics	classical genetics	null Allele: rpl19a-Δ	S288C			Steffen KK, et al. (2008) PMID:18423200
RPL19A	YBR084C-A	replicative lifespan: increased 	classical genetics	classical genetics	null Allele: rpl19a-Δ	S288C			McCormick MA, et al. (2015) PMID:26456335
RPL19A	YBR084C-A	replicative lifespan: increased 	competitive growth	large-scale survey	null Allele: rpl19a-Δ	S288C			Delaney JR, et al. (2011) PMID:21191185
RPL19B	YBL027W	replicative lifespan: increased 	classical genetics	classical genetics	null Allele: rpl19b-Δ	S288C			McCormick MA, et al. (2015) PMID:26456335
RPL1B	YGL135W	replicative lifespan: increased 	classical genetics	classical genetics	null Allele: rpl1b-Δ	S288C			McCormick MA, et al. (2015) PMID:26456335
RPL20A	YMR242C	replicative lifespan: increased 	classical genetics	classical genetics	null Allele: rpl20a-Δ	S288C			Steffen KK, et al. (2012) PMID:22377630
RPL20A	YMR242C	replicative lifespan: increased 	classical genetics	classical genetics	null Allele: rpl20a-Δ	S288C			McCormick MA, et al. (2015) PMID:26456335
RPL20B	YOR312C	replicative lifespan: increased 	large-scale survey	large-scale survey	null Allele: rpl20b-Δ	S288C			Managbanag JR, et al. (2008) PMID:19030232
RPL20B	YOR312C	replicative lifespan: increased 	classical genetics	classical genetics	null Allele: rpl20b-Δ	S288C			Steffen KK, et al. (2008) PMID:18423200
RPL20B	YOR312C	replicative lifespan: increased 	classical genetics	classical genetics	null Allele: rpl20b-Δ	S288C			Steffen KK, et al. (2012) PMID:22377630
RPL20B	YOR312C	replicative lifespan: increased 	classical genetics	classical genetics	null Allele: rpl20b-Δ	S288C			Schleit J, et al. (2013) PMID:23837470
RPL20B	YOR312C	replicative lifespan: increased 	classical genetics	classical genetics	null Allele: rpl20b-Δ	S288C		Treatment: caloric restriction by glucose reduction, 0.05% Details: dietary restriction-induced lifespan extension is increased relative to wt 	Schleit J, et al. (2013) PMID:23837470
RPL20B	YOR312C	replicative lifespan: increased 	systematic mutation set	large-scale survey	null Allele: rpl20b-Δ	S288C			Delaney JR, et al. (2013) PMID:23336757
RPL20B	YOR312C	replicative lifespan: increased 	classical genetics	classical genetics	null Allele: rpl20b-Δ	S288C			McCormick MA, et al. (2015) PMID:26456335
RPL21A	YBR191W	replicative lifespan: increased 	classical genetics	classical genetics	null Allele: rpl21a-Δ	S288C			Steffen KK, et al. (2012) PMID:22377630
RPL21B	YPL079W	replicative lifespan: increased 	classical genetics	classical genetics	null Allele: rpl21b-Δ	S288C			Steffen KK, et al. (2008) PMID:18423200
RPL21B	YPL079W	replicative lifespan: increased 	classical genetics	classical genetics	null Allele: rpl21b-Δ	S288C			Delaney JR, et al. (2011) PMID:21902802
RPL21B	YPL079W	replicative lifespan: increased 	competitive growth	large-scale survey	null Allele: rpl21b-Δ	S288C			Delaney JR, et al. (2011) PMID:21191185
RPL21B	YPL079W	replicative lifespan: increased 	classical genetics	classical genetics	null Allele: rpl21b-Δ	S288C			McCormick MA, et al. (2015) PMID:26456335
RPL22A	YLR061W	replicative lifespan: increased 	classical genetics	classical genetics	null Allele: rpl22a-Δ	S288C			Steffen KK, et al. (2008) PMID:18423200
RPL22A	YLR061W	replicative lifespan: increased 	classical genetics	classical genetics	null Allele: rpl22a-Δ	S288C			Delaney JR, et al. (2011) PMID:21902802
RPL22A	YLR061W	replicative lifespan: increased 	competitive growth	large-scale survey	null Allele: rpl22a-Δ	S288C			Delaney JR, et al. (2011) PMID:21191185
RPL22A	YLR061W	replicative lifespan: increased 	systematic mutation set	large-scale survey	null Allele: rpl22a-Δ	S288C			Delaney JR, et al. (2013) PMID:23336757
RPL22A	YLR061W	replicative lifespan: increased 	classical genetics	classical genetics	null Allele: rpl22a-Δ	S288C			McCormick MA, et al. (2015) PMID:26456335
RPL23A	YBL087C	replicative lifespan: increased 	classical genetics	classical genetics	null Allele: rpl23a-Δ	S288C			Steffen KK, et al. (2008) PMID:18423200
RPL23A	YBL087C	replicative lifespan: increased 	classical genetics	classical genetics	null Allele: rpl23a-Δ	S288C			Delaney JR, et al. (2011) PMID:21902802
RPL23A	YBL087C	replicative lifespan: increased 	competitive growth	large-scale survey	null Allele: rpl23a-Δ	S288C			Delaney JR, et al. (2011) PMID:21191185
RPL23A	YBL087C	replicative lifespan: increased 	classical genetics	classical genetics	null Allele: rpl23a-Δ	S288C			McCormick MA, et al. (2015) PMID:26456335
RPL26A	YLR344W	replicative lifespan: increased 	classical genetics	classical genetics	null Allele: rpl26a-Δ	S288C			McCormick MA, et al. (2015) PMID:26456335
RPL29	YFR032C-A	replicative lifespan: increased 	classical genetics	classical genetics	null Allele: rpl29-Δ	S288C			Steffen KK, et al. (2008) PMID:18423200
RPL29	YFR032C-A	replicative lifespan: increased 	competitive growth	large-scale survey	null Allele: rpl29-Δ	S288C			Delaney JR, et al. (2011) PMID:21191185
RPL29	YFR032C-A	replicative lifespan: increased 	classical genetics	classical genetics	null Allele: rpl29-Δ	S288C			McCormick MA, et al. (2015) PMID:26456335
RPL2B	YIL018W	replicative lifespan: increased 	classical genetics	classical genetics	null Allele: rpl2b-Δ	S288C			Steffen KK, et al. (2012) PMID:22377630
RPL2B	YIL018W	replicative lifespan: increased 	classical genetics	classical genetics	null Allele: rpl2b-Δ	S288C			McCormick MA, et al. (2015) PMID:26456335
RPL31A	YDL075W	replicative lifespan: increased 	classical genetics	classical genetics	null Allele: rpl31a-Δ	S288C			Kaeberlein M, et al. (2005) PMID:16293764
RPL31A	YDL075W	replicative lifespan: increased 	large-scale survey	large-scale survey	null Allele: rpl31a-Δ	S288C			Kaeberlein M, et al. (2005) PMID:16293764
RPL31A	YDL075W	replicative lifespan: increased 	classical genetics	classical genetics	null Allele: rpl31a-Δ	S288C			Steffen KK, et al. (2008) PMID:18423200
RPL31A	YDL075W	replicative lifespan: increased 	competitive growth	large-scale survey	null Allele: rpl31a-Δ	S288C			Delaney JR, et al. (2011) PMID:21191185
RPL31A	YDL075W	replicative lifespan: increased 	classical genetics	classical genetics	null Allele: rpl31a-Δ	S288C		Treatment: caloric restriction by glucose reduction, 0.05% Details: dietary restriction-induced lifespan extension is increased relative to wt 	Schleit J, et al. (2013) PMID:23837470
RPL31A	YDL075W	replicative lifespan: increased 	classical genetics	classical genetics	null Allele: rpl31a-Δ	S288C			McCormick MA, et al. (2015) PMID:26456335
RPL33B	YOR234C	replicative lifespan: increased 	systematic mutation set 	large-scale survey	null Allele: rpl33b-Δ	S288C		Details: long-lived mutants, one standard deviation above the mean, were identified using an HTP sequencing-based yeast replicative lifespan screen (SEBYL), with pooled cells subject to four rounds of biotin-based sorting to enrich old vs young cells, followed by sequencing to ID mutants and abundance 	Yu R, et al. (2021) PMID:33790287
RPL34A	YER056C-A	replicative lifespan: increased 	classical genetics	classical genetics	null Allele: rpl34a-Δ	S288C			Steffen KK, et al. (2012) PMID:22377630
RPL34A	YER056C-A	replicative lifespan: increased 	classical genetics	classical genetics	null Allele: rpl34a-Δ	S288C			McCormick MA, et al. (2015) PMID:26456335
RPL34A	YER056C-A	replicative lifespan: increased 	systematic mutation set 	large-scale survey	null Allele: rpl34a-Δ	S288C		Details: long-lived mutants, one standard deviation above the mean, were identified using an HTP sequencing-based yeast replicative lifespan screen (SEBYL), with pooled cells subject to four rounds of biotin-based sorting to enrich old vs young cells, followed by sequencing to ID mutants and abundance 	Yu R, et al. (2021) PMID:33790287
RPL34B	YIL052C	replicative lifespan: increased 	classical genetics	classical genetics	null Allele: rpl34b-Δ	S288C			Steffen KK, et al. (2008) PMID:18423200
RPL34B	YIL052C	replicative lifespan: increased 	classical genetics	classical genetics	null Allele: rpl34b-Δ	S288C			Steffen KK, et al. (2012) PMID:22377630
RPL34B	YIL052C	replicative lifespan: increased 	classical genetics	classical genetics	null Allele: rpl34b-Δ	S288C			McCormick MA, et al. (2015) PMID:26456335
RPL35A	YDL191W	replicative lifespan: increased 	classical genetics	classical genetics	null Allele: rpl35a-Δ	S288C			Steffen KK, et al. (2012) PMID:22377630
RPL35A	YDL191W	replicative lifespan: increased 	classical genetics	classical genetics	null Allele: rpl35a-Δ	S288C			McCormick MA, et al. (2015) PMID:26456335
RPL37B	YDR500C	replicative lifespan: increased 	large-scale survey	large-scale survey	null Allele: rpl37b-Δ	S288C			Managbanag JR, et al. (2008) PMID:19030232
RPL37B	YDR500C	replicative lifespan: increased 	classical genetics	classical genetics	null Allele: rpl37b-Δ	S288C			Delaney JR, et al. (2011) PMID:21902802
RPL37B	YDR500C	replicative lifespan: increased 	competitive growth	large-scale survey	null Allele: rpl37b-Δ	S288C			Delaney JR, et al. (2011) PMID:21191185
RPL37B	YDR500C	replicative lifespan: increased 	classical genetics	classical genetics	null Allele: rpl37b-Δ	S288C			McCormick MA, et al. (2015) PMID:26456335
RPL40B	YKR094C	replicative lifespan: increased 	systematic mutation set 	large-scale survey	null Allele: rpl40b-Δ	S288C		Details: long-lived mutants, one standard deviation above the mean, were identified using an HTP sequencing-based yeast replicative lifespan screen (SEBYL), with pooled cells subject to four rounds of biotin-based sorting to enrich old vs young cells, followed by sequencing to ID mutants and abundance 	Yu R, et al. (2021) PMID:33790287
RPL41B	YDL133C-A	replicative lifespan: increased 	systematic mutation set 	large-scale survey	null Allele: rpl41b-Δ	S288C		Details: long-lived mutants, one standard deviation above the mean, were identified using an HTP sequencing-based yeast replicative lifespan screen (SEBYL), with pooled cells subject to four rounds of biotin-based sorting to enrich old vs young cells, followed by sequencing to ID mutants and abundance 	Yu R, et al. (2021) PMID:33790287
RPL43B	YJR094W-A	replicative lifespan: increased 	classical genetics	classical genetics	null Allele: rpl43b-Δ	S288C			Steffen KK, et al. (2008) PMID:18423200
RPL43B	YJR094W-A	replicative lifespan: increased 	classical genetics	classical genetics	null Allele: rpl43b-Δ	S288C			Delaney JR, et al. (2011) PMID:21902802
RPL43B	YJR094W-A	replicative lifespan: increased 	competitive growth	large-scale survey	null Allele: rpl43b-Δ	S288C			Delaney JR, et al. (2011) PMID:21191185
RPL43B	YJR094W-A	replicative lifespan: increased 	classical genetics	classical genetics	null Allele: rpl43b-Δ	S288C			McCormick MA, et al. (2015) PMID:26456335
RPL6A	YML073C	replicative lifespan: increased 	classical genetics	classical genetics	null Allele: rpl6a-Δ	S288C			Steffen KK, et al. (2012) PMID:22377630
RPL6A	YML073C	replicative lifespan: increased 	classical genetics	classical genetics	null Allele: rpl6a-Δ	S288C			McCormick MA, et al. (2015) PMID:26456335
RPL6B	YLR448W	replicative lifespan: increased 	large-scale survey	large-scale survey	null Allele: rpl6b-Δ	S288C			Smith ED, et al. (2008) PMID:18340043
RPL6B	YLR448W	replicative lifespan: increased 	classical genetics	classical genetics	null Allele: rpl6b-Δ	S288C			Kaeberlein M, et al. (2005) PMID:16293764
RPL6B	YLR448W	replicative lifespan: increased 	large-scale survey	large-scale survey	null Allele: rpl6b-Δ	S288C			Kaeberlein M, et al. (2005) PMID:16293764
RPL6B	YLR448W	replicative lifespan: increased 	classical genetics	classical genetics	null Allele: rpl6b-Δ	S288C			Steffen KK, et al. (2008) PMID:18423200
RPL6B	YLR448W	replicative lifespan: increased 	competitive growth	large-scale survey	null Allele: rpl6b-Δ	S288C			Delaney JR, et al. (2011) PMID:21191185
RPL6B	YLR448W	replicative lifespan: increased 	classical genetics	classical genetics	null Allele: rpl6b-Δ	S288C			McCormick MA, et al. (2015) PMID:26456335
RPL7A	YGL076C	replicative lifespan: increased 	classical genetics	classical genetics	null Allele: rpl7a-Δ	S288C			Steffen KK, et al. (2008) PMID:18423200
RPL7A	YGL076C	replicative lifespan: increased 	classical genetics	classical genetics	null Allele: rpl7a-Δ	S288C			Delaney JR, et al. (2011) PMID:21902802
RPL7A	YGL076C	replicative lifespan: increased 	competitive growth	large-scale survey	null Allele: rpl7a-Δ	S288C			Delaney JR, et al. (2011) PMID:21191185
RPL7A	YGL076C	replicative lifespan: increased 	classical genetics	classical genetics	null Allele: rpl7a-Δ	S288C			McCormick MA, et al. (2015) PMID:26456335
RPL9A	YGL147C	replicative lifespan: increased 	large-scale survey	large-scale survey	null Allele: rpl9a-Δ	S288C			Smith ED, et al. (2008) PMID:18340043
RPL9A	YGL147C	replicative lifespan: increased 	classical genetics	classical genetics	null Allele: rpl9a-Δ	S288C			Steffen KK, et al. (2008) PMID:18423200
RPL9A	YGL147C	replicative lifespan: increased 	competitive growth	large-scale survey	null Allele: rpl9a-Δ	S288C			Delaney JR, et al. (2011) PMID:21191185
RPL9A	YGL147C	replicative lifespan: increased 	classical genetics	classical genetics	null Allele: rpl9a-Δ	S288C			McCormick MA, et al. (2015) PMID:26456335
RPL9A	YGL147C	replicative lifespan: increased 	systematic mutation set 	large-scale survey	null Allele: rpl9a-Δ	S288C		Details: long-lived mutants, one standard deviation above the mean, were identified using an HTP sequencing-based yeast replicative lifespan screen (SEBYL), with pooled cells subject to four rounds of biotin-based sorting to enrich old vs young cells, followed by sequencing to ID mutants and abundance 	Yu R, et al. (2021) PMID:33790287
RPN4	YDL020C	replicative lifespan: increased 	classical genetics	classical genetics	null Allele: rpn4-Δ	S288C		Treatment: caloric restriction by glucose reduction, 0.05% Details: dietary restriction-induced lifespan extension is increased relative to wt 	Schleit J, et al. (2013) PMID:23837470
RPP1A	YDL081C	replicative lifespan: increased 	classical genetics	classical genetics	null Allele: rpp1a-Δ	S288C			Steffen KK, et al. (2012) PMID:22377630
RPP1A	YDL081C	replicative lifespan: increased 	classical genetics	classical genetics	null Allele: rpp1a-Δ	S288C			McCormick MA, et al. (2015) PMID:26456335
RPP1B	YDL130W	replicative lifespan: increased 	classical genetics	classical genetics	null Allele: rpp1b-Δ	S288C			McCormick MA, et al. (2015) PMID:26456335
RPP1B	YDL130W	replicative lifespan: increased 	systematic mutation set 	large-scale survey	null Allele: rpp1b-Δ	S288C		Details: long-lived mutants, one standard deviation above the mean, were identified using an HTP sequencing-based yeast replicative lifespan screen (SEBYL), with pooled cells subject to four rounds of biotin-based sorting to enrich old vs young cells, followed by sequencing to ID mutants and abundance 	Yu R, et al. (2021) PMID:33790287
RPP2A	YOL039W	replicative lifespan: increased 	classical genetics	classical genetics	null Allele: rpp2a-Δ	S288C			McCormick MA, et al. (2015) PMID:26456335
RPP2B	YDR382W	replicative lifespan: increased 	classical genetics	classical genetics	null Allele: rpp2b-Δ	S288C			Steffen KK, et al. (2008) PMID:18423200
RPP2B	YDR382W	replicative lifespan: increased 	classical genetics	classical genetics	null Allele: rpp2b-Δ	S288C			Delaney JR, et al. (2011) PMID:21902802
RPP2B	YDR382W	replicative lifespan: increased 	competitive growth	large-scale survey	null Allele: rpp2b-Δ	S288C			Delaney JR, et al. (2011) PMID:21191185
RPP2B	YDR382W	replicative lifespan: increased 	classical genetics	classical genetics	null Allele: rpp2b-Δ	S288C			McCormick MA, et al. (2015) PMID:26456335
RPS12	YOR369C	replicative lifespan: increased 	classical genetics	classical genetics	null Allele: rps12-Δ	S288C			McCormick MA, et al. (2015) PMID:26456335
RPS17A	YML024W	replicative lifespan: increased 	classical genetics	classical genetics	null Allele: rps17a-Δ	S288C		Treatment: caloric restriction by glucose reduction, 0.05% Details: dietary restriction-induced lifespan extension is increased relative to wt 	Schleit J, et al. (2013) PMID:23837470
RPS19B	YNL302C	replicative lifespan: increased 	systematic mutation set 	large-scale survey	null Allele: rps19b-Δ	S288C		Details: long-lived mutants, one standard deviation above the mean, were identified using an HTP sequencing-based yeast replicative lifespan screen (SEBYL), with pooled cells subject to four rounds of biotin-based sorting to enrich old vs young cells, followed by sequencing to ID mutants and abundance 	Yu R, et al. (2021) PMID:33790287
RPS1A	YLR441C	replicative lifespan: increased 	large-scale survey	large-scale survey	null Allele: rps1a-Δ	S288C			Kaeberlein M, et al. (2005) PMID:16293764
RPS21A	YKR057W	replicative lifespan: increased 	large-scale survey	large-scale survey	null Allele: rps21a-Δ	S288C			Kaeberlein M, et al. (2005) PMID:16293764
RPS22B	YLR367W	replicative lifespan: increased 	large-scale survey	large-scale survey	null Allele: rps22b-Δ	S288C			Kaeberlein M, et al. (2005) PMID:16293764
RPS23B	YPR132W	replicative lifespan: increased 	systematic mutation set 	large-scale survey	null Allele: rps23b-Δ	S288C		Details: long-lived mutants, one standard deviation above the mean, were identified using an HTP sequencing-based yeast replicative lifespan screen (SEBYL), with pooled cells subject to four rounds of biotin-based sorting to enrich old vs young cells, followed by sequencing to ID mutants and abundance 	Yu R, et al. (2021) PMID:33790287
RPS24A	YER074W	replicative lifespan: increased 	systematic mutation set 	large-scale survey	null Allele: rps24a-Δ	S288C		Details: long-lived mutants, one standard deviation above the mean, were identified using an HTP sequencing-based yeast replicative lifespan screen (SEBYL), with pooled cells subject to four rounds of biotin-based sorting to enrich old vs young cells, followed by sequencing to ID mutants and abundance 	Yu R, et al. (2021) PMID:33790287
RPS6A	YPL090C	replicative lifespan: increased 	classical genetics	classical genetics	null Allele: rps6a-Δ	S288C			McCormick MA, et al. (2015) PMID:26456335
RQC1	YDR333C	replicative lifespan: increased 	systematic mutation set 	large-scale survey	null Allele: rqc1-Δ	S288C		Details: long-lived mutants, one standard deviation above the mean, were identified using an HTP sequencing-based yeast replicative lifespan screen (SEBYL), with pooled cells subject to four rounds of biotin-based sorting to enrich old vs young cells, followed by sequencing to ID mutants and abundance 	Yu R, et al. (2021) PMID:33790287
RRD2	YPL152W	replicative lifespan: increased 	classical genetics	classical genetics	null Allele: rrd2-Δ	S288C			McCormick MA, et al. (2015) PMID:26456335
RRG9	YNL213C	replicative lifespan: increased 	classical genetics	classical genetics	null Allele: rrg9-Δ	S288C			McCormick MA, et al. (2015) PMID:26456335
RRT16	YNL105W	replicative lifespan: increased 	systematic mutation set 	large-scale survey	null Allele: rrt16-Δ	S288C		Details: long-lived mutants, one standard deviation above the mean, were identified using an HTP sequencing-based yeast replicative lifespan screen (SEBYL), with pooled cells subject to four rounds of biotin-based sorting to enrich old vs young cells, followed by sequencing to ID mutants and abundance 	Yu R, et al. (2021) PMID:33790287
RSA3	YLR221C	replicative lifespan: increased 	systematic mutation set 	large-scale survey	null Allele: rsa3-Δ	S288C		Details: long-lived mutants, one standard deviation above the mean, were identified using an HTP sequencing-based yeast replicative lifespan screen (SEBYL), with pooled cells subject to four rounds of biotin-based sorting to enrich old vs young cells, followed by sequencing to ID mutants and abundance 	Yu R, et al. (2021) PMID:33790287
RTA1	YGR213C	replicative lifespan: increased 	systematic mutation set 	large-scale survey	null Allele: rta1-Δ	S288C		Details: long-lived mutants, one standard deviation above the mean, were identified using an HTP sequencing-based yeast replicative lifespan screen (SEBYL), with pooled cells subject to four rounds of biotin-based sorting to enrich old vs young cells, followed by sequencing to ID mutants and abundance 	Yu R, et al. (2021) PMID:33790287
RTG2	YGL252C	replicative lifespan: increased 	classical genetics 	classical genetics	null Allele: rtg2-L56G	Other			Rios-Anjos RM, et al. (2017) PMID:28472157
RTK1	YDL025C	replicative lifespan: increased 	systematic mutation set 	large-scale survey	null Allele: rtk1-Δ	S288C		Details: long-lived mutants, one standard deviation above the mean, were identified using an HTP sequencing-based yeast replicative lifespan screen (SEBYL), with pooled cells subject to four rounds of biotin-based sorting to enrich old vs young cells, followed by sequencing to ID mutants and abundance 	Yu R, et al. (2021) PMID:33790287
RTN1	YDR233C	replicative lifespan: increased 	large-scale survey	large-scale survey	null Allele: rtn1-Δ	S288C			Managbanag JR, et al. (2008) PMID:19030232
RTS1	YOR014W	replicative lifespan: increased 	classical genetics	classical genetics	null Allele: rts1-Δ	S288C			McCormick MA, et al. (2015) PMID:26456335
RTS3	YGR161C	replicative lifespan: increased 	systematic mutation set 	large-scale survey	null Allele: rts3-Δ	S288C		Details: long-lived mutants, one standard deviation above the mean, were identified using an HTP sequencing-based yeast replicative lifespan screen (SEBYL), with pooled cells subject to four rounds of biotin-based sorting to enrich old vs young cells, followed by sequencing to ID mutants and abundance 	Yu R, et al. (2021) PMID:33790287
RUD3	YOR216C	replicative lifespan: increased 	systematic mutation set 	large-scale survey	null Allele: rud3-Δ	S288C		Details: long-lived mutants, one standard deviation above the mean, were identified using an HTP sequencing-based yeast replicative lifespan screen (SEBYL), with pooled cells subject to four rounds of biotin-based sorting to enrich old vs young cells, followed by sequencing to ID mutants and abundance 	Yu R, et al. (2021) PMID:33790287
RUP1	YOR138C	replicative lifespan: increased 	systematic mutation set 	large-scale survey	null Allele: rup1-Δ	S288C		Details: long-lived mutants, one standard deviation above the mean, were identified using an HTP sequencing-based yeast replicative lifespan screen (SEBYL), with pooled cells subject to four rounds of biotin-based sorting to enrich old vs young cells, followed by sequencing to ID mutants and abundance 	Yu R, et al. (2021) PMID:33790287
RVS161	YCR009C	replicative lifespan: increased 	large-scale survey	large-scale survey	null Allele: rvs161-Δ	S288C			Managbanag JR, et al. (2008) PMID:19030232
SAE3	YHR079C-A	replicative lifespan: increased 	systematic mutation set 	large-scale survey	null Allele: sae3-Δ	S288C		Details: long-lived mutants, one standard deviation above the mean, were identified using an HTP sequencing-based yeast replicative lifespan screen (SEBYL), with pooled cells subject to four rounds of biotin-based sorting to enrich old vs young cells, followed by sequencing to ID mutants and abundance 	Yu R, et al. (2021) PMID:33790287
SAM1	YLR180W	replicative lifespan: increased 	large-scale survey	large-scale survey	null Allele: sam1-Δ	S288C			Smith ED, et al. (2008) PMID:18340043
SAM1	YLR180W	replicative lifespan: increased 	classical genetics	classical genetics	null Allele: sam1-Δ	S288C			Delaney JR, et al. (2011) PMID:21902802
SAM1	YLR180W	replicative lifespan: increased 	competitive growth	large-scale survey	null Allele: sam1-Δ	S288C			Delaney JR, et al. (2011) PMID:21191185
SAM1	YLR180W	replicative lifespan: increased 	classical genetics	classical genetics	null Allele: sam1-Δ	S288C			McCormick MA, et al. (2015) PMID:26456335
SAN1	YDR143C	replicative lifespan: increased 	systematic mutation set 	large-scale survey	null Allele: san1-Δ	S288C		Details: long-lived mutants, one standard deviation above the mean, were identified using an HTP sequencing-based yeast replicative lifespan screen (SEBYL), with pooled cells subject to four rounds of biotin-based sorting to enrich old vs young cells, followed by sequencing to ID mutants and abundance 	Yu R, et al. (2021) PMID:33790287
SAP155	YFR040W	replicative lifespan: increased 	classical genetics	classical genetics	null Allele: sap155-Δ	S288C			McCormick MA, et al. (2015) PMID:26456335
SAP185	YJL098W	replicative lifespan: increased 	classical genetics	classical genetics	null Allele: sap185-Δ	S288C			McCormick MA, et al. (2015) PMID:26456335
SAP4	YGL229C	replicative lifespan: increased 	systematic mutation set 	large-scale survey	null Allele: sap4-Δ	S288C		Details: long-lived mutants, one standard deviation above the mean, were identified using an HTP sequencing-based yeast replicative lifespan screen (SEBYL), with pooled cells subject to four rounds of biotin-based sorting to enrich old vs young cells, followed by sequencing to ID mutants and abundance 	Yu R, et al. (2021) PMID:33790287
SAS2	YMR127C	replicative lifespan: increased 	classical genetics	classical genetics	null Allele: sas2-Δ	S288C			Delaney JR, et al. (2011) PMID:21902802
SAS2	YMR127C	replicative lifespan: increased 	classical genetics	classical genetics	null Allele: sas2-Δ	S288C			McCormick MA, et al. (2015) PMID:26456335
SAS3	YBL052C	replicative lifespan: increased 	classical genetics	classical genetics	null Allele: sas3-Δ	S288C			McCormick MA, et al. (2015) PMID:26456335
SCH9	YHR205W	replicative lifespan: increased 	large-scale survey	large-scale survey	null Allele: sch9-Δ	S288C			Smith ED, et al. (2008) PMID:18340043
SCH9	YHR205W	replicative lifespan: increased 	classical genetics	classical genetics	null Allele: sch9-Δ	S288C		Details: caloric restriction fails to further increase lifespan 	Kaeberlein M, et al. (2005) PMID:16293764
SCH9	YHR205W	replicative lifespan: increased 	classical genetics	classical genetics	null Allele: sch9-Δ	S288C			Delaney JR, et al. (2011) PMID:21902802
SCH9	YHR205W	replicative lifespan: increased 	competitive growth	large-scale survey	null Allele: sch9-Δ	S288C			Delaney JR, et al. (2011) PMID:21191185
SCH9	YHR205W	replicative lifespan: increased 	classical genetics	classical genetics	null Allele: sch9-Δ	S288C			Schleit J, et al. (2013) PMID:23837470
SCH9	YHR205W	replicative lifespan: increased 	classical genetics	classical genetics	null Allele: sch9-Δ	S288C			McCormick MA, et al. (2015) PMID:26456335
SCH9	YHR205W	replicative lifespan: increased 	systematic mutation set	large-scale survey	null Allele: sch9-Δ	S288C			Delaney JR, et al. (2013) PMID:23336757
SCP1	YOR367W	replicative lifespan: increased 	classical genetics	classical genetics	null Allele: scp1-Δ	Other			Gourlay CW, et al. (2004) PMID:15024029
SCW4	YGR279C	replicative lifespan: increased 	classical genetics	classical genetics	null Allele: scw4-Δ	S288C			McCormick MA, et al. (2015) PMID:26456335
SDH1	YKL148C	replicative lifespan: increased 	classical genetics	classical genetics	null Allele: sdh1-Δ	S288C			McCormick MA, et al. (2015) PMID:26456335
SDH2	YLL041C	replicative lifespan: increased 	classical genetics	classical genetics	null Allele: sdh2-Δ	S288C			McCormick MA, et al. (2015) PMID:26456335
SDH5	YOL071W	replicative lifespan: increased 	systematic mutation set 	large-scale survey	null Allele: sdh5-Δ	S288C		Details: long-lived mutants, one standard deviation above the mean, were identified using an HTP sequencing-based yeast replicative lifespan screen (SEBYL), with pooled cells subject to four rounds of biotin-based sorting to enrich old vs young cells, followed by sequencing to ID mutants and abundance 	Yu R, et al. (2021) PMID:33790287
SDH5	YOL071W	replicative lifespan: increased 	classical genetics	classical genetics	null Allele: sdh5-Δ	S288C			McCormick MA, et al. (2015) PMID:26456335
SDS3	YIL084C	replicative lifespan: increased 	classical genetics	classical genetics	null Allele: sds3-Δ	S288C			McCormick MA, et al. (2015) PMID:26456335
SEC28	YIL076W	replicative lifespan: increased 	large-scale survey	large-scale survey	null Allele: sec28-Δ	S288C			Managbanag JR, et al. (2008) PMID:19030232
SEY1	YOR165W	replicative lifespan: increased 	systematic mutation set 	large-scale survey	null Allele: sey1-Δ	S288C		Details: long-lived mutants, one standard deviation above the mean, were identified using an HTP sequencing-based yeast replicative lifespan screen (SEBYL), with pooled cells subject to four rounds of biotin-based sorting to enrich old vs young cells, followed by sequencing to ID mutants and abundance 	Yu R, et al. (2021) PMID:33790287
SGF11	YPL047W	replicative lifespan: increased 	classical genetics	classical genetics	null Allele: sgf11-Δ	S288C			McCormick MA, et al. (2015) PMID:26456335
SGF73	YGL066W	replicative lifespan: increased 	classical genetics	classical genetics	null Allele: sgf73-Δ	S288C		Treatment: caloric restriction by glucose reduction, 0.05% Details: dietary restriction-induced lifespan extension is increased relative to wt 	Schleit J, et al. (2013) PMID:23837470
SGF73	YGL066W	replicative lifespan: increased 	classical genetics	classical genetics	null Allele: sgf73-Δ	S288C			McCormick MA, et al. (2015) PMID:26456335
SGS1	YMR190C	replicative lifespan: increased 	classical genetics	classical genetics	null Allele: sgs1-Δ	S288C		Treatment: caloric restriction by glucose reduction, 0.05% Details: dietary restriction-induced lifespan extension is increased relative to wt 	Schleit J, et al. (2013) PMID:23837470
SHE10	YGL228W	replicative lifespan: increased 	systematic mutation set 	large-scale survey	null Allele: she10-Δ	S288C		Details: long-lived mutants, one standard deviation above the mean, were identified using an HTP sequencing-based yeast replicative lifespan screen (SEBYL), with pooled cells subject to four rounds of biotin-based sorting to enrich old vs young cells, followed by sequencing to ID mutants and abundance 	Yu R, et al. (2021) PMID:33790287
SIP2	YGL208W	replicative lifespan: increased 	large-scale survey	large-scale survey	null Allele: sip2-Δ	S288C			Managbanag JR, et al. (2008) PMID:19030232
SIP2	YGL208W	replicative lifespan: increased 	classical genetics	classical genetics	null Allele: sip2-Δ	S288C			Delaney JR, et al. (2011) PMID:21902802
SIP2	YGL208W	replicative lifespan: increased 	competitive growth	large-scale survey	null Allele: sip2-Δ	S288C			Delaney JR, et al. (2011) PMID:21191185
SIR2	YDL042C	replicative lifespan: increased 	classical genetics	classical genetics	overexpression	W303		Details: 30% extension of replicative lifespan relative to wildtype, upon the integration of one additional copy of SIR2 	Kaeberlein M, et al. (1999) PMID:10521401
SIR2	YDL042C	replicative lifespan: increased 	classical genetics	classical genetics	unspecified Allele: sir2-S473A phosphorylation site mutated; mediated by cAMP-PKA and CK2 signaling	S288C		Details: RLS increased by 20% 	Kang WK, et al. (2015) PMID:26329457
SIR4	YDR227W	replicative lifespan: increased 	classical genetics	classical genetics	gain of function Allele: sir4-(uth2-42) also known as sir4-42	Other		Details: semi dominant mutant 	Kennedy BK, et al. (1995) PMID:7859289
SIS2	YKR072C	replicative lifespan: increased 	large-scale survey	large-scale survey	null Allele: sis2-Δ	S288C			Smith ED, et al. (2008) PMID:18340043
SIS2	YKR072C	replicative lifespan: increased 	classical genetics	classical genetics	null Allele: sis2-Δ	S288C			McCormick MA, et al. (2015) PMID:26456335
SIS2	YKR072C	replicative lifespan: increased 	competitive growth	large-scale survey	null Allele: sis2-Δ	S288C			Delaney JR, et al. (2011) PMID:21191185
SIW14	YNL032W	replicative lifespan: increased 	systematic mutation set 	large-scale survey	null Allele: siw14-Δ	S288C		Details: long-lived mutants, one standard deviation above the mean, were identified using an HTP sequencing-based yeast replicative lifespan screen (SEBYL), with pooled cells subject to four rounds of biotin-based sorting to enrich old vs young cells, followed by sequencing to ID mutants and abundance 	Yu R, et al. (2021) PMID:33790287
SKN1	YGR143W	replicative lifespan: increased 	systematic mutation set 	large-scale survey	null Allele: skn1-Δ	S288C		Details: long-lived mutants, one standard deviation above the mean, were identified using an HTP sequencing-based yeast replicative lifespan screen (SEBYL), with pooled cells subject to four rounds of biotin-based sorting to enrich old vs young cells, followed by sequencing to ID mutants and abundance 	Yu R, et al. (2021) PMID:33790287
SKP2	YNL311C	replicative lifespan: increased 	classical genetics	classical genetics	null Allele: skp2-Δ	S288C			McCormick MA, et al. (2015) PMID:26456335
SLM6	YBR266C	replicative lifespan: increased 	systematic mutation set 	large-scale survey	null Allele: slm6-Δ	S288C		Details: long-lived mutants, one standard deviation above the mean, were identified using an HTP sequencing-based yeast replicative lifespan screen (SEBYL), with pooled cells subject to four rounds of biotin-based sorting to enrich old vs young cells, followed by sequencing to ID mutants and abundance 	Yu R, et al. (2021) PMID:33790287
SLM6	YBR266C	replicative lifespan: increased 	competitive growth	large-scale survey	null Allele: slm6-Δ	S288C			Delaney JR, et al. (2011) PMID:21191185
SLM6	YBR266C	replicative lifespan: increased 	classical genetics	classical genetics	null Allele: slm6-Δ	S288C			McCormick MA, et al. (2015) PMID:26456335
SMF2	YHR050W	replicative lifespan: increased 	systematic mutation set 	large-scale survey	null Allele: smf2-Δ	S288C		Details: long-lived mutants, one standard deviation above the mean, were identified using an HTP sequencing-based yeast replicative lifespan screen (SEBYL), with pooled cells subject to four rounds of biotin-based sorting to enrich old vs young cells, followed by sequencing to ID mutants and abundance 	Yu R, et al. (2021) PMID:33790287
SMI1	YGR229C	replicative lifespan: increased 	classical genetics 	classical genetics	null Allele: smi1-Δ	S288C		Details: RLS extended by about 25% relative to wt cells; extension is dependent on SIR2 and HOG1 	Hong S and Huh WK (2021) PMID:33402426
SMI1	YGR229C	replicative lifespan: increased 	classical genetics 	classical genetics	null Allele: smi1-Δ	S288C			Molon M, et al. (2018) PMID:29189912
SMI1	YGR229C	replicative lifespan: increased 	classical genetics 	classical genetics	null Allele: smi1-Δ	S288C		Details: RLS extended in a Sir2-dependent manner by 25% in the null mutant relative to wt cells 	Hong S and Huh WK (2021) PMID:33837734
SMY2	YBR172C	replicative lifespan: increased 	classical genetics	classical genetics	null Allele: smy2-Δ	S288C			McCormick MA, et al. (2015) PMID:26456335
SNF1	YDR477W	replicative lifespan: increased 	large-scale survey	large-scale survey	null Allele: snf1-Δ	S288C			Managbanag JR, et al. (2008) PMID:19030232
SNF1	YDR477W	replicative lifespan: increased 	large-scale survey	large-scale survey	null Allele: snf1-Δ	S288C			Kaeberlein M, et al. (2005) PMID:16293764
SNF4	YGL115W	replicative lifespan: increased 	classical genetics	classical genetics	null Allele: snf4-Δ	S288C		Details: mean lifespan increased to 110-120% that of the wild type 	Ashrafi K, et al. (2000) PMID:10921902
SNO2	YNL334C	replicative lifespan: increased 	systematic mutation set 	large-scale survey	null Allele: sno2-Δ	S288C		Details: long-lived mutants, one standard deviation above the mean, were identified using an HTP sequencing-based yeast replicative lifespan screen (SEBYL), with pooled cells subject to four rounds of biotin-based sorting to enrich old vs young cells, followed by sequencing to ID mutants and abundance 	Yu R, et al. (2021) PMID:33790287
SOK1	YDR006C	replicative lifespan: increased 	large-scale survey	large-scale survey	null Allele: sok1-Δ	S288C			Managbanag JR, et al. (2008) PMID:19030232
SOK1	YDR006C	replicative lifespan: increased 	competitive growth	large-scale survey	null Allele: sok1-Δ	S288C			Delaney JR, et al. (2011) PMID:21191185
SOK1	YDR006C	replicative lifespan: increased 	classical genetics	classical genetics	null Allele: sok1-Δ	S288C			McCormick MA, et al. (2015) PMID:26456335
SOV1	YMR066W	replicative lifespan: increased 	classical genetics	classical genetics	null Allele: sov1-Δ	S288C			Caballero A, et al. (2011) PMID:21549315
SOV1	YMR066W	replicative lifespan: increased 	classical genetics	classical genetics	null Allele: sov1-Δ	S288C			McCormick MA, et al. (2015) PMID:26456335
SPF1	YEL031W	replicative lifespan: increased 	systematic mutation set 	large-scale survey	null Allele: spf1-Δ	S288C		Details: long-lived mutants, one standard deviation above the mean, were identified using an HTP sequencing-based yeast replicative lifespan screen (SEBYL), with pooled cells subject to four rounds of biotin-based sorting to enrich old vs young cells, followed by sequencing to ID mutants and abundance 	Yu R, et al. (2021) PMID:33790287
SPO20	YMR017W	replicative lifespan: increased 	systematic mutation set 	large-scale survey	null Allele: spo20-Δ	S288C		Details: long-lived mutants, one standard deviation above the mean, were identified using an HTP sequencing-based yeast replicative lifespan screen (SEBYL), with pooled cells subject to four rounds of biotin-based sorting to enrich old vs young cells, followed by sequencing to ID mutants and abundance 	Yu R, et al. (2021) PMID:33790287
SPS1	YDR523C	replicative lifespan: increased 	large-scale survey	large-scale survey	null Allele: sps1-Δ	S288C			Managbanag JR, et al. (2008) PMID:19030232
SPS1	YDR523C	replicative lifespan: increased 	classical genetics	classical genetics	null Allele: sps1-Δ	S288C			Delaney JR, et al. (2011) PMID:21902802
SPS1	YDR523C	replicative lifespan: increased 	competitive growth	large-scale survey	null Allele: sps1-Δ	S288C			Delaney JR, et al. (2011) PMID:21191185
SPS1	YDR523C	replicative lifespan: increased 	systematic mutation set	large-scale survey	null Allele: sps1-Δ	S288C			Delaney JR, et al. (2013) PMID:23336757
SPS1	YDR523C	replicative lifespan: increased 	classical genetics	classical genetics	null Allele: sps1-Δ	S288C			McCormick MA, et al. (2015) PMID:26456335
SPT2	YER161C	replicative lifespan: increased 	systematic mutation set 	large-scale survey	null Allele: spt2-Δ	S288C		Details: long-lived mutants, one standard deviation above the mean, were identified using an HTP sequencing-based yeast replicative lifespan screen (SEBYL), with pooled cells subject to four rounds of biotin-based sorting to enrich old vs young cells, followed by sequencing to ID mutants and abundance 	Yu R, et al. (2021) PMID:33790287
SPT23	YKL020C	replicative lifespan: increased 	classical genetics	classical genetics	null Allele: spt23-Δ	S288C			McCormick MA, et al. (2015) PMID:26456335
SPT4	YGR063C	replicative lifespan: increased 	large-scale survey	large-scale survey	null Allele: spt4-Δ	S288C			Smith ED, et al. (2008) PMID:18340043
SPT4	YGR063C	replicative lifespan: increased 	competitive growth	large-scale survey	null Allele: spt4-Δ	S288C			Delaney JR, et al. (2011) PMID:21191185
SPT4	YGR063C	replicative lifespan: increased 	classical genetics	classical genetics	null Allele: spt4-Δ	S288C			McCormick MA, et al. (2015) PMID:26456335
SPT8	YLR055C	replicative lifespan: increased 	classical genetics	classical genetics	null Allele: spt8-Δ	S288C			McCormick MA, et al. (2015) PMID:26456335
SRL1	YOR247W	replicative lifespan: increased 	systematic mutation set 	large-scale survey	null Allele: srl1-Δ	S288C		Details: long-lived mutants, one standard deviation above the mean, were identified using an HTP sequencing-based yeast replicative lifespan screen (SEBYL), with pooled cells subject to four rounds of biotin-based sorting to enrich old vs young cells, followed by sequencing to ID mutants and abundance 	Yu R, et al. (2021) PMID:33790287
SRN2	YLR119W	replicative lifespan: increased 	systematic mutation set 	large-scale survey	null Allele: srn2-Δ	S288C		Details: long-lived mutants, one standard deviation above the mean, were identified using an HTP sequencing-based yeast replicative lifespan screen (SEBYL), with pooled cells subject to four rounds of biotin-based sorting to enrich old vs young cells, followed by sequencing to ID mutants and abundance 	Yu R, et al. (2021) PMID:33790287
SSC1	YJR045C	replicative lifespan: increased 	heterozygous diploid	classical genetics	null Allele: ssc1-Δ	Other			Perić M, et al. (2016) PMID:27346163
SSD1	YDR293C	replicative lifespan: increased 	classical genetics 	classical genetics	overexpression	Other			Hu Z, et al. (2018) PMID:30117416
SSF1	YHR066W	replicative lifespan: increased 	large-scale survey	large-scale survey	null Allele: ssf1-Δ	S288C			Kaeberlein M, et al. (2005) PMID:16293764
SSF1	YHR066W	replicative lifespan: increased 	classical genetics	classical genetics	null Allele: ssf1-Δ	S288C			Steffen KK, et al. (2008) PMID:18423200
SSF1	YHR066W	replicative lifespan: increased 	classical genetics	classical genetics	null Allele: ssf1-Δ	S288C			McCormick MA, et al. (2015) PMID:26456335
SSN2	YDR443C	replicative lifespan: increased 	systematic mutation set 	large-scale survey	null Allele: ssn2-Δ	S288C		Details: long-lived mutants, one standard deviation above the mean, were identified using an HTP sequencing-based yeast replicative lifespan screen (SEBYL), with pooled cells subject to four rounds of biotin-based sorting to enrich old vs young cells, followed by sequencing to ID mutants and abundance 	Yu R, et al. (2021) PMID:33790287
SST2	YLR452C	replicative lifespan: increased 	large-scale survey	large-scale survey	null Allele: sst2-Δ	S288C			Kaeberlein M, et al. (2005) PMID:16293764
STB2	YMR053C	replicative lifespan: increased 	large-scale survey	large-scale survey	null Allele: stb2-Δ	S288C			Kaeberlein M, et al. (2005) PMID:16293764
STP2	YHR006W	replicative lifespan: increased 	systematic mutation set 	large-scale survey	null Allele: stp2-Δ	S288C		Details: long-lived mutants, one standard deviation above the mean, were identified using an HTP sequencing-based yeast replicative lifespan screen (SEBYL), with pooled cells subject to four rounds of biotin-based sorting to enrich old vs young cells, followed by sequencing to ID mutants and abundance 	Yu R, et al. (2021) PMID:33790287
STV1	YMR054W	replicative lifespan: increased 	classical genetics	classical genetics	null Allele: stv1-Δ	S288C		Treatment: caloric restriction by glucose reduction, 0.05% Details: dietary restriction-induced lifespan extension is increased relative to wt 	Schleit J, et al. (2013) PMID:23837470
SUM1	YDR310C	replicative lifespan: increased 	systematic mutation set 	large-scale survey	null Allele: sum1-Δ	S288C		Details: long-lived mutants, one standard deviation above the mean, were identified using an HTP sequencing-based yeast replicative lifespan screen (SEBYL), with pooled cells subject to four rounds of biotin-based sorting to enrich old vs young cells, followed by sequencing to ID mutants and abundance 	Yu R, et al. (2021) PMID:33790287
SUV3	YPL029W	replicative lifespan: increased 	classical genetics	classical genetics	null Allele: suv3-Δ	S288C			Caballero A, et al. (2011) PMID:21549315
SUV3	YPL029W	replicative lifespan: increased 	classical genetics	classical genetics	null Allele: suv3-Δ	S288C			McCormick MA, et al. (2015) PMID:26456335
SWC3	YAL011W	replicative lifespan: increased 	systematic mutation set 	large-scale survey	null Allele: swc3-Δ	S288C		Details: long-lived mutants, one standard deviation above the mean, were identified using an HTP sequencing-based yeast replicative lifespan screen (SEBYL), with pooled cells subject to four rounds of biotin-based sorting to enrich old vs young cells, followed by sequencing to ID mutants and abundance 	Yu R, et al. (2021) PMID:33790287
SWI5	YDR146C	replicative lifespan: increased 	large-scale survey	large-scale survey	null Allele: swi5-Δ	S288C			Managbanag JR, et al. (2008) PMID:19030232
SWT1	YOR166C	replicative lifespan: increased 	systematic mutation set 	large-scale survey	null Allele: swt1-Δ	S288C		Details: long-lived mutants, one standard deviation above the mean, were identified using an HTP sequencing-based yeast replicative lifespan screen (SEBYL), with pooled cells subject to four rounds of biotin-based sorting to enrich old vs young cells, followed by sequencing to ID mutants and abundance 	Yu R, et al. (2021) PMID:33790287
SWT21	YNL187W	replicative lifespan: increased 	systematic mutation set 	large-scale survey	null Allele: swt21-Δ	S288C		Details: long-lived mutants, one standard deviation above the mean, were identified using an HTP sequencing-based yeast replicative lifespan screen (SEBYL), with pooled cells subject to four rounds of biotin-based sorting to enrich old vs young cells, followed by sequencing to ID mutants and abundance 	Yu R, et al. (2021) PMID:33790287
SYH1	YPL105C	replicative lifespan: increased 	classical genetics	classical genetics	null Allele: syh1-Δ	S288C			McCormick MA, et al. (2015) PMID:26456335
SYM1	YLR251W	replicative lifespan: increased 	systematic mutation set 	large-scale survey	null Allele: sym1-Δ	S288C		Details: long-lived mutants, one standard deviation above the mean, were identified using an HTP sequencing-based yeast replicative lifespan screen (SEBYL), with pooled cells subject to four rounds of biotin-based sorting to enrich old vs young cells, followed by sequencing to ID mutants and abundance 	Yu R, et al. (2021) PMID:33790287
SYN8	YAL014C	replicative lifespan: increased 	systematic mutation set 	large-scale survey	null Allele: syn8-Δ	S288C		Details: long-lived mutants, one standard deviation above the mean, were identified using an HTP sequencing-based yeast replicative lifespan screen (SEBYL), with pooled cells subject to four rounds of biotin-based sorting to enrich old vs young cells, followed by sequencing to ID mutants and abundance 	Yu R, et al. (2021) PMID:33790287
SYO1	YDL063C	replicative lifespan: increased 	classical genetics	classical genetics	null Allele: syo1-Δ	S288C			McCormick MA, et al. (2015) PMID:26456335
TAF14	YPL129W	replicative lifespan: increased 	large-scale survey	large-scale survey	null Allele: taf14-Δ	S288C			Managbanag JR, et al. (2008) PMID:19030232
TAT2	YOL020W	replicative lifespan: increased 	classical genetics	classical genetics	null Allele: tat2-Δ	S288C		Details: null mutant exhibits decreased extension of replicative lifespan by ibuprofen treatment 	He C, et al. (2014) PMID:25521617
TAT2	YOL020W	replicative lifespan: increased 	classical genetics	classical genetics	null Allele: tat2-Δ	S288C			McCormick MA, et al. (2015) PMID:26456335
TAT2	YOL020W	replicative lifespan: increased 	systematic mutation set 	large-scale survey	null Allele: tat2-Δ	S288C		Details: long-lived mutants, one standard deviation above the mean, were identified using an HTP sequencing-based yeast replicative lifespan screen (SEBYL), with pooled cells subject to four rounds of biotin-based sorting to enrich old vs young cells, followed by sequencing to ID mutants and abundance 	Yu R, et al. (2021) PMID:33790287
TCD2	YKL027W	replicative lifespan: increased 	systematic mutation set 	large-scale survey	null Allele: tcd2-Δ	S288C		Details: long-lived mutants, one standard deviation above the mean, were identified using an HTP sequencing-based yeast replicative lifespan screen (SEBYL), with pooled cells subject to four rounds of biotin-based sorting to enrich old vs young cells, followed by sequencing to ID mutants and abundance 	Yu R, et al. (2021) PMID:33790287
TDH2	YJR009C	replicative lifespan: increased 	systematic mutation set 	large-scale survey	null Allele: tdh2-Δ	S288C		Details: long-lived mutants, one standard deviation above the mean, were identified using an HTP sequencing-based yeast replicative lifespan screen (SEBYL), with pooled cells subject to four rounds of biotin-based sorting to enrich old vs young cells, followed by sequencing to ID mutants and abundance 	Yu R, et al. (2021) PMID:33790287
TFB6	YOR352W	replicative lifespan: increased 	systematic mutation set 	large-scale survey	null Allele: tfb6-Δ	S288C		Details: long-lived mutants, one standard deviation above the mean, were identified using an HTP sequencing-based yeast replicative lifespan screen (SEBYL), with pooled cells subject to four rounds of biotin-based sorting to enrich old vs young cells, followed by sequencing to ID mutants and abundance 	Yu R, et al. (2021) PMID:33790287
THI21	YPL258C	replicative lifespan: increased 	systematic mutation set 	large-scale survey	null Allele: thi21-Δ	S288C		Details: long-lived mutants, one standard deviation above the mean, were identified using an HTP sequencing-based yeast replicative lifespan screen (SEBYL), with pooled cells subject to four rounds of biotin-based sorting to enrich old vs young cells, followed by sequencing to ID mutants and abundance 	Yu R, et al. (2021) PMID:33790287
THI22	YPR121W	replicative lifespan: increased 	systematic mutation set 	large-scale survey	null Allele: thi22-Δ	S288C		Details: long-lived mutants, one standard deviation above the mean, were identified using an HTP sequencing-based yeast replicative lifespan screen (SEBYL), with pooled cells subject to four rounds of biotin-based sorting to enrich old vs young cells, followed by sequencing to ID mutants and abundance 	Yu R, et al. (2021) PMID:33790287
TIF1	YKR059W	replicative lifespan: increased 	large-scale survey	large-scale survey	null Allele: tif1-Δ	S288C			Smith ED, et al. (2008) PMID:18340043
TIF1	YKR059W	replicative lifespan: increased 	large-scale survey	large-scale survey	null Allele: tif1-Δ	S288C			Kaeberlein M, et al. (2005) PMID:16293764
TIF1	YKR059W	replicative lifespan: increased 	competitive growth	large-scale survey	null Allele: tif1-Δ	S288C			Delaney JR, et al. (2011) PMID:21191185
TIF1	YKR059W	replicative lifespan: increased 	classical genetics	classical genetics	null Allele: tif1-Δ	S288C			McCormick MA, et al. (2015) PMID:26456335
TIF2	YJL138C	replicative lifespan: increased 	large-scale survey	large-scale survey	null Allele: tif2-Δ	S288C			Smith ED, et al. (2008) PMID:18340043
TIF2	YJL138C	replicative lifespan: increased 	classical genetics	classical genetics	null Allele: tif2-Δ	S288C			Delaney JR, et al. (2011) PMID:21902802
TIF2	YJL138C	replicative lifespan: increased 	competitive growth	large-scale survey	null Allele: tif2-Δ	S288C			Delaney JR, et al. (2011) PMID:21191185
TIF2	YJL138C	replicative lifespan: increased 	classical genetics	classical genetics	null Allele: tif2-Δ	S288C			McCormick MA, et al. (2015) PMID:26456335
TIF4631	YGR162W	replicative lifespan: increased 	large-scale survey	large-scale survey	null Allele: tif4631-Δ	S288C			Smith ED, et al. (2008) PMID:18340043
TIF4631	YGR162W	replicative lifespan: increased 	large-scale survey	large-scale survey	null Allele: tif4631-Δ	S288C			Managbanag JR, et al. (2008) PMID:19030232
TIF4631	YGR162W	replicative lifespan: increased 	competitive growth	large-scale survey	null Allele: tif4631-Δ	S288C			Delaney JR, et al. (2011) PMID:21191185
TIF4631	YGR162W	replicative lifespan: increased 	classical genetics	classical genetics	null Allele: tif4631-Δ	S288C			McCormick MA, et al. (2015) PMID:26456335
TIF4631	YGR162W	replicative lifespan: increased 	systematic mutation set 	large-scale survey	null Allele: tif4631-Δ	S288C		Details: long-lived mutants, one standard deviation above the mean, were identified using an HTP sequencing-based yeast replicative lifespan screen (SEBYL), with pooled cells subject to four rounds of biotin-based sorting to enrich old vs young cells, followed by sequencing to ID mutants and abundance 	Yu R, et al. (2021) PMID:33790287
TIP41	YPR040W	replicative lifespan: increased 	classical genetics	classical genetics	null Allele: tip41-Δ	S288C			McCormick MA, et al. (2015) PMID:26456335
TIP41	YPR040W	replicative lifespan: increased 	systematic mutation set 	large-scale survey	null Allele: tip41-Δ	S288C		Details: long-lived mutants, one standard deviation above the mean, were identified using an HTP sequencing-based yeast replicative lifespan screen (SEBYL), with pooled cells subject to four rounds of biotin-based sorting to enrich old vs young cells, followed by sequencing to ID mutants and abundance 	Yu R, et al. (2021) PMID:33790287
TIS11	YLR136C	replicative lifespan: increased 	large-scale survey	large-scale survey	null Allele: tis11-Δ	S288C			Smith ED, et al. (2008) PMID:18340043
TIS11	YLR136C	replicative lifespan: increased 	classical genetics	classical genetics	null Allele: tis11-Δ	S288C			Delaney JR, et al. (2011) PMID:21902802
TIS11	YLR136C	replicative lifespan: increased 	competitive growth	large-scale survey	null Allele: tis11-Δ	S288C			Delaney JR, et al. (2011) PMID:21191185
TIS11	YLR136C	replicative lifespan: increased 	classical genetics	classical genetics	null Allele: tis11-Δ	S288C			McCormick MA, et al. (2015) PMID:26456335
TIS11	YLR136C	replicative lifespan: increased 	classical genetics 	classical genetics	null Allele: tis11-Δ	S288C			Patnaik PK, et al. (2022) PMID:35858543
TIS11	YLR136C	replicative lifespan: increased 	classical genetics 	classical genetics	reduction of function Allele: tis11-C190R	S288C			Patnaik PK, et al. (2022) PMID:35858543
TLC1	YNCB0010W	replicative lifespan: increased 	classical genetics	classical genetics	dominant negative Allele: tTLC1 truncation	W303		Details: increase of 20% relative to W303 parent 	Austriaco NR and Guarente LP (1997) PMID:9275199
TMA19	YKL056C	replicative lifespan: increased 	classical genetics	classical genetics	null Allele: tma19-Δ	S288C		Details: increased lifespan of mother cells 	Rinnerthaler M, et al. (2006) PMID:16806052
TMA19	YKL056C	replicative lifespan: increased 	large-scale survey	large-scale survey	null Allele: tma19-Δ	S288C			Managbanag JR, et al. (2008) PMID:19030232
TMA19	YKL056C	replicative lifespan: increased 	classical genetics	classical genetics	null Allele: tma19-Δ	S288C			Delaney JR, et al. (2011) PMID:21902802
TMA19	YKL056C	replicative lifespan: increased 	competitive growth	large-scale survey	null Allele: tma19-Δ	S288C			Delaney JR, et al. (2011) PMID:21191185
TMA19	YKL056C	replicative lifespan: increased 	classical genetics	classical genetics	null Allele: tma19-Δ	S288C			McCormick MA, et al. (2015) PMID:26456335
TOM1	YDR457W	replicative lifespan: increased 	large-scale survey	large-scale survey	null Allele: tom1-Δ	S288C			Managbanag JR, et al. (2008) PMID:19030232
TOM1	YDR457W	replicative lifespan: increased 	classical genetics	classical genetics	null Allele: tom1-Δ	S288C			Kruegel U, et al. (2011) PMID:21931558
TOM1	YDR457W	replicative lifespan: increased 	classical genetics	classical genetics	null Allele: tom1-Δ	S288C			McCormick MA, et al. (2015) PMID:26456335
TOM1	YDR457W	replicative lifespan: increased 	systematic mutation set 	large-scale survey	null Allele: tom1-Δ	S288C		Details: long-lived mutants, one standard deviation above the mean, were identified using an HTP sequencing-based yeast replicative lifespan screen (SEBYL), with pooled cells subject to four rounds of biotin-based sorting to enrich old vs young cells, followed by sequencing to ID mutants and abundance 	Yu R, et al. (2021) PMID:33790287
TOM70	YNL121C	replicative lifespan: increased 	classical genetics	classical genetics	null Allele: tom70-Δ	S288C		Treatment: caloric restriction by glucose reduction, 0.05% Details: dietary restriction-induced lifespan extension is increased relative to wt 	Schleit J, et al. (2013) PMID:23837470
TOM70	YNL121C	replicative lifespan: increased 	classical genetics 	classical genetics	overexpression	S288C			Liu Q, et al. (2022) PMID:35234609
TOM71	YHR117W	replicative lifespan: increased 	classical genetics	classical genetics	null Allele: tom71-Δ	S288C		Treatment: caloric restriction by glucose reduction, 0.05% Details: dietary restriction-induced lifespan extension is increased relative to wt 	Schleit J, et al. (2013) PMID:23837470
TOR1	YJR066W	replicative lifespan: increased 	large-scale survey	large-scale survey	null Allele: tor1-Δ	S288C			Smith ED, et al. (2008) PMID:18340043
TOR1	YJR066W	replicative lifespan: increased 	classical genetics	classical genetics	null Allele: tor1-Δ	S288C		Details: increases both mean and maximum lifespan by ~20%; caloric restriction fails to further increase lifespan 	Kaeberlein M, et al. (2005) PMID:16293764
TOR1	YJR066W	replicative lifespan: increased 	large-scale survey	large-scale survey	null Allele: tor1-Δ	S288C			Kaeberlein M, et al. (2005) PMID:16293764
TOR1	YJR066W	replicative lifespan: increased 	classical genetics	classical genetics	null Allele: tor1-Δ	S288C			Delaney JR, et al. (2011) PMID:21902802
TOR1	YJR066W	replicative lifespan: increased 	competitive growth	large-scale survey	null Allele: tor1-Δ	S288C			Delaney JR, et al. (2011) PMID:21191185
TOR1	YJR066W	replicative lifespan: increased 	classical genetics	classical genetics	null Allele: tor1-Δ	S288C			Kruegel U, et al. (2011) PMID:21931558
TOR1	YJR066W	replicative lifespan: increased 	systematic mutation set	large-scale survey	null Allele: tor1-Δ	S288C			Delaney JR, et al. (2013) PMID:23336757
TOR1	YJR066W	replicative lifespan: increased 	classical genetics	classical genetics	null Allele: tor1-Δ	S288C			Peng J, et al. (2015) PMID:25822194
TOR1	YJR066W	replicative lifespan: increased 	classical genetics	classical genetics	null Allele: tor1-Δ	S288C			McCormick MA, et al. (2015) PMID:26456335
TOS8	YGL096W	replicative lifespan: increased 	classical genetics	classical genetics	null Allele: tos8-Δ	S288C			McCormick MA, et al. (2015) PMID:26456335
TPO3	YPR156C	replicative lifespan: increased 	classical genetics	classical genetics	null Allele: tpo3-Δ	S288C			McCormick MA, et al. (2015) PMID:26456335
TRK1	YJL129C	replicative lifespan: increased 	classical genetics	classical genetics	null Allele: trk1-Δ	S288C			McCormick MA, et al. (2015) PMID:26456335
TRS65	YGR166W	replicative lifespan: increased 	systematic mutation set 	large-scale survey	null Allele: trs65-Δ	S288C		Details: long-lived mutants, one standard deviation above the mean, were identified using an HTP sequencing-based yeast replicative lifespan screen (SEBYL), with pooled cells subject to four rounds of biotin-based sorting to enrich old vs young cells, followed by sequencing to ID mutants and abundance 	Yu R, et al. (2021) PMID:33790287
TRS85	YDR108W	replicative lifespan: increased 	systematic mutation set 	large-scale survey	null Allele: trs85-Δ	S288C		Details: long-lived mutants, one standard deviation above the mean, were identified using an HTP sequencing-based yeast replicative lifespan screen (SEBYL), with pooled cells subject to four rounds of biotin-based sorting to enrich old vs young cells, followed by sequencing to ID mutants and abundance 	Yu R, et al. (2021) PMID:33790287
TSA1	YML028W	replicative lifespan: increased 	classical genetics	classical genetics	null Allele: tsa1-Δ	S288C		Treatment: caloric restriction by glucose reduction, 0.05% Details: dietary restriction-induced lifespan extension is increased relative to wt 	Schleit J, et al. (2013) PMID:23837470
UBP8	YMR223W	replicative lifespan: increased 	classical genetics	classical genetics	null Allele: ubp8-Δ	S288C			McCormick MA, et al. (2015) PMID:26456335
UBP8	YMR223W	replicative lifespan: increased 	systematic mutation set 	large-scale survey	null Allele: ubp8-Δ	S288C		Details: long-lived mutants, one standard deviation above the mean, were identified using an HTP sequencing-based yeast replicative lifespan screen (SEBYL), with pooled cells subject to four rounds of biotin-based sorting to enrich old vs young cells, followed by sequencing to ID mutants and abundance 	Yu R, et al. (2021) PMID:33790287
UBR2	YLR024C	replicative lifespan: increased 	classical genetics	classical genetics	null Allele: ubr2-Δ	S288C			Kruegel U, et al. (2011) PMID:21931558
UBR2	YLR024C	replicative lifespan: increased 	classical genetics	classical genetics	null Allele: ubr2-Δ	S288C			McCormick MA, et al. (2015) PMID:26456335
UBR2	YLR024C	replicative lifespan: increased 	classical genetics 	classical genetics	null Allele: ubr2-Δ	S288C		Details: lifespan extended by 23%; further extended by mycophenolic acid that can be suppressed by guanine supplementation 	Liu P, et al. (2020) PMID:32232569
UBR2	YLR024C	replicative lifespan: increased 	systematic mutation set 	large-scale survey	null Allele: ubr2-Δ	S288C		Details: long-lived mutants, one standard deviation above the mean, were identified using an HTP sequencing-based yeast replicative lifespan screen (SEBYL), with pooled cells subject to four rounds of biotin-based sorting to enrich old vs young cells, followed by sequencing to ID mutants and abundance 	Yu R, et al. (2021) PMID:33790287
UGA1	YGR019W	replicative lifespan: increased 	classical genetics	classical genetics	null Allele: uga1-Δ	S288C			Kamei Y, et al. (2011) PMID:21371425
UGA3	YDL170W	replicative lifespan: increased 	classical genetics	classical genetics	null Allele: uga3-Δ	S288C			Yoshida R, et al. (2010) PMID:20550517
UGP1	YKL035W	replicative lifespan: increased 	classical genetics 	classical genetics	overexpression	S288C			Yu R, et al. (2021) PMID:33790287
UGX2	YDL169C	replicative lifespan: increased 	classical genetics	classical genetics	null Allele: ugx2-Δ	S288C			McCormick MA, et al. (2015) PMID:26456335
UGX2	YDL169C	replicative lifespan: increased 	systematic mutation set 	large-scale survey	null Allele: ugx2-Δ	S288C		Details: long-lived mutants, one standard deviation above the mean, were identified using an HTP sequencing-based yeast replicative lifespan screen (SEBYL), with pooled cells subject to four rounds of biotin-based sorting to enrich old vs young cells, followed by sequencing to ID mutants and abundance 	Yu R, et al. (2021) PMID:33790287
UNG1	YML021C	replicative lifespan: increased 	classical genetics	classical genetics	null Allele: ung1-Δ	S288C		Treatment: caloric restriction by glucose reduction, 0.05% Details: dietary restriction-induced lifespan extension is increased relative to wt 	Schleit J, et al. (2013) PMID:23837470
URE2	YNL229C	replicative lifespan: increased 	classical genetics	classical genetics	null Allele: ure2-Δ	S288C			Kaeberlein M, et al. (2005) PMID:16293764
URE2	YNL229C	replicative lifespan: increased 	large-scale survey	large-scale survey	null Allele: ure2-Δ	S288C			Kaeberlein M, et al. (2005) PMID:16293764
URE2	YNL229C	replicative lifespan: increased 	classical genetics	classical genetics	null Allele: ure2-Δ	S288C			Delaney JR, et al. (2011) PMID:21902802
URE2	YNL229C	replicative lifespan: increased 	competitive growth	large-scale survey	null Allele: ure2-Δ	S288C			Delaney JR, et al. (2011) PMID:21191185
URE2	YNL229C	replicative lifespan: increased 	classical genetics	classical genetics	null Allele: ure2-Δ	S288C			McCormick MA, et al. (2015) PMID:26456335
URH1	YDR400W	replicative lifespan: increased 	classical genetics	classical genetics	null Allele: urh1-Δ	S288C			Yoshida R, et al. (2010) PMID:20550517
UTH1	YKR042W	replicative lifespan: increased 	classical genetics	classical genetics	null Allele: uth1-Δ	Other			Weng Y, et al. (2010) PMID:20093034
UTH1	YKR042W	replicative lifespan: increased 	classical genetics	classical genetics	unspecified Allele: uth1-328	Other			Kennedy BK, et al. (1995) PMID:7859289
VAB2	YEL005C	replicative lifespan: increased 	systematic mutation set 	large-scale survey	null Allele: vab2-Δ	S288C		Details: long-lived mutants, one standard deviation above the mean, were identified using an HTP sequencing-based yeast replicative lifespan screen (SEBYL), with pooled cells subject to four rounds of biotin-based sorting to enrich old vs young cells, followed by sequencing to ID mutants and abundance 	Yu R, et al. (2021) PMID:33790287
VAB2	YEL005C	replicative lifespan: increased 	classical genetics 	classical genetics	null Allele: vab2-Δ	S288C		Details: long-lived mutant assayed using a microfluidic chip-based lifespan measurement with automatic imaging to validate SEBYL 	Yu R, et al. (2021) PMID:33790287
VBA5	YKR105C	replicative lifespan: increased 	large-scale survey	large-scale survey	null Allele: vba5-Δ	S288C			Kaeberlein M, et al. (2005) PMID:16293764
VFA1	YER128W	replicative lifespan: increased 	classical genetics	classical genetics	null Allele: vfa1-Δ	S288C			McCormick MA, et al. (2015) PMID:26456335
VHS1	YDR247W	replicative lifespan: increased 	large-scale survey	large-scale survey	null Allele: vhs1-Δ	S288C			Kaeberlein M, et al. (2005) PMID:16293764
VID30	YGL227W	replicative lifespan: increased 	systematic mutation set 	large-scale survey	null Allele: vid30-Δ	S288C		Details: long-lived mutants, one standard deviation above the mean, were identified using an HTP sequencing-based yeast replicative lifespan screen (SEBYL), with pooled cells subject to four rounds of biotin-based sorting to enrich old vs young cells, followed by sequencing to ID mutants and abundance 	Yu R, et al. (2021) PMID:33790287
VMA1	YDL185W	replicative lifespan: increased 	homozygous diploid	classical genetics	overexpression	S288C		Details: increase in both median and maximal replicative lifespan 	Hughes AL and Gottschling DE (2012) PMID:23172144
VPH2	YKL119C	replicative lifespan: increased 	homozygous diploid	classical genetics	overexpression	S288C		Details: increase in both median and maximal replicative lifespan 	Hughes AL and Gottschling DE (2012) PMID:23172144
VPS24	YKL041W	replicative lifespan: increased 	classical genetics	classical genetics	null Allele: vps24-Δ	S288C			McCormick MA, et al. (2015) PMID:26456335
VPS3	YDR495C	replicative lifespan: increased 	classical genetics	classical genetics	null Allele: vps3-Δ	S288C		Treatment: caloric restriction by glucose reduction, 0.05% Details: dietary restriction-induced lifespan extension is increased relative to wt 	Schleit J, et al. (2013) PMID:23837470
VPS60	YDR486C	replicative lifespan: increased 	classical genetics	classical genetics	null Allele: vps60-Δ	S288C			McCormick MA, et al. (2015) PMID:26456335
VPS74	YDR372C	replicative lifespan: increased 	systematic mutation set 	large-scale survey	null Allele: vps74-Δ	S288C		Details: long-lived mutants, one standard deviation above the mean, were identified using an HTP sequencing-based yeast replicative lifespan screen (SEBYL), with pooled cells subject to four rounds of biotin-based sorting to enrich old vs young cells, followed by sequencing to ID mutants and abundance 	Yu R, et al. (2021) PMID:33790287
VRL1	YML002W	replicative lifespan: increased 	systematic mutation set 	large-scale survey	null Allele: yml002w-Δ	S288C		Details: long-lived mutants, one standard deviation above the mean, were identified using an HTP sequencing-based yeast replicative lifespan screen (SEBYL), with pooled cells subject to four rounds of biotin-based sorting to enrich old vs young cells, followed by sequencing to ID mutants and abundance 	Yu R, et al. (2021) PMID:33790287
VTC5	YDR089W	replicative lifespan: increased 	systematic mutation set 	large-scale survey	null Allele: vtc5-Δ	S288C		Details: long-lived mutants, one standard deviation above the mean, were identified using an HTP sequencing-based yeast replicative lifespan screen (SEBYL), with pooled cells subject to four rounds of biotin-based sorting to enrich old vs young cells, followed by sequencing to ID mutants and abundance 	Yu R, et al. (2021) PMID:33790287
WHI5	YOR083W	replicative lifespan: increased 	classical genetics	classical genetics	null Allele: whi5-Δ	S288C		Treatment: caloric restriction by glucose reduction, 0.05% Details: dietary restriction-induced lifespan extension is increased relative to wt 	Schleit J, et al. (2013) PMID:23837470
YBL053W	YBL053W	replicative lifespan: increased 	classical genetics	classical genetics	null Allele: ybl053w-Δ	S288C			McCormick MA, et al. (2015) PMID:26456335
YBL053W	YBL053W	replicative lifespan: increased 	systematic mutation set 	large-scale survey	null Allele: ybl053w-Δ	S288C		Details: long-lived mutants, one standard deviation above the mean, were identified using an HTP sequencing-based yeast replicative lifespan screen (SEBYL), with pooled cells subject to four rounds of biotin-based sorting to enrich old vs young cells, followed by sequencing to ID mutants and abundance 	Yu R, et al. (2021) PMID:33790287
YBL081W	YBL081W	replicative lifespan: increased 	systematic mutation set 	large-scale survey	null Allele: ybl081w-Δ	S288C		Details: long-lived mutants, one standard deviation above the mean, were identified using an HTP sequencing-based yeast replicative lifespan screen (SEBYL), with pooled cells subject to four rounds of biotin-based sorting to enrich old vs young cells, followed by sequencing to ID mutants and abundance 	Yu R, et al. (2021) PMID:33790287
YBL083C	YBL083C	replicative lifespan: increased 	classical genetics	classical genetics	null Allele: ybl083c-Δ	S288C			McCormick MA, et al. (2015) PMID:26456335
YBR013C	YBR013C	replicative lifespan: increased 	classical genetics	classical genetics	null Allele: ybr013c-Δ	S288C			McCormick MA, et al. (2015) PMID:26456335
YBR064W	YBR064W	replicative lifespan: increased 	systematic mutation set 	large-scale survey	null Allele: ybr064w-Δ	S288C		Details: long-lived mutants, one standard deviation above the mean, were identified using an HTP sequencing-based yeast replicative lifespan screen (SEBYL), with pooled cells subject to four rounds of biotin-based sorting to enrich old vs young cells, followed by sequencing to ID mutants and abundance 	Yu R, et al. (2021) PMID:33790287
YBR064W	YBR064W	replicative lifespan: increased 	classical genetics 	classical genetics	null Allele: ybr064w-Δ	S288C		Details: long-lived mutant assayed using a microfluidic chip-based lifespan measurement with automatic imaging to validate SEBYL 	Yu R, et al. (2021) PMID:33790287
YBR197C	YBR197C	replicative lifespan: increased 	classical genetics	classical genetics	null Allele: ybr197c-Δ	S288C			McCormick MA, et al. (2015) PMID:26456335
YBR206W	YBR206W	replicative lifespan: increased 	systematic mutation set 	large-scale survey	null Allele: ybr206w-Δ	S288C		Details: long-lived mutants, one standard deviation above the mean, were identified using an HTP sequencing-based yeast replicative lifespan screen (SEBYL), with pooled cells subject to four rounds of biotin-based sorting to enrich old vs young cells, followed by sequencing to ID mutants and abundance 	Yu R, et al. (2021) PMID:33790287
YBR226C	YBR226C	replicative lifespan: increased 	classical genetics	classical genetics	null Allele: ybr226c-Δ	S288C			McCormick MA, et al. (2015) PMID:26456335
YBR238C	YBR238C	replicative lifespan: increased 	classical genetics	classical genetics	null Allele: ybr238c-Δ	S288C			Kaeberlein M, et al. (2005) PMID:16293764
YBR238C	YBR238C	replicative lifespan: increased 	large-scale survey	large-scale survey	null Allele: ybr238c-Δ	S288C			Kaeberlein M, et al. (2005) PMID:16293764
YBR238C	YBR238C	replicative lifespan: increased 	classical genetics	classical genetics	null Allele: ybr238c-Δ	S288C			Delaney JR, et al. (2011) PMID:21902802
YBR238C	YBR238C	replicative lifespan: increased 	competitive growth	large-scale survey	null Allele: ybr238c-Δ	S288C			Delaney JR, et al. (2011) PMID:21191185
YBR238C	YBR238C	replicative lifespan: increased 	classical genetics	classical genetics	null Allele: ybr238c-Δ	S288C			McCormick MA, et al. (2015) PMID:26456335
YCL006C	YCL006C	replicative lifespan: increased 	classical genetics	classical genetics	null Allele: ycl006c-Δ	S288C			McCormick MA, et al. (2015) PMID:26456335
YCL013W	YCL013W	replicative lifespan: increased 	systematic mutation set 	large-scale survey	null Allele: ycl013w-Δ	S288C		Details: long-lived mutants, one standard deviation above the mean, were identified using an HTP sequencing-based yeast replicative lifespan screen (SEBYL), with pooled cells subject to four rounds of biotin-based sorting to enrich old vs young cells, followed by sequencing to ID mutants and abundance 	Yu R, et al. (2021) PMID:33790287
YCR001W	YCR001W	replicative lifespan: increased 	systematic mutation set 	large-scale survey	null Allele: ycr001w-Δ	S288C		Details: long-lived mutants, one standard deviation above the mean, were identified using an HTP sequencing-based yeast replicative lifespan screen (SEBYL), with pooled cells subject to four rounds of biotin-based sorting to enrich old vs young cells, followed by sequencing to ID mutants and abundance 	Yu R, et al. (2021) PMID:33790287
YCR090C	YCR090C	replicative lifespan: increased 	systematic mutation set 	large-scale survey	null Allele: ycr090c-Δ	S288C		Details: long-lived mutants, one standard deviation above the mean, were identified using an HTP sequencing-based yeast replicative lifespan screen (SEBYL), with pooled cells subject to four rounds of biotin-based sorting to enrich old vs young cells, followed by sequencing to ID mutants and abundance 	Yu R, et al. (2021) PMID:33790287
YCR101C	YCR101C	replicative lifespan: increased 	classical genetics	classical genetics	null Allele: ycr101c-Δ	S288C			McCormick MA, et al. (2015) PMID:26456335
YDL062W	YDL062W	replicative lifespan: increased 	classical genetics	classical genetics	null Allele: ydl062w-Δ	S288C			McCormick MA, et al. (2015) PMID:26456335
YDL086W	YDL086W	replicative lifespan: increased 	systematic mutation set 	large-scale survey	null Allele: ydl086w-Δ	S288C		Details: long-lived mutants, one standard deviation above the mean, were identified using an HTP sequencing-based yeast replicative lifespan screen (SEBYL), with pooled cells subject to four rounds of biotin-based sorting to enrich old vs young cells, followed by sequencing to ID mutants and abundance 	Yu R, et al. (2021) PMID:33790287
YDL172C	YDL172C	replicative lifespan: increased 	classical genetics	classical genetics	null Allele: ydl172c-Δ	S288C			McCormick MA, et al. (2015) PMID:26456335
YDL187C	YDL187C	replicative lifespan: increased 	systematic mutation set 	large-scale survey	null Allele: ydl187c-Δ	S288C		Details: long-lived mutants, one standard deviation above the mean, were identified using an HTP sequencing-based yeast replicative lifespan screen (SEBYL), with pooled cells subject to four rounds of biotin-based sorting to enrich old vs young cells, followed by sequencing to ID mutants and abundance 	Yu R, et al. (2021) PMID:33790287
YDL199C	YDL199C	replicative lifespan: increased 	systematic mutation set 	large-scale survey	null Allele: ydl199c-Δ	S288C		Details: long-lived mutants, one standard deviation above the mean, were identified using an HTP sequencing-based yeast replicative lifespan screen (SEBYL), with pooled cells subject to four rounds of biotin-based sorting to enrich old vs young cells, followed by sequencing to ID mutants and abundance 	Yu R, et al. (2021) PMID:33790287
YDR048C	YDR048C	replicative lifespan: increased 	classical genetics	classical genetics	null Allele: ydr048c-Δ	S288C			McCormick MA, et al. (2015) PMID:26456335
YDR124W	YDR124W	replicative lifespan: increased 	large-scale survey	large-scale survey	null Allele: ydr124w-Δ	S288C			Kaeberlein M, et al. (2005) PMID:16293764
YDR133C	YDR133C	replicative lifespan: increased 	systematic mutation set 	large-scale survey	null Allele: ydr133c-Δ	S288C		Details: long-lived mutants, one standard deviation above the mean, were identified using an HTP sequencing-based yeast replicative lifespan screen (SEBYL), with pooled cells subject to four rounds of biotin-based sorting to enrich old vs young cells, followed by sequencing to ID mutants and abundance 	Yu R, et al. (2021) PMID:33790287
YDR157W	YDR157W	replicative lifespan: increased 	systematic mutation set 	large-scale survey	null Allele: ydr157w-Δ	S288C		Details: long-lived mutants, one standard deviation above the mean, were identified using an HTP sequencing-based yeast replicative lifespan screen (SEBYL), with pooled cells subject to four rounds of biotin-based sorting to enrich old vs young cells, followed by sequencing to ID mutants and abundance 	Yu R, et al. (2021) PMID:33790287
YDR248C	YDR248C	replicative lifespan: increased 	large-scale survey	large-scale survey	null Allele: ydr248c-Δ	S288C			Kaeberlein M, et al. (2005) PMID:16293764
YDR274C	YDR274C	replicative lifespan: increased 	large-scale survey	large-scale survey	null Allele: ydr274c-Δ	S288C			Kaeberlein M, et al. (2005) PMID:16293764
YDR340W	YDR340W	replicative lifespan: increased 	systematic mutation set 	large-scale survey	null Allele: ydr340w-Δ	S288C		Details: long-lived mutants, one standard deviation above the mean, were identified using an HTP sequencing-based yeast replicative lifespan screen (SEBYL), with pooled cells subject to four rounds of biotin-based sorting to enrich old vs young cells, followed by sequencing to ID mutants and abundance 	Yu R, et al. (2021) PMID:33790287
YDR391C	YDR391C	replicative lifespan: increased 	systematic mutation set 	large-scale survey	null Allele: ydr391c-Δ	S288C		Details: long-lived mutants, one standard deviation above the mean, were identified using an HTP sequencing-based yeast replicative lifespan screen (SEBYL), with pooled cells subject to four rounds of biotin-based sorting to enrich old vs young cells, followed by sequencing to ID mutants and abundance 	Yu R, et al. (2021) PMID:33790287
YEH1	YLL012W	replicative lifespan: increased 	classical genetics	classical genetics	null Allele: yeh1-Δ	S288C			McCormick MA, et al. (2015) PMID:26456335
YEL1	YBL060W	replicative lifespan: increased 	systematic mutation set 	large-scale survey	null Allele: yel1-Δ	S288C		Details: long-lived mutants, one standard deviation above the mean, were identified using an HTP sequencing-based yeast replicative lifespan screen (SEBYL), with pooled cells subject to four rounds of biotin-based sorting to enrich old vs young cells, followed by sequencing to ID mutants and abundance 	Yu R, et al. (2021) PMID:33790287
YER158C	YER158C	replicative lifespan: increased 	systematic mutation set 	large-scale survey	null Allele: yer158c-Δ	S288C		Details: long-lived mutants, one standard deviation above the mean, were identified using an HTP sequencing-based yeast replicative lifespan screen (SEBYL), with pooled cells subject to four rounds of biotin-based sorting to enrich old vs young cells, followed by sequencing to ID mutants and abundance 	Yu R, et al. (2021) PMID:33790287
YER186C	YER186C	replicative lifespan: increased 	large-scale survey	large-scale survey	null Allele: yer186c-Δ	S288C			Kaeberlein M, et al. (2005) PMID:16293764
YER187W	YER187W	replicative lifespan: increased 	large-scale survey	large-scale survey	null Allele: yer187w-Δ	S288C			Kaeberlein M, et al. (2005) PMID:16293764
YER188W	YER188W	replicative lifespan: increased 	systematic mutation set 	large-scale survey	null Allele: yer188w-Δ	S288C		Details: long-lived mutants, one standard deviation above the mean, were identified using an HTP sequencing-based yeast replicative lifespan screen (SEBYL), with pooled cells subject to four rounds of biotin-based sorting to enrich old vs young cells, followed by sequencing to ID mutants and abundance 	Yu R, et al. (2021) PMID:33790287
YGL041C	YGL041C	replicative lifespan: increased 	systematic mutation set 	large-scale survey	null Allele: ygl041c-Δ	S288C		Details: long-lived mutants, one standard deviation above the mean, were identified using an HTP sequencing-based yeast replicative lifespan screen (SEBYL), with pooled cells subject to four rounds of biotin-based sorting to enrich old vs young cells, followed by sequencing to ID mutants and abundance 	Yu R, et al. (2021) PMID:33790287
YGL042C	YGL042C	replicative lifespan: increased 	classical genetics	classical genetics	null Allele: ygl042c-Δ	S288C			McCormick MA, et al. (2015) PMID:26456335
YGL082W	YGL082W	replicative lifespan: increased 	systematic mutation set 	large-scale survey	null Allele: ygl082w-Δ	S288C		Details: long-lived mutants, one standard deviation above the mean, were identified using an HTP sequencing-based yeast replicative lifespan screen (SEBYL), with pooled cells subject to four rounds of biotin-based sorting to enrich old vs young cells, followed by sequencing to ID mutants and abundance 	Yu R, et al. (2021) PMID:33790287
YGL118C	YGL118C	replicative lifespan: increased 	systematic mutation set 	large-scale survey	null Allele: ygl118c-Δ	S288C		Details: long-lived mutants, one standard deviation above the mean, were identified using an HTP sequencing-based yeast replicative lifespan screen (SEBYL), with pooled cells subject to four rounds of biotin-based sorting to enrich old vs young cells, followed by sequencing to ID mutants and abundance 	Yu R, et al. (2021) PMID:33790287
YGL140C	YGL140C	replicative lifespan: increased 	systematic mutation set 	large-scale survey	null Allele: ygl140c-Δ	S288C		Details: long-lived mutants, one standard deviation above the mean, were identified using an HTP sequencing-based yeast replicative lifespan screen (SEBYL), with pooled cells subject to four rounds of biotin-based sorting to enrich old vs young cells, followed by sequencing to ID mutants and abundance 	Yu R, et al. (2021) PMID:33790287
YGL149W	YGL149W	replicative lifespan: increased 	systematic mutation set 	large-scale survey	null Allele: ygl149w-Δ	S288C		Details: long-lived mutants, one standard deviation above the mean, were identified using an HTP sequencing-based yeast replicative lifespan screen (SEBYL), with pooled cells subject to four rounds of biotin-based sorting to enrich old vs young cells, followed by sequencing to ID mutants and abundance 	Yu R, et al. (2021) PMID:33790287
YGL165C	YGL165C	replicative lifespan: increased 	classical genetics	classical genetics	null Allele: ygl165c-Δ	S288C			McCormick MA, et al. (2015) PMID:26456335
YGL177W	YGL177W	replicative lifespan: increased 	systematic mutation set 	large-scale survey	null Allele: ygl177w-Δ	S288C		Details: long-lived mutants, one standard deviation above the mean, were identified using an HTP sequencing-based yeast replicative lifespan screen (SEBYL), with pooled cells subject to four rounds of biotin-based sorting to enrich old vs young cells, followed by sequencing to ID mutants and abundance 	Yu R, et al. (2021) PMID:33790287
YGR051C	YGR051C	replicative lifespan: increased 	classical genetics	classical genetics	null Allele: ygr051c-Δ	S288C			McCormick MA, et al. (2015) PMID:26456335
YGR130C	YGR130C	replicative lifespan: increased 	large-scale survey	large-scale survey	null Allele: ygr130c-Δ	S288C			Smith ED, et al. (2008) PMID:18340043
YGR153W	YGR153W	replicative lifespan: increased 	systematic mutation set 	large-scale survey	null Allele: ygr153w-Δ	S288C		Details: long-lived mutants, one standard deviation above the mean, were identified using an HTP sequencing-based yeast replicative lifespan screen (SEBYL), with pooled cells subject to four rounds of biotin-based sorting to enrich old vs young cells, followed by sequencing to ID mutants and abundance 	Yu R, et al. (2021) PMID:33790287
YHK8	YHR048W	replicative lifespan: increased 	systematic mutation set 	large-scale survey	null Allele: yhk8-Δ	S288C		Details: long-lived mutants, one standard deviation above the mean, were identified using an HTP sequencing-based yeast replicative lifespan screen (SEBYL), with pooled cells subject to four rounds of biotin-based sorting to enrich old vs young cells, followed by sequencing to ID mutants and abundance 	Yu R, et al. (2021) PMID:33790287
YHR125W	YHR125W	replicative lifespan: increased 	systematic mutation set 	large-scale survey	null Allele: yhr125w-Δ	S288C		Details: long-lived mutants, one standard deviation above the mean, were identified using an HTP sequencing-based yeast replicative lifespan screen (SEBYL), with pooled cells subject to four rounds of biotin-based sorting to enrich old vs young cells, followed by sequencing to ID mutants and abundance 	Yu R, et al. (2021) PMID:33790287
YHR130C	YHR130C	replicative lifespan: increased 	large-scale survey	large-scale survey	null Allele: yhr130c-Δ	S288C			Kaeberlein M, et al. (2005) PMID:16293764
YIL032C	YIL032C	replicative lifespan: increased 	systematic mutation set 	large-scale survey	null Allele: yil032c-Δ	S288C		Details: long-lived mutants, one standard deviation above the mean, were identified using an HTP sequencing-based yeast replicative lifespan screen (SEBYL), with pooled cells subject to four rounds of biotin-based sorting to enrich old vs young cells, followed by sequencing to ID mutants and abundance 	Yu R, et al. (2021) PMID:33790287
YIL089W	YIL089W	replicative lifespan: increased 	classical genetics	classical genetics	null Allele: yil089w-Δ	S288C			McCormick MA, et al. (2015) PMID:26456335
YIP4	YGL198W	replicative lifespan: increased 	systematic mutation set 	large-scale survey	null Allele: yip4-Δ	S288C		Details: long-lived mutants, one standard deviation above the mean, were identified using an HTP sequencing-based yeast replicative lifespan screen (SEBYL), with pooled cells subject to four rounds of biotin-based sorting to enrich old vs young cells, followed by sequencing to ID mutants and abundance 	Yu R, et al. (2021) PMID:33790287
YJL022W	YJL022W	replicative lifespan: increased 	classical genetics	classical genetics	null Allele: yjl022w-Δ	S288C			McCormick MA, et al. (2015) PMID:26456335
YJR084W	YJR084W	replicative lifespan: increased 	systematic mutation set 	large-scale survey	null Allele: yjr084w-Δ	S288C		Details: long-lived mutants, one standard deviation above the mean, were identified using an HTP sequencing-based yeast replicative lifespan screen (SEBYL), with pooled cells subject to four rounds of biotin-based sorting to enrich old vs young cells, followed by sequencing to ID mutants and abundance 	Yu R, et al. (2021) PMID:33790287
YJR154W	YJR154W	replicative lifespan: increased 	systematic mutation set 	large-scale survey	null Allele: yjr154w-Δ	S288C		Details: long-lived mutants, one standard deviation above the mean, were identified using an HTP sequencing-based yeast replicative lifespan screen (SEBYL), with pooled cells subject to four rounds of biotin-based sorting to enrich old vs young cells, followed by sequencing to ID mutants and abundance 	Yu R, et al. (2021) PMID:33790287
YKE4	YIL023C	replicative lifespan: increased 	systematic mutation set 	large-scale survey	null Allele: yke4-Δ	S288C		Details: long-lived mutants, one standard deviation above the mean, were identified using an HTP sequencing-based yeast replicative lifespan screen (SEBYL), with pooled cells subject to four rounds of biotin-based sorting to enrich old vs young cells, followed by sequencing to ID mutants and abundance 	Yu R, et al. (2021) PMID:33790287
YKR073C	YKR073C	replicative lifespan: increased 	classical genetics	classical genetics	null Allele: ykr073c-Δ	S288C			McCormick MA, et al. (2015) PMID:26456335
YLH47	YPR125W	replicative lifespan: increased 	systematic mutation set 	large-scale survey	null Allele: ylh47-Δ	S288C		Details: long-lived mutants, one standard deviation above the mean, were identified using an HTP sequencing-based yeast replicative lifespan screen (SEBYL), with pooled cells subject to four rounds of biotin-based sorting to enrich old vs young cells, followed by sequencing to ID mutants and abundance 	Yu R, et al. (2021) PMID:33790287
YLL017W	YLL017W	replicative lifespan: increased 	systematic mutation set 	large-scale survey	null Allele: yll017w-Δ	S288C		Details: long-lived mutants, one standard deviation above the mean, were identified using an HTP sequencing-based yeast replicative lifespan screen (SEBYL), with pooled cells subject to four rounds of biotin-based sorting to enrich old vs young cells, followed by sequencing to ID mutants and abundance 	Yu R, et al. (2021) PMID:33790287
YLL020C	YLL020C	replicative lifespan: increased 	systematic mutation set 	large-scale survey	null Allele: yll020c-Δ	S288C		Details: long-lived mutants, one standard deviation above the mean, were identified using an HTP sequencing-based yeast replicative lifespan screen (SEBYL), with pooled cells subject to four rounds of biotin-based sorting to enrich old vs young cells, followed by sequencing to ID mutants and abundance 	Yu R, et al. (2021) PMID:33790287
YLR108C	YLR108C	replicative lifespan: increased 	systematic mutation set 	large-scale survey	null Allele: ylr108c-Δ	S288C		Details: long-lived mutants, one standard deviation above the mean, were identified using an HTP sequencing-based yeast replicative lifespan screen (SEBYL), with pooled cells subject to four rounds of biotin-based sorting to enrich old vs young cells, followed by sequencing to ID mutants and abundance 	Yu R, et al. (2021) PMID:33790287
YLR125W	YLR125W	replicative lifespan: increased 	systematic mutation set 	large-scale survey	null Allele: ylr125w-Δ	S288C		Details: long-lived mutants, one standard deviation above the mean, were identified using an HTP sequencing-based yeast replicative lifespan screen (SEBYL), with pooled cells subject to four rounds of biotin-based sorting to enrich old vs young cells, followed by sequencing to ID mutants and abundance 	Yu R, et al. (2021) PMID:33790287
YLR257W	YLR257W	replicative lifespan: increased 	systematic mutation set 	large-scale survey	null Allele: ylr257w-Δ	S288C		Details: long-lived mutants, one standard deviation above the mean, were identified using an HTP sequencing-based yeast replicative lifespan screen (SEBYL), with pooled cells subject to four rounds of biotin-based sorting to enrich old vs young cells, followed by sequencing to ID mutants and abundance 	Yu R, et al. (2021) PMID:33790287
YLR391W	YLR391W	replicative lifespan: increased 	systematic mutation set 	large-scale survey	null Allele: ylr391w-Δ	S288C		Details: long-lived mutants, one standard deviation above the mean, were identified using an HTP sequencing-based yeast replicative lifespan screen (SEBYL), with pooled cells subject to four rounds of biotin-based sorting to enrich old vs young cells, followed by sequencing to ID mutants and abundance 	Yu R, et al. (2021) PMID:33790287
YLR402W	YLR402W	replicative lifespan: increased 	systematic mutation set 	large-scale survey	null Allele: ylr402w-Δ	S288C		Details: long-lived mutants, one standard deviation above the mean, were identified using an HTP sequencing-based yeast replicative lifespan screen (SEBYL), with pooled cells subject to four rounds of biotin-based sorting to enrich old vs young cells, followed by sequencing to ID mutants and abundance 	Yu R, et al. (2021) PMID:33790287
YLR407W	YLR407W	replicative lifespan: increased 	systematic mutation set 	large-scale survey	null Allele: ylr407w-Δ	S288C		Details: long-lived mutants, one standard deviation above the mean, were identified using an HTP sequencing-based yeast replicative lifespan screen (SEBYL), with pooled cells subject to four rounds of biotin-based sorting to enrich old vs young cells, followed by sequencing to ID mutants and abundance 	Yu R, et al. (2021) PMID:33790287
YML009C-A	YML009C-A	replicative lifespan: increased 	systematic mutation set 	large-scale survey	null Allele: yml009c-a-Δ	S288C		Details: long-lived mutants, one standard deviation above the mean, were identified using an HTP sequencing-based yeast replicative lifespan screen (SEBYL), with pooled cells subject to four rounds of biotin-based sorting to enrich old vs young cells, followed by sequencing to ID mutants and abundance 	Yu R, et al. (2021) PMID:33790287
YML009W-B	YML009W-B	replicative lifespan: increased 	systematic mutation set 	large-scale survey	null Allele: yml009w-b-Δ	S288C		Details: long-lived mutants, one standard deviation above the mean, were identified using an HTP sequencing-based yeast replicative lifespan screen (SEBYL), with pooled cells subject to four rounds of biotin-based sorting to enrich old vs young cells, followed by sequencing to ID mutants and abundance 	Yu R, et al. (2021) PMID:33790287
YML018C	YML018C	replicative lifespan: increased 	systematic mutation set 	large-scale survey	null Allele: yml018c-Δ	S288C		Details: long-lived mutants, one standard deviation above the mean, were identified using an HTP sequencing-based yeast replicative lifespan screen (SEBYL), with pooled cells subject to four rounds of biotin-based sorting to enrich old vs young cells, followed by sequencing to ID mutants and abundance 	Yu R, et al. (2021) PMID:33790287
YML033W	YML033W	replicative lifespan: increased 	systematic mutation set 	large-scale survey	null Allele: yml033w-Δ	S288C		Details: long-lived mutants, one standard deviation above the mean, were identified using an HTP sequencing-based yeast replicative lifespan screen (SEBYL), with pooled cells subject to four rounds of biotin-based sorting to enrich old vs young cells, followed by sequencing to ID mutants and abundance 	Yu R, et al. (2021) PMID:33790287
YML116W-A	YML116W-A	replicative lifespan: increased 	systematic mutation set 	large-scale survey	null Allele: yml116w-a-Δ	S288C		Details: long-lived mutants, one standard deviation above the mean, were identified using an HTP sequencing-based yeast replicative lifespan screen (SEBYL), with pooled cells subject to four rounds of biotin-based sorting to enrich old vs young cells, followed by sequencing to ID mutants and abundance 	Yu R, et al. (2021) PMID:33790287
YMR155W	YMR155W	replicative lifespan: increased 	systematic mutation set 	large-scale survey	null Allele: ymr155w-Δ	S288C		Details: long-lived mutants, one standard deviation above the mean, were identified using an HTP sequencing-based yeast replicative lifespan screen (SEBYL), with pooled cells subject to four rounds of biotin-based sorting to enrich old vs young cells, followed by sequencing to ID mutants and abundance 	Yu R, et al. (2021) PMID:33790287
YND1	YER005W	replicative lifespan: increased 	systematic mutation set 	large-scale survey	null Allele: ynd1-Δ	S288C		Details: long-lived mutants, one standard deviation above the mean, were identified using an HTP sequencing-based yeast replicative lifespan screen (SEBYL), with pooled cells subject to four rounds of biotin-based sorting to enrich old vs young cells, followed by sequencing to ID mutants and abundance 	Yu R, et al. (2021) PMID:33790287
YNG1	YOR064C	replicative lifespan: increased 	classical genetics	classical genetics	null Allele: yng1-Δ	S288C			McCormick MA, et al. (2015) PMID:26456335
YNL011C	YNL011C	replicative lifespan: increased 	systematic mutation set 	large-scale survey	null Allele: ynl011c-Δ	S288C		Details: long-lived mutants, one standard deviation above the mean, were identified using an HTP sequencing-based yeast replicative lifespan screen (SEBYL), with pooled cells subject to four rounds of biotin-based sorting to enrich old vs young cells, followed by sequencing to ID mutants and abundance 	Yu R, et al. (2021) PMID:33790287
YNL034W	YNL034W	replicative lifespan: increased 	classical genetics	classical genetics	null Allele: ynl034w-Δ	S288C			McCormick MA, et al. (2015) PMID:26456335
YNL193W	YNL193W	replicative lifespan: increased 	systematic mutation set 	large-scale survey	null Allele: ynl193w-Δ	S288C		Details: long-lived mutants, one standard deviation above the mean, were identified using an HTP sequencing-based yeast replicative lifespan screen (SEBYL), with pooled cells subject to four rounds of biotin-based sorting to enrich old vs young cells, followed by sequencing to ID mutants and abundance 	Yu R, et al. (2021) PMID:33790287
YOL014W	YOL014W	replicative lifespan: increased 	classical genetics	classical genetics	null Allele: yol014w-Δ	S288C			McCormick MA, et al. (2015) PMID:26456335
YOL035C	YOL035C	replicative lifespan: increased 	systematic mutation set 	large-scale survey	null Allele: yol035c-Δ	S288C		Details: long-lived mutants, one standard deviation above the mean, were identified using an HTP sequencing-based yeast replicative lifespan screen (SEBYL), with pooled cells subject to four rounds of biotin-based sorting to enrich old vs young cells, followed by sequencing to ID mutants and abundance 	Yu R, et al. (2021) PMID:33790287
YOL050C	YOL050C	replicative lifespan: increased 	systematic mutation set 	large-scale survey	null Allele: yol050c-Δ	S288C		Details: long-lived mutants, one standard deviation above the mean, were identified using an HTP sequencing-based yeast replicative lifespan screen (SEBYL), with pooled cells subject to four rounds of biotin-based sorting to enrich old vs young cells, followed by sequencing to ID mutants and abundance 	Yu R, et al. (2021) PMID:33790287
YOL079W	YOL079W	replicative lifespan: increased 	systematic mutation set 	large-scale survey	null Allele: yol079w-Δ	S288C		Details: long-lived mutants, one standard deviation above the mean, were identified using an HTP sequencing-based yeast replicative lifespan screen (SEBYL), with pooled cells subject to four rounds of biotin-based sorting to enrich old vs young cells, followed by sequencing to ID mutants and abundance 	Yu R, et al. (2021) PMID:33790287
YOR008C-A	YOR008C-A	replicative lifespan: increased 	systematic mutation set 	large-scale survey	null Allele: yor008c-a-Δ	S288C		Details: long-lived mutants, one standard deviation above the mean, were identified using an HTP sequencing-based yeast replicative lifespan screen (SEBYL), with pooled cells subject to four rounds of biotin-based sorting to enrich old vs young cells, followed by sequencing to ID mutants and abundance 	Yu R, et al. (2021) PMID:33790287
YOR088W	YOR088W	replicative lifespan: increased 	systematic mutation set 	large-scale survey	null Allele: yor088w-Δ	S288C		Details: long-lived mutants, one standard deviation above the mean, were identified using an HTP sequencing-based yeast replicative lifespan screen (SEBYL), with pooled cells subject to four rounds of biotin-based sorting to enrich old vs young cells, followed by sequencing to ID mutants and abundance 	Yu R, et al. (2021) PMID:33790287
YOR105W	YOR105W	replicative lifespan: increased 	systematic mutation set 	large-scale survey	null Allele: yor105w-Δ	S288C		Details: long-lived mutants, one standard deviation above the mean, were identified using an HTP sequencing-based yeast replicative lifespan screen (SEBYL), with pooled cells subject to four rounds of biotin-based sorting to enrich old vs young cells, followed by sequencing to ID mutants and abundance 	Yu R, et al. (2021) PMID:33790287
YOR111W	YOR111W	replicative lifespan: increased 	classical genetics	classical genetics	null Allele: yor111w-Δ	S288C			McCormick MA, et al. (2015) PMID:26456335
YOR139C	YOR139C	replicative lifespan: increased 	systematic mutation set 	large-scale survey	null Allele: yor139c-Δ	S288C		Details: long-lived mutants, one standard deviation above the mean, were identified using an HTP sequencing-based yeast replicative lifespan screen (SEBYL), with pooled cells subject to four rounds of biotin-based sorting to enrich old vs young cells, followed by sequencing to ID mutants and abundance 	Yu R, et al. (2021) PMID:33790287
YOR263C	YOR263C	replicative lifespan: increased 	systematic mutation set 	large-scale survey	null Allele: yor263c-Δ	S288C		Details: long-lived mutants, one standard deviation above the mean, were identified using an HTP sequencing-based yeast replicative lifespan screen (SEBYL), with pooled cells subject to four rounds of biotin-based sorting to enrich old vs young cells, followed by sequencing to ID mutants and abundance 	Yu R, et al. (2021) PMID:33790287
YOR366W	YOR366W	replicative lifespan: increased 	systematic mutation set 	large-scale survey	null Allele: yor366w-Δ	S288C		Details: long-lived mutants, one standard deviation above the mean, were identified using an HTP sequencing-based yeast replicative lifespan screen (SEBYL), with pooled cells subject to four rounds of biotin-based sorting to enrich old vs young cells, followed by sequencing to ID mutants and abundance 	Yu R, et al. (2021) PMID:33790287
YPK3	YBR028C	replicative lifespan: increased 	systematic mutation set 	large-scale survey	null Allele: ypk3-Δ	S288C		Details: long-lived mutants, one standard deviation above the mean, were identified using an HTP sequencing-based yeast replicative lifespan screen (SEBYL), with pooled cells subject to four rounds of biotin-based sorting to enrich old vs young cells, followed by sequencing to ID mutants and abundance 	Yu R, et al. (2021) PMID:33790287
YPR039W	YPR039W	replicative lifespan: increased 	systematic mutation set 	large-scale survey	null Allele: ypr039w-Δ	S288C		Details: long-lived mutants, one standard deviation above the mean, were identified using an HTP sequencing-based yeast replicative lifespan screen (SEBYL), with pooled cells subject to four rounds of biotin-based sorting to enrich old vs young cells, followed by sequencing to ID mutants and abundance 	Yu R, et al. (2021) PMID:33790287
YPR064W	YPR064W	replicative lifespan: increased 	classical genetics	classical genetics	null Allele: ypr064w-Δ	S288C			McCormick MA, et al. (2015) PMID:26456335
YPT52	YKR014C	replicative lifespan: increased 	classical genetics	classical genetics	null Allele: ypt52-Δ	S288C			McCormick MA, et al. (2015) PMID:26456335
YPT6	YLR262C	replicative lifespan: increased 	large-scale survey	large-scale survey	null Allele: ypt6-Δ	S288C			Smith ED, et al. (2008) PMID:18340043
YPT6	YLR262C	replicative lifespan: increased 	competitive growth	large-scale survey	null Allele: ypt6-Δ	S288C			Delaney JR, et al. (2011) PMID:21191185
YPT6	YLR262C	replicative lifespan: increased 	classical genetics	classical genetics	null Allele: ypt6-Δ	S288C			McCormick MA, et al. (2015) PMID:26456335
YRO2	YBR054W	replicative lifespan: increased 	classical genetics	classical genetics	null Allele: yro2-Δ	S288C			McCormick MA, et al. (2015) PMID:26456335
YRO2	YBR054W	replicative lifespan: increased 	systematic mutation set 	large-scale survey	null Allele: yro2-Δ	S288C		Details: long-lived mutants, one standard deviation above the mean, were identified using an HTP sequencing-based yeast replicative lifespan screen (SEBYL), with pooled cells subject to four rounds of biotin-based sorting to enrich old vs young cells, followed by sequencing to ID mutants and abundance 	Yu R, et al. (2021) PMID:33790287
YSC84	YHR016C	replicative lifespan: increased 	large-scale survey	large-scale survey	null Allele: ysc84-Δ	S288C			Managbanag JR, et al. (2008) PMID:19030232
YUR1	YJL139C	replicative lifespan: increased 	systematic mutation set 	large-scale survey	null Allele: yur1-Δ	S288C		Details: long-lived mutants, one standard deviation above the mean, were identified using an HTP sequencing-based yeast replicative lifespan screen (SEBYL), with pooled cells subject to four rounds of biotin-based sorting to enrich old vs young cells, followed by sequencing to ID mutants and abundance 	Yu R, et al. (2021) PMID:33790287
YUR1	YJL139C	replicative lifespan: increased 	classical genetics	classical genetics	null Allele: yur1-Δ	S288C			McCormick MA, et al. (2015) PMID:26456335
YVH1	YIR026C	replicative lifespan: increased 	classical genetics	classical genetics	null Allele: yvh1-Δ	S288C			McCormick MA, et al. (2015) PMID:26456335
ZDS1	YMR273C	replicative lifespan: increased 	classical genetics	classical genetics	null Allele: zds1-Δ	W303		Details: 37% increase in mean life span 	Roy N and Runge KW (2000) PMID:10662670
ZEO1	YOL109W	replicative lifespan: increased 	systematic mutation set 	large-scale survey	null Allele: zeo1-Δ	S288C		Details: long-lived mutants, one standard deviation above the mean, were identified using an HTP sequencing-based yeast replicative lifespan screen (SEBYL), with pooled cells subject to four rounds of biotin-based sorting to enrich old vs young cells, followed by sequencing to ID mutants and abundance 	Yu R, et al. (2021) PMID:33790287
ZNG1	YNR029C	replicative lifespan: increased 	systematic mutation set 	large-scale survey	null Allele: ynr029c-Δ	S288C		Details: long-lived mutants, one standard deviation above the mean, were identified using an HTP sequencing-based yeast replicative lifespan screen (SEBYL), with pooled cells subject to four rounds of biotin-based sorting to enrich old vs young cells, followed by sequencing to ID mutants and abundance 	Yu R, et al. (2021) PMID:33790287
