## Additional File 1 for "Uncharacterized yeast gene *YBR238C,* an effector of TORC1 signaling in a mitochondrial feedback loop, accelerates cellular aging via *HAP4*- and *RMD9*-dependent mechanisms": ReadMe - genome-wide survey for AAGs.pdf

### Genome-wide survey of aging-associated genes (AAGs) in Saccharomyces cerevisiae

The zip-package contains the following files:

- The original downloads from the database SGD
  - chronological_lifespan_decreased_annotations.txt
  - chronological_lifespan_increased_annotations.txt
  - replicative_lifespan_decreased_annotations.txt
  - replicative_lifespan_increased_annotations.txt
- The original download from the database GenAge
  - genage_models_export.tsv
- The outcome after processing the previous 5 files (‘gene-list-CLS+RLS.txt’) with all genes in yeast affecting CLS and/or RLS as reported in SGD and/or GenAge
- The breakdown of the latter file into 15 gene lists with regard to attributes “CLS/RLS increase/decrease” that can be used as input into DAVID (https://david.ncifcrf.gov/home.jsp)
- The list of 944 yeast genes reported in SGD as coding for a protein of unknown function (as of 8$^{th}$ November 2022)
- The list of functionally un- or severely under-characterized yeast genes among the AAG within each of the 15 categories (‘2022-11-18-unknown-func-final’)
- This ReadMe file

**Comments**

**1) Processing of lists of genes affecting life span**

Literature instances about aging with genes of Saccharomyces cerevisiae were downloaded from public databases on the 8th of November 2022.

From SGD (https://www.yeastgenome.org/), four files with such instances about the phenotype of deletion mutants or large-scale surveys were downloaded separately for CLS/RLS and increase/decrease. We found 725 entries (647 genes) for "CLS increased", 941 entries (658 genes) for "RLS increased", 1293 entries (apparently, about 1101 genes – see below) for "CLS decreased" and 546 entries (463 genes) for "RLS decreased" – in total, 3505 instances.

Unique gene names were extracted from these files with a PERL program and 4 files with reduced number of entries (1 gene name occurs only once as a single entry) were created for the categories CLS increased (647 genes), RLS increased (658 genes), CLS decreased (1099 genes), and RLS decreased (463). To note, the original file “CLS decreased” contains an erroneous carriage return in the line for HFP28 making its mutation W303 appear as gene name. Therefore, SGD actually reports the total number of 1100 genes in this file.

In the case of GenAge, all the information can only be downloaded in one file. In total, 1185 instances of gene-literature were found in this database (https://genomics.senescence.info/genes/search.php?organism=Saccharomyces+cerevisiae&show=4).
This files was screened with a PERL program for instances related to an increase/decrease of

chronological/replicative lifespan with the keys "C(c)hronological/R(r)eplicative lifespan (CLS/RLS)increased and/or decreased".

Several entries had to be discarded. Ten entries were without any change in any type of lifespan reported. For another 55, a lifespan increase was reported but the information whether CLS and/or RLS is affected is missing. For further 28, a lifespan decrease was reported but the information whether CLS and/or RLS is affected was not provided.

The GenAge file was split into 4 files CLS increased (121 entries), RLS increased (292 entries), CLS decreased (241 entries), and RLS decreased (438 entries) to be compatible with the SGD file format. Applying the same procedure to remove duplicate gene names, 4 files with reduced number of entries (1 gene name occurs only once as a single entry) were created: CLS increased (116 genes), RLS increased (247 genes), CLS decreased (231 genes), and RLS decreased (408).

To note, there are entries in the "RLS increased", "CLS decreased", and "RLS decreased" files where the locus ID for the given gene name is missing. All the gaps but 4 could be filled with the information from the SGD files. The 4 missing loci (all in the RLS increase file) were added manually:
DLS1 YJL065C
NAT4 YMR069W
RPO31 YOR116C
UBX2 YML013W

Finally, a table (gene-list-CLS+RLS.txt) merging the information from the 8 "reduced" files (each gene name occurs only once in 1 file) was created (2399 gene entries). It contains the gene names, their locus IDs, the information in which DB the entry can be found, and the information regarding the change of lifespan (CLS increased, RLS increased, CLS decreased, RLS decreased).

The splitting into disjunct, non-overlapping categories (CLS/RLS increased/decreased) reveals the following gene numbers (“-“ indicates that the respective property is missing):

```
CLS increased             -             -             - : 328
            - RLS increased             -             - : 349
            -             - CLS decreased             - : 615
            -             -             - RLS decreased : 318
-------------------------------------------------------------
CLS increased RLS increased             -             - : 72
CLS increased             - CLS decreased             - : 108
CLS increased             -             - RLS decreased : 72
            - RLS increased CLS decreased             - : 120
            - RLS increased             - RLS decreased : 59
            -             - CLS decreased RLS decreased : 180
-------------------------------------------------------------
CLS increased RLS increased CLS decreased             - : 29
CLS increased RLS increased             - RLS decreased : 30
CLS increased             - CLS decreased RLS decreased : 45
            - RLS increased CLS decreased RLS decreased : 55
-------------------------------------------------------------
CLS increased RLS increased CLS decreased RLS decreased : 19
```

Total gene numbers per block are: n1 – 1610; n2 – 611; n3 – 159; n4 – 19. The total number of genes in all blocks is 2399.

For completeness, we mention here that there is 8 cases of discrepancy with two genes listed for one and the same locus tag. The complete list is:

| | | | | | | | |
|---|---|---|---|---|---|---|---|
| ADE5,7 | YGL234W | - | GenAge | 1 | 0 | 0 | 0 |
| ADE57 | YGL234W | SGD | - | 0 | 0 | 1 | 0 |
| RPS24A | YER074W | SGD | GenAge | 1 | 1 | 0 | 1 |
| RPS24B | YER074W | - | GenAge | 0 | 0 | 0 | 1 |
| VTC5 | YDR089W | SGD | - | 0 | 1 | 0 | 0 |
| YDR089W | YDR089W | - | GenAge | 0 | 0 | 0 | 1 |
| RCI37 | YIL077C | SGD | - | 1 | 0 | 1 | 0 |
| YIL077C | YIL077C | - | GenAge | 0 | 0 | 0 | 1 |
| SKA1 | YKL023W | SGD | - | 0 | 0 | 1 | 0 |
| YKL023W | YKL023W | - | GenAge | 0 | 0 | 1 | 0 |
| DCK1 | YLR422W | SGD | - | 0 | 1 | 1 | 0 |
| YLR422W | YLR422W | - | GenAge | 0 | 1 | 0 | 0 |
| PEX9 | YMR018W | SGD | - | 0 | 1 | 0 | 0 |
| YMR018W | YMR018W | - | GenAge | 0 | 1 | 0 | 0 |
| TMC1 | YOR052C | SGD | - | 1 | 0 | 1 | 0 |
| YOR052C | YOR052C | - | GenAge | 0 | 0 | 0 | 1 |

Whereas the first case appears an orthographic variation of the same gene name, the lower six cases are apparently due to a divergence of gene name usage between SGD and GenAge. For the gene RPS24B in the second pair, there is a discrepancy between the locus tag names between SGD (YIL069C) and GenAge (YER074W). We did not make manual corrections for those cases in our files for automated processing but this note remains here as alert for future uses of the data.

**2) About functionally un- and under-characterized genes**

Further, these 15 lists were compared with the list of functionally un- or severely under-characterized yeast genes. For this purpose, we downloaded the list of 944 yeast genes annotated as coding for a protein of unknown function from SGD. Further, we added to this category all genes that do not have a true gene name besides the 6-letter locus ID starting with “Y”. The breakdown of the functionally un- or severely under-characterized yeast genes among the 15 categories (CLS/RLS increased/decreased) is as follows:

```
CLS increased             -             -             - : 20
            - RLS increased             -             - : 19
            -             - CLS decreased             - : 22
            -             -             - RLS decreased : 15
-------------------------------------------------------------
CLS increased RLS increased             -             - : 2
CLS increased             - CLS decreased             - : 0
CLS increased             -             - RLS decreased : 3
            - RLS increased CLS decreased             - : 3
            - RLS increased             - RLS decreased : 0
```

```
            -             - CLS decreased RLS decreased : 0
--------------------------------------------------------------
CLS increased RLS increased CLS decreased             - : 1
CLS increased RLS increased             - RLS decreased : 1
CLS increased             - CLS decreased RLS decreased : 1
            - RLS increased CLS decreased RLS decreased : 1
--------------------------------------------------------------
CLS increased RLS increased CLS decreased RLS decreased : 0
```
