## Additional File 2 for "Uncharacterized yeast gene *YBR238C,* an effector of TORC1 signaling in a mitochondrial feedback loop, accelerates cellular aging via *HAP4*- and *RMD9*-dependent mechanisms": ReadMe - rapamycin expression profile.pdf

### Genome-wide survey of aging-associated genes (AAGs) in Saccharomyces cerevisiae

The zip-package contains the following files:

- The assessment of the expression profile under rapamycin treatment 'Expression profile – rapamycin treated for 60 min.xlsx'
- The list of all AAG-related gene names found at the database/literature search with added expression profile information ('final-tab-with-RNASeq-2022-12-07.txt')
- This ReadMe file

#### Comments

##### 1) Assessment of the expression profile

The file in Excel format has several sheets with the following contents:

- Sheet1: summary of expression change under rapamycin treatment (fold change, P-value, false-discovery rate, and logarithm of counts per million reads) obtained with the DBSeq2 method
- Sheet2: the same with the EdgeR method
- Sheet3 and Sheet4: common up-regulated genes (fold change > 1, P-value < 0.05) for the two methods
- Sheet5 and Sheet6: common down-regulated genes (fold change < -1, P-value < 0.05) for the two methods

##### 2) File of all AAGs with expression information under rapamycin treatment

This file contains 2399 lines for all the unique gene names found in the database analyses with 1) gene name, 2) locus tag, 3) and 4) name of the database where the information was taken from (SGD, GenAge), 5), 6), 7) and 8) four columns with either one or zero indicating whether the annotations 'CLS increase', 'RLS increase', 'CLS decrease' or 'RLS decrease' have been found, 9) whether DBSeq2 found the gene up-regulated and it is in the list of common hits (here and below), 10) whether DBSeq2 found the gene down-regulated, 11) whether EdgeR found the gene-up-regulated, 12) whether EdgeR found the gene down-regulated, 13) whether the gene is a RUG, 14) whether the gene is a RDG.

The sign '-' signifies that the property is not found for that gene name. The notion 'na' indicates that the gene name is absence among the gene list for the expression profile.
