## Additional File 3 for "Uncharacterized yeast gene *YBR238C,* an effector of TORC1 signaling in a mitochondrial feedback loop, accelerates cellular aging via *HAP4*- and *RMD9*-dependent mechanisms": AnalysisReport.html

### Metascape Gene List Analysis Report

metascape.org^1^

#### Bar Graph Summary

Figure 1. Bar graph of enriched terms across input gene lists, colored by p-values.

Metascape only visualizes the top 20 clusters. Up to 100 enriched clusters can be viewed here.

The top-level Gene Ontology biological processes can be viewed here.

#### Gene Lists

User-provided gene identifiers are first converted into their corresponding S. cerevisiae Entrez gene IDs using the latest version of the database (last updated on 2022-04-22). If multiple identifiers correspond to the same Entrez gene ID, they will be considered as a single Entrez gene ID in downstream analyses. The gene lists are summarized in Table 1.

Table 1. Statistics of input gene lists.

| Name | Total | Unique |
| --- | --- | --- |
| MyList | 323 | 321 |

#### Gene Annotation

The following are the list of annotations retrieved from the latest version of the database (last updated on 2022-04-22) (Table 2).

Table 2. Gene annotations extracted

| Name | Type | Description |
| --- | --- | --- |
| Gene Symbol | Description | Primary HUGO gene symbol. |
| Description | Description | Short description. |
| Biological Process (GO) | Function/Location | Descriptions summarized based on gene ontology database, where up to three most informative GO terms are kept. |
| Kinase Class (UniProt) | Function/Location | Detailed kinase classes. |
| Protein Function (Protein Atlas) | Function/Location | Protein Function (Protein Atlas) |
| Subcellular Location (Protein Atlas) | Function/Location | Sucellular Location (Protein Atlas) |
| Drug (DrugBank) | Genotype/Phenotype/Disease | Drug information for the given gene as target. |
| Canonical Pathways | Ontology | Canonical Pathways |
| Hallmark Gene Sets | Ontology | Hallmark Gene Sets |

#### Pathway and Process Enrichment Analysis

For each given gene list, pathway and process enrichment analysis has been carried out with the following ontology sources: GO Biological Processes, KEGG Pathway, Reactome Gene Sets, WikiPathways and PANTHER Pathway. All genes in the genome have been used as the enrichment background. Terms with a p-value < 0.01, a minimum count of 3, and an enrichment factor > 1.5 (the enrichment factor is the ratio between the observed counts and the counts expected by chance) are collected and grouped into clusters based on their membership similarities. More specifically, p-values are calculated based on the cumulative hypergeometric distribution^2^, and q-values are calculated using the Benjamini-Hochberg procedure to account for multiple testings^3^. Kappa scores^4^ are used as the similarity metric when performing hierarchical clustering on the enriched terms, and sub-trees with a similarity of > 0.3 are considered a cluster. The most statistically significant term within a cluster is chosen to represent the cluster.

Table 3. Top 20 clusters with their representative enriched terms (one per cluster). "Count" is the number of genes in the user-provided lists with membership in the given ontology term. "%" is the percentage of all of the user-provided genes that are found in the given ontology term (only input genes with at least one ontology term annotation are included in the calculation). "Log10(P)" is the p-value in log base 10. "Log10(q)" is the multi-test adjusted p-value in log base 10.

| GO | Category | Description | Count | % | Log10(P) | Log10(q) |
| --- | --- | --- | --- | --- | --- | --- |
| GO:0046034 | GO Biological Processes | ATP metabolic process | 40 | 13.61 | -27.34 | -23.55 |
| GO:0006754 | GO Biological Processes | ATP biosynthetic process | 15 | 5.10 | -15.62 | -12.83 |
| sce01100 | KEGG Pathway | Metabolic pathways - Saccharomyces cerevisiae (budding yeast) | 86 | 29.25 | -12.72 | -10.24 |
| GO:0005975 | GO Biological Processes | carbohydrate metabolic process | 40 | 13.61 | -11.66 | -9.29 |
| sce00500 | KEGG Pathway | Starch and sucrose metabolism - Saccharomyces cerevisiae (budding yeast) | 15 | 5.10 | -9.42 | -7.20 |
| GO:0005991 | GO Biological Processes | trehalose metabolic process | 8 | 2.72 | -8.31 | -6.19 |
| WP112 | WikiPathways | Principal pathways of carbon metabolism | 18 | 6.12 | -7.45 | -5.37 |
| GO:0006793 | GO Biological Processes | phosphorus metabolic process | 61 | 20.75 | -7.33 | -5.26 |
| GO:0044281 | GO Biological Processes | small molecule metabolic process | 64 | 21.77 | -5.97 | -3.98 |
| GO:0009991 | GO Biological Processes | response to extracellular stimulus | 21 | 7.14 | -5.61 | -3.66 |
| GO:0042407 | GO Biological Processes | cristae formation | 6 | 2.04 | -5.27 | -3.38 |
| WP70 | WikiPathways | Trehalose degradation, low osmolarity | 4 | 1.36 | -4.54 | -2.71 |
| GO:0034219 | GO Biological Processes | carbohydrate transmembrane transport | 9 | 3.06 | -4.44 | -2.62 |
| GO:0005996 | GO Biological Processes | monosaccharide metabolic process | 12 | 4.08 | -4.38 | -2.57 |
| GO:0009408 | GO Biological Processes | response to heat | 11 | 3.74 | -4.20 | -2.39 |
| sce04213 | KEGG Pathway | Longevity regulating pathway - multiple species - Saccharomyces cerevisiae (budding yeast) | 9 | 3.06 | -4.13 | -2.33 |
| GO:0010737 | GO Biological Processes | protein kinase A signaling | 4 | 1.36 | -4.08 | -2.30 |
| GO:0006123 | GO Biological Processes | mitochondrial electron transport, cytochrome c to oxygen | 6 | 2.04 | -3.95 | -2.18 |
| GO:0009269 | GO Biological Processes | response to desiccation | 3 | 1.02 | -3.92 | -2.17 |
| GO:0006109 | GO Biological Processes | regulation of carbohydrate metabolic process | 9 | 3.06 | -3.77 | -2.03 |

To further capture the relationships between the terms, a subset of enriched terms have been selected and rendered as a network plot, where terms with a similarity > 0.3 are connected by edges. We select the terms with the best p-values from each of the 20 clusters, with the constraint that there are no more than 15 terms per cluster and no more than 250 terms in total. The network is visualized using Cytoscape^5^, where each node represents an enriched term and is colored first by its cluster ID (Figure 2.a) and then by its p-value (Figure 2.b). These networks can be interactively viewed in Cytoscape through the .cys files (contained in the Zip package, which also contains a publication-quality version as a PDF) or within a browser by clicking on the web icon. For clarity, term labels are only shown for one term per cluster, so it is recommended to use Cytoscape or a browser to visualize the network in order to inspect all node labels. We can also export the network into a PDF file within Cytoscape, and then edit the labels using Adobe Illustrator for publication purposes. To switch off all labels, delete the "Label" mapping under the "Style" tab within Cytoscape, and then export the network view.

Figure 2. Network of enriched terms: (a) colored by cluster ID, where nodes that share the same cluster ID are typically close to each other; (b) colored by p-value, where terms containing more genes tend to have a more significant p-value.

#### Protein-protein Interaction Enrichment Analysis

For each given gene list, protein-protein interaction enrichment analysis has been carried out with the following databases: STRING^6^, BioGrid^7^, OmniPath^8^, InWeb\_IM^9^.Only physical interactions in STRING (physical score > 0.132) and BioGrid are used (details). The resultant network contains the subset of proteins that form physical interactions with at least one other member in the list. If the network contains between 3 and 500 proteins, the Molecular Complex Detection (MCODE) algorithm^10^ has been applied to identify densely connected network components. The MCODE networks identified for individual gene lists have been gathered and are shown in Figure 3.

Pathway and process enrichment analysis has been applied to each MCODE component independently, and the three best-scoring terms by p-value have been retained as the functional description of the corresponding components, shown in the tables underneath corresponding network plots within Figure 3.

Figure 3. Protein-protein interaction network and MCODE components identified in the gene lists.

| GO | Description | Log10(P) |
| --- | --- | --- |
| GO:0046034 | ATP metabolic process | -29.7 |
| GO:0006091 | generation of precursor metabolites and energy | -29.2 |
| sce00190 | Oxidative phosphorylation - Saccharomyces cerevisiae (budding yeast) | -27.6 |

| Color | MCODE | GO | Description | Log10(P) |
| --- | --- | --- | --- | --- |
|  | MCODE\_1 | sce00190 | Oxidative phosphorylation - Saccharomyces cerevisiae (budding yeast) | -56.9 |
|  | MCODE\_1 | GO:0046034 | ATP metabolic process | -50.4 |
|  | MCODE\_1 | GO:0022904 | respiratory electron transport chain | -31.6 |
|  | MCODE\_2 | sce00500 | Starch and sucrose metabolism - Saccharomyces cerevisiae (budding yeast) | -17.5 |
|  | MCODE\_2 | GO:0005991 | trehalose metabolic process | -14.1 |
|  | MCODE\_2 | GO:0044262 | cellular carbohydrate metabolic process | -12.5 |
|  | MCODE\_3 | GO:0009060 | aerobic respiration | -11.6 |
|  | MCODE\_3 | GO:0045333 | cellular respiration | -11.4 |
|  | MCODE\_3 | R-SCE-1428517 | The citric acid (TCA) cycle and respiratory electron transport | -11.0 |
|  | MCODE\_4 | GO:0006357 | regulation of transcription by RNA polymerase II | -2.3 |
|  | MCODE\_4 | GO:0006355 | regulation of transcription, DNA-templated | -2.1 |
|  | MCODE\_4 | GO:1903506 | regulation of nucleic acid-templated transcription | -2.1 |
|  | MCODE\_5 | sce04213 | Longevity regulating pathway - multiple species - Saccharomyces cerevisiae (budding yeast) | -6.0 |
|  | MCODE\_5 | sce04138 | Autophagy - yeast - Saccharomyces cerevisiae (budding yeast) | -4.9 |
|  | MCODE\_5 | sce04113 | Meiosis - yeast - Saccharomyces cerevisiae (budding yeast) | -4.4 |
|  | MCODE\_7 | GO:0006468 | protein phosphorylation | -4.2 |
|  | MCODE\_7 | GO:0016310 | phosphorylation | -3.6 |
|  | MCODE\_7 | GO:0007154 | cell communication | -2.8 |
|  | MCODE\_8 | sce00630 | Glyoxylate and dicarboxylate metabolism - Saccharomyces cerevisiae (budding yeast) | -6.4 |
|  | MCODE\_8 | sce01110 | Biosynthesis of secondary metabolites - Saccharomyces cerevisiae (budding yeast) | -4.9 |
|  | MCODE\_8 | sce01100 | Metabolic pathways - Saccharomyces cerevisiae (budding yeast) | -3.5 |
|  | MCODE\_9 | R-SCE-170822 | Regulation of Glucokinase by Glucokinase Regulatory Protein | -9.5 |
|  | MCODE\_9 | GO:0033500 | carbohydrate homeostasis | -8.0 |
|  | MCODE\_9 | GO:0001678 | cellular glucose homeostasis | -8.0 |
