## Additional File 3 for "Uncharacterized yeast gene *YBR238C,* an effector of TORC1 signaling in a mitochondrial feedback loop, accelerates cellular aging via *HAP4*- and *RMD9*-dependent mechanisms": AnalysisReport.pptx

### Slide 1
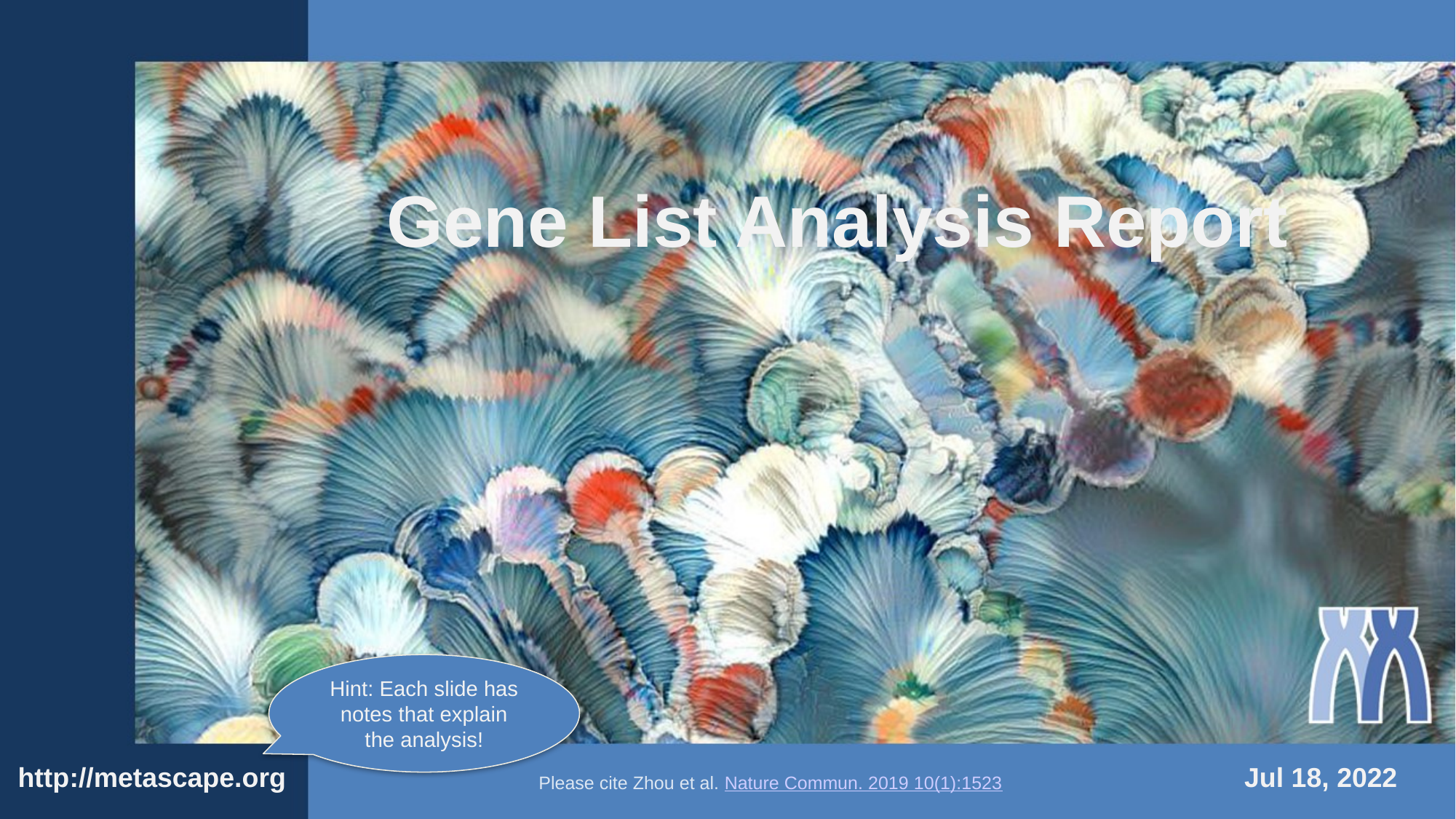

Gene List Analysis Report
Hint: Each slide has notes that explain the analysis!
http://metascape.org
Jul 18, 2022
Please cite Zhou et al. Nature Commun. 2019 10(1):1523

### Slide 2
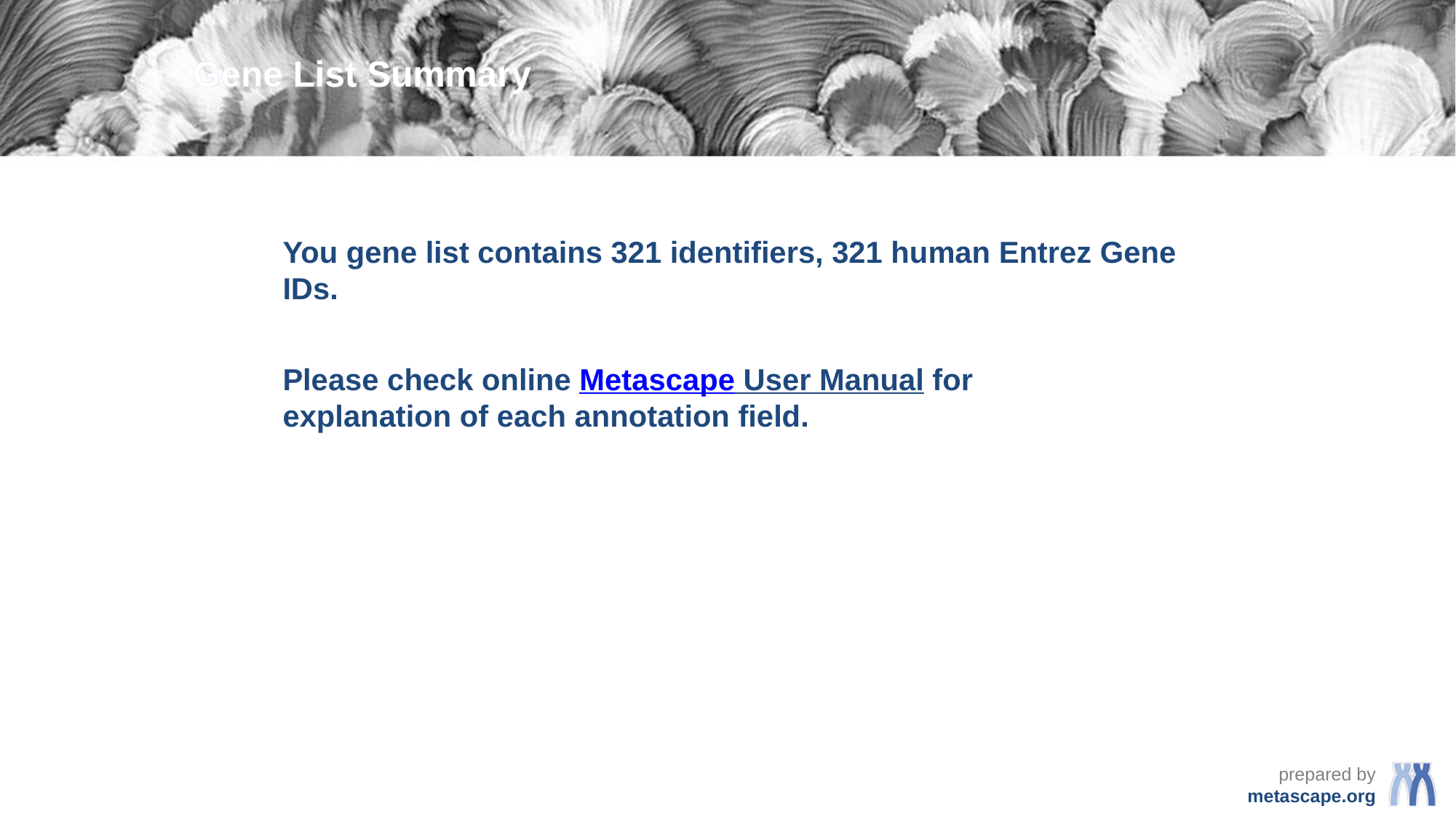

Gene List Summary
You gene list contains 321 identifiers, 321 human Entrez Gene IDs.
Please check online Metascape User Manual for explanation of each annotation field.

### Slide 3
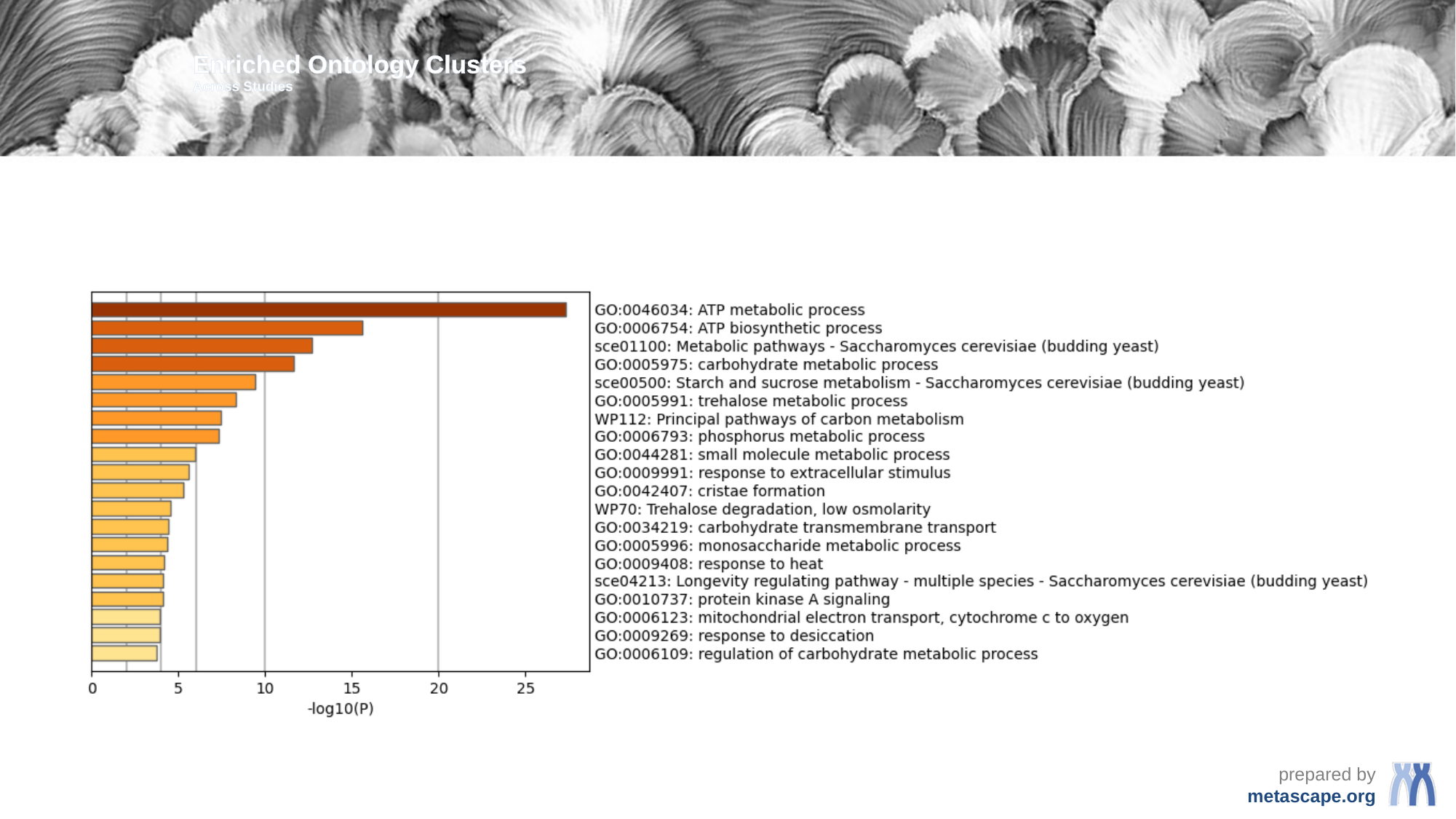

Enriched Ontology Clusters
Across Studies

### Slide 4
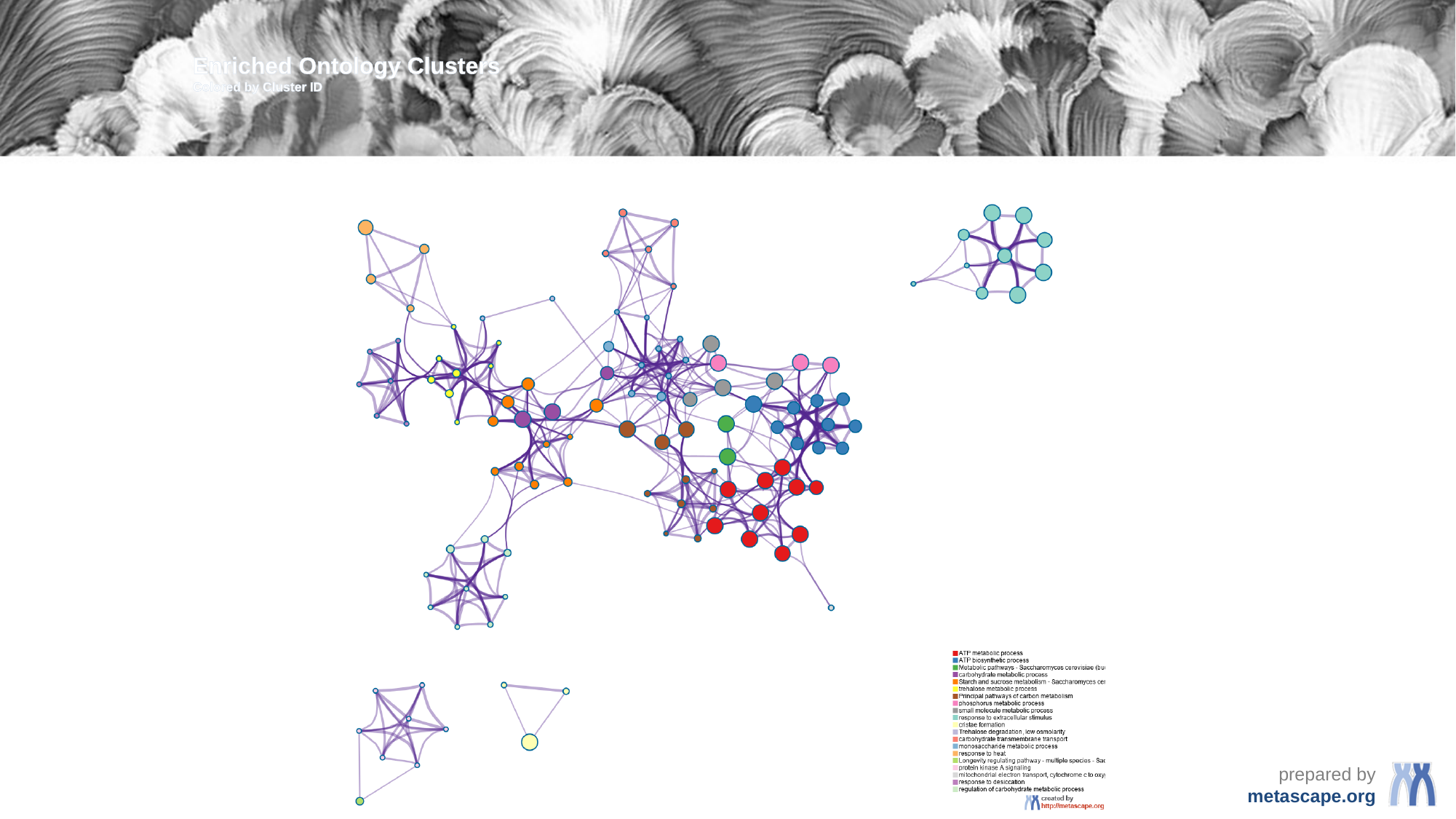

Enriched Ontology Clusters
Colored by Cluster ID

### Slide 5
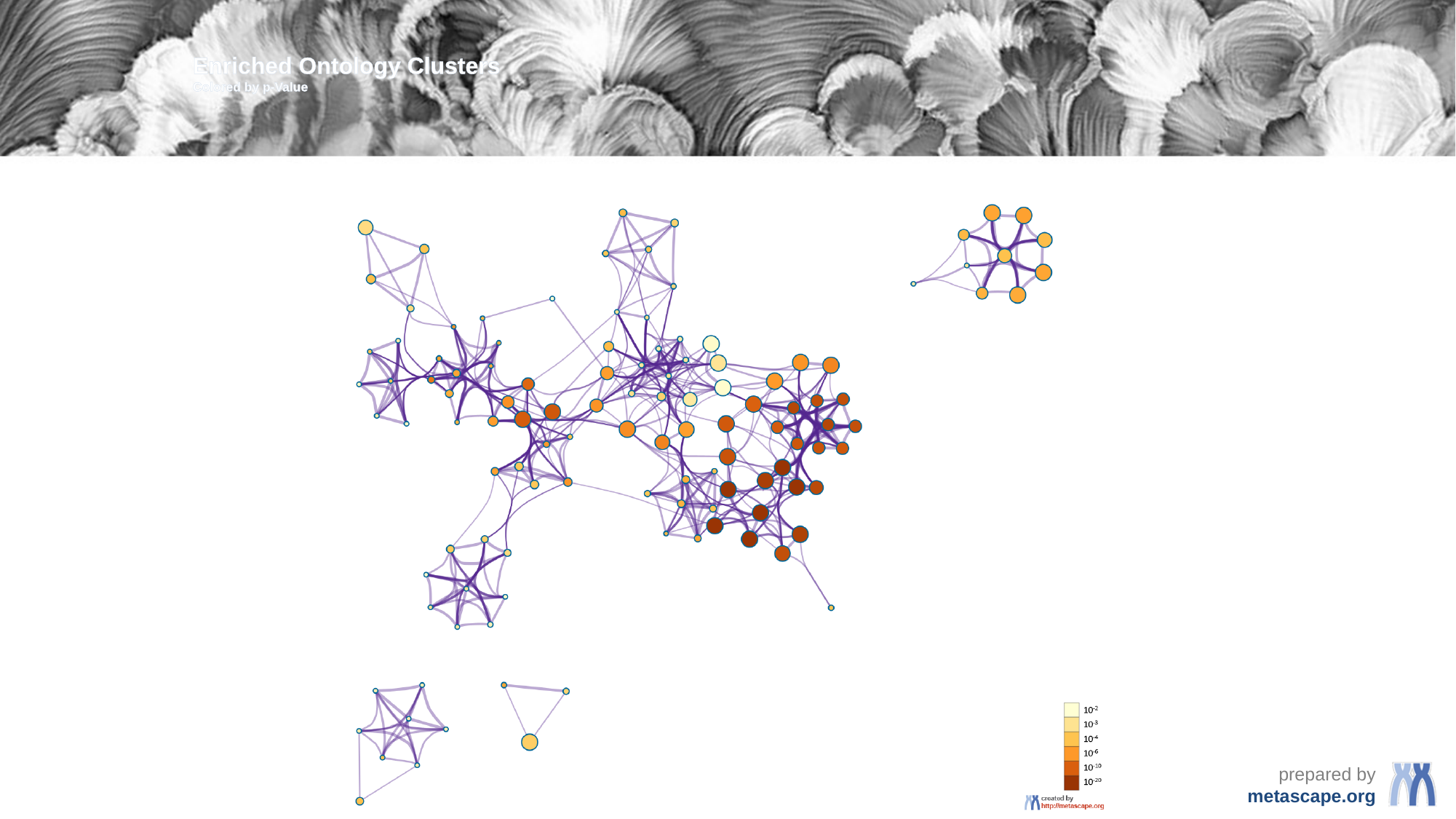

Enriched Ontology Clusters
Colored by p-Value

### Slide 6
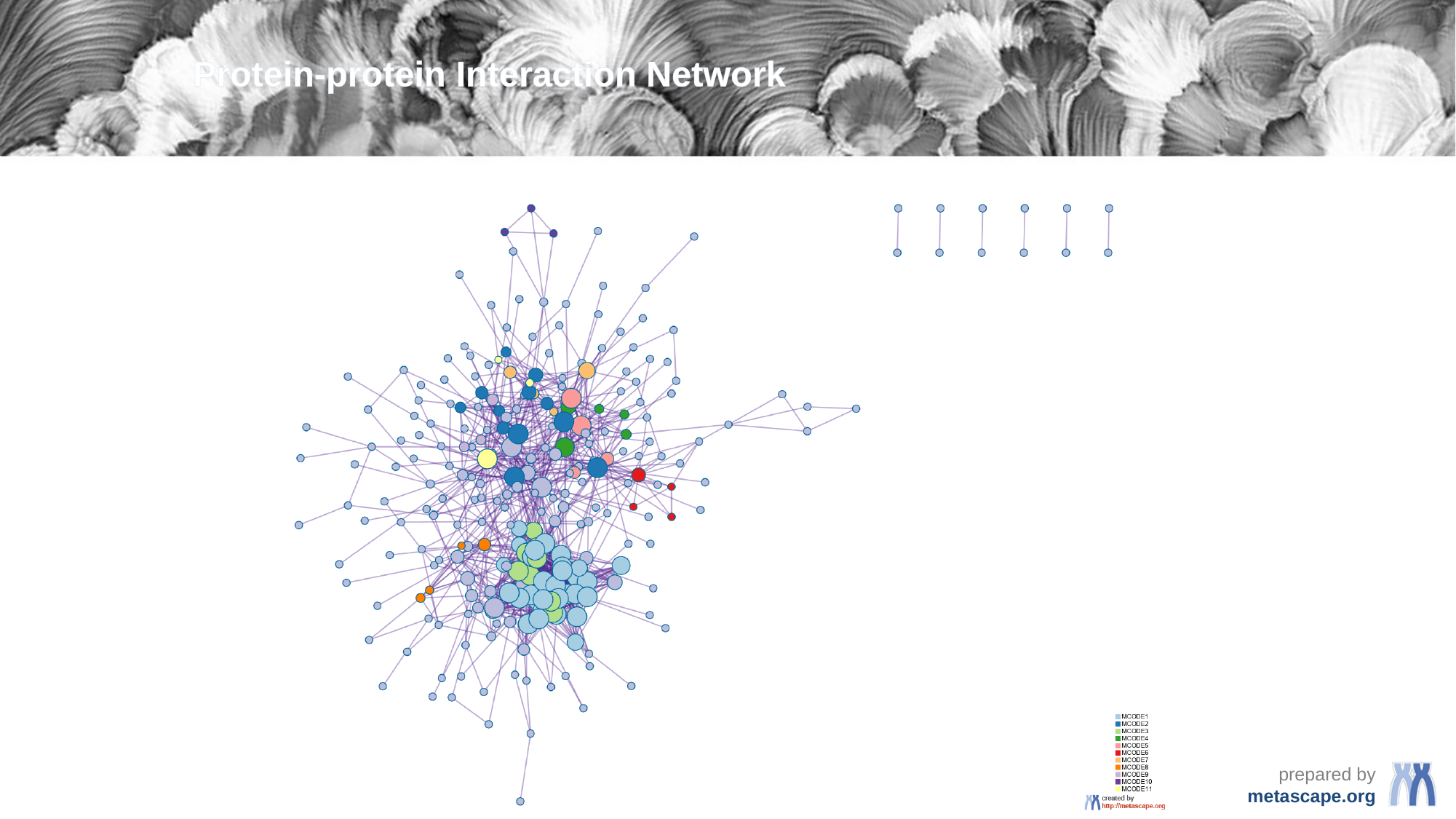

Protein-protein Interaction Network

### Slide 7
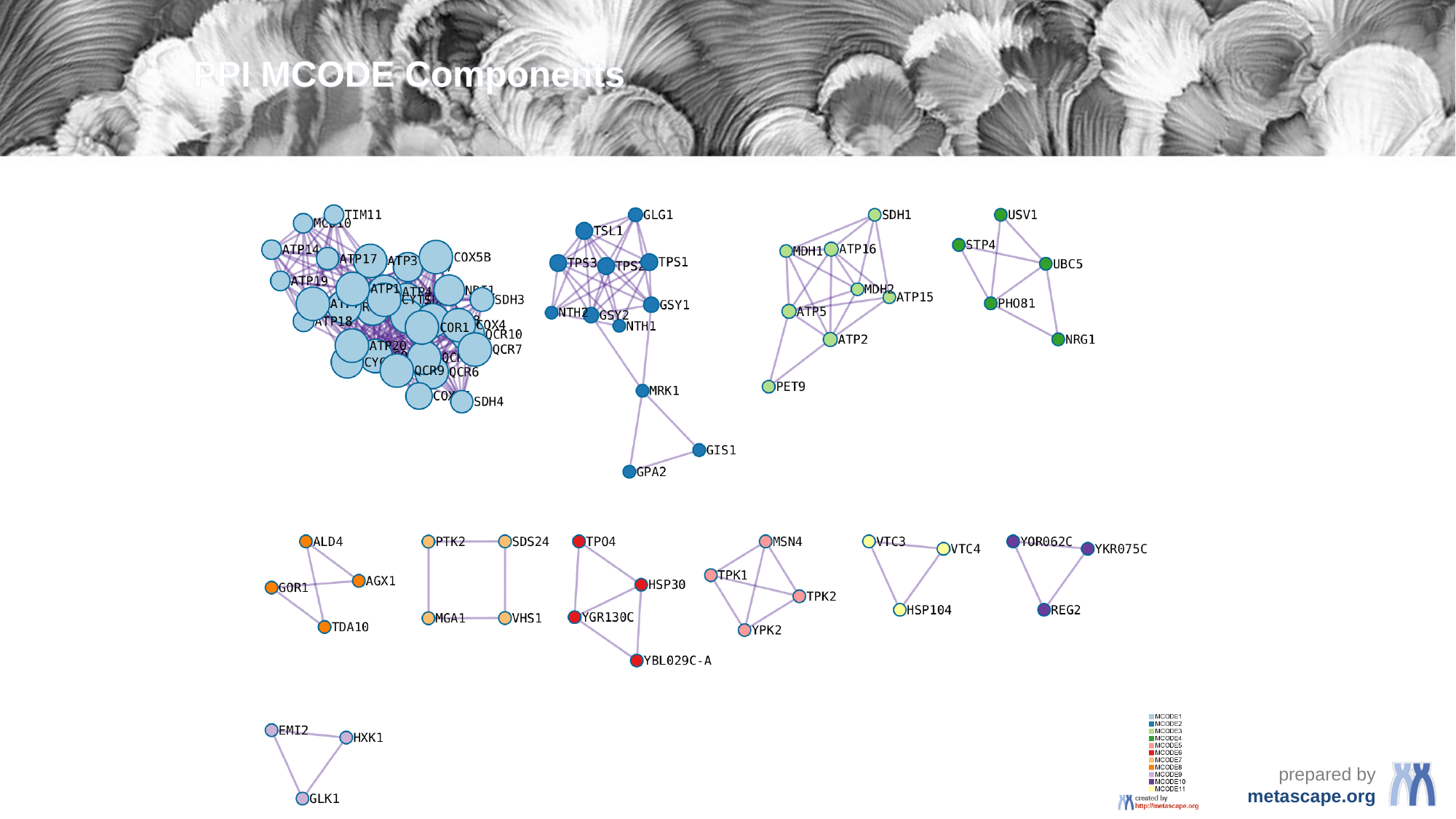

PPI MCODE Components

### Slide 8
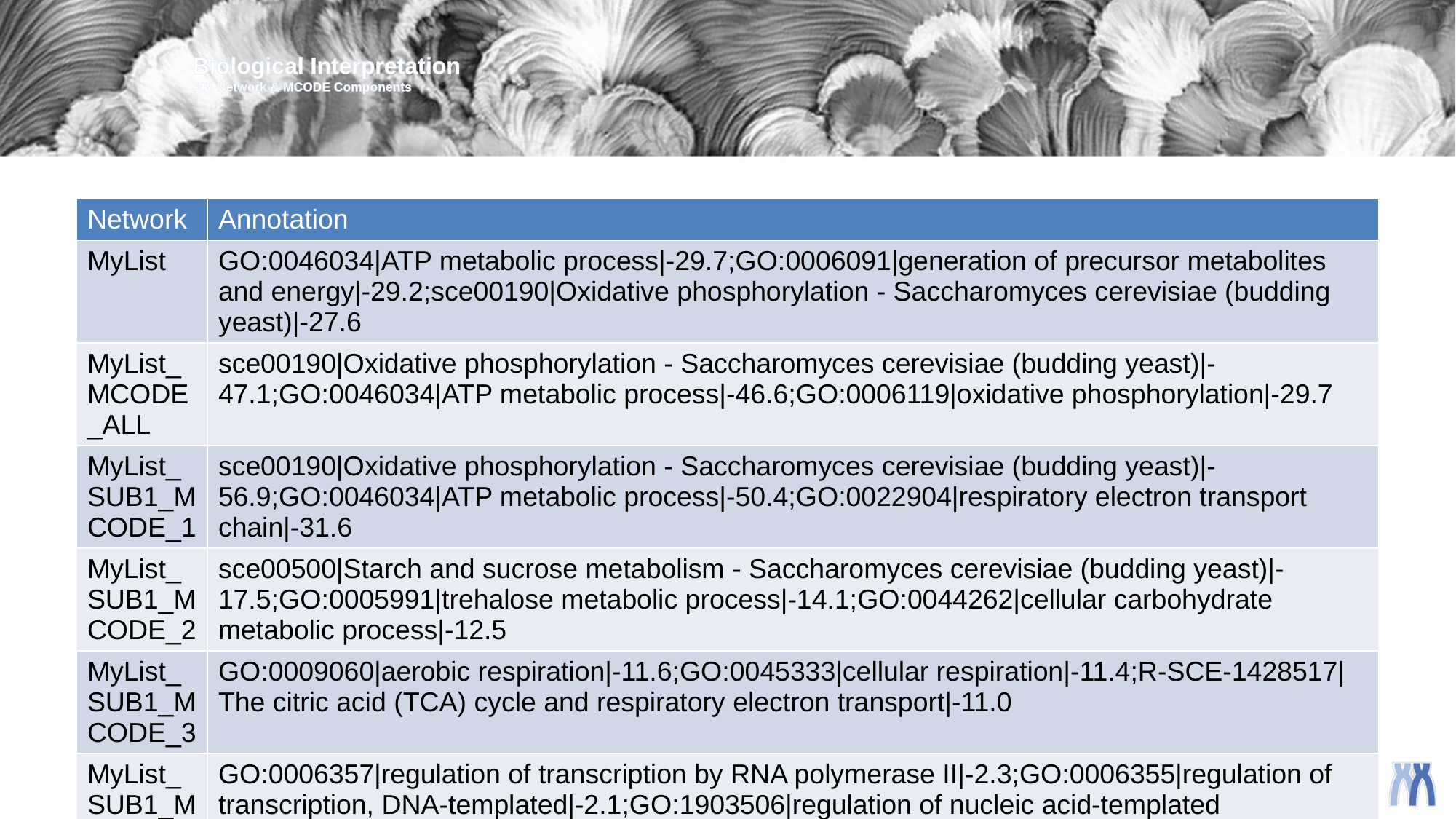

Biological Interpretation
PPI Network & MCODE Components
| Network | Annotation |
| --- | --- |
| MyList | GO:0046034|ATP metabolic process|-29.7;GO:0006091|generation of precursor metabolites and energy|-29.2;sce00190|Oxidative phosphorylation - Saccharomyces cerevisiae (budding yeast)|-27.6 |
| MyList\_MCODE\_ALL | sce00190|Oxidative phosphorylation - Saccharomyces cerevisiae (budding yeast)|-47.1;GO:0046034|ATP metabolic process|-46.6;GO:0006119|oxidative phosphorylation|-29.7 |
| MyList\_SUB1\_MCODE\_1 | sce00190|Oxidative phosphorylation - Saccharomyces cerevisiae (budding yeast)|-56.9;GO:0046034|ATP metabolic process|-50.4;GO:0022904|respiratory electron transport chain|-31.6 |
| MyList\_SUB1\_MCODE\_2 | sce00500|Starch and sucrose metabolism - Saccharomyces cerevisiae (budding yeast)|-17.5;GO:0005991|trehalose metabolic process|-14.1;GO:0044262|cellular carbohydrate metabolic process|-12.5 |
| MyList\_SUB1\_MCODE\_3 | GO:0009060|aerobic respiration|-11.6;GO:0045333|cellular respiration|-11.4;R-SCE-1428517|The citric acid (TCA) cycle and respiratory electron transport|-11.0 |
| MyList\_SUB1\_MCODE\_4 | GO:0006357|regulation of transcription by RNA polymerase II|-2.3;GO:0006355|regulation of transcription, DNA-templated|-2.1;GO:1903506|regulation of nucleic acid-templated transcription|-2.1 |
| MyList\_SUB1\_MCODE\_5 | sce04213|Longevity regulating pathway - multiple species - Saccharomyces cerevisiae (budding yeast)|-6.0;sce04138|Autophagy - yeast - Saccharomyces cerevisiae (budding yeast)|-4.9;sce04113|Meiosis - yeast - Saccharomyces cerevisiae (budding yeast)|-4.4 |
| MyList\_SUB1\_MCODE\_7 | GO:0006468|protein phosphorylation|-4.2;GO:0016310|phosphorylation|-3.6;GO:0007154|cell communication|-2.8 |
| MyList\_SUB1\_MCODE\_8 | sce00630|Glyoxylate and dicarboxylate metabolism - Saccharomyces cerevisiae (budding yeast)|-6.4;sce01110|Biosynthesis of secondary metabolites - Saccharomyces cerevisiae (budding yeast)|-4.9;sce01100|Metabolic pathways - Saccharomyces cerevisiae (budding yeast)|-3.5 |
| MyList\_SUB1\_MCODE\_9 | R-SCE-170822|Regulation of Glucokinase by Glucokinase Regulatory Protein|-9.5;GO:0033500|carbohydrate homeostasis|-8.0;GO:0001678|cellular glucose homeostasis|-8.0 |

### Slide 9
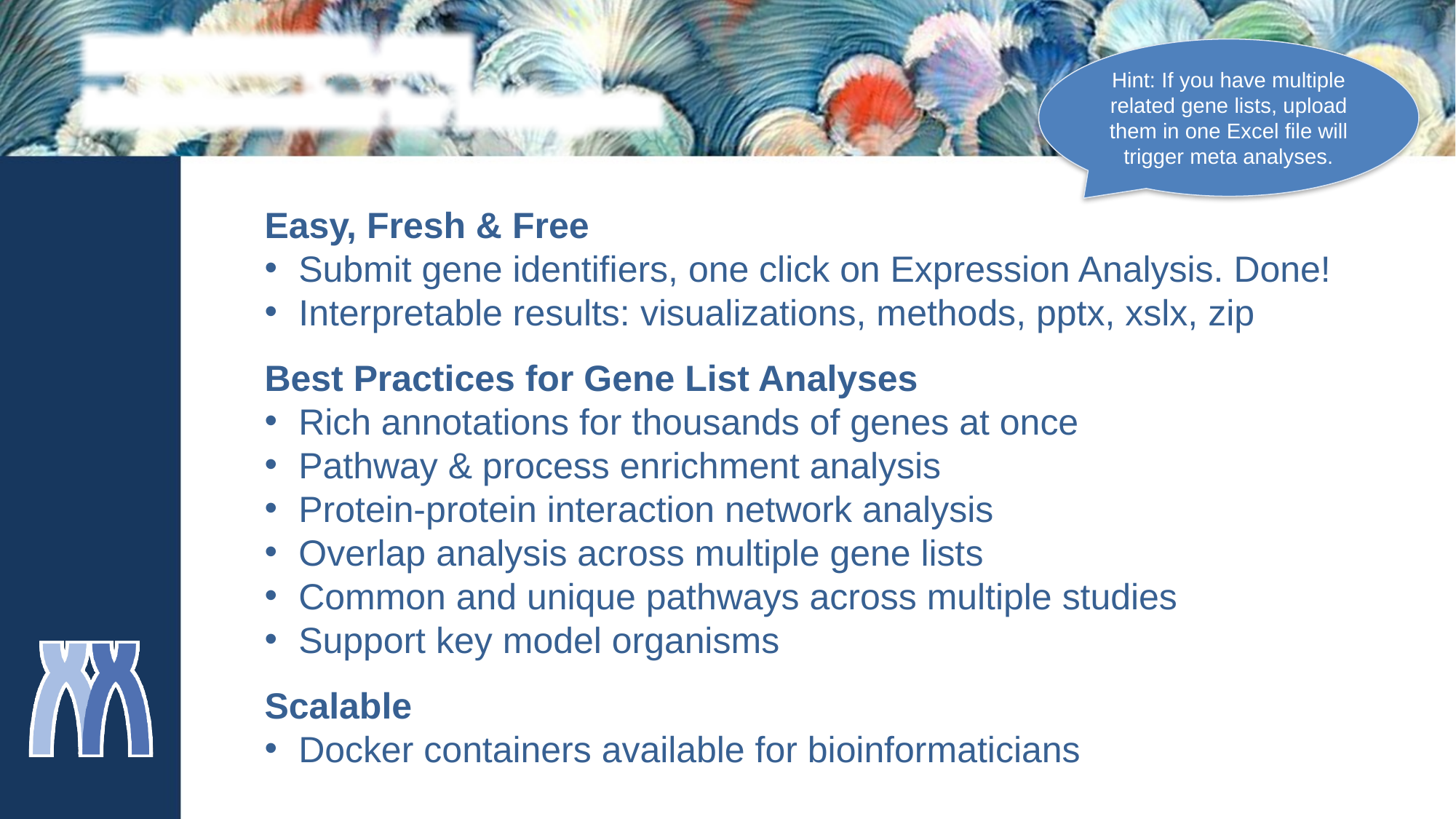

metascape.org
bioinformatics for biologists
Hint: If you have multiple related gene lists, upload them in one Excel file will trigger meta analyses.
Easy, Fresh & Free
Submit gene identifiers, one click on Expression Analysis. Done!
Interpretable results: visualizations, methods, pptx, xslx, zip
Best Practices for Gene List Analyses
Rich annotations for thousands of genes at once
Pathway & process enrichment analysis
Protein-protein interaction network analysis
Overlap analysis across multiple gene lists
Common and unique pathways across multiple studies
Support key model organisms
Scalable
Docker containers available for bioinformaticians
