## Additional File 3 for "Uncharacterized yeast gene *YBR238C,* an effector of TORC1 signaling in a mitochondrial feedback loop, accelerates cellular aging via *HAP4*- and *RMD9*-dependent mechanisms": ColorByCluster.pdf

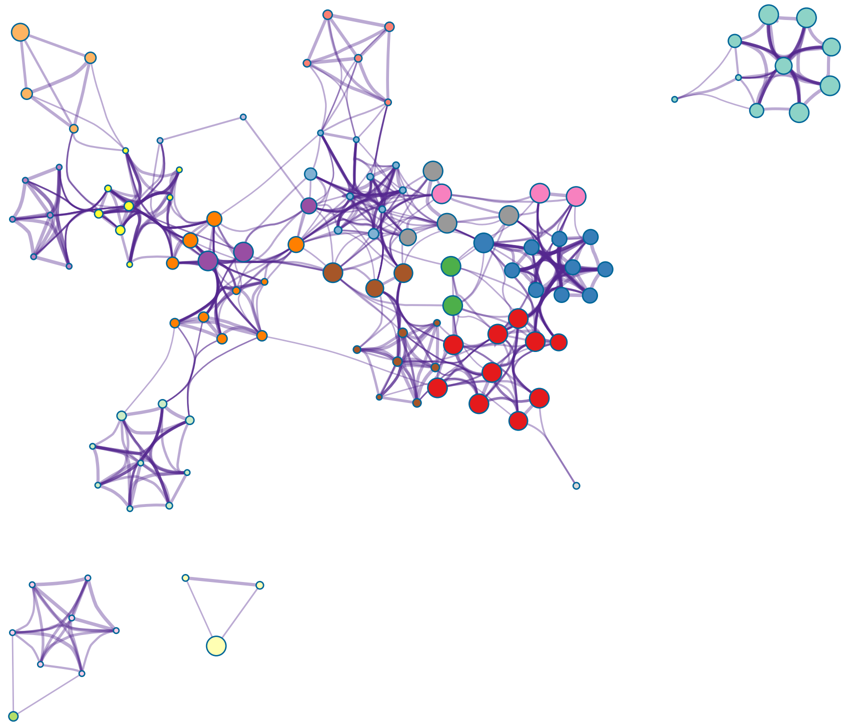

- ATP metabolic process
- ATP biosynthetic process
- Metabolic pathways - Saccharomyces cerevisiae (bu[illegible]
- carbohydrate metabolic process
- Starch and sucrose metabolism - Saccharomyces cer[illegible]
- trehalose metabolic process
- Principal pathways of carbon metabolism
- phosphorus metabolic process
- small molecule metabolic process
- response to extracellular stimulus
- cristae formation
- Trehalose degradation, low osmolarity
- carbohydrate transmembrane transport
- monosaccharide metabolic process
- response to heat
- Longevity regulating pathway - multiple species - Sac[illegible]
- protein kinase A signaling
- mitochondrial electron transport, cytochrome c to oxy[illegible]
- response to desiccation
- regulation of carbohydrate metabolic process

created by
http://metascape.org
