## Additional File 3 for "Uncharacterized yeast gene *YBR238C,* an effector of TORC1 signaling in a mitochondrial feedback loop, accelerates cellular aging via *HAP4*- and *RMD9*-dependent mechanisms": GONetwork.js

```
var networks={"GONetwork": {"format_version": "1.0", "generated_by": "cytoscape-3.9.1", "target_cytoscapejs_version": "~2.1", "data": {"shared_name": "GONetwork", "name": "GONetwork", "SUID": 2741692, "__Annotations": [], "selected": false}, "elements": {"nodes": [{"data": {"id": "2742167", "Enrichment": 14.641929499072354, "BestLogPInGroup": -8.310637859619513, "_PATTERN_": "M1", "_GeneInGOAndHitList": 8.0, "Hits": "HSP104|NTH1|TPS2|NTH2|TPS1|TSL1|PGM2|TPS3", "Description": "trehalose metabolic process", "_GeneInHitList": 294.0, "Category": "GO Biological Processes", "LogP": -8.310637859619513, "_RANK_": 1.0, "FirstInGroupByEnrichment": 1.0, "_LogP_MyList": -8.310637859619513, "_TotalGeneInLibrary": 5919.0, "_GeneInGO": 11.0, "shared_name": "118", "Gene": "GO:0005991", "_MEMBER_MyList": 1.0, "STDV_InGO": 0.9488706521462137, "_COUNT_001": 8.0, "SUID": 2742167, "id_original": "118", "selected": false, "Log_q_value_": -6.18529805792079, "canonicalName": "GO:0005991", "CategoryID": 19.0, "Z_score": 10.352655972315713, "EvidenceCutoff": 0.0, "PARENT_GO": "19_GO:0008152 metabolic process", "FirstInGroupByLogP": 1.0, "GROUP_ID": 6.0, "name": "118", "BestEnrichmentInGroup": 20.132653061224488, "CLUSTER_LABEL": "trehalose metabolic process", "GeneID": "850633|851564|851646|852286|852423|854872|855131|855303", "GiniIndex": 0.0, "_InGO": 2.7210884353741496, "_COUNT_001_PCT": 100.0}, "position": {"x": -50.42886220513253, "y": 217.45008121610908}, "selected": false}, {"data": {"id": "2742164", "Enrichment": 1.8020836306550592, "BestLogPInGroup": -5.971779133371012, "_PATTERN_": "M1", "_GeneInGOAndHitList": 64.0, "Hits": "GLK1|AGX1|HXK1|BNA5|SYM1|ATP14|YLR345W|YDL124W|TMA17|GPM2|ATP16|KGD2|YDR248C|ATP5|TIM11|ATP17|EMI2|ATP1|BNA4|ACH1|ATP3|YBR053C|DSD1|PNC1|XKS1|LSC2|TDH1|VTC4|TES1|YJR096W|ATP2|NNR2|MCR1|MDH1|ATP7|MDH2|BDS1|ATP19|GCY1|PYK2|ALD4|GUT2|XBP1|PKP1|ATP18|PHO84|CYB2|PGM2|HFD1|ALD3|GAD1|GOR1|NNR1|ATP15|ATP4|VTC3|ATP20|ARO9|YEF1|PHM8|GPP2|ICL1|VTC1|MCO10", "Description": "small molecule metabolic process", "_GeneInHitList": 294.0, "Category": "GO Biological Processes", "LogP": -5.971779133371012, "_RANK_": 1.0, "FirstInGroupByEnrichment": 1.0, "_LogP_MyList": -5.971779133371012, "_TotalGeneInLibrary": 5919.0, "_GeneInGO": 715.0, "shared_name": "117", "Gene": "GO:0044281", "_MEMBER_MyList": 1.0, "STDV_InGO": 2.406760689472849, "_COUNT_001": 64.0, "SUID": 2742164, "id_original": "117", "selected": false, "Log_q_value_": -3.9830220494442963, "canonicalName": "GO:0044281", "CategoryID": 19.0, "Z_score": 5.228822656298452, "EvidenceCutoff": 0.0, "PARENT_GO": "19_GO:0008152 metabolic process", "FirstInGroupByLogP": 1.0, "GROUP_ID": 9.0, "name": "117", "BestEnrichmentInGroup": 2.1939429618001043, "CLUSTER_LABEL": "small molecule metabolic process", "GeneID": "850317|850514|850614|850933|850953|851002|851059|851433|851448|851541|851560|851726|851835|851892|851922|851983|852128|852177|852179|852266|852327|852342|852679|852846|853108|853159|853395|853441|853477|853559|853585|853706|853707|853777|853853|853994|854000|854077|854287|854529|854556|854651|854706|854769|854893|854916|854950|855131|855137|855205|855291|855447|855521|855857|856027|856088|856131|856539|856670|856759|856791|856794|856803|1466480", "GiniIndex": 0.0, "_InGO": 21.768707482993197, "_COUNT_001_PCT": 100.0}, "position": {"x": 541.53463877143, "y": 219.90453373075752}, "selected": false}, {"data": {"id": "2742161", "Enrichment": 4.0265306122448985, "BestLogPInGroup": -4.197480086121784, "_PATTERN_": "M1", "_GeneInGOAndHitList": 11.0, "Hits": "HSP12|HSP104|TPS2|HSP42|HSP78|TPS1|YAK1|DDR2|GAC1|HSP82|LSP1", "Description": "response to heat", "_GeneInHitList": 294.0, "Category": "GO Biological Processes", "LogP": -4.197480086121784, "_RANK_": 1.0, "FirstInGroupByEnrichment": 1.0, "_LogP_MyList": -4.197480086121784, "_TotalGeneInLibrary": 5919.0, "_GeneInGO": 55.0, "shared_name": "116", "Gene": "GO:0009408", "_MEMBER_MyList": 1.0, "STDV_InGO": 1.1067985072269073, "_COUNT_001": 11.0, "SUID": 2742161, "id_original": "116", "selected": false, "Log_q_value_": -2.391653685781055, "canonicalName": "GO:0009408", "CategoryID": 19.0, "Z_score": 5.155007686164529, "EvidenceCutoff": 0.0, "PARENT_GO": "19_GO:0050896 response to stimulus", "FirstInGroupByLogP": 1.0, "GROUP_ID": 15.0, "name": "116", "BestEnrichmentInGroup": 4.129774986917844, "CLUSTER_LABEL": "response to heat", "GeneID": "850532|850633|851646|851751|851845|852423|853300|854104|854350|855836|856103", "GiniIndex": 0.0, "_InGO": 3.741496598639456, "_COUNT_001_PCT": 100.0}, "position": {"x": -154.24444448052316, "y": 44.17514835478096}, "selected": false}, {"data": {"id": "2742158", "Enrichment": 3.131746031746032, "BestLogPInGroup": -5.6116974553384775, "_PATTERN_": "M1", "_GeneInGOAndHitList": 21.0, "Hits": "DCS1|NTH1|NRG1|ATG8|PHO5|TPS1|SDS24|STF2|MTL1|YAK1|GSM1|HAP4|MSN4|YKR075C|DDR2|YOR062C|DCS2|CAT8|USV1|PEP4|PHM8", "Description": "response to extracellular stimulus", "_GeneInHitList": 294.0, "Category": "GO Biological Processes", "LogP": -5.6116974553384775, "_RANK_": 1.0, "FirstInGroupByEnrichment": 1.0, "_LogP_MyList": -5.6116974553384775, "_TotalGeneInLibrary": 5919.0, "_GeneInGO": 135.0, "shared_name": "115", "Gene": "GO:0009991", "_MEMBER_MyList": 1.0, "STDV_InGO": 1.502000147334668, "_COUNT_001": 21.0, "SUID": 2742158, "id_original": "115", "selected": false, "Log_q_value_": -3.6624489126954356, "canonicalName": "GO:0009991", "CategoryID": 19.0, "Z_score": 5.72780319727626, "EvidenceCutoff": 0.0, "PARENT_GO": "19_GO:0050896 response to stimulus", "FirstInGroupByLogP": 1.0, "GROUP_ID": 10.0, "name": "115", "BestEnrichmentInGroup": 6.710884353741496, "CLUSTER_LABEL": "response to extracellular stimulus", "GeneID": "850974|851564|851613|852200|852390|852423|852515|852891|852905|853300|853341|853751|853803|853950|854104|854228|854344|855323|855871|855949|856759", "GiniIndex": 0.0, "_InGO": 7.142857142857143, "_COUNT_001_PCT": 100.0}, "position": {"x": 971.0383313983831, "y": -65.44020809053154}, "selected": false}, {"data": {"id": "2742155", "Enrichment": 20.13265306122449, "BestLogPInGroup": -3.915927067855309, "_PATTERN_": "M1", "_GeneInGOAndHitList": 3.0, "Hits": "NTH1|TPS1|STF2", "Description": "response to desiccation", "_GeneInHitList": 294.0, "Category": "GO Biological Processes", "LogP": -3.915927067855309, "_RANK_": 1.0, "FirstInGroupByEnrichment": 1.0, "_LogP_MyList": -3.915927067855309, "_TotalGeneInLibrary": 5919.0, "_GeneInGO": 3.0, "shared_name": "114", "Gene": "GO:0009269", "_MEMBER_MyList": 1.0, "STDV_InGO": 0.5861194403214197, "_COUNT_001": 3.0, "SUID": 2742155, "id_original": "114", "selected": false, "Log_q_value_": -2.165255932415635, "canonicalName": "GO:0009269", "CategoryID": 19.0, "Z_score": 7.577424595277936, "EvidenceCutoff": 0.0, "PARENT_GO": "19_GO:0050896 response to stimulus", "FirstInGroupByLogP": 1.0, "GROUP_ID": 19.0, "name": "114", "BestEnrichmentInGroup": 20.13265306122449, "CLUSTER_LABEL": "response to desiccation", "GeneID": "851564|852423|852891", "GiniIndex": 0.0, "_InGO": 1.0204081632653061, "_COUNT_001_PCT": 100.0}, "position": {"x": -120.42376576958566, "y": 219.52713047148018}, "selected": false}, {"data": {"id": "2742152", "Enrichment": 4.314139941690962, "BestLogPInGroup": -3.769282700434612, "_PATTERN_": "M1", "_GeneInGOAndHitList": 9.0, "Hits": "IGD1|YLR345W|PIB1|VID24|GAC1|FYV10|PCL7|PKP1|UBC8", "Description": "regulation of carbohydrate metabolic process", "_GeneInHitList": 294.0, "Category": "GO Biological Processes", "LogP": -3.769282700434612, "_RANK_": 1.0, "FirstInGroupByEnrichment": 1.0, "_LogP_MyList": -3.769282700434612, "_TotalGeneInLibrary": 5919.0, "_GeneInGO": 42.0, "shared_name": "113", "Gene": "GO:0006109", "_MEMBER_MyList": 1.0, "STDV_InGO": 1.0046682763914156, "_COUNT_001": 9.0, "SUID": 2742152, "id_original": "113", "selected": false, "Log_q_value_": -2.0304030897244965, "canonicalName": "GO:0006109", "CategoryID": 19.0, "Z_score": 4.927396661272298, "EvidenceCutoff": 0.0, "PARENT_GO": "19_GO:0050789 regulation of biological process", "FirstInGroupByLogP": 1.0, "GROUP_ID": 20.0, "name": "113", "BestEnrichmentInGroup": 7.320964749536177, "CLUSTER_LABEL": "regulation of carbohydrate metabolic process", "GeneID": "850572|851059|851908|852402|854350|854710|854761|854769|856704", "GiniIndex": 0.0, "_InGO": 3.061224489795918, "_COUNT_001_PCT": 100.0}, "position": {"x": -17.385191550835657, "y": 509.01433597685127}, "selected": false}, {"data": {"id": "2742149", "Enrichment": 13.421768707482991, "BestLogPInGroup": -4.082697487204731, "_PATTERN_": "M1", "_GeneInGOAndHitList": 4.0, "Hits": "TPK1|YAK1|TPK2|GPA2", "Description": "protein kinase A signaling", "_GeneInHitList": 294.0, "Category": "GO Biological Processes", "LogP": -4.082697487204731, "_RANK_": 1.0, "FirstInGroupByEnrichment": 1.0, "_LogP_MyList": -4.082697487204731, "_TotalGeneInLibrary": 5919.0, "_GeneInGO": 6.0, "shared_name": "112", "Gene": "GO:0010737", "_MEMBER_MyList": 1.0, "STDV_InGO": 0.6756285589444189, "_COUNT_001": 4.0, "SUID": 2742149, "id_original": "112", "selected": false, "Log_q_value_": -2.2989212276070763, "canonicalName": "GO:0010737", "CategoryID": 19.0, "Z_score": 6.959130912757607, "EvidenceCutoff": 0.0, "PARENT_GO": "19_GO:0023052 signaling", "FirstInGroupByLogP": 1.0, "GROUP_ID": 17.0, "name": "112", "BestEnrichmentInGroup": 13.421768707482991, "CLUSTER_LABEL": "protein kinase A signaling", "GeneID": "853275|853300|855898|856741", "GiniIndex": 0.0, "_InGO": 1.3605442176870748, "_COUNT_001_PCT": 100.0}, "position": {"x": -134.52859365044503, "y": 867.6410297024372}, "selected": false}, {"data": {"id": "2742146", "Enrichment": 2.0034124579685058, "BestLogPInGroup": -7.330800802960937, "_PATTERN_": "M1", "_GeneInGOAndHitList": 61.0, "Hits": "GLK1|KIN82|HXK1|KNS1|UPS2|BNA5|ATP14|YLR345W|PHO11|MRK1|GPD1|GPM2|ATP16|KGD2|VHS1|YDR248C|ATP5|TIM11|ATP17|EMI2|ATP1|BNA4|ATP3|PHO5|SDS24|PNC1|FMP48|XKS1|TDA10|PHO81|LSC2|SOL4|TPK1|YAK1|TDH1|VTC4|TES1|PTK2|ATP2|NNR2|ATP7|ATP19|HRK1|PYK2|GUT2|PKP1|ATP18|PHO84|PLB1|YPK2|PGM2|NNR1|ATP15|TPK2|ATP4|VTC3|ATP20|YEF1|PHM8|VTC1|MCO10", "Description": "phosphorus metabolic process", "_GeneInHitList": 294.0, "Category": "GO Biological Processes", "LogP": -7.330800802960937, "_RANK_": 1.0, "FirstInGroupByEnrichment": 1.0, "_LogP_MyList": -7.330800802960937, "_TotalGeneInLibrary": 5919.0, "_GeneInGO": 613.0, "shared_name": "111", "Gene": "GO:0006793", "_MEMBER_MyList": 1.0, "STDV_InGO": 2.36494955959483, "_COUNT_001": 61.0, "SUID": 2742146, "id_original": "111", "selected": false, "Log_q_value_": -5.258706513215438, "canonicalName": "GO:0006793", "CategoryID": 19.0, "Z_score": 5.998269650058398, "EvidenceCutoff": 0.0, "PARENT_GO": "19_GO:0008152 metabolic process", "FirstInGroupByLogP": 1.0, "GROUP_ID": 8.0, "name": "111", "BestEnrichmentInGroup": 2.0034124579685058, "CLUSTER_LABEL": "phosphorus metabolic process", "GeneID": "850317|850453|850614|850641|850865|850933|851002|851059|851299|851480|851539|851541|851560|851726|851834|851835|851892|851922|851983|852128|852177|852179|852327|852390|852515|852846|852943|853108|853119|853148|853159|853163|853275|853300|853395|853441|853477|853522|853585|853706|853853|854077|854441|854529|854651|854769|854893|854916|855020|855130|855131|855521|855857|855898|856027|856088|856131|856670|856759|856803|1466480", "GiniIndex": 0.0, "_InGO": 20.74829931972789, "_COUNT_001_PCT": 100.0}, "position": {"x": 638.5928510517034, "y": 192.49716220976143}, "selected": false}, {"data": {"id": "2742143", "Enrichment": 3.8966425279789343, "BestLogPInGroup": -4.381277688016449, "_PATTERN_": "M1", "_GeneInGOAndHitList": 12.0, "Hits": "GLK1|HXK1|YLR345W|EMI2|YBR053C|XKS1|TDH1|YJR096W|MDH2|GCY1|PKP1|PGM2", "Description": "monosaccharide metabolic process", "_GeneInHitList": 294.0, "Category": "GO Biological Processes", "LogP": -4.381277688016449, "_RANK_": 1.0, "FirstInGroupByEnrichment": 1.0, "_LogP_MyList": -4.381277688016449, "_TotalGeneInLibrary": 5919.0, "_GeneInGO": 62.0, "shared_name": "110", "Gene": "GO:0005996", "_MEMBER_MyList": 1.0, "STDV_InGO": 1.153969104668408, "_COUNT_001": 12.0, "SUID": 2742143, "id_original": "110", "selected": false, "Log_q_value_": -2.56630790823585, "canonicalName": "GO:0005996", "CategoryID": 19.0, "Z_score": 5.2414689520187565, "EvidenceCutoff": 0.0, "PARENT_GO": "19_GO:0008152 metabolic process", "FirstInGroupByLogP": 1.0, "GROUP_ID": 14.0, "name": "110", "BestEnrichmentInGroup": 12.079591836734696, "CLUSTER_LABEL": "monosaccharide metabolic process", "GeneID": "850317|850614|851059|852128|852342|853108|853395|853559|853994|854287|854769|855131", "GiniIndex": 0.0, "_InGO": 4.081632653061225, "_COUNT_001_PCT": 100.0}, "position": {"x": 255.3870405048284, "y": 160.06679569364815}, "selected": false}, {"data": {"id": "2742140", "Enrichment": 7.105642256902762, "BestLogPInGroup": -3.954929368414704, "_PATTERN_": "M1", "_GeneInGOAndHitList": 6.0, "Hits": "COX8|COX4|COX5B|COX7|COX5A|CYC7", "Description": "mitochondrial electron transport, cytochrome c to oxygen", "_GeneInHitList": 294.0, "Category": "GO Biological Processes", "LogP": -3.954929368414704, "_RANK_": 1.0, "FirstInGroupByEnrichment": 1.0, "_LogP_MyList": -3.954929368414704, "_TotalGeneInLibrary": 5919.0, "_GeneInGO": 17.0, "shared_name": "109", "Gene": "GO:0006123", "_MEMBER_MyList": 1.0, "STDV_InGO": 0.8246143220834374, "_COUNT_001": 6.0, "SUID": 2742140, "id_original": "109", "selected": false, "Log_q_value_": -2.183865074333187, "canonicalName": "GO:0006123", "CategoryID": 19.0, "Z_score": 5.763099195053121, "EvidenceCutoff": 0.0, "PARENT_GO": "19_GO:0008152 metabolic process", "FirstInGroupByLogP": 1.0, "GROUP_ID": 18.0, "name": "109", "BestEnrichmentInGroup": 7.105642256902762, "CLUSTER_LABEL": "mitochondrial electron transport, cytochrome c to oxygen", "GeneID": "851111|852688|854695|855298|855675|856672", "GiniIndex": 0.0, "_InGO": 2.0408163265306123, "_COUNT_001_PCT": 100.0}, "position": {"x": 638.9599012226018, "y": 610.0029529202106}, "selected": false}, {"data": {"id": "2742137", "Enrichment": 10.98144712430427, "BestLogPInGroup": -5.272416855992848, "_PATTERN_": "M1", "_GeneInGOAndHitList": 6.0, "Hits": "MIC19|UPS2|ATP14|TIM11|MIC27|ATP20", "Description": "cristae formation", "_GeneInHitList": 294.0, "Category": "GO Biological Processes", "LogP": -5.272416855992848, "_RANK_": 1.0, "FirstInGroupByEnrichment": 1.0, "_LogP_MyList": -5.272416855992848, "_TotalGeneInLibrary": 5919.0, "_GeneInGO": 11.0, "shared_name": "108", "Gene": "GO:0042407", "_MEMBER_MyList": 1.0, "STDV_InGO": 0.8246143220834374, "_COUNT_001": 6.0, "SUID": 2742137, "id_original": "108", "selected": false, "Log_q_value_": -3.376413825303031, "canonicalName": "GO:0042407", "CategoryID": 19.0, "Z_score": 7.574770723658172, "EvidenceCutoff": 0.0, "PARENT_GO": "19_GO:0009987 cellular process", "FirstInGroupByLogP": 1.0, "GROUP_ID": 11.0, "name": "108", "BestEnrichmentInGroup": 10.98144712430427, "CLUSTER_LABEL": "cristae formation", "GeneID": "850563|850865|851002|851922|855623|856131", "GiniIndex": 0.0, "_InGO": 2.0408163265306123, "_COUNT_001_PCT": 100.0}, "position": {"x": 74.86131418646903, "y": 742.9577258694294}, "selected": false}, {"data": {"id": "2742134", "Enrichment": 5.1769679300291545, "BestLogPInGroup": -4.436199066039166, "_PATTERN_": "M1", "_GeneInGOAndHitList": 9.0, "Hits": "GLK1|AQY3|HXK1|MTH1|HXT7|HXT6|MAL31|VPS73|HXT4", "Description": "carbohydrate transmembrane transport", "_GeneInHitList": 294.0, "Category": "GO Biological Processes", "LogP": -4.436199066039166, "_RANK_": 1.0, "FirstInGroupByEnrichment": 1.0, "_LogP_MyList": -4.436199066039166, "_TotalGeneInLibrary": 5919.0, "_GeneInGO": 35.0, "shared_name": "107", "Gene": "GO:0034219", "_MEMBER_MyList": 1.0, "STDV_InGO": 1.0046682763914156, "_COUNT_001": 9.0, "SUID": 2742134, "id_original": "107", "selected": false, "Log_q_value_": -2.616584381212804, "canonicalName": "GO:0034219", "CategoryID": 19.0, "Z_score": 5.6657670348601235, "EvidenceCutoff": 0.0, "PARENT_GO": "19_GO:0051179 localization", "FirstInGroupByLogP": 1.0, "GROUP_ID": 13.0, "name": "107", "BestEnrichmentInGroup": 5.872023809523809, "CLUSTER_LABEL": "carbohydrate transmembrane transport", "GeneID": "850317|850490|850614|851870|851943|851944|852601|852774|856492", "GiniIndex": 0.0, "_InGO": 3.061224489795918, "_COUNT_001_PCT": 100.0}, "position": {"x": 279.99488962592216, "y": -69.8695446383831}, "selected": false}, {"data": {"id": "2742131", "Enrichment": 3.4414791557648696, "BestLogPInGroup": -11.663532448682075, "_PATTERN_": "M1", "_GeneInGOAndHitList": 40.0, "Hits": "GLK1|GSY1|IGD1|HXK1|HSP104|GSY2|YLR345W|GPD1|GPM2|NTH1|TPS2|YDR248C|EMI2|NTH2|YBR053C|MRX18|TPS1|ARA1|MAL31|XKS1|SOL4|TDH1|YJR096W|GLG1|MDH2|GCY1|GAC1|PYK2|GUT2|PCL7|PKP1|EGH1|TSL1|PGM2|TPS3|CAT8|GPH1|GLC3|GPP2|ICL1", "Description": "carbohydrate metabolic process", "_GeneInHitList": 294.0, "Category": "GO Biological Processes", "LogP": -11.663532448682075, "_RANK_": 1.0, "FirstInGroupByEnrichment": 1.0, "_LogP_MyList": -11.663532448682075, "_TotalGeneInLibrary": 5919.0, "_GeneInGO": 234.0, "shared_name": "106", "Gene": "GO:0005975", "_MEMBER_MyList": 1.0, "STDV_InGO": 1.9995204874649775, "_COUNT_001": 40.0, "SUID": 2742131, "id_original": "106", "selected": false, "Log_q_value_": -9.290408163272595, "canonicalName": "GO:0005975", "CategoryID": 19.0, "Z_score": 8.711553282581155, "EvidenceCutoff": 0.0, "PARENT_GO": "19_GO:0008152 metabolic process", "FirstInGroupByLogP": 1.0, "GROUP_ID": 4.0, "name": "106", "BestEnrichmentInGroup": 4.200337689032448, "CLUSTER_LABEL": "carbohydrate metabolic process", "GeneID": "850317|850569|850572|850614|850633|850962|851059|851539|851541|851564|851646|851835|852128|852286|852342|852346|852423|852446|852601|853108|853163|853395|853559|853932|853994|854287|854350|854529|854651|854761|854769|854824|854872|855131|855303|855323|856289|856705|856791|856794", "GiniIndex": 0.0, "_InGO": 13.605442176870747, "_COUNT_001_PCT": 100.0}, "position": {"x": 158.47750986517997, "y": 272.67359005094795}, "selected": false}, {"data": {"id": "2742128", "Enrichment": 16.10612244897959, "BestLogPInGroup": -4.542469332779946, "_PATTERN_": "M1", "_GeneInGOAndHitList": 4.0, "Hits": "GLK1|NTH1|TPS2|NTH2", "Description": "Trehalose degradation, low osmolarity", "_GeneInHitList": 294.0, "Category": "WikiPathways", "LogP": -4.542469332779946, "_RANK_": 1.0, "FirstInGroupByEnrichment": 1.0, "_LogP_MyList": -4.542469332779946, "_TotalGeneInLibrary": 5919.0, "_GeneInGO": 5.0, "shared_name": "105", "Gene": "WP70", "_MEMBER_MyList": 1.0, "STDV_InGO": 0.6756285589444189, "_COUNT_001": 4.0, "SUID": 2742128, "id_original": "105", "selected": false, "Log_q_value_": -2.708614208838974, "canonicalName": "WP70", "CategoryID": 27.0, "Z_score": 7.724977533272773, "EvidenceCutoff": 0.0, "PARENT_GO": "", "FirstInGroupByLogP": 1.0, "GROUP_ID": 12.0, "name": "105", "BestEnrichmentInGroup": 16.10612244897959, "CLUSTER_LABEL": "Trehalose degradation, low osmolarity", "GeneID": "850317|851564|851646|852286", "GiniIndex": 0.0, "_InGO": 1.3605442176870748, "_COUNT_001_PCT": 100.0}, "position": {"x": 38.10354746283622, "y": 111.68717228056221}, "selected": false}, {"data": {"id": "2742125", "Enrichment": 7.365604778496764, "BestLogPInGroup": -9.417680144618705, "_PATTERN_": "M1", "_GeneInGOAndHitList": 15.0, "Hits": "GLK1|GSY1|HXK1|GSY2|NTH1|TPS2|EMI2|NTH2|TPS1|GLG1|TSL1|PGM2|TPS3|GPH1|GLC3", "Description": "Starch and sucrose metabolism - Saccharomyces cerevisiae (budding yeast)", "_GeneInHitList": 294.0, "Category": "KEGG Pathway", "LogP": -9.417680144618705, "_RANK_": 1.0, "FirstInGroupByEnrichment": 1.0, "_LogP_MyList": -9.417680144618705, "_TotalGeneInLibrary": 5919.0, "_GeneInGO": 41.0, "shared_name": "104", "Gene": "sce00500", "_MEMBER_MyList": 1.0, "STDV_InGO": 1.2832956896252923, "_COUNT_001": 15.0, "SUID": 2742125, "id_original": "104", "selected": false, "Log_q_value_": -7.197784235305403, "canonicalName": "sce00500", "CategoryID": 24.0, "Z_score": 9.350109366945773, "EvidenceCutoff": 0.0, "PARENT_GO": "", "FirstInGroupByLogP": 1.0, "GROUP_ID": 5.0, "name": "104", "BestEnrichmentInGroup": 11.504373177842565, "CLUSTER_LABEL": "Starch and sucrose metabolism - Saccharomyces cerevisiae (budding yeast)", "GeneID": "850317|850569|850614|850962|851564|851646|852128|852286|852423|853932|854872|855131|855303|856289|856705", "GiniIndex": 0.0, "_InGO": 5.1020408163265305, "_COUNT_001_PCT": 100.0}, "position": {"x": 116.4863294452581, "y": 224.9177859890583}, "selected": false}, {"data": {"id": "2742122", "Enrichment": 4.5871867734435545, "BestLogPInGroup": -7.45117124911709, "_PATTERN_": "M1", "_GeneInGOAndHitList": 18.0, "Hits": "GLK1|HXK1|SDH2|GPD1|KGD2|SDH4|LSC2|SOL4|TDH1|SDH1|SDH3|MDH1|MDH2|PYK2|ALD4|GUT2|GPP2|ICL1", "Description": "Principal pathways of carbon metabolism", "_GeneInHitList": 294.0, "Category": "WikiPathways", "LogP": -7.45117124911709, "_RANK_": 1.0, "FirstInGroupByEnrichment": 1.0, "_LogP_MyList": -7.45117124911709, "_TotalGeneInLibrary": 5919.0, "_GeneInGO": 79.0, "shared_name": "103", "Gene": "WP112", "_MEMBER_MyList": 1.0, "STDV_InGO": 1.3982016102952535, "_COUNT_001": 18.0, "SUID": 2742122, "id_original": "103", "selected": false, "Log_q_value_": -5.370643791834729, "canonicalName": "WP112", "CategoryID": 27.0, "Z_score": 7.3377176079190285, "EvidenceCutoff": 0.0, "PARENT_GO": "", "FirstInGroupByLogP": 1.0, "GROUP_ID": 7.0, "name": "103", "BestEnrichmentInGroup": 11.184807256235827, "CLUSTER_LABEL": "Principal pathways of carbon metabolism", "GeneID": "850317|850614|850685|851539|851726|851758|853159|853163|853395|853709|853716|853777|853994|854529|854556|854651|856791|856794", "GiniIndex": 0.0, "_InGO": 6.122448979591836, "_COUNT_001_PCT": 100.0}, "position": {"x": 347.98372782172294, "y": 325.0690654385212}, "selected": false}, {"data": {"id": "2742119", "Enrichment": 2.183364644722958, "BestLogPInGroup": -12.723364984945494, "_PATTERN_": "M1", "_GeneInGOAndHitList": 86.0, "Hits": "GLK1|AGX1|GSY1|QCR6|HXK1|SDH2|COX12|BNA5|GSY2|ATP14|COX8|PHO11|GPM2|ATP16|NTH1|TPS2|KGD2|SDH4|YDR248C|ATP5|TIM11|ATP17|EMI2|QCR7|ATP1|BNA4|COR1|ACH1|NTH2|ATP3|RFS1|PHO5|TPS1|ARA1|YPC1|DSD1|COX4|PNC1|QCR9|XKS1|TDA10|LSC2|SOL4|QCR8|TDH1|TES1|ATP2|SDH1|SDH3|MDH1|ATP7|GLG1|MDH2|ATP19|CYT1|GCY1|PYK2|ALD4|COX5B|TSL1|ATP18|CYB2|PGM2|HFD1|ALD3|GAD1|COX7|TPS3|GOR1|COX5A|ATP15|YDC1|ATP4|HST2|ATP20|GPH1|QCR2|QCR10|ARO9|CYC7|RIP1|GLC3|PHM8|GPP2|ICL1|MCO10", "Description": "Metabolic pathways - Saccharomyces cerevisiae (budding yeast)", "_GeneInHitList": 294.0, "Category": "KEGG Pathway", "LogP": -12.723364984945494, "_RANK_": 1.0, "FirstInGroupByEnrichment": 1.0, "_LogP_MyList": -12.723364984945494, "_TotalGeneInLibrary": 5919.0, "_GeneInGO": 793.0, "shared_name": "102", "Gene": "sce01100", "_MEMBER_MyList": 1.0, "STDV_InGO": 2.6531383517257234, "_COUNT_001": 86.0, "SUID": 2742119, "id_original": "102", "selected": false, "Log_q_value_": -10.238620237559852, "canonicalName": "sce01100", "CategoryID": 24.0, "Z_score": 8.185887367118221, "EvidenceCutoff": 0.0, "PARENT_GO": "", "FirstInGroupByLogP": 1.0, "GROUP_ID": 3.0, "name": "102", "BestEnrichmentInGroup": 2.5891216797140144, "CLUSTER_LABEL": "Metabolic pathways - Saccharomyces cerevisiae (budding yeast)", "GeneID": "850317|850514|850569|850593|850614|850685|850727|850933|850962|851002|851111|851299|851541|851560|851564|851646|851726|851758|851835|851892|851922|851983|852128|852142|852177|852179|852235|852266|852286|852327|852341|852390|852423|852446|852481|852679|852688|852846|853095|853108|853119|853159|853163|853273|853395|853477|853585|853709|853716|853777|853853|853932|853994|854077|854231|854287|854529|854556|854695|854872|854893|854950|855131|855137|855205|855291|855298|855303|855447|855675|855857|856018|856027|856092|856131|856289|856321|856390|856539|856672|856689|856705|856759|856791|856794|1466480", "GiniIndex": 0.0, "_InGO": 29.25170068027211, "_COUNT_001_PCT": 100.0}, "position": {"x": 460.43114603461356, "y": 349.8758395779255}, "selected": false}, {"data": {"id": "2742116", "Enrichment": 4.768259935553168, "BestLogPInGroup": -4.130078175744296, "_PATTERN_": "M1", "_GeneInGOAndHitList": 9.0, "Hits": "HSP104|SSA1|GIS1|HSP78|PNC1|TPK1|MSN4|TPK2|HST2", "Description": "Longevity regulating pathway - multiple species - Saccharomyces cerevisiae (budding yeast)", "_GeneInHitList": 294.0, "Category": "KEGG Pathway", "LogP": -4.130078175744296, "_RANK_": 1.0, "FirstInGroupByEnrichment": 1.0, "_LogP_MyList": -4.130078175744296, "_TotalGeneInLibrary": 5919.0, "_GeneInGO": 38.0, "shared_name": "101", "Gene": "sce04213", "_MEMBER_MyList": 1.0, "STDV_InGO": 1.0046682763914156, "_COUNT_001": 9.0, "SUID": 2742116, "id_original": "101", "selected": false, "Log_q_value_": -2.333153628949953, "canonicalName": "sce04213", "CategoryID": 24.0, "Z_score": 5.3272974182583095, "EvidenceCutoff": 0.0, "PARENT_GO": "", "FirstInGroupByLogP": 1.0, "GROUP_ID": 16.0, "name": "101", "BestEnrichmentInGroup": 4.768259935553168, "CLUSTER_LABEL": "Longevity regulating pathway - multiple species - Saccharomyces cerevisiae (budding yeast)", "GeneID": "850633|851259|851670|851845|852846|853275|853803|855898|856092", "GiniIndex": 0.0, "_InGO": 3.061224489795918, "_COUNT_001_PCT": 100.0}, "position": {"x": -173.5293947368708, "y": 942.7381671536091}, "selected": false}, {"data": {"id": "2742113", "Enrichment": 8.47690655209452, "BestLogPInGroup": -27.34010239473319, "_PATTERN_": "M1", "_GeneInGOAndHitList": 40.0, "Hits": "GLK1|QCR6|HXK1|COX12|ATP14|COX8|GPM2|ATP16|SDH4|ATP5|TIM11|ATP17|EMI2|QCR7|ATP1|ATP3|COX4|QCR9|QCR8|TDH1|ATP2|SDH1|SDH3|ATP7|ATP19|CYT1|PYK2|COX5B|ATP18|NDI1|COX7|COX5A|ATP15|ATP4|ATP20|QCR2|QCR10|CYC7|RIP1|MCO10", "Description": "ATP metabolic process", "_GeneInHitList": 294.0, "Category": "GO Biological Processes", "LogP": -27.34010239473319, "_RANK_": 1.0, "FirstInGroupByEnrichment": 1.0, "_LogP_MyList": -27.34010239473319, "_TotalGeneInLibrary": 5919.0, "_GeneInGO": 95.0, "shared_name": "100", "Gene": "GO:0046034", "_MEMBER_MyList": 1.0, "STDV_InGO": 1.9995204874649775, "_COUNT_001": 40.0, "SUID": 2742113, "id_original": "100", "selected": false, "Log_q_value_": -23.552004761352894, "canonicalName": "GO:0046034", "CategoryID": 19.0, "Z_score": 16.794727222905014, "EvidenceCutoff": 0.0, "PARENT_GO": "19_GO:0008152 metabolic process", "FirstInGroupByLogP": 1.0, "GROUP_ID": 1.0, "name": "100", "BestEnrichmentInGroup": 12.079591836734693, "CLUSTER_LABEL": "ATP metabolic process", "GeneID": "850317|850593|850614|850727|851002|851111|851541|851560|851758|851892|851922|851983|852128|852142|852177|852327|852688|853095|853273|853395|853585|853709|853716|853853|854077|854231|854529|854695|854893|854919|855298|855675|855857|856027|856131|856321|856390|856672|856689|1466480", "GiniIndex": 0.0, "_InGO": 13.605442176870747, "_COUNT_001_PCT": 100.0}, "position": {"x": 579.6306318134222, "y": 401.9086993801716}, "selected": false}, {"data": {"id": "2742110", "Enrichment": 15.099489795918368, "BestLogPInGroup": -15.616473896518219, "_PATTERN_": "M1", "_GeneInGOAndHitList": 15.0, "Hits": "ATP14|ATP16|ATP5|TIM11|ATP17|ATP1|ATP3|ATP2|ATP7|ATP19|ATP18|ATP15|ATP4|ATP20|MCO10", "Description": "ATP biosynthetic process", "_GeneInHitList": 294.0, "Category": "GO Biological Processes", "LogP": -15.616473896518219, "_RANK_": 1.0, "FirstInGroupByEnrichment": 1.0, "_LogP_MyList": -15.616473896518219, "_TotalGeneInLibrary": 5919.0, "_GeneInGO": 20.0, "shared_name": "99", "Gene": "GO:0006754", "_MEMBER_MyList": 1.0, "STDV_InGO": 1.2832956896252923, "_COUNT_001": 15.0, "SUID": 2742110, "id_original": "99", "selected": false, "Log_q_value_": -12.82837626313792, "canonicalName": "GO:0006754", "CategoryID": 19.0, "Z_score": 14.438732800724484, "EvidenceCutoff": 0.0, "PARENT_GO": "19_GO:0008152 metabolic process", "FirstInGroupByLogP": 1.0, "GROUP_ID": 2.0, "name": "99", "BestEnrichmentInGroup": 15.099489795918368, "CLUSTER_LABEL": "ATP biosynthetic process", "GeneID": "851002|851560|851892|851922|851983|852177|852327|853585|853853|854077|854893|855857|856027|856131|1466480", "GiniIndex": 0.0, "_InGO": 5.1020408163265305, "_COUNT_001_PCT": 100.0}, "position": {"x": 633.7032178729925, "y": 294.6183327305439}, "selected": false}, {"data": {"id": "2742107", "Enrichment": 5.033163265306122, "BestLogPInGroup": -3.769282700434612, "_PATTERN_": "M1", "_GeneInGOAndHitList": 4.0, "Hits": "PIB1|VID24|FYV10|UBC8", "Description": "negative regulation of small molecule metabolic process", "_GeneInHitList": 294.0, "Category": "GO Biological Processes", "LogP": -2.170626619105343, "_RANK_": 1.0, "FirstInGroupByEnrichment": 0.0, "_LogP_MyList": -2.170626619105343, "_TotalGeneInLibrary": 5919.0, "_GeneInGO": 16.0, "shared_name": "98", "Gene": "GO:0062014", "_MEMBER_MyList": 1.0, "STDV_InGO": 0.6756285589444189, "_COUNT_001": 4.0, "SUID": 2742107, "id_original": "98", "selected": false, "Log_q_value_": -0.7047482804589649, "canonicalName": "GO:0062014", "CategoryID": 19.0, "Z_score": 3.6929147726784044, "EvidenceCutoff": 0.0, "PARENT_GO": "19_GO:0048519 negative regulation of biological process", "FirstInGroupByLogP": 0.0, "GROUP_ID": 20.0, "name": "98", "BestEnrichmentInGroup": 7.320964749536177, "CLUSTER_LABEL": " ", "GeneID": "851908|852402|854710|856704", "GiniIndex": 0.0, "_InGO": 1.3605442176870748, "_COUNT_001_PCT": 100.0}, "position": {"x": -59.26187001763253, "y": 553.0139697659138}, "selected": false}, {"data": {"id": "2742104", "Enrichment": 5.033163265306122, "BestLogPInGroup": -3.769282700434612, "_PATTERN_": "M1", "_GeneInGOAndHitList": 4.0, "Hits": "PIB1|VID24|FYV10|UBC8", "Description": "regulation of gluconeogenesis", "_GeneInHitList": 294.0, "Category": "GO Biological Processes", "LogP": -2.170626619105343, "_RANK_": 1.0, "FirstInGroupByEnrichment": 0.0, "_LogP_MyList": -2.170626619105343, "_TotalGeneInLibrary": 5919.0, "_GeneInGO": 16.0, "shared_name": "97", "Gene": "GO:0006111", "_MEMBER_MyList": 1.0, "STDV_InGO": 0.6756285589444189, "_COUNT_001": 4.0, "SUID": 2742104, "id_original": "97", "selected": false, "Log_q_value_": -0.7047482804589649, "canonicalName": "GO:0006111", "CategoryID": 19.0, "Z_score": 3.6929147726784044, "EvidenceCutoff": 0.0, "PARENT_GO": "19_GO:0050789 regulation of biological process", "FirstInGroupByLogP": 0.0, "GROUP_ID": 20.0, "name": "97", "BestEnrichmentInGroup": 7.320964749536177, "CLUSTER_LABEL": " ", "GeneID": "851908|852402|854710|856704", "GiniIndex": 0.0, "_InGO": 1.3605442176870748, "_COUNT_001_PCT": 100.0}, "position": {"x": 77.24362314642997, "y": 590.9015735256794}, "selected": false}, {"data": {"id": "2742101", "Enrichment": 3.8966425279789343, "BestLogPInGroup": -3.769282700434612, "_PATTERN_": "M1", "_GeneInGOAndHitList": 6.0, "Hits": "PIB1|VID24|GAC1|FYV10|PCL7|UBC8", "Description": "regulation of carbohydrate biosynthetic process", "_GeneInHitList": 294.0, "Category": "GO Biological Processes", "LogP": -2.4358895593388183, "_RANK_": 1.0, "FirstInGroupByEnrichment": 0.0, "_LogP_MyList": -2.4358895593388183, "_TotalGeneInLibrary": 5919.0, "_GeneInGO": 31.0, "shared_name": "96", "Gene": "GO:0043255", "_MEMBER_MyList": 1.0, "STDV_InGO": 0.8246143220834374, "_COUNT_001": 6.0, "SUID": 2742101, "id_original": "96", "selected": false, "Log_q_value_": -0.9288252932062485, "canonicalName": "GO:0043255", "CategoryID": 19.0, "Z_score": 3.696508686588874, "EvidenceCutoff": 0.0, "PARENT_GO": "19_GO:0050789 regulation of biological process", "FirstInGroupByLogP": 0.0, "GROUP_ID": 20.0, "name": "96", "BestEnrichmentInGroup": 7.320964749536177, "CLUSTER_LABEL": " ", "GeneID": "851908|852402|854350|854710|854761|856704", "GiniIndex": 0.0, "_InGO": 2.0408163265306123, "_COUNT_001_PCT": 100.0}, "position": {"x": 51.44717539252372, "y": 638.6755908596638}, "selected": false}, {"data": {"id": "2742098", "Enrichment": 7.320964749536177, "BestLogPInGroup": -3.769282700434612, "_PATTERN_": "M1", "_GeneInGOAndHitList": 4.0, "Hits": "PIB1|VID24|FYV10|UBC8", "Description": "negative regulation of gluconeogenesis", "_GeneInHitList": 294.0, "Category": "GO Biological Processes", "LogP": -2.826596761418444, "_RANK_": 1.0, "FirstInGroupByEnrichment": 0.0, "_LogP_MyList": -2.826596761418444, "_TotalGeneInLibrary": 5919.0, "_GeneInGO": 11.0, "shared_name": "95", "Gene": "GO:0045721", "_MEMBER_MyList": 1.0, "STDV_InGO": 0.6756285589444189, "_COUNT_001": 4.0, "SUID": 2742098, "id_original": "95", "selected": false, "Log_q_value_": -1.2426191106940716, "canonicalName": "GO:0045721", "CategoryID": 19.0, "Z_score": 4.79688547500063, "EvidenceCutoff": 0.0, "PARENT_GO": "19_GO:0048519 negative regulation of biological process", "FirstInGroupByLogP": 0.0, "GROUP_ID": 20.0, "name": "95", "BestEnrichmentInGroup": 7.320964749536177, "CLUSTER_LABEL": " ", "GeneID": "851908|852402|854710|856704", "GiniIndex": 0.0, "_InGO": 1.3605442176870748, "_COUNT_001_PCT": 100.0}, "position": {"x": -5.381987205132532, "y": 642.9577258694294}, "selected": false}, {"data": {"id": "2742095", "Enrichment": 5.921368547418967, "BestLogPInGroup": -3.769282700434612, "_PATTERN_": "M1", "_GeneInGOAndHitList": 5.0, "Hits": "IGD1|PIB1|VID24|FYV10|UBC8", "Description": "negative regulation of carbohydrate metabolic process", "_GeneInHitList": 294.0, "Category": "GO Biological Processes", "LogP": -2.9573916182518474, "_RANK_": 1.0, "FirstInGroupByEnrichment": 0.0, "_LogP_MyList": -2.9573916182518474, "_TotalGeneInLibrary": 5919.0, "_GeneInGO": 17.0, "shared_name": "94", "Gene": "GO:0045912", "_MEMBER_MyList": 1.0, "STDV_InGO": 0.7540721961117447, "_COUNT_001": 5.0, "SUID": 2742095, "id_original": "94", "selected": false, "Log_q_value_": -1.345912500568209, "canonicalName": "GO:0045912", "CategoryID": 19.0, "Z_score": 4.64526644713336, "EvidenceCutoff": 0.0, "PARENT_GO": "19_GO:0048519 negative regulation of biological process", "FirstInGroupByLogP": 0.0, "GROUP_ID": 20.0, "name": "94", "BestEnrichmentInGroup": 7.320964749536177, "CLUSTER_LABEL": " ", "GeneID": "850572|851908|852402|854710|856704", "GiniIndex": 0.0, "_InGO": 1.7006802721088434, "_COUNT_001_PCT": 100.0}, "position": {"x": -51.69848500786691, "y": 609.1261523831013}, "selected": false}, {"data": {"id": "2742092", "Enrichment": 5.921368547418967, "BestLogPInGroup": -3.769282700434612, "_PATTERN_": "M1", "_GeneInGOAndHitList": 5.0, "Hits": "IGD1|PIB1|VID24|FYV10|UBC8", "Description": "negative regulation of cellular carbohydrate metabolic process", "_GeneInHitList": 294.0, "Category": "GO Biological Processes", "LogP": -2.9573916182518474, "_RANK_": 1.0, "FirstInGroupByEnrichment": 0.0, "_LogP_MyList": -2.9573916182518474, "_TotalGeneInLibrary": 5919.0, "_GeneInGO": 17.0, "shared_name": "93", "Gene": "GO:0010677", "_MEMBER_MyList": 1.0, "STDV_InGO": 0.7540721961117447, "_COUNT_001": 5.0, "SUID": 2742092, "id_original": "93", "selected": false, "Log_q_value_": -1.345912500568209, "canonicalName": "GO:0010677", "CategoryID": 19.0, "Z_score": 4.64526644713336, "EvidenceCutoff": 0.0, "PARENT_GO": "19_GO:0048519 negative regulation of biological process", "FirstInGroupByLogP": 0.0, "GROUP_ID": 20.0, "name": "93", "BestEnrichmentInGroup": 7.320964749536177, "CLUSTER_LABEL": " ", "GeneID": "850572|851908|852402|854710|856704", "GiniIndex": 0.0, "_InGO": 1.7006802721088434, "_COUNT_001_PCT": 100.0}, "position": {"x": 10.177247414008093, "y": 577.1806872952106}, "selected": false}, {"data": {"id": "2742089", "Enrichment": 4.129774986917844, "BestLogPInGroup": -3.769282700434612, "_PATTERN_": "M1", "_GeneInGOAndHitList": 8.0, "Hits": "IGD1|PIB1|VID24|GAC1|FYV10|PCL7|PKP1|UBC8", "Description": "regulation of cellular carbohydrate metabolic process", "_GeneInHitList": 294.0, "Category": "GO Biological Processes", "LogP": -3.269539399788446, "_RANK_": 1.0, "FirstInGroupByEnrichment": 0.0, "_LogP_MyList": -3.269539399788446, "_TotalGeneInLibrary": 5919.0, "_GeneInGO": 39.0, "shared_name": "92", "Gene": "GO:0010675", "_MEMBER_MyList": 1.0, "STDV_InGO": 0.9488706521462137, "_COUNT_001": 8.0, "SUID": 2742089, "id_original": "92", "selected": false, "Log_q_value_": -1.5954740397585803, "canonicalName": "GO:0010675", "CategoryID": 19.0, "Z_score": 4.482875091295112, "EvidenceCutoff": 0.0, "PARENT_GO": "19_GO:0050789 regulation of biological process", "FirstInGroupByLogP": 0.0, "GROUP_ID": 20.0, "name": "92", "BestEnrichmentInGroup": 7.320964749536177, "CLUSTER_LABEL": " ", "GeneID": "850572|851908|852402|854350|854710|854761|854769|856704", "GiniIndex": 0.0, "_InGO": 2.7210884353741496, "_COUNT_001_PCT": 100.0}, "position": {"x": 81.07888925971122, "y": 515.3899615872028}, "selected": false}, {"data": {"id": "2742086", "Enrichment": 4.601749271137026, "BestLogPInGroup": -3.769282700434612, "_PATTERN_": "M1", "_GeneInGOAndHitList": 8.0, "Hits": "IGD1|PIB1|VID24|GAC1|FYV10|PCL7|PKP1|UBC8", "Description": "regulation of glucose metabolic process", "_GeneInHitList": 294.0, "Category": "GO Biological Processes", "LogP": -3.611544410401986, "_RANK_": 1.0, "FirstInGroupByEnrichment": 0.0, "_LogP_MyList": -3.611544410401986, "_TotalGeneInLibrary": 5919.0, "_GeneInGO": 35.0, "shared_name": "91", "Gene": "GO:0010906", "_MEMBER_MyList": 1.0, "STDV_InGO": 0.9488706521462137, "_COUNT_001": 8.0, "SUID": 2742086, "id_original": "91", "selected": false, "Log_q_value_": -1.8916326387678504, "canonicalName": "GO:0010906", "CategoryID": 19.0, "Z_score": 4.885522855354363, "EvidenceCutoff": 0.0, "PARENT_GO": "19_GO:0050789 regulation of biological process", "FirstInGroupByLogP": 0.0, "GROUP_ID": 20.0, "name": "91", "BestEnrichmentInGroup": 7.320964749536177, "CLUSTER_LABEL": " ", "GeneID": "850572|851908|852402|854350|854710|854761|854769|856704", "GiniIndex": 0.0, "_InGO": 2.7210884353741496, "_COUNT_001_PCT": 100.0}, "position": {"x": 41.75012339057059, "y": 491.7186664211872}, "selected": false}, {"data": {"id": "2742083", "Enrichment": 7.549744897959185, "BestLogPInGroup": -3.915927067855309, "_PATTERN_": "M1", "_GeneInGOAndHitList": 3.0, "Hits": "NTH1|TPS1|STF2", "Description": "cellular response to water stimulus", "_GeneInHitList": 294.0, "Category": "GO Biological Processes", "LogP": -2.248956556748509, "_RANK_": 1.0, "FirstInGroupByEnrichment": 0.0, "_LogP_MyList": -2.248956556748509, "_TotalGeneInLibrary": 5919.0, "_GeneInGO": 8.0, "shared_name": "90", "Gene": "GO:0071462", "_MEMBER_MyList": 1.0, "STDV_InGO": 0.5861194403214197, "_COUNT_001": 3.0, "SUID": 2742083, "id_original": "90", "selected": false, "Log_q_value_": -0.7683549612814243, "canonicalName": "GO:0071462", "CategoryID": 19.0, "Z_score": 4.2377836522616095, "EvidenceCutoff": 0.0, "PARENT_GO": "19_GO:0050896 response to stimulus", "FirstInGroupByLogP": 0.0, "GROUP_ID": 19.0, "name": "90", "BestEnrichmentInGroup": 20.13265306122449, "CLUSTER_LABEL": " ", "GeneID": "851564|852423|852891", "GiniIndex": 0.0, "_InGO": 1.0204081632653061, "_COUNT_001_PCT": 100.0}, "position": {"x": -174.8902574688044, "y": 225.36776767851143}, "selected": false}, {"data": {"id": "2742080", "Enrichment": 7.549744897959185, "BestLogPInGroup": -3.915927067855309, "_PATTERN_": "M1", "_GeneInGOAndHitList": 3.0, "Hits": "NTH1|TPS1|STF2", "Description": "response to water", "_GeneInHitList": 294.0, "Category": "GO Biological Processes", "LogP": -2.248956556748509, "_RANK_": 1.0, "FirstInGroupByEnrichment": 0.0, "_LogP_MyList": -2.248956556748509, "_TotalGeneInLibrary": 5919.0, "_GeneInGO": 8.0, "shared_name": "89", "Gene": "GO:0009415", "_MEMBER_MyList": 1.0, "STDV_InGO": 0.5861194403214197, "_COUNT_001": 3.0, "SUID": 2742080, "id_original": "89", "selected": false, "Log_q_value_": -0.7683549612814243, "canonicalName": "GO:0009415", "CategoryID": 19.0, "Z_score": 4.2377836522616095, "EvidenceCutoff": 0.0, "PARENT_GO": "19_GO:0050896 response to stimulus", "FirstInGroupByLogP": 0.0, "GROUP_ID": 19.0, "name": "89", "BestEnrichmentInGroup": 20.13265306122449, "CLUSTER_LABEL": " ", "GeneID": "851564|852423|852891", "GiniIndex": 0.0, "_InGO": 1.0204081632653061, "_COUNT_001_PCT": 100.0}, "position": {"x": -93.0974107891169, "y": 293.18283449293403}, "selected": false}, {"data": {"id": "2742077", "Enrichment": 10.066326530612246, "BestLogPInGroup": -3.915927067855309, "_PATTERN_": "M1", "_GeneInGOAndHitList": 3.0, "Hits": "NTH1|TPS1|STF2", "Description": "cellular response to water deprivation", "_GeneInHitList": 294.0, "Category": "GO Biological Processes", "LogP": -2.663750246918336, "_RANK_": 1.0, "FirstInGroupByEnrichment": 0.0, "_LogP_MyList": -2.663750246918336, "_TotalGeneInLibrary": 5919.0, "_GeneInGO": 6.0, "shared_name": "88", "Gene": "GO:0042631", "_MEMBER_MyList": 1.0, "STDV_InGO": 0.5861194403214197, "_COUNT_001": 3.0, "SUID": 2742077, "id_original": "88", "selected": false, "Log_q_value_": -1.113698716666834, "canonicalName": "GO:0042631", "CategoryID": 19.0, "Z_score": 5.079289005464239, "EvidenceCutoff": 0.0, "PARENT_GO": "19_GO:0050896 response to stimulus", "FirstInGroupByLogP": 0.0, "GROUP_ID": 19.0, "name": "88", "BestEnrichmentInGroup": 20.13265306122449, "CLUSTER_LABEL": " ", "GeneID": "851564|852423|852891", "GiniIndex": 0.0, "_InGO": 1.0204081632653061, "_COUNT_001_PCT": 100.0}, "position": {"x": -107.45550405083566, "y": 150.18079410673408}, "selected": false}, {"data": {"id": "2742074", "Enrichment": 10.066326530612246, "BestLogPInGroup": -3.915927067855309, "_PATTERN_": "M1", "_GeneInGOAndHitList": 3.0, "Hits": "NTH1|TPS1|STF2", "Description": "response to water deprivation", "_GeneInHitList": 294.0, "Category": "GO Biological Processes", "LogP": -2.663750246918336, "_RANK_": 1.0, "FirstInGroupByEnrichment": 0.0, "_LogP_MyList": -2.663750246918336, "_TotalGeneInLibrary": 5919.0, "_GeneInGO": 6.0, "shared_name": "87", "Gene": "GO:0009414", "_MEMBER_MyList": 1.0, "STDV_InGO": 0.5861194403214197, "_COUNT_001": 3.0, "SUID": 2742074, "id_original": "87", "selected": false, "Log_q_value_": -1.113698716666834, "canonicalName": "GO:0009414", "CategoryID": 19.0, "Z_score": 5.079289005464239, "EvidenceCutoff": 0.0, "PARENT_GO": "19_GO:0050896 response to stimulus", "FirstInGroupByLogP": 0.0, "GROUP_ID": 19.0, "name": "87", "BestEnrichmentInGroup": 20.13265306122449, "CLUSTER_LABEL": " ", "GeneID": "851564|852423|852891", "GiniIndex": 0.0, "_InGO": 1.0204081632653061, "_COUNT_001_PCT": 100.0}, "position": {"x": -144.35845815239816, "y": 279.3973911869648}, "selected": false}, {"data": {"id": "2742071", "Enrichment": 20.13265306122449, "BestLogPInGroup": -3.915927067855309, "_PATTERN_": "M1", "_GeneInGOAndHitList": 3.0, "Hits": "NTH1|TPS1|STF2", "Description": "cellular response to desiccation", "_GeneInHitList": 294.0, "Category": "GO Biological Processes", "LogP": -3.915927067855309, "_RANK_": 1.0, "FirstInGroupByEnrichment": 0.0, "_LogP_MyList": -3.915927067855309, "_TotalGeneInLibrary": 5919.0, "_GeneInGO": 3.0, "shared_name": "86", "Gene": "GO:0071465", "_MEMBER_MyList": 1.0, "STDV_InGO": 0.5861194403214197, "_COUNT_001": 3.0, "SUID": 2742071, "id_original": "86", "selected": false, "Log_q_value_": -2.165255932415635, "canonicalName": "GO:0071465", "CategoryID": 19.0, "Z_score": 7.577424595277936, "EvidenceCutoff": 0.0, "PARENT_GO": "19_GO:0050896 response to stimulus", "FirstInGroupByLogP": 0.0, "GROUP_ID": 19.0, "name": "86", "BestEnrichmentInGroup": 20.13265306122449, "CLUSTER_LABEL": " ", "GeneID": "851564|852423|852891", "GiniIndex": 0.0, "_InGO": 1.0204081632653061, "_COUNT_001_PCT": 100.0}, "position": {"x": -156.32733022271066, "y": 169.07424198271065}, "selected": false}, {"data": {"id": "2742068", "Enrichment": 6.194662480376765, "BestLogPInGroup": -4.082697487204731, "_PATTERN_": "M1", "_GeneInGOAndHitList": 4.0, "Hits": "TPK1|PLB1|TPK2|GPA2", "Description": "G alpha (i) signalling events", "_GeneInHitList": 294.0, "Category": "Reactome Gene Sets", "LogP": -2.5251297576587057, "_RANK_": 1.0, "FirstInGroupByEnrichment": 0.0, "_LogP_MyList": -2.5251297576587057, "_TotalGeneInLibrary": 5919.0, "_GeneInGO": 13.0, "shared_name": "85", "Gene": "R-SCE-418594", "_MEMBER_MyList": 1.0, "STDV_InGO": 0.6756285589444189, "_COUNT_001": 4.0, "SUID": 2742068, "id_original": "85", "selected": false, "Log_q_value_": -0.9994832140088379, "canonicalName": "R-SCE-418594", "CategoryID": 6.0, "Z_score": 4.286296127783706, "EvidenceCutoff": 0.0, "PARENT_GO": "", "FirstInGroupByLogP": 0.0, "GROUP_ID": 17.0, "name": "85", "BestEnrichmentInGroup": 13.421768707482991, "CLUSTER_LABEL": " ", "GeneID": "853275|855020|855898|856741", "GiniIndex": 0.0, "_InGO": 1.3605442176870748, "_COUNT_001_PCT": 100.0}, "position": {"x": -66.09953748284249, "y": 742.9577258694294}, "selected": false}, {"data": {"id": "2742065", "Enrichment": 6.194662480376765, "BestLogPInGroup": -4.082697487204731, "_PATTERN_": "M1", "_GeneInGOAndHitList": 4.0, "Hits": "TPK1|PLB1|TPK2|GPA2", "Description": "PLC beta mediated events", "_GeneInHitList": 294.0, "Category": "Reactome Gene Sets", "LogP": -2.5251297576587057, "_RANK_": 1.0, "FirstInGroupByEnrichment": 0.0, "_LogP_MyList": -2.5251297576587057, "_TotalGeneInLibrary": 5919.0, "_GeneInGO": 13.0, "shared_name": "84", "Gene": "R-SCE-112043", "_MEMBER_MyList": 1.0, "STDV_InGO": 0.6756285589444189, "_COUNT_001": 4.0, "SUID": 2742065, "id_original": "84", "selected": false, "Log_q_value_": -0.9994832140088379, "canonicalName": "R-SCE-112043", "CategoryID": 6.0, "Z_score": 4.286296127783706, "EvidenceCutoff": 0.0, "PARENT_GO": "", "FirstInGroupByLogP": 0.0, "GROUP_ID": 17.0, "name": "84", "BestEnrichmentInGroup": 13.421768707482991, "CLUSTER_LABEL": " ", "GeneID": "853275|855020|855898|856741", "GiniIndex": 0.0, "_InGO": 1.3605442176870748, "_COUNT_001_PCT": 100.0}, "position": {"x": -25.13868581353097, "y": 818.966652261031}, "selected": false}, {"data": {"id": "2742062", "Enrichment": 6.194662480376765, "BestLogPInGroup": -4.082697487204731, "_PATTERN_": "M1", "_GeneInGOAndHitList": 4.0, "Hits": "TPK1|PLB1|TPK2|GPA2", "Description": "G-protein mediated events", "_GeneInHitList": 294.0, "Category": "Reactome Gene Sets", "LogP": -2.5251297576587057, "_RANK_": 1.0, "FirstInGroupByEnrichment": 0.0, "_LogP_MyList": -2.5251297576587057, "_TotalGeneInLibrary": 5919.0, "_GeneInGO": 13.0, "shared_name": "83", "Gene": "R-SCE-112040", "_MEMBER_MyList": 1.0, "STDV_InGO": 0.6756285589444189, "_COUNT_001": 4.0, "SUID": 2742062, "id_original": "83", "selected": false, "Log_q_value_": -0.9994832140088379, "canonicalName": "R-SCE-112040", "CategoryID": 6.0, "Z_score": 4.286296127783706, "EvidenceCutoff": 0.0, "PARENT_GO": "", "FirstInGroupByLogP": 0.0, "GROUP_ID": 17.0, "name": "83", "BestEnrichmentInGroup": 13.421768707482991, "CLUSTER_LABEL": " ", "GeneID": "853275|855020|855898|856741", "GiniIndex": 0.0, "_InGO": 1.3605442176870748, "_COUNT_001_PCT": 100.0}, "position": {"x": -146.4212099224177, "y": 752.7405017483356}, "selected": false}, {"data": {"id": "2742059", "Enrichment": 6.194662480376765, "BestLogPInGroup": -4.082697487204731, "_PATTERN_": "M1", "_GeneInGOAndHitList": 4.0, "Hits": "TPK1|PLB1|TPK2|GPA2", "Description": "Opioid Signalling", "_GeneInHitList": 294.0, "Category": "Reactome Gene Sets", "LogP": -2.5251297576587057, "_RANK_": 1.0, "FirstInGroupByEnrichment": 0.0, "_LogP_MyList": -2.5251297576587057, "_TotalGeneInLibrary": 5919.0, "_GeneInGO": 13.0, "shared_name": "82", "Gene": "R-SCE-111885", "_MEMBER_MyList": 1.0, "STDV_InGO": 0.6756285589444189, "_COUNT_001": 4.0, "SUID": 2742059, "id_original": "82", "selected": false, "Log_q_value_": -0.9994832140088379, "canonicalName": "R-SCE-111885", "CategoryID": 6.0, "Z_score": 4.286296127783706, "EvidenceCutoff": 0.0, "PARENT_GO": "", "FirstInGroupByLogP": 0.0, "GROUP_ID": 17.0, "name": "82", "BestEnrichmentInGroup": 13.421768707482991, "CLUSTER_LABEL": " ", "GeneID": "853275|855020|855898|856741", "GiniIndex": 0.0, "_InGO": 1.3605442176870748, "_COUNT_001_PCT": 100.0}, "position": {"x": -89.29546414910226, "y": 800.7667773831013}, "selected": false}, {"data": {"id": "2742056", "Enrichment": 10.066326530612246, "BestLogPInGroup": -4.082697487204731, "_PATTERN_": "M1", "_GeneInGOAndHitList": 3.0, "Hits": "AQY3|TPK1|TPK2", "Description": "Aquaporin-mediated transport", "_GeneInHitList": 294.0, "Category": "Reactome Gene Sets", "LogP": -2.663750246918336, "_RANK_": 1.0, "FirstInGroupByEnrichment": 0.0, "_LogP_MyList": -2.663750246918336, "_TotalGeneInLibrary": 5919.0, "_GeneInGO": 6.0, "shared_name": "81", "Gene": "R-SCE-445717", "_MEMBER_MyList": 1.0, "STDV_InGO": 0.5861194403214197, "_COUNT_001": 3.0, "SUID": 2742056, "id_original": "81", "selected": false, "Log_q_value_": -1.113698716666834, "canonicalName": "R-SCE-445717", "CategoryID": 6.0, "Z_score": 5.079289005464239, "EvidenceCutoff": 0.0, "PARENT_GO": "", "FirstInGroupByLogP": 0.0, "GROUP_ID": 17.0, "name": "81", "BestEnrichmentInGroup": 13.421768707482991, "CLUSTER_LABEL": " ", "GeneID": "850490|853275|855898", "GiniIndex": 0.0, "_InGO": 1.0204081632653061, "_COUNT_001_PCT": 100.0}, "position": {"x": -174.8902574688044, "y": 821.9232715237263}, "selected": false}, {"data": {"id": "2742053", "Enrichment": 10.066326530612246, "BestLogPInGroup": -4.082697487204731, "_PATTERN_": "M1", "_GeneInGOAndHitList": 3.0, "Hits": "AQY3|TPK1|TPK2", "Description": "Vasopressin regulates renal water homeostasis via Aquaporins", "_GeneInHitList": 294.0, "Category": "Reactome Gene Sets", "LogP": -2.663750246918336, "_RANK_": 1.0, "FirstInGroupByEnrichment": 0.0, "_LogP_MyList": -2.663750246918336, "_TotalGeneInLibrary": 5919.0, "_GeneInGO": 6.0, "shared_name": "80", "Gene": "R-SCE-432040", "_MEMBER_MyList": 1.0, "STDV_InGO": 0.5861194403214197, "_COUNT_001": 3.0, "SUID": 2742053, "id_original": "80", "selected": false, "Log_q_value_": -1.113698716666834, "canonicalName": "R-SCE-432040", "CategoryID": 6.0, "Z_score": 5.079289005464239, "EvidenceCutoff": 0.0, "PARENT_GO": "", "FirstInGroupByLogP": 0.0, "GROUP_ID": 17.0, "name": "80", "BestEnrichmentInGroup": 13.421768707482991, "CLUSTER_LABEL": " ", "GeneID": "850490|853275|855898", "GiniIndex": 0.0, "_InGO": 1.0204081632653061, "_COUNT_001_PCT": 100.0}, "position": {"x": -74.70549832878976, "y": 881.0023883450153}, "selected": false}, {"data": {"id": "2742050", "Enrichment": 2.353167240922343, "BestLogPInGroup": -4.197480086121784, "_PATTERN_": "M1", "_GeneInGOAndHitList": 18.0, "Hits": "HSP12|HSP104|GPD1|NTH1|NRG1|TPS2|HSP42|HSP78|TPS1|STF2|YAK1|DDR2|GAC1|PHR1|YKU70|HSP82|LSP1|GPP2", "Description": "response to abiotic stimulus", "_GeneInHitList": 294.0, "Category": "GO Biological Processes", "LogP": -3.2458039961180907, "_RANK_": 1.0, "FirstInGroupByEnrichment": 0.0, "_LogP_MyList": -3.2458039961180907, "_TotalGeneInLibrary": 5919.0, "_GeneInGO": 154.0, "shared_name": "79", "Gene": "GO:0009628", "_MEMBER_MyList": 1.0, "STDV_InGO": 1.3982016102952535, "_COUNT_001": 18.0, "SUID": 2742050, "id_original": "79", "selected": false, "Log_q_value_": -1.578280293943643, "canonicalName": "GO:0009628", "CategoryID": 19.0, "Z_score": 3.8896637740928712, "EvidenceCutoff": 0.0, "PARENT_GO": "19_GO:0050896 response to stimulus", "FirstInGroupByLogP": 0.0, "GROUP_ID": 15.0, "name": "79", "BestEnrichmentInGroup": 4.129774986917844, "CLUSTER_LABEL": " ", "GeneID": "850532|850633|851539|851564|851613|851646|851751|851845|852423|852891|853300|854104|854350|854568|855328|855836|856103|856791", "GiniIndex": 0.0, "_InGO": 6.122448979591836, "_COUNT_001_PCT": 100.0}, "position": {"x": -163.5186754375544, "y": -44.58368648408623}, "selected": false}, {"data": {"id": "2742047", "Enrichment": 4.129774986917844, "BestLogPInGroup": -4.197480086121784, "_PATTERN_": "M1", "_GeneInGOAndHitList": 8.0, "Hits": "HSP12|HSP104|TPS2|HSP78|TPS1|YAK1|DDR2|HSP82", "Description": "cellular response to heat", "_GeneInHitList": 294.0, "Category": "GO Biological Processes", "LogP": -3.269539399788446, "_RANK_": 1.0, "FirstInGroupByEnrichment": 0.0, "_LogP_MyList": -3.269539399788446, "_TotalGeneInLibrary": 5919.0, "_GeneInGO": 39.0, "shared_name": "78", "Gene": "GO:0034605", "_MEMBER_MyList": 1.0, "STDV_InGO": 0.9488706521462137, "_COUNT_001": 8.0, "SUID": 2742047, "id_original": "78", "selected": false, "Log_q_value_": -1.5954740397585803, "canonicalName": "GO:0034605", "CategoryID": 19.0, "Z_score": 4.482875091295112, "EvidenceCutoff": 0.0, "PARENT_GO": "19_GO:0050896 response to stimulus", "FirstInGroupByLogP": 0.0, "GROUP_ID": 15.0, "name": "78", "BestEnrichmentInGroup": 4.129774986917844, "CLUSTER_LABEL": " ", "GeneID": "850532|850633|851646|851845|852423|853300|854104|855836", "GiniIndex": 0.0, "_InGO": 2.7210884353741496, "_COUNT_001_PCT": 100.0}, "position": {"x": -86.29592763482003, "y": 94.69089542509346}, "selected": false}, {"data": {"id": "2742044", "Enrichment": 3.8182617874736104, "BestLogPInGroup": -4.197480086121784, "_PATTERN_": "M1", "_GeneInGOAndHitList": 11.0, "Hits": "HSP12|HSP104|TPS2|HSP42|HSP78|TPS1|YAK1|DDR2|GAC1|HSP82|LSP1", "Description": "response to temperature stimulus", "_GeneInHitList": 294.0, "Category": "GO Biological Processes", "LogP": -3.9759715682139274, "_RANK_": 1.0, "FirstInGroupByEnrichment": 0.0, "_LogP_MyList": -3.9759715682139274, "_TotalGeneInLibrary": 5919.0, "_GeneInGO": 58.0, "shared_name": "77", "Gene": "GO:0009266", "_MEMBER_MyList": 1.0, "STDV_InGO": 1.1067985072269073, "_COUNT_001": 11.0, "SUID": 2742044, "id_original": "77", "selected": false, "Log_q_value_": -2.2007111595388023, "canonicalName": "GO:0009266", "CategoryID": 19.0, "Z_score": 4.9307087536160585, "EvidenceCutoff": 0.0, "PARENT_GO": "19_GO:0050896 response to stimulus", "FirstInGroupByLogP": 0.0, "GROUP_ID": 15.0, "name": "77", "BestEnrichmentInGroup": 4.129774986917844, "CLUSTER_LABEL": " ", "GeneID": "850532|850633|851646|851751|851845|852423|853300|854104|854350|855836|856103", "GiniIndex": 0.0, "_InGO": 3.741496598639456, "_COUNT_001_PCT": 100.0}, "position": {"x": -62.30328237114816, "y": -7.675971640336229}, "selected": false}, {"data": {"id": "2742041", "Enrichment": 4.3141399416909625, "BestLogPInGroup": -4.381277688016449, "_PATTERN_": "M1", "_GeneInGOAndHitList": 6.0, "Hits": "GLK1|HXK1|GPM2|EMI2|TDH1|PYK2", "Description": "ADP metabolic process", "_GeneInHitList": 294.0, "Category": "GO Biological Processes", "LogP": -2.6724439906125683, "_RANK_": 1.0, "FirstInGroupByEnrichment": 0.0, "_LogP_MyList": -2.6724439906125683, "_TotalGeneInLibrary": 5919.0, "_GeneInGO": 28.0, "shared_name": "76", "Gene": "GO:0046031", "_MEMBER_MyList": 1.0, "STDV_InGO": 0.8246143220834374, "_COUNT_001": 6.0, "SUID": 2742041, "id_original": "76", "selected": false, "Log_q_value_": -1.113698716666834, "canonicalName": "GO:0046031", "CategoryID": 19.0, "Z_score": 4.01841909960245, "EvidenceCutoff": 0.0, "PARENT_GO": "19_GO:0008152 metabolic process", "FirstInGroupByLogP": 0.0, "GROUP_ID": 14.0, "name": "76", "BestEnrichmentInGroup": 12.079591836734696, "CLUSTER_LABEL": " ", "GeneID": "850317|850614|851541|852128|853395|854529", "GiniIndex": 0.0, "_InGO": 2.0408163265306123, "_COUNT_001_PCT": 100.0}, "position": {"x": 378.65318621100516, "y": 147.42099796415596}, "selected": false}, {"data": {"id": "2742038", "Enrichment": 4.3141399416909625, "BestLogPInGroup": -4.381277688016449, "_PATTERN_": "M1", "_GeneInGOAndHitList": 6.0, "Hits": "GLK1|HXK1|GPM2|EMI2|TDH1|PYK2", "Description": "nucleoside diphosphate phosphorylation", "_GeneInHitList": 294.0, "Category": "GO Biological Processes", "LogP": -2.6724439906125683, "_RANK_": 1.0, "FirstInGroupByEnrichment": 0.0, "_LogP_MyList": -2.6724439906125683, "_TotalGeneInLibrary": 5919.0, "_GeneInGO": 28.0, "shared_name": "75", "Gene": "GO:0006165", "_MEMBER_MyList": 1.0, "STDV_InGO": 0.8246143220834374, "_COUNT_001": 6.0, "SUID": 2742038, "id_original": "75", "selected": false, "Log_q_value_": -1.113698716666834, "canonicalName": "GO:0006165", "CategoryID": 19.0, "Z_score": 4.01841909960245, "EvidenceCutoff": 0.0, "PARENT_GO": "19_GO:0008152 metabolic process", "FirstInGroupByLogP": 0.0, "GROUP_ID": 14.0, "name": "75", "BestEnrichmentInGroup": 12.079591836734696, "CLUSTER_LABEL": " ", "GeneID": "850317|850614|851541|852128|853395|854529", "GiniIndex": 0.0, "_InGO": 2.0408163265306123, "_COUNT_001_PCT": 100.0}, "position": {"x": 341.4157041400823, "y": 163.9888309109333}, "selected": false}, {"data": {"id": "2742035", "Enrichment": 4.645996860282575, "BestLogPInGroup": -4.381277688016449, "_PATTERN_": "M1", "_GeneInGOAndHitList": 6.0, "Hits": "GLK1|HXK1|GPM2|EMI2|TDH1|PYK2", "Description": "ATP generation from ADP", "_GeneInHitList": 294.0, "Category": "GO Biological Processes", "LogP": -2.849936219347253, "_RANK_": 1.0, "FirstInGroupByEnrichment": 0.0, "_LogP_MyList": -2.849936219347253, "_TotalGeneInLibrary": 5919.0, "_GeneInGO": 26.0, "shared_name": "74", "Gene": "GO:0006757", "_MEMBER_MyList": 1.0, "STDV_InGO": 0.8246143220834374, "_COUNT_001": 6.0, "SUID": 2742035, "id_original": "74", "selected": false, "Log_q_value_": -1.2577382383761893, "canonicalName": "GO:0006757", "CategoryID": 19.0, "Z_score": 4.259264651322197, "EvidenceCutoff": 0.0, "PARENT_GO": "19_GO:0008152 metabolic process", "FirstInGroupByLogP": 0.0, "GROUP_ID": 14.0, "name": "74", "BestEnrichmentInGroup": 12.079591836734696, "CLUSTER_LABEL": " ", "GeneID": "850317|850614|851541|852128|853395|854529", "GiniIndex": 0.0, "_InGO": 2.0408163265306123, "_COUNT_001_PCT": 100.0}, "position": {"x": 312.1487592548284, "y": 192.2491305935505}, "selected": false}, {"data": {"id": "2742032", "Enrichment": 4.645996860282575, "BestLogPInGroup": -4.381277688016449, "_PATTERN_": "M1", "_GeneInGOAndHitList": 6.0, "Hits": "GLK1|HXK1|GPM2|EMI2|TDH1|PYK2", "Description": "glycolytic process", "_GeneInHitList": 294.0, "Category": "GO Biological Processes", "LogP": -2.849936219347253, "_RANK_": 1.0, "FirstInGroupByEnrichment": 0.0, "_LogP_MyList": -2.849936219347253, "_TotalGeneInLibrary": 5919.0, "_GeneInGO": 26.0, "shared_name": "73", "Gene": "GO:0006096", "_MEMBER_MyList": 1.0, "STDV_InGO": 0.8246143220834374, "_COUNT_001": 6.0, "SUID": 2742032, "id_original": "73", "selected": false, "Log_q_value_": -1.2577382383761893, "canonicalName": "GO:0006096", "CategoryID": 19.0, "Z_score": 4.259264651322197, "EvidenceCutoff": 0.0, "PARENT_GO": "19_GO:0008152 metabolic process", "FirstInGroupByLogP": 0.0, "GROUP_ID": 14.0, "name": "73", "BestEnrichmentInGroup": 12.079591836734696, "CLUSTER_LABEL": " ", "GeneID": "850317|850614|851541|852128|853395|854529", "GiniIndex": 0.0, "_InGO": 2.0408163265306123, "_COUNT_001_PCT": 100.0}, "position": {"x": 388.4711173862371, "y": 183.39684330107002}, "selected": false}, {"data": {"id": "2742029", "Enrichment": 12.079591836734696, "BestLogPInGroup": -4.381277688016449, "_PATTERN_": "M1", "_GeneInGOAndHitList": 3.0, "Hits": "GLK1|HXK1|EMI2", "Description": "Regulation of Glucokinase by Glucokinase Regulatory Protein", "_GeneInHitList": 294.0, "Category": "Reactome Gene Sets", "LogP": -2.9485363609854973, "_RANK_": 1.0, "FirstInGroupByEnrichment": 0.0, "_LogP_MyList": -2.9485363609854973, "_TotalGeneInLibrary": 5919.0, "_GeneInGO": 5.0, "shared_name": "72", "Gene": "R-SCE-170822", "_MEMBER_MyList": 1.0, "STDV_InGO": 0.5861194403214197, "_COUNT_001": 3.0, "SUID": 2742029, "id_original": "72", "selected": false, "Log_q_value_": -1.345912500568209, "canonicalName": "R-SCE-170822", "CategoryID": 6.0, "Z_score": 5.665888007043756, "EvidenceCutoff": 0.0, "PARENT_GO": "", "FirstInGroupByLogP": 0.0, "GROUP_ID": 14.0, "name": "72", "BestEnrichmentInGroup": 12.079591836734696, "CLUSTER_LABEL": " ", "GeneID": "850317|850614|852128", "GiniIndex": 0.0, "_InGO": 1.0204081632653061, "_COUNT_001_PCT": 100.0}, "position": {"x": 269.70346201361747, "y": 100.7058948147419}, "selected": false}, {"data": {"id": "2742026", "Enrichment": 5.033163265306123, "BestLogPInGroup": -4.381277688016449, "_PATTERN_": "M1", "_GeneInGOAndHitList": 6.0, "Hits": "GLK1|HXK1|YLR345W|EMI2|TDH1|PYK2", "Description": "Glycolysis", "_GeneInHitList": 294.0, "Category": "Reactome Gene Sets", "LogP": -3.046608573232743, "_RANK_": 1.0, "FirstInGroupByEnrichment": 0.0, "_LogP_MyList": -3.046608573232743, "_TotalGeneInLibrary": 5919.0, "_GeneInGO": 24.0, "shared_name": "71", "Gene": "R-SCE-70171", "_MEMBER_MyList": 1.0, "STDV_InGO": 0.8246143220834374, "_COUNT_001": 6.0, "SUID": 2742026, "id_original": "71", "selected": false, "Log_q_value_": -1.4015257401065409, "canonicalName": "R-SCE-70171", "CategoryID": 6.0, "Z_score": 4.52594634689179, "EvidenceCutoff": 0.0, "PARENT_GO": "", "FirstInGroupByLogP": 0.0, "GROUP_ID": 14.0, "name": "71", "BestEnrichmentInGroup": 12.079591836734696, "CLUSTER_LABEL": " ", "GeneID": "850317|850614|851059|852128|853395|854529", "GiniIndex": 0.0, "_InGO": 2.0408163265306123, "_COUNT_001_PCT": 100.0}, "position": {"x": 358.75501001776786, "y": 210.94569431425361}, "selected": false}, {"data": {"id": "2742023", "Enrichment": 4.404017857142857, "BestLogPInGroup": -4.381277688016449, "_PATTERN_": "M1", "_GeneInGOAndHitList": 7.0, "Hits": "GLK1|HXK1|YLR345W|EMI2|TDH1|MDH1|PYK2", "Description": "Glucose metabolism", "_GeneInHitList": 294.0, "Category": "Reactome Gene Sets", "LogP": -3.097716529945372, "_RANK_": 1.0, "FirstInGroupByEnrichment": 0.0, "_LogP_MyList": -3.097716529945372, "_TotalGeneInLibrary": 5919.0, "_GeneInGO": 32.0, "shared_name": "70", "Gene": "R-SCE-70326", "_MEMBER_MyList": 1.0, "STDV_InGO": 0.8891375930475126, "_COUNT_001": 7.0, "SUID": 2742023, "id_original": "70", "selected": false, "Log_q_value_": -1.4494979829663113, "canonicalName": "R-SCE-70326", "CategoryID": 6.0, "Z_score": 4.413871476302858, "EvidenceCutoff": 0.0, "PARENT_GO": "", "FirstInGroupByLogP": 0.0, "GROUP_ID": 14.0, "name": "70", "BestEnrichmentInGroup": 12.079591836734696, "CLUSTER_LABEL": " ", "GeneID": "850317|850614|851059|852128|853395|853777|854529", "GiniIndex": 0.0, "_InGO": 2.380952380952381, "_COUNT_001_PCT": 100.0}, "position": {"x": 295.08786142767997, "y": 241.1922458279255}, "selected": false}, {"data": {"id": "2742020", "Enrichment": 6.710884353741496, "BestLogPInGroup": -4.381277688016449, "_PATTERN_": "M1", "_GeneInGOAndHitList": 5.0, "Hits": "GLK1|HXK1|YLR345W|EMI2|XKS1", "Description": "carbohydrate phosphorylation", "_GeneInHitList": 294.0, "Category": "GO Biological Processes", "LogP": -3.235709728352353, "_RANK_": 1.0, "FirstInGroupByEnrichment": 0.0, "_LogP_MyList": -3.235709728352353, "_TotalGeneInLibrary": 5919.0, "_GeneInGO": 15.0, "shared_name": "69", "Gene": "GO:0046835", "_MEMBER_MyList": 1.0, "STDV_InGO": 0.7540721961117447, "_COUNT_001": 5.0, "SUID": 2742020, "id_original": "69", "selected": false, "Log_q_value_": -1.5714637359391415, "canonicalName": "GO:0046835", "CategoryID": 19.0, "Z_score": 5.062624343261454, "EvidenceCutoff": 0.0, "PARENT_GO": "19_GO:0008152 metabolic process", "FirstInGroupByLogP": 0.0, "GROUP_ID": 14.0, "name": "69", "BestEnrichmentInGroup": 12.079591836734696, "CLUSTER_LABEL": " ", "GeneID": "850317|850614|851059|852128|853108", "GiniIndex": 0.0, "_InGO": 1.7006802721088434, "_COUNT_001_PCT": 100.0}, "position": {"x": 321.3539060443792, "y": 110.41533695341377}, "selected": false}, {"data": {"id": "2742017", "Enrichment": 3.6604823747680886, "BestLogPInGroup": -4.381277688016449, "_PATTERN_": "M1", "_GeneInGOAndHitList": 10.0, "Hits": "GLK1|HXK1|GPM2|EMI2|TDH1|PYK2|ALD4|PGM2|HFD1|ALD3", "Description": "Glycolysis / Gluconeogenesis - Saccharomyces cerevisiae (budding yeast)", "_GeneInHitList": 294.0, "Category": "KEGG Pathway", "LogP": -3.504651605479765, "_RANK_": 1.0, "FirstInGroupByEnrichment": 0.0, "_LogP_MyList": -3.504651605479765, "_TotalGeneInLibrary": 5919.0, "_GeneInGO": 55.0, "shared_name": "68", "Gene": "sce00010", "_MEMBER_MyList": 1.0, "STDV_InGO": 1.057153797996807, "_COUNT_001": 10.0, "SUID": 2742017, "id_original": "68", "selected": false, "Log_q_value_": -1.8029138027742158, "canonicalName": "sce00010", "CategoryID": 24.0, "Z_score": 4.531527629473386, "EvidenceCutoff": 0.0, "PARENT_GO": "", "FirstInGroupByLogP": 0.0, "GROUP_ID": 14.0, "name": "68", "BestEnrichmentInGroup": 12.079591836734696, "CLUSTER_LABEL": " ", "GeneID": "850317|850614|851541|852128|853395|854529|854556|855131|855137|855205", "GiniIndex": 0.0, "_InGO": 3.401360544217687, "_COUNT_001_PCT": 100.0}, "position": {"x": 346.3018049091253, "y": 246.3118518459919}, "selected": false}, {"data": {"id": "2742014", "Enrichment": 5.752186588921283, "BestLogPInGroup": -4.436199066039166, "_PATTERN_": "M1", "_GeneInGOAndHitList": 6.0, "Hits": "GLK1|HXK1|MTH1|HXT7|HXT6|HXT4", "Description": "glucose transmembrane transport", "_GeneInHitList": 294.0, "Category": "GO Biological Processes", "LogP": -3.386314910717348, "_RANK_": 1.0, "FirstInGroupByEnrichment": 0.0, "_LogP_MyList": -3.386314910717348, "_TotalGeneInLibrary": 5919.0, "_GeneInGO": 21.0, "shared_name": "67", "Gene": "GO:1904659", "_MEMBER_MyList": 1.0, "STDV_InGO": 0.8246143220834374, "_COUNT_001": 6.0, "SUID": 2742014, "id_original": "67", "selected": false, "Log_q_value_": -1.6985878224546134, "canonicalName": "GO:1904659", "CategoryID": 19.0, "Z_score": 4.987129318865969, "EvidenceCutoff": 0.0, "PARENT_GO": "19_GO:0051179 localization", "FirstInGroupByLogP": 0.0, "GROUP_ID": 13.0, "name": "67", "BestEnrichmentInGroup": 5.872023809523809, "CLUSTER_LABEL": " ", "GeneID": "850317|850614|851870|851943|851944|856492", "GiniIndex": 0.0, "_InGO": 2.0408163265306123, "_COUNT_001_PCT": 100.0}, "position": {"x": 367.16350305976005, "y": 56.56601749540596}, "selected": false}, {"data": {"id": "2742011", "Enrichment": 4.026530612244898, "BestLogPInGroup": -4.436199066039166, "_PATTERN_": "M1", "_GeneInGOAndHitList": 9.0, "Hits": "GLK1|AQY3|HXK1|MTH1|HXT7|HXT6|MAL31|VPS73|HXT4", "Description": "carbohydrate transport", "_GeneInHitList": 294.0, "Category": "GO Biological Processes", "LogP": -3.5280714868006293, "_RANK_": 1.0, "FirstInGroupByEnrichment": 0.0, "_LogP_MyList": -3.5280714868006293, "_TotalGeneInLibrary": 5919.0, "_GeneInGO": 45.0, "shared_name": "66", "Gene": "GO:0008643", "_MEMBER_MyList": 1.0, "STDV_InGO": 1.0046682763914156, "_COUNT_001": 9.0, "SUID": 2742011, "id_original": "66", "selected": false, "Log_q_value_": -1.8191550994679568, "canonicalName": "GO:0008643", "CategoryID": 19.0, "Z_score": 4.658909121582477, "EvidenceCutoff": 0.0, "PARENT_GO": "19_GO:0051179 localization", "FirstInGroupByLogP": 0.0, "GROUP_ID": 13.0, "name": "66", "BestEnrichmentInGroup": 5.872023809523809, "CLUSTER_LABEL": " ", "GeneID": "850317|850490|850614|851870|851943|851944|852601|852774|856492", "GiniIndex": 0.0, "_InGO": 3.061224489795918, "_COUNT_001_PCT": 100.0}, "position": {"x": 369.2009252399358, "y": -52.419898642289354}, "selected": false}, {"data": {"id": "2742008", "Enrichment": 5.872023809523809, "BestLogPInGroup": -4.436199066039166, "_PATTERN_": "M1", "_GeneInGOAndHitList": 7.0, "Hits": "GLK1|HXK1|MTH1|HXT7|HXT6|VPS73|HXT4", "Description": "monosaccharide transmembrane transport", "_GeneInHitList": 294.0, "Category": "GO Biological Processes", "LogP": -3.936743598280655, "_RANK_": 1.0, "FirstInGroupByEnrichment": 0.0, "_LogP_MyList": -3.936743598280655, "_TotalGeneInLibrary": 5919.0, "_GeneInGO": 24.0, "shared_name": "65", "Gene": "GO:0015749", "_MEMBER_MyList": 1.0, "STDV_InGO": 0.8891375930475126, "_COUNT_001": 7.0, "SUID": 2742008, "id_original": "65", "selected": false, "Log_q_value_": -2.1780297425855673, "canonicalName": "GO:0015749", "CategoryID": 19.0, "Z_score": 5.467301200614909, "EvidenceCutoff": 0.0, "PARENT_GO": "19_GO:0051179 localization", "FirstInGroupByLogP": 0.0, "GROUP_ID": 13.0, "name": "65", "BestEnrichmentInGroup": 5.872023809523809, "CLUSTER_LABEL": " ", "GeneID": "850317|850614|851870|851943|851944|852774|856492", "GiniIndex": 0.0, "_InGO": 2.380952380952381, "_COUNT_001_PCT": 100.0}, "position": {"x": 250.15530527533622, "y": 0.13445316435127097}, "selected": false}, {"data": {"id": "2742005", "Enrichment": 5.872023809523809, "BestLogPInGroup": -4.436199066039166, "_PATTERN_": "M1", "_GeneInGOAndHitList": 7.0, "Hits": "GLK1|HXK1|MTH1|HXT7|HXT6|VPS73|HXT4", "Description": "hexose transmembrane transport", "_GeneInHitList": 294.0, "Category": "GO Biological Processes", "LogP": -3.936743598280655, "_RANK_": 1.0, "FirstInGroupByEnrichment": 0.0, "_LogP_MyList": -3.936743598280655, "_TotalGeneInLibrary": 5919.0, "_GeneInGO": 24.0, "shared_name": "64", "Gene": "GO:0008645", "_MEMBER_MyList": 1.0, "STDV_InGO": 0.8891375930475126, "_COUNT_001": 7.0, "SUID": 2742005, "id_original": "64", "selected": false, "Log_q_value_": -2.1780297425855673, "canonicalName": "GO:0008645", "CategoryID": 19.0, "Z_score": 5.467301200614909, "EvidenceCutoff": 0.0, "PARENT_GO": "19_GO:0051179 localization", "FirstInGroupByLogP": 0.0, "GROUP_ID": 13.0, "name": "64", "BestEnrichmentInGroup": 5.872023809523809, "CLUSTER_LABEL": " ", "GeneID": "850317|850614|851870|851943|851944|852774|856492", "GiniIndex": 0.0, "_InGO": 2.380952380952381, "_COUNT_001_PCT": 100.0}, "position": {"x": 324.17040384711356, "y": -6.993720663773729}, "selected": false}, {"data": {"id": "2742002", "Enrichment": 3.728269085411942, "BestLogPInGroup": -4.542469332779946, "_PATTERN_": "M1", "_GeneInGOAndHitList": 5.0, "Hits": "NTH1|YDR248C|NTH2|GUT2|GPH1", "Description": "cellular carbohydrate catabolic process", "_GeneInHitList": 294.0, "Category": "GO Biological Processes", "LogP": -2.018933858847451, "_RANK_": 1.0, "FirstInGroupByEnrichment": 0.0, "_LogP_MyList": -2.018933858847451, "_TotalGeneInLibrary": 5919.0, "_GeneInGO": 27.0, "shared_name": "63", "Gene": "GO:0044275", "_MEMBER_MyList": 1.0, "STDV_InGO": 0.7540721961117447, "_COUNT_001": 5.0, "SUID": 2742002, "id_original": "63", "selected": false, "Log_q_value_": -0.5752284991522644, "canonicalName": "GO:0044275", "CategoryID": 19.0, "Z_score": 3.2481613711083397, "EvidenceCutoff": 0.0, "PARENT_GO": "19_GO:0008152 metabolic process", "FirstInGroupByLogP": 0.0, "GROUP_ID": 12.0, "name": "63", "BestEnrichmentInGroup": 16.10612244897959, "CLUSTER_LABEL": " ", "GeneID": "851564|851835|852286|854651|856289", "GiniIndex": 0.0, "_InGO": 1.7006802721088434, "_COUNT_001_PCT": 100.0}, "position": {"x": 158.1916517108831, "y": 77.7784045803669}, "selected": false}, {"data": {"id": "2741999", "Enrichment": 5.033163265306122, "BestLogPInGroup": -5.272416855992848, "_PATTERN_": "M1", "_GeneInGOAndHitList": 7.0, "Hits": "MIC19|UPS2|ATP14|TIM11|SDH3|MIC27|ATP20", "Description": "inner mitochondrial membrane organization", "_GeneInHitList": 294.0, "Category": "GO Biological Processes", "LogP": -3.477119776423807, "_RANK_": 1.0, "FirstInGroupByEnrichment": 0.0, "_LogP_MyList": -3.477119776423807, "_TotalGeneInLibrary": 5919.0, "_GeneInGO": 28.0, "shared_name": "62", "Gene": "GO:0007007", "_MEMBER_MyList": 1.0, "STDV_InGO": 0.8891375930475126, "_COUNT_001": 7.0, "SUID": 2741999, "id_original": "62", "selected": false, "Log_q_value_": -1.7789272544829076, "canonicalName": "GO:0007007", "CategoryID": 19.0, "Z_score": 4.890240177622643, "EvidenceCutoff": 0.0, "PARENT_GO": "19_GO:0009987 cellular process", "FirstInGroupByLogP": 0.0, "GROUP_ID": 11.0, "name": "62", "BestEnrichmentInGroup": 10.98144712430427, "CLUSTER_LABEL": " ", "GeneID": "850563|850865|851002|851922|853716|855623|856131", "GiniIndex": 0.0, "_InGO": 2.380952380952381, "_COUNT_001_PCT": 100.0}, "position": {"x": 182.10613573493094, "y": 753.5970080959919}, "selected": false}, {"data": {"id": "2741996", "Enrichment": 2.0424430641821942, "BestLogPInGroup": -5.272416855992848, "_PATTERN_": "M1", "_GeneInGOAndHitList": 28.0, "Hits": "MIC19|COX12|UPS2|ATP14|ATG33|SSA1|HSP78|TIM11|QCR7|ATG8|COR1|OM14|SDH8|QCR9|MEF2|SDH3|MDM32|ATG32|FIS1|ATP18|MIX17|IMP2|MIC27|POR1|HSP82|ATG29|ATP20|AIM17", "Description": "mitochondrion organization", "_GeneInHitList": 294.0, "Category": "GO Biological Processes", "LogP": -3.6683676916916257, "_RANK_": 1.0, "FirstInGroupByEnrichment": 0.0, "_LogP_MyList": -3.6683676916916257, "_TotalGeneInLibrary": 5919.0, "_GeneInGO": 276.0, "shared_name": "61", "Gene": "GO:0007005", "_MEMBER_MyList": 1.0, "STDV_InGO": 1.7119805975670632, "_COUNT_001": 28.0, "SUID": 2741996, "id_original": "61", "selected": false, "Log_q_value_": -1.94096789866494, "canonicalName": "GO:0007005", "CategoryID": 19.0, "Z_score": 4.054634788378833, "EvidenceCutoff": 0.0, "PARENT_GO": "19_GO:0009987 cellular process", "FirstInGroupByLogP": 0.0, "GROUP_ID": 11.0, "name": "61", "BestEnrichmentInGroup": 10.98144712430427, "CLUSTER_LABEL": " ", "GeneID": "850563|850727|850865|851002|851070|851259|851845|851922|852142|852200|852235|852531|852572|853095|853342|853716|854318|854660|854745|854893|855014|855051|855623|855669|855836|855937|856131|856365", "GiniIndex": 0.0, "_InGO": 9.523809523809524, "_COUNT_001_PCT": 100.0}, "position": {"x": 119.25721300543876, "y": 841.1815417873981}, "selected": false}, {"data": {"id": "2741993", "Enrichment": 6.710884353741496, "BestLogPInGroup": -5.6116974553384775, "_PATTERN_": "M1", "_GeneInGOAndHitList": 4.0, "Hits": "VID24|MTL1|YKR075C|YOR062C", "Description": "protein catabolic process in the vacuole", "_GeneInHitList": 294.0, "Category": "GO Biological Processes", "LogP": -2.667682887204678, "_RANK_": 1.0, "FirstInGroupByEnrichment": 0.0, "_LogP_MyList": -2.667682887204678, "_TotalGeneInLibrary": 5919.0, "_GeneInGO": 12.0, "shared_name": "60", "Gene": "GO:0007039", "_MEMBER_MyList": 1.0, "STDV_InGO": 0.6756285589444189, "_COUNT_001": 4.0, "SUID": 2741993, "id_original": "60", "selected": false, "Log_q_value_": -1.113698716666834, "canonicalName": "GO:0007039", "CategoryID": 19.0, "Z_score": 4.526998864350259, "EvidenceCutoff": 0.0, "PARENT_GO": "19_GO:0008152 metabolic process", "FirstInGroupByLogP": 0.0, "GROUP_ID": 10.0, "name": "60", "BestEnrichmentInGroup": 6.710884353741496, "CLUSTER_LABEL": " ", "GeneID": "852402|852905|853950|854228", "GiniIndex": 0.0, "_InGO": 1.3605442176870748, "_COUNT_001_PCT": 100.0}, "position": {"x": 780.7683729022893, "y": 52.30532871366768}, "selected": false}, {"data": {"id": "2741990", "Enrichment": 5.921368547418967, "BestLogPInGroup": -5.6116974553384775, "_PATTERN_": "M1", "_GeneInGOAndHitList": 5.0, "Hits": "SDS24|MTL1|YAK1|YKR075C|YOR062C", "Description": "cellular response to glucose starvation", "_GeneInHitList": 294.0, "Category": "GO Biological Processes", "LogP": -2.9573916182518474, "_RANK_": 1.0, "FirstInGroupByEnrichment": 0.0, "_LogP_MyList": -2.9573916182518474, "_TotalGeneInLibrary": 5919.0, "_GeneInGO": 17.0, "shared_name": "59", "Gene": "GO:0042149", "_MEMBER_MyList": 1.0, "STDV_InGO": 0.7540721961117447, "_COUNT_001": 5.0, "SUID": 2741990, "id_original": "59", "selected": false, "Log_q_value_": -1.345912500568209, "canonicalName": "GO:0042149", "CategoryID": 19.0, "Z_score": 4.64526644713336, "EvidenceCutoff": 0.0, "PARENT_GO": "19_GO:0050896 response to stimulus", "FirstInGroupByLogP": 0.0, "GROUP_ID": 10.0, "name": "59", "BestEnrichmentInGroup": 6.710884353741496, "CLUSTER_LABEL": " ", "GeneID": "852515|852905|853300|853950|854228", "GiniIndex": 0.0, "_InGO": 1.7006802721088434, "_COUNT_001_PCT": 100.0}, "position": {"x": 872.8630994647893, "y": 20.797317849409865}, "selected": false}, {"data": {"id": "2741987", "Enrichment": 2.8521258503401357, "BestLogPInGroup": -5.6116974553384775, "_PATTERN_": "M1", "_GeneInGOAndHitList": 17.0, "Hits": "DCS1|NRG1|ATG8|PHO5|SDS24|MTL1|YAK1|GSM1|HAP4|YKR075C|DDR2|YOR062C|DCS2|CAT8|USV1|PEP4|PHM8", "Description": "cellular response to nutrient levels", "_GeneInHitList": 294.0, "Category": "GO Biological Processes", "LogP": -4.109338376699844, "_RANK_": 1.0, "FirstInGroupByEnrichment": 0.0, "_LogP_MyList": -4.109338376699844, "_TotalGeneInLibrary": 5919.0, "_GeneInGO": 120.0, "shared_name": "58", "Gene": "GO:0031669", "_MEMBER_MyList": 1.0, "STDV_InGO": 1.3612671039915658, "_COUNT_001": 17.0, "SUID": 2741987, "id_original": "58", "selected": false, "Log_q_value_": -2.3212407433195468, "canonicalName": "GO:0031669", "CategoryID": 19.0, "Z_score": 4.685807510772305, "EvidenceCutoff": 0.0, "PARENT_GO": "19_GO:0050896 response to stimulus", "FirstInGroupByLogP": 0.0, "GROUP_ID": 10.0, "name": "58", "BestEnrichmentInGroup": 6.710884353741496, "CLUSTER_LABEL": " ", "GeneID": "850974|851613|852200|852390|852515|852905|853300|853341|853751|853950|854104|854228|854344|855323|855871|855949|856759", "GiniIndex": 0.0, "_InGO": 5.782312925170068, "_COUNT_001_PCT": 100.0}, "position": {"x": 938.0574964374456, "y": 3.9166110623004897}, "selected": false}, {"data": {"id": "2741984", "Enrichment": 2.85344689056725, "BestLogPInGroup": -5.6116974553384775, "_PATTERN_": "M1", "_GeneInGOAndHitList": 18.0, "Hits": "DCS1|NRG1|ATG8|PHO5|SDS24|MTL1|YAK1|GSM1|HAP4|MSN4|YKR075C|DDR2|YOR062C|DCS2|CAT8|USV1|PEP4|PHM8", "Description": "response to nutrient levels", "_GeneInHitList": 294.0, "Category": "GO Biological Processes", "LogP": -4.322654694680918, "_RANK_": 1.0, "FirstInGroupByEnrichment": 0.0, "_LogP_MyList": -4.322654694680918, "_TotalGeneInLibrary": 5919.0, "_GeneInGO": 127.0, "shared_name": "57", "Gene": "GO:0031667", "_MEMBER_MyList": 1.0, "STDV_InGO": 1.3982016102952535, "_COUNT_001": 18.0, "SUID": 2741984, "id_original": "57", "selected": false, "Log_q_value_": -2.5122806665894686, "canonicalName": "GO:0031667", "CategoryID": 19.0, "Z_score": 4.826892338525146, "EvidenceCutoff": 0.0, "PARENT_GO": "19_GO:0050896 response to stimulus", "FirstInGroupByLogP": 0.0, "GROUP_ID": 10.0, "name": "57", "BestEnrichmentInGroup": 6.710884353741496, "CLUSTER_LABEL": " ", "GeneID": "850974|851613|852200|852390|852515|852905|853300|853341|853751|853803|853950|854104|854228|854344|855323|855871|855949|856759", "GiniIndex": 0.0, "_InGO": 6.122448979591836, "_COUNT_001_PCT": 100.0}, "position": {"x": 1007.0105909198675, "y": -23.215308798539354}, "selected": false}, {"data": {"id": "2741981", "Enrichment": 3.738921282798834, "BestLogPInGroup": -5.6116974553384775, "_PATTERN_": "M1", "_GeneInGOAndHitList": 13.0, "Hits": "DCS1|ATG8|PHO5|SDS24|MTL1|YAK1|GSM1|YKR075C|DDR2|YOR062C|DCS2|PEP4|PHM8", "Description": "cellular response to starvation", "_GeneInHitList": 294.0, "Category": "GO Biological Processes", "LogP": -4.500583243544872, "_RANK_": 1.0, "FirstInGroupByEnrichment": 0.0, "_LogP_MyList": -4.500583243544872, "_TotalGeneInLibrary": 5919.0, "_GeneInGO": 70.0, "shared_name": "56", "Gene": "GO:0009267", "_MEMBER_MyList": 1.0, "STDV_InGO": 1.198957642874661, "_COUNT_001": 13.0, "SUID": 2741981, "id_original": "56", "selected": false, "Log_q_value_": -2.67627343751013, "canonicalName": "GO:0009267", "CategoryID": 19.0, "Z_score": 5.269721239321746, "EvidenceCutoff": 0.0, "PARENT_GO": "19_GO:0050896 response to stimulus", "FirstInGroupByLogP": 0.0, "GROUP_ID": 10.0, "name": "56", "BestEnrichmentInGroup": 6.710884353741496, "CLUSTER_LABEL": " ", "GeneID": "850974|852200|852390|852515|852905|853300|853341|853950|854104|854228|854344|855949|856759", "GiniIndex": 0.0, "_InGO": 4.421768707482993, "_COUNT_001_PCT": 100.0}, "position": {"x": 867.5477613300237, "y": -31.97523464082451}, "selected": false}, {"data": {"id": "2741978", "Enrichment": 3.7580952380952377, "BestLogPInGroup": -5.6116974553384775, "_PATTERN_": "M1", "_GeneInGOAndHitList": 14.0, "Hits": "DCS1|ATG8|PHO5|SDS24|MTL1|YAK1|GSM1|MSN4|YKR075C|DDR2|YOR062C|DCS2|PEP4|PHM8", "Description": "response to starvation", "_GeneInHitList": 294.0, "Category": "GO Biological Processes", "LogP": -4.832196272682693, "_RANK_": 1.0, "FirstInGroupByEnrichment": 0.0, "_LogP_MyList": -4.832196272682693, "_TotalGeneInLibrary": 5919.0, "_GeneInGO": 75.0, "shared_name": "55", "Gene": "GO:0042594", "_MEMBER_MyList": 1.0, "STDV_InGO": 1.24200126418405, "_COUNT_001": 14.0, "SUID": 2741978, "id_original": "55", "selected": false, "Log_q_value_": -2.988581311452564, "canonicalName": "GO:0042594", "CategoryID": 19.0, "Z_score": 5.495214745743139, "EvidenceCutoff": 0.0, "PARENT_GO": "19_GO:0050896 response to stimulus", "FirstInGroupByLogP": 0.0, "GROUP_ID": 10.0, "name": "55", "BestEnrichmentInGroup": 6.710884353741496, "CLUSTER_LABEL": " ", "GeneID": "850974|852200|852390|852515|852905|853300|853341|853803|853950|854104|854228|854344|855949|856759", "GiniIndex": 0.0, "_InGO": 4.761904761904762, "_COUNT_001_PCT": 100.0}, "position": {"x": 899.1798719257268, "y": 68.39149509550361}, "selected": false}, {"data": {"id": "2741975", "Enrichment": 2.936011904761905, "BestLogPInGroup": -5.6116974553384775, "_PATTERN_": "M1", "_GeneInGOAndHitList": 21.0, "Hits": "DCS1|NTH1|NRG1|ATG8|PHO5|TPS1|SDS24|STF2|MTL1|YAK1|GSM1|HAP4|MSN4|YKR075C|DDR2|YOR062C|DCS2|CAT8|USV1|PEP4|PHM8", "Description": "response to external stimulus", "_GeneInHitList": 294.0, "Category": "GO Biological Processes", "LogP": -5.153617588317383, "_RANK_": 1.0, "FirstInGroupByEnrichment": 0.0, "_LogP_MyList": -5.153617588317383, "_TotalGeneInLibrary": 5919.0, "_GeneInGO": 144.0, "shared_name": "54", "Gene": "GO:0009605", "_MEMBER_MyList": 1.0, "STDV_InGO": 1.502000147334668, "_COUNT_001": 21.0, "SUID": 2741975, "id_original": "54", "selected": false, "Log_q_value_": -3.2740049738157353, "canonicalName": "GO:0009605", "CategoryID": 19.0, "Z_score": 5.376667221680957, "EvidenceCutoff": 0.0, "PARENT_GO": "19_GO:0050896 response to stimulus", "FirstInGroupByLogP": 0.0, "GROUP_ID": 10.0, "name": "54", "BestEnrichmentInGroup": 6.710884353741496, "CLUSTER_LABEL": " ", "GeneID": "850974|851564|851613|852200|852390|852423|852515|852891|852905|853300|853341|853751|853803|853950|854104|854228|854344|855323|855871|855949|856759", "GiniIndex": 0.0, "_InGO": 7.142857142857143, "_COUNT_001_PCT": 100.0}, "position": {"x": 960.4690870136175, "y": 71.52012382627754}, "selected": false}, {"data": {"id": "2741972", "Enrichment": 3.1457270408163263, "BestLogPInGroup": -5.6116974553384775, "_PATTERN_": "M1", "_GeneInGOAndHitList": 20.0, "Hits": "DCS1|NTH1|NRG1|ATG8|PHO5|TPS1|SDS24|STF2|MTL1|YAK1|GSM1|HAP4|YKR075C|DDR2|YOR062C|DCS2|CAT8|USV1|PEP4|PHM8", "Description": "cellular response to external stimulus", "_GeneInHitList": 294.0, "Category": "GO Biological Processes", "LogP": -5.399707033457556, "_RANK_": 1.0, "FirstInGroupByEnrichment": 0.0, "_LogP_MyList": -5.399707033457556, "_TotalGeneInLibrary": 5919.0, "_GeneInGO": 128.0, "shared_name": "53", "Gene": "GO:0071496", "_MEMBER_MyList": 1.0, "STDV_InGO": 1.4684842185478635, "_COUNT_001": 20.0, "SUID": 2741972, "id_original": "53", "selected": false, "Log_q_value_": -3.4924229923580503, "canonicalName": "GO:0071496", "CategoryID": 19.0, "Z_score": 5.610512817511911, "EvidenceCutoff": 0.0, "PARENT_GO": "19_GO:0050896 response to stimulus", "FirstInGroupByLogP": 0.0, "GROUP_ID": 10.0, "name": "53", "BestEnrichmentInGroup": 6.710884353741496, "CLUSTER_LABEL": " ", "GeneID": "850974|851564|851613|852200|852390|852423|852515|852891|852905|853300|853341|853751|853950|854104|854228|854344|855323|855871|855949|856759", "GiniIndex": 0.0, "_InGO": 6.802721088435374, "_COUNT_001_PCT": 100.0}, "position": {"x": 1004.9940198749456, "y": 32.71288715482979}, "selected": false}, {"data": {"id": "2741969", "Enrichment": 3.1457270408163263, "BestLogPInGroup": -5.6116974553384775, "_PATTERN_": "M1", "_GeneInGOAndHitList": 20.0, "Hits": "DCS1|NTH1|NRG1|ATG8|PHO5|TPS1|SDS24|STF2|MTL1|YAK1|GSM1|HAP4|YKR075C|DDR2|YOR062C|DCS2|CAT8|USV1|PEP4|PHM8", "Description": "cellular response to extracellular stimulus", "_GeneInHitList": 294.0, "Category": "GO Biological Processes", "LogP": -5.399707033457556, "_RANK_": 1.0, "FirstInGroupByEnrichment": 0.0, "_LogP_MyList": -5.399707033457556, "_TotalGeneInLibrary": 5919.0, "_GeneInGO": 128.0, "shared_name": "52", "Gene": "GO:0031668", "_MEMBER_MyList": 1.0, "STDV_InGO": 1.4684842185478635, "_COUNT_001": 20.0, "SUID": 2741969, "id_original": "52", "selected": false, "Log_q_value_": -3.4924229923580503, "canonicalName": "GO:0031668", "CategoryID": 19.0, "Z_score": 5.610512817511911, "EvidenceCutoff": 0.0, "PARENT_GO": "19_GO:0050896 response to stimulus", "FirstInGroupByLogP": 0.0, "GROUP_ID": 10.0, "name": "52", "BestEnrichmentInGroup": 6.710884353741496, "CLUSTER_LABEL": " ", "GeneID": "850974|851564|851613|852200|852390|852423|852515|852891|852905|853300|853341|853751|853950|854104|854228|854344|855323|855871|855949|856759", "GiniIndex": 0.0, "_InGO": 6.802721088435374, "_COUNT_001_PCT": 100.0}, "position": {"x": 916.53952158393, "y": -69.8695446383831}, "selected": false}, {"data": {"id": "2741966", "Enrichment": 1.5523973444799606, "BestLogPInGroup": -5.971779133371012, "_PATTERN_": "M1", "_GeneInGOAndHitList": 32.0, "Hits": "GLK1|AGX1|HXK1|BNA5|TMA17|GPM2|KGD2|YDR248C|EMI2|BNA4|ACH1|YBR053C|DSD1|TDH1|VTC4|TES1|MDH1|MDH2|BDS1|PYK2|ALD4|XBP1|PHO84|CYB2|HFD1|ALD3|GAD1|GOR1|VTC3|ARO9|ICL1|VTC1", "Description": "organic acid metabolic process", "_GeneInHitList": 294.0, "Category": "GO Biological Processes", "LogP": -2.105960109142738, "_RANK_": 1.0, "FirstInGroupByEnrichment": 0.0, "_LogP_MyList": -2.105960109142738, "_TotalGeneInLibrary": 5919.0, "_GeneInGO": 415.0, "shared_name": "51", "Gene": "GO:0006082", "_MEMBER_MyList": 1.0, "STDV_InGO": 1.816371333835593, "_COUNT_001": 32.0, "SUID": 2741966, "id_original": "51", "selected": false, "Log_q_value_": -0.6523162269133715, "canonicalName": "GO:0006082", "CategoryID": 19.0, "Z_score": 2.6676957697490873, "EvidenceCutoff": 0.0, "PARENT_GO": "19_GO:0008152 metabolic process", "FirstInGroupByLogP": 0.0, "GROUP_ID": 9.0, "name": "51", "BestEnrichmentInGroup": 2.1939429618001043, "CLUSTER_LABEL": " ", "GeneID": "850317|850514|850614|850933|851448|851541|851726|851835|852128|852179|852266|852342|852679|853395|853441|853477|853777|853994|854000|854529|854556|854706|854916|854950|855137|855205|855291|855447|856088|856539|856794|856803", "GiniIndex": 0.0, "_InGO": 10.884353741496598, "_COUNT_001_PCT": 100.0}, "position": {"x": 452.3096937030706, "y": 231.08052478910713}, "selected": false}, {"data": {"id": "2741963", "Enrichment": 1.5637012086387951, "BestLogPInGroup": -5.971779133371012, "_PATTERN_": "M1", "_GeneInGOAndHitList": 32.0, "Hits": "GLK1|AGX1|HXK1|BNA5|TMA17|GPM2|KGD2|YDR248C|EMI2|BNA4|ACH1|YBR053C|DSD1|TDH1|VTC4|TES1|MDH1|MDH2|BDS1|PYK2|ALD4|XBP1|PHO84|CYB2|HFD1|ALD3|GAD1|GOR1|VTC3|ARO9|ICL1|VTC1", "Description": "oxoacid metabolic process", "_GeneInHitList": 294.0, "Category": "GO Biological Processes", "LogP": -2.1514956543785657, "_RANK_": 1.0, "FirstInGroupByEnrichment": 0.0, "_LogP_MyList": -2.1514956543785657, "_TotalGeneInLibrary": 5919.0, "_GeneInGO": 412.0, "shared_name": "50", "Gene": "GO:0043436", "_MEMBER_MyList": 1.0, "STDV_InGO": 1.816371333835593, "_COUNT_001": 32.0, "SUID": 2741963, "id_original": "50", "selected": false, "Log_q_value_": -0.6917776244370061, "canonicalName": "GO:0043436", "CategoryID": 19.0, "Z_score": 2.7116892410437106, "EvidenceCutoff": 0.0, "PARENT_GO": "19_GO:0008152 metabolic process", "FirstInGroupByLogP": 0.0, "GROUP_ID": 9.0, "name": "50", "BestEnrichmentInGroup": 2.1939429618001043, "CLUSTER_LABEL": " ", "GeneID": "850317|850514|850614|850933|851448|851541|851726|851835|852128|852179|852266|852342|852679|853395|853441|853477|853777|853994|854000|854529|854556|854706|854916|854950|855137|855205|855291|855447|856088|856539|856794|856803", "GiniIndex": 0.0, "_InGO": 10.884353741496598, "_COUNT_001_PCT": 100.0}, "position": {"x": 432.04861772955985, "y": 155.70903812528877}, "selected": false}, {"data": {"id": "2741960", "Enrichment": 2.1939429618001043, "BestLogPInGroup": -5.971779133371012, "_PATTERN_": "M1", "_GeneInGOAndHitList": 17.0, "Hits": "GLK1|HXK1|BNA5|TMA17|GPM2|YDR248C|EMI2|BNA4|ACH1|TDH1|TES1|PYK2|ALD4|CYB2|ALD3|GOR1|ICL1", "Description": "monocarboxylic acid metabolic process", "_GeneInHitList": 294.0, "Category": "GO Biological Processes", "LogP": -2.753839670069378, "_RANK_": 1.0, "FirstInGroupByEnrichment": 0.0, "_LogP_MyList": -2.753839670069378, "_TotalGeneInLibrary": 5919.0, "_GeneInGO": 156.0, "shared_name": "49", "Gene": "GO:0032787", "_MEMBER_MyList": 1.0, "STDV_InGO": 1.3612671039915658, "_COUNT_001": 17.0, "SUID": 2741960, "id_original": "49", "selected": false, "Log_q_value_": -1.1752570512317115, "canonicalName": "GO:0032787", "CategoryID": 19.0, "Z_score": 3.45478838925973, "EvidenceCutoff": 0.0, "PARENT_GO": "19_GO:0008152 metabolic process", "FirstInGroupByLogP": 0.0, "GROUP_ID": 9.0, "name": "49", "BestEnrichmentInGroup": 2.1939429618001043, "CLUSTER_LABEL": " ", "GeneID": "850317|850614|850933|851448|851541|851835|852128|852179|852266|853395|853477|854529|854556|854950|855205|855447|856794", "GiniIndex": 0.0, "_InGO": 5.782312925170068, "_COUNT_001_PCT": 100.0}, "position": {"x": 395.8272737354192, "y": 251.4088100063923}, "selected": false}, {"data": {"id": "2741957", "Enrichment": 1.9562092043295054, "BestLogPInGroup": -7.330800802960937, "_PATTERN_": "M1", "_GeneInGOAndHitList": 24.0, "Hits": "GLK1|KIN82|HXK1|KNS1|YLR345W|MRK1|GPM2|VHS1|YDR248C|EMI2|SDS24|FMP48|XKS1|TDA10|TPK1|YAK1|TDH1|PTK2|HRK1|PYK2|PKP1|YPK2|TPK2|YEF1", "Description": "phosphorylation", "_GeneInHitList": 294.0, "Category": "GO Biological Processes", "LogP": -2.9469652621171116, "_RANK_": 1.0, "FirstInGroupByEnrichment": 0.0, "_LogP_MyList": -2.9469652621171116, "_TotalGeneInLibrary": 5919.0, "_GeneInGO": 247.0, "shared_name": "48", "Gene": "GO:0016310", "_MEMBER_MyList": 1.0, "STDV_InGO": 1.596858768236636, "_COUNT_001": 24.0, "SUID": 2741957, "id_original": "48", "selected": false, "Log_q_value_": -1.345912500568209, "canonicalName": "GO:0016310", "CategoryID": 19.0, "Z_score": 3.5094023907293472, "EvidenceCutoff": 0.0, "PARENT_GO": "19_GO:0008152 metabolic process", "FirstInGroupByLogP": 0.0, "GROUP_ID": 8.0, "name": "48", "BestEnrichmentInGroup": 2.0034124579685058, "CLUSTER_LABEL": " ", "GeneID": "850317|850453|850614|850641|851059|851480|851541|851834|851835|852128|852515|852943|853108|853119|853275|853300|853395|853522|854441|854529|854769|855130|855898|856670", "GiniIndex": 0.0, "_InGO": 8.16326530612245, "_COUNT_001_PCT": 100.0}, "position": {"x": 444.5527929157171, "y": 188.68518863798408}, "selected": false}, {"data": {"id": "2741954", "Enrichment": 1.9319212533498247, "BestLogPInGroup": -7.330800802960937, "_PATTERN_": "M1", "_GeneInGOAndHitList": 57.0, "Hits": "GLK1|KIN82|HXK1|KNS1|UPS2|BNA5|ATP14|YLR345W|PHO11|MRK1|GPD1|GPM2|ATP16|KGD2|VHS1|YDR248C|ATP5|TIM11|ATP17|EMI2|ATP1|BNA4|ATP3|PHO5|SDS24|PNC1|FMP48|XKS1|TDA10|PHO81|LSC2|SOL4|TPK1|YAK1|TDH1|TES1|PTK2|ATP2|NNR2|ATP7|ATP19|HRK1|PYK2|GUT2|PKP1|ATP18|PLB1|YPK2|PGM2|NNR1|ATP15|TPK2|ATP4|ATP20|YEF1|PHM8|MCO10", "Description": "phosphate-containing compound metabolic process", "_GeneInHitList": 294.0, "Category": "GO Biological Processes", "LogP": -6.302032679943681, "_RANK_": 1.0, "FirstInGroupByEnrichment": 0.0, "_LogP_MyList": -6.302032679943681, "_TotalGeneInLibrary": 5919.0, "_GeneInGO": 594.0, "shared_name": "47", "Gene": "GO:0006796", "_MEMBER_MyList": 1.0, "STDV_InGO": 2.3056352602756656, "_COUNT_001": 57.0, "SUID": 2741954, "id_original": "47", "selected": false, "Log_q_value_": -4.292086296947028, "canonicalName": "GO:0006796", "CategoryID": 19.0, "Z_score": 5.47409733081322, "EvidenceCutoff": 0.0, "PARENT_GO": "19_GO:0008152 metabolic process", "FirstInGroupByLogP": 0.0, "GROUP_ID": 8.0, "name": "47", "BestEnrichmentInGroup": 2.0034124579685058, "CLUSTER_LABEL": " ", "GeneID": "850317|850453|850614|850641|850865|850933|851002|851059|851299|851480|851539|851541|851560|851726|851834|851835|851892|851922|851983|852128|852177|852179|852327|852390|852515|852846|852943|853108|853119|853148|853159|853163|853275|853300|853395|853477|853522|853585|853706|853853|854077|854441|854529|854651|854769|854893|855020|855130|855131|855521|855857|855898|856027|856131|856670|856759|1466480", "GiniIndex": 0.0, "_InGO": 19.387755102040817, "_COUNT_001_PCT": 100.0}, "position": {"x": 586.2846692890081, "y": 187.71742282987861}, "selected": false}, {"data": {"id": "2741951", "Enrichment": 5.872023809523809, "BestLogPInGroup": -7.45117124911709, "_PATTERN_": "M1", "_GeneInGOAndHitList": 7.0, "Hits": "SDH2|KGD2|LSC2|SDH1|SDH3|MDH1|POR1", "Description": "Pyruvate metabolism and Citric Acid (TCA) cycle", "_GeneInHitList": 294.0, "Category": "Reactome Gene Sets", "LogP": -3.936743598280655, "_RANK_": 1.0, "FirstInGroupByEnrichment": 0.0, "_LogP_MyList": -3.936743598280655, "_TotalGeneInLibrary": 5919.0, "_GeneInGO": 24.0, "shared_name": "46", "Gene": "R-SCE-71406", "_MEMBER_MyList": 1.0, "STDV_InGO": 0.8891375930475126, "_COUNT_001": 7.0, "SUID": 2741951, "id_original": "46", "selected": false, "Log_q_value_": -2.1780297425855673, "canonicalName": "R-SCE-71406", "CategoryID": 6.0, "Z_score": 5.467301200614909, "EvidenceCutoff": 0.0, "PARENT_GO": "", "FirstInGroupByLogP": 0.0, "GROUP_ID": 7.0, "name": "46", "BestEnrichmentInGroup": 11.184807256235827, "CLUSTER_LABEL": " ", "GeneID": "850685|851726|853159|853709|853716|853777|855669", "GiniIndex": 0.0, "_InGO": 2.380952380952381, "_COUNT_001_PCT": 100.0}, "position": {"x": 322.3497861713323, "y": 413.3999866116169}, "selected": false}, {"data": {"id": "2741948", "Enrichment": 5.195523370638577, "BestLogPInGroup": -7.45117124911709, "_PATTERN_": "M1", "_GeneInGOAndHitList": 8.0, "Hits": "SDH2|KGD2|SDH4|LSC2|SDH1|SDH3|MDH1|MDH2", "Description": "Citrate cycle (TCA cycle) - Saccharomyces cerevisiae (budding yeast)", "_GeneInHitList": 294.0, "Category": "KEGG Pathway", "LogP": -4.0107904637872265, "_RANK_": 1.0, "FirstInGroupByEnrichment": 0.0, "_LogP_MyList": -4.0107904637872265, "_TotalGeneInLibrary": 5919.0, "_GeneInGO": 31.0, "shared_name": "45", "Gene": "sce00020", "_MEMBER_MyList": 1.0, "STDV_InGO": 0.9488706521462137, "_COUNT_001": 8.0, "SUID": 2741948, "id_original": "45", "selected": false, "Log_q_value_": -2.231293002168847, "canonicalName": "sce00020", "CategoryID": 24.0, "Z_score": 5.354056786279749, "EvidenceCutoff": 0.0, "PARENT_GO": "", "FirstInGroupByLogP": 0.0, "GROUP_ID": 7.0, "name": "45", "BestEnrichmentInGroup": 11.184807256235827, "CLUSTER_LABEL": " ", "GeneID": "850685|851726|851758|853159|853709|853716|853777|853994", "GiniIndex": 0.0, "_InGO": 2.7210884353741496, "_COUNT_001_PCT": 100.0}, "position": {"x": 435.3177160114202, "y": 439.22441898466377}, "selected": false}, {"data": {"id": "2741945", "Enrichment": 7.549744897959185, "BestLogPInGroup": -7.45117124911709, "_PATTERN_": "M1", "_GeneInGOAndHitList": 6.0, "Hits": "SDH2|KGD2|LSC2|SDH1|SDH3|MDH1", "Description": "Citric acid cycle (TCA cycle)", "_GeneInHitList": 294.0, "Category": "Reactome Gene Sets", "LogP": -4.1256160677327, "_RANK_": 1.0, "FirstInGroupByEnrichment": 0.0, "_LogP_MyList": -4.1256160677327, "_TotalGeneInLibrary": 5919.0, "_GeneInGO": 16.0, "shared_name": "44", "Gene": "R-SCE-71403", "_MEMBER_MyList": 1.0, "STDV_InGO": 0.8246143220834374, "_COUNT_001": 6.0, "SUID": 2741945, "id_original": "44", "selected": false, "Log_q_value_": -2.333153628949953, "canonicalName": "R-SCE-71403", "CategoryID": 6.0, "Z_score": 5.997190815213032, "EvidenceCutoff": 0.0, "PARENT_GO": "", "FirstInGroupByLogP": 0.0, "GROUP_ID": 7.0, "name": "44", "BestEnrichmentInGroup": 11.184807256235827, "CLUSTER_LABEL": " ", "GeneID": "850685|851726|853159|853709|853716|853777", "GiniIndex": 0.0, "_InGO": 2.0408163265306123, "_COUNT_001_PCT": 100.0}, "position": {"x": 437.7958349079046, "y": 374.9511340725544}, "selected": false}, {"data": {"id": "2741942", "Enrichment": 11.184807256235827, "BestLogPInGroup": -7.45117124911709, "_PATTERN_": "M1", "_GeneInGOAndHitList": 5.0, "Hits": "SDH2|SDH4|LSC2|SDH1|SDH3", "Description": "Glutamate degradation VII", "_GeneInHitList": 294.0, "Category": "WikiPathways", "LogP": -4.5053524797695825, "_RANK_": 1.0, "FirstInGroupByEnrichment": 0.0, "_LogP_MyList": -4.5053524797695825, "_TotalGeneInLibrary": 5919.0, "_GeneInGO": 9.0, "shared_name": "43", "Gene": "WP559", "_MEMBER_MyList": 1.0, "STDV_InGO": 0.7540721961117447, "_COUNT_001": 5.0, "SUID": 2741942, "id_original": "43", "selected": false, "Log_q_value_": -2.676296238710379, "canonicalName": "WP559", "CategoryID": 27.0, "Z_score": 6.990048853685036, "EvidenceCutoff": 0.0, "PARENT_GO": "", "FirstInGroupByLogP": 0.0, "GROUP_ID": 7.0, "name": "43", "BestEnrichmentInGroup": 11.184807256235827, "CLUSTER_LABEL": " ", "GeneID": "850685|851758|853159|853709|853716", "GiniIndex": 0.0, "_InGO": 1.7006802721088434, "_COUNT_001_PCT": 100.0}, "position": {"x": 354.39067591132255, "y": 482.0858234036091}, "selected": false}, {"data": {"id": "2741939", "Enrichment": 5.662308673469387, "BestLogPInGroup": -7.45117124911709, "_PATTERN_": "M1", "_GeneInGOAndHitList": 9.0, "Hits": "SDH2|KGD2|SDH4|LSC2|SDH1|SDH3|MDH1|MDH2|ICL1", "Description": "TCA cycle", "_GeneInHitList": 294.0, "Category": "WikiPathways", "LogP": -4.77981737523107, "_RANK_": 1.0, "FirstInGroupByEnrichment": 0.0, "_LogP_MyList": -4.77981737523107, "_TotalGeneInLibrary": 5919.0, "_GeneInGO": 32.0, "shared_name": "42", "Gene": "WP490", "_MEMBER_MyList": 1.0, "STDV_InGO": 1.0046682763914156, "_COUNT_001": 9.0, "SUID": 2741939, "id_original": "42", "selected": false, "Log_q_value_": -2.9411097484956854, "canonicalName": "WP490", "CategoryID": 27.0, "Z_score": 6.045453382203662, "EvidenceCutoff": 0.0, "PARENT_GO": "", "FirstInGroupByLogP": 0.0, "GROUP_ID": 7.0, "name": "42", "BestEnrichmentInGroup": 11.184807256235827, "CLUSTER_LABEL": " ", "GeneID": "850685|851726|851758|853159|853709|853716|853777|853994|856794", "GiniIndex": 0.0, "_InGO": 3.061224489795918, "_COUNT_001_PCT": 100.0}, "position": {"x": 380.75093401373954, "y": 430.7409595120075}, "selected": false}, {"data": {"id": "2741936", "Enrichment": 6.039795918367346, "BestLogPInGroup": -7.45117124911709, "_PATTERN_": "M1", "_GeneInGOAndHitList": 9.0, "Hits": "SDH2|KGD2|SDH4|LSC2|SDH1|SDH3|MDH1|MDH2|ICL1", "Description": "tricarboxylic acid cycle", "_GeneInHitList": 294.0, "Category": "GO Biological Processes", "LogP": -5.033928255921912, "_RANK_": 1.0, "FirstInGroupByEnrichment": 0.0, "_LogP_MyList": -5.033928255921912, "_TotalGeneInLibrary": 5919.0, "_GeneInGO": 30.0, "shared_name": "41", "Gene": "GO:0006099", "_MEMBER_MyList": 1.0, "STDV_InGO": 1.0046682763914156, "_COUNT_001": 9.0, "SUID": 2741936, "id_original": "41", "selected": false, "Log_q_value_": -3.164908714917688, "canonicalName": "GO:0006099", "CategoryID": 19.0, "Z_score": 6.326342245632767, "EvidenceCutoff": 0.0, "PARENT_GO": "19_GO:0008152 metabolic process", "FirstInGroupByLogP": 0.0, "GROUP_ID": 7.0, "name": "41", "BestEnrichmentInGroup": 11.184807256235827, "CLUSTER_LABEL": " ", "GeneID": "850685|851726|851758|853159|853709|853716|853777|853994|856794", "GiniIndex": 0.0, "_InGO": 3.061224489795918, "_COUNT_001_PCT": 100.0}, "position": {"x": 388.287978538886, "y": 389.18650089384346}, "selected": false}, {"data": {"id": "2741933", "Enrichment": 7.66958211856171, "BestLogPInGroup": -7.45117124911709, "_PATTERN_": "M1", "_GeneInGOAndHitList": 8.0, "Hits": "SDH2|KGD2|SDH4|LSC2|SDH1|SDH3|MDH1|MDH2", "Description": "TCA cycle - detailed", "_GeneInHitList": 294.0, "Category": "WikiPathways", "LogP": -5.409871404758037, "_RANK_": 1.0, "FirstInGroupByEnrichment": 0.0, "_LogP_MyList": -5.409871404758037, "_TotalGeneInLibrary": 5919.0, "_GeneInGO": 21.0, "shared_name": "40", "Gene": "WP296", "_MEMBER_MyList": 1.0, "STDV_InGO": 0.9488706521462137, "_COUNT_001": 8.0, "SUID": 2741933, "id_original": "40", "selected": false, "Log_q_value_": -3.4924229923580503, "canonicalName": "WP296", "CategoryID": 27.0, "Z_score": 6.999318714801051, "EvidenceCutoff": 0.0, "PARENT_GO": "", "FirstInGroupByLogP": 0.0, "GROUP_ID": 7.0, "name": "40", "BestEnrichmentInGroup": 11.184807256235827, "CLUSTER_LABEL": " ", "GeneID": "850685|851726|851758|853159|853709|853716|853777|853994", "GiniIndex": 0.0, "_InGO": 2.7210884353741496, "_COUNT_001_PCT": 100.0}, "position": {"x": 408.9446152538213, "y": 490.3044055569294}, "selected": false}, {"data": {"id": "2741930", "Enrichment": 3.4153607871720117, "BestLogPInGroup": -7.45117124911709, "_PATTERN_": "M1", "_GeneInGOAndHitList": 19.0, "Hits": "GLK1|AGX1|HXK1|SDH2|GPM2|KGD2|SDH4|YDR248C|EMI2|TDA10|LSC2|SOL4|TDH1|SDH1|SDH3|MDH1|MDH2|PYK2|ICL1", "Description": "Carbon metabolism - Saccharomyces cerevisiae (budding yeast)", "_GeneInHitList": 294.0, "Category": "KEGG Pathway", "LogP": -5.704028685032749, "_RANK_": 1.0, "FirstInGroupByEnrichment": 0.0, "_LogP_MyList": -5.704028685032749, "_TotalGeneInLibrary": 5919.0, "_GeneInGO": 112.0, "shared_name": "39", "Gene": "sce01200", "_MEMBER_MyList": 1.0, "STDV_InGO": 1.4339108527393725, "_COUNT_001": 19.0, "SUID": 2741930, "id_original": "39", "selected": false, "Log_q_value_": -3.7420058543532777, "canonicalName": "sce01200", "CategoryID": 24.0, "Z_score": 5.899497098846316, "EvidenceCutoff": 0.0, "PARENT_GO": "", "FirstInGroupByLogP": 0.0, "GROUP_ID": 7.0, "name": "39", "BestEnrichmentInGroup": 11.184807256235827, "CLUSTER_LABEL": " ", "GeneID": "850317|850514|850614|850685|851541|851726|851758|851835|852128|853119|853159|853163|853395|853709|853716|853777|853994|854529|856794", "GiniIndex": 0.0, "_InGO": 6.462585034013605, "_COUNT_001_PCT": 100.0}, "position": {"x": 389.2847627490911, "y": 303.04686390043526}, "selected": false}, {"data": {"id": "2741927", "Enrichment": 2.3516774231059943, "BestLogPInGroup": -7.45117124911709, "_PATTERN_": "M1", "_GeneInGOAndHitList": 41.0, "Hits": "GLK1|AGX1|GSY1|HXK1|SDH2|GSY2|GPD1|GPM2|NTH1|TPS2|KGD2|SDH4|YDR248C|EMI2|NTH2|RFS1|TPS1|TDA10|LSC2|SOL4|TDH1|TES1|SDH1|SDH3|MDH1|GLG1|MDH2|PYK2|ALD4|GUT2|TSL1|PGM2|HFD1|ALD3|GAD1|TPS3|GOR1|GPH1|ARO9|GLC3|ICL1", "Description": "Biosynthesis of secondary metabolites - Saccharomyces cerevisiae (budding yeast)", "_GeneInHitList": 294.0, "Category": "KEGG Pathway", "LogP": -6.797575375883804, "_RANK_": 1.0, "FirstInGroupByEnrichment": 0.0, "_LogP_MyList": -6.797575375883804, "_TotalGeneInLibrary": 5919.0, "_GeneInGO": 351.0, "shared_name": "38", "Gene": "sce01110", "_MEMBER_MyList": 1.0, "STDV_InGO": 2.0203713127142278, "_COUNT_001": 41.0, "SUID": 2741927, "id_original": "38", "selected": false, "Log_q_value_": -4.740089631794766, "canonicalName": "sce01110", "CategoryID": 24.0, "Z_score": 5.968660641723996, "EvidenceCutoff": 0.0, "PARENT_GO": "", "FirstInGroupByLogP": 0.0, "GROUP_ID": 7.0, "name": "38", "BestEnrichmentInGroup": 11.184807256235827, "CLUSTER_LABEL": " ", "GeneID": "850317|850514|850569|850614|850685|850962|851539|851541|851564|851646|851726|851758|851835|852128|852286|852341|852423|853119|853159|853163|853395|853477|853709|853716|853777|853932|853994|854529|854556|854651|854872|855131|855137|855205|855291|855303|855447|856289|856539|856705|856794", "GiniIndex": 0.0, "_InGO": 13.945578231292517, "_COUNT_001_PCT": 100.0}, "position": {"x": 287.54390848578544, "y": 302.4802726376179}, "selected": false}, {"data": {"id": "2741924", "Enrichment": 12.582908163265305, "BestLogPInGroup": -8.310637859619513, "_PATTERN_": "M1", "_GeneInGOAndHitList": 5.0, "Hits": "TPS2|TPS1|TSL1|PGM2|TPS3", "Description": "disaccharide biosynthetic process", "_GeneInHitList": 294.0, "Category": "GO Biological Processes", "LogP": -4.839538964813974, "_RANK_": 1.0, "FirstInGroupByEnrichment": 0.0, "_LogP_MyList": -4.839538964813974, "_TotalGeneInLibrary": 5919.0, "_GeneInGO": 8.0, "shared_name": "37", "Gene": "GO:0046351", "_MEMBER_MyList": 1.0, "STDV_InGO": 0.7540721961117447, "_COUNT_001": 5.0, "SUID": 2741924, "id_original": "37", "selected": false, "Log_q_value_": -2.990960584052295, "canonicalName": "GO:0046351", "CategoryID": 19.0, "Z_score": 7.494316133629537, "EvidenceCutoff": 0.0, "PARENT_GO": "19_GO:0008152 metabolic process", "FirstInGroupByLogP": 0.0, "GROUP_ID": 6.0, "name": "37", "BestEnrichmentInGroup": 20.132653061224488, "CLUSTER_LABEL": " ", "GeneID": "851646|852423|854872|855131|855303", "GiniIndex": 0.0, "_InGO": 1.7006802721088434, "_COUNT_001_PCT": 100.0}, "position": {"x": 65.95782602728934, "y": 153.9834827053669}, "selected": false}, {"data": {"id": "2741921", "Enrichment": 12.582908163265305, "BestLogPInGroup": -8.310637859619513, "_PATTERN_": "M1", "_GeneInGOAndHitList": 5.0, "Hits": "TPS2|TPS1|TSL1|PGM2|TPS3", "Description": "oligosaccharide biosynthetic process", "_GeneInHitList": 294.0, "Category": "GO Biological Processes", "LogP": -4.839538964813974, "_RANK_": 1.0, "FirstInGroupByEnrichment": 0.0, "_LogP_MyList": -4.839538964813974, "_TotalGeneInLibrary": 5919.0, "_GeneInGO": 8.0, "shared_name": "36", "Gene": "GO:0009312", "_MEMBER_MyList": 1.0, "STDV_InGO": 0.7540721961117447, "_COUNT_001": 5.0, "SUID": 2741921, "id_original": "36", "selected": false, "Log_q_value_": -2.990960584052295, "canonicalName": "GO:0009312", "CategoryID": 19.0, "Z_score": 7.494316133629537, "EvidenceCutoff": 0.0, "PARENT_GO": "19_GO:0008152 metabolic process", "FirstInGroupByLogP": 0.0, "GROUP_ID": 6.0, "name": "36", "BestEnrichmentInGroup": 20.132653061224488, "CLUSTER_LABEL": " ", "GeneID": "851646|852423|854872|855131|855303", "GiniIndex": 0.0, "_InGO": 1.7006802721088434, "_COUNT_001_PCT": 100.0}, "position": {"x": -5.735716453179407, "y": 290.609919177306}, "selected": false}, {"data": {"id": "2741918", "Enrichment": 6.039795918367346, "BestLogPInGroup": -8.310637859619513, "_PATTERN_": "M1", "_GeneInGOAndHitList": 9.0, "Hits": "HSP104|NTH1|TPS2|NTH2|TPS1|MAL31|TSL1|PGM2|TPS3", "Description": "oligosaccharide metabolic process", "_GeneInHitList": 294.0, "Category": "GO Biological Processes", "LogP": -5.033928255921912, "_RANK_": 1.0, "FirstInGroupByEnrichment": 0.0, "_LogP_MyList": -5.033928255921912, "_TotalGeneInLibrary": 5919.0, "_GeneInGO": 30.0, "shared_name": "35", "Gene": "GO:0009311", "_MEMBER_MyList": 1.0, "STDV_InGO": 1.0046682763914156, "_COUNT_001": 9.0, "SUID": 2741918, "id_original": "35", "selected": false, "Log_q_value_": -3.164908714917688, "canonicalName": "GO:0009311", "CategoryID": 19.0, "Z_score": 6.326342245632767, "EvidenceCutoff": 0.0, "PARENT_GO": "19_GO:0008152 metabolic process", "FirstInGroupByLogP": 0.0, "GROUP_ID": 6.0, "name": "35", "BestEnrichmentInGroup": 20.132653061224488, "CLUSTER_LABEL": " ", "GeneID": "850633|851564|851646|852286|852423|852601|854872|855131|855303", "GiniIndex": 0.0, "_InGO": 3.061224489795918, "_COUNT_001_PCT": 100.0}, "position": {"x": -19.215483298882532, "y": 241.14640842558174}, "selected": false}, {"data": {"id": "2741915", "Enrichment": 14.380466472303207, "BestLogPInGroup": -8.310637859619513, "_PATTERN_": "M1", "_GeneInGOAndHitList": 5.0, "Hits": "TPS2|TPS1|TSL1|PGM2|TPS3", "Description": "trehalose biosynthetic process", "_GeneInHitList": 294.0, "Category": "GO Biological Processes", "LogP": -5.247491544844542, "_RANK_": 1.0, "FirstInGroupByEnrichment": 0.0, "_LogP_MyList": -5.247491544844542, "_TotalGeneInLibrary": 5919.0, "_GeneInGO": 7.0, "shared_name": "34", "Gene": "GO:0005992", "_MEMBER_MyList": 1.0, "STDV_InGO": 0.7540721961117447, "_COUNT_001": 5.0, "SUID": 2741915, "id_original": "34", "selected": false, "Log_q_value_": -3.3570210027546863, "canonicalName": "GO:0005992", "CategoryID": 19.0, "Z_score": 8.097537048225105, "EvidenceCutoff": 0.0, "PARENT_GO": "19_GO:0008152 metabolic process", "FirstInGroupByLogP": 0.0, "GROUP_ID": 6.0, "name": "34", "BestEnrichmentInGroup": 20.132653061224488, "CLUSTER_LABEL": " ", "GeneID": "851646|852423|854872|855131|855303", "GiniIndex": 0.0, "_InGO": 1.7006802721088434, "_COUNT_001_PCT": 100.0}, "position": {"x": 52.37832773627372, "y": 193.9078982937458}, "selected": false}, {"data": {"id": "2741912", "Enrichment": 6.710884353741496, "BestLogPInGroup": -8.310637859619513, "_PATTERN_": "M1", "_GeneInGOAndHitList": 9.0, "Hits": "HSP104|NTH1|TPS2|NTH2|TPS1|MAL31|TSL1|PGM2|TPS3", "Description": "disaccharide metabolic process", "_GeneInHitList": 294.0, "Category": "GO Biological Processes", "LogP": -5.461171410690464, "_RANK_": 1.0, "FirstInGroupByEnrichment": 0.0, "_LogP_MyList": -5.461171410690464, "_TotalGeneInLibrary": 5919.0, "_GeneInGO": 27.0, "shared_name": "33", "Gene": "GO:0005984", "_MEMBER_MyList": 1.0, "STDV_InGO": 1.0046682763914156, "_COUNT_001": 9.0, "SUID": 2741912, "id_original": "33", "selected": false, "Log_q_value_": -3.530406273741435, "canonicalName": "GO:0005984", "CategoryID": 19.0, "Z_score": 6.799136511818551, "EvidenceCutoff": 0.0, "PARENT_GO": "19_GO:0008152 metabolic process", "FirstInGroupByLogP": 0.0, "GROUP_ID": 6.0, "name": "33", "BestEnrichmentInGroup": 20.132653061224488, "CLUSTER_LABEL": " ", "GeneID": "850633|851564|851646|852286|852423|852601|854872|855131|855303", "GiniIndex": 0.0, "_InGO": 3.061224489795918, "_COUNT_001_PCT": 100.0}, "position": {"x": -6.934599509820032, "y": 206.28096042753486}, "selected": false}, {"data": {"id": "2741909", "Enrichment": 12.079591836734696, "BestLogPInGroup": -8.310637859619513, "_PATTERN_": "M1", "_GeneInGOAndHitList": 6.0, "Hits": "NTH1|TPS2|TPS1|TSL1|PGM2|TPS3", "Description": "Trehalose anabolism", "_GeneInHitList": 294.0, "Category": "WikiPathways", "LogP": -5.596377654385471, "_RANK_": 1.0, "FirstInGroupByEnrichment": 0.0, "_LogP_MyList": -5.596377654385471, "_TotalGeneInLibrary": 5919.0, "_GeneInGO": 10.0, "shared_name": "32", "Gene": "WP398", "_MEMBER_MyList": 1.0, "STDV_InGO": 0.8246143220834374, "_COUNT_001": 6.0, "SUID": 2741909, "id_original": "32", "selected": false, "Log_q_value_": -3.65337806101943, "canonicalName": "WP398", "CategoryID": 27.0, "Z_score": 8.016165018237352, "EvidenceCutoff": 0.0, "PARENT_GO": "", "FirstInGroupByLogP": 0.0, "GROUP_ID": 6.0, "name": "32", "BestEnrichmentInGroup": 20.132653061224488, "CLUSTER_LABEL": " ", "GeneID": "851564|851646|852423|854872|855131|855303", "GiniIndex": 0.0, "_InGO": 2.0408163265306123, "_COUNT_001_PCT": 100.0}, "position": {"x": -36.97836171685128, "y": 180.87958561064033}, "selected": false}, {"data": {"id": "2741906", "Enrichment": 20.132653061224488, "BestLogPInGroup": -8.310637859619513, "_PATTERN_": "M1", "_GeneInGOAndHitList": 5.0, "Hits": "HSP104|TPS2|TPS1|TSL1|TPS3", "Description": "trehalose metabolism in response to stress", "_GeneInHitList": 294.0, "Category": "GO Biological Processes", "LogP": -6.533618995263603, "_RANK_": 1.0, "FirstInGroupByEnrichment": 0.0, "_LogP_MyList": -6.533618995263603, "_TotalGeneInLibrary": 5919.0, "_GeneInGO": 5.0, "shared_name": "31", "Gene": "GO:0070413", "_MEMBER_MyList": 1.0, "STDV_InGO": 0.7540721961117447, "_COUNT_001": 5.0, "SUID": 2741906, "id_original": "31", "selected": false, "Log_q_value_": -4.501396217555797, "canonicalName": "GO:0070413", "CategoryID": 19.0, "Z_score": 9.784067059501789, "EvidenceCutoff": 0.0, "PARENT_GO": "19_GO:0050896 response to stimulus", "FirstInGroupByLogP": 0.0, "GROUP_ID": 6.0, "name": "31", "BestEnrichmentInGroup": 20.132653061224488, "CLUSTER_LABEL": " ", "GeneID": "850633|851646|852423|854872|855303", "GiniIndex": 0.0, "_InGO": 1.7006802721088434, "_COUNT_001_PCT": 100.0}, "position": {"x": -11.624083152398157, "y": 126.26377140165596}, "selected": false}, {"data": {"id": "2741903", "Enrichment": 4.026530612244898, "BestLogPInGroup": -9.417680144618705, "_PATTERN_": "M1", "_GeneInGOAndHitList": 10.0, "Hits": "GSY1|IGD1|GSY2|MRX18|GLG1|GAC1|PCL7|PGM2|GPH1|GLC3", "Description": "glucan metabolic process", "_GeneInHitList": 294.0, "Category": "GO Biological Processes", "LogP": -3.8632769043878827, "_RANK_": 1.0, "FirstInGroupByEnrichment": 0.0, "_LogP_MyList": -3.8632769043878827, "_TotalGeneInLibrary": 5919.0, "_GeneInGO": 50.0, "shared_name": "30", "Gene": "GO:0044042", "_MEMBER_MyList": 1.0, "STDV_InGO": 1.057153797996807, "_COUNT_001": 10.0, "SUID": 2741903, "id_original": "30", "selected": false, "Log_q_value_": -2.1205022497942427, "canonicalName": "GO:0044042", "CategoryID": 19.0, "Z_score": 4.913012856820789, "EvidenceCutoff": 0.0, "PARENT_GO": "19_GO:0008152 metabolic process", "FirstInGroupByLogP": 0.0, "GROUP_ID": 5.0, "name": "30", "BestEnrichmentInGroup": 11.504373177842565, "CLUSTER_LABEL": " ", "GeneID": "850569|850572|850962|852346|853932|854350|854761|855131|856289|856705", "GiniIndex": 0.0, "_InGO": 3.401360544217687, "_COUNT_001_PCT": 100.0}, "position": {"x": 127.59073008002372, "y": 397.79778324247627}, "selected": false}, {"data": {"id": "2741900", "Enrichment": 4.026530612244898, "BestLogPInGroup": -9.417680144618705, "_PATTERN_": "M1", "_GeneInGOAndHitList": 10.0, "Hits": "GSY1|IGD1|GSY2|MRX18|GLG1|GAC1|PCL7|PGM2|GPH1|GLC3", "Description": "cellular glucan metabolic process", "_GeneInHitList": 294.0, "Category": "GO Biological Processes", "LogP": -3.8632769043878827, "_RANK_": 1.0, "FirstInGroupByEnrichment": 0.0, "_LogP_MyList": -3.8632769043878827, "_TotalGeneInLibrary": 5919.0, "_GeneInGO": 50.0, "shared_name": "29", "Gene": "GO:0006073", "_MEMBER_MyList": 1.0, "STDV_InGO": 1.057153797996807, "_COUNT_001": 10.0, "SUID": 2741900, "id_original": "29", "selected": false, "Log_q_value_": -2.1205022497942427, "canonicalName": "GO:0006073", "CategoryID": 19.0, "Z_score": 4.913012856820789, "EvidenceCutoff": 0.0, "PARENT_GO": "19_GO:0008152 metabolic process", "FirstInGroupByLogP": 0.0, "GROUP_ID": 5.0, "name": "29", "BestEnrichmentInGroup": 11.504373177842565, "CLUSTER_LABEL": " ", "GeneID": "850569|850572|850962|852346|853932|854350|854761|855131|856289|856705", "GiniIndex": 0.0, "_InGO": 3.401360544217687, "_COUNT_001_PCT": 100.0}, "position": {"x": 100.9759534686956, "y": 366.51250492216377}, "selected": false}, {"data": {"id": "2741897", "Enrichment": 7.549744897959185, "BestLogPInGroup": -9.417680144618705, "_PATTERN_": "M1", "_GeneInGOAndHitList": 6.0, "Hits": "GSY1|GSY2|GLG1|GAC1|GPH1|GLC3", "Description": "Glycogen metabolism", "_GeneInHitList": 294.0, "Category": "Reactome Gene Sets", "LogP": -4.1256160677327, "_RANK_": 1.0, "FirstInGroupByEnrichment": 0.0, "_LogP_MyList": -4.1256160677327, "_TotalGeneInLibrary": 5919.0, "_GeneInGO": 16.0, "shared_name": "28", "Gene": "R-SCE-8982491", "_MEMBER_MyList": 1.0, "STDV_InGO": 0.8246143220834374, "_COUNT_001": 6.0, "SUID": 2741897, "id_original": "28", "selected": false, "Log_q_value_": -2.333153628949953, "canonicalName": "R-SCE-8982491", "CategoryID": 6.0, "Z_score": 5.997190815213032, "EvidenceCutoff": 0.0, "PARENT_GO": "", "FirstInGroupByLogP": 0.0, "GROUP_ID": 5.0, "name": "28", "BestEnrichmentInGroup": 11.504373177842565, "CLUSTER_LABEL": " ", "GeneID": "850569|850962|853932|854350|856289|856705", "GiniIndex": 0.0, "_InGO": 2.0408163265306123, "_COUNT_001_PCT": 100.0}, "position": {"x": 188.85255564154716, "y": 315.62920986295967}, "selected": false}, {"data": {"id": "2741894", "Enrichment": 6.9689952904238615, "BestLogPInGroup": -9.417680144618705, "_PATTERN_": "M1", "_GeneInGOAndHitList": 9.0, "Hits": "GSY1|IGD1|GSY2|GLG1|GAC1|PCL7|PGM2|GPH1|GLC3", "Description": "glycogen metabolic process", "_GeneInHitList": 294.0, "Category": "GO Biological Processes", "LogP": -5.618104349538218, "_RANK_": 1.0, "FirstInGroupByEnrichment": 0.0, "_LogP_MyList": -5.618104349538218, "_TotalGeneInLibrary": 5919.0, "_GeneInGO": 26.0, "shared_name": "27", "Gene": "GO:0005977", "_MEMBER_MyList": 1.0, "STDV_InGO": 1.0046682763914156, "_COUNT_001": 9.0, "SUID": 2741894, "id_original": "27", "selected": false, "Log_q_value_": -3.6625156288641567, "canonicalName": "GO:0005977", "CategoryID": 19.0, "Z_score": 6.9729985018255425, "EvidenceCutoff": 0.0, "PARENT_GO": "19_GO:0008152 metabolic process", "FirstInGroupByLogP": 0.0, "GROUP_ID": 5.0, "name": "27", "BestEnrichmentInGroup": 11.504373177842565, "CLUSTER_LABEL": " ", "GeneID": "850569|850572|850962|853932|854350|854761|855131|856289|856705", "GiniIndex": 0.0, "_InGO": 3.061224489795918, "_COUNT_001_PCT": 100.0}, "position": {"x": 59.39889658392997, "y": 375.2296451199177}, "selected": false}, {"data": {"id": "2741891", "Enrichment": 5.251996450754215, "BestLogPInGroup": -9.417680144618705, "_PATTERN_": "M1", "_GeneInGOAndHitList": 12.0, "Hits": "GSY1|IGD1|GSY2|TPS2|TPS1|GLG1|GAC1|TSL1|PGM2|TPS3|GLC3|GPP2", "Description": "cellular carbohydrate biosynthetic process", "_GeneInHitList": 294.0, "Category": "GO Biological Processes", "LogP": -5.814622086748304, "_RANK_": 1.0, "FirstInGroupByEnrichment": 0.0, "_LogP_MyList": -5.814622086748304, "_TotalGeneInLibrary": 5919.0, "_GeneInGO": 46.0, "shared_name": "26", "Gene": "GO:0034637", "_MEMBER_MyList": 1.0, "STDV_InGO": 1.153969104668408, "_COUNT_001": 12.0, "SUID": 2741891, "id_original": "26", "selected": false, "Log_q_value_": -3.8327044273518935, "canonicalName": "GO:0034637", "CategoryID": 19.0, "Z_score": 6.618227486756458, "EvidenceCutoff": 0.0, "PARENT_GO": "19_GO:0008152 metabolic process", "FirstInGroupByLogP": 0.0, "GROUP_ID": 5.0, "name": "26", "BestEnrichmentInGroup": 11.504373177842565, "CLUSTER_LABEL": " ", "GeneID": "850569|850572|850962|851646|852423|853932|854350|854872|855131|855303|856705|856791", "GiniIndex": 0.0, "_InGO": 4.081632653061225, "_COUNT_001_PCT": 100.0}, "position": {"x": 56.21823252142997, "y": 288.6856227982166}, "selected": false}, {"data": {"id": "2741888", "Enrichment": 10.84065934065934, "BestLogPInGroup": -9.417680144618705, "_PATTERN_": "M1", "_GeneInGOAndHitList": 7.0, "Hits": "GSY1|IGD1|GSY2|GLG1|GAC1|PGM2|GLC3", "Description": "glycogen biosynthetic process", "_GeneInHitList": 294.0, "Category": "GO Biological Processes", "LogP": -6.0354431697141715, "_RANK_": 1.0, "FirstInGroupByEnrichment": 0.0, "_LogP_MyList": -6.0354431697141715, "_TotalGeneInLibrary": 5919.0, "_GeneInGO": 13.0, "shared_name": "25", "Gene": "GO:0005978", "_MEMBER_MyList": 1.0, "STDV_InGO": 0.8891375930475126, "_COUNT_001": 7.0, "SUID": 2741888, "id_original": "25", "selected": false, "Log_q_value_": -4.0326753713446415, "canonicalName": "GO:0005978", "CategoryID": 19.0, "Z_score": 8.11986922847149, "EvidenceCutoff": 0.0, "PARENT_GO": "19_GO:0008152 metabolic process", "FirstInGroupByLogP": 0.0, "GROUP_ID": 5.0, "name": "25", "BestEnrichmentInGroup": 11.504373177842565, "CLUSTER_LABEL": " ", "GeneID": "850569|850572|850962|853932|854350|855131|856705", "GiniIndex": 0.0, "_InGO": 2.380952380952381, "_COUNT_001_PCT": 100.0}, "position": {"x": 148.13902414740653, "y": 328.4074405300739}, "selected": false}, {"data": {"id": "2741885", "Enrichment": 7.190233236151603, "BestLogPInGroup": -9.417680144618705, "_PATTERN_": "M1", "_GeneInGOAndHitList": 10.0, "Hits": "GSY1|IGD1|GSY2|GLG1|GAC1|PCL7|PGM2|GPH1|GLC3|RGI1", "Description": "energy reserve metabolic process", "_GeneInHitList": 294.0, "Category": "GO Biological Processes", "LogP": -6.3333871769918355, "_RANK_": 1.0, "FirstInGroupByEnrichment": 0.0, "_LogP_MyList": -6.3333871769918355, "_TotalGeneInLibrary": 5919.0, "_GeneInGO": 28.0, "shared_name": "24", "Gene": "GO:0006112", "_MEMBER_MyList": 1.0, "STDV_InGO": 1.057153797996807, "_COUNT_001": 10.0, "SUID": 2741885, "id_original": "24", "selected": false, "Log_q_value_": -4.316141555253682, "canonicalName": "GO:0006112", "CategoryID": 19.0, "Z_score": 7.505703411683221, "EvidenceCutoff": 0.0, "PARENT_GO": "19_GO:0008152 metabolic process", "FirstInGroupByLogP": 0.0, "GROUP_ID": 5.0, "name": "24", "BestEnrichmentInGroup": 11.504373177842565, "CLUSTER_LABEL": " ", "GeneID": "850569|850572|850962|853932|854350|854761|855131|856289|856705|856797", "GiniIndex": 0.0, "_InGO": 3.401360544217687, "_COUNT_001_PCT": 100.0}, "position": {"x": 185.1824506610784, "y": 393.4878848660114}, "selected": false}, {"data": {"id": "2741882", "Enrichment": 4.412636287391669, "BestLogPInGroup": -9.417680144618705, "_PATTERN_": "M1", "_GeneInGOAndHitList": 16.0, "Hits": "GLK1|GSY1|HXK1|GSY2|YLR345W|EMI2|VPS73|XKS1|TDH1|MDH1|GLG1|GAC1|PYK2|ALD4|GPH1|GLC3", "Description": "Metabolism of carbohydrates", "_GeneInHitList": 294.0, "Category": "Reactome Gene Sets", "LogP": -6.4338233790088575, "_RANK_": 1.0, "FirstInGroupByEnrichment": 0.0, "_LogP_MyList": -6.4338233790088575, "_TotalGeneInLibrary": 5919.0, "_GeneInGO": 73.0, "shared_name": "23", "Gene": "R-SCE-71387", "_MEMBER_MyList": 1.0, "STDV_InGO": 1.3230047202243043, "_COUNT_001": 16.0, "SUID": 2741882, "id_original": "23", "selected": false, "Log_q_value_": -4.409153739191497, "canonicalName": "R-SCE-71387", "CategoryID": 6.0, "Z_score": 6.706901237770429, "EvidenceCutoff": 0.0, "PARENT_GO": "", "FirstInGroupByLogP": 0.0, "GROUP_ID": 5.0, "name": "23", "BestEnrichmentInGroup": 11.504373177842565, "CLUSTER_LABEL": " ", "GeneID": "850317|850569|850614|850962|851059|852128|852774|853108|853395|853777|853932|854350|854529|854556|856289|856705", "GiniIndex": 0.0, "_InGO": 5.442176870748299, "_COUNT_001_PCT": 100.0}, "position": {"x": 234.45982492865653, "y": 261.8542579281452}, "selected": false}, {"data": {"id": "2741879", "Enrichment": 4.793488824101069, "BestLogPInGroup": -9.417680144618705, "_PATTERN_": "M1", "_GeneInGOAndHitList": 15.0, "Hits": "GSY1|IGD1|GSY2|TPS2|YBR053C|TPS1|TDH1|GLG1|MDH2|GAC1|TSL1|PGM2|TPS3|GLC3|GPP2", "Description": "carbohydrate biosynthetic process", "_GeneInHitList": 294.0, "Category": "GO Biological Processes", "LogP": -6.565423416907936, "_RANK_": 1.0, "FirstInGroupByEnrichment": 0.0, "_LogP_MyList": -6.565423416907936, "_TotalGeneInLibrary": 5919.0, "_GeneInGO": 63.0, "shared_name": "22", "Gene": "GO:0016051", "_MEMBER_MyList": 1.0, "STDV_InGO": 1.2832956896252923, "_COUNT_001": 15.0, "SUID": 2741879, "id_original": "22", "selected": false, "Log_q_value_": -4.525513810533839, "canonicalName": "GO:0016051", "CategoryID": 19.0, "Z_score": 6.920037244128552, "EvidenceCutoff": 0.0, "PARENT_GO": "19_GO:0008152 metabolic process", "FirstInGroupByLogP": 0.0, "GROUP_ID": 5.0, "name": "22", "BestEnrichmentInGroup": 11.504373177842565, "CLUSTER_LABEL": " ", "GeneID": "850569|850572|850962|851646|852342|852423|853395|853932|853994|854350|854872|855131|855303|856705|856791", "GiniIndex": 0.0, "_InGO": 5.1020408163265305, "_COUNT_001_PCT": 100.0}, "position": {"x": 81.8717054218206, "y": 255.832260476363}, "selected": false}, {"data": {"id": "2741876", "Enrichment": 3.9768203577727386, "BestLogPInGroup": -11.663532448682075, "_PATTERN_": "M1", "_GeneInGOAndHitList": 16.0, "Hits": "GLK1|HXK1|GPM2|NTH1|YDR248C|EMI2|NTH2|MRX18|XKS1|TDH1|YJR096W|GCY1|PYK2|GUT2|PGM2|GPH1", "Description": "carbohydrate catabolic process", "_GeneInHitList": 294.0, "Category": "GO Biological Processes", "LogP": -5.7867382591633, "_RANK_": 1.0, "FirstInGroupByEnrichment": 0.0, "_LogP_MyList": -5.7867382591633, "_TotalGeneInLibrary": 5919.0, "_GeneInGO": 81.0, "shared_name": "21", "Gene": "GO:0016052", "_MEMBER_MyList": 1.0, "STDV_InGO": 1.3230047202243043, "_COUNT_001": 16.0, "SUID": 2741876, "id_original": "21", "selected": false, "Log_q_value_": -3.814909599374018, "canonicalName": "GO:0016052", "CategoryID": 19.0, "Z_score": 6.1668441561493115, "EvidenceCutoff": 0.0, "PARENT_GO": "19_GO:0008152 metabolic process", "FirstInGroupByLogP": 0.0, "GROUP_ID": 4.0, "name": "21", "BestEnrichmentInGroup": 4.200337689032448, "CLUSTER_LABEL": " ", "GeneID": "850317|850614|851541|851564|851835|852128|852286|852346|853108|853395|853559|854287|854529|854651|855131|856289", "GiniIndex": 0.0, "_InGO": 5.442176870748299, "_COUNT_001_PCT": 100.0}, "position": {"x": 252.8998578876409, "y": 206.01287113310127}, "selected": false}, {"data": {"id": "2741873", "Enrichment": 4.200337689032448, "BestLogPInGroup": -11.663532448682075, "_PATTERN_": "M1", "_GeneInGOAndHitList": 29.0, "Hits": "GLK1|GSY1|IGD1|HXK1|HSP104|GSY2|YLR345W|NTH1|TPS2|YDR248C|EMI2|NTH2|MRX18|TPS1|ARA1|MAL31|XKS1|GLG1|GCY1|GAC1|GUT2|PCL7|TSL1|PGM2|TPS3|GPH1|GLC3|GPP2|ICL1", "Description": "cellular carbohydrate metabolic process", "_GeneInHitList": 294.0, "Category": "GO Biological Processes", "LogP": -10.704885682398865, "_RANK_": 1.0, "FirstInGroupByEnrichment": 0.0, "_LogP_MyList": -10.704885682398865, "_TotalGeneInLibrary": 5919.0, "_GeneInGO": 139.0, "shared_name": "20", "Gene": "GO:0044262", "_MEMBER_MyList": 1.0, "STDV_InGO": 1.7390054388949427, "_COUNT_001": 29.0, "SUID": 2741873, "id_original": "20", "selected": false, "Log_q_value_": -8.435301988896455, "canonicalName": "GO:0044262", "CategoryID": 19.0, "Z_score": 8.728491015089196, "EvidenceCutoff": 0.0, "PARENT_GO": "19_GO:0008152 metabolic process", "FirstInGroupByLogP": 0.0, "GROUP_ID": 4.0, "name": "20", "BestEnrichmentInGroup": 4.200337689032448, "CLUSTER_LABEL": " ", "GeneID": "850317|850569|850572|850614|850633|850962|851059|851564|851646|851835|852128|852286|852346|852423|852446|852601|853108|853932|854287|854350|854651|854761|854872|855131|855303|856289|856705|856791|856794", "GiniIndex": 0.0, "_InGO": 9.863945578231293, "_COUNT_001_PCT": 100.0}, "position": {"x": 107.28634775580497, "y": 285.4519916641834}, "selected": false}, {"data": {"id": "2741870", "Enrichment": 2.5891216797140144, "BestLogPInGroup": -12.723364984945494, "_PATTERN_": "M1", "_GeneInGOAndHitList": 58.0, "Hits": "GLK1|GSY1|QCR6|HXK1|SDH2|BNA5|GSY2|YLR345W|GPD1|ATP16|KGD2|ATP5|EMI2|QCR7|ATP1|BNA4|COR1|ATP3|VPS73|QCR9|XKS1|LSC2|QCR8|TPK1|YJL163C|YJL070C|TDH1|TES1|ATP2|NNR2|SDH1|SDH3|MDH1|ATP7|GLG1|CYT1|GAC1|PYK2|ALD4|GUT2|CYB2|PLB1|HFD1|NNR1|POR1|HSP82|ATP15|TPK2|YDC1|ATP4|ATP20|GPH1|AIM17|LEU5|CYC7|RIP1|GLC3|GPA2", "Description": "Metabolism", "_GeneInHitList": 294.0, "Category": "Reactome Gene Sets", "LogP": -11.328620880518042, "_RANK_": 1.0, "FirstInGroupByEnrichment": 0.0, "_LogP_MyList": -11.328620880518042, "_TotalGeneInLibrary": 5919.0, "_GeneInGO": 451.0, "shared_name": "19", "Gene": "R-SCE-1430728", "_MEMBER_MyList": 1.0, "STDV_InGO": 2.3208603159181043, "_COUNT_001": 58.0, "SUID": 2741870, "id_original": "19", "selected": false, "Log_q_value_": -9.0029212450367, "canonicalName": "R-SCE-1430728", "CategoryID": 6.0, "Z_score": 8.026589503287495, "EvidenceCutoff": 0.0, "PARENT_GO": "", "FirstInGroupByLogP": 0.0, "GROUP_ID": 3.0, "name": "19", "BestEnrichmentInGroup": 2.5891216797140144, "CLUSTER_LABEL": " ", "GeneID": "850317|850569|850593|850614|850685|850933|850962|851059|851539|851560|851726|851892|852128|852142|852177|852179|852235|852327|852774|853095|853108|853159|853273|853275|853276|853375|853395|853477|853585|853706|853709|853716|853777|853853|853932|854231|854350|854529|854556|854651|854950|855020|855137|855521|855669|855836|855857|855898|856018|856027|856131|856289|856365|856391|856672|856689|856705|856741", "GiniIndex": 0.0, "_InGO": 19.727891156462587, "_COUNT_001_PCT": 100.0}, "position": {"x": 457.92758310736747, "y": 293.26085888029365}, "selected": false}, {"data": {"id": "2741867", "Enrichment": 4.645996860282575, "BestLogPInGroup": -15.616473896518219, "_PATTERN_": "M1", "_GeneInGOAndHitList": 24.0, "Hits": "GLK1|HXK1|ATP14|GPM2|ATP16|KGD2|ATP5|TIM11|ATP17|EMI2|ATP1|ATP3|LSC2|TDH1|TES1|ATP2|ATP7|ATP19|PYK2|ATP18|ATP15|ATP4|ATP20|MCO10", "Description": "purine ribonucleotide metabolic process", "_GeneInHitList": 294.0, "Category": "GO Biological Processes", "LogP": -9.894413233381941, "_RANK_": 1.0, "FirstInGroupByEnrichment": 0.0, "_LogP_MyList": -9.894413233381941, "_TotalGeneInLibrary": 5919.0, "_GeneInGO": 104.0, "shared_name": "18", "Gene": "GO:0009150", "_MEMBER_MyList": 1.0, "STDV_InGO": 1.596858768236636, "_COUNT_001": 24.0, "SUID": 2741867, "id_original": "18", "selected": false, "Log_q_value_": -7.65038364435192, "canonicalName": "GO:0009150", "CategoryID": 19.0, "Z_score": 8.57547100142331, "EvidenceCutoff": 0.0, "PARENT_GO": "19_GO:0008152 metabolic process", "FirstInGroupByLogP": 0.0, "GROUP_ID": 2.0, "name": "18", "BestEnrichmentInGroup": 15.099489795918368, "CLUSTER_LABEL": " ", "GeneID": "850317|850614|851002|851541|851560|851726|851892|851922|851983|852128|852177|852327|853159|853395|853477|853585|853853|854077|854529|854893|855857|856027|856131|1466480", "GiniIndex": 0.0, "_InGO": 8.16326530612245, "_COUNT_001_PCT": 100.0}, "position": {"x": 505.1870893329534, "y": 259.43055950285225}, "selected": false}, {"data": {"id": "2741864", "Enrichment": 8.882052821128452, "BestLogPInGroup": -15.616473896518219, "_PATTERN_": "M1", "_GeneInGOAndHitList": 15.0, "Hits": "ATP14|ATP16|ATP5|TIM11|ATP17|ATP1|ATP3|ATP2|ATP7|ATP19|ATP18|ATP15|ATP4|ATP20|MCO10", "Description": "nucleoside triphosphate biosynthetic process", "_GeneInHitList": 294.0, "Category": "GO Biological Processes", "LogP": -10.813767863555837, "_RANK_": 1.0, "FirstInGroupByEnrichment": 0.0, "_LogP_MyList": -10.813767863555837, "_TotalGeneInLibrary": 5919.0, "_GeneInGO": 34.0, "shared_name": "17", "Gene": "GO:0009142", "_MEMBER_MyList": 1.0, "STDV_InGO": 1.2832956896252923, "_COUNT_001": 15.0, "SUID": 2741864, "id_original": "17", "selected": false, "Log_q_value_": -8.530820208495445, "canonicalName": "GO:0009142", "CategoryID": 19.0, "Z_score": 10.536720211539581, "EvidenceCutoff": 0.0, "PARENT_GO": "19_GO:0008152 metabolic process", "FirstInGroupByLogP": 0.0, "GROUP_ID": 2.0, "name": "17", "BestEnrichmentInGroup": 15.099489795918368, "CLUSTER_LABEL": " ", "GeneID": "851002|851560|851892|851922|851983|852177|852327|853585|853853|854077|854893|855857|856027|856131|1466480", "GiniIndex": 0.0, "_InGO": 5.1020408163265305, "_COUNT_001_PCT": 100.0}, "position": {"x": 546.1365369647893, "y": 299.1124700176838}, "selected": false}, {"data": {"id": "2741861", "Enrichment": 9.741606319947334, "BestLogPInGroup": -15.616473896518219, "_PATTERN_": "M1", "_GeneInGOAndHitList": 15.0, "Hits": "ATP14|ATP16|ATP5|TIM11|ATP17|ATP1|ATP3|ATP2|ATP7|ATP19|ATP18|ATP15|ATP4|ATP20|MCO10", "Description": "ribonucleoside triphosphate metabolic process", "_GeneInHitList": 294.0, "Category": "GO Biological Processes", "LogP": -11.545428797132702, "_RANK_": 1.0, "FirstInGroupByEnrichment": 0.0, "_LogP_MyList": -11.545428797132702, "_TotalGeneInLibrary": 5919.0, "_GeneInGO": 31.0, "shared_name": "16", "Gene": "GO:0009199", "_MEMBER_MyList": 1.0, "STDV_InGO": 1.2832956896252923, "_COUNT_001": 15.0, "SUID": 2741861, "id_original": "16", "selected": false, "Log_q_value_": -9.204489195094624, "canonicalName": "GO:0009199", "CategoryID": 19.0, "Z_score": 11.15547513519781, "EvidenceCutoff": 0.0, "PARENT_GO": "19_GO:0008152 metabolic process", "FirstInGroupByLogP": 0.0, "GROUP_ID": 2.0, "name": "16", "BestEnrichmentInGroup": 15.099489795918368, "CLUSTER_LABEL": " ", "GeneID": "851002|851560|851892|851922|851983|852177|852327|853585|853853|854077|854893|855857|856027|856131|1466480", "GiniIndex": 0.0, "_InGO": 5.1020408163265305, "_COUNT_001_PCT": 100.0}, "position": {"x": 658.5037244647893, "y": 335.4160770046833}, "selected": false}, {"data": {"id": "2741858", "Enrichment": 10.066326530612246, "BestLogPInGroup": -15.616473896518219, "_PATTERN_": "M1", "_GeneInGOAndHitList": 15.0, "Hits": "ATP14|ATP16|ATP5|TIM11|ATP17|ATP1|ATP3|ATP2|ATP7|ATP19|ATP18|ATP15|ATP4|ATP20|MCO10", "Description": "ribonucleoside triphosphate biosynthetic process", "_GeneInHitList": 294.0, "Category": "GO Biological Processes", "LogP": -11.812999972380908, "_RANK_": 1.0, "FirstInGroupByEnrichment": 0.0, "_LogP_MyList": -11.812999972380908, "_TotalGeneInLibrary": 5919.0, "_GeneInGO": 30.0, "shared_name": "15", "Gene": "GO:0009201", "_MEMBER_MyList": 1.0, "STDV_InGO": 1.2832956896252923, "_COUNT_001": 15.0, "SUID": 2741858, "id_original": "15", "selected": false, "Log_q_value_": -9.422842347672647, "canonicalName": "GO:0009201", "CategoryID": 19.0, "Z_score": 11.38075538620108, "EvidenceCutoff": 0.0, "PARENT_GO": "19_GO:0008152 metabolic process", "FirstInGroupByLogP": 0.0, "GROUP_ID": 2.0, "name": "15", "BestEnrichmentInGroup": 15.099489795918368, "CLUSTER_LABEL": " ", "GeneID": "851002|851560|851892|851922|851983|852177|852327|853585|853853|854077|854893|855857|856027|856131|1466480", "GiniIndex": 0.0, "_InGO": 5.1020408163265305, "_COUNT_001_PCT": 100.0}, "position": {"x": 580.6873182147893, "y": 327.4429782498005}, "selected": false}, {"data": {"id": "2741855", "Enrichment": 11.184807256235828, "BestLogPInGroup": -15.616473896518219, "_PATTERN_": "M1", "_GeneInGOAndHitList": 15.0, "Hits": "ATP14|ATP16|ATP5|TIM11|ATP17|ATP1|ATP3|ATP2|ATP7|ATP19|ATP18|ATP15|ATP4|ATP20|MCO10", "Description": "purine nucleoside triphosphate metabolic process", "_GeneInHitList": 294.0, "Category": "GO Biological Processes", "LogP": -12.70449857620623, "_RANK_": 1.0, "FirstInGroupByEnrichment": 0.0, "_LogP_MyList": -12.70449857620623, "_TotalGeneInLibrary": 5919.0, "_GeneInGO": 27.0, "shared_name": "14", "Gene": "GO:0009144", "_MEMBER_MyList": 1.0, "STDV_InGO": 1.2832956896252923, "_COUNT_001": 15.0, "SUID": 2741855, "id_original": "14", "selected": false, "Log_q_value_": -10.238620237559852, "canonicalName": "GO:0009144", "CategoryID": 19.0, "Z_score": 12.125599222883867, "EvidenceCutoff": 0.0, "PARENT_GO": "19_GO:0008152 metabolic process", "FirstInGroupByLogP": 0.0, "GROUP_ID": 2.0, "name": "14", "BestEnrichmentInGroup": 15.099489795918368, "CLUSTER_LABEL": " ", "GeneID": "851002|851560|851892|851922|851983|852177|852327|853585|853853|854077|854893|855857|856027|856131|1466480", "GiniIndex": 0.0, "_InGO": 5.1020408163265305, "_COUNT_001_PCT": 100.0}, "position": {"x": 617.631074318305, "y": 334.5195010769489}, "selected": false}, {"data": {"id": "2741852", "Enrichment": 12.079591836734695, "BestLogPInGroup": -15.616473896518219, "_PATTERN_": "M1", "_GeneInGOAndHitList": 15.0, "Hits": "ATP14|ATP16|ATP5|TIM11|ATP17|ATP1|ATP3|ATP2|ATP7|ATP19|ATP18|ATP15|ATP4|ATP20|MCO10", "Description": "purine ribonucleoside triphosphate metabolic process", "_GeneInHitList": 294.0, "Category": "GO Biological Processes", "LogP": -13.390914263964621, "_RANK_": 1.0, "FirstInGroupByEnrichment": 0.0, "_LogP_MyList": -13.390914263964621, "_TotalGeneInLibrary": 5919.0, "_GeneInGO": 25.0, "shared_name": "13", "Gene": "GO:0009205", "_MEMBER_MyList": 1.0, "STDV_InGO": 1.2832956896252923, "_COUNT_001": 15.0, "SUID": 2741852, "id_original": "13", "selected": false, "Log_q_value_": -10.806936613240248, "canonicalName": "GO:0009205", "CategoryID": 19.0, "Z_score": 12.690787800273196, "EvidenceCutoff": 0.0, "PARENT_GO": "19_GO:0008152 metabolic process", "FirstInGroupByLogP": 0.0, "GROUP_ID": 2.0, "name": "13", "BestEnrichmentInGroup": 15.099489795918368, "CLUSTER_LABEL": " ", "GeneID": "851002|851560|851892|851922|851983|852177|852327|853585|853853|854077|854893|855857|856027|856131|1466480", "GiniIndex": 0.0, "_InGO": 5.1020408163265305, "_COUNT_001_PCT": 100.0}, "position": {"x": 680.7683729022893, "y": 297.564269410384}, "selected": false}, {"data": {"id": "2741849", "Enrichment": 12.582908163265307, "BestLogPInGroup": -15.616473896518219, "_PATTERN_": "M1", "_GeneInGOAndHitList": 15.0, "Hits": "ATP14|ATP16|ATP5|TIM11|ATP17|ATP1|ATP3|ATP2|ATP7|ATP19|ATP18|ATP15|ATP4|ATP20|MCO10", "Description": "purine ribonucleoside triphosphate biosynthetic process", "_GeneInHitList": 294.0, "Category": "GO Biological Processes", "LogP": -13.769179548907788, "_RANK_": 1.0, "FirstInGroupByEnrichment": 0.0, "_LogP_MyList": -13.769179548907788, "_TotalGeneInLibrary": 5919.0, "_GeneInGO": 24.0, "shared_name": "12", "Gene": "GO:0009206", "_MEMBER_MyList": 1.0, "STDV_InGO": 1.2832956896252923, "_COUNT_001": 15.0, "SUID": 2741849, "id_original": "12", "selected": false, "Log_q_value_": -11.095025267834327, "canonicalName": "GO:0009206", "CategoryID": 19.0, "Z_score": 12.998140030399856, "EvidenceCutoff": 0.0, "PARENT_GO": "19_GO:0008152 metabolic process", "FirstInGroupByLogP": 0.0, "GROUP_ID": 2.0, "name": "12", "BestEnrichmentInGroup": 15.099489795918368, "CLUSTER_LABEL": " ", "GeneID": "851002|851560|851892|851922|851983|852177|852327|853585|853853|854077|854893|855857|856027|856131|1466480", "GiniIndex": 0.0, "_InGO": 5.1020408163265305, "_COUNT_001_PCT": 100.0}, "position": {"x": 614.4690870136175, "y": 253.67752491117744}, "selected": false}, {"data": {"id": "2741846", "Enrichment": 12.582908163265307, "BestLogPInGroup": -15.616473896518219, "_PATTERN_": "M1", "_GeneInGOAndHitList": 15.0, "Hits": "ATP14|ATP16|ATP5|TIM11|ATP17|ATP1|ATP3|ATP2|ATP7|ATP19|ATP18|ATP15|ATP4|ATP20|MCO10", "Description": "purine nucleoside triphosphate biosynthetic process", "_GeneInHitList": 294.0, "Category": "GO Biological Processes", "LogP": -13.769179548907788, "_RANK_": 1.0, "FirstInGroupByEnrichment": 0.0, "_LogP_MyList": -13.769179548907788, "_TotalGeneInLibrary": 5919.0, "_GeneInGO": 24.0, "shared_name": "11", "Gene": "GO:0009145", "_MEMBER_MyList": 1.0, "STDV_InGO": 1.2832956896252923, "_COUNT_001": 15.0, "SUID": 2741846, "id_original": "11", "selected": false, "Log_q_value_": -11.095025267834327, "canonicalName": "GO:0009145", "CategoryID": 19.0, "Z_score": 12.998140030399856, "EvidenceCutoff": 0.0, "PARENT_GO": "19_GO:0008152 metabolic process", "FirstInGroupByLogP": 0.0, "GROUP_ID": 2.0, "name": "11", "BestEnrichmentInGroup": 15.099489795918368, "CLUSTER_LABEL": " ", "GeneID": "851002|851560|851892|851922|851983|852177|852327|853585|853853|854077|854893|855857|856027|856131|1466480", "GiniIndex": 0.0, "_InGO": 5.1020408163265305, "_COUNT_001_PCT": 100.0}, "position": {"x": 659.5431531757268, "y": 250.91482578398018}, "selected": false}, {"data": {"id": "2741843", "Enrichment": 15.099489795918368, "BestLogPInGroup": -15.616473896518219, "_PATTERN_": "M1", "_GeneInGOAndHitList": 15.0, "Hits": "ATP14|ATP16|ATP5|TIM11|ATP17|ATP1|ATP3|ATP2|ATP7|ATP19|ATP18|ATP15|ATP4|ATP20|MCO10", "Description": "proton motive force-driven ATP synthesis", "_GeneInHitList": 294.0, "Category": "GO Biological Processes", "LogP": -15.616473896518219, "_RANK_": 1.0, "FirstInGroupByEnrichment": 0.0, "_LogP_MyList": -15.616473896518219, "_TotalGeneInLibrary": 5919.0, "_GeneInGO": 20.0, "shared_name": "10", "Gene": "GO:0015986", "_MEMBER_MyList": 1.0, "STDV_InGO": 1.2832956896252923, "_COUNT_001": 15.0, "SUID": 2741843, "id_original": "10", "selected": false, "Log_q_value_": -12.82837626313792, "canonicalName": "GO:0015986", "CategoryID": 19.0, "Z_score": 14.438732800724484, "EvidenceCutoff": 0.0, "PARENT_GO": "19_GO:0008152 metabolic process", "FirstInGroupByLogP": 0.0, "GROUP_ID": 2.0, "name": "10", "BestEnrichmentInGroup": 15.099489795918368, "CLUSTER_LABEL": " ", "GeneID": "851002|851560|851892|851922|851983|852177|852327|853585|853853|854077|854893|855857|856027|856131|1466480", "GiniIndex": 0.0, "_InGO": 5.1020408163265305, "_COUNT_001_PCT": 100.0}, "position": {"x": 574.322007545844, "y": 265.840805398238}, "selected": false}, {"data": {"id": "2741840", "Enrichment": 8.895823445657333, "BestLogPInGroup": -27.34010239473319, "_PATTERN_": "M1", "_GeneInGOAndHitList": 19.0, "Hits": "QCR6|SDH2|COX8|SDH4|QCR7|COX4|QCR9|QCR8|SDH1|SDH3|CYT1|COX5B|NDI1|COX7|COX5A|QCR2|QCR10|CYC7|RIP1", "Description": "respiratory electron transport chain", "_GeneInHitList": 294.0, "Category": "GO Biological Processes", "LogP": -13.588571934877724, "_RANK_": 1.0, "FirstInGroupByEnrichment": 0.0, "_LogP_MyList": -13.588571934877724, "_TotalGeneInLibrary": 5919.0, "_GeneInGO": 43.0, "shared_name": "9", "Gene": "GO:0022904", "_MEMBER_MyList": 1.0, "STDV_InGO": 1.4339108527393725, "_COUNT_001": 19.0, "SUID": 2741840, "id_original": "9", "selected": false, "Log_q_value_": -10.946602337175664, "canonicalName": "GO:0022904", "CategoryID": 19.0, "Z_score": 11.879294393947582, "EvidenceCutoff": 0.0, "PARENT_GO": "19_GO:0008152 metabolic process", "FirstInGroupByLogP": 0.0, "GROUP_ID": 1.0, "name": "9", "BestEnrichmentInGroup": 12.079591836734693, "CLUSTER_LABEL": " ", "GeneID": "850593|850685|851111|851758|852142|852688|853095|853273|853709|853716|854231|854695|854919|855298|855675|856321|856390|856672|856689", "GiniIndex": 0.0, "_InGO": 6.462585034013605, "_COUNT_001_PCT": 100.0}, "position": {"x": 555.1606000751409, "y": 516.3667834866169}, "selected": false}, {"data": {"id": "2741837", "Enrichment": 11.801900070372975, "BestLogPInGroup": -27.34010239473319, "_PATTERN_": "M1", "_GeneInGOAndHitList": 17.0, "Hits": "QCR6|ATP16|ATP5|QCR7|ATP1|COR1|ATP3|QCR9|QCR8|ATP2|ATP7|CYT1|ATP15|ATP4|ATP20|CYC7|RIP1", "Description": "Respiratory electron transport, ATP synthesis by chemiosmotic coupling, and heat production by uncoupling proteins.", "_GeneInHitList": 294.0, "Category": "Reactome Gene Sets", "LogP": -14.882261927206514, "_RANK_": 1.0, "FirstInGroupByEnrichment": 0.0, "_LogP_MyList": -14.882261927206514, "_TotalGeneInLibrary": 5919.0, "_GeneInGO": 29.0, "shared_name": "8", "Gene": "R-SCE-163200", "_MEMBER_MyList": 1.0, "STDV_InGO": 1.3612671039915658, "_COUNT_001": 17.0, "SUID": 2741837, "id_original": "8", "selected": false, "Log_q_value_": -12.135556978984441, "canonicalName": "R-SCE-163200", "CategoryID": 6.0, "Z_score": 13.330344987406589, "EvidenceCutoff": 0.0, "PARENT_GO": "", "FirstInGroupByLogP": 0.0, "GROUP_ID": 1.0, "name": "8", "BestEnrichmentInGroup": 12.079591836734693, "CLUSTER_LABEL": " ", "GeneID": "850593|851560|851892|852142|852177|852235|852327|853095|853273|853585|853853|854231|855857|856027|856131|856672|856689", "GiniIndex": 0.0, "_InGO": 5.782312925170068, "_COUNT_001_PCT": 100.0}, "position": {"x": 613.4162916034612, "y": 402.9549258816364}, "selected": false}, {"data": {"id": "2741834", "Enrichment": 9.842630385487528, "BestLogPInGroup": -27.34010239473319, "_PATTERN_": "M1", "_GeneInGOAndHitList": 22.0, "Hits": "QCR6|COX12|COX8|SDH4|ATP5|QCR7|COX4|QCR9|QCR8|ATP2|SDH1|SDH3|CYT1|COX5B|NDI1|COX7|COX5A|ATP15|QCR2|QCR10|CYC7|RIP1", "Description": "oxidative phosphorylation", "_GeneInHitList": 294.0, "Category": "GO Biological Processes", "LogP": -16.85525518914923, "_RANK_": 1.0, "FirstInGroupByEnrichment": 0.0, "_LogP_MyList": -16.85525518914923, "_TotalGeneInLibrary": 5919.0, "_GeneInGO": 45.0, "shared_name": "7", "Gene": "GO:0006119", "_MEMBER_MyList": 1.0, "STDV_InGO": 1.5345279277391657, "_COUNT_001": 22.0, "SUID": 2741834, "id_original": "7", "selected": false, "Log_q_value_": -13.970247542760875, "canonicalName": "GO:0006119", "CategoryID": 19.0, "Z_score": 13.611959249661368, "EvidenceCutoff": 0.0, "PARENT_GO": "19_GO:0008152 metabolic process", "FirstInGroupByLogP": 0.0, "GROUP_ID": 1.0, "name": "7", "BestEnrichmentInGroup": 12.079591836734693, "CLUSTER_LABEL": " ", "GeneID": "850593|850727|851111|851758|851892|852142|852688|853095|853273|853585|853709|853716|854231|854695|854919|855298|855675|855857|856321|856390|856672|856689", "GiniIndex": 0.0, "_InGO": 7.482993197278912, "_COUNT_001_PCT": 100.0}, "position": {"x": 585.4698194354925, "y": 483.50795780302315}, "selected": false}, {"data": {"id": "2741831", "Enrichment": 9.116673084328072, "BestLogPInGroup": -27.34010239473319, "_PATTERN_": "M1", "_GeneInGOAndHitList": 24.0, "Hits": "QCR6|SDH2|ATP16|KGD2|ATP5|QCR7|ATP1|COR1|ATP3|QCR9|LSC2|QCR8|ATP2|SDH1|SDH3|MDH1|ATP7|CYT1|POR1|ATP15|ATP4|ATP20|CYC7|RIP1", "Description": "The citric acid (TCA) cycle and respiratory electron transport", "_GeneInHitList": 294.0, "Category": "Reactome Gene Sets", "LogP": -17.367712471322072, "_RANK_": 1.0, "FirstInGroupByEnrichment": 0.0, "_LogP_MyList": -17.367712471322072, "_TotalGeneInLibrary": 5919.0, "_GeneInGO": 53.0, "shared_name": "6", "Gene": "R-SCE-1428517", "_MEMBER_MyList": 1.0, "STDV_InGO": 1.596858768236636, "_COUNT_001": 24.0, "SUID": 2741831, "id_original": "6", "selected": false, "Log_q_value_": -14.424712877956031, "canonicalName": "R-SCE-1428517", "CategoryID": 6.0, "Z_score": 13.568906252018738, "EvidenceCutoff": 0.0, "PARENT_GO": "", "FirstInGroupByLogP": 0.0, "GROUP_ID": 1.0, "name": "6", "BestEnrichmentInGroup": 12.079591836734693, "CLUSTER_LABEL": " ", "GeneID": "850593|850685|851560|851726|851892|852142|852177|852235|852327|853095|853159|853273|853585|853709|853716|853777|853853|854231|855669|855857|856027|856131|856672|856689", "GiniIndex": 0.0, "_InGO": 8.16326530612245, "_COUNT_001_PCT": 100.0}, "position": {"x": 525.5026868671331, "y": 391.0632709133747}, "selected": false}, {"data": {"id": "2741828", "Enrichment": 6.902623906705538, "BestLogPInGroup": -27.34010239473319, "_PATTERN_": "M1", "_GeneInGOAndHitList": 36.0, "Hits": "QCR6|SDH2|COX12|COX8|MIX14|KGD2|SDH4|ATP5|QCR7|COR1|PET9|COX4|QCR9|LSC2|QCR8|ATP2|SDH1|SDH3|MBR1|MDH1|PLN1|MDH2|CYT1|COX5B|NDI1|MIX17|AAC1|ISF1|COX7|COX5A|ATP15|QCR2|QCR10|CYC7|RIP1|ICL1", "Description": "cellular respiration", "_GeneInHitList": 294.0, "Category": "GO Biological Processes", "LogP": -20.947216875935336, "_RANK_": 1.0, "FirstInGroupByEnrichment": 0.0, "_LogP_MyList": -20.947216875935336, "_TotalGeneInLibrary": 5919.0, "_GeneInGO": 105.0, "shared_name": "5", "Gene": "GO:0045333", "_MEMBER_MyList": 1.0, "STDV_InGO": 1.9117896572309039, "_COUNT_001": 36.0, "SUID": 2741828, "id_original": "5", "selected": false, "Log_q_value_": -17.937270492938683, "canonicalName": "GO:0045333", "CategoryID": 19.0, "Z_score": 13.95089694218403, "EvidenceCutoff": 0.0, "PARENT_GO": "19_GO:0008152 metabolic process", "FirstInGroupByLogP": 0.0, "GROUP_ID": 1.0, "name": "5", "BestEnrichmentInGroup": 12.079591836734693, "CLUSTER_LABEL": " ", "GeneID": "850593|850685|850727|851111|851595|851726|851758|851892|852142|852235|852250|852688|853095|853159|853273|853585|853709|853716|853769|853777|853920|853994|854231|854695|854919|855014|855078|855105|855298|855675|855857|856321|856390|856672|856689|856794", "GiniIndex": 0.0, "_InGO": 12.244897959183673, "_COUNT_001_PCT": 100.0}, "position": {"x": 498.2838834613714, "y": 491.60874210478096}, "selected": false}, {"data": {"id": "2741825", "Enrichment": 7.2477551020408155, "BestLogPInGroup": -27.34010239473319, "_PATTERN_": "M1", "_GeneInGOAndHitList": 36.0, "Hits": "QCR6|SDH2|COX12|COX8|MIX14|KGD2|SDH4|ATP5|QCR7|COR1|PET9|COX4|QCR9|LSC2|QCR8|ATP2|SDH1|SDH3|MBR1|MDH1|PLN1|MDH2|CYT1|COX5B|NDI1|MIX17|AAC1|ISF1|COX7|COX5A|ATP15|QCR2|QCR10|CYC7|RIP1|ICL1", "Description": "aerobic respiration", "_GeneInHitList": 294.0, "Category": "GO Biological Processes", "LogP": -21.78578080767872, "_RANK_": 1.0, "FirstInGroupByEnrichment": 0.0, "_LogP_MyList": -21.78578080767872, "_TotalGeneInLibrary": 5919.0, "_GeneInGO": 100.0, "shared_name": "4", "Gene": "GO:0009060", "_MEMBER_MyList": 1.0, "STDV_InGO": 1.9117896572309039, "_COUNT_001": 36.0, "SUID": 2741825, "id_original": "4", "selected": false, "Log_q_value_": -18.696653178634442, "canonicalName": "GO:0009060", "CategoryID": 19.0, "Z_score": 14.404550200745925, "EvidenceCutoff": 0.0, "PARENT_GO": "19_GO:0008152 metabolic process", "FirstInGroupByLogP": 0.0, "GROUP_ID": 1.0, "name": "4", "BestEnrichmentInGroup": 12.079591836734693, "CLUSTER_LABEL": " ", "GeneID": "850593|850685|850727|851111|851595|851726|851758|851892|852142|852235|852250|852688|853095|853159|853273|853585|853709|853716|853769|853777|853920|853994|854231|854695|854919|855014|855078|855105|855298|855675|855857|856321|856390|856672|856689|856794", "GiniIndex": 0.0, "_InGO": 12.244897959183673, "_COUNT_001_PCT": 100.0}, "position": {"x": 517.0405820697698, "y": 446.3941272366169}, "selected": false}, {"data": {"id": "2741822", "Enrichment": 6.3000138831042625, "BestLogPInGroup": -27.34010239473319, "_PATTERN_": "M1", "_GeneInGOAndHitList": 46.0, "Hits": "GSY1|IGD1|QCR6|SDH2|COX12|GSY2|COX8|MIX14|KGD2|SDH4|ATP5|QCR7|COR1|PET9|COX4|QCR9|LSC2|QCR8|ATP2|SDH1|SDH3|MBR1|MDH1|PLN1|GLG1|MDH2|CYT1|GAC1|COX5B|PCL7|NDI1|MIX17|AAC1|ISF1|PGM2|COX7|COX5A|ATP15|GPH1|QCR2|QCR10|CYC7|RIP1|GLC3|ICL1|RGI1", "Description": "energy derivation by oxidation of organic compounds", "_GeneInHitList": 294.0, "Category": "GO Biological Processes", "LogP": -24.8606115080577, "_RANK_": 1.0, "FirstInGroupByEnrichment": 0.0, "_LogP_MyList": -24.8606115080577, "_TotalGeneInLibrary": 5919.0, "_GeneInGO": 147.0, "shared_name": "3", "Gene": "GO:0015980", "_MEMBER_MyList": 1.0, "STDV_InGO": 2.118769734469431, "_COUNT_001": 46.0, "SUID": 2741822, "id_original": "3", "selected": false, "Log_q_value_": -21.674573866005368, "canonicalName": "GO:0015980", "CategoryID": 19.0, "Z_score": 14.87552820028148, "EvidenceCutoff": 0.0, "PARENT_GO": "19_GO:0008152 metabolic process", "FirstInGroupByLogP": 0.0, "GROUP_ID": 1.0, "name": "3", "BestEnrichmentInGroup": 12.079591836734693, "CLUSTER_LABEL": " ", "GeneID": "850569|850572|850593|850685|850727|850962|851111|851595|851726|851758|851892|852142|852235|852250|852688|853095|853159|853273|853585|853709|853716|853769|853777|853920|853932|853994|854231|854350|854695|854761|854919|855014|855078|855105|855131|855298|855675|855857|856289|856321|856390|856672|856689|856705|856794|856797", "GiniIndex": 0.0, "_InGO": 15.646258503401361, "_COUNT_001_PCT": 100.0}, "position": {"x": 438.4908536762151, "y": 468.65154300810127}, "selected": false}, {"data": {"id": "2741819", "Enrichment": 9.271616541353385, "BestLogPInGroup": -27.34010239473319, "_PATTERN_": "M1", "_GeneInGOAndHitList": 35.0, "Hits": "QCR6|SDH2|COX12|ATP14|COX8|ATP16|SDH4|ATP5|TIM11|ATP17|QCR7|ATP1|COR1|ATP3|COX4|QCR9|QCR8|ATP2|SDH1|SDH3|ATP7|ATP19|CYT1|COX5B|ATP18|COX7|COX5A|ATP15|ATP4|ATP20|QCR2|QCR10|CYC7|RIP1|MCO10", "Description": "Oxidative phosphorylation - Saccharomyces cerevisiae (budding yeast)", "_GeneInHitList": 294.0, "Category": "KEGG Pathway", "LogP": -25.55237302196449, "_RANK_": 1.0, "FirstInGroupByEnrichment": 0.0, "_LogP_MyList": -25.55237302196449, "_TotalGeneInLibrary": 5919.0, "_GeneInGO": 76.0, "shared_name": "2", "Gene": "sce00190", "_MEMBER_MyList": 1.0, "STDV_InGO": 1.88869968809044, "_COUNT_001": 35.0, "SUID": 2741819, "id_original": "2", "selected": false, "Log_q_value_": -22.241396643303855, "canonicalName": "sce00190", "CategoryID": 24.0, "Z_score": 16.591250796263914, "EvidenceCutoff": 0.0, "PARENT_GO": "", "FirstInGroupByLogP": 0.0, "GROUP_ID": 1.0, "name": "2", "BestEnrichmentInGroup": 12.079591836734693, "CLUSTER_LABEL": " ", "GeneID": "850593|850685|850727|851002|851111|851560|851758|851892|851922|851983|852142|852177|852235|852327|852688|853095|853273|853585|853709|853716|853853|854077|854231|854695|854893|855298|855675|855857|856027|856131|856321|856390|856672|856689|1466480", "GiniIndex": 0.0, "_InGO": 11.904761904761905, "_COUNT_001_PCT": 100.0}, "position": {"x": 555.1145185321722, "y": 368.57917820097236}, "selected": false}, {"data": {"id": "2741720", "Enrichment": 5.620791463793639, "BestLogPInGroup": -27.34010239473319, "_PATTERN_": "M1", "_GeneInGOAndHitList": 55.0, "Hits": "GLK1|GSY1|IGD1|QCR6|HXK1|SDH2|COX12|GSY2|COX8|GPM2|MIX14|KGD2|SDH4|ATP5|EMI2|QCR7|COR1|PET9|COX4|QCR9|LSC2|SOL4|QCR8|TDH1|ATP2|SDH1|SDH3|MBR1|MDH1|PLN1|GLG1|MDH2|CYT1|GAC1|PYK2|ALD4|COX5B|PCL7|NDI1|MIX17|AAC1|ISF1|PGM2|COX7|YMR315W|COX5A|ATP15|GPH1|QCR2|QCR10|CYC7|RIP1|GLC3|ICL1|RGI1", "Description": "generation of precursor metabolites and energy", "_GeneInHitList": 294.0, "Category": "GO Biological Processes", "LogP": -27.030059723427716, "_RANK_": 1.0, "FirstInGroupByEnrichment": 0.0, "_LogP_MyList": -27.030059723427716, "_TotalGeneInLibrary": 5919.0, "_GeneInGO": 197.0, "shared_name": "1", "Gene": "GO:0006091", "_MEMBER_MyList": 1.0, "STDV_InGO": 2.274360476291944, "_COUNT_001": 55.0, "SUID": 2741720, "id_original": "1", "selected": false, "Log_q_value_": -23.5429920857114, "canonicalName": "GO:0006091", "CategoryID": 19.0, "Z_score": 15.07910057310656, "EvidenceCutoff": 0.0, "PARENT_GO": "19_GO:0008152 metabolic process", "FirstInGroupByLogP": 0.0, "GROUP_ID": 1.0, "name": "1", "BestEnrichmentInGroup": 12.079591836734693, "CLUSTER_LABEL": " ", "GeneID": "850317|850569|850572|850593|850614|850685|850727|850962|851111|851541|851595|851726|851758|851892|852128|852142|852235|852250|852688|853095|853159|853163|853273|853395|853585|853709|853716|853769|853777|853920|853932|853994|854231|854350|854529|854556|854695|854761|854919|855014|855078|855105|855131|855298|855363|855675|855857|856289|856321|856390|856672|856689|856705|856794|856797", "GiniIndex": 0.0, "_InGO": 18.707482993197278, "_COUNT_001_PCT": 100.0}, "position": {"x": 461.46698130072684, "y": 406.434685097945}, "selected": false}], "edges": [{"data": {"id": "2743007", "source": "2742167", "target": "2742083", "SCORE": 0.3514910536839404, "shared_interaction": "Direct", "source_original": "118", "Name_B": "GO:0071462", "Name_A": "GO:0005991", "target_original": "90", "Gene_B": "GO:0071462", "interaction": "Direct", "SUID": 2743007, "TYPE": "Direct", "Gene_A": "GO:0005991", "BEND_MAP_ID": 2743007, "selected": false}, "selected": false}, {"data": {"id": "2743004", "source": "2742167", "target": "2742080", "SCORE": 0.3514910536839404, "shared_interaction": "Direct", "source_original": "118", "Name_B": "GO:0009415", "Name_A": "GO:0005991", "target_original": "89", "Gene_B": "GO:0009415", "interaction": "Direct", "SUID": 2743004, "TYPE": "Direct", "Gene_A": "GO:0005991", "BEND_MAP_ID": 2743004, "selected": false}, "selected": false}, {"data": {"id": "2743001", "source": "2742167", "target": "2742077", "SCORE": 0.3514910536839404, "shared_interaction": "Direct", "source_original": "118", "Name_B": "GO:0042631", "Name_A": "GO:0005991", "target_original": "88", "Gene_B": "GO:0042631", "interaction": "Direct", "SUID": 2743001, "TYPE": "Direct", "Gene_A": "GO:0005991", "BEND_MAP_ID": 2743001, "selected": false}, "selected": false}, {"data": {"id": "2742998", "source": "2742167", "target": "2742074", "SCORE": 0.3514910536839404, "shared_interaction": "Direct", "source_original": "118", "Name_B": "GO:0009414", "Name_A": "GO:0005991", "target_original": "87", "Gene_B": "GO:0009414", "interaction": "Direct", "SUID": 2742998, "TYPE": "Direct", "Gene_A": "GO:0005991", "BEND_MAP_ID": 2742998, "selected": false}, "selected": false}, {"data": {"id": "2742995", "source": "2742167", "target": "2742047", "SCORE": 0.3527777777819663, "shared_interaction": "Direct", "source_original": "118", "Name_B": "GO:0034605", "Name_A": "GO:0005991", "target_original": "78", "Gene_B": "GO:0034605", "interaction": "Direct", "SUID": 2742995, "TYPE": "Direct", "Gene_A": "GO:0005991", "BEND_MAP_ID": 2742995, "selected": false}, "selected": false}, {"data": {"id": "2742992", "source": "2742167", "target": "2742071", "SCORE": 0.3514910536839404, "shared_interaction": "Direct", "source_original": "118", "Name_B": "GO:0071465", "Name_A": "GO:0005991", "target_original": "86", "Gene_B": "GO:0071465", "interaction": "Direct", "SUID": 2742992, "TYPE": "Direct", "Gene_A": "GO:0005991", "BEND_MAP_ID": 2742992, "selected": false}, "selected": false}, {"data": {"id": "2742989", "source": "2742167", "target": "2742155", "SCORE": 0.3514910536839404, "shared_interaction": "Direct", "source_original": "118", "Name_B": "GO:0009269", "Name_A": "GO:0005991", "target_original": "114", "Gene_B": "GO:0009269", "interaction": "Direct", "SUID": 2742989, "TYPE": "Direct", "Gene_A": "GO:0005991", "BEND_MAP_ID": 2742989, "selected": false}, "selected": false}, {"data": {"id": "2742986", "source": "2742167", "target": "2742128", "SCORE": 0.48828696925765885, "shared_interaction": "Direct", "source_original": "118", "Name_B": "WP70", "Name_A": "GO:0005991", "target_original": "105", "Gene_B": "WP70", "interaction": "Direct", "SUID": 2742986, "TYPE": "Direct", "Gene_A": "GO:0005991", "BEND_MAP_ID": 2742986, "selected": false}, "selected": false}, {"data": {"id": "2742983", "source": "2742167", "target": "2741924", "SCORE": 0.7629704984759316, "shared_interaction": "Direct", "source_original": "118", "Name_B": "GO:0046351", "Name_A": "GO:0005991", "target_original": "37", "Gene_B": "GO:0046351", "interaction": "Direct", "SUID": 2742983, "TYPE": "Direct", "Gene_A": "GO:0005991", "BEND_MAP_ID": 2742983, "selected": false}, "selected": false}, {"data": {"id": "2742980", "source": "2742167", "target": "2741921", "SCORE": 0.7629704984759316, "shared_interaction": "Direct", "source_original": "118", "Name_B": "GO:0009312", "Name_A": "GO:0005991", "target_original": "36", "Gene_B": "GO:0009312", "interaction": "Direct", "SUID": 2742980, "TYPE": "Direct", "Gene_A": "GO:0005991", "BEND_MAP_ID": 2742980, "selected": false}, "selected": false}, {"data": {"id": "2742977", "source": "2742167", "target": "2741918", "SCORE": 0.9389572963063721, "shared_interaction": "Direct", "source_original": "118", "Name_B": "GO:0009311", "Name_A": "GO:0005991", "target_original": "35", "Gene_B": "GO:0009311", "interaction": "Direct", "SUID": 2742977, "TYPE": "Direct", "Gene_A": "GO:0005991", "BEND_MAP_ID": 2742977, "selected": false}, "selected": false}, {"data": {"id": "2742974", "source": "2742167", "target": "2741915", "SCORE": 0.7629704984759316, "shared_interaction": "Direct", "source_original": "118", "Name_B": "GO:0005992", "Name_A": "GO:0005991", "target_original": "34", "Gene_B": "GO:0005992", "interaction": "Direct", "SUID": 2742974, "TYPE": "Direct", "Gene_A": "GO:0005991", "BEND_MAP_ID": 2742974, "selected": false}, "selected": false}, {"data": {"id": "2742968", "source": "2742167", "target": "2741909", "SCORE": 0.8528111181312159, "shared_interaction": "Direct", "source_original": "118", "Name_B": "WP398", "Name_A": "GO:0005991", "target_original": "32", "Gene_B": "WP398", "interaction": "Direct", "SUID": 2742968, "TYPE": "Direct", "Gene_A": "GO:0005991", "BEND_MAP_ID": 2742968, "selected": false}, "selected": false}, {"data": {"id": "2742965", "source": "2742167", "target": "2741891", "SCORE": 0.4785138764575091, "shared_interaction": "Direct", "source_original": "118", "Name_B": "GO:0034637", "Name_A": "GO:0005991", "target_original": "26", "Gene_B": "GO:0034637", "interaction": "Direct", "SUID": 2742965, "TYPE": "Direct", "Gene_A": "GO:0005991", "BEND_MAP_ID": 2742965, "selected": false}, "selected": false}, {"data": {"id": "2742962", "source": "2742167", "target": "2741906", "SCORE": 0.7629704984759316, "shared_interaction": "Direct", "source_original": "118", "Name_B": "GO:0070413", "Name_A": "GO:0005991", "target_original": "31", "Gene_B": "GO:0070413", "interaction": "Direct", "SUID": 2742962, "TYPE": "Direct", "Gene_A": "GO:0005991", "BEND_MAP_ID": 2742962, "selected": false}, "selected": false}, {"data": {"id": "2742959", "source": "2742167", "target": "2741879", "SCORE": 0.4082828677503002, "shared_interaction": "Direct", "source_original": "118", "Name_B": "GO:0016051", "Name_A": "GO:0005991", "target_original": "22", "Gene_B": "GO:0016051", "interaction": "Direct", "SUID": 2742959, "TYPE": "Direct", "Gene_A": "GO:0005991", "BEND_MAP_ID": 2742959, "selected": false}, "selected": false}, {"data": {"id": "2742896", "source": "2742167", "target": "2742125", "SCORE": 0.5903496776732848, "shared_interaction": "Direct", "source_original": "118", "Name_B": "sce00500", "Name_A": "GO:0005991", "target_original": "104", "Gene_B": "sce00500", "interaction": "Direct", "SUID": 2742896, "TYPE": "Direct", "Gene_A": "GO:0005991", "BEND_MAP_ID": 2742896, "selected": false}, "selected": false}, {"data": {"id": "2742821", "source": "2742167", "target": "2741873", "SCORE": 0.4001471129108714, "shared_interaction": "Direct", "source_original": "118", "Name_B": "GO:0044262", "Name_A": "GO:0005991", "target_original": "20", "Gene_B": "GO:0044262", "interaction": "Direct", "SUID": 2742821, "TYPE": "Direct", "Gene_A": "GO:0005991", "BEND_MAP_ID": 2742821, "selected": false}, "selected": false}, {"data": {"id": "2742698", "source": "2742164", "target": "2742119", "SCORE": 0.43555258541530173, "shared_interaction": "Direct", "source_original": "117", "Name_B": "sce01100", "Name_A": "GO:0044281", "target_original": "102", "Gene_B": "sce01100", "interaction": "Direct", "SUID": 2742698, "TYPE": "Direct", "Gene_A": "GO:0044281", "BEND_MAP_ID": 2742698, "selected": false}, "selected": false}, {"data": {"id": "2743769", "source": "2742161", "target": "2742050", "SCORE": 0.743593774564687, "shared_interaction": "Direct", "source_original": "116", "Name_B": "GO:0009628", "Name_A": "GO:0009408", "target_original": "79", "Gene_B": "GO:0009628", "interaction": "Direct", "SUID": 2743769, "TYPE": "Direct", "Gene_A": "GO:0009408", "BEND_MAP_ID": 2743769, "selected": false}, "selected": false}, {"data": {"id": "2743766", "source": "2742161", "target": "2742047", "SCORE": 0.8355681016240487, "shared_interaction": "Direct", "source_original": "116", "Name_B": "GO:0034605", "Name_A": "GO:0009408", "target_original": "78", "Gene_B": "GO:0034605", "interaction": "Direct", "SUID": 2743766, "TYPE": "Direct", "Gene_A": "GO:0009408", "BEND_MAP_ID": 2743766, "selected": false}, "selected": false}, {"data": {"id": "2743166", "source": "2742161", "target": "2741906", "SCORE": 0.3559977888377568, "shared_interaction": "Direct", "source_original": "116", "Name_B": "GO:0070413", "Name_A": "GO:0009408", "target_original": "31", "Gene_B": "GO:0070413", "interaction": "Direct", "SUID": 2743166, "TYPE": "Direct", "Gene_A": "GO:0009408", "BEND_MAP_ID": 2743166, "selected": false}, "selected": false}, {"data": {"id": "2743451", "source": "2742158", "target": "2741990", "SCORE": 0.3625170998657409, "shared_interaction": "Direct", "source_original": "115", "Name_B": "GO:0042149", "Name_A": "GO:0009991", "target_original": "59", "Gene_B": "GO:0042149", "interaction": "Direct", "SUID": 2743451, "TYPE": "Direct", "Gene_A": "GO:0009991", "BEND_MAP_ID": 2743451, "selected": false}, "selected": false}, {"data": {"id": "2743448", "source": "2742158", "target": "2741987", "SCORE": 0.8855036855040131, "shared_interaction": "Direct", "source_original": "115", "Name_B": "GO:0031669", "Name_A": "GO:0009991", "target_original": "58", "Gene_B": "GO:0031669", "interaction": "Direct", "SUID": 2743448, "TYPE": "Direct", "Gene_A": "GO:0009991", "BEND_MAP_ID": 2743448, "selected": false}, "selected": false}, {"data": {"id": "2743445", "source": "2742158", "target": "2741984", "SCORE": 0.9160965070222008, "shared_interaction": "Direct", "source_original": "115", "Name_B": "GO:0031667", "Name_A": "GO:0009991", "target_original": "57", "Gene_B": "GO:0031667", "interaction": "Direct", "SUID": 2743445, "TYPE": "Direct", "Gene_A": "GO:0009991", "BEND_MAP_ID": 2743445, "selected": false}, "selected": false}, {"data": {"id": "2743439", "source": "2742158", "target": "2741978", "SCORE": 0.7844588344132445, "shared_interaction": "Direct", "source_original": "115", "Name_B": "GO:0042594", "Name_A": "GO:0009991", "target_original": "55", "Gene_B": "GO:0042594", "interaction": "Direct", "SUID": 2743439, "TYPE": "Direct", "Gene_A": "GO:0009991", "BEND_MAP_ID": 2743439, "selected": false}, "selected": false}, {"data": {"id": "2743433", "source": "2742158", "target": "2741972", "SCORE": 0.9732583495926344, "shared_interaction": "Direct", "source_original": "115", "Name_B": "GO:0071496", "Name_A": "GO:0009991", "target_original": "53", "Gene_B": "GO:0071496", "interaction": "Direct", "SUID": 2743433, "TYPE": "Direct", "Gene_A": "GO:0009991", "BEND_MAP_ID": 2743433, "selected": false}, "selected": false}, {"data": {"id": "2743430", "source": "2742158", "target": "2741969", "SCORE": 0.9732583495926344, "shared_interaction": "Direct", "source_original": "115", "Name_B": "GO:0031668", "Name_A": "GO:0009991", "target_original": "52", "Gene_B": "GO:0031668", "interaction": "Direct", "SUID": 2743430, "TYPE": "Direct", "Gene_A": "GO:0009991", "BEND_MAP_ID": 2743430, "selected": false}, "selected": false}, {"data": {"id": "2743859", "source": "2742155", "target": "2742083", "SCORE": 1.0, "shared_interaction": "Direct", "source_original": "114", "Name_B": "GO:0071462", "Name_A": "GO:0009269", "target_original": "90", "Gene_B": "GO:0071462", "interaction": "Direct", "SUID": 2743859, "TYPE": "Direct", "Gene_A": "GO:0009269", "BEND_MAP_ID": 2743859, "selected": false}, "selected": false}, {"data": {"id": "2743856", "source": "2742155", "target": "2742080", "SCORE": 1.0, "shared_interaction": "Direct", "source_original": "114", "Name_B": "GO:0009415", "Name_A": "GO:0009269", "target_original": "89", "Gene_B": "GO:0009415", "interaction": "Direct", "SUID": 2743856, "TYPE": "Direct", "Gene_A": "GO:0009269", "BEND_MAP_ID": 2743856, "selected": false}, "selected": false}, {"data": {"id": "2743853", "source": "2742155", "target": "2742077", "SCORE": 1.0, "shared_interaction": "Direct", "source_original": "114", "Name_B": "GO:0042631", "Name_A": "GO:0009269", "target_original": "88", "Gene_B": "GO:0042631", "interaction": "Direct", "SUID": 2743853, "TYPE": "Direct", "Gene_A": "GO:0009269", "BEND_MAP_ID": 2743853, "selected": false}, "selected": false}, {"data": {"id": "2743850", "source": "2742155", "target": "2742074", "SCORE": 1.0, "shared_interaction": "Direct", "source_original": "114", "Name_B": "GO:0009414", "Name_A": "GO:0009269", "target_original": "87", "Gene_B": "GO:0009414", "interaction": "Direct", "SUID": 2743850, "TYPE": "Direct", "Gene_A": "GO:0009269", "BEND_MAP_ID": 2743850, "selected": false}, "selected": false}, {"data": {"id": "2743847", "source": "2742155", "target": "2742071", "SCORE": 1.0, "shared_interaction": "Direct", "source_original": "114", "Name_B": "GO:0071465", "Name_A": "GO:0009269", "target_original": "86", "Gene_B": "GO:0071465", "interaction": "Direct", "SUID": 2743847, "TYPE": "Direct", "Gene_A": "GO:0009269", "BEND_MAP_ID": 2743847, "selected": false}, "selected": false}, {"data": {"id": "2743637", "source": "2742155", "target": "2741918", "SCORE": 0.3202042304944703, "shared_interaction": "Direct", "source_original": "114", "Name_B": "GO:0009311", "Name_A": "GO:0009269", "target_original": "35", "Gene_B": "GO:0009311", "interaction": "Direct", "SUID": 2743637, "TYPE": "Direct", "Gene_A": "GO:0009269", "BEND_MAP_ID": 2743637, "selected": false}, "selected": false}, {"data": {"id": "2743472", "source": "2742155", "target": "2741909", "SCORE": 0.43474041727955826, "shared_interaction": "Direct", "source_original": "114", "Name_B": "WP398", "Name_A": "GO:0009269", "target_original": "32", "Gene_B": "WP398", "interaction": "Direct", "SUID": 2743472, "TYPE": "Direct", "Gene_A": "GO:0009269", "BEND_MAP_ID": 2743472, "selected": false}, "selected": false}, {"data": {"id": "2743910", "source": "2742152", "target": "2742107", "SCORE": 0.606019614477234, "shared_interaction": "Direct", "source_original": "113", "Name_B": "GO:0062014", "Name_A": "GO:0006109", "target_original": "98", "Gene_B": "GO:0062014", "interaction": "Direct", "SUID": 2743910, "TYPE": "Direct", "Gene_A": "GO:0006109", "BEND_MAP_ID": 2743910, "selected": false}, "selected": false}, {"data": {"id": "2743907", "source": "2742152", "target": "2742104", "SCORE": 0.606019614477234, "shared_interaction": "Direct", "source_original": "113", "Name_B": "GO:0006111", "Name_A": "GO:0006109", "target_original": "97", "Gene_B": "GO:0006111", "interaction": "Direct", "SUID": 2743907, "TYPE": "Direct", "Gene_A": "GO:0006109", "BEND_MAP_ID": 2743907, "selected": false}, "selected": false}, {"data": {"id": "2743904", "source": "2742152", "target": "2742101", "SCORE": 0.7936226749349893, "shared_interaction": "Direct", "source_original": "113", "Name_B": "GO:0043255", "Name_A": "GO:0006109", "target_original": "96", "Gene_B": "GO:0043255", "interaction": "Direct", "SUID": 2743904, "TYPE": "Direct", "Gene_A": "GO:0006109", "BEND_MAP_ID": 2743904, "selected": false}, "selected": false}, {"data": {"id": "2743901", "source": "2742152", "target": "2742098", "SCORE": 0.606019614477234, "shared_interaction": "Direct", "source_original": "113", "Name_B": "GO:0045721", "Name_A": "GO:0006109", "target_original": "95", "Gene_B": "GO:0045721", "interaction": "Direct", "SUID": 2743901, "TYPE": "Direct", "Gene_A": "GO:0006109", "BEND_MAP_ID": 2743901, "selected": false}, "selected": false}, {"data": {"id": "2743898", "source": "2742152", "target": "2742095", "SCORE": 0.706179066836963, "shared_interaction": "Direct", "source_original": "113", "Name_B": "GO:0045912", "Name_A": "GO:0006109", "target_original": "94", "Gene_B": "GO:0045912", "interaction": "Direct", "SUID": 2743898, "TYPE": "Direct", "Gene_A": "GO:0006109", "BEND_MAP_ID": 2743898, "selected": false}, "selected": false}, {"data": {"id": "2743895", "source": "2742152", "target": "2742092", "SCORE": 0.706179066836963, "shared_interaction": "Direct", "source_original": "113", "Name_B": "GO:0010677", "Name_A": "GO:0006109", "target_original": "93", "Gene_B": "GO:0010677", "interaction": "Direct", "SUID": 2743895, "TYPE": "Direct", "Gene_A": "GO:0006109", "BEND_MAP_ID": 2743895, "selected": false}, "selected": false}, {"data": {"id": "2743892", "source": "2742152", "target": "2742089", "SCORE": 0.9389572963063721, "shared_interaction": "Direct", "source_original": "113", "Name_B": "GO:0010675", "Name_A": "GO:0006109", "target_original": "92", "Gene_B": "GO:0010675", "interaction": "Direct", "SUID": 2743892, "TYPE": "Direct", "Gene_A": "GO:0006109", "BEND_MAP_ID": 2743892, "selected": false}, "selected": false}, {"data": {"id": "2743889", "source": "2742152", "target": "2742086", "SCORE": 0.9389572963063721, "shared_interaction": "Direct", "source_original": "113", "Name_B": "GO:0010906", "Name_A": "GO:0006109", "target_original": "91", "Gene_B": "GO:0010906", "interaction": "Direct", "SUID": 2743889, "TYPE": "Direct", "Gene_A": "GO:0006109", "BEND_MAP_ID": 2743889, "selected": false}, "selected": false}, {"data": {"id": "2743808", "source": "2742149", "target": "2742068", "SCORE": 0.7456331877761606, "shared_interaction": "Direct", "source_original": "112", "Name_B": "R-SCE-418594", "Name_A": "GO:0010737", "target_original": "85", "Gene_B": "R-SCE-418594", "interaction": "Direct", "SUID": 2743808, "TYPE": "Direct", "Gene_A": "GO:0010737", "BEND_MAP_ID": 2743808, "selected": false}, "selected": false}, {"data": {"id": "2743805", "source": "2742149", "target": "2742065", "SCORE": 0.7456331877761606, "shared_interaction": "Direct", "source_original": "112", "Name_B": "R-SCE-112043", "Name_A": "GO:0010737", "target_original": "84", "Gene_B": "R-SCE-112043", "interaction": "Direct", "SUID": 2743805, "TYPE": "Direct", "Gene_A": "GO:0010737", "BEND_MAP_ID": 2743805, "selected": false}, "selected": false}, {"data": {"id": "2743802", "source": "2742149", "target": "2742062", "SCORE": 0.7456331877761606, "shared_interaction": "Direct", "source_original": "112", "Name_B": "R-SCE-112040", "Name_A": "GO:0010737", "target_original": "83", "Gene_B": "R-SCE-112040", "interaction": "Direct", "SUID": 2743802, "TYPE": "Direct", "Gene_A": "GO:0010737", "BEND_MAP_ID": 2743802, "selected": false}, "selected": false}, {"data": {"id": "2743799", "source": "2742149", "target": "2742059", "SCORE": 0.7456331877761606, "shared_interaction": "Direct", "source_original": "112", "Name_B": "R-SCE-111885", "Name_A": "GO:0010737", "target_original": "82", "Gene_B": "R-SCE-111885", "interaction": "Direct", "SUID": 2743799, "TYPE": "Direct", "Gene_A": "GO:0010737", "BEND_MAP_ID": 2743799, "selected": false}, "selected": false}, {"data": {"id": "2743796", "source": "2742149", "target": "2742056", "SCORE": 0.5650280024954177, "shared_interaction": "Direct", "source_original": "112", "Name_B": "R-SCE-445717", "Name_A": "GO:0010737", "target_original": "81", "Gene_B": "R-SCE-445717", "interaction": "Direct", "SUID": 2743796, "TYPE": "Direct", "Gene_A": "GO:0010737", "BEND_MAP_ID": 2743796, "selected": false}, "selected": false}, {"data": {"id": "2743793", "source": "2742149", "target": "2742053", "SCORE": 0.5650280024954177, "shared_interaction": "Direct", "source_original": "112", "Name_B": "R-SCE-432040", "Name_A": "GO:0010737", "target_original": "80", "Gene_B": "R-SCE-432040", "interaction": "Direct", "SUID": 2743793, "TYPE": "Direct", "Gene_A": "GO:0010737", "BEND_MAP_ID": 2743793, "selected": false}, "selected": false}, {"data": {"id": "2743067", "source": "2742146", "target": "2741957", "SCORE": 0.48918646678982647, "shared_interaction": "Direct", "source_original": "111", "Name_B": "GO:0016310", "Name_A": "GO:0006793", "target_original": "48", "Gene_B": "GO:0016310", "interaction": "Direct", "SUID": 2743067, "TYPE": "Direct", "Gene_A": "GO:0006793", "BEND_MAP_ID": 2743067, "selected": false}, "selected": false}, {"data": {"id": "2743064", "source": "2742146", "target": "2742164", "SCORE": 0.5299995308913521, "shared_interaction": "Direct", "source_original": "111", "Name_B": "GO:0044281", "Name_A": "GO:0006793", "target_original": "117", "Gene_B": "GO:0044281", "interaction": "Direct", "SUID": 2743064, "TYPE": "Direct", "Gene_A": "GO:0006793", "BEND_MAP_ID": 2743064, "selected": false}, "selected": false}, {"data": {"id": "2743061", "source": "2742146", "target": "2741954", "SCORE": 0.9546251217137808, "shared_interaction": "Direct", "source_original": "111", "Name_B": "GO:0006796", "Name_A": "GO:0006793", "target_original": "47", "Gene_B": "GO:0006796", "interaction": "Direct", "SUID": 2743061, "TYPE": "Direct", "Gene_A": "GO:0006793", "BEND_MAP_ID": 2743061, "selected": false}, "selected": false}, {"data": {"id": "2742866", "source": "2742146", "target": "2741867", "SCORE": 0.48918646678982647, "shared_interaction": "Direct", "source_original": "111", "Name_B": "GO:0009150", "Name_A": "GO:0006793", "target_original": "18", "Gene_B": "GO:0009150", "interaction": "Direct", "SUID": 2742866, "TYPE": "Direct", "Gene_A": "GO:0006793", "BEND_MAP_ID": 2742866, "selected": false}, "selected": false}, {"data": {"id": "2742809", "source": "2742146", "target": "2741864", "SCORE": 0.32497795692251713, "shared_interaction": "Direct", "source_original": "111", "Name_B": "GO:0009142", "Name_A": "GO:0006793", "target_original": "17", "Gene_B": "GO:0009142", "interaction": "Direct", "SUID": 2742809, "TYPE": "Direct", "Gene_A": "GO:0006793", "BEND_MAP_ID": 2742809, "selected": false}, "selected": false}, {"data": {"id": "2742788", "source": "2742146", "target": "2741861", "SCORE": 0.32497795692251713, "shared_interaction": "Direct", "source_original": "111", "Name_B": "GO:0009199", "Name_A": "GO:0006793", "target_original": "16", "Gene_B": "GO:0009199", "interaction": "Direct", "SUID": 2742788, "TYPE": "Direct", "Gene_A": "GO:0006793", "BEND_MAP_ID": 2742788, "selected": false}, "selected": false}, {"data": {"id": "2742731", "source": "2742146", "target": "2741858", "SCORE": 0.32497795692251713, "shared_interaction": "Direct", "source_original": "111", "Name_B": "GO:0009201", "Name_A": "GO:0006793", "target_original": "15", "Gene_B": "GO:0009201", "interaction": "Direct", "SUID": 2742731, "TYPE": "Direct", "Gene_A": "GO:0006793", "BEND_MAP_ID": 2742731, "selected": false}, "selected": false}, {"data": {"id": "2742713", "source": "2742146", "target": "2741855", "SCORE": 0.32497795692251713, "shared_interaction": "Direct", "source_original": "111", "Name_B": "GO:0009144", "Name_A": "GO:0006793", "target_original": "14", "Gene_B": "GO:0009144", "interaction": "Direct", "SUID": 2742713, "TYPE": "Direct", "Gene_A": "GO:0006793", "BEND_MAP_ID": 2742713, "selected": false}, "selected": false}, {"data": {"id": "2742680", "source": "2742146", "target": "2741852", "SCORE": 0.32497795692251713, "shared_interaction": "Direct", "source_original": "111", "Name_B": "GO:0009205", "Name_A": "GO:0006793", "target_original": "13", "Gene_B": "GO:0009205", "interaction": "Direct", "SUID": 2742680, "TYPE": "Direct", "Gene_A": "GO:0006793", "BEND_MAP_ID": 2742680, "selected": false}, "selected": false}, {"data": {"id": "2742650", "source": "2742146", "target": "2741849", "SCORE": 0.32497795692251713, "shared_interaction": "Direct", "source_original": "111", "Name_B": "GO:0009206", "Name_A": "GO:0006793", "target_original": "12", "Gene_B": "GO:0009206", "interaction": "Direct", "SUID": 2742650, "TYPE": "Direct", "Gene_A": "GO:0006793", "BEND_MAP_ID": 2742650, "selected": false}, "selected": false}, {"data": {"id": "2742623", "source": "2742146", "target": "2741846", "SCORE": 0.32497795692251713, "shared_interaction": "Direct", "source_original": "111", "Name_B": "GO:0009145", "Name_A": "GO:0006793", "target_original": "11", "Gene_B": "GO:0009145", "interaction": "Direct", "SUID": 2742623, "TYPE": "Direct", "Gene_A": "GO:0006793", "BEND_MAP_ID": 2742623, "selected": false}, "selected": false}, {"data": {"id": "2742563", "source": "2742146", "target": "2741843", "SCORE": 0.32497795692251713, "shared_interaction": "Direct", "source_original": "111", "Name_B": "GO:0015986", "Name_A": "GO:0006793", "target_original": "10", "Gene_B": "GO:0015986", "interaction": "Direct", "SUID": 2742563, "TYPE": "Direct", "Gene_A": "GO:0006793", "BEND_MAP_ID": 2742563, "selected": false}, "selected": false}, {"data": {"id": "2743754", "source": "2742143", "target": "2742041", "SCORE": 0.4246913580280011, "shared_interaction": "Direct", "source_original": "110", "Name_B": "GO:0046031", "Name_A": "GO:0005996", "target_original": "76", "Gene_B": "GO:0046031", "interaction": "Direct", "SUID": 2743754, "TYPE": "Direct", "Gene_A": "GO:0005996", "BEND_MAP_ID": 2743754, "selected": false}, "selected": false}, {"data": {"id": "2743751", "source": "2742143", "target": "2742038", "SCORE": 0.4246913580280011, "shared_interaction": "Direct", "source_original": "110", "Name_B": "GO:0006165", "Name_A": "GO:0005996", "target_original": "75", "Gene_B": "GO:0006165", "interaction": "Direct", "SUID": 2743751, "TYPE": "Direct", "Gene_A": "GO:0005996", "BEND_MAP_ID": 2743751, "selected": false}, "selected": false}, {"data": {"id": "2743748", "source": "2742143", "target": "2742035", "SCORE": 0.4246913580280011, "shared_interaction": "Direct", "source_original": "110", "Name_B": "GO:0006757", "Name_A": "GO:0005996", "target_original": "74", "Gene_B": "GO:0006757", "interaction": "Direct", "SUID": 2743748, "TYPE": "Direct", "Gene_A": "GO:0005996", "BEND_MAP_ID": 2743748, "selected": false}, "selected": false}, {"data": {"id": "2743745", "source": "2742143", "target": "2742032", "SCORE": 0.4246913580280011, "shared_interaction": "Direct", "source_original": "110", "Name_B": "GO:0006096", "Name_A": "GO:0005996", "target_original": "73", "Gene_B": "GO:0006096", "interaction": "Direct", "SUID": 2743745, "TYPE": "Direct", "Gene_A": "GO:0005996", "BEND_MAP_ID": 2743745, "selected": false}, "selected": false}, {"data": {"id": "2743742", "source": "2742143", "target": "2741957", "SCORE": 0.3438300051222846, "shared_interaction": "Direct", "source_original": "110", "Name_B": "GO:0016310", "Name_A": "GO:0005996", "target_original": "48", "Gene_B": "GO:0016310", "interaction": "Direct", "SUID": 2743742, "TYPE": "Direct", "Gene_A": "GO:0005996", "BEND_MAP_ID": 2743742, "selected": false}, "selected": false}, {"data": {"id": "2743739", "source": "2742143", "target": "2742029", "SCORE": 0.3873794916781408, "shared_interaction": "Direct", "source_original": "110", "Name_B": "R-SCE-170822", "Name_A": "GO:0005996", "target_original": "72", "Gene_B": "R-SCE-170822", "interaction": "Direct", "SUID": 2743739, "TYPE": "Direct", "Gene_A": "GO:0005996", "BEND_MAP_ID": 2743739, "selected": false}, "selected": false}, {"data": {"id": "2743736", "source": "2742143", "target": "2742026", "SCORE": 0.5397530864224009, "shared_interaction": "Direct", "source_original": "110", "Name_B": "R-SCE-70171", "Name_A": "GO:0005996", "target_original": "71", "Gene_B": "R-SCE-70171", "interaction": "Direct", "SUID": 2743736, "TYPE": "Direct", "Gene_A": "GO:0005996", "BEND_MAP_ID": 2743736, "selected": false}, "selected": false}, {"data": {"id": "2743733", "source": "2742143", "target": "2742023", "SCORE": 0.5076308992748236, "shared_interaction": "Direct", "source_original": "110", "Name_B": "R-SCE-70326", "Name_A": "GO:0005996", "target_original": "70", "Gene_B": "R-SCE-70326", "interaction": "Direct", "SUID": 2743733, "TYPE": "Direct", "Gene_A": "GO:0005996", "BEND_MAP_ID": 2743733, "selected": false}, "selected": false}, {"data": {"id": "2743730", "source": "2742143", "target": "2742020", "SCORE": 0.5753709971387387, "shared_interaction": "Direct", "source_original": "110", "Name_B": "GO:0046835", "Name_A": "GO:0005996", "target_original": "69", "Gene_B": "GO:0046835", "interaction": "Direct", "SUID": 2743730, "TYPE": "Direct", "Gene_A": "GO:0005996", "BEND_MAP_ID": 2743730, "selected": false}, "selected": false}, {"data": {"id": "2743727", "source": "2742143", "target": "2742017", "SCORE": 0.42775276299904486, "shared_interaction": "Direct", "source_original": "110", "Name_B": "sce00010", "Name_A": "GO:0005996", "target_original": "68", "Gene_B": "sce00010", "interaction": "Direct", "SUID": 2743727, "TYPE": "Direct", "Gene_A": "GO:0005996", "BEND_MAP_ID": 2743727, "selected": false}, "selected": false}, {"data": {"id": "2743319", "source": "2742143", "target": "2741876", "SCORE": 0.5446254071678518, "shared_interaction": "Direct", "source_original": "110", "Name_B": "GO:0016052", "Name_A": "GO:0005996", "target_original": "21", "Gene_B": "GO:0016052", "interaction": "Direct", "SUID": 2743319, "TYPE": "Direct", "Gene_A": "GO:0005996", "BEND_MAP_ID": 2743319, "selected": false}, "selected": false}, {"data": {"id": "2743190", "source": "2742143", "target": "2741882", "SCORE": 0.39283387622380234, "shared_interaction": "Direct", "source_original": "110", "Name_B": "R-SCE-71387", "Name_A": "GO:0005996", "target_original": "23", "Gene_B": "R-SCE-71387", "interaction": "Direct", "SUID": 2743190, "TYPE": "Direct", "Gene_A": "GO:0005996", "BEND_MAP_ID": 2743190, "selected": false}, "selected": false}, {"data": {"id": "2742662", "source": "2742140", "target": "2741840", "SCORE": 0.45881722351484877, "shared_interaction": "Direct", "source_original": "109", "Name_B": "GO:0022904", "Name_A": "GO:0006123", "target_original": "9", "Gene_B": "GO:0022904", "interaction": "Direct", "SUID": 2742662, "TYPE": "Direct", "Gene_A": "GO:0006123", "BEND_MAP_ID": 2742662, "selected": false}, "selected": false}, {"data": {"id": "2743724", "source": "2742134", "target": "2742029", "SCORE": 0.3202042304944703, "shared_interaction": "Direct", "source_original": "107", "Name_B": "R-SCE-170822", "Name_A": "GO:0034219", "target_original": "72", "Gene_B": "R-SCE-170822", "interaction": "Direct", "SUID": 2743724, "TYPE": "Direct", "Gene_A": "GO:0034219", "BEND_MAP_ID": 2743724, "selected": false}, "selected": false}, {"data": {"id": "2743721", "source": "2742134", "target": "2742014", "SCORE": 0.7936226749349893, "shared_interaction": "Direct", "source_original": "107", "Name_B": "GO:1904659", "Name_A": "GO:0034219", "target_original": "67", "Gene_B": "GO:1904659", "interaction": "Direct", "SUID": 2743721, "TYPE": "Direct", "Gene_A": "GO:0034219", "BEND_MAP_ID": 2743721, "selected": false}, "selected": false}, {"data": {"id": "2742779", "source": "2742131", "target": "2741903", "SCORE": 0.355760368664978, "shared_interaction": "Direct", "source_original": "106", "Name_B": "GO:0044042", "Name_A": "GO:0005975", "target_original": "30", "Gene_B": "GO:0044042", "interaction": "Direct", "SUID": 2742779, "TYPE": "Direct", "Gene_A": "GO:0005975", "BEND_MAP_ID": 2742779, "selected": false}, "selected": false}, {"data": {"id": "2742776", "source": "2742131", "target": "2741900", "SCORE": 0.355760368664978, "shared_interaction": "Direct", "source_original": "106", "Name_B": "GO:0006073", "Name_A": "GO:0005975", "target_original": "29", "Gene_B": "GO:0006073", "interaction": "Direct", "SUID": 2742776, "TYPE": "Direct", "Gene_A": "GO:0005975", "BEND_MAP_ID": 2742776, "selected": false}, "selected": false}, {"data": {"id": "2742773", "source": "2742131", "target": "2742143", "SCORE": 0.41520258157167655, "shared_interaction": "Direct", "source_original": "106", "Name_B": "GO:0005996", "Name_A": "GO:0005975", "target_original": "110", "Gene_B": "GO:0005996", "interaction": "Direct", "SUID": 2742773, "TYPE": "Direct", "Gene_A": "GO:0005975", "BEND_MAP_ID": 2742773, "selected": false}, "selected": false}, {"data": {"id": "2742770", "source": "2742131", "target": "2741918", "SCORE": 0.3247639525115202, "shared_interaction": "Direct", "source_original": "106", "Name_B": "GO:0009311", "Name_A": "GO:0005975", "target_original": "35", "Gene_B": "GO:0009311", "interaction": "Direct", "SUID": 2742770, "TYPE": "Direct", "Gene_A": "GO:0005975", "BEND_MAP_ID": 2742770, "selected": false}, "selected": false}, {"data": {"id": "2742767", "source": "2742131", "target": "2741912", "SCORE": 0.3247639525115202, "shared_interaction": "Direct", "source_original": "106", "Name_B": "GO:0005984", "Name_A": "GO:0005975", "target_original": "33", "Gene_B": "GO:0005984", "interaction": "Direct", "SUID": 2742767, "TYPE": "Direct", "Gene_A": "GO:0005975", "BEND_MAP_ID": 2742767, "selected": false}, "selected": false}, {"data": {"id": "2742764", "source": "2742131", "target": "2741894", "SCORE": 0.3247639525115202, "shared_interaction": "Direct", "source_original": "106", "Name_B": "GO:0005977", "Name_A": "GO:0005975", "target_original": "27", "Gene_B": "GO:0005977", "interaction": "Direct", "SUID": 2742764, "TYPE": "Direct", "Gene_A": "GO:0005975", "BEND_MAP_ID": 2742764, "selected": false}, "selected": false}, {"data": {"id": "2742761", "source": "2742131", "target": "2741876", "SCORE": 0.5248130523462842, "shared_interaction": "Direct", "source_original": "106", "Name_B": "GO:0016052", "Name_A": "GO:0005975", "target_original": "21", "Gene_B": "GO:0016052", "interaction": "Direct", "SUID": 2742761, "TYPE": "Direct", "Gene_A": "GO:0005975", "BEND_MAP_ID": 2742761, "selected": false}, "selected": false}, {"data": {"id": "2742758", "source": "2742131", "target": "2741891", "SCORE": 0.41520258157167655, "shared_interaction": "Direct", "source_original": "106", "Name_B": "GO:0034637", "Name_A": "GO:0005975", "target_original": "26", "Gene_B": "GO:0034637", "interaction": "Direct", "SUID": 2742758, "TYPE": "Direct", "Gene_A": "GO:0005975", "BEND_MAP_ID": 2742758, "selected": false}, "selected": false}, {"data": {"id": "2742755", "source": "2742131", "target": "2741885", "SCORE": 0.3128110599093099, "shared_interaction": "Direct", "source_original": "106", "Name_B": "GO:0006112", "Name_A": "GO:0005975", "target_original": "24", "Gene_B": "GO:0006112", "interaction": "Direct", "SUID": 2742755, "TYPE": "Direct", "Gene_A": "GO:0005975", "BEND_MAP_ID": 2742755, "selected": false}, "selected": false}, {"data": {"id": "2742752", "source": "2742131", "target": "2741882", "SCORE": 0.40601631543285516, "shared_interaction": "Direct", "source_original": "106", "Name_B": "R-SCE-71387", "Name_A": "GO:0005975", "target_original": "23", "Gene_B": "R-SCE-71387", "interaction": "Direct", "SUID": 2742752, "TYPE": "Direct", "Gene_A": "GO:0005975", "BEND_MAP_ID": 2742752, "selected": false}, "selected": false}, {"data": {"id": "2742749", "source": "2742131", "target": "2741879", "SCORE": 0.49849332759463505, "shared_interaction": "Direct", "source_original": "106", "Name_B": "GO:0016051", "Name_A": "GO:0005975", "target_original": "22", "Gene_B": "GO:0016051", "interaction": "Direct", "SUID": 2742749, "TYPE": "Direct", "Gene_A": "GO:0005975", "BEND_MAP_ID": 2742749, "selected": false}, "selected": false}, {"data": {"id": "2742746", "source": "2742131", "target": "2741927", "SCORE": 0.5068941191567684, "shared_interaction": "Direct", "source_original": "106", "Name_B": "sce01110", "Name_A": "GO:0005975", "target_original": "38", "Gene_B": "sce01110", "interaction": "Direct", "SUID": 2742746, "TYPE": "Direct", "Gene_A": "GO:0005975", "BEND_MAP_ID": 2742746, "selected": false}, "selected": false}, {"data": {"id": "2742743", "source": "2742131", "target": "2742125", "SCORE": 0.49849332759463505, "shared_interaction": "Direct", "source_original": "106", "Name_B": "sce00500", "Name_A": "GO:0005975", "target_original": "104", "Gene_B": "sce00500", "interaction": "Direct", "SUID": 2742743, "TYPE": "Direct", "Gene_A": "GO:0005975", "BEND_MAP_ID": 2742743, "selected": false}, "selected": false}, {"data": {"id": "2742740", "source": "2742131", "target": "2741873", "SCORE": 0.8136948462604013, "shared_interaction": "Direct", "source_original": "106", "Name_B": "GO:0044262", "Name_A": "GO:0005975", "target_original": "20", "Gene_B": "GO:0044262", "interaction": "Direct", "SUID": 2742740, "TYPE": "Direct", "Gene_A": "GO:0005975", "BEND_MAP_ID": 2742740, "selected": false}, "selected": false}, {"data": {"id": "2742944", "source": "2742128", "target": "2742125", "SCORE": 0.4049222196456622, "shared_interaction": "Direct", "source_original": "105", "Name_B": "sce00500", "Name_A": "WP70", "target_original": "104", "Gene_B": "sce00500", "interaction": "Direct", "SUID": 2742944, "TYPE": "Direct", "Gene_A": "WP70", "BEND_MAP_ID": 2742944, "selected": false}, "selected": false}, {"data": {"id": "2742899", "source": "2742125", "target": "2741927", "SCORE": 0.487392113725837, "shared_interaction": "Direct", "source_original": "104", "Name_B": "sce01110", "Name_A": "sce00500", "target_original": "38", "Gene_B": "sce01110", "interaction": "Direct", "SUID": 2742899, "TYPE": "Direct", "Gene_A": "sce00500", "BEND_MAP_ID": 2742899, "selected": false}, "selected": false}, {"data": {"id": "2743040", "source": "2742122", "target": "2742017", "SCORE": 0.31959766385721206, "shared_interaction": "Direct", "source_original": "103", "Name_B": "sce00010", "Name_A": "WP112", "target_original": "68", "Gene_B": "sce00010", "interaction": "Direct", "SUID": 2743040, "TYPE": "Direct", "Gene_A": "WP112", "BEND_MAP_ID": 2743040, "selected": false}, "selected": false}, {"data": {"id": "2743034", "source": "2742122", "target": "2741948", "SCORE": 0.5961871750449582, "shared_interaction": "Direct", "source_original": "103", "Name_B": "sce00020", "Name_A": "WP112", "target_original": "45", "Gene_B": "sce00020", "interaction": "Direct", "SUID": 2743034, "TYPE": "Direct", "Gene_A": "WP112", "BEND_MAP_ID": 2743034, "selected": false}, "selected": false}, {"data": {"id": "2743028", "source": "2742122", "target": "2741942", "SCORE": 0.41513805754269667, "shared_interaction": "Direct", "source_original": "103", "Name_B": "WP559", "Name_A": "WP112", "target_original": "43", "Gene_B": "WP559", "interaction": "Direct", "SUID": 2743028, "TYPE": "Direct", "Gene_A": "WP112", "BEND_MAP_ID": 2743028, "selected": false}, "selected": false}, {"data": {"id": "2743025", "source": "2742122", "target": "2741939", "SCORE": 0.6485671191567266, "shared_interaction": "Direct", "source_original": "103", "Name_B": "WP490", "Name_A": "WP112", "target_original": "42", "Gene_B": "WP490", "interaction": "Direct", "SUID": 2743025, "TYPE": "Direct", "Gene_A": "WP112", "BEND_MAP_ID": 2743025, "selected": false}, "selected": false}, {"data": {"id": "2743019", "source": "2742122", "target": "2741933", "SCORE": 0.5961871750449582, "shared_interaction": "Direct", "source_original": "103", "Name_B": "WP296", "Name_A": "WP112", "target_original": "40", "Gene_B": "WP296", "interaction": "Direct", "SUID": 2743019, "TYPE": "Direct", "Gene_A": "WP112", "BEND_MAP_ID": 2743019, "selected": false}, "selected": false}, {"data": {"id": "2743016", "source": "2742122", "target": "2741930", "SCORE": 0.7357943807491691, "shared_interaction": "Direct", "source_original": "103", "Name_B": "sce01200", "Name_A": "WP112", "target_original": "39", "Gene_B": "sce01200", "interaction": "Direct", "SUID": 2743016, "TYPE": "Direct", "Gene_A": "WP112", "BEND_MAP_ID": 2743016, "selected": false}, "selected": false}, {"data": {"id": "2743010", "source": "2742122", "target": "2741927", "SCORE": 0.5253035612428539, "shared_interaction": "Direct", "source_original": "103", "Name_B": "sce01110", "Name_A": "WP112", "target_original": "38", "Gene_B": "sce01110", "interaction": "Direct", "SUID": 2743010, "TYPE": "Direct", "Gene_A": "WP112", "BEND_MAP_ID": 2743010, "selected": false}, "selected": false}, {"data": {"id": "2742695", "source": "2742119", "target": "2741927", "SCORE": 0.49345578774620885, "shared_interaction": "Direct", "source_original": "102", "Name_B": "sce01110", "Name_A": "sce01100", "target_original": "38", "Gene_B": "sce01110", "interaction": "Direct", "SUID": 2742695, "TYPE": "Direct", "Gene_A": "sce01100", "BEND_MAP_ID": 2742695, "selected": false}, "selected": false}, {"data": {"id": "2742248", "source": "2742113", "target": "2741870", "SCORE": 0.33406617566315894, "shared_interaction": "Direct", "source_original": "100", "Name_B": "R-SCE-1430728", "Name_A": "GO:0046034", "target_original": "19", "Gene_B": "R-SCE-1430728", "interaction": "Direct", "SUID": 2742248, "TYPE": "Direct", "Gene_A": "GO:0046034", "BEND_MAP_ID": 2742248, "selected": false}, "selected": false}, {"data": {"id": "2742236", "source": "2742113", "target": "2742119", "SCORE": 0.502446836907714, "shared_interaction": "Direct", "source_original": "100", "Name_B": "sce01100", "Name_A": "GO:0046034", "target_original": "102", "Gene_B": "sce01100", "interaction": "Direct", "SUID": 2742236, "TYPE": "Direct", "Gene_A": "GO:0046034", "BEND_MAP_ID": 2742236, "selected": false}, "selected": false}, {"data": {"id": "2742221", "source": "2742113", "target": "2741837", "SCORE": 0.5113664961002754, "shared_interaction": "Direct", "source_original": "100", "Name_B": "R-SCE-163200", "Name_A": "GO:0046034", "target_original": "8", "Gene_B": "R-SCE-163200", "interaction": "Direct", "SUID": 2742221, "TYPE": "Direct", "Gene_A": "GO:0046034", "BEND_MAP_ID": 2742221, "selected": false}, "selected": false}, {"data": {"id": "2742209", "source": "2742113", "target": "2741831", "SCORE": 0.49784482758710746, "shared_interaction": "Direct", "source_original": "100", "Name_B": "R-SCE-1428517", "Name_A": "GO:0046034", "target_original": "6", "Gene_B": "R-SCE-1428517", "interaction": "Direct", "SUID": 2742209, "TYPE": "Direct", "Gene_A": "GO:0046034", "BEND_MAP_ID": 2742209, "selected": false}, "selected": false}, {"data": {"id": "2742197", "source": "2742113", "target": "2741819", "SCORE": 0.8571039182285063, "shared_interaction": "Direct", "source_original": "100", "Name_B": "sce00190", "Name_A": "GO:0046034", "target_original": "2", "Gene_B": "sce00190", "interaction": "Direct", "SUID": 2742197, "TYPE": "Direct", "Gene_A": "GO:0046034", "BEND_MAP_ID": 2742197, "selected": false}, "selected": false}, {"data": {"id": "2742533", "source": "2742110", "target": "2742164", "SCORE": 0.3075150118284791, "shared_interaction": "Direct", "source_original": "99", "Name_B": "GO:0044281", "Name_A": "GO:0006754", "target_original": "117", "Gene_B": "GO:0044281", "interaction": "Direct", "SUID": 2742533, "TYPE": "Direct", "Gene_A": "GO:0006754", "BEND_MAP_ID": 2742533, "selected": false}, "selected": false}, {"data": {"id": "2742530", "source": "2742110", "target": "2741954", "SCORE": 0.3504579848675916, "shared_interaction": "Direct", "source_original": "99", "Name_B": "GO:0006796", "Name_A": "GO:0006754", "target_original": "47", "Gene_B": "GO:0006796", "interaction": "Direct", "SUID": 2742530, "TYPE": "Direct", "Gene_A": "GO:0006754", "BEND_MAP_ID": 2742530, "selected": false}, "selected": false}, {"data": {"id": "2742527", "source": "2742110", "target": "2742146", "SCORE": 0.32497795692251713, "shared_interaction": "Direct", "source_original": "99", "Name_B": "GO:0006793", "Name_A": "GO:0006754", "target_original": "111", "Gene_B": "GO:0006793", "interaction": "Direct", "SUID": 2742527, "TYPE": "Direct", "Gene_A": "GO:0006754", "BEND_MAP_ID": 2742527, "selected": false}, "selected": false}, {"data": {"id": "2742524", "source": "2742110", "target": "2741867", "SCORE": 0.7493725349594643, "shared_interaction": "Direct", "source_original": "99", "Name_B": "GO:0009150", "Name_A": "GO:0006754", "target_original": "18", "Gene_B": "GO:0009150", "interaction": "Direct", "SUID": 2742524, "TYPE": "Direct", "Gene_A": "GO:0006754", "BEND_MAP_ID": 2742524, "selected": false}, "selected": false}, {"data": {"id": "2742521", "source": "2742110", "target": "2741864", "SCORE": 1.0, "shared_interaction": "Direct", "source_original": "99", "Name_B": "GO:0009142", "Name_A": "GO:0006754", "target_original": "17", "Gene_B": "GO:0009142", "interaction": "Direct", "SUID": 2742521, "TYPE": "Direct", "Gene_A": "GO:0006754", "BEND_MAP_ID": 2742521, "selected": false}, "selected": false}, {"data": {"id": "2742518", "source": "2742110", "target": "2741861", "SCORE": 1.0, "shared_interaction": "Direct", "source_original": "99", "Name_B": "GO:0009199", "Name_A": "GO:0006754", "target_original": "16", "Gene_B": "GO:0009199", "interaction": "Direct", "SUID": 2742518, "TYPE": "Direct", "Gene_A": "GO:0006754", "BEND_MAP_ID": 2742518, "selected": false}, "selected": false}, {"data": {"id": "2742515", "source": "2742110", "target": "2741858", "SCORE": 1.0, "shared_interaction": "Direct", "source_original": "99", "Name_B": "GO:0009201", "Name_A": "GO:0006754", "target_original": "15", "Gene_B": "GO:0009201", "interaction": "Direct", "SUID": 2742515, "TYPE": "Direct", "Gene_A": "GO:0006754", "BEND_MAP_ID": 2742515, "selected": false}, "selected": false}, {"data": {"id": "2742512", "source": "2742110", "target": "2741855", "SCORE": 1.0, "shared_interaction": "Direct", "source_original": "99", "Name_B": "GO:0009144", "Name_A": "GO:0006754", "target_original": "14", "Gene_B": "GO:0009144", "interaction": "Direct", "SUID": 2742512, "TYPE": "Direct", "Gene_A": "GO:0006754", "BEND_MAP_ID": 2742512, "selected": false}, "selected": false}, {"data": {"id": "2742509", "source": "2742110", "target": "2741852", "SCORE": 1.0, "shared_interaction": "Direct", "source_original": "99", "Name_B": "GO:0009205", "Name_A": "GO:0006754", "target_original": "13", "Gene_B": "GO:0009205", "interaction": "Direct", "SUID": 2742509, "TYPE": "Direct", "Gene_A": "GO:0006754", "BEND_MAP_ID": 2742509, "selected": false}, "selected": false}, {"data": {"id": "2742506", "source": "2742110", "target": "2741849", "SCORE": 1.0, "shared_interaction": "Direct", "source_original": "99", "Name_B": "GO:0009206", "Name_A": "GO:0006754", "target_original": "12", "Gene_B": "GO:0009206", "interaction": "Direct", "SUID": 2742506, "TYPE": "Direct", "Gene_A": "GO:0006754", "BEND_MAP_ID": 2742506, "selected": false}, "selected": false}, {"data": {"id": "2742503", "source": "2742110", "target": "2741846", "SCORE": 1.0, "shared_interaction": "Direct", "source_original": "99", "Name_B": "GO:0009145", "Name_A": "GO:0006754", "target_original": "11", "Gene_B": "GO:0009145", "interaction": "Direct", "SUID": 2742503, "TYPE": "Direct", "Gene_A": "GO:0006754", "BEND_MAP_ID": 2742503, "selected": false}, "selected": false}, {"data": {"id": "2742500", "source": "2742110", "target": "2741837", "SCORE": 0.5303771955097815, "shared_interaction": "Direct", "source_original": "99", "Name_B": "R-SCE-163200", "Name_A": "GO:0006754", "target_original": "8", "Gene_B": "R-SCE-163200", "interaction": "Direct", "SUID": 2742500, "TYPE": "Direct", "Gene_A": "GO:0006754", "BEND_MAP_ID": 2742500, "selected": false}, "selected": false}, {"data": {"id": "2742497", "source": "2742110", "target": "2741843", "SCORE": 1.0, "shared_interaction": "Direct", "source_original": "99", "Name_B": "GO:0015986", "Name_A": "GO:0006754", "target_original": "10", "Gene_B": "GO:0015986", "interaction": "Direct", "SUID": 2742497, "TYPE": "Direct", "Gene_A": "GO:0006754", "BEND_MAP_ID": 2742497, "selected": false}, "selected": false}, {"data": {"id": "2742428", "source": "2742110", "target": "2741831", "SCORE": 0.41520258157208373, "shared_interaction": "Direct", "source_original": "99", "Name_B": "R-SCE-1428517", "Name_A": "GO:0006754", "target_original": "6", "Gene_B": "R-SCE-1428517", "interaction": "Direct", "SUID": 2742428, "TYPE": "Direct", "Gene_A": "GO:0006754", "BEND_MAP_ID": 2742428, "selected": false}, "selected": false}, {"data": {"id": "2742308", "source": "2742110", "target": "2741819", "SCORE": 0.5603773584915323, "shared_interaction": "Direct", "source_original": "99", "Name_B": "sce00190", "Name_A": "GO:0006754", "target_original": "2", "Gene_B": "sce00190", "interaction": "Direct", "SUID": 2742308, "TYPE": "Direct", "Gene_A": "GO:0006754", "BEND_MAP_ID": 2742308, "selected": false}, "selected": false}, {"data": {"id": "2742215", "source": "2742110", "target": "2742113", "SCORE": 0.49849332759463505, "shared_interaction": "Direct", "source_original": "99", "Name_B": "GO:0046034", "Name_A": "GO:0006754", "target_original": "100", "Gene_B": "GO:0046034", "interaction": "Direct", "SUID": 2742215, "TYPE": "Direct", "Gene_A": "GO:0006754", "BEND_MAP_ID": 2742215, "selected": false}, "selected": false}, {"data": {"id": "2744258", "source": "2742104", "target": "2742107", "SCORE": 1.0, "shared_interaction": "Direct", "source_original": "97", "Name_B": "GO:0062014", "Name_A": "GO:0006111", "target_original": "98", "Gene_B": "GO:0062014", "interaction": "Direct", "SUID": 2744258, "TYPE": "Direct", "Gene_A": "GO:0006111", "BEND_MAP_ID": 2744258, "selected": false}, "selected": false}, {"data": {"id": "2744249", "source": "2742104", "target": "2742101", "SCORE": 0.7957931638934084, "shared_interaction": "Direct", "source_original": "97", "Name_B": "GO:0043255", "Name_A": "GO:0006111", "target_original": "96", "Gene_B": "GO:0043255", "interaction": "Direct", "SUID": 2744249, "TYPE": "Direct", "Gene_A": "GO:0006111", "BEND_MAP_ID": 2744249, "selected": false}, "selected": false}, {"data": {"id": "2744168", "source": "2742104", "target": "2742098", "SCORE": 1.0, "shared_interaction": "Direct", "source_original": "97", "Name_B": "GO:0045721", "Name_A": "GO:0006111", "target_original": "95", "Gene_B": "GO:0045721", "interaction": "Direct", "SUID": 2744168, "TYPE": "Direct", "Gene_A": "GO:0006111", "BEND_MAP_ID": 2744168, "selected": false}, "selected": false}, {"data": {"id": "2744114", "source": "2742104", "target": "2742095", "SCORE": 0.8867282450182981, "shared_interaction": "Direct", "source_original": "97", "Name_B": "GO:0045912", "Name_A": "GO:0006111", "target_original": "94", "Gene_B": "GO:0045912", "interaction": "Direct", "SUID": 2744114, "TYPE": "Direct", "Gene_A": "GO:0006111", "BEND_MAP_ID": 2744114, "selected": false}, "selected": false}, {"data": {"id": "2744099", "source": "2742104", "target": "2742092", "SCORE": 0.8867282450182981, "shared_interaction": "Direct", "source_original": "97", "Name_B": "GO:0010677", "Name_A": "GO:0006111", "target_original": "93", "Gene_B": "GO:0010677", "interaction": "Direct", "SUID": 2744099, "TYPE": "Direct", "Gene_A": "GO:0006111", "BEND_MAP_ID": 2744099, "selected": false}, "selected": false}, {"data": {"id": "2744012", "source": "2742104", "target": "2742089", "SCORE": 0.6588579795051059, "shared_interaction": "Direct", "source_original": "97", "Name_B": "GO:0010675", "Name_A": "GO:0006111", "target_original": "92", "Gene_B": "GO:0010675", "interaction": "Direct", "SUID": 2744012, "TYPE": "Direct", "Gene_A": "GO:0006111", "BEND_MAP_ID": 2744012, "selected": false}, "selected": false}, {"data": {"id": "2743931", "source": "2742104", "target": "2742086", "SCORE": 0.6588579795051059, "shared_interaction": "Direct", "source_original": "97", "Name_B": "GO:0010906", "Name_A": "GO:0006111", "target_original": "91", "Gene_B": "GO:0010906", "interaction": "Direct", "SUID": 2743931, "TYPE": "Direct", "Gene_A": "GO:0006111", "BEND_MAP_ID": 2743931, "selected": false}, "selected": false}, {"data": {"id": "2744252", "source": "2742101", "target": "2742107", "SCORE": 0.7957931638934084, "shared_interaction": "Direct", "source_original": "96", "Name_B": "GO:0062014", "Name_A": "GO:0043255", "target_original": "98", "Gene_B": "GO:0062014", "interaction": "Direct", "SUID": 2744252, "TYPE": "Direct", "Gene_A": "GO:0043255", "BEND_MAP_ID": 2744252, "selected": false}, "selected": false}, {"data": {"id": "2744165", "source": "2742101", "target": "2742098", "SCORE": 0.7957931638934084, "shared_interaction": "Direct", "source_original": "96", "Name_B": "GO:0045721", "Name_A": "GO:0043255", "target_original": "95", "Gene_B": "GO:0045721", "interaction": "Direct", "SUID": 2744165, "TYPE": "Direct", "Gene_A": "GO:0043255", "BEND_MAP_ID": 2744165, "selected": false}, "selected": false}, {"data": {"id": "2744111", "source": "2742101", "target": "2742095", "SCORE": 0.7207351178611695, "shared_interaction": "Direct", "source_original": "96", "Name_B": "GO:0045912", "Name_A": "GO:0043255", "target_original": "94", "Gene_B": "GO:0045912", "interaction": "Direct", "SUID": 2744111, "TYPE": "Direct", "Gene_A": "GO:0043255", "BEND_MAP_ID": 2744111, "selected": false}, "selected": false}, {"data": {"id": "2744171", "source": "2742098", "target": "2742107", "SCORE": 1.0, "shared_interaction": "Direct", "source_original": "95", "Name_B": "GO:0062014", "Name_A": "GO:0045721", "target_original": "98", "Gene_B": "GO:0062014", "interaction": "Direct", "SUID": 2744171, "TYPE": "Direct", "Gene_A": "GO:0045721", "BEND_MAP_ID": 2744171, "selected": false}, "selected": false}, {"data": {"id": "2744108", "source": "2742098", "target": "2742095", "SCORE": 0.8867282450182981, "shared_interaction": "Direct", "source_original": "95", "Name_B": "GO:0045912", "Name_A": "GO:0045721", "target_original": "94", "Gene_B": "GO:0045912", "interaction": "Direct", "SUID": 2744108, "TYPE": "Direct", "Gene_A": "GO:0045721", "BEND_MAP_ID": 2744108, "selected": false}, "selected": false}, {"data": {"id": "2744117", "source": "2742095", "target": "2742107", "SCORE": 0.8867282450182981, "shared_interaction": "Direct", "source_original": "94", "Name_B": "GO:0062014", "Name_A": "GO:0045912", "target_original": "98", "Gene_B": "GO:0062014", "interaction": "Direct", "SUID": 2744117, "TYPE": "Direct", "Gene_A": "GO:0045912", "BEND_MAP_ID": 2744117, "selected": false}, "selected": false}, {"data": {"id": "2744102", "source": "2742092", "target": "2742107", "SCORE": 0.8867282450182981, "shared_interaction": "Direct", "source_original": "93", "Name_B": "GO:0062014", "Name_A": "GO:0010677", "target_original": "98", "Gene_B": "GO:0062014", "interaction": "Direct", "SUID": 2744102, "TYPE": "Direct", "Gene_A": "GO:0010677", "BEND_MAP_ID": 2744102, "selected": false}, "selected": false}, {"data": {"id": "2744096", "source": "2742092", "target": "2742101", "SCORE": 0.7207351178611695, "shared_interaction": "Direct", "source_original": "93", "Name_B": "GO:0043255", "Name_A": "GO:0010677", "target_original": "96", "Gene_B": "GO:0043255", "interaction": "Direct", "SUID": 2744096, "TYPE": "Direct", "Gene_A": "GO:0010677", "BEND_MAP_ID": 2744096, "selected": false}, "selected": false}, {"data": {"id": "2744093", "source": "2742092", "target": "2742098", "SCORE": 0.8867282450182981, "shared_interaction": "Direct", "source_original": "93", "Name_B": "GO:0045721", "Name_A": "GO:0010677", "target_original": "95", "Gene_B": "GO:0045721", "interaction": "Direct", "SUID": 2744093, "TYPE": "Direct", "Gene_A": "GO:0010677", "BEND_MAP_ID": 2744093, "selected": false}, "selected": false}, {"data": {"id": "2744090", "source": "2742092", "target": "2742095", "SCORE": 1.0, "shared_interaction": "Direct", "source_original": "93", "Name_B": "GO:0045912", "Name_A": "GO:0010677", "target_original": "94", "Gene_B": "GO:0045912", "interaction": "Direct", "SUID": 2744090, "TYPE": "Direct", "Gene_A": "GO:0010677", "BEND_MAP_ID": 2744090, "selected": false}, "selected": false}, {"data": {"id": "2743919", "source": "2742092", "target": "2742086", "SCORE": 0.7629704984759316, "shared_interaction": "Direct", "source_original": "93", "Name_B": "GO:0010906", "Name_A": "GO:0010677", "target_original": "91", "Gene_B": "GO:0010906", "interaction": "Direct", "SUID": 2743919, "TYPE": "Direct", "Gene_A": "GO:0010677", "BEND_MAP_ID": 2743919, "selected": false}, "selected": false}, {"data": {"id": "2744015", "source": "2742089", "target": "2742107", "SCORE": 0.6588579795051059, "shared_interaction": "Direct", "source_original": "92", "Name_B": "GO:0062014", "Name_A": "GO:0010675", "target_original": "98", "Gene_B": "GO:0062014", "interaction": "Direct", "SUID": 2744015, "TYPE": "Direct", "Gene_A": "GO:0010675", "BEND_MAP_ID": 2744015, "selected": false}, "selected": false}, {"data": {"id": "2744009", "source": "2742089", "target": "2742101", "SCORE": 0.8528111181312159, "shared_interaction": "Direct", "source_original": "92", "Name_B": "GO:0043255", "Name_A": "GO:0010675", "target_original": "96", "Gene_B": "GO:0043255", "interaction": "Direct", "SUID": 2744009, "TYPE": "Direct", "Gene_A": "GO:0010675", "BEND_MAP_ID": 2744009, "selected": false}, "selected": false}, {"data": {"id": "2744006", "source": "2742089", "target": "2742098", "SCORE": 0.6588579795051059, "shared_interaction": "Direct", "source_original": "92", "Name_B": "GO:0045721", "Name_A": "GO:0010675", "target_original": "95", "Gene_B": "GO:0045721", "interaction": "Direct", "SUID": 2744006, "TYPE": "Direct", "Gene_A": "GO:0010675", "BEND_MAP_ID": 2744006, "selected": false}, "selected": false}, {"data": {"id": "2744003", "source": "2742089", "target": "2742095", "SCORE": 0.7629704984759316, "shared_interaction": "Direct", "source_original": "92", "Name_B": "GO:0045912", "Name_A": "GO:0010675", "target_original": "94", "Gene_B": "GO:0045912", "interaction": "Direct", "SUID": 2744003, "TYPE": "Direct", "Gene_A": "GO:0010675", "BEND_MAP_ID": 2744003, "selected": false}, "selected": false}, {"data": {"id": "2744000", "source": "2742089", "target": "2742092", "SCORE": 0.7629704984759316, "shared_interaction": "Direct", "source_original": "92", "Name_B": "GO:0010677", "Name_A": "GO:0010675", "target_original": "93", "Gene_B": "GO:0010677", "interaction": "Direct", "SUID": 2744000, "TYPE": "Direct", "Gene_A": "GO:0010675", "BEND_MAP_ID": 2744000, "selected": false}, "selected": false}, {"data": {"id": "2743916", "source": "2742089", "target": "2742086", "SCORE": 1.0, "shared_interaction": "Direct", "source_original": "92", "Name_B": "GO:0010906", "Name_A": "GO:0010675", "target_original": "91", "Gene_B": "GO:0010906", "interaction": "Direct", "SUID": 2743916, "TYPE": "Direct", "Gene_A": "GO:0010675", "BEND_MAP_ID": 2743916, "selected": false}, "selected": false}, {"data": {"id": "2743886", "source": "2742089", "target": "2741903", "SCORE": 0.30689142290930854, "shared_interaction": "Direct", "source_original": "92", "Name_B": "GO:0044042", "Name_A": "GO:0010675", "target_original": "30", "Gene_B": "GO:0044042", "interaction": "Direct", "SUID": 2743886, "TYPE": "Direct", "Gene_A": "GO:0010675", "BEND_MAP_ID": 2743886, "selected": false}, "selected": false}, {"data": {"id": "2743934", "source": "2742086", "target": "2742107", "SCORE": 0.6588579795051059, "shared_interaction": "Direct", "source_original": "91", "Name_B": "GO:0062014", "Name_A": "GO:0010906", "target_original": "98", "Gene_B": "GO:0062014", "interaction": "Direct", "SUID": 2743934, "TYPE": "Direct", "Gene_A": "GO:0010906", "BEND_MAP_ID": 2743934, "selected": false}, "selected": false}, {"data": {"id": "2743928", "source": "2742086", "target": "2742101", "SCORE": 0.8528111181312159, "shared_interaction": "Direct", "source_original": "91", "Name_B": "GO:0043255", "Name_A": "GO:0010906", "target_original": "96", "Gene_B": "GO:0043255", "interaction": "Direct", "SUID": 2743928, "TYPE": "Direct", "Gene_A": "GO:0010906", "BEND_MAP_ID": 2743928, "selected": false}, "selected": false}, {"data": {"id": "2743925", "source": "2742086", "target": "2742098", "SCORE": 0.6588579795051059, "shared_interaction": "Direct", "source_original": "91", "Name_B": "GO:0045721", "Name_A": "GO:0010906", "target_original": "95", "Gene_B": "GO:0045721", "interaction": "Direct", "SUID": 2743925, "TYPE": "Direct", "Gene_A": "GO:0010906", "BEND_MAP_ID": 2743925, "selected": false}, "selected": false}, {"data": {"id": "2743922", "source": "2742086", "target": "2742095", "SCORE": 0.7629704984759316, "shared_interaction": "Direct", "source_original": "91", "Name_B": "GO:0045912", "Name_A": "GO:0010906", "target_original": "94", "Gene_B": "GO:0045912", "interaction": "Direct", "SUID": 2743922, "TYPE": "Direct", "Gene_A": "GO:0010906", "BEND_MAP_ID": 2743922, "selected": false}, "selected": false}, {"data": {"id": "2743883", "source": "2742086", "target": "2741903", "SCORE": 0.30689142290930854, "shared_interaction": "Direct", "source_original": "91", "Name_B": "GO:0044042", "Name_A": "GO:0010906", "target_original": "30", "Gene_B": "GO:0044042", "interaction": "Direct", "SUID": 2743883, "TYPE": "Direct", "Gene_A": "GO:0010906", "BEND_MAP_ID": 2743883, "selected": false}, "selected": false}, {"data": {"id": "2743871", "source": "2742083", "target": "2742071", "SCORE": 1.0, "shared_interaction": "Direct", "source_original": "90", "Name_B": "GO:0071465", "Name_A": "GO:0071462", "target_original": "86", "Gene_B": "GO:0071465", "interaction": "Direct", "SUID": 2743871, "TYPE": "Direct", "Gene_A": "GO:0071462", "BEND_MAP_ID": 2743871, "selected": false}, "selected": false}, {"data": {"id": "2743487", "source": "2742083", "target": "2741909", "SCORE": 0.43474041727955826, "shared_interaction": "Direct", "source_original": "90", "Name_B": "WP398", "Name_A": "GO:0071462", "target_original": "32", "Gene_B": "WP398", "interaction": "Direct", "SUID": 2743487, "TYPE": "Direct", "Gene_A": "GO:0071462", "BEND_MAP_ID": 2743487, "selected": false}, "selected": false}, {"data": {"id": "2744255", "source": "2742080", "target": "2742083", "SCORE": 1.0, "shared_interaction": "Direct", "source_original": "89", "Name_B": "GO:0071462", "Name_A": "GO:0009415", "target_original": "90", "Gene_B": "GO:0071462", "interaction": "Direct", "SUID": 2744255, "TYPE": "Direct", "Gene_A": "GO:0009415", "BEND_MAP_ID": 2744255, "selected": false}, "selected": false}, {"data": {"id": "2744198", "source": "2742080", "target": "2742077", "SCORE": 1.0, "shared_interaction": "Direct", "source_original": "89", "Name_B": "GO:0042631", "Name_A": "GO:0009415", "target_original": "88", "Gene_B": "GO:0042631", "interaction": "Direct", "SUID": 2744198, "TYPE": "Direct", "Gene_A": "GO:0009415", "BEND_MAP_ID": 2744198, "selected": false}, "selected": false}, {"data": {"id": "2743868", "source": "2742080", "target": "2742071", "SCORE": 1.0, "shared_interaction": "Direct", "source_original": "89", "Name_B": "GO:0071465", "Name_A": "GO:0009415", "target_original": "86", "Gene_B": "GO:0071465", "interaction": "Direct", "SUID": 2743868, "TYPE": "Direct", "Gene_A": "GO:0009415", "BEND_MAP_ID": 2743868, "selected": false}, "selected": false}, {"data": {"id": "2743484", "source": "2742080", "target": "2741909", "SCORE": 0.43474041727955826, "shared_interaction": "Direct", "source_original": "89", "Name_B": "WP398", "Name_A": "GO:0009415", "target_original": "32", "Gene_B": "WP398", "interaction": "Direct", "SUID": 2743484, "TYPE": "Direct", "Gene_A": "GO:0009415", "BEND_MAP_ID": 2743484, "selected": false}, "selected": false}, {"data": {"id": "2744201", "source": "2742077", "target": "2742083", "SCORE": 1.0, "shared_interaction": "Direct", "source_original": "88", "Name_B": "GO:0071462", "Name_A": "GO:0042631", "target_original": "90", "Gene_B": "GO:0071462", "interaction": "Direct", "SUID": 2744201, "TYPE": "Direct", "Gene_A": "GO:0042631", "BEND_MAP_ID": 2744201, "selected": false}, "selected": false}, {"data": {"id": "2743865", "source": "2742077", "target": "2742071", "SCORE": 1.0, "shared_interaction": "Direct", "source_original": "88", "Name_B": "GO:0071465", "Name_A": "GO:0042631", "target_original": "86", "Gene_B": "GO:0071465", "interaction": "Direct", "SUID": 2743865, "TYPE": "Direct", "Gene_A": "GO:0042631", "BEND_MAP_ID": 2743865, "selected": false}, "selected": false}, {"data": {"id": "2743481", "source": "2742077", "target": "2741909", "SCORE": 0.43474041727955826, "shared_interaction": "Direct", "source_original": "88", "Name_B": "WP398", "Name_A": "GO:0042631", "target_original": "32", "Gene_B": "WP398", "interaction": "Direct", "SUID": 2743481, "TYPE": "Direct", "Gene_A": "GO:0042631", "BEND_MAP_ID": 2743481, "selected": false}, "selected": false}, {"data": {"id": "2744195", "source": "2742074", "target": "2742083", "SCORE": 1.0, "shared_interaction": "Direct", "source_original": "87", "Name_B": "GO:0071462", "Name_A": "GO:0009414", "target_original": "90", "Gene_B": "GO:0071462", "interaction": "Direct", "SUID": 2744195, "TYPE": "Direct", "Gene_A": "GO:0009414", "BEND_MAP_ID": 2744195, "selected": false}, "selected": false}, {"data": {"id": "2744192", "source": "2742074", "target": "2742080", "SCORE": 1.0, "shared_interaction": "Direct", "source_original": "87", "Name_B": "GO:0009415", "Name_A": "GO:0009414", "target_original": "89", "Gene_B": "GO:0009415", "interaction": "Direct", "SUID": 2744192, "TYPE": "Direct", "Gene_A": "GO:0009414", "BEND_MAP_ID": 2744192, "selected": false}, "selected": false}, {"data": {"id": "2744189", "source": "2742074", "target": "2742077", "SCORE": 1.0, "shared_interaction": "Direct", "source_original": "87", "Name_B": "GO:0042631", "Name_A": "GO:0009414", "target_original": "88", "Gene_B": "GO:0042631", "interaction": "Direct", "SUID": 2744189, "TYPE": "Direct", "Gene_A": "GO:0009414", "BEND_MAP_ID": 2744189, "selected": false}, "selected": false}, {"data": {"id": "2743862", "source": "2742074", "target": "2742071", "SCORE": 1.0, "shared_interaction": "Direct", "source_original": "87", "Name_B": "GO:0071465", "Name_A": "GO:0009414", "target_original": "86", "Gene_B": "GO:0071465", "interaction": "Direct", "SUID": 2743862, "TYPE": "Direct", "Gene_A": "GO:0009414", "BEND_MAP_ID": 2743862, "selected": false}, "selected": false}, {"data": {"id": "2743478", "source": "2742074", "target": "2741909", "SCORE": 0.43474041727955826, "shared_interaction": "Direct", "source_original": "87", "Name_B": "WP398", "Name_A": "GO:0009414", "target_original": "32", "Gene_B": "WP398", "interaction": "Direct", "SUID": 2743478, "TYPE": "Direct", "Gene_A": "GO:0009414", "BEND_MAP_ID": 2743478, "selected": false}, "selected": false}, {"data": {"id": "2743475", "source": "2742071", "target": "2741909", "SCORE": 0.43474041727955826, "shared_interaction": "Direct", "source_original": "86", "Name_B": "WP398", "Name_A": "GO:0071465", "target_original": "32", "Gene_B": "WP398", "interaction": "Direct", "SUID": 2743475, "TYPE": "Direct", "Gene_A": "GO:0071465", "BEND_MAP_ID": 2743475, "selected": false}, "selected": false}, {"data": {"id": "2744228", "source": "2742068", "target": "2742056", "SCORE": 0.5650280024954177, "shared_interaction": "Direct", "source_original": "85", "Name_B": "R-SCE-445717", "Name_A": "R-SCE-418594", "target_original": "81", "Gene_B": "R-SCE-445717", "interaction": "Direct", "SUID": 2744228, "TYPE": "Direct", "Gene_A": "R-SCE-418594", "BEND_MAP_ID": 2744228, "selected": false}, "selected": false}, {"data": {"id": "2744216", "source": "2742068", "target": "2742053", "SCORE": 0.5650280024954177, "shared_interaction": "Direct", "source_original": "85", "Name_B": "R-SCE-432040", "Name_A": "R-SCE-418594", "target_original": "80", "Gene_B": "R-SCE-432040", "interaction": "Direct", "SUID": 2744216, "TYPE": "Direct", "Gene_A": "R-SCE-418594", "BEND_MAP_ID": 2744216, "selected": false}, "selected": false}, {"data": {"id": "2744246", "source": "2742065", "target": "2742068", "SCORE": 1.0, "shared_interaction": "Direct", "source_original": "84", "Name_B": "R-SCE-418594", "Name_A": "R-SCE-112043", "target_original": "85", "Gene_B": "R-SCE-418594", "interaction": "Direct", "SUID": 2744246, "TYPE": "Direct", "Gene_A": "R-SCE-112043", "BEND_MAP_ID": 2744246, "selected": false}, "selected": false}, {"data": {"id": "2744225", "source": "2742065", "target": "2742056", "SCORE": 0.5650280024954177, "shared_interaction": "Direct", "source_original": "84", "Name_B": "R-SCE-445717", "Name_A": "R-SCE-112043", "target_original": "81", "Gene_B": "R-SCE-445717", "interaction": "Direct", "SUID": 2744225, "TYPE": "Direct", "Gene_A": "R-SCE-112043", "BEND_MAP_ID": 2744225, "selected": false}, "selected": false}, {"data": {"id": "2744213", "source": "2742065", "target": "2742053", "SCORE": 0.5650280024954177, "shared_interaction": "Direct", "source_original": "84", "Name_B": "R-SCE-432040", "Name_A": "R-SCE-112043", "target_original": "80", "Gene_B": "R-SCE-432040", "interaction": "Direct", "SUID": 2744213, "TYPE": "Direct", "Gene_A": "R-SCE-112043", "BEND_MAP_ID": 2744213, "selected": false}, "selected": false}, {"data": {"id": "2744243", "source": "2742062", "target": "2742068", "SCORE": 1.0, "shared_interaction": "Direct", "source_original": "83", "Name_B": "R-SCE-418594", "Name_A": "R-SCE-112040", "target_original": "85", "Gene_B": "R-SCE-418594", "interaction": "Direct", "SUID": 2744243, "TYPE": "Direct", "Gene_A": "R-SCE-112040", "BEND_MAP_ID": 2744243, "selected": false}, "selected": false}, {"data": {"id": "2744240", "source": "2742062", "target": "2742065", "SCORE": 1.0, "shared_interaction": "Direct", "source_original": "83", "Name_B": "R-SCE-112043", "Name_A": "R-SCE-112040", "target_original": "84", "Gene_B": "R-SCE-112043", "interaction": "Direct", "SUID": 2744240, "TYPE": "Direct", "Gene_A": "R-SCE-112040", "BEND_MAP_ID": 2744240, "selected": false}, "selected": false}, {"data": {"id": "2744222", "source": "2742062", "target": "2742056", "SCORE": 0.5650280024954177, "shared_interaction": "Direct", "source_original": "83", "Name_B": "R-SCE-445717", "Name_A": "R-SCE-112040", "target_original": "81", "Gene_B": "R-SCE-445717", "interaction": "Direct", "SUID": 2744222, "TYPE": "Direct", "Gene_A": "R-SCE-112040", "BEND_MAP_ID": 2744222, "selected": false}, "selected": false}, {"data": {"id": "2744210", "source": "2742062", "target": "2742053", "SCORE": 0.5650280024954177, "shared_interaction": "Direct", "source_original": "83", "Name_B": "R-SCE-432040", "Name_A": "R-SCE-112040", "target_original": "80", "Gene_B": "R-SCE-432040", "interaction": "Direct", "SUID": 2744210, "TYPE": "Direct", "Gene_A": "R-SCE-112040", "BEND_MAP_ID": 2744210, "selected": false}, "selected": false}, {"data": {"id": "2744237", "source": "2742059", "target": "2742068", "SCORE": 1.0, "shared_interaction": "Direct", "source_original": "82", "Name_B": "R-SCE-418594", "Name_A": "R-SCE-111885", "target_original": "85", "Gene_B": "R-SCE-418594", "interaction": "Direct", "SUID": 2744237, "TYPE": "Direct", "Gene_A": "R-SCE-111885", "BEND_MAP_ID": 2744237, "selected": false}, "selected": false}, {"data": {"id": "2744234", "source": "2742059", "target": "2742065", "SCORE": 1.0, "shared_interaction": "Direct", "source_original": "82", "Name_B": "R-SCE-112043", "Name_A": "R-SCE-111885", "target_original": "84", "Gene_B": "R-SCE-112043", "interaction": "Direct", "SUID": 2744234, "TYPE": "Direct", "Gene_A": "R-SCE-111885", "BEND_MAP_ID": 2744234, "selected": false}, "selected": false}, {"data": {"id": "2744231", "source": "2742059", "target": "2742062", "SCORE": 1.0, "shared_interaction": "Direct", "source_original": "82", "Name_B": "R-SCE-112040", "Name_A": "R-SCE-111885", "target_original": "83", "Gene_B": "R-SCE-112040", "interaction": "Direct", "SUID": 2744231, "TYPE": "Direct", "Gene_A": "R-SCE-111885", "BEND_MAP_ID": 2744231, "selected": false}, "selected": false}, {"data": {"id": "2744219", "source": "2742059", "target": "2742056", "SCORE": 0.5650280024954177, "shared_interaction": "Direct", "source_original": "82", "Name_B": "R-SCE-445717", "Name_A": "R-SCE-111885", "target_original": "81", "Gene_B": "R-SCE-445717", "interaction": "Direct", "SUID": 2744219, "TYPE": "Direct", "Gene_A": "R-SCE-111885", "BEND_MAP_ID": 2744219, "selected": false}, "selected": false}, {"data": {"id": "2744207", "source": "2742059", "target": "2742053", "SCORE": 0.5650280024954177, "shared_interaction": "Direct", "source_original": "82", "Name_B": "R-SCE-432040", "Name_A": "R-SCE-111885", "target_original": "80", "Gene_B": "R-SCE-432040", "interaction": "Direct", "SUID": 2744207, "TYPE": "Direct", "Gene_A": "R-SCE-111885", "BEND_MAP_ID": 2744207, "selected": false}, "selected": false}, {"data": {"id": "2743775", "source": "2742056", "target": "2742116", "SCORE": 0.3202042304944703, "shared_interaction": "Direct", "source_original": "81", "Name_B": "sce04213", "Name_A": "R-SCE-445717", "target_original": "101", "Gene_B": "sce04213", "interaction": "Direct", "SUID": 2743775, "TYPE": "Direct", "Gene_A": "R-SCE-445717", "BEND_MAP_ID": 2743775, "selected": false}, "selected": false}, {"data": {"id": "2744204", "source": "2742053", "target": "2742056", "SCORE": 1.0, "shared_interaction": "Direct", "source_original": "80", "Name_B": "R-SCE-445717", "Name_A": "R-SCE-432040", "target_original": "81", "Gene_B": "R-SCE-445717", "interaction": "Direct", "SUID": 2744204, "TYPE": "Direct", "Gene_A": "R-SCE-432040", "BEND_MAP_ID": 2744204, "selected": false}, "selected": false}, {"data": {"id": "2743772", "source": "2742053", "target": "2742116", "SCORE": 0.3202042304944703, "shared_interaction": "Direct", "source_original": "80", "Name_B": "sce04213", "Name_A": "R-SCE-432040", "target_original": "101", "Gene_B": "sce04213", "interaction": "Direct", "SUID": 2743772, "TYPE": "Direct", "Gene_A": "R-SCE-432040", "BEND_MAP_ID": 2743772, "selected": false}, "selected": false}, {"data": {"id": "2744018", "source": "2742050", "target": "2742047", "SCORE": 0.5961871750449582, "shared_interaction": "Direct", "source_original": "79", "Name_B": "GO:0034605", "Name_A": "GO:0009628", "target_original": "78", "Gene_B": "GO:0034605", "interaction": "Direct", "SUID": 2744018, "TYPE": "Direct", "Gene_A": "GO:0009628", "BEND_MAP_ID": 2744018, "selected": false}, "selected": false}, {"data": {"id": "2743172", "source": "2742047", "target": "2741906", "SCORE": 0.446931163110507, "shared_interaction": "Direct", "source_original": "78", "Name_B": "GO:0070413", "Name_A": "GO:0034605", "target_original": "31", "Gene_B": "GO:0070413", "interaction": "Direct", "SUID": 2743172, "TYPE": "Direct", "Gene_A": "GO:0034605", "BEND_MAP_ID": 2743172, "selected": false}, "selected": false}, {"data": {"id": "2743817", "source": "2742044", "target": "2742050", "SCORE": 0.743593774564687, "shared_interaction": "Direct", "source_original": "77", "Name_B": "GO:0009628", "Name_A": "GO:0009266", "target_original": "79", "Gene_B": "GO:0009628", "interaction": "Direct", "SUID": 2743817, "TYPE": "Direct", "Gene_A": "GO:0009266", "BEND_MAP_ID": 2743817, "selected": false}, "selected": false}, {"data": {"id": "2743814", "source": "2742044", "target": "2742047", "SCORE": 0.8355681016240487, "shared_interaction": "Direct", "source_original": "77", "Name_B": "GO:0034605", "Name_A": "GO:0009266", "target_original": "78", "Gene_B": "GO:0034605", "interaction": "Direct", "SUID": 2743814, "TYPE": "Direct", "Gene_A": "GO:0009266", "BEND_MAP_ID": 2743814, "selected": false}, "selected": false}, {"data": {"id": "2743763", "source": "2742044", "target": "2742161", "SCORE": 1.0, "shared_interaction": "Direct", "source_original": "77", "Name_B": "GO:0009408", "Name_A": "GO:0009266", "target_original": "116", "Gene_B": "GO:0009408", "interaction": "Direct", "SUID": 2743763, "TYPE": "Direct", "Gene_A": "GO:0009266", "BEND_MAP_ID": 2743763, "selected": false}, "selected": false}, {"data": {"id": "2743169", "source": "2742044", "target": "2741906", "SCORE": 0.3559977888377568, "shared_interaction": "Direct", "source_original": "77", "Name_B": "GO:0070413", "Name_A": "GO:0009266", "target_original": "31", "Gene_B": "GO:0070413", "interaction": "Direct", "SUID": 2743169, "TYPE": "Direct", "Gene_A": "GO:0009266", "BEND_MAP_ID": 2743169, "selected": false}, "selected": false}, {"data": {"id": "2744129", "source": "2742041", "target": "2742029", "SCORE": 0.660844250367735, "shared_interaction": "Direct", "source_original": "76", "Name_B": "R-SCE-170822", "Name_A": "GO:0046031", "target_original": "72", "Gene_B": "R-SCE-170822", "interaction": "Direct", "SUID": 2744129, "TYPE": "Direct", "Gene_A": "GO:0046031", "BEND_MAP_ID": 2744129, "selected": false}, "selected": false}, {"data": {"id": "2744087", "source": "2742041", "target": "2742026", "SCORE": 0.8289280469911845, "shared_interaction": "Direct", "source_original": "76", "Name_B": "R-SCE-70171", "Name_A": "GO:0046031", "target_original": "71", "Gene_B": "R-SCE-70171", "interaction": "Direct", "SUID": 2744087, "TYPE": "Direct", "Gene_A": "GO:0046031", "BEND_MAP_ID": 2744087, "selected": false}, "selected": false}, {"data": {"id": "2744066", "source": "2742041", "target": "2742023", "SCORE": 0.7626485568779389, "shared_interaction": "Direct", "source_original": "76", "Name_B": "R-SCE-70326", "Name_A": "GO:0046031", "target_original": "70", "Gene_B": "R-SCE-70326", "interaction": "Direct", "SUID": 2744066, "TYPE": "Direct", "Gene_A": "GO:0046031", "BEND_MAP_ID": 2744066, "selected": false}, "selected": false}, {"data": {"id": "2744042", "source": "2742041", "target": "2742020", "SCORE": 0.5345585297686158, "shared_interaction": "Direct", "source_original": "76", "Name_B": "GO:0046835", "Name_A": "GO:0046031", "target_original": "69", "Gene_B": "GO:0046835", "interaction": "Direct", "SUID": 2744042, "TYPE": "Direct", "Gene_A": "GO:0046031", "BEND_MAP_ID": 2744042, "selected": false}, "selected": false}, {"data": {"id": "2743997", "source": "2742041", "target": "2742014", "SCORE": 0.3157121879647379, "shared_interaction": "Direct", "source_original": "76", "Name_B": "GO:1904659", "Name_A": "GO:0046031", "target_original": "67", "Gene_B": "GO:1904659", "interaction": "Direct", "SUID": 2743997, "TYPE": "Direct", "Gene_A": "GO:0046031", "BEND_MAP_ID": 2743997, "selected": false}, "selected": false}, {"data": {"id": "2743970", "source": "2742041", "target": "2742017", "SCORE": 0.7416851441258369, "shared_interaction": "Direct", "source_original": "76", "Name_B": "sce00010", "Name_A": "GO:0046031", "target_original": "68", "Gene_B": "sce00010", "interaction": "Direct", "SUID": 2743970, "TYPE": "Direct", "Gene_A": "GO:0046031", "BEND_MAP_ID": 2743970, "selected": false}, "selected": false}, {"data": {"id": "2743403", "source": "2742041", "target": "2741930", "SCORE": 0.45881722351484877, "shared_interaction": "Direct", "source_original": "76", "Name_B": "sce01200", "Name_A": "GO:0046031", "target_original": "39", "Gene_B": "sce01200", "interaction": "Direct", "SUID": 2743403, "TYPE": "Direct", "Gene_A": "GO:0046031", "BEND_MAP_ID": 2743403, "selected": false}, "selected": false}, {"data": {"id": "2743229", "source": "2742041", "target": "2741882", "SCORE": 0.43331982164839955, "shared_interaction": "Direct", "source_original": "76", "Name_B": "R-SCE-71387", "Name_A": "GO:0046031", "target_original": "23", "Gene_B": "R-SCE-71387", "interaction": "Direct", "SUID": 2743229, "TYPE": "Direct", "Gene_A": "GO:0046031", "BEND_MAP_ID": 2743229, "selected": false}, "selected": false}, {"data": {"id": "2743058", "source": "2742041", "target": "2742122", "SCORE": 0.3065476190506249, "shared_interaction": "Direct", "source_original": "76", "Name_B": "WP112", "Name_A": "GO:0046031", "target_original": "103", "Gene_B": "WP112", "interaction": "Direct", "SUID": 2743058, "TYPE": "Direct", "Gene_A": "GO:0046031", "BEND_MAP_ID": 2743058, "selected": false}, "selected": false}, {"data": {"id": "2744186", "source": "2742038", "target": "2742041", "SCORE": 1.0, "shared_interaction": "Direct", "source_original": "75", "Name_B": "GO:0046031", "Name_A": "GO:0006165", "target_original": "76", "Gene_B": "GO:0046031", "interaction": "Direct", "SUID": 2744186, "TYPE": "Direct", "Gene_A": "GO:0006165", "BEND_MAP_ID": 2744186, "selected": false}, "selected": false}, {"data": {"id": "2744174", "source": "2742038", "target": "2741960", "SCORE": 0.5028128031060302, "shared_interaction": "Direct", "source_original": "75", "Name_B": "GO:0032787", "Name_A": "GO:0006165", "target_original": "49", "Gene_B": "GO:0032787", "interaction": "Direct", "SUID": 2744174, "TYPE": "Direct", "Gene_A": "GO:0006165", "BEND_MAP_ID": 2744174, "selected": false}, "selected": false}, {"data": {"id": "2744159", "source": "2742038", "target": "2742035", "SCORE": 1.0, "shared_interaction": "Direct", "source_original": "75", "Name_B": "GO:0006757", "Name_A": "GO:0006165", "target_original": "74", "Gene_B": "GO:0006757", "interaction": "Direct", "SUID": 2744159, "TYPE": "Direct", "Gene_A": "GO:0006165", "BEND_MAP_ID": 2744159, "selected": false}, "selected": false}, {"data": {"id": "2744138", "source": "2742038", "target": "2741957", "SCORE": 0.37421665174792307, "shared_interaction": "Direct", "source_original": "75", "Name_B": "GO:0016310", "Name_A": "GO:0006165", "target_original": "48", "Gene_B": "GO:0016310", "interaction": "Direct", "SUID": 2744138, "TYPE": "Direct", "Gene_A": "GO:0006165", "BEND_MAP_ID": 2744138, "selected": false}, "selected": false}, {"data": {"id": "2744126", "source": "2742038", "target": "2742029", "SCORE": 0.660844250367735, "shared_interaction": "Direct", "source_original": "75", "Name_B": "R-SCE-170822", "Name_A": "GO:0006165", "target_original": "72", "Gene_B": "R-SCE-170822", "interaction": "Direct", "SUID": 2744126, "TYPE": "Direct", "Gene_A": "GO:0006165", "BEND_MAP_ID": 2744126, "selected": false}, "selected": false}, {"data": {"id": "2744084", "source": "2742038", "target": "2742026", "SCORE": 0.8289280469911845, "shared_interaction": "Direct", "source_original": "75", "Name_B": "R-SCE-70171", "Name_A": "GO:0006165", "target_original": "71", "Gene_B": "R-SCE-70171", "interaction": "Direct", "SUID": 2744084, "TYPE": "Direct", "Gene_A": "GO:0006165", "BEND_MAP_ID": 2744084, "selected": false}, "selected": false}, {"data": {"id": "2744063", "source": "2742038", "target": "2742023", "SCORE": 0.7626485568779389, "shared_interaction": "Direct", "source_original": "75", "Name_B": "R-SCE-70326", "Name_A": "GO:0006165", "target_original": "70", "Gene_B": "R-SCE-70326", "interaction": "Direct", "SUID": 2744063, "TYPE": "Direct", "Gene_A": "GO:0006165", "BEND_MAP_ID": 2744063, "selected": false}, "selected": false}, {"data": {"id": "2744039", "source": "2742038", "target": "2742020", "SCORE": 0.5345585297686158, "shared_interaction": "Direct", "source_original": "75", "Name_B": "GO:0046835", "Name_A": "GO:0006165", "target_original": "69", "Gene_B": "GO:0046835", "interaction": "Direct", "SUID": 2744039, "TYPE": "Direct", "Gene_A": "GO:0006165", "BEND_MAP_ID": 2744039, "selected": false}, "selected": false}, {"data": {"id": "2743994", "source": "2742038", "target": "2742014", "SCORE": 0.3157121879647379, "shared_interaction": "Direct", "source_original": "75", "Name_B": "GO:1904659", "Name_A": "GO:0006165", "target_original": "67", "Gene_B": "GO:1904659", "interaction": "Direct", "SUID": 2743994, "TYPE": "Direct", "Gene_A": "GO:0006165", "BEND_MAP_ID": 2743994, "selected": false}, "selected": false}, {"data": {"id": "2743967", "source": "2742038", "target": "2742017", "SCORE": 0.7416851441258369, "shared_interaction": "Direct", "source_original": "75", "Name_B": "sce00010", "Name_A": "GO:0006165", "target_original": "68", "Gene_B": "sce00010", "interaction": "Direct", "SUID": 2743967, "TYPE": "Direct", "Gene_A": "GO:0006165", "BEND_MAP_ID": 2743967, "selected": false}, "selected": false}, {"data": {"id": "2743400", "source": "2742038", "target": "2741930", "SCORE": 0.45881722351484877, "shared_interaction": "Direct", "source_original": "75", "Name_B": "sce01200", "Name_A": "GO:0006165", "target_original": "39", "Gene_B": "sce01200", "interaction": "Direct", "SUID": 2743400, "TYPE": "Direct", "Gene_A": "GO:0006165", "BEND_MAP_ID": 2743400, "selected": false}, "selected": false}, {"data": {"id": "2743349", "source": "2742038", "target": "2741876", "SCORE": 0.527766518040333, "shared_interaction": "Direct", "source_original": "75", "Name_B": "GO:0016052", "Name_A": "GO:0006165", "target_original": "21", "Gene_B": "GO:0016052", "interaction": "Direct", "SUID": 2743349, "TYPE": "Direct", "Gene_A": "GO:0006165", "BEND_MAP_ID": 2743349, "selected": false}, "selected": false}, {"data": {"id": "2743226", "source": "2742038", "target": "2741882", "SCORE": 0.43331982164839955, "shared_interaction": "Direct", "source_original": "75", "Name_B": "R-SCE-71387", "Name_A": "GO:0006165", "target_original": "23", "Gene_B": "R-SCE-71387", "interaction": "Direct", "SUID": 2743226, "TYPE": "Direct", "Gene_A": "GO:0006165", "BEND_MAP_ID": 2743226, "selected": false}, "selected": false}, {"data": {"id": "2743055", "source": "2742038", "target": "2742122", "SCORE": 0.3065476190506249, "shared_interaction": "Direct", "source_original": "75", "Name_B": "WP112", "Name_A": "GO:0006165", "target_original": "103", "Gene_B": "WP112", "interaction": "Direct", "SUID": 2743055, "TYPE": "Direct", "Gene_A": "GO:0006165", "BEND_MAP_ID": 2743055, "selected": false}, "selected": false}, {"data": {"id": "2742890", "source": "2742038", "target": "2741867", "SCORE": 0.37421665174792307, "shared_interaction": "Direct", "source_original": "75", "Name_B": "GO:0009150", "Name_A": "GO:0006165", "target_original": "18", "Gene_B": "GO:0009150", "interaction": "Direct", "SUID": 2742890, "TYPE": "Direct", "Gene_A": "GO:0006165", "BEND_MAP_ID": 2742890, "selected": false}, "selected": false}, {"data": {"id": "2744162", "source": "2742035", "target": "2742041", "SCORE": 1.0, "shared_interaction": "Direct", "source_original": "74", "Name_B": "GO:0046031", "Name_A": "GO:0006757", "target_original": "76", "Gene_B": "GO:0046031", "interaction": "Direct", "SUID": 2744162, "TYPE": "Direct", "Gene_A": "GO:0006757", "BEND_MAP_ID": 2744162, "selected": false}, "selected": false}, {"data": {"id": "2744156", "source": "2742035", "target": "2741960", "SCORE": 0.5028128031060302, "shared_interaction": "Direct", "source_original": "74", "Name_B": "GO:0032787", "Name_A": "GO:0006757", "target_original": "49", "Gene_B": "GO:0032787", "interaction": "Direct", "SUID": 2744156, "TYPE": "Direct", "Gene_A": "GO:0006757", "BEND_MAP_ID": 2744156, "selected": false}, "selected": false}, {"data": {"id": "2744135", "source": "2742035", "target": "2741957", "SCORE": 0.37421665174792307, "shared_interaction": "Direct", "source_original": "74", "Name_B": "GO:0016310", "Name_A": "GO:0006757", "target_original": "48", "Gene_B": "GO:0016310", "interaction": "Direct", "SUID": 2744135, "TYPE": "Direct", "Gene_A": "GO:0006757", "BEND_MAP_ID": 2744135, "selected": false}, "selected": false}, {"data": {"id": "2744123", "source": "2742035", "target": "2742029", "SCORE": 0.660844250367735, "shared_interaction": "Direct", "source_original": "74", "Name_B": "R-SCE-170822", "Name_A": "GO:0006757", "target_original": "72", "Gene_B": "R-SCE-170822", "interaction": "Direct", "SUID": 2744123, "TYPE": "Direct", "Gene_A": "GO:0006757", "BEND_MAP_ID": 2744123, "selected": false}, "selected": false}, {"data": {"id": "2744078", "source": "2742035", "target": "2742026", "SCORE": 0.8289280469911845, "shared_interaction": "Direct", "source_original": "74", "Name_B": "R-SCE-70171", "Name_A": "GO:0006757", "target_original": "71", "Gene_B": "R-SCE-70171", "interaction": "Direct", "SUID": 2744078, "TYPE": "Direct", "Gene_A": "GO:0006757", "BEND_MAP_ID": 2744078, "selected": false}, "selected": false}, {"data": {"id": "2744057", "source": "2742035", "target": "2742023", "SCORE": 0.7626485568779389, "shared_interaction": "Direct", "source_original": "74", "Name_B": "R-SCE-70326", "Name_A": "GO:0006757", "target_original": "70", "Gene_B": "R-SCE-70326", "interaction": "Direct", "SUID": 2744057, "TYPE": "Direct", "Gene_A": "GO:0006757", "BEND_MAP_ID": 2744057, "selected": false}, "selected": false}, {"data": {"id": "2744036", "source": "2742035", "target": "2742020", "SCORE": 0.5345585297686158, "shared_interaction": "Direct", "source_original": "74", "Name_B": "GO:0046835", "Name_A": "GO:0006757", "target_original": "69", "Gene_B": "GO:0046835", "interaction": "Direct", "SUID": 2744036, "TYPE": "Direct", "Gene_A": "GO:0006757", "BEND_MAP_ID": 2744036, "selected": false}, "selected": false}, {"data": {"id": "2743991", "source": "2742035", "target": "2742014", "SCORE": 0.3157121879647379, "shared_interaction": "Direct", "source_original": "74", "Name_B": "GO:1904659", "Name_A": "GO:0006757", "target_original": "67", "Gene_B": "GO:1904659", "interaction": "Direct", "SUID": 2743991, "TYPE": "Direct", "Gene_A": "GO:0006757", "BEND_MAP_ID": 2743991, "selected": false}, "selected": false}, {"data": {"id": "2743961", "source": "2742035", "target": "2742017", "SCORE": 0.7416851441258369, "shared_interaction": "Direct", "source_original": "74", "Name_B": "sce00010", "Name_A": "GO:0006757", "target_original": "68", "Gene_B": "sce00010", "interaction": "Direct", "SUID": 2743961, "TYPE": "Direct", "Gene_A": "GO:0006757", "BEND_MAP_ID": 2743961, "selected": false}, "selected": false}, {"data": {"id": "2743394", "source": "2742035", "target": "2741930", "SCORE": 0.45881722351484877, "shared_interaction": "Direct", "source_original": "74", "Name_B": "sce01200", "Name_A": "GO:0006757", "target_original": "39", "Gene_B": "sce01200", "interaction": "Direct", "SUID": 2743394, "TYPE": "Direct", "Gene_A": "GO:0006757", "BEND_MAP_ID": 2743394, "selected": false}, "selected": false}, {"data": {"id": "2743343", "source": "2742035", "target": "2741876", "SCORE": 0.527766518040333, "shared_interaction": "Direct", "source_original": "74", "Name_B": "GO:0016052", "Name_A": "GO:0006757", "target_original": "21", "Gene_B": "GO:0016052", "interaction": "Direct", "SUID": 2743343, "TYPE": "Direct", "Gene_A": "GO:0006757", "BEND_MAP_ID": 2743343, "selected": false}, "selected": false}, {"data": {"id": "2743220", "source": "2742035", "target": "2741882", "SCORE": 0.43331982164839955, "shared_interaction": "Direct", "source_original": "74", "Name_B": "R-SCE-71387", "Name_A": "GO:0006757", "target_original": "23", "Gene_B": "R-SCE-71387", "interaction": "Direct", "SUID": 2743220, "TYPE": "Direct", "Gene_A": "GO:0006757", "BEND_MAP_ID": 2743220, "selected": false}, "selected": false}, {"data": {"id": "2743052", "source": "2742035", "target": "2742122", "SCORE": 0.3065476190506249, "shared_interaction": "Direct", "source_original": "74", "Name_B": "WP112", "Name_A": "GO:0006757", "target_original": "103", "Gene_B": "WP112", "interaction": "Direct", "SUID": 2743052, "TYPE": "Direct", "Gene_A": "GO:0006757", "BEND_MAP_ID": 2743052, "selected": false}, "selected": false}, {"data": {"id": "2742887", "source": "2742035", "target": "2741867", "SCORE": 0.37421665174792307, "shared_interaction": "Direct", "source_original": "74", "Name_B": "GO:0009150", "Name_A": "GO:0006757", "target_original": "18", "Gene_B": "GO:0009150", "interaction": "Direct", "SUID": 2742887, "TYPE": "Direct", "Gene_A": "GO:0006757", "BEND_MAP_ID": 2742887, "selected": false}, "selected": false}, {"data": {"id": "2744153", "source": "2742032", "target": "2742041", "SCORE": 1.0, "shared_interaction": "Direct", "source_original": "73", "Name_B": "GO:0046031", "Name_A": "GO:0006096", "target_original": "76", "Gene_B": "GO:0046031", "interaction": "Direct", "SUID": 2744153, "TYPE": "Direct", "Gene_A": "GO:0006096", "BEND_MAP_ID": 2744153, "selected": false}, "selected": false}, {"data": {"id": "2744150", "source": "2742032", "target": "2742038", "SCORE": 1.0, "shared_interaction": "Direct", "source_original": "73", "Name_B": "GO:0006165", "Name_A": "GO:0006096", "target_original": "75", "Gene_B": "GO:0006165", "interaction": "Direct", "SUID": 2744150, "TYPE": "Direct", "Gene_A": "GO:0006096", "BEND_MAP_ID": 2744150, "selected": false}, "selected": false}, {"data": {"id": "2744147", "source": "2742032", "target": "2741960", "SCORE": 0.5028128031060302, "shared_interaction": "Direct", "source_original": "73", "Name_B": "GO:0032787", "Name_A": "GO:0006096", "target_original": "49", "Gene_B": "GO:0032787", "interaction": "Direct", "SUID": 2744147, "TYPE": "Direct", "Gene_A": "GO:0006096", "BEND_MAP_ID": 2744147, "selected": false}, "selected": false}, {"data": {"id": "2744144", "source": "2742032", "target": "2742035", "SCORE": 1.0, "shared_interaction": "Direct", "source_original": "73", "Name_B": "GO:0006757", "Name_A": "GO:0006096", "target_original": "74", "Gene_B": "GO:0006757", "interaction": "Direct", "SUID": 2744144, "TYPE": "Direct", "Gene_A": "GO:0006096", "BEND_MAP_ID": 2744144, "selected": false}, "selected": false}, {"data": {"id": "2744132", "source": "2742032", "target": "2741957", "SCORE": 0.37421665174792307, "shared_interaction": "Direct", "source_original": "73", "Name_B": "GO:0016310", "Name_A": "GO:0006096", "target_original": "48", "Gene_B": "GO:0016310", "interaction": "Direct", "SUID": 2744132, "TYPE": "Direct", "Gene_A": "GO:0006096", "BEND_MAP_ID": 2744132, "selected": false}, "selected": false}, {"data": {"id": "2744120", "source": "2742032", "target": "2742029", "SCORE": 0.660844250367735, "shared_interaction": "Direct", "source_original": "73", "Name_B": "R-SCE-170822", "Name_A": "GO:0006096", "target_original": "72", "Gene_B": "R-SCE-170822", "interaction": "Direct", "SUID": 2744120, "TYPE": "Direct", "Gene_A": "GO:0006096", "BEND_MAP_ID": 2744120, "selected": false}, "selected": false}, {"data": {"id": "2744075", "source": "2742032", "target": "2742026", "SCORE": 0.8289280469911845, "shared_interaction": "Direct", "source_original": "73", "Name_B": "R-SCE-70171", "Name_A": "GO:0006096", "target_original": "71", "Gene_B": "R-SCE-70171", "interaction": "Direct", "SUID": 2744075, "TYPE": "Direct", "Gene_A": "GO:0006096", "BEND_MAP_ID": 2744075, "selected": false}, "selected": false}, {"data": {"id": "2744054", "source": "2742032", "target": "2742023", "SCORE": 0.7626485568779389, "shared_interaction": "Direct", "source_original": "73", "Name_B": "R-SCE-70326", "Name_A": "GO:0006096", "target_original": "70", "Gene_B": "R-SCE-70326", "interaction": "Direct", "SUID": 2744054, "TYPE": "Direct", "Gene_A": "GO:0006096", "BEND_MAP_ID": 2744054, "selected": false}, "selected": false}, {"data": {"id": "2744033", "source": "2742032", "target": "2742020", "SCORE": 0.5345585297686158, "shared_interaction": "Direct", "source_original": "73", "Name_B": "GO:0046835", "Name_A": "GO:0006096", "target_original": "69", "Gene_B": "GO:0046835", "interaction": "Direct", "SUID": 2744033, "TYPE": "Direct", "Gene_A": "GO:0006096", "BEND_MAP_ID": 2744033, "selected": false}, "selected": false}, {"data": {"id": "2743988", "source": "2742032", "target": "2742014", "SCORE": 0.3157121879647379, "shared_interaction": "Direct", "source_original": "73", "Name_B": "GO:1904659", "Name_A": "GO:0006096", "target_original": "67", "Gene_B": "GO:1904659", "interaction": "Direct", "SUID": 2743988, "TYPE": "Direct", "Gene_A": "GO:0006096", "BEND_MAP_ID": 2743988, "selected": false}, "selected": false}, {"data": {"id": "2743958", "source": "2742032", "target": "2742017", "SCORE": 0.7416851441258369, "shared_interaction": "Direct", "source_original": "73", "Name_B": "sce00010", "Name_A": "GO:0006096", "target_original": "68", "Gene_B": "sce00010", "interaction": "Direct", "SUID": 2743958, "TYPE": "Direct", "Gene_A": "GO:0006096", "BEND_MAP_ID": 2743958, "selected": false}, "selected": false}, {"data": {"id": "2743391", "source": "2742032", "target": "2741930", "SCORE": 0.45881722351484877, "shared_interaction": "Direct", "source_original": "73", "Name_B": "sce01200", "Name_A": "GO:0006096", "target_original": "39", "Gene_B": "sce01200", "interaction": "Direct", "SUID": 2743391, "TYPE": "Direct", "Gene_A": "GO:0006096", "BEND_MAP_ID": 2743391, "selected": false}, "selected": false}, {"data": {"id": "2743340", "source": "2742032", "target": "2741876", "SCORE": 0.527766518040333, "shared_interaction": "Direct", "source_original": "73", "Name_B": "GO:0016052", "Name_A": "GO:0006096", "target_original": "21", "Gene_B": "GO:0016052", "interaction": "Direct", "SUID": 2743340, "TYPE": "Direct", "Gene_A": "GO:0006096", "BEND_MAP_ID": 2743340, "selected": false}, "selected": false}, {"data": {"id": "2743217", "source": "2742032", "target": "2741882", "SCORE": 0.43331982164839955, "shared_interaction": "Direct", "source_original": "73", "Name_B": "R-SCE-71387", "Name_A": "GO:0006096", "target_original": "23", "Gene_B": "R-SCE-71387", "interaction": "Direct", "SUID": 2743217, "TYPE": "Direct", "Gene_A": "GO:0006096", "BEND_MAP_ID": 2743217, "selected": false}, "selected": false}, {"data": {"id": "2743049", "source": "2742032", "target": "2742122", "SCORE": 0.3065476190506249, "shared_interaction": "Direct", "source_original": "73", "Name_B": "WP112", "Name_A": "GO:0006096", "target_original": "103", "Gene_B": "WP112", "interaction": "Direct", "SUID": 2743049, "TYPE": "Direct", "Gene_A": "GO:0006096", "BEND_MAP_ID": 2743049, "selected": false}, "selected": false}, {"data": {"id": "2742884", "source": "2742032", "target": "2741867", "SCORE": 0.37421665174792307, "shared_interaction": "Direct", "source_original": "73", "Name_B": "GO:0009150", "Name_A": "GO:0006096", "target_original": "18", "Gene_B": "GO:0009150", "interaction": "Direct", "SUID": 2742884, "TYPE": "Direct", "Gene_A": "GO:0006096", "BEND_MAP_ID": 2742884, "selected": false}, "selected": false}, {"data": {"id": "2744069", "source": "2742029", "target": "2742026", "SCORE": 0.660844250367735, "shared_interaction": "Direct", "source_original": "72", "Name_B": "R-SCE-70171", "Name_A": "R-SCE-170822", "target_original": "71", "Gene_B": "R-SCE-70171", "interaction": "Direct", "SUID": 2744069, "TYPE": "Direct", "Gene_A": "R-SCE-170822", "BEND_MAP_ID": 2744069, "selected": false}, "selected": false}, {"data": {"id": "2744048", "source": "2742029", "target": "2742023", "SCORE": 0.5926573426614913, "shared_interaction": "Direct", "source_original": "72", "Name_B": "R-SCE-70326", "Name_A": "R-SCE-170822", "target_original": "70", "Gene_B": "R-SCE-70326", "interaction": "Direct", "SUID": 2744048, "TYPE": "Direct", "Gene_A": "R-SCE-170822", "BEND_MAP_ID": 2744048, "selected": false}, "selected": false}, {"data": {"id": "2743952", "source": "2742029", "target": "2742017", "SCORE": 0.4506567868012127, "shared_interaction": "Direct", "source_original": "72", "Name_B": "sce00010", "Name_A": "R-SCE-170822", "target_original": "68", "Gene_B": "sce00010", "interaction": "Direct", "SUID": 2743952, "TYPE": "Direct", "Gene_A": "R-SCE-170822", "BEND_MAP_ID": 2743952, "selected": false}, "selected": false}, {"data": {"id": "2743214", "source": "2742029", "target": "2741882", "SCORE": 0.30062341261055076, "shared_interaction": "Direct", "source_original": "72", "Name_B": "R-SCE-71387", "Name_A": "R-SCE-170822", "target_original": "23", "Gene_B": "R-SCE-71387", "interaction": "Direct", "SUID": 2743214, "TYPE": "Direct", "Gene_A": "R-SCE-170822", "BEND_MAP_ID": 2743214, "selected": false}, "selected": false}, {"data": {"id": "2742956", "source": "2742029", "target": "2742125", "SCORE": 0.31871345029626597, "shared_interaction": "Direct", "source_original": "72", "Name_B": "sce00500", "Name_A": "R-SCE-170822", "target_original": "104", "Gene_B": "sce00500", "interaction": "Direct", "SUID": 2742956, "TYPE": "Direct", "Gene_A": "R-SCE-170822", "BEND_MAP_ID": 2742956, "selected": false}, "selected": false}, {"data": {"id": "2744045", "source": "2742026", "target": "2742023", "SCORE": 0.9208828522926463, "shared_interaction": "Direct", "source_original": "71", "Name_B": "R-SCE-70326", "Name_A": "R-SCE-70171", "target_original": "70", "Gene_B": "R-SCE-70326", "interaction": "Direct", "SUID": 2744045, "TYPE": "Direct", "Gene_A": "R-SCE-70171", "BEND_MAP_ID": 2744045, "selected": false}, "selected": false}, {"data": {"id": "2743949", "source": "2742026", "target": "2742017", "SCORE": 0.6125277161887553, "shared_interaction": "Direct", "source_original": "71", "Name_B": "sce00010", "Name_A": "R-SCE-70171", "target_original": "68", "Gene_B": "sce00010", "interaction": "Direct", "SUID": 2743949, "TYPE": "Direct", "Gene_A": "R-SCE-70171", "BEND_MAP_ID": 2743949, "selected": false}, "selected": false}, {"data": {"id": "2743385", "source": "2742026", "target": "2741930", "SCORE": 0.3755583348248255, "shared_interaction": "Direct", "source_original": "71", "Name_B": "sce01200", "Name_A": "R-SCE-70171", "target_original": "39", "Gene_B": "sce01200", "interaction": "Direct", "SUID": 2743385, "TYPE": "Direct", "Gene_A": "R-SCE-70171", "BEND_MAP_ID": 2743385, "selected": false}, "selected": false}, {"data": {"id": "2743211", "source": "2742026", "target": "2741882", "SCORE": 0.527766518040333, "shared_interaction": "Direct", "source_original": "71", "Name_B": "R-SCE-71387", "Name_A": "R-SCE-70171", "target_original": "23", "Gene_B": "R-SCE-71387", "interaction": "Direct", "SUID": 2743211, "TYPE": "Direct", "Gene_A": "R-SCE-70171", "BEND_MAP_ID": 2743211, "selected": false}, "selected": false}, {"data": {"id": "2743046", "source": "2742026", "target": "2742122", "SCORE": 0.3065476190506249, "shared_interaction": "Direct", "source_original": "71", "Name_B": "WP112", "Name_A": "R-SCE-70171", "target_original": "103", "Gene_B": "WP112", "interaction": "Direct", "SUID": 2743046, "TYPE": "Direct", "Gene_A": "R-SCE-70171", "BEND_MAP_ID": 2743046, "selected": false}, "selected": false}, {"data": {"id": "2743946", "source": "2742023", "target": "2742017", "SCORE": 0.5731483904763008, "shared_interaction": "Direct", "source_original": "70", "Name_B": "sce00010", "Name_A": "R-SCE-70326", "target_original": "68", "Gene_B": "sce00010", "interaction": "Direct", "SUID": 2743946, "TYPE": "Direct", "Gene_A": "R-SCE-70326", "BEND_MAP_ID": 2743946, "selected": false}, "selected": false}, {"data": {"id": "2743382", "source": "2742023", "target": "2741930", "SCORE": 0.43680939226745896, "shared_interaction": "Direct", "source_original": "70", "Name_B": "sce01200", "Name_A": "R-SCE-70326", "target_original": "39", "Gene_B": "sce01200", "interaction": "Direct", "SUID": 2743382, "TYPE": "Direct", "Gene_A": "R-SCE-70326", "BEND_MAP_ID": 2743382, "selected": false}, "selected": false}, {"data": {"id": "2743208", "source": "2742023", "target": "2741882", "SCORE": 0.5916260954254166, "shared_interaction": "Direct", "source_original": "70", "Name_B": "R-SCE-71387", "Name_A": "R-SCE-70326", "target_original": "23", "Gene_B": "R-SCE-71387", "interaction": "Direct", "SUID": 2743208, "TYPE": "Direct", "Gene_A": "R-SCE-70326", "BEND_MAP_ID": 2743208, "selected": false}, "selected": false}, {"data": {"id": "2743043", "source": "2742023", "target": "2742122", "SCORE": 0.3728691907436951, "shared_interaction": "Direct", "source_original": "70", "Name_B": "WP112", "Name_A": "R-SCE-70326", "target_original": "103", "Gene_B": "WP112", "interaction": "Direct", "SUID": 2743043, "TYPE": "Direct", "Gene_A": "R-SCE-70326", "BEND_MAP_ID": 2743043, "selected": false}, "selected": false}, {"data": {"id": "2744027", "source": "2742020", "target": "2742029", "SCORE": 0.7459105779748748, "shared_interaction": "Direct", "source_original": "69", "Name_B": "R-SCE-170822", "Name_A": "GO:0046835", "target_original": "72", "Gene_B": "R-SCE-170822", "interaction": "Direct", "SUID": 2744027, "TYPE": "Direct", "Gene_A": "GO:0046835", "BEND_MAP_ID": 2744027, "selected": false}, "selected": false}, {"data": {"id": "2744024", "source": "2742020", "target": "2742026", "SCORE": 0.7207351178611695, "shared_interaction": "Direct", "source_original": "69", "Name_B": "R-SCE-70171", "Name_A": "GO:0046835", "target_original": "71", "Gene_B": "R-SCE-70171", "interaction": "Direct", "SUID": 2744024, "TYPE": "Direct", "Gene_A": "GO:0046835", "BEND_MAP_ID": 2744024, "selected": false}, "selected": false}, {"data": {"id": "2744021", "source": "2742020", "target": "2742023", "SCORE": 0.658107116657361, "shared_interaction": "Direct", "source_original": "69", "Name_B": "R-SCE-70326", "Name_A": "GO:0046835", "target_original": "70", "Gene_B": "R-SCE-70326", "interaction": "Direct", "SUID": 2744021, "TYPE": "Direct", "Gene_A": "GO:0046835", "BEND_MAP_ID": 2744021, "selected": false}, "selected": false}, {"data": {"id": "2743979", "source": "2742020", "target": "2742014", "SCORE": 0.3483819416760622, "shared_interaction": "Direct", "source_original": "69", "Name_B": "GO:1904659", "Name_A": "GO:0046835", "target_original": "67", "Gene_B": "GO:1904659", "interaction": "Direct", "SUID": 2743979, "TYPE": "Direct", "Gene_A": "GO:0046835", "BEND_MAP_ID": 2743979, "selected": false}, "selected": false}, {"data": {"id": "2743943", "source": "2742020", "target": "2742017", "SCORE": 0.3823269514033555, "shared_interaction": "Direct", "source_original": "69", "Name_B": "sce00010", "Name_A": "GO:0046835", "target_original": "68", "Gene_B": "sce00010", "interaction": "Direct", "SUID": 2743943, "TYPE": "Direct", "Gene_A": "GO:0046835", "BEND_MAP_ID": 2743943, "selected": false}, "selected": false}, {"data": {"id": "2743205", "source": "2742020", "target": "2741882", "SCORE": 0.4584829917626492, "shared_interaction": "Direct", "source_original": "69", "Name_B": "R-SCE-71387", "Name_A": "GO:0046835", "target_original": "23", "Gene_B": "R-SCE-71387", "interaction": "Direct", "SUID": 2743205, "TYPE": "Direct", "Gene_A": "GO:0046835", "BEND_MAP_ID": 2743205, "selected": false}, "selected": false}, {"data": {"id": "2743379", "source": "2742017", "target": "2741930", "SCORE": 0.37886153363877556, "shared_interaction": "Direct", "source_original": "68", "Name_B": "sce01200", "Name_A": "sce00010", "target_original": "39", "Gene_B": "sce01200", "interaction": "Direct", "SUID": 2743379, "TYPE": "Direct", "Gene_A": "sce00010", "BEND_MAP_ID": 2743379, "selected": false}, "selected": false}, {"data": {"id": "2743088", "source": "2742017", "target": "2741927", "SCORE": 0.3471029557999829, "shared_interaction": "Direct", "source_original": "68", "Name_B": "sce01110", "Name_A": "sce00010", "target_original": "38", "Gene_B": "sce01110", "interaction": "Direct", "SUID": 2743088, "TYPE": "Direct", "Gene_A": "sce00010", "BEND_MAP_ID": 2743088, "selected": false}, "selected": false}, {"data": {"id": "2743985", "source": "2742014", "target": "2742029", "SCORE": 0.43474041727955826, "shared_interaction": "Direct", "source_original": "67", "Name_B": "R-SCE-170822", "Name_A": "GO:1904659", "target_original": "72", "Gene_B": "R-SCE-170822", "interaction": "Direct", "SUID": 2743985, "TYPE": "Direct", "Gene_A": "GO:1904659", "BEND_MAP_ID": 2743985, "selected": false}, "selected": false}, {"data": {"id": "2743982", "source": "2742014", "target": "2742026", "SCORE": 0.3157121879647379, "shared_interaction": "Direct", "source_original": "67", "Name_B": "R-SCE-70171", "Name_A": "GO:1904659", "target_original": "71", "Gene_B": "R-SCE-70171", "interaction": "Direct", "SUID": 2743982, "TYPE": "Direct", "Gene_A": "GO:1904659", "BEND_MAP_ID": 2743982, "selected": false}, "selected": false}, {"data": {"id": "2743940", "source": "2742011", "target": "2742029", "SCORE": 0.3202042304944703, "shared_interaction": "Direct", "source_original": "66", "Name_B": "R-SCE-170822", "Name_A": "GO:0008643", "target_original": "72", "Gene_B": "R-SCE-170822", "interaction": "Direct", "SUID": 2743940, "TYPE": "Direct", "Gene_A": "GO:0008643", "BEND_MAP_ID": 2743940, "selected": false}, "selected": false}, {"data": {"id": "2743937", "source": "2742011", "target": "2742014", "SCORE": 0.7936226749349893, "shared_interaction": "Direct", "source_original": "66", "Name_B": "GO:1904659", "Name_A": "GO:0008643", "target_original": "67", "Gene_B": "GO:1904659", "interaction": "Direct", "SUID": 2743937, "TYPE": "Direct", "Gene_A": "GO:0008643", "BEND_MAP_ID": 2743937, "selected": false}, "selected": false}, {"data": {"id": "2743835", "source": "2742011", "target": "2742008", "SCORE": 0.8706274292068336, "shared_interaction": "Direct", "source_original": "66", "Name_B": "GO:0015749", "Name_A": "GO:0008643", "target_original": "65", "Gene_B": "GO:0015749", "interaction": "Direct", "SUID": 2743835, "TYPE": "Direct", "Gene_A": "GO:0008643", "BEND_MAP_ID": 2743835, "selected": false}, "selected": false}, {"data": {"id": "2743823", "source": "2742011", "target": "2742005", "SCORE": 0.8706274292068336, "shared_interaction": "Direct", "source_original": "66", "Name_B": "GO:0008645", "Name_A": "GO:0008643", "target_original": "64", "Gene_B": "GO:0008645", "interaction": "Direct", "SUID": 2743823, "TYPE": "Direct", "Gene_A": "GO:0008643", "BEND_MAP_ID": 2743823, "selected": false}, "selected": false}, {"data": {"id": "2743718", "source": "2742011", "target": "2742134", "SCORE": 1.0, "shared_interaction": "Direct", "source_original": "66", "Name_B": "GO:0034219", "Name_A": "GO:0008643", "target_original": "107", "Gene_B": "GO:0034219", "interaction": "Direct", "SUID": 2743718, "TYPE": "Direct", "Gene_A": "GO:0008643", "BEND_MAP_ID": 2743718, "selected": false}, "selected": false}, {"data": {"id": "2743844", "source": "2742008", "target": "2742029", "SCORE": 0.3889860139922369, "shared_interaction": "Direct", "source_original": "65", "Name_B": "R-SCE-170822", "Name_A": "GO:0015749", "target_original": "72", "Gene_B": "R-SCE-170822", "interaction": "Direct", "SUID": 2743844, "TYPE": "Direct", "Gene_A": "GO:0015749", "BEND_MAP_ID": 2743844, "selected": false}, "selected": false}, {"data": {"id": "2743841", "source": "2742008", "target": "2742020", "SCORE": 0.31621423331472187, "shared_interaction": "Direct", "source_original": "65", "Name_B": "GO:0046835", "Name_A": "GO:0015749", "target_original": "69", "Gene_B": "GO:0046835", "interaction": "Direct", "SUID": 2743841, "TYPE": "Direct", "Gene_A": "GO:0015749", "BEND_MAP_ID": 2743841, "selected": false}, "selected": false}, {"data": {"id": "2743838", "source": "2742008", "target": "2742014", "SCORE": 0.9208828522926463, "shared_interaction": "Direct", "source_original": "65", "Name_B": "GO:1904659", "Name_A": "GO:0015749", "target_original": "67", "Gene_B": "GO:1904659", "interaction": "Direct", "SUID": 2743838, "TYPE": "Direct", "Gene_A": "GO:0015749", "BEND_MAP_ID": 2743838, "selected": false}, "selected": false}, {"data": {"id": "2743715", "source": "2742008", "target": "2742134", "SCORE": 0.8706274292068336, "shared_interaction": "Direct", "source_original": "65", "Name_B": "GO:0034219", "Name_A": "GO:0015749", "target_original": "107", "Gene_B": "GO:0034219", "interaction": "Direct", "SUID": 2743715, "TYPE": "Direct", "Gene_A": "GO:0015749", "BEND_MAP_ID": 2743715, "selected": false}, "selected": false}, {"data": {"id": "2743832", "source": "2742005", "target": "2742029", "SCORE": 0.3889860139922369, "shared_interaction": "Direct", "source_original": "64", "Name_B": "R-SCE-170822", "Name_A": "GO:0008645", "target_original": "72", "Gene_B": "R-SCE-170822", "interaction": "Direct", "SUID": 2743832, "TYPE": "Direct", "Gene_A": "GO:0008645", "BEND_MAP_ID": 2743832, "selected": false}, "selected": false}, {"data": {"id": "2743829", "source": "2742005", "target": "2742020", "SCORE": 0.31621423331472187, "shared_interaction": "Direct", "source_original": "64", "Name_B": "GO:0046835", "Name_A": "GO:0008645", "target_original": "69", "Gene_B": "GO:0046835", "interaction": "Direct", "SUID": 2743829, "TYPE": "Direct", "Gene_A": "GO:0008645", "BEND_MAP_ID": 2743829, "selected": false}, "selected": false}, {"data": {"id": "2743826", "source": "2742005", "target": "2742014", "SCORE": 0.9208828522926463, "shared_interaction": "Direct", "source_original": "64", "Name_B": "GO:1904659", "Name_A": "GO:0008645", "target_original": "67", "Gene_B": "GO:1904659", "interaction": "Direct", "SUID": 2743826, "TYPE": "Direct", "Gene_A": "GO:0008645", "BEND_MAP_ID": 2743826, "selected": false}, "selected": false}, {"data": {"id": "2743820", "source": "2742005", "target": "2742008", "SCORE": 1.0, "shared_interaction": "Direct", "source_original": "64", "Name_B": "GO:0015749", "Name_A": "GO:0008645", "target_original": "65", "Gene_B": "GO:0015749", "interaction": "Direct", "SUID": 2743820, "TYPE": "Direct", "Gene_A": "GO:0008645", "BEND_MAP_ID": 2743820, "selected": false}, "selected": false}, {"data": {"id": "2743712", "source": "2742005", "target": "2742134", "SCORE": 0.8706274292068336, "shared_interaction": "Direct", "source_original": "64", "Name_B": "GO:0034219", "Name_A": "GO:0008645", "target_original": "107", "Gene_B": "GO:0034219", "interaction": "Direct", "SUID": 2743712, "TYPE": "Direct", "Gene_A": "GO:0008645", "BEND_MAP_ID": 2743712, "selected": false}, "selected": false}, {"data": {"id": "2743688", "source": "2742002", "target": "2742128", "SCORE": 0.4336412250914903, "shared_interaction": "Direct", "source_original": "63", "Name_B": "WP70", "Name_A": "GO:0044275", "target_original": "105", "Gene_B": "WP70", "interaction": "Direct", "SUID": 2743688, "TYPE": "Direct", "Gene_A": "GO:0044275", "BEND_MAP_ID": 2743688, "selected": false}, "selected": false}, {"data": {"id": "2743586", "source": "2741999", "target": "2742137", "SCORE": 0.9208828522926463, "shared_interaction": "Direct", "source_original": "62", "Name_B": "GO:0042407", "Name_A": "GO:0007007", "target_original": "108", "Gene_B": "GO:0042407", "interaction": "Direct", "SUID": 2743586, "TYPE": "Direct", "Gene_A": "GO:0007007", "BEND_MAP_ID": 2743586, "selected": false}, "selected": false}, {"data": {"id": "2743913", "source": "2741996", "target": "2741999", "SCORE": 0.36970243462768076, "shared_interaction": "Direct", "source_original": "61", "Name_B": "GO:0007007", "Name_A": "GO:0007005", "target_original": "62", "Gene_B": "GO:0007007", "interaction": "Direct", "SUID": 2743913, "TYPE": "Direct", "Gene_A": "GO:0007005", "BEND_MAP_ID": 2743913, "selected": false}, "selected": false}, {"data": {"id": "2743583", "source": "2741996", "target": "2742137", "SCORE": 0.32428157131765667, "shared_interaction": "Direct", "source_original": "61", "Name_B": "GO:0042407", "Name_A": "GO:0007005", "target_original": "108", "Gene_B": "GO:0042407", "interaction": "Direct", "SUID": 2743583, "TYPE": "Direct", "Gene_A": "GO:0007005", "BEND_MAP_ID": 2743583, "selected": false}, "selected": false}, {"data": {"id": "2744105", "source": "2741993", "target": "2741990", "SCORE": 0.6601847350548942, "shared_interaction": "Direct", "source_original": "60", "Name_B": "GO:0042149", "Name_A": "GO:0007039", "target_original": "59", "Gene_B": "GO:0042149", "interaction": "Direct", "SUID": 2744105, "TYPE": "Direct", "Gene_A": "GO:0007039", "BEND_MAP_ID": 2744105, "selected": false}, "selected": false}, {"data": {"id": "2743709", "source": "2741993", "target": "2741981", "SCORE": 0.3354939071857619, "shared_interaction": "Direct", "source_original": "60", "Name_B": "GO:0009267", "Name_A": "GO:0007039", "target_original": "56", "Gene_B": "GO:0009267", "interaction": "Direct", "SUID": 2743709, "TYPE": "Direct", "Gene_A": "GO:0007039", "BEND_MAP_ID": 2743709, "selected": false}, "selected": false}, {"data": {"id": "2743673", "source": "2741993", "target": "2741978", "SCORE": 0.31504164625574743, "shared_interaction": "Direct", "source_original": "60", "Name_B": "GO:0042594", "Name_A": "GO:0007039", "target_original": "55", "Gene_B": "GO:0042594", "interaction": "Direct", "SUID": 2743673, "TYPE": "Direct", "Gene_A": "GO:0007039", "BEND_MAP_ID": 2743673, "selected": false}, "selected": false}, {"data": {"id": "2743670", "source": "2741990", "target": "2741978", "SCORE": 0.5108467459788656, "shared_interaction": "Direct", "source_original": "59", "Name_B": "GO:0042594", "Name_A": "GO:0042149", "target_original": "55", "Gene_B": "GO:0042594", "interaction": "Direct", "SUID": 2743670, "TYPE": "Direct", "Gene_A": "GO:0042149", "BEND_MAP_ID": 2743670, "selected": false}, "selected": false}, {"data": {"id": "2743580", "source": "2741990", "target": "2741972", "SCORE": 0.3786666666692404, "shared_interaction": "Direct", "source_original": "59", "Name_B": "GO:0071496", "Name_A": "GO:0042149", "target_original": "53", "Gene_B": "GO:0071496", "interaction": "Direct", "SUID": 2743580, "TYPE": "Direct", "Gene_A": "GO:0042149", "BEND_MAP_ID": 2743580, "selected": false}, "selected": false}, {"data": {"id": "2743790", "source": "2741987", "target": "2741990", "SCORE": 0.43583535109224075, "shared_interaction": "Direct", "source_original": "58", "Name_B": "GO:0042149", "Name_A": "GO:0031669", "target_original": "59", "Gene_B": "GO:0042149", "interaction": "Direct", "SUID": 2743790, "TYPE": "Direct", "Gene_A": "GO:0031669", "BEND_MAP_ID": 2743790, "selected": false}, "selected": false}, {"data": {"id": "2743667", "source": "2741987", "target": "2741978", "SCORE": 0.8273306654815508, "shared_interaction": "Direct", "source_original": "58", "Name_B": "GO:0042594", "Name_A": "GO:0031669", "target_original": "55", "Gene_B": "GO:0042594", "interaction": "Direct", "SUID": 2743667, "TYPE": "Direct", "Gene_A": "GO:0031669", "BEND_MAP_ID": 2743667, "selected": false}, "selected": false}, {"data": {"id": "2743577", "source": "2741987", "target": "2741972", "SCORE": 0.9119758216851845, "shared_interaction": "Direct", "source_original": "58", "Name_B": "GO:0071496", "Name_A": "GO:0031669", "target_original": "53", "Gene_B": "GO:0071496", "interaction": "Direct", "SUID": 2743577, "TYPE": "Direct", "Gene_A": "GO:0031669", "BEND_MAP_ID": 2743577, "selected": false}, "selected": false}, {"data": {"id": "2743760", "source": "2741984", "target": "2741990", "SCORE": 0.41513805754269667, "shared_interaction": "Direct", "source_original": "57", "Name_B": "GO:0042149", "Name_A": "GO:0031667", "target_original": "59", "Gene_B": "GO:0042149", "interaction": "Direct", "SUID": 2743760, "TYPE": "Direct", "Gene_A": "GO:0031667", "BEND_MAP_ID": 2743760, "selected": false}, "selected": false}, {"data": {"id": "2743757", "source": "2741984", "target": "2741987", "SCORE": 0.9691104335146121, "shared_interaction": "Direct", "source_original": "57", "Name_B": "GO:0031669", "Name_A": "GO:0031667", "target_original": "58", "Gene_B": "GO:0031669", "interaction": "Direct", "SUID": 2743757, "TYPE": "Direct", "Gene_A": "GO:0031667", "BEND_MAP_ID": 2743757, "selected": false}, "selected": false}, {"data": {"id": "2743664", "source": "2741984", "target": "2741978", "SCORE": 0.865937859609195, "shared_interaction": "Direct", "source_original": "57", "Name_B": "GO:0042594", "Name_A": "GO:0031667", "target_original": "55", "Gene_B": "GO:0042594", "interaction": "Direct", "SUID": 2743664, "TYPE": "Direct", "Gene_A": "GO:0031667", "BEND_MAP_ID": 2743664, "selected": false}, "selected": false}, {"data": {"id": "2743574", "source": "2741984", "target": "2741972", "SCORE": 0.8854192279324649, "shared_interaction": "Direct", "source_original": "57", "Name_B": "GO:0071496", "Name_A": "GO:0031667", "target_original": "53", "Gene_B": "GO:0071496", "interaction": "Direct", "SUID": 2743574, "TYPE": "Direct", "Gene_A": "GO:0031667", "BEND_MAP_ID": 2743574, "selected": false}, "selected": false}, {"data": {"id": "2743556", "source": "2741984", "target": "2741969", "SCORE": 0.8854192279324649, "shared_interaction": "Direct", "source_original": "57", "Name_B": "GO:0031668", "Name_A": "GO:0031667", "target_original": "52", "Gene_B": "GO:0031668", "interaction": "Direct", "SUID": 2743556, "TYPE": "Direct", "Gene_A": "GO:0031667", "BEND_MAP_ID": 2743556, "selected": false}, "selected": false}, {"data": {"id": "2743706", "source": "2741981", "target": "2741990", "SCORE": 0.5413385826797952, "shared_interaction": "Direct", "source_original": "56", "Name_B": "GO:0042149", "Name_A": "GO:0009267", "target_original": "59", "Gene_B": "GO:0042149", "interaction": "Direct", "SUID": 2743706, "TYPE": "Direct", "Gene_A": "GO:0009267", "BEND_MAP_ID": 2743706, "selected": false}, "selected": false}, {"data": {"id": "2743703", "source": "2741981", "target": "2741987", "SCORE": 0.8576664630426567, "shared_interaction": "Direct", "source_original": "56", "Name_B": "GO:0031669", "Name_A": "GO:0009267", "target_original": "58", "Gene_B": "GO:0031669", "interaction": "Direct", "SUID": 2743703, "TYPE": "Direct", "Gene_A": "GO:0009267", "BEND_MAP_ID": 2743703, "selected": false}, "selected": false}, {"data": {"id": "2743700", "source": "2741981", "target": "2741984", "SCORE": 0.8275351591419715, "shared_interaction": "Direct", "source_original": "56", "Name_B": "GO:0031667", "Name_A": "GO:0009267", "target_original": "57", "Gene_B": "GO:0031667", "interaction": "Direct", "SUID": 2743700, "TYPE": "Direct", "Gene_A": "GO:0009267", "BEND_MAP_ID": 2743700, "selected": false}, "selected": false}, {"data": {"id": "2743661", "source": "2741981", "target": "2741978", "SCORE": 0.9606883752321438, "shared_interaction": "Direct", "source_original": "56", "Name_B": "GO:0042594", "Name_A": "GO:0009267", "target_original": "55", "Gene_B": "GO:0042594", "interaction": "Direct", "SUID": 2743661, "TYPE": "Direct", "Gene_A": "GO:0009267", "BEND_MAP_ID": 2743661, "selected": false}, "selected": false}, {"data": {"id": "2743601", "source": "2741981", "target": "2741975", "SCORE": 0.7472885032545943, "shared_interaction": "Direct", "source_original": "56", "Name_B": "GO:0009605", "Name_A": "GO:0009267", "target_original": "54", "Gene_B": "GO:0009605", "interaction": "Direct", "SUID": 2743601, "TYPE": "Direct", "Gene_A": "GO:0009267", "BEND_MAP_ID": 2743601, "selected": false}, "selected": false}, {"data": {"id": "2743571", "source": "2741981", "target": "2741972", "SCORE": 0.7724926768036408, "shared_interaction": "Direct", "source_original": "56", "Name_B": "GO:0071496", "Name_A": "GO:0009267", "target_original": "53", "Gene_B": "GO:0071496", "interaction": "Direct", "SUID": 2743571, "TYPE": "Direct", "Gene_A": "GO:0009267", "BEND_MAP_ID": 2743571, "selected": false}, "selected": false}, {"data": {"id": "2743553", "source": "2741981", "target": "2741969", "SCORE": 0.7724926768036408, "shared_interaction": "Direct", "source_original": "56", "Name_B": "GO:0031668", "Name_A": "GO:0009267", "target_original": "52", "Gene_B": "GO:0031668", "interaction": "Direct", "SUID": 2743553, "TYPE": "Direct", "Gene_A": "GO:0009267", "BEND_MAP_ID": 2743553, "selected": false}, "selected": false}, {"data": {"id": "2743442", "source": "2741981", "target": "2742158", "SCORE": 0.7472885032545943, "shared_interaction": "Direct", "source_original": "56", "Name_B": "GO:0009991", "Name_A": "GO:0009267", "target_original": "115", "Gene_B": "GO:0009991", "interaction": "Direct", "SUID": 2743442, "TYPE": "Direct", "Gene_A": "GO:0009267", "BEND_MAP_ID": 2743442, "selected": false}, "selected": false}, {"data": {"id": "2743568", "source": "2741978", "target": "2741972", "SCORE": 0.746807932627805, "shared_interaction": "Direct", "source_original": "55", "Name_B": "GO:0071496", "Name_A": "GO:0042594", "target_original": "53", "Gene_B": "GO:0071496", "interaction": "Direct", "SUID": 2743568, "TYPE": "Direct", "Gene_A": "GO:0042594", "BEND_MAP_ID": 2743568, "selected": false}, "selected": false}, {"data": {"id": "2743610", "source": "2741975", "target": "2741990", "SCORE": 0.3625170998657409, "shared_interaction": "Direct", "source_original": "54", "Name_B": "GO:0042149", "Name_A": "GO:0009605", "target_original": "59", "Gene_B": "GO:0042149", "interaction": "Direct", "SUID": 2743610, "TYPE": "Direct", "Gene_A": "GO:0009605", "BEND_MAP_ID": 2743610, "selected": false}, "selected": false}, {"data": {"id": "2743607", "source": "2741975", "target": "2741987", "SCORE": 0.8855036855040131, "shared_interaction": "Direct", "source_original": "54", "Name_B": "GO:0031669", "Name_A": "GO:0009605", "target_original": "58", "Gene_B": "GO:0031669", "interaction": "Direct", "SUID": 2743607, "TYPE": "Direct", "Gene_A": "GO:0009605", "BEND_MAP_ID": 2743607, "selected": false}, "selected": false}, {"data": {"id": "2743604", "source": "2741975", "target": "2741984", "SCORE": 0.9160965070222008, "shared_interaction": "Direct", "source_original": "54", "Name_B": "GO:0031667", "Name_A": "GO:0009605", "target_original": "57", "Gene_B": "GO:0031667", "interaction": "Direct", "SUID": 2743604, "TYPE": "Direct", "Gene_A": "GO:0009605", "BEND_MAP_ID": 2743604, "selected": false}, "selected": false}, {"data": {"id": "2743598", "source": "2741975", "target": "2741978", "SCORE": 0.7844588344132445, "shared_interaction": "Direct", "source_original": "54", "Name_B": "GO:0042594", "Name_A": "GO:0009605", "target_original": "55", "Gene_B": "GO:0042594", "interaction": "Direct", "SUID": 2743598, "TYPE": "Direct", "Gene_A": "GO:0009605", "BEND_MAP_ID": 2743598, "selected": false}, "selected": false}, {"data": {"id": "2743565", "source": "2741975", "target": "2741972", "SCORE": 0.9732583495926344, "shared_interaction": "Direct", "source_original": "54", "Name_B": "GO:0071496", "Name_A": "GO:0009605", "target_original": "53", "Gene_B": "GO:0071496", "interaction": "Direct", "SUID": 2743565, "TYPE": "Direct", "Gene_A": "GO:0009605", "BEND_MAP_ID": 2743565, "selected": false}, "selected": false}, {"data": {"id": "2743547", "source": "2741975", "target": "2741969", "SCORE": 0.9732583495926344, "shared_interaction": "Direct", "source_original": "54", "Name_B": "GO:0031668", "Name_A": "GO:0009605", "target_original": "52", "Gene_B": "GO:0031668", "interaction": "Direct", "SUID": 2743547, "TYPE": "Direct", "Gene_A": "GO:0009605", "BEND_MAP_ID": 2743547, "selected": false}, "selected": false}, {"data": {"id": "2743436", "source": "2741975", "target": "2742158", "SCORE": 1.0, "shared_interaction": "Direct", "source_original": "54", "Name_B": "GO:0009991", "Name_A": "GO:0009605", "target_original": "115", "Gene_B": "GO:0009991", "interaction": "Direct", "SUID": 2743436, "TYPE": "Direct", "Gene_A": "GO:0009605", "BEND_MAP_ID": 2743436, "selected": false}, "selected": false}, {"data": {"id": "2743562", "source": "2741969", "target": "2741990", "SCORE": 0.3786666666692404, "shared_interaction": "Direct", "source_original": "52", "Name_B": "GO:0042149", "Name_A": "GO:0031668", "target_original": "59", "Gene_B": "GO:0042149", "interaction": "Direct", "SUID": 2743562, "TYPE": "Direct", "Gene_A": "GO:0031668", "BEND_MAP_ID": 2743562, "selected": false}, "selected": false}, {"data": {"id": "2743559", "source": "2741969", "target": "2741987", "SCORE": 0.9119758216851845, "shared_interaction": "Direct", "source_original": "52", "Name_B": "GO:0031669", "Name_A": "GO:0031668", "target_original": "58", "Gene_B": "GO:0031669", "interaction": "Direct", "SUID": 2743559, "TYPE": "Direct", "Gene_A": "GO:0031668", "BEND_MAP_ID": 2743559, "selected": false}, "selected": false}, {"data": {"id": "2743550", "source": "2741969", "target": "2741978", "SCORE": 0.746807932627805, "shared_interaction": "Direct", "source_original": "52", "Name_B": "GO:0042594", "Name_A": "GO:0031668", "target_original": "55", "Gene_B": "GO:0042594", "interaction": "Direct", "SUID": 2743550, "TYPE": "Direct", "Gene_A": "GO:0031668", "BEND_MAP_ID": 2743550, "selected": false}, "selected": false}, {"data": {"id": "2743544", "source": "2741969", "target": "2741972", "SCORE": 1.0, "shared_interaction": "Direct", "source_original": "52", "Name_B": "GO:0071496", "Name_A": "GO:0031668", "target_original": "53", "Gene_B": "GO:0071496", "interaction": "Direct", "SUID": 2743544, "TYPE": "Direct", "Gene_A": "GO:0031668", "BEND_MAP_ID": 2743544, "selected": false}, "selected": false}, {"data": {"id": "2744261", "source": "2741966", "target": "2741963", "SCORE": 1.0, "shared_interaction": "Direct", "source_original": "51", "Name_B": "GO:0043436", "Name_A": "GO:0006082", "target_original": "50", "Gene_B": "GO:0043436", "interaction": "Direct", "SUID": 2744261, "TYPE": "Direct", "Gene_A": "GO:0006082", "BEND_MAP_ID": 2744261, "selected": false}, "selected": false}, {"data": {"id": "2744183", "source": "2741966", "target": "2741960", "SCORE": 0.6616322974158083, "shared_interaction": "Direct", "source_original": "51", "Name_B": "GO:0032787", "Name_A": "GO:0006082", "target_original": "49", "Gene_B": "GO:0032787", "interaction": "Direct", "SUID": 2744183, "TYPE": "Direct", "Gene_A": "GO:0006082", "BEND_MAP_ID": 2744183, "selected": false}, "selected": false}, {"data": {"id": "2743976", "source": "2741966", "target": "2742017", "SCORE": 0.3885851738480479, "shared_interaction": "Direct", "source_original": "51", "Name_B": "sce00010", "Name_A": "GO:0006082", "target_original": "68", "Gene_B": "sce00010", "interaction": "Direct", "SUID": 2743976, "TYPE": "Direct", "Gene_A": "GO:0006082", "BEND_MAP_ID": 2743976, "selected": false}, "selected": false}, {"data": {"id": "2743409", "source": "2741966", "target": "2741930", "SCORE": 0.41023718008941035, "shared_interaction": "Direct", "source_original": "51", "Name_B": "sce01200", "Name_A": "GO:0006082", "target_original": "39", "Gene_B": "sce01200", "interaction": "Direct", "SUID": 2743409, "TYPE": "Direct", "Gene_A": "GO:0006082", "BEND_MAP_ID": 2743409, "selected": false}, "selected": false}, {"data": {"id": "2743283", "source": "2741966", "target": "2742164", "SCORE": 0.5919439579689967, "shared_interaction": "Direct", "source_original": "51", "Name_B": "GO:0044281", "Name_A": "GO:0006082", "target_original": "117", "Gene_B": "GO:0044281", "interaction": "Direct", "SUID": 2743283, "TYPE": "Direct", "Gene_A": "GO:0006082", "BEND_MAP_ID": 2743283, "selected": false}, "selected": false}, {"data": {"id": "2743097", "source": "2741966", "target": "2741927", "SCORE": 0.4330900243318186, "shared_interaction": "Direct", "source_original": "51", "Name_B": "sce01110", "Name_A": "GO:0006082", "target_original": "38", "Gene_B": "sce01110", "interaction": "Direct", "SUID": 2743097, "TYPE": "Direct", "Gene_A": "GO:0006082", "BEND_MAP_ID": 2743097, "selected": false}, "selected": false}, {"data": {"id": "2743973", "source": "2741963", "target": "2742017", "SCORE": 0.3885851738480479, "shared_interaction": "Direct", "source_original": "50", "Name_B": "sce00010", "Name_A": "GO:0043436", "target_original": "68", "Gene_B": "sce00010", "interaction": "Direct", "SUID": 2743973, "TYPE": "Direct", "Gene_A": "GO:0043436", "BEND_MAP_ID": 2743973, "selected": false}, "selected": false}, {"data": {"id": "2743406", "source": "2741963", "target": "2741930", "SCORE": 0.41023718008941035, "shared_interaction": "Direct", "source_original": "50", "Name_B": "sce01200", "Name_A": "GO:0043436", "target_original": "39", "Gene_B": "sce01200", "interaction": "Direct", "SUID": 2743406, "TYPE": "Direct", "Gene_A": "GO:0043436", "BEND_MAP_ID": 2743406, "selected": false}, "selected": false}, {"data": {"id": "2743280", "source": "2741963", "target": "2742164", "SCORE": 0.5919439579689967, "shared_interaction": "Direct", "source_original": "50", "Name_B": "GO:0044281", "Name_A": "GO:0043436", "target_original": "117", "Gene_B": "GO:0044281", "interaction": "Direct", "SUID": 2743280, "TYPE": "Direct", "Gene_A": "GO:0043436", "BEND_MAP_ID": 2743280, "selected": false}, "selected": false}, {"data": {"id": "2743094", "source": "2741963", "target": "2741927", "SCORE": 0.4330900243318186, "shared_interaction": "Direct", "source_original": "50", "Name_B": "sce01110", "Name_A": "GO:0043436", "target_original": "38", "Gene_B": "sce01110", "interaction": "Direct", "SUID": 2743094, "TYPE": "Direct", "Gene_A": "GO:0043436", "BEND_MAP_ID": 2743094, "selected": false}, "selected": false}, {"data": {"id": "2744180", "source": "2741960", "target": "2741963", "SCORE": 0.6616322974158083, "shared_interaction": "Direct", "source_original": "49", "Name_B": "GO:0043436", "Name_A": "GO:0032787", "target_original": "50", "Gene_B": "GO:0043436", "interaction": "Direct", "SUID": 2744180, "TYPE": "Direct", "Gene_A": "GO:0032787", "BEND_MAP_ID": 2744180, "selected": false}, "selected": false}, {"data": {"id": "2744177", "source": "2741960", "target": "2742041", "SCORE": 0.5028128031060302, "shared_interaction": "Direct", "source_original": "49", "Name_B": "GO:0046031", "Name_A": "GO:0032787", "target_original": "76", "Gene_B": "GO:0046031", "interaction": "Direct", "SUID": 2744177, "TYPE": "Direct", "Gene_A": "GO:0032787", "BEND_MAP_ID": 2744177, "selected": false}, "selected": false}, {"data": {"id": "2744081", "source": "2741960", "target": "2742026", "SCORE": 0.4124151309434901, "shared_interaction": "Direct", "source_original": "49", "Name_B": "R-SCE-70171", "Name_A": "GO:0032787", "target_original": "71", "Gene_B": "R-SCE-70171", "interaction": "Direct", "SUID": 2744081, "TYPE": "Direct", "Gene_A": "GO:0032787", "BEND_MAP_ID": 2744081, "selected": false}, "selected": false}, {"data": {"id": "2744060", "source": "2741960", "target": "2742023", "SCORE": 0.39073589839637607, "shared_interaction": "Direct", "source_original": "49", "Name_B": "R-SCE-70326", "Name_A": "GO:0032787", "target_original": "70", "Gene_B": "R-SCE-70326", "interaction": "Direct", "SUID": 2744060, "TYPE": "Direct", "Gene_A": "GO:0032787", "BEND_MAP_ID": 2744060, "selected": false}, "selected": false}, {"data": {"id": "2743964", "source": "2741960", "target": "2742017", "SCORE": 0.5693160813325551, "shared_interaction": "Direct", "source_original": "49", "Name_B": "sce00010", "Name_A": "GO:0032787", "target_original": "68", "Gene_B": "sce00010", "interaction": "Direct", "SUID": 2743964, "TYPE": "Direct", "Gene_A": "GO:0032787", "BEND_MAP_ID": 2743964, "selected": false}, "selected": false}, {"data": {"id": "2743397", "source": "2741960", "target": "2741930", "SCORE": 0.3980883492655674, "shared_interaction": "Direct", "source_original": "49", "Name_B": "sce01200", "Name_A": "GO:0032787", "target_original": "39", "Gene_B": "sce01200", "interaction": "Direct", "SUID": 2743397, "TYPE": "Direct", "Gene_A": "GO:0032787", "BEND_MAP_ID": 2743397, "selected": false}, "selected": false}, {"data": {"id": "2743277", "source": "2741960", "target": "2742164", "SCORE": 0.344133676709306, "shared_interaction": "Direct", "source_original": "49", "Name_B": "GO:0044281", "Name_A": "GO:0032787", "target_original": "117", "Gene_B": "GO:0044281", "interaction": "Direct", "SUID": 2743277, "TYPE": "Direct", "Gene_A": "GO:0032787", "BEND_MAP_ID": 2743277, "selected": false}, "selected": false}, {"data": {"id": "2743223", "source": "2741960", "target": "2741882", "SCORE": 0.31518544436892326, "shared_interaction": "Direct", "source_original": "49", "Name_B": "R-SCE-71387", "Name_A": "GO:0032787", "target_original": "23", "Gene_B": "R-SCE-71387", "interaction": "Direct", "SUID": 2743223, "TYPE": "Direct", "Gene_A": "GO:0032787", "BEND_MAP_ID": 2743223, "selected": false}, "selected": false}, {"data": {"id": "2743091", "source": "2741960", "target": "2741927", "SCORE": 0.3463696369649529, "shared_interaction": "Direct", "source_original": "49", "Name_B": "sce01110", "Name_A": "GO:0032787", "target_original": "38", "Gene_B": "sce01110", "interaction": "Direct", "SUID": 2743091, "TYPE": "Direct", "Gene_A": "GO:0032787", "BEND_MAP_ID": 2743091, "selected": false}, "selected": false}, {"data": {"id": "2744141", "source": "2741957", "target": "2742041", "SCORE": 0.37421665174792307, "shared_interaction": "Direct", "source_original": "48", "Name_B": "GO:0046031", "Name_A": "GO:0016310", "target_original": "76", "Gene_B": "GO:0046031", "interaction": "Direct", "SUID": 2744141, "TYPE": "Direct", "Gene_A": "GO:0016310", "BEND_MAP_ID": 2744141, "selected": false}, "selected": false}, {"data": {"id": "2744072", "source": "2741957", "target": "2742026", "SCORE": 0.37421665174792307, "shared_interaction": "Direct", "source_original": "48", "Name_B": "R-SCE-70171", "Name_A": "GO:0016310", "target_original": "71", "Gene_B": "R-SCE-70171", "interaction": "Direct", "SUID": 2744072, "TYPE": "Direct", "Gene_A": "GO:0016310", "BEND_MAP_ID": 2744072, "selected": false}, "selected": false}, {"data": {"id": "2744051", "source": "2741957", "target": "2742023", "SCORE": 0.3571947146820062, "shared_interaction": "Direct", "source_original": "48", "Name_B": "R-SCE-70326", "Name_A": "GO:0016310", "target_original": "70", "Gene_B": "R-SCE-70326", "interaction": "Direct", "SUID": 2744051, "TYPE": "Direct", "Gene_A": "GO:0016310", "BEND_MAP_ID": 2744051, "selected": false}, "selected": false}, {"data": {"id": "2744030", "source": "2741957", "target": "2742020", "SCORE": 0.3206997084572391, "shared_interaction": "Direct", "source_original": "48", "Name_B": "GO:0046835", "Name_A": "GO:0016310", "target_original": "69", "Gene_B": "GO:0046835", "interaction": "Direct", "SUID": 2744030, "TYPE": "Direct", "Gene_A": "GO:0016310", "BEND_MAP_ID": 2744030, "selected": false}, "selected": false}, {"data": {"id": "2743955", "source": "2741957", "target": "2742017", "SCORE": 0.31120666487719, "shared_interaction": "Direct", "source_original": "48", "Name_B": "sce00010", "Name_A": "GO:0016310", "target_original": "68", "Gene_B": "sce00010", "interaction": "Direct", "SUID": 2743955, "TYPE": "Direct", "Gene_A": "GO:0016310", "BEND_MAP_ID": 2743955, "selected": false}, "selected": false}, {"data": {"id": "2743388", "source": "2741957", "target": "2741930", "SCORE": 0.3092126935342147, "shared_interaction": "Direct", "source_original": "48", "Name_B": "sce01200", "Name_A": "GO:0016310", "target_original": "39", "Gene_B": "sce01200", "interaction": "Direct", "SUID": 2743388, "TYPE": "Direct", "Gene_A": "GO:0016310", "BEND_MAP_ID": 2743388, "selected": false}, "selected": false}, {"data": {"id": "2743259", "source": "2741954", "target": "2741957", "SCORE": 0.5235173824137759, "shared_interaction": "Direct", "source_original": "47", "Name_B": "GO:0016310", "Name_A": "GO:0006796", "target_original": "48", "Gene_B": "GO:0016310", "interaction": "Direct", "SUID": 2743259, "TYPE": "Direct", "Gene_A": "GO:0006796", "BEND_MAP_ID": 2743259, "selected": false}, "selected": false}, {"data": {"id": "2743256", "source": "2741954", "target": "2742164", "SCORE": 0.47595348614692107, "shared_interaction": "Direct", "source_original": "47", "Name_B": "GO:0044281", "Name_A": "GO:0006796", "target_original": "117", "Gene_B": "GO:0044281", "interaction": "Direct", "SUID": 2743256, "TYPE": "Direct", "Gene_A": "GO:0006796", "BEND_MAP_ID": 2743256, "selected": false}, "selected": false}, {"data": {"id": "2742869", "source": "2741954", "target": "2741867", "SCORE": 0.5235173824137759, "shared_interaction": "Direct", "source_original": "47", "Name_B": "GO:0009150", "Name_A": "GO:0006796", "target_original": "18", "Gene_B": "GO:0009150", "interaction": "Direct", "SUID": 2742869, "TYPE": "Direct", "Gene_A": "GO:0006796", "BEND_MAP_ID": 2742869, "selected": false}, "selected": false}, {"data": {"id": "2742812", "source": "2741954", "target": "2741864", "SCORE": 0.3504579848675916, "shared_interaction": "Direct", "source_original": "47", "Name_B": "GO:0009142", "Name_A": "GO:0006796", "target_original": "17", "Gene_B": "GO:0009142", "interaction": "Direct", "SUID": 2742812, "TYPE": "Direct", "Gene_A": "GO:0006796", "BEND_MAP_ID": 2742812, "selected": false}, "selected": false}, {"data": {"id": "2742803", "source": "2741954", "target": "2741870", "SCORE": 0.31888222761808793, "shared_interaction": "Direct", "source_original": "47", "Name_B": "R-SCE-1430728", "Name_A": "GO:0006796", "target_original": "19", "Gene_B": "R-SCE-1430728", "interaction": "Direct", "SUID": 2742803, "TYPE": "Direct", "Gene_A": "GO:0006796", "BEND_MAP_ID": 2742803, "selected": false}, "selected": false}, {"data": {"id": "2742791", "source": "2741954", "target": "2741861", "SCORE": 0.3504579848675916, "shared_interaction": "Direct", "source_original": "47", "Name_B": "GO:0009199", "Name_A": "GO:0006796", "target_original": "16", "Gene_B": "GO:0009199", "interaction": "Direct", "SUID": 2742791, "TYPE": "Direct", "Gene_A": "GO:0006796", "BEND_MAP_ID": 2742791, "selected": false}, "selected": false}, {"data": {"id": "2742734", "source": "2741954", "target": "2741858", "SCORE": 0.3504579848675916, "shared_interaction": "Direct", "source_original": "47", "Name_B": "GO:0009201", "Name_A": "GO:0006796", "target_original": "15", "Gene_B": "GO:0009201", "interaction": "Direct", "SUID": 2742734, "TYPE": "Direct", "Gene_A": "GO:0006796", "BEND_MAP_ID": 2742734, "selected": false}, "selected": false}, {"data": {"id": "2742716", "source": "2741954", "target": "2741855", "SCORE": 0.3504579848675916, "shared_interaction": "Direct", "source_original": "47", "Name_B": "GO:0009144", "Name_A": "GO:0006796", "target_original": "14", "Gene_B": "GO:0009144", "interaction": "Direct", "SUID": 2742716, "TYPE": "Direct", "Gene_A": "GO:0006796", "BEND_MAP_ID": 2742716, "selected": false}, "selected": false}, {"data": {"id": "2742683", "source": "2741954", "target": "2741852", "SCORE": 0.3504579848675916, "shared_interaction": "Direct", "source_original": "47", "Name_B": "GO:0009205", "Name_A": "GO:0006796", "target_original": "13", "Gene_B": "GO:0009205", "interaction": "Direct", "SUID": 2742683, "TYPE": "Direct", "Gene_A": "GO:0006796", "BEND_MAP_ID": 2742683, "selected": false}, "selected": false}, {"data": {"id": "2742653", "source": "2741954", "target": "2741849", "SCORE": 0.3504579848675916, "shared_interaction": "Direct", "source_original": "47", "Name_B": "GO:0009206", "Name_A": "GO:0006796", "target_original": "12", "Gene_B": "GO:0009206", "interaction": "Direct", "SUID": 2742653, "TYPE": "Direct", "Gene_A": "GO:0006796", "BEND_MAP_ID": 2742653, "selected": false}, "selected": false}, {"data": {"id": "2742626", "source": "2741954", "target": "2741846", "SCORE": 0.3504579848675916, "shared_interaction": "Direct", "source_original": "47", "Name_B": "GO:0009145", "Name_A": "GO:0006796", "target_original": "11", "Gene_B": "GO:0009145", "interaction": "Direct", "SUID": 2742626, "TYPE": "Direct", "Gene_A": "GO:0006796", "BEND_MAP_ID": 2742626, "selected": false}, "selected": false}, {"data": {"id": "2742566", "source": "2741954", "target": "2741843", "SCORE": 0.3504579848675916, "shared_interaction": "Direct", "source_original": "47", "Name_B": "GO:0015986", "Name_A": "GO:0006796", "target_original": "10", "Gene_B": "GO:0015986", "interaction": "Direct", "SUID": 2742566, "TYPE": "Direct", "Gene_A": "GO:0006796", "BEND_MAP_ID": 2742566, "selected": false}, "selected": false}, {"data": {"id": "2743811", "source": "2741951", "target": "2741948", "SCORE": 0.7933786579972849, "shared_interaction": "Direct", "source_original": "46", "Name_B": "sce00020", "Name_A": "R-SCE-71406", "target_original": "45", "Gene_B": "sce00020", "interaction": "Direct", "SUID": 2743811, "TYPE": "Direct", "Gene_A": "R-SCE-71406", "BEND_MAP_ID": 2743811, "selected": false}, "selected": false}, {"data": {"id": "2743697", "source": "2741951", "target": "2741942", "SCORE": 0.658107116657361, "shared_interaction": "Direct", "source_original": "46", "Name_B": "WP559", "Name_A": "R-SCE-71406", "target_original": "43", "Gene_B": "WP559", "interaction": "Direct", "SUID": 2743697, "TYPE": "Direct", "Gene_A": "R-SCE-71406", "BEND_MAP_ID": 2743697, "selected": false}, "selected": false}, {"data": {"id": "2743685", "source": "2741951", "target": "2741939", "SCORE": 0.7412548584136671, "shared_interaction": "Direct", "source_original": "46", "Name_B": "WP490", "Name_A": "R-SCE-71406", "target_original": "42", "Gene_B": "WP490", "interaction": "Direct", "SUID": 2743685, "TYPE": "Direct", "Gene_A": "R-SCE-71406", "BEND_MAP_ID": 2743685, "selected": false}, "selected": false}, {"data": {"id": "2743541", "source": "2741951", "target": "2741933", "SCORE": 0.7933786579972849, "shared_interaction": "Direct", "source_original": "46", "Name_B": "WP296", "Name_A": "R-SCE-71406", "target_original": "40", "Gene_B": "WP296", "interaction": "Direct", "SUID": 2743541, "TYPE": "Direct", "Gene_A": "R-SCE-71406", "BEND_MAP_ID": 2743541, "selected": false}, "selected": false}, {"data": {"id": "2743376", "source": "2741951", "target": "2741930", "SCORE": 0.43680939226745896, "shared_interaction": "Direct", "source_original": "46", "Name_B": "sce01200", "Name_A": "R-SCE-71406", "target_original": "39", "Gene_B": "sce01200", "interaction": "Direct", "SUID": 2743376, "TYPE": "Direct", "Gene_A": "R-SCE-71406", "BEND_MAP_ID": 2743376, "selected": false}, "selected": false}, {"data": {"id": "2743037", "source": "2741951", "target": "2742122", "SCORE": 0.4564866319778691, "shared_interaction": "Direct", "source_original": "46", "Name_B": "WP112", "Name_A": "R-SCE-71406", "target_original": "103", "Gene_B": "WP112", "interaction": "Direct", "SUID": 2743037, "TYPE": "Direct", "Gene_A": "R-SCE-71406", "BEND_MAP_ID": 2743037, "selected": false}, "selected": false}, {"data": {"id": "2743373", "source": "2741948", "target": "2741930", "SCORE": 0.5719057958927636, "shared_interaction": "Direct", "source_original": "45", "Name_B": "sce01200", "Name_A": "sce00020", "target_original": "39", "Gene_B": "sce01200", "interaction": "Direct", "SUID": 2743373, "TYPE": "Direct", "Gene_A": "sce00020", "BEND_MAP_ID": 2743373, "selected": false}, "selected": false}, {"data": {"id": "2743781", "source": "2741945", "target": "2741951", "SCORE": 0.9208828522926463, "shared_interaction": "Direct", "source_original": "44", "Name_B": "R-SCE-71406", "Name_A": "R-SCE-71403", "target_original": "46", "Gene_B": "R-SCE-71406", "interaction": "Direct", "SUID": 2743781, "TYPE": "Direct", "Gene_A": "R-SCE-71403", "BEND_MAP_ID": 2743781, "selected": false}, "selected": false}, {"data": {"id": "2743778", "source": "2741945", "target": "2741948", "SCORE": 0.8528111181312159, "shared_interaction": "Direct", "source_original": "44", "Name_B": "sce00020", "Name_A": "R-SCE-71403", "target_original": "45", "Gene_B": "sce00020", "interaction": "Direct", "SUID": 2743778, "TYPE": "Direct", "Gene_A": "R-SCE-71403", "BEND_MAP_ID": 2743778, "selected": false}, "selected": false}, {"data": {"id": "2743691", "source": "2741945", "target": "2741942", "SCORE": 0.7207351178611695, "shared_interaction": "Direct", "source_original": "44", "Name_B": "WP559", "Name_A": "R-SCE-71403", "target_original": "43", "Gene_B": "WP559", "interaction": "Direct", "SUID": 2743691, "TYPE": "Direct", "Gene_A": "R-SCE-71403", "BEND_MAP_ID": 2743691, "selected": false}, "selected": false}, {"data": {"id": "2743679", "source": "2741945", "target": "2741939", "SCORE": 0.7936226749349893, "shared_interaction": "Direct", "source_original": "44", "Name_B": "WP490", "Name_A": "R-SCE-71403", "target_original": "42", "Gene_B": "WP490", "interaction": "Direct", "SUID": 2743679, "TYPE": "Direct", "Gene_A": "R-SCE-71403", "BEND_MAP_ID": 2743679, "selected": false}, "selected": false}, {"data": {"id": "2743535", "source": "2741945", "target": "2741933", "SCORE": 0.8528111181312159, "shared_interaction": "Direct", "source_original": "44", "Name_B": "WP296", "Name_A": "R-SCE-71403", "target_original": "40", "Gene_B": "WP296", "interaction": "Direct", "SUID": 2743535, "TYPE": "Direct", "Gene_A": "R-SCE-71403", "BEND_MAP_ID": 2743535, "selected": false}, "selected": false}, {"data": {"id": "2743370", "source": "2741945", "target": "2741930", "SCORE": 0.45881722351484877, "shared_interaction": "Direct", "source_original": "44", "Name_B": "sce01200", "Name_A": "R-SCE-71403", "target_original": "39", "Gene_B": "sce01200", "interaction": "Direct", "SUID": 2743370, "TYPE": "Direct", "Gene_A": "R-SCE-71403", "BEND_MAP_ID": 2743370, "selected": false}, "selected": false}, {"data": {"id": "2743031", "source": "2741945", "target": "2742122", "SCORE": 0.47991071428796866, "shared_interaction": "Direct", "source_original": "44", "Name_B": "WP112", "Name_A": "R-SCE-71403", "target_original": "103", "Gene_B": "WP112", "interaction": "Direct", "SUID": 2743031, "TYPE": "Direct", "Gene_A": "R-SCE-71403", "BEND_MAP_ID": 2743031, "selected": false}, "selected": false}, {"data": {"id": "2743694", "source": "2741942", "target": "2741948", "SCORE": 0.7629704984759316, "shared_interaction": "Direct", "source_original": "43", "Name_B": "sce00020", "Name_A": "WP559", "target_original": "45", "Gene_B": "sce00020", "interaction": "Direct", "SUID": 2743694, "TYPE": "Direct", "Gene_A": "WP559", "BEND_MAP_ID": 2743694, "selected": false}, "selected": false}, {"data": {"id": "2743367", "source": "2741942", "target": "2741930", "SCORE": 0.39614957423437086, "shared_interaction": "Direct", "source_original": "43", "Name_B": "sce01200", "Name_A": "WP559", "target_original": "39", "Gene_B": "sce01200", "interaction": "Direct", "SUID": 2743367, "TYPE": "Direct", "Gene_A": "WP559", "BEND_MAP_ID": 2743367, "selected": false}, "selected": false}, {"data": {"id": "2743682", "source": "2741939", "target": "2741948", "SCORE": 0.9389572963063721, "shared_interaction": "Direct", "source_original": "42", "Name_B": "sce00020", "Name_A": "WP490", "target_original": "45", "Gene_B": "sce00020", "interaction": "Direct", "SUID": 2743682, "TYPE": "Direct", "Gene_A": "WP490", "BEND_MAP_ID": 2743682, "selected": false}, "selected": false}, {"data": {"id": "2743676", "source": "2741939", "target": "2741942", "SCORE": 0.706179066836963, "shared_interaction": "Direct", "source_original": "42", "Name_B": "WP559", "Name_A": "WP490", "target_original": "43", "Gene_B": "WP559", "interaction": "Direct", "SUID": 2743676, "TYPE": "Direct", "Gene_A": "WP490", "BEND_MAP_ID": 2743676, "selected": false}, "selected": false}, {"data": {"id": "2743364", "source": "2741939", "target": "2741930", "SCORE": 0.6230993206096381, "shared_interaction": "Direct", "source_original": "42", "Name_B": "sce01200", "Name_A": "WP490", "target_original": "39", "Gene_B": "sce01200", "interaction": "Direct", "SUID": 2743364, "TYPE": "Direct", "Gene_A": "WP490", "BEND_MAP_ID": 2743364, "selected": false}, "selected": false}, {"data": {"id": "2743085", "source": "2741939", "target": "2741927", "SCORE": 0.3167155425234529, "shared_interaction": "Direct", "source_original": "42", "Name_B": "sce01110", "Name_A": "WP490", "target_original": "38", "Gene_B": "sce01110", "interaction": "Direct", "SUID": 2743085, "TYPE": "Direct", "Gene_A": "WP490", "BEND_MAP_ID": 2743085, "selected": false}, "selected": false}, {"data": {"id": "2743625", "source": "2741936", "target": "2741951", "SCORE": 0.7412548584136671, "shared_interaction": "Direct", "source_original": "41", "Name_B": "R-SCE-71406", "Name_A": "GO:0006099", "target_original": "46", "Gene_B": "R-SCE-71406", "interaction": "Direct", "SUID": 2743625, "TYPE": "Direct", "Gene_A": "GO:0006099", "BEND_MAP_ID": 2743625, "selected": false}, "selected": false}, {"data": {"id": "2743622", "source": "2741936", "target": "2741948", "SCORE": 0.9389572963063721, "shared_interaction": "Direct", "source_original": "41", "Name_B": "sce00020", "Name_A": "GO:0006099", "target_original": "45", "Gene_B": "sce00020", "interaction": "Direct", "SUID": 2743622, "TYPE": "Direct", "Gene_A": "GO:0006099", "BEND_MAP_ID": 2743622, "selected": false}, "selected": false}, {"data": {"id": "2743619", "source": "2741936", "target": "2741945", "SCORE": 0.7936226749349893, "shared_interaction": "Direct", "source_original": "41", "Name_B": "R-SCE-71403", "Name_A": "GO:0006099", "target_original": "44", "Gene_B": "R-SCE-71403", "interaction": "Direct", "SUID": 2743619, "TYPE": "Direct", "Gene_A": "GO:0006099", "BEND_MAP_ID": 2743619, "selected": false}, "selected": false}, {"data": {"id": "2743616", "source": "2741936", "target": "2741942", "SCORE": 0.706179066836963, "shared_interaction": "Direct", "source_original": "41", "Name_B": "WP559", "Name_A": "GO:0006099", "target_original": "43", "Gene_B": "WP559", "interaction": "Direct", "SUID": 2743616, "TYPE": "Direct", "Gene_A": "GO:0006099", "BEND_MAP_ID": 2743616, "selected": false}, "selected": false}, {"data": {"id": "2743613", "source": "2741936", "target": "2741939", "SCORE": 1.0, "shared_interaction": "Direct", "source_original": "41", "Name_B": "WP490", "Name_A": "GO:0006099", "target_original": "42", "Gene_B": "WP490", "interaction": "Direct", "SUID": 2743613, "TYPE": "Direct", "Gene_A": "GO:0006099", "BEND_MAP_ID": 2743613, "selected": false}, "selected": false}, {"data": {"id": "2743526", "source": "2741936", "target": "2741933", "SCORE": 0.9389572963063721, "shared_interaction": "Direct", "source_original": "41", "Name_B": "WP296", "Name_A": "GO:0006099", "target_original": "40", "Gene_B": "WP296", "interaction": "Direct", "SUID": 2743526, "TYPE": "Direct", "Gene_A": "GO:0006099", "BEND_MAP_ID": 2743526, "selected": false}, "selected": false}, {"data": {"id": "2743361", "source": "2741936", "target": "2741930", "SCORE": 0.6230993206096381, "shared_interaction": "Direct", "source_original": "41", "Name_B": "sce01200", "Name_A": "GO:0006099", "target_original": "39", "Gene_B": "sce01200", "interaction": "Direct", "SUID": 2743361, "TYPE": "Direct", "Gene_A": "GO:0006099", "BEND_MAP_ID": 2743361, "selected": false}, "selected": false}, {"data": {"id": "2743082", "source": "2741936", "target": "2741927", "SCORE": 0.3167155425234529, "shared_interaction": "Direct", "source_original": "41", "Name_B": "sce01110", "Name_A": "GO:0006099", "target_original": "38", "Gene_B": "sce01110", "interaction": "Direct", "SUID": 2743082, "TYPE": "Direct", "Gene_A": "GO:0006099", "BEND_MAP_ID": 2743082, "selected": false}, "selected": false}, {"data": {"id": "2743022", "source": "2741936", "target": "2742122", "SCORE": 0.6485671191567266, "shared_interaction": "Direct", "source_original": "41", "Name_B": "WP112", "Name_A": "GO:0006099", "target_original": "103", "Gene_B": "WP112", "interaction": "Direct", "SUID": 2743022, "TYPE": "Direct", "Gene_A": "GO:0006099", "BEND_MAP_ID": 2743022, "selected": false}, "selected": false}, {"data": {"id": "2742473", "source": "2741936", "target": "2741831", "SCORE": 0.32575444398728254, "shared_interaction": "Direct", "source_original": "41", "Name_B": "R-SCE-1428517", "Name_A": "GO:0006099", "target_original": "6", "Gene_B": "R-SCE-1428517", "interaction": "Direct", "SUID": 2742473, "TYPE": "Direct", "Gene_A": "GO:0006099", "BEND_MAP_ID": 2742473, "selected": false}, "selected": false}, {"data": {"id": "2742416", "source": "2741936", "target": "2741828", "SCORE": 0.36047575480480853, "shared_interaction": "Direct", "source_original": "41", "Name_B": "GO:0045333", "Name_A": "GO:0006099", "target_original": "5", "Gene_B": "GO:0045333", "interaction": "Direct", "SUID": 2742416, "TYPE": "Direct", "Gene_A": "GO:0006099", "BEND_MAP_ID": 2742416, "selected": false}, "selected": false}, {"data": {"id": "2742389", "source": "2741936", "target": "2741825", "SCORE": 0.36047575480480853, "shared_interaction": "Direct", "source_original": "41", "Name_B": "GO:0009060", "Name_A": "GO:0006099", "target_original": "4", "Gene_B": "GO:0009060", "interaction": "Direct", "SUID": 2742389, "TYPE": "Direct", "Gene_A": "GO:0006099", "BEND_MAP_ID": 2742389, "selected": false}, "selected": false}, {"data": {"id": "2743538", "source": "2741933", "target": "2741948", "SCORE": 1.0, "shared_interaction": "Direct", "source_original": "40", "Name_B": "sce00020", "Name_A": "WP296", "target_original": "45", "Gene_B": "sce00020", "interaction": "Direct", "SUID": 2743538, "TYPE": "Direct", "Gene_A": "WP296", "BEND_MAP_ID": 2743538, "selected": false}, "selected": false}, {"data": {"id": "2743532", "source": "2741933", "target": "2741942", "SCORE": 0.7629704984759316, "shared_interaction": "Direct", "source_original": "40", "Name_B": "WP559", "Name_A": "WP296", "target_original": "43", "Gene_B": "WP559", "interaction": "Direct", "SUID": 2743532, "TYPE": "Direct", "Gene_A": "WP296", "BEND_MAP_ID": 2743532, "selected": false}, "selected": false}, {"data": {"id": "2743529", "source": "2741933", "target": "2741939", "SCORE": 0.9389572963063721, "shared_interaction": "Direct", "source_original": "40", "Name_B": "WP490", "Name_A": "WP296", "target_original": "42", "Gene_B": "WP490", "interaction": "Direct", "SUID": 2743529, "TYPE": "Direct", "Gene_A": "WP296", "BEND_MAP_ID": 2743529, "selected": false}, "selected": false}, {"data": {"id": "2743358", "source": "2741933", "target": "2741930", "SCORE": 0.5719057958927636, "shared_interaction": "Direct", "source_original": "40", "Name_B": "sce01200", "Name_A": "WP296", "target_original": "39", "Gene_B": "sce01200", "interaction": "Direct", "SUID": 2743358, "TYPE": "Direct", "Gene_A": "WP296", "BEND_MAP_ID": 2743358, "selected": false}, "selected": false}, {"data": {"id": "2743079", "source": "2741927", "target": "2741930", "SCORE": 0.5873450330067312, "shared_interaction": "Direct", "source_original": "38", "Name_B": "sce01200", "Name_A": "sce01110", "target_original": "39", "Gene_B": "sce01200", "interaction": "Direct", "SUID": 2743079, "TYPE": "Direct", "Gene_A": "sce01110", "BEND_MAP_ID": 2743079, "selected": false}, "selected": false}, {"data": {"id": "2743466", "source": "2741924", "target": "2741909", "SCORE": 0.9069117059537232, "shared_interaction": "Direct", "source_original": "37", "Name_B": "WP398", "Name_A": "GO:0046351", "target_original": "32", "Gene_B": "WP398", "interaction": "Direct", "SUID": 2743466, "TYPE": "Direct", "Gene_A": "GO:0046351", "BEND_MAP_ID": 2743466, "selected": false}, "selected": false}, {"data": {"id": "2743163", "source": "2741924", "target": "2741906", "SCORE": 0.7956140350898077, "shared_interaction": "Direct", "source_original": "37", "Name_B": "GO:0070413", "Name_A": "GO:0046351", "target_original": "31", "Gene_B": "GO:0070413", "interaction": "Direct", "SUID": 2743163, "TYPE": "Direct", "Gene_A": "GO:0046351", "BEND_MAP_ID": 2743163, "selected": false}, "selected": false}, {"data": {"id": "2742941", "source": "2741924", "target": "2742125", "SCORE": 0.4833702882510058, "shared_interaction": "Direct", "source_original": "37", "Name_B": "sce00500", "Name_A": "GO:0046351", "target_original": "104", "Gene_B": "sce00500", "interaction": "Direct", "SUID": 2742941, "TYPE": "Direct", "Gene_A": "GO:0046351", "BEND_MAP_ID": 2742941, "selected": false}, "selected": false}, {"data": {"id": "2743658", "source": "2741921", "target": "2741924", "SCORE": 1.0, "shared_interaction": "Direct", "source_original": "36", "Name_B": "GO:0046351", "Name_A": "GO:0009312", "target_original": "37", "Gene_B": "GO:0046351", "interaction": "Direct", "SUID": 2743658, "TYPE": "Direct", "Gene_A": "GO:0009312", "BEND_MAP_ID": 2743658, "selected": false}, "selected": false}, {"data": {"id": "2743463", "source": "2741921", "target": "2741909", "SCORE": 0.9069117059537232, "shared_interaction": "Direct", "source_original": "36", "Name_B": "WP398", "Name_A": "GO:0009312", "target_original": "32", "Gene_B": "WP398", "interaction": "Direct", "SUID": 2743463, "TYPE": "Direct", "Gene_A": "GO:0009312", "BEND_MAP_ID": 2743463, "selected": false}, "selected": false}, {"data": {"id": "2743301", "source": "2741921", "target": "2741891", "SCORE": 0.5753709971387387, "shared_interaction": "Direct", "source_original": "36", "Name_B": "GO:0034637", "Name_A": "GO:0009312", "target_original": "26", "Gene_B": "GO:0034637", "interaction": "Direct", "SUID": 2743301, "TYPE": "Direct", "Gene_A": "GO:0009312", "BEND_MAP_ID": 2743301, "selected": false}, "selected": false}, {"data": {"id": "2743160", "source": "2741921", "target": "2741906", "SCORE": 0.7956140350898077, "shared_interaction": "Direct", "source_original": "36", "Name_B": "GO:0070413", "Name_A": "GO:0009312", "target_original": "31", "Gene_B": "GO:0070413", "interaction": "Direct", "SUID": 2743160, "TYPE": "Direct", "Gene_A": "GO:0009312", "BEND_MAP_ID": 2743160, "selected": false}, "selected": false}, {"data": {"id": "2743130", "source": "2741921", "target": "2741879", "SCORE": 0.4833702882510058, "shared_interaction": "Direct", "source_original": "36", "Name_B": "GO:0016051", "Name_A": "GO:0009312", "target_original": "22", "Gene_B": "GO:0016051", "interaction": "Direct", "SUID": 2743130, "TYPE": "Direct", "Gene_A": "GO:0009312", "BEND_MAP_ID": 2743130, "selected": false}, "selected": false}, {"data": {"id": "2742938", "source": "2741921", "target": "2742125", "SCORE": 0.4833702882510058, "shared_interaction": "Direct", "source_original": "36", "Name_B": "sce00500", "Name_A": "GO:0009312", "target_original": "104", "Gene_B": "sce00500", "interaction": "Direct", "SUID": 2742938, "TYPE": "Direct", "Gene_A": "GO:0009312", "BEND_MAP_ID": 2742938, "selected": false}, "selected": false}, {"data": {"id": "2743655", "source": "2741918", "target": "2742083", "SCORE": 0.3202042304944703, "shared_interaction": "Direct", "source_original": "35", "Name_B": "GO:0071462", "Name_A": "GO:0009311", "target_original": "90", "Gene_B": "GO:0071462", "interaction": "Direct", "SUID": 2743655, "TYPE": "Direct", "Gene_A": "GO:0009311", "BEND_MAP_ID": 2743655, "selected": false}, "selected": false}, {"data": {"id": "2743652", "source": "2741918", "target": "2742080", "SCORE": 0.3202042304944703, "shared_interaction": "Direct", "source_original": "35", "Name_B": "GO:0009415", "Name_A": "GO:0009311", "target_original": "89", "Gene_B": "GO:0009415", "interaction": "Direct", "SUID": 2743652, "TYPE": "Direct", "Gene_A": "GO:0009311", "BEND_MAP_ID": 2743652, "selected": false}, "selected": false}, {"data": {"id": "2743649", "source": "2741918", "target": "2742077", "SCORE": 0.3202042304944703, "shared_interaction": "Direct", "source_original": "35", "Name_B": "GO:0042631", "Name_A": "GO:0009311", "target_original": "88", "Gene_B": "GO:0042631", "interaction": "Direct", "SUID": 2743649, "TYPE": "Direct", "Gene_A": "GO:0009311", "BEND_MAP_ID": 2743649, "selected": false}, "selected": false}, {"data": {"id": "2743646", "source": "2741918", "target": "2742074", "SCORE": 0.3202042304944703, "shared_interaction": "Direct", "source_original": "35", "Name_B": "GO:0009414", "Name_A": "GO:0009311", "target_original": "87", "Gene_B": "GO:0009414", "interaction": "Direct", "SUID": 2743646, "TYPE": "Direct", "Gene_A": "GO:0009311", "BEND_MAP_ID": 2743646, "selected": false}, "selected": false}, {"data": {"id": "2743643", "source": "2741918", "target": "2742047", "SCORE": 0.328530259370093, "shared_interaction": "Direct", "source_original": "35", "Name_B": "GO:0034605", "Name_A": "GO:0009311", "target_original": "78", "Gene_B": "GO:0034605", "interaction": "Direct", "SUID": 2743643, "TYPE": "Direct", "Gene_A": "GO:0009311", "BEND_MAP_ID": 2743643, "selected": false}, "selected": false}, {"data": {"id": "2743640", "source": "2741918", "target": "2742071", "SCORE": 0.3202042304944703, "shared_interaction": "Direct", "source_original": "35", "Name_B": "GO:0071465", "Name_A": "GO:0009311", "target_original": "86", "Gene_B": "GO:0071465", "interaction": "Direct", "SUID": 2743640, "TYPE": "Direct", "Gene_A": "GO:0009311", "BEND_MAP_ID": 2743640, "selected": false}, "selected": false}, {"data": {"id": "2743634", "source": "2741918", "target": "2742128", "SCORE": 0.44842746026812763, "shared_interaction": "Direct", "source_original": "35", "Name_B": "WP70", "Name_A": "GO:0009311", "target_original": "105", "Gene_B": "WP70", "interaction": "Direct", "SUID": 2743634, "TYPE": "Direct", "Gene_A": "GO:0009311", "BEND_MAP_ID": 2743634, "selected": false}, "selected": false}, {"data": {"id": "2743631", "source": "2741918", "target": "2741924", "SCORE": 0.706179066836963, "shared_interaction": "Direct", "source_original": "35", "Name_B": "GO:0046351", "Name_A": "GO:0009311", "target_original": "37", "Gene_B": "GO:0046351", "interaction": "Direct", "SUID": 2743631, "TYPE": "Direct", "Gene_A": "GO:0009311", "BEND_MAP_ID": 2743631, "selected": false}, "selected": false}, {"data": {"id": "2743628", "source": "2741918", "target": "2741921", "SCORE": 0.706179066836963, "shared_interaction": "Direct", "source_original": "35", "Name_B": "GO:0009312", "Name_A": "GO:0009311", "target_original": "36", "Gene_B": "GO:0009312", "interaction": "Direct", "SUID": 2743628, "TYPE": "Direct", "Gene_A": "GO:0009311", "BEND_MAP_ID": 2743628, "selected": false}, "selected": false}, {"data": {"id": "2743460", "source": "2741918", "target": "2741909", "SCORE": 0.7936226749349893, "shared_interaction": "Direct", "source_original": "35", "Name_B": "WP398", "Name_A": "GO:0009311", "target_original": "32", "Gene_B": "WP398", "interaction": "Direct", "SUID": 2743460, "TYPE": "Direct", "Gene_A": "GO:0009311", "BEND_MAP_ID": 2743460, "selected": false}, "selected": false}, {"data": {"id": "2743298", "source": "2741918", "target": "2741891", "SCORE": 0.4519991447536391, "shared_interaction": "Direct", "source_original": "35", "Name_B": "GO:0034637", "Name_A": "GO:0009311", "target_original": "26", "Gene_B": "GO:0034637", "interaction": "Direct", "SUID": 2743298, "TYPE": "Direct", "Gene_A": "GO:0009311", "BEND_MAP_ID": 2743298, "selected": false}, "selected": false}, {"data": {"id": "2743157", "source": "2741918", "target": "2741906", "SCORE": 0.706179066836963, "shared_interaction": "Direct", "source_original": "35", "Name_B": "GO:0070413", "Name_A": "GO:0009311", "target_original": "31", "Gene_B": "GO:0070413", "interaction": "Direct", "SUID": 2743157, "TYPE": "Direct", "Gene_A": "GO:0009311", "BEND_MAP_ID": 2743157, "selected": false}, "selected": false}, {"data": {"id": "2743127", "source": "2741918", "target": "2741879", "SCORE": 0.3870725291270728, "shared_interaction": "Direct", "source_original": "35", "Name_B": "GO:0016051", "Name_A": "GO:0009311", "target_original": "22", "Gene_B": "GO:0016051", "interaction": "Direct", "SUID": 2743127, "TYPE": "Direct", "Gene_A": "GO:0009311", "BEND_MAP_ID": 2743127, "selected": false}, "selected": false}, {"data": {"id": "2742935", "source": "2741918", "target": "2742125", "SCORE": 0.5621946636621948, "shared_interaction": "Direct", "source_original": "35", "Name_B": "sce00500", "Name_A": "GO:0009311", "target_original": "104", "Gene_B": "sce00500", "interaction": "Direct", "SUID": 2742935, "TYPE": "Direct", "Gene_A": "GO:0009311", "BEND_MAP_ID": 2742935, "selected": false}, "selected": false}, {"data": {"id": "2742854", "source": "2741918", "target": "2741873", "SCORE": 0.4407105136837532, "shared_interaction": "Direct", "source_original": "35", "Name_B": "GO:0044262", "Name_A": "GO:0009311", "target_original": "20", "Gene_B": "GO:0044262", "interaction": "Direct", "SUID": 2742854, "TYPE": "Direct", "Gene_A": "GO:0009311", "BEND_MAP_ID": 2742854, "selected": false}, "selected": false}, {"data": {"id": "2743595", "source": "2741915", "target": "2741924", "SCORE": 1.0, "shared_interaction": "Direct", "source_original": "34", "Name_B": "GO:0046351", "Name_A": "GO:0005992", "target_original": "37", "Gene_B": "GO:0046351", "interaction": "Direct", "SUID": 2743595, "TYPE": "Direct", "Gene_A": "GO:0005992", "BEND_MAP_ID": 2743595, "selected": false}, "selected": false}, {"data": {"id": "2743592", "source": "2741915", "target": "2741921", "SCORE": 1.0, "shared_interaction": "Direct", "source_original": "34", "Name_B": "GO:0009312", "Name_A": "GO:0005992", "target_original": "36", "Gene_B": "GO:0009312", "interaction": "Direct", "SUID": 2743592, "TYPE": "Direct", "Gene_A": "GO:0005992", "BEND_MAP_ID": 2743592, "selected": false}, "selected": false}, {"data": {"id": "2743589", "source": "2741915", "target": "2741918", "SCORE": 0.706179066836963, "shared_interaction": "Direct", "source_original": "34", "Name_B": "GO:0009311", "Name_A": "GO:0005992", "target_original": "35", "Gene_B": "GO:0009311", "interaction": "Direct", "SUID": 2743589, "TYPE": "Direct", "Gene_A": "GO:0005992", "BEND_MAP_ID": 2743589, "selected": false}, "selected": false}, {"data": {"id": "2743457", "source": "2741915", "target": "2741909", "SCORE": 0.9069117059537232, "shared_interaction": "Direct", "source_original": "34", "Name_B": "WP398", "Name_A": "GO:0005992", "target_original": "32", "Gene_B": "WP398", "interaction": "Direct", "SUID": 2743457, "TYPE": "Direct", "Gene_A": "GO:0005992", "BEND_MAP_ID": 2743457, "selected": false}, "selected": false}, {"data": {"id": "2743295", "source": "2741915", "target": "2741891", "SCORE": 0.5753709971387387, "shared_interaction": "Direct", "source_original": "34", "Name_B": "GO:0034637", "Name_A": "GO:0005992", "target_original": "26", "Gene_B": "GO:0034637", "interaction": "Direct", "SUID": 2743295, "TYPE": "Direct", "Gene_A": "GO:0005992", "BEND_MAP_ID": 2743295, "selected": false}, "selected": false}, {"data": {"id": "2743154", "source": "2741915", "target": "2741906", "SCORE": 0.7956140350898077, "shared_interaction": "Direct", "source_original": "34", "Name_B": "GO:0070413", "Name_A": "GO:0005992", "target_original": "31", "Gene_B": "GO:0070413", "interaction": "Direct", "SUID": 2743154, "TYPE": "Direct", "Gene_A": "GO:0005992", "BEND_MAP_ID": 2743154, "selected": false}, "selected": false}, {"data": {"id": "2743124", "source": "2741915", "target": "2741879", "SCORE": 0.4833702882510058, "shared_interaction": "Direct", "source_original": "34", "Name_B": "GO:0016051", "Name_A": "GO:0005992", "target_original": "22", "Gene_B": "GO:0016051", "interaction": "Direct", "SUID": 2743124, "TYPE": "Direct", "Gene_A": "GO:0005992", "BEND_MAP_ID": 2743124, "selected": false}, "selected": false}, {"data": {"id": "2742932", "source": "2741915", "target": "2742125", "SCORE": 0.4833702882510058, "shared_interaction": "Direct", "source_original": "34", "Name_B": "sce00500", "Name_A": "GO:0005992", "target_original": "104", "Gene_B": "sce00500", "interaction": "Direct", "SUID": 2742932, "TYPE": "Direct", "Gene_A": "GO:0005992", "BEND_MAP_ID": 2742932, "selected": false}, "selected": false}, {"data": {"id": "2743523", "source": "2741912", "target": "2742083", "SCORE": 0.3202042304944703, "shared_interaction": "Direct", "source_original": "33", "Name_B": "GO:0071462", "Name_A": "GO:0005984", "target_original": "90", "Gene_B": "GO:0071462", "interaction": "Direct", "SUID": 2743523, "TYPE": "Direct", "Gene_A": "GO:0005984", "BEND_MAP_ID": 2743523, "selected": false}, "selected": false}, {"data": {"id": "2743520", "source": "2741912", "target": "2742080", "SCORE": 0.3202042304944703, "shared_interaction": "Direct", "source_original": "33", "Name_B": "GO:0009415", "Name_A": "GO:0005984", "target_original": "89", "Gene_B": "GO:0009415", "interaction": "Direct", "SUID": 2743520, "TYPE": "Direct", "Gene_A": "GO:0005984", "BEND_MAP_ID": 2743520, "selected": false}, "selected": false}, {"data": {"id": "2743517", "source": "2741912", "target": "2742077", "SCORE": 0.3202042304944703, "shared_interaction": "Direct", "source_original": "33", "Name_B": "GO:0042631", "Name_A": "GO:0005984", "target_original": "88", "Gene_B": "GO:0042631", "interaction": "Direct", "SUID": 2743517, "TYPE": "Direct", "Gene_A": "GO:0005984", "BEND_MAP_ID": 2743517, "selected": false}, "selected": false}, {"data": {"id": "2743514", "source": "2741912", "target": "2742074", "SCORE": 0.3202042304944703, "shared_interaction": "Direct", "source_original": "33", "Name_B": "GO:0009414", "Name_A": "GO:0005984", "target_original": "87", "Gene_B": "GO:0009414", "interaction": "Direct", "SUID": 2743514, "TYPE": "Direct", "Gene_A": "GO:0005984", "BEND_MAP_ID": 2743514, "selected": false}, "selected": false}, {"data": {"id": "2743511", "source": "2741912", "target": "2742047", "SCORE": 0.328530259370093, "shared_interaction": "Direct", "source_original": "33", "Name_B": "GO:0034605", "Name_A": "GO:0005984", "target_original": "78", "Gene_B": "GO:0034605", "interaction": "Direct", "SUID": 2743511, "TYPE": "Direct", "Gene_A": "GO:0005984", "BEND_MAP_ID": 2743511, "selected": false}, "selected": false}, {"data": {"id": "2743508", "source": "2741912", "target": "2742071", "SCORE": 0.3202042304944703, "shared_interaction": "Direct", "source_original": "33", "Name_B": "GO:0071465", "Name_A": "GO:0005984", "target_original": "86", "Gene_B": "GO:0071465", "interaction": "Direct", "SUID": 2743508, "TYPE": "Direct", "Gene_A": "GO:0005984", "BEND_MAP_ID": 2743508, "selected": false}, "selected": false}, {"data": {"id": "2743505", "source": "2741912", "target": "2742155", "SCORE": 0.3202042304944703, "shared_interaction": "Direct", "source_original": "33", "Name_B": "GO:0009269", "Name_A": "GO:0005984", "target_original": "114", "Gene_B": "GO:0009269", "interaction": "Direct", "SUID": 2743505, "TYPE": "Direct", "Gene_A": "GO:0005984", "BEND_MAP_ID": 2743505, "selected": false}, "selected": false}, {"data": {"id": "2743502", "source": "2741912", "target": "2742128", "SCORE": 0.44842746026812763, "shared_interaction": "Direct", "source_original": "33", "Name_B": "WP70", "Name_A": "GO:0005984", "target_original": "105", "Gene_B": "WP70", "interaction": "Direct", "SUID": 2743502, "TYPE": "Direct", "Gene_A": "GO:0005984", "BEND_MAP_ID": 2743502, "selected": false}, "selected": false}, {"data": {"id": "2743499", "source": "2741912", "target": "2741924", "SCORE": 0.706179066836963, "shared_interaction": "Direct", "source_original": "33", "Name_B": "GO:0046351", "Name_A": "GO:0005984", "target_original": "37", "Gene_B": "GO:0046351", "interaction": "Direct", "SUID": 2743499, "TYPE": "Direct", "Gene_A": "GO:0005984", "BEND_MAP_ID": 2743499, "selected": false}, "selected": false}, {"data": {"id": "2743496", "source": "2741912", "target": "2741921", "SCORE": 0.706179066836963, "shared_interaction": "Direct", "source_original": "33", "Name_B": "GO:0009312", "Name_A": "GO:0005984", "target_original": "36", "Gene_B": "GO:0009312", "interaction": "Direct", "SUID": 2743496, "TYPE": "Direct", "Gene_A": "GO:0005984", "BEND_MAP_ID": 2743496, "selected": false}, "selected": false}, {"data": {"id": "2743493", "source": "2741912", "target": "2741918", "SCORE": 1.0, "shared_interaction": "Direct", "source_original": "33", "Name_B": "GO:0009311", "Name_A": "GO:0005984", "target_original": "35", "Gene_B": "GO:0009311", "interaction": "Direct", "SUID": 2743493, "TYPE": "Direct", "Gene_A": "GO:0005984", "BEND_MAP_ID": 2743493, "selected": false}, "selected": false}, {"data": {"id": "2743490", "source": "2741912", "target": "2741915", "SCORE": 0.706179066836963, "shared_interaction": "Direct", "source_original": "33", "Name_B": "GO:0005992", "Name_A": "GO:0005984", "target_original": "34", "Gene_B": "GO:0005992", "interaction": "Direct", "SUID": 2743490, "TYPE": "Direct", "Gene_A": "GO:0005984", "BEND_MAP_ID": 2743490, "selected": false}, "selected": false}, {"data": {"id": "2743454", "source": "2741912", "target": "2741909", "SCORE": 0.7936226749349893, "shared_interaction": "Direct", "source_original": "33", "Name_B": "WP398", "Name_A": "GO:0005984", "target_original": "32", "Gene_B": "WP398", "interaction": "Direct", "SUID": 2743454, "TYPE": "Direct", "Gene_A": "GO:0005984", "BEND_MAP_ID": 2743454, "selected": false}, "selected": false}, {"data": {"id": "2743292", "source": "2741912", "target": "2741891", "SCORE": 0.4519991447536391, "shared_interaction": "Direct", "source_original": "33", "Name_B": "GO:0034637", "Name_A": "GO:0005984", "target_original": "26", "Gene_B": "GO:0034637", "interaction": "Direct", "SUID": 2743292, "TYPE": "Direct", "Gene_A": "GO:0005984", "BEND_MAP_ID": 2743292, "selected": false}, "selected": false}, {"data": {"id": "2743151", "source": "2741912", "target": "2741906", "SCORE": 0.706179066836963, "shared_interaction": "Direct", "source_original": "33", "Name_B": "GO:0070413", "Name_A": "GO:0005984", "target_original": "31", "Gene_B": "GO:0070413", "interaction": "Direct", "SUID": 2743151, "TYPE": "Direct", "Gene_A": "GO:0005984", "BEND_MAP_ID": 2743151, "selected": false}, "selected": false}, {"data": {"id": "2743121", "source": "2741912", "target": "2741879", "SCORE": 0.3870725291270728, "shared_interaction": "Direct", "source_original": "33", "Name_B": "GO:0016051", "Name_A": "GO:0005984", "target_original": "22", "Gene_B": "GO:0016051", "interaction": "Direct", "SUID": 2743121, "TYPE": "Direct", "Gene_A": "GO:0005984", "BEND_MAP_ID": 2743121, "selected": false}, "selected": false}, {"data": {"id": "2742971", "source": "2741912", "target": "2742167", "SCORE": 0.9389572963063721, "shared_interaction": "Direct", "source_original": "33", "Name_B": "GO:0005991", "Name_A": "GO:0005984", "target_original": "118", "Gene_B": "GO:0005991", "interaction": "Direct", "SUID": 2742971, "TYPE": "Direct", "Gene_A": "GO:0005984", "BEND_MAP_ID": 2742971, "selected": false}, "selected": false}, {"data": {"id": "2742929", "source": "2741912", "target": "2742125", "SCORE": 0.5621946636621948, "shared_interaction": "Direct", "source_original": "33", "Name_B": "sce00500", "Name_A": "GO:0005984", "target_original": "104", "Gene_B": "sce00500", "interaction": "Direct", "SUID": 2742929, "TYPE": "Direct", "Gene_A": "GO:0005984", "BEND_MAP_ID": 2742929, "selected": false}, "selected": false}, {"data": {"id": "2742851", "source": "2741912", "target": "2741873", "SCORE": 0.4407105136837532, "shared_interaction": "Direct", "source_original": "33", "Name_B": "GO:0044262", "Name_A": "GO:0005984", "target_original": "20", "Gene_B": "GO:0044262", "interaction": "Direct", "SUID": 2742851, "TYPE": "Direct", "Gene_A": "GO:0005984", "BEND_MAP_ID": 2742851, "selected": false}, "selected": false}, {"data": {"id": "2743469", "source": "2741909", "target": "2742128", "SCORE": 0.38737949168022523, "shared_interaction": "Direct", "source_original": "32", "Name_B": "WP70", "Name_A": "WP398", "target_original": "105", "Gene_B": "WP70", "interaction": "Direct", "SUID": 2743469, "TYPE": "Direct", "Gene_A": "WP398", "BEND_MAP_ID": 2743469, "selected": false}, "selected": false}, {"data": {"id": "2742926", "source": "2741909", "target": "2742125", "SCORE": 0.5550604710397552, "shared_interaction": "Direct", "source_original": "32", "Name_B": "sce00500", "Name_A": "WP398", "target_original": "104", "Gene_B": "sce00500", "interaction": "Direct", "SUID": 2742926, "TYPE": "Direct", "Gene_A": "WP398", "BEND_MAP_ID": 2742926, "selected": false}, "selected": false}, {"data": {"id": "2743148", "source": "2741906", "target": "2741909", "SCORE": 0.7207351178611695, "shared_interaction": "Direct", "source_original": "31", "Name_B": "WP398", "Name_A": "GO:0070413", "target_original": "32", "Gene_B": "WP398", "interaction": "Direct", "SUID": 2743148, "TYPE": "Direct", "Gene_A": "GO:0070413", "BEND_MAP_ID": 2743148, "selected": false}, "selected": false}, {"data": {"id": "2742905", "source": "2741906", "target": "2742125", "SCORE": 0.38004434590120695, "shared_interaction": "Direct", "source_original": "31", "Name_B": "sce00500", "Name_A": "GO:0070413", "target_original": "104", "Gene_B": "sce00500", "interaction": "Direct", "SUID": 2742905, "TYPE": "Direct", "Gene_A": "GO:0070413", "BEND_MAP_ID": 2742905, "selected": false}, "selected": false}, {"data": {"id": "2743787", "source": "2741903", "target": "2741897", "SCORE": 0.7416851441258369, "shared_interaction": "Direct", "source_original": "30", "Name_B": "R-SCE-8982491", "Name_A": "GO:0044042", "target_original": "28", "Gene_B": "R-SCE-8982491", "interaction": "Direct", "SUID": 2743787, "TYPE": "Direct", "Gene_A": "GO:0044042", "BEND_MAP_ID": 2743787, "selected": false}, "selected": false}, {"data": {"id": "2743199", "source": "2741903", "target": "2741882", "SCORE": 0.4315092366701367, "shared_interaction": "Direct", "source_original": "30", "Name_B": "R-SCE-71387", "Name_A": "GO:0044042", "target_original": "23", "Gene_B": "R-SCE-71387", "interaction": "Direct", "SUID": 2743199, "TYPE": "Direct", "Gene_A": "GO:0044042", "BEND_MAP_ID": 2743199, "selected": false}, "selected": false}, {"data": {"id": "2742953", "source": "2741903", "target": "2742125", "SCORE": 0.45176470588466505, "shared_interaction": "Direct", "source_original": "30", "Name_B": "sce00500", "Name_A": "GO:0044042", "target_original": "104", "Gene_B": "sce00500", "interaction": "Direct", "SUID": 2742953, "TYPE": "Direct", "Gene_A": "GO:0044042", "BEND_MAP_ID": 2742953, "selected": false}, "selected": false}, {"data": {"id": "2742863", "source": "2741903", "target": "2741873", "SCORE": 0.4796050311522422, "shared_interaction": "Direct", "source_original": "30", "Name_B": "GO:0044262", "Name_A": "GO:0044042", "target_original": "20", "Gene_B": "GO:0044262", "interaction": "Direct", "SUID": 2742863, "TYPE": "Direct", "Gene_A": "GO:0044042", "BEND_MAP_ID": 2742863, "selected": false}, "selected": false}, {"data": {"id": "2743880", "source": "2741900", "target": "2742089", "SCORE": 0.30689142290930854, "shared_interaction": "Direct", "source_original": "29", "Name_B": "GO:0010675", "Name_A": "GO:0006073", "target_original": "92", "Gene_B": "GO:0010675", "interaction": "Direct", "SUID": 2743880, "TYPE": "Direct", "Gene_A": "GO:0006073", "BEND_MAP_ID": 2743880, "selected": false}, "selected": false}, {"data": {"id": "2743877", "source": "2741900", "target": "2742086", "SCORE": 0.30689142290930854, "shared_interaction": "Direct", "source_original": "29", "Name_B": "GO:0010906", "Name_A": "GO:0006073", "target_original": "91", "Gene_B": "GO:0010906", "interaction": "Direct", "SUID": 2743877, "TYPE": "Direct", "Gene_A": "GO:0006073", "BEND_MAP_ID": 2743877, "selected": false}, "selected": false}, {"data": {"id": "2743874", "source": "2741900", "target": "2741903", "SCORE": 1.0, "shared_interaction": "Direct", "source_original": "29", "Name_B": "GO:0044042", "Name_A": "GO:0006073", "target_original": "30", "Gene_B": "GO:0044042", "interaction": "Direct", "SUID": 2743874, "TYPE": "Direct", "Gene_A": "GO:0006073", "BEND_MAP_ID": 2743874, "selected": false}, "selected": false}, {"data": {"id": "2743784", "source": "2741900", "target": "2741897", "SCORE": 0.7416851441258369, "shared_interaction": "Direct", "source_original": "29", "Name_B": "R-SCE-8982491", "Name_A": "GO:0006073", "target_original": "28", "Gene_B": "R-SCE-8982491", "interaction": "Direct", "SUID": 2743784, "TYPE": "Direct", "Gene_A": "GO:0006073", "BEND_MAP_ID": 2743784, "selected": false}, "selected": false}, {"data": {"id": "2743310", "source": "2741900", "target": "2741891", "SCORE": 0.6185018419993633, "shared_interaction": "Direct", "source_original": "29", "Name_B": "GO:0034637", "Name_A": "GO:0006073", "target_original": "26", "Gene_B": "GO:0034637", "interaction": "Direct", "SUID": 2743310, "TYPE": "Direct", "Gene_A": "GO:0006073", "BEND_MAP_ID": 2743310, "selected": false}, "selected": false}, {"data": {"id": "2743244", "source": "2741900", "target": "2741885", "SCORE": 0.8955156950678105, "shared_interaction": "Direct", "source_original": "29", "Name_B": "GO:0006112", "Name_A": "GO:0006073", "target_original": "24", "Gene_B": "GO:0006112", "interaction": "Direct", "SUID": 2743244, "TYPE": "Direct", "Gene_A": "GO:0006073", "BEND_MAP_ID": 2743244, "selected": false}, "selected": false}, {"data": {"id": "2743196", "source": "2741900", "target": "2741882", "SCORE": 0.4315092366701367, "shared_interaction": "Direct", "source_original": "29", "Name_B": "R-SCE-71387", "Name_A": "GO:0006073", "target_original": "23", "Gene_B": "R-SCE-71387", "interaction": "Direct", "SUID": 2743196, "TYPE": "Direct", "Gene_A": "GO:0006073", "BEND_MAP_ID": 2743196, "selected": false}, "selected": false}, {"data": {"id": "2743139", "source": "2741900", "target": "2741879", "SCORE": 0.5361085972870243, "shared_interaction": "Direct", "source_original": "29", "Name_B": "GO:0016051", "Name_A": "GO:0006073", "target_original": "22", "Gene_B": "GO:0016051", "interaction": "Direct", "SUID": 2743139, "TYPE": "Direct", "Gene_A": "GO:0006073", "BEND_MAP_ID": 2743139, "selected": false}, "selected": false}, {"data": {"id": "2742950", "source": "2741900", "target": "2742125", "SCORE": 0.45176470588466505, "shared_interaction": "Direct", "source_original": "29", "Name_B": "sce00500", "Name_A": "GO:0006073", "target_original": "104", "Gene_B": "sce00500", "interaction": "Direct", "SUID": 2742950, "TYPE": "Direct", "Gene_A": "GO:0006073", "BEND_MAP_ID": 2742950, "selected": false}, "selected": false}, {"data": {"id": "2742860", "source": "2741900", "target": "2741873", "SCORE": 0.4796050311522422, "shared_interaction": "Direct", "source_original": "29", "Name_B": "GO:0044262", "Name_A": "GO:0006073", "target_original": "20", "Gene_B": "GO:0044262", "interaction": "Direct", "SUID": 2742860, "TYPE": "Direct", "Gene_A": "GO:0006073", "BEND_MAP_ID": 2742860, "selected": false}, "selected": false}, {"data": {"id": "2742947", "source": "2741897", "target": "2742125", "SCORE": 0.4561850201597008, "shared_interaction": "Direct", "source_original": "28", "Name_B": "sce00500", "Name_A": "R-SCE-8982491", "target_original": "104", "Gene_B": "sce00500", "interaction": "Direct", "SUID": 2742947, "TYPE": "Direct", "Gene_A": "R-SCE-8982491", "BEND_MAP_ID": 2742947, "selected": false}, "selected": false}, {"data": {"id": "2743427", "source": "2741894", "target": "2742089", "SCORE": 0.328530259370093, "shared_interaction": "Direct", "source_original": "27", "Name_B": "GO:0010675", "Name_A": "GO:0005977", "target_original": "92", "Gene_B": "GO:0010675", "interaction": "Direct", "SUID": 2743427, "TYPE": "Direct", "Gene_A": "GO:0005977", "BEND_MAP_ID": 2743427, "selected": false}, "selected": false}, {"data": {"id": "2743424", "source": "2741894", "target": "2742086", "SCORE": 0.328530259370093, "shared_interaction": "Direct", "source_original": "27", "Name_B": "GO:0010906", "Name_A": "GO:0005977", "target_original": "91", "Gene_B": "GO:0010906", "interaction": "Direct", "SUID": 2743424, "TYPE": "Direct", "Gene_A": "GO:0005977", "BEND_MAP_ID": 2743424, "selected": false}, "selected": false}, {"data": {"id": "2743421", "source": "2741894", "target": "2742152", "SCORE": 0.30654761905162614, "shared_interaction": "Direct", "source_original": "27", "Name_B": "GO:0006109", "Name_A": "GO:0005977", "target_original": "113", "Gene_B": "GO:0006109", "interaction": "Direct", "SUID": 2743421, "TYPE": "Direct", "Gene_A": "GO:0005977", "BEND_MAP_ID": 2743421, "selected": false}, "selected": false}, {"data": {"id": "2743418", "source": "2741894", "target": "2741903", "SCORE": 0.9451377442903881, "shared_interaction": "Direct", "source_original": "27", "Name_B": "GO:0044042", "Name_A": "GO:0005977", "target_original": "30", "Gene_B": "GO:0044042", "interaction": "Direct", "SUID": 2743418, "TYPE": "Direct", "Gene_A": "GO:0005977", "BEND_MAP_ID": 2743418, "selected": false}, "selected": false}, {"data": {"id": "2743415", "source": "2741894", "target": "2741900", "SCORE": 0.9451377442903881, "shared_interaction": "Direct", "source_original": "27", "Name_B": "GO:0006073", "Name_A": "GO:0005977", "target_original": "29", "Gene_B": "GO:0006073", "interaction": "Direct", "SUID": 2743415, "TYPE": "Direct", "Gene_A": "GO:0005977", "BEND_MAP_ID": 2743415, "selected": false}, "selected": false}, {"data": {"id": "2743412", "source": "2741894", "target": "2741897", "SCORE": 0.7936226749349893, "shared_interaction": "Direct", "source_original": "27", "Name_B": "R-SCE-8982491", "Name_A": "GO:0005977", "target_original": "28", "Gene_B": "R-SCE-8982491", "interaction": "Direct", "SUID": 2743412, "TYPE": "Direct", "Gene_A": "GO:0005977", "BEND_MAP_ID": 2743412, "selected": false}, "selected": false}, {"data": {"id": "2743286", "source": "2741894", "target": "2741891", "SCORE": 0.6512721830250431, "shared_interaction": "Direct", "source_original": "27", "Name_B": "GO:0034637", "Name_A": "GO:0005977", "target_original": "26", "Gene_B": "GO:0034637", "interaction": "Direct", "SUID": 2743286, "TYPE": "Direct", "Gene_A": "GO:0005977", "BEND_MAP_ID": 2743286, "selected": false}, "selected": false}, {"data": {"id": "2743265", "source": "2741894", "target": "2741888", "SCORE": 0.8706274292068336, "shared_interaction": "Direct", "source_original": "27", "Name_B": "GO:0005978", "Name_A": "GO:0005977", "target_original": "25", "Gene_B": "GO:0005978", "interaction": "Direct", "SUID": 2743265, "TYPE": "Direct", "Gene_A": "GO:0005977", "BEND_MAP_ID": 2743265, "selected": false}, "selected": false}, {"data": {"id": "2743238", "source": "2741894", "target": "2741885", "SCORE": 0.9451377442903881, "shared_interaction": "Direct", "source_original": "27", "Name_B": "GO:0006112", "Name_A": "GO:0005977", "target_original": "24", "Gene_B": "GO:0006112", "interaction": "Direct", "SUID": 2743238, "TYPE": "Direct", "Gene_A": "GO:0005977", "BEND_MAP_ID": 2743238, "selected": false}, "selected": false}, {"data": {"id": "2743187", "source": "2741894", "target": "2741882", "SCORE": 0.45295286256325556, "shared_interaction": "Direct", "source_original": "27", "Name_B": "R-SCE-71387", "Name_A": "GO:0005977", "target_original": "23", "Gene_B": "R-SCE-71387", "interaction": "Direct", "SUID": 2743187, "TYPE": "Direct", "Gene_A": "GO:0005977", "BEND_MAP_ID": 2743187, "selected": false}, "selected": false}, {"data": {"id": "2743115", "source": "2741894", "target": "2741879", "SCORE": 0.5621946636621948, "shared_interaction": "Direct", "source_original": "27", "Name_B": "GO:0016051", "Name_A": "GO:0005977", "target_original": "22", "Gene_B": "GO:0016051", "interaction": "Direct", "SUID": 2743115, "TYPE": "Direct", "Gene_A": "GO:0005977", "BEND_MAP_ID": 2743115, "selected": false}, "selected": false}, {"data": {"id": "2742923", "source": "2741894", "target": "2742125", "SCORE": 0.4746335963946338, "shared_interaction": "Direct", "source_original": "27", "Name_B": "sce00500", "Name_A": "GO:0005977", "target_original": "104", "Gene_B": "sce00500", "interaction": "Direct", "SUID": 2742923, "TYPE": "Direct", "Gene_A": "GO:0005977", "BEND_MAP_ID": 2742923, "selected": false}, "selected": false}, {"data": {"id": "2742845", "source": "2741894", "target": "2741873", "SCORE": 0.4407105136837532, "shared_interaction": "Direct", "source_original": "27", "Name_B": "GO:0044262", "Name_A": "GO:0005977", "target_original": "20", "Gene_B": "GO:0044262", "interaction": "Direct", "SUID": 2742845, "TYPE": "Direct", "Gene_A": "GO:0005977", "BEND_MAP_ID": 2742845, "selected": false}, "selected": false}, {"data": {"id": "2743313", "source": "2741891", "target": "2741903", "SCORE": 0.6185018419993633, "shared_interaction": "Direct", "source_original": "26", "Name_B": "GO:0044042", "Name_A": "GO:0034637", "target_original": "30", "Gene_B": "GO:0044042", "interaction": "Direct", "SUID": 2743313, "TYPE": "Direct", "Gene_A": "GO:0034637", "BEND_MAP_ID": 2743313, "selected": false}, "selected": false}, {"data": {"id": "2743307", "source": "2741891", "target": "2741897", "SCORE": 0.5397530864224009, "shared_interaction": "Direct", "source_original": "26", "Name_B": "R-SCE-8982491", "Name_A": "GO:0034637", "target_original": "28", "Gene_B": "R-SCE-8982491", "interaction": "Direct", "SUID": 2743307, "TYPE": "Direct", "Gene_A": "GO:0034637", "BEND_MAP_ID": 2743307, "selected": false}, "selected": false}, {"data": {"id": "2743304", "source": "2741891", "target": "2741924", "SCORE": 0.5753709971387387, "shared_interaction": "Direct", "source_original": "26", "Name_B": "GO:0046351", "Name_A": "GO:0034637", "target_original": "37", "Gene_B": "GO:0046351", "interaction": "Direct", "SUID": 2743304, "TYPE": "Direct", "Gene_A": "GO:0034637", "BEND_MAP_ID": 2743304, "selected": false}, "selected": false}, {"data": {"id": "2743289", "source": "2741891", "target": "2741909", "SCORE": 0.5397530864224009, "shared_interaction": "Direct", "source_original": "26", "Name_B": "WP398", "Name_A": "GO:0034637", "target_original": "32", "Gene_B": "WP398", "interaction": "Direct", "SUID": 2743289, "TYPE": "Direct", "Gene_A": "GO:0034637", "BEND_MAP_ID": 2743289, "selected": false}, "selected": false}, {"data": {"id": "2743181", "source": "2741891", "target": "2741882", "SCORE": 0.31693811075177764, "shared_interaction": "Direct", "source_original": "26", "Name_B": "R-SCE-71387", "Name_A": "GO:0034637", "target_original": "23", "Gene_B": "R-SCE-71387", "interaction": "Direct", "SUID": 2743181, "TYPE": "Direct", "Gene_A": "GO:0034637", "BEND_MAP_ID": 2743181, "selected": false}, "selected": false}, {"data": {"id": "2743145", "source": "2741891", "target": "2741906", "SCORE": 0.454048424892664, "shared_interaction": "Direct", "source_original": "26", "Name_B": "GO:0070413", "Name_A": "GO:0034637", "target_original": "31", "Gene_B": "GO:0070413", "interaction": "Direct", "SUID": 2743145, "TYPE": "Direct", "Gene_A": "GO:0034637", "BEND_MAP_ID": 2743145, "selected": false}, "selected": false}, {"data": {"id": "2742917", "source": "2741891", "target": "2742125", "SCORE": 0.646433990896685, "shared_interaction": "Direct", "source_original": "26", "Name_B": "sce00500", "Name_A": "GO:0034637", "target_original": "104", "Gene_B": "sce00500", "interaction": "Direct", "SUID": 2742917, "TYPE": "Direct", "Gene_A": "GO:0034637", "BEND_MAP_ID": 2742917, "selected": false}, "selected": false}, {"data": {"id": "2742839", "source": "2741891", "target": "2741873", "SCORE": 0.5527831094061669, "shared_interaction": "Direct", "source_original": "26", "Name_B": "GO:0044262", "Name_A": "GO:0034637", "target_original": "20", "Gene_B": "GO:0044262", "interaction": "Direct", "SUID": 2742839, "TYPE": "Direct", "Gene_A": "GO:0034637", "BEND_MAP_ID": 2742839, "selected": false}, "selected": false}, {"data": {"id": "2743274", "source": "2741888", "target": "2741903", "SCORE": 0.8170635959184147, "shared_interaction": "Direct", "source_original": "25", "Name_B": "GO:0044042", "Name_A": "GO:0005978", "target_original": "30", "Gene_B": "GO:0044042", "interaction": "Direct", "SUID": 2743274, "TYPE": "Direct", "Gene_A": "GO:0005978", "BEND_MAP_ID": 2743274, "selected": false}, "selected": false}, {"data": {"id": "2743271", "source": "2741888", "target": "2741900", "SCORE": 0.8170635959184147, "shared_interaction": "Direct", "source_original": "25", "Name_B": "GO:0006073", "Name_A": "GO:0005978", "target_original": "29", "Gene_B": "GO:0006073", "interaction": "Direct", "SUID": 2743271, "TYPE": "Direct", "Gene_A": "GO:0005978", "BEND_MAP_ID": 2743271, "selected": false}, "selected": false}, {"data": {"id": "2743268", "source": "2741888", "target": "2741897", "SCORE": 0.7626485568779389, "shared_interaction": "Direct", "source_original": "25", "Name_B": "R-SCE-8982491", "Name_A": "GO:0005978", "target_original": "28", "Gene_B": "R-SCE-8982491", "interaction": "Direct", "SUID": 2743268, "TYPE": "Direct", "Gene_A": "GO:0005978", "BEND_MAP_ID": 2743268, "selected": false}, "selected": false}, {"data": {"id": "2743262", "source": "2741888", "target": "2741891", "SCORE": 0.7264616107082353, "shared_interaction": "Direct", "source_original": "25", "Name_B": "GO:0034637", "Name_A": "GO:0005978", "target_original": "26", "Gene_B": "GO:0034637", "interaction": "Direct", "SUID": 2743262, "TYPE": "Direct", "Gene_A": "GO:0005978", "BEND_MAP_ID": 2743262, "selected": false}, "selected": false}, {"data": {"id": "2743232", "source": "2741888", "target": "2741885", "SCORE": 0.8170635959184147, "shared_interaction": "Direct", "source_original": "25", "Name_B": "GO:0006112", "Name_A": "GO:0005978", "target_original": "24", "Gene_B": "GO:0006112", "interaction": "Direct", "SUID": 2743232, "TYPE": "Direct", "Gene_A": "GO:0005978", "BEND_MAP_ID": 2743232, "selected": false}, "selected": false}, {"data": {"id": "2743178", "source": "2741888", "target": "2741882", "SCORE": 0.4101265822811573, "shared_interaction": "Direct", "source_original": "25", "Name_B": "R-SCE-71387", "Name_A": "GO:0005978", "target_original": "23", "Gene_B": "R-SCE-71387", "interaction": "Direct", "SUID": 2743178, "TYPE": "Direct", "Gene_A": "GO:0005978", "BEND_MAP_ID": 2743178, "selected": false}, "selected": false}, {"data": {"id": "2743109", "source": "2741888", "target": "2741879", "SCORE": 0.6208299430449216, "shared_interaction": "Direct", "source_original": "25", "Name_B": "GO:0016051", "Name_A": "GO:0005978", "target_original": "22", "Gene_B": "GO:0016051", "interaction": "Direct", "SUID": 2743109, "TYPE": "Direct", "Gene_A": "GO:0005978", "BEND_MAP_ID": 2743109, "selected": false}, "selected": false}, {"data": {"id": "2742914", "source": "2741888", "target": "2742125", "SCORE": 0.4312449145673823, "shared_interaction": "Direct", "source_original": "25", "Name_B": "sce00500", "Name_A": "GO:0005978", "target_original": "104", "Gene_B": "sce00500", "interaction": "Direct", "SUID": 2742914, "TYPE": "Direct", "Gene_A": "GO:0005978", "BEND_MAP_ID": 2742914, "selected": false}, "selected": false}, {"data": {"id": "2742836", "source": "2741888", "target": "2741873", "SCORE": 0.35780506138999774, "shared_interaction": "Direct", "source_original": "25", "Name_B": "GO:0044262", "Name_A": "GO:0005978", "target_original": "20", "Gene_B": "GO:0044262", "interaction": "Direct", "SUID": 2742836, "TYPE": "Direct", "Gene_A": "GO:0005978", "BEND_MAP_ID": 2742836, "selected": false}, "selected": false}, {"data": {"id": "2743253", "source": "2741885", "target": "2742089", "SCORE": 0.30689142290930854, "shared_interaction": "Direct", "source_original": "24", "Name_B": "GO:0010675", "Name_A": "GO:0006112", "target_original": "92", "Gene_B": "GO:0010675", "interaction": "Direct", "SUID": 2743253, "TYPE": "Direct", "Gene_A": "GO:0006112", "BEND_MAP_ID": 2743253, "selected": false}, "selected": false}, {"data": {"id": "2743250", "source": "2741885", "target": "2742086", "SCORE": 0.30689142290930854, "shared_interaction": "Direct", "source_original": "24", "Name_B": "GO:0010906", "Name_A": "GO:0006112", "target_original": "91", "Gene_B": "GO:0010906", "interaction": "Direct", "SUID": 2743250, "TYPE": "Direct", "Gene_A": "GO:0006112", "BEND_MAP_ID": 2743250, "selected": false}, "selected": false}, {"data": {"id": "2743247", "source": "2741885", "target": "2741903", "SCORE": 0.8955156950678105, "shared_interaction": "Direct", "source_original": "24", "Name_B": "GO:0044042", "Name_A": "GO:0006112", "target_original": "30", "Gene_B": "GO:0044042", "interaction": "Direct", "SUID": 2743247, "TYPE": "Direct", "Gene_A": "GO:0006112", "BEND_MAP_ID": 2743247, "selected": false}, "selected": false}, {"data": {"id": "2743241", "source": "2741885", "target": "2741897", "SCORE": 0.7416851441258369, "shared_interaction": "Direct", "source_original": "24", "Name_B": "R-SCE-8982491", "Name_A": "GO:0006112", "target_original": "28", "Gene_B": "R-SCE-8982491", "interaction": "Direct", "SUID": 2743241, "TYPE": "Direct", "Gene_A": "GO:0006112", "BEND_MAP_ID": 2743241, "selected": false}, "selected": false}, {"data": {"id": "2743235", "source": "2741885", "target": "2741891", "SCORE": 0.6185018419993633, "shared_interaction": "Direct", "source_original": "24", "Name_B": "GO:0034637", "Name_A": "GO:0006112", "target_original": "26", "Gene_B": "GO:0034637", "interaction": "Direct", "SUID": 2743235, "TYPE": "Direct", "Gene_A": "GO:0006112", "BEND_MAP_ID": 2743235, "selected": false}, "selected": false}, {"data": {"id": "2743175", "source": "2741885", "target": "2741882", "SCORE": 0.4315092366701367, "shared_interaction": "Direct", "source_original": "24", "Name_B": "R-SCE-71387", "Name_A": "GO:0006112", "target_original": "23", "Gene_B": "R-SCE-71387", "interaction": "Direct", "SUID": 2743175, "TYPE": "Direct", "Gene_A": "GO:0006112", "BEND_MAP_ID": 2743175, "selected": false}, "selected": false}, {"data": {"id": "2743106", "source": "2741885", "target": "2741879", "SCORE": 0.5361085972870243, "shared_interaction": "Direct", "source_original": "24", "Name_B": "GO:0016051", "Name_A": "GO:0006112", "target_original": "22", "Gene_B": "GO:0016051", "interaction": "Direct", "SUID": 2743106, "TYPE": "Direct", "Gene_A": "GO:0006112", "BEND_MAP_ID": 2743106, "selected": false}, "selected": false}, {"data": {"id": "2742911", "source": "2741885", "target": "2742125", "SCORE": 0.45176470588466505, "shared_interaction": "Direct", "source_original": "24", "Name_B": "sce00500", "Name_A": "GO:0006112", "target_original": "104", "Gene_B": "sce00500", "interaction": "Direct", "SUID": 2742911, "TYPE": "Direct", "Gene_A": "GO:0006112", "BEND_MAP_ID": 2742911, "selected": false}, "selected": false}, {"data": {"id": "2742833", "source": "2741885", "target": "2741873", "SCORE": 0.424826613378794, "shared_interaction": "Direct", "source_original": "24", "Name_B": "GO:0044262", "Name_A": "GO:0006112", "target_original": "20", "Gene_B": "GO:0044262", "interaction": "Direct", "SUID": 2742833, "TYPE": "Direct", "Gene_A": "GO:0006112", "BEND_MAP_ID": 2742833, "selected": false}, "selected": false}, {"data": {"id": "2742365", "source": "2741885", "target": "2741822", "SCORE": 0.3083773087084526, "shared_interaction": "Direct", "source_original": "24", "Name_B": "GO:0015980", "Name_A": "GO:0006112", "target_original": "3", "Gene_B": "GO:0015980", "interaction": "Direct", "SUID": 2742365, "TYPE": "Direct", "Gene_A": "GO:0006112", "BEND_MAP_ID": 2742365, "selected": false}, "selected": false}, {"data": {"id": "2743202", "source": "2741882", "target": "2742017", "SCORE": 0.4315092366701367, "shared_interaction": "Direct", "source_original": "23", "Name_B": "sce00010", "Name_A": "R-SCE-71387", "target_original": "68", "Gene_B": "sce00010", "interaction": "Direct", "SUID": 2743202, "TYPE": "Direct", "Gene_A": "R-SCE-71387", "BEND_MAP_ID": 2743202, "selected": false}, "selected": false}, {"data": {"id": "2743193", "source": "2741882", "target": "2741897", "SCORE": 0.527766518040333, "shared_interaction": "Direct", "source_original": "23", "Name_B": "R-SCE-8982491", "Name_A": "R-SCE-71387", "target_original": "28", "Gene_B": "R-SCE-8982491", "interaction": "Direct", "SUID": 2743193, "TYPE": "Direct", "Gene_A": "R-SCE-71387", "BEND_MAP_ID": 2743193, "selected": false}, "selected": false}, {"data": {"id": "2743073", "source": "2741882", "target": "2741927", "SCORE": 0.3575904419763528, "shared_interaction": "Direct", "source_original": "23", "Name_B": "sce01110", "Name_A": "R-SCE-71387", "target_original": "38", "Gene_B": "sce01110", "interaction": "Direct", "SUID": 2743073, "TYPE": "Direct", "Gene_A": "R-SCE-71387", "BEND_MAP_ID": 2743073, "selected": false}, "selected": false}, {"data": {"id": "2743013", "source": "2741882", "target": "2742122", "SCORE": 0.3022052817882191, "shared_interaction": "Direct", "source_original": "23", "Name_B": "WP112", "Name_A": "R-SCE-71387", "target_original": "103", "Gene_B": "WP112", "interaction": "Direct", "SUID": 2743013, "TYPE": "Direct", "Gene_A": "R-SCE-71387", "BEND_MAP_ID": 2743013, "selected": false}, "selected": false}, {"data": {"id": "2742908", "source": "2741882", "target": "2742125", "SCORE": 0.4816847100714928, "shared_interaction": "Direct", "source_original": "23", "Name_B": "sce00500", "Name_A": "R-SCE-71387", "target_original": "104", "Gene_B": "sce00500", "interaction": "Direct", "SUID": 2742908, "TYPE": "Direct", "Gene_A": "R-SCE-71387", "BEND_MAP_ID": 2742908, "selected": false}, "selected": false}, {"data": {"id": "2743142", "source": "2741879", "target": "2741903", "SCORE": 0.5361085972870243, "shared_interaction": "Direct", "source_original": "22", "Name_B": "GO:0044042", "Name_A": "GO:0016051", "target_original": "30", "Gene_B": "GO:0044042", "interaction": "Direct", "SUID": 2743142, "TYPE": "Direct", "Gene_A": "GO:0016051", "BEND_MAP_ID": 2743142, "selected": false}, "selected": false}, {"data": {"id": "2743136", "source": "2741879", "target": "2741897", "SCORE": 0.4561850201597008, "shared_interaction": "Direct", "source_original": "22", "Name_B": "R-SCE-8982491", "Name_A": "GO:0016051", "target_original": "28", "Gene_B": "R-SCE-8982491", "interaction": "Direct", "SUID": 2743136, "TYPE": "Direct", "Gene_A": "GO:0016051", "BEND_MAP_ID": 2743136, "selected": false}, "selected": false}, {"data": {"id": "2743133", "source": "2741879", "target": "2741924", "SCORE": 0.4833702882510058, "shared_interaction": "Direct", "source_original": "22", "Name_B": "GO:0046351", "Name_A": "GO:0016051", "target_original": "37", "Gene_B": "GO:0046351", "interaction": "Direct", "SUID": 2743133, "TYPE": "Direct", "Gene_A": "GO:0016051", "BEND_MAP_ID": 2743133, "selected": false}, "selected": false}, {"data": {"id": "2743118", "source": "2741879", "target": "2741909", "SCORE": 0.4561850201597008, "shared_interaction": "Direct", "source_original": "22", "Name_B": "WP398", "Name_A": "GO:0016051", "target_original": "32", "Gene_B": "WP398", "interaction": "Direct", "SUID": 2743118, "TYPE": "Direct", "Gene_A": "GO:0016051", "BEND_MAP_ID": 2743118, "selected": false}, "selected": false}, {"data": {"id": "2743112", "source": "2741879", "target": "2741891", "SCORE": 0.8821446636322283, "shared_interaction": "Direct", "source_original": "22", "Name_B": "GO:0034637", "Name_A": "GO:0016051", "target_original": "26", "Gene_B": "GO:0034637", "interaction": "Direct", "SUID": 2743112, "TYPE": "Direct", "Gene_A": "GO:0016051", "BEND_MAP_ID": 2743112, "selected": false}, "selected": false}, {"data": {"id": "2743103", "source": "2741879", "target": "2741882", "SCORE": 0.3434672994238909, "shared_interaction": "Direct", "source_original": "22", "Name_B": "R-SCE-71387", "Name_A": "GO:0016051", "target_original": "23", "Gene_B": "R-SCE-71387", "interaction": "Direct", "SUID": 2743103, "TYPE": "Direct", "Gene_A": "GO:0016051", "BEND_MAP_ID": 2743103, "selected": false}, "selected": false}, {"data": {"id": "2743100", "source": "2741879", "target": "2741906", "SCORE": 0.38004434590120695, "shared_interaction": "Direct", "source_original": "22", "Name_B": "GO:0070413", "Name_A": "GO:0016051", "target_original": "31", "Gene_B": "GO:0070413", "interaction": "Direct", "SUID": 2743100, "TYPE": "Direct", "Gene_A": "GO:0016051", "BEND_MAP_ID": 2743100, "selected": false}, "selected": false}, {"data": {"id": "2743070", "source": "2741879", "target": "2741927", "SCORE": 0.3296666102568638, "shared_interaction": "Direct", "source_original": "22", "Name_B": "sce01110", "Name_A": "GO:0016051", "target_original": "38", "Gene_B": "sce01110", "interaction": "Direct", "SUID": 2743070, "TYPE": "Direct", "Gene_A": "GO:0016051", "BEND_MAP_ID": 2743070, "selected": false}, "selected": false}, {"data": {"id": "2742902", "source": "2741879", "target": "2742125", "SCORE": 0.5724770642217064, "shared_interaction": "Direct", "source_original": "22", "Name_B": "sce00500", "Name_A": "GO:0016051", "target_original": "104", "Gene_B": "sce00500", "interaction": "Direct", "SUID": 2742902, "TYPE": "Direct", "Gene_A": "GO:0016051", "BEND_MAP_ID": 2742902, "selected": false}, "selected": false}, {"data": {"id": "2742827", "source": "2741879", "target": "2741873", "SCORE": 0.5033041995322504, "shared_interaction": "Direct", "source_original": "22", "Name_B": "GO:0044262", "Name_A": "GO:0016051", "target_original": "20", "Gene_B": "GO:0044262", "interaction": "Direct", "SUID": 2742827, "TYPE": "Direct", "Gene_A": "GO:0016051", "BEND_MAP_ID": 2742827, "selected": false}, "selected": false}, {"data": {"id": "2743355", "source": "2741876", "target": "2742002", "SCORE": 0.4584829917626492, "shared_interaction": "Direct", "source_original": "21", "Name_B": "GO:0044275", "Name_A": "GO:0016052", "target_original": "63", "Gene_B": "GO:0044275", "interaction": "Direct", "SUID": 2743355, "TYPE": "Direct", "Gene_A": "GO:0016052", "BEND_MAP_ID": 2743355, "selected": false}, "selected": false}, {"data": {"id": "2743352", "source": "2741876", "target": "2742041", "SCORE": 0.527766518040333, "shared_interaction": "Direct", "source_original": "21", "Name_B": "GO:0046031", "Name_A": "GO:0016052", "target_original": "76", "Gene_B": "GO:0046031", "interaction": "Direct", "SUID": 2743352, "TYPE": "Direct", "Gene_A": "GO:0016052", "BEND_MAP_ID": 2743352, "selected": false}, "selected": false}, {"data": {"id": "2743346", "source": "2741876", "target": "2741960", "SCORE": 0.3804058782385496, "shared_interaction": "Direct", "source_original": "21", "Name_B": "GO:0032787", "Name_A": "GO:0016052", "target_original": "49", "Gene_B": "GO:0032787", "interaction": "Direct", "SUID": 2743346, "TYPE": "Direct", "Gene_A": "GO:0016052", "BEND_MAP_ID": 2743346, "selected": false}, "selected": false}, {"data": {"id": "2743337", "source": "2741876", "target": "2741957", "SCORE": 0.3461178671673566, "shared_interaction": "Direct", "source_original": "21", "Name_B": "GO:0016310", "Name_A": "GO:0016052", "target_original": "48", "Gene_B": "GO:0016310", "interaction": "Direct", "SUID": 2743337, "TYPE": "Direct", "Gene_A": "GO:0016052", "BEND_MAP_ID": 2743337, "selected": false}, "selected": false}, {"data": {"id": "2743334", "source": "2741876", "target": "2742029", "SCORE": 0.30062341261055076, "shared_interaction": "Direct", "source_original": "21", "Name_B": "R-SCE-170822", "Name_A": "GO:0016052", "target_original": "72", "Gene_B": "R-SCE-170822", "interaction": "Direct", "SUID": 2743334, "TYPE": "Direct", "Gene_A": "GO:0016052", "BEND_MAP_ID": 2743334, "selected": false}, "selected": false}, {"data": {"id": "2743331", "source": "2741876", "target": "2742026", "SCORE": 0.43331982164839955, "shared_interaction": "Direct", "source_original": "21", "Name_B": "R-SCE-70171", "Name_A": "GO:0016052", "target_original": "71", "Gene_B": "R-SCE-70171", "interaction": "Direct", "SUID": 2743331, "TYPE": "Direct", "Gene_A": "GO:0016052", "BEND_MAP_ID": 2743331, "selected": false}, "selected": false}, {"data": {"id": "2743328", "source": "2741876", "target": "2742023", "SCORE": 0.4101265822811573, "shared_interaction": "Direct", "source_original": "21", "Name_B": "R-SCE-70326", "Name_A": "GO:0016052", "target_original": "70", "Gene_B": "R-SCE-70326", "interaction": "Direct", "SUID": 2743328, "TYPE": "Direct", "Gene_A": "GO:0016052", "BEND_MAP_ID": 2743328, "selected": false}, "selected": false}, {"data": {"id": "2743325", "source": "2741876", "target": "2742020", "SCORE": 0.36002535390131263, "shared_interaction": "Direct", "source_original": "21", "Name_B": "GO:0046835", "Name_A": "GO:0016052", "target_original": "69", "Gene_B": "GO:0046835", "interaction": "Direct", "SUID": 2743325, "TYPE": "Direct", "Gene_A": "GO:0016052", "BEND_MAP_ID": 2743325, "selected": false}, "selected": false}, {"data": {"id": "2743322", "source": "2741876", "target": "2742017", "SCORE": 0.5127222028601172, "shared_interaction": "Direct", "source_original": "21", "Name_B": "sce00010", "Name_A": "GO:0016052", "target_original": "68", "Gene_B": "sce00010", "interaction": "Direct", "SUID": 2743322, "TYPE": "Direct", "Gene_A": "GO:0016052", "BEND_MAP_ID": 2743322, "selected": false}, "selected": false}, {"data": {"id": "2743316", "source": "2741876", "target": "2741930", "SCORE": 0.35166291241753117, "shared_interaction": "Direct", "source_original": "21", "Name_B": "sce01200", "Name_A": "GO:0016052", "target_original": "39", "Gene_B": "sce01200", "interaction": "Direct", "SUID": 2743316, "TYPE": "Direct", "Gene_A": "GO:0016052", "BEND_MAP_ID": 2743316, "selected": false}, "selected": false}, {"data": {"id": "2743184", "source": "2741876", "target": "2741882", "SCORE": 0.39602534562414665, "shared_interaction": "Direct", "source_original": "21", "Name_B": "R-SCE-71387", "Name_A": "GO:0016052", "target_original": "23", "Gene_B": "R-SCE-71387", "interaction": "Direct", "SUID": 2743184, "TYPE": "Direct", "Gene_A": "GO:0016052", "BEND_MAP_ID": 2743184, "selected": false}, "selected": false}, {"data": {"id": "2743076", "source": "2741876", "target": "2741927", "SCORE": 0.3575904419763528, "shared_interaction": "Direct", "source_original": "21", "Name_B": "sce01110", "Name_A": "GO:0016052", "target_original": "38", "Gene_B": "sce01110", "interaction": "Direct", "SUID": 2743076, "TYPE": "Direct", "Gene_A": "GO:0016052", "BEND_MAP_ID": 2743076, "selected": false}, "selected": false}, {"data": {"id": "2742920", "source": "2741876", "target": "2742125", "SCORE": 0.41257600474769185, "shared_interaction": "Direct", "source_original": "21", "Name_B": "sce00500", "Name_A": "GO:0016052", "target_original": "104", "Gene_B": "sce00500", "interaction": "Direct", "SUID": 2742920, "TYPE": "Direct", "Gene_A": "GO:0016052", "BEND_MAP_ID": 2742920, "selected": false}, "selected": false}, {"data": {"id": "2742842", "source": "2741876", "target": "2741873", "SCORE": 0.48801925290487896, "shared_interaction": "Direct", "source_original": "21", "Name_B": "GO:0044262", "Name_A": "GO:0016052", "target_original": "20", "Gene_B": "GO:0044262", "interaction": "Direct", "SUID": 2742842, "TYPE": "Direct", "Gene_A": "GO:0016052", "BEND_MAP_ID": 2742842, "selected": false}, "selected": false}, {"data": {"id": "2742857", "source": "2741873", "target": "2741897", "SCORE": 0.31356474958575564, "shared_interaction": "Direct", "source_original": "20", "Name_B": "R-SCE-8982491", "Name_A": "GO:0044262", "target_original": "28", "Gene_B": "R-SCE-8982491", "interaction": "Direct", "SUID": 2742857, "TYPE": "Direct", "Gene_A": "GO:0044262", "BEND_MAP_ID": 2742857, "selected": false}, "selected": false}, {"data": {"id": "2742848", "source": "2741873", "target": "2741909", "SCORE": 0.31356474958575564, "shared_interaction": "Direct", "source_original": "20", "Name_B": "WP398", "Name_A": "GO:0044262", "target_original": "32", "Gene_B": "WP398", "interaction": "Direct", "SUID": 2742848, "TYPE": "Direct", "Gene_A": "GO:0044262", "BEND_MAP_ID": 2742848, "selected": false}, "selected": false}, {"data": {"id": "2742830", "source": "2741873", "target": "2741882", "SCORE": 0.4392591817529627, "shared_interaction": "Direct", "source_original": "20", "Name_B": "R-SCE-71387", "Name_A": "GO:0044262", "target_original": "23", "Gene_B": "R-SCE-71387", "interaction": "Direct", "SUID": 2742830, "TYPE": "Direct", "Gene_A": "GO:0044262", "BEND_MAP_ID": 2742830, "selected": false}, "selected": false}, {"data": {"id": "2742824", "source": "2741873", "target": "2741927", "SCORE": 0.4313809933974482, "shared_interaction": "Direct", "source_original": "20", "Name_B": "sce01110", "Name_A": "GO:0044262", "target_original": "38", "Gene_B": "sce01110", "interaction": "Direct", "SUID": 2742824, "TYPE": "Direct", "Gene_A": "GO:0044262", "BEND_MAP_ID": 2742824, "selected": false}, "selected": false}, {"data": {"id": "2742818", "source": "2741873", "target": "2742125", "SCORE": 0.6523129396725752, "shared_interaction": "Direct", "source_original": "20", "Name_B": "sce00500", "Name_A": "GO:0044262", "target_original": "104", "Gene_B": "sce00500", "interaction": "Direct", "SUID": 2742818, "TYPE": "Direct", "Gene_A": "GO:0044262", "BEND_MAP_ID": 2742818, "selected": false}, "selected": false}, {"data": {"id": "2742800", "source": "2741870", "target": "2741882", "SCORE": 0.36396724294909777, "shared_interaction": "Direct", "source_original": "19", "Name_B": "R-SCE-71387", "Name_A": "R-SCE-1430728", "target_original": "23", "Gene_B": "R-SCE-71387", "interaction": "Direct", "SUID": 2742800, "TYPE": "Direct", "Gene_A": "R-SCE-1430728", "BEND_MAP_ID": 2742800, "selected": false}, "selected": false}, {"data": {"id": "2742689", "source": "2741870", "target": "2742119", "SCORE": 0.38725822870776544, "shared_interaction": "Direct", "source_original": "19", "Name_B": "sce01100", "Name_A": "R-SCE-1430728", "target_original": "102", "Gene_B": "sce01100", "interaction": "Direct", "SUID": 2742689, "TYPE": "Direct", "Gene_A": "R-SCE-1430728", "BEND_MAP_ID": 2742689, "selected": false}, "selected": false}, {"data": {"id": "2742593", "source": "2741870", "target": "2741837", "SCORE": 0.3837966845136011, "shared_interaction": "Direct", "source_original": "19", "Name_B": "R-SCE-163200", "Name_A": "R-SCE-1430728", "target_original": "8", "Gene_B": "R-SCE-163200", "interaction": "Direct", "SUID": 2742593, "TYPE": "Direct", "Gene_A": "R-SCE-1430728", "BEND_MAP_ID": 2742593, "selected": false}, "selected": false}, {"data": {"id": "2742893", "source": "2741867", "target": "2742041", "SCORE": 0.37421665174792307, "shared_interaction": "Direct", "source_original": "18", "Name_B": "GO:0046031", "Name_A": "GO:0009150", "target_original": "76", "Gene_B": "GO:0046031", "interaction": "Direct", "SUID": 2742893, "TYPE": "Direct", "Gene_A": "GO:0009150", "BEND_MAP_ID": 2742893, "selected": false}, "selected": false}, {"data": {"id": "2742881", "source": "2741867", "target": "2742026", "SCORE": 0.3046851686088034, "shared_interaction": "Direct", "source_original": "18", "Name_B": "R-SCE-70171", "Name_A": "GO:0009150", "target_original": "71", "Gene_B": "R-SCE-70171", "interaction": "Direct", "SUID": 2742881, "TYPE": "Direct", "Gene_A": "GO:0009150", "BEND_MAP_ID": 2742881, "selected": false}, "selected": false}, {"data": {"id": "2742878", "source": "2741867", "target": "2742017", "SCORE": 0.31120666487719, "shared_interaction": "Direct", "source_original": "18", "Name_B": "sce00010", "Name_A": "GO:0009150", "target_original": "68", "Gene_B": "sce00010", "interaction": "Direct", "SUID": 2742878, "TYPE": "Direct", "Gene_A": "GO:0009150", "BEND_MAP_ID": 2742878, "selected": false}, "selected": false}, {"data": {"id": "2742875", "source": "2741867", "target": "2741930", "SCORE": 0.3092126935342147, "shared_interaction": "Direct", "source_original": "18", "Name_B": "sce01200", "Name_A": "GO:0009150", "target_original": "39", "Gene_B": "sce01200", "interaction": "Direct", "SUID": 2742875, "TYPE": "Direct", "Gene_A": "GO:0009150", "BEND_MAP_ID": 2742875, "selected": false}, "selected": false}, {"data": {"id": "2742872", "source": "2741867", "target": "2742164", "SCORE": 0.46535107847707985, "shared_interaction": "Direct", "source_original": "18", "Name_B": "GO:0044281", "Name_A": "GO:0009150", "target_original": "117", "Gene_B": "GO:0044281", "interaction": "Direct", "SUID": 2742872, "TYPE": "Direct", "Gene_A": "GO:0009150", "BEND_MAP_ID": 2742872, "selected": false}, "selected": false}, {"data": {"id": "2742797", "source": "2741867", "target": "2741870", "SCORE": 0.31478985418551425, "shared_interaction": "Direct", "source_original": "18", "Name_B": "R-SCE-1430728", "Name_A": "GO:0009150", "target_original": "19", "Gene_B": "R-SCE-1430728", "interaction": "Direct", "SUID": 2742797, "TYPE": "Direct", "Gene_A": "GO:0009150", "BEND_MAP_ID": 2742797, "selected": false}, "selected": false}, {"data": {"id": "2742785", "source": "2741867", "target": "2741861", "SCORE": 0.7493725349594643, "shared_interaction": "Direct", "source_original": "18", "Name_B": "GO:0009199", "Name_A": "GO:0009150", "target_original": "16", "Gene_B": "GO:0009199", "interaction": "Direct", "SUID": 2742785, "TYPE": "Direct", "Gene_A": "GO:0009150", "BEND_MAP_ID": 2742785, "selected": false}, "selected": false}, {"data": {"id": "2742728", "source": "2741867", "target": "2741858", "SCORE": 0.7493725349594643, "shared_interaction": "Direct", "source_original": "18", "Name_B": "GO:0009201", "Name_A": "GO:0009150", "target_original": "15", "Gene_B": "GO:0009201", "interaction": "Direct", "SUID": 2742728, "TYPE": "Direct", "Gene_A": "GO:0009150", "BEND_MAP_ID": 2742728, "selected": false}, "selected": false}, {"data": {"id": "2742692", "source": "2741867", "target": "2742119", "SCORE": 0.32815552041743346, "shared_interaction": "Direct", "source_original": "18", "Name_B": "sce01100", "Name_A": "GO:0009150", "target_original": "102", "Gene_B": "sce01100", "interaction": "Direct", "SUID": 2742692, "TYPE": "Direct", "Gene_A": "GO:0009150", "BEND_MAP_ID": 2742692, "selected": false}, "selected": false}, {"data": {"id": "2742677", "source": "2741867", "target": "2741852", "SCORE": 0.7493725349594643, "shared_interaction": "Direct", "source_original": "18", "Name_B": "GO:0009205", "Name_A": "GO:0009150", "target_original": "13", "Gene_B": "GO:0009205", "interaction": "Direct", "SUID": 2742677, "TYPE": "Direct", "Gene_A": "GO:0009150", "BEND_MAP_ID": 2742677, "selected": false}, "selected": false}, {"data": {"id": "2742647", "source": "2741867", "target": "2741849", "SCORE": 0.7493725349594643, "shared_interaction": "Direct", "source_original": "18", "Name_B": "GO:0009206", "Name_A": "GO:0009150", "target_original": "12", "Gene_B": "GO:0009206", "interaction": "Direct", "SUID": 2742647, "TYPE": "Direct", "Gene_A": "GO:0009150", "BEND_MAP_ID": 2742647, "selected": false}, "selected": false}, {"data": {"id": "2742599", "source": "2741867", "target": "2741837", "SCORE": 0.38663156690102896, "shared_interaction": "Direct", "source_original": "18", "Name_B": "R-SCE-163200", "Name_A": "GO:0009150", "target_original": "8", "Gene_B": "R-SCE-163200", "interaction": "Direct", "SUID": 2742599, "TYPE": "Direct", "Gene_A": "GO:0009150", "BEND_MAP_ID": 2742599, "selected": false}, "selected": false}, {"data": {"id": "2742560", "source": "2741867", "target": "2741843", "SCORE": 0.7493725349594643, "shared_interaction": "Direct", "source_original": "18", "Name_B": "GO:0015986", "Name_A": "GO:0009150", "target_original": "10", "Gene_B": "GO:0015986", "interaction": "Direct", "SUID": 2742560, "TYPE": "Direct", "Gene_A": "GO:0009150", "BEND_MAP_ID": 2742560, "selected": false}, "selected": false}, {"data": {"id": "2742467", "source": "2741867", "target": "2741831", "SCORE": 0.39613237639693694, "shared_interaction": "Direct", "source_original": "18", "Name_B": "R-SCE-1428517", "Name_A": "GO:0009150", "target_original": "6", "Gene_B": "R-SCE-1428517", "interaction": "Direct", "SUID": 2742467, "TYPE": "Direct", "Gene_A": "GO:0009150", "BEND_MAP_ID": 2742467, "selected": false}, "selected": false}, {"data": {"id": "2742344", "source": "2741867", "target": "2741819", "SCORE": 0.4400430927333262, "shared_interaction": "Direct", "source_original": "18", "Name_B": "sce00190", "Name_A": "GO:0009150", "target_original": "2", "Gene_B": "sce00190", "interaction": "Direct", "SUID": 2742344, "TYPE": "Direct", "Gene_A": "GO:0009150", "BEND_MAP_ID": 2742344, "selected": false}, "selected": false}, {"data": {"id": "2742254", "source": "2741867", "target": "2742113", "SCORE": 0.6054495073898702, "shared_interaction": "Direct", "source_original": "18", "Name_B": "GO:0046034", "Name_A": "GO:0009150", "target_original": "100", "Gene_B": "GO:0046034", "interaction": "Direct", "SUID": 2742254, "TYPE": "Direct", "Gene_A": "GO:0009150", "BEND_MAP_ID": 2742254, "selected": false}, "selected": false}, {"data": {"id": "2742815", "source": "2741864", "target": "2742164", "SCORE": 0.3075150118284791, "shared_interaction": "Direct", "source_original": "17", "Name_B": "GO:0044281", "Name_A": "GO:0009142", "target_original": "117", "Gene_B": "GO:0044281", "interaction": "Direct", "SUID": 2742815, "TYPE": "Direct", "Gene_A": "GO:0009142", "BEND_MAP_ID": 2742815, "selected": false}, "selected": false}, {"data": {"id": "2742806", "source": "2741864", "target": "2741867", "SCORE": 0.7493725349594643, "shared_interaction": "Direct", "source_original": "17", "Name_B": "GO:0009150", "Name_A": "GO:0009142", "target_original": "18", "Gene_B": "GO:0009150", "interaction": "Direct", "SUID": 2742806, "TYPE": "Direct", "Gene_A": "GO:0009142", "BEND_MAP_ID": 2742806, "selected": false}, "selected": false}, {"data": {"id": "2742782", "source": "2741864", "target": "2741861", "SCORE": 1.0, "shared_interaction": "Direct", "source_original": "17", "Name_B": "GO:0009199", "Name_A": "GO:0009142", "target_original": "16", "Gene_B": "GO:0009199", "interaction": "Direct", "SUID": 2742782, "TYPE": "Direct", "Gene_A": "GO:0009142", "BEND_MAP_ID": 2742782, "selected": false}, "selected": false}, {"data": {"id": "2742725", "source": "2741864", "target": "2741858", "SCORE": 1.0, "shared_interaction": "Direct", "source_original": "17", "Name_B": "GO:0009201", "Name_A": "GO:0009142", "target_original": "15", "Gene_B": "GO:0009201", "interaction": "Direct", "SUID": 2742725, "TYPE": "Direct", "Gene_A": "GO:0009142", "BEND_MAP_ID": 2742725, "selected": false}, "selected": false}, {"data": {"id": "2742707", "source": "2741864", "target": "2741855", "SCORE": 1.0, "shared_interaction": "Direct", "source_original": "17", "Name_B": "GO:0009144", "Name_A": "GO:0009142", "target_original": "14", "Gene_B": "GO:0009144", "interaction": "Direct", "SUID": 2742707, "TYPE": "Direct", "Gene_A": "GO:0009142", "BEND_MAP_ID": 2742707, "selected": false}, "selected": false}, {"data": {"id": "2742674", "source": "2741864", "target": "2741852", "SCORE": 1.0, "shared_interaction": "Direct", "source_original": "17", "Name_B": "GO:0009205", "Name_A": "GO:0009142", "target_original": "13", "Gene_B": "GO:0009205", "interaction": "Direct", "SUID": 2742674, "TYPE": "Direct", "Gene_A": "GO:0009142", "BEND_MAP_ID": 2742674, "selected": false}, "selected": false}, {"data": {"id": "2742644", "source": "2741864", "target": "2741849", "SCORE": 1.0, "shared_interaction": "Direct", "source_original": "17", "Name_B": "GO:0009206", "Name_A": "GO:0009142", "target_original": "12", "Gene_B": "GO:0009206", "interaction": "Direct", "SUID": 2742644, "TYPE": "Direct", "Gene_A": "GO:0009142", "BEND_MAP_ID": 2742644, "selected": false}, "selected": false}, {"data": {"id": "2742617", "source": "2741864", "target": "2741846", "SCORE": 1.0, "shared_interaction": "Direct", "source_original": "17", "Name_B": "GO:0009145", "Name_A": "GO:0009142", "target_original": "11", "Gene_B": "GO:0009145", "interaction": "Direct", "SUID": 2742617, "TYPE": "Direct", "Gene_A": "GO:0009142", "BEND_MAP_ID": 2742617, "selected": false}, "selected": false}, {"data": {"id": "2742596", "source": "2741864", "target": "2741837", "SCORE": 0.5303771955097815, "shared_interaction": "Direct", "source_original": "17", "Name_B": "R-SCE-163200", "Name_A": "GO:0009142", "target_original": "8", "Gene_B": "R-SCE-163200", "interaction": "Direct", "SUID": 2742596, "TYPE": "Direct", "Gene_A": "GO:0009142", "BEND_MAP_ID": 2742596, "selected": false}, "selected": false}, {"data": {"id": "2742557", "source": "2741864", "target": "2741843", "SCORE": 1.0, "shared_interaction": "Direct", "source_original": "17", "Name_B": "GO:0015986", "Name_A": "GO:0009142", "target_original": "10", "Gene_B": "GO:0015986", "interaction": "Direct", "SUID": 2742557, "TYPE": "Direct", "Gene_A": "GO:0009142", "BEND_MAP_ID": 2742557, "selected": false}, "selected": false}, {"data": {"id": "2742464", "source": "2741864", "target": "2741831", "SCORE": 0.41520258157208373, "shared_interaction": "Direct", "source_original": "17", "Name_B": "R-SCE-1428517", "Name_A": "GO:0009142", "target_original": "6", "Gene_B": "R-SCE-1428517", "interaction": "Direct", "SUID": 2742464, "TYPE": "Direct", "Gene_A": "GO:0009142", "BEND_MAP_ID": 2742464, "selected": false}, "selected": false}, {"data": {"id": "2742341", "source": "2741864", "target": "2741819", "SCORE": 0.5603773584915323, "shared_interaction": "Direct", "source_original": "17", "Name_B": "sce00190", "Name_A": "GO:0009142", "target_original": "2", "Gene_B": "sce00190", "interaction": "Direct", "SUID": 2742341, "TYPE": "Direct", "Gene_A": "GO:0009142", "BEND_MAP_ID": 2742341, "selected": false}, "selected": false}, {"data": {"id": "2742251", "source": "2741864", "target": "2742113", "SCORE": 0.49849332759463505, "shared_interaction": "Direct", "source_original": "17", "Name_B": "GO:0046034", "Name_A": "GO:0009142", "target_original": "100", "Gene_B": "GO:0046034", "interaction": "Direct", "SUID": 2742251, "TYPE": "Direct", "Gene_A": "GO:0009142", "BEND_MAP_ID": 2742251, "selected": false}, "selected": false}, {"data": {"id": "2742794", "source": "2741861", "target": "2742164", "SCORE": 0.3075150118284791, "shared_interaction": "Direct", "source_original": "16", "Name_B": "GO:0044281", "Name_A": "GO:0009199", "target_original": "117", "Gene_B": "GO:0044281", "interaction": "Direct", "SUID": 2742794, "TYPE": "Direct", "Gene_A": "GO:0009199", "BEND_MAP_ID": 2742794, "selected": false}, "selected": false}, {"data": {"id": "2742722", "source": "2741861", "target": "2741858", "SCORE": 1.0, "shared_interaction": "Direct", "source_original": "16", "Name_B": "GO:0009201", "Name_A": "GO:0009199", "target_original": "15", "Gene_B": "GO:0009201", "interaction": "Direct", "SUID": 2742722, "TYPE": "Direct", "Gene_A": "GO:0009199", "BEND_MAP_ID": 2742722, "selected": false}, "selected": false}, {"data": {"id": "2742671", "source": "2741861", "target": "2741852", "SCORE": 1.0, "shared_interaction": "Direct", "source_original": "16", "Name_B": "GO:0009205", "Name_A": "GO:0009199", "target_original": "13", "Gene_B": "GO:0009205", "interaction": "Direct", "SUID": 2742671, "TYPE": "Direct", "Gene_A": "GO:0009199", "BEND_MAP_ID": 2742671, "selected": false}, "selected": false}, {"data": {"id": "2742641", "source": "2741861", "target": "2741849", "SCORE": 1.0, "shared_interaction": "Direct", "source_original": "16", "Name_B": "GO:0009206", "Name_A": "GO:0009199", "target_original": "12", "Gene_B": "GO:0009206", "interaction": "Direct", "SUID": 2742641, "TYPE": "Direct", "Gene_A": "GO:0009199", "BEND_MAP_ID": 2742641, "selected": false}, "selected": false}, {"data": {"id": "2742590", "source": "2741861", "target": "2741837", "SCORE": 0.5303771955097815, "shared_interaction": "Direct", "source_original": "16", "Name_B": "R-SCE-163200", "Name_A": "GO:0009199", "target_original": "8", "Gene_B": "R-SCE-163200", "interaction": "Direct", "SUID": 2742590, "TYPE": "Direct", "Gene_A": "GO:0009199", "BEND_MAP_ID": 2742590, "selected": false}, "selected": false}, {"data": {"id": "2742554", "source": "2741861", "target": "2741843", "SCORE": 1.0, "shared_interaction": "Direct", "source_original": "16", "Name_B": "GO:0015986", "Name_A": "GO:0009199", "target_original": "10", "Gene_B": "GO:0015986", "interaction": "Direct", "SUID": 2742554, "TYPE": "Direct", "Gene_A": "GO:0009199", "BEND_MAP_ID": 2742554, "selected": false}, "selected": false}, {"data": {"id": "2742458", "source": "2741861", "target": "2741831", "SCORE": 0.41520258157208373, "shared_interaction": "Direct", "source_original": "16", "Name_B": "R-SCE-1428517", "Name_A": "GO:0009199", "target_original": "6", "Gene_B": "R-SCE-1428517", "interaction": "Direct", "SUID": 2742458, "TYPE": "Direct", "Gene_A": "GO:0009199", "BEND_MAP_ID": 2742458, "selected": false}, "selected": false}, {"data": {"id": "2742338", "source": "2741861", "target": "2741819", "SCORE": 0.5603773584915323, "shared_interaction": "Direct", "source_original": "16", "Name_B": "sce00190", "Name_A": "GO:0009199", "target_original": "2", "Gene_B": "sce00190", "interaction": "Direct", "SUID": 2742338, "TYPE": "Direct", "Gene_A": "GO:0009199", "BEND_MAP_ID": 2742338, "selected": false}, "selected": false}, {"data": {"id": "2742245", "source": "2741861", "target": "2742113", "SCORE": 0.49849332759463505, "shared_interaction": "Direct", "source_original": "16", "Name_B": "GO:0046034", "Name_A": "GO:0009199", "target_original": "100", "Gene_B": "GO:0046034", "interaction": "Direct", "SUID": 2742245, "TYPE": "Direct", "Gene_A": "GO:0009199", "BEND_MAP_ID": 2742245, "selected": false}, "selected": false}, {"data": {"id": "2742737", "source": "2741858", "target": "2742164", "SCORE": 0.3075150118284791, "shared_interaction": "Direct", "source_original": "15", "Name_B": "GO:0044281", "Name_A": "GO:0009201", "target_original": "117", "Gene_B": "GO:0044281", "interaction": "Direct", "SUID": 2742737, "TYPE": "Direct", "Gene_A": "GO:0009201", "BEND_MAP_ID": 2742737, "selected": false}, "selected": false}, {"data": {"id": "2742668", "source": "2741858", "target": "2741852", "SCORE": 1.0, "shared_interaction": "Direct", "source_original": "15", "Name_B": "GO:0009205", "Name_A": "GO:0009201", "target_original": "13", "Gene_B": "GO:0009205", "interaction": "Direct", "SUID": 2742668, "TYPE": "Direct", "Gene_A": "GO:0009201", "BEND_MAP_ID": 2742668, "selected": false}, "selected": false}, {"data": {"id": "2742638", "source": "2741858", "target": "2741849", "SCORE": 1.0, "shared_interaction": "Direct", "source_original": "15", "Name_B": "GO:0009206", "Name_A": "GO:0009201", "target_original": "12", "Gene_B": "GO:0009206", "interaction": "Direct", "SUID": 2742638, "TYPE": "Direct", "Gene_A": "GO:0009201", "BEND_MAP_ID": 2742638, "selected": false}, "selected": false}, {"data": {"id": "2742587", "source": "2741858", "target": "2741837", "SCORE": 0.5303771955097815, "shared_interaction": "Direct", "source_original": "15", "Name_B": "R-SCE-163200", "Name_A": "GO:0009201", "target_original": "8", "Gene_B": "R-SCE-163200", "interaction": "Direct", "SUID": 2742587, "TYPE": "Direct", "Gene_A": "GO:0009201", "BEND_MAP_ID": 2742587, "selected": false}, "selected": false}, {"data": {"id": "2742551", "source": "2741858", "target": "2741843", "SCORE": 1.0, "shared_interaction": "Direct", "source_original": "15", "Name_B": "GO:0015986", "Name_A": "GO:0009201", "target_original": "10", "Gene_B": "GO:0015986", "interaction": "Direct", "SUID": 2742551, "TYPE": "Direct", "Gene_A": "GO:0009201", "BEND_MAP_ID": 2742551, "selected": false}, "selected": false}, {"data": {"id": "2742455", "source": "2741858", "target": "2741831", "SCORE": 0.41520258157208373, "shared_interaction": "Direct", "source_original": "15", "Name_B": "R-SCE-1428517", "Name_A": "GO:0009201", "target_original": "6", "Gene_B": "R-SCE-1428517", "interaction": "Direct", "SUID": 2742455, "TYPE": "Direct", "Gene_A": "GO:0009201", "BEND_MAP_ID": 2742455, "selected": false}, "selected": false}, {"data": {"id": "2742335", "source": "2741858", "target": "2741819", "SCORE": 0.5603773584915323, "shared_interaction": "Direct", "source_original": "15", "Name_B": "sce00190", "Name_A": "GO:0009201", "target_original": "2", "Gene_B": "sce00190", "interaction": "Direct", "SUID": 2742335, "TYPE": "Direct", "Gene_A": "GO:0009201", "BEND_MAP_ID": 2742335, "selected": false}, "selected": false}, {"data": {"id": "2742242", "source": "2741858", "target": "2742113", "SCORE": 0.49849332759463505, "shared_interaction": "Direct", "source_original": "15", "Name_B": "GO:0046034", "Name_A": "GO:0009201", "target_original": "100", "Gene_B": "GO:0046034", "interaction": "Direct", "SUID": 2742242, "TYPE": "Direct", "Gene_A": "GO:0009201", "BEND_MAP_ID": 2742242, "selected": false}, "selected": false}, {"data": {"id": "2742719", "source": "2741855", "target": "2742164", "SCORE": 0.3075150118284791, "shared_interaction": "Direct", "source_original": "14", "Name_B": "GO:0044281", "Name_A": "GO:0009144", "target_original": "117", "Gene_B": "GO:0044281", "interaction": "Direct", "SUID": 2742719, "TYPE": "Direct", "Gene_A": "GO:0009144", "BEND_MAP_ID": 2742719, "selected": false}, "selected": false}, {"data": {"id": "2742710", "source": "2741855", "target": "2741867", "SCORE": 0.7493725349594643, "shared_interaction": "Direct", "source_original": "14", "Name_B": "GO:0009150", "Name_A": "GO:0009144", "target_original": "18", "Gene_B": "GO:0009150", "interaction": "Direct", "SUID": 2742710, "TYPE": "Direct", "Gene_A": "GO:0009144", "BEND_MAP_ID": 2742710, "selected": false}, "selected": false}, {"data": {"id": "2742704", "source": "2741855", "target": "2741861", "SCORE": 1.0, "shared_interaction": "Direct", "source_original": "14", "Name_B": "GO:0009199", "Name_A": "GO:0009144", "target_original": "16", "Gene_B": "GO:0009199", "interaction": "Direct", "SUID": 2742704, "TYPE": "Direct", "Gene_A": "GO:0009144", "BEND_MAP_ID": 2742704, "selected": false}, "selected": false}, {"data": {"id": "2742701", "source": "2741855", "target": "2741858", "SCORE": 1.0, "shared_interaction": "Direct", "source_original": "14", "Name_B": "GO:0009201", "Name_A": "GO:0009144", "target_original": "15", "Gene_B": "GO:0009201", "interaction": "Direct", "SUID": 2742701, "TYPE": "Direct", "Gene_A": "GO:0009144", "BEND_MAP_ID": 2742701, "selected": false}, "selected": false}, {"data": {"id": "2742665", "source": "2741855", "target": "2741852", "SCORE": 1.0, "shared_interaction": "Direct", "source_original": "14", "Name_B": "GO:0009205", "Name_A": "GO:0009144", "target_original": "13", "Gene_B": "GO:0009205", "interaction": "Direct", "SUID": 2742665, "TYPE": "Direct", "Gene_A": "GO:0009144", "BEND_MAP_ID": 2742665, "selected": false}, "selected": false}, {"data": {"id": "2742635", "source": "2741855", "target": "2741849", "SCORE": 1.0, "shared_interaction": "Direct", "source_original": "14", "Name_B": "GO:0009206", "Name_A": "GO:0009144", "target_original": "12", "Gene_B": "GO:0009206", "interaction": "Direct", "SUID": 2742635, "TYPE": "Direct", "Gene_A": "GO:0009144", "BEND_MAP_ID": 2742635, "selected": false}, "selected": false}, {"data": {"id": "2742608", "source": "2741855", "target": "2741846", "SCORE": 1.0, "shared_interaction": "Direct", "source_original": "14", "Name_B": "GO:0009145", "Name_A": "GO:0009144", "target_original": "11", "Gene_B": "GO:0009145", "interaction": "Direct", "SUID": 2742608, "TYPE": "Direct", "Gene_A": "GO:0009144", "BEND_MAP_ID": 2742608, "selected": false}, "selected": false}, {"data": {"id": "2742584", "source": "2741855", "target": "2741837", "SCORE": 0.5303771955097815, "shared_interaction": "Direct", "source_original": "14", "Name_B": "R-SCE-163200", "Name_A": "GO:0009144", "target_original": "8", "Gene_B": "R-SCE-163200", "interaction": "Direct", "SUID": 2742584, "TYPE": "Direct", "Gene_A": "GO:0009144", "BEND_MAP_ID": 2742584, "selected": false}, "selected": false}, {"data": {"id": "2742548", "source": "2741855", "target": "2741843", "SCORE": 1.0, "shared_interaction": "Direct", "source_original": "14", "Name_B": "GO:0015986", "Name_A": "GO:0009144", "target_original": "10", "Gene_B": "GO:0015986", "interaction": "Direct", "SUID": 2742548, "TYPE": "Direct", "Gene_A": "GO:0009144", "BEND_MAP_ID": 2742548, "selected": false}, "selected": false}, {"data": {"id": "2742452", "source": "2741855", "target": "2741831", "SCORE": 0.41520258157208373, "shared_interaction": "Direct", "source_original": "14", "Name_B": "R-SCE-1428517", "Name_A": "GO:0009144", "target_original": "6", "Gene_B": "R-SCE-1428517", "interaction": "Direct", "SUID": 2742452, "TYPE": "Direct", "Gene_A": "GO:0009144", "BEND_MAP_ID": 2742452, "selected": false}, "selected": false}, {"data": {"id": "2742332", "source": "2741855", "target": "2741819", "SCORE": 0.5603773584915323, "shared_interaction": "Direct", "source_original": "14", "Name_B": "sce00190", "Name_A": "GO:0009144", "target_original": "2", "Gene_B": "sce00190", "interaction": "Direct", "SUID": 2742332, "TYPE": "Direct", "Gene_A": "GO:0009144", "BEND_MAP_ID": 2742332, "selected": false}, "selected": false}, {"data": {"id": "2742239", "source": "2741855", "target": "2742113", "SCORE": 0.49849332759463505, "shared_interaction": "Direct", "source_original": "14", "Name_B": "GO:0046034", "Name_A": "GO:0009144", "target_original": "100", "Gene_B": "GO:0046034", "interaction": "Direct", "SUID": 2742239, "TYPE": "Direct", "Gene_A": "GO:0009144", "BEND_MAP_ID": 2742239, "selected": false}, "selected": false}, {"data": {"id": "2742686", "source": "2741852", "target": "2742164", "SCORE": 0.3075150118284791, "shared_interaction": "Direct", "source_original": "13", "Name_B": "GO:0044281", "Name_A": "GO:0009205", "target_original": "117", "Gene_B": "GO:0044281", "interaction": "Direct", "SUID": 2742686, "TYPE": "Direct", "Gene_A": "GO:0009205", "BEND_MAP_ID": 2742686, "selected": false}, "selected": false}, {"data": {"id": "2742632", "source": "2741852", "target": "2741849", "SCORE": 1.0, "shared_interaction": "Direct", "source_original": "13", "Name_B": "GO:0009206", "Name_A": "GO:0009205", "target_original": "12", "Gene_B": "GO:0009206", "interaction": "Direct", "SUID": 2742632, "TYPE": "Direct", "Gene_A": "GO:0009205", "BEND_MAP_ID": 2742632, "selected": false}, "selected": false}, {"data": {"id": "2742581", "source": "2741852", "target": "2741837", "SCORE": 0.5303771955097815, "shared_interaction": "Direct", "source_original": "13", "Name_B": "R-SCE-163200", "Name_A": "GO:0009205", "target_original": "8", "Gene_B": "R-SCE-163200", "interaction": "Direct", "SUID": 2742581, "TYPE": "Direct", "Gene_A": "GO:0009205", "BEND_MAP_ID": 2742581, "selected": false}, "selected": false}, {"data": {"id": "2742545", "source": "2741852", "target": "2741843", "SCORE": 1.0, "shared_interaction": "Direct", "source_original": "13", "Name_B": "GO:0015986", "Name_A": "GO:0009205", "target_original": "10", "Gene_B": "GO:0015986", "interaction": "Direct", "SUID": 2742545, "TYPE": "Direct", "Gene_A": "GO:0009205", "BEND_MAP_ID": 2742545, "selected": false}, "selected": false}, {"data": {"id": "2742446", "source": "2741852", "target": "2741831", "SCORE": 0.41520258157208373, "shared_interaction": "Direct", "source_original": "13", "Name_B": "R-SCE-1428517", "Name_A": "GO:0009205", "target_original": "6", "Gene_B": "R-SCE-1428517", "interaction": "Direct", "SUID": 2742446, "TYPE": "Direct", "Gene_A": "GO:0009205", "BEND_MAP_ID": 2742446, "selected": false}, "selected": false}, {"data": {"id": "2742326", "source": "2741852", "target": "2741819", "SCORE": 0.5603773584915323, "shared_interaction": "Direct", "source_original": "13", "Name_B": "sce00190", "Name_A": "GO:0009205", "target_original": "2", "Gene_B": "sce00190", "interaction": "Direct", "SUID": 2742326, "TYPE": "Direct", "Gene_A": "GO:0009205", "BEND_MAP_ID": 2742326, "selected": false}, "selected": false}, {"data": {"id": "2742233", "source": "2741852", "target": "2742113", "SCORE": 0.49849332759463505, "shared_interaction": "Direct", "source_original": "13", "Name_B": "GO:0046034", "Name_A": "GO:0009205", "target_original": "100", "Gene_B": "GO:0046034", "interaction": "Direct", "SUID": 2742233, "TYPE": "Direct", "Gene_A": "GO:0009205", "BEND_MAP_ID": 2742233, "selected": false}, "selected": false}, {"data": {"id": "2742656", "source": "2741849", "target": "2742164", "SCORE": 0.3075150118284791, "shared_interaction": "Direct", "source_original": "12", "Name_B": "GO:0044281", "Name_A": "GO:0009206", "target_original": "117", "Gene_B": "GO:0044281", "interaction": "Direct", "SUID": 2742656, "TYPE": "Direct", "Gene_A": "GO:0009206", "BEND_MAP_ID": 2742656, "selected": false}, "selected": false}, {"data": {"id": "2742575", "source": "2741849", "target": "2741837", "SCORE": 0.5303771955097815, "shared_interaction": "Direct", "source_original": "12", "Name_B": "R-SCE-163200", "Name_A": "GO:0009206", "target_original": "8", "Gene_B": "R-SCE-163200", "interaction": "Direct", "SUID": 2742575, "TYPE": "Direct", "Gene_A": "GO:0009206", "BEND_MAP_ID": 2742575, "selected": false}, "selected": false}, {"data": {"id": "2742542", "source": "2741849", "target": "2741843", "SCORE": 1.0, "shared_interaction": "Direct", "source_original": "12", "Name_B": "GO:0015986", "Name_A": "GO:0009206", "target_original": "10", "Gene_B": "GO:0015986", "interaction": "Direct", "SUID": 2742542, "TYPE": "Direct", "Gene_A": "GO:0009206", "BEND_MAP_ID": 2742542, "selected": false}, "selected": false}, {"data": {"id": "2742440", "source": "2741849", "target": "2741831", "SCORE": 0.41520258157208373, "shared_interaction": "Direct", "source_original": "12", "Name_B": "R-SCE-1428517", "Name_A": "GO:0009206", "target_original": "6", "Gene_B": "R-SCE-1428517", "interaction": "Direct", "SUID": 2742440, "TYPE": "Direct", "Gene_A": "GO:0009206", "BEND_MAP_ID": 2742440, "selected": false}, "selected": false}, {"data": {"id": "2742320", "source": "2741849", "target": "2741819", "SCORE": 0.5603773584915323, "shared_interaction": "Direct", "source_original": "12", "Name_B": "sce00190", "Name_A": "GO:0009206", "target_original": "2", "Gene_B": "sce00190", "interaction": "Direct", "SUID": 2742320, "TYPE": "Direct", "Gene_A": "GO:0009206", "BEND_MAP_ID": 2742320, "selected": false}, "selected": false}, {"data": {"id": "2742227", "source": "2741849", "target": "2742113", "SCORE": 0.49849332759463505, "shared_interaction": "Direct", "source_original": "12", "Name_B": "GO:0046034", "Name_A": "GO:0009206", "target_original": "100", "Gene_B": "GO:0046034", "interaction": "Direct", "SUID": 2742227, "TYPE": "Direct", "Gene_A": "GO:0009206", "BEND_MAP_ID": 2742227, "selected": false}, "selected": false}, {"data": {"id": "2742629", "source": "2741846", "target": "2742164", "SCORE": 0.3075150118284791, "shared_interaction": "Direct", "source_original": "11", "Name_B": "GO:0044281", "Name_A": "GO:0009145", "target_original": "117", "Gene_B": "GO:0044281", "interaction": "Direct", "SUID": 2742629, "TYPE": "Direct", "Gene_A": "GO:0009145", "BEND_MAP_ID": 2742629, "selected": false}, "selected": false}, {"data": {"id": "2742620", "source": "2741846", "target": "2741867", "SCORE": 0.7493725349594643, "shared_interaction": "Direct", "source_original": "11", "Name_B": "GO:0009150", "Name_A": "GO:0009145", "target_original": "18", "Gene_B": "GO:0009150", "interaction": "Direct", "SUID": 2742620, "TYPE": "Direct", "Gene_A": "GO:0009145", "BEND_MAP_ID": 2742620, "selected": false}, "selected": false}, {"data": {"id": "2742614", "source": "2741846", "target": "2741861", "SCORE": 1.0, "shared_interaction": "Direct", "source_original": "11", "Name_B": "GO:0009199", "Name_A": "GO:0009145", "target_original": "16", "Gene_B": "GO:0009199", "interaction": "Direct", "SUID": 2742614, "TYPE": "Direct", "Gene_A": "GO:0009145", "BEND_MAP_ID": 2742614, "selected": false}, "selected": false}, {"data": {"id": "2742611", "source": "2741846", "target": "2741858", "SCORE": 1.0, "shared_interaction": "Direct", "source_original": "11", "Name_B": "GO:0009201", "Name_A": "GO:0009145", "target_original": "15", "Gene_B": "GO:0009201", "interaction": "Direct", "SUID": 2742611, "TYPE": "Direct", "Gene_A": "GO:0009145", "BEND_MAP_ID": 2742611, "selected": false}, "selected": false}, {"data": {"id": "2742605", "source": "2741846", "target": "2741852", "SCORE": 1.0, "shared_interaction": "Direct", "source_original": "11", "Name_B": "GO:0009205", "Name_A": "GO:0009145", "target_original": "13", "Gene_B": "GO:0009205", "interaction": "Direct", "SUID": 2742605, "TYPE": "Direct", "Gene_A": "GO:0009145", "BEND_MAP_ID": 2742605, "selected": false}, "selected": false}, {"data": {"id": "2742602", "source": "2741846", "target": "2741849", "SCORE": 1.0, "shared_interaction": "Direct", "source_original": "11", "Name_B": "GO:0009206", "Name_A": "GO:0009145", "target_original": "12", "Gene_B": "GO:0009206", "interaction": "Direct", "SUID": 2742602, "TYPE": "Direct", "Gene_A": "GO:0009145", "BEND_MAP_ID": 2742602, "selected": false}, "selected": false}, {"data": {"id": "2742572", "source": "2741846", "target": "2741837", "SCORE": 0.5303771955097815, "shared_interaction": "Direct", "source_original": "11", "Name_B": "R-SCE-163200", "Name_A": "GO:0009145", "target_original": "8", "Gene_B": "R-SCE-163200", "interaction": "Direct", "SUID": 2742572, "TYPE": "Direct", "Gene_A": "GO:0009145", "BEND_MAP_ID": 2742572, "selected": false}, "selected": false}, {"data": {"id": "2742539", "source": "2741846", "target": "2741843", "SCORE": 1.0, "shared_interaction": "Direct", "source_original": "11", "Name_B": "GO:0015986", "Name_A": "GO:0009145", "target_original": "10", "Gene_B": "GO:0015986", "interaction": "Direct", "SUID": 2742539, "TYPE": "Direct", "Gene_A": "GO:0009145", "BEND_MAP_ID": 2742539, "selected": false}, "selected": false}, {"data": {"id": "2742437", "source": "2741846", "target": "2741831", "SCORE": 0.41520258157208373, "shared_interaction": "Direct", "source_original": "11", "Name_B": "R-SCE-1428517", "Name_A": "GO:0009145", "target_original": "6", "Gene_B": "R-SCE-1428517", "interaction": "Direct", "SUID": 2742437, "TYPE": "Direct", "Gene_A": "GO:0009145", "BEND_MAP_ID": 2742437, "selected": false}, "selected": false}, {"data": {"id": "2742317", "source": "2741846", "target": "2741819", "SCORE": 0.5603773584915323, "shared_interaction": "Direct", "source_original": "11", "Name_B": "sce00190", "Name_A": "GO:0009145", "target_original": "2", "Gene_B": "sce00190", "interaction": "Direct", "SUID": 2742317, "TYPE": "Direct", "Gene_A": "GO:0009145", "BEND_MAP_ID": 2742317, "selected": false}, "selected": false}, {"data": {"id": "2742224", "source": "2741846", "target": "2742113", "SCORE": 0.49849332759463505, "shared_interaction": "Direct", "source_original": "11", "Name_B": "GO:0046034", "Name_A": "GO:0009145", "target_original": "100", "Gene_B": "GO:0046034", "interaction": "Direct", "SUID": 2742224, "TYPE": "Direct", "Gene_A": "GO:0009145", "BEND_MAP_ID": 2742224, "selected": false}, "selected": false}, {"data": {"id": "2742569", "source": "2741843", "target": "2742164", "SCORE": 0.3075150118284791, "shared_interaction": "Direct", "source_original": "10", "Name_B": "GO:0044281", "Name_A": "GO:0015986", "target_original": "117", "Gene_B": "GO:0044281", "interaction": "Direct", "SUID": 2742569, "TYPE": "Direct", "Gene_A": "GO:0015986", "BEND_MAP_ID": 2742569, "selected": false}, "selected": false}, {"data": {"id": "2742536", "source": "2741843", "target": "2741837", "SCORE": 0.5303771955097815, "shared_interaction": "Direct", "source_original": "10", "Name_B": "R-SCE-163200", "Name_A": "GO:0015986", "target_original": "8", "Gene_B": "R-SCE-163200", "interaction": "Direct", "SUID": 2742536, "TYPE": "Direct", "Gene_A": "GO:0015986", "BEND_MAP_ID": 2742536, "selected": false}, "selected": false}, {"data": {"id": "2742431", "source": "2741843", "target": "2741831", "SCORE": 0.41520258157208373, "shared_interaction": "Direct", "source_original": "10", "Name_B": "R-SCE-1428517", "Name_A": "GO:0015986", "target_original": "6", "Gene_B": "R-SCE-1428517", "interaction": "Direct", "SUID": 2742431, "TYPE": "Direct", "Gene_A": "GO:0015986", "BEND_MAP_ID": 2742431, "selected": false}, "selected": false}, {"data": {"id": "2742311", "source": "2741843", "target": "2741819", "SCORE": 0.5603773584915323, "shared_interaction": "Direct", "source_original": "10", "Name_B": "sce00190", "Name_A": "GO:0015986", "target_original": "2", "Gene_B": "sce00190", "interaction": "Direct", "SUID": 2742311, "TYPE": "Direct", "Gene_A": "GO:0015986", "BEND_MAP_ID": 2742311, "selected": false}, "selected": false}, {"data": {"id": "2742218", "source": "2741843", "target": "2742113", "SCORE": 0.49849332759463505, "shared_interaction": "Direct", "source_original": "10", "Name_B": "GO:0046034", "Name_A": "GO:0015986", "target_original": "100", "Gene_B": "GO:0046034", "interaction": "D
```
