## Additional File 3 for "Uncharacterized yeast gene *YBR238C,* an effector of TORC1 signaling in a mitochondrial feedback loop, accelerates cellular aging via *HAP4*- and *RMD9*-dependent mechanisms": MyList_PPIColorByCluster.pdf

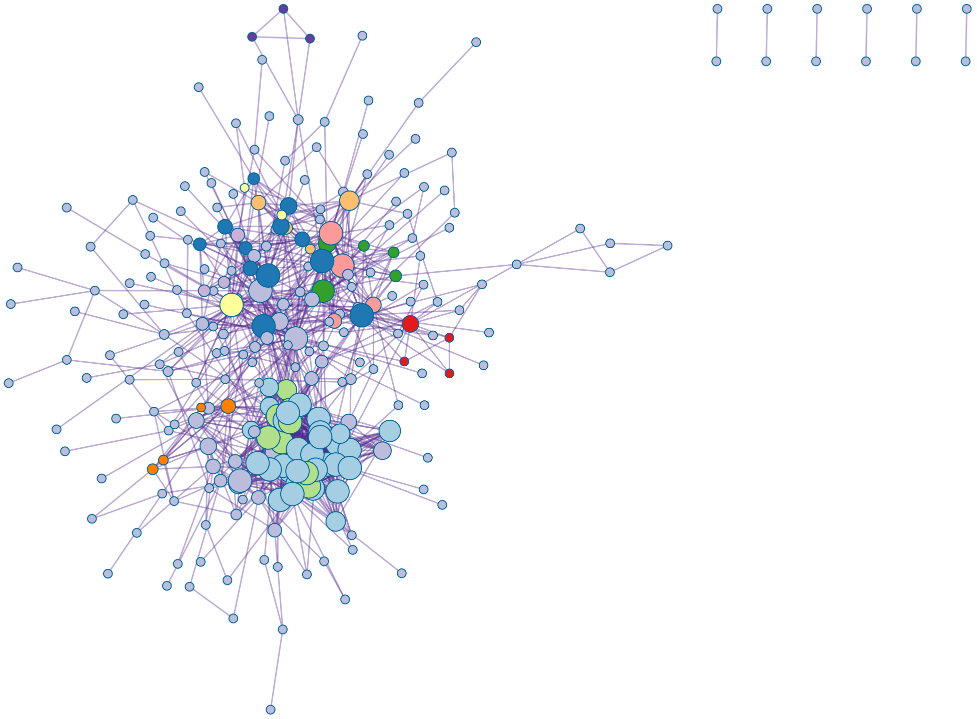

MCODE1
MCODE2
MCODE3
MCODE4
MCODE5
MCODE6
MCODE7
MCODE8
MCODE9
MCODE10
MCODE11

created by
http://metascape.org
