## Additional File 3 for "Uncharacterized yeast gene *YBR238C,* an effector of TORC1 signaling in a mitochondrial feedback loop, accelerates cellular aging via *HAP4*- and *RMD9*-dependent mechanisms": PPINetwork.js

```
var networks={"MyList_MCODE_ALL_PPIColorByCluster": {"format_version": "1.0", "generated_by": "cytoscape-3.9.1", "target_cytoscapejs_version": "~2.1", "data": {"shared_name": "MyList_MCODE_ALL_PPIColorByCluster", "name": "MyList_MCODE_ALL_PPIColorByCluster", "SUID": 2782658, "__Annotations": [], "selected": true}, "elements": {"nodes": [{"data": {"id": "2782953", "shared_name": "856088", "MCODE_SCORE": 1.0, "Gene": "856088", "Symbol": "VTC3", "name": "856088", "MCODE_TYPE": "Clustered", "SUID": 2782953, "DEGREE": 2.0, "id_original": "856088", "MCODE_CLUSTER_ID": 11.0, "selected": false, "canonicalName": "VTC3"}, "position": {"x": 720.8827460328605, "y": 509.0878597849916}, "selected": false}, {"data": {"id": "2782950", "shared_name": "853441", "MCODE_SCORE": 1.0, "Gene": "853441", "Symbol": "VTC4", "name": "853441", "MCODE_TYPE": "Seed", "SUID": 2782950, "DEGREE": 2.0, "id_original": "853441", "MCODE_CLUSTER_ID": 11.0, "selected": false, "canonicalName": "VTC4"}, "position": {"x": 828.1275675813224, "y": 519.7271420115542}, "selected": false}, {"data": {"id": "2782947", "shared_name": "850633", "MCODE_SCORE": 1.0, "Gene": "850633", "Symbol": "HSP104", "name": "850633", "MCODE_TYPE": "Clustered", "SUID": 2782947, "DEGREE": 2.0, "id_original": "850633", "MCODE_CLUSTER_ID": 11.0, "selected": false, "canonicalName": "HSP104"}, "position": {"x": 765.2786448518302, "y": 607.3116757029604}, "selected": false}, {"data": {"id": "2782944", "shared_name": "854228", "MCODE_SCORE": 1.0, "Gene": "854228", "Symbol": "YOR062C", "name": "854228", "MCODE_TYPE": "Clustered", "SUID": 2782944, "DEGREE": 2.0, "id_original": "854228", "MCODE_CLUSTER_ID": 10.0, "selected": false, "canonicalName": "YOR062C"}, "position": {"x": 928.1275675813224, "y": 509.0878597849916}, "selected": false}, {"data": {"id": "2782941", "shared_name": "853950", "MCODE_SCORE": 1.0, "Gene": "853950", "Symbol": "YKR075C", "name": "853950", "MCODE_TYPE": "Clustered", "SUID": 2782941, "DEGREE": 2.0, "id_original": "853950", "MCODE_CLUSTER_ID": 10.0, "selected": false, "canonicalName": "YKR075C"}, "position": {"x": 1035.3723891297843, "y": 519.7271420115542}, "selected": false}, {"data": {"id": "2782938", "shared_name": "852340", "MCODE_SCORE": 1.0, "Gene": "852340", "Symbol": "REG2", "name": "852340", "MCODE_TYPE": "Seed", "SUID": 2782938, "DEGREE": 2.0, "id_original": "852340", "MCODE_CLUSTER_ID": 10.0, "selected": false, "canonicalName": "REG2"}, "position": {"x": 972.5234664002921, "y": 607.3116757029604}, "selected": false}, {"data": {"id": "2782935", "shared_name": "852128", "MCODE_SCORE": 1.0, "Gene": "852128", "Symbol": "EMI2", "name": "852128", "MCODE_TYPE": "Seed", "SUID": 2782935, "DEGREE": 2.0, "id_original": "852128", "MCODE_CLUSTER_ID": 9.0, "selected": false, "canonicalName": "EMI2"}, "position": {"x": -137.71577233872154, "y": 780.1714855784487}, "selected": false}, {"data": {"id": "2782932", "shared_name": "850614", "MCODE_SCORE": 1.0, "Gene": "850614", "Symbol": "HXK1", "name": "850614", "MCODE_TYPE": "Clustered", "SUID": 2782932, "DEGREE": 2.0, "id_original": "850614", "MCODE_CLUSTER_ID": 9.0, "selected": false, "canonicalName": "HXK1"}, "position": {"x": -30.47095079025962, "y": 790.8107678050112}, "selected": false}, {"data": {"id": "2782929", "shared_name": "850317", "MCODE_SCORE": 1.0, "Gene": "850317", "Symbol": "GLK1", "name": "850317", "MCODE_TYPE": "Clustered", "SUID": 2782929, "DEGREE": 2.0, "id_original": "850317", "MCODE_CLUSTER_ID": 9.0, "selected": false, "canonicalName": "GLK1"}, "position": {"x": -93.31987351975181, "y": 878.3953014964175}, "selected": false}, {"data": {"id": "2782926", "shared_name": "855447", "MCODE_SCORE": 1.0, "Gene": "855447", "Symbol": "GOR1", "name": "855447", "MCODE_TYPE": "Seed", "SUID": 2782926, "DEGREE": 2.0, "id_original": "855447", "MCODE_CLUSTER_ID": 8.0, "selected": false, "canonicalName": "GOR1"}, "position": {"x": -137.71577233872154, "y": 575.4371487254214}, "selected": false}, {"data": {"id": "2782923", "shared_name": "854556", "MCODE_SCORE": 1.0, "Gene": "854556", "Symbol": "ALD4", "name": "854556", "MCODE_TYPE": "Clustered", "SUID": 2782923, "DEGREE": 2.0, "id_original": "854556", "MCODE_CLUSTER_ID": 8.0, "selected": false, "canonicalName": "ALD4"}, "position": {"x": -88.39712304672935, "y": 509.0878597849916}, "selected": false}, {"data": {"id": "2782920", "shared_name": "853119", "MCODE_SCORE": 1.0, "Gene": "853119", "Symbol": "TDA10", "name": "853119", "MCODE_TYPE": "Clustered", "SUID": 2782920, "DEGREE": 2.0, "id_original": "853119", "MCODE_CLUSTER_ID": 8.0, "selected": false, "canonicalName": "TDA10"}, "position": {"x": -61.5512368162606, "y": 631.9696656817507}, "selected": false}, {"data": {"id": "2782917", "shared_name": "850514", "MCODE_SCORE": 1.0, "Gene": "850514", "Symbol": "AGX1", "name": "850514", "MCODE_TYPE": "Clustered", "SUID": 2782917, "DEGREE": 2.0, "id_original": "850514", "MCODE_CLUSTER_ID": 8.0, "selected": false, "canonicalName": "AGX1"}, "position": {"x": -12.291928954932473, "y": 565.745593702228}, "selected": false}, {"data": {"id": "2782914", "shared_name": "853522", "MCODE_SCORE": 1.0, "Gene": "853522", "Symbol": "PTK2", "name": "853522", "MCODE_TYPE": "Clustered", "SUID": 2782914, "DEGREE": 2.0, "id_original": "853522", "MCODE_CLUSTER_ID": 7.0, "selected": false, "canonicalName": "PTK2"}, "position": {"x": 87.70807104506753, "y": 509.39265409651506}, "selected": false}, {"data": {"id": "2782911", "shared_name": "853164", "MCODE_SCORE": 1.0, "Gene": "853164", "Symbol": "MGA1", "name": "853164", "MCODE_TYPE": "Clustered", "SUID": 2782911, "DEGREE": 2.0, "id_original": "853164", "MCODE_CLUSTER_ID": 7.0, "selected": false, "canonicalName": "MGA1"}, "position": {"x": 87.78013830580971, "y": 619.483422910602}, "selected": false}, {"data": {"id": "2782908", "shared_name": "852515", "MCODE_SCORE": 1.0, "Gene": "852515", "Symbol": "SDS24", "name": "852515", "MCODE_TYPE": "Clustered", "SUID": 2782908, "DEGREE": 2.0, "id_original": "852515", "MCODE_CLUSTER_ID": 7.0, "selected": false, "canonicalName": "SDS24"}, "position": {"x": 197.7706625978019, "y": 509.0878597849916}, "selected": false}, {"data": {"id": "2782905", "shared_name": "851834", "MCODE_SCORE": 1.0, "Gene": "851834", "Symbol": "VHS1", "name": "851834", "MCODE_TYPE": "Clustered", "SUID": 2782905, "DEGREE": 2.0, "id_original": "851834", "MCODE_CLUSTER_ID": 7.0, "selected": false, "canonicalName": "VHS1"}, "position": {"x": 197.7342856446769, "y": 619.1705376261782}, "selected": false}, {"data": {"id": "2782902", "shared_name": "854447", "MCODE_SCORE": 1.25, "Gene": "854447", "Symbol": "TPO4", "name": "854447", "MCODE_TYPE": "Clustered", "SUID": 2782902, "DEGREE": 2.0, "id_original": "854447", "MCODE_CLUSTER_ID": 6.0, "selected": false, "canonicalName": "TPO4"}, "position": {"x": 304.16923742690346, "y": 509.0878597849916}, "selected": false}, {"data": {"id": "2782899", "shared_name": "853031", "MCODE_SCORE": 1.25, "Gene": "853031", "Symbol": "YGR130C", "name": "853031", "MCODE_TYPE": "Clustered", "SUID": 2782899, "DEGREE": 3.0, "id_original": "853031", "MCODE_CLUSTER_ID": 6.0, "selected": false, "canonicalName": "YGR130C"}, "position": {"x": 297.7706625978019, "y": 617.9175793284487}, "selected": false}, {"data": {"id": "2782896", "shared_name": "852251", "MCODE_SCORE": 1.25, "Gene": "852251", "Symbol": "YBL029C-A", "name": "852251", "MCODE_TYPE": "Seed", "SUID": 2782896, "DEGREE": 2.0, "id_original": "852251", "MCODE_CLUSTER_ID": 6.0, "selected": false, "canonicalName": "YBL029C-A"}, "position": {"x": 387.2258933106925, "y": 680.1714855784487}, "selected": false}, {"data": {"id": "2782893", "shared_name": "850385", "MCODE_SCORE": 1.25, "Gene": "850385", "Symbol": "HSP30", "name": "850385", "MCODE_TYPE": "Clustered", "SUID": 2782893, "DEGREE": 3.0, "id_original": "850385", "MCODE_CLUSTER_ID": 6.0, "selected": false, "canonicalName": "HSP30"}, "position": {"x": 393.6106284181144, "y": 571.3569332713198}, "selected": false}, {"data": {"id": "2782890", "shared_name": "855871", "MCODE_SCORE": 1.4000000000000001, "Gene": "855871", "Symbol": "USV1", "name": "855871", "MCODE_TYPE": "Clustered", "SUID": 2782890, "DEGREE": 2.0, "id_original": "855871", "MCODE_CLUSTER_ID": 4.0, "selected": false, "canonicalName": "USV1"}, "position": {"x": 910.3658488313224, "y": 40.0807263011049}, "selected": false}, {"data": {"id": "2782887", "shared_name": "853148", "MCODE_SCORE": 1.4000000000000001, "Gene": "853148", "Symbol": "PHO81", "name": "853148", "MCODE_TYPE": "Clustered", "SUID": 2782887, "DEGREE": 4.0, "id_original": "853148", "MCODE_CLUSTER_ID": 4.0, "selected": false, "canonicalName": "PHO81"}, "position": {"x": 895.9537394563224, "y": 167.1977917307924}, "selected": false}, {"data": {"id": "2782884", "shared_name": "851631", "MCODE_SCORE": 1.4000000000000001, "Gene": "851631", "Symbol": "UBC5", "name": "851631", "MCODE_TYPE": "Clustered", "SUID": 2782884, "DEGREE": 4.0, "id_original": "851631", "MCODE_CLUSTER_ID": 4.0, "selected": false, "canonicalName": "UBC5"}, "position": {"x": 975.0744364778068, "y": 110.3900524729799}, "selected": false}, {"data": {"id": "2782881", "shared_name": "851613", "MCODE_SCORE": 1.4000000000000001, "Gene": "851613", "Symbol": "NRG1", "name": "851613", "MCODE_TYPE": "Clustered", "SUID": 2782881, "DEGREE": 2.0, "id_original": "851613", "MCODE_CLUSTER_ID": 4.0, "selected": false, "canonicalName": "NRG1"}, "position": {"x": 993.0058253327873, "y": 218.93970457258928}, "selected": false}, {"data": {"id": "2782878", "shared_name": "851512", "MCODE_SCORE": 1.4000000000000001, "Gene": "851512", "Symbol": "STP4", "name": "851512", "MCODE_TYPE": "Clustered", "SUID": 2782878, "DEGREE": 2.0, "id_original": "851512", "MCODE_CLUSTER_ID": 4.0, "selected": false, "canonicalName": "STP4"}, "position": {"x": 850.0485880891349, "y": 83.39475218001115}, "selected": false}, {"data": {"id": "2782875", "shared_name": "855898", "MCODE_SCORE": 1.5, "Gene": "855898", "Symbol": "TPK2", "name": "855898", "MCODE_TYPE": "Seed", "SUID": 2782875, "DEGREE": 3.0, "id_original": "855898", "MCODE_CLUSTER_ID": 5.0, "selected": false, "canonicalName": "TPK2"}, "position": {"x": 620.8827460328605, "y": 587.8973538035464}, "selected": false}, {"data": {"id": "2782872", "shared_name": "855130", "MCODE_SCORE": 1.5, "Gene": "855130", "Symbol": "YPK2", "name": "855130", "MCODE_TYPE": "Clustered", "SUID": 2782872, "DEGREE": 3.0, "id_original": "855130", "MCODE_CLUSTER_ID": 5.0, "selected": false, "canonicalName": "YPK2"}, "position": {"x": 542.106745056298, "y": 636.4463039988589}, "selected": false}, {"data": {"id": "2782869", "shared_name": "853803", "MCODE_SCORE": 1.5, "Gene": "853803", "Symbol": "MSN4", "name": "853803", "MCODE_TYPE": "Clustered", "SUID": 2782869, "DEGREE": 3.0, "id_original": "853803", "MCODE_CLUSTER_ID": 5.0, "selected": false, "canonicalName": "MSN4"}, "position": {"x": 572.3872168580558, "y": 509.0878597849916}, "selected": false}, {"data": {"id": "2782866", "shared_name": "853275", "MCODE_SCORE": 1.5, "Gene": "853275", "Symbol": "TPK1", "name": "853275", "MCODE_TYPE": "Clustered", "SUID": 2782866, "DEGREE": 3.0, "id_original": "853275", "MCODE_CLUSTER_ID": 5.0, "selected": false, "canonicalName": "TPK1"}, "position": {"x": 493.6106284181144, "y": 557.6494137400698}, "selected": false}, {"data": {"id": "2782863", "shared_name": "855857", "MCODE_SCORE": 2.375, "Gene": "855857", "Symbol": "ATP15", "name": "855857", "MCODE_TYPE": "Clustered", "SUID": 2782863, "DEGREE": 4.0, "id_original": "855857", "MCODE_CLUSTER_ID": 3.0, "selected": false, "canonicalName": "ATP15"}, "position": {"x": 750.0485880891349, "y": 158.4558483714174}, "selected": false}, {"data": {"id": "2782860", "shared_name": "853994", "MCODE_SCORE": 2.375, "Gene": "853994", "Symbol": "MDH2", "name": "853994", "MCODE_TYPE": "Clustered", "SUID": 2782860, "DEGREE": 5.0, "id_original": "853994", "MCODE_CLUSTER_ID": 3.0, "selected": false, "canonicalName": "MDH2"}, "position": {"x": 704.4223368684318, "y": 146.94528928938615}, "selected": false}, {"data": {"id": "2782857", "shared_name": "853777", "MCODE_SCORE": 2.375, "Gene": "853777", "Symbol": "MDH1", "name": "853777", "MCODE_TYPE": "Clustered", "SUID": 2782857, "DEGREE": 5.0, "id_original": "853777", "MCODE_CLUSTER_ID": 3.0, "selected": false, "canonicalName": "MDH1"}, "position": {"x": 601.7906840364005, "y": 92.17978635969865}, "selected": false}, {"data": {"id": "2782854", "shared_name": "853709", "MCODE_SCORE": 2.375, "Gene": "853709", "Symbol": "SDH1", "name": "853709", "MCODE_TYPE": "Seed", "SUID": 2782854, "DEGREE": 4.0, "id_original": "853709", "MCODE_CLUSTER_ID": 3.0, "selected": false, "canonicalName": "SDH1"}, "position": {"x": 729.1603282014396, "y": 40.0807263011049}, "selected": false}, {"data": {"id": "2782851", "shared_name": "853585", "MCODE_SCORE": 2.375, "Gene": "853585", "Symbol": "ATP2", "name": "853585", "MCODE_TYPE": "Clustered", "SUID": 2782851, "DEGREE": 6.0, "id_original": "853585", "MCODE_CLUSTER_ID": 3.0, "selected": false, "canonicalName": "ATP2"}, "position": {"x": 665.3381999055412, "y": 219.08957639876115}, "selected": false}, {"data": {"id": "2782848", "shared_name": "852250", "MCODE_SCORE": 2.375, "Gene": "852250", "Symbol": "PET9", "name": "852250", "MCODE_TYPE": "Clustered", "SUID": 2782848, "DEGREE": 2.0, "id_original": "852250", "MCODE_CLUSTER_ID": 3.0, "selected": false, "canonicalName": "PET9"}, "position": {"x": 576.8323710481193, "y": 286.9281384104799}, "selected": false}, {"data": {"id": "2782845", "shared_name": "851892", "MCODE_SCORE": 2.375, "Gene": "851892", "Symbol": "ATP5", "name": "851892", "MCODE_TYPE": "Clustered", "SUID": 2782845, "DEGREE": 6.0, "id_original": "851892", "MCODE_CLUSTER_ID": 3.0, "selected": false, "canonicalName": "ATP5"}, "position": {"x": 605.9933512727287, "y": 178.7065807932924}, "selected": false}, {"data": {"id": "2782842", "shared_name": "851560", "MCODE_SCORE": 2.375, "Gene": "851560", "Symbol": "ATP16", "name": "851560", "MCODE_TYPE": "Clustered", "SUID": 2782842, "DEGREE": 6.0, "id_original": "851560", "MCODE_CLUSTER_ID": 3.0, "selected": false, "canonicalName": "ATP16"}, "position": {"x": 666.7958415071037, "y": 89.40879026594865}, "selected": false}, {"data": {"id": "2782839", "shared_name": "856741", "MCODE_SCORE": 2.9166666666666665, "Gene": "856741", "Symbol": "GPA2", "name": "856741", "MCODE_TYPE": "Clustered", "SUID": 2782839, "DEGREE": 2.0, "id_original": "856741", "MCODE_CLUSTER_ID": 2.0, "selected": false, "canonicalName": "GPA2"}, "position": {"x": 376.5229056397941, "y": 409.0878597849916}, "selected": false}, {"data": {"id": "2782836", "shared_name": "855303", "MCODE_SCORE": 2.9166666666666665, "Gene": "855303", "Symbol": "TPS3", "name": "855303", "MCODE_TYPE": "Clustered", "SUID": 2782836, "DEGREE": 8.0, "id_original": "855303", "MCODE_CLUSTER_ID": 2.0, "selected": false, "canonicalName": "TPS3"}, "position": {"x": 274.08902960219643, "y": 109.22398344221818}, "selected": false}, {"data": {"id": "2782833", "shared_name": "854872", "MCODE_SCORE": 2.9166666666666665, "Gene": "854872", "Symbol": "TSL1", "name": "854872", "MCODE_TYPE": "Clustered", "SUID": 2782833, "DEGREE": 8.0, "id_original": "854872", "MCODE_CLUSTER_ID": 2.0, "selected": false, "canonicalName": "TSL1"}, "position": {"x": 311.73183660903237, "y": 63.223067914874434}, "selected": false}, {"data": {"id": "2782830", "shared_name": "853932", "MCODE_SCORE": 2.9166666666666665, "Gene": "853932", "Symbol": "GLG1", "name": "853932", "MCODE_TYPE": "Clustered", "SUID": 2782830, "DEGREE": 6.0, "id_original": "853932", "MCODE_CLUSTER_ID": 2.0, "selected": false, "canonicalName": "GLG1"}, "position": {"x": 385.4201987306144, "y": 40.0807263011049}, "selected": false}, {"data": {"id": "2782827", "shared_name": "852423", "MCODE_SCORE": 2.9166666666666665, "Gene": "852423", "Symbol": "TPS1", "name": "852423", "MCODE_TYPE": "Clustered", "SUID": 2782827, "DEGREE": 8.0, "id_original": "852423", "MCODE_CLUSTER_ID": 2.0, "selected": false, "canonicalName": "TPS1"}, "position": {"x": 405.2021048585441, "y": 107.99562040510881}, "selected": false}, {"data": {"id": "2782824", "shared_name": "852286", "MCODE_SCORE": 2.9166666666666665, "Gene": "852286", "Symbol": "NTH2", "name": "852286", "MCODE_TYPE": "Clustered", "SUID": 2782824, "DEGREE": 5.0, "id_original": "852286", "MCODE_CLUSTER_ID": 2.0, "selected": false, "canonicalName": "NTH2"}, "position": {"x": 264.72672491469643, "y": 180.62581411543596}, "selected": false}, {"data": {"id": "2782821", "shared_name": "851670", "MCODE_SCORE": 2.9166666666666665, "Gene": "851670", "Symbol": "GIS1", "name": "851670", "MCODE_TYPE": "Clustered", "SUID": 2782821, "DEGREE": 2.0, "id_original": "851670", "MCODE_CLUSTER_ID": 2.0, "selected": false, "canonicalName": "GIS1"}, "position": {"x": 476.8323710481193, "y": 377.9572598094057}, "selected": false}, {"data": {"id": "2782818", "shared_name": "851646", "MCODE_SCORE": 2.9166666666666665, "Gene": "851646", "Symbol": "TPS2", "name": "851646", "MCODE_TYPE": "Seed", "SUID": 2782818, "DEGREE": 8.0, "id_original": "851646", "MCODE_CLUSTER_ID": 2.0, "selected": false, "canonicalName": "TPS2"}, "position": {"x": 343.15172796645425, "y": 113.7327381724428}, "selected": false}, {"data": {"id": "2782815", "shared_name": "851564", "MCODE_SCORE": 2.9166666666666665, "Gene": "851564", "Symbol": "NTH1", "name": "851564", "MCODE_TYPE": "Clustered", "SUID": 2782815, "DEGREE": 5.0, "id_original": "851564", "MCODE_CLUSTER_ID": 2.0, "selected": false, "canonicalName": "NTH1"}, "position": {"x": 361.96283941664956, "y": 199.58985487166154}, "selected": false}, {"data": {"id": "2782812", "shared_name": "851480", "MCODE_SCORE": 2.9166666666666665, "Gene": "851480", "Symbol": "MRK1", "name": "851480", "MCODE_TYPE": "Clustered", "SUID": 2782812, "DEGREE": 4.0, "id_original": "851480", "MCODE_CLUSTER_ID": 2.0, "selected": false, "canonicalName": "MRK1"}, "position": {"x": 395.38185339370034, "y": 292.7763058299135}, "selected": false}, {"data": {"id": "2782809", "shared_name": "850962", "MCODE_SCORE": 2.9166666666666665, "Gene": "850962", "Symbol": "GSY2", "name": "850962", "MCODE_TYPE": "Clustered", "SUID": 2782809, "DEGREE": 7.0, "id_original": "850962", "MCODE_CLUSTER_ID": 2.0, "selected": false, "canonicalName": "GSY2"}, "position": {"x": 321.396539978173, "y": 184.0740753764223}, "selected": false}, {"data": {"id": "2782806", "shared_name": "850569", "MCODE_SCORE": 2.9166666666666665, "Gene": "850569", "Symbol": "GSY1", "name": "850569", "MCODE_TYPE": "Clustered", "SUID": 2782806, "DEGREE": 7.0, "id_original": "850569", "MCODE_CLUSTER_ID": 2.0, "selected": false, "canonicalName": "GSY1"}, "position": {"x": 407.99953680434487, "y": 169.29635206404436}, "selected": false}, {"data": {"id": "2782803", "shared_name": "3799970", "MCODE_SCORE": 9.625, "Gene": "3799970", "Symbol": "COX26", "name": "3799970", "MCODE_TYPE": "Clustered", "SUID": 2782803, "DEGREE": 15.0, "id_original": "3799970", "MCODE_CLUSTER_ID": 1.0, "selected": false, "canonicalName": "COX26"}, "position": {"x": 74.2461698571708, "y": 300.25978819075334}, "selected": false}, {"data": {"id": "2782800", "shared_name": "1466480", "MCODE_SCORE": 9.625, "Gene": "1466480", "Symbol": "MCO10", "name": "1466480", "MCODE_TYPE": "Clustered", "SUID": 2782800, "DEGREE": 10.0, "id_original": "1466480", "MCODE_CLUSTER_ID": 1.0, "selected": false, "canonicalName": "MCO10"}, "position": {"x": -92.28371972641685, "y": 52.201469098956466}, "selected": false}, {"data": {"id": "2782797", "shared_name": "856689", "MCODE_SCORE": 9.625, "Gene": "856689", "Symbol": "RIP1", "name": "856689", "MCODE_TYPE": "Clustered", "SUID": 2782797, "DEGREE": 25.0, "id_original": "856689", "MCODE_CLUSTER_ID": 1.0, "selected": false, "canonicalName": "RIP1"}, "position": {"x": -33.024182983252786, "y": 172.33812681380022}, "selected": false}, {"data": {"id": "2782794", "shared_name": "856672", "MCODE_SCORE": 9.625, "Gene": "856672", "Symbol": "CYC7", "name": "856672", "MCODE_TYPE": "Clustered", "SUID": 2782794, "DEGREE": 19.0, "id_original": "856672", "MCODE_CLUSTER_ID": 1.0, "selected": false, "canonicalName": "CYC7"}, "position": {"x": -28.99595422348716, "y": 251.44777647200334}, "selected": false}, {"data": {"id": "2782791", "shared_name": "856390", "MCODE_SCORE": 9.625, "Gene": "856390", "Symbol": "QCR10", "name": "856390", "MCODE_TYPE": "Clustered", "SUID": 2782791, "DEGREE": 20.0, "id_original": "856390", "MCODE_CLUSTER_ID": 1.0, "selected": false, "canonicalName": "QCR10"}, "position": {"x": 144.1078665772941, "y": 211.49004331770647}, "selected": false}, {"data": {"id": "2782788", "shared_name": "856321", "MCODE_SCORE": 9.625, "Gene": "856321", "Symbol": "QCR2", "name": "856321", "MCODE_TYPE": "Clustered", "SUID": 2782788, "DEGREE": 25.0, "id_original": "856321", "MCODE_CLUSTER_ID": 1.0, "selected": false, "canonicalName": "QCR2"}, "position": {"x": 81.65268021025918, "y": 245.23760191145647}, "selected": false}, {"data": {"id": "2782785", "shared_name": "856131", "MCODE_SCORE": 9.625, "Gene": "856131", "Symbol": "ATP20", "name": "856131", "MCODE_TYPE": "Clustered", "SUID": 2782785, "DEGREE": 24.0, "id_original": "856131", "MCODE_CLUSTER_ID": 1.0, "selected": false, "canonicalName": "ATP20"}, "position": {"x": -22.28483361801841, "y": 227.97829405012834}, "selected": false}, {"data": {"id": "2782782", "shared_name": "856027", "MCODE_SCORE": 9.625, "Gene": "856027", "Symbol": "ATP4", "name": "856027", "MCODE_TYPE": "Clustered", "SUID": 2782782, "DEGREE": 26.0, "id_original": "856027", "MCODE_CLUSTER_ID": 1.0, "selected": false, "canonicalName": "ATP4"}, "position": {"x": 24.80574255385659, "y": 150.69145933333147}, "selected": false}, {"data": {"id": "2782779", "shared_name": "855675", "MCODE_SCORE": 9.625, "Gene": "855675", "Symbol": "COX5A", "name": "855675", "MCODE_TYPE": "Clustered", "SUID": 2782779, "DEGREE": 22.0, "id_original": "855675", "MCODE_CLUSTER_ID": 1.0, "selected": false, "canonicalName": "COX5A"}, "position": {"x": 57.17063932814858, "y": 186.13640562239397}, "selected": false}, {"data": {"id": "2782776", "shared_name": "855298", "MCODE_SCORE": 9.625, "Gene": "855298", "Symbol": "COX7", "name": "855298", "MCODE_TYPE": "Clustered", "SUID": 2782776, "DEGREE": 17.0, "id_original": "855298", "MCODE_CLUSTER_ID": 1.0, "selected": false, "canonicalName": "COX7"}, "position": {"x": 58.259996704247214, "y": 115.15047422590959}, "selected": false}, {"data": {"id": "2782773", "shared_name": "854919", "MCODE_SCORE": 9.625, "Gene": "854919", "Symbol": "NDI1", "name": "854919", "MCODE_TYPE": "Clustered", "SUID": 2782773, "DEGREE": 18.0, "id_original": "854919", "MCODE_CLUSTER_ID": 1.0, "selected": false, "canonicalName": "NDI1"}, "position": {"x": 117.47580938735268, "y": 148.4384380930971}, "selected": false}, {"data": {"id": "2782770", "shared_name": "854893", "MCODE_SCORE": 9.625, "Gene": "854893", "Symbol": "ATP18", "name": "854893", "MCODE_TYPE": "Clustered", "SUID": 2782770, "DEGREE": 11.0, "id_original": "854893", "MCODE_CLUSTER_ID": 1.0, "selected": false, "canonicalName": "ATP18"}, "position": {"x": -91.64329309067466, "y": 192.81182066145647}, "selected": false}, {"data": {"id": "2782767", "shared_name": "854695", "MCODE_SCORE": 9.625, "Gene": "854695", "Symbol": "COX5B", "name": "854695", "MCODE_TYPE": "Clustered", "SUID": 2782767, "DEGREE": 22.0, "id_original": "854695", "MCODE_CLUSTER_ID": 1.0, "selected": false, "canonicalName": "COX5B"}, "position": {"x": 98.46790342726968, "y": 100.85970274153459}, "selected": false}, {"data": {"id": "2782764", "shared_name": "854231", "MCODE_SCORE": 9.625, "Gene": "854231", "Symbol": "CYT1", "name": "854231", "MCODE_TYPE": "Seed", "SUID": 2782764, "DEGREE": 25.0, "id_original": "854231", "MCODE_CLUSTER_ID": 1.0, "selected": false, "canonicalName": "CYT1"}, "position": {"x": 24.132665924217918, "y": 162.1349712962221}, "selected": false}, {"data": {"id": "2782761", "shared_name": "854077", "MCODE_SCORE": 9.625, "Gene": "854077", "Symbol": "ATP19", "name": "854077", "MCODE_TYPE": "Clustered", "SUID": 2782761, "DEGREE": 10.0, "id_original": "854077", "MCODE_CLUSTER_ID": 1.0, "selected": false, "canonicalName": "ATP19"}, "position": {"x": -125.17790002426841, "y": 134.99150816145647}, "selected": false}, {"data": {"id": "2782758", "shared_name": "853853", "MCODE_SCORE": 9.625, "Gene": "853853", "Symbol": "ATP7", "name": "853853", "MCODE_TYPE": "Clustered", "SUID": 2782758, "DEGREE": 22.0, "id_original": "853853", "MCODE_CLUSTER_ID": 1.0, "selected": false, "canonicalName": "ATP7"}, "position": {"x": -78.35426110825279, "y": 167.98747984114397}, "selected": false}, {"data": {"id": "2782755", "shared_name": "853716", "MCODE_SCORE": 9.625, "Gene": "853716", "Symbol": "SDH3", "name": "853716", "MCODE_TYPE": "Clustered", "SUID": 2782755, "DEGREE": 13.0, "id_original": "853716", "MCODE_CLUSTER_ID": 1.0, "selected": false, "canonicalName": "SDH3"}, "position": {"x": 164.72672491469643, "y": 161.75054136458147}, "selected": false}, {"data": {"id": "2782752", "shared_name": "853273", "MCODE_SCORE": 9.625, "Gene": "853273", "Symbol": "QCR8", "name": "853273", "MCODE_TYPE": "Clustered", "SUID": 2782752, "DEGREE": 24.0, "id_original": "853273", "MCODE_CLUSTER_ID": 1.0, "selected": false, "canonicalName": "QCR8"}, "position": {"x": 93.76004057326577, "y": 191.73393980208147}, "selected": false}, {"data": {"id": "2782749", "shared_name": "853095", "MCODE_SCORE": 9.625, "Gene": "853095", "Symbol": "QCR9", "name": "853095", "MCODE_TYPE": "Clustered", "SUID": 2782749, "DEGREE": 25.0, "id_original": "853095", "MCODE_CLUSTER_ID": 1.0, "selected": false, "canonicalName": "QCR9"}, "position": {"x": 42.2818930665519, "y": 263.8041607493471}, "selected": false}, {"data": {"id": "2782746", "shared_name": "852688", "MCODE_SCORE": 9.625, "Gene": "852688", "Symbol": "COX4", "name": "852688", "MCODE_TYPE": "Clustered", "SUID": 2782746, "DEGREE": 22.0, "id_original": "852688", "MCODE_CLUSTER_ID": 1.0, "selected": false, "canonicalName": "COX4"}, "position": {"x": 131.05734472670815, "y": 198.42983213606584}, "selected": false}, {"data": {"id": "2782743", "shared_name": "852327", "MCODE_SCORE": 9.625, "Gene": "852327", "Symbol": "ATP3", "name": "852327", "MCODE_TYPE": "Clustered", "SUID": 2782743, "DEGREE": 21.0, "id_original": "852327", "MCODE_CLUSTER_ID": 1.0, "selected": false, "canonicalName": "ATP3"}, "position": {"x": 4.051561645653464, "y": 106.39086118880022}, "selected": false}, {"data": {"id": "2782740", "shared_name": "852235", "MCODE_SCORE": 9.625, "Gene": "852235", "Symbol": "COR1", "name": "852235", "MCODE_TYPE": "Clustered", "SUID": 2782740, "DEGREE": 26.0, "id_original": "852235", "MCODE_CLUSTER_ID": 1.0, "selected": false, "canonicalName": "COR1"}, "position": {"x": 78.39750891127846, "y": 201.7665630930971}, "selected": false}, {"data": {"id": "2782737", "shared_name": "852177", "MCODE_SCORE": 9.625, "Gene": "852177", "Symbol": "ATP1", "name": "852177", "MCODE_TYPE": "Clustered", "SUID": 2782737, "DEGREE": 23.0, "id_original": "852177", "MCODE_CLUSTER_ID": 1.0, "selected": false, "canonicalName": "ATP1"}, "position": {"x": -21.126706786963723, "y": 146.64092222395647}, "selected": false}, {"data": {"id": "2782734", "shared_name": "852142", "MCODE_SCORE": 9.625, "Gene": "852142", "Symbol": "QCR7", "name": "852142", "MCODE_TYPE": "Clustered", "SUID": 2782734, "DEGREE": 21.0, "id_original": "852142", "MCODE_CLUSTER_ID": 1.0, "selected": false, "canonicalName": "QCR7"}, "position": {"x": 154.42228155532143, "y": 233.52407041731584}, "selected": false}, {"data": {"id": "2782731", "shared_name": "851983", "MCODE_SCORE": 9.625, "Gene": "851983", "Symbol": "ATP17", "name": "851983", "MCODE_TYPE": "Clustered", "SUID": 2782731, "DEGREE": 12.0, "id_original": "851983", "MCODE_CLUSTER_ID": 1.0, "selected": false, "canonicalName": "ATP17"}, "position": {"x": -57.1154458006356, "y": 102.89003721419084}, "selected": false}, {"data": {"id": "2782728", "shared_name": "851922", "MCODE_SCORE": 9.625, "Gene": "851922", "Symbol": "TIM11", "name": "851922", "MCODE_TYPE": "Clustered", "SUID": 2782728, "DEGREE": 10.0, "id_original": "851922", "MCODE_CLUSTER_ID": 1.0, "selected": false, "canonicalName": "TIM11"}, "position": {"x": -48.17296380600669, "y": 40.0807263011049}, "selected": false}, {"data": {"id": "2782725", "shared_name": "851758", "MCODE_SCORE": 9.625, "Gene": "851758", "Symbol": "SDH4", "name": "851758", "MCODE_TYPE": "Clustered", "SUID": 2782725, "DEGREE": 12.0, "id_original": "851758", "MCODE_CLUSTER_ID": 1.0, "selected": false, "canonicalName": "SDH4"}, "position": {"x": 135.72783117690346, "y": 308.48845640364397}, "selected": false}, {"data": {"id": "2782722", "shared_name": "851111", "MCODE_SCORE": 9.625, "Gene": "851111", "Symbol": "COX8", "name": "851111", "MCODE_TYPE": "Clustered", "SUID": 2782722, "DEGREE": 20.0, "id_original": "851111", "MCODE_CLUSTER_ID": 1.0, "selected": false, "canonicalName": "COX8"}, "position": {"x": 7.578089050438621, "y": 174.73293272200334}, "selected": false}, {"data": {"id": "2782719", "shared_name": "851002", "MCODE_SCORE": 9.625, "Gene": "851002", "Symbol": "ATP14", "name": "851002", "MCODE_TYPE": "Clustered", "SUID": 2782719, "DEGREE": 10.0, "id_original": "851002", "MCODE_CLUSTER_ID": 1.0, "selected": false, "canonicalName": "ATP14"}, "position": {"x": -137.71577233872154, "y": 90.15679136458147}, "selected": false}, {"data": {"id": "2782716", "shared_name": "850727", "MCODE_SCORE": 9.625, "Gene": "850727", "Symbol": "COX12", "name": "850727", "MCODE_TYPE": "Clustered", "SUID": 2782716, "DEGREE": 22.0, "id_original": "850727", "MCODE_CLUSTER_ID": 1.0, "selected": false, "canonicalName": "COX12"}, "position": {"x": 11.665392212059714, "y": 242.95378843489397}, "selected": false}, {"data": {"id": "2782713", "shared_name": "850685", "MCODE_SCORE": 9.625, "Gene": "850685", "Symbol": "SDH2", "name": "850685", "MCODE_TYPE": "Clustered", "SUID": 2782713, "DEGREE": 20.0, "id_original": "850685", "MCODE_CLUSTER_ID": 1.0, "selected": false, "canonicalName": "SDH2"}, "position": {"x": 56.49740820327065, "y": 162.39272276106584}, "selected": false}, {"data": {"id": "2782686", "shared_name": "850593", "MCODE_SCORE": 9.625, "Gene": "850593", "Symbol": "QCR6", "name": "850593", "MCODE_TYPE": "Clustered", "SUID": 2782686, "DEGREE": 24.0, "id_original": "850593", "MCODE_CLUSTER_ID": 1.0, "selected": false, "canonicalName": "QCR6"}, "position": {"x": 92.28633051314614, "y": 265.93898740950334}, "selected": false}], "edges": [{"data": {"id": "2783928", "source": "2782950", "target": "2782953", "SCORE": "843", "shared_interaction": "Direct", "source_original": "853441", "Name_B": "VTC3", "Name_A": "VTC4", "target_original": "856088", "Gene_B": "856088", "interaction": "Direct", "SUID": 2783928, "TYPE": "Direct", "Gene_A": "853441", "support": "{\"StringDB\": \"physical\", \"StringEvidence\": \"textmining:0.954,coexpression:0.932,database:0.708,experiments:0.441\", \"BioGrid\": \"physical\", \"BioGrid_PubMed\": \"27587415,25315834\", \"BioGrid_Type\": \"Affinity Capture-Western\", \"String_Score_Physical\": 843, \"String_Score_Combined\": 991}", "selected": false}, "selected": false}, {"data": {"id": "2783175", "source": "2782947", "target": "2782953", "SCORE": "208", "shared_interaction": "Direct", "source_original": "850633", "Name_B": "VTC3", "Name_A": "HSP104", "target_original": "856088", "Gene_B": "856088", "interaction": "Direct", "SUID": 2783175, "TYPE": "Direct", "Gene_A": "850633", "support": "{\"BioGrid\": \"physical\", \"BioGrid_PubMed\": \"27373154\", \"BioGrid_Type\": \"Affinity Capture-MS\"}", "selected": false}, "selected": false}, {"data": {"id": "2783172", "source": "2782947", "target": "2782950", "SCORE": "208", "shared_interaction": "Direct", "source_original": "850633", "Name_B": "VTC4", "Name_A": "HSP104", "target_original": "853441", "Gene_B": "853441", "interaction": "Direct", "SUID": 2783172, "TYPE": "Direct", "Gene_A": "850633", "support": "{\"BioGrid\": \"physical\", \"BioGrid_PubMed\": \"27373154\", \"BioGrid_Type\": \"Affinity Capture-MS\"}", "selected": false}, "selected": false}, {"data": {"id": "2784009", "source": "2782941", "target": "2782944", "SCORE": "540", "shared_interaction": "Direct", "source_original": "853950", "Name_B": "YOR062C", "Name_A": "YKR075C", "target_original": "854228", "Gene_B": "854228", "interaction": "Direct", "SUID": 2784009, "TYPE": "Direct", "Gene_A": "853950", "support": "{\"StringDB\": \"physical\", \"StringEvidence\": \"textmining:0.741,database:0.520\", \"String_Score_Physical\": 540, \"String_Score_Combined\": 579}", "selected": false}, "selected": false}, {"data": {"id": "2783769", "source": "2782938", "target": "2782944", "SCORE": "540", "shared_interaction": "Direct", "source_original": "852340", "Name_B": "YOR062C", "Name_A": "REG2", "target_original": "854228", "Gene_B": "854228", "interaction": "Direct", "SUID": 2783769, "TYPE": "Direct", "Gene_A": "852340", "support": "{\"StringDB\": \"physical\", \"StringEvidence\": \"database:0.520\", \"String_Score_Physical\": 540, \"String_Score_Combined\": 540}", "selected": false}, "selected": false}, {"data": {"id": "2783766", "source": "2782938", "target": "2782941", "SCORE": "540", "shared_interaction": "Direct", "source_original": "852340", "Name_B": "YKR075C", "Name_A": "REG2", "target_original": "853950", "Gene_B": "853950", "interaction": "Direct", "SUID": 2783766, "TYPE": "Direct", "Gene_A": "852340", "support": "{\"StringDB\": \"physical\", \"StringEvidence\": \"database:0.520\", \"String_Score_Physical\": 540, \"String_Score_Combined\": 540}", "selected": false}, "selected": false}, {"data": {"id": "2783169", "source": "2782932", "target": "2782935", "SCORE": "600", "shared_interaction": "Direct", "source_original": "850614", "Name_B": "EMI2", "Name_A": "HXK1", "target_original": "852128", "Gene_B": "852128", "interaction": "Direct", "SUID": 2783169, "TYPE": "Direct", "Gene_A": "850614", "support": "{\"StringDB\": \"physical\", \"StringEvidence\": \"coexpression:0.968,database:0.583,textmining:0.426,cooccurrence:0.421\", \"String_Score_Physical\": 600, \"String_Score_Combined\": 988}", "selected": false}, "selected": false}, {"data": {"id": "2783061", "source": "2782929", "target": "2782935", "SCORE": "905", "shared_interaction": "Direct", "source_original": "850317", "Name_B": "EMI2", "Name_A": "GLK1", "target_original": "852128", "Gene_B": "852128", "interaction": "Direct", "SUID": 2783061, "TYPE": "Direct", "Gene_A": "850317", "support": "{\"StringDB\": \"physical\", \"StringEvidence\": \"coexpression:0.938,experiments:0.764,database:0.583,textmining:0.479,cooccurrence:0.425\", \"BioGrid\": \"genetic\", \"BioGrid_PubMed\": \"18408719\", \"BioGrid_Type\": \"Synthetic Growth Defect\", \"String_Score_Physical\": 905, \"String_Score_Combined\": 994}", "selected": false}, "selected": false}, {"data": {"id": "2783058", "source": "2782929", "target": "2782932", "SCORE": "999", "shared_interaction": "Direct", "source_original": "850317", "Name_B": "HXK1", "Name_A": "GLK1", "target_original": "850614", "Gene_B": "850614", "interaction": "Direct", "SUID": 2783058, "TYPE": "Direct", "Gene_A": "850317", "support": "{\"StringDB\": \"physical\", \"StringEvidence\": \"experiments:0.999,textmining:0.981,coexpression:0.968,database:0.583,cooccurrence:0.422\", \"BioGrid\": \"genetic\", \"BioGrid_PubMed\": \"19525230,16941010,15358789,10217505\", \"BioGrid_Type\": \"Phenotypic Enhancement,Synthetic Lethality\", \"String_Score_Physical\": 999, \"String_Score_Combined\": 999}", "selected": false}, "selected": false}, {"data": {"id": "2783871", "source": "2782920", "target": "2782926", "SCORE": "900", "shared_interaction": "Direct", "source_original": "853119", "Name_B": "GOR1", "Name_A": "TDA10", "target_original": "855447", "Gene_B": "855447", "interaction": "Direct", "SUID": 2783871, "TYPE": "Direct", "Gene_A": "853119", "support": "{\"StringDB\": \"physical\", \"StringEvidence\": \"database:0.896,textmining:0.443,neighborhood:0.117,coexpression:0.043\", \"String_Score_Physical\": 900, \"String_Score_Combined\": 952}", "selected": false}, "selected": false}, {"data": {"id": "2783868", "source": "2782920", "target": "2782923", "SCORE": "900", "shared_interaction": "Direct", "source_original": "853119", "Name_B": "ALD4", "Name_A": "TDA10", "target_original": "854556", "Gene_B": "854556", "interaction": "Direct", "SUID": 2783868, "TYPE": "Direct", "Gene_A": "853119", "support": "{\"StringDB\": \"physical\", \"StringEvidence\": \"database:0.896,textmining:0.083,coexpression:0.059\", \"String_Score_Physical\": 900, \"String_Score_Combined\": 913}", "selected": false}, "selected": false}, {"data": {"id": "2783076", "source": "2782917", "target": "2782926", "SCORE": "900", "shared_interaction": "Direct", "source_original": "850514", "Name_B": "GOR1", "Name_A": "AGX1", "target_original": "855447", "Gene_B": "855447", "interaction": "Direct", "SUID": 2783076, "TYPE": "Direct", "Gene_A": "850514", "support": "{\"StringDB\": \"physical\", \"StringEvidence\": \"database:0.896,textmining:0.888,coexpression:0.273,neighborhood:0.119,fusion:0.026\", \"String_Score_Physical\": 900, \"String_Score_Combined\": 993}", "selected": false}, "selected": false}, {"data": {"id": "2783073", "source": "2782917", "target": "2782923", "SCORE": "273", "shared_interaction": "Direct", "source_original": "850514", "Name_B": "ALD4", "Name_A": "AGX1", "target_original": "854556", "Gene_B": "854556", "interaction": "Direct", "SUID": 2783073, "TYPE": "Direct", "Gene_A": "850514", "support": "{\"StringDB\": \"physical\", \"StringEvidence\": \"coexpression:0.309,experiments:0.242,textmining:0.149\", \"String_Score_Physical\": 273, \"String_Score_Combined\": 572}", "selected": false}, "selected": false}, {"data": {"id": "2783877", "source": "2782911", "target": "2782914", "SCORE": "208", "shared_interaction": "Direct", "source_original": "853164", "Name_B": "PTK2", "Name_A": "MGA1", "target_original": "853522", "Gene_B": "853522", "interaction": "Direct", "SUID": 2783877, "TYPE": "Direct", "Gene_A": "853164", "support": "{\"BioGrid\": \"physical\", \"BioGrid_PubMed\": \"16319894\", \"BioGrid_Type\": \"Biochemical Activity\"}", "selected": false}, "selected": false}, {"data": {"id": "2783781", "source": "2782908", "target": "2782914", "SCORE": "490", "shared_interaction": "Direct", "source_original": "852515", "Name_B": "PTK2", "Name_A": "SDS24", "target_original": "853522", "Gene_B": "853522", "interaction": "Direct", "SUID": 2783781, "TYPE": "Direct", "Gene_A": "852515", "support": "{\"StringDB\": \"physical\", \"StringEvidence\": \"database:0.346,coexpression:0.216,experiments:0.187,textmining:0.047\", \"String_Score_Physical\": 490, \"String_Score_Combined\": 619}", "selected": false}, "selected": false}, {"data": {"id": "2783496", "source": "2782905", "target": "2782911", "SCORE": "208", "shared_interaction": "Direct", "source_original": "851834", "Name_B": "MGA1", "Name_A": "VHS1", "target_original": "853164", "Gene_B": "853164", "interaction": "Direct", "SUID": 2783496, "TYPE": "Direct", "Gene_A": "851834", "support": "{\"BioGrid\": \"physical\", \"BioGrid_PubMed\": \"16319894\", \"BioGrid_Type\": \"Biochemical Activity\"}", "selected": false}, "selected": false}, {"data": {"id": "2783493", "source": "2782905", "target": "2782908", "SCORE": "490", "shared_interaction": "Direct", "source_original": "851834", "Name_B": "SDS24", "Name_A": "VHS1", "target_original": "852515", "Gene_B": "852515", "interaction": "Direct", "SUID": 2783493, "TYPE": "Direct", "Gene_A": "851834", "support": "{\"StringDB\": \"physical\", \"StringEvidence\": \"database:0.346,experiments:0.187,textmining:0.047,coexpression:0.039\", \"String_Score_Physical\": 490, \"String_Score_Combined\": 533}", "selected": false}, "selected": false}, {"data": {"id": "2783823", "source": "2782899", "target": "2782902", "SCORE": "390", "shared_interaction": "Direct", "source_original": "853031", "Name_B": "TPO4", "Name_A": "YGR130C", "target_original": "854447", "Gene_B": "854447", "interaction": "Direct", "SUID": 2783823, "TYPE": "Direct", "Gene_A": "853031", "support": "{\"StringDB\": \"physical\", \"StringEvidence\": \"experiments:0.365,textmining:0.064,coexpression:0.058\", \"BioGrid\": \"physical\", \"BioGrid_PubMed\": \"22615397,18467557\", \"BioGrid_Type\": \"PCA\", \"String_Score_Physical\": 390, \"String_Score_Combined\": 463}", "selected": false}, "selected": false}, {"data": {"id": "2783718", "source": "2782896", "target": "2782899", "SCORE": "286", "shared_interaction": "Direct", "source_original": "852251", "Name_B": "YGR130C", "Name_A": "YBL029C-A", "target_original": "853031", "Gene_B": "853031", "interaction": "Direct", "SUID": 2783718, "TYPE": "Direct", "Gene_A": "852251", "support": "{\"StringDB\": \"physical\", \"StringEvidence\": \"experiments:0.257\", \"String_Score_Physical\": 286, \"String_Score_Combined\": 286}", "selected": false}, "selected": false}, {"data": {"id": "2783070", "source": "2782893", "target": "2782902", "SCORE": "266", "shared_interaction": "Direct", "source_original": "850385", "Name_B": "TPO4", "Name_A": "HSP30", "target_original": "854447", "Gene_B": "854447", "interaction": "Direct", "SUID": 2783070, "TYPE": "Direct", "Gene_A": "850385", "support": "{\"StringDB\": \"physical\", \"StringEvidence\": \"textmining:0.287,experiments:0.235,coexpression:0.059\", \"BioGrid\": \"physical\", \"BioGrid_PubMed\": \"18467557\", \"BioGrid_Type\": \"PCA\", \"String_Score_Physical\": 266, \"String_Score_Combined\": 507}", "selected": false}, "selected": false}, {"data": {"id": "2783067", "source": "2782893", "target": "2782899", "SCORE": "208", "shared_interaction": "Direct", "source_original": "850385", "Name_B": "YGR130C", "Name_A": "HSP30", "target_original": "853031", "Gene_B": "853031", "interaction": "Direct", "SUID": 2783067, "TYPE": "Direct", "Gene_A": "850385", "support": "{\"BioGrid\": \"physical\", \"BioGrid_PubMed\": \"18467557\", \"BioGrid_Type\": \"PCA\"}", "selected": false}, "selected": false}, {"data": {"id": "2783064", "source": "2782893", "target": "2782896", "SCORE": "208", "shared_interaction": "Direct", "source_original": "850385", "Name_B": "YBL029C-A", "Name_A": "HSP30", "target_original": "852251", "Gene_B": "852251", "interaction": "Direct", "SUID": 2783064, "TYPE": "Direct", "Gene_A": "850385", "support": "{\"BioGrid\": \"physical\", \"BioGrid_PubMed\": \"18467557\", \"BioGrid_Type\": \"PCA\"}", "selected": false}, "selected": false}, {"data": {"id": "2783874", "source": "2782887", "target": "2782890", "SCORE": "364", "shared_interaction": "Direct", "source_original": "853148", "Name_B": "USV1", "Name_A": "PHO81", "target_original": "855871", "Gene_B": "855871", "interaction": "Direct", "SUID": 2783874, "TYPE": "Direct", "Gene_A": "853148", "support": "{\"StringDB\": \"physical\", \"StringEvidence\": \"database:0.229,experiments:0.140,coexpression:0.016,textmining:0.001\", \"String_Score_Physical\": 364, \"String_Score_Combined\": 374}", "selected": false}, "selected": false}, {"data": {"id": "2783442", "source": "2782884", "target": "2782890", "SCORE": "295", "shared_interaction": "Direct", "source_original": "851631", "Name_B": "USV1", "Name_A": "UBC5", "target_original": "855871", "Gene_B": "855871", "interaction": "Direct", "SUID": 2783442, "TYPE": "Direct", "Gene_A": "851631", "support": "{\"StringDB\": \"physical\", \"StringEvidence\": \"database:0.194,experiments:0.089,coexpression:0.086,textmining:0.005\", \"String_Score_Physical\": 295, \"String_Score_Combined\": 359}", "selected": false}, "selected": false}, {"data": {"id": "2783439", "source": "2782884", "target": "2782887", "SCORE": "337", "shared_interaction": "Direct", "source_original": "851631", "Name_B": "PHO81", "Name_A": "UBC5", "target_original": "853148", "Gene_B": "853148", "interaction": "Direct", "SUID": 2783439, "TYPE": "Direct", "Gene_A": "851631", "support": "{\"StringDB\": \"physical\", \"StringEvidence\": \"database:0.200,experiments:0.137,textmining:0.034,coexpression:0.001\", \"String_Score_Physical\": 337, \"String_Score_Combined\": 361}", "selected": false}, "selected": false}, {"data": {"id": "2783436", "source": "2782881", "target": "2782887", "SCORE": "364", "shared_interaction": "Direct", "source_original": "851613", "Name_B": "PHO81", "Name_A": "NRG1", "target_original": "853148", "Gene_B": "853148", "interaction": "Direct", "SUID": 2783436, "TYPE": "Direct", "Gene_A": "851613", "support": "{\"StringDB\": \"physical\", \"StringEvidence\": \"database:0.229,experiments:0.140,coexpression:0.016,textmining:0.001\", \"String_Score_Physical\": 364, \"String_Score_Combined\": 374}", "selected": false}, "selected": false}, {"data": {"id": "2783433", "source": "2782881", "target": "2782884", "SCORE": "295", "shared_interaction": "Direct", "source_original": "851613", "Name_B": "UBC5", "Name_A": "NRG1", "target_original": "851631", "Gene_B": "851631", "interaction": "Direct", "SUID": 2783433, "TYPE": "Direct", "Gene_A": "851613", "support": "{\"StringDB\": \"physical\", \"StringEvidence\": \"database:0.194,experiments:0.089,coexpression:0.058,textmining:0.005\", \"String_Score_Physical\": 295, \"String_Score_Combined\": 340}", "selected": false}, "selected": false}, {"data": {"id": "2783397", "source": "2782878", "target": "2782887", "SCORE": "364", "shared_interaction": "Direct", "source_original": "851512", "Name_B": "PHO81", "Name_A": "STP4", "target_original": "853148", "Gene_B": "853148", "interaction": "Direct", "SUID": 2783397, "TYPE": "Direct", "Gene_A": "851512", "support": "{\"StringDB\": \"physical\", \"StringEvidence\": \"database:0.229,experiments:0.140,coexpression:0.016,textmining:0.001\", \"String_Score_Physical\": 364, \"String_Score_Combined\": 374}", "selected": false}, "selected": false}, {"data": {"id": "2783394", "source": "2782878", "target": "2782884", "SCORE": "295", "shared_interaction": "Direct", "source_original": "851512", "Name_B": "UBC5", "Name_A": "STP4", "target_original": "851631", "Gene_B": "851631", "interaction": "Direct", "SUID": 2783394, "TYPE": "Direct", "Gene_A": "851512", "support": "{\"StringDB\": \"physical\", \"StringEvidence\": \"database:0.194,experiments:0.089,textmining:0.005\", \"String_Score_Physical\": 295, \"String_Score_Combined\": 299}", "selected": false}, "selected": false}, {"data": {"id": "2784102", "source": "2782872", "target": "2782875", "SCORE": "229", "shared_interaction": "Direct", "source_original": "855130", "Name_B": "TPK2", "Name_A": "YPK2", "target_original": "855898", "Gene_B": "855898", "interaction": "Direct", "SUID": 2784102, "TYPE": "Direct", "Gene_A": "855130", "support": "{\"StringDB\": \"physical\", \"StringEvidence\": \"textmining:0.285,experiments:0.197,coexpression:0.067\", \"String_Score_Physical\": 229, \"String_Score_Combined\": 308}", "selected": false}, "selected": false}, {"data": {"id": "2783970", "source": "2782869", "target": "2782875", "SCORE": "927", "shared_interaction": "Direct", "source_original": "853803", "Name_B": "TPK2", "Name_A": "MSN4", "target_original": "855898", "Gene_B": "855898", "interaction": "Direct", "SUID": 2783970, "TYPE": "Direct", "Gene_A": "853803", "support": "{\"StringDB\": \"physical\", \"StringEvidence\": \"database:0.833,textmining:0.771,experiments:0.550,coexpression:0.053\", \"BioGrid\": \"genetic,physical\", \"BioGrid_PubMed\": \"9649426,21749328,20489023\", \"BioGrid_Type\": \"Affinity Capture-MS,Synthetic Rescue,Synthetic Growth Defect\", \"BioGrid_Score\": \"1.0\", \"String_Score_Physical\": 927, \"String_Score_Combined\": 984}", "selected": false}, "selected": false}, {"data": {"id": "2783967", "source": "2782869", "target": "2782872", "SCORE": "319", "shared_interaction": "Direct", "source_original": "853803", "Name_B": "YPK2", "Name_A": "MSN4", "target_original": "855130", "Gene_B": "855130", "interaction": "Direct", "SUID": 2783967, "TYPE": "Direct", "Gene_A": "853803", "support": "{\"StringDB\": \"physical\", \"StringEvidence\": \"database:0.267,textmining:0.052,experiments:0.032\", \"String_Score_Physical\": 319, \"String_Score_Combined\": 354}", "selected": false}, "selected": false}, {"data": {"id": "2783925", "source": "2782866", "target": "2782875", "SCORE": "991", "shared_interaction": "Direct", "source_original": "853275", "Name_B": "TPK2", "Name_A": "TPK1", "target_original": "855898", "Gene_B": "855898", "interaction": "Direct", "SUID": 2783925, "TYPE": "Direct", "Gene_A": "853275", "support": "{\"StringDB\": \"physical\", \"StringEvidence\": \"textmining:0.976,experiments:0.912,database:0.896,coexpression:0.831,cooccurrence:0.425\", \"BioGrid\": \"genetic,physical\", \"BioGrid_PubMed\": \"9738892,3036373,23155055,21457714,21108829,20489023,19269370,17916074,16554755,16319894,11805837\", \"BioGrid_Type\": \"Affinity Capture-MS,Synthetic Lethality,Phenotypic Suppression,Phenotypic Enhancement,Negative Genetic,Biochemical Activity\", \"BioGrid_Score\": \"1.0,-2.696966\", \"String_Score_Physical\": 991, \"String_Score_Combined\": 998}", "selected": false}, "selected": false}, {"data": {"id": "2783922", "source": "2782866", "target": "2782872", "SCORE": "229", "shared_interaction": "Direct", "source_original": "853275", "Name_B": "YPK2", "Name_A": "TPK1", "target_original": "855130", "Gene_B": "855130", "interaction": "Direct", "SUID": 2783922, "TYPE": "Direct", "Gene_A": "853275", "support": "{\"StringDB\": \"physical\", \"StringEvidence\": \"textmining:0.461,experiments:0.197,coexpression:0.009\", \"String_Score_Physical\": 229, \"String_Score_Combined\": 280}", "selected": false}, "selected": false}, {"data": {"id": "2783919", "source": "2782866", "target": "2782869", "SCORE": "919", "shared_interaction": "Direct", "source_original": "853275", "Name_B": "MSN4", "Name_A": "TPK1", "target_original": "853803", "Gene_B": "853803", "interaction": "Direct", "SUID": 2783919, "TYPE": "Direct", "Gene_A": "853275", "support": "{\"StringDB\": \"physical\", \"StringEvidence\": \"database:0.833,textmining:0.771,experiments:0.500,coexpression:0.053\", \"BioGrid\": \"genetic,physical\", \"BioGrid_PubMed\": \"21757539,21749328,16319894\", \"BioGrid_Type\": \"Phenotypic Enhancement,Biochemical Activity,Synthetic Growth Defect\", \"String_Score_Physical\": 919, \"String_Score_Combined\": 982}", "selected": false}, "selected": false}, {"data": {"id": "2783964", "source": "2782857", "target": "2782860", "SCORE": "915", "shared_interaction": "Direct", "source_original": "853777", "Name_B": "MDH2", "Name_A": "MDH1", "target_original": "853994", "Gene_B": "853994", "interaction": "Direct", "SUID": 2783964, "TYPE": "Direct", "Gene_A": "853777", "support": "{\"StringDB\": \"physical\", \"StringEvidence\": \"textmining:0.905,database:0.791,experiments:0.579,cooccurrence:0.424\", \"BioGrid\": \"genetic,physical\", \"BioGrid_PubMed\": \"31548421,1986231,1447211\", \"BioGrid_Type\": \"Affinity Capture-MS,Synthetic Growth Defect\", \"String_Score_Physical\": 915, \"String_Score_Combined\": 927}", "selected": false}, "selected": false}, {"data": {"id": "2783946", "source": "2782854", "target": "2782863", "SCORE": "278", "shared_interaction": "Direct", "source_original": "853709", "Name_B": "ATP15", "Name_A": "SDH1", "target_original": "855857", "Gene_B": "855857", "interaction": "Direct", "SUID": 2783946, "TYPE": "Direct", "Gene_A": "853709", "support": "{\"StringDB\": \"physical\", \"StringEvidence\": \"experiments:0.248,textmining:0.161,coexpression:0.085\", \"BioGrid\": \"genetic\", \"BioGrid_PubMed\": \"21623372\", \"BioGrid_Type\": \"Positive Genetic\", \"BioGrid_Score\": \"0.208\", \"String_Score_Physical\": 278, \"String_Score_Combined\": 446}", "selected": false}, "selected": false}, {"data": {"id": "2783943", "source": "2782854", "target": "2782860", "SCORE": "191", "shared_interaction": "Direct", "source_original": "853709", "Name_B": "MDH2", "Name_A": "SDH1", "target_original": "853994", "Gene_B": "853994", "interaction": "Direct", "SUID": 2783943, "TYPE": "Direct", "Gene_A": "853709", "support": "{\"StringDB\": \"physical\", \"StringEvidence\": \"textmining:0.590,experiments:0.156,coexpression:0.155,neighborhood:0.080\", \"String_Score_Physical\": 191, \"String_Score_Combined\": 742}", "selected": false}, "selected": false}, {"data": {"id": "2783940", "source": "2782854", "target": "2782857", "SCORE": "430", "shared_interaction": "Direct", "source_original": "853709", "Name_B": "MDH1", "Name_A": "SDH1", "target_original": "853777", "Gene_B": "853777", "interaction": "Direct", "SUID": 2783940, "TYPE": "Direct", "Gene_A": "853709", "support": "{\"StringDB\": \"physical\", \"StringEvidence\": \"coexpression:0.769,textmining:0.550,experiments:0.406,neighborhood:0.080\", \"BioGrid\": \"physical\", \"BioGrid_PubMed\": \"31548421\", \"BioGrid_Type\": \"Affinity Capture-MS\", \"String_Score_Physical\": 430, \"String_Score_Combined\": 945}", "selected": false}, "selected": false}, {"data": {"id": "2783937", "source": "2782851", "target": "2782863", "SCORE": "999", "shared_interaction": "Direct", "source_original": "853585", "Name_B": "ATP15", "Name_A": "ATP2", "target_original": "855857", "Gene_B": "855857", "interaction": "Direct", "SUID": 2783937, "TYPE": "Direct", "Gene_A": "853585", "support": "{\"StringDB\": \"physical\", \"StringEvidence\": \"experiments:1.000,database:0.896,textmining:0.856,coexpression:0.098\", \"BioGrid\": \"genetic,physical\", \"BioGrid_PubMed\": \"22940862,21748405,21623372,18937496,18722382,15581895\", \"BioGrid_Type\": \"Affinity Capture-MS,Co-purification,Positive Genetic\", \"BioGrid_Score\": \"0.2796\", \"String_Score_Physical\": 999, \"String_Score_Combined\": 999}", "selected": false}, "selected": false}, {"data": {"id": "2783934", "source": "2782851", "target": "2782860", "SCORE": "234", "shared_interaction": "Direct", "source_original": "853585", "Name_B": "MDH2", "Name_A": "ATP2", "target_original": "853994", "Gene_B": "853994", "interaction": "Direct", "SUID": 2783934, "TYPE": "Direct", "Gene_A": "853585", "support": "{\"StringDB\": \"physical\", \"StringEvidence\": \"coexpression:0.629,textmining:0.568,experiments:0.201,neighborhood:0.099\", \"String_Score_Physical\": 234, \"String_Score_Combined\": 889}", "selected": false}, "selected": false}, {"data": {"id": "2783931", "source": "2782851", "target": "2782857", "SCORE": "258", "shared_interaction": "Direct", "source_original": "853585", "Name_B": "MDH1", "Name_A": "ATP2", "target_original": "853777", "Gene_B": "853777", "interaction": "Direct", "SUID": 2783931, "TYPE": "Direct", "Gene_A": "853585", "support": "{\"StringDB\": \"physical\", \"StringEvidence\": \"coexpression:0.923,textmining:0.880,experiments:0.226,neighborhood:0.099\", \"String_Score_Physical\": 258, \"String_Score_Combined\": 993}", "selected": false}, "selected": false}, {"data": {"id": "2783715", "source": "2782848", "target": "2782851", "SCORE": "193", "shared_interaction": "Direct", "source_original": "852250", "Name_B": "ATP2", "Name_A": "PET9", "target_original": "853585", "Gene_B": "853585", "interaction": "Direct", "SUID": 2783715, "TYPE": "Direct", "Gene_A": "852250", "support": "{\"StringDB\": \"physical\", \"StringEvidence\": \"coexpression:0.978,textmining:0.654,experiments:0.158\", \"String_Score_Physical\": 193, \"String_Score_Combined\": 993}", "selected": false}, "selected": false}, {"data": {"id": "2783511", "source": "2782845", "target": "2782863", "SCORE": "999", "shared_interaction": "Direct", "source_original": "851892", "Name_B": "ATP15", "Name_A": "ATP5", "target_original": "855857", "Gene_B": "855857", "interaction": "Direct", "SUID": 2783511, "TYPE": "Direct", "Gene_A": "851892", "support": "{\"StringDB\": \"physical\", \"StringEvidence\": \"experiments:0.999,database:0.896,textmining:0.862,coexpression:0.692\", \"BioGrid\": \"genetic,physical\", \"BioGrid_PubMed\": \"22940862,21623372\", \"BioGrid_Type\": \"Affinity Capture-MS,Positive Genetic\", \"BioGrid_Score\": \"0.2176\", \"String_Score_Physical\": 999, \"String_Score_Combined\": 999}", "selected": false}, "selected": false}, {"data": {"id": "2783508", "source": "2782845", "target": "2782860", "SCORE": "199", "shared_interaction": "Direct", "source_original": "851892", "Name_B": "MDH2", "Name_A": "ATP5", "target_original": "853994", "Gene_B": "853994", "interaction": "Direct", "SUID": 2783508, "TYPE": "Direct", "Gene_A": "851892", "support": "{\"StringDB\": \"physical\", \"StringEvidence\": \"coexpression:0.629,textmining:0.174,experiments:0.165,neighborhood:0.120\", \"String_Score_Physical\": 199, \"String_Score_Combined\": 783}", "selected": false}, "selected": false}, {"data": {"id": "2783505", "source": "2782845", "target": "2782857", "SCORE": "439", "shared_interaction": "Direct", "source_original": "851892", "Name_B": "MDH1", "Name_A": "ATP5", "target_original": "853777", "Gene_B": "853777", "interaction": "Direct", "SUID": 2783505, "TYPE": "Direct", "Gene_A": "851892", "support": "{\"StringDB\": \"physical\", \"StringEvidence\": \"coexpression:0.708,experiments:0.416,textmining:0.232,neighborhood:0.120\", \"BioGrid\": \"genetic\", \"BioGrid_PubMed\": \"27708008,20093466\", \"BioGrid_Type\": \"Negative Genetic\", \"BioGrid_Score\": \"-0.161,-0.1507\", \"String_Score_Physical\": 439, \"String_Score_Combined\": 889}", "selected": false}, "selected": false}, {"data": {"id": "2783502", "source": "2782845", "target": "2782851", "SCORE": "999", "shared_interaction": "Direct", "source_original": "851892", "Name_B": "ATP2", "Name_A": "ATP5", "target_original": "853585", "Gene_B": "853585", "interaction": "Direct", "SUID": 2783502, "TYPE": "Direct", "Gene_A": "851892", "support": "{\"StringDB\": \"physical\", \"StringEvidence\": \"experiments:1.000,coexpression:0.970,textmining:0.941,database:0.896,neighborhood:0.126\", \"BioGrid\": \"genetic,physical\", \"BioGrid_PubMed\": \"29093463,22940862,21748405,21623372,18937496,18722382,15581895\", \"BioGrid_Type\": \"Affinity Capture-MS,Co-purification,Positive Genetic,Affinity Capture-Western\", \"BioGrid_Score\": \"0.2233\", \"String_Score_Physical\": 999, \"String_Score_Combined\": 999}", "selected": false}, "selected": false}, {"data": {"id": "2783499", "source": "2782845", "target": "2782848", "SCORE": "189", "shared_interaction": "Direct", "source_original": "851892", "Name_B": "PET9", "Name_A": "ATP5", "target_original": "852250", "Gene_B": "852250", "interaction": "Direct", "SUID": 2783499, "TYPE": "Direct", "Gene_A": "851892", "support": "{\"StringDB\": \"physical\", \"StringEvidence\": \"coexpression:0.648,textmining:0.180,experiments:0.154\", \"String_Score_Physical\": 189, \"String_Score_Combined\": 765}", "selected": false}, "selected": false}, {"data": {"id": "2783415", "source": "2782842", "target": "2782863", "SCORE": "999", "shared_interaction": "Direct", "source_original": "851560", "Name_B": "ATP15", "Name_A": "ATP16", "target_original": "855857", "Gene_B": "855857", "interaction": "Direct", "SUID": 2783415, "TYPE": "Direct", "Gene_A": "851560", "support": "{\"StringDB\": \"physical\", \"StringEvidence\": \"experiments:0.996,textmining:0.992,coexpression:0.915,database:0.896\", \"String_Score_Physical\": 999, \"String_Score_Combined\": 999}", "selected": false}, "selected": false}, {"data": {"id": "2783412", "source": "2782842", "target": "2782860", "SCORE": "222", "shared_interaction": "Direct", "source_original": "851560", "Name_B": "MDH2", "Name_A": "ATP16", "target_original": "853994", "Gene_B": "853994", "interaction": "Direct", "SUID": 2783412, "TYPE": "Direct", "Gene_A": "851560", "support": "{\"StringDB\": \"physical\", \"StringEvidence\": \"coexpression:0.634,experiments:0.189,neighborhood:0.125,textmining:0.088\", \"String_Score_Physical\": 222, \"String_Score_Combined\": 772}", "selected": false}, "selected": false}, {"data": {"id": "2783409", "source": "2782842", "target": "2782857", "SCORE": "243", "shared_interaction": "Direct", "source_original": "851560", "Name_B": "MDH1", "Name_A": "ATP16", "target_original": "853777", "Gene_B": "853777", "interaction": "Direct", "SUID": 2783409, "TYPE": "Direct", "Gene_A": "851560", "support": "{\"StringDB\": \"physical\", \"StringEvidence\": \"coexpression:0.778,textmining:0.344,experiments:0.212,neighborhood:0.125\", \"String_Score_Physical\": 243, \"String_Score_Combined\": 903}", "selected": false}, "selected": false}, {"data": {"id": "2783406", "source": "2782842", "target": "2782854", "SCORE": "217", "shared_interaction": "Direct", "source_original": "851560", "Name_B": "SDH1", "Name_A": "ATP16", "target_original": "853709", "Gene_B": "853709", "interaction": "Direct", "SUID": 2783406, "TYPE": "Direct", "Gene_A": "851560", "support": "{\"StringDB\": \"physical\", \"StringEvidence\": \"textmining:0.382,coexpression:0.296,experiments:0.184,neighborhood:0.007\", \"String_Score_Physical\": 217, \"String_Score_Combined\": 661}", "selected": false}, "selected": false}, {"data": {"id": "2783403", "source": "2782842", "target": "2782851", "SCORE": "999", "shared_interaction": "Direct", "source_original": "851560", "Name_B": "ATP2", "Name_A": "ATP16", "target_original": "853585", "Gene_B": "853585", "interaction": "Direct", "SUID": 2783403, "TYPE": "Direct", "Gene_A": "851560", "support": "{\"StringDB\": \"physical\", \"StringEvidence\": \"experiments:1.000,textmining:0.980,coexpression:0.917,database:0.896,neighborhood:0.126\", \"BioGrid\": \"physical\", \"BioGrid_PubMed\": \"21748405,18937496,18722382,15581895\", \"BioGrid_Type\": \"Co-purification\", \"String_Score_Physical\": 999, \"String_Score_Combined\": 999}", "selected": false}, "selected": false}, {"data": {"id": "2783400", "source": "2782842", "target": "2782845", "SCORE": "999", "shared_interaction": "Direct", "source_original": "851560", "Name_B": "ATP5", "Name_A": "ATP16", "target_original": "851892", "Gene_B": "851892", "interaction": "Direct", "SUID": 2783400, "TYPE": "Direct", "Gene_A": "851560", "support": "{\"StringDB\": \"physical\", \"StringEvidence\": \"experiments:0.995,coexpression:0.957,textmining:0.914,database:0.896,neighborhood:0.156\", \"String_Score_Physical\": 999, \"String_Score_Combined\": 999}", "selected": false}, "selected": false}, {"data": {"id": "2784078", "source": "2782833", "target": "2782836", "SCORE": "999", "shared_interaction": "Direct", "source_original": "854872", "Name_B": "TPS3", "Name_A": "TSL1", "target_original": "855303", "Gene_B": "855303", "interaction": "Direct", "SUID": 2784078, "TYPE": "Direct", "Gene_A": "854872", "support": "{\"StringDB\": \"physical\", \"StringEvidence\": \"experiments:0.999,database:0.896,textmining:0.801,cooccurrence:0.425,coexpression:0.367\", \"BioGrid\": \"genetic,physical\", \"BioGrid_PubMed\": \"9837904,18408719,16941010,16554755\", \"BioGrid_Type\": \"Phenotypic Enhancement,Synthetic Lethality,Affinity Capture-MS,Synthetic Growth Defect\", \"String_Score_Physical\": 999, \"String_Score_Combined\": 999}", "selected": false}, "selected": false}, {"data": {"id": "2784006", "source": "2782830", "target": "2782836", "SCORE": "650", "shared_interaction": "Direct", "source_original": "853932", "Name_B": "TPS3", "Name_A": "GLG1", "target_original": "855303", "Gene_B": "855303", "interaction": "Direct", "SUID": 2784006, "TYPE": "Direct", "Gene_A": "853932", "support": "{\"StringDB\": \"physical\", \"StringEvidence\": \"database:0.635,textmining:0.036\", \"String_Score_Physical\": 650, \"String_Score_Combined\": 662}", "selected": false}, "selected": false}, {"data": {"id": "2784003", "source": "2782830", "target": "2782833", "SCORE": "650", "shared_interaction": "Direct", "source_original": "853932", "Name_B": "TSL1", "Name_A": "GLG1", "target_original": "854872", "Gene_B": "854872", "interaction": "Direct", "SUID": 2784003, "TYPE": "Direct", "Gene_A": "853932", "support": "{\"StringDB\": \"physical\", \"StringEvidence\": \"database:0.635,coexpression:0.168\", \"String_Score_Physical\": 650, \"String_Score_Combined\": 708}", "selected": false}, "selected": false}, {"data": {"id": "2783778", "source": "2782827", "target": "2782836", "SCORE": "999", "shared_interaction": "Direct", "source_original": "852423", "Name_B": "TPS3", "Name_A": "TPS1", "target_original": "855303", "Gene_B": "855303", "interaction": "Direct", "SUID": 2783778, "TYPE": "Direct", "Gene_A": "852423", "support": "{\"StringDB\": \"physical\", \"StringEvidence\": \"experiments:0.999,database:0.896,textmining:0.878,coexpression:0.626,cooccurrence:0.425\", \"BioGrid\": \"genetic,physical\", \"BioGrid_PubMed\": \"9194697,18719252,16941010,16554755,16429126,11805826\", \"BioGrid_Type\": \"Affinity Capture-MS,Synthetic Lethality,Two-hybrid\", \"String_Score_Physical\": 999, \"String_Score_Combined\": 999}", "selected": false}, "selected": false}, {"data": {"id": "2783775", "source": "2782827", "target": "2782833", "SCORE": "999", "shared_interaction": "Direct", "source_original": "852423", "Name_B": "TSL1", "Name_A": "TPS1", "target_original": "854872", "Gene_B": "854872", "interaction": "Direct", "SUID": 2783775, "TYPE": "Direct", "Gene_A": "852423", "support": "{\"StringDB\": \"physical\", \"StringEvidence\": \"experiments:0.999,coexpression:0.963,database:0.896,textmining:0.777,cooccurrence:0.425\", \"BioGrid\": \"genetic,physical\", \"BioGrid_PubMed\": \"9194697,18719252,16941010,16554755,16429126,11805826\", \"BioGrid_Type\": \"Affinity Capture-MS,Synthetic Lethality,Two-hybrid\", \"String_Score_Physical\": 999, \"String_Score_Combined\": 999}", "selected": false}, "selected": false}, {"data": {"id": "2783772", "source": "2782827", "target": "2782830", "SCORE": "740", "shared_interaction": "Direct", "source_original": "852423", "Name_B": "GLG1", "Name_A": "TPS1", "target_original": "853932", "Gene_B": "853932", "interaction": "Direct", "SUID": 2783772, "TYPE": "Direct", "Gene_A": "852423", "support": "{\"StringDB\": \"physical\", \"StringEvidence\": \"database:0.635,experiments:0.259,coexpression:0.063\", \"BioGrid\": \"genetic\", \"BioGrid_PubMed\": \"16004992\", \"BioGrid_Type\": \"Phenotypic Enhancement\", \"String_Score_Physical\": 740, \"String_Score_Combined\": 756}", "selected": false}, "selected": false}, {"data": {"id": "2783727", "source": "2782824", "target": "2782836", "SCORE": "914", "shared_interaction": "Direct", "source_original": "852286", "Name_B": "TPS3", "Name_A": "NTH2", "target_original": "855303", "Gene_B": "855303", "interaction": "Direct", "SUID": 2783727, "TYPE": "Direct", "Gene_A": "852286", "support": "{\"StringDB\": \"physical\", \"StringEvidence\": \"database:0.896,textmining:0.838,coexpression:0.169,experiments:0.149,neighborhood:0.013\", \"String_Score_Physical\": 914, \"String_Score_Combined\": 988}", "selected": false}, "selected": false}, {"data": {"id": "2783724", "source": "2782824", "target": "2782833", "SCORE": "914", "shared_interaction": "Direct", "source_original": "852286", "Name_B": "TSL1", "Name_A": "NTH2", "target_original": "854872", "Gene_B": "854872", "interaction": "Direct", "SUID": 2783724, "TYPE": "Direct", "Gene_A": "852286", "support": "{\"StringDB\": \"physical\", \"StringEvidence\": \"database:0.896,textmining:0.449,coexpression:0.341,experiments:0.149,neighborhood:0.013\", \"String_Score_Physical\": 914, \"String_Score_Combined\": 969}", "selected": false}, "selected": false}, {"data": {"id": "2783721", "source": "2782824", "target": "2782827", "SCORE": "726", "shared_interaction": "Direct", "source_original": "852286", "Name_B": "TPS1", "Name_A": "NTH2", "target_original": "852423", "Gene_B": "852423", "interaction": "Direct", "SUID": 2783721, "TYPE": "Direct", "Gene_A": "852286", "support": "{\"StringDB\": \"physical\", \"StringEvidence\": \"coexpression:0.909,textmining:0.750,experiments:0.715,neighborhood:0.013\", \"BioGrid\": \"genetic\", \"BioGrid_PubMed\": \"20159575,19180643\", \"BioGrid_Type\": \"Phenotypic Enhancement,Dosage Rescue\", \"String_Score_Physical\": 726, \"String_Score_Combined\": 993}", "selected": false}, "selected": false}, {"data": {"id": "2783460", "source": "2782821", "target": "2782839", "SCORE": "259", "shared_interaction": "Direct", "source_original": "851670", "Name_B": "GPA2", "Name_A": "GIS1", "target_original": "856741", "Gene_B": "856741", "interaction": "Direct", "SUID": 2783460, "TYPE": "Direct", "Gene_A": "851670", "support": "{\"StringDB\": \"physical\", \"StringEvidence\": \"textmining:0.233,database:0.195,experiments:0.041,coexpression:0.009\", \"String_Score_Physical\": 259, \"String_Score_Combined\": 437}", "selected": false}, "selected": false}, {"data": {"id": "2783457", "source": "2782818", "target": "2782836", "SCORE": "999", "shared_interaction": "Direct", "source_original": "851646", "Name_B": "TPS3", "Name_A": "TPS2", "target_original": "855303", "Gene_B": "855303", "interaction": "Direct", "SUID": 2783457, "TYPE": "Direct", "Gene_A": "851646", "support": "{\"StringDB\": \"physical\", \"StringEvidence\": \"experiments:0.999,textmining:0.992,database:0.896,cooccurrence:0.424,coexpression:0.285\", \"BioGrid\": \"physical\", \"BioGrid_PubMed\": \"9194697,20489023,16554755,16429126\", \"BioGrid_Type\": \"Two-hybrid,Affinity Capture-MS\", \"BioGrid_Score\": \"0.959\", \"String_Score_Physical\": 999, \"String_Score_Combined\": 999}", "selected": false}, "selected": false}, {"data": {"id": "2783454", "source": "2782818", "target": "2782833", "SCORE": "977", "shared_interaction": "Direct", "source_original": "851646", "Name_B": "TSL1", "Name_A": "TPS2", "target_original": "854872", "Gene_B": "854872", "interaction": "Direct", "SUID": 2783454, "TYPE": "Direct", "Gene_A": "851646", "support": "{\"StringDB\": \"physical\", \"StringEvidence\": \"coexpression:0.972,database:0.896,textmining:0.789,experiments:0.773,cooccurrence:0.424\", \"BioGrid\": \"genetic,physical\", \"BioGrid_PubMed\": \"9194697,21623372,16554755,16429126\", \"BioGrid_Type\": \"Two-hybrid,Affinity Capture-MS,Positive Genetic\", \"BioGrid_Score\": \"0.0945,0.1279\", \"String_Score_Physical\": 977, \"String_Score_Combined\": 999}", "selected": false}, "selected": false}, {"data": {"id": "2783451", "source": "2782818", "target": "2782830", "SCORE": "650", "shared_interaction": "Direct", "source_original": "851646", "Name_B": "GLG1", "Name_A": "TPS2", "target_original": "853932", "Gene_B": "853932", "interaction": "Direct", "SUID": 2783451, "TYPE": "Direct", "Gene_A": "851646", "support": "{\"StringDB\": \"physical\", \"StringEvidence\": \"database:0.635,coexpression:0.163,textmining:0.034\", \"String_Score_Physical\": 650, \"String_Score_Combined\": 717}", "selected": false}, "selected": false}, {"data": {"id": "2783448", "source": "2782818", "target": "2782827", "SCORE": "992", "shared_interaction": "Direct", "source_original": "851646", "Name_B": "TPS1", "Name_A": "TPS2", "target_original": "852423", "Gene_B": "852423", "interaction": "Direct", "SUID": 2783448, "TYPE": "Direct", "Gene_A": "851646", "support": "{\"StringDB\": \"physical\", \"StringEvidence\": \"textmining:0.937,coexpression:0.932,experiments:0.926,database:0.896,cooccurrence:0.425\", \"BioGrid\": \"genetic,physical\", \"BioGrid_PubMed\": \"9194697,8809751,27708008,21623372,20489023,20093466,19620241,16554755,16429126,11805826\", \"BioGrid_Type\": \"Affinity Capture-MS,Two-hybrid,Positive Genetic,Phenotypic Suppression,Phenotypic Enhancement\", \"BioGrid_Score\": \"0.2055,0.956,0.1113,0.2632\", \"String_Score_Physical\": 992, \"String_Score_Combined\": 999}", "selected": false}, "selected": false}, {"data": {"id": "2783445", "source": "2782818", "target": "2782824", "SCORE": "949", "shared_interaction": "Direct", "source_original": "851646", "Name_B": "NTH2", "Name_A": "TPS2", "target_original": "852286", "Gene_B": "852286", "interaction": "Direct", "SUID": 2783445, "TYPE": "Direct", "Gene_A": "851646", "support": "{\"StringDB\": \"physical\", \"StringEvidence\": \"database:0.896,textmining:0.788,experiments:0.492,coexpression:0.203,neighborhood:0.013\", \"BioGrid\": \"genetic\", \"BioGrid_PubMed\": \"20159575,19180643\", \"BioGrid_Type\": \"Phenotypic Enhancement,Dosage Rescue\", \"String_Score_Physical\": 949, \"String_Score_Combined\": 991}", "selected": false}, "selected": false}, {"data": {"id": "2783430", "source": "2782815", "target": "2782836", "SCORE": "916", "shared_interaction": "Direct", "source_original": "851564", "Name_B": "TPS3", "Name_A": "NTH1", "target_original": "855303", "Gene_B": "855303", "interaction": "Direct", "SUID": 2783430, "TYPE": "Direct", "Gene_A": "851564", "support": "{\"StringDB\": \"physical\", \"StringEvidence\": \"textmining:0.913,database:0.896,coexpression:0.250,experiments:0.162,neighborhood:0.013\", \"BioGrid\": \"genetic\", \"BioGrid_PubMed\": \"21623372\", \"BioGrid_Type\": \"Positive Genetic\", \"BioGrid_Score\": \"0.1069\", \"String_Score_Physical\": 916, \"String_Score_Combined\": 994}", "selected": false}, "selected": false}, {"data": {"id": "2783427", "source": "2782815", "target": "2782833", "SCORE": "914", "shared_interaction": "Direct", "source_original": "851564", "Name_B": "TSL1", "Name_A": "NTH1", "target_original": "854872", "Gene_B": "854872", "interaction": "Direct", "SUID": 2783427, "TYPE": "Direct", "Gene_A": "851564", "support": "{\"StringDB\": \"physical\", \"StringEvidence\": \"database:0.896,coexpression:0.876,textmining:0.449,experiments:0.149,neighborhood:0.013\", \"String_Score_Physical\": 914, \"String_Score_Combined\": 994}", "selected": false}, "selected": false}, {"data": {"id": "2783424", "source": "2782815", "target": "2782827", "SCORE": "764", "shared_interaction": "Direct", "source_original": "851564", "Name_B": "TPS1", "Name_A": "NTH1", "target_original": "852423", "Gene_B": "852423", "interaction": "Direct", "SUID": 2783424, "TYPE": "Direct", "Gene_A": "851564", "support": "{\"StringDB\": \"physical\", \"StringEvidence\": \"coexpression:0.982,experiments:0.754,textmining:0.750,neighborhood:0.013\", \"BioGrid\": \"genetic\", \"BioGrid_PubMed\": \"31347835,20159575,19180643\", \"BioGrid_Type\": \"Phenotypic Enhancement,Dosage Rescue\", \"String_Score_Physical\": 764, \"String_Score_Combined\": 998}", "selected": false}, "selected": false}, {"data": {"id": "2783421", "source": "2782815", "target": "2782824", "SCORE": "959", "shared_interaction": "Direct", "source_original": "851564", "Name_B": "NTH2", "Name_A": "NTH1", "target_original": "852286", "Gene_B": "852286", "interaction": "Direct", "SUID": 2783421, "TYPE": "Direct", "Gene_A": "851564", "support": "{\"StringDB\": \"physical\", \"StringEvidence\": \"textmining:0.917,experiments:0.797,database:0.791,cooccurrence:0.425,coexpression:0.156\", \"BioGrid\": \"genetic,physical\", \"BioGrid_PubMed\": \"19180643,18408719,16554755\", \"BioGrid_Type\": \"Affinity Capture-MS,Synthetic Growth Defect,Phenotypic Enhancement\", \"String_Score_Physical\": 959, \"String_Score_Combined\": 966}", "selected": false}, "selected": false}, {"data": {"id": "2783418", "source": "2782815", "target": "2782818", "SCORE": "949", "shared_interaction": "Direct", "source_original": "851564", "Name_B": "TPS2", "Name_A": "NTH1", "target_original": "851646", "Gene_B": "851646", "interaction": "Direct", "SUID": 2783418, "TYPE": "Direct", "Gene_A": "851564", "support": "{\"StringDB\": \"physical\", \"StringEvidence\": \"database:0.896,textmining:0.850,coexpression:0.824,experiments:0.492,neighborhood:0.013\", \"BioGrid\": \"genetic\", \"BioGrid_PubMed\": \"20159575,19180643\", \"BioGrid_Type\": \"Phenotypic Enhancement,Dosage Rescue\", \"String_Score_Physical\": 949, \"String_Score_Combined\": 998}", "selected": false}, "selected": false}, {"data": {"id": "2783391", "source": "2782812", "target": "2782839", "SCORE": "428", "shared_interaction": "Direct", "source_original": "851480", "Name_B": "GPA2", "Name_A": "MRK1", "target_original": "856741", "Gene_B": "856741", "interaction": "Direct", "SUID": 2783391, "TYPE": "Direct", "Gene_A": "851480", "support": "{\"StringDB\": \"physical\", \"StringEvidence\": \"database:0.229,experiments:0.226,textmining:0.045,coexpression:0.009\", \"String_Score_Physical\": 428, \"String_Score_Combined\": 458}", "selected": false}, "selected": false}, {"data": {"id": "2783388", "source": "2782812", "target": "2782821", "SCORE": "315", "shared_interaction": "Direct", "source_original": "851480", "Name_B": "GIS1", "Name_A": "MRK1", "target_original": "851670", "Gene_B": "851670", "interaction": "Direct", "SUID": 2783388, "TYPE": "Direct", "Gene_A": "851480", "support": "{\"StringDB\": \"physical\", \"StringEvidence\": \"database:0.197,coexpression:0.119,experiments:0.111,textmining:0.088\", \"String_Score_Physical\": 315, \"String_Score_Combined\": 450}", "selected": false}, "selected": false}, {"data": {"id": "2783307", "source": "2782809", "target": "2782836", "SCORE": "650", "shared_interaction": "Direct", "source_original": "850962", "Name_B": "TPS3", "Name_A": "GSY2", "target_original": "855303", "Gene_B": "855303", "interaction": "Direct", "SUID": 2783307, "TYPE": "Direct", "Gene_A": "850962", "support": "{\"StringDB\": \"physical\", \"StringEvidence\": \"textmining:0.673,database:0.635,coexpression:0.245,neighborhood:0.009\", \"String_Score_Physical\": 650, \"String_Score_Combined\": 914}", "selected": false}, "selected": false}, {"data": {"id": "2783304", "source": "2782809", "target": "2782833", "SCORE": "650", "shared_interaction": "Direct", "source_original": "850962", "Name_B": "TSL1", "Name_A": "GSY2", "target_original": "854872", "Gene_B": "854872", "interaction": "Direct", "SUID": 2783304, "TYPE": "Direct", "Gene_A": "850962", "support": "{\"StringDB\": \"physical\", \"StringEvidence\": \"coexpression:0.969,database:0.635,textmining:0.270,neighborhood:0.009\", \"String_Score_Physical\": 650, \"String_Score_Combined\": 992}", "selected": false}, "selected": false}, {"data": {"id": "2783301", "source": "2782809", "target": "2782830", "SCORE": "986", "shared_interaction": "Direct", "source_original": "850962", "Name_B": "GLG1", "Name_A": "GSY2", "target_original": "853932", "Gene_B": "853932", "interaction": "Direct", "SUID": 2783301, "TYPE": "Direct", "Gene_A": "850962", "support": "{\"StringDB\": \"physical\", \"StringEvidence\": \"experiments:0.935,textmining:0.933,database:0.791,coexpression:0.380\", \"BioGrid\": \"physical\", \"BioGrid_PubMed\": \"18467557,16554755\", \"BioGrid_Type\": \"Affinity Capture-MS,PCA\", \"String_Score_Physical\": 986, \"String_Score_Combined\": 999}", "selected": false}, "selected": false}, {"data": {"id": "2783298", "source": "2782809", "target": "2782827", "SCORE": "791", "shared_interaction": "Direct", "source_original": "850962", "Name_B": "TPS1", "Name_A": "GSY2", "target_original": "852423", "Gene_B": "852423", "interaction": "Direct", "SUID": 2783298, "TYPE": "Direct", "Gene_A": "850962", "support": "{\"StringDB\": \"physical\", \"StringEvidence\": \"coexpression:0.901,database:0.635,textmining:0.542,experiments:0.404,neighborhood:0.009\", \"BioGrid\": \"genetic\", \"BioGrid_PubMed\": \"31960994,27923067\", \"BioGrid_Type\": \"Phenotypic Enhancement\", \"String_Score_Physical\": 791, \"String_Score_Combined\": 990}", "selected": false}, "selected": false}, {"data": {"id": "2783295", "source": "2782809", "target": "2782818", "SCORE": "655", "shared_interaction": "Direct", "source_original": "850962", "Name_B": "TPS2", "Name_A": "GSY2", "target_original": "851646", "Gene_B": "851646", "interaction": "Direct", "SUID": 2783295, "TYPE": "Direct", "Gene_A": "850962", "support": "{\"StringDB\": \"physical\", \"StringEvidence\": \"coexpression:0.931,textmining:0.708,database:0.635,experiments:0.015,neighborhood:0.009\", \"BioGrid\": \"genetic\", \"BioGrid_PubMed\": \"21623372\", \"BioGrid_Type\": \"Positive Genetic\", \"BioGrid_Score\": \"0.1022\", \"String_Score_Physical\": 655, \"String_Score_Combined\": 993}", "selected": false}, "selected": false}, {"data": {"id": "2783292", "source": "2782809", "target": "2782812", "SCORE": "527", "shared_interaction": "Direct", "source_original": "850962", "Name_B": "MRK1", "Name_A": "GSY2", "target_original": "851480", "Gene_B": "851480", "interaction": "Direct", "SUID": 2783292, "TYPE": "Direct", "Gene_A": "850962", "support": "{\"StringDB\": \"physical\", \"StringEvidence\": \"database:0.398,experiments:0.181,coexpression:0.110,textmining:0.100,neighborhood:0.014\", \"String_Score_Physical\": 527, \"String_Score_Combined\": 626}", "selected": false}, "selected": false}, {"data": {"id": "2783097", "source": "2782806", "target": "2782836", "SCORE": "650", "shared_interaction": "Direct", "source_original": "850569", "Name_B": "TPS3", "Name_A": "GSY1", "target_original": "855303", "Gene_B": "855303", "interaction": "Direct", "SUID": 2783097, "TYPE": "Direct", "Gene_A": "850569", "support": "{\"StringDB\": \"physical\", \"StringEvidence\": \"textmining:0.734,database:0.635,coexpression:0.142,neighborhood:0.009\", \"String_Score_Physical\": 650, \"String_Score_Combined\": 920}", "selected": false}, "selected": false}, {"data": {"id": "2783094", "source": "2782806", "target": "2782833", "SCORE": "650", "shared_interaction": "Direct", "source_original": "850569", "Name_B": "TSL1", "Name_A": "GSY1", "target_original": "854872", "Gene_B": "854872", "interaction": "Direct", "SUID": 2783094, "TYPE": "Direct", "Gene_A": "850569", "support": "{\"StringDB\": \"physical\", \"StringEvidence\": \"coexpression:0.914,database:0.635,textmining:0.270,neighborhood:0.009\", \"String_Score_Physical\": 650, \"String_Score_Combined\": 978}", "selected": false}, "selected": false}, {"data": {"id": "2783091", "source": "2782806", "target": "2782830", "SCORE": "972", "shared_interaction": "Direct", "source_original": "850569", "Name_B": "GLG1", "Name_A": "GSY1", "target_original": "853932", "Gene_B": "853932", "interaction": "Direct", "SUID": 2783091, "TYPE": "Direct", "Gene_A": "850569", "support": "{\"StringDB\": \"physical\", \"StringEvidence\": \"textmining:0.943,experiments:0.861,database:0.791,coexpression:0.594\", \"BioGrid\": \"physical\", \"BioGrid_PubMed\": \"16554755\", \"BioGrid_Type\": \"Affinity Capture-MS\", \"String_Score_Physical\": 972, \"String_Score_Combined\": 999}", "selected": false}, "selected": false}, {"data": {"id": "2783088", "source": "2782806", "target": "2782827", "SCORE": "740", "shared_interaction": "Direct", "source_original": "850569", "Name_B": "TPS1", "Name_A": "GSY1", "target_original": "852423", "Gene_B": "852423", "interaction": "Direct", "SUID": 2783088, "TYPE": "Direct", "Gene_A": "850569", "support": "{\"StringDB\": \"physical\", \"StringEvidence\": \"coexpression:0.839,database:0.635,textmining:0.542,experiments:0.259,neighborhood:0.009\", \"BioGrid\": \"genetic\", \"BioGrid_PubMed\": \"31960994\", \"BioGrid_Type\": \"Phenotypic Enhancement\", \"String_Score_Physical\": 740, \"String_Score_Combined\": 981}", "selected": false}, "selected": false}, {"data": {"id": "2783085", "source": "2782806", "target": "2782818", "SCORE": "650", "shared_interaction": "Direct", "source_original": "850569", "Name_B": "TPS2", "Name_A": "GSY1", "target_original": "851646", "Gene_B": "851646", "interaction": "Direct", "SUID": 2783085, "TYPE": "Direct", "Gene_A": "850569", "support": "{\"StringDB\": \"physical\", \"StringEvidence\": \"coexpression:0.870,textmining:0.807,database:0.635,neighborhood:0.009\", \"String_Score_Physical\": 650, \"String_Score_Combined\": 991}", "selected": false}, "selected": false}, {"data": {"id": "2783082", "source": "2782806", "target": "2782812", "SCORE": "527", "shared_interaction": "Direct", "source_original": "850569", "Name_B": "MRK1", "Name_A": "GSY1", "target_original": "851480", "Gene_B": "851480", "interaction": "Direct", "SUID": 2783082, "TYPE": "Direct", "Gene_A": "850569", "support": "{\"StringDB\": \"physical\", \"StringEvidence\": \"database:0.398,coexpression:0.252,experiments:0.181,textmining:0.048,neighborhood:0.014\", \"String_Score_Physical\": 527, \"String_Score_Combined\": 668}", "selected": false}, "selected": false}, {"data": {"id": "2783079", "source": "2782806", "target": "2782809", "SCORE": "999", "shared_interaction": "Direct", "source_original": "850569", "Name_B": "GSY2", "Name_A": "GSY1", "target_original": "850962", "Gene_B": "850962", "interaction": "Direct", "SUID": 2783079, "TYPE": "Direct", "Gene_A": "850569", "support": "{\"StringDB\": \"physical\", \"StringEvidence\": \"experiments:0.999,coexpression:0.919,textmining:0.904,database:0.896,cooccurrence:0.425\", \"BioGrid\": \"genetic,physical\", \"BioGrid_PubMed\": \"20826334,18408719,16941010,16554755,11283351\", \"BioGrid_Type\": \"Affinity Capture-MS,Synthetic Lethality,Two-hybrid,Synthetic Growth Defect,Affinity Capture-Western\", \"String_Score_Physical\": 999, \"String_Score_Combined\": 999}", "selected": false}, "selected": false}, {"data": {"id": "2783055", "source": "2782803", "target": "2782797", "SCORE": "958", "shared_interaction": "Direct", "source_original": "3799970", "Name_B": "RIP1", "Name_A": "COX26", "target_original": "856689", "Gene_B": "856689", "interaction": "Direct", "SUID": 2783055, "TYPE": "Direct", "Gene_A": "3799970", "support": "{\"StringDB\": \"physical\", \"StringEvidence\": \"experiments:0.851,database:0.708,coexpression:0.092\", \"BioGrid\": \"physical\", \"BioGrid_PubMed\": \"27083394\", \"BioGrid_Type\": \"Affinity Capture-Western\", \"String_Score_Physical\": 958, \"String_Score_Combined\": 962}", "selected": false}, "selected": false}, {"data": {"id": "2783052", "source": "2782803", "target": "2782791", "SCORE": "941", "shared_interaction": "Direct", "source_original": "3799970", "Name_B": "QCR10", "Name_A": "COX26", "target_original": "856390", "Gene_B": "856390", "interaction": "Direct", "SUID": 2783052, "TYPE": "Direct", "Gene_A": "3799970", "support": "{\"StringDB\": \"physical\", \"StringEvidence\": \"experiments:0.791,database:0.708,coexpression:0.138\", \"String_Score_Physical\": 941, \"String_Score_Combined\": 949}", "selected": false}, "selected": false}, {"data": {"id": "2783049", "source": "2782803", "target": "2782788", "SCORE": "941", "shared_interaction": "Direct", "source_original": "3799970", "Name_B": "QCR2", "Name_A": "COX26", "target_original": "856321", "Gene_B": "856321", "interaction": "Direct", "SUID": 2783049, "TYPE": "Direct", "Gene_A": "3799970", "support": "{\"StringDB\": \"physical\", \"StringEvidence\": \"experiments:0.791,database:0.708,textmining:0.299,coexpression:0.058\", \"String_Score_Physical\": 941, \"String_Score_Combined\": 961}", "selected": false}, "selected": false}, {"data": {"id": "2783046", "source": "2782803", "target": "2782779", "SCORE": "958", "shared_interaction": "Direct", "source_original": "3799970", "Name_B": "COX5A", "Name_A": "COX26", "target_original": "855675", "Gene_B": "855675", "interaction": "Direct", "SUID": 2783046, "TYPE": "Direct", "Gene_A": "3799970", "support": "{\"StringDB\": \"physical\", \"StringEvidence\": \"experiments:0.851,database:0.708,coexpression:0.086\", \"BioGrid\": \"physical\", \"BioGrid_PubMed\": \"27083394\", \"BioGrid_Type\": \"Affinity Capture-Western\", \"String_Score_Physical\": 958, \"String_Score_Combined\": 961}", "selected": false}, "selected": false}, {"data": {"id": "2783043", "source": "2782803", "target": "2782776", "SCORE": "941", "shared_interaction": "Direct", "source_original": "3799970", "Name_B": "COX7", "Name_A": "COX26", "target_original": "855298", "Gene_B": "855298", "interaction": "Direct", "SUID": 2783043, "TYPE": "Direct", "Gene_A": "3799970", "support": "{\"StringDB\": \"physical\", \"StringEvidence\": \"experiments:0.791,database:0.708,textmining:0.131,coexpression:0.086\", \"String_Score_Physical\": 941, \"String_Score_Combined\": 953}", "selected": false}, "selected": false}, {"data": {"id": "2783040", "source": "2782803", "target": "2782767", "SCORE": "958", "shared_interaction": "Direct", "source_original": "3799970", "Name_B": "COX5B", "Name_A": "COX26", "target_original": "854695", "Gene_B": "854695", "interaction": "Direct", "SUID": 2783040, "TYPE": "Direct", "Gene_A": "3799970", "support": "{\"StringDB\": \"physical\", \"StringEvidence\": \"experiments:0.851,database:0.708\", \"BioGrid\": \"physical\", \"BioGrid_PubMed\": \"27083394\", \"BioGrid_Type\": \"Affinity Capture-Western\", \"String_Score_Physical\": 958, \"String_Score_Combined\": 958}", "selected": false}, "selected": false}, {"data": {"id": "2783037", "source": "2782803", "target": "2782764", "SCORE": "941", "shared_interaction": "Direct", "source_original": "3799970", "Name_B": "CYT1", "Name_A": "COX26", "target_original": "854231", "Gene_B": "854231", "interaction": "Direct", "SUID": 2783037, "TYPE": "Direct", "Gene_A": "3799970", "support": "{\"StringDB\": \"physical\", \"StringEvidence\": \"experiments:0.791,database:0.708,textmining:0.506,coexpression:0.205\", \"String_Score_Physical\": 941, \"String_Score_Combined\": 977}", "selected": false}, "selected": false}, {"data": {"id": "2783034", "source": "2782803", "target": "2782752", "SCORE": "958", "shared_interaction": "Direct", "source_original": "3799970", "Name_B": "QCR8", "Name_A": "COX26", "target_original": "853273", "Gene_B": "853273", "interaction": "Direct", "SUID": 2783034, "TYPE": "Direct", "Gene_A": "3799970", "support": "{\"StringDB\": \"physical\", \"StringEvidence\": \"experiments:0.851,database:0.708,textmining:0.108\", \"BioGrid\": \"physical\", \"BioGrid_PubMed\": \"27083394\", \"BioGrid_Type\": \"Affinity Capture-Western\", \"String_Score_Physical\": 958, \"String_Score_Combined\": 962}", "selected": false}, "selected": false}, {"data": {"id": "2783031", "source": "2782803", "target": "2782749", "SCORE": "941", "shared_interaction": "Direct", "source_original": "3799970", "Name_B": "QCR9", "Name_A": "COX26", "target_original": "853095", "Gene_B": "853095", "interaction": "Direct", "SUID": 2783031, "TYPE": "Direct", "Gene_A": "3799970", "support": "{\"StringDB\": \"physical\", \"StringEvidence\": \"experiments:0.791,database:0.708,coexpression:0.209,textmining:0.131\", \"String_Score_Physical\": 941, \"String_Score_Combined\": 959}", "selected": false}, "selected": false}, {"data": {"id": "2783028", "source": "2782803", "target": "2782746", "SCORE": "966", "shared_interaction": "Direct", "source_original": "3799970", "Name_B": "COX4", "Name_A": "COX26", "target_original": "852688", "Gene_B": "852688", "interaction": "Direct", "SUID": 2783028, "TYPE": "Direct", "Gene_A": "3799970", "support": "{\"StringDB\": \"physical\", \"StringEvidence\": \"experiments:0.879,database:0.708,coexpression:0.106\", \"BioGrid\": \"physical\", \"BioGrid_PubMed\": \"27083394,23864706\", \"BioGrid_Type\": \"Affinity Capture-MS,Affinity Capture-Western\", \"String_Score_Physical\": 966, \"String_Score_Combined\": 969}", "selected": false}, "selected": false}, {"data": {"id": "2783025", "source": "2782803", "target": "2782740", "SCORE": "965", "shared_interaction": "Direct", "source_original": "3799970", "Name_B": "COR1", "Name_A": "COX26", "target_original": "852235", "Gene_B": "852235", "interaction": "Direct", "SUID": 2783025, "TYPE": "Direct", "Gene_A": "3799970", "support": "{\"StringDB\": \"physical\", \"StringEvidence\": \"experiments:0.876,database:0.708,textmining:0.088,coexpression:0.058\", \"BioGrid\": \"physical\", \"BioGrid_PubMed\": \"22342701\", \"BioGrid_Type\": \"Affinity Capture-MS\", \"String_Score_Physical\": 965, \"String_Score_Combined\": 970}", "selected": false}, "selected": false}, {"data": {"id": "2783022", "source": "2782803", "target": "2782734", "SCORE": "941", "shared_interaction": "Direct", "source_original": "3799970", "Name_B": "QCR7", "Name_A": "COX26", "target_original": "852142", "Gene_B": "852142", "interaction": "Direct", "SUID": 2783022, "TYPE": "Direct", "Gene_A": "3799970", "support": "{\"StringDB\": \"physical\", \"StringEvidence\": \"experiments:0.791,database:0.708,coexpression:0.106,textmining:0.083\", \"String_Score_Physical\": 941, \"String_Score_Combined\": 952}", "selected": false}, "selected": false}, {"data": {"id": "2783019", "source": "2782803", "target": "2782722", "SCORE": "941", "shared_interaction": "Direct", "source_original": "3799970", "Name_B": "COX8", "Name_A": "COX26", "target_original": "851111", "Gene_B": "851111", "interaction": "Direct", "SUID": 2783019, "TYPE": "Direct", "Gene_A": "3799970", "support": "{\"StringDB\": \"physical\", \"StringEvidence\": \"experiments:0.791,database:0.708,coexpression:0.058\", \"String_Score_Physical\": 941, \"String_Score_Combined\": 945}", "selected": false}, "selected": false}, {"data": {"id": "2783016", "source": "2782803", "target": "2782716", "SCORE": "994", "shared_interaction": "Direct", "source_original": "3799970", "Name_B": "COX12", "Name_A": "COX26", "target_original": "850727", "Gene_B": "850727", "interaction": "Direct", "SUID": 2783016, "TYPE": "Direct", "Gene_A": "3799970", "support": "{\"StringDB\": \"physical\", \"StringEvidence\": \"experiments:0.981,database:0.708,textmining:0.506,coexpression:0.065\", \"String_Score_Physical\": 994, \"String_Score_Combined\": 997}", "selected": false}, "selected": false}, {"data": {"id": "2783013", "source": "2782803", "target": "2782686", "SCORE": "941", "shared_interaction": "Direct", "source_original": "3799970", "Name_B": "QCR6", "Name_A": "COX26", "target_original": "850593", "Gene_B": "850593", "interaction": "Direct", "SUID": 2783013, "TYPE": "Direct", "Gene_A": "3799970", "support": "{\"StringDB\": \"physical\", \"StringEvidence\": \"experiments:0.791,database:0.708,textmining:0.316,coexpression:0.066\", \"String_Score_Physical\": 941, \"String_Score_Combined\": 962}", "selected": false}, "selected": false}, {"data": {"id": "2783010", "source": "2782800", "target": "2782785", "SCORE": "540", "shared_interaction": "Direct", "source_original": "1466480", "Name_B": "ATP20", "Name_A": "MCO10", "target_original": "856131", "Gene_B": "856131", "interaction": "Direct", "SUID": 2783010, "TYPE": "Direct", "Gene_A": "1466480", "support": "{\"StringDB\": \"physical\", \"StringEvidence\": \"database:0.520\", \"String_Score_Physical\": 540, \"String_Score_Combined\": 540}", "selected": false}, "selected": false}, {"data": {"id": "2783007", "source": "2782800", "target": "2782782", "SCORE": "540", "shared_interaction": "Direct", "source_original": "1466480", "Name_B": "ATP4", "Name_A": "MCO10", "target_original": "856027", "Gene_B": "856027", "interaction": "Direct", "SUID": 2783007, "TYPE": "Direct", "Gene_A": "1466480", "support": "{\"StringDB\": \"physical\", \"StringEvidence\": \"database:0.520\", \"String_Score_Physical\": 540, \"String_Score_Combined\": 540}", "selected": false}, "selected": false}, {"data": {"id": "2783004", "source": "2782800", "target": "2782770", "SCORE": "540", "shared_interaction": "Direct", "source_original": "1466480", "Name_B": "ATP18", "Name_A": "MCO10", "target_original": "854893", "Gene_B": "854893", "interaction": "Direct", "SUID": 2783004, "TYPE": "Direct", "Gene_A": "1466480", "support": "{\"StringDB\": \"physical\", \"StringEvidence\": \"database:0.520\", \"String_Score_Physical\": 540, \"String_Score_Combined\": 540}", "selected": false}, "selected": false}, {"data": {"id": "2783001", "source": "2782800", "target": "2782761", "SCORE": "800", "shared_interaction": "Direct", "source_original": "1466480", "Name_B": "ATP19", "Name_A": "MCO10", "target_original": "854077", "Gene_B": "854077", "interaction": "Direct", "SUID": 2783001, "TYPE": "Direct", "Gene_A": "1466480", "support": "{\"StringDB\": \"physical\", \"StringEvidence\": \"database:0.791\", \"String_Score_Physical\": 800, \"String_Score_Combined\": 800}", "selected": false}, "selected": false}, {"data": {"id": "2782998", "source": "2782800", "target": "2782758", "SCORE": "540", "shared_interaction": "Direct", "source_original": "1466480", "Name_B": "ATP7", "Name_A": "MCO10", "target_original": "853853", "Gene_B": "853853", "interaction": "Direct", "SUID": 2782998, "TYPE": "Direct", "Gene_A": "1466480", "support": "{\"StringDB\": \"physical\", \"StringEvidence\": \"database:0.520\", \"String_Score_Physical\": 540, \"String_Score_Combined\": 540}", "selected": false}, "selected": false}, {"data": {"id": "2782995", "source": "2782800", "target": "2782743", "SCORE": "540", "shared_interaction": "Direct", "source_original": "1466480", "Name_B": "ATP3", "Name_A": "MCO10", "target_original": "852327", "Gene_B": "852327", "interaction": "Direct", "SUID": 2782995, "TYPE": "Direct", "Gene_A": "1466480", "support": "{\"StringDB\": \"physical\", \"StringEvidence\": \"database:0.520\", \"String_Score_Physical\": 540, \"String_Score_Combined\": 540}", "selected": false}, "selected": false}, {"data": {"id": "2782992", "source": "2782800", "target": "2782737", "SCORE": "540", "shared_interaction": "Direct", "source_original": "1466480", "Name_B": "ATP1", "Name_A": "MCO10", "target_original": "852177", "Gene_B": "852177", "interaction": "Direct", "SUID": 2782992, "TYPE": "Direct", "Gene_A": "1466480", "support": "{\"StringDB\": \"physical\", \"StringEvidence\": \"database:0.520\", \"String_Score_Physical\": 540, \"String_Score_Combined\": 540}", "selected": false}, "selected": false}, {"data": {"id": "2782989", "source": "2782800", "target": "2782731", "SCORE": "540", "shared_interaction": "Direct", "source_original": "1466480", "Name_B": "ATP17", "Name_A": "MCO10", "target_original": "851983", "Gene_B": "851983", "interaction": "Direct", "SUID": 2782989, "TYPE": "Direct", "Gene_A": "1466480", "support": "{\"StringDB\": \"physical\", \"StringEvidence\": \"database:0.520\", \"String_Score_Physical\": 540, \"String_Score_Combined\": 540}", "selected": false}, "selected": false}, {"data": {"id": "2782986", "source": "2782800", "target": "2782728", "SCORE": "540", "shared_interaction": "Direct", "source_original": "1466480", "Name_B": "TIM11", "Name_A": "MCO10", "target_original": "851922", "Gene_B": "851922", "interaction": "Direct", "SUID": 2782986, "TYPE": "Direct", "Gene_A": "1466480", "support": "{\"StringDB\": \"physical\", \"StringEvidence\": \"database:0.520\", \"String_Score_Physical\": 540, \"String_Score_Combined\": 540}", "selected": false}, "selected": false}, {"data": {"id": "2782956", "source": "2782800", "target": "2782719", "SCORE": "540", "shared_interaction": "Direct", "source_original": "1466480", "Name_B": "ATP14", "Name_A": "MCO10", "target_original": "851002", "Gene_B": "851002", "interaction": "Direct", "SUID": 2782956, "TYPE": "Direct", "Gene_A": "1466480", "support": "{\"StringDB\": \"physical\", \"StringEvidence\": \"database:0.520\", \"String_Score_Physical\": 540, \"String_Score_Combined\": 540}", "selected": false}, "selected": false}, {"data": {"id": "2784171", "source": "2782794", "target": "2782797", "SCORE": "987", "shared_interaction": "Direct", "source_original": "856672", "Name_B": "RIP1", "Name_A": "CYC7", "target_original": "856689", "Gene_B": "856689", "interaction": "Direct", "SUID": 2784171, "TYPE": "Direct", "Gene_A": "856672", "support": "{\"StringDB\": \"physical\", \"StringEvidence\": \"database:0.896,coexpression:0.890,experiments:0.876,textmining:0.746,neighborhood:0.138\", \"String_Score_Physical\": 987, \"String_Score_Combined\": 999}", "selected": false}, "selected": false}, {"data": {"id": "2784168", "source": "2782791", "target": "2782797", "SCORE": "985", "shared_interaction": "Direct", "source_original": "856390", "Name_B": "RIP1", "Name_A": "QCR10", "target_original": "856689", "Gene_B": "856689", "interaction": "Direct", "SUID": 2784168, "TYPE": "Direct", "Gene_A": "856390", "support": "{\"StringDB\": \"physical\", \"StringEvidence\": \"database:0.896,coexpression:0.871,experiments:0.851,textmining:0.429\", \"BioGrid\": \"physical\", \"BioGrid_PubMed\": \"29688796\", \"BioGrid_Type\": \"Affinity Capture-Western\", \"String_Score_Physical\": 985, \"String_Score_Combined\": 998}", "selected": false}, "selected": false}, {"data": {"id": "2784165", "source": "2782788", "target": "2782797", "SCORE": "999", "shared_interaction": "Direct", "source_original": "856321", "Name_B": "RIP1", "Name_A": "QCR2", "target_original": "856689", "Gene_B": "856689", "interaction": "Direct", "SUID": 2784165, "TYPE": "Direct", "Gene_A": "856321", "support": "{\"StringDB\": \"physical\", \"StringEvidence\": \"experiments:0.998,coexpression:0.991,textmining:0.922,database:0.896\", \"BioGrid\": \"physical\", \"BioGrid_PubMed\": \"25683140,24421313,21807901\", \"BioGrid_Type\": \"Affinity Capture-MS,Co-purification,Affinity Capture-Western\", \"String_Score_Physical\": 999, \"String_Score_Combined\": 999}", "selected": false}, "selected": false}, {"data": {"id": "2784162", "source": "2782788", "target": "2782794", "SCORE": "827", "shared_interaction": "Direct", "source_original": "856321", "Name_B": "CYC7", "Name_A": "QCR2", "target_original": "856672", "Gene_B": "856672", "interaction": "Direct", "SUID": 2784162, "TYPE": "Direct", "Gene_A": "856321", "support": "{\"StringDB\": \"physical\", \"StringEvidence\": \"experiments:0.820,textmining:0.473,coexpression:0.223\", \"String_Score_Physical\": 827, \"String_Score_Combined\": 929}", "selected": false}, "selected": false}, {"data": {"id": "2784159", "source": "2782788", "target": "2782791", "SCORE": "988", "shared_interaction": "Direct", "source_original": "856321", "Name_B": "QCR10", "Name_A": "QCR2", "target_original": "856390", "Gene_B": "856390", "interaction": "Direct", "SUID": 2784159, "TYPE": "Direct", "Gene_A": "856321", "support": "{\"StringDB\": \"physical\", \"StringEvidence\": \"textmining:0.897,database:0.896,experiments:0.883,coexpression:0.586\", \"BioGrid\": \"physical\", \"BioGrid_PubMed\": \"29688796,25683140\", \"BioGrid_Type\": \"Affinity Capture-Western\", \"String_Score_Physical\": 988, \"String_Score_Combined\": 999}", "selected": false}, "selected": false}, {"data": {"id": "2784156", "source": "2782785", "target": "2782797", "SCORE": "214", "shared_interaction": "Direct", "source_original": "856131", "Name_B": "RIP1", "Name_A": "ATP20", "target_original": "856689", "Gene_B": "856689", "interaction": "Direct", "SUID": 2784156, "TYPE": "Direct", "Gene_A": "856131", "support": "{\"StringDB\": \"physical\", \"StringEvidence\": \"coexpression:0.923,experiments:0.180,textmining:0.081\", \"String_Score_Physical\": 214, \"String_Score_Combined\": 944}", "selected": false}, "selected": false}, {"data": {"id": "2784153", "source": "2782785", "target": "2782794", "SCORE": "335", "shared_interaction": "Direct", "source_original": "856131", "Name_B": "CYC7", "Name_A": "ATP20", "target_original": "856672", "Gene_B": "856672", "interaction": "Direct", "SUID": 2784153, "TYPE": "Direct", "Gene_A": "856131", "support": "{\"StringDB\": \"physical\", \"StringEvidence\": \"experiments:0.307,coexpression:0.170,textmining:0.014\", \"String_Score_Physical\": 335, \"String_Score_Combined\": 455}", "selected": false}, "selected": false}, {"data": {"id": "2784150", "source": "2782785", "target": "2782788", "SCORE": "962", "shared_interaction": "Direct", "source_original": "856131", "Name_B": "QCR2", "Name_A": "ATP20", "target_original": "856321", "Gene_B": "856321", "interaction": "Direct", "SUID": 2784150, "TYPE": "Direct", "Gene_A": "856131", "support": "{\"StringDB\": \"physical\", \"StringEvidence\": \"experiments:0.960,coexpression:0.745,textmining:0.708\", \"String_Score_Physical\": 962, \"String_Score_Combined\": 997}", "selected": false}, "selected": false}, {"data": {"id": "2784147", "source": "2782782", "target": "2782797", "SCORE": "964", "shared_interaction": "Direct", "source_original": "856027", "Name_B": "RIP1", "Name_A": "ATP4", "target_original": "856689", "Gene_B": "856689", "interaction": "Direct", "SUID": 2784147, "TYPE": "Direct", "Gene_A": "856027", "support": "{\"StringDB\": \"physical\", \"StringEvidence\": \"experiments:0.962,coexpression:0.941,textmining:0.081\", \"String_Score_Physical\": 964, \"String_Score_Combined\": 998}", "selected": false}, "selected": false}, {"data": {"id": "2784144", "source": "2782782", "target": "2782794", "SCORE": "192", "shared_interaction": "Direct", "source_original": "856027", "Name_B": "CYC7", "Name_A": "ATP4", "target_original": "856672", "Gene_B": "856672", "interaction": "Direct", "SUID": 2784144, "TYPE": "Direct", "Gene_A": "856027", "support": "{\"StringDB\": \"physical\", \"StringEvidence\": \"textmining:0.353,coexpression:0.170,experiments:0.157\", \"String_Score_Physical\": 192, \"String_Score_Combined\": 565}", "selected": false}, "selected": false}, {"data": {"id": "2784141", "source": "2782782", "target": "2782791", "SCORE": "315", "shared_interaction": "Direct", "source_original": "856027", "Name_B": "QCR10", "Name_A": "ATP4", "target_original": "856390", "Gene_B": "856390", "interaction": "Direct", "SUID": 2784141, "TYPE": "Direct", "Gene_A": "856027", "support": "{\"StringDB\": \"physical\", \"StringEvidence\": \"experiments:0.286,coexpression:0.215\", \"BioGrid\": \"physical\", \"BioGrid_PubMed\": \"29688796\", \"BioGrid_Type\": \"Affinity Capture-Western\", \"String_Score_Physical\": 315, \"String_Score_Combined\": 462}", "selected": false}, "selected": false}, {"data": {"id": "2784138", "source": "2782782", "target": "2782788", "SCORE": "963", "shared_interaction": "Direct", "source_original": "856027", "Name_B": "QCR2", "Name_A": "ATP4", "target_original": "856321", "Gene_B": "856321", "interaction": "Direct", "SUID": 2784138, "TYPE": "Direct", "Gene_A": "856027", "support": "{\"StringDB\": \"physical\", \"StringEvidence\": \"experiments:0.961,coexpression:0.952,textmining:0.689\", \"String_Score_Physical\": 963, \"String_Score_Combined\": 999}", "selected": false}, "selected": false}, {"data": {"id": "2784135", "source": "2782782", "target": "2782785", "SCORE": "999", "shared_interaction": "Direct", "source_original": "856027", "Name_B": "ATP20", "Name_A": "ATP4", "target_original": "856131", "Gene_B": "856131", "interaction": "Direct", "SUID": 2784135, "TYPE": "Direct", "Gene_A": "856027", "support": "{\"StringDB\": \"physical\", \"StringEvidence\": \"experiments:0.998,coexpression:0.958,textmining:0.911,database:0.791\", \"BioGrid\": \"physical\", \"BioGrid_PubMed\": \"9867807\", \"BioGrid_Type\": \"Co-purification\", \"String_Score_Physical\": 999, \"String_Score_Combined\": 999}", "selected": false}, "selected": false}, {"data": {"id": "2784132", "source": "2782779", "target": "2782797", "SCORE": "998", "shared_interaction": "Direct", "source_original": "855675", "Name_B": "RIP1", "Name_A": "COX5A", "target_original": "856689", "Gene_B": "856689", "interaction": "Direct", "SUID": 2784132, "TYPE": "Direct", "Gene_A": "855675", "support": "{\"StringDB\": \"physical\", \"StringEvidence\": \"coexpression:0.988,experiments:0.987,database:0.896,textmining:0.395\", \"BioGrid\": \"physical\", \"BioGrid_PubMed\": \"30598554\", \"BioGrid_Type\": \"Co-crystal Structure\", \"String_Score_Physical\": 998, \"String_Score_Combined\": 999}", "selected": false}, "selected": false}, {"data": {"id": "2784129", "source": "2782779", "target": "2782794", "SCORE": "569", "shared_interaction": "Direct", "source_original": "855675", "Name_B": "CYC7", "Name_A": "COX5A", "target_original": "856672", "Gene_B": "856672", "interaction": "Direct", "SUID": 2784129, "TYPE": "Direct", "Gene_A": "855675", "support": "{\"StringDB\": \"physical\", \"StringEvidence\": \"textmining:0.715,experiments:0.551,coexpression:0.127\", \"String_Score_Physical\": 569, \"String_Score_Combined\": 892}", "selected": false}, "selected": false}, {"data": {"id": "2784126", "source": "2782779", "target": "2782791", "SCORE": "979", "shared_interaction": "Direct", "source_original": "855675", "Name_B": "QCR10", "Name_A": "COX5A", "target_original": "856390", "Gene_B": "856390", "interaction": "Direct", "SUID": 2784126, "TYPE": "Direct", "Gene_A": "855675", "support": "{\"StringDB\": \"physical\", \"StringEvidence\": \"database:0.896,coexpression:0.827,experiments:0.791,textmining:0.426\", \"String_Score_Physical\": 979, \"String_Score_Combined\": 997}", "selected": false}, "selected": false}, {"data": {"id": "2784123", "source": "2782779", "target": "2782788", "SCORE": "999", "shared_interaction": "Direct", "source_original": "855675", "Name_B": "QCR2", "Name_A": "COX5A", "target_original": "856321", "Gene_B": "856321", "interaction": "Direct", "SUID": 2784123, "TYPE": "Direct", "Gene_A": "855675", "support": "{\"StringDB\": \"physical\", \"StringEvidence\": \"experiments:0.991,database:0.896,coexpression:0.862,textmining:0.694\", \"String_Score_Physical\": 999, \"String_Score_Combined\": 999}", "selected": false}, "selected": false}, {"data": {"id": "2784120", "source": "2782779", "target": "2782785", "SCORE": "696", "shared_interaction": "Direct", "source_original": "855675", "Name_B": "ATP20", "Name_A": "COX5A", "target_original": "856131", "Gene_B": "856131", "interaction": "Direct", "SUID": 2784120, "TYPE": "Direct", "Gene_A": "855675", "support": "{\"StringDB\": \"physical\", \"StringEvidence\": \"coexpression:0.870,experiments:0.683,textmining:0.026\", \"String_Score_Physical\": 696, \"String_Score_Combined\": 961}", "selected": false}, "selected": false}, {"data": {"id": "2784117", "source": "2782779", "target": "2782782", "SCORE": "719", "shared_interaction": "Direct", "source_original": "855675", "Name_B": "ATP4", "Name_A": "COX5A", "target_original": "856027", "Gene_B": "856027", "interaction": "Direct", "SUID": 2784117, "TYPE": "Direct", "Gene_A": "855675", "support": "{\"StringDB\": \"physical\", \"StringEvidence\": \"coexpression:0.731,experiments:0.707,textmining:0.018\", \"String_Score_Physical\": 719, \"String_Score_Combined\": 925}", "selected": false}, "selected": false}, {"data": {"id": "2784114", "source": "2782776", "target": "2782797", "SCORE": "979", "shared_interaction": "Direct", "source_original": "855298", "Name_B": "RIP1", "Name_A": "COX7", "target_original": "856689", "Gene_B": "856689", "interaction": "Direct", "SUID": 2784114, "TYPE": "Direct", "Gene_A": "855298", "support": "{\"StringDB\": \"physical\", \"StringEvidence\": \"database:0.896,experiments:0.791,coexpression:0.331,textmining:0.261\", \"String_Score_Physical\": 979, \"String_Score_Combined\": 989}", "selected": false}, "selected": false}, {"data": {"id": "2784111", "source": "2782776", "target": "2782791", "SCORE": "979", "shared_interaction": "Direct", "source_original": "855298", "Name_B": "QCR10", "Name_A": "COX7", "target_original": "856390", "Gene_B": "856390", "interaction": "Direct", "SUID": 2784111, "TYPE": "Direct", "Gene_A": "855298", "support": "{\"StringDB\": \"physical\", \"StringEvidence\": \"coexpression:0.916,database:0.896,experiments:0.791,textmining:0.062\", \"String_Score_Physical\": 979, \"String_Score_Combined\": 998}", "selected": false}, "selected": false}, {"data": {"id": "2784108", "source": "2782776", "target": "2782788", "SCORE": "979", "shared_interaction": "Direct", "source_original": "855298", "Name_B": "QCR2", "Name_A": "COX7", "target_original": "856321", "Gene_B": "856321", "interaction": "Direct", "SUID": 2784108, "TYPE": "Direct", "Gene_A": "855298", "support": "{\"StringDB\": \"physical\", \"StringEvidence\": \"database:0.896,experiments:0.791,textmining:0.734,coexpression:0.260\", \"String_Score_Physical\": 979, \"String_Score_Combined\": 995}", "selected": false}, "selected": false}, {"data": {"id": "2784105", "source": "2782776", "target": "2782779", "SCORE": "958", "shared_interaction": "Direct", "source_original": "855298", "Name_B": "COX5A", "Name_A": "COX7", "target_original": "855675", "Gene_B": "855675", "interaction": "Direct", "SUID": 2784105, "TYPE": "Direct", "Gene_A": "855298", "support": "{\"StringDB\": \"physical\", \"StringEvidence\": \"textmining:0.904,experiments:0.791,database:0.791,coexpression:0.560\", \"String_Score_Physical\": 958, \"String_Score_Combined\": 998}", "selected": false}, "selected": false}, {"data": {"id": "2784099", "source": "2782773", "target": "2782797", "SCORE": "900", "shared_interaction": "Direct", "source_original": "854919", "Name_B": "RIP1", "Name_A": "NDI1", "target_original": "856689", "Gene_B": "856689", "interaction": "Direct", "SUID": 2784099, "TYPE": "Direct", "Gene_A": "854919", "support": "{\"StringDB\": \"physical\", \"StringEvidence\": \"coexpression:0.926,database:0.896,textmining:0.643\", \"String_Score_Physical\": 900, \"String_Score_Combined\": 997}", "selected": false}, "selected": false}, {"data": {"id": "2784096", "source": "2782773", "target": "2782791", "SCORE": "900", "shared_interaction": "Direct", "source_original": "854919", "Name_B": "QCR10", "Name_A": "NDI1", "target_original": "856390", "Gene_B": "856390", "interaction": "Direct", "SUID": 2784096, "TYPE": "Direct", "Gene_A": "854919", "support": "{\"StringDB\": \"physical\", \"StringEvidence\": \"database:0.896,coexpression:0.321\", \"String_Score_Physical\": 900, \"String_Score_Combined\": 932}", "selected": false}, "selected": false}, {"data": {"id": "2784093", "source": "2782773", "target": "2782788", "SCORE": "900", "shared_interaction": "Direct", "source_original": "854919", "Name_B": "QCR2", "Name_A": "NDI1", "target_original": "856321", "Gene_B": "856321", "interaction": "Direct", "SUID": 2784093, "TYPE": "Direct", "Gene_A": "854919", "support": "{\"StringDB\": \"physical\", \"StringEvidence\": \"database:0.896,coexpression:0.818,textmining:0.460,neighborhood:0.042\", \"String_Score_Physical\": 900, \"String_Score_Combined\": 990}", "selected": false}, "selected": false}, {"data": {"id": "2784090", "source": "2782773", "target": "2782779", "SCORE": "900", "shared_interaction": "Direct", "source_original": "854919", "Name_B": "COX5A", "Name_A": "NDI1", "target_original": "855675", "Gene_B": "855675", "interaction": "Direct", "SUID": 2784090, "TYPE": "Direct", "Gene_A": "854919", "support": "{\"StringDB\": \"physical\", \"StringEvidence\": \"database:0.896,coexpression:0.842,textmining:0.294\", \"String_Score_Physical\": 900, \"String_Score_Combined\": 988}", "selected": false}, "selected": false}, {"data": {"id": "2784087", "source": "2782773", "target": "2782776", "SCORE": "900", "shared_interaction": "Direct", "source_original": "854919", "Name_B": "COX7", "Name_A": "NDI1", "target_original": "855298", "Gene_B": "855298", "interaction": "Direct", "SUID": 2784087, "TYPE": "Direct", "Gene_A": "854919", "support": "{\"StringDB\": \"physical\", \"StringEvidence\": \"database:0.896,coexpression:0.100,textmining:0.093\", \"String_Score_Physical\": 900, \"String_Score_Combined\": 918}", "selected": false}, "selected": false}, {"data": {"id": "2784084", "source": "2782770", "target": "2782785", "SCORE": "795", "shared_interaction": "Direct", "source_original": "854893", "Name_B": "ATP20", "Name_A": "ATP18", "target_original": "856131", "Gene_B": "856131", "interaction": "Direct", "SUID": 2784084, "TYPE": "Direct", "Gene_A": "854893", "support": "{\"StringDB\": \"physical\", \"StringEvidence\": \"textmining:0.735,database:0.708,coexpression:0.311,experiments:0.269\", \"BioGrid\": \"physical\", \"BioGrid_PubMed\": \"10747812\", \"BioGrid_Type\": \"Reconstituted Complex\", \"String_Score_Physical\": 795, \"String_Score_Combined\": 962}", "selected": false}, "selected": false}, {"data": {"id": "2784081", "source": "2782770", "target": "2782782", "SCORE": "999", "shared_interaction": "Direct", "source_original": "854893", "Name_B": "ATP4", "Name_A": "ATP18", "target_original": "856027", "Gene_B": "856027", "interaction": "Direct", "SUID": 2784081, "TYPE": "Direct", "Gene_A": "854893", "support": "{\"StringDB\": \"physical\", \"StringEvidence\": \"experiments:0.999,database:0.896,textmining:0.580,coexpression:0.126\", \"BioGrid\": \"genetic,physical\", \"BioGrid_PubMed\": \"9867807,21623372\", \"BioGrid_Type\": \"Co-purification,Positive Genetic\", \"BioGrid_Score\": \"0.1794\", \"String_Score_Physical\": 999, \"String_Score_Combined\": 999}", "selected": false}, "selected": false}, {"data": {"id": "2784075", "source": "2782767", "target": "2782797", "SCORE": "985", "shared_interaction": "Direct", "source_original": "854695", "Name_B": "RIP1", "Name_A": "COX5B", "target_original": "856689", "Gene_B": "856689", "interaction": "Direct", "SUID": 2784075, "TYPE": "Direct", "Gene_A": "854695", "support": "{\"StringDB\": \"physical\", \"StringEvidence\": \"database:0.896,experiments:0.852,coexpression:0.637,textmining:0.234\", \"String_Score_Physical\": 985, \"String_Score_Combined\": 995}", "selected": false}, "selected": false}, {"data": {"id": "2784072", "source": "2782767", "target": "2782794", "SCORE": "569", "shared_interaction": "Direct", "source_original": "854695", "Name_B": "CYC7", "Name_A": "COX5B", "target_original": "856672", "Gene_B": "856672", "interaction": "Direct", "SUID": 2784072, "TYPE": "Direct", "Gene_A": "854695", "support": "{\"StringDB\": \"physical\", \"StringEvidence\": \"textmining:0.867,experiments:0.551,coexpression:0.308\", \"String_Score_Physical\": 569, \"String_Score_Combined\": 960}", "selected": false}, "selected": false}, {"data": {"id": "2784069", "source": "2782767", "target": "2782791", "SCORE": "979", "shared_interaction": "Direct", "source_original": "854695", "Name_B": "QCR10", "Name_A": "COX5B", "target_original": "856390", "Gene_B": "856390", "interaction": "Direct", "SUID": 2784069, "TYPE": "Direct", "Gene_A": "854695", "support": "{\"StringDB\": \"physical\", \"StringEvidence\": \"database:0.896,experiments:0.791,textmining:0.322\", \"String_Score_Physical\": 979, \"String_Score_Combined\": 985}", "selected": false}, "selected": false}, {"data": {"id": "2784066", "source": "2782767", "target": "2782788", "SCORE": "999", "shared_interaction": "Direct", "source_original": "854695", "Name_B": "QCR2", "Name_A": "COX5B", "target_original": "856321", "Gene_B": "856321", "interaction": "Direct", "SUID": 2784066, "TYPE": "Direct", "Gene_A": "854695", "support": "{\"StringDB\": \"physical\", \"StringEvidence\": \"experiments:0.992,database:0.896,coexpression:0.170,textmining:0.151\", \"BioGrid\": \"genetic\", \"BioGrid_PubMed\": \"21623372\", \"BioGrid_Type\": \"Negative Genetic\", \"BioGrid_Score\": \"-0.1192\", \"String_Score_Physical\": 999, \"String_Score_Combined\": 999}", "selected": false}, "selected": false}, {"data": {"id": "2784063", "source": "2782767", "target": "2782785", "SCORE": "696", "shared_interaction": "Direct", "source_original": "854695", "Name_B": "ATP20", "Name_A": "COX5B", "target_original": "856131", "Gene_B": "856131", "interaction": "Direct", "SUID": 2784063, "TYPE": "Direct", "Gene_A": "854695", "support": "{\"StringDB\": \"physical\", \"StringEvidence\": \"experiments:0.683,coexpression:0.428,textmining:0.026\", \"String_Score_Physical\": 696, \"String_Score_Combined\": 830}", "selected": false}, "selected": false}, {"data": {"id": "2784060", "source": "2782767", "target": "2782782", "SCORE": "719", "shared_interaction": "Direct", "source_original": "854695", "Name_B": "ATP4", "Name_A": "COX5B", "target_original": "856027", "Gene_B": "856027", "interaction": "Direct", "SUID": 2784060, "TYPE": "Direct", "Gene_A": "854695", "support": "{\"StringDB\": \"physical\", \"StringEvidence\": \"experiments:0.707,textmining:0.441,coexpression:0.344\", \"String_Score_Physical\": 719, \"String_Score_Combined\": 896}", "selected": false}, "selected": false}, {"data": {"id": "2784057", "source": "2782767", "target": "2782779", "SCORE": "860", "shared_interaction": "Direct", "source_original": "854695", "Name_B": "COX5A", "Name_A": "COX5B", "target_original": "855675", "Gene_B": "855675", "interaction": "Direct", "SUID": 2784057, "TYPE": "Direct", "Gene_A": "854695", "support": "{\"StringDB\": \"physical\", \"StringEvidence\": \"textmining:0.900,database:0.791,experiments:0.300\", \"BioGrid\": \"genetic\", \"BioGrid_PubMed\": \"27708008,21623372\", \"BioGrid_Type\": \"Negative Genetic\", \"BioGrid_Score\": \"-0.1505,-0.1618\", \"String_Score_Physical\": 860, \"String_Score_Combined\": 864}", "selected": false}, "selected": false}, {"data": {"id": "2784054", "source": "2782767", "target": "2782776", "SCORE": "941", "shared_interaction": "Direct", "source_original": "854695", "Name_B": "COX7", "Name_A": "COX5B", "target_original": "855298", "Gene_B": "855298", "interaction": "Direct", "SUID": 2784054, "TYPE": "Direct", "Gene_A": "854695", "support": "{\"StringDB\": \"physical\", \"StringEvidence\": \"textmining:0.900,experiments:0.791,database:0.708,coexpression:0.006\", \"String_Score_Physical\": 941, \"String_Score_Combined\": 994}", "selected": false}, "selected": false}, {"data": {"id": "2784051", "source": "2782767", "target": "2782773", "SCORE": "900", "shared_interaction": "Direct", "source_original": "854695", "Name_B": "NDI1", "Name_A": "COX5B", "target_original": "854919", "Gene_B": "854919", "interaction": "Direct", "SUID": 2784051, "TYPE": "Direct", "Gene_A": "854695", "support": "{\"StringDB\": \"physical\", \"StringEvidence\": \"database:0.896,textmining:0.049,coexpression:0.016\", \"String_Score_Physical\": 900, \"String_Score_Combined\": 906}", "selected": false}, "selected": false}, {"data": {"id": "2784048", "source": "2782764", "target": "2782797", "SCORE": "999", "shared_interaction": "Direct", "source_original": "854231", "Name_B": "RIP1", "Name_A": "CYT1", "target_original": "856689", "Gene_B": "856689", "interaction": "Direct", "SUID": 2784048, "TYPE": "Direct", "Gene_A": "854231", "support": "{\"StringDB\": \"physical\", \"StringEvidence\": \"experiments:0.998,coexpression:0.992,textmining:0.988,database:0.896,cooccurrence:0.264,neighborhood:0.156,fusion:0.097\", \"BioGrid\": \"genetic\", \"BioGrid_PubMed\": \"20093466\", \"BioGrid_Type\": \"Positive Genetic\", \"BioGrid_Score\": \"0.18\", \"String_Score_Physical\": 999, \"String_Score_Combined\": 999}", "selected": false}, "selected": false}, {"data": {"id": "2784045", "source": "2782764", "target": "2782794", "SCORE": "982", "shared_interaction": "Direct", "source_original": "854231", "Name_B": "CYC7", "Name_A": "CYT1", "target_original": "856672", "Gene_B": "856672", "interaction": "Direct", "SUID": 2784045, "TYPE": "Direct", "Gene_A": "854231", "support": "{\"StringDB\": \"physical\", \"StringEvidence\": \"database:0.896,textmining:0.887,experiments:0.826,coexpression:0.822,cooccurrence:0.212\", \"String_Score_Physical\": 982, \"String_Score_Combined\": 999}", "selected": false}, "selected": false}, {"data": {"id": "2784042", "source": "2782764", "target": "2782791", "SCORE": "988", "shared_interaction": "Direct", "source_original": "854231", "Name_B": "QCR10", "Name_A": "CYT1", "target_original": "856390", "Gene_B": "856390", "interaction": "Direct", "SUID": 2784042, "TYPE": "Direct", "Gene_A": "854231", "support": "{\"StringDB\": \"physical\", \"StringEvidence\": \"textmining:0.897,database:0.896,experiments:0.884,coexpression:0.713\", \"String_Score_Physical\": 988, \"String_Score_Combined\": 999}", "selected": false}, "selected": false}, {"data": {"id": "2784039", "source": "2782764", "target": "2782788", "SCORE": "999", "shared_interaction": "Direct", "source_original": "854231", "Name_B": "QCR2", "Name_A": "CYT1", "target_original": "856321", "Gene_B": "856321", "interaction": "Direct", "SUID": 2784039, "TYPE": "Direct", "Gene_A": "854231", "support": "{\"StringDB\": \"physical\", \"StringEvidence\": \"experiments:0.999,coexpression:0.990,textmining:0.975,database:0.896\", \"BioGrid\": \"physical\", \"BioGrid_PubMed\": \"32119834,25683140,21807901\", \"BioGrid_Type\": \"Affinity Capture-Western\", \"String_Score_Physical\": 999, \"String_Score_Combined\": 999}", "selected": false}, "selected": false}, {"data": {"id": "2784036", "source": "2782764", "target": "2782785", "SCORE": "964", "shared_interaction": "Direct", "source_original": "854231", "Name_B": "ATP20", "Name_A": "CYT1", "target_original": "856131", "Gene_B": "856131", "interaction": "Direct", "SUID": 2784036, "TYPE": "Direct", "Gene_A": "854231", "support": "{\"StringDB\": \"physical\", \"StringEvidence\": \"experiments:0.962,coexpression:0.713,textmining:0.611\", \"String_Score_Physical\": 964, \"String_Score_Combined\": 995}", "selected": false}, "selected": false}, {"data": {"id": "2784033", "source": "2782764", "target": "2782782", "SCORE": "964", "shared_interaction": "Direct", "source_original": "854231", "Name_B": "ATP4", "Name_A": "CYT1", "target_original": "856027", "Gene_B": "856027", "interaction": "Direct", "SUID": 2784033, "TYPE": "Direct", "Gene_A": "854231", "support": "{\"StringDB\": \"physical\", \"StringEvidence\": \"experiments:0.962,coexpression:0.942,textmining:0.845\", \"String_Score_Physical\": 964, \"String_Score_Combined\": 999}", "selected": false}, "selected": false}, {"data": {"id": "2784030", "source": "2782764", "target": "2782779", "SCORE": "999", "shared_interaction": "Direct", "source_original": "854231", "Name_B": "COX5A", "Name_A": "CYT1", "target_original": "855675", "Gene_B": "855675", "interaction": "Direct", "SUID": 2784030, "TYPE": "Direct", "Gene_A": "854231", "support": "{\"StringDB\": \"physical\", \"StringEvidence\": \"experiments:0.994,coexpression:0.959,database:0.896,textmining:0.674\", \"BioGrid\": \"physical\", \"BioGrid_PubMed\": \"30598554\", \"BioGrid_Type\": \"Co-crystal Structure\", \"String_Score_Physical\": 999, \"String_Score_Combined\": 999}", "selected": false}, "selected": false}, {"data": {"id": "2784027", "source": "2782764", "target": "2782776", "SCORE": "979", "shared_interaction": "Direct", "source_original": "854231", "Name_B": "COX7", "Name_A": "CYT1", "target_original": "855298", "Gene_B": "855298", "interaction": "Direct", "SUID": 2784027, "TYPE": "Direct", "Gene_A": "854231", "support": "{\"StringDB\": \"physical\", \"StringEvidence\": \"database:0.896,experiments:0.791,textmining:0.700,coexpression:0.374\", \"String_Score_Physical\": 979, \"String_Score_Combined\": 996}", "selected": false}, "selected": false}, {"data": {"id": "2784024", "source": "2782764", "target": "2782773", "SCORE": "900", "shared_interaction": "Direct", "source_original": "854231", "Name_B": "NDI1", "Name_A": "CYT1", "target_original": "854919", "Gene_B": "854919", "interaction": "Direct", "SUID": 2784024, "TYPE": "Direct", "Gene_A": "854231", "support": "{\"StringDB\": \"physical\", \"StringEvidence\": \"coexpression:0.920,database:0.896,textmining:0.635\", \"String_Score_Physical\": 900, \"String_Score_Combined\": 997}", "selected": false}, "selected": false}, {"data": {"id": "2784021", "source": "2782764", "target": "2782767", "SCORE": "993", "shared_interaction": "Direct", "source_original": "854231", "Name_B": "COX5B", "Name_A": "CYT1", "target_original": "854695", "Gene_B": "854695", "interaction": "Direct", "SUID": 2784021, "TYPE": "Direct", "Gene_A": "854231", "support": "{\"StringDB\": \"physical\", \"StringEvidence\": \"experiments:0.933,database:0.896,coexpression:0.491,textmining:0.422\", \"BioGrid\": \"genetic\", \"BioGrid_PubMed\": \"21623372\", \"BioGrid_Type\": \"Negative Genetic\", \"BioGrid_Score\": \"-0.0817\", \"String_Score_Physical\": 993, \"String_Score_Combined\": 998}", "selected": false}, "selected": false}, {"data": {"id": "2784018", "source": "2782761", "target": "2782785", "SCORE": "720", "shared_interaction": "Direct", "source_original": "854077", "Name_B": "ATP20", "Name_A": "ATP19", "target_original": "856131", "Gene_B": "856131", "interaction": "Direct", "SUID": 2784018, "TYPE": "Direct", "Gene_A": "854077", "support": "{\"StringDB\": \"physical\", \"StringEvidence\": \"coexpression:0.844,textmining:0.771,database:0.708\", \"String_Score_Physical\": 720, \"String_Score_Combined\": 989}", "selected": false}, "selected": false}, {"data": {"id": "2784015", "source": "2782761", "target": "2782782", "SCORE": "941", "shared_interaction": "Direct", "source_original": "854077", "Name_B": "ATP4", "Name_A": "ATP19", "target_original": "856027", "Gene_B": "856027", "interaction": "Direct", "SUID": 2784015, "TYPE": "Direct", "Gene_A": "854077", "support": "{\"StringDB\": \"physical\", \"StringEvidence\": \"experiments:0.791,database:0.708,textmining:0.250,coexpression:0.137\", \"String_Score_Physical\": 941, \"String_Score_Combined\": 962}", "selected": false}, "selected": false}, {"data": {"id": "2784012", "source": "2782761", "target": "2782770", "SCORE": "941", "shared_interaction": "Direct", "source_original": "854077", "Name_B": "ATP18", "Name_A": "ATP19", "target_original": "854893", "Gene_B": "854893", "interaction": "Direct", "SUID": 2784012, "TYPE": "Direct", "Gene_A": "854077", "support": "{\"StringDB\": \"physical\", \"StringEvidence\": \"coexpression:0.852,textmining:0.846,experiments:0.791,database:0.708\", \"String_Score_Physical\": 941, \"String_Score_Combined\": 998}", "selected": false}, "selected": false}, {"data": {"id": "2784000", "source": "2782758", "target": "2782797", "SCORE": "735", "shared_interaction": "Direct", "source_original": "853853", "Name_B": "RIP1", "Name_A": "ATP7", "target_original": "856689", "Gene_B": "856689", "interaction": "Direct", "SUID": 2784000, "TYPE": "Direct", "Gene_A": "853853", "support": "{\"StringDB\": \"physical\", \"StringEvidence\": \"coexpression:0.948,experiments:0.724,textmining:0.462\", \"String_Score_Physical\": 735, \"String_Score_Combined\": 992}", "selected": false}, "selected": false}, {"data": {"id": "2783997", "source": "2782758", "target": "2782794", "SCORE": "192", "shared_interaction": "Direct", "source_original": "853853", "Name_B": "CYC7", "Name_A": "ATP7", "target_original": "856672", "Gene_B": "856672", "interaction": "Direct", "SUID": 2783997, "TYPE": "Direct", "Gene_A": "853853", "support": "{\"StringDB\": \"physical\", \"StringEvidence\": \"textmining:0.526,coexpression:0.219,experiments:0.157\", \"String_Score_Physical\": 192, \"String_Score_Combined\": 701}", "selected": false}, "selected": false}, {"data": {"id": "2783994", "source": "2782758", "target": "2782788", "SCORE": "252", "shared_interaction": "Direct", "source_original": "853853", "Name_B": "QCR2", "Name_A": "ATP7", "target_original": "856321", "Gene_B": "856321", "interaction": "Direct", "SUID": 2783994, "TYPE": "Direct", "Gene_A": "853853", "support": "{\"StringDB\": \"physical\", \"StringEvidence\": \"coexpression:0.969,textmining:0.710,experiments:0.221\", \"String_Score_Physical\": 252, \"String_Score_Combined\": 993}", "selected": false}, "selected": false}, {"data": {"id": "2783991", "source": "2782758", "target": "2782785", "SCORE": "995", "shared_interaction": "Direct", "source_original": "853853", "Name_B": "ATP20", "Name_A": "ATP7", "target_original": "856131", "Gene_B": "856131", "interaction": "Direct", "SUID": 2783991, "TYPE": "Direct", "Gene_A": "853853", "support": "{\"StringDB\": \"physical\", \"StringEvidence\": \"experiments:0.976,coexpression:0.962,database:0.791,textmining:0.680\", \"String_Score_Physical\": 995, \"String_Score_Combined\": 999}", "selected": false}, "selected": false}, {"data": {"id": "2783988", "source": "2782758", "target": "2782782", "SCORE": "999", "shared_interaction": "Direct", "source_original": "853853", "Name_B": "ATP4", "Name_A": "ATP7", "target_original": "856027", "Gene_B": "856027", "interaction": "Direct", "SUID": 2783988, "TYPE": "Direct", "Gene_A": "853853", "support": "{\"StringDB\": \"physical\", \"StringEvidence\": \"experiments:0.999,textmining:0.980,coexpression:0.958,database:0.896\", \"BioGrid\": \"genetic,physical\", \"BioGrid_PubMed\": \"9867807,22940862,21623372\", \"BioGrid_Type\": \"Affinity Capture-MS,Co-purification,Positive Genetic\", \"BioGrid_Score\": \"0.1117\", \"String_Score_Physical\": 999, \"String_Score_Combined\": 999}", "selected": false}, "selected": false}, {"data": {"id": "2783985", "source": "2782758", "target": "2782779", "SCORE": "187", "shared_interaction": "Direct", "source_original": "853853", "Name_B": "COX5A", "Name_A": "ATP7", "target_original": "855675", "Gene_B": "855675", "interaction": "Direct", "SUID": 2783985, "TYPE": "Direct", "Gene_A": "853853", "support": "{\"StringDB\": \"physical\", \"StringEvidence\": \"coexpression:0.853,textmining:0.254,experiments:0.152\", \"String_Score_Physical\": 187, \"String_Score_Combined\": 910}", "selected": false}, "selected": false}, {"data": {"id": "2783982", "source": "2782758", "target": "2782770", "SCORE": "994", "shared_interaction": "Direct", "source_original": "853853", "Name_B": "ATP18", "Name_A": "ATP7", "target_original": "854893", "Gene_B": "854893", "interaction": "Direct", "SUID": 2783982, "TYPE": "Direct", "Gene_A": "853853", "support": "{\"StringDB\": \"physical\", \"StringEvidence\": \"experiments:0.945,database:0.896,textmining:0.506,coexpression:0.279\", \"BioGrid\": \"physical\", \"BioGrid_PubMed\": \"10747812\", \"BioGrid_Type\": \"Reconstituted Complex\", \"String_Score_Physical\": 994, \"String_Score_Combined\": 998}", "selected": false}, "selected": false}, {"data": {"id": "2783979", "source": "2782758", "target": "2782767", "SCORE": "187", "shared_interaction": "Direct", "source_original": "853853", "Name_B": "COX5B", "Name_A": "ATP7", "target_original": "854695", "Gene_B": "854695", "interaction": "Direct", "SUID": 2783979, "TYPE": "Direct", "Gene_A": "853853", "support": "{\"StringDB\": \"physical\", \"StringEvidence\": \"coexpression:0.628,textmining:0.549,experiments:0.152\", \"String_Score_Physical\": 187, \"String_Score_Combined\": 863}", "selected": false}, "selected": false}, {"data": {"id": "2783976", "source": "2782758", "target": "2782764", "SCORE": "272", "shared_interaction": "Direct", "source_original": "853853", "Name_B": "CYT1", "Name_A": "ATP7", "target_original": "854231", "Gene_B": "854231", "interaction": "Direct", "SUID": 2783976, "TYPE": "Direct", "Gene_A": "853853", "support": "{\"StringDB\": \"physical\", \"StringEvidence\": \"coexpression:0.948,textmining:0.351,experiments:0.242\", \"BioGrid\": \"genetic\", \"BioGrid_PubMed\": \"27708008\", \"BioGrid_Type\": \"Negative Genetic\", \"BioGrid_Score\": \"-0.1832\", \"String_Score_Physical\": 272, \"String_Score_Combined\": 975}", "selected": false}, "selected": false}, {"data": {"id": "2783973", "source": "2782758", "target": "2782761", "SCORE": "941", "shared_interaction": "Direct", "source_original": "853853", "Name_B": "ATP19", "Name_A": "ATP7", "target_original": "854077", "Gene_B": "854077", "interaction": "Direct", "SUID": 2783973, "TYPE": "Direct", "Gene_A": "853853", "support": "{\"StringDB\": \"physical\", \"StringEvidence\": \"experiments:0.791,database:0.708,coexpression:0.603,textmining:0.345\", \"String_Score_Physical\": 941, \"String_Score_Combined\": 984}", "selected": false}, "selected": false}, {"data": {"id": "2783961", "source": "2782755", "target": "2782797", "SCORE": "904", "shared_interaction": "Direct", "source_original": "853716", "Name_B": "RIP1", "Name_A": "SDH3", "target_original": "856689", "Gene_B": "856689", "interaction": "Direct", "SUID": 2783961, "TYPE": "Direct", "Gene_A": "853716", "support": "{\"StringDB\": \"physical\", \"StringEvidence\": \"database:0.896,coexpression:0.818,textmining:0.643,experiments:0.045\", \"BioGrid\": \"genetic\", \"BioGrid_PubMed\": \"16941010\", \"BioGrid_Type\": \"Synthetic Lethality\", \"String_Score_Physical\": 904, \"String_Score_Combined\": 993}", "selected": false}, "selected": false}, {"data": {"id": "2783958", "source": "2782755", "target": "2782791", "SCORE": "900", "shared_interaction": "Direct", "source_original": "853716", "Name_B": "QCR10", "Name_A": "SDH3", "target_original": "856390", "Gene_B": "856390", "interaction": "Direct", "SUID": 2783958, "TYPE": "Direct", "Gene_A": "853716", "support": "{\"StringDB\": \"physical\", \"StringEvidence\": \"database:0.896,coexpression:0.518,textmining:0.039\", \"String_Score_Physical\": 900, \"String_Score_Combined\": 953}", "selected": false}, "selected": false}, {"data": {"id": "2783955", "source": "2782755", "target": "2782788", "SCORE": "900", "shared_interaction": "Direct", "source_original": "853716", "Name_B": "QCR2", "Name_A": "SDH3", "target_original": "856321", "Gene_B": "856321", "interaction": "Direct", "SUID": 2783955, "TYPE": "Direct", "Gene_A": "853716", "support": "{\"StringDB\": \"physical\", \"StringEvidence\": \"database:0.896,coexpression:0.755,textmining:0.474\", \"String_Score_Physical\": 900, \"String_Score_Combined\": 987}", "selected": false}, "selected": false}, {"data": {"id": "2783952", "source": "2782755", "target": "2782773", "SCORE": "900", "shared_interaction": "Direct", "source_original": "853716", "Name_B": "NDI1", "Name_A": "SDH3", "target_original": "854919", "Gene_B": "854919", "interaction": "Direct", "SUID": 2783952, "TYPE": "Direct", "Gene_A": "853716", "support": "{\"StringDB\": \"physical\", \"StringEvidence\": \"database:0.896,coexpression:0.457,textmining:0.034\", \"String_Score_Physical\": 900, \"String_Score_Combined\": 947}", "selected": false}, "selected": false}, {"data": {"id": "2783949", "source": "2782755", "target": "2782764", "SCORE": "929", "shared_interaction": "Direct", "source_original": "853716", "Name_B": "CYT1", "Name_A": "SDH3", "target_original": "854231", "Gene_B": "854231", "interaction": "Direct", "SUID": 2783949, "TYPE": "Direct", "Gene_A": "853716", "support": "{\"StringDB\": \"physical\", \"StringEvidence\": \"database:0.896,textmining:0.436,coexpression:0.411,experiments:0.297\", \"BioGrid\": \"physical\", \"BioGrid_PubMed\": \"14672930\", \"BioGrid_Type\": \"Co-localization\", \"String_Score_Physical\": 929, \"String_Score_Combined\": 976}", "selected": false}, "selected": false}, {"data": {"id": "2783916", "source": "2782752", "target": "2782797", "SCORE": "999", "shared_interaction": "Direct", "source_original": "853273", "Name_B": "RIP1", "Name_A": "QCR8", "target_original": "856689", "Gene_B": "856689", "interaction": "Direct", "SUID": 2783916, "TYPE": "Direct", "Gene_A": "853273", "support": "{\"StringDB\": \"physical\", \"StringEvidence\": \"experiments:0.995,coexpression:0.950,textmining:0.938,database:0.896\", \"String_Score_Physical\": 999, \"String_Score_Combined\": 999}", "selected": false}, "selected": false}, {"data": {"id": "2783913", "source": "2782752", "target": "2782794", "SCORE": "981", "shared_interaction": "Direct", "source_original": "853273", "Name_B": "CYC7", "Name_A": "QCR8", "target_original": "856672", "Gene_B": "856672", "interaction": "Direct", "SUID": 2783913, "TYPE": "Direct", "Gene_A": "853273", "support": "{\"StringDB\": \"physical\", \"StringEvidence\": \"database:0.896,experiments:0.819,textmining:0.574,coexpression:0.328\", \"String_Score_Physical\": 981, \"String_Score_Combined\": 994}", "selected": false}, "selected": false}, {"data": {"id": "2783910", "source": "2782752", "target": "2782791", "SCORE": "979", "shared_interaction": "Direct", "source_original": "853273", "Name_B": "QCR10", "Name_A": "QCR8", "target_original": "856390", "Gene_B": "856390", "interaction": "Direct", "SUID": 2783910, "TYPE": "Direct", "Gene_A": "853273", "support": "{\"StringDB\": \"physical\", \"StringEvidence\": \"textmining:0.897,database:0.896,coexpression:0.800,experiments:0.791\", \"String_Score_Physical\": 979, \"String_Score_Combined\": 999}", "selected": false}, "selected": false}, {"data": {"id": "2783907", "source": "2782752", "target": "2782788", "SCORE": "999", "shared_interaction": "Direct", "source_original": "853273", "Name_B": "QCR2", "Name_A": "QCR8", "target_original": "856321", "Gene_B": "856321", "interaction": "Direct", "SUID": 2783907, "TYPE": "Direct", "Gene_A": "853273", "support": "{\"StringDB\": \"physical\", \"StringEvidence\": \"experiments:0.998,textmining:0.967,database:0.896,coexpression:0.798\", \"BioGrid\": \"physical\", \"BioGrid_PubMed\": \"25683140,22940862,21807901\", \"BioGrid_Type\": \"Affinity Capture-MS,Affinity Capture-Western\", \"String_Score_Physical\": 999, \"String_Score_Combined\": 999}", "selected": false}, "selected": false}, {"data": {"id": "2783904", "source": "2782752", "target": "2782785", "SCORE": "770", "shared_interaction": "Direct", "source_original": "853273", "Name_B": "ATP20", "Name_A": "QCR8", "target_original": "856131", "Gene_B": "856131", "interaction": "Direct", "SUID": 2783904, "TYPE": "Direct", "Gene_A": "853273", "support": "{\"StringDB\": \"physical\", \"StringEvidence\": \"coexpression:0.960,experiments:0.760,textmining:0.515\", \"String_Score_Physical\": 770, \"String_Score_Combined\": 995}", "selected": false}, "selected": false}, {"data": {"id": "2783901", "source": "2782752", "target": "2782782", "SCORE": "763", "shared_interaction": "Direct", "source_original": "853273", "Name_B": "ATP4", "Name_A": "QCR8", "target_original": "856027", "Gene_B": "856027", "interaction": "Direct", "SUID": 2783901, "TYPE": "Direct", "Gene_A": "853273", "support": "{\"StringDB\": \"physical\", \"StringEvidence\": \"coexpression:0.901,experiments:0.753,textmining:0.566\", \"String_Score_Physical\": 763, \"String_Score_Combined\": 989}", "selected": false}, "selected": false}, {"data": {"id": "2783898", "source": "2782752", "target": "2782779", "SCORE": "998", "shared_interaction": "Direct", "source_original": "853273", "Name_B": "COX5A", "Name_A": "QCR8", "target_original": "855675", "Gene_B": "855675", "interaction": "Direct", "SUID": 2783898, "TYPE": "Direct", "Gene_A": "853273", "support": "{\"StringDB\": \"physical\", \"StringEvidence\": \"experiments:0.986,database:0.896,coexpression:0.579,textmining:0.065\", \"BioGrid\": \"physical\", \"BioGrid_PubMed\": \"30598554\", \"BioGrid_Type\": \"Co-crystal Structure\", \"String_Score_Physical\": 998, \"String_Score_Combined\": 999}", "selected": false}, "selected": false}, {"data": {"id": "2783895", "source": "2782752", "target": "2782776", "SCORE": "979", "shared_interaction": "Direct", "source_original": "853273", "Name_B": "COX7", "Name_A": "QCR8", "target_original": "855298", "Gene_B": "855298", "interaction": "Direct", "SUID": 2783895, "TYPE": "Direct", "Gene_A": "853273", "support": "{\"StringDB\": \"physical\", \"StringEvidence\": \"database:0.896,experiments:0.797,coexpression:0.762,textmining:0.429\", \"BioGrid\": \"genetic\", \"BioGrid_PubMed\": \"27708008,20093466\", \"BioGrid_Type\": \"Positive Genetic\", \"BioGrid_Score\": \"0.2542,0.2734\", \"String_Score_Physical\": 979, \"String_Score_Combined\": 997}", "selected": false}, "selected": false}, {"data": {"id": "2783892", "source": "2782752", "target": "2782773", "SCORE": "900", "shared_interaction": "Direct", "source_original": "853273", "Name_B": "NDI1", "Name_A": "QCR8", "target_original": "854919", "Gene_B": "854919", "interaction": "Direct", "SUID": 2783892, "TYPE": "Direct", "Gene_A": "853273", "support": "{\"StringDB\": \"physical\", \"StringEvidence\": \"database:0.896,coexpression:0.200,textmining:0.163\", \"String_Score_Physical\": 900, \"String_Score_Combined\": 933}", "selected": false}, "selected": false}, {"data": {"id": "2783889", "source": "2782752", "target": "2782767", "SCORE": "984", "shared_interaction": "Direct", "source_original": "853273", "Name_B": "COX5B", "Name_A": "QCR8", "target_original": "854695", "Gene_B": "854695", "interaction": "Direct", "SUID": 2783889, "TYPE": "Direct", "Gene_A": "853273", "support": "{\"StringDB\": \"physical\", \"StringEvidence\": \"database:0.896,experiments:0.845,textmining:0.502,coexpression:0.231\", \"String_Score_Physical\": 984, \"String_Score_Combined\": 994}", "selected": false}, "selected": false}, {"data": {"id": "2783886", "source": "2782752", "target": "2782764", "SCORE": "999", "shared_interaction": "Direct", "source_original": "853273", "Name_B": "CYT1", "Name_A": "QCR8", "target_original": "854231", "Gene_B": "854231", "interaction": "Direct", "SUID": 2783886, "TYPE": "Direct", "Gene_A": "853273", "support": "{\"StringDB\": \"physical\", \"StringEvidence\": \"experiments:0.997,textmining:0.987,coexpression:0.929,database:0.896\", \"String_Score_Physical\": 999, \"String_Score_Combined\": 999}", "selected": false}, "selected": false}, {"data": {"id": "2783883", "source": "2782752", "target": "2782758", "SCORE": "197", "shared_interaction": "Direct", "source_original": "853273", "Name_B": "ATP7", "Name_A": "QCR8", "target_original": "853853", "Gene_B": "853853", "interaction": "Direct", "SUID": 2783883, "TYPE": "Direct", "Gene_A": "853273", "support": "{\"StringDB\": \"physical\", \"StringEvidence\": \"coexpression:0.970,experiments:0.163,textmining:0.139\", \"BioGrid\": \"genetic\", \"BioGrid_PubMed\": \"21623372\", \"BioGrid_Type\": \"Negative Genetic\", \"BioGrid_Score\": \"-0.1175\", \"String_Score_Physical\": 197, \"String_Score_Combined\": 979}", "selected": false}, "selected": false}, {"data": {"id": "2783880", "source": "2782752", "target": "2782755", "SCORE": "900", "shared_interaction": "Direct", "source_original": "853273", "Name_B": "SDH3", "Name_A": "QCR8", "target_original": "853716", "Gene_B": "853716", "interaction": "Direct", "SUID": 2783880, "TYPE": "Direct", "Gene_A": "853273", "support": "{\"StringDB\": \"physical\", \"StringEvidence\": \"coexpression:0.907,database:0.896,textmining:0.188\", \"String_Score_Physical\": 900, \"String_Score_Combined\": 992}", "selected": false}, "selected": false}, {"data": {"id": "2783865", "source": "2782749", "target": "2782797", "SCORE": "999", "shared_interaction": "Direct", "source_original": "853095", "Name_B": "RIP1", "Name_A": "QCR9", "target_original": "856689", "Gene_B": "856689", "interaction": "Direct", "SUID": 2783865, "TYPE": "Direct", "Gene_A": "853095", "support": "{\"StringDB\": \"physical\", \"StringEvidence\": \"experiments:0.995,textmining:0.955,coexpression:0.948,database:0.896\", \"String_Score_Physical\": 999, \"String_Score_Combined\": 999}", "selected": false}, "selected": false}, {"data": {"id": "2783862", "source": "2782749", "target": "2782794", "SCORE": "982", "shared_interaction": "Direct", "source_original": "853095", "Name_B": "CYC7", "Name_A": "QCR9", "target_original": "856672", "Gene_B": "856672", "interaction": "Direct", "SUID": 2783862, "TYPE": "Direct", "Gene_A": "853095", "support": "{\"StringDB\": \"physical\", \"StringEvidence\": \"database:0.896,experiments:0.826,textmining:0.603,coexpression:0.251\", \"String_Score_Physical\": 982, \"String_Score_Combined\": 994}", "selected": false}, "selected": false}, {"data": {"id": "2783859", "source": "2782749", "target": "2782791", "SCORE": "979", "shared_interaction": "Direct", "source_original": "853095", "Name_B": "QCR10", "Name_A": "QCR9", "target_original": "856390", "Gene_B": "856390", "interaction": "Direct", "SUID": 2783859, "TYPE": "Direct", "Gene_A": "853095", "support": "{\"StringDB\": \"physical\", \"StringEvidence\": \"coexpression:0.919,textmining:0.897,database:0.896,experiments:0.799\", \"BioGrid\": \"genetic\", \"BioGrid_PubMed\": \"21623372\", \"BioGrid_Type\": \"Positive Genetic\", \"BioGrid_Score\": \"0.1131\", \"String_Score_Physical\": 979, \"String_Score_Combined\": 999}", "selected": false}, "selected": false}, {"data": {"id": "2783856", "source": "2782749", "target": "2782788", "SCORE": "999", "shared_interaction": "Direct", "source_original": "853095", "Name_B": "QCR2", "Name_A": "QCR9", "target_original": "856321", "Gene_B": "856321", "interaction": "Direct", "SUID": 2783856, "TYPE": "Direct", "Gene_A": "853095", "support": "{\"StringDB\": \"physical\", \"StringEvidence\": \"experiments:0.997,textmining:0.966,database:0.896,coexpression:0.647\", \"BioGrid\": \"physical\", \"BioGrid_PubMed\": \"25683140,21807901\", \"BioGrid_Type\": \"Affinity Capture-MS,Affinity Capture-Western\", \"String_Score_Physical\": 999, \"String_Score_Combined\": 999}", "selected": false}, "selected": false}, {"data": {"id": "2783853", "source": "2782749", "target": "2782785", "SCORE": "240", "shared_interaction": "Direct", "source_original": "853095", "Name_B": "ATP20", "Name_A": "QCR9", "target_original": "856131", "Gene_B": "856131", "interaction": "Direct", "SUID": 2783853, "TYPE": "Direct", "Gene_A": "853095", "support": "{\"StringDB\": \"physical\", \"StringEvidence\": \"coexpression:0.957,textmining:0.442,experiments:0.209\", \"String_Score_Physical\": 240, \"String_Score_Combined\": 981}", "selected": false}, "selected": false}, {"data": {"id": "2783850", "source": "2782749", "target": "2782782", "SCORE": "275", "shared_interaction": "Direct", "source_original": "853095", "Name_B": "ATP4", "Name_A": "QCR9", "target_original": "856027", "Gene_B": "856027", "interaction": "Direct", "SUID": 2783850, "TYPE": "Direct", "Gene_A": "853095", "support": "{\"StringDB\": \"physical\", \"StringEvidence\": \"coexpression:0.358,textmining:0.281,experiments:0.244\", \"String_Score_Physical\": 275, \"String_Score_Combined\": 665}", "selected": false}, "selected": false}, {"data": {"id": "2783847", "source": "2782749", "target": "2782779", "SCORE": "989", "shared_interaction": "Direct", "source_original": "853095", "Name_B": "COX5A", "Name_A": "QCR9", "target_original": "855675", "Gene_B": "855675", "interaction": "Direct", "SUID": 2783847, "TYPE": "Direct", "Gene_A": "853095", "support": "{\"StringDB\": \"physical\", \"StringEvidence\": \"coexpression:0.917,database:0.896,experiments:0.893,textmining:0.334\", \"String_Score_Physical\": 989, \"String_Score_Combined\": 999}", "selected": false}, "selected": false}, {"data": {"id": "2783844", "source": "2782749", "target": "2782776", "SCORE": "979", "shared_interaction": "Direct", "source_original": "853095", "Name_B": "COX7", "Name_A": "QCR9", "target_original": "855298", "Gene_B": "855298", "interaction": "Direct", "SUID": 2783844, "TYPE": "Direct", "Gene_A": "853095", "support": "{\"StringDB\": \"physical\", \"StringEvidence\": \"database:0.896,coexpression:0.858,experiments:0.791,textmining:0.552\", \"String_Score_Physical\": 979, \"String_Score_Combined\": 998}", "selected": false}, "selected": false}, {"data": {"id": "2783841", "source": "2782749", "target": "2782773", "SCORE": "900", "shared_interaction": "Direct", "source_original": "853095", "Name_B": "NDI1", "Name_A": "QCR9", "target_original": "854919", "Gene_B": "854919", "interaction": "Direct", "SUID": 2783841, "TYPE": "Direct", "Gene_A": "853095", "support": "{\"StringDB\": \"physical\", \"StringEvidence\": \"database:0.896,coexpression:0.293\", \"String_Score_Physical\": 900, \"String_Score_Combined\": 929}", "selected": false}, "selected": false}, {"data": {"id": "2783838", "source": "2782749", "target": "2782767", "SCORE": "989", "shared_interaction": "Direct", "source_original": "853095", "Name_B": "COX5B", "Name_A": "QCR9", "target_original": "854695", "Gene_B": "854695", "interaction": "Direct", "SUID": 2783838, "TYPE": "Direct", "Gene_A": "853095", "support": "{\"StringDB\": \"physical\", \"StringEvidence\": \"database:0.896,experiments:0.893,textmining:0.221,coexpression:0.133\", \"String_Score_Physical\": 989, \"String_Score_Combined\": 992}", "selected": false}, "selected": false}, {"data": {"id": "2783835", "source": "2782749", "target": "2782764", "SCORE": "999", "shared_interaction": "Direct", "source_original": "853095", "Name_B": "CYT1", "Name_A": "QCR9", "target_original": "854231", "Gene_B": "854231", "interaction": "Direct", "SUID": 2783835, "TYPE": "Direct", "Gene_A": "853095", "support": "{\"StringDB\": \"physical\", \"StringEvidence\": \"experiments:0.999,textmining:0.981,coexpression:0.979,database:0.896\", \"BioGrid\": \"genetic\", \"BioGrid_PubMed\": \"27708008,21623372,20093466,11717439\", \"BioGrid_Type\": \"Positive Genetic,Synthetic Growth Defect\", \"BioGrid_Score\": \"0.1026,0.2432,0.2515\", \"String_Score_Physical\": 999, \"String_Score_Combined\": 999}", "selected": false}, "selected": false}, {"data": {"id": "2783832", "source": "2782749", "target": "2782758", "SCORE": "275", "shared_interaction": "Direct", "source_original": "853095", "Name_B": "ATP7", "Name_A": "QCR9", "target_original": "853853", "Gene_B": "853853", "interaction": "Direct", "SUID": 2783832, "TYPE": "Direct", "Gene_A": "853095", "support": "{\"StringDB\": \"physical\", \"StringEvidence\": \"coexpression:0.956,experiments:0.244,textmining:0.079\", \"String_Score_Physical\": 275, \"String_Score_Combined\": 970}", "selected": false}, "selected": false}, {"data": {"id": "2783829", "source": "2782749", "target": "2782755", "SCORE": "902", "shared_interaction": "Direct", "source_original": "853095", "Name_B": "SDH3", "Name_A": "QCR9", "target_original": "853716", "Gene_B": "853716", "interaction": "Direct", "SUID": 2783829, "TYPE": "Direct", "Gene_A": "853095", "support": "{\"StringDB\": \"physical\", \"StringEvidence\": \"database:0.896,textmining:0.350,coexpression:0.308,experiments:0.028\", \"BioGrid\": \"genetic\", \"BioGrid_PubMed\": \"21623372\", \"BioGrid_Type\": \"Negative Genetic\", \"BioGrid_Score\": \"-0.1105\", \"String_Score_Physical\": 902, \"String_Score_Combined\": 956}", "selected": false}, "selected": false}, {"data": {"id": "2783826", "source": "2782749", "target": "2782752", "SCORE": "999", "shared_interaction": "Direct", "source_original": "853095", "Name_B": "QCR8", "Name_A": "QCR9", "target_original": "853273", "Gene_B": "853273", "interaction": "Direct", "SUID": 2783826, "TYPE": "Direct", "Gene_A": "853095", "support": "{\"StringDB\": \"physical\", \"StringEvidence\": \"experiments:0.995,textmining:0.987,coexpression:0.984,database:0.896\", \"String_Score_Physical\": 999, \"String_Score_Combined\": 999}", "selected": false}, "selected": false}, {"data": {"id": "2783820", "source": "2782746", "target": "2782797", "SCORE": "999", "shared_interaction": "Direct", "source_original": "852688", "Name_B": "RIP1", "Name_A": "COX4", "target_original": "856689", "Gene_B": "856689", "interaction": "Direct", "SUID": 2783820, "TYPE": "Direct", "Gene_A": "852688", "support": "{\"StringDB\": \"physical\", \"StringEvidence\": \"experiments:0.999,coexpression:0.996,database:0.896,textmining:0.863\", \"BioGrid\": \"physical\", \"BioGrid_PubMed\": \"29386296,27662906,27083394,24928273,23864706,22342701\", \"BioGrid_Type\": \"Affinity Capture-MS,Affinity Capture-Western\", \"String_Score_Physical\": 999, \"String_Score_Combined\": 999}", "selected": false}, "selected": false}, {"data": {"id": "2783817", "source": "2782746", "target": "2782794", "SCORE": "788", "shared_interaction": "Direct", "source_original": "852688", "Name_B": "CYC7", "Name_A": "COX4", "target_original": "856672", "Gene_B": "856672", "interaction": "Direct", "SUID": 2783817, "TYPE": "Direct", "Gene_A": "852688", "support": "{\"StringDB\": \"physical\", \"StringEvidence\": \"experiments:0.779,textmining:0.697,coexpression:0.230\", \"BioGrid\": \"physical\", \"BioGrid_PubMed\": \"23864706\", \"BioGrid_Type\": \"Affinity Capture-MS\", \"String_Score_Physical\": 788, \"String_Score_Combined\": 950}", "selected": false}, "selected": false}, {"data": {"id": "2783814", "source": "2782746", "target": "2782791", "SCORE": "987", "shared_interaction": "Direct", "source_original": "852688", "Name_B": "QCR10", "Name_A": "COX4", "target_original": "856390", "Gene_B": "856390", "interaction": "Direct", "SUID": 2783814, "TYPE": "Direct", "Gene_A": "852688", "support": "{\"StringDB\": \"physical\", \"StringEvidence\": \"coexpression:0.938,database:0.896,experiments:0.879,textmining:0.594\", \"BioGrid\": \"physical\", \"BioGrid_PubMed\": \"29688796,23864706\", \"BioGrid_Type\": \"Affinity Capture-MS,Affinity Capture-Western\", \"String_Score_Physical\": 987, \"String_Score_Combined\": 999}", "selected": false}, "selected": false}, {"data": {"id": "2783811", "source": "2782746", "target": "2782788", "SCORE": "999", "shared_interaction": "Direct", "source_original": "852688", "Name_B": "QCR2", "Name_A": "COX4", "target_original": "856321", "Gene_B": "856321", "interaction": "Direct", "SUID": 2783811, "TYPE": "Direct", "Gene_A": "852688", "support": "{\"StringDB\": \"physical\", \"StringEvidence\": \"experiments:0.995,coexpression:0.967,database:0.896,textmining:0.827\", \"BioGrid\": \"physical\", \"BioGrid_PubMed\": \"23864706\", \"BioGrid_Type\": \"Affinity Capture-MS\", \"String_Score_Physical\": 999, \"String_Score_Combined\": 999}", "selected": false}, "selected": false}, {"data": {"id": "2783808", "source": "2782746", "target": "2782785", "SCORE": "819", "shared_interaction": "Direct", "source_original": "852688", "Name_B": "ATP20", "Name_A": "COX4", "target_original": "856131", "Gene_B": "856131", "interaction": "Direct", "SUID": 2783808, "TYPE": "Direct", "Gene_A": "852688", "support": "{\"StringDB\": \"physical\", \"StringEvidence\": \"coexpression:0.989,experiments:0.811,textmining:0.248\", \"String_Score_Physical\": 819, \"String_Score_Combined\": 998}", "selected": false}, "selected": false}, {"data": {"id": "2783805", "source": "2782746", "target": "2782782", "SCORE": "819", "shared_interaction": "Direct", "source_original": "852688", "Name_B": "ATP4", "Name_A": "COX4", "target_original": "856027", "Gene_B": "856027", "interaction": "Direct", "SUID": 2783805, "TYPE": "Direct", "Gene_A": "852688", "support": "{\"StringDB\": \"physical\", \"StringEvidence\": \"coexpression:0.983,experiments:0.811,textmining:0.449\", \"String_Score_Physical\": 819, \"String_Score_Combined\": 998}", "selected": false}, "selected": false}, {"data": {"id": "2783802", "source": "2782746", "target": "2782779", "SCORE": "999", "shared_interaction": "Direct", "source_original": "852688", "Name_B": "COX5A", "Name_A": "COX4", "target_original": "855675", "Gene_B": "855675", "interaction": "Direct", "SUID": 2783802, "TYPE": "Direct", "Gene_A": "852688", "support": "{\"StringDB\": \"physical\", \"StringEvidence\": \"experiments:0.991,coexpression:0.990,textmining:0.954,database:0.896\", \"BioGrid\": \"physical\", \"BioGrid_PubMed\": \"27083394,23864706,16554755\", \"BioGrid_Type\": \"Affinity Capture-MS,Affinity Capture-Western\", \"String_Score_Physical\": 999, \"String_Score_Combined\": 999}", "selected": false}, "selected": false}, {"data": {"id": "2783799", "source": "2782746", "target": "2782776", "SCORE": "968", "shared_interaction": "Direct", "source_original": "852688", "Name_B": "COX7", "Name_A": "COX4", "target_original": "855298", "Gene_B": "855298", "interaction": "Direct", "SUID": 2783799, "TYPE": "Direct", "Gene_A": "852688", "support": "{\"StringDB\": \"physical\", \"StringEvidence\": \"textmining:0.900,coexpression:0.896,experiments:0.845,database:0.791\", \"BioGrid\": \"physical\", \"BioGrid_PubMed\": \"23864706\", \"BioGrid_Type\": \"Affinity Capture-MS\", \"String_Score_Physical\": 968, \"String_Score_Combined\": 999}", "selected": false}, "selected": false}, {"data": {"id": "2783796", "source": "2782746", "target": "2782773", "SCORE": "925", "shared_interaction": "Direct", "source_original": "852688", "Name_B": "NDI1", "Name_A": "COX4", "target_original": "854919", "Gene_B": "854919", "interaction": "Direct", "SUID": 2783796, "TYPE": "Direct", "Gene_A": "852688", "support": "{\"StringDB\": \"physical\", \"StringEvidence\": \"database:0.896,coexpression:0.811,textmining:0.296,experiments:0.258\", \"BioGrid\": \"physical\", \"BioGrid_PubMed\": \"23864706\", \"BioGrid_Type\": \"Affinity Capture-MS\", \"String_Score_Physical\": 925, \"String_Score_Combined\": 990}", "selected": false}, "selected": false}, {"data": {"id": "2783793", "source": "2782746", "target": "2782767", "SCORE": "996", "shared_interaction": "Direct", "source_original": "852688", "Name_B": "COX5B", "Name_A": "COX4", "target_original": "854695", "Gene_B": "854695", "interaction": "Direct", "SUID": 2783793, "TYPE": "Direct", "Gene_A": "852688", "support": "{\"StringDB\": \"physical\", \"StringEvidence\": \"experiments:0.987,textmining:0.955,database:0.708,coexpression:0.627\", \"BioGrid\": \"physical\", \"BioGrid_PubMed\": \"27083394,23864706\", \"BioGrid_Type\": \"Affinity Capture-MS,Affinity Capture-Western\", \"String_Score_Physical\": 996, \"String_Score_Combined\": 999}", "selected": false}, "selected": false}, {"data": {"id": "2783790", "source": "2782746", "target": "2782764", "SCORE": "999", "shared_interaction": "Direct", "source_original": "852688", "Name_B": "CYT1", "Name_A": "COX4", "target_original": "854231", "Gene_B": "854231", "interaction": "Direct", "SUID": 2783790, "TYPE": "Direct", "Gene_A": "852688", "support": "{\"StringDB\": \"physical\", \"StringEvidence\": \"experiments:0.998,coexpression:0.992,textmining:0.937,database:0.896\", \"BioGrid\": \"physical\", \"BioGrid_PubMed\": \"29386296,23864706,22310663\", \"BioGrid_Type\": \"Affinity Capture-MS,Affinity Capture-Western\", \"String_Score_Physical\": 999, \"String_Score_Combined\": 999}", "selected": false}, "selected": false}, {"data": {"id": "2783787", "source": "2782746", "target": "2782752", "SCORE": "999", "shared_interaction": "Direct", "source_original": "852688", "Name_B": "QCR8", "Name_A": "COX4", "target_original": "853273", "Gene_B": "853273", "interaction": "Direct", "SUID": 2783787, "TYPE": "Direct", "Gene_A": "852688", "support": "{\"StringDB\": \"physical\", \"StringEvidence\": \"experiments:0.998,coexpression:0.978,database:0.896,textmining:0.609\", \"BioGrid\": \"physical\", \"BioGrid_PubMed\": \"27083394,23864706,22342701\", \"BioGrid_Type\": \"Affinity Capture-MS,Affinity Capture-Western\", \"String_Score_Physical\": 999, \"String_Score_Combined\": 999}", "selected": false}, "selected": false}, {"data": {"id": "2783784", "source": "2782746", "target": "2782749", "SCORE": "993", "shared_interaction": "Direct", "source_original": "852688", "Name_B": "QCR9", "Name_A": "COX4", "target_original": "853095", "Gene_B": "853095", "interaction": "Direct", "SUID": 2783784, "TYPE": "Direct", "Gene_A": "852688", "support": "{\"StringDB\": \"physical\", \"StringEvidence\": \"coexpression:0.996,experiments:0.938,database:0.896,textmining:0.581\", \"BioGrid\": \"physical\", \"BioGrid_PubMed\": \"23864706\", \"BioGrid_Type\": \"Affinity Capture-MS\", \"String_Score_Physical\": 993, \"String_Score_Combined\": 999}", "selected": false}, "selected": false}, {"data": {"id": "2783763", "source": "2782743", "target": "2782797", "SCORE": "254", "shared_interaction": "Direct", "source_original": "852327", "Name_B": "RIP1", "Name_A": "ATP3", "target_original": "856689", "Gene_B": "856689", "interaction": "Direct", "SUID": 2783763, "TYPE": "Direct", "Gene_A": "852327", "support": "{\"StringDB\": \"physical\", \"StringEvidence\": \"coexpression:0.974,textmining:0.632,experiments:0.223\", \"String_Score_Physical\": 254, \"String_Score_Combined\": 992}", "selected": false}, "selected": false}, {"data": {"id": "2783760", "source": "2782743", "target": "2782794", "SCORE": "193", "shared_interaction": "Direct", "source_original": "852327", "Name_B": "CYC7", "Name_A": "ATP3", "target_original": "856672", "Gene_B": "856672", "interaction": "Direct", "SUID": 2783760, "TYPE": "Direct", "Gene_A": "852327", "support": "{\"StringDB\": \"physical\", \"StringEvidence\": \"coexpression:0.628,experiments:0.158,textmining:0.043\", \"String_Score_Physical\": 193, \"String_Score_Combined\": 712}", "selected": false}, "selected": false}, {"data": {"id": "2783757", "source": "2782743", "target": "2782788", "SCORE": "365", "shared_interaction": "Direct", "source_original": "852327", "Name_B": "QCR2", "Name_A": "ATP3", "target_original": "856321", "Gene_B": "856321", "interaction": "Direct", "SUID": 2783757, "TYPE": "Direct", "Gene_A": "852327", "support": "{\"StringDB\": \"physical\", \"StringEvidence\": \"coexpression:0.956,textmining:0.581,experiments:0.338\", \"String_Score_Physical\": 365, \"String_Score_Combined\": 988}", "selected": false}, "selected": false}, {"data": {"id": "2783754", "source": "2782743", "target": "2782785", "SCORE": "988", "shared_interaction": "Direct", "source_original": "852327", "Name_B": "ATP20", "Name_A": "ATP3", "target_original": "856131", "Gene_B": "856131", "interaction": "Direct", "SUID": 2783754, "TYPE": "Direct", "Gene_A": "852327", "support": "{\"StringDB\": \"physical\", \"StringEvidence\": \"experiments:0.960,coexpression:0.941,textmining:0.790,database:0.708\", \"String_Score_Physical\": 988, \"String_Score_Combined\": 999}", "selected": false}, "selected": false}, {"data": {"id": "2783751", "source": "2782743", "target": "2782782", "SCORE": "999", "shared_interaction": "Direct", "source_original": "852327", "Name_B": "ATP4", "Name_A": "ATP3", "target_original": "856027", "Gene_B": "856027", "interaction": "Direct", "SUID": 2783751, "TYPE": "Direct", "Gene_A": "852327", "support": "{\"StringDB\": \"physical\", \"StringEvidence\": \"experiments:0.999,textmining:0.946,coexpression:0.941,database:0.896\", \"BioGrid\": \"physical\", \"BioGrid_PubMed\": \"9867807\", \"BioGrid_Type\": \"Co-purification\", \"String_Score_Physical\": 999, \"String_Score_Combined\": 999}", "selected": false}, "selected": false}, {"data": {"id": "2783748", "source": "2782743", "target": "2782779", "SCORE": "190", "shared_interaction": "Direct", "source_original": "852327", "Name_B": "COX5A", "Name_A": "ATP3", "target_original": "855675", "Gene_B": "855675", "interaction": "Direct", "SUID": 2783748, "TYPE": "Direct", "Gene_A": "852327", "support": "{\"StringDB\": \"physical\", \"StringEvidence\": \"coexpression:0.730,textmining:0.478,experiments:0.155\", \"String_Score_Physical\": 190, \"String_Score_Combined\": 885}", "selected": false}, "selected": false}, {"data": {"id": "2783745", "source": "2782743", "target": "2782770", "SCORE": "979", "shared_interaction": "Direct", "source_original": "852327", "Name_B": "ATP18", "Name_A": "ATP3", "target_original": "854893", "Gene_B": "854893", "interaction": "Direct", "SUID": 2783745, "TYPE": "Direct", "Gene_A": "852327", "support": "{\"StringDB\": \"physical\", \"StringEvidence\": \"database:0.896,experiments:0.791,textmining:0.600,coexpression:0.081\", \"String_Score_Physical\": 979, \"String_Score_Combined\": 992}", "selected": false}, "selected": false}, {"data": {"id": "2783742", "source": "2782743", "target": "2782767", "SCORE": "190", "shared_interaction": "Direct", "source_original": "852327", "Name_B": "COX5B", "Name_A": "ATP3", "target_original": "854695", "Gene_B": "854695", "interaction": "Direct", "SUID": 2783742, "TYPE": "Direct", "Gene_A": "852327", "support": "{\"StringDB\": \"physical\", \"StringEvidence\": \"coexpression:0.302,textmining:0.179,experiments:0.155\", \"String_Score_Physical\": 190, \"String_Score_Combined\": 536}", "selected": false}, "selected": false}, {"data": {"id": "2783739", "source": "2782743", "target": "2782764", "SCORE": "258", "shared_interaction": "Direct", "source_original": "852327", "Name_B": "CYT1", "Name_A": "ATP3", "target_original": "854231", "Gene_B": "854231", "interaction": "Direct", "SUID": 2783739, "TYPE": "Direct", "Gene_A": "852327", "support": "{\"StringDB\": \"physical\", \"StringEvidence\": \"coexpression:0.914,textmining:0.485,experiments:0.226\", \"String_Score_Physical\": 258, \"String_Score_Combined\": 967}", "selected": false}, "selected": false}, {"data": {"id": "2783736", "source": "2782743", "target": "2782761", "SCORE": "941", "shared_interaction": "Direct", "source_original": "852327", "Name_B": "ATP19", "Name_A": "ATP3", "target_original": "854077", "Gene_B": "854077", "interaction": "Direct", "SUID": 2783736, "TYPE": "Direct", "Gene_A": "852327", "support": "{\"StringDB\": \"physical\", \"StringEvidence\": \"experiments:0.791,database:0.708,coexpression:0.231\", \"String_Score_Physical\": 941, \"String_Score_Combined\": 955}", "selected": false}, "selected": false}, {"data": {"id": "2783733", "source": "2782743", "target": "2782758", "SCORE": "999", "shared_interaction": "Direct", "source_original": "852327", "Name_B": "ATP7", "Name_A": "ATP3", "target_original": "853853", "Gene_B": "853853", "interaction": "Direct", "SUID": 2783733, "TYPE": "Direct", "Gene_A": "852327", "support": "{\"StringDB\": \"physical\", \"StringEvidence\": \"experiments:0.996,coexpression:0.969,textmining:0.913,database:0.896\", \"BioGrid\": \"physical\", \"BioGrid_PubMed\": \"22940862\", \"BioGrid_Type\": \"Affinity Capture-MS\", \"String_Score_Physical\": 999, \"String_Score_Combined\": 999}", "selected": false}, "selected": false}, {"data": {"id": "2783730", "source": "2782743", "target": "2782749", "SCORE": "353", "shared_interaction": "Direct", "source_original": "852327", "Name_B": "QCR9", "Name_A": "ATP3", "target_original": "853095", "Gene_B": "853095", "interaction": "Direct", "SUID": 2783730, "TYPE": "Direct", "Gene_A": "852327", "support": "{\"StringDB\": \"physical\", \"StringEvidence\": \"coexpression:0.360,experiments:0.325,textmining:0.202\", \"String_Score_Physical\": 353, \"String_Score_Combined\": 669}", "selected": false}, "selected": false}, {"data": {"id": "2783712", "source": "2782740", "target": "2782797", "SCORE": "999", "shared_interaction": "Direct", "source_original": "852235", "Name_B": "RIP1", "Name_A": "COR1", "target_original": "856689", "Gene_B": "856689", "interaction": "Direct", "SUID": 2783712, "TYPE": "Direct", "Gene_A": "852235", "support": "{\"StringDB\": \"physical\", \"StringEvidence\": \"experiments:0.999,coexpression:0.991,database:0.896,textmining:0.779\", \"BioGrid\": \"physical\", \"BioGrid_PubMed\": \"29386296,27662906,27030670,24928273,22613836,22342701,17882259\", \"BioGrid_Type\": \"Affinity Capture-MS,Affinity Capture-Western\", \"String_Score_Physical\": 999, \"String_Score_Combined\": 999}", "selected": false}, "selected": false}, {"data": {"id": "2783709", "source": "2782740", "target": "2782794", "SCORE": "981", "shared_interaction": "Direct", "source_original": "852235", "Name_B": "CYC7", "Name_A": "COR1", "target_original": "856672", "Gene_B": "856672", "interaction": "Direct", "SUID": 2783709, "TYPE": "Direct", "Gene_A": "852235", "support": "{\"StringDB\": \"physical\", \"StringEvidence\": \"database:0.896,experiments:0.817,textmining:0.494,coexpression:0.137\", \"String_Score_Physical\": 981, \"String_Score_Combined\": 991}", "selected": false}, "selected": false}, {"data": {"id": "2783706", "source": "2782740", "target": "2782791", "SCORE": "979", "shared_interaction": "Direct", "source_original": "852235", "Name_B": "QCR10", "Name_A": "COR1", "target_original": "856390", "Gene_B": "856390", "interaction": "Direct", "SUID": 2783706, "TYPE": "Direct", "Gene_A": "852235", "support": "{\"StringDB\": \"physical\", \"StringEvidence\": \"textmining:0.897,database:0.896,experiments:0.791,coexpression:0.561\", \"String_Score_Physical\": 979, \"String_Score_Combined\": 999}", "selected": false}, "selected": false}, {"data": {"id": "2783703", "source": "2782740", "target": "2782788", "SCORE": "999", "shared_interaction": "Direct", "source_original": "852235", "Name_B": "QCR2", "Name_A": "COR1", "target_original": "856321", "Gene_B": "856321", "interaction": "Direct", "SUID": 2783703, "TYPE": "Direct", "Gene_A": "852235", "support": "{\"StringDB\": \"physical\", \"StringEvidence\": \"experiments:0.999,coexpression:0.990,textmining:0.969,database:0.896,cooccurrence:0.373\", \"BioGrid\": \"physical\", \"BioGrid_PubMed\": \"32119834,25683140,22342701,21807901,17113393,16554755,16429126,11805826,11283351\", \"BioGrid_Type\": \"Affinity Capture-MS,Co-fractionation,Affinity Capture-Western,Two-hybrid\", \"String_Score_Physical\": 999, \"String_Score_Combined\": 999}", "selected": false}, "selected": false}, {"data": {"id": "2783700", "source": "2782740", "target": "2782785", "SCORE": "262", "shared_interaction": "Direct", "source_original": "852235", "Name_B": "ATP20", "Name_A": "COR1", "target_original": "856131", "Gene_B": "856131", "interaction": "Direct", "SUID": 2783700, "TYPE": "Direct", "Gene_A": "852235", "support": "{\"StringDB\": \"physical\", \"StringEvidence\": \"coexpression:0.363,textmining:0.253,experiments:0.230\", \"String_Score_Physical\": 262, \"String_Score_Combined\": 648}", "selected": false}, "selected": false}, {"data": {"id": "2783697", "source": "2782740", "target": "2782782", "SCORE": "267", "shared_interaction": "Direct", "source_original": "852235", "Name_B": "ATP4", "Name_A": "COR1", "target_original": "856027", "Gene_B": "856027", "interaction": "Direct", "SUID": 2783697, "TYPE": "Direct", "Gene_A": "852235", "support": "{\"StringDB\": \"physical\", \"StringEvidence\": \"coexpression:0.807,textmining:0.397,experiments:0.236\", \"String_Score_Physical\": 267, \"String_Score_Combined\": 914}", "selected": false}, "selected": false}, {"data": {"id": "2783694", "source": "2782740", "target": "2782779", "SCORE": "999", "shared_interaction": "Direct", "source_original": "852235", "Name_B": "COX5A", "Name_A": "COR1", "target_original": "855675", "Gene_B": "855675", "interaction": "Direct", "SUID": 2783694, "TYPE": "Direct", "Gene_A": "852235", "support": "{\"StringDB\": \"physical\", \"StringEvidence\": \"experiments:0.999,database:0.896,coexpression:0.867,textmining:0.651\", \"BioGrid\": \"physical\", \"BioGrid_PubMed\": \"30598554,22342701\", \"BioGrid_Type\": \"Affinity Capture-MS,Co-crystal Structure,Affinity Capture-Western\", \"String_Score_Physical\": 999, \"String_Score_Combined\": 999}", "selected": false}, "selected": false}, {"data": {"id": "2783691", "source": "2782740", "target": "2782776", "SCORE": "987", "shared_interaction": "Direct", "source_original": "852235", "Name_B": "COX7", "Name_A": "COR1", "target_original": "855298", "Gene_B": "855298", "interaction": "Direct", "SUID": 2783691, "TYPE": "Direct", "Gene_A": "852235", "support": "{\"StringDB\": \"physical\", \"StringEvidence\": \"database:0.896,experiments:0.876,textmining:0.624,coexpression:0.216\", \"BioGrid\": \"physical\", \"BioGrid_PubMed\": \"22342701\", \"BioGrid_Type\": \"Affinity Capture-MS\", \"String_Score_Physical\": 987, \"String_Score_Combined\": 996}", "selected": false}, "selected": false}, {"data": {"id": "2783688", "source": "2782740", "target": "2782773", "SCORE": "900", "shared_interaction": "Direct", "source_original": "852235", "Name_B": "NDI1", "Name_A": "COR1", "target_original": "854919", "Gene_B": "854919", "interaction": "Direct", "SUID": 2783688, "TYPE": "Direct", "Gene_A": "852235", "support": "{\"StringDB\": \"physical\", \"StringEvidence\": \"database:0.896,coexpression:0.845,textmining:0.548,neighborhood:0.042\", \"String_Score_Physical\": 900, \"String_Score_Combined\": 993}", "selected": false}, "selected": false}, {"data": {"id": "2783685", "source": "2782740", "target": "2782767", "SCORE": "990", "shared_interaction": "Direct", "source_original": "852235", "Name_B": "COX5B", "Name_A": "COR1", "target_original": "854695", "Gene_B": "854695", "interaction": "Direct", "SUID": 2783685, "TYPE": "Direct", "Gene_A": "852235", "support": "{\"StringDB\": \"physical\", \"StringEvidence\": \"experiments:0.905,database:0.896,textmining:0.248,coexpression:0.169\", \"BioGrid\": \"physical\", \"BioGrid_PubMed\": \"22342701\", \"BioGrid_Type\": \"Affinity Capture-MS\", \"String_Score_Physical\": 990, \"String_Score_Combined\": 994}", "selected": false}, "selected": false}, {"data": {"id": "2783682", "source": "2782740", "target": "2782764", "SCORE": "999", "shared_interaction": "Direct", "source_original": "852235", "Name_B": "CYT1", "Name_A": "COR1", "target_original": "854231", "Gene_B": "854231", "interaction": "Direct", "SUID": 2783682, "TYPE": "Direct", "Gene_A": "852235", "support": "{\"StringDB\": \"physical\", \"StringEvidence\": \"experiments:0.999,coexpression:0.991,textmining:0.924,database:0.896\", \"BioGrid\": \"physical\", \"BioGrid_PubMed\": \"29386296,22342701,22310663,20876281,17882259\", \"BioGrid_Type\": \"Affinity Capture-MS,Affinity Capture-Western\", \"String_Score_Physical\": 999, \"String_Score_Combined\": 999}", "selected": false}, "selected": false}, {"data": {"id": "2783679", "source": "2782740", "target": "2782758", "SCORE": "418", "shared_interaction": "Direct", "source_original": "852235", "Name_B": "ATP7", "Name_A": "COR1", "target_original": "853853", "Gene_B": "853853", "interaction": "Direct", "SUID": 2783679, "TYPE": "Direct", "Gene_A": "852235", "support": "{\"StringDB\": \"physical\", \"StringEvidence\": \"coexpression:0.697,textmining:0.613,experiments:0.394\", \"String_Score_Physical\": 418, \"String_Score_Combined\": 932}", "selected": false}, "selected": false}, {"data": {"id": "2783676", "source": "2782740", "target": "2782755", "SCORE": "900", "shared_interaction": "Direct", "source_original": "852235", "Name_B": "SDH3", "Name_A": "COR1", "target_original": "853716", "Gene_B": "853716", "interaction": "Direct", "SUID": 2783676, "TYPE": "Direct", "Gene_A": "852235", "support": "{\"StringDB\": \"physical\", \"StringEvidence\": \"database:0.896,coexpression:0.404,textmining:0.265\", \"String_Score_Physical\": 900, \"String_Score_Combined\": 956}", "selected": false}, "selected": false}, {"data": {"id": "2783673", "source": "2782740", "target": "2782752", "SCORE": "999", "shared_interaction": "Direct", "source_original": "852235", "Name_B": "QCR8", "Name_A": "COR1", "target_original": "853273", "Gene_B": "853273", "interaction": "Direct", "SUID": 2783673, "TYPE": "Direct", "Gene_A": "852235", "support": "{\"StringDB\": \"physical\", \"StringEvidence\": \"experiments:0.998,textmining:0.931,database:0.896,coexpression:0.540\", \"BioGrid\": \"physical\", \"BioGrid_PubMed\": \"29386296,27030670,22613836,22342701\", \"BioGrid_Type\": \"Affinity Capture-MS,Affinity Capture-Western\", \"String_Score_Physical\": 999, \"String_Score_Combined\": 999}", "selected": false}, "selected": false}, {"data": {"id": "2783670", "source": "2782740", "target": "2782749", "SCORE": "999", "shared_interaction": "Direct", "source_original": "852235", "Name_B": "QCR9", "Name_A": "COR1", "target_original": "853095", "Gene_B": "853095", "interaction": "Direct", "SUID": 2783670, "TYPE": "Direct", "Gene_A": "852235", "support": "{\"StringDB\": \"physical\", \"StringEvidence\": \"experiments:0.995,textmining:0.909,database:0.896,coexpression:0.601\", \"BioGrid\": \"physical\", \"BioGrid_PubMed\": \"22342701\", \"BioGrid_Type\": \"Affinity Capture-MS\", \"String_Score_Physical\": 999, \"String_Score_Combined\": 999}", "selected": false}, "selected": false}, {"data": {"id": "2783667", "source": "2782740", "target": "2782746", "SCORE": "997", "shared_interaction": "Direct", "source_original": "852235", "Name_B": "COX4", "Name_A": "COR1", "target_original": "852688", "Gene_B": "852688", "interaction": "Direct", "SUID": 2783667, "TYPE": "Direct", "Gene_A": "852235", "support": "{\"StringDB\": \"physical\", \"StringEvidence\": \"experiments:0.977,coexpression:0.928,database:0.896,textmining:0.811\", \"BioGrid\": \"physical\", \"BioGrid_PubMed\": \"29386296,23864706,22342701,20876281,17882259,17113393\", \"BioGrid_Type\": \"Affinity Capture-MS,Affinity Capture-Western\", \"String_Score_Physical\": 997, \"String_Score_Combined\": 999}", "selected": false}, "selected": false}, {"data": {"id": "2783664", "source": "2782740", "target": "2782743", "SCORE": "216", "shared_interaction": "Direct", "source_original": "852235", "Name_B": "ATP3", "Name_A": "COR1", "target_original": "852327", "Gene_B": "852327", "interaction": "Direct", "SUID": 2783664, "TYPE": "Direct", "Gene_A": "852235", "support": "{\"StringDB\": \"physical\", \"StringEvidence\": \"coexpression:0.989,textmining:0.554,experiments:0.182\", \"String_Score_Physical\": 216, \"String_Score_Combined\": 996}", "selected": false}, "selected": false}, {"data": {"id": "2783661", "source": "2782737", "target": "2782797", "SCORE": "244", "shared_interaction": "Direct", "source_original": "852177", "Name_B": "RIP1", "Name_A": "ATP1", "target_original": "856689", "Gene_B": "856689", "interaction": "Direct", "SUID": 2783661, "TYPE": "Direct", "Gene_A": "852177", "support": "{\"StringDB\": \"physical\", \"StringEvidence\": \"coexpression:0.943,textmining:0.789,experiments:0.213\", \"String_Score_Physical\": 244, \"String_Score_Combined\": 990}", "selected": false}, "selected": false}, {"data": {"id": "2783658", "source": "2782737", "target": "2782794", "SCORE": "190", "shared_interaction": "Direct", "source_original": "852177", "Name_B": "CYC7", "Name_A": "ATP1", "target_original": "856672", "Gene_B": "856672", "interaction": "Direct", "SUID": 2783658, "TYPE": "Direct", "Gene_A": "852177", "support": "{\"StringDB\": \"physical\", \"StringEvidence\": \"textmining:0.511,experiments:0.155,coexpression:0.076\", \"String_Score_Physical\": 190, \"String_Score_Combined\": 633}", "selected": false}, "selected": false}, {"data": {"id": "2783655", "source": "2782737", "target": "2782788", "SCORE": "494", "shared_interaction": "Direct", "source_original": "852177", "Name_B": "QCR2", "Name_A": "ATP1", "target_original": "856321", "Gene_B": "856321", "interaction": "Direct", "SUID": 2783655, "TYPE": "Direct", "Gene_A": "852177", "support": "{\"StringDB\": \"physical\", \"StringEvidence\": \"coexpression:0.949,textmining:0.899,experiments:0.473\", \"String_Score_Physical\": 494, \"String_Score_Combined\": 997}", "selected": false}, "selected": false}, {"data": {"id": "2783652", "source": "2782737", "target": "2782785", "SCORE": "999", "shared_interaction": "Direct", "source_original": "852177", "Name_B": "ATP20", "Name_A": "ATP1", "target_original": "856131", "Gene_B": "856131", "interaction": "Direct", "SUID": 2783652, "TYPE": "Direct", "Gene_A": "852177", "support": "{\"StringDB\": \"physical\", \"StringEvidence\": \"experiments:0.998,textmining:0.787,database:0.708,coexpression:0.259\", \"String_Score_Physical\": 999, \"String_Score_Combined\": 999}", "selected": false}, "selected": false}, {"data": {"id": "2783649", "source": "2782737", "target": "2782782", "SCORE": "999", "shared_interaction": "Direct", "source_original": "852177", "Name_B": "ATP4", "Name_A": "ATP1", "target_original": "856027", "Gene_B": "856027", "interaction": "Direct", "SUID": 2783649, "TYPE": "Direct", "Gene_A": "852177", "support": "{\"StringDB\": \"physical\", \"StringEvidence\": \"experiments:0.999,coexpression:0.940,textmining:0.917,database:0.896\", \"BioGrid\": \"physical\", \"BioGrid_PubMed\": \"9867807,22940862\", \"BioGrid_Type\": \"Affinity Capture-MS,Co-purification\", \"String_Score_Physical\": 999, \"String_Score_Combined\": 999}", "selected": false}, "selected": false}, {"data": {"id": "2783646", "source": "2782737", "target": "2782779", "SCORE": "369", "shared_interaction": "Direct", "source_original": "852177", "Name_B": "COX5A", "Name_A": "ATP1", "target_original": "855675", "Gene_B": "855675", "interaction": "Direct", "SUID": 2783646, "TYPE": "Direct", "Gene_A": "852177", "support": "{\"StringDB\": \"physical\", \"StringEvidence\": \"textmining:0.658,coexpression:0.344,experiments:0.342\", \"String_Score_Physical\": 369, \"String_Score_Combined\": 858}", "selected": false}, "selected": false}, {"data": {"id": "2783643", "source": "2782737", "target": "2782776", "SCORE": "305", "shared_interaction": "Direct", "source_original": "852177", "Name_B": "COX7", "Name_A": "ATP1", "target_original": "855298", "Gene_B": "855298", "interaction": "Direct", "SUID": 2783643, "TYPE": "Direct", "Gene_A": "852177", "support": "{\"StringDB\": \"physical\", \"StringEvidence\": \"textmining:0.429,experiments:0.275\", \"String_Score_Physical\": 305, \"String_Score_Combined\": 602}", "selected": false}, "selected": false}, {"data": {"id": "2783640", "source": "2782737", "target": "2782770", "SCORE": "999", "shared_interaction": "Direct", "source_original": "852177", "Name_B": "ATP18", "Name_A": "ATP1", "target_original": "854893", "Gene_B": "854893", "interaction": "Direct", "SUID": 2783640, "TYPE": "Direct", "Gene_A": "852177", "support": "{\"StringDB\": \"physical\", \"StringEvidence\": \"experiments:0.999,database:0.896,textmining:0.638,coexpression:0.059\", \"BioGrid\": \"genetic,physical\", \"BioGrid_PubMed\": \"22940862,21623372\", \"BioGrid_Type\": \"Affinity Capture-MS,Positive Genetic\", \"BioGrid_Score\": \"0.1814\", \"String_Score_Physical\": 999, \"String_Score_Combined\": 999}", "selected": false}, "selected": false}, {"data": {"id": "2783637", "source": "2782737", "target": "2782767", "SCORE": "405", "shared_interaction": "Direct", "source_original": "852177", "Name_B": "COX5B", "Name_A": "ATP1", "target_original": "854695", "Gene_B": "854695", "interaction": "Direct", "SUID": 2783637, "TYPE": "Direct", "Gene_A": "852177", "support": "{\"StringDB\": \"physical\", \"StringEvidence\": \"experiments:0.380,textmining:0.260,coexpression:0.145\", \"BioGrid\": \"genetic\", \"BioGrid_PubMed\": \"21623372\", \"BioGrid_Type\": \"Negative Genetic\", \"BioGrid_Score\": \"-0.0864\", \"String_Score_Physical\": 405, \"String_Score_Combined\": 623}", "selected": false}, "selected": false}, {"data": {"id": "2783634", "source": "2782737", "target": "2782764", "SCORE": "944", "shared_interaction": "Direct", "source_original": "852177", "Name_B": "CYT1", "Name_A": "ATP1", "target_original": "854231", "Gene_B": "854231", "interaction": "Direct", "SUID": 2783634, "TYPE": "Direct", "Gene_A": "852177", "support": "{\"StringDB\": \"physical\", \"StringEvidence\": \"coexpression:0.948,database:0.896,textmining:0.535,experiments:0.442\", \"String_Score_Physical\": 944, \"String_Score_Combined\": 998}", "selected": false}, "selected": false}, {"data": {"id": "2783631", "source": "2782737", "target": "2782761", "SCORE": "941", "shared_interaction": "Direct", "source_original": "852177", "Name_B": "ATP19", "Name_A": "ATP1", "target_original": "854077", "Gene_B": "854077", "interaction": "Direct", "SUID": 2783631, "TYPE": "Direct", "Gene_A": "852177", "support": "{\"StringDB\": \"physical\", \"StringEvidence\": \"experiments:0.791,database:0.708,coexpression:0.060\", \"String_Score_Physical\": 941, \"String_Score_Combined\": 945}", "selected": false}, "selected": false}, {"data": {"id": "2783628", "source": "2782737", "target": "2782758", "SCORE": "999", "shared_interaction": "Direct", "source_original": "852177", "Name_B": "ATP7", "Name_A": "ATP1", "target_original": "853853", "Gene_B": "853853", "interaction": "Direct", "SUID": 2783628, "TYPE": "Direct", "Gene_A": "852177", "support": "{\"StringDB\": \"physical\", \"StringEvidence\": \"experiments:0.999,coexpression:0.914,database:0.896,textmining:0.885\", \"BioGrid\": \"genetic,physical\", \"BioGrid_PubMed\": \"22940862,21623372\", \"BioGrid_Type\": \"Affinity Capture-MS,Positive Genetic\", \"BioGrid_Score\": \"0.098\", \"String_Score_Physical\": 999, \"String_Score_Combined\": 999}", "selected": false}, "selected": false}, {"data": {"id": "2783625", "source": "2782737", "target": "2782755", "SCORE": "319", "shared_interaction": "Direct", "source_original": "852177", "Name_B": "SDH3", "Name_A": "ATP1", "target_original": "853716", "Gene_B": "853716", "interaction": "Direct", "SUID": 2783625, "TYPE": "Direct", "Gene_A": "852177", "support": "{\"StringDB\": \"physical\", \"StringEvidence\": \"textmining:0.458,coexpression:0.436,experiments:0.290,neighborhood:0.111\", \"BioGrid\": \"genetic\", \"BioGrid_PubMed\": \"21623372,16941010\", \"BioGrid_Type\": \"Synthetic Lethality,Positive Genetic\", \"BioGrid_Score\": \"0.1059\", \"String_Score_Physical\": 319, \"String_Score_Combined\": 814}", "selected": false}, "selected": false}, {"data": {"id": "2783622", "source": "2782737", "target": "2782752", "SCORE": "250", "shared_interaction": "Direct", "source_original": "852177", "Name_B": "QCR8", "Name_A": "ATP1", "target_original": "853273", "Gene_B": "853273", "interaction": "Direct", "SUID": 2783622, "TYPE": "Direct", "Gene_A": "852177", "support": "{\"StringDB\": \"physical\", \"StringEvidence\": \"coexpression:0.537,textmining:0.228,experiments:0.218\", \"BioGrid\": \"genetic\", \"BioGrid_PubMed\": \"21623372\", \"BioGrid_Type\": \"Negative Genetic\", \"BioGrid_Score\": \"-0.2114\", \"String_Score_Physical\": 250, \"String_Score_Combined\": 732}", "selected": false}, "selected": false}, {"data": {"id": "2783619", "source": "2782737", "target": "2782749", "SCORE": "308", "shared_interaction": "Direct", "source_original": "852177", "Name_B": "QCR9", "Name_A": "ATP1", "target_original": "853095", "Gene_B": "853095", "interaction": "Direct", "SUID": 2783619, "TYPE": "Direct", "Gene_A": "852177", "support": "{\"StringDB\": \"physical\", \"StringEvidence\": \"textmining:0.326,experiments:0.279,coexpression:0.134\", \"BioGrid\": \"genetic\", \"BioGrid_PubMed\": \"21623372\", \"BioGrid_Type\": \"Negative Genetic\", \"BioGrid_Score\": \"-0.1451\", \"String_Score_Physical\": 308, \"String_Score_Combined\": 596}", "selected": false}, "selected": false}, {"data": {"id": "2783616", "source": "2782737", "target": "2782746", "SCORE": "223", "shared_interaction": "Direct", "source_original": "852177", "Name_B": "COX4", "Name_A": "ATP1", "target_original": "852688", "Gene_B": "852688", "interaction": "Direct", "SUID": 2783616, "TYPE": "Direct", "Gene_A": "852177", "support": "{\"StringDB\": \"physical\", \"StringEvidence\": \"textmining:0.896,coexpression:0.716,experiments:0.190\", \"String_Score_Physical\": 223, \"String_Score_Combined\": 976}", "selected": false}, "selected": false}, {"data": {"id": "2783613", "source": "2782737", "target": "2782743", "SCORE": "999", "shared_interaction": "Direct", "source_original": "852177", "Name_B": "ATP3", "Name_A": "ATP1", "target_original": "852327", "Gene_B": "852327", "interaction": "Direct", "SUID": 2783613, "TYPE": "Direct", "Gene_A": "852177", "support": "{\"StringDB\": \"physical\", \"StringEvidence\": \"experiments:0.999,textmining:0.991,coexpression:0.967,database:0.896,cooccurrence:0.412,neighborhood:0.156\", \"BioGrid\": \"physical\", \"BioGrid_PubMed\": \"22940862\", \"BioGrid_Type\": \"Affinity Capture-MS\", \"String_Score_Physical\": 999, \"String_Score_Combined\": 999}", "selected": false}, "selected": false}, {"data": {"id": "2783610", "source": "2782737", "target": "2782740", "SCORE": "420", "shared_interaction": "Direct", "source_original": "852177", "Name_B": "COR1", "Name_A": "ATP1", "target_original": "852235", "Gene_B": "852235", "interaction": "Direct", "SUID": 2783610, "TYPE": "Direct", "Gene_A": "852177", "support": "{\"StringDB\": \"physical\", \"StringEvidence\": \"coexpression:0.988,textmining:0.882,experiments:0.395\", \"String_Score_Physical\": 420, \"String_Score_Combined\": 999}", "selected": false}, "selected": false}, {"data": {"id": "2783607", "source": "2782734", "target": "2782797", "SCORE": "999", "shared_interaction": "Direct", "source_original": "852142", "Name_B": "RIP1", "Name_A": "QCR7", "target_original": "856689", "Gene_B": "856689", "interaction": "Direct", "SUID": 2783607, "TYPE": "Direct", "Gene_A": "852142", "support": "{\"StringDB\": \"physical\", \"StringEvidence\": \"experiments:0.995,coexpression:0.982,textmining:0.946,database:0.896\", \"String_Score_Physical\": 999, \"String_Score_Combined\": 999}", "selected": false}, "selected": false}, {"data": {"id": "2783604", "source": "2782734", "target": "2782794", "SCORE": "979", "shared_interaction": "Direct", "source_original": "852142", "Name_B": "CYC7", "Name_A": "QCR7", "target_original": "856672", "Gene_B": "856672", "interaction": "Direct", "SUID": 2783604, "TYPE": "Direct", "Gene_A": "852142", "support": "{\"StringDB\": \"physical\", \"StringEvidence\": \"database:0.896,experiments:0.791,coexpression:0.629,textmining:0.495\", \"String_Score_Physical\": 979, \"String_Score_Combined\": 996}", "selected": false}, "selected": false}, {"data": {"id": "2783601", "source": "2782734", "target": "2782791", "SCORE": "979", "shared_interaction": "Direct", "source_original": "852142", "Name_B": "QCR10", "Name_A": "QCR7", "target_original": "856390", "Gene_B": "856390", "interaction": "Direct", "SUID": 2783601, "TYPE": "Direct", "Gene_A": "852142", "support": "{\"StringDB\": \"physical\", \"StringEvidence\": \"textmining:0.897,database:0.896,coexpression:0.854,experiments:0.794\", \"BioGrid\": \"genetic\", \"BioGrid_PubMed\": \"21623372\", \"BioGrid_Type\": \"Negative Genetic\", \"BioGrid_Score\": \"-0.1054\", \"String_Score_Physical\": 979, \"String_Score_Combined\": 999}", "selected": false}, "selected": false}, {"data": {"id": "2783598", "source": "2782734", "target": "2782788", "SCORE": "999", "shared_interaction": "Direct", "source_original": "852142", "Name_B": "QCR2", "Name_A": "QCR7", "target_original": "856321", "Gene_B": "856321", "interaction": "Direct", "SUID": 2783598, "TYPE": "Direct", "Gene_A": "852142", "support": "{\"StringDB\": \"physical\", \"StringEvidence\": \"experiments:0.999,textmining:0.980,coexpression:0.964,database:0.896\", \"BioGrid\": \"physical\", \"BioGrid_PubMed\": \"32119834,25683140,21807901\", \"BioGrid_Type\": \"Affinity Capture-Western\", \"String_Score_Physical\": 999, \"String_Score_Combined\": 999}", "selected": false}, "selected": false}, {"data": {"id": "2783595", "source": "2782734", "target": "2782782", "SCORE": "963", "shared_interaction": "Direct", "source_original": "852142", "Name_B": "ATP4", "Name_A": "QCR7", "target_original": "856027", "Gene_B": "856027", "interaction": "Direct", "SUID": 2783595, "TYPE": "Direct", "Gene_A": "852142", "support": "{\"StringDB\": \"physical\", \"StringEvidence\": \"experiments:0.961,coexpression:0.940,textmining:0.755\", \"String_Score_Physical\": 963, \"String_Score_Combined\": 999}", "selected": false}, "selected": false}, {"data": {"id": "2783592", "source": "2782734", "target": "2782779", "SCORE": "984", "shared_interaction": "Direct", "source_original": "852142", "Name_B": "COX5A", "Name_A": "QCR7", "target_original": "855675", "Gene_B": "855675", "interaction": "Direct", "SUID": 2783592, "TYPE": "Direct", "Gene_A": "852142", "support": "{\"StringDB\": \"physical\", \"StringEvidence\": \"database:0.896,experiments:0.844,coexpression:0.786,textmining:0.729\", \"String_Score_Physical\": 984, \"String_Score_Combined\": 999}", "selected": false}, "selected": false}, {"data": {"id": "2783589", "source": "2782734", "target": "2782776", "SCORE": "979", "shared_interaction": "Direct", "source_original": "852142", "Name_B": "COX7", "Name_A": "QCR7", "target_original": "855298", "Gene_B": "855298", "interaction": "Direct", "SUID": 2783589, "TYPE": "Direct", "Gene_A": "852142", "support": "{\"StringDB\": \"physical\", \"StringEvidence\": \"database:0.896,coexpression:0.872,experiments:0.791,textmining:0.550\", \"String_Score_Physical\": 979, \"String_Score_Combined\": 998}", "selected": false}, "selected": false}, {"data": {"id": "2783586", "source": "2782734", "target": "2782773", "SCORE": "900", "shared_interaction": "Direct", "source_original": "852142", "Name_B": "NDI1", "Name_A": "QCR7", "target_original": "854919", "Gene_B": "854919", "interaction": "Direct", "SUID": 2783586, "TYPE": "Direct", "Gene_A": "852142", "support": "{\"StringDB\": \"physical\", \"StringEvidence\": \"database:0.896,textmining:0.555,coexpression:0.318\", \"String_Score_Physical\": 900, \"String_Score_Combined\": 969}", "selected": false}, "selected": false}, {"data": {"id": "2783583", "source": "2782734", "target": "2782767", "SCORE": "984", "shared_interaction": "Direct", "source_original": "852142", "Name_B": "COX5B", "Name_A": "QCR7", "target_original": "854695", "Gene_B": "854695", "interaction": "Direct", "SUID": 2783583, "TYPE": "Direct", "Gene_A": "852142", "support": "{\"StringDB\": \"physical\", \"StringEvidence\": \"database:0.896,experiments:0.844,textmining:0.315,coexpression:0.231\", \"String_Score_Physical\": 984, \"String_Score_Combined\": 991}", "selected": false}, "selected": false}, {"data": {"id": "2783580", "source": "2782734", "target": "2782764", "SCORE": "999", "shared_interaction": "Direct", "source_original": "852142", "Name_B": "CYT1", "Name_A": "QCR7", "target_original": "854231", "Gene_B": "854231", "interaction": "Direct", "SUID": 2783580, "TYPE": "Direct", "Gene_A": "852142", "support": "{\"StringDB\": \"physical\", \"StringEvidence\": \"experiments:0.997,textmining:0.993,coexpression:0.978,database:0.896\", \"BioGrid\": \"physical\", \"BioGrid_PubMed\": \"32119834\", \"BioGrid_Type\": \"Affinity Capture-Western\", \"String_Score_Physical\": 999, \"String_Score_Combined\": 999}", "selected": false}, "selected": false}, {"data": {"id": "2783577", "source": "2782734", "target": "2782755", "SCORE": "900", "shared_interaction": "Direct", "source_original": "852142", "Name_B": "SDH3", "Name_A": "QCR7", "target_original": "853716", "Gene_B": "853716", "interaction": "Direct", "SUID": 2783577, "TYPE": "Direct", "Gene_A": "852142", "support": "{\"StringDB\": \"physical\", \"StringEvidence\": \"database:0.896,coexpression:0.378,textmining:0.209\", \"String_Score_Physical\": 900, \"String_Score_Combined\": 950}", "selected": false}, "selected": false}, {"data": {"id": "2783574", "source": "2782734", "target": "2782752", "SCORE": "999", "shared_interaction": "Direct", "source_original": "852142", "Name_B": "QCR8", "Name_A": "QCR7", "target_original": "853273", "Gene_B": "853273", "interaction": "Direct", "SUID": 2783574, "TYPE": "Direct", "Gene_A": "852142", "support": "{\"StringDB\": \"physical\", \"StringEvidence\": \"textmining:0.996,experiments:0.995,coexpression:0.984,database:0.896\", \"String_Score_Physical\": 999, \"String_Score_Combined\": 999}", "selected": false}, "selected": false}, {"data": {"id": "2783571", "source": "2782734", "target": "2782749", "SCORE": "999", "shared_interaction": "Direct", "source_original": "852142", "Name_B": "QCR9", "Name_A": "QCR7", "target_original": "853095", "Gene_B": "853095", "interaction": "Direct", "SUID": 2783571, "TYPE": "Direct", "Gene_A": "852142", "support": "{\"StringDB\": \"physical\", \"StringEvidence\": \"experiments:0.995,coexpression:0.988,textmining:0.981,database:0.896\", \"String_Score_Physical\": 999, \"String_Score_Combined\": 999}", "selected": false}, "selected": false}, {"data": {"id": "2783568", "source": "2782734", "target": "2782746", "SCORE": "999", "shared_interaction": "Direct", "source_original": "852142", "Name_B": "COX4", "Name_A": "QCR7", "target_original": "852688", "Gene_B": "852688", "interaction": "Direct", "SUID": 2783568, "TYPE": "Direct", "Gene_A": "852142", "support": "{\"StringDB\": \"physical\", \"StringEvidence\": \"experiments:0.995,coexpression:0.991,database:0.896,textmining:0.781\", \"BioGrid\": \"physical\", \"BioGrid_PubMed\": \"23864706\", \"BioGrid_Type\": \"Affinity Capture-MS\", \"String_Score_Physical\": 999, \"String_Score_Combined\": 999}", "selected": false}, "selected": false}, {"data": {"id": "2783565", "source": "2782734", "target": "2782740", "SCORE": "999", "shared_interaction": "Direct", "source_original": "852142", "Name_B": "COR1", "Name_A": "QCR7", "target_original": "852235", "Gene_B": "852235", "interaction": "Direct", "SUID": 2783565, "TYPE": "Direct", "Gene_A": "852142", "support": "{\"StringDB\": \"physical\", \"StringEvidence\": \"experiments:0.998,textmining:0.915,database:0.896,coexpression:0.760\", \"BioGrid\": \"physical\", \"BioGrid_PubMed\": \"32119834,22342701\", \"BioGrid_Type\": \"Affinity Capture-MS,Affinity Capture-Western\", \"String_Score_Physical\": 999, \"String_Score_Combined\": 999}", "selected": false}, "selected": false}, {"data": {"id": "2783562", "source": "2782731", "target": "2782785", "SCORE": "720", "shared_interaction": "Direct", "source_original": "851983", "Name_B": "ATP20", "Name_A": "ATP17", "target_original": "856131", "Gene_B": "856131", "interaction": "Direct", "SUID": 2783562, "TYPE": "Direct", "Gene_A": "851983", "support": "{\"StringDB\": \"physical\", \"StringEvidence\": \"coexpression:0.787,database:0.708,textmining:0.426\", \"String_Score_Physical\": 720, \"String_Score_Combined\": 965}", "selected": false}, "selected": false}, {"data": {"id": "2783559", "source": "2782731", "target": "2782782", "SCORE": "999", "shared_interaction": "Direct", "source_original": "851983", "Name_B": "ATP4", "Name_A": "ATP17", "target_original": "856027", "Gene_B": "856027", "interaction": "Direct", "SUID": 2783559, "TYPE": "Direct", "Gene_A": "851983", "support": "{\"StringDB\": \"physical\", \"StringEvidence\": \"experiments:0.999,coexpression:0.806,textmining:0.788,database:0.708\", \"BioGrid\": \"physical\", \"BioGrid_PubMed\": \"9893937,9867807,9425084\", \"BioGrid_Type\": \"Reconstituted Complex,Co-purification\", \"String_Score_Physical\": 999, \"String_Score_Combined\": 999}", "selected": false}, "selected": false}, {"data": {"id": "2783556", "source": "2782731", "target": "2782770", "SCORE": "985", "shared_interaction": "Direct", "source_original": "851983", "Name_B": "ATP18", "Name_A": "ATP17", "target_original": "854893", "Gene_B": "854893", "interaction": "Direct", "SUID": 2783556, "TYPE": "Direct", "Gene_A": "851983", "support": "{\"StringDB\": \"physical\", \"StringEvidence\": \"experiments:0.950,database:0.708,textmining:0.575,coexpression:0.323\", \"BioGrid\": \"genetic,physical\", \"BioGrid_PubMed\": \"21623372,10747812\", \"BioGrid_Type\": \"Reconstituted Complex,Positive Genetic\", \"BioGrid_Score\": \"0.1181\", \"String_Score_Physical\": 985, \"String_Score_Combined\": 996}", "selected": false}, "selected": false}, {"data": {"id": "2783553", "source": "2782731", "target": "2782761", "SCORE": "941", "shared_interaction": "Direct", "source_original": "851983", "Name_B": "ATP19", "Name_A": "ATP17", "target_original": "854077", "Gene_B": "854077", "interaction": "Direct", "SUID": 2783553, "TYPE": "Direct", "Gene_A": "851983", "support": "{\"StringDB\": \"physical\", \"StringEvidence\": \"experiments:0.791,coexpression:0.758,database:0.708,textmining:0.430\", \"String_Score_Physical\": 941, \"String_Score_Combined\": 991}", "selected": false}, "selected": false}, {"data": {"id": "2783550", "source": "2782731", "target": "2782758", "SCORE": "963", "shared_interaction": "Direct", "source_original": "851983", "Name_B": "ATP7", "Name_A": "ATP17", "target_original": "853853", "Gene_B": "853853", "interaction": "Direct", "SUID": 2783550, "TYPE": "Direct", "Gene_A": "851983", "support": "{\"StringDB\": \"physical\", \"StringEvidence\": \"textmining:0.934,experiments:0.870,coexpression:0.814,database:0.708\", \"BioGrid\": \"genetic,physical\", \"BioGrid_PubMed\": \"21623372,10747812\", \"BioGrid_Type\": \"Reconstituted Complex,Positive Genetic\", \"BioGrid_Score\": \"0.1036\", \"String_Score_Physical\": 963, \"String_Score_Combined\": 999}", "selected": false}, "selected": false}, {"data": {"id": "2783547", "source": "2782731", "target": "2782746", "SCORE": "288", "shared_interaction": "Direct", "source_original": "851983", "Name_B": "COX4", "Name_A": "ATP17", "target_original": "852688", "Gene_B": "852688", "interaction": "Direct", "SUID": 2783547, "TYPE": "Direct", "Gene_A": "851983", "support": "{\"StringDB\": \"physical\", \"StringEvidence\": \"coexpression:0.922,textmining:0.460,experiments:0.258\", \"BioGrid\": \"physical\", \"BioGrid_PubMed\": \"23864706\", \"BioGrid_Type\": \"Affinity Capture-MS\", \"String_Score_Physical\": 288, \"String_Score_Combined\": 969}", "selected": false}, "selected": false}, {"data": {"id": "2783544", "source": "2782731", "target": "2782743", "SCORE": "941", "shared_interaction": "Direct", "source_original": "851983", "Name_B": "ATP3", "Name_A": "ATP17", "target_original": "852327", "Gene_B": "852327", "interaction": "Direct", "SUID": 2783544, "TYPE": "Direct", "Gene_A": "851983", "support": "{\"StringDB\": \"physical\", \"StringEvidence\": \"experiments:0.791,database:0.708,textmining:0.684,coexpression:0.284\", \"String_Score_Physical\": 941, \"String_Score_Combined\": 986}", "selected": false}, "selected": false}, {"data": {"id": "2783541", "source": "2782731", "target": "2782740", "SCORE": "432", "shared_interaction": "Direct", "source_original": "851983", "Name_B": "COR1", "Name_A": "ATP17", "target_original": "852235", "Gene_B": "852235", "interaction": "Direct", "SUID": 2783541, "TYPE": "Direct", "Gene_A": "851983", "support": "{\"StringDB\": \"physical\", \"StringEvidence\": \"experiments:0.408,coexpression:0.251\", \"BioGrid\": \"physical\", \"BioGrid_PubMed\": \"22342701\", \"BioGrid_Type\": \"Affinity Capture-MS\", \"String_Score_Physical\": 432, \"String_Score_Combined\": 574}", "selected": false}, "selected": false}, {"data": {"id": "2783538", "source": "2782731", "target": "2782737", "SCORE": "994", "shared_interaction": "Direct", "source_original": "851983", "Name_B": "ATP1", "Name_A": "ATP17", "target_original": "852177", "Gene_B": "852177", "interaction": "Direct", "SUID": 2783538, "TYPE": "Direct", "Gene_A": "851983", "support": "{\"StringDB\": \"physical\", \"StringEvidence\": \"experiments:0.981,database:0.708,textmining:0.565,coexpression:0.168\", \"String_Score_Physical\": 994, \"String_Score_Combined\": 998}", "selected": false}, "selected": false}, {"data": {"id": "2783535", "source": "2782728", "target": "2782785", "SCORE": "865", "shared_interaction": "Direct", "source_original": "851922", "Name_B": "ATP20", "Name_A": "TIM11", "target_original": "856131", "Gene_B": "856131", "interaction": "Direct", "SUID": 2783535, "TYPE": "Direct", "Gene_A": "851922", "support": "{\"StringDB\": \"physical\", \"StringEvidence\": \"textmining:0.974,coexpression:0.879,database:0.708,experiments:0.520\", \"BioGrid\": \"genetic,physical\", \"BioGrid_PubMed\": \"21623372,19635484,15886192\", \"BioGrid_Type\": \"Positive Genetic,Affinity Capture-Western\", \"BioGrid_Score\": \"0.1279\", \"String_Score_Physical\": 865, \"String_Score_Combined\": 999}", "selected": false}, "selected": false}, {"data": {"id": "2783532", "source": "2782728", "target": "2782782", "SCORE": "976", "shared_interaction": "Direct", "source_original": "851922", "Name_B": "ATP4", "Name_A": "TIM11", "target_original": "856027", "Gene_B": "856027", "interaction": "Direct", "SUID": 2783532, "TYPE": "Direct", "Gene_A": "851922", "support": "{\"StringDB\": \"physical\", \"StringEvidence\": \"experiments:0.914,database:0.708,textmining:0.635,coexpression:0.407\", \"BioGrid\": \"physical\", \"BioGrid_PubMed\": \"9867807\", \"BioGrid_Type\": \"Co-purification\", \"String_Score_Physical\": 976, \"String_Score_Combined\": 994}", "selected": false}, "selected": false}, {"data": {"id": "2783529", "source": "2782728", "target": "2782770", "SCORE": "800", "shared_interaction": "Direct", "source_original": "851922", "Name_B": "ATP18", "Name_A": "TIM11", "target_original": "854893", "Gene_B": "854893", "interaction": "Direct", "SUID": 2783529, "TYPE": "Direct", "Gene_A": "851922", "support": "{\"StringDB\": \"physical\", \"StringEvidence\": \"textmining:0.736,database:0.708,coexpression:0.449,experiments:0.286\", \"BioGrid\": \"physical\", \"BioGrid_PubMed\": \"12206911\", \"BioGrid_Type\": \"Affinity Capture-Western\", \"String_Score_Physical\": 800, \"String_Score_Combined\": 970}", "selected": false}, "selected": false}, {"data": {"id": "2783526", "source": "2782728", "target": "2782761", "SCORE": "720", "shared_interaction": "Direct", "source_original": "851922", "Name_B": "ATP19", "Name_A": "TIM11", "target_original": "854077", "Gene_B": "854077", "interaction": "Direct", "SUID": 2783526, "TYPE": "Direct", "Gene_A": "851922", "support": "{\"StringDB\": \"physical\", \"StringEvidence\": \"textmining:0.738,database:0.708,coexpression:0.557\", \"String_Score_Physical\": 720, \"String_Score_Combined\": 967}", "selected": false}, "selected": false}, {"data": {"id": "2783523", "source": "2782728", "target": "2782758", "SCORE": "720", "shared_interaction": "Direct", "source_original": "851922", "Name_B": "ATP7", "Name_A": "TIM11", "target_original": "853853", "Gene_B": "853853", "interaction": "Direct", "SUID": 2783523, "TYPE": "Direct", "Gene_A": "851922", "support": "{\"StringDB\": \"physical\", \"StringEvidence\": \"database:0.708,coexpression:0.340,textmining:0.249\", \"String_Score_Physical\": 720, \"String_Score_Combined\": 861}", "selected": false}, "selected": false}, {"data": {"id": "2783520", "source": "2782728", "target": "2782743", "SCORE": "720", "shared_interaction": "Direct", "source_original": "851922", "Name_B": "ATP3", "Name_A": "TIM11", "target_original": "852327", "Gene_B": "852327", "interaction": "Direct", "SUID": 2783520, "TYPE": "Direct", "Gene_A": "851922", "support": "{\"StringDB\": \"physical\", \"StringEvidence\": \"database:0.708,textmining:0.426,coexpression:0.244\", \"String_Score_Physical\": 720, \"String_Score_Combined\": 878}", "selected": false}, "selected": false}, {"data": {"id": "2783517", "source": "2782728", "target": "2782737", "SCORE": "805", "shared_interaction": "Direct", "source_original": "851922", "Name_B": "ATP1", "Name_A": "TIM11", "target_original": "852177", "Gene_B": "852177", "interaction": "Direct", "SUID": 2783517, "TYPE": "Direct", "Gene_A": "851922", "support": "{\"StringDB\": \"physical\", \"StringEvidence\": \"database:0.708,experiments:0.304,textmining:0.180,coexpression:0.086\", \"BioGrid\": \"physical\", \"BioGrid_PubMed\": \"9271204\", \"BioGrid_Type\": \"Affinity Capture-Western\", \"String_Score_Physical\": 805, \"String_Score_Combined\": 854}", "selected": false}, "selected": false}, {"data": {"id": "2783514", "source": "2782728", "target": "2782731", "SCORE": "720", "shared_interaction": "Direct", "source_original": "851922", "Name_B": "ATP17", "Name_A": "TIM11", "target_original": "851983", "Gene_B": "851983", "interaction": "Direct", "SUID": 2783514, "TYPE": "Direct", "Gene_A": "851922", "support": "{\"StringDB\": \"physical\", \"StringEvidence\": \"coexpression:0.716,database:0.708,textmining:0.594\", \"String_Score_Physical\": 720, \"String_Score_Combined\": 967}", "selected": false}, "selected": false}, {"data": {"id": "2783490", "source": "2782725", "target": "2782797", "SCORE": "900", "shared_interaction": "Direct", "source_original": "851758", "Name_B": "RIP1", "Name_A": "SDH4", "target_original": "856689", "Gene_B": "856689", "interaction": "Direct", "SUID": 2783490, "TYPE": "Direct", "Gene_A": "851758", "support": "{\"StringDB\": \"physical\", \"StringEvidence\": \"coexpression:0.937,database:0.896,textmining:0.351\", \"String_Score_Physical\": 900, \"String_Score_Combined\": 995}", "selected": false}, "selected": false}, {"data": {"id": "2783487", "source": "2782725", "target": "2782791", "SCORE": "900", "shared_interaction": "Direct", "source_original": "851758", "Name_B": "QCR10", "Name_A": "SDH4", "target_original": "856390", "Gene_B": "856390", "interaction": "Direct", "SUID": 2783487, "TYPE": "Direct", "Gene_A": "851758", "support": "{\"StringDB\": \"physical\", \"StringEvidence\": \"database:0.896,coexpression:0.528,textmining:0.039\", \"String_Score_Physical\": 900, \"String_Score_Combined\": 954}", "selected": false}, "selected": false}, {"data": {"id": "2783484", "source": "2782725", "target": "2782788", "SCORE": "914", "shared_interaction": "Direct", "source_original": "851758", "Name_B": "QCR2", "Name_A": "SDH4", "target_original": "856321", "Gene_B": "856321", "interaction": "Direct", "SUID": 2783484, "TYPE": "Direct", "Gene_A": "851758", "support": "{\"StringDB\": \"physical\", \"StringEvidence\": \"database:0.896,coexpression:0.418,textmining:0.263,experiments:0.144\", \"BioGrid\": \"genetic\", \"BioGrid_PubMed\": \"27453043\", \"BioGrid_Type\": \"Positive Genetic\", \"BioGrid_Score\": \"2.4\", \"String_Score_Physical\": 914, \"String_Score_Combined\": 963}", "selected": false}, "selected": false}, {"data": {"id": "2783481", "source": "2782725", "target": "2782773", "SCORE": "900", "shared_interaction": "Direct", "source_original": "851758", "Name_B": "NDI1", "Name_A": "SDH4", "target_original": "854919", "Gene_B": "854919", "interaction": "Direct", "SUID": 2783481, "TYPE": "Direct", "Gene_A": "851758", "support": "{\"StringDB\": \"physical\", \"StringEvidence\": \"database:0.896,coexpression:0.867,textmining:0.063\", \"String_Score_Physical\": 900, \"String_Score_Combined\": 987}", "selected": false}, "selected": false}, {"data": {"id": "2783478", "source": "2782725", "target": "2782764", "SCORE": "940", "shared_interaction": "Direct", "source_original": "851758", "Name_B": "CYT1", "Name_A": "SDH4", "target_original": "854231", "Gene_B": "854231", "interaction": "Direct", "SUID": 2783478, "TYPE": "Direct", "Gene_A": "851758", "support": "{\"StringDB\": \"physical\", \"StringEvidence\": \"database:0.896,coexpression:0.639,experiments:0.407,textmining:0.322\", \"BioGrid\": \"physical\", \"BioGrid_PubMed\": \"14672930\", \"BioGrid_Type\": \"Co-localization\", \"String_Score_Physical\": 940, \"String_Score_Combined\": 985}", "selected": false}, "selected": false}, {"data": {"id": "2783475", "source": "2782725", "target": "2782755", "SCORE": "971", "shared_interaction": "Direct", "source_original": "851758", "Name_B": "SDH3", "Name_A": "SDH4", "target_original": "853716", "Gene_B": "853716", "interaction": "Direct", "SUID": 2783475, "TYPE": "Direct", "Gene_A": "851758", "support": "{\"StringDB\": \"physical\", \"StringEvidence\": \"textmining:0.983,coexpression:0.962,database:0.896,experiments:0.715\", \"BioGrid\": \"physical\", \"BioGrid_PubMed\": \"22152483,18313953\", \"BioGrid_Type\": \"Affinity Capture-Western,PCA\", \"String_Score_Physical\": 971, \"String_Score_Combined\": 999}", "selected": false}, "selected": false}, {"data": {"id": "2783472", "source": "2782725", "target": "2782752", "SCORE": "900", "shared_interaction": "Direct", "source_original": "851758", "Name_B": "QCR8", "Name_A": "SDH4", "target_original": "853273", "Gene_B": "853273", "interaction": "Direct", "SUID": 2783472, "TYPE": "Direct", "Gene_A": "851758", "support": "{\"StringDB\": \"physical\", \"StringEvidence\": \"database:0.896,textmining:0.509,coexpression:0.277\", \"String_Score_Physical\": 900, \"String_Score_Combined\": 964}", "selected": false}, "selected": false}, {"data": {"id": "2783469", "source": "2782725", "target": "2782749", "SCORE": "900", "shared_interaction": "Direct", "source_original": "851758", "Name_B": "QCR9", "Name_A": "SDH4", "target_original": "853095", "Gene_B": "853095", "interaction": "Direct", "SUID": 2783469, "TYPE": "Direct", "Gene_A": "851758", "support": "{\"StringDB\": \"physical\", \"StringEvidence\": \"database:0.896,coexpression:0.364,textmining:0.262\", \"String_Score_Physical\": 900, \"String_Score_Combined\": 953}", "selected": false}, "selected": false}, {"data": {"id": "2783466", "source": "2782725", "target": "2782740", "SCORE": "914", "shared_interaction": "Direct", "source_original": "851758", "Name_B": "COR1", "Name_A": "SDH4", "target_original": "852235", "Gene_B": "852235", "interaction": "Direct", "SUID": 2783466, "TYPE": "Direct", "Gene_A": "851758", "support": "{\"StringDB\": \"physical\", \"StringEvidence\": \"database:0.896,coexpression:0.604,textmining:0.592,experiments:0.149\", \"String_Score_Physical\": 914, \"String_Score_Combined\": 986}", "selected": false}, "selected": false}, {"data": {"id": "2783463", "source": "2782725", "target": "2782734", "SCORE": "900", "shared_interaction": "Direct", "source_original": "851758", "Name_B": "QCR7", "Name_A": "SDH4", "target_original": "852142", "Gene_B": "852142", "interaction": "Direct", "SUID": 2783463, "TYPE": "Direct", "Gene_A": "851758", "support": "{\"StringDB\": \"physical\", \"StringEvidence\": \"database:0.896,textmining:0.566,coexpression:0.441\", \"String_Score_Physical\": 900, \"String_Score_Combined\": 975}", "selected": false}, "selected": false}, {"data": {"id": "2783385", "source": "2782722", "target": "2782797", "SCORE": "997", "shared_interaction": "Direct", "source_original": "851111", "Name_B": "RIP1", "Name_A": "COX8", "target_original": "856689", "Gene_B": "856689", "interaction": "Direct", "SUID": 2783385, "TYPE": "Direct", "Gene_A": "851111", "support": "{\"StringDB\": \"physical\", \"StringEvidence\": \"experiments:0.980,database:0.896,coexpression:0.868,textmining:0.452\", \"BioGrid\": \"physical\", \"BioGrid_PubMed\": \"29688796\", \"BioGrid_Type\": \"Affinity Capture-Western\", \"String_Score_Physical\": 997, \"String_Score_Combined\": 999}", "selected": false}, "selected": false}, {"data": {"id": "2783382", "source": "2782722", "target": "2782794", "SCORE": "408", "shared_interaction": "Direct", "source_original": "851111", "Name_B": "CYC7", "Name_A": "COX8", "target_original": "856672", "Gene_B": "856672", "interaction": "Direct", "SUID": 2783382, "TYPE": "Direct", "Gene_A": "851111", "support": "{\"StringDB\": \"physical\", \"StringEvidence\": \"textmining:0.517,experiments:0.384,coexpression:0.057\", \"String_Score_Physical\": 408, \"String_Score_Combined\": 730}", "selected": false}, "selected": false}, {"data": {"id": "2783379", "source": "2782722", "target": "2782791", "SCORE": "979", "shared_interaction": "Direct", "source_original": "851111", "Name_B": "QCR10", "Name_A": "COX8", "target_original": "856390", "Gene_B": "856390", "interaction": "Direct", "SUID": 2783379, "TYPE": "Direct", "Gene_A": "851111", "support": "{\"StringDB\": \"physical\", \"StringEvidence\": \"coexpression:0.912,database:0.896,experiments:0.791\", \"String_Score_Physical\": 979, \"String_Score_Combined\": 998}", "selected": false}, "selected": false}, {"data": {"id": "2783376", "source": "2782722", "target": "2782788", "SCORE": "998", "shared_interaction": "Direct", "source_original": "851111", "Name_B": "QCR2", "Name_A": "COX8", "target_original": "856321", "Gene_B": "856321", "interaction": "Direct", "SUID": 2783376, "TYPE": "Direct", "Gene_A": "851111", "support": "{\"StringDB\": \"physical\", \"StringEvidence\": \"experiments:0.983,database:0.896,coexpression:0.486,textmining:0.311\", \"BioGrid\": \"physical\", \"BioGrid_PubMed\": \"29688796\", \"BioGrid_Type\": \"Affinity Capture-Western\", \"String_Score_Physical\": 998, \"String_Score_Combined\": 999}", "selected": false}, "selected": false}, {"data": {"id": "2783373", "source": "2782722", "target": "2782785", "SCORE": "239", "shared_interaction": "Direct", "source_original": "851111", "Name_B": "ATP20", "Name_A": "COX8", "target_original": "856131", "Gene_B": "856131", "interaction": "Direct", "SUID": 2783373, "TYPE": "Direct", "Gene_A": "851111", "support": "{\"StringDB\": \"physical\", \"StringEvidence\": \"coexpression:0.989,textmining:0.772,experiments:0.208\", \"String_Score_Physical\": 239, \"String_Score_Combined\": 998}", "selected": false}, "selected": false}, {"data": {"id": "2783370", "source": "2782722", "target": "2782782", "SCORE": "315", "shared_interaction": "Direct", "source_original": "851111", "Name_B": "ATP4", "Name_A": "COX8", "target_original": "856027", "Gene_B": "856027", "interaction": "Direct", "SUID": 2783370, "TYPE": "Direct", "Gene_A": "851111", "support": "{\"StringDB\": \"physical\", \"StringEvidence\": \"coexpression:0.676,textmining:0.493,experiments:0.286\", \"BioGrid\": \"genetic,physical\", \"BioGrid_PubMed\": \"29688796,21623372\", \"BioGrid_Type\": \"Affinity Capture-Western,Negative Genetic\", \"BioGrid_Score\": \"-0.1755\", \"String_Score_Physical\": 315, \"String_Score_Combined\": 887}", "selected": false}, "selected": false}, {"data": {"id": "2783367", "source": "2782722", "target": "2782779", "SCORE": "983", "shared_interaction": "Direct", "source_original": "851111", "Name_B": "COX5A", "Name_A": "COX8", "target_original": "855675", "Gene_B": "855675", "interaction": "Direct", "SUID": 2783367, "TYPE": "Direct", "Gene_A": "851111", "support": "{\"StringDB\": \"physical\", \"StringEvidence\": \"textmining:0.930,coexpression:0.920,experiments:0.920,database:0.791\", \"String_Score_Physical\": 983, \"String_Score_Combined\": 999}", "selected": false}, "selected": false}, {"data": {"id": "2783364", "source": "2782722", "target": "2782776", "SCORE": "958", "shared_interaction": "Direct", "source_original": "851111", "Name_B": "COX7", "Name_A": "COX8", "target_original": "855298", "Gene_B": "855298", "interaction": "Direct", "SUID": 2783364, "TYPE": "Direct", "Gene_A": "851111", "support": "{\"StringDB\": \"physical\", \"StringEvidence\": \"coexpression:0.919,textmining:0.904,experiments:0.791,database:0.791\", \"String_Score_Physical\": 958, \"String_Score_Combined\": 999}", "selected": false}, "selected": false}, {"data": {"id": "2783361", "source": "2782722", "target": "2782773", "SCORE": "900", "shared_interaction": "Direct", "source_original": "851111", "Name_B": "NDI1", "Name_A": "COX8", "target_original": "854919", "Gene_B": "854919", "interaction": "Direct", "SUID": 2783361, "TYPE": "Direct", "Gene_A": "851111", "support": "{\"StringDB\": \"physical\", \"StringEvidence\": \"database:0.896,coexpression:0.169\", \"String_Score_Physical\": 900, \"String_Score_Combined\": 916}", "selected": false}, "selected": false}, {"data": {"id": "2783358", "source": "2782722", "target": "2782770", "SCORE": "230", "shared_interaction": "Direct", "source_original": "851111", "Name_B": "ATP18", "Name_A": "COX8", "target_original": "854893", "Gene_B": "854893", "interaction": "Direct", "SUID": 2783358, "TYPE": "Direct", "Gene_A": "851111", "support": "{\"StringDB\": \"physical\", \"StringEvidence\": \"textmining:0.548,coexpression:0.518,experiments:0.198\", \"BioGrid\": \"genetic\", \"BioGrid_PubMed\": \"27708008,20093466\", \"BioGrid_Type\": \"Negative Genetic\", \"BioGrid_Score\": \"-0.3508,-0.3825\", \"String_Score_Physical\": 230, \"String_Score_Combined\": 832}", "selected": false}, "selected": false}, {"data": {"id": "2783355", "source": "2782722", "target": "2782767", "SCORE": "977", "shared_interaction": "Direct", "source_original": "851111", "Name_B": "COX5B", "Name_A": "COX8", "target_original": "854695", "Gene_B": "854695", "interaction": "Direct", "SUID": 2783355, "TYPE": "Direct", "Gene_A": "851111", "support": "{\"StringDB\": \"physical\", \"StringEvidence\": \"textmining:0.927,experiments:0.920,database:0.708,coexpression:0.087\", \"String_Score_Physical\": 977, \"String_Score_Combined\": 998}", "selected": false}, "selected": false}, {"data": {"id": "2783352", "source": "2782722", "target": "2782764", "SCORE": "991", "shared_interaction": "Direct", "source_original": "851111", "Name_B": "CYT1", "Name_A": "COX8", "target_original": "854231", "Gene_B": "854231", "interaction": "Direct", "SUID": 2783352, "TYPE": "Direct", "Gene_A": "851111", "support": "{\"StringDB\": \"physical\", \"StringEvidence\": \"experiments:0.920,database:0.896,coexpression:0.639,textmining:0.481\", \"String_Score_Physical\": 991, \"String_Score_Combined\": 998}", "selected": false}, "selected": false}, {"data": {"id": "2783349", "source": "2782722", "target": "2782752", "SCORE": "997", "shared_interaction": "Direct", "source_original": "851111", "Name_B": "QCR8", "Name_A": "COX8", "target_original": "853273", "Gene_B": "853273", "interaction": "Direct", "SUID": 2783349, "TYPE": "Direct", "Gene_A": "851111", "support": "{\"StringDB\": \"physical\", \"StringEvidence\": \"experiments:0.977,coexpression:0.973,database:0.896,textmining:0.589\", \"String_Score_Physical\": 997, \"String_Score_Combined\": 999}", "selected": false}, "selected": false}, {"data": {"id": "2783346", "source": "2782722", "target": "2782749", "SCORE": "997", "shared_interaction": "Direct", "source_original": "851111", "Name_B": "QCR9", "Name_A": "COX8", "target_original": "853095", "Gene_B": "853095", "interaction": "Direct", "SUID": 2783346, "TYPE": "Direct", "Gene_A": "851111", "support": "{\"StringDB\": \"physical\", \"StringEvidence\": \"experiments:0.972,database:0.896,coexpression:0.889,textmining:0.216\", \"BioGrid\": \"genetic\", \"BioGrid_PubMed\": \"21623372\", \"BioGrid_Type\": \"Positive Genetic\", \"BioGrid_Score\": \"0.0821\", \"String_Score_Physical\": 997, \"String_Score_Combined\": 999}", "selected": false}, "selected": false}, {"data": {"id": "2783343", "source": "2782722", "target": "2782746", "SCORE": "995", "shared_interaction": "Direct", "source_original": "851111", "Name_B": "COX4", "Name_A": "COX8", "target_original": "852688", "Gene_B": "852688", "interaction": "Direct", "SUID": 2783343, "TYPE": "Direct", "Gene_A": "851111", "support": "{\"StringDB\": \"physical\", \"StringEvidence\": \"coexpression:0.992,experiments:0.980,textmining:0.975,database:0.791\", \"BioGrid\": \"physical\", \"BioGrid_PubMed\": \"29688796,23864706\", \"BioGrid_Type\": \"Affinity Capture-MS,Affinity Capture-Western\", \"String_Score_Physical\": 995, \"String_Score_Combined\": 999}", "selected": false}, "selected": false}, {"data": {"id": "2783340", "source": "2782722", "target": "2782740", "SCORE": "992", "shared_interaction": "Direct", "source_original": "851111", "Name_B": "COR1", "Name_A": "COX8", "target_original": "852235", "Gene_B": "852235", "interaction": "Direct", "SUID": 2783340, "TYPE": "Direct", "Gene_A": "851111", "support": "{\"StringDB\": \"physical\", \"StringEvidence\": \"experiments:0.925,database:0.896,coexpression:0.466,textmining:0.276\", \"BioGrid\": \"physical\", \"BioGrid_PubMed\": \"22342701\", \"BioGrid_Type\": \"Affinity Capture-MS\", \"String_Score_Physical\": 992, \"String_Score_Combined\": 997}", "selected": false}, "selected": false}, {"data": {"id": "2783337", "source": "2782722", "target": "2782734", "SCORE": "991", "shared_interaction": "Direct", "source_original": "851111", "Name_B": "QCR7", "Name_A": "COX8", "target_original": "852142", "Gene_B": "852142", "interaction": "Direct", "SUID": 2783337, "TYPE": "Direct", "Gene_A": "851111", "support": "{\"StringDB\": \"physical\", \"StringEvidence\": \"coexpression:0.979,experiments:0.917,database:0.896,textmining:0.360\", \"String_Score_Physical\": 991, \"String_Score_Combined\": 999}", "selected": false}, "selected": false}, {"data": {"id": "2783334", "source": "2782719", "target": "2782785", "SCORE": "720", "shared_interaction": "Direct", "source_original": "851002", "Name_B": "ATP20", "Name_A": "ATP14", "target_original": "856131", "Gene_B": "856131", "interaction": "Direct", "SUID": 2783334, "TYPE": "Direct", "Gene_A": "851002", "support": "{\"StringDB\": \"physical\", \"StringEvidence\": \"coexpression:0.750,database:0.708,textmining:0.550\", \"String_Score_Physical\": 720, \"String_Score_Combined\": 968}", "selected": false}, "selected": false}, {"data": {"id": "2783331", "source": "2782719", "target": "2782782", "SCORE": "999", "shared_interaction": "Direct", "source_original": "851002", "Name_B": "ATP4", "Name_A": "ATP14", "target_original": "856027", "Gene_B": "856027", "interaction": "Direct", "SUID": 2783331, "TYPE": "Direct", "Gene_A": "851002", "support": "{\"StringDB\": \"physical\", \"StringEvidence\": \"experiments:0.999,textmining:0.803,database:0.708,coexpression:0.487\", \"BioGrid\": \"genetic,physical\", \"BioGrid_PubMed\": \"9867807,21623372\", \"BioGrid_Type\": \"Co-purification,Positive Genetic\", \"BioGrid_Score\": \"0.1201\", \"String_Score_Physical\": 999, \"String_Score_Combined\": 999}", "selected": false}, "selected": false}, {"data": {"id": "2783328", "source": "2782719", "target": "2782770", "SCORE": "983", "shared_interaction": "Direct", "source_original": "851002", "Name_B": "ATP18", "Name_A": "ATP14", "target_original": "854893", "Gene_B": "854893", "interaction": "Direct", "SUID": 2783328, "TYPE": "Direct", "Gene_A": "851002", "support": "{\"StringDB\": \"physical\", \"StringEvidence\": \"experiments:0.941,database:0.708,textmining:0.698,coexpression:0.244\", \"BioGrid\": \"genetic\", \"BioGrid_PubMed\": \"21623372\", \"BioGrid_Type\": \"Positive Genetic\", \"BioGrid_Score\": \"0.1835\", \"String_Score_Physical\": 983, \"String_Score_Combined\": 996}", "selected": false}, "selected": false}, {"data": {"id": "2783325", "source": "2782719", "target": "2782761", "SCORE": "720", "shared_interaction": "Direct", "source_original": "851002", "Name_B": "ATP19", "Name_A": "ATP14", "target_original": "854077", "Gene_B": "854077", "interaction": "Direct", "SUID": 2783325, "TYPE": "Direct", "Gene_A": "851002", "support": "{\"StringDB\": \"physical\", \"StringEvidence\": \"database:0.708,coexpression:0.447,textmining:0.258\", \"String_Score_Physical\": 720, \"String_Score_Combined\": 885}", "selected": false}, "selected": false}, {"data": {"id": "2783322", "source": "2782719", "target": "2782758", "SCORE": "987", "shared_interaction": "Direct", "source_original": "851002", "Name_B": "ATP7", "Name_A": "ATP14", "target_original": "853853", "Gene_B": "853853", "interaction": "Direct", "SUID": 2783322, "TYPE": "Direct", "Gene_A": "851002", "support": "{\"StringDB\": \"physical\", \"StringEvidence\": \"experiments:0.955,coexpression:0.899,textmining:0.846,database:0.708\", \"BioGrid\": \"genetic,physical\", \"BioGrid_PubMed\": \"22940862,21623372\", \"BioGrid_Type\": \"Affinity Capture-MS,Positive Genetic\", \"BioGrid_Score\": \"0.1471\", \"String_Score_Physical\": 987, \"String_Score_Combined\": 999}", "selected": false}, "selected": false}, {"data": {"id": "2783319", "source": "2782719", "target": "2782743", "SCORE": "980", "shared_interaction": "Direct", "source_original": "851002", "Name_B": "ATP3", "Name_A": "ATP14", "target_original": "852327", "Gene_B": "852327", "interaction": "Direct", "SUID": 2783319, "TYPE": "Direct", "Gene_A": "851002", "support": "{\"StringDB\": \"physical\", \"StringEvidence\": \"experiments:0.929,textmining:0.733,database:0.708,coexpression:0.390\", \"String_Score_Physical\": 980, \"String_Score_Combined\": 996}", "selected": false}, "selected": false}, {"data": {"id": "2783316", "source": "2782719", "target": "2782737", "SCORE": "999", "shared_interaction": "Direct", "source_original": "851002", "Name_B": "ATP1", "Name_A": "ATP14", "target_original": "852177", "Gene_B": "852177", "interaction": "Direct", "SUID": 2783316, "TYPE": "Direct", "Gene_A": "851002", "support": "{\"StringDB\": \"physical\", \"StringEvidence\": \"experiments:0.999,database:0.708,textmining:0.705,coexpression:0.138\", \"BioGrid\": \"physical\", \"BioGrid_PubMed\": \"22940862\", \"BioGrid_Type\": \"Affinity Capture-MS\", \"String_Score_Physical\": 999, \"String_Score_Combined\": 999}", "selected": false}, "selected": false}, {"data": {"id": "2783313", "source": "2782719", "target": "2782731", "SCORE": "980", "shared_interaction": "Direct", "source_original": "851002", "Name_B": "ATP17", "Name_A": "ATP14", "target_original": "851983", "Gene_B": "851983", "interaction": "Direct", "SUID": 2783313, "TYPE": "Direct", "Gene_A": "851002", "support": "{\"StringDB\": \"physical\", \"StringEvidence\": \"experiments:0.930,coexpression:0.888,database:0.708,textmining:0.639\", \"BioGrid\": \"genetic\", \"BioGrid_PubMed\": \"21623372\", \"BioGrid_Type\": \"Positive Genetic\", \"BioGrid_Score\": \"0.1113\", \"String_Score_Physical\": 980, \"String_Score_Combined\": 999}", "selected": false}, "selected": false}, {"data": {"id": "2783310", "source": "2782719", "target": "2782728", "SCORE": "727", "shared_interaction": "Direct", "source_original": "851002", "Name_B": "TIM11", "Name_A": "ATP14", "target_original": "851922", "Gene_B": "851922", "interaction": "Direct", "SUID": 2783310, "TYPE": "Direct", "Gene_A": "851002", "support": "{\"StringDB\": \"physical\", \"StringEvidence\": \"database:0.708,textmining:0.600,coexpression:0.214,experiments:0.025\", \"BioGrid\": \"genetic\", \"BioGrid_PubMed\": \"21623372\", \"BioGrid_Type\": \"Positive Genetic\", \"BioGrid_Score\": \"0.1082\", \"String_Score_Physical\": 727, \"String_Score_Combined\": 914}", "selected": false}, "selected": false}, {"data": {"id": "2783289", "source": "2782716", "target": "2782797", "SCORE": "999", "shared_interaction": "Direct", "source_original": "850727", "Name_B": "RIP1", "Name_A": "COX12", "target_original": "856689", "Gene_B": "856689", "interaction": "Direct", "SUID": 2783289, "TYPE": "Direct", "Gene_A": "850727", "support": "{\"StringDB\": \"physical\", \"StringEvidence\": \"experiments:0.999,coexpression:0.925,database:0.896,textmining:0.448\", \"String_Score_Physical\": 999, \"String_Score_Combined\": 999}", "selected": false}, "selected": false}, {"data": {"id": "2783286", "source": "2782716", "target": "2782794", "SCORE": "618", "shared_interaction": "Direct", "source_original": "850727", "Name_B": "CYC7", "Name_A": "COX12", "target_original": "856672", "Gene_B": "856672", "interaction": "Direct", "SUID": 2783286, "TYPE": "Direct", "Gene_A": "850727", "support": "{\"StringDB\": \"physical\", \"StringEvidence\": \"experiments:0.602,textmining:0.312,coexpression:0.191\", \"String_Score_Physical\": 618, \"String_Score_Combined\": 787}", "selected": false}, "selected": false}, {"data": {"id": "2783283", "source": "2782716", "target": "2782791", "SCORE": "999", "shared_interaction": "Direct", "source_original": "850727", "Name_B": "QCR10", "Name_A": "COX12", "target_original": "856390", "Gene_B": "856390", "interaction": "Direct", "SUID": 2783283, "TYPE": "Direct", "Gene_A": "850727", "support": "{\"StringDB\": \"physical\", \"StringEvidence\": \"experiments:0.999,database:0.896,coexpression:0.886\", \"String_Score_Physical\": 999, \"String_Score_Combined\": 999}", "selected": false}, "selected": false}, {"data": {"id": "2783280", "source": "2782716", "target": "2782788", "SCORE": "999", "shared_interaction": "Direct", "source_original": "850727", "Name_B": "QCR2", "Name_A": "COX12", "target_original": "856321", "Gene_B": "856321", "interaction": "Direct", "SUID": 2783280, "TYPE": "Direct", "Gene_A": "850727", "support": "{\"StringDB\": \"physical\", \"StringEvidence\": \"experiments:0.999,database:0.896,textmining:0.612,coexpression:0.422\", \"String_Score_Physical\": 999, \"String_Score_Combined\": 999}", "selected": false}, "selected": false}, {"data": {"id": "2783277", "source": "2782716", "target": "2782785", "SCORE": "963", "shared_interaction": "Direct", "source_original": "850727", "Name_B": "ATP20", "Name_A": "COX12", "target_original": "856131", "Gene_B": "856131", "interaction": "Direct", "SUID": 2783277, "TYPE": "Direct", "Gene_A": "850727", "support": "{\"StringDB\": \"physical\", \"StringEvidence\": \"coexpression:0.980,experiments:0.961,textmining:0.181\", \"String_Score_Physical\": 963, \"String_Score_Combined\": 999}", "selected": false}, "selected": false}, {"data": {"id": "2783274", "source": "2782716", "target": "2782782", "SCORE": "964", "shared_interaction": "Direct", "source_original": "850727", "Name_B": "ATP4", "Name_A": "COX12", "target_original": "856027", "Gene_B": "856027", "interaction": "Direct", "SUID": 2783274, "TYPE": "Direct", "Gene_A": "850727", "support": "{\"StringDB\": \"physical\", \"StringEvidence\": \"experiments:0.962,coexpression:0.519,textmining:0.192\", \"String_Score_Physical\": 964, \"String_Score_Combined\": 986}", "selected": false}, "selected": false}, {"data": {"id": "2783271", "source": "2782716", "target": "2782779", "SCORE": "999", "shared_interaction": "Direct", "source_original": "850727", "Name_B": "COX5A", "Name_A": "COX12", "target_original": "855675", "Gene_B": "855675", "interaction": "Direct", "SUID": 2783271, "TYPE": "Direct", "Gene_A": "850727", "support": "{\"StringDB\": \"physical\", \"StringEvidence\": \"experiments:1.000,coexpression:0.936,textmining:0.923,database:0.791\", \"String_Score_Physical\": 999, \"String_Score_Combined\": 999}", "selected": false}, "selected": false}, {"data": {"id": "2783268", "source": "2782716", "target": "2782776", "SCORE": "999", "shared_interaction": "Direct", "source_original": "850727", "Name_B": "COX7", "Name_A": "COX12", "target_original": "855298", "Gene_B": "855298", "interaction": "Direct", "SUID": 2783268, "TYPE": "Direct", "Gene_A": "850727", "support": "{\"StringDB\": \"physical\", \"StringEvidence\": \"experiments:0.999,coexpression:0.919,textmining:0.900,database:0.791\", \"String_Score_Physical\": 999, \"String_Score_Combined\": 999}", "selected": false}, "selected": false}, {"data": {"id": "2783265", "source": "2782716", "target": "2782773", "SCORE": "900", "shared_interaction": "Direct", "source_original": "850727", "Name_B": "NDI1", "Name_A": "COX12", "target_original": "854919", "Gene_B": "854919", "interaction": "Direct", "SUID": 2783265, "TYPE": "Direct", "Gene_A": "850727", "support": "{\"StringDB\": \"physical\", \"StringEvidence\": \"database:0.896,textmining:0.556,coexpression:0.202\", \"String_Score_Physical\": 900, \"String_Score_Combined\": 964}", "selected": false}, "selected": false}, {"data": {"id": "2783262", "source": "2782716", "target": "2782767", "SCORE": "975", "shared_interaction": "Direct", "source_original": "850727", "Name_B": "COX5B", "Name_A": "COX12", "target_original": "854695", "Gene_B": "854695", "interaction": "Direct", "SUID": 2783262, "TYPE": "Direct", "Gene_A": "850727", "support": "{\"StringDB\": \"physical\", \"StringEvidence\": \"textmining:0.924,experiments:0.913,database:0.708,coexpression:0.220\", \"String_Score_Physical\": 975, \"String_Score_Combined\": 998}", "selected": false}, "selected": false}, {"data": {"id": "2783259", "source": "2782716", "target": "2782764", "SCORE": "999", "shared_interaction": "Direct", "source_original": "850727", "Name_B": "CYT1", "Name_A": "COX12", "target_original": "854231", "Gene_B": "854231", "interaction": "Direct", "SUID": 2783259, "TYPE": "Direct", "Gene_A": "850727", "support": "{\"StringDB\": \"physical\", \"StringEvidence\": \"experiments:0.999,coexpression:0.951,database:0.896,textmining:0.743\", \"String_Score_Physical\": 999, \"String_Score_Combined\": 999}", "selected": false}, "selected": false}, {"data": {"id": "2783256", "source": "2782716", "target": "2782758", "SCORE": "257", "shared_interaction": "Direct", "source_original": "850727", "Name_B": "ATP7", "Name_A": "COX12", "target_original": "853853", "Gene_B": "853853", "interaction": "Direct", "SUID": 2783256, "TYPE": "Direct", "Gene_A": "850727", "support": "{\"StringDB\": \"physical\", \"StringEvidence\": \"coexpression:0.950,experiments:0.226,textmining:0.036\", \"BioGrid\": \"genetic\", \"BioGrid_PubMed\": \"27708008\", \"BioGrid_Type\": \"Negative Genetic\", \"BioGrid_Score\": \"-0.2994\", \"String_Score_Physical\": 257, \"String_Score_Combined\": 964}", "selected": false}, "selected": false}, {"data": {"id": "2783253", "source": "2782716", "target": "2782752", "SCORE": "999", "shared_interaction": "Direct", "source_original": "850727", "Name_B": "QCR8", "Name_A": "COX12", "target_original": "853273", "Gene_B": "853273", "interaction": "Direct", "SUID": 2783253, "TYPE": "Direct", "Gene_A": "850727", "support": "{\"StringDB\": \"physical\", \"StringEvidence\": \"experiments:0.999,coexpression:0.960,database:0.896,textmining:0.801\", \"BioGrid\": \"genetic\", \"BioGrid_PubMed\": \"21623372\", \"BioGrid_Type\": \"Positive Genetic\", \"BioGrid_Score\": \"0.0925\", \"String_Score_Physical\": 999, \"String_Score_Combined\": 999}", "selected": false}, "selected": false}, {"data": {"id": "2783250", "source": "2782716", "target": "2782749", "SCORE": "999", "shared_interaction": "Direct", "source_original": "850727", "Name_B": "QCR9", "Name_A": "COX12", "target_original": "853095", "Gene_B": "853095", "interaction": "Direct", "SUID": 2783250, "TYPE": "Direct", "Gene_A": "850727", "support": "{\"StringDB\": \"physical\", \"StringEvidence\": \"experiments:0.999,coexpression:0.944,database:0.896,textmining:0.544\", \"String_Score_Physical\": 999, \"String_Score_Combined\": 999}", "selected": false}, "selected": false}, {"data": {"id": "2783247", "source": "2782716", "target": "2782746", "SCORE": "999", "shared_interaction": "Direct", "source_original": "850727", "Name_B": "COX4", "Name_A": "COX12", "target_original": "852688", "Gene_B": "852688", "interaction": "Direct", "SUID": 2783247, "TYPE": "Direct", "Gene_A": "850727", "support": "{\"StringDB\": \"physical\", \"StringEvidence\": \"experiments:0.999,coexpression:0.991,textmining:0.965,database:0.791\", \"BioGrid\": \"physical\", \"BioGrid_PubMed\": \"23864706\", \"BioGrid_Type\": \"Affinity Capture-MS\", \"String_Score_Physical\": 999, \"String_Score_Combined\": 999}", "selected": false}, "selected": false}, {"data": {"id": "2783244", "source": "2782716", "target": "2782743", "SCORE": "243", "shared_interaction": "Direct", "source_original": "850727", "Name_B": "ATP3", "Name_A": "COX12", "target_original": "852327", "Gene_B": "852327", "interaction": "Direct", "SUID": 2783244, "TYPE": "Direct", "Gene_A": "850727", "support": "{\"StringDB\": \"physical\", \"StringEvidence\": \"coexpression:0.907,experiments:0.212,textmining:0.199\", \"String_Score_Physical\": 243, \"String_Score_Combined\": 943}", "selected": false}, "selected": false}, {"data": {"id": "2783241", "source": "2782716", "target": "2782740", "SCORE": "999", "shared_interaction": "Direct", "source_original": "850727", "Name_B": "COR1", "Name_A": "COX12", "target_original": "852235", "Gene_B": "852235", "interaction": "Direct", "SUID": 2783241, "TYPE": "Direct", "Gene_A": "850727", "support": "{\"StringDB\": \"physical\", \"StringEvidence\": \"experiments:0.999,database:0.896,textmining:0.470,coexpression:0.447\", \"BioGrid\": \"genetic,physical\", \"BioGrid_PubMed\": \"22342701,21623372\", \"BioGrid_Type\": \"Affinity Capture-MS,Positive Genetic\", \"BioGrid_Score\": \"0.1078\", \"String_Score_Physical\": 999, \"String_Score_Combined\": 999}", "selected": false}, "selected": false}, {"data": {"id": "2783238", "source": "2782716", "target": "2782734", "SCORE": "999", "shared_interaction": "Direct", "source_original": "850727", "Name_B": "QCR7", "Name_A": "COX12", "target_original": "852142", "Gene_B": "852142", "interaction": "Direct", "SUID": 2783238, "TYPE": "Direct", "Gene_A": "850727", "support": "{\"StringDB\": \"physical\", \"StringEvidence\": \"experiments:0.999,coexpression:0.989,database:0.896,textmining:0.707\", \"String_Score_Physical\": 999, \"String_Score_Combined\": 999}", "selected": false}, "selected": false}, {"data": {"id": "2783235", "source": "2782716", "target": "2782722", "SCORE": "999", "shared_interaction": "Direct", "source_original": "850727", "Name_B": "COX8", "Name_A": "COX12", "target_original": "851111", "Gene_B": "851111", "interaction": "Direct", "SUID": 2783235, "TYPE": "Direct", "Gene_A": "850727", "support": "{\"StringDB\": \"physical\", \"StringEvidence\": \"experiments:0.999,coexpression:0.976,textmining:0.975,database:0.791\", \"String_Score_Physical\": 999, \"String_Score_Combined\": 999}", "selected": false}, "selected": false}, {"data": {"id": "2783232", "source": "2782713", "target": "2782797", "SCORE": "923", "shared_interaction": "Direct", "source_original": "850685", "Name_B": "RIP1", "Name_A": "SDH2", "target_original": "856689", "Gene_B": "856689", "interaction": "Direct", "SUID": 2783232, "TYPE": "Direct", "Gene_A": "850685", "support": "{\"StringDB\": \"physical\", \"StringEvidence\": \"coexpression:0.992,database:0.896,textmining:0.789,experiments:0.230,neighborhood:0.109\", \"String_Score_Physical\": 923, \"String_Score_Combined\": 999}", "selected": false}, "selected": false}, {"data": {"id": "2783229", "source": "2782713", "target": "2782794", "SCORE": "193", "shared_interaction": "Direct", "source_original": "850685", "Name_B": "CYC7", "Name_A": "SDH2", "target_original": "856672", "Gene_B": "856672", "interaction": "Direct", "SUID": 2783229, "TYPE": "Direct", "Gene_A": "850685", "support": "{\"StringDB\": \"physical\", \"StringEvidence\": \"textmining:0.608,experiments:0.158,coexpression:0.157\", \"String_Score_Physical\": 193, \"String_Score_Combined\": 733}", "selected": false}, "selected": false}, {"data": {"id": "2783226", "source": "2782713", "target": "2782791", "SCORE": "900", "shared_interaction": "Direct", "source_original": "850685", "Name_B": "QCR10", "Name_A": "SDH2", "target_original": "856390", "Gene_B": "856390", "interaction": "Direct", "SUID": 2783226, "TYPE": "Direct", "Gene_A": "850685", "support": "{\"StringDB\": \"physical\", \"StringEvidence\": \"database:0.896,coexpression:0.715,textmining:0.187\", \"String_Score_Physical\": 900, \"String_Score_Combined\": 976}", "selected": false}, "selected": false}, {"data": {"id": "2783223", "source": "2782713", "target": "2782788", "SCORE": "996", "shared_interaction": "Direct", "source_original": "850685", "Name_B": "QCR2", "Name_A": "SDH2", "target_original": "856321", "Gene_B": "856321", "interaction": "Direct", "SUID": 2783223, "TYPE": "Direct", "Gene_A": "850685", "support": "{\"StringDB\": \"physical\", \"StringEvidence\": \"experiments:0.965,coexpression:0.941,database:0.896,textmining:0.893\", \"String_Score_Physical\": 996, \"String_Score_Combined\": 999}", "selected": false}, "selected": false}, {"data": {"id": "2783220", "source": "2782713", "target": "2782785", "SCORE": "962", "shared_interaction": "Direct", "source_original": "850685", "Name_B": "ATP20", "Name_A": "SDH2", "target_original": "856131", "Gene_B": "856131", "interaction": "Direct", "SUID": 2783220, "TYPE": "Direct", "Gene_A": "850685", "support": "{\"StringDB\": \"physical\", \"StringEvidence\": \"experiments:0.960,coexpression:0.327,textmining:0.163\", \"String_Score_Physical\": 962, \"String_Score_Combined\": 978}", "selected": false}, "selected": false}, {"data": {"id": "2783217", "source": "2782713", "target": "2782782", "SCORE": "963", "shared_interaction": "Direct", "source_original": "850685", "Name_B": "ATP4", "Name_A": "SDH2", "target_original": "856027", "Gene_B": "856027", "interaction": "Direct", "SUID": 2783217, "TYPE": "Direct", "Gene_A": "850685", "support": "{\"StringDB\": \"physical\", \"StringEvidence\": \"experiments:0.961,coexpression:0.714,textmining:0.410\", \"String_Score_Physical\": 963, \"String_Score_Combined\": 993}", "selected": false}, "selected": false}, {"data": {"id": "2783214", "source": "2782713", "target": "2782773", "SCORE": "900", "shared_interaction": "Direct", "source_original": "850685", "Name_B": "NDI1", "Name_A": "SDH2", "target_original": "854919", "Gene_B": "854919", "interaction": "Direct", "SUID": 2783214, "TYPE": "Direct", "Gene_A": "850685", "support": "{\"StringDB\": \"physical\", \"StringEvidence\": \"coexpression:0.925,database:0.896,textmining:0.572\", \"String_Score_Physical\": 900, \"String_Score_Combined\": 996}", "selected": false}, "selected": false}, {"data": {"id": "2783211", "source": "2782713", "target": "2782764", "SCORE": "922", "shared_interaction": "Direct", "source_original": "850685", "Name_B": "CYT1", "Name_A": "SDH2", "target_original": "854231", "Gene_B": "854231", "interaction": "Direct", "SUID": 2783211, "TYPE": "Direct", "Gene_A": "850685", "support": "{\"StringDB\": \"physical\", \"StringEvidence\": \"coexpression:0.982,database:0.896,textmining:0.671,experiments:0.224\", \"String_Score_Physical\": 922, \"String_Score_Combined\": 999}", "selected": false}, "selected": false}, {"data": {"id": "2783208", "source": "2782713", "target": "2782758", "SCORE": "248", "shared_interaction": "Direct", "source_original": "850685", "Name_B": "ATP7", "Name_A": "SDH2", "target_original": "853853", "Gene_B": "853853", "interaction": "Direct", "SUID": 2783208, "TYPE": "Direct", "Gene_A": "850685", "support": "{\"StringDB\": \"physical\", \"StringEvidence\": \"coexpression:0.952,textmining:0.686,experiments:0.217\", \"String_Score_Physical\": 248, \"String_Score_Combined\": 988}", "selected": false}, "selected": false}, {"data": {"id": "2783205", "source": "2782713", "target": "2782755", "SCORE": "995", "shared_interaction": "Direct", "source_original": "850685", "Name_B": "SDH3", "Name_A": "SDH2", "target_original": "853716", "Gene_B": "853716", "interaction": "Direct", "SUID": 2783205, "TYPE": "Direct", "Gene_A": "850685", "support": "{\"StringDB\": \"physical\", \"StringEvidence\": \"coexpression:0.989,textmining:0.970,experiments:0.958,database:0.896,neighborhood:0.138\", \"String_Score_Physical\": 995, \"String_Score_Combined\": 999}", "selected": false}, "selected": false}, {"data": {"id": "2783202", "source": "2782713", "target": "2782752", "SCORE": "900", "shared_interaction": "Direct", "source_original": "850685", "Name_B": "QCR8", "Name_A": "SDH2", "target_original": "853273", "Gene_B": "853273", "interaction": "Direct", "SUID": 2783202, "TYPE": "Direct", "Gene_A": "850685", "support": "{\"StringDB\": \"physical\", \"StringEvidence\": \"database:0.896,coexpression:0.646,textmining:0.549\", \"String_Score_Physical\": 900, \"String_Score_Combined\": 984}", "selected": false}, "selected": false}, {"data": {"id": "2783199", "source": "2782713", "target": "2782749", "SCORE": "922", "shared_interaction": "Direct", "source_original": "850685", "Name_B": "QCR9", "Name_A": "SDH2", "target_original": "853095", "Gene_B": "853095", "interaction": "Direct", "SUID": 2783199, "TYPE": "Direct", "Gene_A": "850685", "support": "{\"StringDB\": \"physical\", \"StringEvidence\": \"database:0.896,coexpression:0.771,textmining:0.270,experiments:0.228\", \"String_Score_Physical\": 922, \"String_Score_Combined\": 987}", "selected": false}, "selected": false}, {"data": {"id": "2783196", "source": "2782713", "target": "2782746", "SCORE": "553", "shared_interaction": "Direct", "source_original": "850685", "Name_B": "COX4", "Name_A": "SDH2", "target_original": "852688", "Gene_B": "852688", "interaction": "Direct", "SUID": 2783196, "TYPE": "Direct", "Gene_A": "850685", "support": "{\"StringDB\": \"physical\", \"StringEvidence\": \"coexpression:0.990,textmining:0.599,experiments:0.534\", \"String_Score_Physical\": 553, \"String_Score_Combined\": 998}", "selected": false}, "selected": false}, {"data": {"id": "2783193", "source": "2782713", "target": "2782743", "SCORE": "254", "shared_interaction": "Direct", "source_original": "850685", "Name_B": "ATP3", "Name_A": "SDH2", "target_original": "852327", "Gene_B": "852327", "interaction": "Direct", "SUID": 2783193, "TYPE": "Direct", "Gene_A": "850685", "support": "{\"StringDB\": \"physical\", \"StringEvidence\": \"coexpression:0.934,textmining:0.647,experiments:0.223,neighborhood:0.141\", \"String_Score_Physical\": 254, \"String_Score_Combined\": 984}", "selected": false}, "selected": false}, {"data": {"id": "2783190", "source": "2782713", "target": "2782740", "SCORE": "924", "shared_interaction": "Direct", "source_original": "850685", "Name_B": "COR1", "Name_A": "SDH2", "target_original": "852235", "Gene_B": "852235", "interaction": "Direct", "SUID": 2783190, "TYPE": "Direct", "Gene_A": "850685", "support": "{\"StringDB\": \"physical\", \"StringEvidence\": \"coexpression:0.922,database:0.896,textmining:0.790,experiments:0.242\", \"String_Score_Physical\": 924, \"String_Score_Combined\": 998}", "selected": false}, "selected": false}, {"data": {"id": "2783187", "source": "2782713", "target": "2782737", "SCORE": "244", "shared_interaction": "Direct", "source_original": "850685", "Name_B": "ATP1", "Name_A": "SDH2", "target_original": "852177", "Gene_B": "852177", "interaction": "Direct", "SUID": 2783187, "TYPE": "Direct", "Gene_A": "850685", "support": "{\"StringDB\": \"physical\", \"StringEvidence\": \"textmining:0.776,coexpression:0.457,experiments:0.213,neighborhood:0.129\", \"String_Score_Physical\": 244, \"String_Score_Combined\": 919}", "selected": false}, "selected": false}, {"data": {"id": "2783184", "source": "2782713", "target": "2782734", "SCORE": "996", "shared_interaction": "Direct", "source_original": "850685", "Name_B": "QCR7", "Name_A": "SDH2", "target_original": "852142", "Gene_B": "852142", "interaction": "Direct", "SUID": 2783184, "TYPE": "Direct", "Gene_A": "850685", "support": "{\"StringDB\": \"physical\", \"StringEvidence\": \"experiments:0.961,database:0.896,coexpression:0.654,textmining:0.629\", \"BioGrid\": \"genetic\", \"BioGrid_PubMed\": \"21623372\", \"BioGrid_Type\": \"Negative Genetic\", \"BioGrid_Score\": \"-0.1452\", \"String_Score_Physical\": 996, \"String_Score_Combined\": 999}", "selected": false}, "selected": false}, {"data": {"id": "2783181", "source": "2782713", "target": "2782725", "SCORE": "980", "shared_interaction": "Direct", "source_original": "850685", "Name_B": "SDH4", "Name_A": "SDH2", "target_original": "851758", "Gene_B": "851758", "interaction": "Direct", "SUID": 2783181, "TYPE": "Direct", "Gene_A": "850685", "support": "{\"StringDB\": \"physical\", \"StringEvidence\": \"coexpression:0.986,textmining:0.964,database:0.896,experiments:0.805\", \"BioGrid\": \"physical\", \"BioGrid_PubMed\": \"29386296,22152483\", \"BioGrid_Type\": \"Affinity Capture-Western\", \"String_Score_Physical\": 980, \"String_Score_Combined\": 999}", "selected": false}, "selected": false}, {"data": {"id": "2783178", "source": "2782713", "target": "2782716", "SCORE": "962", "shared_interaction": "Direct", "source_original": "850685", "Name_B": "COX12", "Name_A": "SDH2", "target_original": "850727", "Gene_B": "850727", "interaction": "Direct", "SUID": 2783178, "TYPE": "Direct", "Gene_A": "850685", "support": "{\"StringDB\": \"physical\", \"StringEvidence\": \"experiments:0.960,coexpression:0.505,textmining:0.162\", \"String_Score_Physical\": 962, \"String_Score_Combined\": 984}", "selected": false}, "selected": false}, {"data": {"id": "2783166", "source": "2782686", "target": "2782797", "SCORE": "999", "shared_interaction": "Direct", "source_original": "850593", "Name_B": "RIP1", "Name_A": "QCR6", "target_original": "856689", "Gene_B": "856689", "interaction": "Direct", "SUID": 2783166, "TYPE": "Direct", "Gene_A": "850593", "support": "{\"StringDB\": \"physical\", \"StringEvidence\": \"experiments:0.995,textmining:0.950,coexpression:0.942,database:0.896\", \"String_Score_Physical\": 999, \"String_Score_Combined\": 999}", "selected": false}, "selected": false}, {"data": {"id": "2783163", "source": "2782686", "target": "2782794", "SCORE": "982", "shared_interaction": "Direct", "source_original": "850593", "Name_B": "CYC7", "Name_A": "QCR6", "target_original": "856672", "Gene_B": "856672", "interaction": "Direct", "SUID": 2783163, "TYPE": "Direct", "Gene_A": "850593", "support": "{\"StringDB\": \"physical\", \"StringEvidence\": \"database:0.896,coexpression:0.891,experiments:0.824,textmining:0.718\", \"String_Score_Physical\": 982, \"String_Score_Combined\": 999}", "selected": false}, "selected": false}, {"data": {"id": "2783160", "source": "2782686", "target": "2782791", "SCORE": "987", "shared_interaction": "Direct", "source_original": "850593", "Name_B": "QCR10", "Name_A": "QCR6", "target_original": "856390", "Gene_B": "856390", "interaction": "Direct", "SUID": 2783160, "TYPE": "Direct", "Gene_A": "850593", "support": "{\"StringDB\": \"physical\", \"StringEvidence\": \"textmining:0.897,database:0.896,experiments:0.878,coexpression:0.318\", \"BioGrid\": \"genetic\", \"BioGrid_PubMed\": \"8175712,21623372,15009199\", \"BioGrid_Type\": \"Negative Genetic,Synthetic Growth Defect\", \"BioGrid_Score\": \"-0.1649,-0.1246\", \"String_Score_Physical\": 987, \"String_Score_Combined\": 999}", "selected": false}, "selected": false}, {"data": {"id": "2783157", "source": "2782686", "target": "2782788", "SCORE": "999", "shared_interaction": "Direct", "source_original": "850593", "Name_B": "QCR2", "Name_A": "QCR6", "target_original": "856321", "Gene_B": "856321", "interaction": "Direct", "SUID": 2783157, "TYPE": "Direct", "Gene_A": "850593", "support": "{\"StringDB\": \"physical\", \"StringEvidence\": \"experiments:0.997,textmining:0.981,coexpression:0.927,database:0.896\", \"BioGrid\": \"physical\", \"BioGrid_PubMed\": \"25683140,21807901\", \"BioGrid_Type\": \"Affinity Capture-Western\", \"String_Score_Physical\": 999, \"String_Score_Combined\": 999}", "selected": false}, "selected": false}, {"data": {"id": "2783154", "source": "2782686", "target": "2782785", "SCORE": "963", "shared_interaction": "Direct", "source_original": "850593", "Name_B": "ATP20", "Name_A": "QCR6", "target_original": "856131", "Gene_B": "856131", "interaction": "Direct", "SUID": 2783154, "TYPE": "Direct", "Gene_A": "850593", "support": "{\"StringDB\": \"physical\", \"StringEvidence\": \"experiments:0.961,coexpression:0.909,textmining:0.561\", \"String_Score_Physical\": 963, \"String_Score_Combined\": 998}", "selected": false}, "selected": false}, {"data": {"id": "2783151", "source": "2782686", "target": "2782782", "SCORE": "964", "shared_interaction": "Direct", "source_original": "850593", "Name_B": "ATP4", "Name_A": "QCR6", "target_original": "856027", "Gene_B": "856027", "interaction": "Direct", "SUID": 2783151, "TYPE": "Direct", "Gene_A": "850593", "support": "{\"StringDB\": \"physical\", \"StringEvidence\": \"experiments:0.962,coexpression:0.910,textmining:0.393\", \"String_Score_Physical\": 964, \"String_Score_Combined\": 998}", "selected": false}, "selected": false}, {"data": {"id": "2783148", "source": "2782686", "target": "2782779", "SCORE": "998", "shared_interaction": "Direct", "source_original": "850593", "Name_B": "COX5A", "Name_A": "QCR6", "target_original": "855675", "Gene_B": "855675", "interaction": "Direct", "SUID": 2783148, "TYPE": "Direct", "Gene_A": "850593", "support": "{\"StringDB\": \"physical\", \"StringEvidence\": \"experiments:0.986,database:0.896,coexpression:0.661,textmining:0.557\", \"BioGrid\": \"physical\", \"BioGrid_PubMed\": \"30598554\", \"BioGrid_Type\": \"Co-crystal Structure\", \"String_Score_Physical\": 998, \"String_Score_Combined\": 999}", "selected": false}, "selected": false}, {"data": {"id": "2783145", "source": "2782686", "target": "2782776", "SCORE": "979", "shared_interaction": "Direct", "source_original": "850593", "Name_B": "COX7", "Name_A": "QCR6", "target_original": "855298", "Gene_B": "855298", "interaction": "Direct", "SUID": 2783145, "TYPE": "Direct", "Gene_A": "850593", "support": "{\"StringDB\": \"physical\", \"StringEvidence\": \"database:0.896,experiments:0.791,textmining:0.656,coexpression:0.231\", \"String_Score_Physical\": 979, \"String_Score_Combined\": 994}", "selected": false}, "selected": false}, {"data": {"id": "2783142", "source": "2782686", "target": "2782773", "SCORE": "900", "shared_interaction": "Direct", "source_original": "850593", "Name_B": "NDI1", "Name_A": "QCR6", "target_original": "854919", "Gene_B": "854919", "interaction": "Direct", "SUID": 2783142, "TYPE": "Direct", "Gene_A": "850593", "support": "{\"StringDB\": \"physical\", \"StringEvidence\": \"database:0.896,textmining:0.293,coexpression:0.231\", \"String_Score_Physical\": 900, \"String_Score_Combined\": 945}", "selected": false}, "selected": false}, {"data": {"id": "2783139", "source": "2782686", "target": "2782767", "SCORE": "984", "shared_interaction": "Direct", "source_original": "850593", "Name_B": "COX5B", "Name_A": "QCR6", "target_original": "854695", "Gene_B": "854695", "interaction": "Direct", "SUID": 2783139, "TYPE": "Direct", "Gene_A": "850593", "support": "{\"StringDB\": \"physical\", \"StringEvidence\": \"database:0.896,experiments:0.845,coexpression:0.170,textmining:0.139\", \"String_Score_Physical\": 984, \"String_Score_Combined\": 988}", "selected": false}, "selected": false}, {"data": {"id": "2783136", "source": "2782686", "target": "2782764", "SCORE": "999", "shared_interaction": "Direct", "source_original": "850593", "Name_B": "CYT1", "Name_A": "QCR6", "target_original": "854231", "Gene_B": "854231", "interaction": "Direct", "SUID": 2783136, "TYPE": "Direct", "Gene_A": "850593", "support": "{\"StringDB\": \"physical\", \"StringEvidence\": \"experiments:0.997,textmining:0.969,coexpression:0.956,database:0.896\", \"String_Score_Physical\": 999, \"String_Score_Combined\": 999}", "selected": false}, "selected": false}, {"data": {"id": "2783133", "source": "2782686", "target": "2782758", "SCORE": "248", "shared_interaction": "Direct", "source_original": "850593", "Name_B": "ATP7", "Name_A": "QCR6", "target_original": "853853", "Gene_B": "853853", "interaction": "Direct", "SUID": 2783133, "TYPE": "Direct", "Gene_A": "850593", "support": "{\"StringDB\": \"physical\", \"StringEvidence\": \"coexpression:0.914,textmining:0.680,experiments:0.217\", \"String_Score_Physical\": 248, \"String_Score_Combined\": 979}", "selected": false}, "selected": false}, {"data": {"id": "2783130", "source": "2782686", "target": "2782755", "SCORE": "909", "shared_interaction": "Direct", "source_original": "850593", "Name_B": "SDH3", "Name_A": "QCR6", "target_original": "853716", "Gene_B": "853716", "interaction": "Direct", "SUID": 2783130, "TYPE": "Direct", "Gene_A": "850593", "support": "{\"StringDB\": \"physical\", \"StringEvidence\": \"database:0.896,coexpression:0.282,textmining:0.243,experiments:0.097\", \"BioGrid\": \"genetic\", \"BioGrid_PubMed\": \"21623372\", \"BioGrid_Type\": \"Negative Genetic\", \"BioGrid_Score\": \"-0.1122\", \"String_Score_Physical\": 909, \"String_Score_Combined\": 950}", "selected": false}, "selected": false}, {"data": {"id": "2783127", "source": "2782686", "target": "2782752", "SCORE": "999", "shared_interaction": "Direct", "source_original": "850593", "Name_B": "QCR8", "Name_A": "QCR6", "target_original": "853273", "Gene_B": "853273", "interaction": "Direct", "SUID": 2783127, "TYPE": "Direct", "Gene_A": "850593", "support": "{\"StringDB\": \"physical\", \"StringEvidence\": \"experiments:0.995,textmining:0.975,coexpression:0.914,database:0.896\", \"String_Score_Physical\": 999, \"String_Score_Combined\": 999}", "selected": false}, "selected": false}, {"data": {"id": "2783124", "source": "2782686", "target": "2782749", "SCORE": "999", "shared_interaction": "Direct", "source_original": "850593", "Name_B": "QCR9", "Name_A": "QCR6", "target_original": "853095", "Gene_B": "853095", "interaction": "Direct", "SUID": 2783124, "TYPE": "Direct", "Gene_A": "850593", "support": "{\"StringDB\": \"physical\", \"StringEvidence\": \"experiments:0.996,textmining:0.992,coexpression:0.934,database:0.896\", \"BioGrid\": \"genetic\", \"BioGrid_PubMed\": \"21623372\", \"BioGrid_Type\": \"Negative Genetic\", \"BioGrid_Score\": \"-0.1266\", \"String_Score_Physical\": 999, \"String_Score_Combined\": 999}", "selected": false}, "selected": false}, {"data": {"id": "2783121", "source": "2782686", "target": "2782746", "SCORE": "993", "shared_interaction": "Direct", "source_original": "850593", "Name_B": "COX4", "Name_A": "QCR6", "target_original": "852688", "Gene_B": "852688", "interaction": "Direct", "SUID": 2783121, "TYPE": "Direct", "Gene_A": "850593", "support": "{\"StringDB\": \"physical\", \"StringEvidence\": \"experiments:0.938,database:0.896,coexpression:0.881,textmining:0.744\", \"BioGrid\": \"physical\", \"BioGrid_PubMed\": \"23864706\", \"BioGrid_Type\": \"Affinity Capture-MS\", \"String_Score_Physical\": 993, \"String_Score_Combined\": 999}", "selected": false}, "selected": false}, {"data": {"id": "2783118", "source": "2782686", "target": "2782743", "SCORE": "246", "shared_interaction": "Direct", "source_original": "850593", "Name_B": "ATP3", "Name_A": "QCR6", "target_original": "852327", "Gene_B": "852327", "interaction": "Direct", "SUID": 2783118, "TYPE": "Direct", "Gene_A": "850593", "support": "{\"StringDB\": \"physical\", \"StringEvidence\": \"coexpression:0.906,textmining:0.507,experiments:0.215\", \"String_Score_Physical\": 246, \"String_Score_Combined\": 965}", "selected": false}, "selected": false}, {"data": {"id": "2783115", "source": "2782686", "target": "2782740", "SCORE": "999", "shared_interaction": "Direct", "source_original": "850593", "Name_B": "COR1", "Name_A": "QCR6", "target_original": "852235", "Gene_B": "852235", "interaction": "Direct", "SUID": 2783115, "TYPE": "Direct", "Gene_A": "850593", "support": "{\"StringDB\": \"physical\", \"StringEvidence\": \"experiments:0.997,textmining:0.958,database:0.896,coexpression:0.527\", \"BioGrid\": \"physical\", \"BioGrid_PubMed\": \"22342701,17882259\", \"BioGrid_Type\": \"Affinity Capture-MS,Affinity Capture-Western\", \"String_Score_Physical\": 999, \"String_Score_Combined\": 999}", "selected": false}, "selected": false}, {"data": {"id": "2783112", "source": "2782686", "target": "2782734", "SCORE": "999", "shared_interaction": "Direct", "source_original": "850593", "Name_B": "QCR7", "Name_A": "QCR6", "target_original": "852142", "Gene_B": "852142", "interaction": "Direct", "SUID": 2783112, "TYPE": "Direct", "Gene_A": "850593", "support": "{\"StringDB\": \"physical\", \"StringEvidence\": \"experiments:0.996,textmining:0.989,coexpression:0.969,database:0.896\", \"BioGrid\": \"genetic\", \"BioGrid_PubMed\": \"21623372,15009199\", \"BioGrid_Type\": \"Negative Genetic,Synthetic Growth Defect\", \"BioGrid_Score\": \"-0.1167\", \"String_Score_Physical\": 999, \"String_Score_Combined\": 999}", "selected": false}, "selected": false}, {"data": {"id": "2783109", "source": "2782686", "target": "2782725", "SCORE": "900", "shared_interaction": "Direct", "source_original": "850593", "Name_B": "SDH4", "Name_A": "QCR6", "target_original": "851758", "Gene_B": "851758", "interaction": "Direct", "SUID": 2783109, "TYPE": "Direct", "Gene_A": "850593", "support": "{\"StringDB\": \"physical\", \"StringEvidence\": \"database:0.896,coexpression:0.186,textmining:0.082\", \"String_Score_Physical\": 900, \"String_Score_Combined\": 925}", "selected": false}, "selected": false}, {"data": {"id": "2783106", "source": "2782686", "target": "2782722", "SCORE": "997", "shared_interaction": "Direct", "source_original": "850593", "Name_B": "COX8", "Name_A": "QCR6", "target_original": "851111", "Gene_B": "851111", "interaction": "Direct", "SUID": 2783106, "TYPE": "Direct", "Gene_A": "850593", "support": "{\"StringDB\": \"physical\", \"StringEvidence\": \"experiments:0.972,coexpression:0.922,database:0.896,textmining:0.756\", \"String_Score_Physical\": 997, \"String_Score_Combined\": 999}", "selected": false}, "selected": false}, {"data": {"id": "2783103", "source": "2782686", "target": "2782716", "SCORE": "999", "shared_interaction": "Direct", "source_original": "850593", "Name_B": "COX12", "Name_A": "QCR6", "target_original": "850727", "Gene_B": "850727", "interaction": "Direct", "SUID": 2783103, "TYPE": "Direct", "Gene_A": "850593", "support": "{\"StringDB\": \"physical\", \"StringEvidence\": \"experiments:0.999,textmining:0.933,coexpression:0.925,database:0.896\", \"String_Score_Physical\": 999, \"String_Score_Combined\": 999}", "selected": false}, "selected": false}, {"data": {"id": "2783100", "source": "2782686", "target": "2782713", "SCORE": "921", "shared_interaction": "Direct", "source_original": "850593", "Name_B": "SDH2", "Name_A": "QCR6", "target_original": "850685", "Gene_B": "850685", "interaction": "Direct", "SUID": 2783100, "TYPE": "Direct", "Gene_A": "850593", "support": "{\"StringDB\": \"physical\", \"StringEvidence\": \"database:0.896,textmining:0.679,coexpression:0.584,experiments:0.216\", \"String_Score_Physical\": 921, \"String_Score_Combined\": 989}", "selected": false}, "selected": false}]}}, "MyList_PPIColorByCluster": {"format_version": "1.0", "generated_by": "cytoscape-3.9.1", "target_cytoscapejs_version": "~2.1", "data": {"shared_name": "MyList_PPIColorByCluster", "name": "MyList_PPIColorByCluster", "SUID": 2785255, "__Annotations": [], "selected": true}, "elements": {"nodes": [{"data": {"id": "2786107", "shared_name": "5142379", "MCODE_SCORE": 0.0, "Gene": "5142379", "Symbol": "MIN8", "name": "5142379", "MCODE_TYPE": "", "SUID": 2786107, "DEGREE": 3.0, "id_original": "5142379", "MCODE_CLUSTER_ID": 0.0, "selected": false, "canonicalName": "MIN8"}, "position": {"x": 75.16728353947099, "y": 406.61599837497187}, "selected": false}, {"data": {"id": "2786104", "shared_name": "3799970", "MCODE_SCORE": 9.625, "Gene": "3799970", "Symbol": "COX26", "name": "3799970", "MCODE_TYPE": "Clustered", "SUID": 2786104, "DEGREE": 16.0, "id_original": "3799970", "MCODE_CLUSTER_ID": 1.0, "selected": false, "canonicalName": "COX26"}, "position": {"x": 361.3916602179378, "y": 691.324753868136}, "selected": false}, {"data": {"id": "2786101", "shared_name": "1466480", "MCODE_SCORE": 9.625, "Gene": "1466480", "Symbol": "MCO10", "name": "1466480", "MCODE_TYPE": "Clustered", "SUID": 2786101, "DEGREE": 14.0, "id_original": "1466480", "MCODE_CLUSTER_ID": 1.0, "selected": false, "canonicalName": "MCO10"}, "position": {"x": 187.71299887150224, "y": 503.6469889755578}, "selected": false}, {"data": {"id": "2786098", "shared_name": "856903", "MCODE_SCORE": 0.0, "Gene": "856903", "Symbol": "YER158C", "name": "856903", "MCODE_TYPE": "", "SUID": 2786098, "DEGREE": 1.0, "id_original": "856903", "MCODE_CLUSTER_ID": 0.0, "selected": false, "canonicalName": "YER158C"}, "position": {"x": -17.58831643611495, "y": 189.1286030884118}, "selected": false}, {"data": {"id": "2786095", "shared_name": "856835", "MCODE_SCORE": 0.0, "Gene": "856835", "Symbol": "UBP9", "name": "856835", "MCODE_TYPE": "", "SUID": 2786095, "DEGREE": 3.0, "id_original": "856835", "MCODE_CLUSTER_ID": 0.0, "selected": false, "canonicalName": "UBP9"}, "position": {"x": 518.805955414471, "y": 109.78716384128046}, "selected": false}, {"data": {"id": "2786092", "shared_name": "856803", "MCODE_SCORE": 0.0, "Gene": "856803", "Symbol": "VTC1", "name": "856803", "MCODE_TYPE": "", "SUID": 2786092, "DEGREE": 2.0, "id_original": "856803", "MCODE_CLUSTER_ID": 0.0, "selected": false, "canonicalName": "VTC1"}, "position": {"x": 156.78915548771317, "y": -125.79048050686407}, "selected": false}, {"data": {"id": "2786089", "shared_name": "856797", "MCODE_SCORE": 0.0, "Gene": "856797", "Symbol": "RGI1", "name": "856797", "MCODE_TYPE": "", "SUID": 2786089, "DEGREE": 2.0, "id_original": "856797", "MCODE_CLUSTER_ID": 0.0, "selected": false, "canonicalName": "RGI1"}, "position": {"x": 110.05690145939286, "y": 291.1855326767297}, "selected": false}, {"data": {"id": "2786086", "shared_name": "856794", "MCODE_SCORE": 0.0, "Gene": "856794", "Symbol": "ICL1", "name": "856794", "MCODE_TYPE": "", "SUID": 2786086, "DEGREE": 8.0, "id_original": "856794", "MCODE_CLUSTER_ID": 0.0, "selected": false, "canonicalName": "ICL1"}, "position": {"x": 194.38929892033036, "y": 506.94572554782343}, "selected": false}, {"data": {"id": "2786083", "shared_name": "856791", "MCODE_SCORE": 0.0, "Gene": "856791", "Symbol": "GPP2", "name": "856791", "MCODE_TYPE": "", "SUID": 2786083, "DEGREE": 3.0, "id_original": "856791", "MCODE_CLUSTER_ID": 0.0, "selected": false, "canonicalName": "GPP2"}, "position": {"x": -190.4042954400212, "y": 359.679749595675}, "selected": false}, {"data": {"id": "2786080", "shared_name": "856779", "MCODE_SCORE": 0.0, "Gene": "856779", "Symbol": "PIC2", "name": "856779", "MCODE_TYPE": "", "SUID": 2786080, "DEGREE": 7.0, "id_original": "856779", "MCODE_CLUSTER_ID": 0.0, "selected": false, "canonicalName": "PIC2"}, "position": {"x": 392.5011305853694, "y": 399.1734782333703}, "selected": false}, {"data": {"id": "2786077", "shared_name": "856755", "MCODE_SCORE": 0.0, "Gene": "856755", "Symbol": "ZRG8", "name": "856755", "MCODE_TYPE": "", "SUID": 2786077, "DEGREE": 1.0, "id_original": "856755", "MCODE_CLUSTER_ID": 0.0, "selected": false, "canonicalName": "ZRG8"}, "position": {"x": 543.5651717230647, "y": 452.66887007907343}, "selected": false}, {"data": {"id": "2786074", "shared_name": "856741", "MCODE_SCORE": 2.9166666666666665, "Gene": "856741", "Symbol": "GPA2", "name": "856741", "MCODE_TYPE": "Clustered", "SUID": 2786074, "DEGREE": 26.0, "id_original": "856741", "MCODE_CLUSTER_ID": 2.0, "selected": false, "canonicalName": "GPA2"}, "position": {"x": 414.8382353827327, "y": 268.22134505465937}, "selected": false}, {"data": {"id": "2786071", "shared_name": "856705", "MCODE_SCORE": 0.0, "Gene": "856705", "Symbol": "GLC3", "name": "856705", "MCODE_TYPE": "", "SUID": 2786071, "DEGREE": 5.0, "id_original": "856705", "MCODE_CLUSTER_ID": 0.0, "selected": false, "canonicalName": "GLC3"}, "position": {"x": 329.0980935141414, "y": 71.44430648044062}, "selected": false}, {"data": {"id": "2786068", "shared_name": "856704", "MCODE_SCORE": 0.0, "Gene": "856704", "Symbol": "UBC8", "name": "856704", "MCODE_TYPE": "", "SUID": 2786068, "DEGREE": 2.0, "id_original": "856704", "MCODE_CLUSTER_ID": 0.0, "selected": false, "canonicalName": "UBC8"}, "position": {"x": 39.012238983806924, "y": 343.20510970309687}, "selected": false}, {"data": {"id": "2786065", "shared_name": "856689", "MCODE_SCORE": 9.625, "Gene": "856689", "Symbol": "RIP1", "name": "856689", "MCODE_TYPE": "Clustered", "SUID": 2786065, "DEGREE": 37.0, "id_original": "856689", "MCODE_CLUSTER_ID": 1.0, "selected": false, "canonicalName": "RIP1"}, "position": {"x": 344.7310576483577, "y": 574.2628947372766}, "selected": false}, {"data": {"id": "2786062", "shared_name": "856672", "MCODE_SCORE": 9.625, "Gene": "856672", "Symbol": "CYC7", "name": "856672", "MCODE_TYPE": "Clustered", "SUID": 2786062, "DEGREE": 23.0, "id_original": "856672", "MCODE_CLUSTER_ID": 1.0, "selected": false, "canonicalName": "CYC7"}, "position": {"x": 247.00377035587724, "y": 646.9455729599329}, "selected": false}, {"data": {"id": "2786059", "shared_name": "856670", "MCODE_SCORE": 0.0, "Gene": "856670", "Symbol": "YEF1", "name": "856670", "MCODE_TYPE": "", "SUID": 2786059, "DEGREE": 4.0, "id_original": "856670", "MCODE_CLUSTER_ID": 0.0, "selected": false, "canonicalName": "YEF1"}, "position": {"x": -11.488249297443076, "y": 465.87907515719843}, "selected": false}, {"data": {"id": "2786056", "shared_name": "856581", "MCODE_SCORE": 0.0, "Gene": "856581", "Symbol": "FMO1", "name": "856581", "MCODE_TYPE": "", "SUID": 2786056, "DEGREE": 1.0, "id_original": "856581", "MCODE_CLUSTER_ID": 0.0, "selected": false, "canonicalName": "FMO1"}, "position": {"x": 43.529206757244424, "y": 54.687226402315616}, "selected": false}, {"data": {"id": "2786053", "shared_name": "856540", "MCODE_SCORE": 0.0, "Gene": "856540", "Symbol": "YHR138C", "name": "856540", "MCODE_TYPE": "", "SUID": 2786053, "DEGREE": 1.0, "id_original": "856540", "MCODE_CLUSTER_ID": 0.0, "selected": false, "canonicalName": "YHR138C"}, "position": {"x": 1142.6048598334164, "y": -252.84245194241095}, "selected": false}, {"data": {"id": "2786050", "shared_name": "856539", "MCODE_SCORE": 0.0, "Gene": "856539", "Symbol": "ARO9", "name": "856539", "MCODE_TYPE": "", "SUID": 2786050, "DEGREE": 1.0, "id_original": "856539", "MCODE_CLUSTER_ID": 0.0, "selected": false, "canonicalName": "ARO9"}, "position": {"x": 37.3178725287288, "y": 778.3067484970422}, "selected": false}, {"data": {"id": "2786047", "shared_name": "856538", "MCODE_SCORE": 0.0, "Gene": "856538", "Symbol": "SPL2", "name": "856538", "MCODE_TYPE": "", "SUID": 2786047, "DEGREE": 1.0, "id_original": "856538", "MCODE_CLUSTER_ID": 0.0, "selected": false, "canonicalName": "SPL2"}, "position": {"x": -149.6854233697087, "y": 396.6902934189172}, "selected": false}, {"data": {"id": "2786044", "shared_name": "856497", "MCODE_SCORE": 0.0, "Gene": "856497", "Symbol": "PAL2", "name": "856497", "MCODE_TYPE": "", "SUID": 2786044, "DEGREE": 3.0, "id_original": "856497", "MCODE_CLUSTER_ID": 0.0, "selected": false, "canonicalName": "PAL2"}, "position": {"x": 133.45010519474442, "y": 341.27909957126093}, "selected": false}, {"data": {"id": "2786041", "shared_name": "856492", "MCODE_SCORE": 0.0, "Gene": "856492", "Symbol": "HXT4", "name": "856492", "MCODE_TYPE": "", "SUID": 2786041, "DEGREE": 3.0, "id_original": "856492", "MCODE_CLUSTER_ID": 0.0, "selected": false, "canonicalName": "HXT4"}, "position": {"x": 58.064546112713174, "y": 113.15563307956171}, "selected": false}, {"data": {"id": "2786038", "shared_name": "856487", "MCODE_SCORE": 0.0, "Gene": "856487", "Symbol": "RTC3", "name": "856487", "MCODE_TYPE": "", "SUID": 2786038, "DEGREE": 1.0, "id_original": "856487", "MCODE_CLUSTER_ID": 0.0, "selected": false, "canonicalName": "RTC3"}, "position": {"x": 580.0993666693538, "y": 657.2011881943079}, "selected": false}, {"data": {"id": "2786035", "shared_name": "856409", "MCODE_SCORE": 0.0, "Gene": "856409", "Symbol": "YSC84", "name": "856409", "MCODE_TYPE": "", "SUID": 2786035, "DEGREE": 3.0, "id_original": "856409", "MCODE_CLUSTER_ID": 0.0, "selected": false, "canonicalName": "YSC84"}, "position": {"x": 252.6834273382991, "y": 912.7999735946985}, "selected": false}, {"data": {"id": "2786032", "shared_name": "856391", "MCODE_SCORE": 0.0, "Gene": "856391", "Symbol": "LEU5", "name": "856391", "MCODE_TYPE": "", "SUID": 2786032, "DEGREE": 2.0, "id_original": "856391", "MCODE_CLUSTER_ID": 0.0, "selected": false, "canonicalName": "LEU5"}, "position": {"x": 61.51422262638505, "y": 825.165330040011}, "selected": false}, {"data": {"id": "2786029", "shared_name": "856390", "MCODE_SCORE": 9.625, "Gene": "856390", "Symbol": "QCR10", "name": "856390", "MCODE_TYPE": "Clustered", "SUID": 2786029, "DEGREE": 24.0, "id_original": "856390", "MCODE_CLUSTER_ID": 1.0, "selected": false, "canonicalName": "QCR10"}, "position": {"x": 314.6961054846614, "y": 623.7472392196985}, "selected": false}, {"data": {"id": "2786026", "shared_name": "856361", "MCODE_SCORE": 0.0, "Gene": "856361", "Symbol": "RIM4", "name": "856361", "MCODE_TYPE": "", "SUID": 2786026, "DEGREE": 2.0, "id_original": "856361", "MCODE_CLUSTER_ID": 0.0, "selected": false, "canonicalName": "RIM4"}, "position": {"x": 118.16571188419755, "y": 46.93054305270624}, "selected": false}, {"data": {"id": "2786023", "shared_name": "856349", "MCODE_SCORE": 0.0, "Gene": "856349", "Symbol": "MUP3", "name": "856349", "MCODE_TYPE": "", "SUID": 2786023, "DEGREE": 1.0, "id_original": "856349", "MCODE_CLUSTER_ID": 0.0, "selected": false, "canonicalName": "MUP3"}, "position": {"x": 92.2379164740413, "y": 173.4053469772668}, "selected": false}, {"data": {"id": "2786020", "shared_name": "856321", "MCODE_SCORE": 9.625, "Gene": "856321", "Symbol": "QCR2", "name": "856321", "MCODE_TYPE": "Clustered", "SUID": 2786020, "DEGREE": 39.0, "id_original": "856321", "MCODE_CLUSTER_ID": 1.0, "selected": false, "canonicalName": "QCR2"}, "position": {"x": 284.1561408087581, "y": 543.007126915011}, "selected": false}, {"data": {"id": "2786017", "shared_name": "856289", "MCODE_SCORE": 0.0, "Gene": "856289", "Symbol": "GPH1", "name": "856289", "MCODE_TYPE": "", "SUID": 2786017, "DEGREE": 7.0, "id_original": "856289", "MCODE_CLUSTER_ID": 0.0, "selected": false, "canonicalName": "GPH1"}, "position": {"x": 386.5010771796077, "y": 184.88113032535028}, "selected": false}, {"data": {"id": "2786014", "shared_name": "856280", "MCODE_SCORE": 0.0, "Gene": "856280", "Symbol": "TDA6", "name": "856280", "MCODE_TYPE": "", "SUID": 2786014, "DEGREE": 1.0, "id_original": "856280", "MCODE_CLUSTER_ID": 0.0, "selected": false, "canonicalName": "TDA6"}, "position": {"x": 664.8329787299007, "y": 370.85879622653437}, "selected": false}, {"data": {"id": "2786011", "shared_name": "856159", "MCODE_SCORE": 0.0, "Gene": "856159", "Symbol": "MCM16", "name": "856159", "MCODE_TYPE": "", "SUID": 2786011, "DEGREE": 5.0, "id_original": "856159", "MCODE_CLUSTER_ID": 0.0, "selected": false, "canonicalName": "MCM16"}, "position": {"x": 732.7644057318538, "y": 163.80636321261835}, "selected": false}, {"data": {"id": "2786008", "shared_name": "856142", "MCODE_SCORE": 0.0, "Gene": "856142", "Symbol": "CSR2", "name": "856142", "MCODE_TYPE": "", "SUID": 2786008, "DEGREE": 3.0, "id_original": "856142", "MCODE_CLUSTER_ID": 0.0, "selected": false, "canonicalName": "CSR2"}, "position": {"x": -55.097258086505576, "y": 31.58293257907343}, "selected": false}, {"data": {"id": "2786005", "shared_name": "856131", "MCODE_SCORE": 9.625, "Gene": "856131", "Symbol": "ATP20", "name": "856131", "MCODE_TYPE": "Clustered", "SUID": 2786005, "DEGREE": 34.0, "id_original": "856131", "MCODE_CLUSTER_ID": 1.0, "selected": false, "canonicalName": "ATP20"}, "position": {"x": 225.8686766669124, "y": 584.4204264755579}, "selected": false}, {"data": {"id": "2786002", "shared_name": "856103", "MCODE_SCORE": 0.0, "Gene": "856103", "Symbol": "LSP1", "name": "856103", "MCODE_TYPE": "", "SUID": 2786002, "DEGREE": 5.0, "id_original": "856103", "MCODE_CLUSTER_ID": 0.0, "selected": false, "canonicalName": "LSP1"}, "position": {"x": 405.99672079533036, "y": 250.99524794772577}, "selected": false}, {"data": {"id": "2785999", "shared_name": "856092", "MCODE_SCORE": 0.0, "Gene": "856092", "Symbol": "HST2", "name": "856092", "MCODE_TYPE": "", "SUID": 2785999, "DEGREE": 5.0, "id_original": "856092", "MCODE_CLUSTER_ID": 0.0, "selected": false, "canonicalName": "HST2"}, "position": {"x": 278.6364340826839, "y": 374.7557231064172}, "selected": false}, {"data": {"id": "2785996", "shared_name": "856089", "MCODE_SCORE": 0.0, "Gene": "856089", "Symbol": "CTF19", "name": "856089", "MCODE_TYPE": "", "SUID": 2785996, "DEGREE": 3.0, "id_original": "856089", "MCODE_CLUSTER_ID": 0.0, "selected": false, "canonicalName": "CTF19"}, "position": {"x": 924.1461195990413, "y": 179.80577765658808}, "selected": false}, {"data": {"id": "2785993", "shared_name": "856088", "MCODE_SCORE": 1.0, "Gene": "856088", "Symbol": "VTC3", "name": "856088", "MCODE_TYPE": "Clustered", "SUID": 2785993, "DEGREE": 6.0, "id_original": "856088", "MCODE_CLUSTER_ID": 11.0, "selected": false, "canonicalName": "VTC3"}, "position": {"x": 250.72834158390458, "y": 62.516923057589054}, "selected": false}, {"data": {"id": "2785990", "shared_name": "856083", "MCODE_SCORE": 0.0, "Gene": "856083", "Symbol": "RMI1", "name": "856083", "MCODE_TYPE": "", "SUID": 2785990, "DEGREE": 1.0, "id_original": "856083", "MCODE_CLUSTER_ID": 0.0, "selected": false, "canonicalName": "RMI1"}, "position": {"x": 471.0090651556819, "y": -61.299345863309384}, "selected": false}, {"data": {"id": "2785987", "shared_name": "856027", "MCODE_SCORE": 9.625, "Gene": "856027", "Symbol": "ATP4", "name": "856027", "MCODE_TYPE": "Clustered", "SUID": 2785987, "DEGREE": 41.0, "id_original": "856027", "MCODE_CLUSTER_ID": 1.0, "selected": false, "canonicalName": "ATP4"}, "position": {"x": 329.5732092902034, "y": 518.249558555636}, "selected": false}, {"data": {"id": "2785984", "shared_name": "856018", "MCODE_SCORE": 0.0, "Gene": "856018", "Symbol": "YDC1", "name": "856018", "MCODE_TYPE": "", "SUID": 2785984, "DEGREE": 1.0, "id_original": "856018", "MCODE_CLUSTER_ID": 0.0, "selected": false, "canonicalName": "YDC1"}, "position": {"x": 1244.9264502570004, "y": -252.84245194241095}, "selected": false}, {"data": {"id": "2785981", "shared_name": "855980", "MCODE_SCORE": 0.0, "Gene": "855980", "Symbol": "RNY1", "name": "855980", "MCODE_TYPE": "", "SUID": 2785981, "DEGREE": 3.0, "id_original": "855980", "MCODE_CLUSTER_ID": 0.0, "selected": false, "canonicalName": "RNY1"}, "position": {"x": 541.6749129340022, "y": 205.2896014328332}, "selected": false}, {"data": {"id": "2785978", "shared_name": "855969", "MCODE_SCORE": 0.0, "Gene": "855969", "Symbol": "ODC1", "name": "855969", "MCODE_TYPE": "", "SUID": 2785978, "DEGREE": 2.0, "id_original": "855969", "MCODE_CLUSTER_ID": 0.0, "selected": false, "canonicalName": "ODC1"}, "position": {"x": 139.1415572211116, "y": 811.816941368136}, "selected": false}, {"data": {"id": "2785975", "shared_name": "855949", "MCODE_SCORE": 0.0, "Gene": "855949", "Symbol": "PEP4", "name": "855949", "MCODE_TYPE": "", "SUID": 2785975, "DEGREE": 4.0, "id_original": "855949", "MCODE_CLUSTER_ID": 0.0, "selected": false, "canonicalName": "PEP4"}, "position": {"x": 5.362763886150674, "y": 634.149460899386}, "selected": false}, {"data": {"id": "2785972", "shared_name": "855898", "MCODE_SCORE": 1.5, "Gene": "855898", "Symbol": "TPK2", "name": "855898", "MCODE_TYPE": "Seed", "SUID": 2785972, "DEGREE": 26.0, "id_original": "855898", "MCODE_CLUSTER_ID": 5.0, "selected": false, "canonicalName": "TPK2"}, "position": {"x": 351.6881995245784, "y": 99.84351454928827}, "selected": false}, {"data": {"id": "2785969", "shared_name": "855879", "MCODE_SCORE": 0.0, "Gene": "855879", "Symbol": "FMP40", "name": "855879", "MCODE_TYPE": "", "SUID": 2785969, "DEGREE": 1.0, "id_original": "855879", "MCODE_CLUSTER_ID": 0.0, "selected": false, "canonicalName": "FMP40"}, "position": {"x": 1347.2480406805844, "y": -252.84245194241095}, "selected": false}, {"data": {"id": "2785966", "shared_name": "855871", "MCODE_SCORE": 1.4000000000000001, "Gene": "855871", "Symbol": "USV1", "name": "855871", "MCODE_TYPE": "Clustered", "SUID": 2785966, "DEGREE": 7.0, "id_original": "855871", "MCODE_CLUSTER_ID": 4.0, "selected": false, "canonicalName": "USV1"}, "position": {"x": 480.2832350775569, "y": 139.09853469088984}, "selected": false}, {"data": {"id": "2785963", "shared_name": "855857", "MCODE_SCORE": 2.375, "Gene": "855857", "Symbol": "ATP15", "name": "855857", "MCODE_TYPE": "Clustered", "SUID": 2785963, "DEGREE": 25.0, "id_original": "855857", "MCODE_CLUSTER_ID": 3.0, "selected": false, "canonicalName": "ATP15"}, "position": {"x": 250.58192587345536, "y": 528.5603190536829}, "selected": false}, {"data": {"id": "2785960", "shared_name": "855836", "MCODE_SCORE": 0.0, "Gene": "855836", "Symbol": "HSP82", "name": "855836", "MCODE_TYPE": "", "SUID": 2785960, "DEGREE": 45.0, "id_original": "855836", "MCODE_CLUSTER_ID": 0.0, "selected": false, "canonicalName": "HSP82"}, "position": {"x": 206.87358999699052, "y": 217.71516524508905}, "selected": false}, {"data": {"id": "2785957", "shared_name": "855829", "MCODE_SCORE": 0.0, "Gene": "855829", "Symbol": "YPL247C", "name": "855829", "MCODE_TYPE": "", "SUID": 2785957, "DEGREE": 4.0, "id_original": "855829", "MCODE_CLUSTER_ID": 0.0, "selected": false, "canonicalName": "YPL247C"}, "position": {"x": 394.3486113592952, "y": 209.9870997543664}, "selected": false}, {"data": {"id": "2785954", "shared_name": "855770", "MCODE_SCORE": 0.0, "Gene": "855770", "Symbol": "EGO4", "name": "855770", "MCODE_TYPE": "", "SUID": 2785954, "DEGREE": 1.0, "id_original": "855770", "MCODE_CLUSTER_ID": 0.0, "selected": false, "canonicalName": "EGO4"}, "position": {"x": 35.78717184513505, "y": 216.1465121383996}, "selected": false}, {"data": {"id": "2785951", "shared_name": "855717", "MCODE_SCORE": 0.0, "Gene": "855717", "Symbol": "PBI2", "name": "855717", "MCODE_TYPE": "", "SUID": 2785951, "DEGREE": 1.0, "id_original": "855717", "MCODE_CLUSTER_ID": 0.0, "selected": false, "canonicalName": "PBI2"}, "position": {"x": 1144.9264502570004, "y": -360.5818013076453}, "selected": false}, {"data": {"id": "2785948", "shared_name": "855675", "MCODE_SCORE": 9.625, "Gene": "855675", "Symbol": "COX5A", "name": "855675", "MCODE_TYPE": "Clustered", "SUID": 2785948, "DEGREE": 31.0, "id_original": "855675", "MCODE_CLUSTER_ID": 1.0, "selected": false, "canonicalName": "COX5A"}, "position": {"x": 389.8851332709163, "y": 544.4878703232141}, "selected": false}, {"data": {"id": "2785945", "shared_name": "855669", "MCODE_SCORE": 0.0, "Gene": "855669", "Symbol": "POR1", "name": "855669", "MCODE_TYPE": "", "SUID": 2785945, "DEGREE": 23.0, "id_original": "855669", "MCODE_CLUSTER_ID": 0.0, "selected": false, "canonicalName": "POR1"}, "position": {"x": 165.0789809271663, "y": 607.4974528427454}, "selected": false}, {"data": {"id": "2785942", "shared_name": "855623", "MCODE_SCORE": 0.0, "Gene": "855623", "Symbol": "MIC27", "name": "855623", "MCODE_TYPE": "", "SUID": 2785942, "DEGREE": 2.0, "id_original": "855623", "MCODE_CLUSTER_ID": 0.0, "selected": false, "canonicalName": "MIC27"}, "position": {"x": 380.7753691717952, "y": 851.231980430636}, "selected": false}, {"data": {"id": "2785939", "shared_name": "855562", "MCODE_SCORE": 0.0, "Gene": "855562", "Symbol": "YGP1", "name": "855562", "MCODE_TYPE": "", "SUID": 2785939, "DEGREE": 1.0, "id_original": "855562", "MCODE_CLUSTER_ID": 0.0, "selected": false, "canonicalName": "YGP1"}, "position": {"x": 242.51222372501786, "y": 784.5178080673547}, "selected": false}, {"data": {"id": "2785936", "shared_name": "855521", "MCODE_SCORE": 0.0, "Gene": "855521", "Symbol": "NNR1", "name": "855521", "MCODE_TYPE": "", "SUID": 2785936, "DEGREE": 1.0, "id_original": "855521", "MCODE_CLUSTER_ID": 0.0, "selected": false, "canonicalName": "NNR1"}, "position": {"x": 584.8294386908382, "y": 12.013383018526554}, "selected": false}, {"data": {"id": "2785933", "shared_name": "855447", "MCODE_SCORE": 1.0, "Gene": "855447", "Symbol": "GOR1", "name": "855447", "MCODE_TYPE": "Seed", "SUID": 2785933, "DEGREE": 5.0, "id_original": "855447", "MCODE_CLUSTER_ID": 8.0, "selected": false, "canonicalName": "GOR1"}, "position": {"x": 85.17692709415849, "y": 457.4221354599328}, "selected": false}, {"data": {"id": "2785930", "shared_name": "855363", "MCODE_SCORE": 0.0, "Gene": "855363", "Symbol": "YMR315W", "name": "855363", "MCODE_TYPE": "", "SUID": 2785930, "DEGREE": 2.0, "id_original": "855363", "MCODE_CLUSTER_ID": 0.0, "selected": false, "canonicalName": "YMR315W"}, "position": {"x": -138.8593735650212, "y": 685.7672282333704}, "selected": false}, {"data": {"id": "2785927", "shared_name": "855328", "MCODE_SCORE": 0.0, "Gene": "855328", "Symbol": "YKU70", "name": "855328", "MCODE_TYPE": "", "SUID": 2785927, "DEGREE": 4.0, "id_original": "855328", "MCODE_CLUSTER_ID": 0.0, "selected": false, "canonicalName": "YKU70"}, "position": {"x": 307.36480951755937, "y": 342.34782515719843}, "selected": false}, {"data": {"id": "2785924", "shared_name": "855303", "MCODE_SCORE": 2.9166666666666665, "Gene": "855303", "Symbol": "TPS3", "name": "855303", "MCODE_TYPE": "Clustered", "SUID": 2785924, "DEGREE": 11.0, "id_original": "855303", "MCODE_CLUSTER_ID": 2.0, "selected": false, "canonicalName": "TPS3"}, "position": {"x": 292.71406317203935, "y": 112.15761672213984}, "selected": false}, {"data": {"id": "2785921", "shared_name": "855298", "MCODE_SCORE": 9.625, "Gene": "855298", "Symbol": "COX7", "name": "855298", "MCODE_TYPE": "Clustered", "SUID": 2785921, "DEGREE": 18.0, "id_original": "855298", "MCODE_CLUSTER_ID": 1.0, "selected": false, "canonicalName": "COX7"}, "position": {"x": 472.3854537054866, "y": 505.56413375094843}, "selected": false}, {"data": {"id": "2785918", "shared_name": "855291", "MCODE_SCORE": 0.0, "Gene": "855291", "Symbol": "GAD1", "name": "855291", "MCODE_TYPE": "", "SUID": 2785918, "DEGREE": 7.0, "id_original": "855291", "MCODE_CLUSTER_ID": 0.0, "selected": false, "canonicalName": "GAD1"}, "position": {"x": 157.36683798283036, "y": 677.0294962997766}, "selected": false}, {"data": {"id": "2785915", "shared_name": "855246", "MCODE_SCORE": 0.0, "Gene": "855246", "Symbol": "YMR206W", "name": "855246", "MCODE_TYPE": "", "SUID": 2785915, "DEGREE": 1.0, "id_original": "855246", "MCODE_CLUSTER_ID": 0.0, "selected": false, "canonicalName": "YMR206W"}, "position": {"x": 1449.5696311041684, "y": -252.84245194241095}, "selected": false}, {"data": {"id": "2785912", "shared_name": "855205", "MCODE_SCORE": 0.0, "Gene": "855205", "Symbol": "ALD3", "name": "855205", "MCODE_TYPE": "", "SUID": 2785912, "DEGREE": 5.0, "id_original": "855205", "MCODE_CLUSTER_ID": 0.0, "selected": false, "canonicalName": "ALD3"}, "position": {"x": 29.758576874431924, "y": 649.4557353134485}, "selected": false}, {"data": {"id": "2785909", "shared_name": "855137", "MCODE_SCORE": 0.0, "Gene": "855137", "Symbol": "HFD1", "name": "855137", "MCODE_TYPE": "", "SUID": 2785909, "DEGREE": 13.0, "id_original": "855137", "MCODE_CLUSTER_ID": 0.0, "selected": false, "canonicalName": "HFD1"}, "position": {"x": 99.89129782169755, "y": 536.8683940048547}, "selected": false}, {"data": {"id": "2785906", "shared_name": "855131", "MCODE_SCORE": 0.0, "Gene": "855131", "Symbol": "PGM2", "name": "855131", "MCODE_TYPE": "", "SUID": 2785906, "DEGREE": 9.0, "id_original": "855131", "MCODE_CLUSTER_ID": 0.0, "selected": false, "canonicalName": "PGM2"}, "position": {"x": 220.53020620792802, "y": 314.8045817489953}, "selected": false}, {"data": {"id": "2785903", "shared_name": "855130", "MCODE_SCORE": 1.5, "Gene": "855130", "Symbol": "YPK2", "name": "855130", "MCODE_TYPE": "Clustered", "SUID": 2785903, "DEGREE": 12.0, "id_original": "855130", "MCODE_CLUSTER_ID": 5.0, "selected": false, "canonicalName": "YPK2"}, "position": {"x": 438.73387289493974, "y": 246.74046431735468}, "selected": false}, {"data": {"id": "2785900", "shared_name": "855105", "MCODE_SCORE": 0.0, "Gene": "855105", "Symbol": "ISF1", "name": "855105", "MCODE_TYPE": "", "SUID": 2785900, "DEGREE": 3.0, "id_original": "855105", "MCODE_CLUSTER_ID": 0.0, "selected": false, "canonicalName": "ISF1"}, "position": {"x": 426.1632323309749, "y": -21.49066056057501}, "selected": false}, {"data": {"id": "2785897", "shared_name": "855078", "MCODE_SCORE": 0.0, "Gene": "855078", "Symbol": "AAC1", "name": "855078", "MCODE_TYPE": "", "SUID": 2785897, "DEGREE": 10.0, "id_original": "855078", "MCODE_CLUSTER_ID": 0.0, "selected": false, "canonicalName": "AAC1"}, "position": {"x": 155.31997824161942, "y": 567.9643412704797}, "selected": false}, {"data": {"id": "2785894", "shared_name": "855051", "MCODE_SCORE": 0.0, "Gene": "855051", "Symbol": "IMP2", "name": "855051", "MCODE_TYPE": "", "SUID": 2785894, "DEGREE": 2.0, "id_original": "855051", "MCODE_CLUSTER_ID": 0.0, "selected": false, "canonicalName": "IMP2"}, "position": {"x": 30.737275604900674, "y": 492.0849772568078}, "selected": false}, {"data": {"id": "2785891", "shared_name": "855020", "MCODE_SCORE": 0.0, "Gene": "855020", "Symbol": "PLB1", "name": "855020", "MCODE_TYPE": "", "SUID": 2785891, "DEGREE": 1.0, "id_original": "855020", "MCODE_CLUSTER_ID": 0.0, "selected": false, "canonicalName": "PLB1"}, "position": {"x": 502.26805258243974, "y": -23.801298988309384}, "selected": false}, {"data": {"id": "2785888", "shared_name": "855014", "MCODE_SCORE": 0.0, "Gene": "855014", "Symbol": "MIX17", "name": "855014", "MCODE_TYPE": "", "SUID": 2785888, "DEGREE": 1.0, "id_original": "855014", "MCODE_CLUSTER_ID": 0.0, "selected": false, "canonicalName": "MIX17"}, "position": {"x": 1551.8912215277524, "y": -252.84245194241095}, "selected": false}, {"data": {"id": "2785885", "shared_name": "854950", "MCODE_SCORE": 0.0, "Gene": "854950", "Symbol": "CYB2", "name": "854950", "MCODE_TYPE": "", "SUID": 2785885, "DEGREE": 12.0, "id_original": "854950", "MCODE_CLUSTER_ID": 0.0, "selected": false, "canonicalName": "CYB2"}, "position": {"x": 75.32211447208817, "y": 484.17253218844843}, "selected": false}, {"data": {"id": "2785882", "shared_name": "854919", "MCODE_SCORE": 9.625, "Gene": "854919", "Symbol": "NDI1", "name": "854919", "MCODE_TYPE": "Clustered", "SUID": 2785882, "DEGREE": 25.0, "id_original": "854919", "MCODE_CLUSTER_ID": 1.0, "selected": false, "canonicalName": "NDI1"}, "position": {"x": 263.188279633221, "y": 468.16218672946405}, "selected": false}, {"data": {"id": "2785879", "shared_name": "854916", "MCODE_SCORE": 0.0, "Gene": "854916", "Symbol": "PHO84", "name": "854916", "MCODE_TYPE": "", "SUID": 2785879, "DEGREE": 5.0, "id_original": "854916", "MCODE_CLUSTER_ID": 0.0, "selected": false, "canonicalName": "PHO84"}, "position": {"x": 0.2356581732600489, "y": 368.63078414157343}, "selected": false}, {"data": {"id": "2785876", "shared_name": "854911", "MCODE_SCORE": 0.0, "Gene": "854911", "Symbol": "MSC1", "name": "854911", "MCODE_TYPE": "", "SUID": 2785876, "DEGREE": 1.0, "id_original": "854911", "MCODE_CLUSTER_ID": 0.0, "selected": false, "canonicalName": "MSC1"}, "position": {"x": -61.4514145806462, "y": 202.33509551242304}, "selected": false}, {"data": {"id": "2785873", "shared_name": "854893", "MCODE_SCORE": 9.625, "Gene": "854893", "Symbol": "ATP18", "name": "854893", "MCODE_TYPE": "Clustered", "SUID": 2785873, "DEGREE": 15.0, "id_original": "854893", "MCODE_CLUSTER_ID": 1.0, "selected": false, "canonicalName": "ATP18"}, "position": {"x": 264.7896971747249, "y": 591.2024088974329}, "selected": false}, {"data": {"id": "2785870", "shared_name": "854872", "MCODE_SCORE": 2.9166666666666665, "Gene": "854872", "Symbol": "TSL1", "name": "854872", "MCODE_TYPE": "Clustered", "SUID": 2785870, "DEGREE": 11.0, "id_original": "854872", "MCODE_CLUSTER_ID": 2.0, "selected": false, "canonicalName": "TSL1"}, "position": {"x": 134.25137472599442, "y": 86.49015914157343}, "selected": false}, {"data": {"id": "2785867", "shared_name": "854857", "MCODE_SCORE": 0.0, "Gene": "854857", "Symbol": "YPS6", "name": "854857", "MCODE_TYPE": "", "SUID": 2785867, "DEGREE": 1.0, "id_original": "854857", "MCODE_CLUSTER_ID": 0.0, "selected": false, "canonicalName": "YPS6"}, "position": {"x": -291.5303940728337, "y": 170.08458911136103}, "selected": false}, {"data": {"id": "2785864", "shared_name": "854853", "MCODE_SCORE": 0.0, "Gene": "854853", "Symbol": "NRE1", "name": "854853", "MCODE_TYPE": "", "SUID": 2785864, "DEGREE": 1.0, "id_original": "854853", "MCODE_CLUSTER_ID": 0.0, "selected": false, "canonicalName": "NRE1"}, "position": {"x": 15.025697235760049, "y": 823.308640586886}, "selected": false}, {"data": {"id": "2785861", "shared_name": "854833", "MCODE_SCORE": 0.0, "Gene": "854833", "Symbol": "YIR016W", "name": "854833", "MCODE_TYPE": "", "SUID": 2785861, "DEGREE": 1.0, "id_original": "854833", "MCODE_CLUSTER_ID": 0.0, "selected": false, "canonicalName": "YIR016W"}, "position": {"x": 225.2276625677913, "y": -140.51884354397345}, "selected": false}, {"data": {"id": "2785858", "shared_name": "854824", "MCODE_SCORE": 0.0, "Gene": "854824", "Symbol": "EGH1", "name": "854824", "MCODE_TYPE": "", "SUID": 2785858, "DEGREE": 1.0, "id_original": "854824", "MCODE_CLUSTER_ID": 0.0, "selected": false, "canonicalName": "EGH1"}, "position": {"x": -211.4704575493962, "y": 502.3228617782922}, "selected": false}, {"data": {"id": "2785855", "shared_name": "854771", "MCODE_SCORE": 0.0, "Gene": "854771", "Symbol": "APQ12", "name": "854771", "MCODE_TYPE": "", "SUID": 2785855, "DEGREE": 2.0, "id_original": "854771", "MCODE_CLUSTER_ID": 0.0, "selected": false, "canonicalName": "APQ12"}, "position": {"x": 863.2238783881038, "y": 90.10793410495234}, "selected": false}, {"data": {"id": "2785852", "shared_name": "854769", "MCODE_SCORE": 0.0, "Gene": "854769", "Symbol": "PKP1", "name": "854769", "MCODE_TYPE": "", "SUID": 2785852, "DEGREE": 4.0, "id_original": "854769", "MCODE_CLUSTER_ID": 0.0, "selected": false, "canonicalName": "PKP1"}, "position": {"x": 438.89549398868974, "y": 378.64712630465937}, "selected": false}, {"data": {"id": "2785849", "shared_name": "854761", "MCODE_SCORE": 0.0, "Gene": "854761", "Symbol": "PCL7", "name": "854761", "MCODE_TYPE": "", "SUID": 2785849, "DEGREE": 1.0, "id_original": "854761", "MCODE_CLUSTER_ID": 0.0, "selected": false, "canonicalName": "PCL7"}, "position": {"x": 485.37152243107255, "y": 34.764542686495304}, "selected": false}, {"data": {"id": "2785846", "shared_name": "854745", "MCODE_SCORE": 0.0, "Gene": "854745", "Symbol": "FIS1", "name": "854745", "MCODE_TYPE": "", "SUID": 2785846, "DEGREE": 11.0, "id_original": "854745", "MCODE_CLUSTER_ID": 0.0, "selected": false, "canonicalName": "FIS1"}, "position": {"x": 109.75917196720536, "y": 578.2838908310266}, "selected": false}, {"data": {"id": "2785843", "shared_name": "854733", "MCODE_SCORE": 0.0, "Gene": "854733", "Symbol": "RCI37", "name": "854733", "MCODE_TYPE": "", "SUID": 2785843, "DEGREE": 2.0, "id_original": "854733", "MCODE_CLUSTER_ID": 0.0, "selected": false, "canonicalName": "RCI37"}, "position": {"x": 396.3661208197444, "y": 749.560349571261}, "selected": false}, {"data": {"id": "2785840", "shared_name": "854710", "MCODE_SCORE": 0.0, "Gene": "854710", "Symbol": "FYV10", "name": "854710", "MCODE_TYPE": "", "SUID": 2785840, "DEGREE": 1.0, "id_original": "854710", "MCODE_CLUSTER_ID": 0.0, "selected": false, "canonicalName": "FYV10"}, "position": {"x": -89.24923562556808, "y": 480.13594161227655}, "selected": false}, {"data": {"id": "2785837", "shared_name": "854706", "MCODE_SCORE": 0.0, "Gene": "854706", "Symbol": "XBP1", "name": "854706", "MCODE_TYPE": "", "SUID": 2785837, "DEGREE": 2.0, "id_original": "854706", "MCODE_CLUSTER_ID": 0.0, "selected": false, "canonicalName": "XBP1"}, "position": {"x": 561.0626997992366, "y": 309.61364852145624}, "selected": false}, {"data": {"id": "2785834", "shared_name": "854695", "MCODE_SCORE": 9.625, "Gene": "854695", "Symbol": "COX5B", "name": "854695", "MCODE_TYPE": "Clustered", "SUID": 2785834, "DEGREE": 28.0, "id_original": "854695", "MCODE_CLUSTER_ID": 1.0, "selected": false, "canonicalName": "COX5B"}, "position": {"x": 254.06166601627763, "y": 576.7765666122766}, "selected": false}, {"data": {"id": "2785831", "shared_name": "854670", "MCODE_SCORE": 0.0, "Gene": "854670", "Symbol": "OM45", "name": "854670", "MCODE_TYPE": "", "SUID": 2785831, "DEGREE": 5.0, "id_original": "854670", "MCODE_CLUSTER_ID": 0.0, "selected": false, "canonicalName": "OM45"}, "position": {"x": 204.26864767521317, "y": 406.89770613864374}, "selected": false}, {"data": {"id": "2785828", "shared_name": "854660", "MCODE_SCORE": 0.0, "Gene": "854660", "Symbol": "ATG32", "name": "854660", "MCODE_TYPE": "", "SUID": 2785828, "DEGREE": 3.0, "id_original": "854660", "MCODE_CLUSTER_ID": 0.0, "selected": false, "canonicalName": "ATG32"}, "position": {"x": 392.3256621405452, "y": 589.4234171982141}, "selected": false}, {"data": {"id": "2785825", "shared_name": "854651", "MCODE_SCORE": 0.0, "Gene": "854651", "Symbol": "GUT2", "name": "854651", "MCODE_TYPE": "", "SUID": 2785825, "DEGREE": 4.0, "id_original": "854651", "MCODE_CLUSTER_ID": 0.0, "selected": false, "canonicalName": "GUT2"}, "position": {"x": 101.7023940131038, "y": 622.9054728622766}, "selected": false}, {"data": {"id": "2785822", "shared_name": "854568", "MCODE_SCORE": 0.0, "Gene": "854568", "Symbol": "PHR1", "name": "854568", "MCODE_TYPE": "", "SUID": 2785822, "DEGREE": 5.0, "id_original": "854568", "MCODE_CLUSTER_ID": 0.0, "selected": false, "canonicalName": "PHR1"}, "position": {"x": 330.0882983252376, "y": 50.563019859346866}, "selected": false}, {"data": {"id": "2785819", "shared_name": "854556", "MCODE_SCORE": 1.0, "Gene": "854556", "Symbol": "ALD4", "name": "854556", "MCODE_TYPE": "Clustered", "SUID": 2785819, "DEGREE": 11.0, "id_original": "854556", "MCODE_CLUSTER_ID": 8.0, "selected": false, "canonicalName": "ALD4"}, "position": {"x": 140.859178070721, "y": 454.62776290133905}, "selected": false}, {"data": {"id": "2785816", "shared_name": "854529", "MCODE_SCORE": 0.0, "Gene": "854529", "Symbol": "PYK2", "name": "854529", "MCODE_TYPE": "", "SUID": 2785816, "DEGREE": 4.0, "id_original": "854529", "MCODE_CLUSTER_ID": 0.0, "selected": false, "canonicalName": "PYK2"}, "position": {"x": 348.1614184424251, "y": 281.45535384372187}, "selected": false}, {"data": {"id": "2785813", "shared_name": "854463", "MCODE_SCORE": 0.0, "Gene": "854463", "Symbol": "YOR289W", "name": "854463", "MCODE_TYPE": "", "SUID": 2785813, "DEGREE": 1.0, "id_original": "854463", "MCODE_CLUSTER_ID": 0.0, "selected": false, "canonicalName": "YOR289W"}, "position": {"x": -31.0404038384587, "y": 246.1338931198449}, "selected": false}, {"data": {"id": "2785810", "shared_name": "854447", "MCODE_SCORE": 1.25, "Gene": "854447", "Symbol": "TPO4", "name": "854447", "MCODE_TYPE": "Clustered", "SUID": 2785810, "DEGREE": 4.0, "id_original": "854447", "MCODE_CLUSTER_ID": 6.0, "selected": false, "canonicalName": "TPO4"}, "position": {"x": 502.1530318304866, "y": 363.00627242282343}, "selected": false}, {"data": {"id": "2785807", "shared_name": "854441", "MCODE_SCORE": 0.0, "Gene": "854441", "Symbol": "HRK1", "name": "854441", "MCODE_TYPE": "", "SUID": 2785807, "DEGREE": 3.0, "id_original": "854441", "MCODE_CLUSTER_ID": 0.0, "selected": false, "canonicalName": "HRK1"}, "position": {"x": 125.62625265568192, "y": 120.99629317477655}, "selected": false}, {"data": {"id": "2785804", "shared_name": "854357", "MCODE_SCORE": 0.0, "Gene": "854357", "Symbol": "GSP2", "name": "854357", "MCODE_TYPE": "", "SUID": 2785804, "DEGREE": 6.0, "id_original": "854357", "MCODE_CLUSTER_ID": 0.0, "selected": false, "canonicalName": "GSP2"}, "position": {"x": 336.3721633002132, "y": 330.88591109469843}, "selected": false}, {"data": {"id": "2785801", "shared_name": "854350", "MCODE_SCORE": 0.0, "Gene": "854350", "Symbol": "GAC1", "name": "854350", "MCODE_TYPE": "", "SUID": 2785801, "DEGREE": 6.0, "id_original": "854350", "MCODE_CLUSTER_ID": 0.0, "selected": false, "canonicalName": "GAC1"}, "position": {"x": 284.4508490607112, "y": -133.30207718655157}, "selected": false}, {"data": {"id": "2785798", "shared_name": "854344", "MCODE_SCORE": 0.0, "Gene": "854344", "Symbol": "DCS2", "name": "854344", "MCODE_TYPE": "", "SUID": 2785798, "DEGREE": 3.0, "id_original": "854344", "MCODE_CLUSTER_ID": 0.0, "selected": false, "canonicalName": "DCS2"}, "position": {"x": 147.62819052189286, "y": 176.849050628695}, "selected": false}, {"data": {"id": "2785795", "shared_name": "854318", "MCODE_SCORE": 0.0, "Gene": "854318", "Symbol": "MDM32", "name": "854318", "MCODE_TYPE": "", "SUID": 2785795, "DEGREE": 2.0, "id_original": "854318", "MCODE_CLUSTER_ID": 0.0, "selected": false, "canonicalName": "MDM32"}, "position": {"x": -29.54131936580245, "y": 142.63480101779413}, "selected": false}, {"data": {"id": "2785792", "shared_name": "854287", "MCODE_SCORE": 0.0, "Gene": "854287", "Symbol": "GCY1", "name": "854287", "MCODE_TYPE": "", "SUID": 2785792, "DEGREE": 1.0, "id_original": "854287", "MCODE_CLUSTER_ID": 0.0, "selected": false, "canonicalName": "GCY1"}, "position": {"x": -309.6434311822087, "y": 407.4508219833703}, "selected": false}, {"data": {"id": "2785789", "shared_name": "854231", "MCODE_SCORE": 9.625, "Gene": "854231", "Symbol": "CYT1", "name": "854231", "MCODE_TYPE": "Seed", "SUID": 2785789, "DEGREE": 39.0, "id_original": "854231", "MCODE_CLUSTER_ID": 1.0, "selected": false, "canonicalName": "CYT1"}, "position": {"x": 283.01254797428544, "y": 587.8630534286829}, "selected": false}, {"data": {"id": "2785786", "shared_name": "854228", "MCODE_SCORE": 1.0, "Gene": "854228", "Symbol": "YOR062C", "name": "854228", "MCODE_TYPE": "Clustered", "SUID": 2785786, "DEGREE": 3.0, "id_original": "854228", "MCODE_CLUSTER_ID": 10.0, "selected": false, "canonicalName": "YOR062C"}, "position": {"x": 189.8512969061702, "y": -303.2690571670203}, "selected": false}, {"data": {"id": "2785783", "shared_name": "854217", "MCODE_SCORE": 0.0, "Gene": "854217", "Symbol": "TMC1", "name": "854217", "MCODE_TYPE": "", "SUID": 2785783, "DEGREE": 1.0, "id_original": "854217", "MCODE_CLUSTER_ID": 0.0, "selected": false, "canonicalName": "TMC1"}, "position": {"x": 538.8434157416194, "y": 387.3557749863}, "selected": false}, {"data": {"id": "2785780", "shared_name": "854109", "MCODE_SCORE": 0.0, "Gene": "854109", "Symbol": "RRT8", "name": "854109", "MCODE_TYPE": "", "SUID": 2785780, "DEGREE": 1.0, "id_original": "854109", "MCODE_CLUSTER_ID": 0.0, "selected": false, "canonicalName": "RRT8"}, "position": {"x": -106.0512680962712, "y": 798.4061747665735}, "selected": false}, {"data": {"id": "2785777", "shared_name": "854104", "MCODE_SCORE": 0.0, "Gene": "854104", "Symbol": "DDR2", "name": "854104", "MCODE_TYPE": "", "SUID": 2785777, "DEGREE": 1.0, "id_original": "854104", "MCODE_CLUSTER_ID": 0.0, "selected": false, "canonicalName": "DDR2"}, "position": {"x": -13.145628447833701, "y": 67.85814009860468}, "selected": false}, {"data": {"id": "2785774", "shared_name": "854077", "MCODE_SCORE": 9.625, "Gene": "854077", "Symbol": "ATP19", "name": "854077", "MCODE_TYPE": "Clustered", "SUID": 2785774, "DEGREE": 14.0, "id_original": "854077", "MCODE_CLUSTER_ID": 1.0, "selected": false, "canonicalName": "ATP19"}, "position": {"x": 277.29137945621903, "y": 612.909470665011}, "selected": false}, {"data": {"id": "2785771", "shared_name": "854071", "MCODE_SCORE": 0.0, "Gene": "854071", "Symbol": "ATG34", "name": "854071", "MCODE_TYPE": "", "SUID": 2785771, "DEGREE": 2.0, "id_original": "854071", "MCODE_CLUSTER_ID": 0.0, "selected": false, "canonicalName": "ATG34"}, "position": {"x": -101.95599221736495, "y": 350.07315169528437}, "selected": false}, {"data": {"id": "2785768", "shared_name": "854010", "MCODE_SCORE": 0.0, "Gene": "854010", "Symbol": "HPF1", "name": "854010", "MCODE_TYPE": "", "SUID": 2785768, "DEGREE": 1.0, "id_original": "854010", "MCODE_CLUSTER_ID": 0.0, "selected": false, "canonicalName": "HPF1"}, "position": {"x": 594.6896987006038, "y": 88.5093165512414}, "selected": false}, {"data": {"id": "2785765", "shared_name": "853994", "MCODE_SCORE": 2.375, "Gene": "853994", "Symbol": "MDH2", "name": "853994", "MCODE_TYPE": "Clustered", "SUID": 2785765, "DEGREE": 17.0, "id_original": "853994", "MCODE_CLUSTER_ID": 3.0, "selected": false, "canonicalName": "MDH2"}, "position": {"x": 260.286309723553, "y": 421.35359297946405}, "selected": false}, {"data": {"id": "2785762", "shared_name": "853973", "MCODE_SCORE": 0.0, "Gene": "853973", "Symbol": "UBP11", "name": "853973", "MCODE_TYPE": "", "SUID": 2785762, "DEGREE": 2.0, "id_original": "853973", "MCODE_CLUSTER_ID": 0.0, "selected": false, "canonicalName": "UBP11"}, "position": {"x": 542.6194777533382, "y": 4.671982872042179}, "selected": false}, {"data": {"id": "2785759", "shared_name": "853951", "MCODE_SCORE": 0.0, "Gene": "853951", "Symbol": "ECM4", "name": "853951", "MCODE_TYPE": "", "SUID": 2785759, "DEGREE": 1.0, "id_original": "853951", "MCODE_CLUSTER_ID": 0.0, "selected": false, "canonicalName": "ECM4"}, "position": {"x": -305.3762192681462, "y": 245.18005477145624}, "selected": false}, {"data": {"id": "2785756", "shared_name": "853950", "MCODE_SCORE": 1.0, "Gene": "853950", "Symbol": "YKR075C", "name": "853950", "MCODE_TYPE": "Clustered", "SUID": 2785756, "DEGREE": 3.0, "id_original": "853950", "MCODE_CLUSTER_ID": 10.0, "selected": false, "canonicalName": "YKR075C"}, "position": {"x": 254.05112219303544, "y": -360.5818013076453}, "selected": false}, {"data": {"id": "2785753", "shared_name": "853932", "MCODE_SCORE": 2.9166666666666665, "Gene": "853932", "Symbol": "GLG1", "name": "853932", "MCODE_TYPE": "Clustered", "SUID": 2785753, "DEGREE": 8.0, "id_original": "853932", "MCODE_CLUSTER_ID": 2.0, "selected": false, "canonicalName": "GLG1"}, "position": {"x": 193.2234892889827, "y": -12.201781166043759}, "selected": false}, {"data": {"id": "2785750", "shared_name": "853920", "MCODE_SCORE": 0.0, "Gene": "853920", "Symbol": "PLN1", "name": "853920", "MCODE_TYPE": "", "SUID": 2785750, "DEGREE": 10.0, "id_original": "853920", "MCODE_CLUSTER_ID": 0.0, "selected": false, "canonicalName": "PLN1"}, "position": {"x": 202.79159689396317, "y": 641.9644633407922}, "selected": false}, {"data": {"id": "2785747", "shared_name": "853853", "MCODE_SCORE": 9.625, "Gene": "853853", "Symbol": "ATP7", "name": "853853", "MCODE_TYPE": "Clustered", "SUID": 2785747, "DEGREE": 33.0, "id_original": "853853", "MCODE_CLUSTER_ID": 1.0, "selected": false, "canonicalName": "ATP7"}, "position": {"x": 361.4865965887874, "y": 574.4352275009485}, "selected": false}, {"data": {"id": "2785744", "shared_name": "853842", "MCODE_SCORE": 0.0, "Gene": "853842", "Symbol": "GPX1", "name": "853842", "MCODE_TYPE": "", "SUID": 2785744, "DEGREE": 1.0, "id_original": "853842", "MCODE_CLUSTER_ID": 0.0, "selected": false, "canonicalName": "GPX1"}, "position": {"x": -173.93368386775558, "y": 260.1571665878625}, "selected": false}, {"data": {"id": "2785741", "shared_name": "853804", "MCODE_SCORE": 0.0, "Gene": "853804", "Symbol": "BLI1", "name": "853804", "MCODE_TYPE": "", "SUID": 2785741, "DEGREE": 3.0, "id_original": "853804", "MCODE_CLUSTER_ID": 0.0, "selected": false, "canonicalName": "BLI1"}, "position": {"x": 508.79249716251786, "y": 59.897614585909366}, "selected": false}, {"data": {"id": "2785738", "shared_name": "853803", "MCODE_SCORE": 1.5, "Gene": "853803", "Symbol": "MSN4", "name": "853803", "MCODE_TYPE": "Clustered", "SUID": 2785738, "DEGREE": 11.0, "id_original": "853803", "MCODE_CLUSTER_ID": 5.0, "selected": false, "canonicalName": "MSN4"}, "position": {"x": 358.55494833439286, "y": 279.75978957370234}, "selected": false}, {"data": {"id": "2785735", "shared_name": "853777", "MCODE_SCORE": 2.375, "Gene": "853777", "Symbol": "MDH1", "name": "853777", "MCODE_TYPE": "Clustered", "SUID": 2785735, "DEGREE": 22.0, "id_original": "853777", "MCODE_CLUSTER_ID": 3.0, "selected": false, "canonicalName": "MDH1"}, "position": {"x": 242.74187612980302, "y": 475.05549727633905}, "selected": false}, {"data": {"id": "2785732", "shared_name": "853769", "MCODE_SCORE": 0.0, "Gene": "853769", "Symbol": "MBR1", "name": "853769", "MCODE_TYPE": "", "SUID": 2785732, "DEGREE": 2.0, "id_original": "853769", "MCODE_CLUSTER_ID": 0.0, "selected": false, "canonicalName": "MBR1"}, "position": {"x": 322.30815744846757, "y": -76.75056351467657}, "selected": false}, {"data": {"id": "2785729", "shared_name": "853751", "MCODE_SCORE": 0.0, "Gene": "853751", "Symbol": "HAP4", "name": "853751", "MCODE_TYPE": "", "SUID": 2785729, "DEGREE": 2.0, "id_original": "853751", "MCODE_CLUSTER_ID": 0.0, "selected": false, "canonicalName": "HAP4"}, "position": {"x": 214.81878042667802, "y": 770.158249961886}, "selected": false}, {"data": {"id": "2785726", "shared_name": "853738", "MCODE_SCORE": 0.0, "Gene": "853738", "Symbol": "DGR2", "name": "853738", "MCODE_TYPE": "", "SUID": 2785726, "DEGREE": 2.0, "id_original": "853738", "MCODE_CLUSTER_ID": 0.0, "selected": false, "canonicalName": "DGR2"}, "position": {"x": 531.584275726971, "y": -167.65495194241095}, "selected": false}, {"data": {"id": "2785723", "shared_name": "853716", "MCODE_SCORE": 9.625, "Gene": "853716", "Symbol": "SDH3", "name": "853716", "MCODE_TYPE": "Clustered", "SUID": 2785723, "DEGREE": 16.0, "id_original": "853716", "MCODE_CLUSTER_ID": 1.0, "selected": false, "canonicalName": "SDH3"}, "position": {"x": 370.3911833807796, "y": 511.16618453219843}, "selected": false}, {"data": {"id": "2785720", "shared_name": "853715", "MCODE_SCORE": 0.0, "Gene": "853715", "Symbol": "MRP8", "name": "853715", "MCODE_TYPE": "", "SUID": 2785720, "DEGREE": 2.0, "id_original": "853715", "MCODE_CLUSTER_ID": 0.0, "selected": false, "canonicalName": "MRP8"}, "position": {"x": 605.7046828314632, "y": 57.65458785251093}, "selected": false}, {"data": {"id": "2785717", "shared_name": "853709", "MCODE_SCORE": 2.375, "Gene": "853709", "Symbol": "SDH1", "name": "853709", "MCODE_TYPE": "Seed", "SUID": 2785717, "DEGREE": 26.0, "id_original": "853709", "MCODE_CLUSTER_ID": 3.0, "selected": false, "canonicalName": "SDH1"}, "position": {"x": 306.6608023688167, "y": 619.310837852511}, "selected": false}, {"data": {"id": "2785714", "shared_name": "853707", "MCODE_SCORE": 0.0, "Gene": "853707", "Symbol": "MCR1", "name": "853707", "MCODE_TYPE": "", "SUID": 2785714, "DEGREE": 3.0, "id_original": "853707", "MCODE_CLUSTER_ID": 0.0, "selected": false, "canonicalName": "MCR1"}, "position": {"x": 302.311157707867, "y": 799.9149638290735}, "selected": false}, {"data": {"id": "2785711", "shared_name": "853706", "MCODE_SCORE": 0.0, "Gene": "853706", "Symbol": "NNR2", "name": "853706", "MCODE_TYPE": "", "SUID": 2785711, "DEGREE": 1.0, "id_original": "853706", "MCODE_CLUSTER_ID": 0.0, "selected": false, "canonicalName": "NNR2"}, "position": {"x": 110.71017599552567, "y": 218.34254180148554}, "selected": false}, {"data": {"id": "2785708", "shared_name": "853663", "MCODE_SCORE": 0.0, "Gene": "853663", "Symbol": "JEN1", "name": "853663", "MCODE_TYPE": "", "SUID": 2785708, "DEGREE": 1.0, "id_original": "853663", "MCODE_CLUSTER_ID": 0.0, "selected": false, "canonicalName": "JEN1"}, "position": {"x": 417.37518454044755, "y": -103.68217362209845}, "selected": false}, {"data": {"id": "2785705", "shared_name": "853585", "MCODE_SCORE": 2.375, "Gene": "853585", "Symbol": "ATP2", "name": "853585", "MCODE_TYPE": "Clustered", "SUID": 2785705, "DEGREE": 40.0, "id_original": "853585", "MCODE_CLUSTER_ID": 3.0, "selected": false, "canonicalName": "ATP2"}, "position": {"x": 268.06007528751786, "y": 487.4350749130578}, "selected": false}, {"data": {"id": "2785702", "shared_name": "853559", "MCODE_SCORE": 0.0, "Gene": "853559", "Symbol": "YJR096W", "name": "853559", "MCODE_TYPE": "", "SUID": 2785702, "DEGREE": 2.0, "id_original": "853559", "MCODE_CLUSTER_ID": 0.0, "selected": false, "canonicalName": "YJR096W"}, "position": {"x": 10.132050995525674, "y": 161.37941466525507}, "selected": false}, {"data": {"id": "2785699", "shared_name": "853540", "MCODE_SCORE": 0.0, "Gene": "853540", "Symbol": "MIR1", "name": "853540", "MCODE_TYPE": "", "SUID": 2785699, "DEGREE": 10.0, "id_original": "853540", "MCODE_CLUSTER_ID": 0.0, "selected": false, "canonicalName": "MIR1"}, "position": {"x": 311.8725314184993, "y": 397.89824019626093}, "selected": false}, {"data": {"id": "2785696", "shared_name": "853522", "MCODE_SCORE": 1.0, "Gene": "853522", "Symbol": "PTK2", "name": "853522", "MCODE_TYPE": "Clustered", "SUID": 2785696, "DEGREE": 16.0, "id_original": "853522", "MCODE_CLUSTER_ID": 7.0, "selected": false, "canonicalName": "PTK2"}, "position": {"x": 389.49937582462724, "y": 32.990555870089054}, "selected": false}, {"data": {"id": "2785693", "shared_name": "853477", "MCODE_SCORE": 0.0, "Gene": "853477", "Symbol": "TES1", "name": "853477", "MCODE_TYPE": "", "SUID": 2785693, "DEGREE": 1.0, "id_original": "853477", "MCODE_CLUSTER_ID": 0.0, "selected": false, "canonicalName": "TES1"}, "position": {"x": 515.1946120306819, "y": 240.0456362839074}, "selected": false}, {"data": {"id": "2785690", "shared_name": "853441", "MCODE_SCORE": 1.0, "Gene": "853441", "Symbol": "VTC4", "name": "853441", "MCODE_TYPE": "Seed", "SUID": 2785690, "DEGREE": 4.0, "id_original": "853441", "MCODE_CLUSTER_ID": 11.0, "selected": false, "canonicalName": "VTC4"}, "position": {"x": 174.4909224554866, "y": 6.795151817354679}, "selected": false}, {"data": {"id": "2785687", "shared_name": "853436", "MCODE_SCORE": 0.0, "Gene": "853436", "Symbol": "TPH3", "name": "853436", "MCODE_TYPE": "", "SUID": 2785687, "DEGREE": 1.0, "id_original": "853436", "MCODE_CLUSTER_ID": 0.0, "selected": false, "canonicalName": "TPH3"}, "position": {"x": 676.1181349799007, "y": 303.63505660251093}, "selected": false}, {"data": {"id": "2785684", "shared_name": "853399", "MCODE_SCORE": 0.0, "Gene": "853399", "Symbol": "UBX6", "name": "853399", "MCODE_TYPE": "", "SUID": 2785684, "DEGREE": 1.0, "id_original": "853399", "MCODE_CLUSTER_ID": 0.0, "selected": false, "canonicalName": "UBX6"}, "position": {"x": 649.6740126654475, "y": -292.17671097561407}, "selected": false}, {"data": {"id": "2785681", "shared_name": "853395", "MCODE_SCORE": 0.0, "Gene": "853395", "Symbol": "TDH1", "name": "853395", "MCODE_TYPE": "", "SUID": 2785681, "DEGREE": 7.0, "id_original": "853395", "MCODE_CLUSTER_ID": 0.0, "selected": false, "canonicalName": "TDH1"}, "position": {"x": 195.82960653751786, "y": 333.2509928818078}, "selected": false}, {"data": {"id": "2785678", "shared_name": "853375", "MCODE_SCORE": 0.0, "Gene": "853375", "Symbol": "YJL070C", "name": "853375", "MCODE_TYPE": "", "SUID": 2785678, "DEGREE": 1.0, "id_original": "853375", "MCODE_CLUSTER_ID": 0.0, "selected": false, "canonicalName": "YJL070C"}, "position": {"x": 477.3569045111507, "y": 228.3431559677453}, "selected": false}, {"data": {"id": "2785675", "shared_name": "853342", "MCODE_SCORE": 0.0, "Gene": "853342", "Symbol": "MEF2", "name": "853342", "MCODE_TYPE": "", "SUID": 2785675, "DEGREE": 6.0, "id_original": "853342", "MCODE_CLUSTER_ID": 0.0, "selected": false, "canonicalName": "MEF2"}, "position": {"x": 369.5940070196956, "y": 265.7846079941125}, "selected": false}, {"data": {"id": "2785672", "shared_name": "853300", "MCODE_SCORE": 0.0, "Gene": "853300", "Symbol": "YAK1", "name": "853300", "MCODE_TYPE": "", "SUID": 2785672, "DEGREE": 14.0, "id_original": "853300", "MCODE_CLUSTER_ID": 0.0, "selected": false, "canonicalName": "YAK1"}, "position": {"x": 246.0076460882991, "y": 280.71016036227655}, "selected": false}, {"data": {"id": "2785669", "shared_name": "853275", "MCODE_SCORE": 1.5, "Gene": "853275", "Symbol": "TPK1", "name": "853275", "MCODE_TYPE": "Clustered", "SUID": 2785669, "DEGREE": 28.0, "id_original": "853275", "MCODE_CLUSTER_ID": 5.0, "selected": false, "canonicalName": "TPK1"}, "position": {"x": 375.02707434147294, "y": 166.34190474704218}, "selected": false}, {"data": {"id": "2785666", "shared_name": "853273", "MCODE_SCORE": 9.625, "Gene": "853273", "Symbol": "QCR8", "name": "853273", "MCODE_TYPE": "Clustered", "SUID": 2785666, "DEGREE": 31.0, "id_original": "853273", "MCODE_CLUSTER_ID": 1.0, "selected": false, "canonicalName": "QCR8"}, "position": {"x": 365.0732626959651, "y": 629.5714579696985}, "selected": false}, {"data": {"id": "2785663", "shared_name": "853240", "MCODE_SCORE": 0.0, "Gene": "853240", "Symbol": "MBB1", "name": "853240", "MCODE_TYPE": "", "SUID": 2785663, "DEGREE": 1.0, "id_original": "853240", "MCODE_CLUSTER_ID": 0.0, "selected": false, "canonicalName": "MBB1"}, "position": {"x": 297.9134879156917, "y": -9.482222450223446}, "selected": false}, {"data": {"id": "2785660", "shared_name": "853219", "MCODE_SCORE": 0.0, "Gene": "853219", "Symbol": "PEX2", "name": "853219", "MCODE_TYPE": "", "SUID": 2785660, "DEGREE": 2.0, "id_original": "853219", "MCODE_CLUSTER_ID": 0.0, "selected": false, "canonicalName": "PEX2"}, "position": {"x": 151.2042708441585, "y": 890.2360697861047}, "selected": false}, {"data": {"id": "2785657", "shared_name": "853165", "MCODE_SCORE": 0.0, "Gene": "853165", "Symbol": "RIE1", "name": "853165", "MCODE_TYPE": "", "SUID": 2785657, "DEGREE": 5.0, "id_original": "853165", "MCODE_CLUSTER_ID": 0.0, "selected": false, "canonicalName": "RIE1"}, "position": {"x": 159.9390425726741, "y": 233.11266051486444}, "selected": false}, {"data": {"id": "2785654", "shared_name": "853164", "MCODE_SCORE": 1.0, "Gene": "853164", "Symbol": "MGA1", "name": "853164", "MCODE_TYPE": "Clustered", "SUID": 2785654, "DEGREE": 6.0, "id_original": "853164", "MCODE_CLUSTER_ID": 7.0, "selected": false, "canonicalName": "MGA1"}, "position": {"x": 309.25535345524247, "y": 132.22549925998163}, "selected": false}, {"data": {"id": "2785651", "shared_name": "853163", "MCODE_SCORE": 0.0, "Gene": "853163", "Symbol": "SOL4", "name": "853163", "MCODE_TYPE": "", "SUID": 2785651, "DEGREE": 2.0, "id_original": "853163", "MCODE_CLUSTER_ID": 0.0, "selected": false, "canonicalName": "SOL4"}, "position": {"x": 257.5294661566585, "y": -49.201338661160946}, "selected": false}, {"data": {"id": "2785648", "shared_name": "853159", "MCODE_SCORE": 0.0, "Gene": "853159", "Symbol": "LSC2", "name": "853159", "MCODE_TYPE": "", "SUID": 2785648, "DEGREE": 14.0, "id_original": "853159", "MCODE_CLUSTER_ID": 0.0, "selected": false, "canonicalName": "LSC2"}, "position": {"x": 457.2804884955257, "y": 546.0734110946985}, "selected": false}, {"data": {"id": "2785645", "shared_name": "853158", "MCODE_SCORE": 0.0, "Gene": "853158", "Symbol": "MPC3", "name": "853158", "MCODE_TYPE": "", "SUID": 2785645, "DEGREE": 10.0, "id_original": "853158", "MCODE_CLUSTER_ID": 0.0, "selected": false, "canonicalName": "MPC3"}, "position": {"x": 236.3023467108577, "y": 709.183884727511}, "selected": false}, {"data": {"id": "2785642", "shared_name": "853148", "MCODE_SCORE": 1.4000000000000001, "Gene": "853148", "Symbol": "PHO81", "name": "853148", "MCODE_TYPE": "Clustered", "SUID": 2785642, "DEGREE": 19.0, "id_original": "853148", "MCODE_CLUSTER_ID": 4.0, "selected": false, "canonicalName": "PHO81"}, "position": {"x": 335.51173544376786, "y": 218.5236102218957}, "selected": false}, {"data": {"id": "2785639", "shared_name": "853119", "MCODE_SCORE": 1.0, "Gene": "853119", "Symbol": "TDA10", "name": "853119", "MCODE_TYPE": "Clustered", "SUID": 2785639, "DEGREE": 7.0, "id_original": "853119", "MCODE_CLUSTER_ID": 8.0, "selected": false, "canonicalName": "TDA10"}, "position": {"x": -14.205473418536826, "y": 584.1144877548547}, "selected": false}, {"data": {"id": "2785636", "shared_name": "853108", "MCODE_SCORE": 0.0, "Gene": "853108", "Symbol": "XKS1", "name": "853108", "MCODE_TYPE": "", "SUID": 2785636, "DEGREE": 1.0, "id_original": "853108", "MCODE_CLUSTER_ID": 0.0, "selected": false, "canonicalName": "XKS1"}, "position": {"x": 1349.5696311041684, "y": -360.5818013076453}, "selected": false}, {"data": {"id": "2785633", "shared_name": "853095", "MCODE_SCORE": 9.625, "Gene": "853095", "Symbol": "QCR9", "name": "853095", "MCODE_TYPE": "Clustered", "SUID": 2785633, "DEGREE": 31.0, "id_original": "853095", "MCODE_CLUSTER_ID": 1.0, "selected": false, "canonicalName": "QCR9"}, "position": {"x": 326.36024427860673, "y": 480.65068160251093}, "selected": false}, {"data": {"id": "2785630", "shared_name": "853039", "MCODE_SCORE": 0.0, "Gene": "853039", "Symbol": "TPO2", "name": "853039", "MCODE_TYPE": "", "SUID": 2785630, "DEGREE": 3.0, "id_original": "853039", "MCODE_CLUSTER_ID": 0.0, "selected": false, "canonicalName": "TPO2"}, "position": {"x": 661.5949111029475, "y": 204.71734915927362}, "selected": false}, {"data": {"id": "2785627", "shared_name": "853031", "MCODE_SCORE": 1.25, "Gene": "853031", "Symbol": "YGR130C", "name": "853031", "MCODE_TYPE": "Clustered", "SUID": 2785627, "DEGREE": 13.0, "id_original": "853031", "MCODE_CLUSTER_ID": 6.0, "selected": false, "canonicalName": "YGR130C"}, "position": {"x": 514.6865553900569, "y": 285.75288497165155}, "selected": false}, {"data": {"id": "2785624", "shared_name": "852961", "MCODE_SCORE": 0.0, "Gene": "852961", "Symbol": "ROM1", "name": "852961", "MCODE_TYPE": "", "SUID": 2785624, "DEGREE": 6.0, "id_original": "852961", "MCODE_CLUSTER_ID": 0.0, "selected": false, "canonicalName": "ROM1"}, "position": {"x": 377.186212067303, "y": 15.227082359346866}, "selected": false}, {"data": {"id": "2785621", "shared_name": "852943", "MCODE_SCORE": 0.0, "Gene": "852943", "Symbol": "FMP48", "name": "852943", "MCODE_TYPE": "", "SUID": 2785621, "DEGREE": 9.0, "id_original": "852943", "MCODE_CLUSTER_ID": 0.0, "selected": false, "canonicalName": "FMP48"}, "position": {"x": 194.0933394476741, "y": 146.29907332614374}, "selected": false}, {"data": {"id": "2785618", "shared_name": "852905", "MCODE_SCORE": 0.0, "Gene": "852905", "Symbol": "MTL1", "name": "852905", "MCODE_TYPE": "", "SUID": 2785618, "DEGREE": 1.0, "id_original": "852905", "MCODE_CLUSTER_ID": 0.0, "selected": false, "canonicalName": "MTL1"}, "position": {"x": 428.6741042181819, "y": -172.63090409084845}, "selected": false}, {"data": {"id": "2785615", "shared_name": "852891", "MCODE_SCORE": 0.0, "Gene": "852891", "Symbol": "STF2", "name": "852891", "MCODE_TYPE": "", "SUID": 2785615, "DEGREE": 1.0, "id_original": "852891", "MCODE_CLUSTER_ID": 0.0, "selected": false, "canonicalName": "STF2"}, "position": {"x": 1451.8912215277524, "y": -360.5818013076453}, "selected": false}, {"data": {"id": "2785612", "shared_name": "852878", "MCODE_SCORE": 0.0, "Gene": "852878", "Symbol": "PMC1", "name": "852878", "MCODE_TYPE": "", "SUID": 2785612, "DEGREE": 1.0, "id_original": "852878", "MCODE_CLUSTER_ID": 0.0, "selected": false, "canonicalName": "PMC1"}, "position": {"x": 496.92637777775224, "y": 797.530076133761}, "selected": false}, {"data": {"id": "2785609", "shared_name": "852846", "MCODE_SCORE": 0.0, "Gene": "852846", "Symbol": "PNC1", "name": "852846", "MCODE_TYPE": "", "SUID": 2785609, "DEGREE": 4.0, "id_original": "852846", "MCODE_CLUSTER_ID": 0.0, "selected": false, "canonicalName": "PNC1"}, "position": {"x": 171.860887055096, "y": 348.9225016708703}, "selected": false}, {"data": {"id": "2785606", "shared_name": "852783", "MCODE_SCORE": 0.0, "Gene": "852783", "Symbol": "TOS8", "name": "852783", "MCODE_TYPE": "", "SUID": 2785606, "DEGREE": 1.0, "id_original": "852783", "MCODE_CLUSTER_ID": 0.0, "selected": false, "canonicalName": "TOS8"}, "position": {"x": -118.89993142634933, "y": 603.1153117294641}, "selected": false}, {"data": {"id": "2785603", "shared_name": "852774", "MCODE_SCORE": 0.0, "Gene": "852774", "Symbol": "VPS73", "name": "852774", "MCODE_TYPE": "", "SUID": 2785603, "DEGREE": 2.0, "id_original": "852774", "MCODE_CLUSTER_ID": 0.0, "selected": false, "canonicalName": "VPS73"}, "position": {"x": 263.16999578922685, "y": 329.4685984726281}, "selected": false}, {"data": {"id": "2785600", "shared_name": "852688", "MCODE_SCORE": 9.625, "Gene": "852688", "Symbol": "COX4", "name": "852688", "MCODE_TYPE": "Clustered", "SUID": 2785600, "DEGREE": 37.0, "id_original": "852688", "MCODE_CLUSTER_ID": 1.0, "selected": false, "canonicalName": "COX4"}, "position": {"x": 272.47764349430497, "y": 634.8657084579797}, "selected": false}, {"data": {"id": "2785597", "shared_name": "852679", "MCODE_SCORE": 0.0, "Gene": "852679", "Symbol": "DSD1", "name": "852679", "MCODE_TYPE": "", "SUID": 2785597, "DEGREE": 1.0, "id_original": "852679", "MCODE_CLUSTER_ID": 0.0, "selected": false, "canonicalName": "DSD1"}, "position": {"x": 416.15814352482255, "y": -305.45295609280157}, "selected": false}, {"data": {"id": "2785594", "shared_name": "852641", "MCODE_SCORE": 0.0, "Gene": "852641", "Symbol": "HFM1", "name": "852641", "MCODE_TYPE": "", "SUID": 2785594, "DEGREE": 2.0, "id_original": "852641", "MCODE_CLUSTER_ID": 0.0, "selected": false, "canonicalName": "HFM1"}, "position": {"x": 471.7213912054866, "y": 83.07491408542109}, "selected": false}, {"data": {"id": "2785591", "shared_name": "852601", "MCODE_SCORE": 0.0, "Gene": "852601", "Symbol": "MAL31", "name": "852601", "MCODE_TYPE": "", "SUID": 2785591, "DEGREE": 1.0, "id_original": "852601", "MCODE_CLUSTER_ID": 0.0, "selected": false, "canonicalName": "MAL31"}, "position": {"x": 541.8869948431819, "y": 625.5424052353235}, "selected": false}, {"data": {"id": "2785588", "shared_name": "852572", "MCODE_SCORE": 0.0, "Gene": "852572", "Symbol": "SDH8", "name": "852572", "MCODE_TYPE": "", "SUID": 2785588, "DEGREE": 2.0, "id_original": "852572", "MCODE_CLUSTER_ID": 0.0, "selected": false, "canonicalName": "SDH8"}, "position": {"x": 490.09173727482255, "y": 452.6895609970422}, "selected": false}, {"data": {"id": "2785585", "shared_name": "852558", "MCODE_SCORE": 0.0, "Gene": "852558", "Symbol": "RCF3", "name": "852558", "MCODE_TYPE": "", "SUID": 2785585, "DEGREE": 5.0, "id_original": "852558", "MCODE_CLUSTER_ID": 0.0, "selected": false, "canonicalName": "RCF3"}, "position": {"x": 394.422540192303, "y": 719.5912943954797}, "selected": false}, {"data": {"id": "2785582", "shared_name": "852531", "MCODE_SCORE": 0.0, "Gene": "852531", "Symbol": "OM14", "name": "852531", "MCODE_TYPE": "", "SUID": 2785582, "DEGREE": 13.0, "id_original": "852531", "MCODE_CLUSTER_ID": 0.0, "selected": false, "canonicalName": "OM14"}, "position": {"x": 233.57994986027177, "y": 550.1878520126672}, "selected": false}, {"data": {"id": "2785579", "shared_name": "852515", "MCODE_SCORE": 1.0, "Gene": "852515", "Symbol": "SDS24", "name": "852515", "MCODE_TYPE": "Clustered", "SUID": 2785579, "DEGREE": 9.0, "id_original": "852515", "MCODE_CLUSTER_ID": 7.0, "selected": false, "canonicalName": "SDS24"}, "position": {"x": 259.7218031927913, "y": 88.76086531833124}, "selected": false}, {"data": {"id": "2785576", "shared_name": "852481", "MCODE_SCORE": 0.0, "Gene": "852481", "Symbol": "YPC1", "name": "852481", "MCODE_TYPE": "", "SUID": 2785576, "DEGREE": 1.0, "id_original": "852481", "MCODE_CLUSTER_ID": 0.0, "selected": false, "canonicalName": "YPC1"}, "position": {"x": 1247.2480406805844, "y": -360.5818013076453}, "selected": false}, {"data": {"id": "2785573", "shared_name": "852467", "MCODE_SCORE": 0.0, "Gene": "852467", "Symbol": "SSE2", "name": "852467", "MCODE_TYPE": "", "SUID": 2785573, "DEGREE": 8.0, "id_original": "852467", "MCODE_CLUSTER_ID": 0.0, "selected": false, "canonicalName": "SSE2"}, "position": {"x": 253.9327406927913, "y": 245.6608401413293}, "selected": false}, {"data": {"id": "2785570", "shared_name": "852446", "MCODE_SCORE": 0.0, "Gene": "852446", "Symbol": "ARA1", "name": "852446", "MCODE_TYPE": "", "SUID": 2785570, "DEGREE": 1.0, "id_original": "852446", "MCODE_CLUSTER_ID": 0.0, "selected": false, "canonicalName": "ARA1"}, "position": {"x": 151.263719086346, "y": 19.080109703096866}, "selected": false}, {"data": {"id": "2785567", "shared_name": "852436", "MCODE_SCORE": 0.0, "Gene": "852436", "Symbol": "ATG42", "name": "852436", "MCODE_TYPE": "", "SUID": 2785567, "DEGREE": 5.0, "id_original": "852436", "MCODE_CLUSTER_ID": 0.0, "selected": false, "canonicalName": "ATG42"}, "position": {"x": -132.83395242244308, "y": 217.5830355758996}, "selected": false}, {"data": {"id": "2785564", "shared_name": "852423", "MCODE_SCORE": 2.9166666666666665, "Gene": "852423", "Symbol": "TPS1", "name": "852423", "MCODE_TYPE": "Clustered", "SUID": 2785564, "DEGREE": 20.0, "id_original": "852423", "MCODE_CLUSTER_ID": 2.0, "selected": false, "canonicalName": "TPS1"}, "position": {"x": 222.7301497504085, "y": 186.5464783783288}, "selected": false}, {"data": {"id": "2785561", "shared_name": "852402", "MCODE_SCORE": 0.0, "Gene": "852402", "Symbol": "VID24", "name": "852402", "MCODE_TYPE": "", "SUID": 2785561, "DEGREE": 8.0, "id_original": "852402", "MCODE_CLUSTER_ID": 0.0, "selected": false, "canonicalName": "VID24"}, "position": {"x": 100.57545614689286, "y": 458.97285567477655}, "selected": false}, {"data": {"id": "2785558", "shared_name": "852390", "MCODE_SCORE": 0.0, "Gene": "852390", "Symbol": "PHO5", "name": "852390", "MCODE_TYPE": "", "SUID": 2785558, "DEGREE": 5.0, "id_original": "852390", "MCODE_CLUSTER_ID": 0.0, "selected": false, "canonicalName": "PHO5"}, "position": {"x": 134.6748824164241, "y": 399.49646102145624}, "selected": false}, {"data": {"id": "2785555", "shared_name": "852381", "MCODE_SCORE": 0.0, "Gene": "852381", "Symbol": "YBR085C-A", "name": "852381", "MCODE_TYPE": "", "SUID": 2785555, "DEGREE": 1.0, "id_original": "852381", "MCODE_CLUSTER_ID": 0.0, "selected": false, "canonicalName": "YBR085C-A"}, "position": {"x": 411.0213179632991, "y": 364.7531443710656}, "selected": false}, {"data": {"id": "2785552", "shared_name": "852346", "MCODE_SCORE": 0.0, "Gene": "852346", "Symbol": "MRX18", "name": "852346", "MCODE_TYPE": "", "SUID": 2785552, "DEGREE": 1.0, "id_original": "852346", "MCODE_CLUSTER_ID": 0.0, "selected": false, "canonicalName": "MRX18"}, "position": {"x": -46.801298613849326, "y": 714.609910118136}, "selected": false}, {"data": {"id": "2785549", "shared_name": "852342", "MCODE_SCORE": 0.0, "Gene": "852342", "Symbol": "YBR053C", "name": "852342", "MCODE_TYPE": "", "SUID": 2785549, "DEGREE": 1.0, "id_original": "852342", "MCODE_CLUSTER_ID": 0.0, "selected": false, "canonicalName": "YBR053C"}, "position": {"x": 106.3320326849788, "y": -3.2455281142859462}, "selected": false}, {"data": {"id": "2785546", "shared_name": "852341", "MCODE_SCORE": 0.0, "Gene": "852341", "Symbol": "RFS1", "name": "852341", "MCODE_TYPE": "", "SUID": 2785546, "DEGREE": 3.0, "id_original": "852341", "MCODE_CLUSTER_ID": 0.0, "selected": false, "canonicalName": "RFS1"}, "position": {"x": 615.4115920111507, "y": 257.78212844088984}, "selected": false}, {"data": {"id": "2785543", "shared_name": "852340", "MCODE_SCORE": 1.0, "Gene": "852340", "Symbol": "REG2", "name": "852340", "MCODE_TYPE": "Seed", "SUID": 2785543, "DEGREE": 3.0, "id_original": "852340", "MCODE_CLUSTER_ID": 10.0, "selected": false, "canonicalName": "REG2"}, "position": {"x": 308.6290307089656, "y": -299.48908890530157}, "selected": false}, {"data": {"id": "2785540", "shared_name": "852335", "MCODE_SCORE": 0.0, "Gene": "852335", "Symbol": "ZTA1", "name": "852335", "MCODE_TYPE": "", "SUID": 2785540, "DEGREE": 2.0, "id_original": "852335", "MCODE_CLUSTER_ID": 0.0, "selected": false, "canonicalName": "ZTA1"}, "position": {"x": 237.4962172552913, "y": 92.37474929049921}, "selected": false}, {"data": {"id": "2785537", "shared_name": "852327", "MCODE_SCORE": 9.625, "Gene": "852327", "Symbol": "ATP3", "name": "852327", "MCODE_TYPE": "Clustered", "SUID": 2785537, "DEGREE": 39.0, "id_original": "852327", "MCODE_CLUSTER_ID": 1.0, "selected": false, "canonicalName": "ATP3"}, "position": {"x": 256.8053908392757, "y": 484.01664839938593}, "selected": false}, {"data": {"id": "2785534", "shared_name": "852286", "MCODE_SCORE": 2.9166666666666665, "Gene": "852286", "Symbol": "NTH2", "name": "852286", "MCODE_TYPE": "Clustered", "SUID": 2785534, "DEGREE": 9.0, "id_original": "852286", "MCODE_CLUSTER_ID": 2.0, "selected": false, "canonicalName": "NTH2"}, "position": {"x": 82.51106405704911, "y": 122.4579020614953}, "selected": false}, {"data": {"id": "2785531", "shared_name": "852266", "MCODE_SCORE": 0.0, "Gene": "852266", "Symbol": "ACH1", "name": "852266", "MCODE_TYPE": "", "SUID": 2785531, "DEGREE": 9.0, "id_original": "852266", "MCODE_CLUSTER_ID": 0.0, "selected": false, "canonicalName": "ACH1"}, "position": {"x": 125.19972372501786, "y": 607.616501915011}, "selected": false}, {"data": {"id": "2785528", "shared_name": "852251", "MCODE_SCORE": 1.25, "Gene": "852251", "Symbol": "YBL029C-A", "name": "852251", "MCODE_TYPE": "Seed", "SUID": 2785528, "DEGREE": 3.0, "id_original": "852251", "MCODE_CLUSTER_ID": 6.0, "selected": false, "canonicalName": "YBL029C-A"}, "position": {"x": 594.8743910834163, "y": 387.41130171969843}, "selected": false}, {"data": {"id": "2785525", "shared_name": "852250", "MCODE_SCORE": 2.375, "Gene": "852250", "Symbol": "PET9", "name": "852250", "MCODE_TYPE": "Clustered", "SUID": 2785525, "DEGREE": 26.0, "id_original": "852250", "MCODE_CLUSTER_ID": 3.0, "selected": false, "canonicalName": "PET9"}, "position": {"x": 223.21538687199052, "y": 519.0248576279016}, "selected": false}, {"data": {"id": "2785522", "shared_name": "852237", "MCODE_SCORE": 0.0, "Gene": "852237", "Symbol": "ECM13", "name": "852237", "MCODE_TYPE": "", "SUID": 2785522, "DEGREE": 2.0, "id_original": "852237", "MCODE_CLUSTER_ID": 0.0, "selected": false, "canonicalName": "ECM13"}, "position": {"x": 1042.6048598334164, "y": 125.12912093356562}, "selected": false}, {"data": {"id": "2785519", "shared_name": "852235", "MCODE_SCORE": 9.625, "Gene": "852235", "Symbol": "COR1", "name": "852235", "MCODE_TYPE": "Clustered", "SUID": 2785519, "DEGREE": 39.0, "id_original": "852235", "MCODE_CLUSTER_ID": 1.0, "selected": false, "canonicalName": "COR1"}, "position": {"x": 320.2740912482112, "y": 585.0470439071985}, "selected": false}, {"data": {"id": "2785516", "shared_name": "852214", "MCODE_SCORE": 0.0, "Gene": "852214", "Symbol": "SEF1", "name": "852214", "MCODE_TYPE": "", "SUID": 2785516, "DEGREE": 2.0, "id_original": "852214", "MCODE_CLUSTER_ID": 0.0, "selected": false, "canonicalName": "SEF1"}, "position": {"x": 92.45407247990067, "y": -25.904036415067196}, "selected": false}, {"data": {"id": "2785513", "shared_name": "852200", "MCODE_SCORE": 0.0, "Gene": "852200", "Symbol": "ATG8", "name": "852200", "MCODE_TYPE": "", "SUID": 2785513, "DEGREE": 5.0, "id_original": "852200", "MCODE_CLUSTER_ID": 0.0, "selected": false, "canonicalName": "ATG8"}, "position": {"x": 18.62677145451005, "y": 504.7926188583703}, "selected": false}, {"data": {"id": "2785510", "shared_name": "852181", "MCODE_SCORE": 0.0, "Gene": "852181", "Symbol": "MRX3", "name": "852181", "MCODE_TYPE": "", "SUID": 2785510, "DEGREE": 1.0, "id_original": "852181", "MCODE_CLUSTER_ID": 0.0, "selected": false, "canonicalName": "MRX3"}, "position": {"x": 550.3913588568538, "y": 560.5555583114954}, "selected": false}, {"data": {"id": "2785507", "shared_name": "852179", "MCODE_SCORE": 0.0, "Gene": "852179", "Symbol": "BNA4", "name": "852179", "MCODE_TYPE": "", "SUID": 2785507, "DEGREE": 1.0, "id_original": "852179", "MCODE_CLUSTER_ID": 0.0, "selected": false, "canonicalName": "BNA4"}, "position": {"x": 210.4494643256038, "y": -256.0784748916297}, "selected": false}, {"data": {"id": "2785504", "shared_name": "852177", "MCODE_SCORE": 9.625, "Gene": "852177", "Symbol": "ATP1", "name": "852177", "MCODE_TYPE": "Clustered", "SUID": 2785504, "DEGREE": 42.0, "id_original": "852177", "MCODE_CLUSTER_ID": 1.0, "selected": false, "canonicalName": "ATP1"}, "position": {"x": 287.29908132999833, "y": 451.79194747165155}, "selected": false}, {"data": {"id": "2785501", "shared_name": "852142", "MCODE_SCORE": 9.625, "Gene": "852142", "Symbol": "QCR7", "name": "852142", "MCODE_TYPE": "Clustered", "SUID": 2785501, "DEGREE": 26.0, "id_original": "852142", "MCODE_CLUSTER_ID": 1.0, "selected": false, "canonicalName": "QCR7"}, "position": {"x": 200.9562010809749, "y": 571.5932780380579}, "selected": false}, {"data": {"id": "2785498", "shared_name": "852138", "MCODE_SCORE": 0.0, "Gene": "852138", "Symbol": "SNA2", "name": "852138", "MCODE_TYPE": "", "SUID": 2785498, "DEGREE": 1.0, "id_original": "852138", "MCODE_CLUSTER_ID": 0.0, "selected": false, "canonicalName": "SNA2"}, "position": {"x": 170.71441793888505, "y": 646.418503868136}, "selected": false}, {"data": {"id": "2785495", "shared_name": "852128", "MCODE_SCORE": 1.0, "Gene": "852128", "Symbol": "EMI2", "name": "852128", "MCODE_TYPE": "Seed", "SUID": 2785495, "DEGREE": 8.0, "id_original": "852128", "MCODE_CLUSTER_ID": 9.0, "selected": false, "canonicalName": "EMI2"}, "position": {"x": 132.2000746771663, "y": 200.64103995517206}, "selected": false}, {"data": {"id": "2785492", "shared_name": "852064", "MCODE_SCORE": 0.0, "Gene": "852064", "Symbol": "TSA2", "name": "852064", "MCODE_TYPE": "", "SUID": 2785492, "DEGREE": 6.0, "id_original": "852064", "MCODE_CLUSTER_ID": 0.0, "selected": false, "canonicalName": "TSA2"}, "position": {"x": 9.363862518963174, "y": 307.52775679782343}, "selected": false}, {"data": {"id": "2785489", "shared_name": "851983", "MCODE_SCORE": 9.625, "Gene": "851983", "Symbol": "ATP17", "name": "851983", "MCODE_TYPE": "Clustered", "SUID": 2785489, "DEGREE": 16.0, "id_original": "851983", "MCODE_CLUSTER_ID": 1.0, "selected": false, "canonicalName": "ATP17"}, "position": {"x": 161.80479574650224, "y": 614.2383891220422}, "selected": false}, {"data": {"id": "2785486", "shared_name": "851960", "MCODE_SCORE": 0.0, "Gene": "851960", "Symbol": "GGA1", "name": "851960", "MCODE_TYPE": "", "SUID": 2785486, "DEGREE": 1.0, "id_original": "851960", "MCODE_CLUSTER_ID": 0.0, "selected": false, "canonicalName": "GGA1"}, "position": {"x": 80.16775656193192, "y": -199.9033039931922}, "selected": false}, {"data": {"id": "2785483", "shared_name": "851944", "MCODE_SCORE": 0.0, "Gene": "851944", "Symbol": "HXT6", "name": "851944", "MCODE_TYPE": "", "SUID": 2785483, "DEGREE": 3.0, "id_original": "851944", "MCODE_CLUSTER_ID": 0.0, "selected": false, "canonicalName": "HXT6"}, "position": {"x": -19.24847268611495, "y": 105.07956038669062}, "selected": false}, {"data": {"id": "2785480", "shared_name": "851943", "MCODE_SCORE": 0.0, "Gene": "851943", "Symbol": "HXT7", "name": "851943", "MCODE_TYPE": "", "SUID": 2785480, "DEGREE": 4.0, "id_original": "851943", "MCODE_CLUSTER_ID": 0.0, "selected": false, "canonicalName": "HXT7"}, "position": {"x": 55.9112563177913, "y": 264.0787974472375}, "selected": false}, {"data": {"id": "2785477", "shared_name": "851922", "MCODE_SCORE": 9.625, "Gene": "851922", "Symbol": "TIM11", "name": "851922", "MCODE_TYPE": "Clustered", "SUID": 2785477, "DEGREE": 15.0, "id_original": "851922", "MCODE_CLUSTER_ID": 1.0, "selected": false, "canonicalName": "TIM11"}, "position": {"x": 225.56434011906083, "y": 455.4081889267297}, "selected": false}, {"data": {"id": "2785474", "shared_name": "851919", "MCODE_SCORE": 0.0, "Gene": "851919", "Symbol": "DAD4", "name": "851919", "MCODE_TYPE": "", "SUID": 2785474, "DEGREE": 2.0, "id_original": "851919", "MCODE_CLUSTER_ID": 0.0, "selected": false, "canonicalName": "DAD4"}, "position": {"x": 924.7023940131038, "y": 120.50867568210077}, "selected": false}, {"data": {"id": "2785471", "shared_name": "851908", "MCODE_SCORE": 0.0, "Gene": "851908", "Symbol": "PIB1", "name": "851908", "MCODE_TYPE": "", "SUID": 2785471, "DEGREE": 3.0, "id_original": "851908", "MCODE_CLUSTER_ID": 0.0, "selected": false, "canonicalName": "PIB1"}, "position": {"x": 117.17454672306474, "y": 345.66670333102655}, "selected": false}, {"data": {"id": "2785468", "shared_name": "851892", "MCODE_SCORE": 2.375, "Gene": "851892", "Symbol": "ATP5", "name": "851892", "MCODE_TYPE": "Clustered", "SUID": 2785468, "DEGREE": 30.0, "id_original": "851892", "MCODE_CLUSTER_ID": 3.0, "selected": false, "canonicalName": "ATP5"}, "position": {"x": 267.81394720524247, "y": 462.28288375094843}, "selected": false}, {"data": {"id": "2785465", "shared_name": "851870", "MCODE_SCORE": 0.0, "Gene": "851870", "Symbol": "MTH1", "name": "851870", "MCODE_TYPE": "", "SUID": 2785465, "DEGREE": 4.0, "id_original": "851870", "MCODE_CLUSTER_ID": 0.0, "selected": false, "canonicalName": "MTH1"}, "position": {"x": 305.0359606787532, "y": 168.0538903351159}, "selected": false}, {"data": {"id": "2785462", "shared_name": "851862", "MCODE_SCORE": 0.0, "Gene": "851862", "Symbol": "CCC2", "name": "851862", "MCODE_TYPE": "", "SUID": 2785462, "DEGREE": 1.0, "id_original": "851862", "MCODE_CLUSTER_ID": 0.0, "selected": false, "canonicalName": "CCC2"}, "position": {"x": 227.72843313663896, "y": 1077.366257774386}, "selected": false}, {"data": {"id": "2785459", "shared_name": "851846", "MCODE_SCORE": 0.0, "Gene": "851846", "Symbol": "YAP6", "name": "851846", "MCODE_TYPE": "", "SUID": 2785459, "DEGREE": 2.0, "id_original": "851846", "MCODE_CLUSTER_ID": 0.0, "selected": false, "canonicalName": "YAP6"}, "position": {"x": 535.08673239201, "y": 146.29025374606562}, "selected": false}, {"data": {"id": "2785456", "shared_name": "851845", "MCODE_SCORE": 0.0, "Gene": "851845", "Symbol": "HSP78", "name": "851845", "MCODE_TYPE": "", "SUID": 2785456, "DEGREE": 6.0, "id_original": "851845", "MCODE_CLUSTER_ID": 0.0, "selected": false, "canonicalName": "HSP78"}, "position": {"x": 288.41077566593583, "y": 220.16567336276484}, "selected": false}, {"data": {"id": "2785453", "shared_name": "851835", "MCODE_SCORE": 0.0, "Gene": "851835", "Symbol": "YDR248C", "name": "851835", "MCODE_TYPE": "", "SUID": 2785453, "DEGREE": 3.0, "id_original": "851835", "MCODE_CLUSTER_ID": 0.0, "selected": false, "canonicalName": "YDR248C"}, "position": {"x": 338.7695803686946, "y": -128.70014847561407}, "selected": false}, {"data": {"id": "2785450", "shared_name": "851834", "MCODE_SCORE": 1.0, "Gene": "851834", "Symbol": "VHS1", "name": "851834", "MCODE_TYPE": "Clustered", "SUID": 2785450, "DEGREE": 11.0, "id_original": "851834", "MCODE_CLUSTER_ID": 7.0, "selected": false, "canonicalName": "VHS1"}, "position": {"x": 202.98253012150224, "y": 37.02331649020624}, "selected": false}, {"data": {"id": "2785447", "shared_name": "851758", "MCODE_SCORE": 9.625, "Gene": "851758", "Symbol": "SDH4", "name": "851758", "MCODE_TYPE": "Clustered", "SUID": 2785447, "DEGREE": 16.0, "id_original": "851758", "MCODE_CLUSTER_ID": 1.0, "selected": false, "canonicalName": "SDH4"}, "position": {"x": 277.97325277775224, "y": 465.1275797958703}, "selected": false}, {"data": {"id": "2785444", "shared_name": "851751", "MCODE_SCORE": 0.0, "Gene": "851751", "Symbol": "HSP42", "name": "851751", "MCODE_TYPE": "", "SUID": 2785444, "DEGREE": 9.0, "id_original": "851751", "MCODE_CLUSTER_ID": 0.0, "selected": false, "canonicalName": "HSP42"}, "position": {"x": 87.72793722599442, "y": 285.6642695541711}, "selected": false}, {"data": {"id": "2785441", "shared_name": "851726", "MCODE_SCORE": 0.0, "Gene": "851726", "Symbol": "KGD2", "name": "851726", "MCODE_TYPE": "", "SUID": 2785441, "DEGREE": 12.0, "id_original": "851726", "MCODE_CLUSTER_ID": 0.0, "selected": false, "canonicalName": "KGD2"}, "position": {"x": 388.1703887030452, "y": 487.1186381454797}, "selected": false}, {"data": {"id": "2785438", "shared_name": "851670", "MCODE_SCORE": 2.9166666666666665, "Gene": "851670", "Symbol": "GIS1", "name": "851670", "MCODE_TYPE": "Clustered", "SUID": 2785438, "DEGREE": 20.0, "id_original": "851670", "MCODE_CLUSTER_ID": 2.0, "selected": false, "canonicalName": "GIS1"}, "position": {"x": 213.35538626163896, "y": 290.7286997909875}, "selected": false}, {"data": {"id": "2785435", "shared_name": "851646", "MCODE_SCORE": 2.9166666666666665, "Gene": "851646", "Symbol": "TPS2", "name": "851646", "MCODE_TYPE": "Seed", "SUID": 2785435, "DEGREE": 9.0, "id_original": "851646", "MCODE_CLUSTER_ID": 2.0, "selected": false, "canonicalName": "TPS2"}, "position": {"x": 176.63864279240067, "y": 130.25319777682734}, "selected": false}, {"data": {"id": "2785432", "shared_name": "851642", "MCODE_SCORE": 0.0, "Gene": "851642", "Symbol": "FMP16", "name": "851642", "MCODE_TYPE": "", "SUID": 2785432, "DEGREE": 1.0, "id_original": "851642", "MCODE_CLUSTER_ID": 0.0, "selected": false, "canonicalName": "FMP16"}, "position": {"x": 525.1457686468929, "y": -93.88368119045782}, "selected": false}, {"data": {"id": "2785429", "shared_name": "851631", "MCODE_SCORE": 1.4000000000000001, "Gene": "851631", "Symbol": "UBC5", "name": "851631", "MCODE_TYPE": "Clustered", "SUID": 2785429, "DEGREE": 14.0, "id_original": "851631", "MCODE_CLUSTER_ID": 4.0, "selected": false, "canonicalName": "UBC5"}, "position": {"x": 344.0015196844905, "y": 120.91108046725702}, "selected": false}, {"data": {"id": "2785426", "shared_name": "851613", "MCODE_SCORE": 1.4000000000000001, "Gene": "851613", "Symbol": "NRG1", "name": "851613", "MCODE_TYPE": "Clustered", "SUID": 2785426, "DEGREE": 7.0, "id_original": "851613", "MCODE_CLUSTER_ID": 4.0, "selected": false, "canonicalName": "NRG1"}, "position": {"x": 419.22986746281083, "y": 125.60513411715937}, "selected": false}, {"data": {"id": "2785423", "shared_name": "851597", "MCODE_SCORE": 0.0, "Gene": "851597", "Symbol": "MRH1", "name": "851597", "MCODE_TYPE": "", "SUID": 2785423, "DEGREE": 1.0, "id_original": "851597", "MCODE_CLUSTER_ID": 0.0, "selected": false, "canonicalName": "MRH1"}, "position": {"x": -74.49786233455245, "y": 266.009224998507}, "selected": false}, {"data": {"id": "2785420", "shared_name": "851595", "MCODE_SCORE": 0.0, "Gene": "851595", "Symbol": "MIX14", "name": "851595", "MCODE_TYPE": "", "SUID": 2785420, "DEGREE": 1.0, "id_original": "851595", "MCODE_CLUSTER_ID": 0.0, "selected": false, "canonicalName": "MIX14"}, "position": {"x": 1554.2128119513363, "y": -360.5818013076453}, "selected": false}, {"data": {"id": "2785417", "shared_name": "851564", "MCODE_SCORE": 2.9166666666666665, "Gene": "851564", "Symbol": "NTH1", "name": "851564", "MCODE_TYPE": "Clustered", "SUID": 2785417, "DEGREE": 11.0, "id_original": "851564", "MCODE_CLUSTER_ID": 2.0, "selected": false, "canonicalName": "NTH1"}, "position": {"x": 186.36854696720536, "y": 171.8899499054284}, "selected": false}, {"data": {"id": "2785414", "shared_name": "851560", "MCODE_SCORE": 2.375, "Gene": "851560", "Symbol": "ATP16", "name": "851560", "MCODE_TYPE": "Clustered", "SUID": 2785414, "DEGREE": 30.0, "id_original": "851560", "MCODE_CLUSTER_ID": 3.0, "selected": false, "canonicalName": "ATP16"}, "position": {"x": 301.6217055365413, "y": 592.1027079696985}, "selected": false}, {"data": {"id": "2785411", "shared_name": "851541", "MCODE_SCORE": 0.0, "Gene": "851541", "Symbol": "GPM2", "name": "851541", "MCODE_TYPE": "", "SUID": 2785411, "DEGREE": 3.0, "id_original": "851541", "MCODE_CLUSTER_ID": 0.0, "selected": false, "canonicalName": "GPM2"}, "position": {"x": 190.18971395939286, "y": 364.5946971054406}, "selected": false}, {"data": {"id": "2785408", "shared_name": "851539", "MCODE_SCORE": 0.0, "Gene": "851539", "Symbol": "GPD1", "name": "851539", "MCODE_TYPE": "", "SUID": 2785408, "DEGREE": 6.0, "id_original": "851539", "MCODE_CLUSTER_ID": 0.0, "selected": false, "canonicalName": "GPD1"}, "position": {"x": 17.4563002631038, "y": 383.5219431991906}, "selected": false}, {"data": {"id": "2785405", "shared_name": "851512", "MCODE_SCORE": 1.4000000000000001, "Gene": "851512", "Symbol": "STP4", "name": "851512", "MCODE_TYPE": "Clustered", "SUID": 2785405, "DEGREE": 8.0, "id_original": "851512", "MCODE_CLUSTER_ID": 4.0, "selected": false, "canonicalName": "STP4"}, "position": {"x": 484.5988326117366, "y": 187.30443154529047}, "selected": false}, {"data": {"id": "2785402", "shared_name": "851480", "MCODE_SCORE": 2.9166666666666665, "Gene": "851480", "Symbol": "MRK1", "name": "851480", "MCODE_TYPE": "Clustered", "SUID": 2785402, "DEGREE": 26.0, "id_original": "851480", "MCODE_CLUSTER_ID": 2.0, "selected": false, "canonicalName": "MRK1"}, "position": {"x": 333.45008326023515, "y": 156.9437476272912}, "selected": false}, {"data": {"id": "2785399", "shared_name": "851448", "MCODE_SCORE": 0.0, "Gene": "851448", "Symbol": "TMA17", "name": "851448", "MCODE_TYPE": "", "SUID": 2785399, "DEGREE": 1.0, "id_original": "851448", "MCODE_CLUSTER_ID": 0.0, "selected": false, "canonicalName": "TMA17"}, "position": {"x": 1654.2128119513363, "y": -252.84245194241095}, "selected": false}, {"data": {"id": "2785396", "shared_name": "851426", "MCODE_SCORE": 0.0, "Gene": "851426", "Symbol": "STF1", "name": "851426", "MCODE_TYPE": "", "SUID": 2785396, "DEGREE": 3.0, "id_original": "851426", "MCODE_CLUSTER_ID": 0.0, "selected": false, "canonicalName": "STF1"}, "position": {"x": 570.3421340033382, "y": 240.65606414035273}, "selected": false}, {"data": {"id": "2785393", "shared_name": "851411", "MCODE_SCORE": 0.0, "Gene": "851411", "Symbol": "YDL144C", "name": "851411", "MCODE_TYPE": "", "SUID": 2785393, "DEGREE": 1.0, "id_original": "851411", "MCODE_CLUSTER_ID": 0.0, "selected": false, "canonicalName": "YDL144C"}, "position": {"x": 432.8025832220882, "y": 180.18201838687372}, "selected": false}, {"data": {"id": "2785390", "shared_name": "851347", "MCODE_SCORE": 0.0, "Gene": "851347", "Symbol": "INH1", "name": "851347", "MCODE_TYPE": "", "SUID": 2785390, "DEGREE": 9.0, "id_original": "851347", "MCODE_CLUSTER_ID": 0.0, "selected": false, "canonicalName": "INH1"}, "position": {"x": 332.7072768256038, "y": 363.2702799911828}, "selected": false}, {"data": {"id": "2785387", "shared_name": "851323", "MCODE_SCORE": 0.0, "Gene": "851323", "Symbol": "RTN2", "name": "851323", "MCODE_TYPE": "", "SUID": 2785387, "DEGREE": 4.0, "id_original": "851323", "MCODE_CLUSTER_ID": 0.0, "selected": false, "canonicalName": "RTN2"}, "position": {"x": 378.0871062323421, "y": 302.9450541611047}, "selected": false}, {"data": {"id": "2785384", "shared_name": "851299", "MCODE_SCORE": 0.0, "Gene": "851299", "Symbol": "PHO11", "name": "851299", "MCODE_TYPE": "", "SUID": 2785384, "DEGREE": 1.0, "id_original": "851299", "MCODE_CLUSTER_ID": 0.0, "selected": false, "canonicalName": "PHO11"}, "position": {"x": -61.5134262993962, "y": 400.4456492538781}, "selected": false}, {"data": {"id": "2785381", "shared_name": "851259", "MCODE_SCORE": 0.0, "Gene": "851259", "Symbol": "SSA1", "name": "851259", "MCODE_TYPE": "", "SUID": 2785381, "DEGREE": 48.0, "id_original": "851259", "MCODE_CLUSTER_ID": 0.0, "selected": false, "canonicalName": "SSA1"}, "position": {"x": 279.63016653507646, "y": 315.7148295517297}, "selected": false}, {"data": {"id": "2785378", "shared_name": "851111", "MCODE_SCORE": 9.625, "Gene": "851111", "Symbol": "COX8", "name": "851111", "MCODE_TYPE": "Clustered", "SUID": 2785378, "DEGREE": 22.0, "id_original": "851111", "MCODE_CLUSTER_ID": 1.0, "selected": false, "canonicalName": "COX8"}, "position": {"x": 365.00181723087724, "y": 532.3109294052454}, "selected": false}, {"data": {"id": "2785375", "shared_name": "851059", "MCODE_SCORE": 0.0, "Gene": "851059", "Symbol": "YLR345W", "name": "851059", "MCODE_TYPE": "", "SUID": 2785375, "DEGREE": 6.0, "id_original": "851059", "MCODE_CLUSTER_ID": 0.0, "selected": false, "canonicalName": "YLR345W"}, "position": {"x": 218.97258902042802, "y": 126.34352599338007}, "selected": false}, {"data": {"id": "2785372", "shared_name": "851037", "MCODE_SCORE": 0.0, "Gene": "851037", "Symbol": "TMA10", "name": "851037", "MCODE_TYPE": "", "SUID": 2785372, "DEGREE": 1.0, "id_original": "851037", "MCODE_CLUSTER_ID": 0.0, "selected": false, "canonicalName": "TMA10"}, "position": {"x": 84.33255148380692, "y": 774.143113243136}, "selected": false}, {"data": {"id": "2785369", "shared_name": "851021", "MCODE_SCORE": 0.0, "Gene": "851021", "Symbol": "ATG39", "name": "851021", "MCODE_TYPE": "", "SUID": 2785369, "DEGREE": 1.0, "id_original": "851021", "MCODE_CLUSTER_ID": 0.0, "selected": false, "canonicalName": "ATG39"}, "position": {"x": -194.20883035213058, "y": 547.3441935654016}, "selected": false}, {"data": {"id": "2785366", "shared_name": "851002", "MCODE_SCORE": 9.625, "Gene": "851002", "Symbol": "ATP14", "name": "851002", "MCODE_TYPE": "Clustered", "SUID": 2785366, "DEGREE": 15.0, "id_original": "851002", "MCODE_CLUSTER_ID": 1.0, "selected": false, "canonicalName": "ATP14"}, "position": {"x": 224.9322981879085, "y": 416.0355082626672}, "selected": false}, {"data": {"id": "2785363", "shared_name": "850974", "MCODE_SCORE": 0.0, "Gene": "850974", "Symbol": "DCS1", "name": "850974", "MCODE_TYPE": "", "SUID": 2785363, "DEGREE": 6.0, "id_original": "850974", "MCODE_CLUSTER_ID": 0.0, "selected": false, "canonicalName": "DCS1"}, "position": {"x": 130.815476898846, "y": 306.6783305282922}, "selected": false}, {"data": {"id": "2785360", "shared_name": "850962", "MCODE_SCORE": 2.9166666666666665, "Gene": "850962", "Symbol": "GSY2", "name": "850962", "MCODE_TYPE": "Clustered", "SUID": 2785360, "DEGREE": 13.0, "id_original": "850962", "MCODE_CLUSTER_ID": 2.0, "selected": false, "canonicalName": "GSY2"}, "position": {"x": 264.85023642033036, "y": 43.762833702120304}, "selected": false}, {"data": {"id": "2785357", "shared_name": "850953", "MCODE_SCORE": 0.0, "Gene": "850953", "Symbol": "SYM1", "name": "850953", "MCODE_TYPE": "", "SUID": 2785357, "DEGREE": 3.0, "id_original": "850953", "MCODE_CLUSTER_ID": 0.0, "selected": false, "canonicalName": "SYM1"}, "position": {"x": 94.8740248724788, "y": 698.4000102157922}, "selected": false}, {"data": {"id": "2785354", "shared_name": "850933", "MCODE_SCORE": 0.0, "Gene": "850933", "Symbol": "BNA5", "name": "850933", "MCODE_TYPE": "", "SUID": 2785354, "DEGREE": 3.0, "id_original": "850933", "MCODE_CLUSTER_ID": 0.0, "selected": false, "canonicalName": "BNA5"}, "position": {"x": 194.84610891788896, "y": -71.71015824123907}, "selected": false}, {"data": {"id": "2785351", "shared_name": "850875", "MCODE_SCORE": 0.0, "Gene": "850875", "Symbol": "TFS1", "name": "850875", "MCODE_TYPE": "", "SUID": 2785351, "DEGREE": 5.0, "id_original": "850875", "MCODE_CLUSTER_ID": 0.0, "selected": false, "canonicalName": "TFS1"}, "position": {"x": 489.2705855414241, "y": 305.5538493271203}, "selected": false}, {"data": {"id": "2785348", "shared_name": "850865", "MCODE_SCORE": 0.0, "Gene": "850865", "Symbol": "UPS2", "name": "850865", "MCODE_TYPE": "", "SUID": 2785348, "DEGREE": 1.0, "id_original": "850865", "MCODE_CLUSTER_ID": 0.0, "selected": false, "canonicalName": "UPS2"}, "position": {"x": -190.61038064509933, "y": 47.23710738376093}, "selected": false}, {"data": {"id": "2785345", "shared_name": "850727", "MCODE_SCORE": 9.625, "Gene": "850727", "Symbol": "COX12", "name": "850727", "MCODE_TYPE": "Clustered", "SUID": 2785345, "DEGREE": 29.0, "id_original": "850727", "MCODE_CLUSTER_ID": 1.0, "selected": false, "canonicalName": "COX12"}, "position": {"x": 313.5277295157283, "y": 553.9008341904016}, "selected": false}, {"data": {"id": "2785342", "shared_name": "850685", "MCODE_SCORE": 9.625, "Gene": "850685", "Symbol": "SDH2", "name": "850685", "MCODE_TYPE": "Clustered", "SUID": 2785342, "DEGREE": 31.0, "id_original": "850685", "MCODE_CLUSTER_ID": 1.0, "selected": false, "canonicalName": "SDH2"}, "position": {"x": 329.4146409079402, "y": 508.6909342880578}, "selected": false}, {"data": {"id": "2785339", "shared_name": "850641", "MCODE_SCORE": 0.0, "Gene": "850641", "Symbol": "KNS1", "name": "850641", "MCODE_TYPE": "", "SUID": 2785339, "DEGREE": 11.0, "id_original": "850641", "MCODE_CLUSTER_ID": 0.0, "selected": false, "canonicalName": "KNS1"}, "position": {"x": 312.8012309119075, "y": 235.5537387008996}, "selected": false}, {"data": {"id": "2785336", "shared_name": "850633", "MCODE_SCORE": 1.0, "Gene": "850633", "Symbol": "HSP104", "name": "850633", "MCODE_TYPE": "Clustered", "SUID": 2785336, "DEGREE": 28.0, "id_original": "850633", "MCODE_CLUSTER_ID": 11.0, "selected": false, "canonicalName": "HSP104"}, "position": {"x": 147.64296102970536, "y": 246.8714991684289}, "selected": false}, {"data": {"id": "2785333", "shared_name": "850614", "MCODE_SCORE": 1.0, "Gene": "850614", "Symbol": "HXK1", "name": "850614", "MCODE_TYPE": "Clustered", "SUID": 2785333, "DEGREE": 10.0, "id_original": "850614", "MCODE_CLUSTER_ID": 9.0, "selected": false, "canonicalName": "HXK1"}, "position": {"x": 160.4812789007991, "y": 103.57756911471796}, "selected": false}, {"data": {"id": "2785330", "shared_name": "850593", "MCODE_SCORE": 9.625, "Gene": "850593", "Symbol": "QCR6", "name": "850593", "MCODE_TYPE": "Clustered", "SUID": 2785330, "DEGREE": 28.0, "id_original": "850593", "MCODE_CLUSTER_ID": 1.0, "selected": false, "canonicalName": "QCR6"}, "position": {"x": 390.23018789738114, "y": 581.5488139267297}, "selected": false}, {"data": {"id": "2785327", "shared_name": "850569", "MCODE_SCORE": 2.9166666666666665, "Gene": "850569", "Symbol": "GSY1", "name": "850569", "MCODE_TYPE": "Clustered", "SUID": 2785327, "DEGREE": 13.0, "id_original": "850569", "MCODE_CLUSTER_ID": 2.0, "selected": false, "canonicalName": "GSY1"}, "position": {"x": 248.9922881171077, "y": 85.33502303317499}, "selected": false}, {"data": {"id": "2785324", "shared_name": "850563", "MCODE_SCORE": 0.0, "Gene": "850563", "Symbol": "MIC19", "name": "850563", "MCODE_TYPE": "", "SUID": 2785324, "DEGREE": 2.0, "id_original": "850563", "MCODE_CLUSTER_ID": 0.0, "selected": false, "canonicalName": "MIC19"}, "position": {"x": 337.65051603763993, "y": 772.9982157821985}, "selected": false}, {"data": {"id": "2785321", "shared_name": "850532", "MCODE_SCORE": 0.0, "Gene": "850532", "Symbol": "HSP12", "name": "850532", "MCODE_TYPE": "", "SUID": 2785321, "DEGREE": 1.0, "id_original": "850532", "MCODE_CLUSTER_ID": 0.0, "selected": false, "canonicalName": "HSP12"}, "position": {"x": 51.960542206463174, "y": 3.1479197616906163}, "selected": false}, {"data": {"id": "2785318", "shared_name": "850514", "MCODE_SCORE": 1.0, "Gene": "850514", "Symbol": "AGX1", "name": "850514", "MCODE_TYPE": "Clustered", "SUID": 2785318, "DEGREE": 6.0, "id_original": "850514", "MCODE_CLUSTER_ID": 8.0, "selected": false, "canonicalName": "AGX1"}, "position": {"x": 7.735566620525674, "y": 565.294968711886}, "selected": false}, {"data": {"id": "2785315", "shared_name": "850490", "MCODE_SCORE": 0.0, "Gene": "850490", "Symbol": "AQY3", "name": "850490", "MCODE_TYPE": "", "SUID": 2785315, "DEGREE": 2.0, "id_original": "850490", "MCODE_CLUSTER_ID": 0.0, "selected": false, "canonicalName": "AQY3"}, "position": {"x": 599.6923537299007, "y": -65.7584065322547}, "selected": false}, {"data": {"id": "2785312", "shared_name": "850489", "MCODE_SCORE": 0.0, "Gene": "850489", "Symbol": "AGP3", "name": "850489", "MCODE_TYPE": "", "SUID": 2785312, "DEGREE": 1.0, "id_original": "850489", "MCODE_CLUSTER_ID": 0.0, "selected": false, "canonicalName": "AGP3"}, "position": {"x": 1656.5344023749203, "y": -360.5818013076453}, "selected": false}, {"data": {"id": "2785309", "shared_name": "850453", "MCODE_SCORE": 0.0, "Gene": "850453", "Symbol": "KIN82", "name": "850453", "MCODE_TYPE": "", "SUID": 2785309, "DEGREE": 2.0, "id_original": "850453", "MCODE_CLUSTER_ID": 0.0, "selected": false, "canonicalName": "KIN82"}, "position": {"x": 374.83152533024247, "y": 404.9984294052453}, "selected": false}, {"data": {"id": "2785306", "shared_name": "850444", "MCODE_SCORE": 0.0, "Gene": "850444", "Symbol": "TRX3", "name": "850444", "MCODE_TYPE": "", "SUID": 2785306, "DEGREE": 2.0, "id_original": "850444", "MCODE_CLUSTER_ID": 0.0, "selected": false, "canonicalName": "TRX3"}, "position": {"x": -141.55380105525558, "y": 127.33751021579218}, "selected": false}, {"data": {"id": "2785303", "shared_name": "850385", "MCODE_SCORE": 1.25, "Gene": "850385", "Symbol": "HSP30", "name": "850385", "MCODE_TYPE": "Clustered", "SUID": 2785303, "DEGREE": 5.0, "id_original": "850385", "MCODE_CLUSTER_ID": 6.0, "selected": false, "canonicalName": "HSP30"}, "position": {"x": 594.6048903509944, "y": 314.3218699570031}, "selected": false}, {"data": {"id": "2785300", "shared_name": "850317", "MCODE_SCORE": 1.0, "Gene": "850317", "Symbol": "GLK1", "name": "850317", "MCODE_TYPE": "Clustered", "SUID": 2785300, "DEGREE": 8.0, "id_original": "850317", "MCODE_CLUSTER_ID": 9.0, "selected": false, "canonicalName": "GLK1"}, "position": {"x": 91.66972494572099, "y": 217.80863677218866}, "selected": false}], "edges": [{"data": {"id": "2786206", "source": "2786107", "target": "2785621", "SCORE": "213", "shared_interaction": "Direct", "source_original": "5142379", "Name_B": "FMP48", "Name_A": "MIN8", "target_original": "852943", "Gene_B": "852943", "interaction": "Direct", "SUID": 2786206, "TYPE": "Direct", "Gene_A": "5142379", "support": "{\"StringDB\": \"physical\", \"StringEvidence\": \"experiments:0.179\", \"String_Score_Physical\": 213, \"String_Score_Combined\": 213}", "selected": false}, "selected": false}, {"data": {"id": "2786203", "source": "2786107", "target": "2785519", "SCORE": "432", "shared_interaction": "Direct", "source_original": "5142379", "Name_B": "COR1", "Name_A": "MIN8", "target_original": "852235", "Gene_B": "852235", "interaction": "Direct", "SUID": 2786203, "TYPE": "Direct", "Gene_A": "5142379", "support": "{\"StringDB\": \"physical\", \"StringEvidence\": \"experiments:0.408,coexpression:0.281\", \"BioGrid\": \"physical\", \"BioGrid_PubMed\": \"22342701\", \"BioGrid_Type\": \"Affinity Capture-MS\", \"String_Score_Physical\": 432, \"String_Score_Combined\": 591}", "selected": false}, "selected": false}, {"data": {"id": "2786200", "source": "2786107", "target": "2785336", "SCORE": "208", "shared_interaction": "Direct", "source_original": "5142379", "Name_B": "HSP104", "Name_A": "MIN8", "target_original": "850633", "Gene_B": "850633", "interaction": "Direct", "SUID": 2786200, "TYPE": "Direct", "Gene_A": "5142379", "support": "{\"BioGrid\": \"physical\", \"BioGrid_PubMed\": \"27373154\", \"BioGrid_Type\": \"Affinity Capture-MS\"}", "selected": false}, "selected": false}, {"data": {"id": "2786197", "source": "2786104", "target": "2786065", "SCORE": "958", "shared_interaction": "Direct", "source_original": "3799970", "Name_B": "RIP1", "Name_A": "COX26", "target_original": "856689", "Gene_B": "856689", "interaction": "Direct", "SUID": 2786197, "TYPE": "Direct", "Gene_A": "3799970", "support": "{\"StringDB\": \"physical\", \"StringEvidence\": \"experiments:0.851,database:0.708,coexpression:0.092\", \"BioGrid\": \"physical\", \"BioGrid_PubMed\": \"27083394\", \"BioGrid_Type\": \"Affinity Capture-Western\", \"String_Score_Physical\": 958, \"String_Score_Combined\": 962}", "selected": false}, "selected": false}, {"data": {"id": "2786194", "source": "2786104", "target": "2786029", "SCORE": "941", "shared_interaction": "Direct", "source_original": "3799970", "Name_B": "QCR10", "Name_A": "COX26", "target_original": "856390", "Gene_B": "856390", "interaction": "Direct", "SUID": 2786194, "TYPE": "Direct", "Gene_A": "3799970", "support": "{\"StringDB\": \"physical\", \"StringEvidence\": \"experiments:0.791,database:0.708,coexpression:0.138\", \"String_Score_Physical\": 941, \"String_Score_Combined\": 949}", "selected": false}, "selected": false}, {"data": {"id": "2786191", "source": "2786104", "target": "2786020", "SCORE": "941", "shared_interaction": "Direct", "source_original": "3799970", "Name_B": "QCR2", "Name_A": "COX26", "target_original": "856321", "Gene_B": "856321", "interaction": "Direct", "SUID": 2786191, "TYPE": "Direct", "Gene_A": "3799970", "support": "{\"StringDB\": \"physical\", \"StringEvidence\": \"experiments:0.791,database:0.708,textmining:0.299,coexpression:0.058\", \"String_Score_Physical\": 941, \"String_Score_Combined\": 961}", "selected": false}, "selected": false}, {"data": {"id": "2786188", "source": "2786104", "target": "2785948", "SCORE": "958", "shared_interaction": "Direct", "source_original": "3799970", "Name_B": "COX5A", "Name_A": "COX26", "target_original": "855675", "Gene_B": "855675", "interaction": "Direct", "SUID": 2786188, "TYPE": "Direct", "Gene_A": "3799970", "support": "{\"StringDB\": \"physical\", \"StringEvidence\": \"experiments:0.851,database:0.708,coexpression:0.086\", \"BioGrid\": \"physical\", \"BioGrid_PubMed\": \"27083394\", \"BioGrid_Type\": \"Affinity Capture-Western\", \"String_Score_Physical\": 958, \"String_Score_Combined\": 961}", "selected": false}, "selected": false}, {"data": {"id": "2786185", "source": "2786104", "target": "2785921", "SCORE": "941", "shared_interaction": "Direct", "source_original": "3799970", "Name_B": "COX7", "Name_A": "COX26", "target_original": "855298", "Gene_B": "855298", "interaction": "Direct", "SUID": 2786185, "TYPE": "Direct", "Gene_A": "3799970", "support": "{\"StringDB\": \"physical\", \"StringEvidence\": \"experiments:0.791,database:0.708,textmining:0.131,coexpression:0.086\", \"String_Score_Physical\": 941, \"String_Score_Combined\": 953}", "selected": false}, "selected": false}, {"data": {"id": "2786182", "source": "2786104", "target": "2785834", "SCORE": "958", "shared_interaction": "Direct", "source_original": "3799970", "Name_B": "COX5B", "Name_A": "COX26", "target_original": "854695", "Gene_B": "854695", "interaction": "Direct", "SUID": 2786182, "TYPE": "Direct", "Gene_A": "3799970", "support": "{\"StringDB\": \"physical\", \"StringEvidence\": \"experiments:0.851,database:0.708\", \"BioGrid\": \"physical\", \"BioGrid_PubMed\": \"27083394\", \"BioGrid_Type\": \"Affinity Capture-Western\", \"String_Score_Physical\": 958, \"String_Score_Combined\": 958}", "selected": false}, "selected": false}, {"data": {"id": "2786179", "source": "2786104", "target": "2785789", "SCORE": "941", "shared_interaction": "Direct", "source_original": "3799970", "Name_B": "CYT1", "Name_A": "COX26", "target_original": "854231", "Gene_B": "854231", "interaction": "Direct", "SUID": 2786179, "TYPE": "Direct", "Gene_A": "3799970", "support": "{\"StringDB\": \"physical\", \"StringEvidence\": \"experiments:0.791,database:0.708,textmining:0.506,coexpression:0.205\", \"String_Score_Physical\": 941, \"String_Score_Combined\": 977}", "selected": false}, "selected": false}, {"data": {"id": "2786176", "source": "2786104", "target": "2785666", "SCORE": "958", "shared_interaction": "Direct", "source_original": "3799970", "Name_B": "QCR8", "Name_A": "COX26", "target_original": "853273", "Gene_B": "853273", "interaction": "Direct", "SUID": 2786176, "TYPE": "Direct", "Gene_A": "3799970", "support": "{\"StringDB\": \"physical\", \"StringEvidence\": \"experiments:0.851,database:0.708,textmining:0.108\", \"BioGrid\": \"physical\", \"BioGrid_PubMed\": \"27083394\", \"BioGrid_Type\": \"Affinity Capture-Western\", \"String_Score_Physical\": 958, \"String_Score_Combined\": 962}", "selected": false}, "selected": false}, {"data": {"id": "2786173", "source": "2786104", "target": "2785633", "SCORE": "941", "shared_interaction": "Direct", "source_original": "3799970", "Name_B": "QCR9", "Name_A": "COX26", "target_original": "853095", "Gene_B": "853095", "interaction": "Direct", "SUID": 2786173, "TYPE": "Direct", "Gene_A": "3799970", "support": "{\"StringDB\": \"physical\", \"StringEvidence\": \"experiments:0.791,database:0.708,coexpression:0.209,textmining:0.131\", \"String_Score_Physical\": 941, \"String_Score_Combined\": 959}", "selected": false}, "selected": false}, {"data": {"id": "2786170", "source": "2786104", "target": "2785612", "SCORE": "311", "shared_interaction": "Direct", "source_original": "3799970", "Name_B": "PMC1", "Name_A": "COX26", "target_original": "852878", "Gene_B": "852878", "interaction": "Direct", "SUID": 2786170, "TYPE": "Direct", "Gene_A": "3799970", "support": "{\"StringDB\": \"physical\", \"StringEvidence\": \"experiments:0.283\", \"String_Score_Physical\": 311, \"String_Score_Combined\": 311}", "selected": false}, "selected": false}, {"data": {"id": "2786167", "source": "2786104", "target": "2785600", "SCORE": "966", "shared_interaction": "Direct", "source_original": "3799970", "Name_B": "COX4", "Name_A": "COX26", "target_original": "852688", "Gene_B": "852688", "interaction": "Direct", "SUID": 2786167, "TYPE": "Direct", "Gene_A": "3799970", "support": "{\"StringDB\": \"physical\", \"StringEvidence\": \"experiments:0.879,database:0.708,coexpression:0.106\", \"BioGrid\": \"physical\", \"BioGrid_PubMed\": \"27083394,23864706\", \"BioGrid_Type\": \"Affinity Capture-MS,Affinity Capture-Western\", \"String_Score_Physical\": 966, \"String_Score_Combined\": 969}", "selected": false}, "selected": false}, {"data": {"id": "2786164", "source": "2786104", "target": "2785519", "SCORE": "965", "shared_interaction": "Direct", "source_original": "3799970", "Name_B": "COR1", "Name_A": "COX26", "target_original": "852235", "Gene_B": "852235", "interaction": "Direct", "SUID": 2786164, "TYPE": "Direct", "Gene_A": "3799970", "support": "{\"StringDB\": \"physical\", \"StringEvidence\": \"experiments:0.876,database:0.708,textmining:0.088,coexpression:0.058\", \"BioGrid\": \"physical\", \"BioGrid_PubMed\": \"22342701\", \"BioGrid_Type\": \"Affinity Capture-MS\", \"String_Score_Physical\": 965, \"String_Score_Combined\": 970}", "selected": false}, "selected": false}, {"data": {"id": "2786161", "source": "2786104", "target": "2785501", "SCORE": "941", "shared_interaction": "Direct", "source_original": "3799970", "Name_B": "QCR7", "Name_A": "COX26", "target_original": "852142", "Gene_B": "852142", "interaction": "Direct", "SUID": 2786161, "TYPE": "Direct", "Gene_A": "3799970", "support": "{\"StringDB\": \"physical\", \"StringEvidence\": \"experiments:0.791,database:0.708,coexpression:0.106,textmining:0.083\", \"String_Score_Physical\": 941, \"String_Score_Combined\": 952}", "selected": false}, "selected": false}, {"data": {"id": "2786158", "source": "2786104", "target": "2785378", "SCORE": "941", "shared_interaction": "Direct", "source_original": "3799970", "Name_B": "COX8", "Name_A": "COX26", "target_original": "851111", "Gene_B": "851111", "interaction": "Direct", "SUID": 2786158, "TYPE": "Direct", "Gene_A": "3799970", "support": "{\"StringDB\": \"physical\", \"StringEvidence\": \"experiments:0.791,database:0.708,coexpression:0.058\", \"String_Score_Physical\": 941, \"String_Score_Combined\": 945}", "selected": false}, "selected": false}, {"data": {"id": "2786155", "source": "2786104", "target": "2785345", "SCORE": "994", "shared_interaction": "Direct", "source_original": "3799970", "Name_B": "COX12", "Name_A": "COX26", "target_original": "850727", "Gene_B": "850727", "interaction": "Direct", "SUID": 2786155, "TYPE": "Direct", "Gene_A": "3799970", "support": "{\"StringDB\": \"physical\", \"StringEvidence\": \"experiments:0.981,database:0.708,textmining:0.506,coexpression:0.065\", \"String_Score_Physical\": 994, \"String_Score_Combined\": 997}", "selected": false}, "selected": false}, {"data": {"id": "2786152", "source": "2786104", "target": "2785330", "SCORE": "941", "shared_interaction": "Direct", "source_original": "3799970", "Name_B": "QCR6", "Name_A": "COX26", "target_original": "850593", "Gene_B": "850593", "interaction": "Direct", "SUID": 2786152, "TYPE": "Direct", "Gene_A": "3799970", "support": "{\"StringDB\": \"physical\", \"StringEvidence\": \"experiments:0.791,database:0.708,textmining:0.316,coexpression:0.066\", \"String_Score_Physical\": 941, \"String_Score_Combined\": 962}", "selected": false}, "selected": false}, {"data": {"id": "2786149", "source": "2786101", "target": "2786005", "SCORE": "540", "shared_interaction": "Direct", "source_original": "1466480", "Name_B": "ATP20", "Name_A": "MCO10", "target_original": "856131", "Gene_B": "856131", "interaction": "Direct", "SUID": 2786149, "TYPE": "Direct", "Gene_A": "1466480", "support": "{\"StringDB\": \"physical\", \"StringEvidence\": \"database:0.520\", \"String_Score_Physical\": 540, \"String_Score_Combined\": 540}", "selected": false}, "selected": false}, {"data": {"id": "2786146", "source": "2786101", "target": "2785987", "SCORE": "540", "shared_interaction": "Direct", "source_original": "1466480", "Name_B": "ATP4", "Name_A": "MCO10", "target_original": "856027", "Gene_B": "856027", "interaction": "Direct", "SUID": 2786146, "TYPE": "Direct", "Gene_A": "1466480", "support": "{\"StringDB\": \"physical\", \"StringEvidence\": \"database:0.520\", \"String_Score_Physical\": 540, \"String_Score_Combined\": 540}", "selected": false}, "selected": false}, {"data": {"id": "2786143", "source": "2786101", "target": "2785963", "SCORE": "540", "shared_interaction": "Direct", "source_original": "1466480", "Name_B": "ATP15", "Name_A": "MCO10", "target_original": "855857", "Gene_B": "855857", "interaction": "Direct", "SUID": 2786143, "TYPE": "Direct", "Gene_A": "1466480", "support": "{\"StringDB\": \"physical\", \"StringEvidence\": \"database:0.520\", \"String_Score_Physical\": 540, \"String_Score_Combined\": 540}", "selected": false}, "selected": false}, {"data": {"id": "2786140", "source": "2786101", "target": "2785873", "SCORE": "540", "shared_interaction": "Direct", "source_original": "1466480", "Name_B": "ATP18", "Name_A": "MCO10", "target_original": "854893", "Gene_B": "854893", "interaction": "Direct", "SUID": 2786140, "TYPE": "Direct", "Gene_A": "1466480", "support": "{\"StringDB\": \"physical\", \"StringEvidence\": \"database:0.520\", \"String_Score_Physical\": 540, \"String_Score_Combined\": 540}", "selected": false}, "selected": false}, {"data": {"id": "2786137", "source": "2786101", "target": "2785774", "SCORE": "800", "shared_interaction": "Direct", "source_original": "1466480", "Name_B": "ATP19", "Name_A": "MCO10", "target_original": "854077", "Gene_B": "854077", "interaction": "Direct", "SUID": 2786137, "TYPE": "Direct", "Gene_A": "1466480", "support": "{\"StringDB\": \"physical\", \"StringEvidence\": \"database:0.791\", \"String_Score_Physical\": 800, \"String_Score_Combined\": 800}", "selected": false}, "selected": false}, {"data": {"id": "2786134", "source": "2786101", "target": "2785747", "SCORE": "540", "shared_interaction": "Direct", "source_original": "1466480", "Name_B": "ATP7", "Name_A": "MCO10", "target_original": "853853", "Gene_B": "853853", "interaction": "Direct", "SUID": 2786134, "TYPE": "Direct", "Gene_A": "1466480", "support": "{\"StringDB\": \"physical\", \"StringEvidence\": \"database:0.520\", \"String_Score_Physical\": 540, \"String_Score_Combined\": 540}", "selected": false}, "selected": false}, {"data": {"id": "2786131", "source": "2786101", "target": "2785705", "SCORE": "540", "shared_interaction": "Direct", "source_original": "1466480", "Name_B": "ATP2", "Name_A": "MCO10", "target_original": "853585", "Gene_B": "853585", "interaction": "Direct", "SUID": 2786131, "TYPE": "Direct", "Gene_A": "1466480", "support": "{\"StringDB\": \"physical\", \"StringEvidence\": \"database:0.520,textmining:0.131\", \"String_Score_Physical\": 540, \"String_Score_Combined\": 600}", "selected": false}, "selected": false}, {"data": {"id": "2786128", "source": "2786101", "target": "2785537", "SCORE": "540", "shared_interaction": "Direct", "source_original": "1466480", "Name_B": "ATP3", "Name_A": "MCO10", "target_original": "852327", "Gene_B": "852327", "interaction": "Direct", "SUID": 2786128, "TYPE": "Direct", "Gene_A": "1466480", "support": "{\"StringDB\": \"physical\", \"StringEvidence\": \"database:0.520\", \"String_Score_Physical\": 540, \"String_Score_Combined\": 540}", "selected": false}, "selected": false}, {"data": {"id": "2786125", "source": "2786101", "target": "2785504", "SCORE": "540", "shared_interaction": "Direct", "source_original": "1466480", "Name_B": "ATP1", "Name_A": "MCO10", "target_original": "852177", "Gene_B": "852177", "interaction": "Direct", "SUID": 2786125, "TYPE": "Direct", "Gene_A": "1466480", "support": "{\"StringDB\": \"physical\", \"StringEvidence\": \"database:0.520\", \"String_Score_Physical\": 540, \"String_Score_Combined\": 540}", "selected": false}, "selected": false}, {"data": {"id": "2786122", "source": "2786101", "target": "2785489", "SCORE": "540", "shared_interaction": "Direct", "source_original": "1466480", "Name_B": "ATP17", "Name_A": "MCO10", "target_original": "851983", "Gene_B": "851983", "interaction": "Direct", "SUID": 2786122, "TYPE": "Direct", "Gene_A": "1466480", "support": "{\"StringDB\": \"physical\", \"StringEvidence\": \"database:0.520\", \"String_Score_Physical\": 540, \"String_Score_Combined\": 540}", "selected": false}, "selected": false}, {"data": {"id": "2786119", "source": "2786101", "target": "2785477", "SCORE": "540", "shared_interaction": "Direct", "source_original": "1466480", "Name_B": "TIM11", "Name_A": "MCO10", "target_original": "851922", "Gene_B": "851922", "interaction": "Direct", "SUID": 2786119, "TYPE": "Direct", "Gene_A": "1466480", "support": "{\"StringDB\": \"physical\", \"StringEvidence\": \"database:0.520\", \"String_Score_Physical\": 540, \"String_Score_Combined\": 540}", "selected": false}, "selected": false}, {"data": {"id": "2786116", "source": "2786101", "target": "2785468", "SCORE": "540", "shared_interaction": "Direct", "source_original": "1466480", "Name_B": "ATP5", "Name_A": "MCO10", "target_original": "851892", "Gene_B": "851892", "interaction": "Direct", "SUID": 2786116, "TYPE": "Direct", "Gene_A": "1466480", "support": "{\"StringDB\": \"physical\", \"StringEvidence\": \"database:0.520\", \"String_Score_Physical\": 540, \"String_Score_Combined\": 540}", "selected": false}, "selected": false}, {"data": {"id": "2786113", "source": "2786101", "target": "2785414", "SCORE": "540", "shared_interaction": "Direct", "source_original": "1466480", "Name_B": "ATP16", "Name_A": "MCO10", "target_original": "851560", "Gene_B": "851560", "interaction": "Direct", "SUID": 2786113, "TYPE": "Direct", "Gene_A": "1466480", "support": "{\"StringDB\": \"physical\", \"StringEvidence\": \"database:0.520\", \"String_Score_Physical\": 540, \"String_Score_Combined\": 540}", "selected": false}, "selected": false}, {"data": {"id": "2786110", "source": "2786101", "target": "2785366", "SCORE": "540", "shared_interaction": "Direct", "source_original": "1466480", "Name_B": "ATP14", "Name_A": "MCO10", "target_original": "851002", "Gene_B": "851002", "interaction": "Direct", "SUID": 2786110, "TYPE": "Direct", "Gene_A": "1466480", "support": "{\"StringDB\": \"physical\", \"StringEvidence\": \"database:0.520\", \"String_Score_Physical\": 540, \"String_Score_Combined\": 540}", "selected": false}, "selected": false}, {"data": {"id": "2789539", "source": "2786074", "target": "2786080", "SCORE": "208", "shared_interaction": "Direct", "source_original": "856741", "Name_B": "PIC2", "Name_A": "GPA2", "target_original": "856779", "Gene_B": "856779", "interaction": "Direct", "SUID": 2789539, "TYPE": "Direct", "Gene_A": "856741", "support": "{\"StringDB\": \"physical\", \"StringEvidence\": \"experiments:0.174,coexpression:0.019\", \"String_Score_Physical\": 208, \"String_Score_Combined\": 222}", "selected": false}, "selected": false}, {"data": {"id": "2789536", "source": "2786065", "target": "2786074", "SCORE": "252", "shared_interaction": "Direct", "source_original": "856689", "Name_B": "GPA2", "Name_A": "RIP1", "target_original": "856741", "Gene_B": "856741", "interaction": "Direct", "SUID": 2789536, "TYPE": "Direct", "Gene_A": "856689", "support": "{\"StringDB\": \"physical\", \"StringEvidence\": \"experiments:0.221,textmining:0.048\", \"String_Score_Physical\": 252, \"String_Score_Combined\": 288}", "selected": false}, "selected": false}, {"data": {"id": "2789533", "source": "2786062", "target": "2786065", "SCORE": "987", "shared_interaction": "Direct", "source_original": "856672", "Name_B": "RIP1", "Name_A": "CYC7", "target_original": "856689", "Gene_B": "856689", "interaction": "Direct", "SUID": 2789533, "TYPE": "Direct", "Gene_A": "856672", "support": "{\"StringDB\": \"physical\", \"StringEvidence\": \"database:0.896,coexpression:0.890,experiments:0.876,textmining:0.746,neighborhood:0.138\", \"String_Score_Physical\": 987, \"String_Score_Combined\": 999}", "selected": false}, "selected": false}, {"data": {"id": "2789530", "source": "2786029", "target": "2786065", "SCORE": "985", "shared_interaction": "Direct", "source_original": "856390", "Name_B": "RIP1", "Name_A": "QCR10", "target_original": "856689", "Gene_B": "856689", "interaction": "Direct", "SUID": 2789530, "TYPE": "Direct", "Gene_A": "856390", "support": "{\"StringDB\": \"physical\", \"StringEvidence\": \"database:0.896,coexpression:0.871,experiments:0.851,textmining:0.429\", \"BioGrid\": \"physical\", \"BioGrid_PubMed\": \"29688796\", \"BioGrid_Type\": \"Affinity Capture-Western\", \"String_Score_Physical\": 985, \"String_Score_Combined\": 998}", "selected": false}, "selected": false}, {"data": {"id": "2789527", "source": "2786020", "target": "2786065", "SCORE": "999", "shared_interaction": "Direct", "source_original": "856321", "Name_B": "RIP1", "Name_A": "QCR2", "target_original": "856689", "Gene_B": "856689", "interaction": "Direct", "SUID": 2789527, "TYPE": "Direct", "Gene_A": "856321", "support": "{\"StringDB\": \"physical\", \"StringEvidence\": \"experiments:0.998,coexpression:0.991,textmining:0.922,database:0.896\", \"BioGrid\": \"physical\", \"BioGrid_PubMed\": \"25683140,24421313,21807901\", \"BioGrid_Type\": \"Affinity Capture-MS,Co-purification,Affinity Capture-Western\", \"String_Score_Physical\": 999, \"String_Score_Combined\": 999}", "selected": false}, "selected": false}, {"data": {"id": "2789524", "source": "2786020", "target": "2786062", "SCORE": "827", "shared_interaction": "Direct", "source_original": "856321", "Name_B": "CYC7", "Name_A": "QCR2", "target_original": "856672", "Gene_B": "856672", "interaction": "Direct", "SUID": 2789524, "TYPE": "Direct", "Gene_A": "856321", "support": "{\"StringDB\": \"physical\", \"StringEvidence\": \"experiments:0.820,textmining:0.473,coexpression:0.223\", \"String_Score_Physical\": 827, \"String_Score_Combined\": 929}", "selected": false}, "selected": false}, {"data": {"id": "2789521", "source": "2786020", "target": "2786029", "SCORE": "988", "shared_interaction": "Direct", "source_original": "856321", "Name_B": "QCR10", "Name_A": "QCR2", "target_original": "856390", "Gene_B": "856390", "interaction": "Direct", "SUID": 2789521, "TYPE": "Direct", "Gene_A": "856321", "support": "{\"StringDB\": \"physical\", \"StringEvidence\": \"textmining:0.897,database:0.896,experiments:0.883,coexpression:0.586\", \"BioGrid\": \"physical\", \"BioGrid_PubMed\": \"29688796,25683140\", \"BioGrid_Type\": \"Affinity Capture-Western\", \"String_Score_Physical\": 988, \"String_Score_Combined\": 999}", "selected": false}, "selected": false}, {"data": {"id": "2789518", "source": "2786017", "target": "2786071", "SCORE": "907", "shared_interaction": "Direct", "source_original": "856289", "Name_B": "GLC3", "Name_A": "GPH1", "target_original": "856705", "Gene_B": "856705", "interaction": "Direct", "SUID": 2789518, "TYPE": "Direct", "Gene_A": "856289", "support": "{\"StringDB\": \"physical\", \"StringEvidence\": \"coexpression:0.991,textmining:0.973,database:0.896,neighborhood:0.156,experiments:0.072\", \"String_Score_Physical\": 907, \"String_Score_Combined\": 999}", "selected": false}, "selected": false}, {"data": {"id": "2789515", "source": "2786014", "target": "2786074", "SCORE": "188", "shared_interaction": "Direct", "source_original": "856280", "Name_B": "GPA2", "Name_A": "TDA6", "target_original": "856741", "Gene_B": "856741", "interaction": "Direct", "SUID": 2789515, "TYPE": "Direct", "Gene_A": "856280", "support": "{\"StringDB\": \"physical\", \"StringEvidence\": \"experiments:0.153\", \"String_Score_Physical\": 188, \"String_Score_Combined\": 188}", "selected": false}, "selected": false}, {"data": {"id": "2789512", "source": "2786005", "target": "2786065", "SCORE": "214", "shared_interaction": "Direct", "source_original": "856131", "Name_B": "RIP1", "Name_A": "ATP20", "target_original": "856689", "Gene_B": "856689", "interaction": "Direct", "SUID": 2789512, "TYPE": "Direct", "Gene_A": "856131", "support": "{\"StringDB\": \"physical\", \"StringEvidence\": \"coexpression:0.923,experiments:0.180,textmining:0.081\", \"String_Score_Physical\": 214, \"String_Score_Combined\": 944}", "selected": false}, "selected": false}, {"data": {"id": "2789509", "source": "2786005", "target": "2786062", "SCORE": "335", "shared_interaction": "Direct", "source_original": "856131", "Name_B": "CYC7", "Name_A": "ATP20", "target_original": "856672", "Gene_B": "856672", "interaction": "Direct", "SUID": 2789509, "TYPE": "Direct", "Gene_A": "856131", "support": "{\"StringDB\": \"physical\", \"StringEvidence\": \"experiments:0.307,coexpression:0.170,textmining:0.014\", \"String_Score_Physical\": 335, \"String_Score_Combined\": 455}", "selected": false}, "selected": false}, {"data": {"id": "2789506", "source": "2786005", "target": "2786020", "SCORE": "962", "shared_interaction": "Direct", "source_original": "856131", "Name_B": "QCR2", "Name_A": "ATP20", "target_original": "856321", "Gene_B": "856321", "interaction": "Direct", "SUID": 2789506, "TYPE": "Direct", "Gene_A": "856131", "support": "{\"StringDB\": \"physical\", \"StringEvidence\": \"experiments:0.960,coexpression:0.745,textmining:0.708\", \"String_Score_Physical\": 962, \"String_Score_Combined\": 997}", "selected": false}, "selected": false}, {"data": {"id": "2789503", "source": "2785996", "target": "2786011", "SCORE": "930", "shared_interaction": "Direct", "source_original": "856089", "Name_B": "MCM16", "Name_A": "CTF19", "target_original": "856159", "Gene_B": "856159", "interaction": "Direct", "SUID": 2789503, "TYPE": "Direct", "Gene_A": "856089", "support": "{\"StringDB\": \"physical\", \"StringEvidence\": \"textmining:0.929,experiments:0.927\", \"BioGrid\": \"physical\", \"BioGrid_PubMed\": \"22561346\", \"BioGrid_Type\": \"Affinity Capture-MS\", \"String_Score_Physical\": 930, \"String_Score_Combined\": 995}", "selected": false}, "selected": false}, {"data": {"id": "2789500", "source": "2785993", "target": "2786092", "SCORE": "843", "shared_interaction": "Direct", "source_original": "856088", "Name_B": "VTC1", "Name_A": "VTC3", "target_original": "856803", "Gene_B": "856803", "interaction": "Direct", "SUID": 2789500, "TYPE": "Direct", "Gene_A": "856088", "support": "{\"StringDB\": \"physical\", \"StringEvidence\": \"textmining:0.956,coexpression:0.914,database:0.708,experiments:0.441\", \"BioGrid\": \"physical\", \"BioGrid_PubMed\": \"25315834,11823419\", \"BioGrid_Type\": \"Affinity Capture-MS,Affinity Capture-Western\", \"String_Score_Physical\": 843, \"String_Score_Combined\": 989}", "selected": false}, "selected": false}, {"data": {"id": "2789497", "source": "2785987", "target": "2786065", "SCORE": "964", "shared_interaction": "Direct", "source_original": "856027", "Name_B": "RIP1", "Name_A": "ATP4", "target_original": "856689", "Gene_B": "856689", "interaction": "Direct", "SUID": 2789497, "TYPE": "Direct", "Gene_A": "856027", "support": "{\"StringDB\": \"physical\", \"StringEvidence\": \"experiments:0.962,coexpression:0.941,textmining:0.081\", \"String_Score_Physical\": 964, \"String_Score_Combined\": 998}", "selected": false}, "selected": false}, {"data": {"id": "2789494", "source": "2785987", "target": "2786062", "SCORE": "192", "shared_interaction": "Direct", "source_original": "856027", "Name_B": "CYC7", "Name_A": "ATP4", "target_original": "856672", "Gene_B": "856672", "interaction": "Direct", "SUID": 2789494, "TYPE": "Direct", "Gene_A": "856027", "support": "{\"StringDB\": \"physical\", \"StringEvidence\": \"textmining:0.353,coexpression:0.170,experiments:0.157\", \"String_Score_Physical\": 192, \"String_Score_Combined\": 565}", "selected": false}, "selected": false}, {"data": {"id": "2789491", "source": "2785987", "target": "2786029", "SCORE": "315", "shared_interaction": "Direct", "source_original": "856027", "Name_B": "QCR10", "Name_A": "ATP4", "target_original": "856390", "Gene_B": "856390", "interaction": "Direct", "SUID": 2789491, "TYPE": "Direct", "Gene_A": "856027", "support": "{\"StringDB\": \"physical\", \"StringEvidence\": \"experiments:0.286,coexpression:0.215\", \"BioGrid\": \"physical\", \"BioGrid_PubMed\": \"29688796\", \"BioGrid_Type\": \"Affinity Capture-Western\", \"String_Score_Physical\": 315, \"String_Score_Combined\": 462}", "selected": false}, "selected": false}, {"data": {"id": "2789488", "source": "2785987", "target": "2786020", "SCORE": "963", "shared_interaction": "Direct", "source_original": "856027", "Name_B": "QCR2", "Name_A": "ATP4", "target_original": "856321", "Gene_B": "856321", "interaction": "Direct", "SUID": 2789488, "TYPE": "Direct", "Gene_A": "856027", "support": "{\"StringDB\": \"physical\", \"StringEvidence\": \"experiments:0.961,coexpression:0.952,textmining:0.689\", \"String_Score_Physical\": 963, \"String_Score_Combined\": 999}", "selected": false}, "selected": false}, {"data": {"id": "2789485", "source": "2785987", "target": "2786005", "SCORE": "999", "shared_interaction": "Direct", "source_original": "856027", "Name_B": "ATP20", "Name_A": "ATP4", "target_original": "856131", "Gene_B": "856131", "interaction": "Direct", "SUID": 2789485, "TYPE": "Direct", "Gene_A": "856027", "support": "{\"StringDB\": \"physical\", \"StringEvidence\": \"experiments:0.998,coexpression:0.958,textmining:0.911,database:0.791\", \"BioGrid\": \"physical\", \"BioGrid_PubMed\": \"9867807\", \"BioGrid_Type\": \"Co-purification\", \"String_Score_Physical\": 999, \"String_Score_Combined\": 999}", "selected": false}, "selected": false}, {"data": {"id": "2789482", "source": "2785981", "target": "2786017", "SCORE": "300", "shared_interaction": "Direct", "source_original": "855980", "Name_B": "GPH1", "Name_A": "RNY1", "target_original": "856289", "Gene_B": "856289", "interaction": "Direct", "SUID": 2789482, "TYPE": "Direct", "Gene_A": "855980", "support": "{\"StringDB\": \"physical\", \"StringEvidence\": \"database:0.270,coexpression:0.171\", \"String_Score_Physical\": 300, \"String_Score_Combined\": 419}", "selected": false}, "selected": false}, {"data": {"id": "2789479", "source": "2785972", "target": "2786074", "SCORE": "546", "shared_interaction": "Direct", "source_original": "855898", "Name_B": "GPA2", "Name_A": "TPK2", "target_original": "856741", "Gene_B": "856741", "interaction": "Direct", "SUID": 2789479, "TYPE": "Direct", "Gene_A": "855898", "support": "{\"StringDB\": \"physical\", \"StringEvidence\": \"textmining:0.921,experiments:0.318,database:0.307,coexpression:0.039\", \"BioGrid\": \"genetic\", \"BioGrid_PubMed\": \"10373537\", \"BioGrid_Type\": \"Synthetic Rescue\", \"String_Score_Physical\": 546, \"String_Score_Combined\": 965}", "selected": false}, "selected": false}, {"data": {"id": "2789476", "source": "2785966", "target": "2786074", "SCORE": "259", "shared_interaction": "Direct", "source_original": "855871", "Name_B": "GPA2", "Name_A": "USV1", "target_original": "856741", "Gene_B": "856741", "interaction": "Direct", "SUID": 2789476, "TYPE": "Direct", "Gene_A": "855871", "support": "{\"StringDB\": \"physical\", \"StringEvidence\": \"database:0.195,experiments:0.041,textmining:0.004\", \"String_Score_Physical\": 259, \"String_Score_Combined\": 262}", "selected": false}, "selected": false}, {"data": {"id": "2789473", "source": "2785966", "target": "2785972", "SCORE": "335", "shared_interaction": "Direct", "source_original": "855871", "Name_B": "TPK2", "Name_A": "USV1", "target_original": "855898", "Gene_B": "855898", "interaction": "Direct", "SUID": 2789473, "TYPE": "Direct", "Gene_A": "855871", "support": "{\"StringDB\": \"physical\", \"StringEvidence\": \"coexpression:0.256,database:0.197,experiments:0.137,textmining:0.019\", \"String_Score_Physical\": 335, \"String_Score_Combined\": 514}", "selected": false}, "selected": false}, {"data": {"id": "2789470", "source": "2785963", "target": "2786065", "SCORE": "248", "shared_interaction": "Direct", "source_original": "855857", "Name_B": "RIP1", "Name_A": "ATP15", "target_original": "856689", "Gene_B": "856689", "interaction": "Direct", "SUID": 2789470, "TYPE": "Direct", "Gene_A": "855857", "support": "{\"StringDB\": \"physical\", \"StringEvidence\": \"textmining:0.339,experiments:0.217,coexpression:0.116\", \"String_Score_Physical\": 248, \"String_Score_Combined\": 560}", "selected": false}, "selected": false}, {"data": {"id": "2789467", "source": "2785963", "target": "2786020", "SCORE": "252", "shared_interaction": "Direct", "source_original": "855857", "Name_B": "QCR2", "Name_A": "ATP15", "target_original": "856321", "Gene_B": "856321", "interaction": "Direct", "SUID": 2789467, "TYPE": "Direct", "Gene_A": "855857", "support": "{\"StringDB\": \"physical\", \"StringEvidence\": \"experiments:0.221,coexpression:0.142,textmining:0.019\", \"String_Score_Physical\": 252, \"String_Score_Combined\": 371}", "selected": false}, "selected": false}, {"data": {"id": "2789464", "source": "2785963", "target": "2786005", "SCORE": "992", "shared_interaction": "Direct", "source_original": "855857", "Name_B": "ATP20", "Name_A": "ATP15", "target_original": "856131", "Gene_B": "856131", "interaction": "Direct", "SUID": 2789464, "TYPE": "Direct", "Gene_A": "855857", "support": "{\"StringDB\": \"physical\", \"StringEvidence\": \"experiments:0.973,coexpression:0.927,database:0.708,textmining:0.509\", \"String_Score_Physical\": 992, \"String_Score_Combined\": 999}", "selected": false}, "selected": false}, {"data": {"id": "2789461", "source": "2785963", "target": "2785987", "SCORE": "999", "shared_interaction": "Direct", "source_original": "855857", "Name_B": "ATP4", "Name_A": "ATP15", "target_original": "856027", "Gene_B": "856027", "interaction": "Direct", "SUID": 2789461, "TYPE": "Direct", "Gene_A": "855857", "support": "{\"StringDB\": \"physical\", \"StringEvidence\": \"experiments:0.999,textmining:0.960,database:0.896,coexpression:0.261\", \"BioGrid\": \"physical\", \"BioGrid_PubMed\": \"9867807,22940862\", \"BioGrid_Type\": \"Affinity Capture-MS,Co-purification\", \"String_Score_Physical\": 999, \"String_Score_Combined\": 999}", "selected": false}, "selected": false}, {"data": {"id": "2789458", "source": "2785960", "target": "2786089", "SCORE": "208", "shared_interaction": "Direct", "source_original": "855836", "Name_B": "RGI1", "Name_A": "HSP82", "target_original": "856797", "Gene_B": "856797", "interaction": "Direct", "SUID": 2789458, "TYPE": "Direct", "Gene_A": "855836", "support": "{\"BioGrid\": \"physical\", \"BioGrid_PubMed\": \"26804907,26217697,25452130\", \"BioGrid_Type\": \"Affinity Capture-MS\"}", "selected": false}, "selected": false}, {"data": {"id": "2789455", "source": "2785960", "target": "2786068", "SCORE": "208", "shared_interaction": "Direct", "source_original": "855836", "Name_B": "UBC8", "Name_A": "HSP82", "target_original": "856704", "Gene_B": "856704", "interaction": "Direct", "SUID": 2789455, "TYPE": "Direct", "Gene_A": "855836", "support": "{\"BioGrid\": \"physical\", \"BioGrid_PubMed\": \"31399574\", \"BioGrid_Type\": \"Affinity Capture-MS\"}", "selected": false}, "selected": false}, {"data": {"id": "2789452", "source": "2785960", "target": "2786056", "SCORE": "208", "shared_interaction": "Direct", "source_original": "855836", "Name_B": "FMO1", "Name_A": "HSP82", "target_original": "856581", "Gene_B": "856581", "interaction": "Direct", "SUID": 2789452, "TYPE": "Direct", "Gene_A": "855836", "support": "{\"BioGrid\": \"genetic,physical\", \"BioGrid_PubMed\": \"17923092,15766533\", \"BioGrid_Type\": \"Two-hybrid,Synthetic Growth Defect\"}", "selected": false}, "selected": false}, {"data": {"id": "2789449", "source": "2785960", "target": "2786044", "SCORE": "208", "shared_interaction": "Direct", "source_original": "855836", "Name_B": "PAL2", "Name_A": "HSP82", "target_original": "856497", "Gene_B": "856497", "interaction": "Direct", "SUID": 2789449, "TYPE": "Direct", "Gene_A": "855836", "support": "{\"BioGrid\": \"physical\", \"BioGrid_PubMed\": \"26804907,26217697,25452130\", \"BioGrid_Type\": \"Affinity Capture-MS\"}", "selected": false}, "selected": false}, {"data": {"id": "2789446", "source": "2785960", "target": "2786026", "SCORE": "208", "shared_interaction": "Direct", "source_original": "855836", "Name_B": "RIM4", "Name_A": "HSP82", "target_original": "856361", "Gene_B": "856361", "interaction": "Direct", "SUID": 2789446, "TYPE": "Direct", "Gene_A": "855836", "support": "{\"BioGrid\": \"genetic,physical\", \"BioGrid_PubMed\": \"26804907,17923092\", \"BioGrid_Type\": \"Affinity Capture-MS,Synthetic Growth Defect\"}", "selected":
```
