## Additional File 3 for "Uncharacterized yeast gene *YBR238C,* an effector of TORC1 signaling in a mitochondrial feedback loop, accelerates cellular aging via *HAP4*- and *RMD9*-dependent mechanisms": PPINetwork.style.js

```
var styles=[{"format_version": "1.0", "generated_by": "cytoscape-3.9.1", "target_cytoscapejs_version": "~2.1", "title": "PPIColorByCluster", "style": [{"selector": "node", "css": {"font-family": "Dialog.plain", "font-weight": "normal", "font-size": 18, "color": "rgb(0,0,0)", "text-opacity": 1.0, "border-opacity": 1.0, "width": 35.0, "shape": "ellipse", "background-opacity": 1.0, "background-color": "rgb(0,153,204)", "border-color": "rgb(0,102,153)", "border-width": 4.0, "text-valign": "center", "text-halign": "right", "height": 35.0, "content": "data(Symbol)"}}, {"selector": "node[MCODE_CLUSTER_ID = 0.0]", "css": {"background-color": "rgb(188,189,220)"}}, {"selector": "node[MCODE_CLUSTER_ID = 2.0]", "css": {"background-color": "rgb(31,120,180)"}}, {"selector": "node[MCODE_CLUSTER_ID = 8.0]", "css": {"background-color": "rgb(255,127,0)"}}, {"selector": "node[MCODE_CLUSTER_ID = 9.0]", "css": {"background-color": "rgb(202,178,214)"}}, {"selector": "node[MCODE_CLUSTER_ID = 10.0]", "css": {"background-color": "rgb(106,61,154)"}}, {"selector": "node[MCODE_CLUSTER_ID = 11.0]", "css": {"background-color": "rgb(255,255,153)"}}, {"selector": "node[MCODE_CLUSTER_ID = 3.0]", "css": {"background-color": "rgb(178,223,138)"}}, {"selector": "node[MCODE_CLUSTER_ID = 1.0]", "css": {"background-color": "rgb(166,206,227)"}}, {"selector": "node[MCODE_CLUSTER_ID = 4.0]", "css": {"background-color": "rgb(51,160,44)"}}, {"selector": "node[MCODE_CLUSTER_ID = 5.0]", "css": {"background-color": "rgb(251,154,153)"}}, {"selector": "node[MCODE_CLUSTER_ID = 6.0]", "css": {"background-color": "rgb(227,26,28)"}}, {"selector": "node[MCODE_CLUSTER_ID = 7.0]", "css": {"background-color": "rgb(253,191,111)"}}, {"selector": "node:selected", "css": {"background-color": "rgb(255,255,0)"}}, {"selector": "edge", "css": {"text-opacity": 1.0, "content": "", "color": "rgb(0,0,0)", "font-family": "Dialog.plain", "font-weight": "normal", "font-size": 10, "line-style": "solid", "target-arrow-color": "rgb(0,0,0)", "opacity": 0.39215686274509803, "target-arrow-shape": "none", "width": 3.0, "source-arrow-color": "rgb(0,0,0)", "source-arrow-shape": "none", "line-color": "rgb(84,39,143)"}}, {"selector": "edge[SCORE > 1]", "css": {"width": 5.0}}, {"selector": "edge[SCORE = 1]", "css": {"width": 5.0}}, {"selector": "edge[SCORE > 0.3][SCORE < 1]", "css": {"width": "mapData(SCORE,0.3,1,2.0,5.0)"}}, {"selector": "edge[SCORE = 0.3]", "css": {"width": 2.0}}, {"selector": "edge[SCORE < 0.3]", "css": {"width": 2.0}}, {"selector": "edge:selected", "css": {"line-color": "rgb(255,0,0)"}}, {"selector": "node[DEGREE<=5]", "css": {"width": 20.0, "height": 20.0}}, {"selector": "node[DEGREE>5][DEGREE<20]", "css": {"width": "mapData(DEGREE,5,20,35.0,50.0)", "height": "mapData(DEGREE,5,20,35.0,50.0)"}}, {"selector": "node[DEGREE>=20]", "css": {"width": 50.0, "height": 50.0}}]}, {"format_version": "1.0", "generated_by": "cytoscape-3.9.1", "target_cytoscapejs_version": "~2.1", "title": "PPIColorByClusterNoLabel", "style": [{"selector": "node", "css": {"font-family": "Dialog.plain", "font-weight": "normal", "font-size": 20, "content": "", "color": "rgb(0,0,0)", "text-opacity": 1.0, "border-opacity": 1.0, "width": 35.0, "shape": "ellipse", "background-opacity": 1.0, "background-color": "rgb(0,153,204)", "border-color": "rgb(0,102,153)", "border-width": 4.0, "text-valign": "center", "text-halign": "right", "height": 35.0}}, {"selector": "node[MCODE_CLUSTER_ID = 0.0]", "css": {"background-color": "rgb(188,189,220)"}}, {"selector": "node[MCODE_CLUSTER_ID = 2.0]", "css": {"background-color": "rgb(31,120,180)"}}, {"selector": "node[MCODE_CLUSTER_ID = 8.0]", "css": {"background-color": "rgb(255,127,0)"}}, {"selector": "node[MCODE_CLUSTER_ID = 9.0]", "css": {"background-color": "rgb(202,178,214)"}}, {"selector": "node[MCODE_CLUSTER_ID = 10.0]", "css": {"background-color": "rgb(106,61,154)"}}, {"selector": "node[MCODE_CLUSTER_ID = 11.0]", "css": {"background-color": "rgb(255,255,153)"}}, {"selector": "node[MCODE_CLUSTER_ID = 3.0]", "css": {"background-color": "rgb(178,223,138)"}}, {"selector": "node[MCODE_CLUSTER_ID = 1.0]", "css": {"background-color": "rgb(166,206,227)"}}, {"selector": "node[MCODE_CLUSTER_ID = 4.0]", "css": {"background-color": "rgb(51,160,44)"}}, {"selector": "node[MCODE_CLUSTER_ID = 5.0]", "css": {"background-color": "rgb(251,154,153)"}}, {"selector": "node[MCODE_CLUSTER_ID = 6.0]", "css": {"background-color": "rgb(227,26,28)"}}, {"selector": "node[MCODE_CLUSTER_ID = 7.0]", "css": {"background-color": "rgb(253,191,111)"}}, {"selector": "node:selected", "css": {"background-color": "rgb(255,255,0)"}}, {"selector": "edge", "css": {"text-opacity": 1.0, "content": "", "color": "rgb(0,0,0)", "font-family": "Dialog.plain", "font-weight": "normal", "font-size": 10, "line-style": "solid", "target-arrow-color": "rgb(0,0,0)", "opacity": 0.39215686274509803, "target-arrow-shape": "none", "width": 3.0, "source-arrow-color": "rgb(0,0,0)", "source-arrow-shape": "none", "line-color": "rgb(84,39,143)"}}, {"selector": "edge[SCORE > 1]", "css": {"width": 5.0}}, {"selector": "edge[SCORE = 1]", "css": {"width": 5.0}}, {"selector": "edge[SCORE > 0.3][SCORE < 1]", "css": {"width": "mapData(SCORE,0.3,1,2.0,5.0)"}}, {"selector": "edge[SCORE = 0.3]", "css": {"width": 2.0}}, {"selector": "edge[SCORE < 0.3]", "css": {"width": 2.0}}, {"selector": "edge:selected", "css": {"line-color": "rgb(255,0,0)"}}, {"selector": "node[DEGREE<=5]", "css": {"width": 20.0, "height": 20.0}}, {"selector": "node[DEGREE>5][DEGREE<20]", "css": {"width": "mapData(DEGREE,5,20,35.0,50.0)", "height": "mapData(DEGREE,5,20,35.0,50.0)"}}, {"selector": "node[DEGREE>=20]", "css": {"width": 50.0, "height": 50.0}}]}, {"format_version": "1.0", "generated_by": "cytoscape-3.3.0", "target_cytoscapejs_version": "~2.1", "title": "default", "style": [{"selector": "node", "css": {"text-opacity": 1.0, "text-valign": "center", "text-halign": "right", "color": "rgb(0,0,0)", "font-family": "Dialog.plain", "font-weight": "normal", "border-opacity": 1.0, "border-color": "rgb(0,102,153)", "shape": "ellipse", "font-size": 20, "content": "data(Symbol)", "background-color": "rgb(153,204,255)", "height": 35.0, "background-opacity": 1.0, "width": 35.0, "border-width": 4.0}}, {"selector": "node[_GeneInGOAndHitList > 20]", "css": {"width": 50.0}}, {"selector": "node[_GeneInGOAndHitList = 20]", "css": {"width": 50.0}}, {"selector": "node[_GeneInGOAndHitList > 5][_GeneInGOAndHitList < 20]", "css": {"width": "mapData(_GeneInGOAndHitList,5,20,20.0,50.0)"}}, {"selector": "node[_GeneInGOAndHitList = 5]", "css": {"width": 20.0}}, {"selector": "node[_GeneInGOAndHitList < 5]", "css": {"width": 20.0}}, {"selector": "node[_GeneInGOAndHitList > 20]", "css": {"height": 50.0}}, {"selector": "node[_GeneInGOAndHitList = 20]", "css": {"height": 50.0}}, {"selector": "node[_GeneInGOAndHitList > 5][_GeneInGOAndHitList < 20]", "css": {"height": "mapData(_GeneInGOAndHitList,5,20,20.0,50.0)"}}, {"selector": "node[_GeneInGOAndHitList = 5]", "css": {"height": 20.0}}, {"selector": "node[_GeneInGOAndHitList < 5]", "css": {"height": 20.0}}, {"selector": "node:selected", "css": {"background-color": "rgb(255,255,0)"}}, {"selector": "edge", "css": {"font-size": 10, "line-style": "solid", "opacity": 0.39215686274509803, "color": "rgb(0,0,0)", "target-arrow-color": "rgb(0,0,0)", "source-arrow-color": "rgb(0,0,0)", "content": "", "text-opacity": 1.0, "target-arrow-shape": "none", "source-arrow-shape": "none", "font-family": "Dialog.plain", "font-weight": "normal", "width": 3.0, "line-color": "rgb(84,39,143)"}}, {"selector": "edge[SCORE > 1]", "css": {"width": 10.0}}, {"selector": "edge[SCORE = 1]", "css": {"width": 10.0}}, {"selector": "edge[SCORE > 0.3][SCORE < 1]", "css": {"width": "mapData(SCORE,0.3,1,2.0,10.0)"}}, {"selector": "edge[SCORE = 0.3]", "css": {"width": 2.0}}, {"selector": "edge[SCORE < 0.3]", "css": {"width": 2.0}}, {"selector": "edge:selected", "css": {"line-color": "rgb(255,0,0)"}}, {"selector": "node[DEGREE<=5]", "css": {"width": 20.0, "height": 20.0}}, {"selector": "node[DEGREE>5][DEGREE<20]", "css": {"width": "mapData(DEGREE,5,20,35.0,50.0)", "height": "mapData(DEGREE,5,20,35.0,50.0)"}}, {"selector": "node[DEGREE>=20]", "css": {"width": 50.0, "height": 50.0}}]}];
```
