## Supplementary figures and images for "Uncharacterized yeast gene *YBR238C,* an effector of TORC1 signaling in a mitochondrial feedback loop, accelerates cellular aging via *HAP4*- and *RMD9*-dependent mechanisms"

### ColorByCluster.png

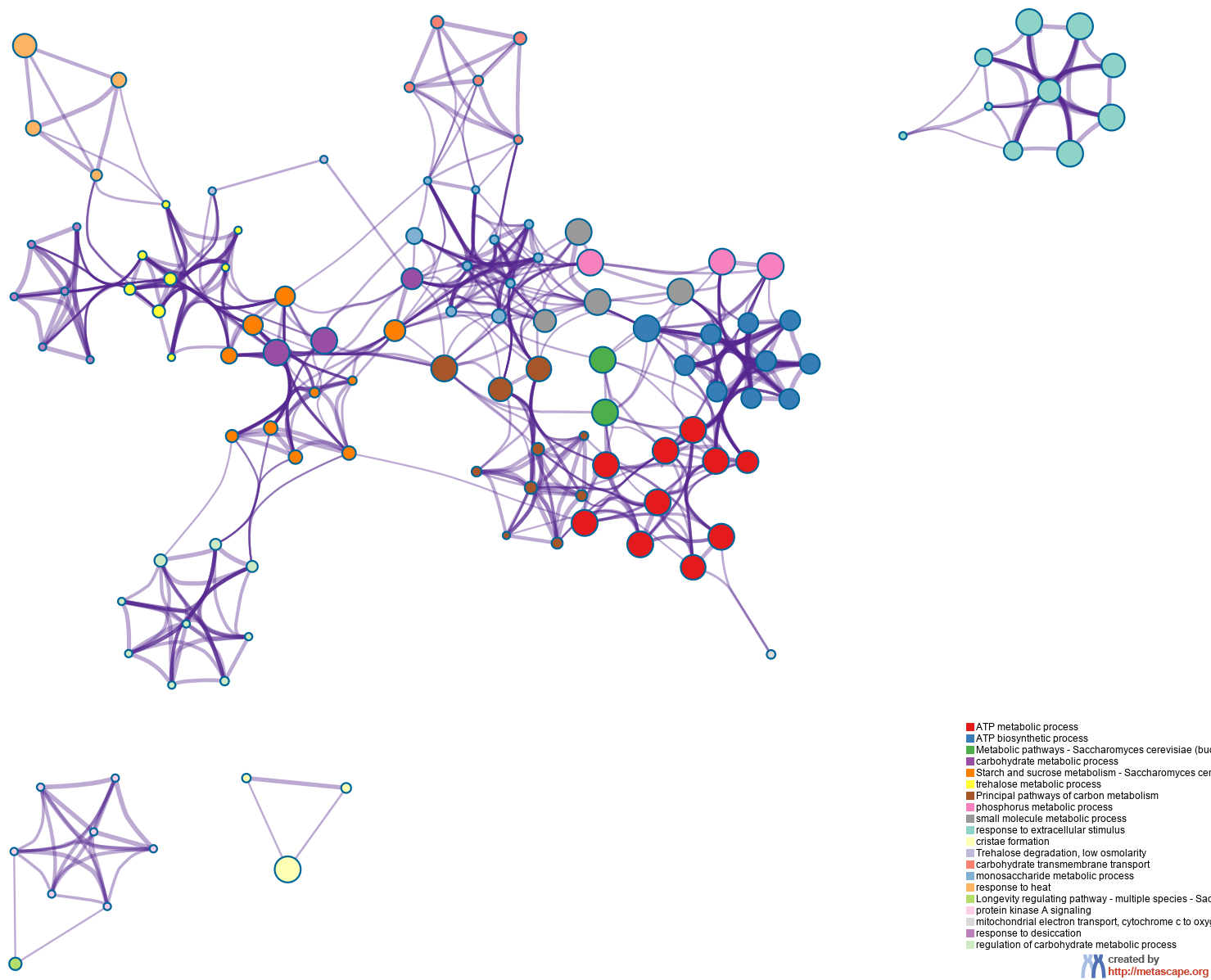

### ColorByPValue.pdf

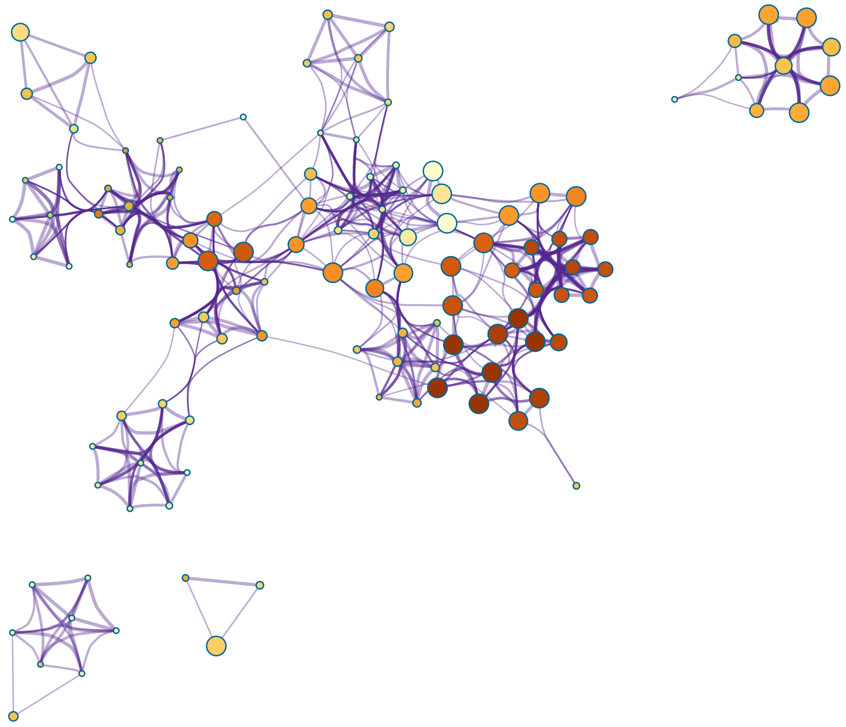

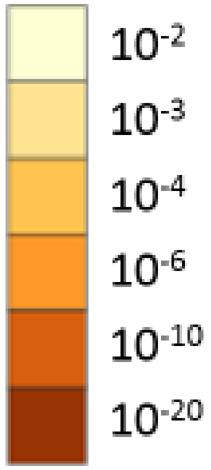


created by
http://metascape.org

### ColorByPValue.png

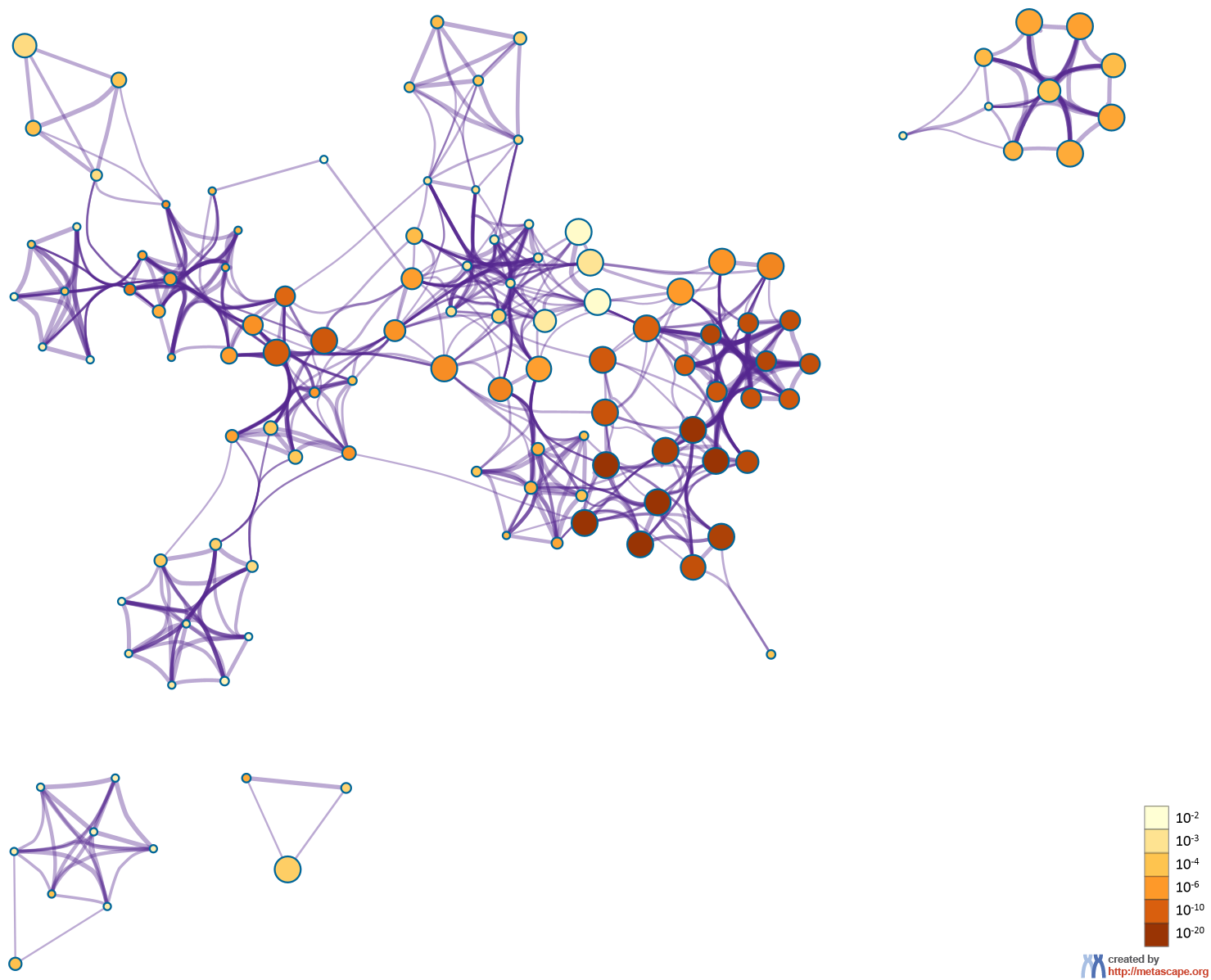

### CYS48.png

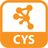

### HeatmapSelectedGO.pdf

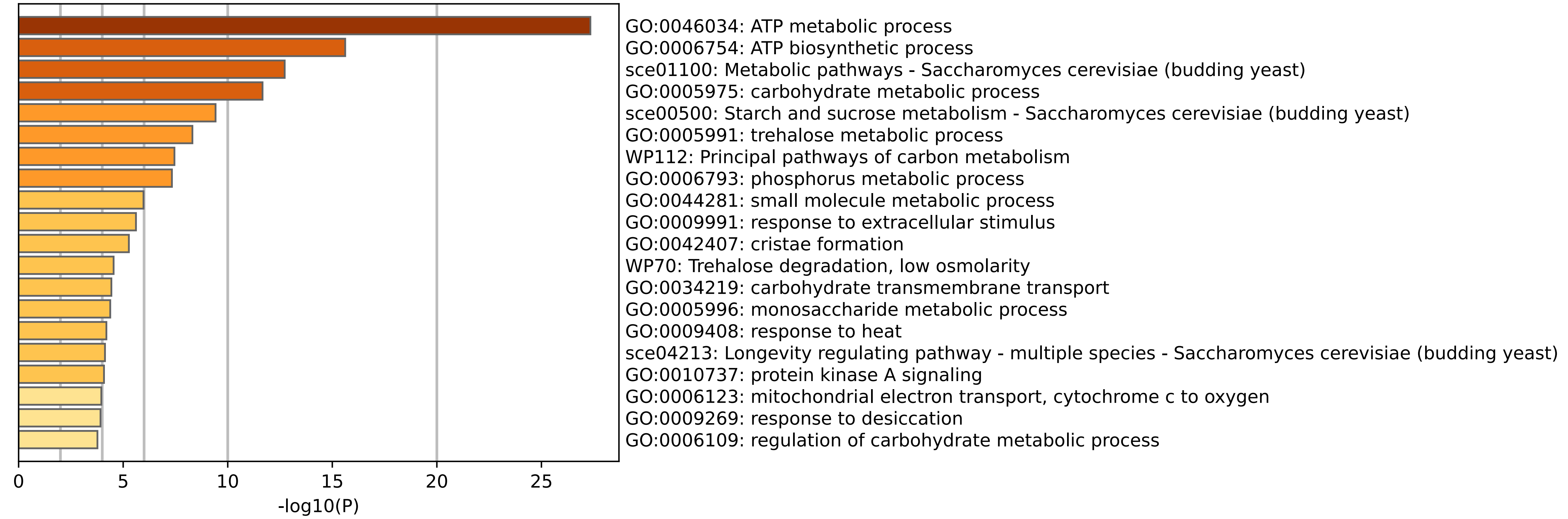

### HeatmapSelectedGO.png

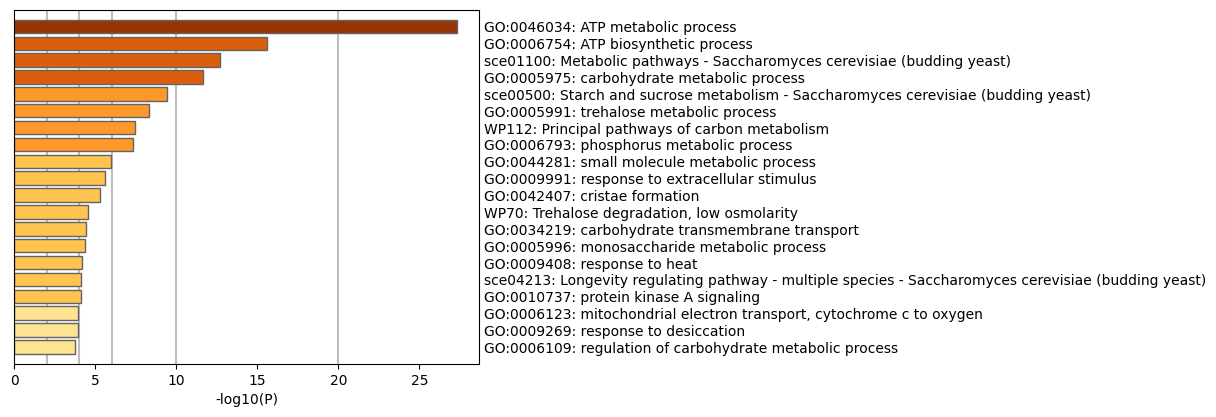

### HeatmapSelectedGOParent.pdf

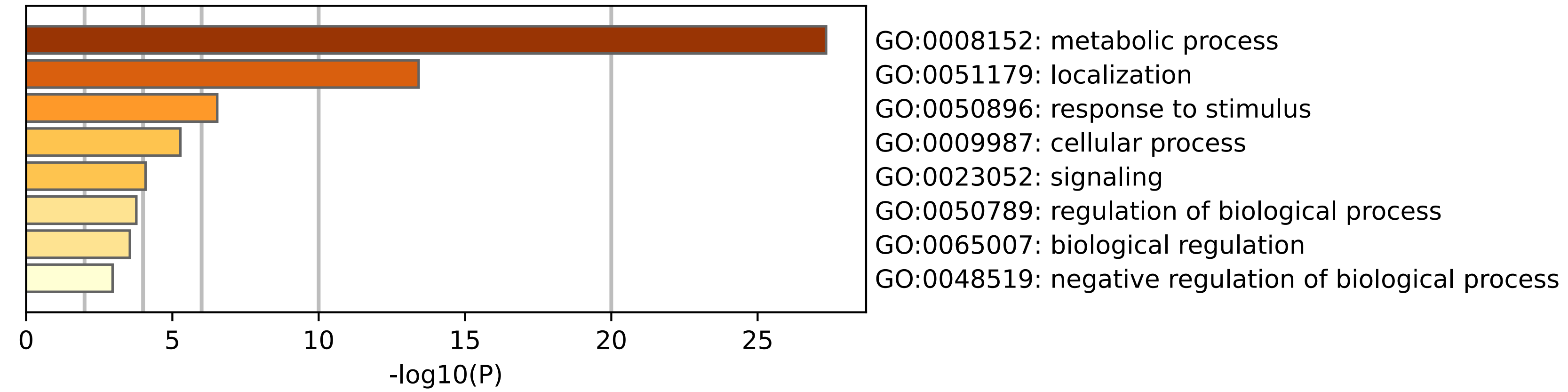

### HeatmapSelectedGOParent.png

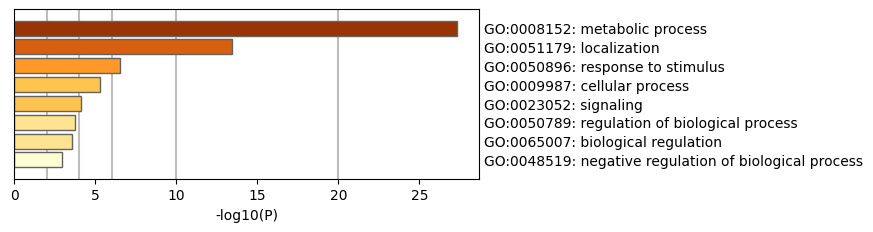

### HeatmapSelectedGOTop100.pdf

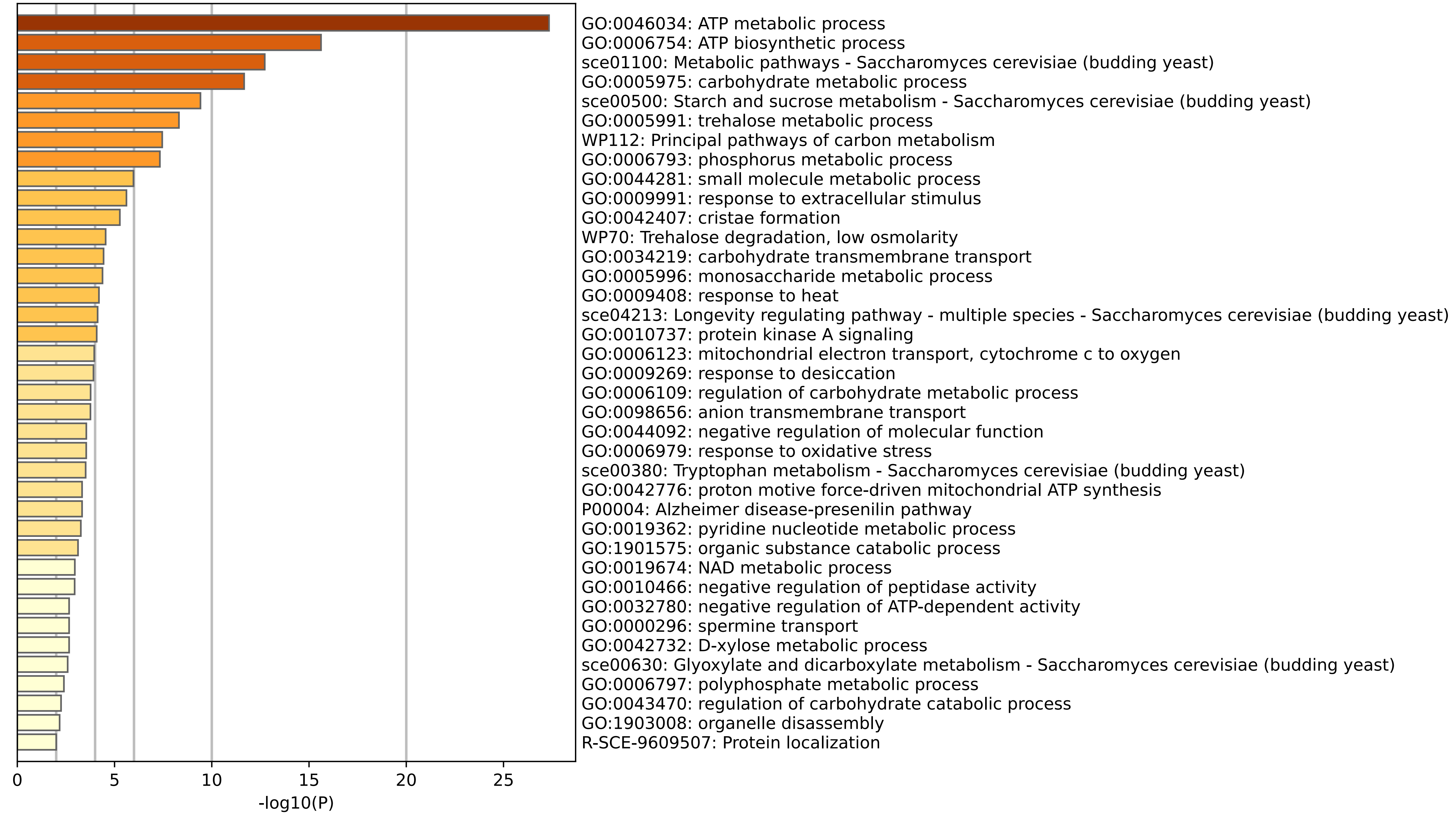

### HeatmapSelectedGOTop100.png

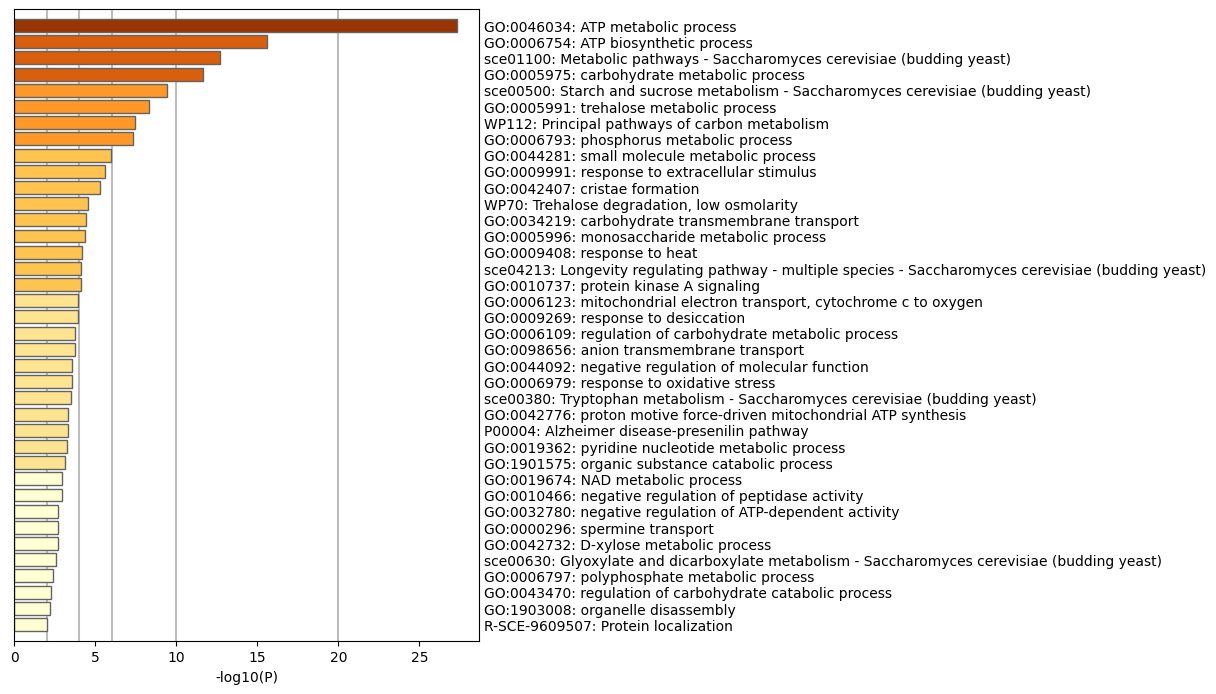

### MyList_MCODE_ALL_PPIColorByCluster.pdf

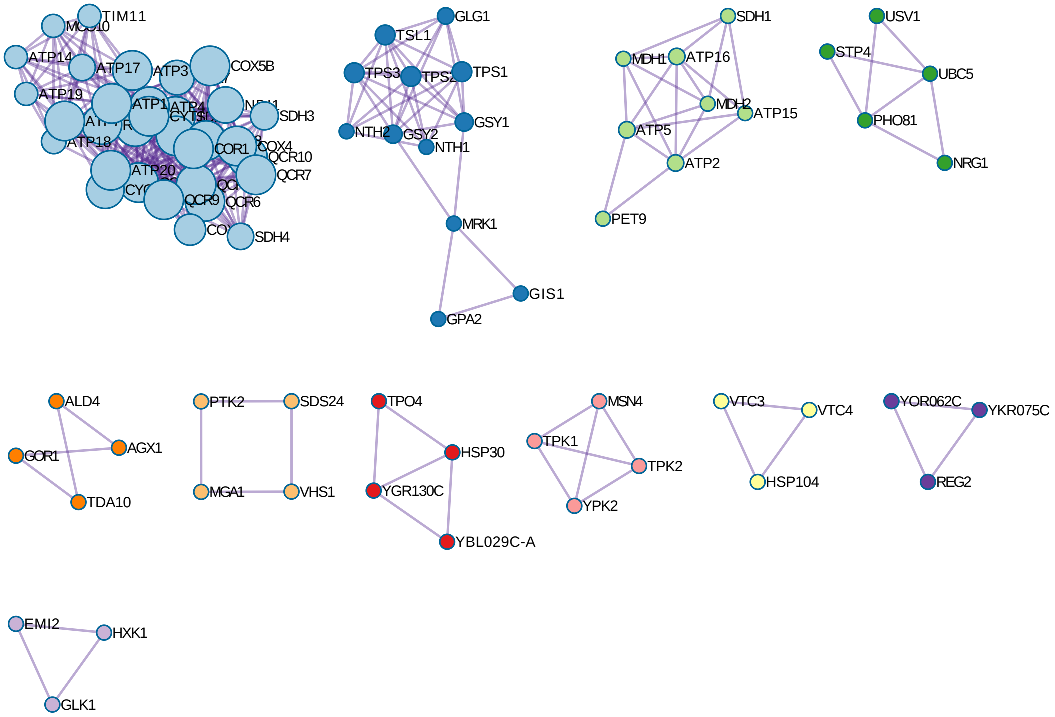

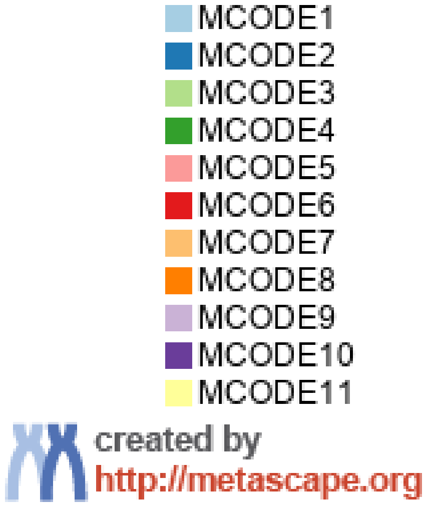

### MyList_MCODE_ALL_PPIColorByCluster.png

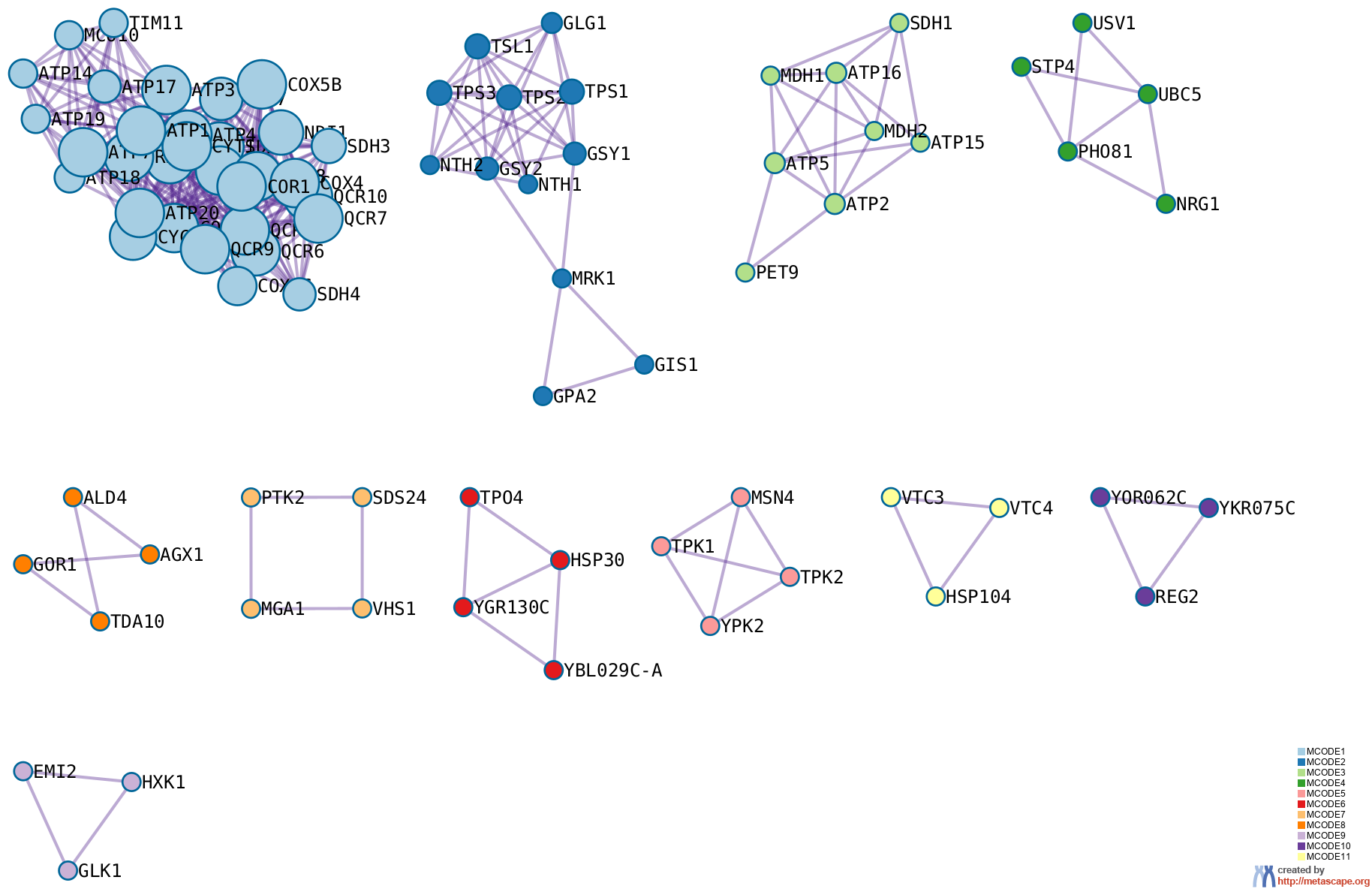

### MyList_PPIColorByCluster.png

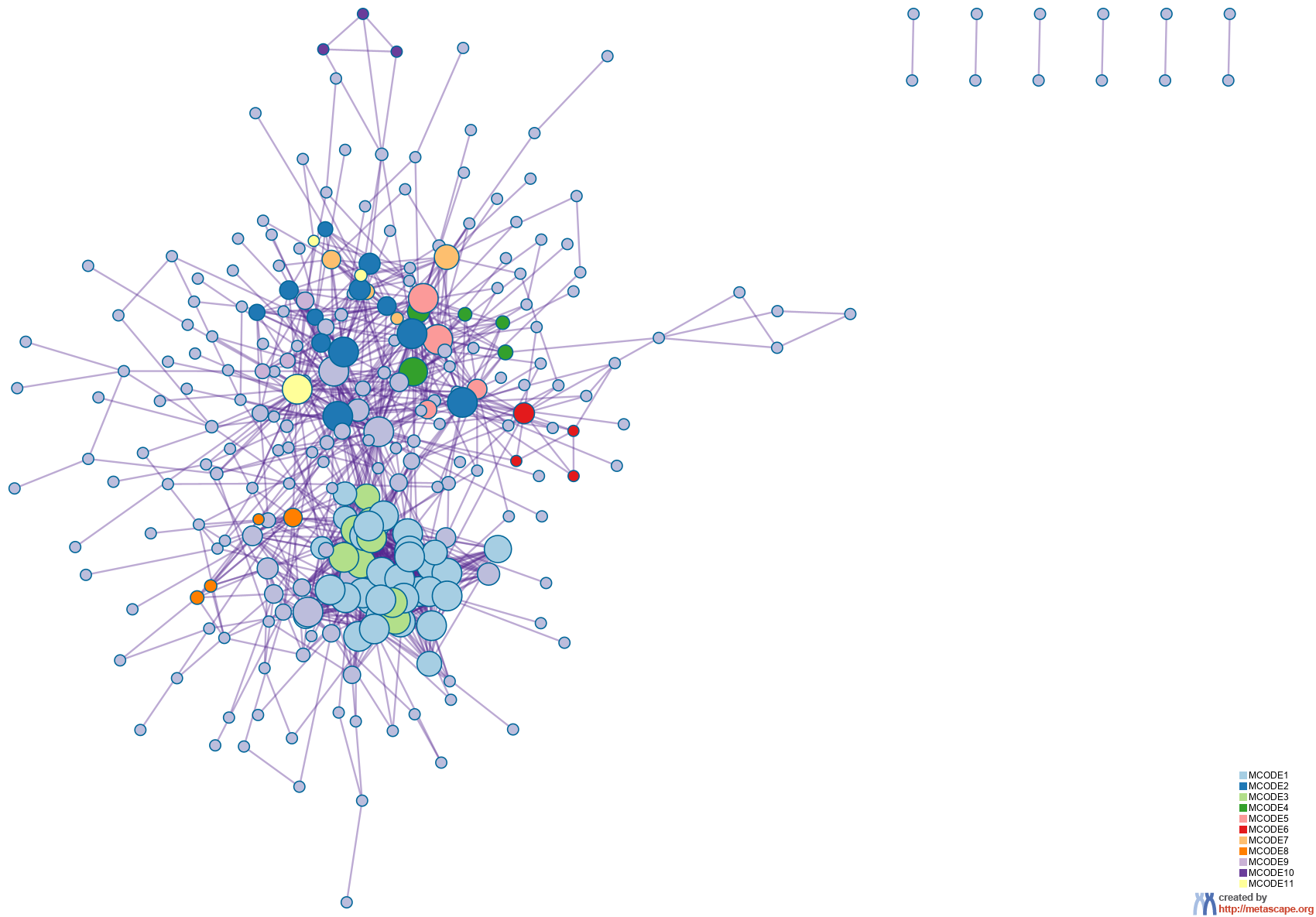

### PDF48.png

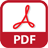

### SVG48.png

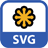

### WEB_CYS48.png

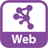
